# Polyomavirus ALTOs, but not MTs, downregulate viral early gene expression by activating the NF-κB pathway

**DOI:** 10.1101/2024.05.24.595774

**Authors:** Nicholas J. H. Salisbury, Supriya Amonkar, Joselyn Landazuri Vinueza, Joseph J. Carter, Ann Roman, Denise A. Galloway

## Abstract

Polyomaviruses are small, circular dsDNA viruses that can cause cancer. Alternative splicing of polyomavirus early transcripts generates large and small tumor antigens (LT, ST) that play essential roles in viral replication and tumorigenesis. Some polyomaviruses also express middle tumor antigens (MTs) or <u>A</u>lternate <u>LT O</u>RFs (ALTOs), which are evolutionarily related but have distinct gene structures. MTs are a splice variant of the early transcript whereas ALTOs are overprinted on the second exon of the LT transcript in an alternate reading frame and are translated via an alternative start codon. Merkel cell polyomavirus (MCPyV), the only human polyomavirus that causes cancer, encodes an ALTO but its role in the viral lifecycle and tumorigenesis has remained elusive. Here, we show MCPyV ALTO acts as a tumor suppressor and is silenced in Merkel cell carcinoma (MCC). Rescuing ALTO in MCC cells induces growth arrest and activates NF-κB signaling. ALTO activates NF-κB by binding SQSTM1 and TRAF2&3 via two <u>N</u>-<u>T</u>erminal <u>A</u>ctivating <u>R</u>egions (NTAR1+2), resembling Epstein-Barr virus (EBV) Latent Membrane Protein 1 (LMP1).. Following activation, NF-κB dimers bind the MCPyV non-coding control region (NCCR) and downregulate early transcription. Beyond MCPyV, NTAR motifs are conserved in other polyomavirus ALTOs, which activate NF-κB signaling, but are lacking in MTs that do not. Furthermore, polyomavirus ALTOs downregulate their respective viral early transcription in an NF-κB and NTAR dependent manner. Our findings suggest that ALTOs evolved to suppress viral replication and promote viral latency and that MCPyV ALTO must be silenced for MCC to develop.

## Introduction

Polyomaviruses are non-enveloped viruses with ∼5 kb double-stranded DNA genomes that infect fish, birds, and mammals and can cause cancer. Simian virus 40 (SV40) and murine polyomavirus (MuPyV) have been extensively studied due to their ability to transform human and rodent cells and have contributed to identifying key proteins that are involved in tumorigenesis such as p53(1), pRb(2), tyrosine kinases(3), and phosphoinositide-3 kinase (PI3K)(4). To date, fifteen human polyomaviruses have been identified. They typically cause asymptomatic, persistent infections, with morbidity primarily associated with immunosuppressed populations. Merkel cell polyomavirus (MCPyV) is the only human polyomavirus that causes cancer and causes ∼80% of Merkel cell carcinoma (MCC) cases (5).

Polyomavirus genomes have a tri-partite structure. The early region encodes two major regulatory proteins, the small and large tumor antigens (ST and LT, respectively), which are generated by alternative splicing of the early transcript and play important roles in the viral lifecycle and tumorigenesis. The late region encodes the structural proteins: VP1, VP2, and sometimes VP3, which form the viral capsid. The early and late regions are separated by a non-coding control region (NCCR) that contains the origin of replication and promoter elements that control early and late transcription. Some polyomaviruses encode other regulatory proteins such as middle tumor antigens (MTs), <u>A</u>lternative <u>LT O</u>RFs (ALTOs), and agnoproteins.

In virus-positive MCC, MCPyV LT and ST drive tumorigenesis by inactivating pRb and p53, respectively (6, 7). Meanwhile in virus-negative MCC, TP53 and RB1 are the two most commonly mutated genes. MCPyV LT contains an origin binding domain (OBD) that binds 5’-GRGGC-3’ repeats in the NCCR and a helicase/ATPase domain at the C terminus that is important for viral replication(8). In MCC, MCPyV LT is truncated (LT-t) by deletions and missense mutations that abrogate viral replication but retains its **L**x**C**x**E** motif that binds to and inhibits pRb(9, 10). MCPyV ST regulates LT stability, potentiates viral replication and has recently been shown to regulate early and late transcription(11-13). In MCC, ST remains full length and inactivates p53 by forming a complex with L-Myc that upregulates MDM2 and MDM4 via CK1α(14, 15).

In addition to inactivating p53, MCPyV ST has been shown to manipulate the NF-κB pathway, a master regulator of cell growth, inflammation, and antiviral responses(16, 17). The NF-κB pathway has two arms – canonical NF-κB mediated by RelA/NFKB1 p50 dimers and non-canonical NF-κB mediated by RelB/NFKB2 p52 dimers. Stimulation of the canonical NF-κB pathway activates the IκB kinase (IKK) complex, which allows RelA/p50 dimers to translocate to the nucleus, bind κB DNA sequences and regulate target gene expression. The IKK complex phosphorylates RelA on S536(18), found within RelA’s transcriptional activation domain, which enhances transactivation through increased binding to the histone acetyltransferase CBP (p300)(19). Stimulation of the non-canonical NF-κB pathway upregulates RelB, which is typically expressed at low levels in unstimulated cells, and induces proteasomal processing of NFKB2 p100 isoform to p52. MCPyV ST inactivates canonical NF-κB signaling by binding to and inhibiting NEMO, the regulatory subunit of the IKK complex(16) and has been shown to activate non-canonical NF-κB signaling in fibroblasts (17). BK and JC polyomaviruses (BKPyV and JCPyV) contain κB sequences in their non-coding control regions (NCCRs) and NF-κB signaling has been shown to upregulate their early gene expression (20, 21). Similarly, Epstein-Barr virus (EBV), a herpesvirus and DNA tumor virus, can inhibit NF-κB via its BZLF1 protein which binds directly to RelA and inhibits its transcriptional activity to promote lytic replication(22). EBV also activates NF-κB signaling via its Latent Membrane Protein 1 (LMP1) to promote viral latency and tumorigenesis in B cells(23, 24). LMP1 binds NF-κB pathway proteins via two C Terminal Activating Regions (CTAR1&2), which act upstream of the IKK complex(25-28). In summary, DNA viruses manipulate NF-κB signaling and its activation can regulate viral gene expression and promote tumorigenesis.

We previously showed that, in addition to LT and ST, MCPyV encodes an ALTO(29). ALTOs are evolutionarily related to but distinct from MTs, which are a third splice variant of the early transcript containing two exons (Fig S1). ALTOs are wholly encoded within the second exon of their respective LT and are translated using an alternative start codon found near the beginning of LT’s second exon (Fig 1A). We also identified ALTOs in gorilla, chimp, bat, and raccoon polyomaviruses(29). MuPyV and hamster polyomavirus MTs, have been relatively well characterized in contrast to ALTOs, in part due to their ability to transform cells independently of their respective LT and ST(30). MTs bear a resemblance to a constitutively active receptor tyrosine kinase, despite lacking any intrinsic enzymatic activity. MuPyV MT recruits Src tyrosine kinase family members (SFKs) which phosphorylate MT on tyrosine residues providing binding sites for SHC1, PI3K p85 and PLCγ1, which are subsequently tyrosine phosphorylated and activated, leading to activation of Ras/MAPK signaling and AKT/mTOR signaling. Recently, it has been reported that MCPyV ALTO can also bind SFKs, is phosphorylated on Y114 and can bind and activate PLCγ1 signaling(31). Mutant MCPyV genomes with ALTO Y114F, deficient for PLCγ1 activation, displayed increased viral replication compared to wild type(31). However, the role of MCPyV ALTO in tumorigenesis remains unexplored and the role of other polyomavirus ALTOs remains to be determined.

**Figure 1.**
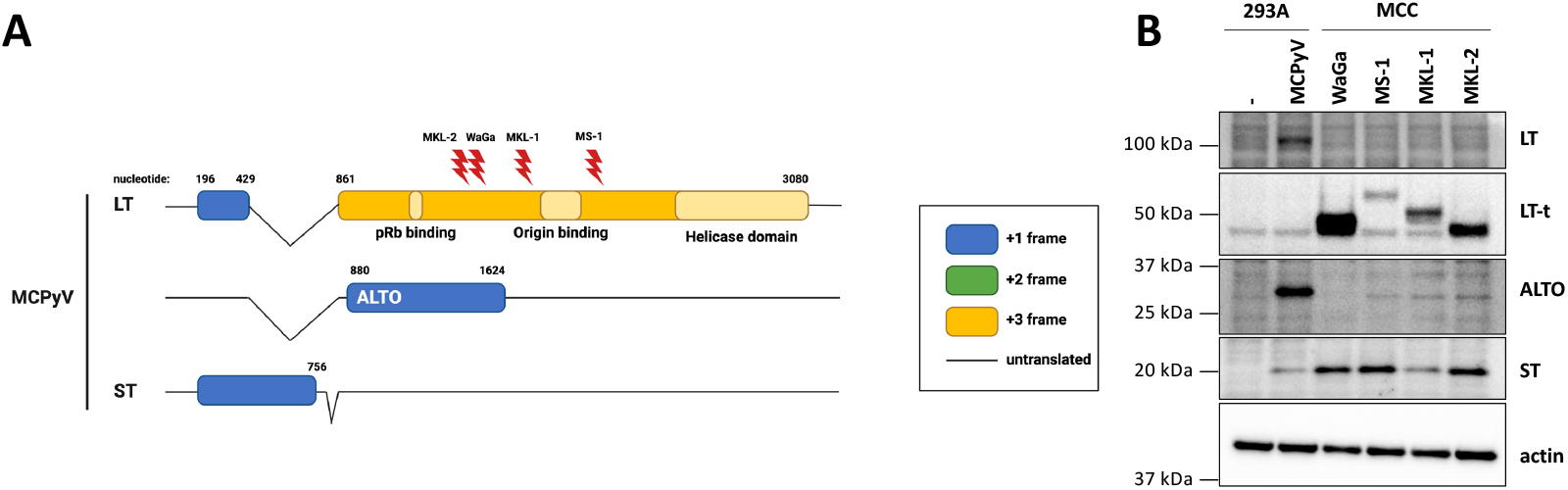
MCPyV ALTO is silenced in MCC. **A**) MCPyV early transcripts with MCC LT-truncating mutations for WaGa, MS-1, MKL-1, and MKL-2 cell lines indicated. Reading frames of coding sequences are indicated by blue (+1), green (+2), and yellow (+3). Untranslated sequences are represented by a solid black line. **B**) T antigen expression in 293A cells transfected with wt MCPyV genome and MCC cell lines.

Here, we show, in contrast to oncogenic MuPyV MT, MCPyV ALTO acts as a tumor suppressor and is silenced in MCC cell lines. When rescued, ALTO arrests cell growth and activates NF-κB signaling. MCPyV ALTO activates the NF-κB pathway by binding to SQSTM1 and TRAF2/3 via two <u>N</u>-<u>T</u>erminal <u>A</u>ctivating <u>R</u>egions, resembling EBV LMP1. Furthermore, we found that other PyV ALTOs, but not MTs, activate NF-κB using conserved NTAR motifs and repress their own viral early gene transcription in an NF-κB and NTAR dependent manner.Our findings reveal a novel regulatory network between PyV early proteins, whereby ALTOs activate NF-κB signaling to repress LT and ST expression and suggest that MCPyV ALTO must be silenced for MCC to develop.

## Results

### MCPyV ALTO is not expressed in virus-positive Merkel cell carcinoma

We previously showed that MCPyV ALTO is expressed during the viral lifecycle but is not required for viral replication. However, ALTO’s contribution to MCC development remained unknown. Since MCPyV ALTO is evolutionarily related to oncogenic MuPyV MT, we hypothesized that ALTO contributes to the development of MCC. As a first step, we analyzed ALTO protein expression in 293A cells transfected with wt MCPyV genome and in MKL-1, MKL-2, MS-1, and WaGa MCC cell lines (Fig. 1B). In genome-transfected cells we readily detected full-length LT, ST, and ALTO. In the four MCC cell lines, we detected truncated LT (LT-t) and ST but not ALTO.

### ALTO requires its C-terminal hydrophobic domain for robust expression

Since ALTO is overprinted on the second exon of LT, it is possible that LT-truncating mutations could impact the ALTO reading frame and prevent ALTO expression in MCC tumors. Therefore, we analyzed the ALTO coding sequences from the four MCC cell lines (Fig. S2A&B). In WaGa and MS-1, LT-truncating mutations have no impact on the ALTO reading frame and ALTO is wild type. In MKL-1, the LT-truncating deletion causes a frame shift in ALTO’s reading frame, resulting in substitution of the C terminal two residues (KQ) with five alternate amino acids (SYRVI). In MKL-2, the LT-truncating point mutation is silent in the ALTO reading frame; however, there is a downstream C->T point mutation in the ALTO reading frame that introduces a premature stop codon and truncates ALTO.

To assess the impact of these mutations on ALTO expression levels we subcloned the ALTO variants into vectors for transfection in 293A cells and compared ALTO protein levels (Fig. S2C). As previously reported(29, 31), we observed multiple ALTO isoforms by Western blotting with our polyclonal ALTO antisera in cells transfected with wt ALTO plasmid due to the presence of multiple translational start codons within the ALTO reading frame. Like wt ALTO, ALTO MKL-1 and its isoforms were also readily detectable by Western blotting. However, ALTO MKL-2 proteins were barely detectable. ALTO MKL-2 lacks the C-terminal hydrophobic domain (residues 224-246) which is conserved across polyomavirus MTs and ALTOs and is important for MuPyV MT’s localization to the plasma membrane. In fact, ALTO lacking just the C-terminal hydrophobic domain (ALTO ΔHD) was also poorly expressed in 293A cells, but we were able to rescue ALTO MKL-2 expression by adding back just the hydrophobic domain to ALTO MKL-2 (Fig. S2D). Thus, one mechanism to silence ALTO expression in MCC is to truncate ALTO’s C terminal hydrophobic domain.

### ALTO suppresses growth of MCC cell lines by activating the NF-κB pathway

As ALTO is not expressed in MCC cell lines, we predicted that it may function as a tumor suppressor. Upon rescuing ALTO expression in MKL-1 and MKL-2 cells via a doxycycline-inducible lentiviral vector, we observed a significant reduction in cell proliferation compared to control cells with doxycycline-inducible RFP expression (Fig. 2A&B), confirming that ALTO acts as a tumor suppressor in MCC. ALTO had a stronger effect on growth and higher expression in MKL-2 cells compared to MKL-1 cells (Fig. 2C).

**Figure 2.**
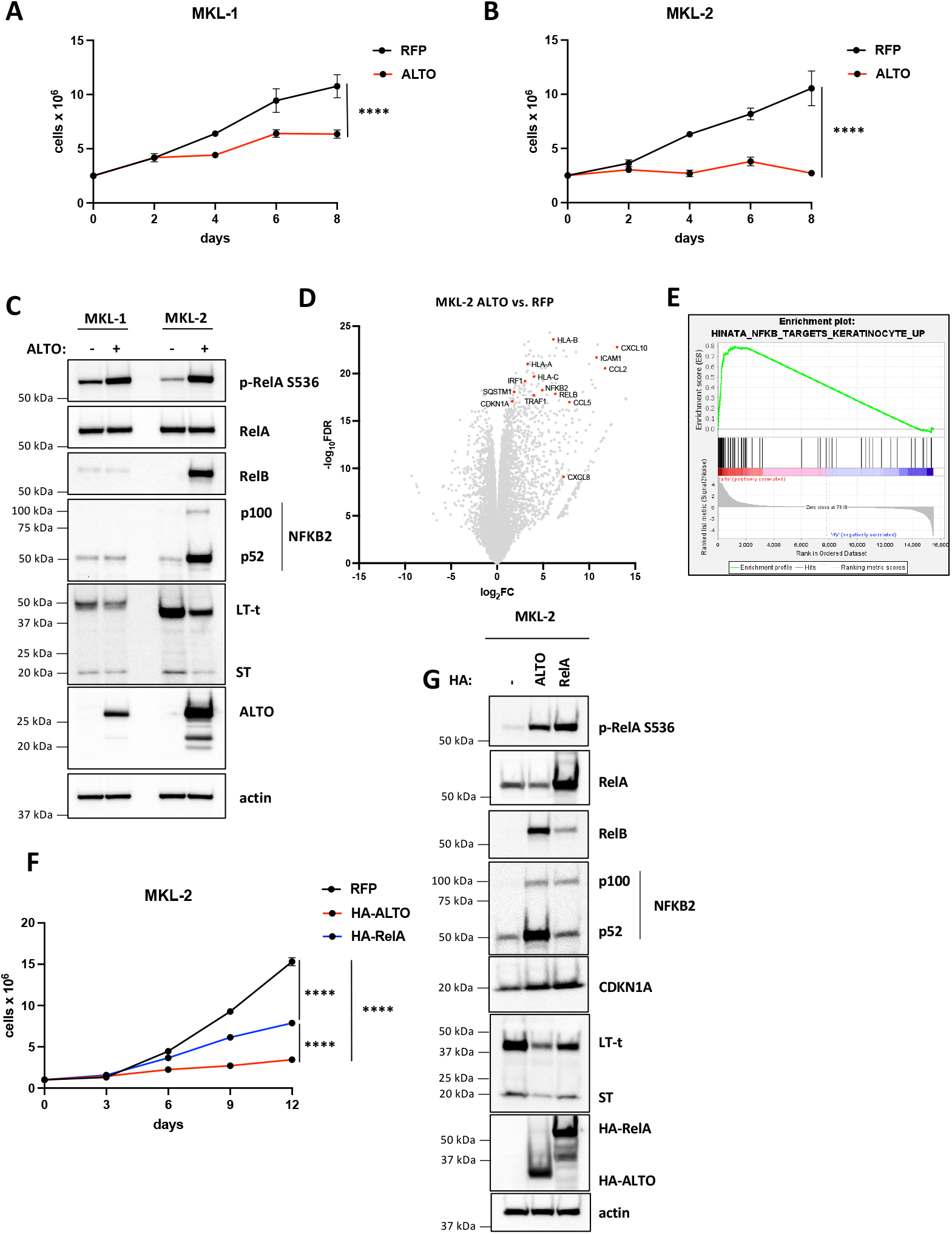
ALTO induces growth arrest in in MCC cells and activates NF-κB signaling. **A&B**) Growth curves of MKL-1 and MKL-2 cells expressing RFP or ALTO following doxycycline treatment. Each data point represents the mean ± standard deviation of three or more technical replicates. Statistical significance was calculated by 2-way ANOVA. **** P<0.0001. **C)** Western blots of MKL-1 and MKL-2 cells expressing RFP or ALTO. Cells were harvested after 6 days of doxycycline treatment. **D**) Volcano plot of differentially expressed genes in MKL-2 ALTO vs. RFP cells treated with doxycycline for 48 hours from RNA sequencing of extracts from three replicates. A subset of NF-κB target genes is highlighted. **E**) Enrichment plot for Hinata et al. (32) NF-κB targets upregulated in keratinocytes gene set. **F**) Growth curves of MKL-2 cells expressing RFP, HA-ALTO or HA-RelA following treatment with doxycycline. Each data point represents the mean ± standard deviation of three or more technical replicates. Statistical significance was calculated by 2-way ANOVA. ***** P<0.0001. **G**) Western blots of MKL-2 RFP, HA-ALTO and HA-RelA cells treated with doxycycline for 6 days.

To understand how ALTO induces growth arrest, we performed bulk RNA sequencing in MKL-2 cells expressing ALTO or RFP. Notably, ALTO upregulated CDKN1A expression 2.9 fold (P = 6x10^-20^) compared to RFP cells (Fig 2D). ALTO also upregulated numerous genes involved in antiviral responses such as chemokines and cytokines, antigen presentation and NF-κB pathway proteins (Fig. 2D). We performed Gene Set Enrichment Analysis (GSEA) and found a significant overlap (NES = 1.7, q= 1.1x10^-4^) of ALTO upregulated genes, including CDKN1A, with NF-κB target genes previously identified in keratinocytes by overexpressing RelA(32) (Fig. 2E). In agreement with the GSEA, ALTO induced RelA S536 phosphorylation in MKL-1 and MKL-2 cells, confirming NF-κB activation (Fig 2C). To further validate this finding, we performed NF-κB luciferase assays in 293A cells (Fig. S3A). Cotransfection of the NF-κB luciferase reporter with either HA-ALTO or HA-RelA plasmid upregulated luciferase activity 22-fold and 92-fold, respectively, compared to empty vector control cells . Furthermore, ALTO also induced RelA S536 phosphorylation in 293A cells (Fig. S3B).

Since CDKN1A appeared to be a RelA target gene in MKL-2 cells, we postulated that canonical NF-κB signaling contributes to growth arrest by ALTO. As expected, overexpressing HA-RelA in MKL-2 cells induced growth arrest (Fig. 2F), although milder than HA-ALTO, and upregulated CDKN1A expression to similar levels as HA-ALTO (Fig. 2G). Thus, canonical NF-κB activation and CDKN1A appear to contribute to ALTO-mediated growth arrest but ALTO also appears to regulate other pathways and genes that contribute to growth arrest. Besides CDKN1A, ALTO also upregulated expression of non-canonical NF-κB genes, NFKB2 and RelB, in MKL-2 cells (Fig. 2D). NFKB2 and RelB are direct target genes of RelA (32). As expected, both ALTO and RelA upregulated RelB and NFKB2 p100 and p52 expression in MKL-2 cells (Fig. 2C,G), although RelA did not upregulate RelB and NFKB2 to the same level as ALTO. Notably, ALTO does not activate non-canonical signaling in MKL-1 cells and causes a milder growth arrest in MKL-1 cells than in MKL-2 cells (Fig. 2A, B, C). Taken together, these data suggest that non-canonical NF-κB signaling makes an important contribution to ALTO-mediated growth arrest distinct from that of canonical NF-κB.

### ALTO binds SQSTM1 and TRAF2/3 to activate NF-κB signaling

ALTO has been reported to activate canonical NF-κB signaling in 293 cells by binding and activating PLCγ1, via phosphorylated Y114(31). While ALTO does indeed bind PLCγ1 via p-Y114 in MKL-2 cells, we did not observe any reduction in RelA S536 phosphorylation with ALTO Y114F compared to wt ALTO (Fig S4A&B). In 293A cells, ALTO Y114F was not deficient in its ability to induce RelA S536 phosphorylation or activate the NF-κB luciferase reporter (Fig S4C&D). Furthermore, wt PLCγ1 or constitutively active PLCγ1 S345F overexpression did not upregulate RelA S536 phosphorylation or NF-κB luciferase activity in 293A cells (Fig S4C&D). These findings suggest that other ALTO interactors mediate NF-κB activation.

To understand how ALTO activates NF-κB signaling in MKL-2 cells we performed proximity labeling mass spectrometry to identify additional ALTO interactors. We transduced MKL-2 cells with lentiviral vectors encoding the TurboID (TID) biotin ligase fused to the N terminus of ALTO (TID-ALTO) and confirmed TID-ALTO’s ability to activate NF-κB signaling (Fig 3A). MKL-2 TID-ALTO cells were labeled with biotin and biotinylated proteins were purified from cell lysates using streptavidin beads and analyzed by mass spectrometry (Fig. 3B).

**Figure 3.**
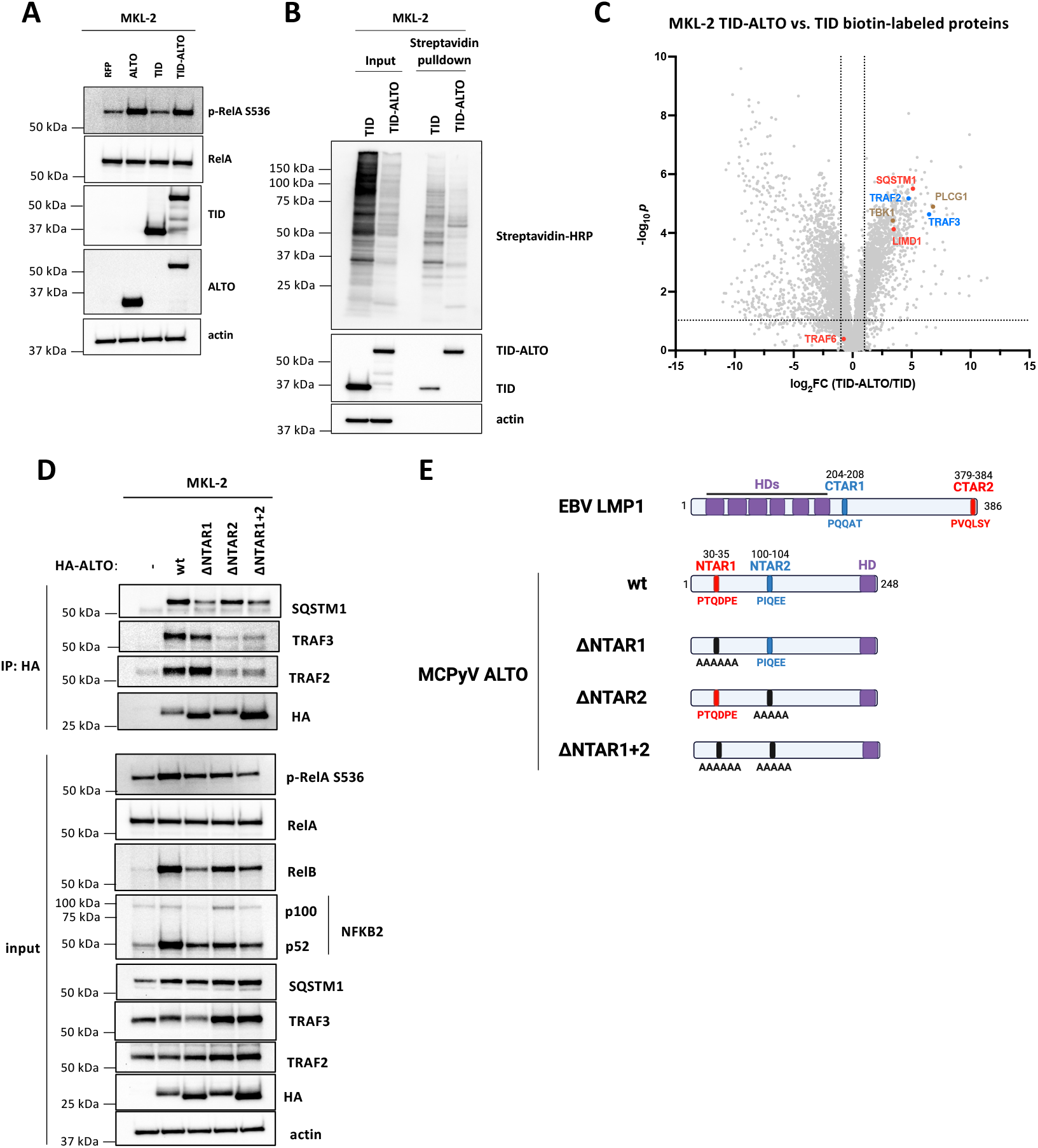
ALTO binds SQSTM1 and TRAF2&3 via NTAR1+2 motifs to activate NF-κB signaling. **A**) Western blots of MKL-2 cells expressing RFP, ALTO, TID or TID-ALTO. **B**) Western blots of streptavidin pulldown using lysates from MKL-2 TID and TID-ALTO cells biotinylated for 24 h prior to lysis. **C**) Volcano plot of proteins identified by mass spectrometry enriched in MKL-2 TID-ALTO vs TID samples. Proteins associated with LMP1’s CTAR1 and CTAR2 are highlighted in blue and red, respectively. **D**) Western blots of MKL-2 HA-ALTO variant cell lysates and anti-HA immunoprecipitations. **E**) Cartoon of EBV LMP1 and MCPyV ALTO variants. LMP1 CTAR2 and ALTO NTAR1 are highlighted in red. LMP1 CTAR1 and ALTO NTAR2 are highlighted in blue. Hydrophobic domains are highlighted in purple.

As expected, PLCγ1 was strongly enriched in TID-ALTO samples compared to TID only (log_2_FC = 6.8, Fig. 5C). EBV LMP1 interactors SQSTM1, LIMD1, TRAF2, TRAF3, were identified among the proteins most strongly enriched in TID-ALTO samples compared to TID control samples (Fig 3C) and we were able to co-immunoprecipitate (coIP) SQSTM1, TRAF2 and TRAF3 with HA-ALTO from MKL-2 cell lysates (Fig. 3D). We were unable to coIP LIMD1 (Fig. S4E), suggesting that it does not bind directly to ALTO but perhaps via a bridging interaction with SQSTM1, as reported for LMP1 (26). In contrast to LMP1 which binds directly to TRAF6 (28), TRAF6 was not enriched in TID-ALTO samples compared to TID only. As expected, we were unable to coIP TRAF6 with HA-ALTO (Fig S4E).

Next, we sought to identify ALTO mutants that were deficient for SQSTM1 and TRAF2/3 binding to assess their importance for NF-κB activation. LMP1 binds directly to TRAF6 via LMP1 379-384 (_379_**P**V**Q**LS**Y**_384_) contained within LMP1’s CTAR2 (28). LMP1 379-384 resembles the consensus TRAF6 binding sequence **P**x**E**xx**Z** (where Z is aromatic or acidic residue), except for a mismatch of glutamine in place of glutamate in the central position (**P**x**Q**xx**Z**). While we could not coIP TRAF6 with HA-ALTO in MKL-2 cells ALTO contains a **P**x**Q**xx**Z** motif (_30_**P**T**Q**DP**E**_35_), which we termed the <u>N</u>-<u>T</u>erminal <u>A</u>ctivating <u>R</u>egion 1 (NTAR1, Fig. 3E) and predicted may be important for SQSTM1 binding. HA-ALTO 30-35A (ΔNTAR1) was deficient for SQSTM1 binding, and induction of RelA S536 phosphorylation, RelB, and NFKB2 compared to wt ALTO(Fig. 3D), indicating that ALTO activates NF-κB in part by binding to SQSTM1.

TRAF2 and TRAF3 bind to LMP1 CTAR1 (_204_PQQAT_208_). Although ALTO does not contain a PQQAT motif, ALTO _100_PIQEE_104_ is identical to the TRAF2/3 binding site in human OX40(33). Thus, we named ALTO 100-104 the <u>N</u>-<u>T</u>erminal <u>A</u>ctivating <u>R</u>egion 2 (NTAR2, Fig. 3E). As predicted, HA-ALTO 100-104A (ΔNTAR2) was deficient for binding to TRAF2&3 andand induction of RelA S536 phosphorylation, RelB, and NFKB2 (Fig. 3D), indicating that ALTO activates NF-κB in part by binding to TRAF2&3.

Mutation of NTAR1 and NTAR2 alone caused only a partial reduction in NF-κB activation,suggesting that both NTAR1 and NTAR2 contribute to activation of NF-κB signaling independently. Thus, we assessed the ability of the double ΔNTAR1+2 mutant (ALTO 30-35A+100-104A) to activate NF-κB signaling. As expected, HA-ALTO ΔNTAR1+2 was deficient for binding to SQSTM1 and TRAF2&3. ALTO ΔNTAR1+2 completely failed to induce RelA S536 phosphorylation, but did not show any further reduction in RelB and NFKB2 induction compared to ALTO ΔNTAR1. Therefore, both SQSTM1 and TRAF2&3 binding appear to be necessary and sufficient for canonical NF-κB activation. While for non-canonical NF-κB activation, both SQSTM1 and TRAF2/3 binding are required for full activation but other interactors may also contribute. It also appears that SQSTM1 binding plays a more important role than TRAF2&3 binding, since ALTO ΔNTAR2 is less deficient in non-canonical NF-κB activation than ALTO ΔNTAR1 and ALTO ΔNTAR1&2 is no more deficient that ALTO ΔNTAR1.

TBK1 was also identified as a putative ALTO interactor (Fig 3C) and has been reported to play a role in canonical NF-κB activation (34). While we were able to coIP TBK1 with HA-ALTO wt, we could also coIP TBK1 to similar levels with the ALTO NTAR mutants (Fig. S4E). Since ALTO NTAR1+2 is completely deficient in its ability to activate canonical NF-κB but still retains TBK1 binding, it appears that TBK1 does not play a role in canonical NF-κB activation in MKL-2 cells.

### ALTO downregulates MCPyV early gene expression via NF-κB signaling

Next, we sought to better understand how ALTO and NF-κB signaling induces growth arrest in MCC cells. Since ALTO activates NF-κB in an analogous manner to LMP1, we postulated that ALTO may have evolved a similar function to LMP1. LMP1 represses the EBV lytic transcription factor BZLF1 via NF-κB(24). Therefore, we predicted that ALTO downregulates expression of MCPyV LT and ST, which control viral replication during the normal viral lifecycle and are essential for cell growth in MCC cells, via NF-κB signaling. Yang et al. recently showed that RelA binds to the MCPyV NCCR and upregulates MCPyV LT and ST expression (35). To test whether NF-κB proteins bind to the MCPyV NCCR in MCC cells, we performed DNA pulldowns using MKL-2 HA-ALTO lysates, which contain activated phosphorylated RelA, NFKB1 p50, RelB, and NKB2 p52 (Fig. 4A). We were able to readily detect enriched binding of RelB, p52 and p50 to the MCPyV NCCR compared to a similar size (∼500 bp) fragment of the coding region of the ampicillin resistance gene (AmpR), which we used as a negative control. In contrast to Yang et al. (35) who overexpressed RelA in their DNA pulldowns, we were only able to detect very weak RelA binding to MCPyV NCCR with MKL-2 cell lysates expressing endogenous levels of RelA. To test further whether RelA/p50 complexes bind to the MCPyV NCCR, we performed electromobility shift assays (EMSAs) using *in vitro* transcribed and translated RelA/p50 wheat germ extracts (Fig S5). We readily detected binding of RelA/p50 to probes derived from the BKPyV and JCPyV NCCRs which contain consensus κB (GGGRNYYYCC) motifs but did not detect any binding of RelA/p50 to probes spanning the entire MCPyV NCCR, which lacks a consensus κB motif. Since wheat germ extracts do not contain any other mammalian proteins besides RelA and NFKB1 p50, we conclude that RelA does not bind directly to the MCPyV NCCR but may be recruited as part of a complex in MKL-2 cells. Meanwhile, it appears that RelB binds the MCPyV NCCR in a complex with either p52 or p50.

**Figure 4.**
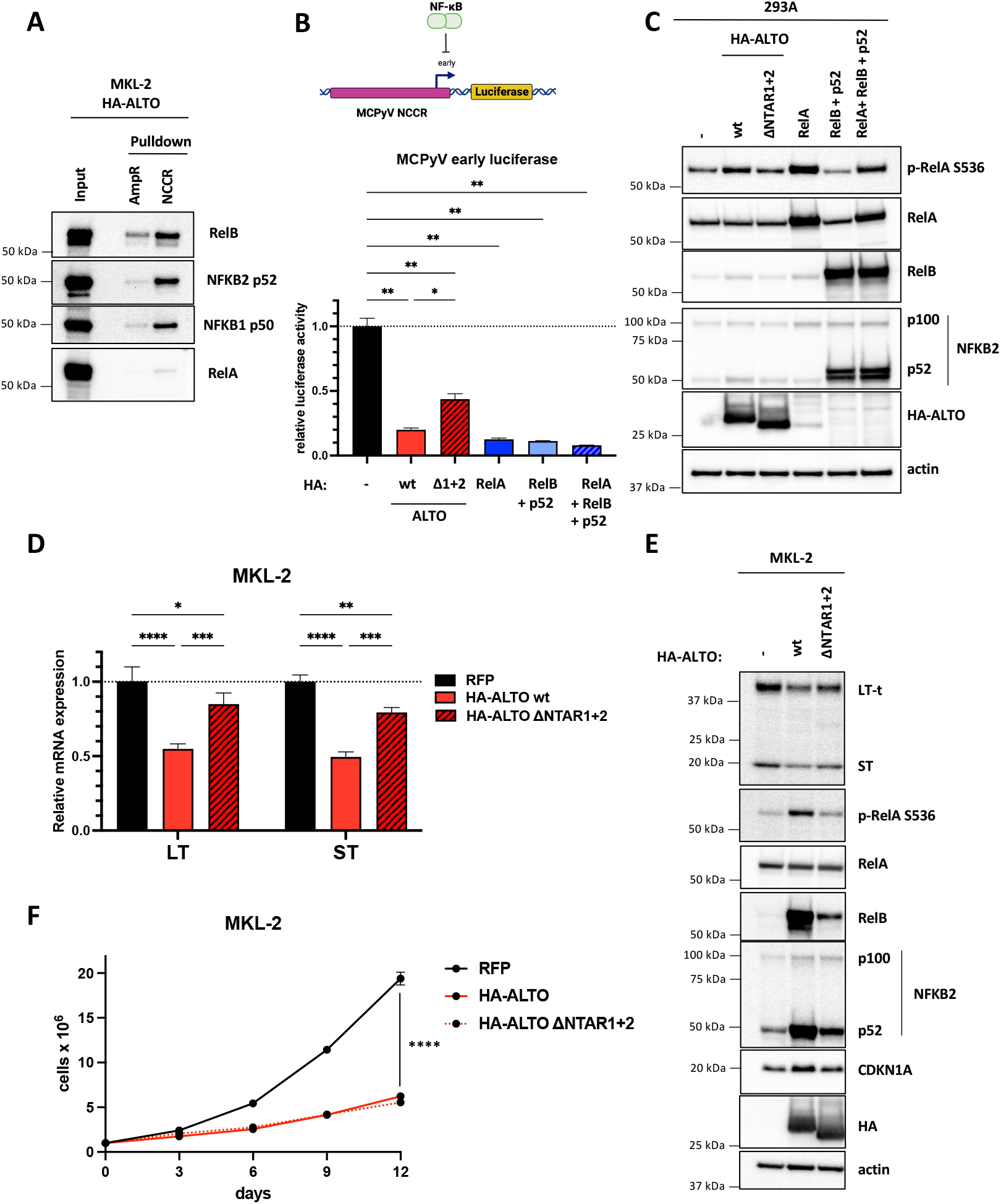
ALTO downregulates MCPyV early gene expression via NF-κB signaling. **A**) DNA pulldowns with MKL-2 HA-ALTO cell lysates incubated with biotinylated MCPyV NCCR or AmpR dsDNA coupled to streptavidin beads. **B**) MCPyV early luciferase in 293A cells transfected with HA-ALTO wt, HA-ALTO ΔNTAR1+2 or HA-RelA plasmids. Luciferase activity was normalized to CMV-driven renilla activity. Statistical significance was calculated by one-way ANOVA. * P<0.03, ** P<0.002. **C**) Western blots from 293A cells transfected with either HA-ALTO wt; HA-ALTO ΔNTAR1+2; HA-RelA; HA-RelB and FLAG-p52; or HA-RelA, HA-RelB and FLAG p52 plasmids. **D**) qRT-PCR analysis of relative LT and ST expression normalized to 36B4 expression in MKL-2 RFP, HA-ALTO wt or ΔNTAR1+2 cells treated with doxycycline for 6 days. The data show the mean ± the standard deviation of three technical replicates. Statistical significance was calculated by 2-way ANOVA. * P<0.03, ** P<0.003, *** P<0.0002, **** P<0.0001. **E**) Western blots of MKL-2 cells expressing RFP, HA-ALTO wt or ΔNTAR1+2 harvested after 6 days doxycycline induction. **F**) Growth curves of MKL-2 cells expressing RFP, HA-ALTO wt or ΔNTAR1+2. Each data point represents the mean ± standard deviation of three or more technical replicates. Statistical significance was calculated by 2-way ANOVA. **** P<0.0001.

To test whether ALTO and NF-κB regulate LT and ST transcription, we generated a MCPyV early luciferase reporter plasmid and cotransfected it with HA-ALTO; HA-ALTO ΔNTAR1&2; HA-RelA; HA-RelB and FLAG-p52; or HA-RelA, HA-RelB and FLAG-p52 plasmids into 293A cells (Fig 4B). Activation of NF-κB signaling either by ALTO or RelA or RelB+p52 or RelA+RelB+p52 overexpression caused downregulation of MCPyV early luciferase activity compared to empty vector transfected cells. In contrast, ALTO ΔNTAR1+2, which is deficient for both canonical and non-canonical NF-κB signaling (Fig 4C), showed a partial rescue in luciferase activity, indicating that NF-κB activation is important for ALTO to downregulate MCPyV early gene expression but that other pathways may contribute.

To validate that ALTO downregulates MCPyV early gene expression via NF-κB, we analyzed LT and ST mRNA and protein levels in MKL-2 cells expressing HA-ALTO wt or ΔNTAR1+2 (Fig 4D&E). In agreement with the luciferase experiments, ALTO downregulated LT and ST mRNAs and proteins compared to RFP control cells. Mutating ALTO NTAR1&2 caused a partial rescue of LT and ST mRNA and protein, consistent with the luciferase assay results.

We also assessed the effect of ALTO ΔNTAR1+2 on MKL-2 cell growth (Fig 4F). While ALTO ΔNTAR1+2 failed to upregulate CDKN1A, consistent with it being a RelA target gene, ALTO ΔNTAR1+2 showed no deficit in its ability to induce growth arrest. ALTO ΔNTAR1&2 still mildly activated non-canonical NF-κB signaling and still significantly downregulated LT and ST, suggesting that growth arrest is strongly dependent on LT and ST expression and that LT and ST expression is highly dependent on RelB which bind directly to the MCPyV NCCR.

### ALTO/NF-κB/early transcription signaling network is conserved in ALTO-encoding PyVs

As far as we are aware, NF-κB activation and SQSTM1 and TRAF2/3 binding have not been reported for MuPyV MT. We analyzed the MT coding sequence for NTAR1-like (**P**x**Q**x**E**/**T**) and NTAR2-like (**P**x**Q**xx**Z**) motifs but did not identify any such sequences (Fig. 5A). As predicted based on the lack of NTAR motifs, MT-HA failed to activate NF-κB signaling (Fig. 5B) and did not activate the NF-κB luciferase reporter in 293A cells (Fig S6A). However, MT was expressed at lower levels than both LMP1 and ALTO. We also generated a MuPyV ALTO construct from MuPyV MT’s second exon (amino acids 208-421). MuPyV HA-ALTO expressed at a higher level than MCPyV HA-ALTO (Fig. 5B) but still failed to activate NF-κB signaling or activate the NF-κB luciferase reporter (Fig S6A), indicating that the presence of NTAR motifs is critical for NF-κB activation by an ALTO or MT protein.

**Figure 5.**
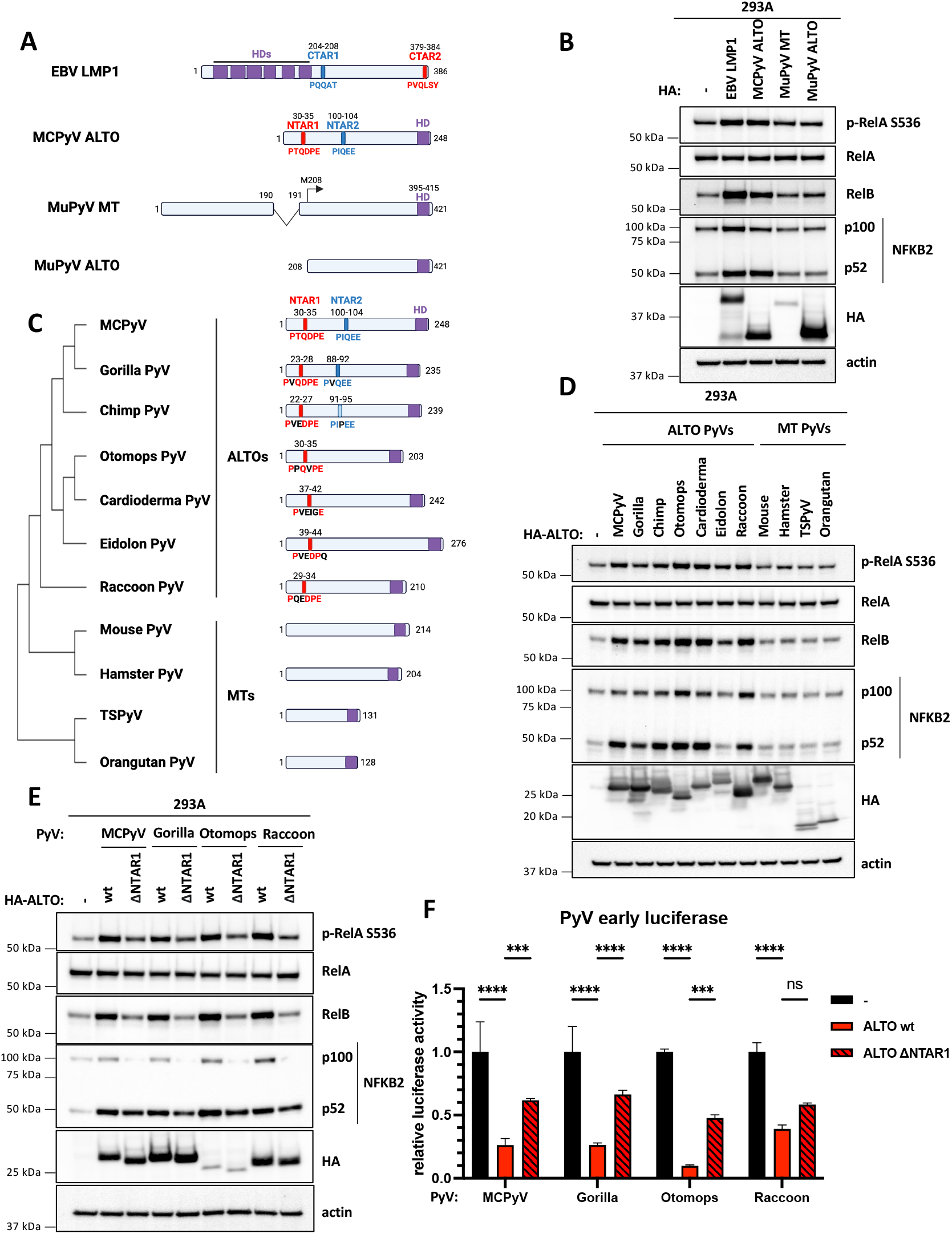
ALTO/NF-κB/early gene signaling network is conserved in ALTO-encoding polyomaviruses. **A**) EBV LMP1, MCPyV ALTO, MuPyV MT and MuPyV ALTO constructs transfected in 293A cells in B). NTAR1/CTAR2 and NTAR2/CTAR1 motifs are highlighted in red and blue, respectively. C terminal hydrophobic domains are highlighted in purple. **B**) Western blots of 293A cells transfected with plasmids encoding EBV HA-LMP1, MCPyV HA-ALTO, MuPyV MT-HA or MuPyV HA-ALTO. **C**) Cartoon schematic comparing ALTOs and MTs from almipolyomaviruses. Amino acid sequences were aligned using Clustal Omega to create a phylogenetic tree. NTAR1 and NTAR2 motifs are highlighted in red and blue, respectively. C terminal hydrophobic domains are highlighted in purple. **D**) Western blots of 293A cells transfected with plasmids encoding PyV HA-ALTOs. **E**) Western blots of 293A cells transfected with plasmids encoding PyV HA-ALTO wt and ΔNTAR1 mutants. **F**) PyV early luciferase assays in 293A cells transfected with HA-ALTO wt or ΔNTAR1 plasmids. Luciferase activity was normalized to CMV-driven renilla activity. Statistical significance compared to empty vector control was determined by 2-way ANOVA. *** P<0.0002, **** P<0.0001.

Since MuPyV MT failed to activate NF-κB signaling, we examined whether the presence of NTAR1&2 motifs and NF-κB activation might be another distinguishing characteristic of ALTOs and MTs, besides their gene structure. We searched the coding sequences of MTs and ALTOs from a range of mammalian polyomaviruses for NTAR1- and NTAR2-like sequences (Fig 5C). NTAR1-like and NTAR2-like sequences were identified in ALTOs from gorilla and chimp polyomaviruses that have high sequence identity to MCPyV ALTO while only NTAR1-like sequences were found in ALTOs from more sequence-divergent bat polyomaviruses (otomops, cardioderma and eidolon) and raccoon polyomavirus. No NTAR1-or NTAR2-like sequences were found in any of the four MT-encoding polyomaviruses (mouse, hamster, human TSPyV and orangutan). All six ALTOs containing NTAR1-or NTAR2-like motifs induced RelA S536 phosphorylation, RelB, and NFKB2 and activated the NF-κB luciferase reporter (Fig 5D, S6B). In contrast, ALTOs derived from hamster, TSPyV and orangutan MTs failed to activate NF-κB signaling, consistent with the lack of NTAR motifs. Next, we mutated the NTAR1 motifs to alanine in a subset of the polyomavirus ALTOs (gorilla, otomops and raccoon ALTOs). As expected, we observed a dramatic reduction in RelA S536 phosphorylation, RelB, and NFKB2 and NF-κB luciferase activity (Fig 5E & S6C). Comparison of the NTAR1 motifs from these four ALTOs suggests that the minimal NTAR1 motif is **P**x(**E**/**Q**)xx**E**.

Since these other PyV ALTOs appear to activate NF-κB signaling in a similar manner to MCPyV ALTO, we predicted that they may also regulate their early gene expression in an NF-κB dependent manner.. To test this hypothesis, we generated early luciferase reporter plasmids derived from the NCCRs of gorilla, otomops, and raccoon polyomaviruses. Consistent with MCPyV, each ALTO downregulated its respective early luciferase reporter (Fig. 5F). Furthermore, each ALTO ΔNTAR1 mutant was partially deficient in its ability to downregulate luciferase activity, consistent with NF-κB playing an important role in control of early gene expression in ALTO-encoding polyomaviruses.

## Discussion

While the existence of polyomavirus ALTOs has been known for over a decade, their function has remained elusive until now. As the MCPyV host cell and MCC cell of origin has yet to be conclusively determined, we made use of MCC cell lines to study the function of ALTO in tumorigenesis and to gain insights into its role in the viral lifecycle. In contrast to MuPyV MT’s well-known role in oncogenesis, ALTO is not expressed in MCC cell lines and rescuing ALTO expression suppressed cell growth. Recruitment of SFKs, tyrosine phosphorylation, and activation of Ras/MAPK and AKT/mTOR signaling via SHC1 and PI3K p85 binding are critical for MT’s transforming activity (Fig 6A). While it has been reported that MCPyV ALTO binds SFKs, is tyrosine phosphorylated and binds and activates PLCγ1 in a similar manner to MuPyV MT, we saw no evidence of binding to SHC1 or PI3K p85 in our mass spectrometry data. ALTO contains one other tyrosine residue (Y79) that could potentially be tyrosine phosphorylated. However, the amino acids surrounding Y79 (_75_EDPI**<u>Y</u>**LPNTM_84_), which dictate binding partner specificity, do not match the consensus SHC1 and p85 binding motifs found in MT (_247_NPT**Y**_250, 315_**Y**MPM_318_). The other polyomavirus ALTOs which encode NTAR1&2 motifs, like MCPyV ALTO, contain binding sites for PLCγ1 (**Y**L[D/E][I/V]) but not SHC1 or p85, suggesting that SFK and PLCγ1 binding evolved early before the divergence of MT and ALTO, with SHC1/p85 binding motifs and NTAR1/2 motifs arising after.

**Fig 6.**
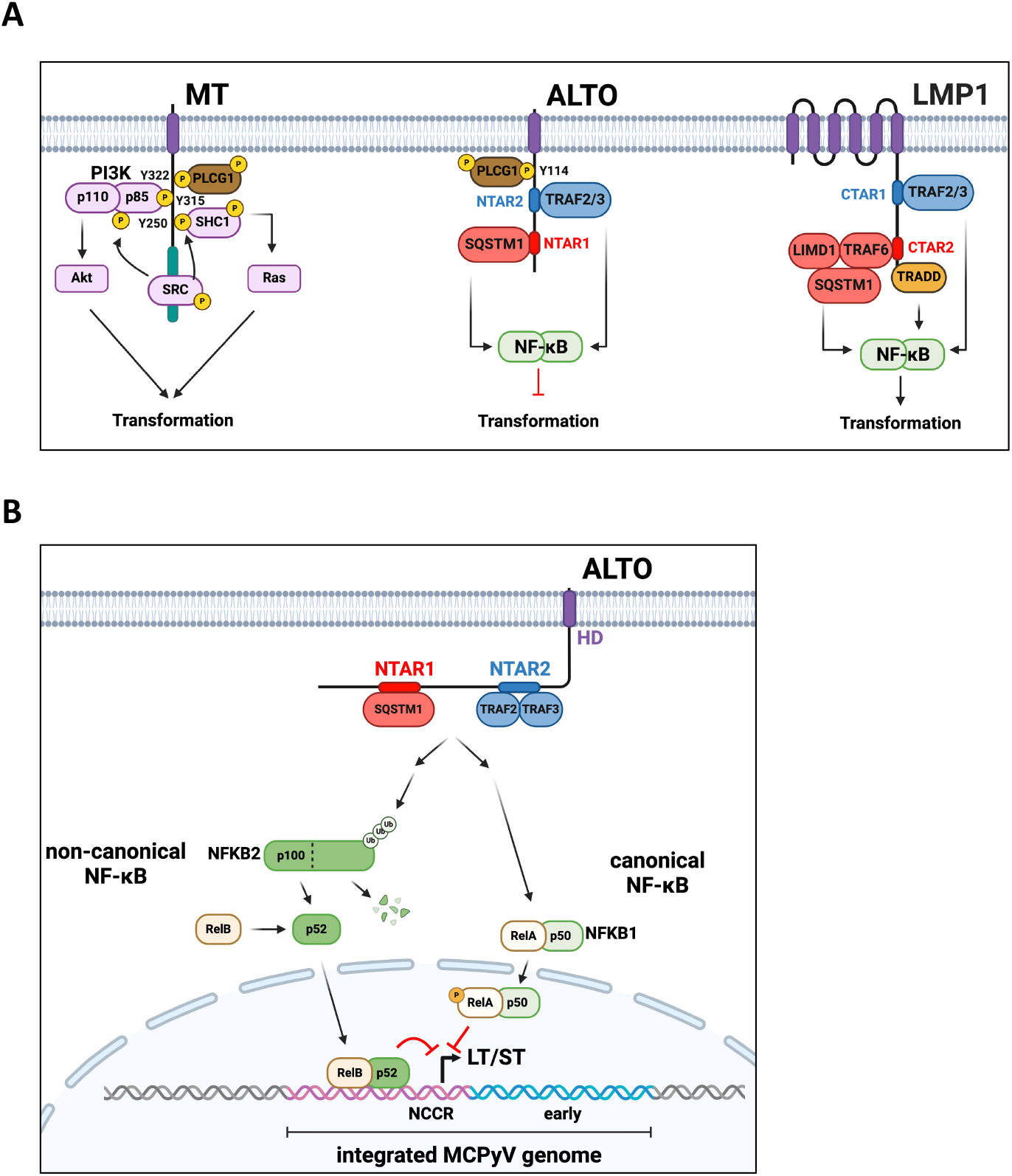
MCPyV ALTO activates the NF-κB pathway and suppresses tumorigenesis by downregulating LT and ST. **A)** MuPyV MT, MCPyV ALTO, and EBV LMP1 binding partners, activated pathways, and their effect on tumorigenesis. **B)** ALTO activates NF-κB signaling by binding SQSTM1 and TRAF2&3 via two N-Terminal Activating Regions (NTAR1&2). Canonical NF-κB signaling via RelA appears to indirectly downregulate MCPyV early transcription. Meanwhile, non-canonical NF-κB signaling directly inhibits MCPyV early transcription with RelB binding to MCPyV NCCR.

We discovered that MCPyV ALTO activates the NF-κB pathway in a similar manner to EBV LMP1 (Fig 6A), highlighting convergent evolution of two proteins from disparate DNA viruses. Both LMP1 and ALTO contain two similar NF-κB activating motifs that recruit similar complexes and resemble constitutively active TNFR family members. While both ALTO and LMP1 bind TRAF2&3, ALTO does not appear to associate with TRAF6, despite the presence of a putative TRAF6 binding motif (_30_**P**T**Q**DP**E**_35_). Despite their similarities, LMP1 and ALTO’s contributions to tumorigenesis are exactly opposite. Like MuPyV MT but not MCPyV ALTO, LMP1 has also been shown to activate both Ras/MAPK and PI3K/Akt signaling(36, 37). Furthermore, the host cells for EBV and MCPyV are distinctly different. EBV primarily infects B cells and causes lymphomas, although it is also associated with nasopharyngeal and gastric cancers(38), whereas MCPyV causes a skin cancer. In B cells, NF-κB drives cellular proliferation and is activated following B cell receptor stimulation(39). In contrast, NF-κB activation in epithelial keratinocytes causes growth arrest and induction of CDKN1A(32). In mesenchymal fibroblasts, NF-κB activation has no effect on cell growth or CDKN1A induction (32). The differing outcomes on cell proliferation following NF-κB activation can likely be attributed to differences in expression of other transcriptional coactivators or corepressors that bind to RelA’s transcriptional activation domain in cells of differing lineages. In our RNAseq data from MKL-2 cells, we observed ALTO induces CDKN1A expression (Fig 2D).

MCC cells, therefore, behave similarly to keratinocytes with respect to their response to NF-κB activation, suggesting that MCC may have arisen from a cell with epithelial characteristics.

While the MCPyV host cell remain elusive, our data from MCC cells also provide insights into ALTO’s role in the normal viral lifecycle. We showed that MCPyV ALTO downregulates LT and ST in an NF-κB dependent manner (Fig 6B). Canonical NF-kB activation via RelA overexpression in MKL-2 cells downregulated LT and ST transcripts and proteins, although not to the same level as ALTO (Fig 4E), which also activates non-canonical NF-κB signaling. We showed that RelB binds strongly to MCPyV NCCR, likely in a complex with either NFKB1 p50 or NFKB2 p52, although the exact DNA sequence bound remains to be identified. During the normal EBV lifecycle, LMP1 promotes and maintains EBV latency via NF-κB activation, which represses EBV’s lytic master regulator BZLF1. While we previously did not see any difference in viral replication in 293 cells transfected with ALTO KO or wt MCPyV genome(29), Peng et al. showed that ALTO Y114F, deficient for NF-κB activation in their system, displayed enhanced viral replication(31), supporting the hypothesis that ALTO acts to suppress viral replication during the normal viral lifecycle. Therefore, it appears that both EBV and MCPyV have evolved membrane proteins that can activate NF-κB to repress viral replication. Similarly, both viruses have proteins (BZLF1 and ST) that promote viral replication and repress NF-κB(16, 22), indicating that both the EBV and MCPyV life cycles are intimately linked to NF-κB signaling.

## Materials and Methods

### Cell lines

MKL-1 and MKL-2 were gifts from Drs. Yuan Chan and Patrick Moore, MS-1 from Dr. Masahiro Shuda (University of Pittsburgh, PA, USA) and WaGa from Dr. Jurgen Becker (University of Duisberg-Essen, Germany). MCC cell lines were maintained in RPMI 1640 HEPES media (Gibco) supplemented with 10% tetracycline negative fetal bovine serum (GeminiBio) and 50 units/mL penicillin and 50 ug/mL streptomycin (Gibco). 293A and 293T cells were purchased from Invitrogen and maintained in DMEM (Gibco) supplemented with 10% fetal bovine serum (Corning), 1x glutaMAX (Gibco), 1x MEM non-essential amino acids (Gibco) and 50 units/mL penicillin and 50 ug/mL streptomycin (Gibco). All cell lines were cultured at 37°C and 5% CO_2_.

### Subcloning and plasmids

PyV and EBV sequences used for ALTO, MT and LMP1 cloning are detailed in Table S1&2. MCPyV ALTO variants were codon optimized to remove the LT and miR-M1 reading frame and synthesized as dsDNA fragments (IDT) and cloned into pcDNA3 (Invitrogen) or pTRIPZ (gift from Sandra Demaria – Addgene #127696) vectors using In-Fusion cloning (Takara). HA-RelA was codon-optimized and synthesized as dsDNA fragment (IDT) and cloned into pcDNDA using In-Fusion cloning.

### Transfection and lentiviral transduction to induce ALTO expression

For expression analysis and luciferase assays in 293A cells, cells were transfected with TransIT-293 (Mirus Bio) following the manufacturer’s instructions. To generate lentiviral particles, 293T cells were transfected with psPAX2 and pMD2.G (gifts from Didier Trono – Addgene #12260, #12259) and pTRIPZ lentiviral vectors. Supernatants containing lentiviral particles were harvested 48 hours post transfection, 0.45 uM filtered, and diluted 1:2 with fresh DMEM and then supplemented with 6 ug/mL polybrene. Target cells were incubated with lentiviral supernatants for 24 h followed by 24 h in RPMI 1640 media. Transduced cells were selected with 0.5 ug/mL puromycin for 4 days prior to expansion in puromycin-free media.

### Cell growth assays

In brief, MCC cells were seeded at 0.1x10^6^ or 0.25x10^6^ cells per mL in media supplemented with 0.5 μg/mL doxycycline. Media was refreshed every 3 days. Prior to cell counting with 0.2% trypan blue, cell clumps were digested with accutase (Stem Cell Technologies) for 10 mins at room temperature to produce single cell suspensions.

### RNA extraction and qRT-PCR

Total RNA was extracted from cells using RNeasy Plus Mini kit (Qiagen). RNA concentration and purity was measured by Nanodrop. cDNA was synthesized from 1 μg total RNA using the SuperScript VILO cDNA Synthesis Kit (Thermo Fisher Scientific) per manufacturer’s instructions. qRT-PCR reactions were performed using PowerUp SYBR Green Master Mix (ThermoFisher Scientific) and StepOnePlus Real-Time PCR system (Applied Biosystems). Primer sequences are provided in Table S3. Differential LT and ST expression was determined using 2^-ΔΔCt^ method and normalized to housekeeping gene 36B4 expression.

### RNA sequencing and Gene Set Enrichment Analysis

Total RNA was extracted from cells using RNeasy Plus Mini kit (Qiagen). Prior to RNA-seq library prep, RNA integrity was verified by TapeStation analysis (Agilent) and quantified using a Trinean DropSense96 spectrophotometer (Caliper Life Sciences). RNA-seq libraries were prepared using the TruSeq stranded mRNA kit (Illumina) and 500 ng total RNA as templates. Library size distribution was validated by TapeStation analysis. Additional library QC, blending of pooled indexed libraries, and cluster optimization was performed using Qubit 2.0 Fluorometer (Invitrogen). RNA-seq libraries were pooled (30-plex) and clustered onto an SP flow cell. Sequencing was performed using an Illumina NovaSeq 6000 employing a paired-end, 50 base read length (PE50) sequencing strategy. Image analysis and base calling was performed using Illumina’s Real Time Analysis v3.4.4 software, followed by demultiplexing of indexed reads and generation of FASTQ files using Illumina’s bcl2fastq Conversion Software v2.20. Alignment was performed against GRCm38 reference using STAR v-2.7.7a(40) in the two-pass alignment mode. Gene abundance quantification was performed using STAR2’s --quantMode and Gencode v38 gene definitions. Differential expression comparisons between relevant sample groups were performed using Bioconductor’s edgeR(41). A significance threshold of log2FC >= 1 or log2FC <= -1 at 5% FDR was used to define the genes of interest in each comparison. Gene set enrichment analysis was performed against MSigDB collections C2 (curated), C3 (regulatory target) and C6 (oncogenic signature) to identify pathways and gene-sets enriched in the data(42, 43).

### Luciferase assays

NF-κB luciferase vector pGL4.32 and CMV renilla pGL4.75 plasmids were obtained from Promega. To generate PyV early luciferase vectors, NCCR sequences spanning between the ATG start codons for VP2 and LT/ST were synthesized as dsDNA (IDT) and cloned into pGL4.11 (Promega). NCCR sequences are detailed in Table S4. 2x10^4^ 293A cells were seeded in each well of 96 well plate and transfected with 100 ng pGL4 NCCR-luc plasmid, 100 ng CMV renilla plasmid and 100 ng pcDNA3 plasmid. Cells were lysed and luciferase/renilla activity was analyzed using Dual-Glo luciferase assay system (Promega) following manufacturer’s instructions. Luciferase and renilla activity were measured using a Veritas microplate luminometer (Turner BioSystems).

### Electromobility shift assays

HA-RelA and FLAG-NFKB1 p50 sequences were cloned into pCS2 vector downstream of SP6 promoter. RelA and p50 were *in vitro* transcribed and translated using TnT SP6 coupled wheat germ extract system (Promega) following manufacturer’s instructions for the non-radioactive reaction. Briefly, 0.5 ug pCS2 HA-RelA and 0.5 ug pCS2 FLAG-p50 were added to wheat germ extract supplemented with amino acid mixtures lacking leucine and methionine. Reactions were incubated at 30°C for 2 h. Binding reactions were performed using the Odyssey infrared EMSA kit (LI-COR), 10 uL RelA/p50 or empty vector wheat-germ extract and 10 ng IRDye700 labelled probes and incubated at room temperature for 30 mins. Products of the binding reaction were separated on 6% native polyacrylamide gels using 0.5x TBE (Invitrogen). BKPyV, JCPyV and MCPyV probes (Table S5) were synthesized and 5’-IRDye700 labeled by IDT.

### Cell lysis, Western blotting, and immunoprecipitations

Cells were lysed by sonication in NP-40 buffer (50 mM Tris HCl pH 8.0, 150 mM NaCl, 1% NP-40) supplemented with fresh protease and phosphatase inhibitor cocktail (Pierce). For immunoprecipitations, typically 200 ug cell lysates were used as input and incubated with 50 uL anti-HA magnetic beads (Pierce) overnight at 4°C. Beads were then washed five times with NP-40 buffer. Bound proteins were eluted in 100 uL of 1x Bolt LDS sample buffer (Novex) supplemented with 100 mM DTT and incubated at 70°C for 10 mins under rapid agitation. Typically, 20 ug of cell lysate or 20% of IP eluate were separated on 8-16% denaturing tris-glycine polyacrylamide gels and then transferred to PVDF membranes. Blots were blocked in 3% BSA in TBS containing 0.1% Tween-20 for 30 min and then incubated with primary antibodies overnight at 4°C, followed by HRP-conjugated secondary antibodies for 1 h at room temperature. Blots were developed using Clarity Max Western ECL chemiluminescent substrate (Bio-Rad) and imaged on ChemiDoc (Bio-Rad).

The following antibodies were used: ALTO polyclonal rabbit antisera (as previously described(29)); HA (16B12, BioLegend); actin-HRP [#5125], HA [#3724], NFKB2 [#3017], PLCγ1 [#2822], p-PLCγ1 Y783 [#2821] RelA [#8242], p-RelA S536 [#3033], RelB [#10544], SQSTM1 [#39749], streptavidin-HRP [#3999], TBK1 [#3054], TRAF2 [#4724], TRAF3 [#61095], TRAF6 [#8028], V5 [#13202] (Cell Signaling Technologies). LIMD1 (H-4, sc-271448).

### Proximity labeling mass spectrometry

MKL-2 TurboID (TID) and TID-ALTO cells were cultured in biotin-free RPMI 1640 media (USBiological, R9002-01) supplemented with 10% tetracycline-free fetal bovine serum, pen/strep and 5 ug/mL doxycycline for 3 days prior to addition of 0.5 mM biotin for 24 hours. Cells were then pelleted and washed three times with ice-cold PBS to terminate the biotinylation reaction. Cells were lysed in RIPA buffer (25 mM Tris-HCl pH 7.5, 150 mM NaCl, 0.1% SDS, 1% NP-40, 1% sodium deoxycholate) supplemented with protease inhibitor cocktail (Pierce) by sonication. BCA assays were performed to determine protein concentration of samples. 3 mg cell lysate was incubated with 300 uL streptavidin magnetic beads (Pierce) overnight at 4°C. Beads were then washed twice with RIPA buffer, once with 1 M KCl, once with 0.1 M Na_2_CO_3_ and then once with 2 M urea, 10 mM Tris-HCl pH 7.5. 10% of beads were removed and bound proteins eluted with 1xLDS+DTT by heating at 70°C for 10 mins. The remaining beads were resuspended in 100 mM HEPES pH 8.5. Proteins were reduced using TCEP and alkylated with chloroacetamide. Proteins were then digested with Lys-C and trypsin. Peptides were analysed by liquid chromatography coupled to mass spectrometry (LC-MS) using a Easy1200 nLC (Thermo Scientific) coupled to a tribrid Orbitrap Eclipse with FAIMSpro mass spectrometer (Thermo Scientific).

Data analysis was performed using Proteome Discoverer 2.5 (Thermo Scientific, San Jose, CA). The data were searched against a protein database combined from human (Swiss Prot UP000005640), Merkel cell polyomavirus (Swiss Prot UP000152665), and common contaminants (Global Proteome Machine cRAP) databases as well as a custom database containing LT, small T antigen, TID, and TID-ALTO amino acid sequences. Searches were performed with settings for the proteolytic enzyme trypsin. Maximum missed cleavages were set to 2. The precursor ion tolerance was set to 10 ppm and the fragment ion tolerance was set to 0.6 Da. Dynamic peptide modifications included oxidation (+15.995 Da on M). Dynamic modifications on the protein terminus included acetyl (+42.-11 Da on N-terminus), Met-loss (-131.040 Da on M) and Met-loss+Acetyl (-89.030 Da on M) and static modification of carbamidomethyl (+57.021 on C). All search results were run through Percolator for peptide validation and false discovery rate estimation. Label free quantification was carried using the Precursor Ions Quantifier node in Proteome Discoverer. Raw data were normalized by conditions (TID or TID-ALTO). Normalized intensity was log2 transformed and missing values were imputed by 50% of the global minimum intensity. Uncertain proteins, such as protein identified with only single peptide or that were assigned with median or low confidence by Proteome Discoverer, were eliminated prior to statistical tests. P-values for pairwise comparisons on normalized intensity were calculated by t-test.

### DNA pulldowns

MCPyV NCCR and AmpR fragments were PCR amplified using 5’ biotinylated primers. Sequences are provided in Table S3. PCR products were analyzed by gel electrophoresis to confirm molecular weight and purity of PCR products. PCR products were further analyzed by Sanger sequencing. For each primer set, only a single 500 bp product was produced per 50 uL PCR reaction. Accordingly, 5 PCRs were pooled and purified on one Qiaquick spin column (Qiagen) following manufacturer’s instructions.

MKL-2 HA-ALTO cells were treated with 0.5 μg/mL doxycycline for 6 days and then lysed by sonication in 20 mM Tris HCl pH 7.5, 50 mM KCl, 0.25% Tween 20, 0.05% NP-40 buffer supplemented with protease and phosphatase inhibitors (Pierce). Cell lysates were clarified by centrifugation at 20,000*g* for 15 mins at 4°C. 500 μg cell lysate was diluted to 500 μL in lysis buffer supplemented with 1 mM DTT and 25 μg poly(dI:dC) (Roche) and precleared by incubating with 50 μL streptavidin beads for 1 hour at 4°C on rotating wheel. Meanwhile, 5 μg biotinylated PCR product (MCPyV NCCR or AmpR) was diluted to 500 μL with lysis buffer and incubated with 50 μL streptavidin beads at room temperature on rotating wheel to bind DNA to the beads. After 1 hour, NCCR/AmpR-beads were washed once to removed unbound dsDNA and incubated with precleared cell lysates overnight at 4°C. Next morning, beads were washed five times with 500 μL lysis buffer. Bound proteins were eluted by incubating beads in 1x LDS supplemented with 25 mM DTT at 70°C for 10 mins. Eluted proteins were then analyzed by Western blotting.

## Acknowledgments

This research was funded by National Institutes of Health/National Cancer Institute (R35 CA209979) to DAG and a Brave Fellowship awarded by the Brave Like Gabe Foundation to NJHS. This research was supported by the Genomics & Bioinformatics Shared Resource, RRID:SCR_022606, of the Fred Hutch/University of Washington/Seattle Children’s Cancer Consortium (P30 CA015704). Assistance with Turbo-ID experiments was provided by Lisa Jones, Chenwei Lin, and Phil Gafken and the Fred Hutch Proteomics & Metabolomics Resource, which is funded in part through NIH/NCI Cancer Center Support Grant P30 CA015704. We would like to acknowledge Dr. Richard Yang (University of Texas Southwestern) for helpful discussions at the DNA Tumor Virus Meeting 2023 in Montreal, Canada, and Dr. Frank Szuzlewsky (Fred Hutch Cancer Center) for help with luciferase assays.

## References

1. D. P. Lane, L. V. Crawford, T antigen is bound to a host protein in SV40-transformed cells. Nature 278, 261–263 (1979).

2. J. A. DeCaprio et al., SV40 large tumor antigen forms a specific complex with the product of the retinoblastoma susceptibility gene. Cell 54, 275–283 (1988).

3. S. A. Courtneidge, A. E. Smith, Polyoma virus transforming protein associates with the product of the c-src cellular gene. Nature 303, 435–439 (1983).

4. M. Whitman, D. R. Kaplan, B. Schaffhausen, L. Cantley, T. M. Roberts, Association of phosphatidylinositol kinase activity with polyoma middle-T competent for transformation. Nature 315, 239–242 (1985).

5. H. Feng, M. Shuda, Y. Chang, P. S. Moore, Clonal integration of a polyomavirus in human Merkel cell carcinoma. Science 319, 1096–1100 (2008).

6. P. W. Harms et al., The Distinctive Mutational Spectra of Polyomavirus-Negative Merkel Cell Carcinoma. Cancer Res 75, 3720–3727 (2015).

7. S. Q. Wong et al., UV-Associated Mutations Underlie the Etiology of MCV-Negative Merkel Cell Carcinomas. Cancer Res 75, 5228–5234 (2015).

8. H. J. Kwun et al., The minimum replication origin of merkel cell polyomavirus has a unique large T-antigen loading architecture and requires small T-antigen expression for optimal replication. J Virol 83, 12118–12128 (2009).

9. M. Shuda et al., T antigen mutations are a human tumor-specific signature for Merkel cell polyomavirus. Proc Natl Acad Sci U S A 105, 16272–16277 (2008).

10. R. Houben et al., An intact retinoblastoma protein-binding site in Merkel cell polyomavirus large T antigen is required for promoting growth of Merkel cell carcinoma cells. Int J Cancer 130, 847–856 (2012).

11. H. Feng et al., Cellular and viral factors regulating Merkel cell polyomavirus replication. PLoS One 6, e22468 (2011).

12. H. J. Kwun et al., Merkel cell polyomavirus small T antigen controls viral replication and oncoprotein expression by targeting the cellular ubiquitin ligase SCFFbw7. Cell Host Microbe 14, 125–135 (2013).

13. K. Rapchak, S. D. Yagobian, J. Moore, M. Khattri, M. Shuda, Merkel cell polyomavirus small T antigen is a viral transcription activator that is essential for viral genome maintenance. PLoS Pathog 18, e1011039 (2022).

14. J. Cheng et al., Merkel cell polyomavirus recruits MYCL to the EP400 complex to promote oncogenesis. PLoS Pathog 13, e1006668 (2017).

15. D. E. Park et al., Dual inhibition of MDM2 and MDM4 in virus-positive Merkel cell carcinoma enhances the p53 response. Proc Natl Acad Sci U S A 116, 1027–1032 (2019).

16. D. A. Griffiths et al., Merkel cell polyomavirus small T antigen targets the NEMO adaptor protein to disrupt inflammatory signaling. J Virol 87, 13853–13867 (2013).

17. J. Zhao et al., Merkel Cell Polyomavirus Small T Antigen Activates Noncanonical NF-kappaB Signaling to Promote Tumorigenesis. Mol Cancer Res 18, 1623–1637 (2020).

18. H. Buss et al., Constitutive and interleukin-1-inducible phosphorylation of p65 NF-kappaB at serine 536 is mediated by multiple protein kinases including IkappaB kinase (IKK)-alpha, IKKbeta, IKKepsilon, TRAF family member-associated (TANK)-binding kinase 1 (TBK1), and an unknown kinase and couples p65 to TATA-binding protein-associated factor II31-mediated interleukin-8 transcription. J Biol Chem 279, 55633–55643 (2004).

19. L. F. Chen et al., NF-kappaB RelA phosphorylation regulates RelA acetylation. Mol Cell Biol 25, 7966–7975 (2005).

20. T. S. Gorrill, K. Khalili, Cooperative interaction of p65 and C/EBPbeta modulates transcription of BKV early promoter. Virology 335, 1–9 (2005).

21. P. N. Ranganathan, K. Khalili, The transcriptional enhancer element, kappa B, regulates promoter activity of the human neurotropic virus, JCV, in cells derived from the CNS. Nucleic Acids Res 21, 1959–1964 (1993).

22. T. E. Morrison, S. C. Kenney, BZLF1, an Epstein-Barr virus immediate-early protein, induces p65 nuclear translocation while inhibiting p65 transcriptional function. Virology 328, 219–232 (2004).

23. K. M. Kaye, K. M. Izumi, E. Kieff, Epstein-Barr virus latent membrane protein 1 is essential for B-lymphocyte growth transformation. Proc Natl Acad Sci U S A 90, 9150–9154 (1993).

24. S. Prince et al., Latent membrane protein 1 inhibits Epstein-Barr virus lytic cycle induction and progress via different mechanisms. J Virol 77, 5000–5007 (2003).

25. O. Devergne et al., Association of TRAF1, TRAF2, and TRAF3 with an Epstein-Barr virus LMP1 domain important for B-lymphocyte transformation: role in NF-kappaB activation. Mol Cell Biol 16, 7098–7108 (1996).

26. L. Wang et al., The Ubiquitin Sensor and Adaptor Protein p62 Mediates Signal Transduction of a Viral Oncogenic Pathway. mBio 12, e0109721 (2021).

27. K. M. Izumi, E. D. Kieff, The Epstein-Barr virus oncogene product latent membrane protein 1 engages the tumor necrosis factor receptor-associated death domain protein to mediate B lymphocyte growth transformation and activate NF-kappaB. Proc Natl Acad Sci U S A 94, 12592–12597 (1997).

28. F. Giehler et al., Epstein-Barr virus-driven B cell lymphoma mediated by a direct LMP1-TRAF6 complex. Nat Commun 15, 414 (2024).

29. J. J. Carter et al., Identification of an overprinting gene in Merkel cell polyomavirus provides evolutionary insight into the birth of viral genes. Proc Natl Acad Sci U S A 110, 12744–12749 (2013).

30. K. A. Gottlieb, L. P. Villarreal, Natural biology of polyomavirus middle T antigen. Microbiol Mol Biol Rev 65, 288–318 ; second and third pages, table of contents (2001).

31. W. Y. Peng et al., Membrane-bound Merkel cell polyomavirus middle T protein constitutively activates PLCgamma1 signaling through Src-family kinases. Proc Natl Acad Sci U S A 120, e2316467120 (2023).

32. K. Hinata, A. M. Gervin, Y. Jennifer Zhang, P. A. Khavari, Divergent gene regulation and growth effects by NF-kappa B in epithelial and mesenchymal cells of human skin. Oncogene 22, 1955–1964 (2003).

33. R. H. Arch, C. B. Thompson, 4-1BB and Ox40 are members of a tumor necrosis factor (TNF)-nerve growth factor receptor subfamily that bind TNF receptor-associated factors and activate nuclear factor kappaB. Mol Cell Biol 18, 558–565 (1998).

34. T. Abe, G. N. Barber, Cytosolic-DNA-mediated, STING-dependent proinflammatory gene induction necessitates canonical NF-kappaB activation through TBK1. J Virol 88, 5328–5341 (2014).

35. J. F. Yang, W. Liu, J. You, Characterization of molecular mechanisms driving Merkel cell polyomavirus oncogene transcription and tumorigenic potential. PLoS Pathog 19, e1011598 (2023).

36. M. L. Roberts, N. R. Cooper, Activation of a ras-MAPK-dependent pathway by Epstein-Barr virus latent membrane protein 1 is essential for cellular transformation. Virology 240, 93–99 (1998).

37. C. W. Dawson, G. Tramountanis, A. G. Eliopoulos, L. S. Young, Epstein-Barr virus latent membrane protein 1 (LMP1) activates the phosphatidylinositol 3-kinase/Akt pathway to promote cell survival and induce actin filament remodeling. J Biol Chem 278, 3694–3704 (2003).

38. S. Han et al., Epstein-Barr Virus Epithelial Cancers-A Comprehensive Understanding to Drive Novel Therapies. Front Immunol 12, 734293 (2021).

39. Y. Sasaki, K. Iwai, Roles of the NF-kappaB Pathway in B-Lymphocyte Biology. Curr Top Microbiol Immunol 393, 177–209 (2016).

40. A. Dobin et al., STAR: ultrafast universal RNA-seq aligner. Bioinformatics 29, 15–21 (2013).

41. M. D. Robinson, D. J. McCarthy, G. K. Smyth, edgeR: a Bioconductor package for differential expression analysis of digital gene expression data. Bioinformatics 26, 139–140 (2010).

42. A. Subramanian et al., Gene set enrichment analysis: a knowledge-based approach for interpreting genome-wide expression profiles. Proc Natl Acad Sci U S A 102, 15545–15550 (2005).

43. V. K. Mootha et al., PGC-1alpha-responsive genes involved in oxidative phosphorylation are coordinately downregulated in human diabetes. Nat Genet 34, 267–273 (2003).

