## Supplementary Figures 1-6 for "Polyomavirus ALTOs, but not MTs, downregulate viral early gene expression by activating the NF-κB pathway"

**This PDF file includes:**

Figures S1 to S6
Tables S1 to S5
Legends for Datasets S1 to S3

**Other supporting materials for this manuscript include the following:**

Datasets S1 to S3

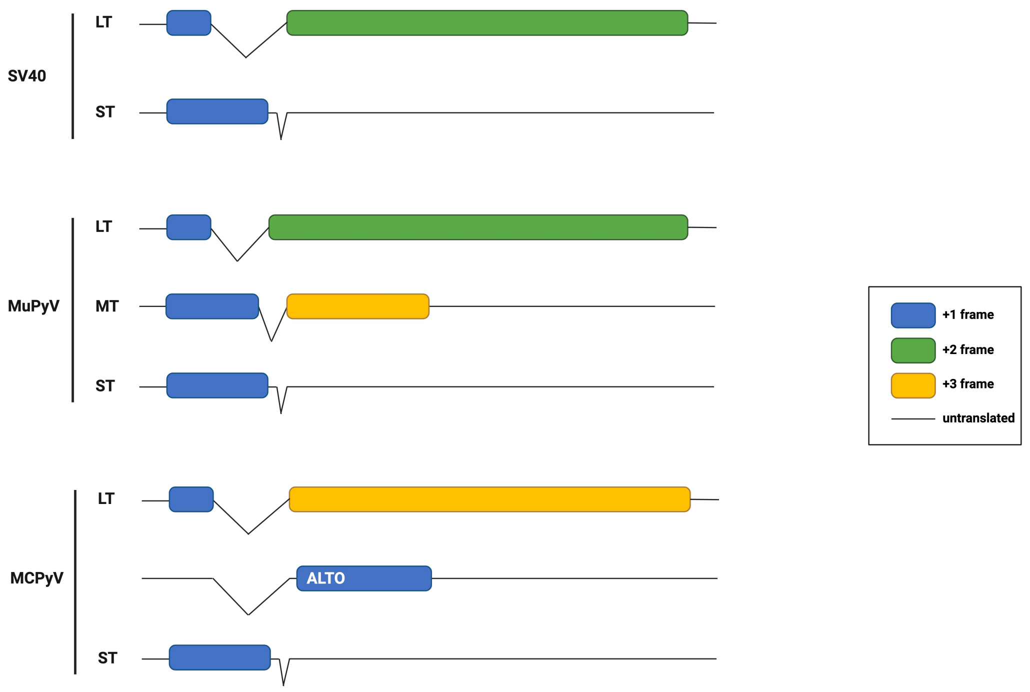


**Figure S1. SV40, MuPyV and MCPyV early transcripts.** Relative reading frames of coding sequences are indicated by blue (+1), green (+2), and yellow (+3). Untranslated sequences are represented by a solid black line.

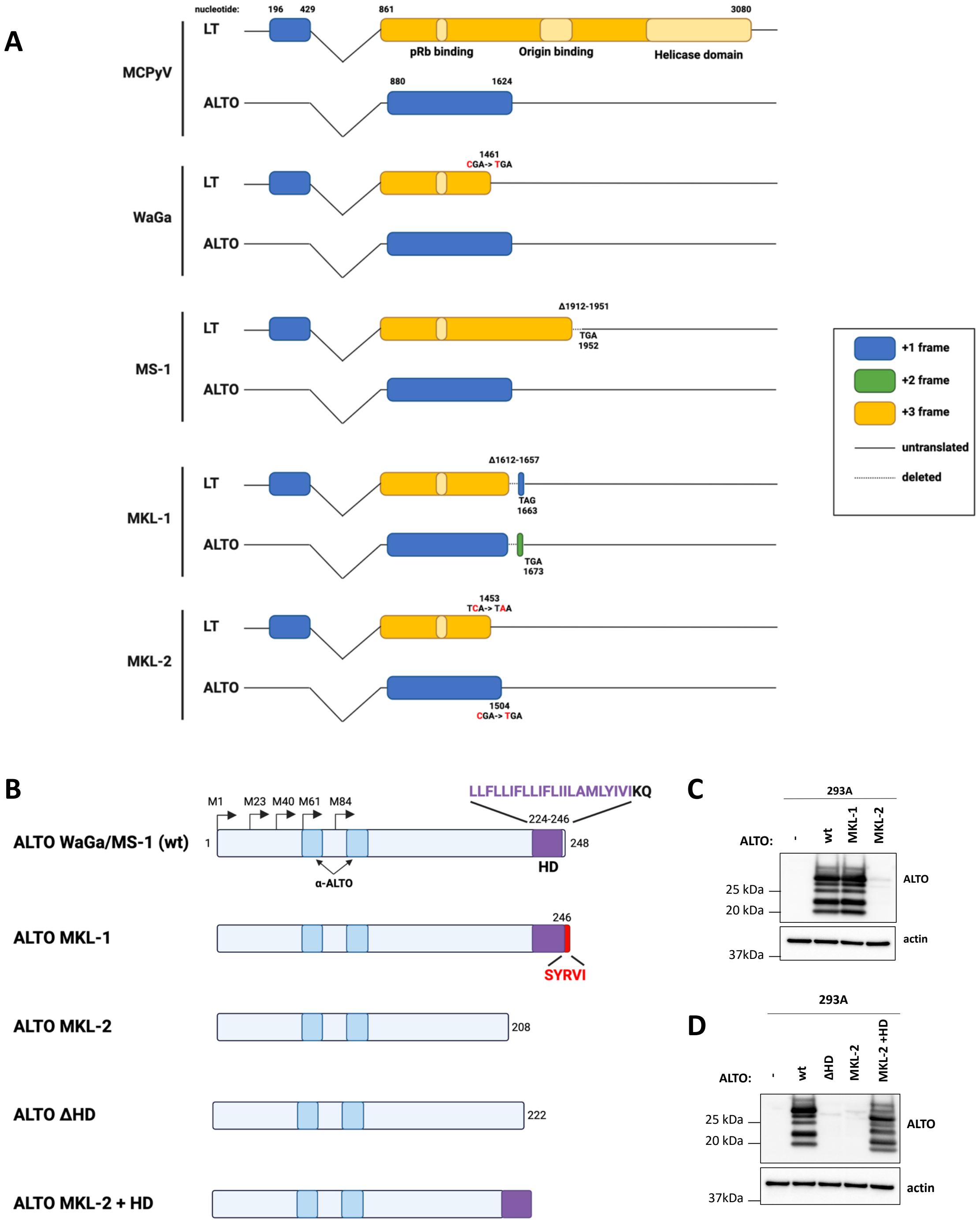


**Figure S2. Effect of LT truncating mutations on ALTO reading frame in MCC cell lines. A**) Point mutations and deletions observed in MCPyV genomes from MCC cells lines relative to MCPyV HF (GenBank #: JF813003) are indicated. The relative reading frames of coding sequences are indicated by blue (+1), green (+2), and yellow (+3). Untranslated sequences are represented by a solid black line. Deleted sequences are indicated by dashed lines. **B**) ALTO constructs transfected in 293A cells in C) and D). Multiple start codons are found within the ALTO reading frame indicated by arrows. ALTO's hydrophobic domain (HD) is highlighted in purple. The peptides used to generate the ALTO antiserum are indicated (α-ALTO). **C&D**) Western blotting analysis of ALTO expression in transfected 293A cells.

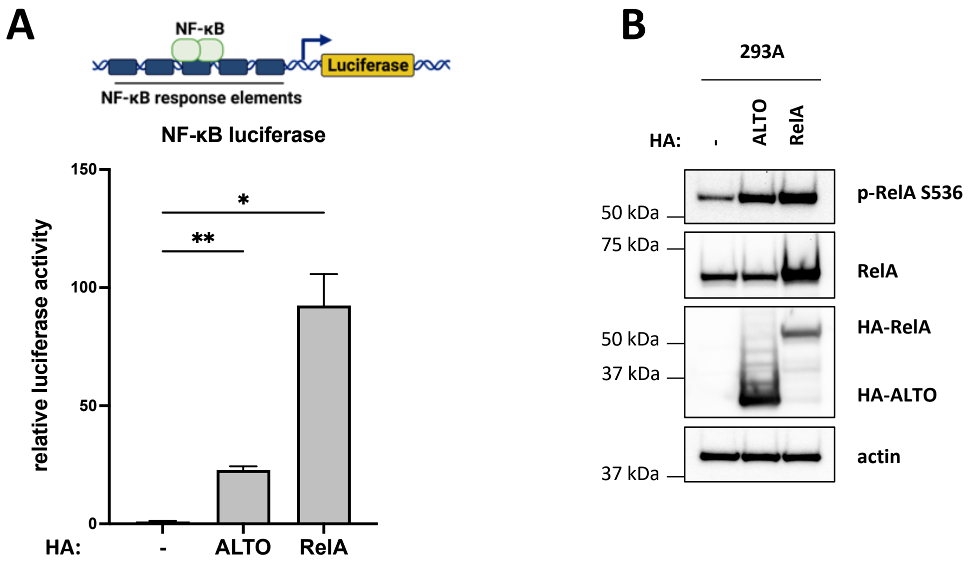


**Figure S3. MCPyV ALTO activates NF-κB signaling in 293A cells. A)** NF-κB luciferase assay in 293A cells transfected with HA-ALTO or HA-RelA plasmids. Luciferase activity was normalized to CMV-driven renilla activity. Statistical significance compared to empty vector control was determined by one-way ANOVA. * $P<0.03$, ** $P<0.002$. **B)** Western blots of 293A cells transfected with HA-ALTO and HA-RelA plasmids. Cells were harvested 2 days post-transfection.

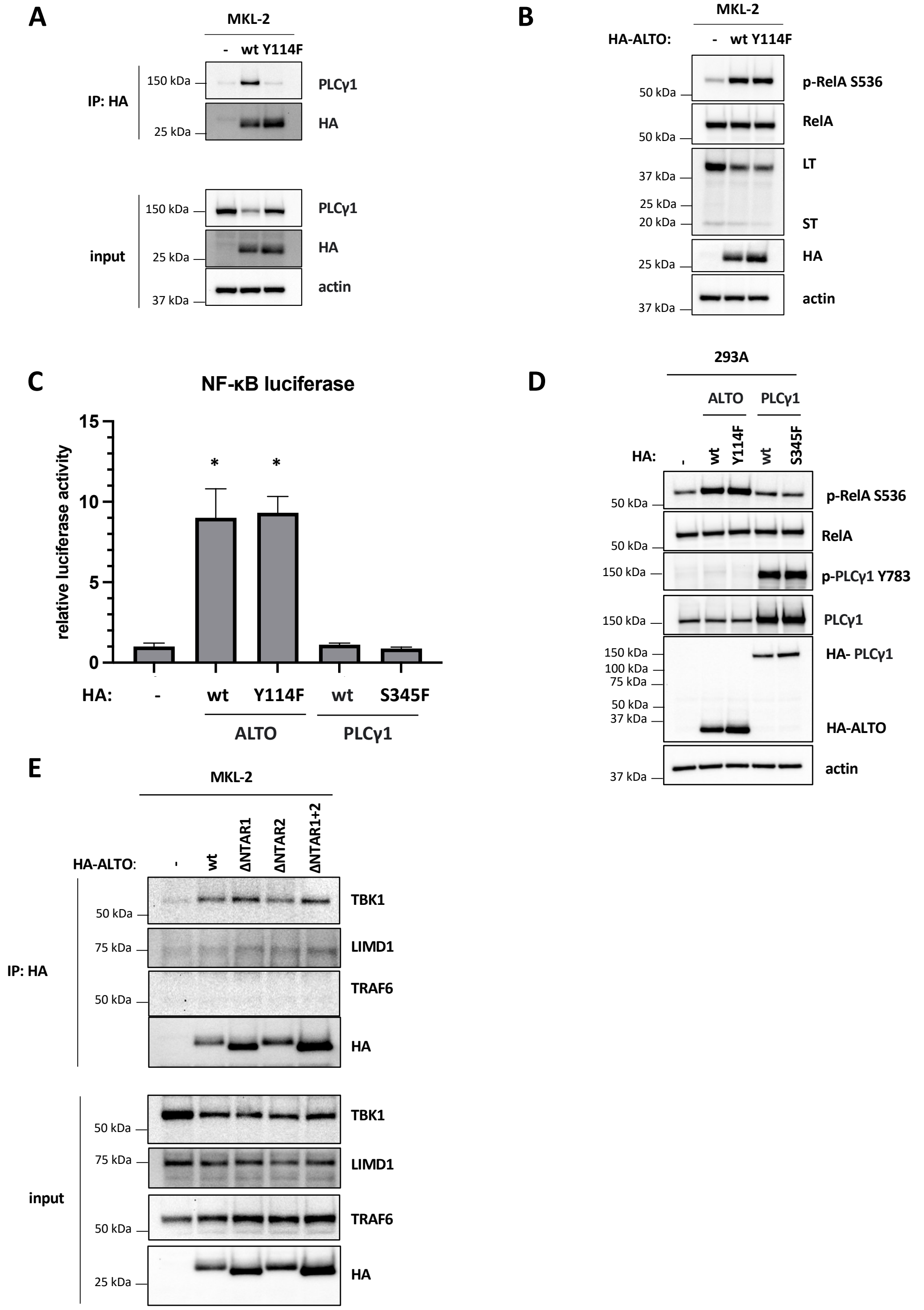


**Figure S4. PLCγ1 and TBK1 bind ALTO but do not contribute to NF-κB activation in MKL-2 cells. LIMD1 and TRAF6 do not appear to bind directly to ALTO. A**) Western blots of co-immunoprecipitations using anti-HA antibody to pull down HA-ALTO wt or Y114F expressed in MKL-2 cells. **B**) Western blots of MKL-2 cells expressing HA-ALTO wt or Y114F. **C**) NF-κB luciferase assays performed in 293A cells transfected with HA-ALTO wt or Y114F or PLCγ1 wt or S345F plasmids. Luciferase activity was normalized to CMV-driven renilla activity. Statistical significance compared to empty vector control was determined by one-way ANOVA (* P<0.03). **D**) Western blots of 293A cells transfected with HA-ALTO wt or Y114F or PLCγ1 wt or S345F plasmids. **E**) Western blots of MKL-2 HA-ALTO variant cell lysates and anti-HA immunoprecipitations. The same cell lysates and IP samples were blotted as in Fig 3D.

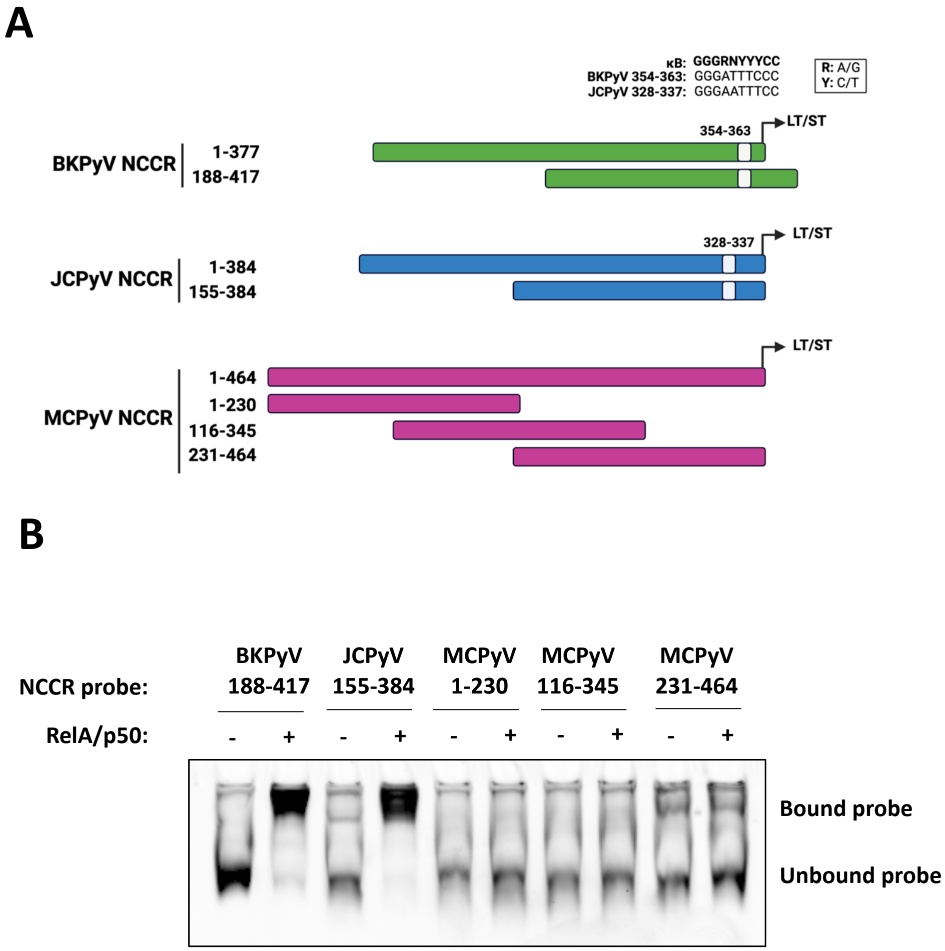


**Figure S5. RelA/p50 dimers do not bind directly to MCPyV NCCR. A**) Cartoons of BKPyV, JCPyV and MCPyV NCCRs with consensus κB sequences highlighted. NCCR probes used for EMSA are shown. **B**) EMSA using BKPyV, JCPyV and MCPyV NCCR probes incubated with RelA/p50 or empty vector in vitro transcribed and translated wheat germ extracts.

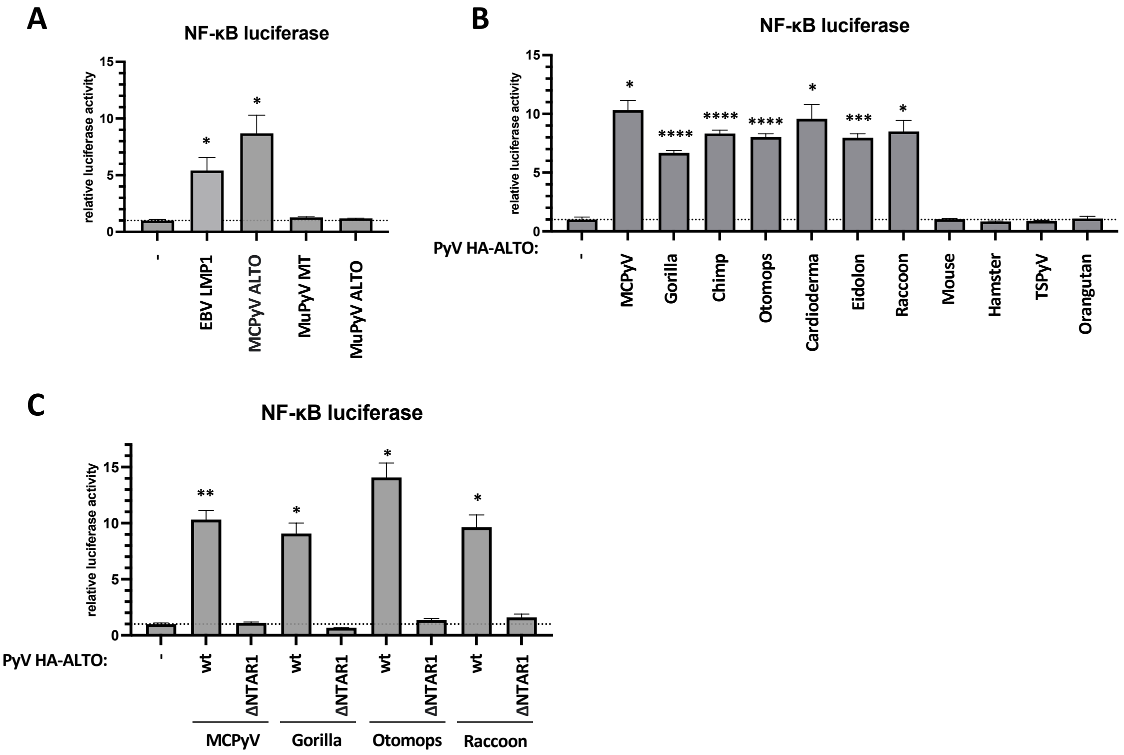


**Figure S6. NF-κB luciferase assays with EBV LMP1 and PyV ALTOs and MuPyV MT. A**) NF-κB luciferase assays performed in 293A cells transfected with plasmids encoding EBV HA-LMP1, MCPyV HA-ALTO, MuPyV MT-HA or MuPyV HA-ALTO. Luciferase activity was normalized to CMV-driven renilla activity. Statistical significance compared to empty vector control was determined by one-way ANOVA. * $P<0.03$. **B**) NF-κB luciferase assays performed in 293A cells transfected with plasmids encoding PyV HA-ALTOs. Luciferase activity was normalized to CMV-driven renilla activity. Statistical significance compared to empty vector control was determined by one-way ANOVA. * $P<0.03$, *** $P<0.0002$, **** $P<0.0001$. **C**) NF-κB luciferase assays performed in 293A cells transfected with plasmids encoding PyV HA-ALTO wt and ΔNTAR1 mutants. Luciferase activity was normalized to CMV-driven renilla activity. Statistical significance compared to empty vector control was determined by one-way ANOVA. * $P<0.03$, ** $P<0.002$.

**Table S1. NCBI accession numbers for viral genomes**

| Virus | NCBI accession number |
|---|---|
| MCPyV | JF813003 |
| BKPyV | AB365141 |
| JCPyV | OP549862 |
| GorillaPyV | HQ385752 |
| ChimpPyV | HQ385746 |
| OtomopsPyV | NC020071 |
| CardiodermaPyV | NC020067 |
| EidolonPyV | NC020068 |
| RaccoonPyV | JQ178241 |
| Mouse (MuPyV) | AF442959 |
| HamsterPyV | JX416853 |
| TSPyV | GU989205 |
| OrangutanPyV | FN356900 |

**Table S2. Viral protein construct sequences.** NTAR1/CTAR2 motifs are highlighted in red. NTAR2/CTAR1 motifs are highlighted in blue. Hydrophobic domains are highlighted in purple.

| Protein | Amino acid sequences |
|---|---|
| MCPyV HA-ALTO | MAYPYDVPDYASLGPLNSKNGGDQEDSASGRHTNMGPIHTG**PTQDPE**<br>SLPPMHPGEPPVEAHHPTARALPLGMGPSQRPRLQTPSPEDPIYLPNT<br>MRNPPHPLDPVAERRP**PIQEE**NPAHPMEPVYLEILPERMAPGRISSAM<br>NHFPPLSLPRPLRSLRSPPPQEARPGSPRLPLPRRPRHLSLQMRNTDP<br>PPSPPRRPLLHSQESENLGGPEALQALLVQQVLQALHQSQKRTEK**LLF**<br>**LLIFLLIFLIILAMLYIVI**KQ |
| EBV HA-LMP1 | MAYPYDVPDYASLEHDLERGPPGPRRPPRGPPLSSS**LGLALLLLLLAL**<br>**LFWLYIVM**SDWTGG**ALLVLYSFALMLIIIILIIFIF**RRDLLCP**LGALCILLLM**<br>**ITLLLIALW**NLHGQA**LFLGIVLFIFGCLLVLGIWIYLLEML**WRLGAT**IWQL**<br>**LAFFLAFFLDLILLIIAL**YLQQNWWT**LLVDLLWLLLFLAILIWM**YYHGQR<br>HSDEHHHDDSLPH**PQQAT**DDSGHESDSNSNEGRHHLLVSGAGDGPP<br>LCSQNLGAPGGGPDNGPQDPDNTDDNGPQDPDNTDDNGPHDPLPQD<br>PDNTDDNGPQDPDNTDDNGPHDPLPHSPSDSAGNDGGPPQLTEEVE<br>NKGGDQGPPLMTDGGGGHSHDSGHGGGDPHLPTLLLGSSGSGGDD<br>DDPHG**PVQLSY**YD |
| MuPyV MT-HA | MDRVLSRADKERLLELLKLPRQLWGDFGRMQQAYKQQSLLLHPDKGG<br>SHALMQELNSLWGTFKTEVYNLRMNLGGTGFQVRRLHADGWNLSTKD<br>TFGDRYYQRFCRMPLTCLVNVKYSSCSCILCLLRKQHRELKDKCDARC<br>LVLGECFCLECYMQWFGTPTRDVLNLYADFIASMPIDWLDLDVHSVYN<br>PKRRSEELRRAATVHYTMTTGHSAMEASTSQGNGMISSESGTPATSR<br>RLRLPSLLSNPTYSVMRSHSYPPTRVLQQIHPHILLEEDEILVLLSPMTA<br>YPRTPPELLYPESDQDQLEPLEEEEEEYMPMEDLYLDILPGEQVPQLIP<br>PPIIPRAGLSPWEGLILRDLQRAHFDPILDASQRMRATHRAALRAHSMQ<br>RHLRRLGRT**LLLVTFLAALLGICLMLFILI**KRSRHFGGGGSAYPYDVPD<br>YASL |
| MuPyV HA-ALTO | MAYPYDVPDYASLTTGHSAMEASTSQGNGMISSESGTPATSRRLRLPS<br>LLSNPTYSVMRSHSYPPTRVLQQIHPHILLEEDEILVLLSPMTAYPRTPP<br>ELLYPESDQDQLEPLEEEEEEYMPMEDLYLDILPEEQVPQLIPPPIIPRA<br>GLSPWEGLILRDLQRAHFDPILDASQRMRATHRAALRAHSMQRHLRRL<br>GRT**LLLVTFLAALLGICLMLFILI**KRSRHF |
| GorillaPyV HA-ALTO | MAYPYDVPDYASLGQQNLKSGGTPEAMALGRRTR**PVQDPE**SPPPMH<br>PGEPPVEPPLQPAARALPPAMGPSQRPRLQTPSPEEPVYLPNTLQRLP<br>MEPRP**PVQEE**NQAHSQEPVYLEVLPEVMAPGMICSVMNPFRRQSPPR<br>PQRNLRSPPSQQGAAPSSPRLPLPKRPRHLSLQMRNVEPPSPPQRPL<br>LPPESENSGGPEVLQALLVQQVLQVLHQSQRRTER**LLFLLIFLLIFLIFL**<br>**AMLYIVI**KQ |

| Protein | Amino acid sequences |
|---|---|
| ChimpPyV HA-ALTO | MAYPYDVPDYASLAQQNLSSGSLPEGMALGRHT**PVEDPE**SPRQTHPGGPPVTEPLLPAARALPLDMGQAHSHRPRLQTPSPEDPIYLPNSLQNPPHPLQPRP**PIPEE**NPAHPAEPVYLEVLPGAVAPGMISTAMSPSPHQSLLPPLRSPRSPPAPEGRRSSPRIPLPRRPPHLNLQMRNAEDLHSPHQRPPLPSPESENSGGTEALPVLLVQQVLAALHQSQKRTEK**LLFLLISLLICLIILAMLFIVI**KQ |
| OtomopsPyV HA-ALTO | MAYPYDVPDYASLMMRIPRFMALQPSERGGEGSNMAIGEQRP**PQVPEE**ESLPIQETPAAMAVVNSPRGPQNFQSHPIPETPPERIYLPHLLPSSREGTYLEIQQDVPLPRPRGMTSPSAVMRPCLHPNPRPPRRSPSLNHQRGPPLPARNPMRTLSPPPEVHPILIEGEGLQDLLHQVILAHHQSQKSLR**ILLILVIFLQIVLIILA**TLSIVIRL |
| CardiodermaPyV HA-ALTO | MAYPYDVPDYASLILKKKTSPHLMAPHNLDIGGLSSSTMGLNLQLMQG**PVEIGE**ESLPLPTQEQGGAPRGPLPDILQRPRLPTPPQTPTEPIYLPQPLMMRRQDPLPRPPGLPFPTLNPPIQEGTEEAIYLEILQEDPSQIGMIPYTATSLCPPSPRPPPRRMRHFRSPSPRPPSPRPRPREPALPLPRRPPSPLMNPSPCHQAHHDEENGTLLQHRLTLALLQNQRKTGK**MLLLMTFLLISLNILVMLYILI**KH |
| EidolonPyV HA-ALTO | MAYPYDVPDYASLKRRMIPLFMGPPNSRPGGTRSTPAGARGAMTLPHLNH**PVEDPQ**DVPDRTVGTRPRSRTPTPPLFHRPRLPTPQMDSPSSGSLPSIPEEPLYLPRDFNPPTRDTIMDLDLRSHQVDGGRRSAPPAMLPRENSGGEEAIYLEIAPEYPHLIGMIHCDAMSPCPRPRSLQVRRTPRRAQSPPRNTPRIPLPKRPPSRRNSRRSTSTLDPPALPSQLQEPLLVEEESSLDPRDFLVLRMQIAIQALRRGQRNQKR**MIFLMIFLLILLTILVML**STAIRR |
| RaccoonPyV HA-ALTO | MAYPYDVPDYASLEPPPLESGGSTSRTYTPGVMMARPPQV**PQEDPE**DGIPLGIMENSPEARGPAGGTGYIPQSQVPLCLPTDPTPQPIYTEIIPRTHSSVKRHLIHHHHHHHHRLLTPTPSPPLSPLLPPRLSDSPVRHRPRPMPPREIPGPPDPIPTLLRLADENTERIIQQVLMLREEVSQAHLQNRRSQG**ILLVLLIFLVIFLTASLALLLVI**RLRMPS |
| HamsterPyV HA-ALTO | MAYPYDVPDYASLLPPPPADPESSTILTQEDTGPTLMDQQDTLTSRRNTGKSFSLSGMLMRTSPAKKNYHHQKTNSPPGIPIPPPPLFLFPVTAPVPPVMRNTQETQAERENEYMPMAPQIHLYSQIREPTHQEEEPQYEEIPIYLELLPENPNQHLALTSTARRSLRRKYHKHNSHIITQHQRNRLR**WLVLMIFLLSLGGFFLTLFFLI**KRKMHL |
| TSPyV HA-ALTO | MAYPYDVPDYASLMFQPRMEEIYLPMGTPPGPAGGKASIKNGTTCLTPCRTQISSAMNPPFPLMNLDLQAPLRDPLLNLARRIQEEEELPHQRTPPAAPRAPSLPPPQSQKNLSMTLS**LMIFLICCGLFFLMLSIVI**KLYHLF |
| OrangutanPyV HA-ALTO | MAYPYDVPDYASLMFQPRTEETYLPMGTPPGPPGGKASIGIGTTSLKTSKTQTFSVMNPPFPLMNLDLQAPLRAPLHNLARRIQEEEEVRTPRLAASPPSQQPHQNQRNLSMTLS**LMIFLMSCGLFFLLLSIVI**RLYHLS |

**Table S3. qRT-PCR and PCR primers.**

| Target | Nucleotide sequence (5’ --> 3’) |
|---|---|
| MCPyV LT | F: GATGGAATTGAACACCCTTTGG<br>R: CTCCTGATCTCCACCATTCTTT |
| MCPyV ST | F: GGATTTCCTCCTACTTGGGAAAG<br>R: TGCAGATGCAGTAAGCAGTAG |
| 36B4 | F: TGCCAGTGTCTGTCTGCAGA<br>R: ACAAAGGCAGATGGATCAGC |
| MCPyV NCCR | F: CCCCATCCTGAAAAATAAATAAG<br>R: GACTAAATCCATCTTGTCTATATGC |
| AmpR | F: TTACCAGTGCTTGATCAGTGAGG<br>R: TAGTGCTGCCATTACCATGAGC |

**Table S4. PyV early luciferase reporter inserts.** The following sequences were cloned into pGL4.11 to generate early luciferase reporter plasmids.

| **PyV** | **NCCR early nucleotide sequence (5' --> 3')** |
|---|---|
| MCPyV | CCTGAAAAATAAATAAGGATACTTACTCTTTTAATGTCCTCCTCCCTTTGTAAGAGAAA<br>AAAAAGCCTCCGGGCCTCCCTTGTTGAAAAAAAGTTAAGAGTCTTCCGTCTCCCTCC<br>CAAACAGAAAGAAAAAAAGTTTTGTTTATCAGTCAAACTCCGCCTCTCCAGGAAATGA<br>GTCAATGCCAGAAACCCTGCAGCAATAAAAGTTCAATCATGTAACCACAACTTGGCTG<br>CCTAGGTGACTTTTTTTTTTTCAAGTTGGCAGAGGCTTGGGGCTCCTAGCCTCCGAGG<br>CCTCTGGAAAAAAAAGAGAGAGGCCTCTGAGGCTTAAGAGGCTTAATTAGCAAAAAA<br>GGCAGTATCTAAGGGCAGATCCCAAGGGCGGGAAACTGCAGTATAAAAACCACTCCT<br>TAGTGAGGTAGCTCATTTGCTCCTCTGCTCTTTCTGCAAACTCCTTCTGCATATAGAC<br>AAG |
| GorillaPyV | CCTGAAAAATAAATAAGGATTACTTACTCAGCCTTGTCCTCCTCCCTTTGTAAGAGAA<br>AAAAAAGGAGTCTTCTCGCTTCCCTCCTCCCTTTTGAAGAAAAAAAAATGCTGCGTCG<br>CTCTCCCCGCTTGTCGCCTCCCTTTGTGTTGAAAAAAAGTTGTGTTAAGAGTCTACTT<br>CCTCCCTCCCAACTAGATTTAAAAAAATTGTTTATTATATAACTCCGCCTCTCCAGGAT<br>ATGAGTCAATGCCAAGAAGCCTGCAGCAATAAAAGTTCAATCAGAGTAAAACCACAA<br>GCTGTCTGCCAGACCACAAGCGTTGCCTAGGCAGCCTATTTTTTTTTACAAATTAGTG<br>CGAGGCTTGGGGCTCCTAGCCTCCGAGGCCTCTGGAAAAAATAGTGAGAGGCCTCT<br>GAGGCCTCTAACAGCTTAATTAGCAGAACCATTCCTGGGCGGGAAACTGCAGTATAA<br>AAGCCACTCCTAAGTGATGTAGCTCATTTTGCTTGCAGAGCTCCACCAAC |
| OtomopsPyV | TCTGAAAAAAAAAAAACATGTACTCACTTTTAATGCCTCCGCCCGTTCAGAAAGAAAA<br>AAATCCACTCGGCGCTGGGGCTCCCGCCCGCTCTGTTTAAAAAAAATGTTTGAAATG<br>GTTGCTGACCTCCTCCCTTCGTGCTTAGAAAAAATCCTACTCATCATGACTAACCCCG<br>CCCGCAGAGACAGAAAAAAACAATTTAAAAGGCTGCAGTAAGGAAATGACTCATTCG<br>TGCCGGCGCCTGAACCAAATGACAGGGGGAGCTCTTTTTTTTTTCAAGTATGCAGAG<br>GCTAGAGGCCCTTAGCCCCTGAGGCTTTCACAGAAAAAGTAGAGAGGCCCTGGGAG<br>GCTTTTTTTAAATTATAGCCGTTAATAGGCGGGAAGGGCTGGTATAAAAGCCTGTTAT<br>TCTCCTCCTCAGGATCTCTAGAGCCTCTTCAGCC |
| RaccoonPyV | CTGTAAAAAAGGAGAGGTTACTTTAAGAAAAAAACTTACAGTGAGAAGTCTTTTCTTCT<br>GAAAAAAAGAGCTTCTGACCTCACTCGCCTCTTGTCCCTGGAAACTGAGAAAAAAAA<br>GTTCTTACTTCCTGGCGAGAGTTGCGGGTGCGGCCGGAAATGTTCTCAACTCTCTCA<br>GAAAATTGCCTGCGGGTGCAGTTTTTTCTCCAAGTGGGCGGAGAAACTCGCCAGTTG<br>CCTAGCAATATCTGAGTCAGAACATGTCAGTACATTCTTGGCTAACCGCAACTTTGGC<br>AGTATTTTTTCTTAAGTATACTGGGGGCCTGAGGCTCCTTGCCTCTTATACTCTTAAG<br>AAAAAGAGGGAGAGGCTGCTGCTGCTACATCTGGTCTGGAGCAACTATAAAAGGCTC<br>GGGCACTATTTTTAACTGCAGATAGGGGACA |

**Table S5. PyV NCCR EMSA probes.** Consensus κB sequences in BKPyV and JCPyV probes are highlighted.

| **PyV** | **Nucleotide sequence (5' --> 3')** |
|---|---|
| BKPyV 188-417 | AGGTCATGGTTTGGCTGCATTCCATGGGAAAGCAGCTCCTCCCTGTGGCCTTTTTTTTTAT<br>AATATATAAGAGGCCGAGGCCGCCTCTGCCTCCACCCTTTCTCTCAAGTAGTAAGGGTGT<br>GGAGGCTTTTTCTGAGGCCTAGCAAAACTATTTG**GGGAAATCCC**TAATCTTTTGCAATTTT<br>TTGCAAAAATGGATAAAGTGCTAAACAGGGAAGAATCCATGGAGCTCA |
| JCPyV 145-374 | GCTGTCAGCTGGTTGGCTCCCTAGGTATGAGCTCATGCTTGGCTGGCAGCCATCCAGTTT<br>TAGCCAGCTCCTCCCTACCTTCCCTTTTTTTTATATATACAGGAGGCCGAGGCCGCCTCCG<br>CCTCCAAGCTTACTCAGAAGTAGTAAGGGCGTGGAGGCTTTTTAGGAGGCCA**GGGAAATT**<br>**CC**CTTGTTTTTCCCTTTTTTGCACTAATTTTTTGCTGCAAAAAGCTAAA |
| MCPyV 1-230 | CCTGAAAAATAAATAAGGATACTTACTCTTTTAATGTCCTCCTCCCTTTGTAAGAGAAAAAA<br>AAGCCTCCGGGCCTCCCTTGTTGAAAAAAAGTTAAGAGTCTTCCGTCTCCCTCCCAAACA<br>GAAAGAAAAAAAGTTTTGTTTATCAGTCAAACTCCGCCTCTCCAGGAAATGAGTCAATGCC<br>AGAAACCCTGCAGCAATAAAAGTTCAATCATGTAACCACAACTTGGC |
| MCPyV 116-345 | CCAAACAGAAAGAAAAAAAGTTTTGTTTATCAGTCAAACTCCGCCTCTCCAGGAAATGAGT<br>CAATGCCAGAAACCCTGCAGCAATAAAAGTTCAATCATGTAACCACAACTTGGCTGCCTAG<br>GTGACTTTTTTTTTTCAAGTTGGCAGAGGCTTGGGGCTCCTAGCCTCCGAGGCCTCTGGA<br>AAAAAAAGAGAGAGGCCTCTGAGGCTTAAGAGGCTTAATTAGCAAAAA |
| MCPyV 231-464 | TGCCTAGGTGACTTTTTTTTTTCAAGTTGGCAGAGGCTTGGGGCTCCTAGCCTCCGAGGC<br>CTCTGGAAAAAAAAGAGAGAGGCCTCTGAGGCTTAAGAGGCTTAATTAGCAAAAAAGGCA<br>GTATCTAAGGGCAGATCCCAAGGGCGGGAAACTGCAGTATAAAAACCACTCCTTAGTGAG<br>GTAGCTCATTTGCTCCTCTGCTCTTTCTGCAAACTCCTTCTGCATATAGACAAG |

**Dataset S1 (separate file).** Differential gene expression analysis from RNA sequencing of MKL-2 ALTO vs. RFP cells.

**Dataset S2 (separate file).** Gene Set Enrichment Analysis of MKL-2 ALTO vs. RFP upregulated genes.

**Dataset S3 (separate file).** Differential enrichment of biotinylated proteins from MKL-2 TID-ALTO vs. TID cells.
