## Supplementary dataset 1 for "Polyomavirus ALTOs, but not MTs, downregulate viral early gene expression by activating the NF-κB pathway"

| GeneName | GeneType | log2FC | - log10(P) |
|---|---|---|---|
| XAF1 | protein_coding | 14.0437884 | 19.4859357 |
| IFI44 | protein_coding | 13.9492122 | 19.597105 |
| **CXCL10** | protein_coding | 13.0241569 | 26.1501811 |
| CHI3L1 | protein_coding | 12.4315306 | 16.4797267 |
| IL27 | protein_coding | 12.3050029 | 16.3635857 |
| TRIM22 | protein_coding | 11.8507622 | 15.6384935 |
| **CCL2** | protein_coding | 11.72616 | 23.2556158 |
| MARCO | protein_coding | 11.6175244 | 14.8390054 |
| CSF2RB | protein_coding | 11.4965983 | 14.8094506 |
| VCAM1 | protein_coding | 11.3968217 | 14.5215705 |
| CX3CL1 | protein_coding | 11.2646667 | 14.3891465 |
| GBP4 | protein_coding | 11.0012702 | 13.5592764 |
| CD74 | protein_coding | 10.9271411 | 26.4911026 |
| **ICAM1** | protein_coding | 10.7962362 | 24.5985902 |
| HLA-DRB1 | protein_coding | 10.7061342 | 13.2778282 |
| IFI44L | protein_coding | 10.6728516 | 20.8836755 |
| IL17REL | protein_coding | 10.4370651 | 11.9160453 |
| HLA-DRA | protein_coding | 10.429563 | 17.9300128 |
| IFITM3 | protein_coding | 10.3672855 | 21.2661045 |
| MX2 | protein_coding | 10.2444575 | 21.8777507 |
| REM1 | protein_coding | 10.1328218 | 12.0418785 |
| TNFRSF9 | protein_coding | 10.0890872 | 16.4099126 |
| OAS2 | protein_coding | 9.98368159 | 23.7976435 |
| SFTPA1 | protein_coding | 9.89496419 | 11.4460873 |
| MAP3K8 | protein_coding | 9.88579343 | 11.0302539 |
| SELE | protein_coding | 9.83598024 | 10.2026221 |
| TNFAIP6 | protein_coding | 9.63676658 | 15.77013 |
| GJB3 | protein_coding | 9.54538115 | 10.8893708 |
| RIPK2 | protein_coding | 9.54101031 | 10.4006283 |
| CIITA | protein_coding | 9.32523344 | 20.365196 |
| SAMD9L | protein_coding | 9.2092609 | 14.2453347 |
| CEACAM20 | protein_coding | 9.18416136 | 9.85785912 |
| C2CD2 | protein_coding | 9.16299702 | 9.65852165 |
| CLEC7A | protein_coding | 9.12746467 | 10.0982812 |
| HLA-DPB1 | protein_coding | 9.12099435 | 9.78772164 |
| CD70 | protein_coding | 8.96176323 | 9.47439571 |
| SLC26A9 | protein_coding | 8.91804393 | 16.9893076 |
| ICAM2 | protein_coding | 8.88685413 | 9.3454882 |
| RP11-115D1( | lncRNA | 8.87160462 | 9.07728569 |
| APOBEC3B | protein_coding | 8.86195467 | 9.49150276 |
| HLA-DQA1 | protein_coding | 8.85365734 | 9.34675314 |
| IFI16 | protein_coding | 8.84360125 | 8.8835027 |
| CXCL9 | protein_coding | 8.80122892 | 8.71859752 |
| TRIM5 | protein_coding | 8.79196642 | 9.07105394 |

| | | | |
|---|---|---|---|
| SLCO2B1 | protein_coding | 8.77366372 | 9.37335384 |
| SOCS3-DT | lncRNA | 8.72664771 | 8.93387463 |
| TNC | protein_coding | 8.68115098 | 15.4987081 |
| IL32 | protein_coding | 8.65781495 | 18.2819356 |
| NCF4 | protein_coding | 8.588744 | 8.85293079 |
| EBI3 | protein_coding | 8.58704991 | 14.4664227 |
| IFIT3 | protein_coding | 8.56571055 | 25.4293411 |
| IL2RA | protein_coding | 8.52749692 | 7.77109435 |
| WNT2 | protein_coding | 8.52523995 | 8.72102753 |
| DAPP1 | protein_coding | 8.51619088 | 8.20765578 |
| PTGES | protein_coding | 8.49513737 | 18.1100675 |
| HLA-DPA1 | protein_coding | 8.48696782 | 13.1955675 |
| HLA-DRB5 | protein_coding | 8.48213586 | 8.68229871 |
| CXCL11 | protein_coding | 8.41286969 | 20.3977897 |
| IFNL1 | protein_coding | 8.37831441 | 8.44959165 |
| PDZK1IP1 | protein_coding | 8.35454778 | 8.00913195 |
| NEURL3 | protein_coding | 8.31587909 | 8.19914873 |
| SAA1 | protein_coding | 8.25316721 | 8.24859899 |
| CAMK1G | protein_coding | 8.23943653 | 8.18246205 |
| CCL17 | protein_coding | 8.23895042 | 8.11986559 |
| SLC15A3 | protein_coding | 8.18238168 | 16.6426039 |
| IFITM1 | protein_coding | 8.16825973 | 17.1376249 |
| SP100 | protein_coding | 8.16778104 | 16.5920809 |
| IL2RG | protein_coding | 8.14377034 | 8.09019837 |
| IFNB1 | protein_coding | 7.98095375 | 7.48338891 |
| ALPK2 | protein_coding | 7.88460161 | 14.9692162 |
| CASP1 | protein_coding | 7.88260895 | 7.4213973 |
| VIT | protein_coding | 7.86537731 | 7.60521376 |
| **CCL5** | protein_coding | 7.85747379 | 19.1761034 |
| LINC01822 | lncRNA | 7.79476887 | 7.43795379 |
| TYMP | protein_coding | 7.79210847 | 18.1031456 |
| GBP1 | protein_coding | 7.76088886 | 14.2727109 |
| CD80 | protein_coding | 7.75848512 | 7.41474376 |
| MYBPH | protein_coding | 7.7584188 | 7.29610909 |
| HLA-DQB1 | protein_coding | 7.75797366 | 10.967588 |
| NKX2-2 | protein_coding | 7.65386574 | 12.917812 |
| LINC02089 | lncRNA | 7.64539781 | 5.84049007 |
| IL15 | protein_coding | 7.62440628 | 6.75628717 |
| CEACAM1 | protein_coding | 7.61431103 | 10.4186221 |
| LCN2 | protein_coding | 7.60339777 | 6.54888042 |
| RP11-466I1.1 | lncRNA | 7.56282886 | 6.3692961 |
| TNFAIP3 | protein_coding | 7.54468247 | 18.2479534 |
| MX1 | protein_coding | 7.52183549 | 24.7208274 |
| TCIM | protein_coding | 7.51803626 | 6.53524759 |
| RTP4 | protein_coding | 7.49739573 | 6.71227606 |

| | | | |
|---|---|---|---|
| ACHE | protein_coding | 7.4891631 | 16.3847604 |
| RGS1 | protein_coding | 7.47506879 | 6.80495626 |
| CXCL3 | protein_coding | 7.47435467 | 6.79781362 |
| EGOT | lncRNA | 7.47426634 | 6.70751414 |
| IKBKE | protein_coding | 7.464227 | 15.3451219 |
| CAV1 | protein_coding | 7.45239289 | 6.8281743 |
| GBP3 | protein_coding | 7.38169516 | 6.55850005 |
| TMPRSS2 | protein_coding | 7.35927215 | 5.8075654 |
| SAA2 | protein_coding | 7.3579239 | 6.43208829 |
| CYP24A1 | protein_coding | 7.3393761 | 14.6738276 |
| DDO | protein_coding | 7.28350582 | 6.36341901 |
| VSTM1 | protein_coding | 7.25711902 | 5.72367076 |
| RIN3 | protein_coding | 7.25686334 | 5.96471461 |
| SOCS3 | protein_coding | 7.25210618 | 17.5216356 |
| RP11-325K1[illegible] | lncRNA | 7.23086933 | 6.35698295 |
| DNTT | protein_coding | 7.23069011 | 15.0244948 |
| **CXCL8** | protein_coding | 7.22380809 | 10.1725323 |
| IFI27 | protein_coding | 7.22345411 | 25.6218278 |
| GJB4 | protein_coding | 7.14933142 | 5.51812456 |
| KRT14 | protein_coding | 7.13245027 | 11.6815292 |
| BCL3 | protein_coding | 7.1240721 | 18.2390556 |
| TNFRSF8 | protein_coding | 7.12078126 | 6.08177706 |
| ANKRD22 | protein_coding | 7.11517468 | 9.24479922 |
| APOBEC3G | protein_coding | 7.09891267 | 10.3204037 |
| USP30-AS1 | lncRNA | 7.09754273 | 9.61400866 |
| C3 | protein_coding | 7.07534825 | 11.3105945 |
| KRT74 | protein_coding | 7.06313751 | 5.5699951 |
| XKR4 | protein_coding | 7.05735365 | 9.98774914 |
| BCL2L14 | protein_coding | 7.05617721 | 9.41419366 |
| ARHGAP42 | protein_coding | 7.04901682 | 10.1955085 |
| CYTH4 | protein_coding | 7.03189774 | 5.96049974 |
| SFTPA2 | protein_coding | 7.01707099 | 11.1479068 |
| LINC02605 | lncRNA | 7.00127586 | 5.92411589 |
| RP11-471J7.[illegible] | lncRNA | 6.97016992 | 5.95124885 |
| LINC01094 | lncRNA | 6.96999118 | 5.968706 |
| ESM1 | protein_coding | 6.93843535 | 5.67031062 |
| TNFSF8 | protein_coding | 6.92556153 | 9.78674435 |
| ZBP1 | protein_coding | 6.90554836 | 5.59758228 |
| MYBPC2 | protein_coding | 6.85397076 | 13.3020778 |
| OAS1 | protein_coding | 6.84137737 | 26.8182426 |
| HLA-DOA | protein_coding | 6.80177371 | 5.51017786 |
| HLA-DQB2 | protein_coding | 6.80176657 | 5.63241211 |
| LINC00173 | lncRNA | 6.80152142 | 9.60343418 |
| BIRC3 | protein_coding | 6.79188387 | 18.0501378 |
| HSH2D | protein_coding | 6.77150678 | 14.2494262 |

| | | | |
|---|---|---|---|
| S100A14 | protein_coding | 6.76967448 | 13.6998674 |
| NCCRP1 | protein_coding | 6.76170979 | 8.6608107 |
| FOXN1 | protein_coding | 6.7603573 | 9.15650583 |
| TFPI | protein_coding | 6.7284487 | 5.47556818 |
| FEZF2 | protein_coding | 6.7283631 | 5.43478996 |
| OASL | protein_coding | 6.71092989 | 19.3787616 |
| GPR132 | protein_coding | 6.69445333 | 10.5927475 |
| LTB | protein_coding | 6.69037738 | 15.4809601 |
| AIRE | protein_coding | 6.687325 | 8.90168614 |
| HLA-DMB | protein_coding | 6.67407579 | 10.9502944 |
| ODF3B | protein_coding | 6.67181425 | 10.8196458 |
| CCL18 | protein_coding | 6.65082856 | 5.32607879 |
| GPR39 | protein_coding | 6.6105993 | 5.33054115 |
| MSN | protein_coding | 6.61033209 | 8.72491661 |
| ITGAM | protein_coding | 6.5938544 | 11.84152 |
| ICOSLG | protein_coding | 6.53849441 | 8.63040292 |
| STAR | protein_coding | 6.53446724 | 10.2506199 |
| CLIC2 | protein_coding | 6.53113348 | 17.5353061 |
| SPIB | protein_coding | 6.4915256 | 15.5129491 |
| KRT5 | protein_coding | 6.48914849 | 22.0869464 |
| ITGB8 | protein_coding | 6.48601913 | 11.0874603 |
| TNFSF10 | protein_coding | 6.46656381 | 8.18845859 |
| WNT10B | protein_coding | 6.46643291 | 17.616505 |
| TRPV3 | protein_coding | 6.4477812 | 10.6011839 |
| DDX60 | protein_coding | 6.44315047 | 25.9832195 |
| TMEM140 | protein_coding | 6.43233016 | 12.1094832 |
| GAL | protein_coding | 6.40901355 | 11.7861287 |
| IL21R | protein_coding | 6.38939787 | 16.6423209 |
| ISG15 | protein_coding | 6.36539019 | 21.5067627 |
| EGFR | protein_coding | 6.36143222 | 8.50768653 |
| **RELB** | protein_coding | 6.32939237 | 20.1511804 |
| SPRY4 | protein_coding | 6.26885956 | 13.1454658 |
| RSAD2 | protein_coding | 6.25575536 | 24.9553349 |
| KRT2 | protein_coding | 6.2289944 | 14.6139939 |
| SH3TC1 | protein_coding | 6.19663529 | 17.7532972 |
| CTD-3064M3 | lncRNA | 6.15230775 | 9.90958063 |
| LUCAT1 | lncRNA | 6.14206345 | 8.08794521 |
| **HLA-B** | protein_coding | 6.11733427 | 27.4978552 |
| COL19A1 | protein_coding | 6.11543636 | 9.72712252 |
| COL5A3 | protein_coding | 6.10415317 | 10.0877897 |
| CDH5 | protein_coding | 6.09109169 | 9.19789273 |
| UBASH3A | protein_coding | 6.008784 | 7.5800349 |
| DDX58 | protein_coding | 5.98564224 | 13.2075574 |
| IFI35 | protein_coding | 5.97123439 | 21.5103077 |
| CALHM5 | protein_coding | 5.88227024 | 6.96792172 |

| | | | |
|---|---|---|---|
| ARHGEF40 | protein_coding | 5.87985174 | 14.4589961 |
| RP11-199F6. | lncRNA | 5.84471776 | 7.51004472 |
| SLFN5 | protein_coding | 5.82720119 | 8.44666898 |
| CHIT1 | protein_coding | 5.81216689 | 13.2855179 |
| NCF2 | protein_coding | 5.81085921 | 9.3233637 |
| KRT16 | protein_coding | 5.81057367 | 12.8640292 |
| WNT5A | protein_coding | 5.78233836 | 18.0491133 |
| CASC8 | lncRNA | 5.76173271 | 9.46857865 |
| IL34 | protein_coding | 5.74467166 | 8.10315828 |
| CCND1 | protein_coding | 5.72333716 | 7.24307857 |
| UNQ6494 | lncRNA | 5.72283442 | 10.8576278 |
| LT | protein_coding | 5.7137505 | 28.5127263 |
| ATP10D | protein_coding | 5.70255423 | 7.02471557 |
| TNFRSF4 | protein_coding | 5.69057701 | 8.28007059 |
| S100A16 | protein_coding | 5.6803303 | 6.94305106 |
| PARP14 | protein_coding | 5.67716338 | 24.9301187 |
| ATP2C2-AS1 | lncRNA | 5.6706453 | 11.0151444 |
| TNF | protein_coding | 5.64499042 | 22.4019679 |
| KLHL6 | protein_coding | 5.63740503 | 10.9984754 |
| CD40 | protein_coding | 5.62477743 | 15.5308954 |
| TNFSF15 | protein_coding | 5.60073205 | 17.0154957 |
| CXCL5 | protein_coding | 5.58984456 | 6.403425 |
| SLC5A5 | protein_coding | 5.58552625 | 15.779253 |
| NRIR | lncRNA | 5.56820727 | 6.48465344 |
| RP11-450B1! | lncRNA | 5.54241168 | 5.76279011 |
| CLDN14 | protein_coding | 5.53018649 | 10.4155518 |
| RP11-134L1( | lncRNA | 5.522904 | 8.28381946 |
| LINC02091 | lncRNA | 5.51965356 | 6.72340355 |
| CASP4 | protein_coding | 5.49524961 | 6.64573695 |
| CCDC194 | protein_coding | 5.49523679 | 6.7127051 |
| SLC28A3 | protein_coding | 5.48433534 | 17.4456197 |
| SERPINE1 | protein_coding | 5.47461356 | 9.06324042 |
| KRT6A | protein_coding | 5.47101591 | 5.97895441 |
| CSF1 | protein_coding | 5.46372716 | 8.40135806 |
| DTX4 | protein_coding | 5.45690413 | 19.0325245 |
| IER3 | protein_coding | 5.41449561 | 20.0429915 |
| CPXM1 | protein_coding | 5.40068056 | 12.0434913 |
| PLEKHS1 | protein_coding | 5.39371356 | 5.90079994 |
| RP11-310I9.2 | lncRNA | 5.37193509 | 9.27940966 |
| CXCL12 | protein_coding | 5.36069549 | 10.5728691 |
| FLVCR2 | protein_coding | 5.33913255 | 6.17239243 |
| LINC02717 | lncRNA | 5.31072163 | 8.58530285 |
| HERC6 | protein_coding | 5.27648578 | 23.4557852 |
| NFKBIZ | protein_coding | 5.26378882 | 11.9923918 |
| SRGN | protein_coding | 5.25326247 | 6.13911621 |

| | | | |
|---|---|---|---|
| SAMD9 | protein_coding | 5.25108434 | 23.2454368 |
| PLAU | protein_coding | 5.2465489 | 17.7445196 |
| NUPR1 | protein_coding | 5.2282303 | 8.49434606 |
| IFIT1 | protein_coding | 5.22591781 | 23.635097 |
| CTD-2062A1. | lncRNA | 5.19331427 | 5.78580982 |
| KLHDC7B | protein_coding | 5.18141238 | 14.073509 |
| ITGA5 | protein_coding | 5.15829941 | 11.3757847 |
| ELF3-AS1 | lncRNA | 5.1387898 | 8.26492539 |
| ENPP6 | protein_coding | 5.12919776 | 10.5867167 |
| HSPB8 | protein_coding | 5.10614519 | 14.0744887 |
| PLSCR1 | protein_coding | 5.0840833 | 23.4344685 |
| FAS | protein_coding | 5.07734813 | 12.6539341 |
| TSPAN11 | protein_coding | 5.06653213 | 5.78228693 |
| CCK | protein_coding | 5.0509146 | 13.1665179 |
| LILRB4 | protein_coding | 5.03303435 | 5.46976023 |
| KLHL5 | protein_coding | 5.03240606 | 7.99398507 |
| FBLN5 | protein_coding | 5.00293314 | 6.96216976 |
| CTD-2284J15 | lncRNA | 5.00260217 | 7.21041442 |
| SEMA5A | protein_coding | 4.99875535 | 5.62808126 |
| LGALS1 | protein_coding | 4.99847663 | 21.7615861 |
| PLEKHA4 | protein_coding | 4.99113136 | 10.2948543 |
| MYO18B | protein_coding | 4.983873 | 18.7518346 |
| CHST2 | protein_coding | 4.96770004 | 8.74880811 |
| CTA-270D13. | lncRNA | 4.96310076 | 5.51306019 |
| KRT75 | protein_coding | 4.96284752 | 5.3989134 |
| CLDN19 | protein_coding | 4.96078694 | 6.47290936 |
| STAT5A | protein_coding | 4.95266216 | 16.611091 |
| IL4I1 | protein_coding | 4.93855271 | 15.3719534 |
| PLA2G4C | protein_coding | 4.9275996 | 18.5363061 |
| FCAMR | protein_coding | 4.92664793 | 5.3721364 |
| **NFKB2** | protein_coding | 4.91522505 | 20.6046503 |
| RAI2 | protein_coding | 4.90074357 | 11.3478097 |
| SLCO2A1 | protein_coding | 4.89487067 | 9.38271098 |
| PRDM1 | protein_coding | 4.86983731 | 10.2783487 |
| BCAN | protein_coding | 4.85408382 | 8.85743279 |
| OTOF | protein_coding | 4.84575742 | 17.5023083 |
| TMEM163 | protein_coding | 4.84303769 | 8.7732641 |
| BATF2 | protein_coding | 4.83176731 | 15.1193975 |
| BST2 | protein_coding | 4.83122233 | 24.6563303 |
| HLA-F | protein_coding | 4.82661755 | 19.5539639 |
| PARP9 | protein_coding | 4.82138542 | 23.2339247 |
| LRRC55 | protein_coding | 4.81225042 | 5.1500686 |
| ANXA3 | protein_coding | 4.80433316 | 10.4201541 |
| GBP5 | protein_coding | 4.7780816 | 9.00467798 |
| ANKRD29 | protein_coding | 4.76999834 | 9.20632057 |

| | | | |
|---|---|---|---|
| SERPING1 | protein_coding | 4.71367431 | 5.85882744 |
| LAPTM5 | protein_coding | 4.70811131 | 18.0418235 |
| DDX60L | protein_coding | 4.6829414 | 23.8625372 |
| ATP2C2 | protein_coding | 4.67505848 | 15.5044186 |
| ELF3 | protein_coding | 4.67135303 | 13.9284013 |
| ADA2 | protein_coding | 4.66720003 | 14.099579 |
| RP11-283G6. | lncRNA | 4.65139946 | 6.9804458 |
| LPAR1 | protein_coding | 4.63763746 | 12.1807794 |
| LGI2 | protein_coding | 4.60712662 | 11.6888232 |
| RASGRP3 | protein_coding | 4.58314283 | 6.26220281 |
| TUNAR | protein_coding | 4.57972508 | 17.9020492 |
| CH507-9B2.1 | protein_coding | 4.56301515 | 14.3286472 |
| MYOSLID | lncRNA | 4.55796516 | 7.3226743 |
| PDGFRA | protein_coding | 4.55730085 | 7.31645651 |
| CDH16 | protein_coding | 4.55681451 | 6.94700174 |
| LGALS9 | protein_coding | 4.54463506 | 15.048498 |
| TWIST1 | protein_coding | 4.53706224 | 12.0780201 |
| BTN2A2 | protein_coding | 4.52414178 | 16.8288332 |
| CTSS | protein_coding | 4.51189862 | 10.1383537 |
| CLU | protein_coding | 4.50099778 | 21.3964466 |
| WAKMAR2 | lncRNA | 4.49967415 | 6.02987822 |
| TRPV4 | protein_coding | 4.4993896 | 5.21964201 |
| TAP1 | protein_coding | 4.49675548 | 24.8579064 |
| LYPD1 | protein_coding | 4.49398491 | 15.8938144 |
| KLK10 | protein_coding | 4.47692321 | 5.6740859 |
| PLEK | protein_coding | 4.47103696 | 12.1361515 |
| TP63 | protein_coding | 4.44812409 | 9.52969573 |
| IRF4 | protein_coding | 4.44194365 | 12.4036602 |
| TRIM21 | protein_coding | 4.41167435 | 18.3909393 |
| KRT7 | protein_coding | 4.40539457 | 13.8029507 |
| WIPF3 | protein_coding | 4.39204715 | 9.57738956 |
| MUCL1 | protein_coding | 4.36675285 | 7.67494053 |
| HLA-G | protein_coding | 4.36024823 | 10.9741982 |
| PSMB9 | protein_coding | 4.34796495 | 21.0538635 |
| SP110 | protein_coding | 4.339055 | 19.2862673 |
| RPRM | protein_coding | 4.33869351 | 14.2001507 |
| GALNT15 | protein_coding | 4.33303576 | 12.322339 |
| FILIP1L | protein_coding | 4.30399883 | 6.2066317 |
| HLA-DOB | protein_coding | 4.26661182 | 8.15265402 |
| LINC02723 | lncRNA | 4.24069689 | 6.59763661 |
| TLR3 | protein_coding | 4.23987341 | 8.97041649 |
| NPHS1 | protein_coding | 4.23884568 | 17.5169262 |
| PLAAT4 | protein_coding | 4.23767561 | 12.6778429 |
| FAP | protein_coding | 4.23310324 | 5.59407546 |
| MAML2 | protein_coding | 4.2256123 | 7.78684539 |

| | | | |
|---|---|---|---|
| PITPNC1 | protein_coding | 4.21497515 | 13.7250989 |
| KCNF1 | protein_coding | 4.21403139 | 10.0260176 |
| CYP4B1 | protein_coding | 4.20222953 | 6.1317247 |
| KLHDC7B-DT | lncRNA | 4.18275122 | 5.49544521 |
| IFIT2 | protein_coding | 4.17508496 | 18.6207891 |
| RASGEF1A | protein_coding | 4.16945217 | 10.2425749 |
| APOL6 | protein_coding | 4.16172856 | 22.3176496 |
| TNNI2 | protein_coding | 4.15849863 | 13.806093 |
| RP11-1136L8 | lncRNA | 4.14559925 | 9.51975642 |
| DRAM1 | protein_coding | 4.13288623 | 10.5257181 |
| PIK3CD | protein_coding | 4.12939509 | 14.942821 |
| PSMB10 | protein_coding | 4.11467532 | 14.6273481 |
| CHRNA6 | protein_coding | 4.10707103 | 4.73568702 |
| STING1 | protein_coding | 4.09831951 | 16.7395074 |
| COL8A2 | protein_coding | 4.09016831 | 8.27336242 |
| DHX58 | protein_coding | 4.08285743 | 19.8852914 |
| KCNN3 | protein_coding | 4.08057584 | 17.3013255 |
| PRODH2 | protein_coding | 4.07849716 | 5.85436103 |
| ISG20 | protein_coding | 4.07115116 | 14.290205 |
| CEMIP | protein_coding | 4.05138335 | 5.71659529 |
| GALNT14 | protein_coding | 4.04729556 | 9.09432004 |
| EDARADD | protein_coding | 4.0418246 | 5.73707708 |
| SEMA4A | protein_coding | 4.03521661 | 9.84711765 |
| LINC01619 | lncRNA | 4.02956538 | 4.5410474 |
| CPED1 | protein_coding | 4.02321958 | 6.14820907 |
| KLK11 | protein_coding | 4.02124236 | 10.5695335 |
| CHST15 | protein_coding | 4.0143304 | 12.558224 |
| **TRAF1** | protein_coding | 3.99710015 | 19.9704724 |
| **HLA-C** | protein_coding | 3.9953546 | 22.2605725 |
| CREB3L3 | protein_coding | 3.985325 | 6.16244599 |
| CTD-3064M3 | lncRNA | 3.97026952 | 6.46520913 |
| POTEH | protein_coding | 3.96640105 | 4.92824012 |
| ADPRH | protein_coding | 3.95487812 | 6.75429461 |
| ANO9 | protein_coding | 3.95394734 | 24.7726519 |
| PDE6G | protein_coding | 3.95233519 | 8.31225136 |
| CITED4 | protein_coding | 3.94795308 | 12.6201021 |
| DTX3L | protein_coding | 3.93881729 | 22.651712 |
| COL17A1 | protein_coding | 3.93673002 | 12.5936148 |
| PAPPA | protein_coding | 3.93415223 | 13.4440166 |
| RXFP4 | protein_coding | 3.92680963 | 6.51619988 |
| GADD45B | protein_coding | 3.91859388 | 14.4932123 |
| EPDR1 | protein_coding | 3.90821889 | 6.53497638 |
| PKIG | protein_coding | 3.90772894 | 7.31791537 |
| APOL1 | protein_coding | 3.89506626 | 18.9510201 |
| CMPK2 | protein_coding | 3.89428294 | 22.7107272 |

| | | | |
|---|---|---|---|
| NRG4 | protein_coding | 3.88805295 | 11.7895007 |
| RARG | protein_coding | 3.88033835 | 17.838066 |
| BCL6 | protein_coding | 3.8803295 | 9.10464956 |
| SMIM35 | protein_coding | 3.8741962 | 12.0106184 |
| CXCL13 | protein_coding | 3.86131497 | 9.37411182 |
| SLC35F3 | protein_coding | 3.85721151 | 12.2556672 |
| SNCAIP | protein_coding | 3.85716569 | 6.36225117 |
| KCNIP4 | protein_coding | 3.85032281 | 7.65624023 |
| UBA7 | protein_coding | 3.84437147 | 17.1558251 |
| RDH16 | protein_coding | 3.83858043 | 11.4356271 |
| S100A3 | protein_coding | 3.82005208 | 5.70025435 |
| RAPGEF3 | protein_coding | 3.81813722 | 10.8799918 |
| SOCS1 | protein_coding | 3.81544254 | 14.0512122 |
| RAMP3 | protein_coding | 3.8121496 | 5.2077006 |
| TNFRSF14 | protein_coding | 3.80151252 | 14.2547682 |
| CCL19 | protein_coding | 3.79802954 | 9.89200901 |
| CHST3 | protein_coding | 3.79569803 | 9.30684654 |
| HES1 | protein_coding | 3.77881733 | 12.6077942 |
| GPR55 | protein_coding | 3.76589461 | 7.68860637 |
| RERG | protein_coding | 3.76584615 | 9.56754127 |
| ZNF385C | protein_coding | 3.7653638 | 19.9659276 |
| ANXA13 | protein_coding | 3.76216329 | 9.51914384 |
| CBR3 | protein_coding | 3.75932192 | 9.16480758 |
| IFFO2 | protein_coding | 3.75345549 | 17.8999602 |
| HERC5 | protein_coding | 3.7513203 | 21.2661894 |
| FABP6 | protein_coding | 3.75014945 | 12.6049658 |
| PATL2 | protein_coding | 3.7453303 | 8.13532241 |
| NGFR | protein_coding | 3.72456921 | 15.4927775 |
| GNAL | protein_coding | 3.71175233 | 4.88799318 |
| UBE2L6 | protein_coding | 3.69735931 | 23.6610757 |
| ACSL5 | protein_coding | 3.69692333 | 12.0104448 |
| CXCL1 | protein_coding | 3.68045255 | 6.79416707 |
| RP11-378I6.1 | lncRNA | 3.66833345 | 7.34419617 |
| TRIM56 | protein_coding | 3.64826237 | 20.4911482 |
| FAM83B | protein_coding | 3.63173151 | 6.12665266 |
| HCAR3 | protein_coding | 3.62340903 | 6.92834104 |
| SPRY3 | protein_coding | 3.62333682 | 4.73649563 |
| RNF213 | protein_coding | 3.61943735 | 25.680355 |
| EGFL6 | protein_coding | 3.61569653 | 9.20366353 |
| MUC4 | protein_coding | 3.60986157 | 15.1018908 |
| APOL3 | protein_coding | 3.60149654 | 17.0732194 |
| KIAA0040 | protein_coding | 3.58795764 | 7.48058008 |
| IFIH1 | protein_coding | 3.58051521 | 24.1068425 |
| CRB2 | protein_coding | 3.57645154 | 6.86488186 |
| PKDCC | protein_coding | 3.57385391 | 8.17247143 |

| | | | |
|---|---|---|---|
| ANXA6 | protein_coding | 3.55273985 | 23.8467422 |
| TEX101 | protein_coding | 3.5353378 | 8.79938198 |
| APOE | protein_coding | 3.52564124 | 13.5484764 |
| TGM4 | protein_coding | 3.52471259 | 4.04349032 |
| FRMD1 | protein_coding | 3.52101933 | 11.7523121 |
| NLRP1 | protein_coding | 3.49426218 | 18.4420975 |
| HMOX1 | protein_coding | 3.48112887 | 16.1324987 |
| IRAK2 | protein_coding | 3.4798044 | 6.85448887 |
| LINC02822 | lncRNA | 3.46626398 | 6.07193184 |
| CD274 | protein_coding | 3.46381837 | 13.309026 |
| LIF | protein_coding | 3.45719335 | 7.49129073 |
| CCN3 | protein_coding | 3.45678371 | 7.35219974 |
| B2M | protein_coding | 3.45206445 | 25.1471731 |
| GATA6 | protein_coding | 3.44720596 | 9.30955532 |
| LTBP2 | protein_coding | 3.44477517 | 18.7904043 |
| ESRRB | protein_coding | 3.44216379 | 20.3941838 |
| CLIP3 | protein_coding | 3.44081185 | 19.0591756 |
| RP11-37B2.1 | lncRNA | 3.42969404 | 6.68312829 |
| BCO1 | protein_coding | 3.41250523 | 12.2533718 |
| COLCA1 | lncRNA | 3.40690307 | 15.6997273 |
| LINC02362 | lncRNA | 3.40576343 | 4.49947789 |
| OAS3 | protein_coding | 3.40040999 | 27.0945521 |
| ICAM4 | protein_coding | 3.38454341 | 5.96216529 |
| LRG1 | protein_coding | 3.38252726 | 5.02687191 |
| LRRC32 | protein_coding | 3.36610146 | 9.5964617 |
| ANKRD6 | protein_coding | 3.35589516 | 16.2716539 |
| PLXNA4 | protein_coding | 3.34341215 | 15.3483609 |
| LINC02137 | lncRNA | 3.34183113 | 8.31048069 |
| LAMC2 | protein_coding | 3.34142938 | 11.7974304 |
| BISPR | lncRNA | 3.33490861 | 12.5482889 |
| SERPINE2 | protein_coding | 3.32613815 | 12.7763439 |
| PTX3 | protein_coding | 3.32547654 | 6.27090546 |
| NLRC5 | protein_coding | 3.32146084 | 23.0017411 |
| C1QTNF1 | protein_coding | 3.31713839 | 15.2038105 |
| NFKBIA | protein_coding | 3.31542999 | 23.6262974 |
| **HLA-A** | protein_coding | 3.30796165 | 23.8801245 |
| BDKRB2 | protein_coding | 3.30556876 | 10.7728195 |
| TENM2 | protein_coding | 3.29859921 | 14.9306078 |
| IL23A | protein_coding | 3.26814152 | 11.1958106 |
| UMODL1 | protein_coding | 3.26798768 | 6.15156041 |
| KREMEN2 | protein_coding | 3.25537629 | 15.2835991 |
| PECAM1 | protein_coding | 3.24958866 | 6.80341265 |
| RFTN1 | protein_coding | 3.24729704 | 16.1877671 |
| REC8 | protein_coding | 3.24584524 | 17.6595911 |
| GGT5 | protein_coding | 3.24160001 | 7.54024707 |

| | | | |
|---|---|---|---|
| BTN3A3 | protein_coding | 3.23755515 | 14.688507 |
| COL16A1 | protein_coding | 3.23288642 | 18.4006948 |
| HELZ2 | protein_coding | 3.22962896 | 17.4674133 |
| SIGLEC15 | protein_coding | 3.22835267 | 6.2584154 |
| NOX4 | protein_coding | 3.20721791 | 10.8997599 |
| HMCN2 | protein_coding | 3.20309292 | 22.8195014 |
| UNC5C-AS1 | lncRNA | 3.19149323 | 4.76072342 |
| PRELP | protein_coding | 3.18222346 | 14.3824471 |
| NECTIN4 | protein_coding | 3.1685846 | 5.03118005 |
| PAPLN | protein_coding | 3.16768379 | 12.7150915 |
| CA4 | protein_coding | 3.15804542 | 7.06031806 |
| STAT1 | protein_coding | 3.15525994 | 24.9134852 |
| RP11-536O1 | lncRNA | 3.15316573 | 6.55215461 |
| GABARAPL1 | protein_coding | 3.14266676 | 13.6148993 |
| DST | protein_coding | 3.1414218 | 13.8734333 |
| CA5A | protein_coding | 3.13491439 | 9.81765096 |
| LINC01227 | lncRNA | 3.13081829 | 7.47494071 |
| LCAL1 | lncRNA | 3.12539585 | 19.5621627 |
| UNC5C | protein_coding | 3.11285906 | 13.158946 |
| CEACAM16-A | lncRNA | 3.11079641 | 4.42839081 |
| BMP6 | protein_coding | 3.11059758 | 13.1669007 |
| RP11-134L1C | lncRNA | 3.1087376 | 5.26316252 |
| COL27A1 | protein_coding | 3.08833341 | 13.9689197 |
| FBXO6 | protein_coding | 3.0839878 | 12.5636799 |
| CTD-2213F21 | lncRNA | 3.0762133 | 4.59680751 |
| AATBC | lncRNA | 3.07325315 | 13.1450644 |
| USP18 | protein_coding | 3.07304378 | 21.7675338 |
| SSTR2 | protein_coding | 3.07292227 | 17.9300971 |
| KCNA2 | protein_coding | 3.06040796 | 15.7379402 |
| **IRF1** | protein_coding | 3.03007362 | 21.711844 |
| MCAM | protein_coding | 3.02828746 | 19.9663173 |
| NFE2L3 | protein_coding | 3.01671967 | 18.2439415 |
| IL6 | protein_coding | 3.0076571 | 6.42370046 |
| FAAH2 | protein_coding | 3.00721841 | 5.51334927 |
| PARP12 | protein_coding | 3.00456692 | 20.7464252 |
| NECAB1 | protein_coding | 3.00194156 | 8.48716863 |
| HCP5 | lncRNA | 3.00102025 | 18.4816226 |
| GPR37L1 | protein_coding | 2.99854852 | 8.1907966 |
| PHF11 | protein_coding | 2.99211327 | 13.1153982 |
| CHL1 | protein_coding | 2.98988127 | 8.81984924 |
| SGPP2 | protein_coding | 2.98820409 | 18.357901 |
| THBS4 | protein_coding | 2.98625702 | 17.6032051 |
| CCR10 | protein_coding | 2.97701518 | 9.33949595 |
| C1R | protein_coding | 2.97426348 | 10.1196086 |
| ADAM8 | protein_coding | 2.97342938 | 10.6040412 |

| | | | |
|---|---|---|---|
| SMAD3 | protein_coding | 2.97122705 | 21.3031805 |
| RND1 | protein_coding | 2.97047232 | 15.0217382 |
| MYC | protein_coding | 2.96926343 | 14.9855705 |
| IFI6 | protein_coding | 2.96808651 | 15.4855906 |
| HLA-E | protein_coding | 2.96034666 | 26.5184133 |
| APOBEC3F | protein_coding | 2.95834883 | 10.7360651 |
| ATXN1 | protein_coding | 2.95070462 | 8.9896796 |
| PTGER4 | protein_coding | 2.94523301 | 5.13342984 |
| OPTN | protein_coding | 2.93960219 | 20.5194122 |
| PTN | protein_coding | 2.92890818 | 7.59565237 |
| NFKB1 | protein_coding | 2.92788975 | 16.5081575 |
| PYCARD | protein_coding | 2.921812 | 7.49819859 |
| TMC1 | protein_coding | 2.91632917 | 6.88891912 |
| TNFAIP8L3 | protein_coding | 2.91543988 | 7.71509502 |
| COX6B2 | protein_coding | 2.91521043 | 10.5548644 |
| LAMP3 | protein_coding | 2.9012549 | 21.0795662 |
| PROSER2 | protein_coding | 2.89885643 | 4.97368284 |
| PARP10 | protein_coding | 2.88884463 | 18.6395997 |
| LINC02499 | lncRNA | 2.87587164 | 4.52580678 |
| SLC28A1 | protein_coding | 2.8681056 | 5.98997732 |
| CRISPLD2 | protein_coding | 2.86729281 | 15.2080524 |
| PTP4A3 | protein_coding | 2.865212 | 18.6766931 |
| NEDD9 | protein_coding | 2.86355432 | 6.36808211 |
| TVP23A | protein_coding | 2.86323445 | 4.22256902 |
| AMBP | protein_coding | 2.8630566 | 6.49018696 |
| NCEH1 | protein_coding | 2.85579969 | 5.86892191 |
| KIFC3 | protein_coding | 2.85567411 | 4.37143343 |
| STAP2 | protein_coding | 2.8546466 | 17.8674226 |
| IRF9 | protein_coding | 2.85444607 | 13.4138957 |
| TAP2 | protein_coding | 2.85417458 | 21.3903932 |
| ADAD2 | protein_coding | 2.84103463 | 4.68994657 |
| C2CD4B | protein_coding | 2.83182052 | 4.81954976 |
| ALOX5 | protein_coding | 2.82140282 | 8.42557254 |
| ADRA2A | protein_coding | 2.8170273 | 7.98494054 |
| MIR23AHG | lncRNA | 2.81598964 | 7.02530669 |
| RASSF6 | protein_coding | 2.815902 | 10.2647909 |
| LCAT | protein_coding | 2.81009092 | 5.48063325 |
| SHFL | protein_coding | 2.80966959 | 19.9813169 |
| ADAM12 | protein_coding | 2.80712796 | 9.54774768 |
| SH2D3C | protein_coding | 2.80645908 | 11.7161022 |
| C1QTNF1-AS | lncRNA | 2.79986113 | 11.6481961 |
| C2CD4A | protein_coding | 2.79951886 | 14.5268436 |
| KCND3 | protein_coding | 2.78695709 | 7.57040044 |
| CCNJL | protein_coding | 2.78592165 | 9.7540322 |
| IZUMO4 | protein_coding | 2.7769118 | 10.5562056 |

| | | | |
|---|---|---|---|
| ESPNL | protein_coding | 2.75950038 | 10.1400881 |
| BNIP5 | protein_coding | 2.7362172 | 6.84664705 |
| ANKRD53 | protein_coding | 2.73046327 | 6.39596251 |
| LSP1 | protein_coding | 2.72884706 | 15.3934363 |
| RP11-118H4. | lncRNA | 2.7249906 | 4.05730901 |
| RP11-182J23 | lncRNA | 2.71500508 | 4.86315991 |
| FCGBP | protein_coding | 2.71138116 | 7.55641676 |
| SLC12A4 | protein_coding | 2.70597425 | 10.5028686 |
| ABLIM3 | protein_coding | 2.70475996 | 7.93603211 |
| PSMB8 | protein_coding | 2.70271369 | 21.4711929 |
| STAC | protein_coding | 2.67611316 | 11.3940159 |
| RUFY4 | protein_coding | 2.67191332 | 11.6195508 |
| GFRA1 | protein_coding | 2.66638952 | 14.2545185 |
| CTSO | protein_coding | 2.66520386 | 10.6319239 |
| FAM78B | protein_coding | 2.6552913 | 6.6432086 |
| ITGA9 | protein_coding | 2.65424757 | 4.00160438 |
| COLQ | protein_coding | 2.65218368 | 11.993585 |
| ADM2 | protein_coding | 2.6512437 | 20.7940431 |
| RNF207 | protein_coding | 2.64605192 | 7.1034837 |
| TMC8 | protein_coding | 2.64489977 | 6.70451618 |
| ABCC3 | protein_coding | 2.64272403 | 6.13041886 |
| BTN3A2 | protein_coding | 2.64048576 | 15.8109603 |
| MIR22HG | lncRNA | 2.63930205 | 9.39567341 |
| RASSF2 | protein_coding | 2.63421753 | 16.7992173 |
| MDGA1 | protein_coding | 2.62175429 | 17.3856839 |
| CBLN3 | protein_coding | 2.61970218 | 9.0314406 |
| IFIT5 | protein_coding | 2.61824367 | 18.6080875 |
| GPR142 | protein_coding | 2.6165295 | 4.98283457 |
| ETV7-AS1 | lncRNA | 2.6132311 | 6.24709604 |
| GABRP | protein_coding | 2.60871605 | 4.41124786 |
| S100A10 | protein_coding | 2.60087502 | 9.1080883 |
| POPDC2 | protein_coding | 2.58739602 | 6.17525449 |
| MIR4500HG | lncRNA | 2.57948729 | 5.95257901 |
| BTN3A1 | protein_coding | 2.57847311 | 16.7706648 |
| DPP4 | protein_coding | 2.56664881 | 4.68353302 |
| SLC12A1 | protein_coding | 2.56517179 | 5.99044624 |
| SCNN1G | protein_coding | 2.56101929 | 16.8305314 |
| LTA | protein_coding | 2.55434875 | 5.38774847 |
| IKZF3 | protein_coding | 2.54427585 | 8.05344955 |
| WNT7B | protein_coding | 2.54057454 | 18.1413581 |
| HCAR2 | protein_coding | 2.54041581 | 7.62181345 |
| SEMA7A | protein_coding | 2.53933508 | 13.6069112 |
| DUSP10 | protein_coding | 2.53861293 | 13.7957327 |
| BDNF | protein_coding | 2.53574335 | 12.9632463 |
| RP11-177H1: | lncRNA | 2.53408277 | 6.77661005 |

| | | | |
|---|---|---|---|
| CGNL1 | protein_coding | 2.52457664 | 9.86668676 |
| RGS8 | protein_coding | 2.51563051 | 4.37171606 |
| PARM1 | protein_coding | 2.51230147 | 10.3136602 |
| C15orf62 | protein_coding | 2.51079815 | 8.71511522 |
| TNFSF9 | protein_coding | 2.50834687 | 4.15746348 |
| CYP1A1 | protein_coding | 2.50797975 | 4.53409699 |
| LINC02574 | lncRNA | 2.5035698 | 5.74741463 |
| DGAT2 | protein_coding | 2.50285485 | 20.2515965 |
| SAMD4A | protein_coding | 2.49320894 | 6.88045921 |
| PIK3R6 | protein_coding | 2.48898263 | 11.5038057 |
| RP11-29H23. | lncRNA | 2.48235003 | 5.47956944 |
| LINC01348 | lncRNA | 2.47795072 | 5.1481691 |
| PDGFRB | protein_coding | 2.47773365 | 3.9880442 |
| AC010226.4 | lncRNA | 2.47340437 | 6.47262353 |
| C9orf139 | lncRNA | 2.47128828 | 4.47484345 |
| PLXNC1 | protein_coding | 2.46736011 | 3.67324148 |
| ST18 | protein_coding | 2.46565826 | 16.1226045 |
| SERPINB1 | protein_coding | 2.46181512 | 11.9495321 |
| SV2C | protein_coding | 2.46147566 | 10.8942828 |
| MIR7-3HG | lncRNA | 2.45581401 | 6.12514359 |
| CREB5 | protein_coding | 2.45016667 | 11.0335175 |
| CXCR5 | protein_coding | 2.44655906 | 7.58614881 |
| P2RY6 | protein_coding | 2.43801314 | 6.72462593 |
| SLC24A2 | protein_coding | 2.43701208 | 14.8236416 |
| TAPBP | protein_coding | 2.43607488 | 20.7472619 |
| FRK | protein_coding | 2.43204442 | 13.3291632 |
| SAMHD1 | protein_coding | 2.42919687 | 20.3645579 |
| SORCS1 | protein_coding | 2.4285065 | 10.7785083 |
| MGLL | protein_coding | 2.42783294 | 22.0035475 |
| MVP | protein_coding | 2.42575379 | 17.6859747 |
| PPP1R18 | protein_coding | 2.4067612 | 19.1681587 |
| TMEFF2 | protein_coding | 2.40663984 | 8.36654401 |
| MAP1A | protein_coding | 2.40326526 | 19.1030551 |
| HLA-DMA | protein_coding | 2.40166029 | 9.01569867 |
| PRSS8 | protein_coding | 2.40074111 | 17.7677682 |
| FN1 | protein_coding | 2.396153 | 15.5791345 |
| MPZ | protein_coding | 2.39143821 | 15.6069449 |
| RP4-739H11. | lncRNA | 2.37971927 | 6.78338807 |
| NUP153-AS1 | lncRNA | 2.37938558 | 8.543629 |
| CDK18 | protein_coding | 2.37921611 | 15.0071092 |
| LINC01894 | lncRNA | 2.37344313 | 4.00599106 |
| CTD-2095E4. | lncRNA | 2.37004474 | 11.2224541 |
| TRIM69 | protein_coding | 2.36983779 | 21.0893007 |
| CALB1 | protein_coding | 2.35857914 | 9.95668927 |
| ARSJ | protein_coding | 2.35745595 | 11.9288942 |

| | | | |
|---|---|---|---|
| CCL25 | protein_coding | 2.35456001 | 5.13669898 |
| ERAP2 | protein_coding | 2.35424448 | 18.6139839 |
| FBN1 | protein_coding | 2.35225603 | 9.71913389 |
| RP11-525K1( | lncRNA | 2.33945753 | 7.95200628 |
| IRF1-AS1 | lncRNA | 2.33474567 | 12.4219546 |
| MIOX | protein_coding | 2.33332513 | 5.17475208 |
| ADAM19 | protein_coding | 2.32774957 | 18.5724881 |
| VWC2L | protein_coding | 2.31865615 | 6.67525571 |
| PIK3R5 | protein_coding | 2.3184266 | 13.6977331 |
| THSD7B | protein_coding | 2.31790169 | 5.7647054 |
| CAPG | protein_coding | 2.3148147 | 5.23717708 |
| KCNIP3 | protein_coding | 2.30901124 | 15.7536273 |
| SGK1 | protein_coding | 2.30280304 | 15.5936731 |
| ZNF750 | protein_coding | 2.30225704 | 8.11288225 |
| PAX2 | protein_coding | 2.30161358 | 4.41018749 |
| CCND2 | protein_coding | 2.3009087 | 9.63564205 |
| GPBAR1 | protein_coding | 2.29567097 | 10.0785647 |
| ETV7 | protein_coding | 2.29389446 | 17.9247361 |
| SPATS2L | protein_coding | 2.29004317 | 7.22519573 |
| LINC00877 | lncRNA | 2.28930039 | 3.01708383 |
| CNTNAP1 | protein_coding | 2.28359666 | 17.6327897 |
| CDC42EP1 | protein_coding | 2.27692872 | 9.49589839 |
| TRIM2 | protein_coding | 2.27509418 | 3.28456603 |
| ZYX | protein_coding | 2.27048527 | 11.1856497 |
| CXXC5 | protein_coding | 2.26268491 | 16.2404128 |
| EPHA2 | protein_coding | 2.26196782 | 10.0188438 |
| SNX10 | protein_coding | 2.26064981 | 16.9288411 |
| MGC15885 | lncRNA | 2.25951636 | 3.56562083 |
| AKR1C3 | protein_coding | 2.25481844 | 10.6819189 |
| ABTB2 | protein_coding | 2.23857005 | 13.6582534 |
| RP11-731I21 | lncRNA | 2.23768142 | 5.14823727 |
| EPHA4 | protein_coding | 2.23638867 | 15.9828015 |
| IL6R | protein_coding | 2.2339808 | 9.58440421 |
| CLIP2 | protein_coding | 2.23019585 | 19.7489591 |
| RP11-844P9. | lncRNA | 2.21736797 | 3.78309459 |
| MIR646HG | lncRNA | 2.21721266 | 5.74987465 |
| BRINP2 | protein_coding | 2.20622086 | 5.67687236 |
| OSGIN1 | protein_coding | 2.2041092 | 10.7040068 |
| RP11-396B1 | lncRNA | 2.20323966 | 4.89457141 |
| ELMOD1 | protein_coding | 2.19117591 | 9.17746218 |
| CR1L | protein_coding | 2.18998927 | 5.31916483 |
| EMID1 | protein_coding | 2.18931518 | 12.3539889 |
| IRX6 | protein_coding | 2.18809047 | 11.3843675 |
| AC008079.1( | lncRNA | 2.18450017 | 9.13018894 |
| VSIR | protein_coding | 2.1833414 | 4.4230061 |

| | | | |
|---|---|---|---|
| RHBDF2 | protein_coding | 2.18166063 | 13.6835418 |
| MATN2 | protein_coding | 2.17652133 | 5.33231311 |
| RPH3AL-AS1 | lncRNA | 2.17185782 | 4.29309362 |
| GEM | protein_coding | 2.17118289 | 6.99492477 |
| NFATC2 | protein_coding | 2.17053305 | 7.60271869 |
| ST6GALNAC2 | protein_coding | 2.1677538 | 7.90542764 |
| FZD9 | protein_coding | 2.16535525 | 9.2796928 |
| DSC2 | protein_coding | 2.16349529 | 11.6770154 |
| ANO2 | protein_coding | 2.14370508 | 3.90580224 |
| PIWIL2 | protein_coding | 2.13924679 | 4.70123222 |
| FERMT1 | protein_coding | 2.1388842 | 4.8559579 |
| ARHGEF16 | protein_coding | 2.13652452 | 15.6851979 |
| CTD-2201G3. | lncRNA | 2.12933002 | 3.76082421 |
| RP11-473M2 | lncRNA | 2.12299939 | 4.11531297 |
| LHFPL6 | protein_coding | 2.11891072 | 4.4458175 |
| NAT1 | protein_coding | 2.11799489 | 6.95014002 |
| DIO3 | protein_coding | 2.11551281 | 7.81237863 |
| ITPR1 | protein_coding | 2.11434772 | 12.2075222 |
| CLN3 | protein_coding | 2.11209488 | 6.31862867 |
| MOB3C | protein_coding | 2.10318861 | 11.3392748 |
| PPM1K | protein_coding | 2.09942034 | 13.4349713 |
| CNN3-DT | lncRNA | 2.08826486 | 9.87975463 |
| PDPN | protein_coding | 2.08336333 | 5.30630783 |
| AZIN2 | protein_coding | 2.07969554 | 12.880228 |
| LINC02520 | lncRNA | 2.07815823 | 4.85359492 |
| COL18A1 | protein_coding | 2.07724802 | 7.57360221 |
| INSYN2A | protein_coding | 2.07642058 | 11.7328343 |
| RRAD | protein_coding | 2.07458623 | 7.55391071 |
| TRIM50 | protein_coding | 2.06952773 | 4.51156671 |
| MSX1 | protein_coding | 2.06892199 | 6.76185012 |
| CREB3L1 | protein_coding | 2.06847198 | 11.6861749 |
| ETV5 | protein_coding | 2.06739455 | 12.5735828 |
| ADAMTS18 | protein_coding | 2.06687923 | 4.87561524 |
| RFLNA | protein_coding | 2.06431014 | 4.94787201 |
| GAS6-DT | lncRNA | 2.06221487 | 11.3164647 |
| TLR5 | protein_coding | 2.05811645 | 6.83442467 |
| RP11-245P1( | lncRNA | 2.05654816 | 6.59160617 |
| C3orf52 | protein_coding | 2.05618781 | 6.28313583 |
| NCKAP5 | protein_coding | 2.05549475 | 10.6160165 |
| SLC4A11 | protein_coding | 2.05361207 | 13.1992718 |
| FNBP1 | protein_coding | 2.05337762 | 17.3989684 |
| TNIP1 | protein_coding | 2.05120951 | 19.0801605 |
| LGALS3BP | protein_coding | 2.04443256 | 22.6908408 |
| BAG3 | protein_coding | 2.04405253 | 6.66669522 |
| WNT4 | protein_coding | 2.04283628 | 10.1808784 |

| | | | |
|---|---|---|---|
| TVP23C | protein_coding | 2.04147315 | 7.10568915 |
| PSMB8-AS1 | lncRNA | 2.03888981 | 12.4497571 |
| RAB24 | protein_coding | 2.0357458 | 8.21557861 |
| PSME2 | protein_coding | 2.03463443 | 21.3806619 |
| WNT10A | protein_coding | 2.02766635 | 18.2370013 |
| SEMA5B | protein_coding | 2.02454131 | 4.33111489 |
| LRRIQ1 | protein_coding | 2.02445058 | 14.0451234 |
| LAMA4 | protein_coding | 2.01632109 | 5.43536086 |
| SLCO5A1 | protein_coding | 2.01518996 | 13.863464 |
| BHLHE41 | protein_coding | 2.01453352 | 5.21118483 |
| INSRR | protein_coding | 2.01449559 | 13.0873261 |
| ZBTB46 | protein_coding | 2.0144168 | 12.6499573 |
| PDE6B | protein_coding | 2.01347242 | 12.8476086 |
| ZC3H12A | protein_coding | 2.01322456 | 13.280311 |
| TAPBPL | protein_coding | 2.00773724 | 12.9597506 |
| FAM167A | protein_coding | 2.00328548 | 15.1646012 |
| ADAMTS9 | protein_coding | 2.00316838 | 10.6804359 |
| MOV10 | protein_coding | 2.00159723 | 19.0871396 |
| TNFAIP2 | protein_coding | 1.99875521 | 21.1610621 |
| BRINP3 | protein_coding | 1.99677012 | 6.18643064 |
| C3orf80 | protein_coding | 1.98833766 | 14.264842 |
| GADD45A | protein_coding | 1.981695 | 12.824663 |
| SERPINB5 | protein_coding | 1.98141736 | 9.00018447 |
| SCARA3 | protein_coding | 1.97360542 | 6.60730794 |
| EXT1 | protein_coding | 1.97150129 | 16.7660103 |
| ZNF44 | protein_coding | 1.96240806 | 7.28755339 |
| RENBP | protein_coding | 1.96052441 | 8.69818028 |
| IL12A-AS1 | lncRNA | 1.95939558 | 9.39433582 |
| BDKRB1 | protein_coding | 1.95358683 | 3.82728021 |
| TLE4 | protein_coding | 1.95289917 | 9.96489493 |
| EVX2 | protein_coding | 1.95221937 | 7.19192064 |
| FAXDC2 | protein_coding | 1.95160329 | 12.290959 |
| RP11-386M2 | lncRNA | 1.9490514 | 4.15337827 |
| LAP3 | protein_coding | 1.94805534 | 21.3012695 |
| SYNPO | protein_coding | 1.94510155 | 18.3605923 |
| IL17RC | protein_coding | 1.94479148 | 4.50753329 |
| TGIF2 | protein_coding | 1.94183244 | 12.2563679 |
| LURAP1L | protein_coding | 1.9365089 | 13.1181472 |
| CYP2J2 | protein_coding | 1.93499951 | 13.3328331 |
| RP1-288H2.2 | lncRNA | 1.93103517 | 7.83458163 |
| LIPG | protein_coding | 1.93050427 | 7.46773226 |
| IQSEC1 | protein_coding | 1.92955862 | 16.7046351 |
| CCNA1 | protein_coding | 1.92804905 | 8.99972816 |
| FGF21 | protein_coding | 1.92776887 | 5.70587462 |
| PRKCG | protein_coding | 1.92735632 | 11.1905103 |

| | | | |
|---|---|---|---|
| RP11-215D1( | lncRNA | 1.90954466 | 3.70140905 |
| VAT1L | protein_coding | 1.90880056 | 3.76414457 |
| GDF15 | protein_coding | 1.90872108 | 3.47076446 |
| RDH12 | protein_coding | 1.90846568 | 4.97072228 |
| CDC42EP4 | protein_coding | 1.90629513 | 18.8981471 |
| SLC6A19 | protein_coding | 1.89841759 | 5.12119085 |
| CREG1 | protein_coding | 1.89665864 | 20.4645901 |
| ETV6 | protein_coding | 1.89210545 | 17.2582992 |
| EIF2AK2 | protein_coding | 1.89195861 | 20.8595846 |
| PAXIP1-AS2 | lncRNA | 1.8827857 | 15.7198143 |
| P2RX7 | protein_coding | 1.86987734 | 2.62468794 |
| MYLK | protein_coding | 1.86634625 | 12.4395709 |
| THCAT155 | lncRNA | 1.86367584 | 3.64530468 |
| MYO3B | protein_coding | 1.86353754 | 7.10742752 |
| NMI | protein_coding | 1.86199053 | 17.9788659 |
| LINC00514 | protein_coding | 1.85360408 | 11.5670032 |
| HACD4 | protein_coding | 1.85258991 | 11.6719431 |
| PROM1 | protein_coding | 1.84919075 | 5.08448713 |
| **SQSTM1** | protein_coding | 1.84397922 | 20.3742323 |
| APOL2 | protein_coding | 1.84270845 | 18.661974 |
| HDX | protein_coding | 1.84242187 | 8.7186238 |
| ARHGAP6 | protein_coding | 1.82900635 | 4.70758822 |
| CACNG5 | protein_coding | 1.8260463 | 8.71893647 |
| MTMR11 | protein_coding | 1.82357709 | 3.56230663 |
| IGDCC4 | protein_coding | 1.81936831 | 9.62480816 |
| PPFIBP2 | protein_coding | 1.81721543 | 17.101878 |
| SMARCD3 | protein_coding | 1.8145404 | 4.32504457 |
| POPDC3 | protein_coding | 1.81007629 | 5.34700833 |
| LY6E-DT | lncRNA | 1.80797595 | 8.00818342 |
| LCP1 | protein_coding | 1.80634184 | 10.0881152 |
| JAM2 | protein_coding | 1.80329689 | 5.65021998 |
| PTAFR | protein_coding | 1.80251206 | 5.44805474 |
| RP11-513I15 | lncRNA | 1.80197812 | 7.23080771 |
| PRSS12 | protein_coding | 1.80026005 | 11.8319218 |
| IL11RA | protein_coding | 1.79940185 | 14.3381667 |
| SLC1A2 | protein_coding | 1.79884754 | 6.55490336 |
| AASS | protein_coding | 1.79654473 | 5.7008446 |
| BMP2 | protein_coding | 1.79583896 | 6.21774738 |
| RP11-468E2. | lncRNA | 1.79542042 | 4.97943592 |
| RP11-301L8.: | lncRNA | 1.7946141 | 5.06123097 |
| NPL | protein_coding | 1.79340903 | 12.9804843 |
| EFNA1 | protein_coding | 1.79279087 | 19.4418078 |
| TP73-AS3 | lncRNA | 1.79237522 | 5.85611535 |
| RP11-266O8. | lncRNA | 1.79046586 | 7.69883176 |
| SPATA1 | protein_coding | 1.79007039 | 3.54606653 |

| | | | |
|---|---|---|---|
| BARX2 | protein_coding | 1.7899683 | 3.21141948 |
| PIRT | protein_coding | 1.78990689 | 3.06065374 |
| RP11-439E19 | lncRNA | 1.78490333 | 5.43560414 |
| NIBAN1 | protein_coding | 1.78241279 | 16.667656 |
| RP11-541G9. | lncRNA | 1.78164387 | 3.16806688 |
| HSD17B14 | protein_coding | 1.7806522 | 4.86366842 |
| MFSD6L | protein_coding | 1.78016834 | 4.73627731 |
| BEND6 | protein_coding | 1.77868364 | 4.20941266 |
| RHCG | protein_coding | 1.77724335 | 4.24951128 |
| LINC01099 | lncRNA | 1.77570963 | 2.94396484 |
| SCNN1B | protein_coding | 1.77531942 | 10.1701711 |
| TMEM150C | protein_coding | 1.77522985 | 12.3692281 |
| RP11-92C13. | lncRNA | 1.76521423 | 3.14083228 |
| LINC00511 | lncRNA | 1.76174237 | 12.1611333 |
| TRAF3 | protein_coding | 1.76045085 | 19.5956256 |
| AC107218.3 | lncRNA | 1.75926544 | 8.2909277 |
| NRTN | protein_coding | 1.75555638 | 9.36487747 |
| HS3ST1 | protein_coding | 1.754774 | 6.80555067 |
| RHOV | protein_coding | 1.7533799 | 8.0051992 |
| SLC7A8 | protein_coding | 1.74637271 | 9.24345019 |
| EDN1 | protein_coding | 1.74625639 | 3.72172753 |
| PML | protein_coding | 1.73860577 | 17.8129751 |
| MOV10L1 | protein_coding | 1.73672627 | 5.36490677 |
| NFKBID | protein_coding | 1.73358926 | 13.3723985 |
| RP11-5C23.2 | lncRNA | 1.73111752 | 4.48860022 |
| RERG-AS1 | lncRNA | 1.72955668 | 4.28778609 |
| FBXO40 | protein_coding | 1.7286595 | 3.26223887 |
| PEX11G | protein_coding | 1.7273035 | 9.93349593 |
| EMP3 | protein_coding | 1.71975871 | 12.8316687 |
| SEMA3C | protein_coding | 1.71802527 | 17.5648303 |
| MICAL2 | protein_coding | 1.71661114 | 14.9890279 |
| H2BC18 | protein_coding | 1.71638452 | 4.38072938 |
| ANKRD33B | protein_coding | 1.71188963 | 18.162143 |
| CAMK2A | protein_coding | 1.71177514 | 9.93288258 |
| LY6E | protein_coding | 1.7049574 | 19.9952175 |
| PLEKHG1 | protein_coding | 1.70196318 | 9.1467597 |
| SHISA2 | protein_coding | 1.70101074 | 15.3058891 |
| KIAA1522 | protein_coding | 1.70019099 | 17.1446907 |
| SLC2A3 | protein_coding | 1.69898543 | 7.04943191 |
| RP11-423H2. | lncRNA | 1.69713163 | 10.457083 |
| SERPINF1 | protein_coding | 1.69691527 | 6.2219819 |
| COLCA2 | protein_coding | 1.69197813 | 9.76892545 |
| TMEM108-A: | lncRNA | 1.6910006 | 3.49691147 |
| B3GNT5 | protein_coding | 1.69096418 | 3.79317989 |
| RAB19 | protein_coding | 1.68960137 | 4.2822921 |

| | | | |
|---|---|---|---|
| RP11-1069G[illegible] | lncRNA | 1.68920404 | 4.59075906 |
| TMEM229B | protein_coding | 1.68864079 | 3.99004902 |
| JADE2 | protein_coding | 1.68289279 | 12.623791 |
| RP11-115H1[illegible] | lncRNA | 1.68279716 | 10.4972526 |
| APOL4 | protein_coding | 1.68074658 | 6.24208108 |
| HHIPL2 | protein_coding | 1.67882814 | 6.36387971 |
| RP11-96L14.[illegible] | lncRNA | 1.67796091 | 7.97239387 |
| SCLY | protein_coding | 1.67626885 | 11.1523403 |
| IRF2 | protein_coding | 1.67512995 | 13.8495791 |
| LINC02328 | lncRNA | 1.6741239 | 4.37676397 |
| EGF | protein_coding | 1.67226313 | 11.1888375 |
| TRIM47 | protein_coding | 1.66918928 | 14.3874487 |
| DIO3OS | lncRNA | 1.66381898 | 4.78641069 |
| TRIM25 | protein_coding | 1.6619889 | 19.546481 |
| **CDKN1A** | protein_coding | 1.65671528 | 19.2551279 |
| WNT5B | protein_coding | 1.65453379 | 4.03011088 |
| ALDH4A1 | protein_coding | 1.65154138 | 15.4506866 |
| RRAS | protein_coding | 1.64955047 | 6.52929327 |
| ERAP1 | protein_coding | 1.64889424 | 18.1789072 |
| TMC6 | protein_coding | 1.6444784 | 8.07802468 |
| SERPINB6 | protein_coding | 1.64211564 | 3.79877999 |
| TMEM40 | protein_coding | 1.64000927 | 12.5180354 |
| NOTCH4 | protein_coding | 1.63841931 | 6.81331115 |
| TGFB3 | protein_coding | 1.63778734 | 9.30020301 |
| FHAD1 | protein_coding | 1.6361822 | 6.68497703 |
| AC009133.1[illegible] | lncRNA | 1.63528578 | 7.19944731 |
| RASSF4 | protein_coding | 1.6348686 | 8.1930005 |
| BBC3 | protein_coding | 1.63013471 | 9.89754136 |
| SH3KBP1 | protein_coding | 1.62963301 | 14.2835168 |
| ARHGAP26 | protein_coding | 1.62568138 | 15.8601537 |
| DUSP16 | protein_coding | 1.62512536 | 15.9749768 |
| CYFIP1 | protein_coding | 1.6204684 | 9.97591442 |
| CLDN6 | protein_coding | 1.62039945 | 4.78798043 |
| MST1R | protein_coding | 1.61995319 | 13.1506764 |
| COLEC12 | protein_coding | 1.61531753 | 4.06769444 |
| NUAK2 | protein_coding | 1.61482269 | 14.5211801 |
| FGF18 | protein_coding | 1.61293913 | 4.23165473 |
| CRACDL | protein_coding | 1.6080802 | 11.8832518 |
| RAPGEFL1 | protein_coding | 1.60258679 | 17.7366795 |
| KRT17 | protein_coding | 1.60171373 | 15.5895375 |
| NCKAP5-AS2 | lncRNA | 1.59982358 | 4.28724053 |
| ADAMTS14 | protein_coding | 1.59930326 | 11.8666642 |
| RNASEL | protein_coding | 1.59681606 | 6.32534532 |
| TMEM150A | protein_coding | 1.59607781 | 3.18755994 |
| DSCR9 | lncRNA | 1.59315149 | 3.49554185 |

| | | | |
|---|---|---|---|
| FAM83F | protein_coding | 1.59157127 | 9.00501138 |
| SYT2 | protein_coding | 1.5901993 | 12.2495403 |
| AGBL2 | protein_coding | 1.58830806 | 4.85348053 |
| PTGIR | protein_coding | 1.58691015 | 12.7063541 |
| EMILIN2 | protein_coding | 1.58660718 | 8.3312352 |
| LITAF | protein_coding | 1.58505373 | 17.4401982 |
| RP11-259O2. | lncRNA | 1.58367692 | 5.56273625 |
| CTB-113D17. | lncRNA | 1.58210608 | 2.6928834 |
| PNRC1 | protein_coding | 1.58180269 | 15.5761742 |
| RP11-807C2( | protein_coding | 1.57838788 | 6.74955025 |
| ZNF516 | protein_coding | 1.5778662 | 4.89747927 |
| DUSP8 | protein_coding | 1.57656702 | 15.2230742 |
| PYY | protein_coding | 1.57485854 | 3.49826527 |
| RBFOX3 | protein_coding | 1.57252748 | 6.12955005 |
| BNC2 | protein_coding | 1.57240486 | 7.22362827 |
| AEBP1 | protein_coding | 1.5710737 | 2.48542172 |
| FRMPD3 | protein_coding | 1.56886143 | 14.4022826 |
| RP4-794I6.4 | lncRNA | 1.56875693 | 5.78135005 |
| AL132709.1 | lncRNA | 1.56826057 | 6.91631213 |
| FGF17 | protein_coding | 1.56444874 | 6.68861233 |
| RP1-90J20.1! | lncRNA | 1.56211261 | 3.01194235 |
| ADAR | protein_coding | 1.56143758 | 22.0666016 |
| MYD88 | protein_coding | 1.56093363 | 13.4644492 |
| NES | protein_coding | 1.5580894 | 11.6509534 |
| KRTAP5-AS1 | lncRNA | 1.55276944 | 7.7965749 |
| LINC00930 | lncRNA | 1.55139201 | 3.92422905 |
| PRKG1 | protein_coding | 1.55075366 | 8.43766052 |
| STAT2 | protein_coding | 1.54817229 | 18.2409539 |
| PHLDB2 | protein_coding | 1.54702292 | 12.6624694 |
| TRIL | protein_coding | 1.5467844 | 7.98229488 |
| ELFN1 | protein_coding | 1.54632066 | 3.17255306 |
| ALLC | protein_coding | 1.54540598 | 2.59147446 |
| CCDC28B | protein_coding | 1.54246287 | 16.2645589 |
| WDFY4 | protein_coding | 1.53714149 | 3.65507956 |
| NID1 | protein_coding | 1.53700759 | 13.7161648 |
| SLC9A3R2 | protein_coding | 1.53335765 | 14.629279 |
| HOXC13 | protein_coding | 1.53180662 | 6.22548368 |
| CCDC159 | protein_coding | 1.53110323 | 8.24037758 |
| CPNE6 | protein_coding | 1.52945727 | 11.2421316 |
| CTD-2308L22 | lncRNA | 1.52763218 | 4.09944855 |
| GOLGA7B | protein_coding | 1.52701225 | 9.24860221 |
| MIIP | protein_coding | 1.52595337 | 12.6429363 |
| STX4 | protein_coding | 1.52028975 | 14.5185604 |
| RP11-624C23 | lncRNA | 1.51703947 | 3.11421053 |
| SLC7A11 | protein_coding | 1.51674753 | 16.7456021 |

| | | | |
|---|---|---|---|
| RP11-1252I4 | lncRNA | 1.51501487 | 3.03808116 |
| PPP1R15A | protein_coding | 1.50989353 | 15.0958121 |
| ATF3 | protein_coding | 1.50801833 | 14.3377738 |
| SERTAD2 | protein_coding | 1.50782655 | 13.5421123 |
| AHNAK | protein_coding | 1.50560734 | 17.7566584 |
| DCDC2 | protein_coding | 1.50402346 | 6.64117795 |
| NTRK1 | protein_coding | 1.50184179 | 9.46867875 |
| PLOD3 | protein_coding | 1.50165267 | 13.8186314 |
| LINC00092 | lncRNA | 1.50130444 | 3.12474513 |
| H2AC6 | protein_coding | 1.49962302 | 9.24259056 |
| BGN | protein_coding | 1.49947845 | 10.5518285 |
| UNC5B | protein_coding | 1.49859157 | 15.7727756 |
| ICAM5 | protein_coding | 1.49754852 | 9.16225104 |
| CD82 | protein_coding | 1.49552347 | 12.3477825 |
| SLC6A7 | protein_coding | 1.49532752 | 7.45053387 |
| HEATR4 | protein_coding | 1.49224108 | 4.34496849 |
| LOXL2 | protein_coding | 1.49216507 | 4.37790198 |
| RELN | protein_coding | 1.49137842 | 19.280584 |
| DMD | protein_coding | 1.49070594 | 14.294244 |
| SLC17A6 | protein_coding | 1.49038694 | 13.3150909 |
| ATF5 | protein_coding | 1.48805057 | 20.6428728 |
| GAS6 | protein_coding | 1.48786263 | 15.0610395 |
| DOC2B | protein_coding | 1.48731828 | 16.5148405 |
| RRH | protein_coding | 1.48618554 | 2.39894357 |
| EPS8L2 | protein_coding | 1.4854173 | 10.6454181 |
| TDRD7 | protein_coding | 1.48423933 | 18.0225677 |
| BCL9L | protein_coding | 1.47915867 | 14.1754076 |
| CRYZ | protein_coding | 1.47790443 | 12.0932665 |
| CEBPB | protein_coding | 1.47635005 | 9.02533972 |
| ZNFX1 | protein_coding | 1.47487132 | 18.3905175 |
| RYR2 | protein_coding | 1.47298195 | 8.91835015 |
| MERTK | protein_coding | 1.47123728 | 6.35177384 |
| LA16c-306A4 | lncRNA | 1.46991636 | 3.48974003 |
| SCG5 | protein_coding | 1.46825449 | 18.0414716 |
| TTC30B | protein_coding | 1.46813932 | 11.4398004 |
| LGALS3 | protein_coding | 1.46486504 | 4.42579935 |
| RP11-572M1 | lncRNA | 1.46380184 | 2.33279292 |
| NQO1 | protein_coding | 1.46316775 | 14.5224001 |
| BEST1 | protein_coding | 1.45779646 | 8.7759765 |
| CTD-3088G3. | protein_coding | 1.45740782 | 3.83549198 |
| DNAI2 | protein_coding | 1.45587863 | 3.19225811 |
| PTPRB | protein_coding | 1.44991012 | 8.01621873 |
| UST | protein_coding | 1.44707494 | 11.9898943 |
| TAMALIN | protein_coding | 1.44507432 | 14.6828807 |
| NOXRED1 | protein_coding | 1.44354053 | 2.92277367 |

| | | | |
|---|---|---|---|
| NRP1 | protein_coding | 1.43861424 | 13.9118107 |
| TTC39A | protein_coding | 1.43469085 | 13.3739467 |
| L3MBTL2-AS | lncRNA | 1.43464762 | 8.07214873 |
| PEA15 | protein_coding | 1.43185382 | 11.6055735 |
| PTK2B | protein_coding | 1.42791402 | 13.6575817 |
| LINC02586 | lncRNA | 1.42704555 | 5.92291713 |
| GALNT6 | protein_coding | 1.42620296 | 11.7881754 |
| PIWIL4 | protein_coding | 1.42596448 | 6.19946937 |
| GSN | protein_coding | 1.42226673 | 17.4129724 |
| TGFA | protein_coding | 1.42183403 | 3.79736552 |
| PID1 | protein_coding | 1.42101263 | 3.45899083 |
| CTD-2002H8. | lncRNA | 1.42038195 | 2.49611887 |
| SEC14L5 | protein_coding | 1.41981687 | 9.94796798 |
| TBL1Y | protein_coding | 1.41922371 | 7.89873711 |
| ATP8B4 | protein_coding | 1.41819282 | 7.28112157 |
| PARP8 | protein_coding | 1.41611804 | 12.3015217 |
| CD8A | protein_coding | 1.41394019 | 4.19171564 |
| RP11-567M1 | lncRNA | 1.412437 | 5.03354541 |
| HAPLN3 | protein_coding | 1.40858794 | 5.5573304 |
| GRINA | protein_coding | 1.40830674 | 15.5137268 |
| ALDH2 | protein_coding | 1.40793623 | 13.2782264 |
| DDR1 | protein_coding | 1.40645782 | 19.64932 |
| MT2A | protein_coding | 1.40339237 | 14.9610862 |
| RP11-174G6. | lncRNA | 1.3983471 | 5.87853706 |
| IL15RA | protein_coding | 1.39775939 | 2.72394883 |
| INPP4A | protein_coding | 1.39771069 | 16.761275 |
| ANXA2 | protein_coding | 1.39669644 | 10.1215164 |
| SCARF1 | protein_coding | 1.39431754 | 2.77468594 |
| PALM2AKAP2 | protein_coding | 1.39190186 | 14.1528611 |
| CABP1 | protein_coding | 1.39023977 | 9.76291712 |
| FLOT2 | protein_coding | 1.3882338 | 17.2547836 |
| GDPD5 | protein_coding | 1.38667418 | 12.5928139 |
| NINJ1 | protein_coding | 1.38035979 | 14.0205618 |
| RUNX1 | protein_coding | 1.37999184 | 6.87076813 |
| DGKG | protein_coding | 1.37912198 | 3.63185186 |
| FBLN1 | protein_coding | 1.37839871 | 14.5712068 |
| VDR | protein_coding | 1.37611266 | 15.0828503 |
| BBX | protein_coding | 1.37511705 | 12.0642734 |
| INHBE | protein_coding | 1.37371346 | 10.7183734 |
| CRYZL2P-SEC | lncRNA | 1.37198519 | 2.78352883 |
| GPX2 | protein_coding | 1.37185825 | 7.13031878 |
| EGR1 | protein_coding | 1.37183429 | 5.00633745 |
| NTSR1 | protein_coding | 1.37006848 | 6.57036698 |
| DAPK1 | protein_coding | 1.36935283 | 9.42986452 |
| IGSF9 | protein_coding | 1.36587227 | 15.3169768 |

| | | | |
|---|---|---|---|
| ARFGEF1 | protein_coding | 1.35824838 | 14.7708041 |
| AC002057.2 | lncRNA | 1.35649233 | 3.66541198 |
| VASN | protein_coding | 1.35380622 | 5.45559416 |
| PNPT1 | protein_coding | 1.34849904 | 17.8603666 |
| DEPP1 | protein_coding | 1.34730283 | 11.8893662 |
| SLC47A2 | protein_coding | 1.34588904 | 3.85561716 |
| CFLAR | protein_coding | 1.34387338 | 16.6594356 |
| PEAK1 | protein_coding | 1.34325125 | 16.7677893 |
| NIBAN3 | protein_coding | 1.34297262 | 9.11723255 |
| PCDHB8 | protein_coding | 1.34156145 | 5.80394685 |
| JDP2 | protein_coding | 1.34114977 | 6.88711291 |
| DCT | protein_coding | 1.33507241 | 2.92464749 |
| PHYHD1 | protein_coding | 1.33456597 | 3.31252136 |
| ICA1 | protein_coding | 1.33451061 | 12.7107252 |
| GALT | protein_coding | 1.33022055 | 11.6390739 |
| PMAIP1 | protein_coding | 1.32887931 | 14.9146656 |
| CCDC102A | protein_coding | 1.32783214 | 2.31339388 |
| RP11-416N4. | lncRNA | 1.32689253 | 2.28462993 |
| SPATA32 | protein_coding | 1.32685307 | 3.12356451 |
| ALCAM | protein_coding | 1.32635035 | 16.805757 |
| RP1-102E24. | lncRNA | 1.32502399 | 3.80666767 |
| XCL1 | protein_coding | 1.32398574 | 6.3234312 |
| GABRQ | protein_coding | 1.32222533 | 13.7336481 |
| NFIA-AS2 | lncRNA | 1.31999344 | 3.94845755 |
| PILRA | protein_coding | 1.31928421 | 5.30218494 |
| LTBP1 | protein_coding | 1.31758394 | 11.9533978 |
| GRID2 | protein_coding | 1.31665637 | 8.60291925 |
| RP11-511P7. | protein_coding | 1.31467262 | 10.6896702 |
| SH2B2 | protein_coding | 1.31320952 | 11.1382505 |
| TMSB4X | protein_coding | 1.31143267 | 16.3644513 |
| GPR155 | protein_coding | 1.30726771 | 7.93563488 |
| PPP1R13B | protein_coding | 1.30471094 | 15.46381 |
| SYTL1 | protein_coding | 1.30345374 | 8.44799726 |
| FAM131C | protein_coding | 1.30030965 | 7.4865311 |
| NOXO1 | protein_coding | 1.29921657 | 4.06809002 |
| MAFB | protein_coding | 1.29813257 | 13.8058622 |
| SLCO4C1 | protein_coding | 1.29667073 | 5.90010472 |
| DENND2B | protein_coding | 1.29583961 | 15.364717 |
| CLDN3 | protein_coding | 1.29523405 | 7.17000428 |
| N4BP2L1 | protein_coding | 1.29217333 | 10.1993844 |
| TRIM67 | protein_coding | 1.29211905 | 8.10939004 |
| PCDHB7 | protein_coding | 1.29049049 | 2.64099015 |
| LINC02527 | lncRNA | 1.28981874 | 4.60854045 |
| MED14OS | lncRNA | 1.28729632 | 2.06181566 |
| SIX4 | protein_coding | 1.28672064 | 3.52282273 |

| | | | |
|---|---|---|---|
| DNAH10 | protein_coding | 1.28647204 | 3.67644847 |
| MATN1-AS1 | lncRNA | 1.28625942 | 8.67747527 |
| WNT9A | protein_coding | 1.28590759 | 4.85561379 |
| LINC02885 | lncRNA | 1.27958854 | 2.04204331 |
| RP11-305F14 | lncRNA | 1.27842082 | 2.66333467 |
| AC007566.10 | lncRNA | 1.27803107 | 3.74420241 |
| CTD-2516F10 | lncRNA | 1.27717354 | 7.45401887 |
| HSPG2 | protein_coding | 1.2759272 | 12.4609161 |
| EHD1 | protein_coding | 1.27447231 | 13.9117966 |
| GALNT16 | protein_coding | 1.27155234 | 11.4902834 |
| BIRC2 | protein_coding | 1.26684947 | 12.8131208 |
| MORN3 | protein_coding | 1.2665662 | 7.94842438 |
| SUOX | protein_coding | 1.26501992 | 8.81812029 |
| VANGL2 | protein_coding | 1.26497541 | 16.9713815 |
| RP11-379L18 | lncRNA | 1.26492362 | 4.43851735 |
| SETD7 | protein_coding | 1.26457875 | 13.7770115 |
| KCNQ3 | protein_coding | 1.26154953 | 17.8747057 |
| RP11-111H13 | protein_coding | 1.26003739 | 1.71017391 |
| ABCA1 | protein_coding | 1.25835935 | 12.966294 |
| TRIM16 | protein_coding | 1.25818417 | 10.9369935 |
| TPST1 | protein_coding | 1.25769839 | 11.2770754 |
| CASR | protein_coding | 1.25569807 | 2.63359881 |
| TENT5B | protein_coding | 1.25532418 | 3.39452082 |
| RP11-89H19. | lncRNA | 1.25418322 | 3.0790152 |
| AC093326.3 | lncRNA | 1.25403483 | 1.62868196 |
| JUNB | protein_coding | 1.25161973 | 12.2983722 |
| GMPR | protein_coding | 1.25084799 | 8.19948289 |
| UNC93B1 | protein_coding | 1.25039908 | 11.1838496 |
| PHTF1 | protein_coding | 1.24843872 | 10.4600658 |
| H4-16 | protein_coding | 1.24782707 | 3.67683177 |
| HIP1 | protein_coding | 1.24573299 | 18.0781538 |
| ROM1 | protein_coding | 1.24414271 | 8.79832202 |
| ACTR3C | protein_coding | 1.24054766 | 5.71132797 |
| SEMA3B | protein_coding | 1.24009308 | 3.54628728 |
| PSME1 | protein_coding | 1.23966319 | 18.3625105 |
| RP11-85O21. | lncRNA | 1.23903981 | 4.00992497 |
| ZBED3 | protein_coding | 1.23864327 | 5.02077936 |
| DOCK4 | protein_coding | 1.23710028 | 13.7399432 |
| MLKL | protein_coding | 1.23620021 | 8.81338672 |
| MEAK7 | protein_coding | 1.2354566 | 9.42523731 |
| ZNF502 | protein_coding | 1.233974 | 4.61625817 |
| THEMIS2 | protein_coding | 1.23390619 | 7.87730469 |
| CLEC2D | protein_coding | 1.23279936 | 5.41037724 |
| ZMAT1 | protein_coding | 1.23262555 | 2.69107991 |
| SLC26A7 | protein_coding | 1.23218781 | 10.3892455 |

| | | | |
|---|---|---|---|
| SYN1 | protein_coding | 1.2306473 | 11.7072819 |
| LOXL1 | protein_coding | 1.23032336 | 11.0126058 |
| NFKBIE | protein_coding | 1.22966375 | 11.2219915 |
| SLC35F2 | protein_coding | 1.2284378 | 9.20452918 |
| GRHL3 | protein_coding | 1.22141244 | 2.61566769 |
| RNF31 | protein_coding | 1.22034067 | 3.8947019 |
| DGKQ | protein_coding | 1.21935567 | 13.6385624 |
| SLC16A9 | protein_coding | 1.21931327 | 10.1054084 |
| AKNA | protein_coding | 1.2191564 | 15.0818532 |
| EYA4 | protein_coding | 1.21903454 | 4.6976512 |
| STX17-AS1 | lncRNA | 1.21821963 | 1.91508981 |
| CAMKK1 | protein_coding | 1.21596791 | 12.7606073 |
| KIF19 | protein_coding | 1.21416251 | 15.6428011 |
| SEMA6D | protein_coding | 1.20916673 | 10.3270883 |
| DDIT3 | protein_coding | 1.20826744 | 14.3994886 |
| TMEM266 | protein_coding | 1.20729928 | 8.96078851 |
| MTUS1 | protein_coding | 1.20711092 | 13.4694511 |
| LSAMP | protein_coding | 1.20569401 | 4.19140586 |
| ROBO1 | protein_coding | 1.20458674 | 15.404263 |
| HOXA10-AS | lncRNA | 1.20386336 | 9.67237491 |
| ANXA1 | protein_coding | 1.20384723 | 7.95818107 |
| RFX8 | protein_coding | 1.20273701 | 11.2433375 |
| FUT2 | protein_coding | 1.20222601 | 8.48761109 |
| RP1-298M8.[illegible] | lncRNA | 1.20167114 | 4.29222107 |
| CORO2B | protein_coding | 1.20140675 | 5.09868396 |
| FZD5 | protein_coding | 1.20043162 | 9.83210955 |
| KRT9 | protein_coding | 1.19987468 | 13.9156193 |
| PPP1R14D | protein_coding | 1.19928062 | 2.30309495 |
| CDH1 | protein_coding | 1.19916945 | 13.6180296 |
| MAT1A | protein_coding | 1.19907243 | 2.96513498 |
| INSC | protein_coding | 1.19588295 | 4.41367678 |
| GPT | protein_coding | 1.1944526 | 3.05507213 |
| CILP | protein_coding | 1.19396817 | 2.8561155 |
| SESN2 | protein_coding | 1.19381193 | 12.636718 |
| MBNL2 | protein_coding | 1.19358256 | 16.3745718 |
| ILDR1 | protein_coding | 1.19223158 | 2.77763807 |
| OCA2 | protein_coding | 1.18868273 | 5.98901998 |
| LCN15 | protein_coding | 1.18788934 | 3.94332619 |
| MCF2 | protein_coding | 1.1874794 | 1.97346026 |
| H3C6 | protein_coding | 1.18688736 | 3.94684675 |
| FGD2 | protein_coding | 1.1864192 | 2.29906931 |
| LRFN2 | protein_coding | 1.18563252 | 2.71487532 |
| DTX1 | protein_coding | 1.18484788 | 4.67719163 |
| LDLRAD3 | protein_coding | 1.18471515 | 16.6979886 |
| CYB561D2 | lncRNA | 1.18298501 | 1.84487111 |

| | | | |
|---|---|---|---|
| AP001062.7 | lncRNA | 1.18102373 | 11.1359287 |
| RP11-736N1 | lncRNA | 1.1792078 | 3.80328287 |
| PIPOX | protein_coding | 1.17869588 | 3.83402086 |
| CTD-2014B1 | lncRNA | 1.17753756 | 2.64123629 |
| LY75 | protein_coding | 1.17745422 | 4.57462686 |
| BID | protein_coding | 1.17552739 | 13.9137474 |
| SNX33 | protein_coding | 1.17500504 | 8.96178647 |
| IL27RA | protein_coding | 1.17406422 | 9.38001642 |
| SOD2 | protein_coding | 1.17254816 | 17.3200321 |
| LINC00052 | lncRNA | 1.17246258 | 4.19479589 |
| CLK4 | protein_coding | 1.17119511 | 10.9273358 |
| DNAH7 | protein_coding | 1.16936945 | 3.59018025 |
| TMEM92 | protein_coding | 1.16920912 | 2.00147461 |
| NUTM2D | protein_coding | 1.16870457 | 5.01830841 |
| RD3 | protein_coding | 1.16686346 | 6.20408799 |
| GPR85 | protein_coding | 1.16578423 | 2.905524 |
| FAT3 | protein_coding | 1.16506925 | 2.91431299 |
| REL | protein_coding | 1.16476966 | 12.6479958 |
| RP6-109B7.2 | lncRNA | 1.16269476 | 2.00919052 |
| AC004538.3 | lncRNA | 1.16268865 | 2.23841826 |
| PFKM | protein_coding | 1.16257373 | 10.5448549 |
| CPNE5 | protein_coding | 1.16045353 | 6.04967415 |
| GS1-57L11.3 | lncRNA | 1.15915072 | 2.07971494 |
| PAX6 | protein_coding | 1.15841724 | 15.6172809 |
| SMTNL2 | protein_coding | 1.15774892 | 4.48344073 |
| PNPLA1 | protein_coding | 1.15560461 | 2.86841725 |
| LOXL4 | protein_coding | 1.15481604 | 3.90583046 |
| C1S | protein_coding | 1.15426312 | 6.61769529 |
| RP11-16N10 | protein_coding | 1.15353376 | 1.84979558 |
| CD63 | protein_coding | 1.15292858 | 17.0910805 |
| SYNPO2 | protein_coding | 1.14961296 | 8.17459232 |
| IL22RA1 | protein_coding | 1.14883762 | 8.9433864 |
| CDC14A | protein_coding | 1.14880294 | 10.2654346 |
| PCDHAC1 | protein_coding | 1.14682741 | 7.11884366 |
| COL1A1 | protein_coding | 1.14635456 | 9.2259365 |
| PNKD | protein_coding | 1.1458886 | 14.8500335 |
| ANXA4 | protein_coding | 1.14555228 | 8.61600267 |
| EFNB2 | protein_coding | 1.14529899 | 12.7848712 |
| GRIA4 | protein_coding | 1.1441148 | 12.5298149 |
| PPP1R36 | protein_coding | 1.13990952 | 2.29980486 |
| LINC01278 | lncRNA | 1.13935441 | 7.47078252 |
| DNPEP | protein_coding | 1.13924241 | 16.1000982 |
| TRANK1 | protein_coding | 1.13837876 | 4.79644273 |
| PKHD1L1 | protein_coding | 1.13738016 | 6.7451328 |
| H2BC5 | protein_coding | 1.13310428 | 8.87261302 |

| | | | |
|---|---|---|---|
| TMSB4Y | protein_coding | 1.13281042 | 3.06729819 |
| GGACT | protein_coding | 1.13213159 | 4.26154383 |
| ZNF600 | protein_coding | 1.13151917 | 4.35555943 |
| SPARCL1 | protein_coding | 1.13015052 | 5.17060506 |
| RP11-526I2.5 | lncRNA | 1.12980224 | 3.17839847 |
| RAB30 | protein_coding | 1.1295141 | 10.9451353 |
| CDHR3 | protein_coding | 1.12872777 | 10.6965298 |
| TPO | protein_coding | 1.1251511 | 4.21468085 |
| JSRP1 | protein_coding | 1.12267963 | 2.18141958 |
| LACC1 | protein_coding | 1.1225698 | 4.5603929 |
| RP11-400F19 | lncRNA | 1.1224396 | 5.66833386 |
| RP11-143K11 | lncRNA | 1.12235972 | 2.03172579 |
| NECTIN1 | protein_coding | 1.1223256 | 17.5915422 |
| GRK3 | protein_coding | 1.1220007 | 12.2743258 |
| GCNT2 | protein_coding | 1.12027292 | 13.1757227 |
| GPRC5A | protein_coding | 1.11973321 | 4.14127574 |
| FBXL2 | protein_coding | 1.11775948 | 1.92475822 |
| H2BC8 | protein_coding | 1.11772326 | 5.16320137 |
| RP11-1105G | lncRNA | 1.11631465 | 2.35171509 |
| STUM | protein_coding | 1.11424813 | 15.8700451 |
| SCN11A | protein_coding | 1.114239 | 2.88319606 |
| PXYLP1 | protein_coding | 1.11297719 | 12.2751734 |
| RNASEK | protein_coding | 1.11180617 | 3.29438304 |
| SLC25A22 | protein_coding | 1.11058919 | 14.9160624 |
| FAM222A | protein_coding | 1.11054025 | 11.6935765 |
| KIRREL2 | protein_coding | 1.10944433 | 13.05999 |
| NUB1 | protein_coding | 1.10803367 | 15.0632485 |
| ESPN | protein_coding | 1.10770244 | 10.9113618 |
| CASP8 | protein_coding | 1.1042587 | 10.9346433 |
| NECTIN2 | protein_coding | 1.10262981 | 8.30278234 |
| PPP4R4 | protein_coding | 1.1015327 | 13.628226 |
| TFE3 | protein_coding | 1.09974267 | 14.7442661 |
| ARRB1 | protein_coding | 1.09889511 | 5.36890516 |
| RP11-429A24 | lncRNA | 1.09833053 | 2.79366654 |
| TCN2 | protein_coding | 1.0971387 | 4.52232252 |
| ARHGEF2 | protein_coding | 1.0967061 | 16.3605956 |
| PIM1 | protein_coding | 1.09559458 | 13.0302241 |
| RP5-820A21. | lncRNA | 1.09516524 | 2.03144763 |
| CHRM1 | protein_coding | 1.0939831 | 2.93807321 |
| UGGT2 | protein_coding | 1.09303338 | 5.96865766 |
| HVCN1 | protein_coding | 1.09298758 | 2.52601157 |
| PRDM9 | protein_coding | 1.09199553 | 2.38689798 |
| ECHDC2 | protein_coding | 1.08941777 | 4.65196144 |
| NUTM2B | protein_coding | 1.08820421 | 2.20248862 |
| HS3ST4 | protein_coding | 1.08784975 | 2.00108007 |

| | | | |
|---|---|---|---|
| ADM | protein_coding | 1.0856286 | 12.6246713 |
| SERPINB8 | protein_coding | 1.08496855 | 4.97892742 |
| ADAMTS4 | protein_coding | 1.0830793 | 12.5095835 |
| TENT5C | protein_coding | 1.08139382 | 3.51653221 |
| ZAP70 | protein_coding | 1.08066595 | 7.70366371 |
| SHISA5 | protein_coding | 1.08057241 | 15.3514811 |
| PDE4D | protein_coding | 1.0798816 | 9.07055577 |
| ETS2 | protein_coding | 1.07952599 | 17.0305069 |
| ASNS | protein_coding | 1.0793307 | 7.2215176 |
| GNG3 | protein_coding | 1.07800576 | 2.50475351 |
| CMTM3 | protein_coding | 1.07660389 | 9.1235778 |
| RP11-66N24. | lncRNA | 1.0757274 | 8.90189767 |
| STRA6 | protein_coding | 1.07460736 | 16.8338835 |
| COL25A1 | protein_coding | 1.07439399 | 12.309211 |
| AC005606.14 | lncRNA | 1.07435943 | 2.61599781 |
| NUTM2A | protein_coding | 1.07435256 | 3.06796103 |
| NMB | protein_coding | 1.07152793 | 9.51946848 |
| S100A6 | protein_coding | 1.07135545 | 12.1757567 |
| EPHX2 | protein_coding | 1.06998166 | 10.7672715 |
| PLXNA1 | protein_coding | 1.06850782 | 14.6130713 |
| RP11-20G13. | lncRNA | 1.06681906 | 4.07990993 |
| SLC9A7 | protein_coding | 1.06603997 | 13.7641667 |
| SLC6A5 | protein_coding | 1.06362384 | 2.54895447 |
| USP41 | protein_coding | 1.06238693 | 3.93092104 |
| VGF | protein_coding | 1.06236941 | 7.4121319 |
| TLCD1 | protein_coding | 1.06233412 | 11.7436787 |
| IL6ST | protein_coding | 1.0620188 | 10.5848198 |
| ZNF441 | protein_coding | 1.06076784 | 5.09265647 |
| SMOX | protein_coding | 1.05625411 | 4.02210997 |
| ZC3H7B | protein_coding | 1.05516112 | 17.7784375 |
| CYP4X1 | protein_coding | 1.05228176 | 1.94517478 |
| PCK2 | protein_coding | 1.05176771 | 14.1779544 |
| B4GALT6 | protein_coding | 1.0506532 | 11.0200339 |
| PRKD2 | protein_coding | 1.05025736 | 13.3523121 |
| STOM | protein_coding | 1.05006002 | 14.4006537 |
| RP11-817I4.2 | lncRNA | 1.05000973 | 1.63909042 |
| TRIB3 | protein_coding | 1.04990529 | 14.891169 |
| ORAI1 | protein_coding | 1.04938769 | 10.9276632 |
| MAN2B1 | protein_coding | 1.04910509 | 12.3712023 |
| B4GALNT1 | protein_coding | 1.04897429 | 15.5198984 |
| RP11-235E17 | lncRNA | 1.04874436 | 1.43039497 |
| EBLN2 | protein_coding | 1.04750272 | 1.78288522 |
| CRTAC1 | protein_coding | 1.04691533 | 15.7369154 |
| DSCAM | protein_coding | 1.04567484 | 12.1428149 |
| ZBTB10 | protein_coding | 1.04466417 | 13.7752568 |

| | | | |
|---|---|---|---|
| LINC01235 | lncRNA | 1.04460328 | 3.11829595 |
| TGM7 | protein_coding | 1.04350241 | 2.18645525 |
| ARHGAP45 | protein_coding | 1.04311332 | 7.68116194 |
| CTD-2587H24 | lncRNA | 1.0417575 | 3.07897117 |
| GBX2 | protein_coding | 1.04111056 | 6.04432633 |
| STARD4-AS1 | lncRNA | 1.04080202 | 4.40598976 |
| ADGRE5 | protein_coding | 1.03931888 | 2.5224615 |
| ZSCAN9 | protein_coding | 1.03889166 | 5.34844547 |
| LINC02754 | lncRNA | 1.03800734 | 2.9624067 |
| SNAP25-AS1 | lncRNA | 1.03614431 | 10.4306498 |
| RIPOR3 | protein_coding | 1.03486313 | 2.76301141 |
| PRODH | protein_coding | 1.03471238 | 9.92848623 |
| VSTM4 | protein_coding | 1.03433985 | 10.5542336 |
| BVES | protein_coding | 1.03427075 | 8.69430872 |
| NXPH1 | protein_coding | 1.03356252 | 3.05853473 |
| ZXDA | protein_coding | 1.03233284 | 3.84064188 |
| RP11-666A8. | lncRNA | 1.03093696 | 1.86481185 |
| CHAC1 | protein_coding | 1.02779285 | 13.4163532 |
| NBN | protein_coding | 1.02668434 | 15.4116343 |
| H2BC17 | protein_coding | 1.02626951 | 1.59493427 |
| TCTN1 | protein_coding | 1.02384834 | 9.44909484 |
| ARHGEF28 | protein_coding | 1.0218097 | 5.94183029 |
| CYP4V2 | protein_coding | 1.0212321 | 6.26760986 |
| SDCBP | protein_coding | 1.02015317 | 7.68259448 |
| RBMS3-AS3 | lncRNA | 1.01825727 | 1.80538623 |
| RBMS3 | protein_coding | 1.01803412 | 9.17148933 |
| SCML4 | protein_coding | 1.0155743 | 7.47823104 |
| SLC7A2 | protein_coding | 1.01535971 | 2.73833356 |
| BLVRB | protein_coding | 1.01478014 | 9.86423742 |
| RP3-382I10.7 | protein_coding | 1.01401888 | 1.70722995 |
| AJ011932.1 | lncRNA | 1.01376342 | 1.80732484 |
| CTC-479C5.1 | protein_coding | 1.01351734 | 3.49706002 |
| ATP7B | protein_coding | 1.01334952 | 9.0876357 |
| DCN | protein_coding | 1.01276167 | 5.24498988 |
| INPP1 | protein_coding | 1.01211947 | 14.0675485 |
| RP11-443G1 | lncRNA | 1.01177928 | 1.51477979 |
| LINC00472 | lncRNA | 1.01149312 | 9.31441557 |
| TRIB2 | protein_coding | 1.01093306 | 14.8313977 |
| ZFP36L1 | protein_coding | 1.01036555 | 8.03095998 |
| GSTM2 | protein_coding | 1.00908955 | 4.52200139 |
| COL5A1 | protein_coding | 1.00803121 | 5.78219078 |
| EYS | protein_coding | 1.00766529 | 4.73068162 |
| SLC41A2 | protein_coding | 1.00761665 | 9.26381598 |
| NAMPT | protein_coding | 1.00749014 | 12.2827975 |
| GLOD5 | protein_coding | 1.00633019 | 2.78613535 |

| | | | |
|---|---|---|---|
| RP11-303E16 | protein_coding | 1.00616074 | 2.6745704 |
| STX16-NPEPI | protein_coding | 1.00592042 | 2.32407168 |
| HPGD | protein_coding | 1.00542931 | 3.80136497 |
| KLF4 | protein_coding | 1.00394239 | 10.730485 |
| TEKT5 | protein_coding | 1.00251591 | 3.34065583 |
| FBXO2 | protein_coding | 1.00242361 | 1.90757533 |
| CD24 | protein_coding | 1.00174006 | 12.9797675 |
| SRPK1 | protein_coding | 1.00119304 | 17.814412 |
| SEPTIN4-AS1 | lncRNA | 1.00093213 | 2.69736756 |
| CSRP1 | protein_coding | 1.00072925 | 17.3302761 |
| LINC01238 | lncRNA | 1.00053043 | 5.82966243 |
| CDH18 | protein_coding | 1.00006051 | 7.59894676 |
| ZFHX2 | protein_coding | 1.00002169 | 9.51764318 |
| CTNNA2 | protein_coding | 0.99847227 | 7.45701837 |
| RP11-552F3. | lncRNA | 0.99727647 | 4.86867146 |
| TJP2 | protein_coding | 0.99351723 | 4.73179021 |
| PDE7B | protein_coding | 0.99247224 | 2.41538074 |
| RELT | protein_coding | 0.99088401 | 14.6455364 |
| MMEL1 | protein_coding | 0.99077463 | 3.24840572 |
| H4C5 | protein_coding | 0.99068497 | 1.58491032 |
| RP11-30K9.5 | lncRNA | 0.99040113 | 1.57810554 |
| RP11-325K4. | lncRNA | 0.98915765 | 2.30187781 |
| MYO1C | protein_coding | 0.98911597 | 15.6332408 |
| HPCA | protein_coding | 0.98863216 | 16.2195008 |
| IFT57 | protein_coding | 0.98801432 | 3.98474401 |
| SSPN | protein_coding | 0.98709976 | 3.34684037 |
| RAP1GAP | protein_coding | 0.98620341 | 14.6622532 |
| FUT1 | protein_coding | 0.98470536 | 9.94177081 |
| PHKG1 | protein_coding | 0.98399562 | 2.86829808 |
| AC003090.1 | lncRNA | 0.98372158 | 2.77737182 |
| PLEKHM1 | protein_coding | 0.98334336 | 12.7904771 |
| TTC39A-AS1 | lncRNA | 0.98245779 | 3.24662622 |
| EHD3 | protein_coding | 0.98181002 | 2.44547289 |
| PDIA5 | protein_coding | 0.98095142 | 9.25495455 |
| NT5DC1 | protein_coding | 0.98086865 | 11.1466282 |
| RP11-574K11 | lncRNA | 0.98081656 | 1.6336258 |
| TPPP3 | protein_coding | 0.97996933 | 11.6033967 |
| AMMECR1 | protein_coding | 0.97958927 | 11.4007636 |
| SERINC4 | protein_coding | 0.97943793 | 4.5667615 |
| PLA2G3 | protein_coding | 0.97890471 | 2.64908722 |
| SLC25A18 | protein_coding | 0.97872267 | 7.22919457 |
| CTC-499B15. | lncRNA | 0.97850037 | 2.84577851 |
| ABCC11 | protein_coding | 0.97806739 | 3.74083567 |
| IRF2-DT | lncRNA | 0.9767898 | 5.19314201 |
| LDLRAD4 | protein_coding | 0.97664985 | 8.06748463 |

| | | | |
|---|---|---|---|
| CHROMR | lncRNA | 0.97655297 | 11.089487 |
| KCNJ5 | protein_coding | 0.97517837 | 11.1594975 |
| SMCO2 | protein_coding | 0.97504571 | 1.96789761 |
| LINC01436 | lncRNA | 0.97354874 | 6.02100399 |
| CTD-2587H2 | lncRNA | 0.97057205 | 3.25181111 |
| H1-2 | protein_coding | 0.97048808 | 8.64093358 |
| NEAT1 | lncRNA | 0.96802214 | 15.1501137 |
| HCST | protein_coding | 0.96705691 | 2.18682082 |
| TEX14 | protein_coding | 0.9652689 | 3.58067459 |
| AC097721.2 | lncRNA | 0.96518881 | 1.53954559 |
| PPP1R26-AS | lncRNA | 0.96509462 | 7.59058837 |
| TNFRSF11A | protein_coding | 0.9645823 | 8.51982338 |
| H2AW | protein_coding | 0.96212873 | 10.5118189 |
| CCDC32 | protein_coding | 0.96172653 | 9.87995465 |
| SLC23A3 | protein_coding | 0.96093848 | 1.77717177 |
| ITGA3 | protein_coding | 0.96020156 | 12.9429088 |
| IGDCC3 | protein_coding | 0.95875274 | 11.0743745 |
| GALNT8 | protein_coding | 0.95824655 | 1.75588983 |
| TAS2R4 | protein_coding | 0.9580901 | 2.59529002 |
| AMER1 | protein_coding | 0.95645569 | 9.99948752 |
| LA16c-321D | lncRNA | 0.95623695 | 7.24404834 |
| HYOU1 | protein_coding | 0.95450545 | 18.2203887 |
| E2F5 | protein_coding | 0.95305767 | 7.20644531 |
| FAM153CP | lncRNA | 0.95232867 | 2.24585937 |
| LGMN | protein_coding | 0.95119975 | 11.7387052 |
| CTB-12A17.3 | lncRNA | 0.95108062 | 4.45306874 |
| SLC4A5 | protein_coding | 0.95055242 | 1.78976114 |
| RFX5 | protein_coding | 0.9504055 | 14.0636408 |
| LINC01686 | lncRNA | 0.94961455 | 4.2568852 |
| ITPKC | protein_coding | 0.9484954 | 9.84978048 |
| PAK3 | protein_coding | 0.94645053 | 2.76775412 |
| H2BC21 | protein_coding | 0.94437584 | 12.1578186 |
| DENND3 | protein_coding | 0.94281791 | 14.5203337 |
| SDC4 | protein_coding | 0.9409153 | 13.220921 |
| HCN4 | protein_coding | 0.94021365 | 13.5607957 |
| FNDC5 | protein_coding | 0.93932953 | 15.4882823 |
| CSDC2 | protein_coding | 0.93903747 | 10.6419196 |
| CARD8-AS1 | lncRNA | 0.93885903 | 3.76170561 |
| CTD-2014B1 | lncRNA | 0.93857881 | 1.21953712 |
| RP11-644F5. | lncRNA | 0.93641822 | 6.00740285 |
| LAMB3 | protein_coding | 0.93576028 | 7.39320234 |
| PCDHB13 | protein_coding | 0.93567848 | 4.50765775 |
| RILPL2 | protein_coding | 0.93550667 | 7.01688562 |
| STK17A | protein_coding | 0.93517986 | 8.07260928 |
| TRIP10 | protein_coding | 0.93387127 | 11.3749404 |

| | | | |
|---|---|---|---|
| ARHGAP22 | protein_coding | 0.9334772 | 2.75624358 |
| MAP3K14 | protein_coding | 0.93292325 | 15.2113236 |
| RP11-35G9.3 | lncRNA | 0.93201451 | 2.82503422 |
| DAND5 | protein_coding | 0.93130182 | 6.97244991 |
| RHPN2 | protein_coding | 0.93071595 | 7.02064354 |
| NOD1 | protein_coding | 0.93039862 | 7.43999849 |
| TP53 | protein_coding | 0.93035492 | 10.0182667 |
| FSTL4 | protein_coding | 0.93006373 | 12.982123 |
| WHRN | protein_coding | 0.92880468 | 11.9022063 |
| RP11-762I7.5 | protein_coding | 0.92732951 | 1.60005975 |
| ITGA10 | protein_coding | 0.92662831 | 3.13735732 |
| DAPK2 | protein_coding | 0.92599238 | 10.8000485 |
| CBX4 | protein_coding | 0.92552076 | 10.3533329 |
| KCNMB2-AS1 | lncRNA | 0.92451476 | 3.97210966 |
| KIF3C | protein_coding | 0.9244907 | 15.0097798 |
| AC128709.2 | lncRNA | 0.92433947 | 2.66353595 |
| TCF7L1 | protein_coding | 0.92377223 | 1.70995377 |
| RP11-817I4.1 | lncRNA | 0.92065975 | 2.93088368 |
| MRGPRE | protein_coding | 0.91758731 | 4.21124465 |
| OXTR | protein_coding | 0.91604008 | 9.99962425 |
| LHX2 | protein_coding | 0.91564526 | 11.6345714 |
| CD83 | protein_coding | 0.91544442 | 14.0489802 |
| H2BC4 | protein_coding | 0.91444275 | 2.41574692 |
| PRR5-ARHGA | protein_coding | 0.91274126 | 1.87615548 |
| SMAD7 | protein_coding | 0.9126772 | 6.63724981 |
| TRIM14 | protein_coding | 0.910721 | 14.7072756 |
| BCAS4 | protein_coding | 0.91032561 | 3.50795835 |
| DYNC2H1 | protein_coding | 0.90983223 | 2.11045848 |
| AIFM2 | protein_coding | 0.90925897 | 9.22253298 |
| UNKL | protein_coding | 0.90901878 | 11.9911621 |
| SGCE | protein_coding | 0.90702594 | 4.24456095 |
| TRIM66 | protein_coding | 0.90644576 | 9.48539168 |
| RP4-660H19. | lncRNA | 0.90550224 | 1.36720606 |
| ZNF347 | protein_coding | 0.9054559 | 3.17046695 |
| COTL1 | protein_coding | 0.90501625 | 12.9744211 |
| TSPO | protein_coding | 0.90447334 | 10.382545 |
| ID1 | protein_coding | 0.90326024 | 4.22098305 |
| HEATR5A | protein_coding | 0.90263513 | 9.40129101 |
| PLXDC1 | protein_coding | 0.90260328 | 3.08908248 |
| SKIL | protein_coding | 0.9014221 | 9.33734186 |
| ABCA5 | protein_coding | 0.90109284 | 13.3214323 |
| FBXO48 | protein_coding | 0.90041587 | 5.84274688 |
| ASCC3 | protein_coding | 0.9002975 | 11.8923785 |
| ZNF267 | protein_coding | 0.90026682 | 9.79391294 |
| JAK2 | protein_coding | 0.89996521 | 8.64456413 |

| | | | |
|---|---|---|---|
| NHSL2 | protein_coding | 0.89815495 | 13.9071934 |
| EPB41L4A-DT | lncRNA | 0.89749491 | 3.26608101 |
| AJUBA | protein_coding | 0.8964287 | 9.878602 |
| ALX4 | protein_coding | 0.89573729 | 3.08863557 |
| SAT1 | protein_coding | 0.89568085 | 6.45831026 |
| PCDHAC2 | protein_coding | 0.89560997 | 3.75625864 |
| AC005281.1 | protein_coding | 0.89501033 | 1.66913825 |
| PARP3 | protein_coding | 0.89488382 | 6.33616399 |
| AC137932.6 | lncRNA | 0.89483744 | 1.673651 |
| BTN2A1 | protein_coding | 0.89339473 | 9.8205409 |
| SH2D6 | protein_coding | 0.89271444 | 1.57454174 |
| KIAA1671 | protein_coding | 0.89258736 | 10.4552579 |
| NQO2 | protein_coding | 0.89174511 | 10.0330133 |
| RP11-482D24 | lncRNA | 0.8914118 | 1.42058059 |
| PSMA3 | protein_coding | 0.89110952 | 14.9730237 |
| PIK3R1 | protein_coding | 0.8907952 | 12.6862189 |
| TAL2 | protein_coding | 0.89024103 | 6.28079344 |
| ABCD1 | protein_coding | 0.89023401 | 8.60621855 |
| RIPOR1 | protein_coding | 0.88999021 | 10.8175204 |
| JAG1 | protein_coding | 0.88915885 | 9.81628157 |
| CPEB2 | protein_coding | 0.88734974 | 9.23458473 |
| TYK2 | protein_coding | 0.88622729 | 14.6693947 |
| TYW3 | protein_coding | 0.88599018 | 10.7947634 |
| RP11-334C17 | lncRNA | 0.88579119 | 4.39922681 |
| ARHGAP23 | protein_coding | 0.88523789 | 13.1264311 |
| CTD-2650P22 | lncRNA | 0.88402024 | 2.49046462 |
| CHRNA1 | protein_coding | 0.88396424 | 12.9125554 |
| CDKL1 | protein_coding | 0.88215325 | 1.85360967 |
| HAUS7 | protein_coding | 0.88208557 | 4.80287446 |
| TNFRSF1B | protein_coding | 0.88062296 | 7.92188243 |
| LRATD2 | protein_coding | 0.88018092 | 5.63020768 |
| SLC37A1 | protein_coding | 0.87940473 | 11.4942938 |
| PCYT1B | protein_coding | 0.87919923 | 7.05417118 |
| ADGRF5 | protein_coding | 0.87875478 | 14.6941433 |
| RIMS3 | protein_coding | 0.87859934 | 17.294858 |
| PLLP | protein_coding | 0.87858679 | 3.04460013 |
| RP11-881M1 | lncRNA | 0.87848614 | 1.4794372 |
| AC010729.1 | lncRNA | 0.8784306 | 1.28581692 |
| IRGQ | protein_coding | 0.87634886 | 9.24636377 |
| B4GALT1 | protein_coding | 0.87630823 | 14.7811893 |
| CRAT | protein_coding | 0.87550422 | 10.9690276 |
| DDR2 | protein_coding | 0.87548507 | 4.14807213 |
| RBM43 | protein_coding | 0.87499918 | 7.23139845 |
| RP11-286B14 | lncRNA | 0.87498028 | 1.22594045 |
| SH2B3 | protein_coding | 0.87416624 | 11.995945 |

| | | | |
|---|---|---|---|
| RASL11A | protein_coding | 0.87375825 | 9.53424315 |
| RP11-549B18 | lncRNA | 0.87341927 | 6.39053589 |
| GRPR | protein_coding | 0.87127193 | 3.99899713 |
| MTMR4 | protein_coding | 0.87076937 | 16.8232237 |
| WDR49 | protein_coding | 0.87043432 | 2.1886113 |
| PTPRR | protein_coding | 0.86591594 | 5.45480901 |
| EPGN | protein_coding | 0.86559901 | 9.17869706 |
| STAC2 | protein_coding | 0.86486836 | 4.42795709 |
| SEMA4C | protein_coding | 0.86414539 | 14.3622799 |
| LINC01252 | lncRNA | 0.86216862 | 2.60377784 |
| NPIPA1 | protein_coding | 0.86188282 | 1.56305838 |
| TUBE1 | protein_coding | 0.86142523 | 9.54512567 |
| PIK3R2 | protein_coding | 0.86106169 | 1.66156769 |
| ARMH4 | protein_coding | 0.86084425 | 6.36797044 |
| RPH3AL | protein_coding | 0.86064201 | 4.74342636 |
| PLAC8L1 | protein_coding | 0.85764226 | 2.77028418 |
| RYBP | protein_coding | 0.85730169 | 12.127116 |
| C1RL-AS1 | lncRNA | 0.85726817 | 3.61707148 |
| SLC7A5 | protein_coding | 0.85653312 | 15.5978496 |
| SYNGR2 | protein_coding | 0.85534581 | 14.6860168 |
| PCDHA11 | protein_coding | 0.85402057 | 8.24770069 |
| CDH22 | protein_coding | 0.85161167 | 11.833548 |
| LARP6 | protein_coding | 0.85045461 | 6.96676952 |
| PCDHA4 | protein_coding | 0.84878023 | 5.87645502 |
| MRO | protein_coding | 0.84876293 | 2.28330509 |
| LMO3 | protein_coding | 0.84837408 | 7.13514953 |
| CDH12 | protein_coding | 0.8482066 | 6.6023275 |
| ARRDC3 | protein_coding | 0.84672379 | 13.3363624 |
| VSIG10L | protein_coding | 0.84627032 | 4.10349993 |
| TTC38 | protein_coding | 0.84460762 | 6.48381607 |
| PDE2A | protein_coding | 0.84386592 | 8.94010871 |
| GAD1 | protein_coding | 0.84338247 | 5.24300446 |
| NRIP2 | protein_coding | 0.84240734 | 11.3709433 |
| KLF12 | protein_coding | 0.84003408 | 13.2009767 |
| NTHL1 | protein_coding | 0.83991349 | 10.3720131 |
| MYO1E | protein_coding | 0.83991129 | 10.1150028 |
| QSOX1 | protein_coding | 0.8395746 | 11.1830917 |
| CTC-366B18. | lncRNA | 0.839036 | 1.87088764 |
| IGFBP4 | protein_coding | 0.83897402 | 2.82862133 |
| SDK1 | protein_coding | 0.83852423 | 12.4383358 |
| UBR5-AS1 | lncRNA | 0.83784826 | 1.22427511 |
| ZC2HC1C | protein_coding | 0.83704239 | 8.71487341 |
| SSUH2 | protein_coding | 0.83610983 | 2.97817837 |
| ETHE1 | protein_coding | 0.83557687 | 6.99124226 |
| PCDHB10 | protein_coding | 0.83455396 | 3.06936612 |

| | | | |
|---|---|---|---|
| WDR91 | protein_coding | 0.83438254 | 12.0680163 |
| RASSF5 | protein_coding | 0.83141125 | 11.6492127 |
| KIAA2012 | protein_coding | 0.8309766 | 1.68601616 |
| PCDHB14 | protein_coding | 0.83093827 | 6.51644735 |
| KIAA0513 | protein_coding | 0.83081809 | 11.7121868 |
| RPS6KL1 | protein_coding | 0.82980216 | 12.5414055 |
| KCNV1 | protein_coding | 0.82943867 | 7.59250465 |
| PRRT3-AS1 | lncRNA | 0.82787188 | 1.08990669 |
| DRGX | protein_coding | 0.82763958 | 10.4237438 |
| RP11-72M17 | lncRNA | 0.82757749 | 2.17251478 |
| NEDD4 | protein_coding | 0.82708721 | 10.2823105 |
| ACADVL | protein_coding | 0.82690958 | 11.9671815 |
| TRMT11 | protein_coding | 0.82668294 | 9.74923422 |
| AC005220.3 | lncRNA | 0.82665133 | 1.06032759 |
| ELL3 | protein_coding | 0.8258523 | 3.4208613 |
| C2orf92 | protein_coding | 0.82563993 | 4.5188391 |
| RP11-1391J7 | lncRNA | 0.82420258 | 2.24153375 |
| ZSCAN31 | protein_coding | 0.82359548 | 11.9510314 |
| PALM3 | protein_coding | 0.82331248 | 2.23096826 |
| SH3BP5 | protein_coding | 0.8233035 | 8.52481122 |
| FAM178B | protein_coding | 0.82283595 | 4.23549314 |
| TNK1 | protein_coding | 0.82277386 | 7.23853633 |
| NFE2L1 | protein_coding | 0.82260875 | 15.8036506 |
| DDX47 | protein_coding | 0.82197305 | 2.34372298 |
| ANXA7 | protein_coding | 0.82010671 | 12.8727698 |
| INTS6L | protein_coding | 0.8200958 | 10.7309222 |
| RAP2C-AS1 | lncRNA | 0.81872531 | 2.3668346 |
| HOXD13 | protein_coding | 0.81827166 | 3.09737485 |
| NPB | protein_coding | 0.81783153 | 1.43693981 |
| LAMC1 | protein_coding | 0.81736218 | 11.6128534 |
| MMD | protein_coding | 0.81693545 | 9.51080897 |
| C15orf65 | protein_coding | 0.8168032 | 2.0506516 |
| ZBED6CL | protein_coding | 0.81679441 | 10.3001993 |
| PCDHA12 | protein_coding | 0.81607148 | 3.37067025 |
| ADAMTS7 | protein_coding | 0.81529733 | 3.06814706 |
| CEP135 | protein_coding | 0.81524945 | 7.64791385 |
| H4C8 | protein_coding | 0.81489065 | 2.43194557 |
| PCDHA7 | protein_coding | 0.81424394 | 1.6016011 |
| RP11-244H3. | protein_coding | 0.81403351 | 1.75145365 |
| FAM219A | protein_coding | 0.81375958 | 12.767185 |
| RP11-161H2: | lncRNA | 0.81261928 | 1.71338318 |
| PSMA6 | protein_coding | 0.81229814 | 9.36784074 |
| MET | protein_coding | 0.81209226 | 10.0322638 |
| SV2B | protein_coding | 0.81158035 | 12.0259762 |
| RP11-531A2 | lncRNA | 0.81129808 | 5.56052536 |

| | | | |
|---|---|---|---|
| RCBTB1 | protein_coding | 0.81099566 | 13.0631596 |
| P2RX6 | protein_coding | 0.81038347 | 3.05665659 |
| PSAP | protein_coding | 0.81017143 | 16.7236207 |
| ZNF880 | protein_coding | 0.80972826 | 3.35901475 |
| KCND1 | protein_coding | 0.80970817 | 7.03499805 |
| MEGF11 | protein_coding | 0.80881737 | 8.11464689 |
| TPH1 | protein_coding | 0.80873783 | 1.48063982 |
| PIK3IP1 | protein_coding | 0.80810949 | 6.53381623 |
| TRAM2 | protein_coding | 0.80785885 | 9.4596431 |
| RP11-542K23 | lncRNA | 0.80626028 | 1.26056578 |
| CMTR1 | protein_coding | 0.80575973 | 14.3724447 |
| RP5-1057I20 | lncRNA | 0.80543975 | 2.30206675 |
| LENG9 | protein_coding | 0.80471383 | 3.05931929 |
| NMNAT2 | protein_coding | 0.80358952 | 8.71710174 |
| PCDHA5 | protein_coding | 0.80248492 | 6.3189491 |
| EXOC3L1 | protein_coding | 0.80210098 | 4.3937086 |
| THSD4 | protein_coding | 0.80204408 | 7.28461187 |
| BANK1 | protein_coding | 0.80169359 | 2.69931626 |
| RUFY1-AS1 | lncRNA | 0.80070495 | 0.86195992 |
| RP11-589P10 | lncRNA | 0.80063889 | 1.25270457 |
| KHNYN | protein_coding | 0.8006202 | 12.6672668 |
| VWA7 | protein_coding | 0.80059602 | 2.35349993 |
| SOX11 | protein_coding | 0.79927016 | 11.8501744 |
| EPHA3 | protein_coding | 0.7991853 | 7.54143991 |
| TTC34 | protein_coding | 0.79913385 | 2.76566511 |
| H2BU1 | protein_coding | 0.79895888 | 6.81249689 |
| TRPM2 | protein_coding | 0.79880405 | 11.7018894 |
| TP53AIP1 | protein_coding | 0.79872979 | 1.90239165 |
| MFGE8 | protein_coding | 0.7980422 | 11.0440089 |
| ACAD11 | protein_coding | 0.79801721 | 2.94241011 |
| DIRAS2 | protein_coding | 0.79714494 | 3.30856021 |
| CMIP | protein_coding | 0.79705189 | 15.6696459 |
| OVGP1 | protein_coding | 0.79622013 | 3.47969149 |
| TEX19 | protein_coding | 0.79598896 | 1.91935321 |
| PTCH2 | protein_coding | 0.79529667 | 3.92931625 |
| DOCK6 | protein_coding | 0.79521124 | 1.00566092 |
| BHLHE40 | protein_coding | 0.79487512 | 10.4442179 |
| THBS3-AS1 | lncRNA | 0.79346373 | 3.1127655 |
| LCA5 | protein_coding | 0.79298867 | 8.69775665 |
| LINC02525 | lncRNA | 0.79286803 | 5.94050918 |
| AC005609.16 | lncRNA | 0.7923693 | 4.44408378 |
| BIN1 | protein_coding | 0.79106182 | 6.85814973 |
| ANKDD1A | protein_coding | 0.79045074 | 1.90897161 |
| EPB41L4A | protein_coding | 0.79003272 | 12.3826515 |
| ERO1B | protein_coding | 0.78978616 | 11.9736619 |

| | | | |
|---|---|---|---|
| USP45 | protein_coding | 0.78918155 | 10.627721 |
| CFAP94 | protein_coding | 0.78905577 | 1.3757343 |
| PDGFD | protein_coding | 0.78848483 | 8.2736018 |
| MCC | protein_coding | 0.78790387 | 11.9970789 |
| CYLD | protein_coding | 0.78673232 | 12.9477717 |
| TMCC2 | protein_coding | 0.78635546 | 10.622061 |
| FAM111A-DT | lncRNA | 0.7861027 | 4.84315577 |
| C17orf49 | protein_coding | 0.78541731 | 1.23954936 |
| PGAM2 | protein_coding | 0.78527417 | 1.2351356 |
| C6orf141 | protein_coding | 0.78504309 | 10.4158206 |
| TMC5 | protein_coding | 0.78424843 | 1.22823914 |
| KCNS2 | protein_coding | 0.78405532 | 3.01763045 |
| ARSI | protein_coding | 0.78298352 | 10.8729222 |
| RBM20 | protein_coding | 0.78238136 | 7.22023662 |
| ADAMTSL2 | protein_coding | 0.78171344 | 5.04611524 |
| NOL4 | protein_coding | 0.78129719 | 14.6449236 |
| HRK | protein_coding | 0.78121927 | 5.5349716 |
| H2AC8 | protein_coding | 0.78109919 | 1.98562377 |
| SNAP91 | protein_coding | 0.78034609 | 1.70773095 |
| RHOQ | protein_coding | 0.779769 | 11.1256841 |
| LINC01126 | lncRNA | 0.77820975 | 1.83557578 |
| SPECC1 | protein_coding | 0.77811281 | 7.63019434 |
| MSI1 | protein_coding | 0.77802148 | 3.29214063 |
| HIF3A | protein_coding | 0.77764793 | 4.45929716 |
| CPT1A | protein_coding | 0.77746725 | 13.4326082 |
| PCDHGB1 | protein_coding | 0.77692827 | 2.33258906 |
| GALNT2 | protein_coding | 0.77687759 | 10.0282146 |
| CNN3 | protein_coding | 0.77589295 | 14.5221546 |
| ZBTB7B | protein_coding | 0.77573355 | 13.2136178 |
| ZNF165 | protein_coding | 0.77563339 | 6.82688974 |
| RAB39A | protein_coding | 0.77538332 | 9.63053199 |
| GCLM | protein_coding | 0.77501742 | 9.68887245 |
| MAP1B | protein_coding | 0.77448557 | 17.5106751 |
| OGFR | protein_coding | 0.77436231 | 10.9064404 |
| LL22NC03-12 | lncRNA | 0.77331912 | 1.22676101 |
| RUSC1-AS1 | lncRNA | 0.77212573 | 1.41405976 |
| SECTM1 | protein_coding | 0.77155289 | 10.4548311 |
| N4BP1 | protein_coding | 0.77076015 | 11.826716 |
| ZBTB8B | protein_coding | 0.77023061 | 7.399576 |
| CTD-2017F17 | lncRNA | 0.76951725 | 1.44982238 |
| KCNN1 | protein_coding | 0.76777403 | 8.32171615 |
| SPRYD7 | protein_coding | 0.7673602 | 4.81488031 |
| DAB2IP | protein_coding | 0.76729616 | 12.1823428 |
| RP11-514P8. | lncRNA | 0.76696179 | 1.10073157 |
| DNHD1 | protein_coding | 0.76649355 | 6.1335965 |

| | | | |
|---|---|---|---|
| PROM2 | protein_coding | 0.76597697 | 0.98236848 |
| LRP1B | protein_coding | 0.76511161 | 9.64721331 |
| NBPF15 | protein_coding | 0.76432638 | 2.72224913 |
| SULT2B1 | protein_coding | 0.76399489 | 1.31594721 |
| KCNC4 | protein_coding | 0.76370487 | 8.74421357 |
| LRRTM4 | protein_coding | 0.7623355 | 1.80689024 |
| ALPK1 | protein_coding | 0.76204621 | 1.26159296 |
| CRHR2 | protein_coding | 0.76111859 | 2.0602911 |
| YAF2 | protein_coding | 0.76095533 | 5.74864337 |
| RP1-257A7.4 | lncRNA | 0.76084951 | 2.71161419 |
| OAF | protein_coding | 0.76063322 | 1.01669267 |
| LDB3 | protein_coding | 0.75914761 | 6.02837774 |
| ITM2B | protein_coding | 0.75904453 | 13.2063956 |
| TRAF4 | protein_coding | 0.75820186 | 14.9044006 |
| U73166.2 | lncRNA | 0.75799032 | 1.10294287 |
| SLC27A2 | protein_coding | 0.75724013 | 10.6161424 |
| FAM47E | protein_coding | 0.75545717 | 1.07998736 |
| GCHFR | protein_coding | 0.75401403 | 1.32308792 |
| CTD-2008P7. | lncRNA | 0.75360021 | 1.96470078 |
| FLRT3 | protein_coding | 0.7533998 | 4.56215291 |
| ST7-AS1 | lncRNA | 0.75165878 | 5.37810894 |
| JUN | protein_coding | 0.75164966 | 9.19466273 |
| C4orf33 | protein_coding | 0.75161135 | 8.1558278 |
| PDC-AS1 | lncRNA | 0.75107505 | 1.53255415 |
| ST3GAL1 | protein_coding | 0.75087175 | 14.8199622 |
| ZNF608 | protein_coding | 0.75030835 | 9.45157329 |
| CTF1 | protein_coding | 0.75017572 | 2.31563632 |
| ALDH1B1 | protein_coding | 0.75012732 | 7.25299758 |
| RP11-798G7. | lncRNA | 0.74961878 | 1.25234602 |
| SPX | protein_coding | 0.74911421 | 5.61503203 |
| RP11-475M2 | lncRNA | 0.74840761 | 1.87650267 |
| PTPRO | protein_coding | 0.74750875 | 3.51653776 |
| CACNG4 | protein_coding | 0.7474739 | 10.3561983 |
| ASPHD2 | protein_coding | 0.74704358 | 5.40710108 |
| TBCE | protein_coding | 0.74672817 | 2.43506128 |
| SLITRK5 | protein_coding | 0.74619955 | 8.59025396 |
| CATSPERG | protein_coding | 0.74592591 | 5.28298814 |
| GSPT2 | protein_coding | 0.74542234 | 1.83818291 |
| ZFP36 | protein_coding | 0.74472105 | 5.93927927 |
| ODAD4 | protein_coding | 0.74440612 | 3.03095912 |
| C3orf62 | protein_coding | 0.74415947 | 7.20351433 |
| ATP6V1B1 | protein_coding | 0.74414238 | 1.07000981 |
| NEU1 | protein_coding | 0.74395034 | 10.6753443 |
| RP11-284N8. | lncRNA | 0.74277432 | 5.70289403 |
| SLC12A7 | protein_coding | 0.74273858 | 9.42527493 |

| | | | |
|---|---|---|---|
| CHKB | protein_coding | 0.74239306 | 2.19269221 |
| HACD2 | protein_coding | 0.74171797 | 13.8682039 |
| RP11-48B3.4 | lncRNA | 0.74145449 | 1.08494337 |
| NOCT | protein_coding | 0.74131683 | 7.30718658 |
| TVP23B | protein_coding | 0.74035562 | 5.6460988 |
| KCNN2 | protein_coding | 0.73955 | 10.4358957 |
| AGAP4 | protein_coding | 0.73935845 | 1.95306602 |
| LINC00365 | lncRNA | 0.73929693 | 2.95008915 |
| GPR108 | protein_coding | 0.73893202 | 10.1041575 |
| ZSCAN23 | protein_coding | 0.73855743 | 2.33836736 |
| MR1 | protein_coding | 0.73754443 | 7.86325339 |
| PPARGC1B | protein_coding | 0.73728058 | 11.8319968 |
| SUCLG2-AS1 | lncRNA | 0.73614901 | 2.75107023 |
| LINC01664 | lncRNA | 0.73563184 | 1.73267974 |
| RP11-499E18 | lncRNA | 0.73415744 | 1.03940728 |
| WDFY1 | protein_coding | 0.73412706 | 9.52536219 |
| ARHGAP27 | protein_coding | 0.73386458 | 7.0938874 |
| WDR41 | protein_coding | 0.73354495 | 10.978088 |
| VEGFD | protein_coding | 0.73297829 | 1.57438085 |
| ADAMTS1 | protein_coding | 0.73276851 | 2.04912416 |
| RFFL | protein_coding | 0.73275189 | 8.6308596 |
| AGTRAP | protein_coding | 0.73268697 | 6.94481248 |
| MLLT11 | protein_coding | 0.73187318 | 11.7114569 |
| GPR179 | protein_coding | 0.73178871 | 2.28302352 |
| DTX2 | protein_coding | 0.73129666 | 7.47357849 |
| MAPRE3 | protein_coding | 0.73106691 | 8.29960944 |
| IL18BP | protein_coding | 0.72994462 | 12.3653683 |
| SP6 | protein_coding | 0.7296375 | 11.3806077 |
| GATAD1 | protein_coding | 0.7286754 | 9.34531568 |
| PODXL | protein_coding | 0.72785422 | 15.8475819 |
| SH3BGR | protein_coding | 0.72773927 | 1.37489529 |
| BAHD1 | protein_coding | 0.72721188 | 11.2268566 |
| LINC00364 | lncRNA | 0.72686821 | 8.63254892 |
| ATXN7L1 | protein_coding | 0.72618487 | 8.41628204 |
| PCDHGB2 | protein_coding | 0.72513507 | 8.9690117 |
| H1-0 | protein_coding | 0.72470189 | 15.3320953 |
| ZNF773 | protein_coding | 0.72446703 | 7.83876871 |
| RELL1 | protein_coding | 0.72397561 | 11.5615108 |
| ZNF726 | protein_coding | 0.7238789 | 2.61257523 |
| TTC27 | protein_coding | 0.72379098 | 9.43299703 |
| RBM34 | protein_coding | 0.7230691 | 1.79085823 |
| GOLGA6L10 | protein_coding | 0.72230344 | 1.97979565 |
| RP11-774D14 | lncRNA | 0.72215655 | 1.52790123 |
| DEXI | protein_coding | 0.72212775 | 11.2640739 |
| CALCOCO2 | protein_coding | 0.72153214 | 11.8333474 |

| | | | |
|---|---|---|---|
| STK40 | protein_coding | 0.72140618 | 12.0143628 |
| ALPK3 | protein_coding | 0.72031047 | 11.1886373 |
| UNC119 | protein_coding | 0.71941966 | 7.6963435 |
| PRICKLE3 | protein_coding | 0.71881068 | 7.56372101 |
| APBA3 | protein_coding | 0.71812686 | 8.98994141 |
| KIAA0408 | protein_coding | 0.71792883 | 0.73927063 |
| UBE2O | protein_coding | 0.71667009 | 14.3084037 |
| SNHG26 | lncRNA | 0.71578991 | 5.52062699 |
| RASGRF2 | protein_coding | 0.71578107 | 2.05661592 |
| JAK1 | protein_coding | 0.71517509 | 13.196206 |
| CD47 | protein_coding | 0.71482062 | 12.5973521 |
| CLEC18A | protein_coding | 0.7148127 | 1.99861841 |
| SCGB1D1 | protein_coding | 0.71464399 | 1.74746685 |
| BICD1 | protein_coding | 0.71426906 | 8.55985624 |
| PLEKHO2 | protein_coding | 0.71375063 | 3.98257319 |
| FAM117A | protein_coding | 0.71329067 | 10.5442446 |
| SLC43A1 | protein_coding | 0.71325155 | 11.1062817 |
| PCDHB16 | protein_coding | 0.71240162 | 2.87335171 |
| PAG1 | protein_coding | 0.71134742 | 10.4517006 |
| REPIN1 | protein_coding | 0.71094531 | 9.61691305 |
| CARS1 | protein_coding | 0.71048302 | 13.9744587 |
| ABHD16A | protein_coding | 0.71014504 | 2.29231818 |
| HES2 | protein_coding | 0.70929824 | 5.08333914 |
| LINC01569 | lncRNA | 0.70889551 | 3.06548428 |
| ZSCAN5A | protein_coding | 0.70873421 | 7.27296087 |
| CDKL2 | protein_coding | 0.70730288 | 5.51502813 |
| STOX1 | protein_coding | 0.70720071 | 6.725755 |
| A1BG | protein_coding | 0.706726 | 1.34117135 |
| GNA15 | protein_coding | 0.70649156 | 1.76017163 |
| WARS1 | protein_coding | 0.70599111 | 14.9671606 |
| LBHD1 | protein_coding | 0.70522194 | 5.24763847 |
| RP11-107G2 | lncRNA | 0.70444761 | 1.20978006 |
| TMBIM1 | protein_coding | 0.70379068 | 12.6403535 |
| HAS3 | protein_coding | 0.70331217 | 8.61396257 |
| HOXC4 | protein_coding | 0.7025658 | 5.33892923 |
| EN2 | protein_coding | 0.70231643 | 6.05359394 |
| SLC6A18 | protein_coding | 0.70221538 | 4.35155787 |
| TGIF1 | protein_coding | 0.70218489 | 1.5637289 |
| MDK | protein_coding | 0.70174116 | 10.6015866 |
| CEBPG | protein_coding | 0.70151798 | 15.014824 |
| CFAP57 | protein_coding | 0.70102468 | 1.693985 |
| CYP27C1 | protein_coding | 0.70053188 | 3.87536843 |
| ITPKB | protein_coding | 0.70049117 | 5.96636257 |
| GALNT10 | protein_coding | 0.70024671 | 7.671362 |
| NPTN-IT1 | lncRNA | 0.70023694 | 4.76924905 |

| | | | |
|---|---|---|---|
| ARMC12 | protein_coding | 0.69798488 | 1.99713266 |
| PDE8A | protein_coding | 0.69769676 | 10.4783597 |
| AC005754.8 | lncRNA | 0.69725546 | 0.99319669 |
| IFRD1 | protein_coding | 0.69722042 | 6.94619095 |
| SCPEP1 | protein_coding | 0.69614948 | 11.1291833 |
| TMEM216 | protein_coding | 0.69592468 | 7.3849931 |
| CTA-315H11. | lncRNA | 0.69495529 | 0.99769509 |
| PTPRZ1 | protein_coding | 0.69492836 | 13.4506343 |
| NUCB1 | protein_coding | 0.69483438 | 12.761654 |
| IRF2BPL | protein_coding | 0.69460147 | 7.87785215 |
| TMC4 | protein_coding | 0.69439951 | 1.36294981 |
| PCDHB2 | protein_coding | 0.69392287 | 8.03956945 |
| MT1G | protein_coding | 0.69362923 | 4.99492956 |
| CXorf38 | protein_coding | 0.69360436 | 9.47760384 |
| LINC00992 | lncRNA | 0.69236489 | 1.38200776 |
| TRIP6 | protein_coding | 0.69235114 | 2.45993071 |
| TMEM63A | protein_coding | 0.69168736 | 3.51478519 |
| SMCHD1 | protein_coding | 0.69087321 | 14.6587665 |
| RP11-155D1 | lncRNA | 0.68998907 | 1.70438004 |
| TMEM131L | protein_coding | 0.68913431 | 9.92034854 |
| FKBP11 | protein_coding | 0.68902286 | 5.81051284 |
| MEIS1 | protein_coding | 0.68869943 | 7.82281818 |
| SASH3 | protein_coding | 0.68862133 | 2.59491081 |
| SMKR1 | protein_coding | 0.68849498 | 2.05929573 |
| NIPAL2 | protein_coding | 0.68787905 | 7.604029 |
| PCDHGA1 | protein_coding | 0.68769271 | 3.45820514 |
| RASIP1 | protein_coding | 0.68685209 | 3.70127602 |
| FLG-AS1 | lncRNA | 0.68461786 | 4.37902231 |
| TSPEAR | protein_coding | 0.68336775 | 1.74262695 |
| GAS2 | protein_coding | 0.68295245 | 1.75641173 |
| TP53BP1 | protein_coding | 0.68260939 | 13.5030845 |
| EDIL3-DT | lncRNA | 0.68211572 | 1.29644081 |
| MATN1 | protein_coding | 0.68197569 | 3.48172905 |
| SLC8B1 | protein_coding | 0.68175548 | 8.244009 |
| DAG1 | protein_coding | 0.68151126 | 12.8316111 |
| MLC1 | protein_coding | 0.68090106 | 1.35022999 |
| SPG11 | protein_coding | 0.67999211 | 11.7534319 |
| ST20 | protein_coding | 0.67899389 | 1.55158928 |
| TMEM178B | protein_coding | 0.67894891 | 14.0592311 |
| H2BC7 | protein_coding | 0.6789278 | 1.27568751 |
| DLGAP4 | protein_coding | 0.67880144 | 10.128481 |
| TPM1 | protein_coding | 0.67864316 | 14.453485 |
| IRAK1 | protein_coding | 0.67862521 | 13.42193 |
| MIA2 | protein_coding | 0.6774939 | 10.8528078 |
| NIT2 | protein_coding | 0.67720009 | 7.25217621 |

| | | | |
|---|---|---|---|
| ACCS | protein_coding | 0.67712131 | 5.22841413 |
| SNAP23 | protein_coding | 0.67692635 | 5.61406212 |
| NBR2 | lncRNA | 0.67662696 | 7.02365076 |
| RFX3-AS1 | lncRNA | 0.67648982 | 7.15083742 |
| ZNF619 | protein_coding | 0.67595386 | 4.16283468 |
| ACTN2 | protein_coding | 0.67594424 | 0.93296123 |
| RAB39B | protein_coding | 0.67584568 | 6.45777466 |
| SEMA6A-AS1 | lncRNA | 0.67551435 | 3.59301966 |
| AC128709.3 | lncRNA | 0.67520052 | 1.73129115 |
| GPR135 | protein_coding | 0.67448554 | 2.3157876 |
| UVRAG | protein_coding | 0.67379967 | 9.49781352 |
| RP11-631N1( | lncRNA | 0.67377423 | 3.31007934 |
| STAT3 | protein_coding | 0.67364125 | 15.7549653 |
| ECPAS | protein_coding | 0.67315417 | 13.6919347 |
| TENT5A | protein_coding | 0.67304465 | 11.3772085 |
| PSMA2 | protein_coding | 0.67300359 | 3.2030804 |
| DTNA | protein_coding | 0.67299093 | 8.43905307 |
| GOLGA6L9 | protein_coding | 0.67265676 | 5.50585685 |
| TRIOBP | protein_coding | 0.67263012 | 11.3440949 |
| ZNF222 | protein_coding | 0.67252947 | 5.7053667 |
| RP11-794G2 | lncRNA | 0.67073673 | 2.63303977 |
| SHB | protein_coding | 0.67029368 | 6.17663527 |
| FCMR | protein_coding | 0.6690844 | 1.35400577 |
| ATXN7 | protein_coding | 0.66883495 | 7.95527689 |
| LYST | protein_coding | 0.66860602 | 10.0949701 |
| NABP1 | protein_coding | 0.6684008 | 3.6995277 |
| RP11-472N1 | lncRNA | 0.66835566 | 1.18828449 |
| SFR1 | protein_coding | 0.66795188 | 4.96935219 |
| RP11-348P1( | lncRNA | 0.66757402 | 1.38906111 |
| CSRNP3 | protein_coding | 0.66713319 | 11.1262682 |
| HSPA4L | protein_coding | 0.66689026 | 11.404899 |
| CTD-3193O1 | lncRNA | 0.66552284 | 1.74165779 |
| HIVEP1 | protein_coding | 0.66482177 | 10.728458 |
| GALNT3 | protein_coding | 0.66434419 | 7.00477673 |
| RP11-170M1 | lncRNA | 0.66225203 | 0.84060521 |
| TMEM9B-AS | lncRNA | 0.66217877 | 4.24428948 |
| ZFHX4-AS1 | lncRNA | 0.6614279 | 1.17041276 |
| S100A13 | protein_coding | 0.66141124 | 2.24084075 |
| SYTL5 | protein_coding | 0.66121457 | 2.06058037 |
| AFDN-DT | lncRNA | 0.66098456 | 2.32893368 |
| NFATC1 | protein_coding | 0.66095001 | 6.22364641 |
| CEBPB-AS1 | lncRNA | 0.65897916 | 2.89258227 |
| DDIT4L | protein_coding | 0.65874177 | 2.16481839 |
| KCNH8 | protein_coding | 0.65871853 | 8.14365718 |
| CALB2 | protein_coding | 0.65839237 | 8.41299993 |

| | | | |
|---|---|---|---|
| CNKSR1 | protein_coding | 0.65789614 | 3.95822265 |
| FBXO16 | protein_coding | 0.65642703 | 3.02454596 |
| AFAP1 | protein_coding | 0.65625905 | 12.0125547 |
| CRABP1 | protein_coding | 0.65564377 | 8.79407991 |
| RP4-548D19. | lncRNA | 0.65541676 | 1.76701781 |
| PDLIM3 | protein_coding | 0.65536865 | 7.22119901 |
| RP11-34P13. | lncRNA | 0.65534156 | 0.90079543 |
| DOCK9 | protein_coding | 0.65512185 | 10.2703475 |
| NOB1 | protein_coding | 0.65502646 | 10.7471671 |
| RBCK1 | protein_coding | 0.65358209 | 11.7468352 |
| STX16 | protein_coding | 0.65329578 | 11.8422216 |
| LINC01616 | lncRNA | 0.65247762 | 1.24128388 |
| PPTC7 | protein_coding | 0.65075276 | 9.62158089 |
| GNRH1 | protein_coding | 0.65059449 | 1.21669935 |
| WDSUB1 | protein_coding | 0.65050685 | 7.99402975 |
| KCNB2 | protein_coding | 0.65023872 | 6.43960369 |
| ADGRL1-AS1 | lncRNA | 0.65021521 | 1.90783854 |
| PCDH8 | protein_coding | 0.64928096 | 3.81536316 |
| SRGAP1 | protein_coding | 0.64925449 | 10.9382672 |
| FAM53B | protein_coding | 0.64905281 | 8.20981573 |
| SYNGR3 | protein_coding | 0.64844608 | 9.35740752 |
| PCDHB12 | protein_coding | 0.64733446 | 2.86643719 |
| ZEB1 | protein_coding | 0.64722342 | 6.88638827 |
| CXCR4 | protein_coding | 0.64600766 | 10.2925972 |
| PCDHB9 | protein_coding | 0.64584311 | 1.55295611 |
| SEMA3E | protein_coding | 0.64522668 | 11.5065182 |
| ARL2BP | protein_coding | 0.64517645 | 11.0634875 |
| LAMP2 | protein_coding | 0.64306909 | 10.1614114 |
| CCDC33 | protein_coding | 0.6428373 | 5.84631525 |
| CAPN12 | protein_coding | 0.64266994 | 7.02454443 |
| TBC1D9 | protein_coding | 0.64266837 | 3.99688333 |
| DPP6 | protein_coding | 0.64250222 | 7.63768913 |
| ARHGAP18 | protein_coding | 0.64240947 | 11.1247869 |
| GRHPR | protein_coding | 0.64181031 | 11.9418705 |
| SCAMP3 | protein_coding | 0.64174222 | 12.0556524 |
| FRY | protein_coding | 0.64171228 | 11.5231436 |
| FAM184B | protein_coding | 0.64149346 | 10.6567544 |
| H2BC6 | protein_coding | 0.64146695 | 3.18905984 |
| ARF3 | protein_coding | 0.64121495 | 14.5654468 |
| RHBDD3 | protein_coding | 0.64082539 | 8.85123604 |
| PLEKHG6 | protein_coding | 0.64044817 | 1.84316246 |
| MASTL | protein_coding | 0.6402724 | 8.37862369 |
| NOMO2 | protein_coding | 0.64021585 | 7.5023109 |
| USP2 | protein_coding | 0.63968077 | 11.8179174 |
| TMOD4 | protein_coding | 0.63952571 | 2.42366845 |

| | | | |
|---|---|---|---|
| MYH14 | protein_coding | 0.63887485 | 9.56772532 |
| SEC22A | protein_coding | 0.63830817 | 6.85583848 |
| LRRN1 | protein_coding | 0.63811523 | 6.23115561 |
| RP11-680A1: | lncRNA | 0.63773807 | 1.3473665 |
| NPLOC4 | protein_coding | 0.63599732 | 12.5432717 |
| RBBP9 | protein_coding | 0.63573616 | 9.19514622 |
| KIF5C-AS1 | lncRNA | 0.63534464 | 1.21213039 |
| TUSC1 | protein_coding | 0.63506781 | 5.2791338 |
| NPAS1 | protein_coding | 0.63489965 | 2.3626934 |
| XDH | protein_coding | 0.63481442 | 4.63253157 |
| GPT2 | protein_coding | 0.63446418 | 9.62799008 |
| CCDC122 | protein_coding | 0.63375249 | 2.66144545 |
| RNF128 | protein_coding | 0.63266009 | 6.67137311 |
| IRAK3 | protein_coding | 0.63251138 | 2.55727026 |
| ARAP2 | protein_coding | 0.63199646 | 4.81441212 |
| NEK8 | protein_coding | 0.63153896 | 7.03229689 |
| RP11-7F17.8 | lncRNA | 0.6314971 | 2.87702974 |
| WFIKKN1 | protein_coding | 0.63110999 | 1.87543946 |
| LIMCH1 | protein_coding | 0.630837 | 8.98680817 |
| RP3-525N10. | lncRNA | 0.63057491 | 1.3942617 |
| LZTS3 | protein_coding | 0.63024455 | 8.84195972 |
| VANGL1 | protein_coding | 0.6299145 | 12.0225647 |
| CACNA1E | protein_coding | 0.62894004 | 2.25751871 |
| MAD1L1 | protein_coding | 0.62805405 | 1.65298982 |
| AC009264.1 | lncRNA | 0.62745918 | 3.01866069 |
| ACVR1 | protein_coding | 0.62724088 | 2.67676512 |
| CAPS2 | protein_coding | 0.627216 | 1.82717709 |
| LINC01806 | lncRNA | 0.6271505 | 4.60683312 |
| MITD1 | protein_coding | 0.62678942 | 6.8615528 |
| IFNLR1 | protein_coding | 0.62617339 | 9.30433828 |
| RBPMS | protein_coding | 0.62611803 | 1.05788612 |
| AC127904.2 | lncRNA | 0.62605987 | 0.82051023 |
| POGLUT1 | protein_coding | 0.62579153 | 7.57662255 |
| GBA2 | protein_coding | 0.62551561 | 12.0998697 |
| NBPF20 | protein_coding | 0.62530859 | 3.45582421 |
| DLX2 | protein_coding | 0.6252509 | 1.4198869 |
| CASP7 | protein_coding | 0.62523731 | 9.01757529 |
| PCDHGB6 | protein_coding | 0.6248771 | 5.9931055 |
| PDCD6IP-DT | lncRNA | 0.6243292 | 0.98752344 |
| ZNRF3 | protein_coding | 0.62409944 | 9.45904697 |
| LEMD1 | protein_coding | 0.62404908 | 0.89774328 |
| BTBD9 | protein_coding | 0.62404651 | 5.90539795 |
| VASH2 | protein_coding | 0.6239192 | 8.75438189 |
| C1orf162 | protein_coding | 0.62338099 | 0.93565638 |
| ZNF462 | protein_coding | 0.6231479 | 7.83702321 |

| | | | |
|---|---|---|---|
| FUT4 | protein_coding | 0.6229608 | 8.19982638 |
| KCNA3 | protein_coding | 0.62245313 | 5.28585498 |
| PCDHB4 | protein_coding | 0.62224598 | 1.25551316 |
| CTD-2201E18 | lncRNA | 0.62176609 | 5.76075615 |
| TMEM132A | protein_coding | 0.62152991 | 10.9486133 |
| FGD6 | protein_coding | 0.62136649 | 12.2671333 |
| ABLIM2 | protein_coding | 0.62132996 | 10.9249991 |
| PLXNA2 | protein_coding | 0.62115783 | 10.9660732 |
| MAP4 | protein_coding | 0.62087879 | 13.3296886 |
| JUP | protein_coding | 0.62084351 | 12.6474063 |
| TCEA1 | protein_coding | 0.62067504 | 9.89084289 |
| MYO6 | protein_coding | 0.61990061 | 12.0752595 |
| RAPGEF1 | protein_coding | 0.61966535 | 13.6126861 |
| MREG | protein_coding | 0.61947981 | 10.6526556 |
| KCNS3 | protein_coding | 0.61906305 | 1.37060434 |
| RCN1 | protein_coding | 0.61904285 | 11.0821981 |
| SMAD9 | protein_coding | 0.61903591 | 3.47142771 |
| NIM1K | protein_coding | 0.61894761 | 10.879378 |
| RP11-225H2 | lncRNA | 0.61877379 | 0.81616475 |
| ZNF524 | protein_coding | 0.61872027 | 4.76704003 |
| NLN | protein_coding | 0.61817021 | 11.6147894 |
| CA10 | protein_coding | 0.61773487 | 4.39229799 |
| ZMIZ2 | protein_coding | 0.61761574 | 14.0673801 |
| LNX1 | protein_coding | 0.61735261 | 2.1030383 |
| RP5-1086D1 | lncRNA | 0.61689986 | 6.94405341 |
| STPG3 | protein_coding | 0.6163855 | 2.39009797 |
| SRSF12 | protein_coding | 0.6159379 | 6.35030888 |
| PGM5 | protein_coding | 0.61453409 | 7.42882626 |
| AKIP1 | protein_coding | 0.61411854 | 10.9570826 |
| UXS1 | protein_coding | 0.61410753 | 10.4395693 |
| TERT | protein_coding | 0.61398259 | 4.77375896 |
| STPG1 | protein_coding | 0.61370739 | 7.98055068 |
| LINC02891 | lncRNA | 0.61344331 | 1.88216693 |
| GCC1 | protein_coding | 0.61341741 | 10.0344919 |
| RP4-673M15 | lncRNA | 0.61306304 | 0.98322717 |
| SAMD10 | protein_coding | 0.6121482 | 5.9463532 |
| H2BC15 | protein_coding | 0.6112624 | 1.88299645 |
| RP1-278C19. | lncRNA | 0.61123689 | 1.22582933 |
| RP4-669H2.1 | lncRNA | 0.61104128 | 1.63018644 |
| BCL11A | protein_coding | 0.61096213 | 3.2399431 |
| SPOCK2 | protein_coding | 0.61093305 | 11.5936248 |
| DGLUCY | protein_coding | 0.61067919 | 10.3120718 |
| HSPB7 | protein_coding | 0.6103296 | 1.25446978 |
| GLRX | protein_coding | 0.61008529 | 5.03589242 |
| TIGD4 | protein_coding | 0.60935518 | 2.09825875 |

| | | | |
|---|---|---|---|
| ACOT9 | protein_coding | 0.60922133 | 4.9828656 |
| PCGF5 | protein_coding | 0.60871913 | 8.94691919 |
| KHDRBS3 | protein_coding | 0.60868985 | 10.4787665 |
| ZNF546 | protein_coding | 0.60813376 | 4.64743887 |
| RHEBL1 | protein_coding | 0.60787161 | 4.9117447 |
| GTPBP2 | protein_coding | 0.60769346 | 11.8421874 |
| ZSCAN2 | protein_coding | 0.60752218 | 4.81313539 |
| PSMA5 | protein_coding | 0.6074252 | 12.2976016 |
| SALL2 | protein_coding | 0.60730326 | 13.4207688 |
| L3HYPDH | protein_coding | 0.60725518 | 4.08374155 |
| ITGB4 | protein_coding | 0.60706946 | 3.37506548 |
| GOLT1A | protein_coding | 0.60651079 | 4.30918849 |
| IFI27L2 | protein_coding | 0.60627703 | 9.26007093 |
| LINC02724 | lncRNA | 0.60536421 | 1.07389217 |
| LINC01772 | lncRNA | 0.60531305 | 1.29073243 |
| CLSTN3 | protein_coding | 0.60468292 | 9.1127486 |
| RP11-215I16 | lncRNA | 0.60442193 | 1.5171283 |
| AF131216.5 | lncRNA | 0.60440437 | 2.61569596 |
| CTBS | protein_coding | 0.60419297 | 3.40797446 |
| POLR2J4 | lncRNA | 0.60395407 | 2.78441255 |
| PCDHB5 | protein_coding | 0.60339365 | 5.81051239 |
| RP1-63G5.7 | lncRNA | 0.60335658 | 0.78598977 |
| NKD2 | protein_coding | 0.60310697 | 2.24336443 |
| RP11-1038A1 | lncRNA | 0.60282302 | 1.01622808 |
| FOXA1 | protein_coding | 0.60252753 | 1.1159096 |
| CTD-2523D13 | lncRNA | 0.60242846 | 0.89883086 |
| LRRC61 | protein_coding | 0.6022972 | 6.89895105 |
| AGRN | protein_coding | 0.6022635 | 6.5483741 |
| ZNF296 | protein_coding | 0.60213857 | 2.74077749 |
| HOXD4 | protein_coding | 0.60118865 | 6.1837228 |
| FAM171B | protein_coding | 0.60078391 | 11.3551545 |
| IGF2R | protein_coding | 0.60042652 | 12.156878 |
| GPNMB | protein_coding | 0.60029469 | 5.90567277 |
| MTRF1L | protein_coding | 0.59968485 | 5.16565633 |
| UBE2E3 | protein_coding | 0.59952236 | 12.5413554 |
| RP11-996F15 | lncRNA | 0.59857649 | 0.7463724 |
| TULP4 | protein_coding | 0.59836608 | 8.16858322 |
| LRCH1 | protein_coding | 0.59786107 | 7.33283899 |
| CH507-9B2.9 | protein_coding | 0.59727864 | 2.1653364 |
| RAB27B | protein_coding | 0.59677124 | 1.27548845 |
| CSMD3 | protein_coding | 0.59660045 | 3.4632837 |
| FAM102A | protein_coding | 0.59618381 | 12.7156182 |
| SCGN | protein_coding | 0.59614051 | 8.58445214 |
| RP11-638I2.4 | lncRNA | 0.5959899 | 0.71801042 |
| TANK | protein_coding | 0.5951959 | 9.69145656 |

| | | | |
|---|---|---|---|
| DUSP7 | protein_coding | 0.59506016 | 8.66672267 |
| C10orf143 | protein_coding | 0.59370953 | 1.98542646 |
| C6orf62 | protein_coding | 0.59310806 | 12.3119002 |
| SLC5A4-AS1 | lncRNA | 0.59251068 | 1.968476 |
| MIRLET7BHG | lncRNA | 0.59025284 | 3.44880141 |
| PLEKHA8 | protein_coding | 0.59004799 | 6.24631703 |
| HIRA | protein_coding | 0.59003619 | 8.8301546 |
| FYCO1 | protein_coding | 0.58993759 | 4.8880121 |
| ABCC1 | protein_coding | 0.58847625 | 8.50452622 |
| PATL1 | protein_coding | 0.58841932 | 11.9107346 |
| EOGT | protein_coding | 0.58743455 | 2.73262072 |
| HOXC5 | protein_coding | 0.58697139 | 0.89481363 |
| GH2 | protein_coding | 0.58683594 | 4.36129321 |
| RP11-297C4. | lncRNA | 0.58680196 | 0.88715987 |
| RP11-14H3.7 | lncRNA | 0.58670034 | 0.87195569 |
| PRR5 | protein_coding | 0.58613829 | 8.39297059 |
| ACP2 | protein_coding | 0.58574022 | 9.52364711 |
| DIMT1 | protein_coding | 0.58573651 | 7.39110843 |
| CYP20A1 | protein_coding | 0.58549076 | 9.06088883 |
| TAOK3 | protein_coding | 0.58477338 | 7.13489706 |
| CDK11A | protein_coding | 0.58447081 | 1.54814516 |
| ATP2B4 | protein_coding | 0.5830281 | 10.9949611 |
| PALLD | protein_coding | 0.58287834 | 11.4552641 |
| LIMD1-AS1 | lncRNA | 0.58239222 | 2.67447174 |
| CCDC144A | protein_coding | 0.58236407 | 2.19782489 |
| SGSM3 | protein_coding | 0.5820379 | 1.72645071 |
| FANCE | protein_coding | 0.58194217 | 9.74096326 |
| RP1-197B17. | lncRNA | 0.58168138 | 1.0665008 |
| PGM5-AS1 | lncRNA | 0.58155541 | 1.09342512 |
| ZNF503 | protein_coding | 0.58104669 | 4.07747738 |
| DCLRE1C | protein_coding | 0.58102629 | 6.81715493 |
| SLC30A2 | protein_coding | 0.5807857 | 10.8067554 |
| RP3-467K16. | lncRNA | 0.58059311 | 1.02933574 |
| PILRB | protein_coding | 0.58009515 | 4.35042732 |
| CDK17 | protein_coding | 0.58009209 | 8.90574834 |
| ERFE | protein_coding | 0.57934223 | 3.00723644 |
| TSPAN33 | protein_coding | 0.57929589 | 7.84847989 |
| LINC01004 | lncRNA | 0.57894663 | 2.15738876 |
| TYW5 | protein_coding | 0.57882611 | 8.1395458 |
| GPR63 | protein_coding | 0.57862919 | 3.9038287 |
| SGCD | protein_coding | 0.57824997 | 6.96816643 |
| RP11-305L7. | lncRNA | 0.57822879 | 1.24188777 |
| CTD-2054N2 | lncRNA | 0.57799008 | 2.56558862 |
| RNF114 | protein_coding | 0.57778647 | 12.776717 |
| RGS10 | protein_coding | 0.5766838 | 8.06164772 |

| | | | |
|---|---|---|---|
| PSMC2 | protein_coding | 0.57655266 | 11.8494519 |
| RFPL1S | lncRNA | 0.57602049 | 8.23851321 |
| FOS | protein_coding | 0.57600147 | 1.67379038 |
| ST7 | protein_coding | 0.57597392 | 7.59950603 |
| MAPK11 | protein_coding | 0.57596523 | 5.67223959 |
| TSPAN4 | protein_coding | 0.57583062 | 9.55969272 |
| HESX1 | protein_coding | 0.57582221 | 1.72077414 |
| ELOVL7 | protein_coding | 0.57572157 | 9.47271374 |
| SNHG27 | lncRNA | 0.57560252 | 1.36115466 |
| RP11-715J22 | lncRNA | 0.57524794 | 1.47182721 |
| LRP8 | protein_coding | 0.5749176 | 13.6206772 |
| DOCK10 | protein_coding | 0.57470092 | 10.8857688 |
| ZNF583 | protein_coding | 0.57450403 | 5.60043303 |
| CARD8 | protein_coding | 0.57419938 | 7.94504094 |
| IFNGR1 | protein_coding | 0.57386557 | 5.19085422 |
| SAMD14 | protein_coding | 0.57335706 | 1.69729946 |
| ZDHHC1 | protein_coding | 0.57291615 | 3.81817615 |
| STX17 | protein_coding | 0.57268233 | 10.946085 |
| SUCLG2 | protein_coding | 0.57262478 | 7.07068493 |
| CTH | protein_coding | 0.57206563 | 2.67273199 |
| FAM13C | protein_coding | 0.57175863 | 5.35052698 |
| MB | protein_coding | 0.57175013 | 2.98708417 |
| GPC4 | protein_coding | 0.57161102 | 1.35478352 |
| CES3 | protein_coding | 0.57159431 | 9.2084943 |
| SCAMP1-AS1 | lncRNA | 0.57122495 | 6.99974484 |
| PHLDA1 | protein_coding | 0.57085629 | 8.7419515 |
| SLC31A2 | protein_coding | 0.57066905 | 4.6290835 |
| RP11-55K13. | lncRNA | 0.5705348 | 0.60102067 |
| MON2 | protein_coding | 0.57048939 | 11.5422353 |
| BACH1 | protein_coding | 0.5702445 | 9.89029213 |
| DYRK3 | protein_coding | 0.57013456 | 5.47021822 |
| DMBX1 | protein_coding | 0.57006786 | 1.03791161 |
| PCDHGA4 | protein_coding | 0.56988951 | 1.42543726 |
| LINC01176 | lncRNA | 0.56968285 | 2.65090389 |
| WSCD2 | protein_coding | 0.56939447 | 9.66955377 |
| ITSN2 | protein_coding | 0.56914208 | 10.8763041 |
| AF064858.8 | lncRNA | 0.56897017 | 1.17688493 |
| UCHL1 | protein_coding | 0.56894993 | 14.5858558 |
| AC008269.2 | lncRNA | 0.56894605 | 0.71550684 |
| CTC-444N24. | lncRNA | 0.56876398 | 4.95890692 |
| PATJ | protein_coding | 0.56866183 | 6.83421896 |
| PAUPAR | lncRNA | 0.56806861 | 2.91984657 |
| KLHDC9 | protein_coding | 0.56782365 | 2.09199505 |
| LYPD5 | protein_coding | 0.56767257 | 5.03186443 |
| CHN1 | protein_coding | 0.5676207 | 11.6890791 |

| | | | |
|---|---|---|---|
| RNF19B | protein_coding | 0.56742363 | 9.75089666 |
| LRRC49 | protein_coding | 0.5668122 | 5.31142105 |
| CTD-2006H1· | lncRNA | 0.56664743 | 1.94206846 |
| CACNA2D3 | protein_coding | 0.5664564 | 1.09586729 |
| PSMD11 | protein_coding | 0.56616197 | 13.2211344 |
| RP11-315D1( | protein_coding | 0.56552387 | 3.3005559 |
| LSR | protein_coding | 0.5653548 | 10.5251526 |
| FFAR2 | protein_coding | 0.5653448 | 1.09603276 |
| CYP4F26P | lncRNA | 0.56363756 | 6.57088563 |
| LIN7A | protein_coding | 0.56300484 | 3.35087946 |
| TOM1L1 | protein_coding | 0.562659 | 9.98427888 |
| FTL | protein_coding | 0.56227748 | 15.049956 |
| PAX9 | protein_coding | 0.56227671 | 3.87928492 |
| C4orf50 | protein_coding | 0.56210464 | 4.37745115 |
| RAB12 | protein_coding | 0.56190237 | 8.34073727 |
| AFF1 | protein_coding | 0.56177197 | 11.5793446 |
| SLC9A3R1 | protein_coding | 0.56150979 | 9.11170077 |
| NLRP11 | protein_coding | 0.56142196 | 5.51172669 |
| ZNF425 | protein_coding | 0.5612289 | 3.79084697 |
| COL22A1 | protein_coding | 0.56093877 | 11.3994527 |
| ADCY7 | protein_coding | 0.56085324 | 3.86043209 |
| MIDEAS | protein_coding | 0.56049649 | 12.4452542 |
| MYH3 | protein_coding | 0.56021682 | 2.10150222 |
| LINC00698 | lncRNA | 0.56021522 | 2.84975178 |
| GLTP | protein_coding | 0.56018139 | 9.00052873 |
| ACTRT3 | protein_coding | 0.56016545 | 1.87491711 |
| RP11-274B2: | lncRNA | 0.55997887 | 2.36541349 |
| GSEC | lncRNA | 0.55992614 | 3.19583283 |
| MICALL2 | protein_coding | 0.55943838 | 8.63033688 |
| TPP1 | protein_coding | 0.55864447 | 9.1135725 |
| PPP3CB-AS1 | lncRNA | 0.55726799 | 1.68909495 |
| CYB561 | protein_coding | 0.55706697 | 9.95304273 |
| LYG1 | protein_coding | 0.55697602 | 2.23558995 |
| PSAT1 | protein_coding | 0.55680439 | 12.8406326 |
| ZCCHC2 | protein_coding | 0.55637501 | 9.96702768 |
| KDM4D | protein_coding | 0.55626489 | 1.87396766 |
| ACTN1 | protein_coding | 0.55601871 | 12.1010017 |
| FTH1 | protein_coding | 0.55560307 | 9.40189795 |
| ZXDB | protein_coding | 0.5555065 | 6.05738666 |
| HPSE2 | protein_coding | 0.55531434 | 0.80981343 |
| TSC22D1 | protein_coding | 0.55519749 | 11.3446671 |
| UBE2H | protein_coding | 0.55430491 | 10.996921 |
| FRMD4B | protein_coding | 0.55429155 | 4.27997222 |
| LUNAR1 | lncRNA | 0.55412789 | 1.12600834 |
| SCARB2 | protein_coding | 0.55403592 | 14.0545557 |

| | | | |
|---|---|---|---|
| ZNF43 | protein_coding | 0.55400665 | 6.18872821 |
| SLC25A12 | protein_coding | 0.55396639 | 9.48795742 |
| ATP8B2 | protein_coding | 0.55361621 | 14.4379396 |
| NFKBIB | protein_coding | 0.55350501 | 7.8362847 |
| PCDHA1 | protein_coding | 0.55312071 | 0.78991985 |
| ANXA9 | protein_coding | 0.55277125 | 1.85332859 |
| PKNOX2 | protein_coding | 0.55267388 | 1.47348389 |
| PCDHA2 | protein_coding | 0.55261459 | 3.6793185 |
| RAP1GAP2 | protein_coding | 0.55246752 | 11.1597068 |
| PCNX2 | protein_coding | 0.55228512 | 7.99877819 |
| SLC6A2 | protein_coding | 0.5517032 | 5.27915153 |
| ADAM17 | protein_coding | 0.55163173 | 9.69264547 |
| RP11-344B2: | lncRNA | 0.55087707 | 1.12990294 |
| HOXD10 | protein_coding | 0.54987298 | 9.34026485 |
| RP11-421A1( | lncRNA | 0.54969548 | 1.61874829 |
| SLC35F6 | protein_coding | 0.54945726 | 9.09372064 |
| RP1-80N2.2 | lncRNA | 0.54903653 | 1.70748838 |
| ADCY5 | protein_coding | 0.54825557 | 4.19981396 |
| TAGAP-AS1 | lncRNA | 0.54729184 | 1.67876768 |
| MTX3 | protein_coding | 0.54699446 | 10.1661023 |
| TNRC6C | protein_coding | 0.54640356 | 9.98111358 |
| GCNA | protein_coding | 0.54584089 | 1.81598252 |
| SMIM29 | protein_coding | 0.54542785 | 8.28496137 |
| NMRK1 | protein_coding | 0.54534259 | 5.86047394 |
| PLAA | protein_coding | 0.54528288 | 11.5825182 |
| RP3-368A4.6 | lncRNA | 0.54485561 | 2.66538492 |
| TMEM268 | protein_coding | 0.54477831 | 9.54353287 |
| SPHK1 | protein_coding | 0.5446664 | 7.19743526 |
| ABCA3 | protein_coding | 0.54455028 | 10.7460719 |
| CSRNP1 | protein_coding | 0.54408769 | 7.46987125 |
| LRRC37B | protein_coding | 0.54369642 | 7.50621749 |
| KCNJ14 | protein_coding | 0.54360298 | 2.34542034 |
| AC025171.1 | lncRNA | 0.54307708 | 5.81236079 |
| AGBL3 | protein_coding | 0.54289191 | 1.24099599 |
| TATDN1 | protein_coding | 0.5419732 | 7.47964005 |
| FAM227A | protein_coding | 0.54160719 | 3.07400037 |
| VWDE | protein_coding | 0.54115764 | 8.10310948 |
| ZBED6 | protein_coding | 0.54077095 | 0.84367111 |
| CCDC93 | protein_coding | 0.54048096 | 10.6286325 |
| SH3BP4 | protein_coding | 0.54031476 | 12.4467171 |
| RP11-122G1 | lncRNA | 0.54024457 | 4.44715274 |
| RP11-421L21 | lncRNA | 0.54010649 | 4.63462345 |
| SGPL1 | protein_coding | 0.53992248 | 9.5910921 |
| CNR1 | protein_coding | 0.53990671 | 8.05824394 |
| SLC2A6 | protein_coding | 0.53975203 | 5.88256477 |

| | | | |
|---|---|---|---|
| PLEC | protein_coding | 0.53885749 | 5.65042197 |
| RAB9A | protein_coding | 0.53870884 | 7.10063673 |
| CHRNB4 | protein_coding | 0.53859207 | 2.06341882 |
| SETDB2 | protein_coding | 0.53838482 | 5.98557255 |
| AP000692.10 | lncRNA | 0.53832643 | 0.65176744 |
| C2 | protein_coding | 0.53819218 | 1.205038 |
| DZIP3 | protein_coding | 0.53800393 | 7.79952491 |
| RAB36 | protein_coding | 0.53776943 | 4.93168502 |
| ZC3H12C | protein_coding | 0.53767587 | 7.95032444 |
| MIR217HG | lncRNA | 0.5372786 | 1.33098978 |
| DYNC1H1 | protein_coding | 0.53721708 | 14.8213291 |
| FAM106A | lncRNA | 0.53690771 | 2.46143036 |
| RP11-187K14 | lncRNA | 0.5369067 | 1.39251571 |
| UNC79 | protein_coding | 0.53683389 | 11.6418854 |
| AGAP2 | protein_coding | 0.5364345 | 5.72189634 |
| ENC1 | protein_coding | 0.53618333 | 11.5074238 |
| CCDC106 | protein_coding | 0.53576515 | 6.78564398 |
| ZNF547 | protein_coding | 0.53537216 | 4.38562248 |
| NPIPB4 | protein_coding | 0.53500529 | 2.64915635 |
| CTD-2655K5. | lncRNA | 0.53498872 | 0.76418505 |
| RP11-98F14. | lncRNA | 0.53450506 | 1.05345764 |
| WDR13 | protein_coding | 0.53441093 | 8.81984337 |
| MAFG | protein_coding | 0.53435301 | 9.97088452 |
| GCA | protein_coding | 0.5342326 | 8.47884837 |
| MINPP1 | protein_coding | 0.53401007 | 5.43306782 |
| SLC2A5 | protein_coding | 0.53391979 | 9.2230095 |
| ADAM9 | protein_coding | 0.53370853 | 8.94271704 |
| LYPD6B | protein_coding | 0.53331338 | 6.2131055 |
| RP1-197B17. | lncRNA | 0.53276988 | 0.79440634 |
| HR | protein_coding | 0.53145611 | 7.66496624 |
| RP1-239B22. | lncRNA | 0.53116511 | 3.48880933 |
| RP11-248J18 | lncRNA | 0.5307097 | 0.92256651 |
| PSMA4 | protein_coding | 0.53031443 | 11.8028178 |
| RAB27A | protein_coding | 0.5301594 | 9.17551625 |
| C3orf18 | protein_coding | 0.53003028 | 7.23157804 |
| PPIC | protein_coding | 0.52977276 | 8.07930154 |
| PWWP2B | protein_coding | 0.52973259 | 4.1409396 |
| CILP2 | protein_coding | 0.52940261 | 2.42316772 |
| PELI1 | protein_coding | 0.52938829 | 6.02940569 |
| KCNK12 | protein_coding | 0.52932247 | 3.91426206 |
| IBA57 | protein_coding | 0.52910271 | 8.57124695 |
| CTD-3138B18 | lncRNA | 0.52868307 | 1.13656859 |
| GPCPD1 | protein_coding | 0.52838606 | 6.62614979 |
| ABAT | protein_coding | 0.52837803 | 2.2839843 |
| MTMR10 | protein_coding | 0.52830705 | 8.26392369 |

| | | | |
|---|---|---|---|
| FRMD4A | protein_coding | 0.52787157 | 5.98264281 |
| PEAR1 | protein_coding | 0.5276099 | 2.23467386 |
| PFKL | protein_coding | 0.52729955 | 9.8656406 |
| GXYLT2 | protein_coding | 0.52677604 | 1.58088808 |
| ZNF217 | protein_coding | 0.52675292 | 8.92689784 |
| LINC02615 | lncRNA | 0.5265786 | 1.36032731 |
| TRIM44 | protein_coding | 0.52643495 | 13.5241944 |
| PCDHA10 | protein_coding | 0.5256214 | 1.23679229 |
| PRH1 | protein_coding | 0.52545856 | 0.89229379 |
| PCDHB3 | protein_coding | 0.52502819 | 1.10852513 |
| ZNF528 | protein_coding | 0.5248464 | 0.99652628 |
| PACS1 | protein_coding | 0.52472613 | 11.0995612 |
| ETFA | protein_coding | 0.52368159 | 11.9891621 |
| RP3-508I15.1 | lncRNA | 0.52187442 | 0.90594199 |
| TMEM255A | protein_coding | 0.52174133 | 1.92755988 |
| CCDC180 | protein_coding | 0.52080065 | 1.52299173 |
| ATP5MK | protein_coding | 0.52059087 | 9.50972769 |
| PVR | protein_coding | 0.52028785 | 8.5781127 |
| TRPM4 | protein_coding | 0.52023739 | 2.50549086 |
| ZNF699 | protein_coding | 0.52009209 | 5.10020298 |
| BMP2K | protein_coding | 0.51840387 | 8.88111839 |
| ASAH2 | protein_coding | 0.51805602 | 2.21234806 |
| SNHG4 | lncRNA | 0.51800122 | 3.28192435 |
| CTXN2 | protein_coding | 0.51745412 | 1.6752913 |
| RAC2 | protein_coding | 0.51743838 | 1.2065628 |
| ZNF558 | protein_coding | 0.51722135 | 6.61720734 |
| ZBTB22 | protein_coding | 0.51707224 | 6.31932878 |
| EVC | protein_coding | 0.51697439 | 8.95696812 |
| PCDHGA10 | protein_coding | 0.51672792 | 5.54400691 |
| VOPP1 | protein_coding | 0.51585145 | 11.7723295 |
| NKAIN2 | protein_coding | 0.51582517 | 3.82975929 |
| SNCG | protein_coding | 0.51582325 | 5.86227514 |
| SYTL3 | protein_coding | 0.51570021 | 4.99165093 |
| INO80C | protein_coding | 0.51508379 | 7.05105213 |
| GPR107 | protein_coding | 0.51426393 | 12.7527365 |
| ZNF333 | protein_coding | 0.51424571 | 4.96655585 |
| XXbac-B562F | lncRNA | 0.51421557 | 1.17179352 |
| CAPN1 | protein_coding | 0.51399913 | 10.2659687 |
| UBR2 | protein_coding | 0.51368335 | 9.75417483 |
| NPIPA5 | protein_coding | 0.51311907 | 4.03412318 |
| FAM133A | protein_coding | 0.51311553 | 1.28247736 |
| TAS2R5 | protein_coding | 0.5130774 | 0.74107899 |
| NIPAL1 | protein_coding | 0.51303865 | 1.02209755 |
| IQCH | protein_coding | 0.51296262 | 4.27647018 |
| ARL14EP | protein_coding | 0.51280786 | 8.64103607 |

| | | | |
|---|---|---|---|
| TARBP1 | protein_coding | 0.51271595 | 9.76271345 |
| CH17-264L24 | lncRNA | 0.51263891 | 0.89693019 |
| SSH3 | protein_coding | 0.51247244 | 2.62033798 |
| PCDHGA11 | protein_coding | 0.51243364 | 1.04527127 |
| NRG1 | protein_coding | 0.51242809 | 3.71255081 |
| B3GALNT1 | protein_coding | 0.5121467 | 2.84556807 |
| PVT1 | lncRNA | 0.51198731 | 4.56281364 |
| RP11-69L16. | lncRNA | 0.51182727 | 0.70810006 |
| DRICH1 | protein_coding | 0.51156142 | 0.92840308 |
| LMX1B | protein_coding | 0.51140621 | 2.18312077 |
| BCO2 | protein_coding | 0.51114706 | 0.75933501 |
| RP1-178F15. | lncRNA | 0.51109348 | 0.8921204 |
| EGFL8 | protein_coding | 0.51102258 | 0.86245141 |
| LINC00963 | lncRNA | 0.510958 | 5.85588777 |
| RHOF | protein_coding | 0.51092733 | 5.55610933 |
| NRAV | lncRNA | 0.51079789 | 4.49523 |
| ALPG | protein_coding | 0.51065216 | 4.89782126 |
| RP4-564F22. | lncRNA | 0.51044199 | 0.76142913 |
| MESTIT1 | lncRNA | 0.51028619 | 0.64009109 |
| GABRR2 | protein_coding | 0.51005002 | 0.50524329 |
| ZNF506 | protein_coding | 0.50978507 | 7.4283606 |
| SMPD3 | protein_coding | 0.50917797 | 3.02620451 |
| AGK | protein_coding | 0.50886149 | 9.32451433 |
| CHGA | protein_coding | 0.50842379 | 9.48555626 |
| SLC9A8 | protein_coding | 0.50736799 | 7.46575884 |
| PLIN5 | protein_coding | 0.50724275 | 1.05998546 |
| FHIT | protein_coding | 0.50661445 | 1.69878189 |
| SERTAD1 | protein_coding | 0.5061545 | 2.10549925 |
| MIPOL1 | protein_coding | 0.50606648 | 4.73046799 |
| TTYH2 | protein_coding | 0.50605372 | 6.21342044 |
| LMO4 | protein_coding | 0.50561588 | 8.14300973 |
| KDM1B | protein_coding | 0.50555024 | 11.3805797 |
| MCCC2 | protein_coding | 0.50532788 | 9.44953252 |
| CMYA5 | protein_coding | 0.5051459 | 1.10741209 |
| RP11-499P2( | lncRNA | 0.50492287 | 1.29335306 |
| CTD-2540F1: | lncRNA | 0.50414229 | 1.70023083 |
| KANSL1L | protein_coding | 0.50409141 | 5.22694408 |
| TPK1 | protein_coding | 0.50403138 | 3.17638512 |
| HYPK | protein_coding | 0.5034493 | 1.92741504 |
| NDRG4 | protein_coding | 0.50335228 | 11.7215213 |
| RP11-342K6. | lncRNA | 0.50298649 | 1.01205303 |
| DIABLO | protein_coding | 0.50280592 | 2.1412946 |
| LRP10 | protein_coding | 0.50216795 | 5.88476252 |
| SDCBP2-AS1 | lncRNA | 0.5020584 | 3.61100353 |
| TMEM88 | protein_coding | 0.50187014 | 1.31731122 |

| | | | |
|---|---|---|---|
| ZGLP1 | protein_coding | 0.50180826 | 2.36258707 |
| ZMYM5 | protein_coding | 0.50130068 | 5.25224053 |
| CCDC146 | protein_coding | 0.50116867 | 2.35360667 |
| SH3BP5-AS1 | lncRNA | 0.5010355 | 1.99439006 |
| GNPDA1 | protein_coding | 0.50095372 | 9.71218397 |
| PIM2 | protein_coding | 0.50047669 | 5.38371041 |
| RP11-488C13 | lncRNA | 0.50040026 | 1.08139078 |
| NRIP1 | protein_coding | 0.50027476 | 6.86220784 |
| PHYH | protein_coding | 0.49992591 | 4.64355127 |
| ALS2 | protein_coding | 0.49986786 | 10.0621765 |
| MALAT1 | lncRNA | 0.49956501 | 12.0617054 |
| GPR158-AS1 | lncRNA | 0.49892836 | 0.65573293 |
| ERICD | lncRNA | 0.4987912 | 1.17826326 |
| QRSL1 | protein_coding | 0.49874696 | 6.4009648 |
| C1orf56 | protein_coding | 0.49874546 | 1.12006953 |
| PCED1B | protein_coding | 0.49871633 | 2.66437406 |
| KIF1A | protein_coding | 0.49866754 | 13.0082323 |
| DCX | protein_coding | 0.49861226 | 8.89645579 |
| NKILA | lncRNA | 0.49851413 | 1.35041851 |
| YY2 | protein_coding | 0.4981424 | 0.83989786 |
| LA16c-306E5 | protein_coding | 0.49788322 | 1.18412763 |
| MAPKBP1 | protein_coding | 0.49781102 | 9.32562911 |
| ZNF385D | protein_coding | 0.49752493 | 0.74050476 |
| MICALL1 | protein_coding | 0.49743944 | 8.59157213 |
| DOCK9-DT | lncRNA | 0.49682019 | 1.69040668 |
| RP11-73E17. | lncRNA | 0.49597746 | 2.78507616 |
| RP11-422P24 | lncRNA | 0.49597167 | 1.34451634 |
| NXN | protein_coding | 0.49552721 | 9.74780122 |
| AC019118.3 | lncRNA | 0.49550992 | 0.64808955 |
| DAP3 | protein_coding | 0.4953776 | 11.1643514 |
| CTD-2201E18 | lncRNA | 0.49510063 | 0.91935057 |
| PLPP7 | protein_coding | 0.49500611 | 3.15664955 |
| SLC5A3 | protein_coding | 0.49491304 | 7.37345199 |
| MIOS | protein_coding | 0.4947268 | 7.81305241 |
| ATF4 | protein_coding | 0.49471939 | 14.1182131 |
| TMEM121B | protein_coding | 0.49443833 | 8.66095304 |
| LINC01117 | lncRNA | 0.49443398 | 1.73230689 |
| RASAL1 | protein_coding | 0.49422193 | 7.63256835 |
| ADPRM | protein_coding | 0.4930139 | 3.12424867 |
| WASHC4 | protein_coding | 0.49299532 | 7.68289864 |
| NOMO3 | protein_coding | 0.49288229 | 4.28789316 |
| CRYBG3 | protein_coding | 0.49263993 | 13.7012883 |
| LINC02709 | lncRNA | 0.49255119 | 4.42599693 |
| ALDH3B1 | protein_coding | 0.49237188 | 4.19581904 |
| NCS1 | protein_coding | 0.49230676 | 11.2209234 |

| | | | |
|---|---|---|---|
| DNAJB5 | protein_coding | 0.49227875 | 6.25803389 |
| ARNTL2 | protein_coding | 0.49202556 | 6.89509697 |
| MYO1D | protein_coding | 0.49191784 | 11.7152122 |
| MED13L | protein_coding | 0.49187792 | 11.3583681 |
| SEPTIN11 | protein_coding | 0.49175963 | 12.0610211 |
| EPB41L3 | protein_coding | 0.49140401 | 9.5202955 |
| CNTN2 | protein_coding | 0.49082677 | 11.6140792 |
| PPM1M | protein_coding | 0.49020375 | 4.96948779 |
| SLC27A1 | protein_coding | 0.48929615 | 1.52794533 |
| ZNF827 | protein_coding | 0.4888462 | 8.30607105 |
| HOXD11 | protein_coding | 0.48866827 | 4.53663891 |
| CTC-459F4.1 | lncRNA | 0.4886382 | 0.77882396 |
| MEX3A | protein_coding | 0.48861185 | 11.1167635 |
| USF1 | protein_coding | 0.48855513 | 9.54712817 |
| BLVRA | protein_coding | 0.48784698 | 9.9127328 |
| CTC-260F20.: | protein_coding | 0.48766327 | 0.9454647 |
| DND1 | protein_coding | 0.48764461 | 2.02736588 |
| DUSP5 | protein_coding | 0.48717179 | 1.0743245 |
| TAGLN2 | protein_coding | 0.48687599 | 12.7265326 |
| PARD6B | protein_coding | 0.48638996 | 3.6241997 |
| EEF2K | protein_coding | 0.48616776 | 10.3393874 |
| MYO10 | protein_coding | 0.48616182 | 11.5761628 |
| EBF1 | protein_coding | 0.48557494 | 9.41416434 |
| ENOX1 | protein_coding | 0.48531139 | 2.73270087 |
| PBX1 | protein_coding | 0.48512289 | 8.02304543 |
| SBF2 | protein_coding | 0.48420568 | 10.4514953 |
| ANKRD23 | protein_coding | 0.48409842 | 1.7570093 |
| RP11-845M1 | lncRNA | 0.48399324 | 0.86483903 |
| RP11-506M1 | lncRNA | 0.48378153 | 0.79545495 |
| FIRRE | lncRNA | 0.48368223 | 6.10944624 |
| MIOS-DT | lncRNA | 0.48353998 | 0.82554592 |
| STXBP5 | protein_coding | 0.4835247 | 5.41586522 |
| RP11-195F1( | lncRNA | 0.48336146 | 3.12595726 |
| RP5-858B6.3 | lncRNA | 0.48278978 | 0.78568398 |
| HIVEP3 | protein_coding | 0.48250271 | 6.25420947 |
| RRP12 | protein_coding | 0.48224401 | 7.65551595 |
| IL10RA | protein_coding | 0.48214182 | 1.10074983 |
| HM13-IT1 | lncRNA | 0.48205036 | 0.63953564 |
| RPS6KA2 | protein_coding | 0.48187304 | 0.61117952 |
| AC138035.2 | lncRNA | 0.48142993 | 0.84906306 |
| TGFBR3 | protein_coding | 0.4813776 | 5.86539464 |
| COLEC11 | protein_coding | 0.481303 | 0.6196861 |
| RP11-17A1.3 | lncRNA | 0.48117152 | 0.49618537 |
| SNHG15 | lncRNA | 0.48096739 | 9.09998057 |
| INO80 | protein_coding | 0.4804491 | 9.08949303 |

| | | | |
|---|---|---|---|
| LYPLAL1 | protein_coding | 0.48029392 | 6.27997973 |
| CERKL | protein_coding | 0.48007765 | 1.33063338 |
| PELO | protein_coding | 0.47954167 | 7.48859487 |
| MEGF10 | protein_coding | 0.4793455 | 1.28541397 |
| TLK2 | protein_coding | 0.47932203 | 9.07504389 |
| DOCK2 | protein_coding | 0.4786327 | 0.76248734 |
| BRAF | protein_coding | 0.47851608 | 9.5252786 |
| KLC2 | protein_coding | 0.47826953 | 10.6477528 |
| TMEM270 | protein_coding | 0.47780708 | 0.74968201 |
| SYNJ2BP | protein_coding | 0.47739742 | 10.0624037 |
| KB-1836B5.1 | lncRNA | 0.47731887 | 0.61593174 |
| NOMO1 | protein_coding | 0.47715035 | 8.84376645 |
| IRAK1BP1 | protein_coding | 0.4767352 | 4.91994224 |
| SNORC | protein_coding | 0.47665002 | 1.56610233 |
| MIR137HG | lncRNA | 0.47651064 | 2.33409153 |
| ZNF493 | protein_coding | 0.47648672 | 2.1388895 |
| PLEKHF2 | protein_coding | 0.47631007 | 5.65604422 |
| SLC47A1 | protein_coding | 0.47560655 | 7.35304654 |
| PCF11-AS1 | lncRNA | 0.47555256 | 0.88920355 |
| CYSRT1 | protein_coding | 0.47540464 | 1.24992469 |
| EHD4 | protein_coding | 0.47538473 | 5.10913858 |
| TET2 | protein_coding | 0.47499652 | 9.3853917 |
| NUDT16 | protein_coding | 0.47493517 | 7.70503404 |
| ABCB6 | protein_coding | 0.47390203 | 3.73915609 |
| RP5-1090P18 | lncRNA | 0.47384028 | 1.74633832 |
| ATP1B2 | protein_coding | 0.47370678 | 1.32034261 |
| ZCCHC14 | protein_coding | 0.47350504 | 10.6361042 |
| GRAMD2B | protein_coding | 0.47302873 | 4.56162124 |
| PLXNB1 | protein_coding | 0.47302828 | 7.55916048 |
| RP5-894A10. | lncRNA | 0.47265693 | 2.36518082 |
| HOXA9 | protein_coding | 0.47247849 | 8.50930547 |
| FAM24B | protein_coding | 0.47239658 | 3.32063736 |
| RP11-23P13. | lncRNA | 0.47210828 | 3.37141571 |
| SPIDR | protein_coding | 0.4720765 | 10.0808215 |
| RFX5-AS1 | lncRNA | 0.47174653 | 2.34950894 |
| MATN4 | protein_coding | 0.47102606 | 0.83293094 |
| RARA-AS1 | lncRNA | 0.47070111 | 0.69218333 |
| EVC2 | protein_coding | 0.47058332 | 5.84956493 |
| PANX2 | protein_coding | 0.47050854 | 4.6005543 |
| ALDH1L2 | protein_coding | 0.47044512 | 1.05988968 |
| GPATCH2 | protein_coding | 0.47036746 | 5.34583211 |
| ZFP64 | protein_coding | 0.47024683 | 7.54827925 |
| RP11-298J23 | protein_coding | 0.47013805 | 1.41910584 |
| RP5-1182A14 | lncRNA | 0.46980823 | 2.02030037 |
| ZFAND5 | protein_coding | 0.46953338 | 11.413453 |

| | | | |
|---|---|---|---|
| AKAP3 | protein_coding | 0.46948929 | 0.62396286 |
| RP5-1119A7. | lncRNA | 0.46931773 | 1.22200718 |
| LMF1 | protein_coding | 0.46917608 | 5.3379825 |
| PCDHGA7 | protein_coding | 0.46914557 | 0.87755913 |
| YAE1 | protein_coding | 0.46883558 | 6.43918441 |
| NPIPB11 | protein_coding | 0.46815626 | 0.40297798 |
| RP11-423P1( | lncRNA | 0.46770676 | 0.81745422 |
| ATXN2L | protein_coding | 0.46751036 | 12.73074 |
| ZNF724 | protein_coding | 0.46726743 | 2.76536102 |
| RP11-497E1( | lncRNA | 0.46701514 | 0.94455998 |
| MUCL3 | protein_coding | 0.46674678 | 0.93329252 |
| SLC39A1 | protein_coding | 0.46616526 | 2.05142786 |
| ZNF780B | protein_coding | 0.46615229 | 6.72175192 |
| TPMT | protein_coding | 0.4652501 | 8.60849732 |
| NSFL1C | protein_coding | 0.46457767 | 9.60494756 |
| ZNF592 | protein_coding | 0.46421469 | 11.85624 |
| FBXO10 | protein_coding | 0.46393205 | 3.56407583 |
| RP5-991G20. | lncRNA | 0.46378704 | 0.68531264 |
| FREM2 | protein_coding | 0.46374665 | 1.64421715 |
| ZDHHC14 | protein_coding | 0.46360162 | 6.60683828 |
| HHIPL1 | protein_coding | 0.46281684 | 1.274405 |
| PLCB4 | protein_coding | 0.46253044 | 4.75365517 |
| SNHG31 | lncRNA | 0.46178017 | 3.79379486 |
| KLK1 | protein_coding | 0.46121409 | 1.63941205 |
| CFAP44 | protein_coding | 0.46023823 | 1.67361435 |
| LRRC23 | protein_coding | 0.46014937 | 1.82816928 |
| AZI2 | protein_coding | 0.45984637 | 7.15991045 |
| DMGDH | protein_coding | 0.45965755 | 1.09173264 |
| PPIP5K1 | protein_coding | 0.45943123 | 7.52254542 |
| RP11-337N6. | lncRNA | 0.45907392 | 0.58794381 |
| FGF5 | protein_coding | 0.45882644 | 6.61284809 |
| ZNF221 | protein_coding | 0.45870719 | 1.85843023 |
| SAPCD1 | protein_coding | 0.45846341 | 0.87319493 |
| MSLNL | protein_coding | 0.4583439 | 2.25171557 |
| LINC00634 | protein_coding | 0.45809057 | 2.90800598 |
| ARHGAP27P: | lncRNA | 0.45804877 | 1.7456474 |
| USP46 | protein_coding | 0.45801772 | 0.97048378 |
| TAF4B | protein_coding | 0.45768015 | 5.68558188 |
| SCN3B | protein_coding | 0.45700436 | 7.10251565 |
| NHLH1 | protein_coding | 0.45680277 | 12.9437997 |
| ARHGEF17 | protein_coding | 0.4564105 | 9.90737323 |
| NIN | protein_coding | 0.45622476 | 10.3268105 |
| TMEM200C | protein_coding | 0.45601422 | 4.1230458 |
| MSRB1 | protein_coding | 0.4557875 | 8.17507517 |
| VAV1 | protein_coding | 0.45563189 | 6.65576862 |

| | | | |
|---|---|---|---|
| EMX2 | protein_coding | 0.45526972 | 6.08823287 |
| FBXO30 | protein_coding | 0.45517421 | 4.54805322 |
| FTX | lncRNA | 0.455014 | 4.65521713 |
| DUSP22 | protein_coding | 0.45471899 | 9.14471305 |
| OTUD7B | protein_coding | 0.454587 | 6.81671118 |
| ZNF674 | protein_coding | 0.45357174 | 3.28068494 |
| NPRL2 | protein_coding | 0.45333789 | 4.66912258 |
| MAP4K4 | protein_coding | 0.45280763 | 14.0341598 |
| PMEPA1 | protein_coding | 0.45270109 | 4.44881499 |
| KHDRBS2-OT | lncRNA | 0.45258026 | 0.59245945 |
| KLHL3 | protein_coding | 0.45247007 | 4.36760622 |
| AC011330.13 | lncRNA | 0.4523967 | 0.59769826 |
| PKD1P6-NPIF | lncRNA | 0.4523933 | 1.88383774 |
| ZNF582-DT | lncRNA | 0.45235212 | 1.52763832 |
| FLCN | protein_coding | 0.4517809 | 5.31041686 |
| ZNF429 | protein_coding | 0.45170842 | 0.59124236 |
| C17orf67 | protein_coding | 0.45170645 | 2.84156755 |
| FAIM2 | protein_coding | 0.45141899 | 4.69954671 |
| HCN3 | protein_coding | 0.4512625 | 7.46785364 |
| LY6G5C | protein_coding | 0.45075588 | 1.08294494 |
| CTA-217C2.2 | lncRNA | 0.45054436 | 0.65512897 |
| DYNC2I1 | protein_coding | 0.45030981 | 5.55638829 |
| BCAR3 | protein_coding | 0.45028132 | 1.89286289 |
| DBF4B | protein_coding | 0.45020604 | 6.078152 |
| STK33 | protein_coding | 0.45011954 | 4.31295675 |
| NR1D1 | protein_coding | 0.44963322 | 4.17686719 |
| TICAM1 | protein_coding | 0.44963151 | 5.6222629 |
| MAPT-AS1 | lncRNA | 0.44954536 | 1.62268305 |
| ADGRB2 | protein_coding | 0.44942582 | 7.63762991 |
| RARB | protein_coding | 0.44939553 | 2.73756851 |
| RIF1 | protein_coding | 0.44931289 | 12.5667828 |
| PARP6 | protein_coding | 0.44915341 | 10.4545958 |
| RP11-134G8. | lncRNA | 0.44886975 | 2.55132016 |
| TMEM202-A: | lncRNA | 0.44802599 | 1.00449329 |
| DCTN1 | protein_coding | 0.44795813 | 12.8402167 |
| GPR137B | protein_coding | 0.44780939 | 2.47375052 |
| GIPR | protein_coding | 0.44767052 | 2.69953919 |
| MTCL1 | protein_coding | 0.44759151 | 6.67033123 |
| SLC3A2 | protein_coding | 0.44734187 | 11.2805737 |
| MZF1 | protein_coding | 0.44719033 | 4.08432226 |
| RCHY1 | protein_coding | 0.44696958 | 6.87520664 |
| RASAL2-AS1 | lncRNA | 0.44693483 | 0.70100878 |
| MPP5 | protein_coding | 0.44691552 | 6.7408154 |
| KIAA1109 | protein_coding | 0.44664239 | 10.2401351 |
| AARSD1 | protein_coding | 0.44642521 | 1.60405888 |

| | | | |
|---|---|---|---|
| SHANK3 | protein_coding | 0.44606402 | 7.17358109 |
| TEN1-CDK3 | protein_coding | 0.44555706 | 1.19558808 |
| FAM3C | protein_coding | 0.44554591 | 6.50922118 |
| ZRSR2 | protein_coding | 0.44551655 | 4.42908264 |
| NOS3 | protein_coding | 0.44549956 | 1.20645343 |
| DNAH17 | protein_coding | 0.44534907 | 5.21661871 |
| SCO2 | protein_coding | 0.44502342 | 4.01831741 |
| SENP6 | protein_coding | 0.44493117 | 10.5700225 |
| CLDN4 | protein_coding | 0.4448325 | 8.64708727 |
| CTD-2396E7. | lncRNA | 0.44461816 | 3.39141771 |
| HS1BP3 | protein_coding | 0.44439204 | 6.45972438 |
| APOA2 | protein_coding | 0.4443754 | 0.70625384 |
| SLC22A4 | protein_coding | 0.44431223 | 0.74840427 |
| LINC02253 | lncRNA | 0.44413384 | 2.99908882 |
| LINC02055 | lncRNA | 0.44360162 | 4.97735405 |
| NCAM2 | protein_coding | 0.44360011 | 1.82873795 |
| MTHFR | protein_coding | 0.44355433 | 5.42178765 |
| MAT2B | protein_coding | 0.44295135 | 6.65062841 |
| ATAD2B | protein_coding | 0.44268268 | 9.25685768 |
| CEP43 | protein_coding | 0.44246103 | 2.25640793 |
| SPRY2 | protein_coding | 0.44244563 | 0.72953595 |
| BTD | protein_coding | 0.44243531 | 2.18913168 |
| CHMP5 | protein_coding | 0.44237664 | 9.9739422 |
| NUP210 | protein_coding | 0.44226606 | 11.5369375 |
| MGP | protein_coding | 0.44224803 | 5.55895762 |
| LINC02609 | lncRNA | 0.44163142 | 1.39450449 |
| ANKFY1 | protein_coding | 0.44154687 | 9.80071074 |
| GLUL | protein_coding | 0.44142618 | 12.4622238 |
| DNAH2 | protein_coding | 0.441398 | 8.78151103 |
| SLC6A16 | protein_coding | 0.44104651 | 3.21787031 |
| GORAB | protein_coding | 0.44077548 | 3.01285092 |
| CCDC150 | protein_coding | 0.44058518 | 3.46195298 |
| INPP5E | protein_coding | 0.44044349 | 4.93388907 |
| ADAM23 | protein_coding | 0.44014606 | 9.68849716 |
| PCDHGA9 | protein_coding | 0.44005961 | 1.753374 |
| PAX8-AS1 | lncRNA | 0.43995876 | 8.69659106 |
| KIAA0319 | protein_coding | 0.43981859 | 4.97222666 |
| B3GNT2 | protein_coding | 0.43977472 | 8.09270702 |
| TBK1 | protein_coding | 0.43969078 | 7.63882649 |
| XYLT2 | protein_coding | 0.43947937 | 6.86469279 |
| CACNA1F | protein_coding | 0.43946877 | 2.22481914 |
| LRRC69 | protein_coding | 0.43941033 | 0.69405263 |
| CDC42SE2 | protein_coding | 0.43847514 | 9.94703937 |
| XRCC2 | protein_coding | 0.438224 | 7.26526566 |
| RP3-434P1.8 | lncRNA | 0.43800228 | 1.6308729 |

| | | | |
|---|---|---|---|
| CAMK2D | protein_coding | 0.43770586 | 8.81139286 |
| AC010967.2 | lncRNA | 0.43760462 | 1.13886334 |
| TPM1-AS | lncRNA | 0.43733435 | 0.58097504 |
| RP11-461O7. | lncRNA | 0.43733042 | 3.22650579 |
| PLXND1 | protein_coding | 0.43699947 | 9.19421222 |
| NHLRC4 | protein_coding | 0.43673207 | 1.19941016 |
| VAV2 | protein_coding | 0.43671445 | 10.0564457 |
| LCN12 | protein_coding | 0.43662904 | 0.65061823 |
| PRKG2 | protein_coding | 0.43659115 | 7.72864013 |
| SLC38A4 | protein_coding | 0.43611621 | 0.7779196 |
| NREP | protein_coding | 0.43586574 | 11.2699325 |
| OTUD4 | protein_coding | 0.43578671 | 10.0097131 |
| DCDC1 | protein_coding | 0.43474202 | 1.06725086 |
| MIA3 | protein_coding | 0.43434534 | 9.30801544 |
| GAP43 | protein_coding | 0.43401728 | 0.68482486 |
| VRK2 | protein_coding | 0.43347425 | 7.60318299 |
| ZNF460 | protein_coding | 0.43338787 | 3.27541957 |
| C9orf116 | protein_coding | 0.4333529 | 2.69336368 |
| SFT2D1 | protein_coding | 0.43325174 | 3.04933314 |
| FCHSD2 | protein_coding | 0.43276917 | 11.6677043 |
| PIM3 | protein_coding | 0.43267119 | 7.33255317 |
| CPT1B | protein_coding | 0.43243118 | 1.74972945 |
| TBC1D32 | protein_coding | 0.43225164 | 2.36595992 |
| LINC01572 | lncRNA | 0.43145342 | 1.17646347 |
| KCNQ4 | protein_coding | 0.43120586 | 6.47765028 |
| IGF2BP3 | protein_coding | 0.43119539 | 9.07507068 |
| IRF8 | protein_coding | 0.4309753 | 4.23914028 |
| ARHGEF9 | protein_coding | 0.43078152 | 11.6951667 |
| CTNND1 | protein_coding | 0.4305921 | 4.6857496 |
| ZNF703 | protein_coding | 0.43022382 | 3.07059043 |
| DENND4C | protein_coding | 0.43000932 | 9.33665887 |
| NAGLU | protein_coding | 0.42991024 | 7.09830457 |
| SPICE1 | protein_coding | 0.42988293 | 0.49522852 |
| RP11-251G2: | lncRNA | 0.42964205 | 2.5070345 |
| PMFBP1 | protein_coding | 0.42958694 | 0.85616691 |
| NCR3LG1 | protein_coding | 0.42948743 | 5.90397108 |
| PSMA1 | protein_coding | 0.42895876 | 5.69572698 |
| COQ8B | protein_coding | 0.42855343 | 7.01876273 |
| ARHGAP31 | protein_coding | 0.42833599 | 5.37071494 |
| PCDHGA2 | protein_coding | 0.42818135 | 1.20190766 |
| PDE4B | protein_coding | 0.42792053 | 4.75107455 |
| RP11-385F5. | lncRNA | 0.42758013 | 0.52789484 |
| ZNF410 | protein_coding | 0.42742119 | 1.19374826 |
| CRY2 | protein_coding | 0.42741083 | 6.91890323 |
| SEMA6C | protein_coding | 0.42720338 | 7.9865464 |

| | | | |
|---|---|---|---|
| RAB33B | protein_coding | 0.42719965 | 5.29379354 |
| CADPS2 | protein_coding | 0.42719379 | 0.4943724 |
| SGTB | protein_coding | 0.42691474 | 5.19360784 |
| ZNF598 | protein_coding | 0.42683334 | 6.97712391 |
| SLC38A6 | protein_coding | 0.42680818 | 2.35939604 |
| ZSCAN12 | protein_coding | 0.42671406 | 5.54476179 |
| C12orf56 | protein_coding | 0.42662621 | 3.87495378 |
| RP5-1024G6. | lncRNA | 0.42628733 | 0.72931562 |
| RP11-390P24 | lncRNA | 0.42627836 | 1.53576606 |
| SNHG20 | lncRNA | 0.426073 | 4.64928776 |
| SAT2 | protein_coding | 0.42606963 | 2.78908359 |
| MIR3659HG | lncRNA | 0.42588077 | 1.4292524 |
| RP11-467L13 | lncRNA | 0.42558115 | 0.99522964 |
| TNS1 | protein_coding | 0.42530977 | 7.11804339 |
| DUSP14 | protein_coding | 0.42512569 | 6.41476461 |
| SYT11 | protein_coding | 0.42512046 | 7.2949366 |
| COLGALT1 | protein_coding | 0.42480637 | 11.1157973 |
| SEMA4D | protein_coding | 0.42457299 | 8.46390087 |
| SYT7 | protein_coding | 0.4244977 | 11.2133792 |
| CSRNP2 | protein_coding | 0.42440421 | 8.22650088 |
| ZNF100 | protein_coding | 0.42404196 | 4.23765794 |
| RP11-134O2 | lncRNA | 0.42388919 | 2.88783327 |
| MED14 | protein_coding | 0.42347605 | 11.2244891 |
| ZFP36L2 | protein_coding | 0.42347048 | 5.08732665 |
| TMTC2 | protein_coding | 0.42302535 | 9.0880985 |
| ST3GAL5-AS | lncRNA | 0.42300448 | 1.17652772 |
| PIK3C2B | protein_coding | 0.4228551 | 7.89887813 |
| BTG1 | protein_coding | 0.42261427 | 6.25304672 |
| MYADM | protein_coding | 0.42240767 | 5.38715733 |
| CDC37L1 | protein_coding | 0.42238664 | 6.33274943 |
| ACOT11 | protein_coding | 0.4222171 | 2.95019673 |
| PER1 | protein_coding | 0.42218753 | 4.82129404 |
| AGPAT3 | protein_coding | 0.422158 | 9.83562623 |
| OSTM1 | protein_coding | 0.42204531 | 3.39473238 |
| SCN3A | protein_coding | 0.42197763 | 1.59148564 |
| ILK | protein_coding | 0.42185708 | 9.48738261 |
| RANBP10 | protein_coding | 0.42165292 | 7.56407226 |
| LRP1 | protein_coding | 0.42137518 | 6.04592895 |
| C10orf90 | protein_coding | 0.42127029 | 2.5247921 |
| DUSP12 | protein_coding | 0.42115427 | 6.59177588 |
| ZNF626 | protein_coding | 0.42113231 | 2.8975571 |
| GPRC5C | protein_coding | 0.42089024 | 0.99644358 |
| RP11-422P24 | lncRNA | 0.42076971 | 0.65100287 |
| CXCL16 | protein_coding | 0.42063431 | 0.93625803 |
| CYP27B1 | protein_coding | 0.42062531 | 2.51360735 |

| | | | |
|---|---|---|---|
| PDCD5 | protein_coding | 0.42059701 | 8.51751759 |
| SMG1 | protein_coding | 0.41991883 | 12.1988886 |
| UEVLD | protein_coding | 0.4197215 | 3.19849154 |
| CFAP58-DT | lncRNA | 0.4196409 | 0.72617773 |
| RP11-350N1! | lncRNA | 0.4194019 | 0.75251256 |
| SLC2A12 | protein_coding | 0.41933365 | 0.64068172 |
| TNKS1BP1 | protein_coding | 0.41895881 | 10.0654719 |
| HSPA9 | protein_coding | 0.41872806 | 12.7393887 |
| ZNF419 | protein_coding | 0.41869997 | 5.84751189 |
| MARK1 | protein_coding | 0.41817806 | 4.73706529 |
| KMT2C | protein_coding | 0.41811661 | 11.3424807 |
| MAS1 | protein_coding | 0.41802207 | 2.4508024 |
| ZNF277 | protein_coding | 0.41777041 | 7.40993058 |
| JADE3 | protein_coding | 0.41754626 | 7.93898216 |
| CTTNBP2NL | protein_coding | 0.41744657 | 6.34310906 |
| RCOR3 | protein_coding | 0.4172744 | 8.42710172 |
| CCDC191 | protein_coding | 0.41721362 | 0.64511888 |
| KREMEN1 | protein_coding | 0.41700477 | 6.46107156 |
| PSMB4 | protein_coding | 0.41647726 | 11.3757868 |
| ZNF484 | protein_coding | 0.41597777 | 5.03477911 |
| RP11-385F5. | lncRNA | 0.41592412 | 0.66311937 |
| TSPAN18 | protein_coding | 0.41588868 | 3.49475657 |
| NPIPB3 | protein_coding | 0.41584628 | 1.04691723 |
| FNIP1 | protein_coding | 0.41571473 | 6.85958583 |
| ZNF286A | protein_coding | 0.41565602 | 4.5398358 |
| FMN1 | protein_coding | 0.41513792 | 7.73907712 |
| HDHD3 | protein_coding | 0.41509102 | 5.52233629 |
| ATP6V1G2 | protein_coding | 0.41480739 | 6.11684336 |
| DCLK2 | protein_coding | 0.41475115 | 1.78235296 |
| RP11-468E2. | lncRNA | 0.41440676 | 1.58674142 |
| SOCS7 | protein_coding | 0.41439349 | 10.5571453 |
| MSH2-OT1 | lncRNA | 0.41419372 | 1.46841633 |
| WEE2-AS1 | lncRNA | 0.41413562 | 1.96075676 |
| DLD | protein_coding | 0.41413365 | 10.1638867 |
| KCTD15 | protein_coding | 0.413851 | 5.20128042 |
| DPY19L2 | protein_coding | 0.41376992 | 0.656244 |
| LRRC75A | protein_coding | 0.41361766 | 4.36126748 |
| NUDT17 | protein_coding | 0.41358411 | 1.42563296 |
| LINC01003 | lncRNA | 0.41340778 | 3.42069324 |
| TSFM | protein_coding | 0.41323768 | 0.681651 |
| SAXO2 | protein_coding | 0.41320232 | 0.6895964 |
| RIDA | protein_coding | 0.41319138 | 4.31151796 |
| MAIP1 | protein_coding | 0.41301131 | 4.91360002 |
| CFAP45 | protein_coding | 0.41286493 | 0.57958042 |
| LAMA3 | protein_coding | 0.4128309 | 8.41787296 |

| | | | |
|---|---|---|---|
| AHCYL2 | protein_coding | 0.41256721 | 5.15110894 |
| NOP2 | protein_coding | 0.41240858 | 1.17764255 |
| USP15 | protein_coding | 0.41232159 | 8.02804424 |
| ASB6 | protein_coding | 0.41227429 | 5.87884689 |
| LINC01297 | lncRNA | 0.41165132 | 0.63711335 |
| FBXL14 | protein_coding | 0.41079559 | 4.23209138 |
| NFE2L2 | protein_coding | 0.41056915 | 9.37327505 |
| XRN1 | protein_coding | 0.41041664 | 7.38821735 |
| CSGALNACT2 | lncRNA | 0.41024306 | 0.69852815 |
| MARS1 | protein_coding | 0.40980082 | 11.9406688 |
| CTSD | protein_coding | 0.40974263 | 4.91672538 |
| ENGASE | protein_coding | 0.40973986 | 7.6151317 |
| LINC01232 | lncRNA | 0.40954233 | 0.82754722 |
| CTD-2639E6. | lncRNA | 0.40927947 | 0.55831111 |
| RPS6KA6 | protein_coding | 0.40909457 | 6.08014003 |
| NDE1 | protein_coding | 0.40864976 | 9.19508436 |
| PCMTD1 | protein_coding | 0.40860793 | 3.90433667 |
| HERPUD1 | protein_coding | 0.40819883 | 10.7572168 |
| RTCA | protein_coding | 0.40769342 | 7.01863652 |
| IRF2BP2 | protein_coding | 0.40761996 | 9.85741887 |
| CRABP2 | protein_coding | 0.40758379 | 1.10285154 |
| ZNF641 | protein_coding | 0.40735514 | 4.89772884 |
| RALY-AS1 | lncRNA | 0.40724542 | 2.59375949 |
| RP11-267M2 | lncRNA | 0.40712972 | 0.86838541 |
| FCSK | protein_coding | 0.40709034 | 5.50628336 |
| RP1-228H13. | lncRNA | 0.40702395 | 3.7897184 |
| RP5-1047A19 | lncRNA | 0.40695388 | 1.10223158 |
| GSG1 | protein_coding | 0.40669839 | 0.44682539 |
| SMG7 | protein_coding | 0.40603774 | 11.8210692 |
| RP11-96K19. | lncRNA | 0.40599185 | 0.99153462 |
| ZNF544 | protein_coding | 0.40589746 | 8.64170366 |
| EIF2S2 | protein_coding | 0.40583453 | 11.3596774 |
| ADCY10 | protein_coding | 0.40581736 | 0.65964831 |
| GRAMD1B | protein_coding | 0.40571388 | 5.54477306 |
| DNASE2 | protein_coding | 0.40529904 | 7.77897174 |
| RP11-728G15 | lncRNA | 0.40523407 | 2.05379583 |
| SPSB2 | protein_coding | 0.4052275 | 2.40974425 |
| RP4-635E18. | lncRNA | 0.40516259 | 0.53499842 |
| UNC80 | protein_coding | 0.40509689 | 8.39303751 |
| CPVL | protein_coding | 0.40509582 | 6.96048336 |
| PCNX1 | protein_coding | 0.40496896 | 9.97460959 |
| ATF6 | protein_coding | 0.4048854 | 9.15490045 |
| ZNF609 | protein_coding | 0.40484352 | 8.67406734 |
| USH1G | protein_coding | 0.4048024 | 3.61382 |
| TULP2 | protein_coding | 0.40461493 | 1.39032005 |

| | | | |
|---|---|---|---|
| GARS1-DT | lncRNA | 0.40453619 | 5.89848645 |
| RCBTB2 | protein_coding | 0.40441091 | 2.37503309 |
| SH2D3A | protein_coding | 0.40436883 | 3.3806631 |
| ABRACL | protein_coding | 0.40417952 | 2.9321987 |
| RP11-83M8.: | lncRNA | 0.40402385 | 0.7916802 |
| RP11-539G1 | lncRNA | 0.40393206 | 1.03148867 |
| MIR200CHG | lncRNA | 0.40372338 | 1.60237562 |
| FKBP9 | protein_coding | 0.40367288 | 6.57774285 |
| KYAT3 | protein_coding | 0.40329452 | 7.04797204 |
| USP40 | protein_coding | 0.40326853 | 6.66568088 |
| POU3F1 | protein_coding | 0.40309684 | 5.23659273 |
| SLC37A2 | protein_coding | 0.40290842 | 0.56496413 |
| PLXNB2 | protein_coding | 0.40275106 | 8.98820439 |
| CELF2 | protein_coding | 0.40266809 | 5.57975281 |
| LINC01473 | lncRNA | 0.40260533 | 0.46027547 |
| VSTM2L | protein_coding | 0.40253767 | 2.73894197 |
| ALK | protein_coding | 0.40245384 | 8.56313083 |
| TRAM2-AS1 | lncRNA | 0.40239958 | 2.72348361 |
| ANKRD36 | protein_coding | 0.40216005 | 4.04103301 |
| CCDC18-AS1 | lncRNA | 0.4019692 | 4.29951859 |
| PRLR | protein_coding | 0.40193821 | 3.75281428 |
| LCN8 | protein_coding | 0.40155237 | 1.20146922 |
| TDRKH-AS1 | lncRNA | 0.40140292 | 1.24197593 |
| BCL2L13 | protein_coding | 0.4012521 | 10.0922001 |
| UTP15 | protein_coding | 0.4011276 | 6.8338101 |
| SLIT1 | protein_coding | 0.40087483 | 7.60159134 |
| MTO1 | protein_coding | 0.40032622 | 7.5363094 |
| TAF1D | protein_coding | 0.40003903 | 7.98404355 |
| CYREN | protein_coding | 0.39978569 | 7.86123472 |
| ADAM22 | protein_coding | 0.39973655 | 7.66277623 |
| PDE9A | protein_coding | 0.39965409 | 6.24112276 |
| LIG4 | protein_coding | 0.39944643 | 5.23765253 |
| GRM3 | protein_coding | 0.39942215 | 0.38955987 |
| GLIPR2 | protein_coding | 0.39941079 | 1.22064862 |
| OTOGL | protein_coding | 0.39902449 | 4.30368341 |
| GTF2IRD2 | protein_coding | 0.39890728 | 0.89536942 |
| KLC3 | protein_coding | 0.39881886 | 2.43187795 |
| PSD4 | protein_coding | 0.3987753 | 1.31232632 |
| RP5-1111F2 | lncRNA | 0.3987651 | 0.57823069 |
| TSC2 | protein_coding | 0.39873503 | 9.39997739 |
| PSMD14 | protein_coding | 0.39871945 | 10.0404525 |
| ARNT2 | protein_coding | 0.39841584 | 5.97923591 |
| PRRG4 | protein_coding | 0.39840653 | 1.49488386 |
| EFR3A | protein_coding | 0.39786226 | 7.28223053 |
| CMTR2 | protein_coding | 0.39777784 | 5.3574921 |

| | | | |
|---|---|---|---|
| PTPRN2 | protein_coding | 0.39769002 | 4.5300946 |
| TASOR2 | protein_coding | 0.39758855 | 9.41806271 |
| KCNC2 | protein_coding | 0.39740667 | 2.5138016 |
| SEL1L | protein_coding | 0.39723966 | 8.94808961 |
| PPARD | protein_coding | 0.39714043 | 5.24605755 |
| PCDH9 | protein_coding | 0.39710788 | 10.9379595 |
| BTG2 | protein_coding | 0.39703278 | 11.4799795 |
| RBM14-RBM | protein_coding | 0.39696232 | 0.65896951 |
| OGT | protein_coding | 0.3968826 | 10.2484052 |
| ISCU | protein_coding | 0.39664238 | 9.33859795 |
| H4C14 | protein_coding | 0.3964999 | 0.40796141 |
| OTULIN | protein_coding | 0.39631587 | 7.86737581 |
| PEF1 | protein_coding | 0.39605234 | 9.48913134 |
| AP4E1 | protein_coding | 0.39595655 | 7.85627622 |
| NPPC | protein_coding | 0.39571204 | 2.25582385 |
| CIP2A | protein_coding | 0.39560104 | 6.8815311 |
| CSRP2 | protein_coding | 0.39551732 | 11.2088875 |
| MAP1LC3A | protein_coding | 0.3950072 | 3.11144245 |
| LGALS8 | protein_coding | 0.39446924 | 5.0246525 |
| CHGB | protein_coding | 0.39446261 | 8.75194395 |
| OGFRP1 | lncRNA | 0.39438269 | 1.43326905 |
| ACAP2-IT1 | lncRNA | 0.39425727 | 0.68385986 |
| CEACAM19 | protein_coding | 0.39415848 | 0.78423203 |
| PAX5 | protein_coding | 0.39366419 | 11.0205068 |
| RRAGC | protein_coding | 0.39365284 | 5.50101798 |
| RTN4IP1 | protein_coding | 0.39352576 | 3.4998417 |
| TARS1 | protein_coding | 0.39266214 | 12.162516 |
| RP1-223B1.1 | lncRNA | 0.39252335 | 0.94741972 |
| GPATCH2L | protein_coding | 0.39244676 | 9.50214519 |
| ANKHD1-EIF4 | protein_coding | 0.39189391 | 0.43072542 |
| RP11-472H7. | lncRNA | 0.39169939 | 0.62073997 |
| TNIP2 | protein_coding | 0.3915633 | 6.14249159 |
| FZD6 | protein_coding | 0.39153202 | 7.57855391 |
| PRC1-AS1 | lncRNA | 0.39101485 | 0.9603545 |
| UBAP1L | protein_coding | 0.39087304 | 2.21819516 |
| MAMDC4 | protein_coding | 0.39062867 | 3.57431624 |
| SYCP2L | protein_coding | 0.39052577 | 5.31442771 |
| PRR22 | protein_coding | 0.39026099 | 1.84425359 |
| TRIM45 | protein_coding | 0.39001247 | 8.63799272 |
| VPS13C | protein_coding | 0.38992232 | 10.824489 |
| GPD1 | protein_coding | 0.38987956 | 3.17654702 |
| MRPL39 | protein_coding | 0.38987692 | 6.04280925 |
| DNAJC10 | protein_coding | 0.38977947 | 9.71197352 |
| CTC-301O7.4 | lncRNA | 0.38944819 | 1.38710041 |
| SNX14 | protein_coding | 0.38936785 | 1.25461256 |

| | | | |
|---|---|---|---|
| LINC01515 | lncRNA | 0.38921016 | 2.70240208 |
| ZCCHC7 | protein_coding | 0.38914718 | 7.56987611 |
| NCOA7 | protein_coding | 0.38895059 | 7.02025249 |
| RGMA | protein_coding | 0.38883946 | 6.82721578 |
| DPH2 | protein_coding | 0.38869363 | 9.66939747 |
| NFKBIL1 | protein_coding | 0.38833488 | 4.8724726 |
| KIAA1958 | protein_coding | 0.38823006 | 6.16415364 |
| SLC39A8 | protein_coding | 0.38811007 | 5.73283571 |
| CD55 | protein_coding | 0.38805318 | 5.98385855 |
| SLC26A11 | protein_coding | 0.38803154 | 2.74608474 |
| PCYT1A | protein_coding | 0.38796102 | 9.10502817 |
| DNAJA1 | protein_coding | 0.3877153 | 11.6972202 |
| IQCA1 | protein_coding | 0.38768524 | 0.71491301 |
| ALKBH8 | protein_coding | 0.38756249 | 2.83785534 |
| CELSR1 | protein_coding | 0.38741376 | 1.14172205 |
| SRRD | protein_coding | 0.38736612 | 6.37097649 |
| THBS3 | protein_coding | 0.38704701 | 7.19621857 |
| RRAGD | protein_coding | 0.38693022 | 10.3196497 |
| ZNF621 | protein_coding | 0.38677813 | 5.87946932 |
| LINC00863 | lncRNA | 0.38667084 | 2.14179271 |
| DYNC1LI2 | protein_coding | 0.38626981 | 10.7140977 |
| FBLL1 | protein_coding | 0.38592032 | 3.83942986 |
| KIAA1217 | protein_coding | 0.38540321 | 8.85366874 |
| SLC46A1 | protein_coding | 0.38501938 | 6.7337805 |
| CARMIL3 | protein_coding | 0.38472946 | 5.222539 |
| SNHG17 | lncRNA | 0.38437695 | 6.11805728 |
| TRNP1 | protein_coding | 0.38431833 | 1.24989399 |
| LRRC8D | protein_coding | 0.38429326 | 7.20215168 |
| USP2-AS1 | lncRNA | 0.38409661 | 1.41950666 |
| NTRK3 | protein_coding | 0.38400512 | 8.28011912 |
| RAD23B | protein_coding | 0.38383312 | 11.4043352 |
| SUSD4 | protein_coding | 0.38350952 | 0.78162742 |
| RP11-181L23 | lncRNA | 0.3829386 | 0.61292992 |
| RP3-508I15.2 | lncRNA | 0.38283668 | 1.31230243 |
| SLC25A13 | protein_coding | 0.38275387 | 3.68790661 |
| IGSF9B | protein_coding | 0.38245428 | 9.60738849 |
| MVK | protein_coding | 0.38243727 | 7.56825861 |
| ZDHHC11 | protein_coding | 0.38211292 | 3.68928836 |
| RAB18 | protein_coding | 0.38202243 | 6.33624943 |
| ARHGEF3 | protein_coding | 0.38181244 | 4.57317941 |
| DLX3 | protein_coding | 0.38178243 | 9.15045771 |
| TNFAIP8L1 | protein_coding | 0.38128919 | 6.6874776 |
| ACOT12 | protein_coding | 0.38076325 | 0.74930273 |
| ARHGAP11A | protein_coding | 0.38056799 | 10.8756289 |
| PPP1R26 | protein_coding | 0.38056582 | 6.05657355 |

| | | | |
|---|---|---|---|
| CLN5 | protein_coding | 0.38054225 | 5.56371949 |
| ARSA | protein_coding | 0.38037703 | 4.86602391 |
| CTA-228A9.3 | lncRNA | 0.38024731 | 2.0437105 |
| RP11-218F1( | lncRNA | 0.37970634 | 0.62670843 |
| ZNF440 | protein_coding | 0.37963882 | 3.83552016 |
| CMTM4 | protein_coding | 0.37952468 | 9.25691454 |
| ZKSCAN3 | protein_coding | 0.37912549 | 2.86382662 |
| RBMXL1 | protein_coding | 0.37887655 | 7.07333629 |
| MOB3B | protein_coding | 0.3784672 | 3.910118 |
| ALAS1 | protein_coding | 0.37794185 | 6.06704271 |
| MIB1 | protein_coding | 0.37784717 | 9.53791206 |
| DENND5A | protein_coding | 0.3776716 | 9.93975048 |
| CIZ1 | protein_coding | 0.37722376 | 10.1741765 |
| TOGARAM1 | protein_coding | 0.37716491 | 5.45239553 |
| ZNF630 | protein_coding | 0.37710566 | 2.42566177 |
| LINC00649 | lncRNA | 0.37692856 | 3.57907821 |
| INE1 | lncRNA | 0.37692163 | 1.18858461 |
| RPL36A | protein_coding | 0.37687564 | 2.86701335 |
| LTB4R2 | protein_coding | 0.37687428 | 2.1046516 |
| CNOT4 | protein_coding | 0.37625079 | 6.29754104 |
| ZFHX4 | protein_coding | 0.37599822 | 6.61239793 |
| ASF1A | protein_coding | 0.37585896 | 5.50796592 |
| DNAJC2 | protein_coding | 0.3758198 | 7.39607971 |
| RP11-448A1! | lncRNA | 0.37556999 | 3.8114787 |
| UBIAD1 | protein_coding | 0.37550046 | 5.45093111 |
| MAPK8IP2 | protein_coding | 0.37488794 | 4.92318912 |
| CITF22-49E9. | lncRNA | 0.37446655 | 0.52300473 |
| RP5-1157M2 | lncRNA | 0.37416491 | 1.26499576 |
| INO80D | protein_coding | 0.37410623 | 6.05620254 |
| CH17-174L2( | lncRNA | 0.37390724 | 0.92782815 |
| BBS1 | protein_coding | 0.37389547 | 0.89169292 |
| RP5-1085F1; | lncRNA | 0.37389239 | 0.97282582 |
| UCHL3 | protein_coding | 0.37359478 | 2.4991461 |
| PARP4 | protein_coding | 0.37343068 | 9.9515724 |
| MCCC1 | protein_coding | 0.37318553 | 7.35249014 |
| RAB29 | protein_coding | 0.37291193 | 6.4610799 |
| GPX4 | protein_coding | 0.37267824 | 8.93020561 |
| BAZ2A | protein_coding | 0.37235552 | 10.8016436 |
| LINC01355 | lncRNA | 0.37230314 | 1.68923375 |
| CES2 | protein_coding | 0.37217918 | 6.12593794 |
| KDM5D | protein_coding | 0.37214101 | 7.00753167 |
| CEND1 | protein_coding | 0.37206539 | 0.99771529 |
| EPHA7 | protein_coding | 0.37178041 | 3.41815091 |
| ULBP1 | protein_coding | 0.37176079 | 1.95094303 |
| RFXAP | protein_coding | 0.37174584 | 3.74054436 |

| | | | |
|---|---|---|---|
| NEK3 | protein_coding | 0.37174575 | 1.39238852 |
| SNX29 | protein_coding | 0.37108822 | 7.74160428 |
| LAD1 | protein_coding | 0.37097722 | 0.75957151 |
| MFSD11 | protein_coding | 0.37051254 | 5.94711158 |
| UBE2J1 | protein_coding | 0.37027787 | 7.24686564 |
| SATB2 | protein_coding | 0.36945193 | 8.40607087 |
| FANCF | protein_coding | 0.36924892 | 6.30426207 |
| HGSNAT | protein_coding | 0.36862472 | 6.36180287 |
| METTL22 | protein_coding | 0.36856733 | 4.64998646 |
| KCNA6 | protein_coding | 0.368192 | 0.85661576 |
| PCDHA9 | protein_coding | 0.36818516 | 0.57555843 |
| NSUN6 | protein_coding | 0.36785524 | 3.33828821 |
| GGT6 | protein_coding | 0.36740374 | 4.10952943 |
| MAN2C1 | protein_coding | 0.36729068 | 7.49521859 |
| IDE | protein_coding | 0.3671892 | 6.22768263 |
| C17orf113 | protein_coding | 0.36679334 | 2.36779968 |
| ZNF823 | protein_coding | 0.36678271 | 4.36805747 |
| PSMD7 | protein_coding | 0.36670008 | 10.2838301 |
| CTD-2292N2. | lncRNA | 0.36656324 | 1.04437839 |
| ACBD4 | protein_coding | 0.36650232 | 2.76624797 |
| RP11-14C10. | lncRNA | 0.36643627 | 0.78456263 |
| AKAP11 | protein_coding | 0.36640016 | 7.70100277 |
| C8orf37 | protein_coding | 0.36626515 | 2.45148527 |
| NADK | protein_coding | 0.36621936 | 6.60529286 |
| HCG27 | lncRNA | 0.36600871 | 1.23692508 |
| MYL5 | protein_coding | 0.36583891 | 2.18601935 |
| SLC22A23 | protein_coding | 0.36564724 | 4.5690339 |
| EPB41L4A-A! | lncRNA | 0.36543489 | 5.36555404 |
| ZDHHC11B | protein_coding | 0.3652062 | 2.59526995 |
| AARS1 | protein_coding | 0.36511112 | 12.052178 |
| ADGRB1 | protein_coding | 0.36487908 | 1.10546787 |
| SAP30 | protein_coding | 0.36486574 | 6.3666611 |
| ZC3HAV1 | protein_coding | 0.36450407 | 8.74363333 |
| SLK | protein_coding | 0.36416935 | 5.37434389 |
| SLC5A8 | protein_coding | 0.36410641 | 2.0663345 |
| NFASC | protein_coding | 0.36406672 | 10.1568433 |
| L1CAM | protein_coding | 0.36383981 | 7.00240361 |
| HSD17B1 | protein_coding | 0.36378272 | 1.6195872 |
| SLC35F1 | protein_coding | 0.36337231 | 1.07800975 |
| SATB2-AS1 | lncRNA | 0.36330746 | 4.55160211 |
| CACNA1C | protein_coding | 0.36319333 | 5.75828484 |
| MAP7 | protein_coding | 0.36306655 | 2.91316102 |
| MFHAS1 | protein_coding | 0.36305584 | 7.93968867 |
| HEXD | protein_coding | 0.36283523 | 4.8967271 |
| KCNMB3 | protein_coding | 0.36281777 | 1.99206658 |

| | | | |
|---|---|---|---|
| GANAB | protein_coding | 0.36243565 | 11.9277889 |
| SLMAP | protein_coding | 0.36240443 | 6.62792594 |
| ZNF280C | protein_coding | 0.3621578 | 3.00211787 |
| DENND4B | protein_coding | 0.36213229 | 8.76319546 |
| RGPD8 | protein_coding | 0.36178213 | 1.51458544 |
| RP4-773N10. | lncRNA | 0.36176836 | 4.57846551 |
| FLACC1 | protein_coding | 0.36174613 | 0.74883451 |
| ZNF436 | protein_coding | 0.36169399 | 7.72459489 |
| CCNG2 | protein_coding | 0.36165717 | 8.96870362 |
| TIMM9 | protein_coding | 0.36106293 | 6.25875388 |
| KCNH4 | protein_coding | 0.36095955 | 4.90641067 |
| SOCS2 | protein_coding | 0.36088699 | 4.25924602 |
| SNHG16 | lncRNA | 0.36050187 | 8.02886596 |
| MAPT | protein_coding | 0.36036677 | 10.4248658 |
| C4A | protein_coding | 0.36028542 | 0.75605624 |
| APLP1 | protein_coding | 0.36023419 | 9.26740704 |
| FBXW8 | protein_coding | 0.36014186 | 5.62605844 |
| PHC3 | protein_coding | 0.36011498 | 7.60663773 |
| ANGPTL2 | protein_coding | 0.35984308 | 0.49209597 |
| CDH11 | protein_coding | 0.35957522 | 5.6068301 |
| GPR1 | protein_coding | 0.35956313 | 5.26683788 |
| TREX2 | protein_coding | 0.35945299 | 0.77651473 |
| HTT | protein_coding | 0.35933293 | 10.1831836 |
| MAPK6-DT | lncRNA | 0.35925792 | 0.41822248 |
| XXbac-BPGBI | lncRNA | 0.35913667 | 1.14191653 |
| C12orf75 | protein_coding | 0.35899794 | 5.78789949 |
| TTLL3 | protein_coding | 0.35839963 | 1.71491035 |
| CTD-2528L19 | lncRNA | 0.35830281 | 0.75085388 |
| CCDC125 | protein_coding | 0.35822813 | 6.51295401 |
| NSRP1 | protein_coding | 0.35804252 | 5.96111922 |
| MMP25-AS1 | lncRNA | 0.35773064 | 1.28511272 |
| NUTM2E | protein_coding | 0.35771677 | 0.33905059 |
| FZD7 | protein_coding | 0.35701593 | 4.23252681 |
| RP5-1120P11 | lncRNA | 0.35692801 | 2.18554882 |
| RINT1 | protein_coding | 0.35622307 | 6.55879027 |
| RASAL2 | protein_coding | 0.35614524 | 5.43654403 |
| VAMP8 | protein_coding | 0.35607855 | 0.61458067 |
| AAK1 | protein_coding | 0.35596703 | 7.41237139 |
| GARS1 | protein_coding | 0.3558761 | 10.6018747 |
| BAG2 | protein_coding | 0.35568512 | 2.43077648 |
| APBB3 | protein_coding | 0.35529769 | 6.34926028 |
| RGL3 | protein_coding | 0.35504266 | 3.01238092 |
| MGST1 | protein_coding | 0.3549661 | 0.79036564 |
| ACAA1 | protein_coding | 0.35490879 | 4.75137292 |
| IL17RB | protein_coding | 0.35480037 | 8.01985116 |

| | | | |
|---|---|---|---|
| RBM26 | protein_coding | 0.35464047 | 7.56417358 |
| SKIDA1 | protein_coding | 0.35447651 | 2.46164558 |
| FANCA | protein_coding | 0.35441423 | 8.93996816 |
| PCID2 | protein_coding | 0.35425277 | 7.48688791 |
| QTRT1 | protein_coding | 0.35425111 | 5.38200378 |
| TSEN2 | protein_coding | 0.35419064 | 3.85027436 |
| RSKR | protein_coding | 0.35412892 | 4.36910626 |
| FAM228B | protein_coding | 0.35411005 | 2.92119683 |
| HMGN3-AS1 | lncRNA | 0.3538177 | 0.75104498 |
| NEK1 | protein_coding | 0.35379217 | 6.40889199 |
| ABCC2 | protein_coding | 0.35374379 | 0.47175766 |
| CHD6 | protein_coding | 0.35372039 | 9.19084375 |
| TMEM187 | protein_coding | 0.35364988 | 1.94876066 |
| PLD5P1 | protein_coding | 0.35351005 | 0.55809793 |
| HAX1 | protein_coding | 0.35338814 | 9.09987626 |
| IARS1 | protein_coding | 0.35336736 | 11.669081 |
| PKD1 | protein_coding | 0.35315606 | 6.18257739 |
| RP1-30M3.5 | lncRNA | 0.35310409 | 1.16835197 |
| RFPL3S | lncRNA | 0.35291344 | 0.9182086 |
| LINC00265 | lncRNA | 0.35286303 | 2.07169146 |
| WDFY3 | protein_coding | 0.35279659 | 7.57944654 |
| RFX7 | protein_coding | 0.35274092 | 8.29325106 |
| NSUN4 | protein_coding | 0.35267196 | 7.33647199 |
| ZNF140 | protein_coding | 0.35262104 | 6.94561094 |
| BECN2 | protein_coding | 0.35242446 | 1.27945517 |
| PSMB1 | protein_coding | 0.35199725 | 8.13592389 |
| PPP2R5B | protein_coding | 0.35198375 | 5.27175774 |
| AC159540.1 | lncRNA | 0.3516118 | 1.58144386 |
| DHDH | protein_coding | 0.35151706 | 0.58725045 |
| PRR34 | lncRNA | 0.35113595 | 0.59935667 |
| EXOSC8 | protein_coding | 0.35107333 | 6.06137978 |
| RP11-631N1( | lncRNA | 0.35105528 | 1.04557537 |
| DCTD | protein_coding | 0.35104222 | 8.43572782 |
| CTSC | protein_coding | 0.35093294 | 6.05978547 |
| KIF9 | protein_coding | 0.35089607 | 1.62482817 |
| SLCO3A1 | protein_coding | 0.35088468 | 8.95628439 |
| MYO18A | protein_coding | 0.35088195 | 7.6674312 |
| KB-1125A3.1 | lncRNA | 0.35052752 | 0.61718187 |
| GRHL1 | protein_coding | 0.35024371 | 3.37990836 |
| SRCIN1 | protein_coding | 0.3498737 | 4.36824232 |
| TTC4 | protein_coding | 0.34971328 | 0.98845347 |
| WI2-87327B | lncRNA | 0.34963144 | 1.80540203 |
| TMEM205 | protein_coding | 0.34956388 | 8.0998567 |
| EFEMP2 | protein_coding | 0.34919728 | 0.81190131 |
| RP11-1C8.7 | lncRNA | 0.34904982 | 0.65601025 |

| | | | |
|---|---|---|---|
| CTC1 | protein_coding | 0.34895868 | 5.7472816 |
| CTB-89H12.4 | lncRNA | 0.34861576 | 3.79919862 |
| RP11-416I2.3 | lncRNA | 0.3484884 | 0.53722986 |
| XPOT | protein_coding | 0.3482467 | 11.9426064 |
| C15orf39 | protein_coding | 0.34810069 | 7.08317431 |
| TMEM269 | protein_coding | 0.34808905 | 1.06403666 |
| CD27-AS1 | lncRNA | 0.34785854 | 5.25001445 |
| DNAAF2 | protein_coding | 0.34784315 | 5.40501853 |
| CAB39 | protein_coding | 0.34779704 | 8.50524926 |
| STS | protein_coding | 0.34769955 | 3.99489421 |
| GPR89A | protein_coding | 0.34754107 | 3.01823972 |
| LRRFIP2 | protein_coding | 0.34749401 | 5.41733491 |
| RP11-68I18.1 | lncRNA | 0.34740155 | 1.0855318 |
| MTHFD1L | protein_coding | 0.34738625 | 5.55367534 |
| RP11-250B2. | lncRNA | 0.3473131 | 0.66661181 |
| USP9X | protein_coding | 0.34721836 | 10.3586671 |
| RP11-95O2.5 | lncRNA | 0.34696655 | 0.66587057 |
| CCZ1 | protein_coding | 0.34682523 | 4.98427303 |
| RBM5 | protein_coding | 0.34673622 | 7.38987711 |
| MDN1 | protein_coding | 0.34644818 | 9.46210178 |
| THUMPD3-A: | lncRNA | 0.34641197 | 3.48897759 |
| EGLN3 | protein_coding | 0.3463402 | 4.35027541 |
| AACS | protein_coding | 0.34627761 | 7.41070708 |
| ARL6IP4 | protein_coding | 0.34622542 | 0.49297191 |
| TRAFD1 | protein_coding | 0.34618659 | 8.51181047 |
| HROB | protein_coding | 0.34606035 | 3.42100167 |
| RALA | protein_coding | 0.34556404 | 8.08198544 |
| TBC1D1 | protein_coding | 0.34549679 | 7.57329815 |
| NEK11 | protein_coding | 0.34508644 | 0.53110211 |
| C2orf27A | lncRNA | 0.34496188 | 0.66259046 |
| RP11-284F21 | lncRNA | 0.34496118 | 3.40960195 |
| ABCF2 | protein_coding | 0.34471847 | 4.69287274 |
| ASH2L | protein_coding | 0.34463141 | 7.52811251 |
| MIR193BHG | lncRNA | 0.34452019 | 1.08085171 |
| UBR4 | protein_coding | 0.34447478 | 11.1539126 |
| LYSMD2 | protein_coding | 0.34433517 | 6.03879127 |
| RNF11 | protein_coding | 0.34429386 | 6.92579901 |
| DUSP3 | protein_coding | 0.34424762 | 6.44322445 |
| CDK20 | protein_coding | 0.34423226 | 3.56521521 |
| PCDHA6 | protein_coding | 0.34423148 | 0.7150683 |
| ZFAND2A | protein_coding | 0.34421756 | 4.90469398 |
| RNF165 | protein_coding | 0.34396657 | 5.207063 |
| FBXO8 | protein_coding | 0.3438607 | 3.28491564 |
| ANAPC10 | protein_coding | 0.34370285 | 2.38708566 |
| RP11-774O3. | lncRNA | 0.34340114 | 0.4622925 |

| | | | |
|---|---|---|---|
| EML4 | protein_coding | 0.34338105 | 9.93763648 |
| RPF2 | protein_coding | 0.34328696 | 5.90151489 |
| AGAP6 | protein_coding | 0.34324091 | 0.77447733 |
| CDH26 | protein_coding | 0.34315702 | 1.54371738 |
| FKBP5 | protein_coding | 0.34291635 | 9.22397057 |
| ASXL3 | protein_coding | 0.34290391 | 5.20976247 |
| EXOC3L4 | protein_coding | 0.34285084 | 4.28634317 |
| PLD6 | protein_coding | 0.34280035 | 4.09054878 |
| PPP1R9B | protein_coding | 0.34279457 | 6.44251542 |
| FKTN | protein_coding | 0.34258925 | 6.89582143 |
| NUP93 | protein_coding | 0.34236655 | 9.49804084 |
| GSDMB | protein_coding | 0.34215215 | 1.76115047 |
| NEB | protein_coding | 0.34209567 | 0.93925799 |
| MSANTD3 | protein_coding | 0.3419303 | 5.85351861 |
| PYROXD1 | protein_coding | 0.34183132 | 3.76161869 |
| NLGN3 | protein_coding | 0.34141538 | 0.48651151 |
| DNAH1 | protein_coding | 0.34124387 | 1.60528981 |
| MEX3C | protein_coding | 0.34108478 | 7.57517906 |
| PROSER1 | protein_coding | 0.34091457 | 6.44606272 |
| SEC61A2 | protein_coding | 0.34079368 | 3.49361414 |
| UNC50 | protein_coding | 0.3405397 | 5.77700107 |
| MANBA | protein_coding | 0.34049088 | 4.76619565 |
| PUS7 | protein_coding | 0.34024421 | 5.79004432 |
| PIGB | protein_coding | 0.34021596 | 2.77743715 |
| RP2 | protein_coding | 0.33975641 | 4.1472935 |
| PJA1 | protein_coding | 0.33955151 | 6.13259956 |
| FAM27C | lncRNA | 0.33932437 | 0.94936825 |
| SLC6A6 | protein_coding | 0.33927043 | 7.20608442 |
| MIATNB | lncRNA | 0.33849852 | 1.25211702 |
| RP11-496H1. | lncRNA | 0.33829503 | 0.84340955 |
| GNL1 | protein_coding | 0.33807376 | 9.1786072 |
| HADHB | protein_coding | 0.3380669 | 6.05827731 |
| ACSF2 | protein_coding | 0.33773833 | 4.81804955 |
| U2AF1L4 | protein_coding | 0.33756974 | 2.32306627 |
| PHACTR1 | protein_coding | 0.33733969 | 2.50038404 |
| CBS | protein_coding | 0.33727534 | 4.81583953 |
| RP11-159D1. | lncRNA | 0.33718436 | 3.28686748 |
| ACOT13 | protein_coding | 0.33692966 | 6.15337378 |
| RP5-991G20. | lncRNA | 0.33648906 | 2.96475184 |
| ZNF136 | protein_coding | 0.33631346 | 5.63925698 |
| AP001063.1 | lncRNA | 0.33629748 | 0.84383012 |
| TXN | protein_coding | 0.33592208 | 9.79205477 |
| LRRC75B | protein_coding | 0.33591248 | 0.85332484 |
| PLGRKT | protein_coding | 0.33577372 | 5.14995283 |
| ITGA2 | protein_coding | 0.33572388 | 0.65434842 |

| | | | |
|---|---|---|---|
| ARL4D | protein_coding | 0.33538129 | 3.8337123 |
| GNMT | protein_coding | 0.33520045 | 0.75628862 |
| RWDD2B | protein_coding | 0.33511412 | 5.57335766 |
| PI4KB | protein_coding | 0.33498907 | 8.57102403 |
| XPO4 | protein_coding | 0.33498876 | 5.49484107 |
| SPATA2 | protein_coding | 0.33497779 | 4.9452599 |
| RP13-254B1( | lncRNA | 0.33487669 | 0.54570331 |
| PAQR6 | protein_coding | 0.33483638 | 1.4944578 |
| CGN | protein_coding | 0.3345758 | 6.16155559 |
| SPTBN5 | protein_coding | 0.33449041 | 2.91780104 |
| NXPE3 | protein_coding | 0.33446195 | 6.85840107 |
| EMC1 | protein_coding | 0.33435511 | 9.44223153 |
| ATM | protein_coding | 0.33413786 | 7.59278198 |
| FCHO1 | protein_coding | 0.3340784 | 7.14108942 |
| PDCD2L | protein_coding | 0.33389245 | 4.32092612 |
| HARBI1 | protein_coding | 0.33381029 | 3.17790312 |
| TJP1 | protein_coding | 0.33355493 | 7.77521929 |
| COG5 | protein_coding | 0.33347793 | 6.95503596 |
| GK | protein_coding | 0.33334521 | 5.36190858 |
| ABITRAM | protein_coding | 0.3332149 | 5.37404377 |
| ING3 | protein_coding | 0.33307504 | 5.16159702 |
| PAK4 | protein_coding | 0.33274121 | 6.09647509 |
| OLFM1 | protein_coding | 0.33222572 | 9.62742608 |
| TCF7L2 | protein_coding | 0.33216136 | 2.52504949 |
| EIF4E3 | protein_coding | 0.33213541 | 5.49272479 |
| GRIPAP1 | protein_coding | 0.33202962 | 7.67819024 |
| RP11-63L7.6 | lncRNA | 0.33197069 | 1.31759725 |
| GTF2H1 | protein_coding | 0.33185428 | 7.48483181 |
| EIF3CL | protein_coding | 0.33177228 | 0.8643236 |
| CCNJ | protein_coding | 0.33165901 | 7.52672743 |
| LRRC8C-DT | lncRNA | 0.3313398 | 2.6798284 |
| BCL2L2 | protein_coding | 0.33132809 | 6.52553728 |
| UFL1-AS1 | lncRNA | 0.3313066 | 0.61055783 |
| NECAP2 | protein_coding | 0.33123594 | 6.15295728 |
| COMMD8 | protein_coding | 0.3310842 | 2.34423892 |
| LOXHD1 | protein_coding | 0.33073944 | 2.8091326 |
| TLE3 | protein_coding | 0.33067997 | 5.35794065 |
| POGLUT2 | protein_coding | 0.33058357 | 4.65318753 |
| BBS4 | protein_coding | 0.33047899 | 5.84492869 |
| MMP2 | protein_coding | 0.33028997 | 0.57622025 |
| TBC1D23 | protein_coding | 0.33009492 | 4.75973414 |
| FAM126B | protein_coding | 0.33006911 | 5.79230282 |
| EML2 | protein_coding | 0.32994869 | 7.06888679 |
| SH3RF1 | protein_coding | 0.32973414 | 6.78601696 |
| WBP4 | protein_coding | 0.32970124 | 3.65464048 |

| | | | |
|---|---|---|---|
| E2F6 | protein_coding | 0.3295862 | 5.6646135 |
| AAMP | protein_coding | 0.32943511 | 9.92760752 |
| SMIM17 | protein_coding | 0.32915226 | 0.53907378 |
| DGCR11 | lncRNA | 0.32914393 | 0.67391436 |
| TRIO | protein_coding | 0.32901072 | 7.06525678 |
| CCDC157 | protein_coding | 0.32888232 | 1.12956776 |
| EGLN2 | protein_coding | 0.32885791 | 2.11971462 |
| RABEPK | protein_coding | 0.32877152 | 6.60474663 |
| ZNF675 | protein_coding | 0.32866638 | 3.95530247 |
| PRRT2 | protein_coding | 0.32862058 | 0.63620357 |
| THG1L | protein_coding | 0.32853993 | 4.43162951 |
| LRRC56 | protein_coding | 0.32843703 | 1.57495226 |
| CCDC24 | protein_coding | 0.32833715 | 2.72796338 |
| FCGRT | protein_coding | 0.32828278 | 7.19421439 |
| CSNK1A1 | protein_coding | 0.32826994 | 9.39882694 |
| ST3GAL4 | protein_coding | 0.32812376 | 7.18626508 |
| TTLL7 | protein_coding | 0.32777655 | 6.77997944 |
| KCNMB4 | protein_coding | 0.3273878 | 4.96212172 |
| SMPDL3A | protein_coding | 0.32729744 | 0.6364572 |
| PABPC1L | protein_coding | 0.32718145 | 6.088908 |
| C11orf45 | protein_coding | 0.32713987 | 1.96510621 |
| KCNJ4 | protein_coding | 0.32674307 | 4.29670211 |
| CDH2 | protein_coding | 0.32673502 | 9.99537379 |
| BCL7A | protein_coding | 0.32648877 | 8.36777261 |
| AL136419.6 | lncRNA | 0.32641339 | 0.39505882 |
| ACAP2 | protein_coding | 0.32630467 | 8.8098683 |
| STAT4 | protein_coding | 0.32626152 | 0.57956061 |
| MKLN1-AS | lncRNA | 0.32623993 | 1.65913099 |
| BRF2 | protein_coding | 0.3261437 | 4.26103641 |
| LPXN | protein_coding | 0.32611102 | 1.69679946 |
| GPRASP1 | protein_coding | 0.3260678 | 1.70812207 |
| POLQ | protein_coding | 0.32600497 | 5.24514963 |
| UBXN4 | protein_coding | 0.32596691 | 8.83571758 |
| RP11-421J10 | lncRNA | 0.32590359 | 0.46202791 |
| AC009495.2 | lncRNA | 0.32589719 | 0.33905634 |
| PLCG2 | protein_coding | 0.32553738 | 2.96598972 |
| ZNF283 | protein_coding | 0.32550935 | 2.93291419 |
| NPAS3 | protein_coding | 0.3252882 | 0.68226971 |
| SLC48A1 | protein_coding | 0.32512976 | 6.45745313 |
| SNHG12 | lncRNA | 0.32461996 | 3.40166561 |
| RASGEF1B | protein_coding | 0.32433622 | 7.72728569 |
| RIOX2 | protein_coding | 0.32420909 | 7.45223584 |
| ACSS1 | protein_coding | 0.32415962 | 5.6109215 |
| GTF2IRD2B | protein_coding | 0.32403812 | 1.87359076 |
| NBPF11 | protein_coding | 0.323865 | 2.65640697 |

| | | | |
|---|---|---|---|
| COBLL1 | protein_coding | 0.32365962 | 8.33618509 |
| DIRC3-AS1 | lncRNA | 0.32350762 | 3.78942685 |
| CTA-941F9.1 | lncRNA | 0.3234852 | 0.52038718 |
| ADM5 | protein_coding | 0.32335442 | 1.11960642 |
| FBXL6 | protein_coding | 0.32335102 | 5.23501458 |
| PIGBOS1 | protein_coding | 0.3229793 | 1.1078272 |
| GCFC2 | protein_coding | 0.32266975 | 6.3484904 |
| PURB | protein_coding | 0.32258903 | 6.82495609 |
| GRM4 | protein_coding | 0.32256069 | 5.95676428 |
| EDIL3 | protein_coding | 0.32253352 | 2.82702093 |
| PTCD2 | protein_coding | 0.322361 | 4.13813114 |
| BCAN-AS1 | lncRNA | 0.32217871 | 1.9632113 |
| EMP2 | protein_coding | 0.32211097 | 8.47910944 |
| DCP1A | protein_coding | 0.32180074 | 6.18380533 |
| SCGB1D2 | protein_coding | 0.3217145 | 0.60384745 |
| PHC1 | protein_coding | 0.32154989 | 3.57296319 |
| MAN2B2 | protein_coding | 0.32150315 | 5.33617058 |
| WASHC2A | protein_coding | 0.32142326 | 4.75408435 |
| PLEKHG3 | protein_coding | 0.321367 | 7.227924 |
| GABBR1 | protein_coding | 0.32128374 | 3.32422944 |
| USP53 | protein_coding | 0.32124768 | 4.74023552 |
| SHLD3 | protein_coding | 0.32116608 | 2.48264587 |
| SENP1 | protein_coding | 0.32103739 | 7.62761749 |
| SRD5A1 | protein_coding | 0.32093154 | 6.81651606 |
| TIRAP | protein_coding | 0.32089017 | 1.19364443 |
| PCOTH | lncRNA | 0.32077805 | 0.84797 |
| LARS1 | protein_coding | 0.32076273 | 10.8730514 |
| PSMD13 | protein_coding | 0.32067643 | 9.39248381 |
| PDF | protein_coding | 0.32065059 | 0.52540365 |
| TRAPPC10 | protein_coding | 0.32064435 | 7.50849945 |
| SCG3 | protein_coding | 0.32061869 | 10.180634 |
| ADGRF3 | protein_coding | 0.3203591 | 0.60082116 |
| CMTM6 | protein_coding | 0.32028864 | 4.97289157 |
| DEDD | protein_coding | 0.32023219 | 6.39097555 |
| ENOSF1 | protein_coding | 0.32004805 | 6.07745891 |
| GPS2 | protein_coding | 0.31978929 | 0.74418185 |
| EME2 | protein_coding | 0.3196871 | 6.32168277 |
| KIF13A | protein_coding | 0.31959782 | 2.31256826 |
| ME3 | protein_coding | 0.3195433 | 4.51858723 |
| SHROOM1 | protein_coding | 0.31949974 | 1.62608672 |
| SERP2 | protein_coding | 0.31948386 | 0.40316142 |
| FLRT2 | protein_coding | 0.31944134 | 6.31616053 |
| COMMD5 | protein_coding | 0.31888931 | 5.17188987 |
| CA13 | protein_coding | 0.31888644 | 0.64484179 |
| PLD2 | protein_coding | 0.31882302 | 1.32534337 |

| | | | |
|---|---|---|---|
| CC2D2A | protein_coding | 0.31859075 | 2.71928495 |
| GPATCH4 | protein_coding | 0.31825102 | 5.60143556 |
| LRIF1 | protein_coding | 0.31820198 | 6.85738979 |
| RP11-338N1( | lncRNA | 0.31806795 | 0.46966763 |
| JARID2 | protein_coding | 0.31806698 | 6.73190777 |
| MIR1915HG | lncRNA | 0.31801557 | 0.55750265 |
| RP4-681L3.3 | lncRNA | 0.31788805 | 1.39528344 |
| S1PR2 | protein_coding | 0.31780013 | 2.41704515 |
| EIF2S3 | protein_coding | 0.31762622 | 9.74471342 |
| POLR3C | protein_coding | 0.31752157 | 7.12455522 |
| RP11-399B1 | lncRNA | 0.31739666 | 2.09798549 |
| MYH9 | protein_coding | 0.31739535 | 9.83713005 |
| PNPLA6 | protein_coding | 0.31739225 | 4.96932365 |
| QTRT2 | protein_coding | 0.31737228 | 3.94500896 |
| THUMPD2 | protein_coding | 0.31683125 | 5.44002598 |
| TIMM44 | protein_coding | 0.3166677 | 7.74562193 |
| LATS1 | protein_coding | 0.31664524 | 6.77065615 |
| EFCAB2 | protein_coding | 0.31660722 | 2.91421983 |
| SUPV3L1 | protein_coding | 0.31660474 | 4.61120816 |
| ZNF234 | protein_coding | 0.31649941 | 4.64087002 |
| UBE2D4 | protein_coding | 0.31616884 | 4.85514902 |
| UBR3 | protein_coding | 0.31616771 | 7.95285216 |
| NFU1 | protein_coding | 0.31611464 | 5.20829756 |
| CMSS1 | protein_coding | 0.31609397 | 6.17047979 |
| H3C10 | protein_coding | 0.31605206 | 1.40753745 |
| PDXK | protein_coding | 0.31586801 | 6.59869333 |
| PSMD1 | protein_coding | 0.31583078 | 9.8892654 |
| SLC6A9 | protein_coding | 0.31566196 | 6.20988484 |
| RAB4B | protein_coding | 0.31548644 | 0.4887778 |
| POLB | protein_coding | 0.31529292 | 4.5377527 |
| POLR2J3 | protein_coding | 0.31522233 | 0.79670024 |
| ASS1 | protein_coding | 0.31518279 | 7.61947108 |
| TAPT1-AS1 | lncRNA | 0.31487956 | 1.83585235 |
| KIT | protein_coding | 0.31465706 | 7.53326 |
| SPRED1 | protein_coding | 0.314631 | 0.5563989 |
| ANKLE1 | protein_coding | 0.31461176 | 0.76285375 |
| SLC30A7 | protein_coding | 0.31448975 | 4.00343206 |
| RP11-159H1( | lncRNA | 0.31440907 | 0.6519303 |
| NORAD | lncRNA | 0.31428897 | 10.2412671 |
| ZNF233 | protein_coding | 0.31421973 | 2.42254062 |
| ESRRG | protein_coding | 0.31405914 | 4.65189533 |
| RP11-540O1 | lncRNA | 0.31396697 | 0.4958223 |
| KIF12 | protein_coding | 0.31382859 | 2.66922206 |
| EXT2 | protein_coding | 0.31375354 | 7.19730494 |
| SLC18B1 | protein_coding | 0.3135131 | 3.85154782 |

| | | | |
|---|---|---|---|
| LRRC41 | protein_coding | 0.31339221 | 6.21979866 |
| WASHC2C | protein_coding | 0.31316721 | 3.29207878 |
| AL022341.3 | lncRNA | 0.31310053 | 0.48117709 |
| FJX1 | protein_coding | 0.3129937 | 2.59339138 |
| AC005306.3 | lncRNA | 0.31292164 | 1.03949083 |
| KRT80 | protein_coding | 0.31234576 | 3.52656255 |
| SLIT3 | protein_coding | 0.31221002 | 1.44358638 |
| LIX1L | protein_coding | 0.31199814 | 5.55399231 |
| CSGALNACT1 | protein_coding | 0.31193088 | 5.57470033 |
| KHDC4 | protein_coding | 0.31160896 | 7.46706351 |
| TBC1D19 | protein_coding | 0.31149452 | 1.95335822 |
| CAMSAP2 | protein_coding | 0.3114375 | 6.22019559 |
| PRMT8 | protein_coding | 0.31122263 | 0.37150052 |
| FIBCD1 | protein_coding | 0.31108518 | 0.60859025 |
| USP38 | protein_coding | 0.31103324 | 7.25888281 |
| TSR1 | protein_coding | 0.31102667 | 7.20331856 |
| DSCAML1 | protein_coding | 0.31098616 | 4.95712002 |
| ZFAND1 | protein_coding | 0.31082561 | 6.36814357 |
| CHST12 | protein_coding | 0.3107036 | 5.07343314 |
| C1QTNF6 | protein_coding | 0.31054108 | 2.57924044 |
| ATF1 | protein_coding | 0.31014117 | 3.5305141 |
| SLC9A3 | protein_coding | 0.3100422 | 0.33722403 |
| ZNF783 | protein_coding | 0.31004142 | 4.73228713 |
| C21orf91 | protein_coding | 0.31001765 | 3.87854808 |
| OSBPL5 | protein_coding | 0.3100078 | 1.45867186 |
| ZNF780A | protein_coding | 0.30992562 | 4.70877693 |
| PSME4 | protein_coding | 0.30985421 | 8.01893167 |
| RIPK1 | protein_coding | 0.30978031 | 5.85927021 |
| ITGAV | protein_coding | 0.30972523 | 4.71393877 |
| TM9SF4 | protein_coding | 0.30968534 | 8.17939626 |
| MSANTD1 | protein_coding | 0.30958612 | 0.46866458 |
| PARD3B | protein_coding | 0.30946519 | 0.90760755 |
| ONECUT2 | protein_coding | 0.3093406 | 6.8172461 |
| PPEF1 | protein_coding | 0.3092216 | 1.76307769 |
| GTF2H3 | protein_coding | 0.30918499 | 6.05756848 |
| FAM174B | protein_coding | 0.30918249 | 6.77512094 |
| COL7A1 | protein_coding | 0.3091778 | 3.70753224 |
| PTPRK | protein_coding | 0.30912248 | 8.22825521 |
| DNMT3A | protein_coding | 0.3090869 | 7.78395492 |
| ZNF41 | protein_coding | 0.30895528 | 5.32380558 |
| VCP | protein_coding | 0.3089279 | 9.71334951 |
| MAX | protein_coding | 0.30887606 | 7.2130588 |
| STK38L | protein_coding | 0.30877554 | 5.15310974 |
| CAPRIN2 | protein_coding | 0.30874824 | 3.42577347 |
| SPIRE2 | protein_coding | 0.30869942 | 5.41358223 |

| | | | |
|---|---|---|---|
| TRMT10B | protein_coding | 0.30861056 | 2.9265358 |
| EIF4A1 | protein_coding | 0.30860106 | 1.31992338 |
| PRMT1 | protein_coding | 0.30857734 | 9.66234949 |
| COLGALT2 | protein_coding | 0.30851813 | 7.64623684 |
| FAM53A | protein_coding | 0.3084861 | 1.69506083 |
| RP11-98D18. | lncRNA | 0.30839868 | 0.55891131 |
| RFX3 | protein_coding | 0.3081263 | 5.82039157 |
| RPP40 | protein_coding | 0.30800951 | 3.87969348 |
| RNF135 | protein_coding | 0.30793801 | 0.9452829 |
| RP3-467N11. | lncRNA | 0.30784619 | 0.87560961 |
| TSLP | protein_coding | 0.30767361 | 0.58353707 |
| SEZ6L2 | protein_coding | 0.30753933 | 8.02260665 |
| EFR3B | protein_coding | 0.30751707 | 3.65902695 |
| CLUL1 | protein_coding | 0.30718298 | 0.76121835 |
| LINC00665 | lncRNA | 0.30710564 | 6.71323944 |
| HUWE1 | protein_coding | 0.30707709 | 11.007286 |
| ZZEF1 | protein_coding | 0.30676228 | 7.7436838 |
| MYCN | protein_coding | 0.30669689 | 0.3683276 |
| SOCS2-AS1 | lncRNA | 0.30661489 | 0.79884826 |
| RP11-248J18 | lncRNA | 0.30633765 | 0.3477511 |
| RNF185 | protein_coding | 0.30611182 | 5.61288627 |
| NDUFAF7 | protein_coding | 0.30610559 | 3.37133915 |
| PSMC6 | protein_coding | 0.30539789 | 7.49976895 |
| AC093843.1 | lncRNA | 0.30533048 | 0.38614631 |
| RRM2B | protein_coding | 0.3051201 | 4.92085719 |
| PRR15 | protein_coding | 0.30478889 | 0.48324615 |
| FBF1 | protein_coding | 0.30465889 | 2.36739638 |
| LINC02604 | lncRNA | 0.30453609 | 2.46199533 |
| TBC1D3L | protein_coding | 0.30447721 | 0.86964049 |
| CCDC18 | protein_coding | 0.30442423 | 3.72794849 |
| C16orf86 | protein_coding | 0.30435284 | 1.16901823 |
| MARCKS | protein_coding | 0.30402797 | 6.10742992 |
| EXOC6B | protein_coding | 0.30398808 | 4.10871676 |
| ANKRD36B | protein_coding | 0.30397745 | 2.53757833 |
| ELF1 | protein_coding | 0.3038997 | 3.09054815 |
| YY1AP1 | protein_coding | 0.30368099 | 6.24508746 |
| CCDC59 | protein_coding | 0.30358686 | 5.35619007 |
| ZNF543 | protein_coding | 0.30356129 | 3.83914889 |
| PEX3 | protein_coding | 0.30312346 | 1.51462782 |
| SLC35D3 | protein_coding | 0.30311106 | 3.81660208 |
| ERC2 | protein_coding | 0.30309318 | 1.27116018 |
| RASA2 | protein_coding | 0.30290371 | 4.63308025 |
| KIF3B | protein_coding | 0.30278827 | 7.32055051 |
| AC007292.7 | lncRNA | 0.30269826 | 0.45467296 |
| SLC39A14 | protein_coding | 0.30262013 | 6.04304897 |

| | | | |
|---|---|---|---|
| FECH | protein_coding | 0.3026017 | 5.92876543 |
| GCLC | protein_coding | 0.30233747 | 5.63625731 |
| HMCN1 | protein_coding | 0.30227062 | 1.22482125 |
| RIT1 | protein_coding | 0.30187708 | 3.20978898 |
| ZNF772 | protein_coding | 0.30181604 | 5.57440657 |
| CDK7 | protein_coding | 0.30166199 | 4.00843348 |
| AP000254.8 | lncRNA | 0.30154386 | 0.61225078 |
| PKP3 | protein_coding | 0.30139257 | 3.81084709 |
| NFIA | protein_coding | 0.30135319 | 7.09346806 |
| METTL3 | protein_coding | 0.30111653 | 6.41427733 |
| RGS5 | protein_coding | 0.30103396 | 5.73839995 |
| NOM1 | protein_coding | 0.30101415 | 6.84863243 |
| ADAT2 | protein_coding | 0.30089363 | 2.44986996 |
| RAP2C | protein_coding | 0.30082051 | 2.70065333 |
| ST3GAL2 | protein_coding | 0.30069194 | 4.21455061 |
| ZC3HC1 | protein_coding | 0.30059149 | 4.40330097 |
| CDC42BPB | protein_coding | 0.30059 | 8.69568695 |
| CTD-3234P18 | lncRNA | 0.30034305 | 0.35333092 |
| POU6F1 | protein_coding | 0.30021096 | 5.55651815 |
| IKBKB | protein_coding | 0.30016879 | 4.57477476 |
| LINC00535 | lncRNA | 0.30014818 | 1.00501402 |
| SLC25A25-AS | lncRNA | 0.30008639 | 3.68911818 |
| SSBP3 | protein_coding | 0.30004223 | 7.84758822 |
| CCDC74A | protein_coding | 0.30001557 | 0.42715588 |
| ZDHHC22 | protein_coding | 0.29983489 | 2.21910124 |
| METTL5 | protein_coding | 0.29972913 | 5.73580729 |
| CBX6 | protein_coding | 0.29961132 | 7.24098026 |
| PORCN | protein_coding | 0.29959153 | 1.55114126 |
| PCDHGB3 | protein_coding | 0.29931148 | 0.73752807 |
| CYB5R1 | protein_coding | 0.29928482 | 8.98566334 |
| RHBDL3 | protein_coding | 0.29926309 | 7.84588242 |
| NR5A2 | protein_coding | 0.29922772 | 2.01085029 |
| DUSP1 | protein_coding | 0.2991247 | 1.74769808 |
| CTC-471J1.11 | lncRNA | 0.29901162 | 0.46489828 |
| DCUN1D3 | protein_coding | 0.29880658 | 1.91111681 |
| CAP2 | protein_coding | 0.29870994 | 2.20196876 |
| DRAM2 | protein_coding | 0.29869404 | 4.09350958 |
| DCAF8 | protein_coding | 0.29867998 | 5.47360841 |
| SNHG8 | lncRNA | 0.29858688 | 5.70382424 |
| SIRT4 | protein_coding | 0.2984709 | 0.66216557 |
| BRCA2 | protein_coding | 0.29840999 | 4.51579224 |
| FA2H | protein_coding | 0.29836252 | 4.07839836 |
| LINC00663 | lncRNA | 0.29825151 | 0.90042744 |
| PAXIP1 | protein_coding | 0.29791712 | 6.92903145 |
| VEZT | protein_coding | 0.29790503 | 8.1526966 |

| | | | |
|---|---|---|---|
| SPATA6 | protein_coding | 0.29754102 | 2.37240887 |
| LRRIQ4 | protein_coding | 0.29750052 | 0.35442141 |
| NSUN7 | protein_coding | 0.29703784 | 1.76646635 |
| SPPL2C | protein_coding | 0.29700846 | 0.36872279 |
| CTD-2619J13 | lncRNA | 0.29672611 | 0.90001348 |
| LINC01132 | lncRNA | 0.29669254 | 0.42266609 |
| FBXO4 | protein_coding | 0.29651918 | 0.9605266 |
| ZNF418 | protein_coding | 0.29650012 | 4.38445415 |
| RAB10 | protein_coding | 0.29649137 | 7.36518203 |
| SERF1B | protein_coding | 0.29624119 | 0.42494371 |
| TSPAN12 | protein_coding | 0.29593303 | 3.50305887 |
| KCNH7 | protein_coding | 0.29591218 | 3.53179975 |
| VGLL4 | protein_coding | 0.29579331 | 6.02331955 |
| ZNF862 | protein_coding | 0.29566028 | 3.01174249 |
| ATP6V0A1 | protein_coding | 0.29563209 | 9.44536516 |
| LNPEP | protein_coding | 0.29537224 | 7.48262724 |
| B3GALNT2 | protein_coding | 0.29524891 | 6.12150485 |
| ADAMTS13 | protein_coding | 0.29516707 | 2.16306052 |
| COQ9 | protein_coding | 0.29505795 | 6.4555105 |
| JMY | protein_coding | 0.29484559 | 5.15436702 |
| RP11-76P17. | lncRNA | 0.29483085 | 0.72000335 |
| SMG5 | protein_coding | 0.29477546 | 9.18345451 |
| ATG2A | protein_coding | 0.29477388 | 5.6667298 |
| LINC00630 | lncRNA | 0.29411231 | 1.67391939 |
| LINC01122 | lncRNA | 0.2940529 | 0.7782915 |
| ABCC10 | protein_coding | 0.2933271 | 3.5420943 |
| CISD1 | protein_coding | 0.29330866 | 3.93386398 |
| DLG3 | protein_coding | 0.29306645 | 6.14889153 |
| PPP2R3A | protein_coding | 0.2929571 | 0.67900222 |
| SPART | protein_coding | 0.29290281 | 0.78435556 |
| ZNF184 | protein_coding | 0.29277306 | 5.58115009 |
| LIN37 | protein_coding | 0.29271405 | 0.93474539 |
| CTD-3220F14 | lncRNA | 0.29269853 | 0.92570413 |
| LRRC37A | protein_coding | 0.29260809 | 0.51959241 |
| GABRG3 | protein_coding | 0.29248077 | 1.17641429 |
| MAP1LC3B2 | protein_coding | 0.29246704 | 0.70417646 |
| REXO5 | protein_coding | 0.29234245 | 3.15457368 |
| ACADSB | protein_coding | 0.29223224 | 3.88795919 |
| TBX18 | protein_coding | 0.29218366 | 0.44945682 |
| VPS35L | protein_coding | 0.29213921 | 6.43140267 |
| EIF5A2 | protein_coding | 0.29212705 | 0.60822628 |
| TBPL1 | protein_coding | 0.29212238 | 3.41049839 |
| CTD-2622I13 | lncRNA | 0.29181655 | 0.36105397 |
| ZSCAN26 | protein_coding | 0.2916634 | 2.73440655 |
| INTS12 | protein_coding | 0.29165166 | 4.22253349 |

| | | | |
|---|---|---|---|
| UHRF1BP1 | protein_coding | 0.29134197 | 9.00208014 |
| ARHGAP11B | protein_coding | 0.29127111 | 1.06592872 |
| RUBCNL | protein_coding | 0.29126694 | 1.69783482 |
| PPP1CB | protein_coding | 0.29119939 | 4.41397815 |
| SCLT1 | protein_coding | 0.29096017 | 4.36287712 |
| ALPL | protein_coding | 0.29087566 | 0.99136487 |
| PLCD1 | protein_coding | 0.29081905 | 1.02659715 |
| NUBP1 | protein_coding | 0.29063652 | 4.39791196 |
| CCDC47 | protein_coding | 0.29058009 | 9.08198713 |
| SMAD1 | protein_coding | 0.29052486 | 6.28298003 |
| ZNF777 | protein_coding | 0.29024638 | 3.90638265 |
| ZNF582 | protein_coding | 0.29017206 | 2.24596648 |
| RABGAP1L | protein_coding | 0.29016936 | 5.30224133 |
| OTULIN-DT | lncRNA | 0.29015887 | 0.60880728 |
| CHRAC1 | protein_coding | 0.29001813 | 5.58869958 |
| RP11-138I1.4 | lncRNA | 0.28980491 | 0.33909669 |
| AC007563.5 | lncRNA | 0.28978352 | 0.33906744 |
| NET1 | protein_coding | 0.28970226 | 3.97285283 |
| UBE3D | protein_coding | 0.28956297 | 3.31094102 |
| TMEM128 | protein_coding | 0.2895075 | 3.80636438 |
| RPS6KA1 | protein_coding | 0.28916304 | 8.14646898 |
| ADCY6 | protein_coding | 0.28915983 | 3.65408654 |
| SUPT7L | protein_coding | 0.28909952 | 5.80312931 |
| RMDN3 | protein_coding | 0.28905085 | 5.48992465 |
| CELSR3 | protein_coding | 0.28891157 | 8.04630013 |
| BMPR2 | protein_coding | 0.28850656 | 5.93522282 |
| TOP1 | protein_coding | 0.28839101 | 8.53058416 |
| USE1 | protein_coding | 0.28809937 | 3.92903052 |
| ORC5 | protein_coding | 0.28786572 | 6.10935365 |
| SDC3 | protein_coding | 0.28782652 | 8.98504866 |
| CASK | protein_coding | 0.28777197 | 6.45561765 |
| SLC16A1-AS1 | lncRNA | 0.287684 | 3.0953781 |
| RXRA | protein_coding | 0.2876609 | 4.13771587 |
| MAP3K2 | protein_coding | 0.28756398 | 7.09938704 |
| FBXL12 | protein_coding | 0.28755158 | 3.60933362 |
| NME6 | protein_coding | 0.28738752 | 2.92876557 |
| SLC17A7 | protein_coding | 0.28720965 | 2.08208369 |
| VPS13D | protein_coding | 0.28718062 | 5.33322276 |
| WWP1 | protein_coding | 0.28708814 | 5.46049684 |
| ERLIN2 | protein_coding | 0.28687288 | 4.62018849 |
| TK2 | protein_coding | 0.28647148 | 2.15686648 |
| CADPS | protein_coding | 0.28641333 | 2.54965029 |
| MFSD5 | protein_coding | 0.28633067 | 3.80893801 |
| ARHGEF6 | protein_coding | 0.28626076 | 1.7787266 |
| NF1 | protein_coding | 0.28619768 | 7.94826242 |

| | | | |
|---|---|---|---|
| PLK2 | protein_coding | 0.28617541 | 1.30820498 |
| LINC00910 | lncRNA | 0.28605847 | 2.82821641 |
| PFDN2 | protein_coding | 0.28590128 | 6.63180476 |
| GLRX2 | protein_coding | 0.2858443 | 3.611394 |
| ZNF260 | protein_coding | 0.28582399 | 6.61695873 |
| MARF1 | protein_coding | 0.28553998 | 7.49711031 |
| TCIRG1 | protein_coding | 0.28552743 | 2.8054852 |
| MRPS5 | protein_coding | 0.28536311 | 7.28967068 |
| PTCHD4 | protein_coding | 0.28496937 | 1.9517289 |
| OGFRL1 | protein_coding | 0.28477227 | 4.83255365 |
| ZFP41 | protein_coding | 0.28476794 | 3.03164604 |
| SLC25A28 | protein_coding | 0.28474144 | 3.5496034 |
| SPATA6L | protein_coding | 0.28420675 | 1.14921393 |
| CHMP2B | protein_coding | 0.28417775 | 3.44406781 |
| TENT2 | protein_coding | 0.28407398 | 4.7776549 |
| LINC01977 | lncRNA | 0.28403844 | 2.15769986 |
| FMR1 | protein_coding | 0.28397192 | 6.06547262 |
| ZNF254 | protein_coding | 0.28383358 | 4.53537932 |
| ARL13B | protein_coding | 0.28383165 | 2.63633541 |
| ZNF117 | protein_coding | 0.28359833 | 1.97721583 |
| VPS13B | protein_coding | 0.28359374 | 7.0620407 |
| GHDC | protein_coding | 0.28356165 | 3.74196086 |
| GOLGB1 | protein_coding | 0.28354024 | 4.7775351 |
| COG7 | protein_coding | 0.28342176 | 3.50933144 |
| PITRM1 | protein_coding | 0.28338407 | 6.13589371 |
| TENM4 | protein_coding | 0.28306483 | 7.81431634 |
| CSTB | protein_coding | 0.28302989 | 5.15491749 |
| PTK7 | protein_coding | 0.28296155 | 9.47770862 |
| FAM81A | protein_coding | 0.28286458 | 2.55978081 |
| MSH3 | protein_coding | 0.2828608 | 5.82490126 |
| AC074286.1 | lncRNA | 0.28284464 | 0.46926836 |
| CHD9 | protein_coding | 0.28281646 | 7.40343759 |
| PTPRS | protein_coding | 0.28255064 | 5.29657653 |
| FAM217B | protein_coding | 0.28237003 | 3.73755082 |
| ALOX12-AS1 | lncRNA | 0.28231623 | 0.77605375 |
| CHD4 | protein_coding | 0.28226475 | 4.3744646 |
| ZBTB21 | protein_coding | 0.28224454 | 5.62452847 |
| LNC-LBCS | lncRNA | 0.28221889 | 0.42816392 |
| ANKRD10 | protein_coding | 0.28211256 | 5.75162833 |
| RP11-304L19 | lncRNA | 0.28209423 | 0.73454592 |
| PSMC4 | protein_coding | 0.28206122 | 8.42799367 |
| MCFD2 | protein_coding | 0.28202935 | 7.924062 |
| EXOC7 | protein_coding | 0.28177271 | 7.40433119 |
| PSMD3 | protein_coding | 0.28170162 | 7.48661394 |
| IBTK | protein_coding | 0.28150716 | 6.71528948 |

| | | | |
|---|---|---|---|
| SLC12A5-AS1 | lncRNA | 0.28125487 | 1.3958809 |
| RAB3D | protein_coding | 0.28106037 | 6.43678579 |
| SLC7A1 | protein_coding | 0.28101202 | 10.8188615 |
| LL0XNC01-7F | lncRNA | 0.28099942 | 1.50693966 |
| KIAA0319L | protein_coding | 0.28094967 | 6.79781029 |
| CYB5RL | protein_coding | 0.28090168 | 5.17045299 |
| LL0XNC01-23 | lncRNA | 0.28064109 | 0.93559539 |
| SDAD1 | protein_coding | 0.28032639 | 6.5038297 |
| CCDC88B | protein_coding | 0.28009156 | 3.27672678 |
| TBL3 | protein_coding | 0.28001203 | 5.42289272 |
| TBCK | protein_coding | 0.27995626 | 3.31389915 |
| LRSAM1 | protein_coding | 0.27984506 | 4.17966599 |
| PATE2 | protein_coding | 0.27910638 | 0.46095833 |
| MYCBP2 | protein_coding | 0.27909826 | 8.40494843 |
| LINC00324 | lncRNA | 0.27901184 | 0.35129384 |
| PGBD4 | protein_coding | 0.27898481 | 1.71540798 |
| ADI1 | protein_coding | 0.27892272 | 7.24589283 |
| RAB3GAP1 | protein_coding | 0.27890614 | 7.95568089 |
| CUL4A | protein_coding | 0.27882891 | 9.12342552 |
| MTIF2 | protein_coding | 0.27872944 | 6.17665154 |
| TEAD2 | protein_coding | 0.27847437 | 0.69848691 |
| MAP3K5 | protein_coding | 0.27817994 | 4.49336162 |
| DNAJC27 | protein_coding | 0.27816041 | 2.91016245 |
| MAPK6 | protein_coding | 0.27813343 | 7.44873832 |
| MANEA | protein_coding | 0.27804037 | 2.3059976 |
| CBWD6 | protein_coding | 0.27796975 | 0.97504981 |
| HMGN4 | protein_coding | 0.27783031 | 6.44260331 |
| USP24 | protein_coding | 0.27775669 | 6.42145889 |
| SMCR8 | protein_coding | 0.27774303 | 4.36824475 |
| KDM6B | protein_coding | 0.27768578 | 6.2836419 |
| JUND | protein_coding | 0.27760449 | 2.31049287 |
| EPRS1 | protein_coding | 0.27753033 | 9.37292272 |
| IL20RB | protein_coding | 0.27747946 | 0.65070854 |
| RBM28 | protein_coding | 0.27734359 | 7.21407302 |
| CAB39L | protein_coding | 0.27733602 | 2.06612639 |
| PSMC1 | protein_coding | 0.27716132 | 6.84747459 |
| OTX2 | protein_coding | 0.27708907 | 1.48188551 |
| ASH1L-AS1 | lncRNA | 0.27692706 | 1.56482828 |
| RAD51D | protein_coding | 0.27681296 | 5.30718803 |
| SHLD1 | protein_coding | 0.27677256 | 1.69338894 |
| FAM135A | protein_coding | 0.2764394 | 4.63328314 |
| MOGS | protein_coding | 0.276232 | 5.05959083 |
| CBLL1 | protein_coding | 0.27593249 | 4.72170345 |
| FAM199X | protein_coding | 0.27584882 | 3.43188837 |
| ZFP90 | protein_coding | 0.2755846 | 6.51378967 |

| | | | |
|---|---|---|---|
| LRRC37A3 | protein_coding | 0.27555054 | 1.21100029 |
| WDR74 | protein_coding | 0.27553302 | 5.17663959 |
| LINC01011 | lncRNA | 0.27536764 | 0.36637368 |
| MED12 | protein_coding | 0.27522604 | 6.00749234 |
| NUP50-DT | lncRNA | 0.27513665 | 1.86671398 |
| HSPA2 | protein_coding | 0.27506444 | 8.67074966 |
| RESF1 | protein_coding | 0.27502089 | 3.67616146 |
| UBE2Z | protein_coding | 0.27501738 | 7.70540608 |
| AXIN2 | protein_coding | 0.27501422 | 3.18880406 |
| DECR2 | protein_coding | 0.27493252 | 2.40900749 |
| NOC3L | protein_coding | 0.27485811 | 5.09159731 |
| MUTYH | protein_coding | 0.27485152 | 1.20273118 |
| RP11-807H2 | protein_coding | 0.27484444 | 0.29587466 |
| ST3GAL1-DT | lncRNA | 0.27464906 | 0.67436463 |
| PTPN6 | protein_coding | 0.27449592 | 3.5015977 |
| IDH1 | protein_coding | 0.27386339 | 5.49240507 |
| EHBP1 | protein_coding | 0.27372137 | 6.75249856 |
| RP11-1081L1 | lncRNA | 0.27350135 | 0.44527303 |
| USP42 | protein_coding | 0.27345448 | 4.82540576 |
| MYOZ3 | protein_coding | 0.2733477 | 0.48465775 |
| ABCD4 | protein_coding | 0.27318028 | 6.01116693 |
| CASQ2 | protein_coding | 0.27307222 | 1.09989297 |
| RP11-517B1 | lncRNA | 0.27280127 | 1.13545928 |
| PDCD6 | protein_coding | 0.27263616 | 2.22396607 |
| DENND11 | protein_coding | 0.27263407 | 6.29343088 |
| UBA5 | protein_coding | 0.27263076 | 4.94148929 |
| TTC13 | protein_coding | 0.27259788 | 3.63822875 |
| FDXR | protein_coding | 0.27255871 | 4.03472849 |
| CLIP4 | protein_coding | 0.27243445 | 2.97580295 |
| ZNF800 | protein_coding | 0.27243271 | 3.87130421 |
| WASH6P | protein_coding | 0.27242046 | 1.83751257 |
| EAF2 | protein_coding | 0.27239029 | 1.15471348 |
| TMEM131 | protein_coding | 0.27238806 | 8.09086426 |
| PLCB1 | protein_coding | 0.27237585 | 5.01007706 |
| NAT16 | protein_coding | 0.27234526 | 2.27662927 |
| STAC3 | protein_coding | 0.2722944 | 1.01744558 |
| STOML1 | protein_coding | 0.27229042 | 5.47661826 |
| NOS1AP | protein_coding | 0.27215753 | 3.3709314 |
| ARMC10 | protein_coding | 0.27213882 | 5.43740853 |
| ANKRD24 | protein_coding | 0.2718864 | 1.82114309 |
| ZNF382 | protein_coding | 0.27186242 | 4.76982476 |
| CCDC57 | protein_coding | 0.27182273 | 4.32146633 |
| FASTK | protein_coding | 0.27149597 | 5.68816153 |
| DENND1A | protein_coding | 0.27139712 | 6.0926415 |
| FEM1B | protein_coding | 0.27132185 | 7.55664267 |

| | | | |
|---|---|---|---|
| NCSTN | protein_coding | 0.27128096 | 6.3439292 |
| HMBOX1 | protein_coding | 0.27100645 | 2.82853772 |
| AFDN | protein_coding | 0.27083218 | 6.91746095 |
| ADD2 | protein_coding | 0.2701755 | 6.56872295 |
| ADGRB3 | protein_coding | 0.26999972 | 2.53919407 |
| CEP83 | protein_coding | 0.26981515 | 4.75267987 |
| ZNF585B | protein_coding | 0.26975405 | 3.55949971 |
| PLPP1 | protein_coding | 0.26975045 | 1.90240355 |
| RP11-441O1! | lncRNA | 0.26966705 | 0.90828807 |
| GOLPH3-DT | lncRNA | 0.26966386 | 0.91776731 |
| TXNRD1 | protein_coding | 0.26962352 | 9.35516557 |
| IPO4 | protein_coding | 0.26962078 | 0.40161716 |
| LMO1 | protein_coding | 0.26951824 | 3.89657313 |
| MLLT6 | protein_coding | 0.26936881 | 8.49852735 |
| PSMB6 | protein_coding | 0.26933489 | 6.67752824 |
| CCZ1B | protein_coding | 0.2692918 | 4.99334114 |
| SNX16 | protein_coding | 0.26918317 | 2.35052928 |
| LINC00923 | lncRNA | 0.26901583 | 1.57182389 |
| HOOK2 | protein_coding | 0.26900569 | 6.18735881 |
| SPAG8 | protein_coding | 0.26899722 | 0.42729295 |
| SNX11 | protein_coding | 0.26893589 | 4.76414206 |
| FAM83G | protein_coding | 0.26889769 | 5.46016504 |
| EFL1 | protein_coding | 0.26860894 | 6.68632701 |
| PHLPP2 | protein_coding | 0.26854858 | 5.2199771 |
| C3orf14 | protein_coding | 0.26854668 | 5.75825983 |
| MCL1 | protein_coding | 0.26851467 | 8.15283664 |
| ZNRF1 | protein_coding | 0.26851446 | 3.52163177 |
| N4BP2L2 | protein_coding | 0.2682846 | 5.76915379 |
| DNAJC13 | protein_coding | 0.26814391 | 6.77471476 |
| TRMT13 | protein_coding | 0.26805287 | 3.61709617 |
| ZNF786 | protein_coding | 0.26787883 | 2.69388077 |
| CCNL1 | protein_coding | 0.26783565 | 5.99761195 |
| GGPS1 | protein_coding | 0.26779707 | 2.4587544 |
| VLDLR-AS1 | lncRNA | 0.26768977 | 0.68376213 |
| NOL12 | protein_coding | 0.26758822 | 1.5241781 |
| AK6 | protein_coding | 0.26756057 | 3.13299912 |
| MTRR | protein_coding | 0.26754473 | 4.56695921 |
| ADRM1 | protein_coding | 0.26730562 | 5.63355794 |
| HSPA1A | protein_coding | 0.26724815 | 3.69164588 |
| SMAD2 | protein_coding | 0.2672003 | 8.07954794 |
| ZNF229 | protein_coding | 0.26705872 | 3.62414669 |
| UBFD1 | protein_coding | 0.26704175 | 7.06064597 |
| ARSF | protein_coding | 0.2668566 | 1.24728935 |
| MLLT3 | protein_coding | 0.26671501 | 5.50913059 |
| CHPF2 | protein_coding | 0.26657763 | 4.61224012 |

| | | | |
|---|---|---|---|
| DHRSX | protein_coding | 0.26635514 | 2.42372929 |
| SLC29A2 | protein_coding | 0.26578249 | 4.38013956 |
| INTS8 | protein_coding | 0.26576335 | 5.74267711 |
| NDUFAF2 | protein_coding | 0.26559566 | 3.58845168 |
| USP30 | protein_coding | 0.26557365 | 4.48509403 |
| CISH | protein_coding | 0.26550252 | 2.83010937 |
| GLRA3 | protein_coding | 0.26545388 | 1.22102667 |
| RP11-305F18 | lncRNA | 0.26538793 | 1.49754409 |
| STAM2 | protein_coding | 0.2652296 | 4.64821961 |
| EML6 | protein_coding | 0.26522479 | 3.83424508 |
| BAAT | protein_coding | 0.26511934 | 0.28366469 |
| MORC3 | protein_coding | 0.26510902 | 5.45126931 |
| PCTP | protein_coding | 0.26478717 | 1.7736112 |
| THTPA | protein_coding | 0.26468439 | 2.17337513 |
| MED1 | protein_coding | 0.26440901 | 9.01899462 |
| HERC1 | protein_coding | 0.26439717 | 8.60899206 |
| ZNF771 | protein_coding | 0.2641073 | 1.27133776 |
| CTD-2541M1 | lncRNA | 0.26410362 | 0.46359529 |
| CEROX1 | lncRNA | 0.26407175 | 0.5076122 |
| WDR59 | protein_coding | 0.26351914 | 5.11642168 |
| TES | protein_coding | 0.26339526 | 8.03984137 |
| ZNF627 | protein_coding | 0.26278767 | 3.55744714 |
| DCAF10 | protein_coding | 0.26248983 | 6.2467229 |
| HELB | protein_coding | 0.26242563 | 1.71905976 |
| CTB-1H10.1 | lncRNA | 0.26242299 | 0.25846545 |
| EPB41 | protein_coding | 0.26228536 | 9.89512883 |
| THOC7 | protein_coding | 0.261776 | 4.44168516 |
| NDUFA7 | protein_coding | 0.26172801 | 0.40386495 |
| COG1 | protein_coding | 0.26140397 | 5.95118381 |
| IQCH-AS1 | lncRNA | 0.26139698 | 2.76110691 |
| LINC01250 | lncRNA | 0.26137778 | 0.41008706 |
| APCDD1 | protein_coding | 0.26129382 | 0.68078023 |
| CNIH4 | protein_coding | 0.26126582 | 5.05281061 |
| CAPNS1 | protein_coding | 0.26123041 | 8.17852015 |
| SZT2 | protein_coding | 0.26109711 | 6.32266152 |
| NUAK1 | protein_coding | 0.26103193 | 3.51969941 |
| RP11-747H7. | lncRNA | 0.26100745 | 0.37880667 |
| KIF21A | protein_coding | 0.26081245 | 8.35931239 |
| MRPL38 | protein_coding | 0.26077871 | 1.18164273 |
| RP5-894A10. | lncRNA | 0.26051858 | 0.58390484 |
| KALRN | protein_coding | 0.26049214 | 1.37865592 |
| HCG18 | lncRNA | 0.26026769 | 4.93785715 |
| PSMA7 | protein_coding | 0.26019617 | 7.93439307 |
| TSGA10 | protein_coding | 0.25996514 | 2.18581418 |
| HMGCS1 | protein_coding | 0.25976017 | 8.06620704 |

| | | | |
|---|---|---|---|
| TEAD1 | protein_coding | 0.25975381 | 7.26197061 |
| TET3 | protein_coding | 0.25972634 | 6.38540895 |
| CAMK2N1 | protein_coding | 0.25964465 | 4.47618729 |
| CLSTN1 | protein_coding | 0.25960447 | 6.17189496 |
| NIPSNAP3B | protein_coding | 0.25946114 | 2.86662732 |
| ZEB2 | protein_coding | 0.25916682 | 4.45356566 |
| ELMOD3 | protein_coding | 0.25890071 | 2.74961013 |
| CCNG1 | protein_coding | 0.25882314 | 4.9624321 |
| NBPF12 | protein_coding | 0.25879872 | 2.50646518 |
| VPS8 | protein_coding | 0.25879312 | 5.91241044 |
| NAA25 | protein_coding | 0.25869617 | 6.1726531 |
| ST3GAL3 | protein_coding | 0.25857229 | 0.79353654 |
| AC073343.13 | lncRNA | 0.25855029 | 0.3432866 |
| ZNF250 | protein_coding | 0.25847717 | 4.60616503 |
| NDUFAF4 | protein_coding | 0.25820967 | 3.27044658 |
| MDM4 | protein_coding | 0.25820034 | 4.6760851 |
| KDM2A | protein_coding | 0.2581203 | 7.40811185 |
| HMBS | protein_coding | 0.25798675 | 5.26528582 |
| ACADM | protein_coding | 0.25798321 | 5.71179729 |
| CEP120 | protein_coding | 0.25780855 | 6.00910509 |
| AGA | protein_coding | 0.2577979 | 3.20310674 |
| NBPF26 | protein_coding | 0.25760988 | 0.3703389 |
| JCAD | protein_coding | 0.25756487 | 2.38254885 |
| GRIK2 | protein_coding | 0.25748974 | 4.62950491 |
| DRAP1 | protein_coding | 0.25708519 | 6.37150852 |
| BIVM | protein_coding | 0.25691008 | 3.32733272 |
| RP11-752G15 | lncRNA | 0.2568313 | 0.78491172 |
| CC2D1B | protein_coding | 0.256681 | 5.67555383 |
| LINC02802 | lncRNA | 0.25651796 | 0.39743592 |
| PUS10 | protein_coding | 0.25633835 | 2.03584699 |
| GABRB3 | protein_coding | 0.25628946 | 5.95988108 |
| CKAP4 | protein_coding | 0.25622709 | 5.37558649 |
| HHLA3 | lncRNA | 0.25620074 | 1.19166265 |
| SEZ6 | protein_coding | 0.25614729 | 6.0239498 |
| GUCY1A1 | protein_coding | 0.25584014 | 5.15290778 |
| WDR31 | protein_coding | 0.25580193 | 1.33500587 |
| SCN8A | protein_coding | 0.25576771 | 4.94674149 |
| USP27X | protein_coding | 0.25564251 | 1.26140849 |
| IGF1R | protein_coding | 0.25550876 | 5.38573253 |
| SNX19 | protein_coding | 0.25548195 | 6.03282928 |
| KIF18A | protein_coding | 0.25527341 | 4.68830265 |
| CD164L2 | protein_coding | 0.25523347 | 0.66360912 |
| VARS1 | protein_coding | 0.25513398 | 6.22210114 |
| LUZP2 | protein_coding | 0.25475846 | 1.57151109 |
| SGF29 | protein_coding | 0.2547088 | 3.73105896 |

| | | | |
|---|---|---|---|
| ATAT1 | protein_coding | 0.25459626 | 3.49383694 |
| C11orf24 | protein_coding | 0.25439205 | 4.57012032 |
| FAM161B | protein_coding | 0.25402997 | 4.55021025 |
| WDR27 | protein_coding | 0.25388254 | 1.80139425 |
| GTF2H2C | protein_coding | 0.25384189 | 2.34068081 |
| RP11-491F9. | lncRNA | 0.25382301 | 1.29257419 |
| CIAO2A | protein_coding | 0.25375078 | 5.60133515 |
| FCN2 | protein_coding | 0.25357308 | 0.78522676 |
| PCNX4 | protein_coding | 0.25355875 | 6.78569203 |
| DERL1 | protein_coding | 0.25351296 | 6.67188822 |
| EXOSC10 | protein_coding | 0.25347571 | 5.29368537 |
| ADCY1 | protein_coding | 0.25316304 | 10.2881625 |
| GPR176 | protein_coding | 0.25288214 | 2.23465346 |
| TM9SF1 | protein_coding | 0.25270945 | 1.52206895 |
| ZCCHC8 | protein_coding | 0.2524888 | 5.98871644 |
| CNOT1 | protein_coding | 0.2522752 | 9.43991256 |
| PLEKHM2 | protein_coding | 0.25225124 | 6.79974826 |
| MTOR | protein_coding | 0.25214161 | 6.65301955 |
| RHOU | protein_coding | 0.25202718 | 4.11236854 |
| RHBDD1 | protein_coding | 0.25199701 | 6.11542598 |
| LINC01237 | lncRNA | 0.25199129 | 0.58991617 |
| OFD1 | protein_coding | 0.25179203 | 5.36446591 |
| MAN2A1 | protein_coding | 0.25173528 | 4.3098895 |
| SOX8 | protein_coding | 0.25170095 | 0.34559735 |
| CEP85L | protein_coding | 0.25169627 | 2.24382816 |
| SPTBN1 | protein_coding | 0.25168535 | 8.99939067 |
| KPNA4 | protein_coding | 0.25159065 | 4.74719849 |
| LRIG2 | protein_coding | 0.25093264 | 4.28453328 |
| FNBP4 | protein_coding | 0.25081161 | 6.19402539 |
| ARHGAP5 | protein_coding | 0.2507433 | 6.555686 |
| MSH5 | protein_coding | 0.25064234 | 1.3666881 |
| PLAGL2 | protein_coding | 0.25058408 | 4.55150544 |
| AHDC1 | protein_coding | 0.25043433 | 3.8497572 |
| PDLIM2 | protein_coding | 0.2504326 | 0.54311889 |
| CLK1 | protein_coding | 0.25040147 | 5.98185159 |
| PBX2 | protein_coding | 0.25036209 | 5.50848649 |
| PRKRIP1 | protein_coding | 0.25035724 | 4.73985033 |
| ZNF671 | protein_coding | 0.25027697 | 2.11328519 |
| SPTSSB | protein_coding | 0.25027642 | 1.05538012 |
| SPNS1 | protein_coding | 0.25012685 | 0.38245351 |
| MED24 | protein_coding | 0.24996133 | 6.54426204 |
| PPIL3 | protein_coding | 0.24974719 | 3.16768597 |
| GPR89B | protein_coding | 0.24973645 | 1.0472181 |
| PTOV1-AS2 | lncRNA | 0.24942049 | 3.13344245 |
| ZNF461 | protein_coding | 0.24915065 | 2.27795401 |

| | | | |
|---|---|---|---|
| EML1 | protein_coding | 0.24912023 | 5.40968216 |
| SBF1 | protein_coding | 0.24907059 | 6.86215989 |
| ICE2 | protein_coding | 0.24896686 | 6.38217476 |
| POLG2 | protein_coding | 0.24890856 | 3.03856466 |
| TBX19 | protein_coding | 0.24888611 | 0.81323963 |
| CUX1 | protein_coding | 0.24885078 | 6.86799962 |
| TBC1D20 | protein_coding | 0.24879958 | 5.77273314 |
| SPPL2A | protein_coding | 0.24876981 | 5.53314434 |
| SBNO2 | protein_coding | 0.24875267 | 2.46665835 |
| RAB3GAP2 | protein_coding | 0.24875007 | 5.35966385 |
| CPNE1 | protein_coding | 0.24869921 | 5.54411254 |
| ZSCAN29 | protein_coding | 0.24849732 | 5.43305345 |
| RGS9 | protein_coding | 0.24827104 | 0.85167683 |
| POLI | protein_coding | 0.24810501 | 4.43333551 |
| ADCY3 | protein_coding | 0.24808155 | 7.00038758 |
| CREB3 | protein_coding | 0.24806059 | 4.0094176 |
| LYAR | protein_coding | 0.24787 | 4.18568761 |
| POC5 | protein_coding | 0.24774992 | 3.93891898 |
| ST14 | protein_coding | 0.247684 | 3.76652907 |
| ORC3 | protein_coding | 0.24738484 | 4.81798721 |
| KLF15 | protein_coding | 0.24730723 | 1.53209769 |
| TTPAL | protein_coding | 0.24708561 | 5.39492245 |
| U95743.1 | lncRNA | 0.24699023 | 1.54146147 |
| RP11-1348G | lncRNA | 0.24671138 | 0.37631922 |
| GAS8 | protein_coding | 0.2466839 | 4.01233831 |
| UTP14A | protein_coding | 0.2465062 | 4.3393655 |
| CCNYL1 | protein_coding | 0.24650112 | 2.93556246 |
| PPA2 | protein_coding | 0.24601933 | 4.30634585 |
| ZNF322 | protein_coding | 0.24589191 | 2.01768144 |
| TRIP12 | protein_coding | 0.2456857 | 8.35984645 |
| FAM160B1 | protein_coding | 0.24563408 | 2.57244621 |
| H6PD | protein_coding | 0.24556721 | 2.52108065 |
| ASMTL-AS1 | lncRNA | 0.24556563 | 0.71519051 |
| KCTD20 | protein_coding | 0.24528082 | 7.07348079 |
| CTHRC1 | protein_coding | 0.24527988 | 4.05936426 |
| ZNF8-ERVK3 | lncRNA | 0.24518975 | 0.57766645 |
| GAL3ST4 | protein_coding | 0.24515279 | 1.1737532 |
| ICAM3 | protein_coding | 0.24512046 | 0.39831922 |
| DIP2A | protein_coding | 0.24479871 | 5.91791524 |
| PGAP3 | protein_coding | 0.24476723 | 2.65191851 |
| MISP | protein_coding | 0.24471622 | 3.4334505 |
| AC005618.6 | lncRNA | 0.24469127 | 0.77021875 |
| ZNF554 | protein_coding | 0.24455897 | 2.18526091 |
| ZNF320 | protein_coding | 0.24451181 | 4.7291158 |
| PTPRG-AS1 | lncRNA | 0.24447685 | 2.68636523 |

| | | | |
|---|---|---|---|
| KSR1 | protein_coding | 0.24439646 | 5.26921209 |
| WASHC1 | protein_coding | 0.24434592 | 1.94699124 |
| ZBTB39 | protein_coding | 0.2440695 | 4.61590591 |
| SMURF1 | protein_coding | 0.24387842 | 4.60862474 |
| ZNF227 | protein_coding | 0.24385072 | 4.71044252 |
| TRIP4 | protein_coding | 0.24365009 | 3.68238284 |
| PACRGL | protein_coding | 0.24336621 | 4.11800574 |
| MON1A | protein_coding | 0.24328618 | 2.55076469 |
| SLC30A1 | protein_coding | 0.24327982 | 3.88160778 |
| KMT2D | protein_coding | 0.24323784 | 8.76917961 |
| MSANTD2 | protein_coding | 0.24313079 | 1.99643097 |
| COX10 | protein_coding | 0.2430641 | 2.27773681 |
| LINC01694 | lncRNA | 0.2430614 | 0.480143 |
| ZNF720 | protein_coding | 0.2430036 | 2.89542022 |
| DELE1 | protein_coding | 0.24297233 | 3.99095049 |
| RP11-884K1( | lncRNA | 0.2427358 | 0.54426133 |
| C9orf147 | lncRNA | 0.24252027 | 0.42993376 |
| YME1L1 | protein_coding | 0.24251788 | 7.24763597 |
| FBXO31 | protein_coding | 0.24233834 | 4.40271342 |
| RNF17 | protein_coding | 0.24227094 | 0.30973557 |
| LINC00106 | lncRNA | 0.24225897 | 0.46192103 |
| POGK | protein_coding | 0.24209078 | 7.72332485 |
| SLC25A42 | protein_coding | 0.24201748 | 5.09919705 |
| NRAS | protein_coding | 0.24200348 | 4.52663278 |
| ARVCF | protein_coding | 0.24179407 | 3.91983404 |
| SUDS3 | protein_coding | 0.24177006 | 6.89255315 |
| TEP1 | protein_coding | 0.24167356 | 2.99749009 |
| UBLCP1 | protein_coding | 0.24160872 | 4.49176154 |
| RP11-283I3.( | lncRNA | 0.24155279 | 1.94121792 |
| SGSH | protein_coding | 0.24128954 | 1.52806029 |
| RP11-629G1: | lncRNA | 0.24112453 | 0.34651231 |
| RP11-534C1: | lncRNA | 0.24109445 | 0.26443176 |
| ATP13A1 | protein_coding | 0.24104583 | 6.56237742 |
| TMX3 | protein_coding | 0.24099671 | 4.56111767 |
| NDEL1 | protein_coding | 0.24087227 | 2.45988379 |
| HEXA | protein_coding | 0.24082393 | 3.01766214 |
| PJVK | protein_coding | 0.24082344 | 1.20063566 |
| RP11-87C12. | lncRNA | 0.24067882 | 0.82141271 |
| TM7SF3 | protein_coding | 0.24060899 | 7.75895701 |
| SETD4 | protein_coding | 0.24050812 | 2.72324298 |
| ISY1 | protein_coding | 0.2404866 | 2.9931647 |
| CTD-2619J13 | lncRNA | 0.24033846 | 0.88672131 |
| FADS3 | protein_coding | 0.2403323 | 2.44882756 |
| ATR | protein_coding | 0.24027677 | 4.10849039 |
| GNG8 | protein_coding | 0.24019807 | 4.32648293 |

| | | | |
|---|---|---|---|
| NRXN3 | protein_coding | 0.24018509 | 6.50771636 |
| DNAJC24 | protein_coding | 0.24011568 | 4.29003435 |
| LINC00899 | lncRNA | 0.24004779 | 1.85667385 |
| RBM15B | protein_coding | 0.23997989 | 6.09682108 |
| MXD1 | protein_coding | 0.23996162 | 5.47360253 |
| TMEM65 | protein_coding | 0.23972538 | 2.55597668 |
| COG2 | protein_coding | 0.23967706 | 4.76445301 |
| PHF21A | protein_coding | 0.23965702 | 5.83050623 |
| MCEE | protein_coding | 0.23960986 | 2.36932204 |
| DPY19L1 | protein_coding | 0.23949971 | 2.51028194 |
| PEX2 | protein_coding | 0.23948447 | 2.36575565 |
| PFDN4 | protein_coding | 0.23946531 | 3.27580527 |
| DCP1B | protein_coding | 0.23902203 | 2.5632208 |
| UTP6 | protein_coding | 0.23894829 | 6.10666595 |
| CTB-193M12 | lncRNA | 0.23884559 | 2.35797347 |
| H2AJ | protein_coding | 0.23869175 | 3.04460007 |
| RP11-531A2: | lncRNA | 0.23866218 | 0.59257669 |
| YTHDC2 | protein_coding | 0.23809606 | 5.1649888 |
| EBLN3P | lncRNA | 0.23809026 | 5.07064813 |
| NUTM2A-AS | lncRNA | 0.23797941 | 2.44447324 |
| LHX3 | protein_coding | 0.23786255 | 3.32882705 |
| RP11-440D1: | lncRNA | 0.23765448 | 1.97685266 |
| UBE3C | protein_coding | 0.23762717 | 7.45127098 |
| ARL5B | protein_coding | 0.23747761 | 3.60457989 |
| INSIG1 | protein_coding | 0.2374687 | 6.43225609 |
| TBCA | protein_coding | 0.23735037 | 5.91538946 |
| WNK2 | protein_coding | 0.23733867 | 3.07277319 |
| SRPRA | protein_coding | 0.23721782 | 7.82833149 |
| GPR180 | protein_coding | 0.2371852 | 3.49539473 |
| BICRAL | protein_coding | 0.23714636 | 3.64200577 |
| GGT7 | protein_coding | 0.23709076 | 3.97361464 |
| RP11-359J14 | lncRNA | 0.23704992 | 0.23350282 |
| ADARB1 | protein_coding | 0.23699471 | 2.460233 |
| SNUPN | protein_coding | 0.23679546 | 2.76525535 |
| RP11-66D17. | lncRNA | 0.23676486 | 0.25269642 |
| CNOT2 | protein_coding | 0.23667758 | 5.68006849 |
| FKBP15 | protein_coding | 0.23659747 | 5.38158422 |
| CD46 | protein_coding | 0.23655716 | 4.89399079 |
| CLEC16A | protein_coding | 0.23649825 | 5.45241302 |
| FAHD2B | protein_coding | 0.23643781 | 1.67289928 |
| RP5-823G15. | lncRNA | 0.23641773 | 0.48475993 |
| GLG1 | protein_coding | 0.23635402 | 6.59746996 |
| FAM229A | protein_coding | 0.23633254 | 1.29051185 |
| TRIM26 | protein_coding | 0.23631564 | 5.79990804 |
| DDX39B | protein_coding | 0.2362262 | 6.48595278 |

| | | | |
|---|---|---|---|
| SYT5 | protein_coding | 0.23622471 | 0.94708654 |
| AC079305.1( | lncRNA | 0.23604229 | 0.26561504 |
| TAF9 | protein_coding | 0.235981 | 4.60838073 |
| ZNF692 | protein_coding | 0.23588839 | 4.29223779 |
| DPH3 | protein_coding | 0.23579875 | 2.7599385 |
| HECTD4 | protein_coding | 0.23578653 | 7.37596722 |
| LAG3 | protein_coding | 0.23575713 | 0.61067225 |
| LRRCC1 | protein_coding | 0.23564203 | 3.97683532 |
| CCNI | protein_coding | 0.23552477 | 8.62111482 |
| SLC9A5 | protein_coding | 0.2354059 | 0.59199463 |
| CEBPA | protein_coding | 0.23503488 | 0.23131454 |
| CCDC167 | protein_coding | 0.23501695 | 2.69493724 |
| PPP1R12B | protein_coding | 0.23474861 | 3.98818727 |
| USP35 | protein_coding | 0.23474123 | 1.78823253 |
| TMEM41A | protein_coding | 0.23469344 | 4.02043034 |
| ITFG1 | protein_coding | 0.23468624 | 5.72732119 |
| ZNF211 | protein_coding | 0.2346612 | 3.19400143 |
| MCOLN3 | protein_coding | 0.2346152 | 4.50734077 |
| TMEM42 | protein_coding | 0.23452903 | 1.40147659 |
| PSMB3 | protein_coding | 0.23452536 | 7.06368932 |
| SYNE2 | protein_coding | 0.23449818 | 9.02839756 |
| ETV3 | protein_coding | 0.23434173 | 4.29755347 |
| GS1-57L11.1 | lncRNA | 0.23425055 | 0.92513158 |
| NFX1 | protein_coding | 0.23420081 | 6.61573593 |
| ADSL | protein_coding | 0.23418357 | 2.18933892 |
| DDX17 | protein_coding | 0.234095 | 9.26245769 |
| ROCK2 | protein_coding | 0.23386057 | 5.71971747 |
| TAF1A | protein_coding | 0.23378569 | 2.16831893 |
| DUS4L | protein_coding | 0.23335999 | 1.47166285 |
| TTF2 | protein_coding | 0.23334759 | 5.49659939 |
| SNX6 | protein_coding | 0.23328117 | 5.79994969 |
| NDUFA9 | protein_coding | 0.23325723 | 2.81458087 |
| STK10 | protein_coding | 0.23324987 | 0.87334939 |
| DCTN4 | protein_coding | 0.23317142 | 6.77162979 |
| ESF1 | protein_coding | 0.23304966 | 3.84918864 |
| AKR7A3 | protein_coding | 0.23304222 | 0.47723314 |
| DENND2D | protein_coding | 0.23292565 | 1.53354394 |
| RALGAPA1 | protein_coding | 0.23283286 | 4.55207896 |
| RNF10 | protein_coding | 0.23281247 | 5.96923334 |
| POLR3D | protein_coding | 0.23277936 | 4.30321036 |
| LIMK1 | protein_coding | 0.23242249 | 4.21470304 |
| CDC40 | protein_coding | 0.23237719 | 3.53268574 |
| ZNF594 | protein_coding | 0.23223528 | 3.08874199 |
| MAT2A | protein_coding | 0.23218759 | 8.08269728 |
| SETDB1 | protein_coding | 0.23218474 | 5.81568627 |

| | | | |
|---|---|---|---|
| ZNF594-DT | lncRNA | 0.23192184 | 0.62612347 |
| CCDC112 | protein_coding | 0.23168366 | 3.32331467 |
| GFY | protein_coding | 0.23156413 | 1.14811849 |
| CBX7 | protein_coding | 0.23152729 | 3.24800353 |
| BRPF3 | protein_coding | 0.23151048 | 7.04217748 |
| UBE2F | protein_coding | 0.23138638 | 1.53389827 |
| PDCD11 | protein_coding | 0.23125599 | 6.19748834 |
| ATP7A | protein_coding | 0.23120886 | 3.62498104 |
| NEK10 | protein_coding | 0.23078643 | 0.47248894 |
| MYSM1 | protein_coding | 0.23054886 | 3.62639163 |
| COMTD1 | protein_coding | 0.23052685 | 1.46152044 |
| RP11-302L19 | lncRNA | 0.23045532 | 0.75857792 |
| SRPK3 | protein_coding | 0.23039782 | 3.01074904 |
| CTC-459F4.9 | lncRNA | 0.2303294 | 0.37212344 |
| CIPC | protein_coding | 0.22995076 | 5.85834221 |
| LINC00621 | lncRNA | 0.22994003 | 0.60020621 |
| LONRF2 | protein_coding | 0.22980953 | 8.30570345 |
| OXNAD1 | protein_coding | 0.22976708 | 2.53737251 |
| MLH3 | protein_coding | 0.22973291 | 4.48833151 |
| SETD5 | protein_coding | 0.22948791 | 6.09890389 |
| ENKD1 | protein_coding | 0.22933339 | 2.1210654 |
| FXN | protein_coding | 0.22932186 | 1.62038577 |
| TRPM7 | protein_coding | 0.22914992 | 5.8907785 |
| RBM24 | protein_coding | 0.22890463 | 6.50608624 |
| KDM6A | protein_coding | 0.22883518 | 4.08665389 |
| TMEM69 | protein_coding | 0.2288226 | 4.40555692 |
| ST6GAL2 | protein_coding | 0.22862819 | 0.42520557 |
| PIGN | protein_coding | 0.22853438 | 4.43876076 |
| ATG16L2 | protein_coding | 0.22842422 | 3.1560692 |
| SLC10A7 | protein_coding | 0.2284188 | 2.03018735 |
| SRM | protein_coding | 0.22840057 | 4.5569581 |
| R3HDM2 | protein_coding | 0.22828175 | 5.17848633 |
| CFL2 | protein_coding | 0.22827937 | 3.38591167 |
| SEPTIN8 | protein_coding | 0.22823114 | 7.37617623 |
| PBRM1 | protein_coding | 0.22817439 | 3.84082708 |
| RP11-431H16 | lncRNA | 0.22815842 | 0.28116548 |
| U2AF1L5 | protein_coding | 0.22809739 | 0.73385004 |
| UBE2D1 | protein_coding | 0.2279632 | 1.85863706 |
| KCNK9 | protein_coding | 0.2279293 | 3.60942148 |
| ZFAND3 | protein_coding | 0.22791047 | 5.51559505 |
| AC004076.5 | lncRNA | 0.2277987 | 0.69384445 |
| ABCB9 | protein_coding | 0.2277465 | 4.87983425 |
| LTB4R | protein_coding | 0.22771968 | 2.82434044 |
| ADAM10 | protein_coding | 0.2275656 | 4.84570688 |
| ZNF274 | protein_coding | 0.22744478 | 3.98359649 |

| | | | |
|---|---|---|---|
| SYT9 | protein_coding | 0.22742291 | 4.83554887 |
| VWA8 | protein_coding | 0.22736142 | 3.14018758 |
| C11orf68 | protein_coding | 0.22730773 | 4.29771907 |
| SCGB2A1 | protein_coding | 0.227211 | 3.11021344 |
| RP5-1092A3. | lncRNA | 0.22718225 | 0.54933026 |
| IQCK | protein_coding | 0.22717808 | 1.99282851 |
| SNRPN | protein_coding | 0.22707874 | 1.14786711 |
| PSMD5 | protein_coding | 0.22694914 | 5.74215487 |
| CDK5R2 | protein_coding | 0.22694401 | 4.17061938 |
| BRI3 | protein_coding | 0.2268973 | 3.98430053 |
| IP6K1 | protein_coding | 0.22662971 | 4.03509456 |
| HEATR1 | protein_coding | 0.22655863 | 5.79268465 |
| LPIN1 | protein_coding | 0.22638797 | 4.64331584 |
| LINC02287 | lncRNA | 0.22618509 | 0.44274052 |
| LRP12 | protein_coding | 0.22610767 | 4.59388468 |
| ZNF517 | protein_coding | 0.22608698 | 1.31382439 |
| EPOR | protein_coding | 0.2259923 | 1.25102021 |
| UBE2B | protein_coding | 0.2259804 | 2.95838455 |
| BBS2 | protein_coding | 0.22573882 | 2.75242417 |
| GIN1 | protein_coding | 0.22565101 | 1.82119706 |
| TBRG1 | protein_coding | 0.22558604 | 4.6648882 |
| MNAT1 | protein_coding | 0.22544115 | 2.42724548 |
| SLC41A1 | protein_coding | 0.22541704 | 4.94834863 |
| PGGT1B | protein_coding | 0.22539333 | 3.67495555 |
| ZNF146 | protein_coding | 0.22529023 | 8.01141691 |
| PCED1A | protein_coding | 0.22527905 | 3.35895384 |
| RP11-263K19 | lncRNA | 0.22521936 | 0.2365334 |
| CFAP61 | protein_coding | 0.22512774 | 2.23532831 |
| XBP1 | protein_coding | 0.22508543 | 5.28398748 |
| EPC2 | protein_coding | 0.22497789 | 4.78984629 |
| TIMM23B | protein_coding | 0.22455831 | 0.6696492 |
| CCDC25 | protein_coding | 0.22454952 | 4.49876551 |
| FNDC3B | protein_coding | 0.22448813 | 3.00012201 |
| PKP2 | protein_coding | 0.22441407 | 6.28891869 |
| ABCC5 | protein_coding | 0.22438798 | 7.11617876 |
| CBSL | protein_coding | 0.22430811 | 0.25607206 |
| STK36 | protein_coding | 0.22406356 | 2.2630702 |
| ALDH5A1 | protein_coding | 0.22397437 | 4.84403923 |
| TKT | protein_coding | 0.22393363 | 7.04772635 |
| NFXL1 | protein_coding | 0.22387003 | 4.05855634 |
| HACL1 | protein_coding | 0.22374167 | 1.99897009 |
| SUPT20H | protein_coding | 0.2237035 | 4.77447811 |
| BACE1-AS | lncRNA | 0.22369508 | 2.57216697 |
| SLC9B2 | protein_coding | 0.22367965 | 3.15603143 |
| MORN4 | protein_coding | 0.22344045 | 2.62601246 |

| | | | |
|---|---|---|---|
| SCFD1 | protein_coding | 0.22323571 | 4.92948106 |
| ZNF444 | protein_coding | 0.22309927 | 1.75954248 |
| SPATA13 | protein_coding | 0.22308581 | 4.85083454 |
| AVEN | protein_coding | 0.22303769 | 1.02018869 |
| POT1 | protein_coding | 0.22293711 | 4.54741381 |
| RAB2B | protein_coding | 0.22272899 | 4.40965934 |
| DDX59 | protein_coding | 0.22270424 | 3.41506892 |
| NR1H3 | protein_coding | 0.22264673 | 1.8165063 |
| CTD-2035E11 | lncRNA | 0.22251199 | 0.53324515 |
| NUP88 | protein_coding | 0.22245062 | 4.91632336 |
| TRUB1 | protein_coding | 0.22242547 | 3.29311164 |
| KRBA2 | protein_coding | 0.22238028 | 1.53638237 |
| RP11-48B3.5 | lncRNA | 0.22236368 | 0.53862524 |
| FITM1 | protein_coding | 0.22221748 | 0.25789638 |
| BAIAP2L2 | protein_coding | 0.22215081 | 0.46316624 |
| UBQLN4 | protein_coding | 0.22211414 | 5.59619099 |
| WDR19 | protein_coding | 0.22209363 | 3.07696005 |
| TGOLN2 | protein_coding | 0.22188332 | 7.80874933 |
| ATP12A | protein_coding | 0.2217745 | 0.37009274 |
| ZNF620 | protein_coding | 0.22174107 | 2.45154813 |
| AGER | protein_coding | 0.22169083 | 1.33198814 |
| LMOD2 | protein_coding | 0.221665 | 1.09756549 |
| IGIP | protein_coding | 0.22161749 | 1.18675426 |
| PRPF39 | protein_coding | 0.22158127 | 4.83696499 |
| ZNF264 | protein_coding | 0.22149672 | 4.3508431 |
| KIF2A | protein_coding | 0.22148495 | 7.93048936 |
| PSMD2 | protein_coding | 0.22147237 | 8.86068569 |
| SEPTIN4 | protein_coding | 0.22107323 | 0.92827167 |
| AP5Z1 | protein_coding | 0.22104986 | 3.06620012 |
| GABPB1-AS1 | lncRNA | 0.22093368 | 4.59669226 |
| ZNF607 | protein_coding | 0.2208656 | 3.39203422 |
| THSD7A | protein_coding | 0.22082015 | 1.80338012 |
| EML3 | protein_coding | 0.2204708 | 4.47898141 |
| ZNF235 | protein_coding | 0.22044155 | 2.15556334 |
| MYO9A | protein_coding | 0.22038967 | 4.3433549 |
| TMEM170A | protein_coding | 0.22026568 | 3.0283764 |
| CBWD2 | protein_coding | 0.22020544 | 2.67263365 |
| ZNF606 | protein_coding | 0.22017706 | 2.73488869 |
| RABGGTB | protein_coding | 0.21995425 | 3.93682754 |
| ZNF331 | protein_coding | 0.2198862 | 3.59268917 |
| HUS1 | protein_coding | 0.21985609 | 2.56139802 |
| TAF1C | protein_coding | 0.21983563 | 5.26294974 |
| TXNDC16 | protein_coding | 0.21980717 | 4.94919446 |
| CREBBP | protein_coding | 0.21967073 | 6.6773171 |
| MED26 | protein_coding | 0.21964799 | 2.03931677 |

| | | | |
|---|---|---|---|
| CAMKK2 | protein_coding | 0.2196357 | 5.81310456 |
| CYBB | protein_coding | 0.21954225 | 0.68399153 |
| ASL | protein_coding | 0.21948904 | 1.0908816 |
| RP11-250B2. | lncRNA | 0.21940799 | 0.77465897 |
| KDM4A | protein_coding | 0.21922053 | 5.13096655 |
| MAP3K12 | protein_coding | 0.21905901 | 2.00608689 |
| AL133243.1 | lncRNA | 0.21899452 | 0.39554165 |
| UPF3A | protein_coding | 0.21883303 | 5.24220454 |
| LUC7L | protein_coding | 0.21876176 | 4.99483541 |
| TCTE1 | protein_coding | 0.21849375 | 0.49581609 |
| SRP19 | protein_coding | 0.21841234 | 2.97698078 |
| MMRN2 | protein_coding | 0.21830871 | 0.80424688 |
| ATP2C1 | protein_coding | 0.21823589 | 7.26212807 |
| RAB11FIP1 | protein_coding | 0.21805373 | 3.98622743 |
| TNFAIP1 | protein_coding | 0.2179771 | 4.96047932 |
| TMEM62 | protein_coding | 0.2179757 | 4.57326107 |
| LINC00960 | lncRNA | 0.21778099 | 1.70633727 |
| MAP4K2 | protein_coding | 0.21776678 | 7.24711369 |
| EMX2OS | lncRNA | 0.21776467 | 3.55923649 |
| RIOK1 | protein_coding | 0.21765465 | 5.55124087 |
| CD59 | protein_coding | 0.21762219 | 2.75478257 |
| LTV1 | protein_coding | 0.21747142 | 2.11096726 |
| ZNF182 | protein_coding | 0.21735725 | 2.85883108 |
| RLIM | protein_coding | 0.21729844 | 5.42183454 |
| RP11-692D1: | lncRNA | 0.21697413 | 0.26023124 |
| STX18-AS1 | lncRNA | 0.21692599 | 1.86980647 |
| SLC35D1 | protein_coding | 0.21687534 | 2.62010461 |
| KCP | protein_coding | 0.21677548 | 0.35343372 |
| CWC25 | protein_coding | 0.21664354 | 3.77818289 |
| OTUB2 | protein_coding | 0.21660034 | 1.97350002 |
| LOXL1-AS1 | lncRNA | 0.21657024 | 2.65480896 |
| UBR5 | protein_coding | 0.21637113 | 7.34932557 |
| EIF3J | protein_coding | 0.21631275 | 6.65024488 |
| TMEM43 | protein_coding | 0.21624365 | 3.20285848 |
| RP11-424G1 | lncRNA | 0.21618378 | 0.24996272 |
| CRELD2 | protein_coding | 0.21598189 | 4.64652367 |
| TPT1-AS1 | lncRNA | 0.21597264 | 0.83107095 |
| MMP9 | protein_coding | 0.21595458 | 1.05920838 |
| EFCAB7 | protein_coding | 0.21593939 | 2.35364195 |
| N4BP2L2-IT2 | lncRNA | 0.21557937 | 0.33603302 |
| FAM43A | protein_coding | 0.21556511 | 1.42785411 |
| ADAM15 | protein_coding | 0.21541827 | 3.97451132 |
| IFNGR2 | protein_coding | 0.21539594 | 4.12845477 |
| BTBD3 | protein_coding | 0.21532472 | 5.04813671 |
| SNX15 | protein_coding | 0.21523443 | 0.6503713 |

| | | | |
|---|---|---|---|
| ATAD3C | protein_coding | 0.2151862 | 1.10746939 |
| TGDS | protein_coding | 0.21518235 | 2.25555401 |
| ENTR1 | protein_coding | 0.21515535 | 5.18119462 |
| OAZ1 | protein_coding | 0.21501457 | 6.67283366 |
| TUBGCP6 | protein_coding | 0.21501348 | 5.22705439 |
| PARD3 | protein_coding | 0.21495694 | 2.97558839 |
| BOLA3 | protein_coding | 0.21495025 | 3.09963684 |
| C2CD6 | protein_coding | 0.21489461 | 0.8263117 |
| POLR3G | protein_coding | 0.21458028 | 1.87334445 |
| ELFN2 | lncRNA | 0.21454996 | 0.62326036 |
| EIF2AK4 | protein_coding | 0.21444624 | 3.92953454 |
| PHACTR4 | protein_coding | 0.21439924 | 5.19926331 |
| TLN2 | protein_coding | 0.21429347 | 4.41368812 |
| CYTH1 | protein_coding | 0.21426006 | 3.71164951 |
| CLIC1 | protein_coding | 0.21410423 | 6.47989042 |
| PAK1IP1 | protein_coding | 0.2139649 | 4.90375256 |
| RP11-819C21 | lncRNA | 0.21391909 | 0.7171343 |
| MPV17 | protein_coding | 0.21390808 | 3.23081188 |
| SNX24 | protein_coding | 0.21386518 | 1.08230369 |
| CTD-2319I12 | lncRNA | 0.21353495 | 1.11804879 |
| TMEM123 | protein_coding | 0.21350292 | 5.1966468 |
| PPM1A | protein_coding | 0.2134971 | 5.50891069 |
| SNHG32 | lncRNA | 0.21344327 | 2.56055225 |
| RNF14 | protein_coding | 0.21331978 | 5.90716781 |
| PCSK6 | protein_coding | 0.2132153 | 6.70424342 |
| COL9A3 | protein_coding | 0.21321208 | 1.49232316 |
| TRAPPC8 | protein_coding | 0.21306455 | 5.28302754 |
| DUS3L | protein_coding | 0.21305961 | 1.74317931 |
| MAN2A2 | protein_coding | 0.21288173 | 6.93664557 |
| RRP9 | protein_coding | 0.21266773 | 2.5701287 |
| RP11-443B2( | lncRNA | 0.21264755 | 0.61840418 |
| TSC22D4 | protein_coding | 0.21261731 | 2.78926621 |
| UBA3 | protein_coding | 0.2125813 | 2.54436921 |
| USP47 | protein_coding | 0.21258053 | 5.14662549 |
| ABHD13 | protein_coding | 0.21251277 | 2.03597495 |
| MRPL15 | protein_coding | 0.21248058 | 3.57364197 |
| TAF15 | protein_coding | 0.2122429 | 7.42509479 |
| NOTCH2 | protein_coding | 0.21216261 | 2.73040655 |
| EDEM1 | protein_coding | 0.21212731 | 2.24052505 |
| ZNF226 | protein_coding | 0.21211444 | 3.08609562 |
| ZNF285 | protein_coding | 0.21201971 | 1.54694292 |
| SYT4 | protein_coding | 0.21189203 | 4.89920628 |
| LARP1 | protein_coding | 0.2118877 | 7.35014982 |
| PSMD4 | protein_coding | 0.21185891 | 7.09367553 |
| NEDD1 | protein_coding | 0.21183232 | 4.67895327 |

| | | | |
|---|---|---|---|
| MRPS6 | protein_coding | 0.21168341 | 3.0027761 |
| GALNT18 | protein_coding | 0.21165188 | 3.0079198 |
| CDK5RAP1 | protein_coding | 0.21156891 | 3.41618037 |
| UBAP1 | protein_coding | 0.21153229 | 4.87047163 |
| PRICKLE1 | protein_coding | 0.21147745 | 1.61999453 |
| FAM214A | protein_coding | 0.21136769 | 3.94850781 |
| ILKAP | protein_coding | 0.21129155 | 4.39435213 |
| CD200 | protein_coding | 0.21108487 | 3.1709995 |
| LAMB1 | protein_coding | 0.21090962 | 7.20935096 |
| CNTF | protein_coding | 0.21083938 | 0.5845003 |
| HIPK2 | protein_coding | 0.21078048 | 6.86834589 |
| POU3F3 | protein_coding | 0.21077455 | 1.78291747 |
| GXYLT1 | protein_coding | 0.21074714 | 4.79399666 |
| ZNF205 | protein_coding | 0.21050309 | 2.49799418 |
| RSPH3 | protein_coding | 0.21046344 | 1.24660276 |
| ZNF587 | protein_coding | 0.21015968 | 2.90128147 |
| C7orf31 | protein_coding | 0.21008306 | 1.37496589 |
| RBM33 | protein_coding | 0.20987729 | 5.81110864 |
| AP001429.1 | lncRNA | 0.20983608 | 0.1817515 |
| RBFA | protein_coding | 0.20968002 | 2.86542029 |
| PRRC2C | protein_coding | 0.20965048 | 7.39930727 |
| STK19 | protein_coding | 0.2096349 | 2.91197647 |
| RP4-756H11. | lncRNA | 0.20962282 | 1.30750626 |
| TUBG2 | protein_coding | 0.20951396 | 3.28337036 |
| FPGT-TNNI3 | protein_coding | 0.20944827 | 0.28175679 |
| SOGA1 | protein_coding | 0.20943301 | 7.5530221 |
| SCRT1 | protein_coding | 0.20929399 | 0.49914361 |
| LINC01409 | lncRNA | 0.20928651 | 0.2651402 |
| IQCJ-SCHIP1 | protein_coding | 0.20923476 | 0.64444354 |
| PABIR3 | protein_coding | 0.20922803 | 1.29831137 |
| TENT4B | protein_coding | 0.20907584 | 4.47284214 |
| OSGEP | protein_coding | 0.2086234 | 3.43283805 |
| ST8SIA3 | protein_coding | 0.2085968 | 4.80857181 |
| ZNF579 | protein_coding | 0.20857254 | 1.03585936 |
| LONP1 | protein_coding | 0.20833044 | 5.76997417 |
| GTF3C3 | protein_coding | 0.20826379 | 4.56943393 |
| DNAJA3 | protein_coding | 0.20819367 | 7.00279927 |
| BSN | protein_coding | 0.20815316 | 4.46055369 |
| INTS7 | protein_coding | 0.20814865 | 6.02163312 |
| IDUA | protein_coding | 0.20807743 | 0.94619647 |
| TENM3 | protein_coding | 0.20790168 | 6.70646349 |
| ARMC5 | protein_coding | 0.20788784 | 2.70776675 |
| SETX | protein_coding | 0.20778679 | 6.15780772 |
| GNAI2 | protein_coding | 0.2076844 | 6.14580384 |
| CBY2 | protein_coding | 0.20749628 | 0.24360854 |

| | | | |
|---|---|---|---|
| PI4K2B | protein_coding | 0.20728752 | 4.27613755 |
| ARMCX5 | protein_coding | 0.2072854 | 1.93771847 |
| KMT2A | protein_coding | 0.20710036 | 7.03909004 |
| RP11-37C7.3 | lncRNA | 0.20707026 | 0.35310149 |
| MAP3K6 | protein_coding | 0.20696714 | 3.80029569 |
| TAGLN | protein_coding | 0.20696384 | 0.78532119 |
| GSTK1 | protein_coding | 0.20690505 | 2.18566688 |
| HMGN5 | protein_coding | 0.20689504 | 3.8712151 |
| NANS | protein_coding | 0.20667466 | 4.55861185 |
| GPBP1 | protein_coding | 0.20664068 | 5.98646617 |
| SAP30BP | protein_coding | 0.20657463 | 4.97275028 |
| GLB1 | protein_coding | 0.20654821 | 2.67445948 |
| PFKFB2 | protein_coding | 0.20635864 | 4.44989166 |
| S100A2 | protein_coding | 0.20630026 | 0.28734364 |
| PARP1 | protein_coding | 0.20627318 | 8.76826188 |
| ZNF787 | protein_coding | 0.20623209 | 1.42145816 |
| COL11A2 | protein_coding | 0.20619157 | 0.8456173 |
| ZFYVE26 | protein_coding | 0.2060477 | 3.71487116 |
| C6orf89 | protein_coding | 0.20596572 | 7.02286939 |
| PRPH | protein_coding | 0.20587324 | 1.71139805 |
| CHDH | protein_coding | 0.20571707 | 3.3558815 |
| SH3GLB1 | protein_coding | 0.20566544 | 5.32641411 |
| PPP1R15B | protein_coding | 0.2055163 | 6.22675327 |
| CLCN4 | protein_coding | 0.20532031 | 1.73990991 |
| TIMMDC1 | protein_coding | 0.20521875 | 3.82374264 |
| CCDC183-AS1 | lncRNA | 0.20515396 | 1.28512846 |
| FOXO3 | protein_coding | 0.20508592 | 3.8997187 |
| GALNT7 | protein_coding | 0.2050473 | 2.51715717 |
| ZCCHC4 | protein_coding | 0.20502602 | 2.73177469 |
| LIN54 | protein_coding | 0.20496813 | 3.04644127 |
| XIAP | protein_coding | 0.20492471 | 3.69380408 |
| TCEANC | protein_coding | 0.20477871 | 1.21520744 |
| RP1-191J18.6 | lncRNA | 0.20470286 | 0.2948834 |
| C15orf40 | protein_coding | 0.20469709 | 2.5067918 |
| RAB28 | protein_coding | 0.2046106 | 3.03751609 |
| CENPT | protein_coding | 0.20460558 | 3.81682631 |
| ATG2B | protein_coding | 0.20448449 | 5.51960327 |
| GRPEL2 | protein_coding | 0.20446198 | 3.80814701 |
| UBXN8 | protein_coding | 0.20416908 | 2.18136329 |
| ZNF439 | protein_coding | 0.20406222 | 0.22648156 |
| NUFIP1 | protein_coding | 0.20382953 | 1.49490013 |
| TRAPPC13 | protein_coding | 0.20382227 | 3.79411728 |
| CRB3 | protein_coding | 0.20374707 | 1.79562937 |
| C5orf34 | protein_coding | 0.20371485 | 1.50535807 |
| ZFAS1 | lncRNA | 0.203683 | 2.95330293 |

| | | | |
|---|---|---|---|
| TMEM167B | protein_coding | 0.20365057 | 3.66909201 |
| BAZ1A | protein_coding | 0.20355262 | 6.90397065 |
| TMEM126A | protein_coding | 0.20334378 | 3.01475994 |
| THAP3 | protein_coding | 0.20309401 | 1.51735706 |
| SIPA1L1 | protein_coding | 0.20302861 | 5.28236762 |
| PTPRU | protein_coding | 0.20287501 | 1.42252988 |
| PLAC8 | protein_coding | 0.20279937 | 0.39587916 |
| PMPCB | protein_coding | 0.20271304 | 5.02619836 |
| KAT2A | protein_coding | 0.20270096 | 5.64165983 |
| PWP1 | protein_coding | 0.20261905 | 4.39639823 |
| FOXA3 | protein_coding | 0.20261108 | 2.50129389 |
| AKAP13 | protein_coding | 0.20258868 | 5.02464094 |
| RP11-35G9.5 | lncRNA | 0.20248855 | 0.9028326 |
| RAD18 | protein_coding | 0.20247595 | 2.54461382 |
| C3orf35 | lncRNA | 0.2022718 | 0.2619718 |
| KIAA0100 | protein_coding | 0.20216466 | 7.47412976 |
| CASC15 | lncRNA | 0.20212423 | 0.1859727 |
| CCDC22 | protein_coding | 0.20202229 | 2.68079116 |
| MAL2 | protein_coding | 0.20201661 | 4.03854642 |
| MIPEP | protein_coding | 0.20198667 | 3.12572481 |
| RGP1 | protein_coding | 0.20193859 | 5.76221269 |
| PDE4DIP | protein_coding | 0.20186611 | 2.64438508 |
| AKNAD1 | protein_coding | 0.20179459 | 1.52176875 |
| STRIP1 | protein_coding | 0.20175354 | 4.32413266 |
| TSPAN1 | protein_coding | 0.20169311 | 0.19363326 |
| CDK12 | protein_coding | 0.20168558 | 6.01839068 |
| PCSK2 | protein_coding | 0.20164718 | 6.76205201 |
| MRPL37 | protein_coding | 0.20163617 | 5.89759138 |
| LRRC8B | protein_coding | 0.2015555 | 4.03266957 |
| DNAJB14 | protein_coding | 0.2013419 | 3.119789 |
| DCLRE1A | protein_coding | 0.20129654 | 3.36429086 |
| WDR24 | protein_coding | 0.20106421 | 1.99093921 |
| RPAP2 | protein_coding | 0.20105468 | 4.31551592 |
| BSDC1 | protein_coding | 0.20103159 | 6.11189807 |
| AIMP2 | protein_coding | 0.20084551 | 4.59031157 |
| PLEKHH3 | protein_coding | 0.20077447 | 2.78878988 |
| VAMP3 | protein_coding | 0.20072818 | 2.69707133 |
| AFTPH | protein_coding | 0.20061291 | 4.6201297 |
| ADA | protein_coding | 0.20056566 | 0.3636213 |
| MLLT10 | protein_coding | 0.20055877 | 3.12632697 |
| GEMIN2 | protein_coding | 0.20045003 | 3.23110589 |
| MEGF6 | protein_coding | 0.20036696 | 0.72797319 |
| USP34 | protein_coding | 0.20022642 | 6.55014248 |
| UBB | protein_coding | 0.20018015 | 7.3410774 |
| ZBTB40 | protein_coding | 0.20013302 | 4.6005709 |

| | | | |
|---|---|---|---|
| PLEKHA1 | protein_coding | 0.20004382 | 3.0483023 |
| MKNK2 | protein_coding | 0.20002245 | 5.02319322 |
| RP11-338N1( | lncRNA | 0.19999131 | 0.33603757 |
| CA5B | protein_coding | 0.19978438 | 1.11574551 |
| BMF | protein_coding | 0.19976988 | 3.48274288 |
| DXO | protein_coding | 0.19959404 | 2.93056955 |
| SLC44A1 | protein_coding | 0.19955774 | 5.11871747 |
| AKIRIN2 | protein_coding | 0.19950556 | 3.38799958 |
| TOR3A | protein_coding | 0.19947035 | 2.88468592 |
| FOXO3B | protein_coding | 0.19938617 | 1.24277653 |
| FAIM | protein_coding | 0.19936517 | 1.97070501 |
| PLXNB3 | protein_coding | 0.19931956 | 0.64855436 |
| ZDHHC7 | protein_coding | 0.19928072 | 4.97817255 |
| TRIM39 | protein_coding | 0.19928032 | 2.14117894 |
| VCL | protein_coding | 0.1991924 | 3.68165109 |
| KRTCAP2 | protein_coding | 0.19911366 | 1.1837032 |
| MTERF4 | protein_coding | 0.19906686 | 4.45672353 |
| ARMC9 | protein_coding | 0.19901083 | 0.91728241 |
| CAMK2G | protein_coding | 0.19894387 | 3.11264278 |
| PRKCD | protein_coding | 0.19886017 | 1.40369941 |
| COMMD6 | protein_coding | 0.19867715 | 2.38034178 |
| QRICH2 | protein_coding | 0.19862947 | 1.97278412 |
| PGLS-DT | lncRNA | 0.19853534 | 0.2247518 |
| GRM2 | protein_coding | 0.19845828 | 4.40121055 |
| NLRP14 | protein_coding | 0.1984391 | 0.63993197 |
| CTD-2349P21 | lncRNA | 0.19838875 | 0.34356641 |
| FAM71E1 | protein_coding | 0.19838544 | 1.10716613 |
| BSPRY | protein_coding | 0.19834472 | 3.94153993 |
| MRPL47 | protein_coding | 0.1981597 | 4.89765365 |
| SLC7A4 | protein_coding | 0.19809049 | 1.26257555 |
| CATSPER2 | protein_coding | 0.19799738 | 0.73088482 |
| MPPED2 | protein_coding | 0.19797626 | 0.83371465 |
| GAB3 | protein_coding | 0.19785665 | 0.34356641 |
| POLR1H | protein_coding | 0.19779631 | 2.14918731 |
| NAP1L1 | protein_coding | 0.1976581 | 6.16351277 |
| NOL6 | protein_coding | 0.19762147 | 5.42283115 |
| NCOR1 | protein_coding | 0.19744433 | 5.89940062 |
| ALAD | protein_coding | 0.19731316 | 3.92317431 |
| NRDC | protein_coding | 0.19721409 | 7.33185428 |
| PDE12 | protein_coding | 0.19715606 | 3.05703684 |
| BRD2 | protein_coding | 0.19713235 | 8.16573531 |
| MRPS10 | protein_coding | 0.19694453 | 4.27709633 |
| LINC00461 | lncRNA | 0.1969429 | 2.83850533 |
| MTF1 | protein_coding | 0.19690464 | 3.12691919 |
| GAS5 | lncRNA | 0.19687684 | 5.43650175 |

| | | | |
|---|---|---|---|
| GOLM1 | protein_coding | 0.19673593 | 4.0660606 |
| AC009404.2 | lncRNA | 0.19671424 | 0.88429143 |
| HPS4 | protein_coding | 0.19664323 | 5.02769251 |
| SMG7-AS1 | lncRNA | 0.19663985 | 0.17991278 |
| ATP6V0E2-A! | lncRNA | 0.19656555 | 1.25620431 |
| RP11-817O1: | lncRNA | 0.19652682 | 0.19643843 |
| CTC-459F4.3 | lncRNA | 0.19632802 | 1.67566028 |
| GRIP1 | protein_coding | 0.19632329 | 2.34614335 |
| DHRS11 | protein_coding | 0.19625283 | 1.6606225 |
| ZBED1 | protein_coding | 0.19619506 | 2.65695103 |
| WDR11 | protein_coding | 0.1960781 | 3.90158586 |
| FSD1L | protein_coding | 0.19594277 | 2.01579977 |
| BRMS1L | protein_coding | 0.1959189 | 2.59212206 |
| OSER1 | protein_coding | 0.19559361 | 3.21745756 |
| TTC37 | protein_coding | 0.19543874 | 6.25034352 |
| CLCN6 | protein_coding | 0.19537337 | 2.28274407 |
| CHRNB1 | protein_coding | 0.19522382 | 0.85784556 |
| GEN1 | protein_coding | 0.19510891 | 3.55213035 |
| CEP95 | protein_coding | 0.19507185 | 3.78823235 |
| USP34-DT | lncRNA | 0.19505717 | 0.37647012 |
| ARFGEF2 | protein_coding | 0.19489773 | 4.63350447 |
| C1orf131 | protein_coding | 0.19474404 | 2.52891446 |
| METTL15 | protein_coding | 0.19466699 | 2.26094027 |
| UBA6 | protein_coding | 0.1944602 | 4.97622502 |
| DDX21 | protein_coding | 0.19445582 | 6.44619214 |
| BTBD17 | protein_coding | 0.19444036 | 1.21754574 |
| ACTR10 | protein_coding | 0.19439363 | 5.17919834 |
| ZNF585A | protein_coding | 0.19432447 | 1.96658437 |
| MAK16 | protein_coding | 0.19430697 | 3.03164916 |
| NARS1 | protein_coding | 0.19411902 | 7.18573647 |
| PDIA3 | protein_coding | 0.19411644 | 7.04643019 |
| COCH | protein_coding | 0.19400451 | 5.99340954 |
| MGAT4A | protein_coding | 0.19394749 | 4.84595727 |
| RP11-651L5.: | lncRNA | 0.1938882 | 0.14026339 |
| EMG1 | protein_coding | 0.19385376 | 4.34113107 |
| NHLRC3 | protein_coding | 0.19381632 | 1.31074851 |
| ADGRG1 | protein_coding | 0.19375494 | 4.02939334 |
| DUXAP8 | lncRNA | 0.1936957 | 2.0127666 |
| ZNF112 | protein_coding | 0.19365294 | 2.46447892 |
| P2RX5 | protein_coding | 0.19347789 | 0.33245182 |
| CCDC82 | protein_coding | 0.19345494 | 2.77056768 |
| UQCRFS1-DT | lncRNA | 0.19345272 | 0.16196677 |
| RP11-816J6.: | lncRNA | 0.19336993 | 0.29765942 |
| HEATR3 | protein_coding | 0.1933657 | 4.46085204 |
| SFRP1 | protein_coding | 0.19335017 | 0.60281664 |

| | | | |
|---|---|---|---|
| BABAM1 | protein_coding | 0.19332627 | 2.11476856 |
| PTGES3L | protein_coding | 0.19323605 | 0.4849453 |
| ERCC8 | protein_coding | 0.19321256 | 2.21784998 |
| FSCN1 | protein_coding | 0.19314151 | 2.93066641 |
| JOSD1 | protein_coding | 0.19284929 | 5.08902511 |
| FLYWCH1 | protein_coding | 0.19242304 | 1.9906441 |
| NUP42 | protein_coding | 0.1923961 | 2.59248503 |
| DCAF5 | protein_coding | 0.1923814 | 5.18996257 |
| AEN | protein_coding | 0.19235273 | 5.18038036 |
| COQ2 | protein_coding | 0.19223685 | 2.66313153 |
| RNF166 | protein_coding | 0.19208599 | 2.65924601 |
| LYRM4-AS1 | lncRNA | 0.19197931 | 1.02321888 |
| CCDC144NL-/ | lncRNA | 0.19196817 | 1.88580712 |
| DYRK2 | protein_coding | 0.19185721 | 4.47750639 |
| RSL1D1 | protein_coding | 0.19182134 | 7.12087788 |
| MRPL58 | protein_coding | 0.19181251 | 2.63545679 |
| RP1-80N2.3 | lncRNA | 0.19175398 | 1.79086908 |
| RP11-290F2⁄ | lncRNA | 0.19172258 | 0.34837808 |
| PLPP4 | protein_coding | 0.19165246 | 0.23855359 |
| ABHD3 | protein_coding | 0.19130522 | 4.67743734 |
| PRIMPOL | protein_coding | 0.19123522 | 2.22992551 |
| SIK3 | protein_coding | 0.19103307 | 3.70588344 |
| ZNF37A | protein_coding | 0.1910019 | 3.62742211 |
| MDC1 | protein_coding | 0.19095841 | 5.99643246 |
| C3orf33 | protein_coding | 0.19094735 | 0.29405534 |
| RREB1 | protein_coding | 0.1909327 | 3.71987523 |
| ZNF224 | protein_coding | 0.19080343 | 3.24654216 |
| SNN | protein_coding | 0.19066736 | 5.93014677 |
| CTC-559E9.5 | lncRNA | 0.19038364 | 0.43998357 |
| RTCA-AS1 | lncRNA | 0.19036004 | 1.14210757 |
| REV3L | protein_coding | 0.19031227 | 4.83655564 |
| RP9 | protein_coding | 0.19013942 | 2.19432221 |
| ZNF302 | protein_coding | 0.19010828 | 2.98748778 |
| TANC1 | protein_coding | 0.19010329 | 3.76994714 |
| LIN28B | protein_coding | 0.19009035 | 1.97004502 |
| SDHD | protein_coding | 0.19008619 | 2.66605392 |
| ILRUN | protein_coding | 0.1898072 | 6.97536307 |
| YIF1A | protein_coding | 0.18947458 | 4.09656975 |
| NRBF2 | protein_coding | 0.18946167 | 1.22381717 |
| TMEM39B | protein_coding | 0.18945709 | 2.87332939 |
| ID2 | protein_coding | 0.18941311 | 1.15216103 |
| SNX13 | protein_coding | 0.18925646 | 5.43829788 |
| GINS3 | protein_coding | 0.18925182 | 3.40435976 |
| SLC2A13 | protein_coding | 0.18886872 | 1.69113216 |
| ATP5PO | protein_coding | 0.1888079 | 3.57492363 |

| | | | |
|---|---|---|---|
| MOSPD2 | protein_coding | 0.18854176 | 2.60925932 |
| MAP3K1 | protein_coding | 0.18849699 | 2.2712502 |
| MPHOSPH10 | protein_coding | 0.18847991 | 4.24053083 |
| PRR3 | protein_coding | 0.18839051 | 3.79212049 |
| PKN2 | protein_coding | 0.1883277 | 4.25268461 |
| PSMD12 | protein_coding | 0.18822077 | 4.87137598 |
| AGAP9 | protein_coding | 0.18811138 | 0.65629789 |
| SARNP | protein_coding | 0.18809625 | 0.39566527 |
| NIPSNAP2 | protein_coding | 0.18799563 | 4.0867721 |
| WIPF1 | protein_coding | 0.18791179 | 0.79134443 |
| LINC00327 | lncRNA | 0.18786193 | 0.28468971 |
| RP11-127I20 | lncRNA | 0.18774652 | 0.16769853 |
| ANKRD17 | protein_coding | 0.18771652 | 6.68561283 |
| DDX42 | protein_coding | 0.18769428 | 6.62802538 |
| TFEB | protein_coding | 0.18748206 | 2.05785344 |
| ICA1L | protein_coding | 0.18732104 | 0.83484476 |
| COMMD2 | protein_coding | 0.18728495 | 2.62235895 |
| APBA2 | protein_coding | 0.18713354 | 2.49565931 |
| MAP1S | protein_coding | 0.18708982 | 2.09451799 |
| IPO13 | protein_coding | 0.18698868 | 5.16367477 |
| CRAMP1 | protein_coding | 0.186982 | 3.37927387 |
| TUBA8 | protein_coding | 0.18695724 | 0.26031782 |
| ZNF107 | protein_coding | 0.18692598 | 3.55050444 |
| DENND4A | protein_coding | 0.18673289 | 4.29769057 |
| ACOT1 | protein_coding | 0.18672477 | 0.61363885 |
| TMCC3 | protein_coding | 0.18667786 | 2.02288586 |
| COA1 | protein_coding | 0.18664749 | 4.60337275 |
| SH3D21 | protein_coding | 0.18664626 | 1.64408946 |
| ULK3 | protein_coding | 0.18657008 | 5.03261023 |
| CRIP2 | protein_coding | 0.18654938 | 1.21867407 |
| CTIF | protein_coding | 0.18651671 | 4.65938476 |
| RBFADN | lncRNA | 0.1864764 | 0.22546931 |
| POLDIP2 | protein_coding | 0.18646766 | 5.69664213 |
| ZBTB2 | protein_coding | 0.18634594 | 2.10394642 |
| PBLD | protein_coding | 0.18630763 | 0.95897051 |
| MED23 | protein_coding | 0.18622396 | 3.12208193 |
| ELAC1 | protein_coding | 0.18621583 | 1.04750905 |
| USP4 | protein_coding | 0.18593275 | 3.18239293 |
| EIF5 | protein_coding | 0.18591047 | 7.85405815 |
| CRLF3 | protein_coding | 0.18579821 | 3.46496225 |
| ZNF513 | protein_coding | 0.18576703 | 2.16550797 |
| TSTD2 | protein_coding | 0.1855749 | 4.54366194 |
| PRKAB2 | protein_coding | 0.18554514 | 4.07184428 |
| CDC42BPA | protein_coding | 0.18548894 | 4.96250426 |
| ZNHIT6 | protein_coding | 0.18538973 | 2.91164131 |

| | | | |
|---|---|---|---|
| TRADD | protein_coding | 0.18535333 | 0.62949316 |
| SEC23B | protein_coding | 0.18534697 | 5.65959694 |
| MAD2L1BP | protein_coding | 0.18529403 | 3.6300467 |
| CC2D1A | protein_coding | 0.18523369 | 4.23682092 |
| ACO2 | protein_coding | 0.18521625 | 6.58023089 |
| MRPS33 | protein_coding | 0.18490099 | 3.98084718 |
| RP11-417L19 | lncRNA | 0.18487612 | 0.28458356 |
| VPS41 | protein_coding | 0.18480643 | 3.95503542 |
| ACACA | protein_coding | 0.18471281 | 5.67994524 |
| ZNF470-DT | lncRNA | 0.18468031 | 0.24670731 |
| ALDH3A2 | protein_coding | 0.18467027 | 4.97811116 |
| DLG2 | protein_coding | 0.18454328 | 0.88798464 |
| SLC30A6 | protein_coding | 0.1845217 | 3.73895444 |
| PRRC2A | protein_coding | 0.18448443 | 7.2007975 |
| SYS1 | protein_coding | 0.1843551 | 2.37876625 |
| C18orf54 | protein_coding | 0.18434644 | 2.00133187 |
| ZNF569 | protein_coding | 0.18428874 | 2.81188456 |
| TNPO3 | protein_coding | 0.18421013 | 5.4984099 |
| ZNF697 | protein_coding | 0.18416613 | 1.7322769 |
| SGO1-AS1 | lncRNA | 0.18397969 | 0.28876537 |
| METAP1D | protein_coding | 0.18383817 | 1.48336178 |
| STX3 | protein_coding | 0.18371219 | 3.49391316 |
| TMEM260 | protein_coding | 0.18361437 | 2.60579714 |
| GFOD1 | protein_coding | 0.18350907 | 1.70555541 |
| PPFIA4 | protein_coding | 0.18342843 | 2.5934758 |
| ZNF90 | protein_coding | 0.1833981 | 0.90547498 |
| TBC1D13 | protein_coding | 0.18337274 | 3.30616667 |
| CTD-2649C14 | lncRNA | 0.1833024 | 0.25667858 |
| RP11-381K20 | lncRNA | 0.18325239 | 0.81549294 |
| NUP205 | protein_coding | 0.18320528 | 6.54384224 |
| CDKN2B-AS1 | lncRNA | 0.18308396 | 2.22869022 |
| NAE1 | protein_coding | 0.18303015 | 4.55899994 |
| RP11-367N14 | lncRNA | 0.18295289 | 0.34580548 |
| CPD | protein_coding | 0.18269185 | 5.42869714 |
| HMGXB3 | protein_coding | 0.18255028 | 5.4638344 |
| PSMB5 | protein_coding | 0.18237939 | 6.33272024 |
| METTL14 | protein_coding | 0.18231378 | 3.18872023 |
| PSMB7 | protein_coding | 0.18223278 | 6.11139905 |
| RP11-255O19 | lncRNA | 0.18216343 | 0.1926461 |
| DCC | protein_coding | 0.18215582 | 1.19518727 |
| RAB38 | protein_coding | 0.18201209 | 0.20372878 |
| ATMIN | protein_coding | 0.18200475 | 5.98998552 |
| TBC1D2B | protein_coding | 0.18182579 | 3.09700679 |
| TMEM87B | protein_coding | 0.18177961 | 3.45873971 |
| FAM237A | protein_coding | 0.18174476 | 0.21856287 |

| | | | |
|---|---|---|---|
| ZNF700 | protein_coding | 0.18168452 | 2.32496609 |
| GRN | protein_coding | 0.18158494 | 4.32930312 |
| POGZ | protein_coding | 0.18157513 | 5.54559162 |
| MRPS18C | protein_coding | 0.18156503 | 3.47608723 |
| LSM6 | protein_coding | 0.18141471 | 2.1153048 |
| CLPTM1L | protein_coding | 0.18122676 | 6.44636957 |
| PHC2 | protein_coding | 0.18100895 | 6.87439122 |
| MTHFS | protein_coding | 0.18087142 | 0.28934054 |
| STARD4 | protein_coding | 0.18078477 | 2.66903562 |
| TMEM161B-/ | lncRNA | 0.18077558 | 2.26759363 |
| PAXIP1-DT | lncRNA | 0.18077266 | 1.65984092 |
| HOXD8 | protein_coding | 0.18070234 | 2.93664087 |
| PIK3CB | protein_coding | 0.18069104 | 3.89371497 |
| ARHGAP8 | protein_coding | 0.18045589 | 0.74435474 |
| NXT2 | protein_coding | 0.18035555 | 1.01486176 |
| VMAC | protein_coding | 0.18033879 | 0.49897744 |
| NDUFB8 | protein_coding | 0.18029024 | 0.84632984 |
| AMACR | protein_coding | 0.18004577 | 0.53134038 |
| HSF2BP | protein_coding | 0.18003539 | 0.37571193 |
| OXSR1 | protein_coding | 0.18002449 | 3.30019389 |
| ERP44 | protein_coding | 0.1798942 | 4.04692284 |
| NEURL2 | protein_coding | 0.17980714 | 0.27073133 |
| PIKFYVE | protein_coding | 0.17975772 | 3.44346813 |
| LACTB | protein_coding | 0.17967659 | 2.30278941 |
| C5 | protein_coding | 0.17962238 | 1.53539827 |
| C5AR1 | protein_coding | 0.17952276 | 0.22658577 |
| ANKAR | protein_coding | 0.17951917 | 0.97603067 |
| CMC1 | protein_coding | 0.17951661 | 1.43361131 |
| CCDC130 | protein_coding | 0.17950732 | 3.3962601 |
| MINDY1 | protein_coding | 0.17950566 | 2.44248024 |
| AGAP1 | protein_coding | 0.17946373 | 4.25716795 |
| EXOC2 | protein_coding | 0.17913748 | 3.45363421 |
| CAST | protein_coding | 0.17894943 | 2.3374849 |
| ZNF503-AS2 | lncRNA | 0.17885796 | 1.45964545 |
| EAF1 | protein_coding | 0.17859919 | 4.20505283 |
| IQCB1 | protein_coding | 0.17851028 | 2.85572307 |
| BLOC1S3 | protein_coding | 0.17849219 | 1.17763139 |
| RAD23A | protein_coding | 0.17840917 | 6.22766567 |
| SLC25A44 | protein_coding | 0.1783908 | 4.60247933 |
| CTD-2227E11 | lncRNA | 0.17836046 | 0.36320451 |
| SCNM1 | protein_coding | 0.17821825 | 2.79666433 |
| SNHG6 | lncRNA | 0.17821456 | 2.71433482 |
| CHEK2 | protein_coding | 0.17815483 | 2.49371488 |
| DUSP6 | protein_coding | 0.17811933 | 0.19614889 |
| PAWR | protein_coding | 0.177972 | 3.47162594 |

| | | | |
|---|---|---|---|
| CUL7 | protein_coding | 0.177956 | 1.99576208 |
| PRUNE2 | protein_coding | 0.17781391 | 6.21313947 |
| PTPRF | protein_coding | 0.17769629 | 5.04166334 |
| ZNF93 | protein_coding | 0.17763671 | 1.95210898 |
| SDR42E1 | protein_coding | 0.17763003 | 3.47944008 |
| RP11-305E6. | lncRNA | 0.17755636 | 0.41335035 |
| C12orf29 | protein_coding | 0.17716215 | 2.6453527 |
| NHSL1 | protein_coding | 0.17705054 | 1.5229841 |
| POLR1A | protein_coding | 0.17695585 | 4.98730446 |
| CCDC97 | protein_coding | 0.17688836 | 3.42112355 |
| TTN-AS1 | lncRNA | 0.17686789 | 1.25273335 |
| PXK | protein_coding | 0.17677445 | 1.41535104 |
| MUC20 | protein_coding | 0.17674465 | 0.37633662 |
| NBPF1 | protein_coding | 0.17650319 | 3.61465035 |
| GPAT3 | protein_coding | 0.17649473 | 1.47056823 |
| WDR45B | protein_coding | 0.17639888 | 5.64602669 |
| DNTTIP1 | protein_coding | 0.17607503 | 2.90226146 |
| NCKAP1 | protein_coding | 0.17585887 | 4.61681024 |
| KIF15 | protein_coding | 0.1758119 | 1.96249848 |
| C5orf51 | protein_coding | 0.17575711 | 3.20400982 |
| WDR62 | protein_coding | 0.17571225 | 4.65410988 |
| METAP2 | protein_coding | 0.17567655 | 5.26797615 |
| MICB | protein_coding | 0.17566287 | 1.27136799 |
| SLC45A4 | protein_coding | 0.17563225 | 2.06613674 |
| BCL9 | protein_coding | 0.17561236 | 1.8873069 |
| C14orf93 | protein_coding | 0.17560678 | 2.19695514 |
| LRRC1 | protein_coding | 0.17550959 | 1.76933406 |
| B9D1 | protein_coding | 0.17548481 | 1.06304057 |
| MRPL1 | protein_coding | 0.17542493 | 2.67361881 |
| FAM104A | protein_coding | 0.1751089 | 3.82616232 |
| BIRC6 | protein_coding | 0.17506829 | 6.41961757 |
| BDP1 | protein_coding | 0.17505952 | 5.38813564 |
| OSGEPL1-AS | lncRNA | 0.17501878 | 0.28977194 |
| FRMPD4 | protein_coding | 0.175009 | 1.27252109 |
| TUBB2B | protein_coding | 0.17498397 | 4.66034348 |
| RBM7 | protein_coding | 0.17497585 | 1.12688848 |
| STON2 | protein_coding | 0.17495827 | 2.25578419 |
| SIDT2 | protein_coding | 0.174895 | 2.11481678 |
| ESYT2 | protein_coding | 0.17483719 | 5.71064084 |
| METTL1 | protein_coding | 0.17477874 | 1.45559895 |
| MRPL40 | protein_coding | 0.17450206 | 2.78241837 |
| TBC1D22B | protein_coding | 0.1744232 | 2.74971688 |
| MEST | protein_coding | 0.17422693 | 5.61653317 |
| C10orf88 | protein_coding | 0.17420177 | 2.01238484 |
| RPP38 | protein_coding | 0.1740969 | 1.56440707 |

| | | | |
|---|---|---|---|
| PRKCSH | protein_coding | 0.17405915 | 6.47328247 |
| CBWD3 | protein_coding | 0.17398301 | 0.38861362 |
| BMS1 | protein_coding | 0.17386689 | 3.66291016 |
| ERCC4 | protein_coding | 0.17383726 | 3.06994367 |
| HPF1 | protein_coding | 0.1737442 | 2.70715003 |
| BRD9 | protein_coding | 0.17360477 | 3.68197236 |
| TXNL1 | protein_coding | 0.17344621 | 4.58360568 |
| PLGLB1 | protein_coding | 0.17336618 | 0.22806683 |
| MARCHF7 | protein_coding | 0.17335823 | 4.52247861 |
| ATXN1L | protein_coding | 0.17333723 | 3.90840208 |
| CTC-453G23. | lncRNA | 0.17333219 | 0.24170951 |
| PPP4R2 | protein_coding | 0.17316481 | 2.77193785 |
| KBTBD2 | protein_coding | 0.17309311 | 4.84941478 |
| UNC13A | protein_coding | 0.17300707 | 6.37998674 |
| TOM1 | protein_coding | 0.17300303 | 2.43346933 |
| CEP164 | protein_coding | 0.17299443 | 3.08779214 |
| CLTC | protein_coding | 0.17298035 | 7.01163401 |
| LPGAT1 | protein_coding | 0.17296159 | 5.96919264 |
| NFIC | protein_coding | 0.1728301 | 3.42510053 |
| C9orf64 | protein_coding | 0.17258521 | 3.14821394 |
| AMMECR1L | protein_coding | 0.17226606 | 4.13885506 |
| NIP7 | protein_coding | 0.17222946 | 3.13885278 |
| LINC01341 | lncRNA | 0.17219658 | 0.4324857 |
| TXNRD2 | protein_coding | 0.17219144 | 2.56511464 |
| VPS54 | protein_coding | 0.17218362 | 3.35379211 |
| SH3BP1 | protein_coding | 0.17216829 | 1.16508964 |
| RTEL1-TNFRS | protein_coding | 0.17211265 | 0.5028314 |
| DCAF4L1 | protein_coding | 0.17182278 | 0.26001587 |
| ATRX | protein_coding | 0.17142538 | 5.26376619 |
| SFXN5 | protein_coding | 0.17141954 | 2.27976983 |
| IFT20 | protein_coding | 0.1713941 | 1.51915058 |
| SMG6 | protein_coding | 0.17134791 | 2.79032797 |
| PRCP | protein_coding | 0.17133782 | 2.89997859 |
| ARFIP1 | protein_coding | 0.17123991 | 1.76213702 |
| DOCK1 | protein_coding | 0.17111863 | 1.61936489 |
| ZHX2 | protein_coding | 0.17101186 | 1.39480667 |
| GAA | protein_coding | 0.17069632 | 4.18136309 |
| MTHFD2 | protein_coding | 0.17050538 | 6.54056299 |
| TCTN3 | protein_coding | 0.17050266 | 1.52686755 |
| PKD2 | protein_coding | 0.17032186 | 2.13271015 |
| TTBK2 | protein_coding | 0.1702574 | 4.07378066 |
| PWWP2A | protein_coding | 0.17019117 | 2.91917619 |
| LRCH3 | protein_coding | 0.1701028 | 4.14870703 |
| KCNH3 | protein_coding | 0.16999 | 0.90769151 |
| RP11-34F13. | lncRNA | 0.16998424 | 0.26793954 |

| | | | |
|---|---|---|---|
| GLA | protein_coding | 0.16992867 | 1.73729468 |
| PFN4 | protein_coding | 0.16967475 | 0.19061835 |
| AKAP8L | protein_coding | 0.1696098 | 3.36804335 |
| WDR75 | protein_coding | 0.16959202 | 4.40547388 |
| NCAPG2 | protein_coding | 0.16958592 | 6.53742928 |
| FMC1 | protein_coding | 0.16951612 | 0.25418677 |
| CPB2-AS1 | lncRNA | 0.16933296 | 0.39379335 |
| RWDD2A | protein_coding | 0.16923105 | 1.1661973 |
| TIAF1 | protein_coding | 0.16922168 | 0.44384521 |
| CALCB | protein_coding | 0.16921015 | 0.72800413 |
| AKAP6 | protein_coding | 0.16915766 | 1.75989906 |
| FAM27E3 | lncRNA | 0.16914087 | 0.34927195 |
| IRF6 | protein_coding | 0.16900221 | 1.52784879 |
| CCDC7 | protein_coding | 0.16898573 | 0.39887293 |
| STMN2 | protein_coding | 0.16896365 | 4.72719564 |
| RAD54L2 | protein_coding | 0.16895207 | 2.79053335 |
| PHF14 | protein_coding | 0.16889597 | 5.57964123 |
| SYNJ2 | protein_coding | 0.16889029 | 1.0773169 |
| ZDHHC23 | protein_coding | 0.16885802 | 1.71604133 |
| ABCA2 | protein_coding | 0.16884726 | 4.01395816 |
| TRIM46 | protein_coding | 0.16882009 | 2.20683752 |
| RP11-977G19 | lncRNA | 0.16877886 | 0.20355634 |
| NT5C3A | protein_coding | 0.16873045 | 3.71967101 |
| HSPA1B | protein_coding | 0.1687239 | 1.55826731 |
| HECTD1 | protein_coding | 0.16866944 | 6.15521281 |
| ASNSD1 | protein_coding | 0.16866241 | 2.59309261 |
| CHMP4A | protein_coding | 0.16863187 | 0.37174943 |
| DDN-AS1 | lncRNA | 0.16857211 | 1.25805324 |
| PPID | protein_coding | 0.16850072 | 3.38669353 |
| IVNS1ABP | protein_coding | 0.16847995 | 5.97486776 |
| MRPL9 | protein_coding | 0.16845678 | 3.85433421 |
| ZBTB48 | protein_coding | 0.16843016 | 1.50498588 |
| B3GALT9 | protein_coding | 0.16842908 | 1.16081087 |
| ZKSCAN4 | protein_coding | 0.16838078 | 0.69787489 |
| DSE | protein_coding | 0.16836864 | 0.48068161 |
| RP11-332M2 | lncRNA | 0.16818226 | 0.75495187 |
| PCDHGA6 | protein_coding | 0.16816265 | 0.62775629 |
| RPL18A | protein_coding | 0.16812944 | 6.19303853 |
| ACOT2 | protein_coding | 0.16809679 | 1.01461248 |
| CREBRF | protein_coding | 0.16803998 | 0.82240692 |
| FSTL3 | protein_coding | 0.16792952 | 1.14446911 |
| SLC2A11 | protein_coding | 0.16790657 | 1.17201444 |
| AUTS2 | protein_coding | 0.16777086 | 3.27954763 |
| NUFIP2 | protein_coding | 0.16762793 | 4.55154962 |
| ZZZ3 | protein_coding | 0.16761917 | 4.41218729 |

| | | | |
|---|---|---|---|
| GNL3 | protein_coding | 0.1675873 | 4.02927117 |
| PIK3CA | protein_coding | 0.16754527 | 4.14182237 |
| TRMT5 | protein_coding | 0.1674979 | 2.88956855 |
| FOSL2 | protein_coding | 0.16748508 | 2.97016115 |
| CTD-3222D1¦ | lncRNA | 0.1674466 | 0.52572838 |
| TRMT44 | protein_coding | 0.16741921 | 1.82180439 |
| IDH3A | protein_coding | 0.16740404 | 3.80046302 |
| EPHB2 | protein_coding | 0.16701745 | 1.2590068 |
| ZBTB26 | protein_coding | 0.16698277 | 2.6371619 |
| AGL | protein_coding | 0.16696858 | 3.97747931 |
| VAMP4 | protein_coding | 0.16695355 | 1.54108263 |
| NBEAL1 | protein_coding | 0.16684294 | 1.81449304 |
| RP11-80H5.2 | lncRNA | 0.1668231 | 0.3524804 |
| TMEM86B | protein_coding | 0.16680737 | 0.43887181 |
| RP11-341G2 | lncRNA | 0.16680235 | 0.5338855 |
| FAM98A | protein_coding | 0.16678185 | 3.58102277 |
| EPC1 | protein_coding | 0.16676144 | 2.92169545 |
| CHCHD3 | protein_coding | 0.16668697 | 5.12035303 |
| NPM3 | protein_coding | 0.16656964 | 2.23018501 |
| HPS5 | protein_coding | 0.16647908 | 2.65720828 |
| OPA3 | protein_coding | 0.16647293 | 3.82576129 |
| LSM1 | protein_coding | 0.16644462 | 2.08402054 |
| USP20 | protein_coding | 0.1664367 | 3.19103407 |
| IER2 | protein_coding | 0.16629265 | 1.8024102 |
| DNAJB2 | protein_coding | 0.16616055 | 4.97700299 |
| LIPT1 | protein_coding | 0.16608928 | 0.87818424 |
| ZNF174 | protein_coding | 0.16607249 | 1.54202723 |
| ARHGAP4 | protein_coding | 0.16598241 | 2.27493895 |
| UTY | protein_coding | 0.16582985 | 2.20526852 |
| ODR4 | protein_coding | 0.16577219 | 2.49445768 |
| FAM156A | protein_coding | 0.16569307 | 0.19020207 |
| BRD4 | protein_coding | 0.1656304 | 4.72289225 |
| NOXA1 | protein_coding | 0.16556551 | 1.23669599 |
| C14orf132 | protein_coding | 0.16547706 | 5.45841695 |
| MTHFSD | protein_coding | 0.16546963 | 2.88161279 |
| TRMT12 | protein_coding | 0.16546052 | 2.01413356 |
| AIMP1 | protein_coding | 0.16544679 | 3.77287934 |
| UQCRC2 | protein_coding | 0.16542182 | 6.11157962 |
| IGHMBP2 | protein_coding | 0.16538611 | 1.91151723 |
| DNAJB6 | protein_coding | 0.16535018 | 4.48808348 |
| CCDC92 | protein_coding | 0.1653032 | 3.46704182 |
| ADNP-AS1 | lncRNA | 0.16530272 | 0.27445793 |
| LENG8-AS1 | lncRNA | 0.16517725 | 0.60056472 |
| H2AC13 | protein_coding | 0.16506589 | 0.21033945 |
| CTC-479C5.1 | lncRNA | 0.16495526 | 0.21130362 |

| | | | |
|---|---|---|---|
| RP11-44F14. | lncRNA | 0.16491888 | 0.26602239 |
| UBE3A | protein_coding | 0.16480643 | 5.38745038 |
| GPR137C | protein_coding | 0.16477965 | 1.45678218 |
| ZNF678 | protein_coding | 0.16451427 | 2.29865268 |
| HERC2 | protein_coding | 0.16449041 | 6.53363971 |
| TEFM | protein_coding | 0.16448002 | 1.46055586 |
| SLC66A1 | protein_coding | 0.16441833 | 1.93960457 |
| USP9Y | protein_coding | 0.16437514 | 4.14922892 |
| LIAS | protein_coding | 0.16436765 | 1.6696686 |
| ANP32E | protein_coding | 0.16421502 | 2.84854965 |
| STAT5B | protein_coding | 0.16402239 | 3.50066972 |
| NDUFAF1 | protein_coding | 0.16399158 | 2.08333329 |
| NUMBL | protein_coding | 0.163868 | 2.84041632 |
| ARID4A | protein_coding | 0.16380538 | 2.20591074 |
| TMEM238 | protein_coding | 0.16377821 | 0.31398024 |
| C16orf72 | protein_coding | 0.16364402 | 3.20095475 |
| MRM2 | protein_coding | 0.16345978 | 0.15662462 |
| B4GALT4 | protein_coding | 0.1634199 | 1.29040035 |
| LANCL2 | protein_coding | 0.16324482 | 3.51145376 |
| CASD1 | protein_coding | 0.16321287 | 1.91734329 |
| UBP1 | protein_coding | 0.16309144 | 3.44903944 |
| YJEFN3 | protein_coding | 0.16303276 | 0.77236202 |
| TOR1A | protein_coding | 0.1629905 | 3.59182956 |
| BLZF1 | protein_coding | 0.16297532 | 2.555158 |
| TCF12 | protein_coding | 0.16292784 | 5.33121789 |
| GNAO1 | protein_coding | 0.16287977 | 5.72056574 |
| NCOA6 | protein_coding | 0.16279594 | 4.92761212 |
| CWF19L2 | protein_coding | 0.16278508 | 1.41042393 |
| EZH1 | protein_coding | 0.16272175 | 3.10464801 |
| ALG9 | protein_coding | 0.16268069 | 0.96808817 |
| RADX | protein_coding | 0.16260835 | 1.29596965 |
| GOSR1 | protein_coding | 0.16256073 | 3.97550964 |
| AC091133.1 | lncRNA | 0.16243793 | 0.35724598 |
| PLEKHG4 | protein_coding | 0.16241494 | 4.38725057 |
| FAM149A | protein_coding | 0.16232047 | 2.731765 |
| STC2 | protein_coding | 0.1622472 | 4.36602616 |
| SHOC2 | protein_coding | 0.16221537 | 2.62906572 |
| ATP6V0A2 | protein_coding | 0.16218237 | 3.44263426 |
| ZNF507 | protein_coding | 0.16207096 | 3.29298808 |
| SLC35B3 | protein_coding | 0.16207027 | 1.5880784 |
| PRMT3 | protein_coding | 0.16199395 | 2.8678512 |
| TIGD7 | protein_coding | 0.16172943 | 1.91995426 |
| TMEM196 | protein_coding | 0.16172207 | 1.947448 |
| FAN1 | protein_coding | 0.16161443 | 3.99351118 |
| ERVK13-1 | lncRNA | 0.16154718 | 1.37381021 |

| | | | |
|---|---|---|---|
| ACKR3 | protein_coding | 0.16153628 | 0.54725601 |
| DNAJC7 | protein_coding | 0.16151421 | 4.66847801 |
| DGKI | protein_coding | 0.16149781 | 0.65659188 |
| DPM1 | protein_coding | 0.16143415 | 2.52983812 |
| FBXO22 | protein_coding | 0.16136795 | 3.31355253 |
| STARD7-AS1 | lncRNA | 0.16132888 | 1.27086516 |
| TMEM108 | protein_coding | 0.16130849 | 3.00834503 |
| TFAP2A | protein_coding | 0.16124791 | 4.85959327 |
| FANCD2 | protein_coding | 0.16123997 | 2.45372869 |
| AC005562.1 | lncRNA | 0.16105098 | 0.20964872 |
| IFFO1 | protein_coding | 0.16102053 | 1.42388171 |
| GLMN | protein_coding | 0.16097274 | 2.22232499 |
| BCLAF3 | protein_coding | 0.1608948 | 2.03129526 |
| TOMM5 | protein_coding | 0.1608159 | 0.65071422 |
| LMNA | protein_coding | 0.16074508 | 4.49674461 |
| RP5-890O3.9 | lncRNA | 0.16073704 | 0.27862635 |
| SRFBP1 | protein_coding | 0.16072813 | 1.62807146 |
| ELAPOR2 | protein_coding | 0.16063016 | 3.67895835 |
| ZDHHC4 | protein_coding | 0.16061491 | 2.69372957 |
| ACBD7 | protein_coding | 0.16060693 | 1.42770958 |
| WHAMM | protein_coding | 0.16057485 | 2.55043212 |
| HMG20A | protein_coding | 0.16027657 | 3.73964326 |
| RASA4 | protein_coding | 0.16004222 | 0.71422446 |
| NIT1 | protein_coding | 0.15995135 | 2.46802 |
| ZNF3 | protein_coding | 0.15992412 | 4.25685267 |
| SNRPA1 | protein_coding | 0.15987013 | 3.5176363 |
| LINC00894 | lncRNA | 0.15959308 | 0.39201053 |
| ZNF768 | protein_coding | 0.15948101 | 3.06874057 |
| EPS8 | protein_coding | 0.15940073 | 2.49863531 |
| RP11-377D9. | lncRNA | 0.15939888 | 0.21008821 |
| FAM114A2 | protein_coding | 0.15938375 | 2.49662992 |
| IKZF4 | protein_coding | 0.15936975 | 1.19341028 |
| TRIM59 | protein_coding | 0.15935118 | 1.15911613 |
| ANKRD10-IT1 | lncRNA | 0.15933084 | 0.50374818 |
| MCM3AP | protein_coding | 0.15928986 | 4.49235313 |
| PBDC1 | protein_coding | 0.15923027 | 1.70625229 |
| WDR81 | protein_coding | 0.15919348 | 1.83662317 |
| RERE | protein_coding | 0.15909148 | 4.08330696 |
| UTP25 | protein_coding | 0.15901324 | 2.42934027 |
| FZD4 | protein_coding | 0.15877097 | 0.66196613 |
| WRN | protein_coding | 0.15876543 | 2.59617058 |
| PDP2 | protein_coding | 0.15855948 | 3.37156693 |
| RP11-428J1.4 | lncRNA | 0.15852843 | 0.26191127 |
| VASH1-AS1 | lncRNA | 0.15852775 | 1.6378606 |
| ATRNL1 | protein_coding | 0.15850174 | 2.44280965 |

| | | | |
|---|---|---|---|
| ZNF75D | protein_coding | 0.15846874 | 1.4348501 |
| RP11-1406H1 | lncRNA | 0.15845454 | 2.60726223 |
| RP11-649A18 | lncRNA | 0.15845215 | 0.18244943 |
| PABPN1 | protein_coding | 0.15830076 | 1.14057559 |
| PRPF40A | protein_coding | 0.15827824 | 5.60883193 |
| MT-CO2 | protein_coding | 0.15819356 | 5.68378608 |
| SMPDL3B | protein_coding | 0.15814242 | 0.94665621 |
| NRSN2-AS1 | lncRNA | 0.15813944 | 1.06968213 |
| NAT10 | protein_coding | 0.15810719 | 4.49749092 |
| PHGDH | protein_coding | 0.15792982 | 5.84657057 |
| SLC35A3 | protein_coding | 0.15768247 | 1.31948751 |
| C18orf25 | protein_coding | 0.15762739 | 4.07086388 |
| PPIL6 | protein_coding | 0.15754637 | 0.28770021 |
| NAA30 | protein_coding | 0.15752547 | 2.15958705 |
| PSMB2 | protein_coding | 0.15751542 | 5.94561891 |
| LMAN1 | protein_coding | 0.15750686 | 4.88120643 |
| UBE2D3 | protein_coding | 0.15745313 | 4.34196282 |
| PCM1 | protein_coding | 0.15741836 | 4.73131101 |
| C19orf12 | protein_coding | 0.15732334 | 2.13221243 |
| ARTN | protein_coding | 0.15729377 | 0.23627682 |
| PLPP6 | protein_coding | 0.15725114 | 2.06030677 |
| XX-FW83563 | lncRNA | 0.1572506 | 0.35614594 |
| ZNF225 | protein_coding | 0.15692847 | 1.37421321 |
| ADAT1 | protein_coding | 0.15690598 | 4.07100978 |
| CHP1 | protein_coding | 0.15684278 | 3.20598818 |
| CUL9 | protein_coding | 0.15682382 | 3.09973815 |
| CCNL2 | protein_coding | 0.15664971 | 3.96048523 |
| AP2B1 | protein_coding | 0.15664726 | 6.15514402 |
| IFT43 | protein_coding | 0.15647129 | 1.70663071 |
| ANKRD40 | protein_coding | 0.15646435 | 4.03355398 |
| VPS36 | protein_coding | 0.15643696 | 2.57717022 |
| GGA3 | protein_coding | 0.15637499 | 3.40047909 |
| ZNF121 | protein_coding | 0.15634105 | 3.3854054 |
| GLYR1 | protein_coding | 0.15623696 | 5.6498741 |
| USP48 | protein_coding | 0.15622872 | 5.00354548 |
| HTATIP2 | protein_coding | 0.1561157 | 4.36083865 |
| ACTR8 | protein_coding | 0.15577681 | 2.04110013 |
| NUP153 | protein_coding | 0.15576998 | 5.69590519 |
| C6orf3 | lncRNA | 0.15576015 | 0.37501546 |
| ANKRD28 | protein_coding | 0.15575519 | 2.98013273 |
| DPH1 | protein_coding | 0.15571251 | 1.55144299 |
| THAP6 | protein_coding | 0.15560495 | 2.09591018 |
| BRCA1 | protein_coding | 0.15531961 | 3.21287966 |
| KIF1B | protein_coding | 0.15517788 | 4.15615233 |
| SLC4A1AP | protein_coding | 0.15491191 | 3.03670112 |

| | | | |
|---|---|---|---|
| PRELID1 | protein_coding | 0.15476876 | 4.16572856 |
| DNAJA4 | protein_coding | 0.15476208 | 3.14565974 |
| KLF9 | protein_coding | 0.15474623 | 2.62697202 |
| VPS11 | protein_coding | 0.15459794 | 3.19529181 |
| AHI1 | protein_coding | 0.15454883 | 1.36181706 |
| SAMD4B | protein_coding | 0.15451239 | 5.02216439 |
| CYTH2 | protein_coding | 0.15432937 | 3.46793933 |
| APOLD1 | protein_coding | 0.15426332 | 1.58842493 |
| ANKS1A | protein_coding | 0.15423436 | 3.56222657 |
| USP8 | protein_coding | 0.15419895 | 3.6592335 |
| PIGL | protein_coding | 0.15418738 | 0.78107104 |
| GRIK5 | protein_coding | 0.15418207 | 1.95074459 |
| TSC1 | protein_coding | 0.15413453 | 2.58968881 |
| CLPX | protein_coding | 0.15408029 | 3.11169881 |
| WARS2-AS1 | lncRNA | 0.15396794 | 1.44510202 |
| GNE | protein_coding | 0.15395902 | 3.40143803 |
| RAB21 | protein_coding | 0.15390455 | 2.41672926 |
| CHORDC1 | protein_coding | 0.15382972 | 2.73667165 |
| DDX1 | protein_coding | 0.15381957 | 5.27633875 |
| MMADHC | protein_coding | 0.15376913 | 4.75640849 |
| SETD1B | protein_coding | 0.1537435 | 2.85308282 |
| RPUSD4 | protein_coding | 0.15364475 | 2.28964297 |
| CCDC51 | protein_coding | 0.15358995 | 1.96747897 |
| HOXD3 | protein_coding | 0.15349217 | 1.07969762 |
| EDC3 | protein_coding | 0.15344739 | 3.89623725 |
| ZFP3 | protein_coding | 0.15343587 | 1.74687904 |
| ZSCAN30 | protein_coding | 0.15337673 | 2.48593266 |
| RAB30-DT | lncRNA | 0.15334164 | 1.33287923 |
| RP11-378A1: | lncRNA | 0.15332576 | 0.13875752 |
| GET4 | protein_coding | 0.15331806 | 0.40649155 |
| LMO2 | protein_coding | 0.15330726 | 3.78917846 |
| CHST8 | protein_coding | 0.15319639 | 1.86832565 |
| PANX1 | protein_coding | 0.15315589 | 3.68340306 |
| RP11-54O7.1 | lncRNA | 0.15310932 | 0.49799107 |
| ME2 | protein_coding | 0.15306956 | 3.99190493 |
| SRRM2 | protein_coding | 0.15303264 | 7.08511403 |
| BUD31 | protein_coding | 0.1529664 | 4.44872752 |
| ZNF343 | protein_coding | 0.15295636 | 1.94493837 |
| TAF8 | protein_coding | 0.15281689 | 2.63008524 |
| PHEX | protein_coding | 0.15281317 | 0.23264116 |
| SLTM | protein_coding | 0.1527335 | 5.02796202 |
| OS9 | protein_coding | 0.15272349 | 5.22648048 |
| ZNF189 | protein_coding | 0.15253209 | 2.76839172 |
| EXOG | protein_coding | 0.15249304 | 1.16458528 |
| TXNDC15 | protein_coding | 0.15246969 | 2.58645093 |

| | | | |
|---|---|---|---|
| PNO1 | protein_coding | 0.15245253 | 3.0070031 |
| C11orf71 | protein_coding | 0.15244141 | 0.89820284 |
| FASTKD2 | protein_coding | 0.1523917 | 3.30712276 |
| C1orf226 | protein_coding | 0.15233516 | 1.7472678 |
| PDGFA | protein_coding | 0.15229145 | 0.62991031 |
| APC | protein_coding | 0.15225743 | 3.9365003 |
| ZNF713 | protein_coding | 0.15220316 | 0.76274616 |
| RP5-984P4.6 | lncRNA | 0.15214162 | 0.30695414 |
| POMP | protein_coding | 0.15203975 | 4.7351425 |
| ELMO1 | protein_coding | 0.15203424 | 0.44902206 |
| CBR1 | protein_coding | 0.15199897 | 2.13967886 |
| UBXN1 | protein_coding | 0.15197289 | 4.73912165 |
| THAP8 | protein_coding | 0.15190037 | 1.18560438 |
| SLC1A4 | protein_coding | 0.15188817 | 5.04738565 |
| WDR73 | protein_coding | 0.15185713 | 2.19758542 |
| C8orf76 | protein_coding | 0.15185131 | 0.36078313 |
| AREL1 | protein_coding | 0.15174885 | 4.81238621 |
| RP11-1070N | lncRNA | 0.15172801 | 0.2625945 |
| NINJ2-AS1 | lncRNA | 0.15163075 | 0.54149413 |
| TAB2 | protein_coding | 0.15152443 | 2.23286959 |
| GDAP1 | protein_coding | 0.15137791 | 3.19733277 |
| CCDC66 | protein_coding | 0.15132941 | 1.21017105 |
| SRRM5 | protein_coding | 0.15132873 | 0.24035355 |
| FRS2 | protein_coding | 0.15124414 | 2.26012121 |
| C1orf53 | protein_coding | 0.1512059 | 0.43953652 |
| SFXN4 | protein_coding | 0.15105128 | 2.76408552 |
| LINC01089 | lncRNA | 0.15101796 | 2.19160032 |
| PIBF1 | protein_coding | 0.15088027 | 2.26712192 |
| DPAGT1 | protein_coding | 0.1508409 | 2.06725175 |
| NPC1 | protein_coding | 0.15083625 | 2.8020057 |
| ZNF383 | protein_coding | 0.15080609 | 1.63734169 |
| EDEM2 | protein_coding | 0.15028894 | 1.31512601 |
| ZBTB25 | protein_coding | 0.15027913 | 2.09439217 |
| MTMR1 | protein_coding | 0.15022393 | 3.28914993 |
| MAP3K20-AS | lncRNA | 0.15021974 | 2.55675355 |
| RALGDS | protein_coding | 0.15010042 | 0.96300783 |
| NMNAT1 | protein_coding | 0.15007337 | 0.73124993 |
| CRTC2 | protein_coding | 0.15000733 | 2.58422448 |
| DHX33 | protein_coding | 0.14999832 | 2.34607855 |
| ANKS3 | protein_coding | 0.14999064 | 1.4507426 |
| PBK | protein_coding | 0.14997817 | 2.15244372 |
| TMEM17 | protein_coding | 0.1499684 | 0.6874137 |
| ZSCAN16 | protein_coding | 0.14995105 | 0.74341484 |
| RSL24D1 | protein_coding | 0.14942853 | 3.19213806 |
| BRD8 | protein_coding | 0.14941704 | 3.5026595 |

| | | | |
|---|---|---|---|
| CD164 | protein_coding | 0.1493728 | 2.04318079 |
| GJC1 | protein_coding | 0.14922558 | 3.31302357 |
| TFRC | protein_coding | 0.14916987 | 6.48169914 |
| MCOLN2 | protein_coding | 0.14901378 | 0.40240084 |
| ACRV1 | protein_coding | 0.14900849 | 0.41413723 |
| PLAG1 | protein_coding | 0.14873846 | 0.40701351 |
| SRRM3 | protein_coding | 0.14871745 | 2.23536986 |
| DUSP9 | protein_coding | 0.14867797 | 1.1016908 |
| NUP54 | protein_coding | 0.14862147 | 2.7437986 |
| RNF187 | protein_coding | 0.14858716 | 5.90057753 |
| GALC | protein_coding | 0.14856969 | 1.23084829 |
| TMLHE | protein_coding | 0.14852033 | 0.12355323 |
| CGRRF1 | protein_coding | 0.14850174 | 0.43996528 |
| SETBP1-DT | lncRNA | 0.14847494 | 0.21577837 |
| MUS81 | protein_coding | 0.14836967 | 2.67381241 |
| NR3C1 | protein_coding | 0.14829442 | 3.32821283 |
| CLUH | protein_coding | 0.14822499 | 3.04262457 |
| RP11-809O1 | lncRNA | 0.14819424 | 0.29390181 |
| MTERF2 | protein_coding | 0.14810886 | 1.53641504 |
| ARIH2 | protein_coding | 0.14810088 | 3.67431434 |
| ADNP | protein_coding | 0.14808291 | 5.2262074 |
| RPS6KC1 | protein_coding | 0.14800046 | 1.96161329 |
| NAGK | protein_coding | 0.1479115 | 2.80712596 |
| TIGD6 | protein_coding | 0.14785471 | 1.09616163 |
| TTC9C | protein_coding | 0.1478104 | 2.15011964 |
| LINC02321 | lncRNA | 0.14772664 | 0.14410127 |
| TMOD1 | protein_coding | 0.14757651 | 1.89942685 |
| HEXB | protein_coding | 0.14753603 | 3.15401115 |
| TIAM2 | protein_coding | 0.1473971 | 0.59699786 |
| HPS3 | protein_coding | 0.14736859 | 0.96321687 |
| PSMC5 | protein_coding | 0.14721384 | 4.92118722 |
| ROBO2 | protein_coding | 0.14700621 | 2.990234 |
| C5orf15 | protein_coding | 0.14698138 | 2.25100828 |
| RP4-621N11 | lncRNA | 0.14695068 | 0.15219772 |
| SRA1 | protein_coding | 0.14691032 | 2.99668141 |
| C12orf73 | protein_coding | 0.14679381 | 1.58813982 |
| GPR3 | protein_coding | 0.14679049 | 0.62995852 |
| PLEKHA3 | protein_coding | 0.1466171 | 2.1155233 |
| RAB23 | protein_coding | 0.14658355 | 3.32098442 |
| RP11-342M1 | lncRNA | 0.14655958 | 0.6093685 |
| GNA13 | protein_coding | 0.14639281 | 2.96891565 |
| EPCAM-DT | lncRNA | 0.14621788 | 0.37426335 |
| CLEC18C | protein_coding | 0.14619105 | 0.20758273 |
| PUM3 | protein_coding | 0.14613027 | 3.12875317 |
| RPH3A | protein_coding | 0.14605267 | 0.22566435 |

| | | | |
|---|---|---|---|
| SSBP1 | protein_coding | 0.14600675 | 3.36609324 |
| GFRA4 | protein_coding | 0.14594892 | 0.18142182 |
| AKAP9 | protein_coding | 0.1459399 | 3.91223254 |
| CCDC43 | protein_coding | 0.14589983 | 2.58178637 |
| PRDM4 | protein_coding | 0.14586256 | 2.78163191 |
| COA7 | protein_coding | 0.14581965 | 4.05293927 |
| TAF13 | protein_coding | 0.14573701 | 1.68168116 |
| TM2D3 | protein_coding | 0.1457167 | 3.21878558 |
| CENATAC | protein_coding | 0.14563214 | 1.29945778 |
| MRRF | protein_coding | 0.14549528 | 1.59894833 |
| TATDN2 | protein_coding | 0.14546254 | 1.94118932 |
| ZNF84 | protein_coding | 0.14541301 | 3.99676774 |
| UBN1 | protein_coding | 0.1453772 | 4.66443036 |
| NOP58 | protein_coding | 0.14535462 | 4.49863129 |
| COQ6 | protein_coding | 0.14534349 | 3.29269847 |
| RORA-AS1 | lncRNA | 0.14526003 | 0.33567215 |
| SELENOM | protein_coding | 0.14509304 | 1.63873483 |
| DNAJC22 | protein_coding | 0.14506268 | 0.13913845 |
| UBE2E2 | protein_coding | 0.14500577 | 1.14499449 |
| CLASP1 | protein_coding | 0.1449858 | 3.92099889 |
| SCUBE1 | protein_coding | 0.1449846 | 0.97825923 |
| EEF1AKMT3 | protein_coding | 0.14496584 | 0.92585239 |
| ERMARD | protein_coding | 0.1449277 | 0.9566532 |
| RWDD3 | protein_coding | 0.14489764 | 0.2804646 |
| RPAP3 | protein_coding | 0.14486618 | 3.38112286 |
| ZBTB17 | protein_coding | 0.14483386 | 2.73958112 |
| ZNF581 | protein_coding | 0.14481643 | 1.74736258 |
| KIAA2026 | protein_coding | 0.14467726 | 2.66599155 |
| POU4F3 | protein_coding | 0.14467595 | 4.7516261 |
| TMEM120B | protein_coding | 0.14462873 | 4.3772448 |
| PDCD6IP | protein_coding | 0.14460942 | 2.59043712 |
| TRPC1 | protein_coding | 0.14457455 | 1.53576402 |
| OBSCN | protein_coding | 0.14455585 | 2.5656482 |
| ZNF793 | protein_coding | 0.14454227 | 1.81481501 |
| RABL3 | protein_coding | 0.14444608 | 1.61271405 |
| ZUP1 | protein_coding | 0.14433886 | 1.15658941 |
| CRISP2 | protein_coding | 0.14411737 | 0.23174628 |
| ECI1 | protein_coding | 0.14401757 | 2.70702553 |
| USP32 | protein_coding | 0.14385509 | 2.80013233 |
| P4HTM | protein_coding | 0.14382409 | 2.49057184 |
| TBCEL | protein_coding | 0.14373692 | 1.1439786 |
| METTL17 | protein_coding | 0.14363374 | 3.12627508 |
| STN1 | protein_coding | 0.14362463 | 1.45083829 |
| TRAF5 | protein_coding | 0.14342903 | 0.25290916 |
| RP11-715J22 | lncRNA | 0.1434158 | 0.21752759 |

| | | | |
|---|---|---|---|
| TAPT1 | protein_coding | 0.14337244 | 3.19961459 |
| RUFY1 | protein_coding | 0.14330884 | 1.84242939 |
| ANKIB1 | protein_coding | 0.14315485 | 2.04465428 |
| ASIC3 | protein_coding | 0.14308508 | 0.72794104 |
| PGAP6 | protein_coding | 0.14308325 | 3.02076491 |
| INPP5J | protein_coding | 0.14303207 | 1.56375578 |
| SMARCD1 | protein_coding | 0.14298146 | 4.99908684 |
| BX470102.3 | lncRNA | 0.14291579 | 0.15374098 |
| ARHGEF10L | protein_coding | 0.14289397 | 2.71040156 |
| ST20-AS1 | lncRNA | 0.14283799 | 0.54378125 |
| GOLPH3 | protein_coding | 0.14278221 | 3.97355641 |
| PLXNA3 | protein_coding | 0.14277873 | 3.78015117 |
| POLR1C | protein_coding | 0.14277276 | 1.99866478 |
| WDR36 | protein_coding | 0.14267627 | 3.30779668 |
| FIGN | protein_coding | 0.14249023 | 1.6474589 |
| OPLAH | protein_coding | 0.14246237 | 0.15578307 |
| NECAB3 | protein_coding | 0.14243819 | 2.06721274 |
| RC3H1 | protein_coding | 0.14240285 | 1.56672068 |
| CD79B | protein_coding | 0.14227584 | 0.78192637 |
| TMBIM6 | protein_coding | 0.14205159 | 5.70963717 |
| ZBTB43 | protein_coding | 0.1419055 | 1.8595108 |
| ERBB4 | protein_coding | 0.14177249 | 0.77866086 |
| COASY | protein_coding | 0.14169136 | 4.12792645 |
| CDADC1 | protein_coding | 0.14163753 | 0.88472004 |
| JMJD1C | protein_coding | 0.14161953 | 2.84242998 |
| MED30 | protein_coding | 0.14154642 | 1.35753224 |
| BCS1L | protein_coding | 0.14153676 | 3.2759435 |
| PALB2 | protein_coding | 0.1414143 | 2.05430077 |
| TXNDC9 | protein_coding | 0.14132947 | 2.06712578 |
| ANAPC7 | protein_coding | 0.14124836 | 4.24800347 |
| ZNF28 | protein_coding | 0.1410969 | 2.34240935 |
| ESCO1 | protein_coding | 0.14102984 | 3.46089734 |
| RSRC2 | protein_coding | 0.14100104 | 3.32175523 |
| VPS16 | protein_coding | 0.14099033 | 3.26221587 |
| IMMP2L | protein_coding | 0.14090892 | 1.0104942 |
| KIAA0586 | protein_coding | 0.14084047 | 2.69411844 |
| REX1BD | protein_coding | 0.14075542 | 0.72413352 |
| MYL12A | protein_coding | 0.14072079 | 2.29623246 |
| CPLANE1 | protein_coding | 0.14063406 | 2.09497032 |
| DPH7 | protein_coding | 0.14059375 | 2.77394512 |
| SNAPC5 | protein_coding | 0.1405743 | 1.56721412 |
| RP11-26J3.3 | lncRNA | 0.14055967 | 0.14217537 |
| RP11-16E12. | lncRNA | 0.14047641 | 0.29058658 |
| CIB2 | protein_coding | 0.14045234 | 1.4769429 |
| MFSD8 | protein_coding | 0.14036755 | 1.55500407 |

| | | | |
|---|---|---|---|
| SPG21 | protein_coding | 0.14031746 | 3.63307367 |
| ANKRD49 | protein_coding | 0.14029354 | 1.19257169 |
| OLA1 | protein_coding | 0.14003423 | 4.6626821 |
| E2F4 | protein_coding | 0.1399501 | 2.83926547 |
| PIK3C2A | protein_coding | 0.13994937 | 3.05180808 |
| MAP3K2-DT | lncRNA | 0.13991618 | 0.35447435 |
| ZNF404 | protein_coding | 0.13988509 | 0.36008285 |
| RP13-216E22 | lncRNA | 0.13988337 | 0.13817273 |
| MYLK4 | protein_coding | 0.13982341 | 0.46160262 |
| MAP3K11 | protein_coding | 0.13980902 | 2.52074984 |
| UGDH | protein_coding | 0.13979469 | 1.42478876 |
| SOCS4 | protein_coding | 0.13970797 | 2.04794985 |
| SRSF6 | protein_coding | 0.139671 | 3.17533123 |
| KEAP1 | protein_coding | 0.13964869 | 3.57598475 |
| UAP1L1 | protein_coding | 0.13961067 | 2.01162434 |
| RP11-158H5. | lncRNA | 0.13955982 | 0.33800617 |
| CTD-3113P16 | lncRNA | 0.13955174 | 0.16969954 |
| HECW2-AS1 | lncRNA | 0.1394927 | 0.21603675 |
| RNF181 | protein_coding | 0.13943851 | 2.68957464 |
| DHX34 | protein_coding | 0.13942775 | 1.79421272 |
| AMIGO1 | protein_coding | 0.13938409 | 1.35125432 |
| CHURC1 | protein_coding | 0.13916338 | 2.57787183 |
| SNHG1 | lncRNA | 0.13914866 | 3.62744572 |
| MRPS11 | protein_coding | 0.13908241 | 2.86261931 |
| ZFP91 | protein_coding | 0.1390801 | 2.76206682 |
| RP11-156E8. | lncRNA | 0.13905089 | 0.14893799 |
| VBP1 | protein_coding | 0.13885704 | 2.77064773 |
| PTGS1 | protein_coding | 0.13875753 | 1.88056006 |
| CBX2 | protein_coding | 0.1387381 | 4.3274301 |
| GOLGA8K | protein_coding | 0.13865605 | 0.16270671 |
| PPP2R1B | protein_coding | 0.1385205 | 2.67611843 |
| TBC1D17 | protein_coding | 0.13848499 | 1.99301641 |
| RP11-111G1: | lncRNA | 0.13847126 | 0.25963723 |
| METTL6 | protein_coding | 0.13835606 | 1.4370073 |
| RPAIN | protein_coding | 0.13824773 | 2.99699522 |
| GUSB | protein_coding | 0.13824636 | 3.05798852 |
| PLBD2 | protein_coding | 0.13813149 | 3.21422106 |
| SHOC1 | protein_coding | 0.13808704 | 0.27981322 |
| ZCWPW1 | protein_coding | 0.13796711 | 0.47147983 |
| ZSCAN20 | protein_coding | 0.13796033 | 1.51061617 |
| SARS1 | protein_coding | 0.13791086 | 4.44652132 |
| MRPS22 | protein_coding | 0.13791044 | 1.55161433 |
| ABTB1 | protein_coding | 0.13790679 | 0.76749872 |
| DHX16 | protein_coding | 0.13783633 | 2.58981393 |
| GUSBP11 | lncRNA | 0.13781643 | 0.96358532 |

| | | | |
|---|---|---|---|
| MRPL4 | protein_coding | 0.1377822 | 2.11853718 |
| HSD3B7 | protein_coding | 0.13774894 | 1.2909377 |
| METTL25 | protein_coding | 0.13759142 | 0.61027352 |
| ORC4 | protein_coding | 0.13755658 | 2.15219272 |
| INTS3 | protein_coding | 0.13750811 | 3.96755773 |
| BVES-AS1 | lncRNA | 0.13747174 | 0.20153297 |
| RP3-329A5.8 | lncRNA | 0.13742649 | 0.33708796 |
| UBE3B | protein_coding | 0.13730683 | 2.87664348 |
| NAA20 | protein_coding | 0.13715698 | 3.06392515 |
| C5orf22 | protein_coding | 0.13714566 | 2.29700477 |
| ING5 | protein_coding | 0.13703265 | 3.27422022 |
| ZNF26 | protein_coding | 0.13696505 | 1.73031512 |
| TOMM34 | protein_coding | 0.13680825 | 2.7435289 |
| COPRS | protein_coding | 0.13680639 | 2.31048928 |
| ZNF195 | protein_coding | 0.13676365 | 2.12482627 |
| VPS29 | protein_coding | 0.13670386 | 2.78222996 |
| AVL9 | protein_coding | 0.13665482 | 4.25403268 |
| PIP5K1A | protein_coding | 0.13660942 | 4.12133839 |
| SNHG11 | lncRNA | 0.13660184 | 1.45579414 |
| CHKA | protein_coding | 0.1365522 | 3.14572578 |
| AC133680.1 | lncRNA | 0.1365344 | 0.1668093 |
| RBIS | protein_coding | 0.13650272 | 1.65669595 |
| ZNF551 | protein_coding | 0.13638259 | 2.64905942 |
| SMC6 | protein_coding | 0.13610855 | 3.80485867 |
| LETMD1 | protein_coding | 0.13596036 | 3.47551469 |
| LINC01563 | lncRNA | 0.13592819 | 0.6827063 |
| MAN1A2 | protein_coding | 0.13587196 | 3.90008278 |
| EIF4G3 | protein_coding | 0.13585092 | 4.66862262 |
| KCTD9 | protein_coding | 0.13577262 | 1.83679511 |
| DDIAS | protein_coding | 0.13573895 | 1.76957796 |
| ZNF280B | protein_coding | 0.13553363 | 1.18466876 |
| PDCD4 | protein_coding | 0.13535745 | 2.89051492 |
| VPS72 | protein_coding | 0.13515115 | 3.11494842 |
| TAF6L | protein_coding | 0.13499401 | 1.95716611 |
| ZNF580 | protein_coding | 0.1349933 | 1.08893523 |
| SHTN1 | protein_coding | 0.13492533 | 3.18834098 |
| RP11-398C13 | protein_coding | 0.13490586 | 1.8843409 |
| GOLGA8B | protein_coding | 0.13490018 | 2.15463351 |
| TTC1 | protein_coding | 0.13481827 | 2.90801976 |
| PCBP3 | protein_coding | 0.1347881 | 2.72489429 |
| TTC19 | protein_coding | 0.13475761 | 2.11322425 |
| DDX19A | protein_coding | 0.13467747 | 1.3633835 |
| SCAF11 | protein_coding | 0.13466841 | 4.48121508 |
| POLK | protein_coding | 0.13457958 | 2.34014668 |
| LINC01535 | lncRNA | 0.13456127 | 0.28286944 |

| | | | |
|---|---|---|---|
| EXOSC7 | protein_coding | 0.13435084 | 1.07176619 |
| DNAJB4 | protein_coding | 0.13407968 | 1.21365918 |
| ZNF75A | protein_coding | 0.13407341 | 2.58121543 |
| CCS | protein_coding | 0.13405326 | 1.24034028 |
| ZNF529 | protein_coding | 0.13404794 | 2.17185804 |
| NARF-AS2 | lncRNA | 0.13404069 | 0.23447531 |
| PRNP | protein_coding | 0.13402879 | 2.90154521 |
| C1orf52 | protein_coding | 0.13401524 | 2.13903495 |
| GTPBP10 | protein_coding | 0.13384018 | 1.42660959 |
| EIF2AK3 | protein_coding | 0.13381905 | 1.8809826 |
| NMD3 | protein_coding | 0.13381807 | 2.37173482 |
| ANTKMT | protein_coding | 0.13349894 | 0.49650567 |
| SOS2 | protein_coding | 0.13330924 | 2.06141482 |
| PRKAG2 | protein_coding | 0.13305851 | 2.3485418 |
| PHOSPHO2 | protein_coding | 0.13301697 | 0.43604736 |
| ZNF253 | protein_coding | 0.13295442 | 2.12223496 |
| NEU3 | protein_coding | 0.13288679 | 2.34887543 |
| GTF3C4 | protein_coding | 0.13284091 | 3.7235673 |
| CLIP1 | protein_coding | 0.1328015 | 2.74038147 |
| SLFNL1-AS1 | lncRNA | 0.13266515 | 0.48014548 |
| TWF1 | protein_coding | 0.13257541 | 1.62641018 |
| ECD | protein_coding | 0.13249621 | 1.47228247 |
| ZBTB38 | protein_coding | 0.13246895 | 3.09710897 |
| LDLR | protein_coding | 0.13244443 | 3.05760331 |
| RP11-176H8. | protein_coding | 0.1323942 | 0.17790759 |
| CNTNAP5 | protein_coding | 0.13209137 | 1.06798475 |
| SECISBP2L | protein_coding | 0.13208914 | 3.01372591 |
| FMNL3 | protein_coding | 0.1320793 | 2.68526378 |
| ARL1 | protein_coding | 0.13206282 | 2.10836309 |
| TMEM9B | protein_coding | 0.13200775 | 2.9345388 |
| ZNF416 | protein_coding | 0.13184281 | 1.00958988 |
| PPARA | protein_coding | 0.13176976 | 2.34075811 |
| MADCAM1-A | lncRNA | 0.13163875 | 0.15906567 |
| RP11-161M6 | lncRNA | 0.13162134 | 0.24650411 |
| CEPT1 | protein_coding | 0.1316202 | 1.83763744 |
| PTGES2 | protein_coding | 0.13157959 | 2.95093549 |
| TMEM126B | protein_coding | 0.13156046 | 1.44864024 |
| SUCO | protein_coding | 0.13153348 | 4.0315897 |
| RNF214 | protein_coding | 0.13147313 | 2.01578648 |
| ZFYVE19 | protein_coding | 0.13133534 | 1.85747703 |
| CALR | protein_coding | 0.13124942 | 6.24172108 |
| ZNF529-AS1 | lncRNA | 0.13096333 | 0.44864976 |
| TAF10 | protein_coding | 0.13091337 | 0.8700243 |
| ZGRF1 | protein_coding | 0.13089734 | 1.85918892 |
| TBC1D5 | protein_coding | 0.13082425 | 3.23909126 |

| | | | |
|---|---|---|---|
| PDIA4 | protein_coding | 0.13081844 | 3.9932706 |
| TAF3 | protein_coding | 0.13076427 | 1.09365927 |
| CTD-2517M2 | lncRNA | 0.13073509 | 0.48638193 |
| ZNF518B | protein_coding | 0.13068662 | 2.04852168 |
| UBQLN1 | protein_coding | 0.13065961 | 4.2163766 |
| RMI2 | protein_coding | 0.13055017 | 2.67217694 |
| GPR27 | protein_coding | 0.13041149 | 1.83741083 |
| GDPGP1 | protein_coding | 0.13031812 | 0.37399412 |
| DOK6 | protein_coding | 0.1302208 | 1.2700633 |
| C2CD3 | protein_coding | 0.1301451 | 3.34376763 |
| LINC02348 | lncRNA | 0.13012742 | 0.57524874 |
| BECN1 | protein_coding | 0.13009405 | 2.31502641 |
| MARK2 | protein_coding | 0.12995867 | 3.24761519 |
| ATIC | protein_coding | 0.12988649 | 4.41246039 |
| HERC4 | protein_coding | 0.1298339 | 1.43182127 |
| AAGAB | protein_coding | 0.12970386 | 3.82829625 |
| FAM185A | protein_coding | 0.12967 | 0.57908926 |
| CSPP1 | protein_coding | 0.12962547 | 1.11250561 |
| ESCO2 | protein_coding | 0.12958569 | 0.98501324 |
| RP11-472I20 | lncRNA | 0.12955288 | 0.27823776 |
| ZNF527 | protein_coding | 0.12948096 | 1.27347162 |
| DYNLT3 | protein_coding | 0.12945428 | 1.10656024 |
| NUP160 | protein_coding | 0.12927448 | 4.60837636 |
| ASAH2B | protein_coding | 0.1292649 | 0.64674644 |
| TFB2M | protein_coding | 0.12921256 | 1.40362933 |
| SIKE1 | protein_coding | 0.1290228 | 2.50307176 |
| SDCCAG8 | protein_coding | 0.12901215 | 2.22964244 |
| NXT1 | protein_coding | 0.1289054 | 1.56747953 |
| ZNF776 | protein_coding | 0.12878691 | 2.10041298 |
| PEX5L | protein_coding | 0.12870171 | 4.20683696 |
| GFER | protein_coding | 0.12862685 | 1.96307098 |
| ARGLU1 | protein_coding | 0.1285749 | 3.77354386 |
| SLC25A40 | protein_coding | 0.1285731 | 1.45761664 |
| RNASEH2B | protein_coding | 0.12857258 | 2.09327509 |
| PTPDC1 | protein_coding | 0.12848707 | 1.94594125 |
| RBM22 | protein_coding | 0.12848048 | 3.61400979 |
| TAB3 | protein_coding | 0.12828517 | 2.52005481 |
| FAM234A | protein_coding | 0.12827747 | 3.25629369 |
| WDR26 | protein_coding | 0.12826885 | 3.169806 |
| NMT1 | protein_coding | 0.12825512 | 4.42688607 |
| COPS5 | protein_coding | 0.12821466 | 1.68198394 |
| STXBP3 | protein_coding | 0.1281741 | 2.23169023 |
| ERCC5 | protein_coding | 0.12797626 | 0.64878961 |
| CD99 | protein_coding | 0.12782682 | 1.56631858 |
| SEPTIN6 | protein_coding | 0.12781445 | 1.40444521 |

| | | | |
|---|---|---|---|
| PSMD6 | protein_coding | 0.12773689 | 1.87935116 |
| CSNK1D | protein_coding | 0.12772686 | 2.94969304 |
| CCDC174 | protein_coding | 0.12772426 | 1.104551 |
| KIF9-AS1 | lncRNA | 0.12762692 | 0.43319891 |
| SKA1 | protein_coding | 0.12761321 | 1.90430533 |
| GALK2 | protein_coding | 0.12759887 | 2.17848182 |
| CCPG1 | protein_coding | 0.12756076 | 0.45629388 |
| MYBBP1A | protein_coding | 0.12752274 | 2.21816936 |
| PIR | protein_coding | 0.1275176 | 2.9070306 |
| APH1A | protein_coding | 0.12748207 | 3.79261548 |
| FNTA | protein_coding | 0.12744552 | 1.90383859 |
| MTG2 | protein_coding | 0.12736858 | 2.39447865 |
| DSTYK | protein_coding | 0.12720727 | 1.36648893 |
| RP11-73M18 | lncRNA | 0.12719022 | 0.78496706 |
| CCDC17 | protein_coding | 0.12715781 | 0.24265455 |
| THOC1 | protein_coding | 0.12705457 | 2.4713109 |
| PPP2CB | protein_coding | 0.12704408 | 2.00447438 |
| FAHD2A | protein_coding | 0.12699506 | 1.58268268 |
| NAPB | protein_coding | 0.12697744 | 3.18038812 |
| EMILIN1 | protein_coding | 0.12694514 | 0.32086406 |
| CRTC3-AS1 | lncRNA | 0.12692837 | 0.18820839 |
| CDKN2A | protein_coding | 0.1268783 | 1.37638212 |
| TAF7 | protein_coding | 0.12677071 | 3.67861252 |
| ZNF316 | protein_coding | 0.12673913 | 1.25745134 |
| CTU2 | protein_coding | 0.12671881 | 1.65215166 |
| TRPV1 | protein_coding | 0.12660703 | 0.49615897 |
| RASEF | protein_coding | 0.12659435 | 1.93439149 |
| PPA1 | protein_coding | 0.12656831 | 3.60123156 |
| DCAF4 | protein_coding | 0.12656039 | 1.30351267 |
| ZKSCAN1 | protein_coding | 0.12652272 | 4.13934596 |
| CORO2A | protein_coding | 0.12650932 | 4.62919988 |
| HDAC9 | protein_coding | 0.12648233 | 2.1857581 |
| SEPHS2 | protein_coding | 0.12645816 | 3.13535088 |
| AP5S1 | protein_coding | 0.12642669 | 1.05871366 |
| P2RX4 | protein_coding | 0.1264259 | 1.24530835 |
| TACO1 | protein_coding | 0.12641179 | 1.7543372 |
| SMIM10L1 | protein_coding | 0.12636931 | 2.05466958 |
| GLS | protein_coding | 0.1263155 | 1.91878036 |
| COG6 | protein_coding | 0.12629096 | 1.02910585 |
| FOXP4 | protein_coding | 0.12617522 | 3.29111887 |
| CNTLN | protein_coding | 0.12615809 | 2.70524297 |
| ARNILA | lncRNA | 0.12596437 | 0.25879531 |
| THOC5 | protein_coding | 0.12595211 | 2.08079091 |
| SEC24A | protein_coding | 0.12593571 | 2.47500585 |
| PCNP | protein_coding | 0.12586893 | 1.81654896 |

| | | | |
|---|---|---|---|
| RBM41 | protein_coding | 0.12574479 | 1.70411805 |
| TNKS | protein_coding | 0.12572813 | 3.06982335 |
| HIKESHI | protein_coding | 0.12570545 | 2.10571084 |
| MKLN1 | protein_coding | 0.12563815 | 3.26699265 |
| ARHGAP12 | protein_coding | 0.12562976 | 1.65056314 |
| MAN1B1-DT | lncRNA | 0.12554051 | 0.41395485 |
| JMJD6 | protein_coding | 0.12535548 | 2.05088731 |
| SPEG | protein_coding | 0.12532939 | 1.81969344 |
| RNF115 | protein_coding | 0.12513396 | 2.13375034 |
| OAT | protein_coding | 0.1250037 | 1.82575672 |
| TUBA1C | protein_coding | 0.12491397 | 3.85136042 |
| C2CD2L | protein_coding | 0.12483842 | 1.39823294 |
| ABCB7 | protein_coding | 0.12482834 | 1.85441578 |
| GPKOW | protein_coding | 0.12473426 | 1.47405897 |
| KRBA1 | protein_coding | 0.1246078 | 1.24432678 |
| NAPA | protein_coding | 0.12434072 | 3.71834812 |
| ZNF711 | protein_coding | 0.12433154 | 3.3156189 |
| MRPL16 | protein_coding | 0.12432798 | 1.67206553 |
| FAM193A | protein_coding | 0.12429525 | 2.81431613 |
| MAGT1 | protein_coding | 0.12417405 | 2.50943777 |
| THNSL1 | protein_coding | 0.12415842 | 0.8122994 |
| XPO5 | protein_coding | 0.12411272 | 4.30605935 |
| JPT1 | protein_coding | 0.12406068 | 4.08897193 |
| NDUFS4 | protein_coding | 0.12405785 | 2.09858683 |
| AP3S2 | protein_coding | 0.12360515 | 1.23400513 |
| MTMR9 | protein_coding | 0.1235397 | 1.32817951 |
| ZFP69B | protein_coding | 0.12327899 | 1.78799834 |
| RMND1 | protein_coding | 0.12320494 | 1.22218796 |
| GPRC5B | protein_coding | 0.12318748 | 2.78078256 |
| NEK9 | protein_coding | 0.12315658 | 3.92711237 |
| GOLGA3 | protein_coding | 0.12315066 | 2.74259268 |
| CNPY2 | protein_coding | 0.12313412 | 0.81105314 |
| MAST1 | protein_coding | 0.12304404 | 1.94458452 |
| PCGF6 | protein_coding | 0.12302385 | 0.95742201 |
| RPAP1 | protein_coding | 0.12299116 | 2.47408729 |
| PDE4A | protein_coding | 0.12291918 | 0.15193245 |
| DNAJC21 | protein_coding | 0.12290173 | 3.11657129 |
| METTL21A | protein_coding | 0.12282531 | 2.02194006 |
| DHX36 | protein_coding | 0.12279409 | 3.6344746 |
| CREM | protein_coding | 0.12270447 | 0.71388267 |
| KLF6 | protein_coding | 0.12266467 | 1.24829177 |
| ETFBKMT | protein_coding | 0.12264931 | 0.16120069 |
| ZNF24 | protein_coding | 0.1223739 | 3.40544501 |
| TMEM39A | protein_coding | 0.12237379 | 0.90828206 |
| SPRTN | protein_coding | 0.12236311 | 1.30674232 |

| | | | |
|---|---|---|---|
| SSC4D | protein_coding | 0.12231175 | 0.25870462 |
| SCYL3 | protein_coding | 0.1222583 | 0.95594371 |
| KMT5A | protein_coding | 0.12225675 | 2.82646246 |
| WTAP | protein_coding | 0.12225447 | 3.106869 |
| CCNT2 | protein_coding | 0.12211583 | 2.16388813 |
| GTF2H2 | protein_coding | 0.12196302 | 0.7186526 |
| ZFX | protein_coding | 0.12178691 | 1.79798722 |
| ZNF280D | protein_coding | 0.12171514 | 1.14166669 |
| TM2D2 | protein_coding | 0.12170933 | 1.15284877 |
| TP53BP2 | protein_coding | 0.12162837 | 2.99732858 |
| LUC7L3 | protein_coding | 0.12156873 | 3.75570727 |
| RP11-384L8. | lncRNA | 0.1215613 | 0.19705815 |
| AC067956.1 | lncRNA | 0.12152159 | 0.194262 |
| CENPW | protein_coding | 0.12146755 | 1.14567595 |
| IKBKG | protein_coding | 0.12127642 | 0.63817391 |
| MICU1 | protein_coding | 0.12126082 | 1.48052364 |
| CLINT1 | protein_coding | 0.12125274 | 3.14724774 |
| PRMT7 | protein_coding | 0.12117887 | 2.23978164 |
| CERS6 | protein_coding | 0.12114692 | 4.08090323 |
| POLR1F | protein_coding | 0.12110892 | 1.4016466 |
| CTTN | protein_coding | 0.12109593 | 3.7401523 |
| THAP2 | protein_coding | 0.12099771 | 0.83387566 |
| PTRH2 | protein_coding | 0.12096548 | 1.85280279 |
| NKTR | protein_coding | 0.12093529 | 2.29469843 |
| HSPBAP1 | protein_coding | 0.12088637 | 0.69448317 |
| CIBAR1 | protein_coding | 0.12082048 | 0.90124443 |
| MTCH2 | protein_coding | 0.12069662 | 3.78479726 |
| SMARCB1 | protein_coding | 0.12058524 | 3.66866919 |
| RPIA | protein_coding | 0.12055517 | 1.73047823 |
| ZNF334 | protein_coding | 0.12051251 | 1.7460071 |
| LYSMD1 | protein_coding | 0.1204939 | 1.31702747 |
| FBRS | protein_coding | 0.12047354 | 1.79884944 |
| ZNF451 | protein_coding | 0.1204081 | 2.07019802 |
| IFNAR2 | protein_coding | 0.12027091 | 0.99767967 |
| MRPS31 | protein_coding | 0.1202476 | 0.67587139 |
| USP6NL | protein_coding | 0.12023719 | 1.1225418 |
| CDC42SE1 | protein_coding | 0.12018808 | 2.13148512 |
| PRPF40B | protein_coding | 0.12018054 | 0.99025585 |
| ZNF14 | protein_coding | 0.12011042 | 1.35210522 |
| CCDC142 | protein_coding | 0.12004914 | 1.67963573 |
| ZNF181 | protein_coding | 0.11991842 | 1.05227996 |
| AFG1L | protein_coding | 0.11987218 | 0.47481157 |
| ERBIN | protein_coding | 0.11986295 | 3.14493821 |
| TRMT10A | protein_coding | 0.1198226 | 0.31386226 |
| KIN | protein_coding | 0.11980625 | 0.81851722 |

| | | | |
|---|---|---|---|
| CHFR | protein_coding | 0.11970692 | 2.53124272 |
| OGFOD3 | protein_coding | 0.11965132 | 1.46003538 |
| PLRG1 | protein_coding | 0.11964371 | 2.46179892 |
| C5orf63 | protein_coding | 0.11963974 | 0.44036678 |
| BAD | protein_coding | 0.1194954 | 2.07158579 |
| C9orf24 | protein_coding | 0.11942443 | 0.19357614 |
| CTD-3162L10 | lncRNA | 0.11939568 | 0.2209961 |
| NFAM1 | protein_coding | 0.11928335 | 0.30445213 |
| CWC27 | protein_coding | 0.11903388 | 2.33757364 |
| EP300 | protein_coding | 0.11899373 | 4.15666651 |
| RP11-394O4. | lncRNA | 0.11885769 | 0.18008333 |
| ZHX1 | protein_coding | 0.11857734 | 2.98212044 |
| RDH11 | protein_coding | 0.11854243 | 3.05324339 |
| MYL6B | protein_coding | 0.11845562 | 2.14292588 |
| CUL5 | protein_coding | 0.11845085 | 2.05816961 |
| TESC | protein_coding | 0.11841595 | 1.28677216 |
| RHBDL1 | protein_coding | 0.11833279 | 0.60876045 |
| BLOC1S2 | protein_coding | 0.11830391 | 1.66674699 |
| RAB11FIP5 | protein_coding | 0.1181401 | 1.77277248 |
| BCL10 | protein_coding | 0.11808207 | 1.30040176 |
| NUDT9 | protein_coding | 0.11804159 | 1.8989503 |
| RAB6A | protein_coding | 0.11802942 | 3.43529498 |
| ABHD18 | protein_coding | 0.11798036 | 1.29175741 |
| ZNF708 | protein_coding | 0.11795262 | 1.05171047 |
| LRRC27 | protein_coding | 0.11766788 | 0.35301338 |
| GPC2 | protein_coding | 0.11765328 | 0.98309175 |
| C12orf4 | protein_coding | 0.11764961 | 2.19148044 |
| LINC01915 | lncRNA | 0.11763416 | 0.18599904 |
| PARG | protein_coding | 0.11762803 | 1.24658748 |
| SENP7 | protein_coding | 0.11758512 | 1.74792671 |
| INTS10 | protein_coding | 0.11745094 | 2.46975589 |
| ZNF335 | protein_coding | 0.11744585 | 2.2482237 |
| TACC2 | protein_coding | 0.11744026 | 0.82428795 |
| DTWD1 | protein_coding | 0.11730012 | 1.43913161 |
| NBR1 | protein_coding | 0.11726692 | 3.68711002 |
| ZNF749 | protein_coding | 0.11724102 | 1.00958817 |
| SEZ6L | protein_coding | 0.11722323 | 3.4754139 |
| ARIH1 | protein_coding | 0.11721834 | 3.58612539 |
| FBXL5 | protein_coding | 0.11712138 | 2.39864799 |
| ESYT1 | protein_coding | 0.11704037 | 3.77669151 |
| TMEM272 | protein_coding | 0.11703882 | 0.35728873 |
| DDX10 | protein_coding | 0.11703286 | 1.27906227 |
| KIF20B | protein_coding | 0.11699743 | 1.67928807 |
| HMGN3 | protein_coding | 0.11696373 | 2.76105829 |
| DOP1A | protein_coding | 0.11672631 | 2.80823051 |

| | | | |
|---|---|---|---|
| ZNF587B | protein_coding | 0.11671229 | 1.25330561 |
| ABCF1 | protein_coding | 0.11670604 | 4.3333177 |
| USP28 | protein_coding | 0.11651842 | 0.70105459 |
| PPFIA3 | protein_coding | 0.11644588 | 2.17568788 |
| RPRD1B | protein_coding | 0.11635978 | 3.24706931 |
| THOC2 | protein_coding | 0.1163405 | 3.01760958 |
| RP11-513M1 | lncRNA | 0.11626749 | 0.21057925 |
| B4GALT5 | protein_coding | 0.11610627 | 4.08259746 |
| HIF1A | protein_coding | 0.11602469 | 2.44851292 |
| BCL7C | protein_coding | 0.11598736 | 1.52792407 |
| ATG4B | protein_coding | 0.11598605 | 2.85270252 |
| PCBP1-AS1 | lncRNA | 0.11597232 | 1.72358804 |
| MAGOH | protein_coding | 0.11597141 | 1.87894449 |
| CYC1 | protein_coding | 0.11573868 | 2.80242642 |
| NGLY1 | protein_coding | 0.11566479 | 1.30691821 |
| ZNF397 | protein_coding | 0.11562118 | 2.34904843 |
| ZNF467 | protein_coding | 0.11551503 | 0.67525269 |
| KCNAB2 | protein_coding | 0.11518619 | 1.3339439 |
| CBR4 | protein_coding | 0.11512236 | 2.32314543 |
| TRIM4 | protein_coding | 0.11506942 | 1.42927716 |
| UHMK1 | protein_coding | 0.11505538 | 2.51892915 |
| RIC3 | protein_coding | 0.11503154 | 1.28846936 |
| TRIM37 | protein_coding | 0.11497564 | 3.54355844 |
| CCDC14 | protein_coding | 0.11491195 | 2.37068814 |
| FLAD1 | protein_coding | 0.11490556 | 2.88079183 |
| MRPL3 | protein_coding | 0.11486734 | 3.65876284 |
| ZNF821 | protein_coding | 0.11483593 | 1.3037054 |
| RAB34 | protein_coding | 0.11478989 | 0.43149045 |
| LARGE1 | protein_coding | 0.11474193 | 1.10694332 |
| PPP4R3B | protein_coding | 0.11462169 | 3.62151957 |
| GTPBP4 | protein_coding | 0.11461417 | 2.60353615 |
| MRPL32 | protein_coding | 0.11447713 | 1.71145307 |
| UBAC2 | protein_coding | 0.11446659 | 3.10612394 |
| SNAPC4 | protein_coding | 0.11436248 | 1.8665823 |
| ZNF358 | protein_coding | 0.11435544 | 0.56905489 |
| ATF6B | protein_coding | 0.11432498 | 2.91978071 |
| NBEA | protein_coding | 0.11427156 | 2.21962149 |
| CHCHD10 | protein_coding | 0.11423005 | 1.00791389 |
| LTO1 | protein_coding | 0.11404475 | 1.4928346 |
| ANKMY1 | protein_coding | 0.11398712 | 1.13036607 |
| CLDN12 | protein_coding | 0.11398646 | 1.1208253 |
| STK3 | protein_coding | 0.11396586 | 1.12420323 |
| PEX7 | protein_coding | 0.11392431 | 0.75528921 |
| TBCC | protein_coding | 0.1138723 | 2.36300984 |
| ASAH1 | protein_coding | 0.1138688 | 1.20167454 |

| | | | |
|---|---|---|---|
| ZNF500 | protein_coding | 0.11384201 | 1.00877815 |
| FAM222B | protein_coding | 0.11377905 | 1.57447389 |
| AGO4 | protein_coding | 0.11376834 | 1.59756442 |
| DALRD3 | protein_coding | 0.11375679 | 0.46709398 |
| POLM | protein_coding | 0.11374878 | 1.2219439 |
| MTRF1 | protein_coding | 0.11366048 | 0.53815644 |
| RP11-53O19. | lncRNA | 0.11331415 | 0.56151854 |
| ALKBH3 | protein_coding | 0.11315949 | 1.35720792 |
| MCUR1 | protein_coding | 0.11304886 | 2.1532514 |
| C18orf21 | protein_coding | 0.11291745 | 0.89544269 |
| GOLGA4-AS1 | lncRNA | 0.1127508 | 0.23935382 |
| NRSN2 | protein_coding | 0.11275061 | 2.60009832 |
| MED13 | protein_coding | 0.11259261 | 3.4166975 |
| KLK15 | protein_coding | 0.11235437 | 0.92006552 |
| GMDS-DT | lncRNA | 0.1123304 | 0.38607994 |
| MYPOP | protein_coding | 0.11230278 | 0.82870698 |
| SNX2 | protein_coding | 0.11228232 | 2.70757216 |
| RING1 | protein_coding | 0.11223633 | 2.23106626 |
| NAA35 | protein_coding | 0.1121415 | 1.93129822 |
| RP5-864K19. | lncRNA | 0.11207992 | 0.24656948 |
| RAB11FIP2 | protein_coding | 0.11190557 | 1.55896529 |
| ZFHX2-AS1 | lncRNA | 0.11183779 | 0.27438558 |
| HBP1 | protein_coding | 0.11176852 | 1.25293469 |
| P4HB | protein_coding | 0.11168898 | 4.31378997 |
| APTX | protein_coding | 0.11163073 | 2.26849804 |
| SLC35A1 | protein_coding | 0.111496 | 0.92441803 |
| RNF26 | protein_coding | 0.11128149 | 2.4823331 |
| BPTF | protein_coding | 0.11122676 | 3.44805491 |
| MAF | protein_coding | 0.11119109 | 0.95631752 |
| NGDN | protein_coding | 0.11117071 | 1.86088033 |
| UTP20 | protein_coding | 0.11111666 | 2.24353411 |
| RPL22L1 | protein_coding | 0.11098054 | 2.3981504 |
| MT1X | protein_coding | 0.11097829 | 0.43047328 |
| MAGOHB | protein_coding | 0.11086595 | 1.70195039 |
| CLDN15 | protein_coding | 0.11078162 | 1.24292124 |
| DGCR8 | protein_coding | 0.11071413 | 2.40558043 |
| TMCC1-DT | lncRNA | 0.11065462 | 0.28369955 |
| LENG8 | protein_coding | 0.11063864 | 2.83724186 |
| SFMBT1 | protein_coding | 0.11056415 | 1.37816218 |
| SRCAP | protein_coding | 0.11052405 | 0.57585323 |
| CHIC2 | protein_coding | 0.11038922 | 1.04184416 |
| STK38 | protein_coding | 0.11037556 | 3.15988208 |
| PYCR1 | protein_coding | 0.11037323 | 3.95380979 |
| KARS1 | protein_coding | 0.11036807 | 3.57962639 |
| DHRS13 | protein_coding | 0.11029239 | 1.7369642 |

| | | | |
|---|---|---|---|
| DPP8 | protein_coding | 0.11027665 | 2.6030318 |
| SLC16A2 | protein_coding | 0.1102743 | 0.61970666 |
| SECISBP2 | protein_coding | 0.11025921 | 2.56921584 |
| RRP15 | protein_coding | 0.11014284 | 1.50463583 |
| TRAF2 | protein_coding | 0.11007796 | 1.67354373 |
| ANKRD11 | protein_coding | 0.11004655 | 2.79790072 |
| ATG5 | protein_coding | 0.10994689 | 1.1229146 |
| ZFYVE16 | protein_coding | 0.10991964 | 1.52796583 |
| ZNF131 | protein_coding | 0.10987429 | 2.59224315 |
| C7orf26 | protein_coding | 0.10981492 | 1.4566912 |
| KBTBD8 | protein_coding | 0.10965911 | 0.87729557 |
| MED20 | protein_coding | 0.10961665 | 1.48601074 |
| SYNCRIP | protein_coding | 0.10961023 | 1.41284904 |
| ASPSCR1 | protein_coding | 0.10959822 | 1.66108038 |
| KIF1C | protein_coding | 0.10955001 | 0.93673566 |
| AP1S3 | protein_coding | 0.109527 | 0.62046629 |
| TMOD3 | protein_coding | 0.10943724 | 2.4229519 |
| NAP1L5 | protein_coding | 0.10941471 | 1.13048053 |
| UBN2 | protein_coding | 0.10936577 | 2.16919725 |
| PAN2 | protein_coding | 0.10929408 | 2.48799282 |
| EIF4EBP1 | protein_coding | 0.10926092 | 2.60746325 |
| FRAS1 | protein_coding | 0.10925297 | 3.97124321 |
| TDRD3 | protein_coding | 0.10924129 | 1.995212 |
| ZNF33B | protein_coding | 0.10922164 | 1.73571435 |
| ZNF106 | protein_coding | 0.10919747 | 4.02500747 |
| KLHL7-DT | lncRNA | 0.1090483 | 0.29683499 |
| TDP2 | protein_coding | 0.10894958 | 1.70465669 |
| MOB3A | protein_coding | 0.10891277 | 1.05920477 |
| RGS16 | protein_coding | 0.10888799 | 2.04162561 |
| TCF25 | protein_coding | 0.10883806 | 1.92321826 |
| NOL8 | protein_coding | 0.10879792 | 2.24308787 |
| ERLIN1 | protein_coding | 0.10855566 | 1.11295024 |
| NKAP | protein_coding | 0.10838094 | 1.48272265 |
| TMA16 | protein_coding | 0.10833581 | 1.51342537 |
| RP1-283E3.8 | lncRNA | 0.10832942 | 0.35029456 |
| RCC2 | protein_coding | 0.108013 | 4.31572524 |
| CD276 | protein_coding | 0.10791761 | 0.50162227 |
| GNS | protein_coding | 0.10791709 | 3.62077472 |
| PCGF1 | protein_coding | 0.10783618 | 0.94273732 |
| C1QBP | protein_coding | 0.10779798 | 2.09420791 |
| CDON | protein_coding | 0.10777045 | 0.86587042 |
| RNF44 | protein_coding | 0.10773815 | 2.12828079 |
| STX18 | protein_coding | 0.10772823 | 1.46446794 |
| PPRC1 | protein_coding | 0.10771955 | 2.45688609 |
| TJAP1 | protein_coding | 0.1076176 | 1.75581844 |

| | | | |
|---|---|---|---|
| NCK1 | protein_coding | 0.10754987 | 1.23472923 |
| CHD2 | protein_coding | 0.10752475 | 4.02516562 |
| ADGRV1 | protein_coding | 0.10750637 | 1.75326391 |
| SLC1A5 | protein_coding | 0.10728623 | 2.72227095 |
| ZNF623 | protein_coding | 0.1072753 | 2.18267976 |
| ANKH | protein_coding | 0.10725003 | 2.55355253 |
| RICTOR | protein_coding | 0.10711622 | 1.34968465 |
| ATP2B3 | protein_coding | 0.10708969 | 2.02325545 |
| TMEM52 | protein_coding | 0.10704181 | 0.17363692 |
| CTD-3099C6. | protein_coding | 0.10691601 | 1.14412105 |
| MRPL21 | protein_coding | 0.10690979 | 2.01446818 |
| HINT1 | protein_coding | 0.10683425 | 3.99701904 |
| PLEKHM3 | protein_coding | 0.10679103 | 1.54501975 |
| ACOXL | protein_coding | 0.10677748 | 0.39244012 |
| FAM177A1 | protein_coding | 0.10675279 | 2.31062717 |
| RP11-81A1.6 | lncRNA | 0.1067416 | 0.41373508 |
| WAC | protein_coding | 0.10672687 | 2.79754147 |
| NT5C2 | protein_coding | 0.10671403 | 1.64966511 |
| NDUFB2 | protein_coding | 0.1066327 | 2.03535997 |
| ACAT1 | protein_coding | 0.10659678 | 1.24975824 |
| HASPIN | protein_coding | 0.10652964 | 0.60888783 |
| YIPF2 | protein_coding | 0.10641587 | 1.9936359 |
| ROBO3 | protein_coding | 0.10639746 | 0.97373586 |
| GON4L | protein_coding | 0.10628125 | 2.55353672 |
| PSTK | protein_coding | 0.10624701 | 0.34613287 |
| CCDC110 | protein_coding | 0.10624388 | 0.66448191 |
| CWC22 | protein_coding | 0.10622302 | 1.7370242 |
| NDUFA6 | protein_coding | 0.10621935 | 3.37313356 |
| FIZ1 | protein_coding | 0.10612488 | 1.15035733 |
| AP5B1 | protein_coding | 0.10590618 | 1.12365995 |
| RARS1 | protein_coding | 0.10588817 | 2.39342048 |
| ARHGAP9 | protein_coding | 0.10582927 | 0.63796695 |
| ZNF655 | protein_coding | 0.10582568 | 2.42468093 |
| ERCC6 | protein_coding | 0.10576268 | 1.30520888 |
| RAB40C | protein_coding | 0.10561035 | 1.70688265 |
| ZKSCAN2-DT | lncRNA | 0.10557354 | 0.33553874 |
| PGD | protein_coding | 0.10556247 | 2.60315409 |
| RBM48 | protein_coding | 0.10555519 | 0.9819061 |
| ZBTB41 | protein_coding | 0.10544372 | 1.50434516 |
| ANKRA2 | protein_coding | 0.10541306 | 0.82225461 |
| RRNAD1 | protein_coding | 0.10534869 | 1.11800802 |
| ZNF669 | protein_coding | 0.1053385 | 0.68737312 |
| POC1B | protein_coding | 0.10532979 | 0.5769941 |
| SNX5 | protein_coding | 0.10524137 | 2.86981094 |
| C1orf109 | protein_coding | 0.10518257 | 2.18969218 |

| | | | |
|---|---|---|---|
| ZNRF2 | protein_coding | 0.10501326 | 0.8720624 |
| A1BG-AS1 | lncRNA | 0.10488572 | 0.79589923 |
| KTN1-AS1 | lncRNA | 0.10485771 | 0.59898599 |
| ABCB8 | protein_coding | 0.10478992 | 1.71728592 |
| GTF3C1 | protein_coding | 0.10477301 | 3.85280247 |
| SIAH2 | protein_coding | 0.10472668 | 1.78647513 |
| ACTR1B | protein_coding | 0.1047253 | 2.34848942 |
| CENPH | protein_coding | 0.10468759 | 1.87106636 |
| MICAL3 | protein_coding | 0.10454648 | 1.86599337 |
| SETD6 | protein_coding | 0.10453349 | 1.78305982 |
| FAM118B | protein_coding | 0.10448142 | 1.16764552 |
| LEMD2 | protein_coding | 0.10447072 | 1.88805793 |
| ARF6 | protein_coding | 0.10438671 | 2.79525301 |
| USP10 | protein_coding | 0.10429296 | 2.86246665 |
| CRNDE | lncRNA | 0.10417099 | 1.15397709 |
| DDX50 | protein_coding | 0.10413052 | 2.01750844 |
| AP4B1 | protein_coding | 0.10408426 | 1.77194901 |
| ABHD2 | protein_coding | 0.10400577 | 2.56018841 |
| HSPA5 | protein_coding | 0.1037648 | 4.24136394 |
| COPG2 | protein_coding | 0.10365196 | 1.38804801 |
| PGS1 | protein_coding | 0.10346507 | 1.29096484 |
| TOMM70 | protein_coding | 0.10342828 | 3.87845081 |
| PHIP | protein_coding | 0.10337729 | 2.57572163 |
| CDH24 | protein_coding | 0.10335882 | 2.40093782 |
| ZBED5 | protein_coding | 0.10324397 | 1.78768639 |
| MRPL45 | protein_coding | 0.10321982 | 1.58637986 |
| ADCYAP1 | protein_coding | 0.1031821 | 0.98550421 |
| CCDC149 | protein_coding | 0.1030747 | 1.26069656 |
| SIPA1L3 | protein_coding | 0.10304401 | 1.6315878 |
| CPSF2 | protein_coding | 0.10302212 | 2.95127841 |
| ABCA12 | protein_coding | 0.10298813 | 0.17515044 |
| FAM76B | protein_coding | 0.10290799 | 1.4188668 |
| ZPR1 | protein_coding | 0.10289992 | 2.58022811 |
| TSEN15 | protein_coding | 0.10288719 | 2.2106081 |
| MAPKAPK3 | protein_coding | 0.10275992 | 0.9642812 |
| OSBPL7 | protein_coding | 0.10272583 | 0.9901922 |
| LRRC3 | protein_coding | 0.10272435 | 0.87216571 |
| ERCC6L2 | protein_coding | 0.1025447 | 2.27350524 |
| NCOA1 | protein_coding | 0.10238801 | 3.00510533 |
| ZNF433-AS1 | lncRNA | 0.10232902 | 0.23747334 |
| RP11-689P11 | lncRNA | 0.10225096 | 0.14771862 |
| MATR3 | protein_coding | 0.10220593 | 0.2543623 |
| C1GALT1 | protein_coding | 0.10205268 | 1.34530197 |
| FAM90A1 | protein_coding | 0.10203381 | 0.4220508 |
| PHF20L1 | protein_coding | 0.10198838 | 2.34629685 |

| | | | |
|---|---|---|---|
| ANAPC1 | protein_coding | 0.10194742 | 2.16878067 |
| FAR1 | protein_coding | 0.10194142 | 2.68199284 |
| DCAF17 | protein_coding | 0.10192839 | 1.70921265 |
| UNC5CL | protein_coding | 0.10187256 | 0.22839397 |
| PIP4K2C | protein_coding | 0.10185034 | 2.13149796 |
| ANK1 | protein_coding | 0.1017568 | 1.66340218 |
| C6orf163 | protein_coding | 0.10162465 | 0.13875033 |
| OSBPL1A | protein_coding | 0.10161616 | 0.99367429 |
| MECR | protein_coding | 0.1015581 | 1.32897513 |
| SELENOO | protein_coding | 0.10154097 | 1.27034133 |
| CEP290 | protein_coding | 0.10149167 | 1.47611936 |
| CRYGS | protein_coding | 0.10147982 | 0.20683196 |
| FBXW4 | protein_coding | 0.10143055 | 1.23943988 |
| RP11-158K1. | lncRNA | 0.10141631 | 0.16722765 |
| YARS2 | protein_coding | 0.10140528 | 1.27586855 |
| SNX8 | protein_coding | 0.10139967 | 1.52845363 |
| SLC11A2 | protein_coding | 0.10130702 | 1.94813658 |
| CD7 | protein_coding | 0.10130577 | 0.12118109 |
| VN1R1 | protein_coding | 0.10129236 | 0.18163727 |
| RP11-158H5. | lncRNA | 0.1012796 | 0.13575569 |
| MYO5C | protein_coding | 0.10125602 | 1.24868704 |
| ATP6V1A | protein_coding | 0.10125451 | 1.78068287 |
| DUS4L-BCAP | protein_coding | 0.10124855 | 0.13441536 |
| USP21 | protein_coding | 0.10123584 | 1.66118325 |
| TRMT6 | protein_coding | 0.10123542 | 2.37728618 |
| RFNG | protein_coding | 0.10123121 | 1.14086831 |
| PITX2 | protein_coding | 0.10120242 | 0.17842328 |
| EIF3M | protein_coding | 0.10114359 | 2.68885951 |
| TRIM73 | protein_coding | 0.10111051 | 0.09180429 |
| FAM227B | protein_coding | 0.10099537 | 0.29509686 |
| TMEM267 | protein_coding | 0.1009663 | 0.64498159 |
| RP11-260L6.: | lncRNA | 0.10096245 | 0.09176761 |
| DNM1L | protein_coding | 0.1009173 | 2.88982626 |
| CTD-2325M2 | lncRNA | 0.10091062 | 0.12187645 |
| AC053503.4 | lncRNA | 0.10086762 | 0.12623452 |
| TAF2 | protein_coding | 0.10086585 | 2.33850523 |
| RP11-849H4. | lncRNA | 0.10082833 | 0.20134897 |
| PAM16 | protein_coding | 0.10079987 | 0.30757974 |
| NAV2-AS6 | lncRNA | 0.10077498 | 0.17019147 |
| RP11-363E6. | lncRNA | 0.10076789 | 0.21294303 |
| RP1-228P16. | lncRNA | 0.10076774 | 0.14844671 |
| WNK1 | protein_coding | 0.10076468 | 4.51653181 |
| DTX3 | protein_coding | 0.1007525 | 1.99822228 |
| USP16 | protein_coding | 0.10067111 | 2.08563964 |
| SIN3B | protein_coding | 0.10066866 | 1.76192165 |

| | | | |
|---|---|---|---|
| RP11-452L6. | lncRNA | 0.10064279 | 0.19944549 |
| POC1B-AS1 | lncRNA | 0.10059917 | 0.13922888 |
| HPCAL1 | protein_coding | 0.1005871 | 1.35578904 |
| RBM39 | protein_coding | 0.10055641 | 3.21982466 |
| NUDT13 | protein_coding | 0.10050233 | 0.13305321 |
| SIX3-AS1 | lncRNA | 0.10049834 | 0.11772401 |
| GTF2F2 | protein_coding | 0.10048974 | 1.24290296 |
| MLXIP | protein_coding | 0.10046184 | 2.53170815 |
| CTC-281F24. | protein_coding | 0.10045713 | 0.29530728 |
| SFN | protein_coding | 0.10045428 | 2.22655753 |
| CHODL | protein_coding | 0.10044754 | 1.02070326 |
| MYMX | protein_coding | 0.10032646 | 0.68801178 |
| MTPAP | protein_coding | 0.10032026 | 1.26521052 |
| KLHDC1 | protein_coding | 0.10031983 | 0.14620445 |
| ORC2 | protein_coding | 0.10031088 | 1.86984699 |
| FBXO30-DT | lncRNA | 0.10030505 | 0.11738586 |
| RP11-343L5. | lncRNA | 0.10027275 | 0.10083204 |
| RTKN | protein_coding | 0.10026752 | 0.52202582 |
| PANK2-AS1 | lncRNA | 0.1001834 | 0.11758718 |
| SOS1 | protein_coding | 0.10015615 | 2.01524959 |
| EMX1 | protein_coding | 0.10013204 | 0.35195814 |
| PARD6G-AS1 | lncRNA | 0.10012903 | 0.11374426 |
| FAM124A | protein_coding | 0.10010079 | 0.5039889 |
| LEAP2 | protein_coding | 0.10006925 | 0.13504591 |
| RAD17 | protein_coding | 0.10001861 | 2.06521979 |
| KYAT1 | protein_coding | 0.09994355 | 0.1159661 |
| SMIM30 | protein_coding | 0.09987284 | 0.97121357 |
| INTS13 | protein_coding | 0.09981772 | 1.83837622 |
| GABPB1 | protein_coding | 0.0997679 | 1.56831104 |
| BOD1 | protein_coding | 0.09976553 | 2.07439895 |
| SPAG4 | protein_coding | 0.09974835 | 0.23686314 |
| RP11-124N1 | lncRNA | 0.09973048 | 0.10998184 |
| AFAP1L1 | protein_coding | 0.09966476 | 0.10457759 |
| ATL2 | protein_coding | 0.09959846 | 2.16019215 |
| DDX19B | protein_coding | 0.09956777 | 0.94582727 |
| DHDDS | protein_coding | 0.09953441 | 2.02730702 |
| SH2D5 | protein_coding | 0.09952749 | 2.05464515 |
| RP11-359E3. | lncRNA | 0.09949669 | 0.14072046 |
| RP11-706O1 | lncRNA | 0.09949636 | 0.81496775 |
| TIA1 | protein_coding | 0.099365 | 1.6604122 |
| B3GALT2 | protein_coding | 0.09936094 | 0.10871362 |
| HPS1 | protein_coding | 0.09934603 | 0.99845293 |
| PDE6D | protein_coding | 0.09927296 | 1.79001278 |
| RBM18 | protein_coding | 0.09923792 | 2.203867 |
| ABCE1 | protein_coding | 0.09923061 | 2.52371831 |

| | | | |
|---|---|---|---|
| PGBD2 | protein_coding | 0.09920948 | 0.59711975 |
| WDR5B | protein_coding | 0.09919992 | 0.77328244 |
| TEX22 | protein_coding | 0.09906326 | 0.19597508 |
| CTD-3091F23 | lncRNA | 0.09905248 | 0.09243554 |
| MAPKAPK5 | protein_coding | 0.09901609 | 1.55355412 |
| PPP2R2A | protein_coding | 0.09899576 | 1.62349489 |
| RGS3 | protein_coding | 0.09895884 | 1.80591142 |
| RP11-534K14 | lncRNA | 0.09887332 | 0.72884359 |
| DPF2 | protein_coding | 0.09885587 | 2.21406118 |
| FBXW5 | protein_coding | 0.09876412 | 1.09969535 |
| WDR44 | protein_coding | 0.09864506 | 0.95453447 |
| CTB-58E17.3 | lncRNA | 0.09859507 | 0.08518739 |
| AFF4 | protein_coding | 0.09854458 | 2.45546909 |
| KANSL2 | protein_coding | 0.098454 | 1.94395155 |
| ECHDC1 | protein_coding | 0.09837865 | 1.35880013 |
| GDF11 | protein_coding | 0.09835118 | 2.08870555 |
| C17orf75 | protein_coding | 0.0983002 | 1.58606602 |
| ARAF | protein_coding | 0.09814022 | 1.93923504 |
| ACTR6 | protein_coding | 0.09810127 | 1.63045545 |
| SP1 | protein_coding | 0.09802075 | 3.06101685 |
| SMG8 | protein_coding | 0.09800643 | 1.62532474 |
| TOB1 | protein_coding | 0.09794504 | 1.6998245 |
| SYNRG | protein_coding | 0.09792842 | 2.68185499 |
| ZFAT | protein_coding | 0.09767801 | 0.98822572 |
| CCNC | protein_coding | 0.09764538 | 0.9593588 |
| MBD6 | protein_coding | 0.09756429 | 2.24424866 |
| PHAX | protein_coding | 0.09754764 | 1.93107096 |
| GHITM | protein_coding | 0.09753565 | 3.05027035 |
| COA5 | protein_coding | 0.09749808 | 0.99119948 |
| LINC01357 | lncRNA | 0.09747805 | 0.79990344 |
| ARF4 | protein_coding | 0.09730632 | 1.34103758 |
| ZNF318 | protein_coding | 0.09717365 | 2.206975 |
| TMEM116 | protein_coding | 0.09707852 | 0.98394208 |
| IP6K2 | protein_coding | 0.09686626 | 2.12290888 |
| CHST14 | protein_coding | 0.0968321 | 1.06821417 |
| CDC16 | protein_coding | 0.09681618 | 2.32908519 |
| TSPAN15 | protein_coding | 0.09651728 | 1.09335209 |
| KIF27 | protein_coding | 0.0963585 | 0.57619534 |
| PIGS | protein_coding | 0.09631356 | 3.20803377 |
| CYSTM1 | protein_coding | 0.09626359 | 0.86100533 |
| KIAA1328 | protein_coding | 0.09626069 | 1.02534478 |
| ERGIC3 | protein_coding | 0.09615781 | 2.47598502 |
| ZBTB37 | protein_coding | 0.09609544 | 0.90534994 |
| RNF25 | protein_coding | 0.09601898 | 0.89976234 |
| CSDE1 | protein_coding | 0.09599368 | 4.39326137 |

| | | | |
|---|---|---|---|
| NHEJ1 | protein_coding | 0.09596234 | 0.76407329 |
| INVS | protein_coding | 0.0959428 | 0.89136704 |
| CAPZA1 | protein_coding | 0.09584149 | 2.87418038 |
| ERVK3-1 | protein_coding | 0.09571662 | 0.77906975 |
| EIF2A | protein_coding | 0.09570239 | 2.39280632 |
| ALG6 | protein_coding | 0.09569194 | 1.05631441 |
| PTPN23 | protein_coding | 0.09537852 | 1.44393907 |
| WDR3 | protein_coding | 0.0952289 | 1.9970808 |
| PCDHA3 | protein_coding | 0.09520173 | 0.39072564 |
| EIF4G1 | protein_coding | 0.09506472 | 4.45392086 |
| RBKS | protein_coding | 0.09496588 | 0.26955081 |
| SEC31B | protein_coding | 0.09485664 | 0.40163074 |
| TRAPPC6B | protein_coding | 0.09482167 | 1.99512964 |
| HSCB | protein_coding | 0.09478248 | 0.63583266 |
| CIR1 | protein_coding | 0.09477746 | 1.15916122 |
| PSMG3-AS1 | lncRNA | 0.09477043 | 0.61902834 |
| AC006946.16 | lncRNA | 0.09475601 | 0.43832018 |
| EWSR1 | protein_coding | 0.09470793 | 3.59757466 |
| PICALM | protein_coding | 0.09466262 | 2.5784162 |
| MFAP2 | protein_coding | 0.09458857 | 0.2880693 |
| MED15 | protein_coding | 0.09456608 | 1.77319497 |
| C19orf44 | protein_coding | 0.09453636 | 0.5089834 |
| RPTOR | protein_coding | 0.0944926 | 1.49112555 |
| KLF10 | protein_coding | 0.09444715 | 1.57796831 |
| EEF1E1 | protein_coding | 0.09435029 | 0.57044377 |
| PRR14 | protein_coding | 0.09426712 | 1.7824865 |
| PPP2R5E | protein_coding | 0.09423808 | 2.10111751 |
| MLST8 | protein_coding | 0.09417267 | 2.16121677 |
| CASC3 | protein_coding | 0.09413533 | 3.47503084 |
| TMEM161B | protein_coding | 0.09400667 | 1.2745603 |
| CDKN1B | protein_coding | 0.09393979 | 2.53957414 |
| SUGP2 | protein_coding | 0.09387846 | 3.19143185 |
| LINC02731 | lncRNA | 0.09386177 | 0.25621577 |
| PPWD1 | protein_coding | 0.09384338 | 1.25982141 |
| KLHL20 | protein_coding | 0.093742 | 1.02915415 |
| SETD2 | protein_coding | 0.09370204 | 2.13541804 |
| VPS28 | protein_coding | 0.09349961 | 1.76600162 |
| DPP7 | protein_coding | 0.09341272 | 1.26143679 |
| MPDZ | protein_coding | 0.09327763 | 1.83064238 |
| APLP2 | protein_coding | 0.09324128 | 4.288578 |
| RP11-445N1[illegible] | lncRNA | 0.09319618 | 0.30873512 |
| HERPUD2 | protein_coding | 0.0931913 | 0.74589858 |
| ZNF408 | protein_coding | 0.09318998 | 0.95075959 |
| USP14 | protein_coding | 0.0929654 | 2.83989661 |
| TESMIN | protein_coding | 0.09283576 | 0.45436606 |

| | | | |
|---|---|---|---|
| FIG4 | protein_coding | 0.09275542 | 0.83039242 |
| KCTD21 | protein_coding | 0.09271848 | 0.38421801 |
| HM13 | protein_coding | 0.09261075 | 2.46730942 |
| ZFAND4 | protein_coding | 0.09253035 | 0.70516174 |
| NEK4 | protein_coding | 0.09242787 | 1.07561718 |
| POLR3E | protein_coding | 0.09233074 | 1.69791653 |
| MBD5 | protein_coding | 0.09231225 | 1.47863731 |
| PMS2 | protein_coding | 0.09217738 | 0.91622262 |
| ENTPD6 | protein_coding | 0.09215335 | 2.10896722 |
| TBC1D15 | protein_coding | 0.092128 | 1.29021027 |
| MSANTD4 | protein_coding | 0.092054 | 1.2689037 |
| DNTTIP2 | protein_coding | 0.09199276 | 2.10065587 |
| ZBTB1 | protein_coding | 0.09196517 | 2.0688289 |
| HYLS1 | protein_coding | 0.0919142 | 1.29359199 |
| MEF2A | protein_coding | 0.09183611 | 1.59753195 |
| SLC25A45 | protein_coding | 0.0918358 | 0.48235495 |
| RP5-1061H2( | lncRNA | 0.09180077 | 0.63942064 |
| CPSF7 | protein_coding | 0.09155491 | 2.93044289 |
| CARF | protein_coding | 0.09151534 | 0.51341096 |
| ZNF549 | protein_coding | 0.09150703 | 1.13671584 |
| BARD1 | protein_coding | 0.09142983 | 1.45838936 |
| COMMD1 | protein_coding | 0.09127905 | 0.88030142 |
| POFUT1 | protein_coding | 0.09122872 | 2.25548011 |
| RBL1 | protein_coding | 0.09115769 | 1.27217055 |
| PRKRA | protein_coding | 0.09115747 | 2.03410542 |
| ZNF548 | protein_coding | 0.09105022 | 1.03770918 |
| ZC3H15 | protein_coding | 0.09092775 | 2.31259404 |
| ST13 | protein_coding | 0.09087195 | 2.47575645 |
| ELP2 | protein_coding | 0.09072154 | 2.17317614 |
| LSM10 | protein_coding | 0.09069566 | 0.93207906 |
| CTDSP2 | protein_coding | 0.0906298 | 3.4644538 |
| SLC25A20 | protein_coding | 0.09055312 | 0.59907976 |
| MRPS2 | protein_coding | 0.0905478 | 1.91317383 |
| NFAT5 | protein_coding | 0.09051315 | 1.70605957 |
| KCMF1 | protein_coding | 0.09042504 | 1.65099259 |
| ACSL1 | protein_coding | 0.09026744 | 1.74074712 |
| ACIN1 | protein_coding | 0.09023145 | 3.11665882 |
| EXOC3 | protein_coding | 0.09013859 | 1.20834966 |
| STK11 | protein_coding | 0.09012505 | 0.77064725 |
| MAMDC2-AS | lncRNA | 0.09010704 | 0.20142689 |
| ZBTB24 | protein_coding | 0.09003645 | 0.99637382 |
| NR2C1 | protein_coding | 0.08993798 | 1.15310725 |
| PARL | protein_coding | 0.089891 | 0.64962799 |
| ZDHHC6 | protein_coding | 0.08973004 | 1.33748342 |
| SLC38A7 | protein_coding | 0.08968863 | 1.02069764 |

| | | | |
|---|---|---|---|
| STRBP | protein_coding | 0.08967076 | 2.0243721 |
| RPS6KB2 | protein_coding | 0.0896548 | 1.79027311 |
| ZNF628 | protein_coding | 0.08961213 | 0.34719426 |
| MGRN1 | protein_coding | 0.08948999 | 1.69403022 |
| ZCCHC3 | protein_coding | 0.08941412 | 1.70953095 |
| GIGYF2 | protein_coding | 0.08928313 | 2.36061415 |
| DERL2 | protein_coding | 0.08917095 | 0.66373102 |
| PCDH1 | protein_coding | 0.08912083 | 0.35434412 |
| RIC1 | protein_coding | 0.08910364 | 1.01278682 |
| SILC1 | lncRNA | 0.08907571 | 0.20783763 |
| FAM200B | protein_coding | 0.08893381 | 0.6244646 |
| LTBP4 | protein_coding | 0.08890931 | 1.30951272 |
| HARS1 | protein_coding | 0.08886545 | 2.24438751 |
| FAM50A | protein_coding | 0.08874548 | 1.88882899 |
| DBT | protein_coding | 0.08873015 | 1.35429005 |
| NUP98 | protein_coding | 0.08867179 | 2.56454949 |
| FASTKD3 | protein_coding | 0.08864564 | 0.6979048 |
| LINC01137 | lncRNA | 0.0885633 | 0.25667637 |
| C1orf50 | protein_coding | 0.08853779 | 0.70666498 |
| MT1F | protein_coding | 0.08851438 | 0.28932113 |
| TMEM138 | protein_coding | 0.08851146 | 0.99530397 |
| NEIL2 | protein_coding | 0.08845918 | 0.16148448 |
| NDUFB9 | protein_coding | 0.08840743 | 2.30424704 |
| RP11-973H7. | lncRNA | 0.08835973 | 0.19466797 |
| MBD2 | protein_coding | 0.08835473 | 1.44435651 |
| RAB25 | protein_coding | 0.08829615 | 1.28867999 |
| DDX31 | protein_coding | 0.08828173 | 1.02281338 |
| ZNF202 | protein_coding | 0.08827399 | 1.27080648 |
| AC009403.2 | lncRNA | 0.08815232 | 0.22348502 |
| ARL4A | protein_coding | 0.08813471 | 0.68714489 |
| ZNF605 | protein_coding | 0.0880677 | 1.08448582 |
| ATP11C | protein_coding | 0.0880495 | 1.98908194 |
| C12orf43 | protein_coding | 0.08794457 | 1.08174698 |
| CCDC28A | protein_coding | 0.08787908 | 0.39109791 |
| GFI1 | protein_coding | 0.08786415 | 2.54045225 |
| RSPRY1 | protein_coding | 0.08780509 | 2.11425908 |
| SEC22C | protein_coding | 0.0877537 | 1.29082586 |
| AC005519.4 | lncRNA | 0.08770783 | 0.18721092 |
| ZNF330 | protein_coding | 0.08765412 | 1.44116209 |
| ERAL1 | protein_coding | 0.0875724 | 1.90644591 |
| WDR35 | protein_coding | 0.08752251 | 1.27467177 |
| NEMF | protein_coding | 0.0874694 | 1.39803675 |
| CTD-2647L4. | lncRNA | 0.08739752 | 0.16619518 |
| WASL | protein_coding | 0.0873858 | 1.53441225 |
| PCSK7 | protein_coding | 0.08720723 | 1.9299232 |

| | | | |
|---|---|---|---|
| ZNF18 | protein_coding | 0.08705652 | 0.50107393 |
| RFT1 | protein_coding | 0.08704138 | 0.95339704 |
| GTF3C6 | protein_coding | 0.0870089 | 1.25918437 |
| CEP350 | protein_coding | 0.08698753 | 2.25109739 |
| NSUN2 | protein_coding | 0.0869595 | 2.5018781 |
| ATF2 | protein_coding | 0.08687403 | 1.5297684 |
| UBXN2B | protein_coding | 0.08680424 | 1.13343393 |
| TADA3 | protein_coding | 0.08679059 | 1.27655717 |
| EIF4G2 | protein_coding | 0.0866192 | 3.76055246 |
| LINC00342 | lncRNA | 0.08648812 | 1.10016885 |
| PRKAB1 | protein_coding | 0.08644161 | 1.6226404 |
| NINL | protein_coding | 0.08641626 | 0.96908778 |
| GSAP | protein_coding | 0.08640938 | 0.15999928 |
| PRMT5 | protein_coding | 0.08621958 | 2.77563733 |
| YARS1 | protein_coding | 0.08611111 | 3.66673256 |
| RP11-582E3. | lncRNA | 0.08605654 | 0.34920743 |
| CEP170 | protein_coding | 0.08605159 | 2.41525067 |
| COQ5 | protein_coding | 0.08603201 | 0.86245355 |
| ADORA1 | protein_coding | 0.0859922 | 0.66588487 |
| VPS50 | protein_coding | 0.08588453 | 1.23199513 |
| MAP2K5 | protein_coding | 0.08569625 | 1.2458383 |
| PRSS23 | protein_coding | 0.08561836 | 0.75331876 |
| PDSS2 | protein_coding | 0.08556432 | 0.66013504 |
| TWNK | protein_coding | 0.0855549 | 0.84400087 |
| ZNF790-AS1 | lncRNA | 0.08553507 | 0.34733344 |
| RBSN | protein_coding | 0.08552505 | 1.16289147 |
| TTLL1 | protein_coding | 0.08545814 | 0.49733578 |
| GFM1 | protein_coding | 0.08540191 | 1.90855517 |
| NFATC3 | protein_coding | 0.08535863 | 1.66779967 |
| HELLPAR | lncRNA | 0.08525775 | 0.24078491 |
| MTX2 | protein_coding | 0.08520173 | 1.77167199 |
| NVL | protein_coding | 0.08513688 | 1.51717358 |
| APPBP2 | protein_coding | 0.08512517 | 1.64846638 |
| LINC02482 | lncRNA | 0.08509954 | 0.29512505 |
| CILK1 | protein_coding | 0.08509672 | 1.07163054 |
| NOL7 | protein_coding | 0.08500582 | 1.39583207 |
| CISD3 | protein_coding | 0.08494453 | 0.95116137 |
| NDUFA13 | protein_coding | 0.08494176 | 0.18258635 |
| TRIM23 | protein_coding | 0.08478347 | 0.94125676 |
| TMEM132B | protein_coding | 0.08466796 | 0.13933782 |
| RP11-727A2: | lncRNA | 0.08464835 | 0.16454906 |
| ZNF764 | protein_coding | 0.08464275 | 0.753586 |
| MTMR3 | protein_coding | 0.08447829 | 1.91272142 |
| SIAH1 | protein_coding | 0.08444478 | 1.3244987 |
| ZNF791 | protein_coding | 0.08440148 | 1.36590137 |

| | | | |
|---|---|---|---|
| KRT18 | protein_coding | 0.08435533 | 2.06379198 |
| ZNF12 | protein_coding | 0.08422715 | 1.4446918 |
| XRCC1 | protein_coding | 0.08421423 | 1.2753729 |
| THBS1 | protein_coding | 0.08418446 | 0.80944856 |
| TOPORS | protein_coding | 0.08418197 | 1.46195663 |
| HSP90AB1 | protein_coding | 0.08408439 | 3.7586815 |
| GDAP2 | protein_coding | 0.08408316 | 0.7353868 |
| CSNK1G1 | protein_coding | 0.08407938 | 1.50759577 |
| PITPNB | protein_coding | 0.0840187 | 2.01464532 |
| CSNK2A2 | protein_coding | 0.0840125 | 1.86080987 |
| PANK3 | protein_coding | 0.08400161 | 1.19430098 |
| NBPF9 | protein_coding | 0.08397472 | 0.20374375 |
| WDR45 | protein_coding | 0.08391356 | 0.56735709 |
| DARS1 | protein_coding | 0.08391059 | 2.59427437 |
| RRP8 | protein_coding | 0.08379271 | 1.02767636 |
| USP33 | protein_coding | 0.08375868 | 1.95532128 |
| ARID5A | protein_coding | 0.08364562 | 0.43150966 |
| ARMT1 | protein_coding | 0.08352637 | 0.56456359 |
| MAP7D1 | protein_coding | 0.08348668 | 1.14144348 |
| MAP3K10 | protein_coding | 0.08348187 | 0.53479312 |
| LRRC8C | protein_coding | 0.08338912 | 1.00257479 |
| EEA1 | protein_coding | 0.08338696 | 0.76484801 |
| ALDH7A1 | protein_coding | 0.08333833 | 1.37548415 |
| PDCD7 | protein_coding | 0.08329261 | 1.14934733 |
| NR2F1 | protein_coding | 0.0832002 | 0.10429874 |
| GSR | protein_coding | 0.08314133 | 1.54898913 |
| NOL11 | protein_coding | 0.0830622 | 1.50394599 |
| OMA1 | protein_coding | 0.08304636 | 0.49097229 |
| RP11-1275H1 | lncRNA | 0.08292723 | 0.20567917 |
| NUP107 | protein_coding | 0.0828901 | 1.8343564 |
| EWSAT1 | lncRNA | 0.0828857 | 0.74546825 |
| ZNF747 | protein_coding | 0.08288186 | 0.75542395 |
| WDR70 | protein_coding | 0.08283738 | 1.29862199 |
| CDC27 | protein_coding | 0.08274678 | 1.96924897 |
| RP11-359B1 | lncRNA | 0.08270069 | 0.23693442 |
| WDPCP | protein_coding | 0.08257116 | 0.48464522 |
| EIF2B4 | protein_coding | 0.08249747 | 1.26027345 |
| YLPM1 | protein_coding | 0.08246958 | 2.8051531 |
| GSDME | protein_coding | 0.08235066 | 0.12209043 |
| C20orf27 | protein_coding | 0.08234669 | 1.37429123 |
| JMJD7-PLA2 | protein_coding | 0.08230407 | 0.31137836 |
| TOX4 | protein_coding | 0.08230185 | 1.89901165 |
| ZSCAN21 | protein_coding | 0.08225837 | 0.84294505 |
| COPS2 | protein_coding | 0.08225287 | 1.54228643 |
| GOSR2 | protein_coding | 0.08224023 | 1.20796044 |

| | | | |
|---|---|---|---|
| AUP1 | protein_coding | 0.08200024 | 2.72316438 |
| PCBD1 | protein_coding | 0.08198021 | 0.61590041 |
| TMEM38B | protein_coding | 0.08193158 | 1.72163016 |
| PRPSAP1 | protein_coding | 0.08191332 | 1.1842294 |
| TUBA3FP | lncRNA | 0.08190812 | 0.26016983 |
| TBC1D24 | protein_coding | 0.08187322 | 1.69540221 |
| ZC2HC1A | protein_coding | 0.0817063 | 0.27087414 |
| WASHC3 | protein_coding | 0.08163542 | 0.95091357 |
| AMBRA1 | protein_coding | 0.08136871 | 1.22634405 |
| GPR75 | protein_coding | 0.08136086 | 0.1984567 |
| PINX1 | protein_coding | 0.0813146 | 0.56160066 |
| TROAP | protein_coding | 0.08130492 | 1.81448628 |
| C17orf58 | protein_coding | 0.08121351 | 0.40755447 |
| TCF3 | protein_coding | 0.08120007 | 1.63123091 |
| DRG2 | protein_coding | 0.08116975 | 1.01783626 |
| RNF216 | protein_coding | 0.08113557 | 1.43456891 |
| CAD | protein_coding | 0.08110149 | 2.23420565 |
| ABRAXAS2 | protein_coding | 0.08109711 | 0.69725184 |
| SLC39A11 | protein_coding | 0.08104891 | 0.98802453 |
| CNOT10 | protein_coding | 0.0810426 | 1.050813 |
| JPX | lncRNA | 0.08102311 | 1.0381459 |
| LRP6 | protein_coding | 0.08097031 | 1.07412414 |
| TM9SF2 | protein_coding | 0.0809606 | 2.48047666 |
| ANGEL1 | protein_coding | 0.08094912 | 1.72498811 |
| PRR13 | protein_coding | 0.08084702 | 0.67769814 |
| UFM1 | protein_coding | 0.08079373 | 0.97745064 |
| EEF1A2 | protein_coding | 0.08074912 | 1.62436766 |
| SELENOI | protein_coding | 0.08068003 | 1.59528554 |
| IKZF5 | protein_coding | 0.08061132 | 0.61660772 |
| EXTL3 | protein_coding | 0.0805936 | 1.17861052 |
| MFSD4B | protein_coding | 0.08054441 | 0.52453287 |
| RP11-108M9 | lncRNA | 0.08050133 | 0.12175439 |
| RWDD1 | protein_coding | 0.0803827 | 0.92579101 |
| RAD52 | protein_coding | 0.08033259 | 0.66072451 |
| DNAH10OS | lncRNA | 0.08029625 | 0.31308043 |
| MARK3 | protein_coding | 0.08029026 | 2.6668411 |
| RABGGTA | protein_coding | 0.08015582 | 1.09111873 |
| EIF3C | protein_coding | 0.08006515 | 0.58271181 |
| ABHD6 | protein_coding | 0.08000771 | 0.62306981 |
| MRPL44 | protein_coding | 0.07999899 | 1.32156221 |
| PEX26 | protein_coding | 0.0799458 | 1.5518704 |
| LRRC14 | protein_coding | 0.07986791 | 0.98574574 |
| ATG7 | protein_coding | 0.07984387 | 0.65100756 |
| FAM53C | protein_coding | 0.07982245 | 1.25557319 |
| MTMR14 | protein_coding | 0.07971575 | 0.76014836 |

| | | | |
|---|---|---|---|
| PHF23 | protein_coding | 0.07965766 | 0.72238973 |
| P3H1 | protein_coding | 0.0795713 | 1.2417946 |
| BSN-DT | lncRNA | 0.07953391 | 0.10659762 |
| MBNL1-AS1 | lncRNA | 0.07949023 | 0.23776327 |
| SHC3 | protein_coding | 0.07942095 | 0.21803018 |
| ZNF644 | protein_coding | 0.079409 | 1.44169018 |
| FIP1L1 | protein_coding | 0.07939893 | 1.19810281 |
| SOX13 | protein_coding | 0.07939617 | 0.21956228 |
| TSC22D3 | protein_coding | 0.07931763 | 0.63724152 |
| PEX10 | protein_coding | 0.0792785 | 0.59799748 |
| TMEM230 | protein_coding | 0.07926005 | 1.79651969 |
| MARCHF6 | protein_coding | 0.07907602 | 2.24060469 |
| ZNF214 | protein_coding | 0.07898122 | 0.3509907 |
| USF3 | protein_coding | 0.07895965 | 0.61096579 |
| HMGXB4 | protein_coding | 0.07891955 | 1.73678734 |
| NOL9 | protein_coding | 0.07891791 | 0.75757058 |
| ATP13A2 | protein_coding | 0.07880141 | 1.12640048 |
| NME1 | protein_coding | 0.07880056 | 0.9378961 |
| FAM219B | protein_coding | 0.07878349 | 1.54209536 |
| DDRGK1 | protein_coding | 0.07874808 | 1.12715323 |
| UBE2E1 | protein_coding | 0.07873231 | 1.10632805 |
| TNRC6A | protein_coding | 0.07872989 | 1.72037197 |
| TRMT2B | protein_coding | 0.07871835 | 1.478704 |
| ETNK1 | protein_coding | 0.07860498 | 1.27926531 |
| MTRNR2L12 | protein_coding | 0.07860093 | 0.11785627 |
| TUBA1A | protein_coding | 0.07848454 | 3.1765667 |
| ABCG8 | protein_coding | 0.0784114 | 0.90475166 |
| ZFP82 | protein_coding | 0.07840287 | 0.92015195 |
| RP11-482M8 | lncRNA | 0.07839369 | 0.11551508 |
| C1orf43 | protein_coding | 0.07838443 | 2.92542747 |
| RP11-395N3. | lncRNA | 0.07835915 | 0.25192843 |
| AGO3 | protein_coding | 0.07835658 | 1.3286407 |
| MTFR1 | protein_coding | 0.07830153 | 0.49105746 |
| PQBP1 | protein_coding | 0.07823212 | 1.59371759 |
| XK | protein_coding | 0.07809476 | 0.54497436 |
| ARHGEF37 | protein_coding | 0.07805306 | 0.66671535 |
| PCDHGC3 | protein_coding | 0.07800687 | 0.20941452 |
| CNTROB | protein_coding | 0.07797074 | 1.17839049 |
| HSP90B1 | protein_coding | 0.07794737 | 3.27431645 |
| ABC12-4666 | lncRNA | 0.07791569 | 0.32056306 |
| CNNM4 | protein_coding | 0.07780435 | 0.95078406 |
| LRBA | protein_coding | 0.07775855 | 3.03899504 |
| CHCHD4 | protein_coding | 0.07772772 | 0.48811208 |
| LRRC40 | protein_coding | 0.07771744 | 1.14418206 |
| NKAPD1 | protein_coding | 0.07762014 | 0.9277306 |

| | | | |
|---|---|---|---|
| DNAJC15 | protein_coding | 0.0776004 | 0.64379053 |
| WWC3 | protein_coding | 0.07756949 | 0.63674591 |
| TMUB2 | protein_coding | 0.07752966 | 1.08217371 |
| ETNK2 | protein_coding | 0.07750252 | 0.98077618 |
| GATD3A | protein_coding | 0.07745178 | 0.31690878 |
| ZC3H6 | protein_coding | 0.07745092 | 0.66660349 |
| SNHG7 | lncRNA | 0.07742383 | 1.11540008 |
| RCN3 | protein_coding | 0.07724243 | 0.31410642 |
| ANKRD52 | protein_coding | 0.0771677 | 1.83745836 |
| MRPS9 | protein_coding | 0.07706479 | 1.61567918 |
| ABCF3 | protein_coding | 0.07701183 | 1.48956794 |
| ZNF875 | protein_coding | 0.07692769 | 1.3423277 |
| SLC35E2B | protein_coding | 0.07692247 | 1.2448266 |
| ARID2 | protein_coding | 0.07687575 | 1.90015181 |
| GPN1 | protein_coding | 0.07681284 | 1.01234783 |
| ZNF45 | protein_coding | 0.07679501 | 1.03888231 |
| CHIC1 | protein_coding | 0.07677432 | 0.36028256 |
| GNPTAB | protein_coding | 0.0766742 | 1.76476018 |
| INIP | protein_coding | 0.07663332 | 1.32554825 |
| CCT2 | protein_coding | 0.07645037 | 2.79323983 |
| CBL | protein_coding | 0.07643817 | 1.54759196 |
| COQ10B | protein_coding | 0.07642058 | 0.60888291 |
| CMTM1 | protein_coding | 0.0763648 | 0.3092681 |
| PIAS4 | protein_coding | 0.07634274 | 1.09709854 |
| TIPIN | protein_coding | 0.0763074 | 0.95779233 |
| FEM1A | protein_coding | 0.07607558 | 1.14119498 |
| RP11-260M2 | lncRNA | 0.07606602 | 0.10026762 |
| PAPOLG | protein_coding | 0.07605972 | 1.11190328 |
| CTA-390C10. | lncRNA | 0.07604671 | 0.15736491 |
| SGO1 | protein_coding | 0.07603529 | 0.56149435 |
| ITPR2 | protein_coding | 0.07602087 | 0.50162908 |
| RP11-795F1§ | protein_coding | 0.07601824 | 0.08264495 |
| C12orf66 | protein_coding | 0.07593502 | 0.57111637 |
| LRRC37A2 | protein_coding | 0.07582813 | 0.37737523 |
| FAM168A | protein_coding | 0.07560904 | 2.07284287 |
| ZNF597 | protein_coding | 0.07558533 | 0.5722749 |
| KIAA0753 | protein_coding | 0.0755042 | 0.53275404 |
| MRPL19 | protein_coding | 0.07543956 | 1.28345238 |
| MRPL46 | protein_coding | 0.07543635 | 0.29639929 |
| MAP4K3-DT | lncRNA | 0.0753942 | 0.34237146 |
| MYO7A | protein_coding | 0.07527482 | 0.15528664 |
| UBE2A | protein_coding | 0.07526054 | 1.31151835 |
| STYXL1 | protein_coding | 0.07509761 | 0.74851069 |
| TNFRSF12A | protein_coding | 0.07502573 | 0.4887902 |
| CCDC71L | protein_coding | 0.07488089 | 0.13176994 |

| | | | |
|---|---|---|---|
| KIZ | protein_coding | 0.07458339 | 0.69282042 |
| GPATCH1 | protein_coding | 0.07454274 | 0.70567027 |
| LRRFIP1 | protein_coding | 0.07446377 | 1.37679465 |
| RP11-680H2( | lncRNA | 0.07437388 | 0.16972483 |
| ACSS2 | protein_coding | 0.07436261 | 0.63700598 |
| MPC1 | protein_coding | 0.07435645 | 0.42292604 |
| USP19 | protein_coding | 0.07434435 | 1.40388456 |
| RELL2 | protein_coding | 0.07429633 | 0.4009771 |
| AC073043.1 | lncRNA | 0.07428667 | 0.10157407 |
| TPRG1L | protein_coding | 0.07415038 | 0.60386546 |
| DDX3X | protein_coding | 0.07412266 | 2.29831844 |
| STAM-AS1 | lncRNA | 0.0740939 | 0.09145652 |
| TBC1D16 | protein_coding | 0.0740226 | 1.90854098 |
| GASAL1 | lncRNA | 0.07401434 | 0.13699049 |
| RP11-156K23 | lncRNA | 0.07397271 | 0.16625784 |
| TMEM51 | protein_coding | 0.07381791 | 0.72440325 |
| LSM5 | protein_coding | 0.07377803 | 0.59844497 |
| THAP9 | protein_coding | 0.07377328 | 0.41486822 |
| SLC7A6OS | protein_coding | 0.07374671 | 0.99401734 |
| SRRM4 | protein_coding | 0.07365983 | 1.15954176 |
| BMPER | protein_coding | 0.07365567 | 0.13138544 |
| SPG7 | protein_coding | 0.07363802 | 1.49902037 |
| VPS37A | protein_coding | 0.07343518 | 0.80789799 |
| BMPR1A | protein_coding | 0.07328677 | 0.34630703 |
| NSMCE4A | protein_coding | 0.07321787 | 0.91260454 |
| ZNF536 | protein_coding | 0.07318984 | 0.7987995 |
| MKS1 | protein_coding | 0.073143 | 1.0079564 |
| PHF1 | protein_coding | 0.07311192 | 1.22380937 |
| RANGRF | protein_coding | 0.07310943 | 0.48608352 |
| RTCB | protein_coding | 0.0729845 | 1.7114899 |
| RP11-723O4. | lncRNA | 0.07293437 | 0.23054429 |
| RP11-384P7. | lncRNA | 0.07282829 | 0.2540652 |
| ABL2 | protein_coding | 0.07262608 | 1.1956893 |
| NSUN3 | protein_coding | 0.07257853 | 0.58247998 |
| TXNL4B | protein_coding | 0.07257575 | 1.18690277 |
| MGAT4B | protein_coding | 0.07253147 | 0.73043401 |
| RP11-53G8.3 | lncRNA | 0.07245954 | 0.09045464 |
| ARID1B | protein_coding | 0.07233627 | 1.26388863 |
| NDUFA5 | protein_coding | 0.07230624 | 0.84724839 |
| EMC2 | protein_coding | 0.07217404 | 0.67280355 |
| ERI1 | protein_coding | 0.07204897 | 0.65803026 |
| HCFC2 | protein_coding | 0.07201091 | 0.75166623 |
| SMARCE1 | protein_coding | 0.07195926 | 0.55097822 |
| SFXN2 | protein_coding | 0.07195777 | 0.57747511 |
| PUS1 | protein_coding | 0.07190612 | 0.78059877 |

| | | | |
|---|---|---|---|
| ZMYM2 | protein_coding | 0.07178371 | 1.87620943 |
| RRP7A | protein_coding | 0.07173933 | 1.42433428 |
| OSGIN2 | protein_coding | 0.0717276 | 0.76227579 |
| RNF7 | protein_coding | 0.07147883 | 0.97589503 |
| TASOR | protein_coding | 0.07123074 | 1.24293734 |
| HCCS | protein_coding | 0.07113142 | 0.70862166 |
| RDH14 | protein_coding | 0.07108275 | 0.66645847 |
| GATB | protein_coding | 0.07106039 | 0.73630694 |
| HELQ | protein_coding | 0.07105252 | 0.53904772 |
| ZNF71 | protein_coding | 0.0708873 | 0.92008309 |
| MEGF8 | protein_coding | 0.07088517 | 0.96657917 |
| HECTD2 | protein_coding | 0.07082204 | 0.73497082 |
| CTA-38K21.6 | lncRNA | 0.07079028 | 0.08277878 |
| TIPARP | protein_coding | 0.07075067 | 0.93130683 |
| ZNF142 | protein_coding | 0.07074776 | 1.33256063 |
| CAV3 | protein_coding | 0.0707453 | 0.40394705 |
| UBE2M | protein_coding | 0.07062489 | 1.38776721 |
| COL6A1 | protein_coding | 0.07054796 | 0.43388343 |
| NOP14 | protein_coding | 0.07041276 | 1.49663066 |
| TSPAN14 | protein_coding | 0.07038184 | 0.81629619 |
| KCNC3 | protein_coding | 0.07028608 | 0.54017735 |
| TRAPPC11 | protein_coding | 0.07017777 | 1.46013641 |
| ATG14 | protein_coding | 0.07010676 | 0.56913741 |
| RP5-827C21. | lncRNA | 0.07009334 | 0.0904547 |
| SETD9 | protein_coding | 0.07004029 | 0.25565652 |
| GAREM1 | protein_coding | 0.07002528 | 0.53541498 |
| G3BP1 | protein_coding | 0.07002501 | 1.74692335 |
| DIS3L2 | protein_coding | 0.06996239 | 0.81550421 |
| APIP | protein_coding | 0.06981423 | 0.92430677 |
| HSPB6 | protein_coding | 0.06949651 | 0.05772223 |
| ZNF268 | protein_coding | 0.06945496 | 1.14502793 |
| NR1D2 | protein_coding | 0.06937223 | 1.11079989 |
| ARNT | protein_coding | 0.06935008 | 1.16127862 |
| PNPLA3 | protein_coding | 0.06911986 | 0.12540659 |
| RSAD1 | protein_coding | 0.06909889 | 0.92842393 |
| TRAPPC2 | protein_coding | 0.06904561 | 0.60726739 |
| TTC23 | protein_coding | 0.06898776 | 0.38337819 |
| NADK2 | protein_coding | 0.06894278 | 0.85666326 |
| COX4I1 | protein_coding | 0.06887574 | 2.51282515 |
| ARL6IP5 | protein_coding | 0.06883362 | 0.54031512 |
| KIAA0930 | protein_coding | 0.06871591 | 0.97894005 |
| ETF1 | protein_coding | 0.06868435 | 2.15150306 |
| CLOCK | protein_coding | 0.06867613 | 0.62697093 |
| GTPBP1 | protein_coding | 0.06866598 | 2.08966596 |
| EXOC4 | protein_coding | 0.06866464 | 1.55582275 |

| | | | |
|---|---|---|---|
| RP5-1074L1.4 | lncRNA | 0.06863776 | 0.23983892 |
| CIAO3 | protein_coding | 0.06859704 | 0.87091849 |
| VPS9D1 | protein_coding | 0.06855453 | 0.42240221 |
| CA11 | protein_coding | 0.06852547 | 1.19457276 |
| UGGT1 | protein_coding | 0.0684755 | 1.75847242 |
| TBL1X | protein_coding | 0.0684518 | 1.21467569 |
| ACACB | protein_coding | 0.06834692 | 0.54223047 |
| RP11-379F4. | lncRNA | 0.06834001 | 0.08153542 |
| MIGA2 | protein_coding | 0.06831637 | 0.90855396 |
| ZNF682 | protein_coding | 0.06829277 | 0.2770042 |
| ZNF746 | protein_coding | 0.0682923 | 0.69765267 |
| G6PD | protein_coding | 0.06827492 | 1.3873506 |
| EIF1 | protein_coding | 0.06826964 | 2.60963452 |
| RP11-262H14 | lncRNA | 0.06826573 | 0.06961941 |
| ZNF213 | protein_coding | 0.0682433 | 0.62307175 |
| PPP1R11 | protein_coding | 0.06821806 | 1.30021007 |
| THAP9-AS1 | lncRNA | 0.06817854 | 1.20577636 |
| NPY | protein_coding | 0.06815037 | 0.61264241 |
| TTLL11 | protein_coding | 0.06812838 | 0.4530463 |
| ZBTB49 | protein_coding | 0.06803851 | 0.37233589 |
| SLIRP | protein_coding | 0.06803563 | 1.51323015 |
| SH3BP2 | protein_coding | 0.06800003 | 1.11264725 |
| C2CD5 | protein_coding | 0.06789812 | 1.31918155 |
| GLRX3 | protein_coding | 0.06776201 | 0.82918836 |
| CNEP1R1 | protein_coding | 0.06771185 | 0.44536258 |
| DMWD | protein_coding | 0.06768072 | 0.89872496 |
| HELZ | protein_coding | 0.06764992 | 1.54622736 |
| ZRANB2 | protein_coding | 0.06760833 | 0.98010394 |
| MACIR | protein_coding | 0.06756597 | 0.64746441 |
| ATAD3B | protein_coding | 0.0675206 | 0.64959218 |
| ASH1L | protein_coding | 0.06751437 | 1.79432832 |
| TUBD1 | protein_coding | 0.06745203 | 0.53507716 |
| KDM2B | protein_coding | 0.06741696 | 1.68112663 |
| IFT81 | protein_coding | 0.06730866 | 0.49273686 |
| TDRKH | protein_coding | 0.06729635 | 0.69391543 |
| TRAPPC14 | protein_coding | 0.0672864 | 0.4679642 |
| CBWD1 | protein_coding | 0.06727201 | 0.38472282 |
| RIPK4 | protein_coding | 0.06722351 | 0.52631474 |
| LRRC14B | protein_coding | 0.06719844 | 0.17379862 |
| MTFMT | protein_coding | 0.06706414 | 0.5753362 |
| RASA4B | protein_coding | 0.06704881 | 0.34970285 |
| MAP3K7 | protein_coding | 0.06703655 | 0.69151633 |
| CTNS | protein_coding | 0.0669606 | 0.36572879 |
| PAIP1 | protein_coding | 0.06691465 | 1.00987441 |
| C2orf42 | protein_coding | 0.06688935 | 0.47402659 |

| | | | |
|---|---|---|---|
| SKIV2L | protein_coding | 0.06682232 | 1.17544703 |
| EIF5A | protein_coding | 0.06674438 | 1.76072471 |
| SLC37A3 | protein_coding | 0.06669045 | 1.07378907 |
| NECAP1 | protein_coding | 0.06664958 | 1.31334197 |
| PCP4 | protein_coding | 0.06657466 | 1.34254921 |
| PM20D2 | protein_coding | 0.06639095 | 1.07441204 |
| RCL1 | protein_coding | 0.06639058 | 0.99428245 |
| RPL12 | protein_coding | 0.06636596 | 2.63597007 |
| DIS3L | protein_coding | 0.0663429 | 1.10405549 |
| CYP7B1 | protein_coding | 0.06626488 | 0.34289499 |
| ZNF784 | protein_coding | 0.06624143 | 0.40130078 |
| ANKS6 | protein_coding | 0.06611591 | 1.1877741 |
| AKAP17A | protein_coding | 0.066071 | 0.72372417 |
| ZFAND2A-DT | lncRNA | 0.06607049 | 0.43841431 |
| LMBR1L | protein_coding | 0.06585754 | 0.66114001 |
| TSPAN13 | protein_coding | 0.06584835 | 0.81388946 |
| CENPL | protein_coding | 0.06581202 | 0.85294925 |
| H3C3 | protein_coding | 0.06577798 | 0.06585221 |
| FAM160A1 | protein_coding | 0.06575278 | 0.92645761 |
| TTC7A | protein_coding | 0.06565721 | 0.79902231 |
| SELENOW | protein_coding | 0.06560846 | 0.74328223 |
| HDAC8 | protein_coding | 0.06553017 | 0.23616288 |
| NEURL4 | protein_coding | 0.06548221 | 0.91610038 |
| RBM17 | protein_coding | 0.06545271 | 1.10958111 |
| TOR1B | protein_coding | 0.06543775 | 1.01878264 |
| COPA | protein_coding | 0.06540903 | 2.0911923 |
| ATP2A1 | protein_coding | 0.06539408 | 0.15403592 |
| SURF2 | protein_coding | 0.06535308 | 0.65753356 |
| PMS1 | protein_coding | 0.06532208 | 0.51629226 |
| TMEM63C | protein_coding | 0.06516588 | 1.59660933 |
| KAT6A | protein_coding | 0.06515094 | 1.36919592 |
| ZCCHC9 | protein_coding | 0.06508653 | 0.82024796 |
| TRRAP | protein_coding | 0.06504607 | 1.9035428 |
| CTD-2589H1 | lncRNA | 0.06496496 | 0.1084119 |
| BACH1-IT1 | lncRNA | 0.06495407 | 0.06131902 |
| B4GALT3 | protein_coding | 0.06491504 | 1.4258023 |
| CDK5RAP3 | protein_coding | 0.06484692 | 0.30474955 |
| UFSP2 | protein_coding | 0.06465421 | 0.46747007 |
| RBBP6 | protein_coding | 0.06459603 | 1.53562085 |
| CTNNB1 | protein_coding | 0.06456192 | 1.62996525 |
| FNDC3A | protein_coding | 0.06455829 | 0.77270317 |
| THAP4 | protein_coding | 0.06455753 | 0.73100131 |
| MAP2K1 | protein_coding | 0.06453616 | 1.23765947 |
| WAPL | protein_coding | 0.06450232 | 1.14031459 |
| GTPBP6 | protein_coding | 0.06449163 | 0.53412284 |

| | | | |
|---|---|---|---|
| ARMC8 | protein_coding | 0.0644336 | 0.97069561 |
| RNF24 | protein_coding | 0.06442479 | 0.93346521 |
| MECP2 | protein_coding | 0.0644193 | 1.53126736 |
| RIMS2 | protein_coding | 0.06441461 | 1.56100811 |
| DUSP11 | protein_coding | 0.06440282 | 0.47388385 |
| MLX | protein_coding | 0.06436661 | 1.64292918 |
| PNPO | protein_coding | 0.06429844 | 0.77145018 |
| ELAPOR1 | protein_coding | 0.06404139 | 0.07483012 |
| RP11-368L12 | lncRNA | 0.06396129 | 0.06706334 |
| AP3M2 | protein_coding | 0.0639498 | 0.84978253 |
| UAP1 | protein_coding | 0.06390126 | 0.78642405 |
| GPR158 | protein_coding | 0.06381444 | 0.50671132 |
| POLL | protein_coding | 0.06378407 | 0.54388967 |
| PTEN | protein_coding | 0.06374707 | 0.84134478 |
| ANGPTL6 | protein_coding | 0.0636091 | 0.10401782 |
| CMAS | protein_coding | 0.06357225 | 1.10941357 |
| CIART | protein_coding | 0.06351281 | 0.39771319 |
| GCN1 | protein_coding | 0.06347685 | 1.78909712 |
| DDX18 | protein_coding | 0.06338747 | 1.67014338 |
| SNAPC3 | protein_coding | 0.06329938 | 1.16886725 |
| LINC02408 | lncRNA | 0.06328727 | 0.0667805 |
| TNXB | protein_coding | 0.06316388 | 0.09529859 |
| ATP6V0E1 | protein_coding | 0.0630921 | 0.90431379 |
| PSMF1 | protein_coding | 0.06309156 | 1.59942558 |
| TUBGCP3 | protein_coding | 0.06306884 | 1.39298141 |
| KDM5C | protein_coding | 0.06302805 | 1.79992968 |
| CLASP2 | protein_coding | 0.06302695 | 0.62128877 |
| FKRP | protein_coding | 0.06298647 | 0.52100851 |
| CELF3 | protein_coding | 0.06295143 | 0.65434714 |
| AP2A2 | protein_coding | 0.06293 | 1.1679946 |
| LUC7L2 | protein_coding | 0.06287626 | 0.72031325 |
| TMEM129 | protein_coding | 0.06285426 | 0.87924763 |
| ZNF770 | protein_coding | 0.06283332 | 1.24913524 |
| RHOT1 | protein_coding | 0.06276813 | 1.35316426 |
| ZNF438 | protein_coding | 0.06273568 | 0.20357252 |
| SNW1 | protein_coding | 0.06256208 | 1.5688392 |
| RP11-95O2.1 | lncRNA | 0.06253455 | 0.12713394 |
| SPATA5L1 | protein_coding | 0.06247044 | 0.41123627 |
| SNAP47 | protein_coding | 0.06241091 | 0.85203772 |
| DEFB1 | protein_coding | 0.06238012 | 0.25770106 |
| DPYD | protein_coding | 0.06234187 | 0.78435112 |
| MRPS35 | protein_coding | 0.06233202 | 0.97397596 |
| MFN2 | protein_coding | 0.06231357 | 1.42680766 |
| NEFH | protein_coding | 0.0623064 | 1.49508434 |
| MBD4 | protein_coding | 0.06224396 | 1.18904473 |

| | | | |
|---|---|---|---|
| SUSD6 | protein_coding | 0.06223307 | 0.99823212 |
| ZSWIM8 | protein_coding | 0.06221942 | 0.66250009 |
| DNAJC11 | protein_coding | 0.06205629 | 1.12826779 |
| MT-ND1 | protein_coding | 0.06194831 | 1.83037414 |
| TMEM87A | protein_coding | 0.06192698 | 1.40308305 |
| RP11-672L1C | lncRNA | 0.06164205 | 0.08528341 |
| GMPR2 | protein_coding | 0.06163962 | 0.67798913 |
| STOML2 | protein_coding | 0.06162647 | 1.22796813 |
| TMEM80 | protein_coding | 0.06159291 | 0.59119957 |
| ZNF155 | protein_coding | 0.0615921 | 0.34951473 |
| FAR2 | protein_coding | 0.06154258 | 0.44652285 |
| MNX1 | protein_coding | 0.06149308 | 0.33349447 |
| BET1 | protein_coding | 0.06144342 | 0.28348124 |
| BAK1 | protein_coding | 0.06141367 | 0.65876053 |
| MSS51 | protein_coding | 0.06133697 | 0.08276084 |
| PCDHGA5 | protein_coding | 0.06127051 | 0.1111111 |
| SNX9 | protein_coding | 0.061267 | 0.87227364 |
| HMGCR | protein_coding | 0.061214 | 1.59448229 |
| AC084149.2 | lncRNA | 0.06117475 | 0.05760593 |
| SYT14 | protein_coding | 0.06107021 | 0.33507542 |
| CTPS2 | protein_coding | 0.06104087 | 0.77037934 |
| OXSM | protein_coding | 0.0609505 | 0.32253543 |
| ZSWIM7 | protein_coding | 0.06092325 | 0.34138091 |
| BFAR | protein_coding | 0.06089441 | 0.87269965 |
| PCBP3-OT1 | lncRNA | 0.06074195 | 0.11724583 |
| RP13-968A2. | lncRNA | 0.06071134 | 0.05719653 |
| ANAPC2 | protein_coding | 0.06070729 | 0.99800203 |
| ZNF337 | protein_coding | 0.060707 | 0.73934358 |
| TSC22D2 | protein_coding | 0.06066562 | 0.432628 |
| RHOBTB3 | protein_coding | 0.06062158 | 0.82258874 |
| PNISR | protein_coding | 0.06060123 | 1.08329296 |
| SLC35B1 | protein_coding | 0.06052662 | 1.00929988 |
| NARS2 | protein_coding | 0.06043167 | 0.95094345 |
| MFAP4 | protein_coding | 0.06039658 | 1.27198123 |
| BLACAT1 | protein_coding | 0.06021937 | 0.13416229 |
| CFAP97 | protein_coding | 0.06020603 | 1.06150292 |
| XXbac-B444F | lncRNA | 0.06011819 | 0.10124068 |
| CDYL | protein_coding | 0.06005728 | 0.63492002 |
| CIT | protein_coding | 0.05999442 | 0.99331259 |
| MED8 | protein_coding | 0.05997115 | 0.91162959 |
| DNASE1L1 | protein_coding | 0.05994828 | 0.7039373 |
| PSTPIP1 | protein_coding | 0.05984867 | 0.44231338 |
| OCRL | protein_coding | 0.05982646 | 1.15346194 |
| PRAF2 | protein_coding | 0.05980212 | 0.24555635 |
| HARS2 | protein_coding | 0.05978646 | 0.87453381 |

| | | | |
|---|---|---|---|
| PNKP | protein_coding | 0.05971444 | 0.9795386 |
| CLBA1 | protein_coding | 0.05967116 | 0.30057412 |
| COPS7B | protein_coding | 0.05963439 | 0.85013921 |
| FAM155B | protein_coding | 0.05963288 | 0.57771674 |
| RBM12B | protein_coding | 0.05960776 | 1.11966891 |
| KANSL1 | protein_coding | 0.05959873 | 1.20479291 |
| GAR1 | protein_coding | 0.05949004 | 0.58136684 |
| POU2F1 | protein_coding | 0.05943297 | 0.94300289 |
| TXNDC11 | protein_coding | 0.05938171 | 0.63384393 |
| EZH2 | protein_coding | 0.05920173 | 1.41847263 |
| NELFE | protein_coding | 0.05913553 | 1.2853047 |
| RP11-867G2: | lncRNA | 0.05913381 | 0.05231584 |
| SLA | protein_coding | 0.0590468 | 0.31624245 |
| RABEP1 | protein_coding | 0.05903389 | 1.07645249 |
| STEAP3 | protein_coding | 0.05881111 | 1.29110133 |
| KRBOX4 | protein_coding | 0.05876492 | 0.38750773 |
| TRIM9 | protein_coding | 0.05865156 | 0.73404309 |
| SELENOK | protein_coding | 0.05862088 | 0.52207035 |
| IMPA1 | protein_coding | 0.05861647 | 0.65119339 |
| YTHDF3 | protein_coding | 0.05859011 | 0.77446978 |
| NAXD | protein_coding | 0.05855577 | 0.86064342 |
| C5orf66 | lncRNA | 0.05853194 | 0.09865665 |
| TRMT61B | protein_coding | 0.05841672 | 0.5003692 |
| C17orf80 | protein_coding | 0.05837273 | 0.81861913 |
| ZKSCAN8 | protein_coding | 0.0583666 | 0.90100154 |
| RFK | protein_coding | 0.05833674 | 0.51570207 |
| FGGY | protein_coding | 0.05830582 | 0.24859172 |
| IPO11 | protein_coding | 0.0582565 | 0.66695869 |
| ZNF691 | protein_coding | 0.05825366 | 0.5258431 |
| URI1 | protein_coding | 0.0582388 | 1.33413427 |
| MIEF1 | protein_coding | 0.05815268 | 1.44637992 |
| COA6 | protein_coding | 0.05802235 | 0.60242906 |
| CTD-2587H2 | lncRNA | 0.0579856 | 0.15794682 |
| LIME1 | protein_coding | 0.05792036 | 0.16518205 |
| AP001626.2 | lncRNA | 0.05791526 | 0.10433506 |
| STIMATE | protein_coding | 0.05791436 | 0.08274856 |
| ZFP37 | protein_coding | 0.05791369 | 0.08516364 |
| FMNL1-DT | lncRNA | 0.05789058 | 0.05749813 |
| ZNF717 | protein_coding | 0.05783923 | 0.35724507 |
| DUS2 | protein_coding | 0.05779575 | 0.37056815 |
| ATG101 | protein_coding | 0.05779363 | 0.71217436 |
| DAAM1 | protein_coding | 0.05762628 | 1.56026098 |
| CRY1 | protein_coding | 0.05759799 | 0.75886855 |
| CH507-9B2.5 | protein_coding | 0.05759438 | 0.20864491 |
| CENPO | protein_coding | 0.05757245 | 0.83838121 |

| | | | |
|---|---|---|---|
| GCC2 | protein_coding | 0.05756294 | 0.72532478 |
| U2SURP | protein_coding | 0.05753324 | 1.51161923 |
| TFCP2 | protein_coding | 0.05748313 | 1.04000161 |
| THEM4 | protein_coding | 0.05735195 | 0.33588148 |
| ARHGAP35 | protein_coding | 0.05735141 | 1.82021617 |
| POU3F2 | protein_coding | 0.05731432 | 0.25325476 |
| ZNF736 | protein_coding | 0.05726905 | 0.60736767 |
| CUL2 | protein_coding | 0.05719797 | 0.84825541 |
| AIDA | protein_coding | 0.05711783 | 0.66461345 |
| USP54 | protein_coding | 0.05702788 | 0.50116386 |
| AUH | protein_coding | 0.05700473 | 0.30626829 |
| TRMT1 | protein_coding | 0.05688391 | 0.63321601 |
| IFRD2 | protein_coding | 0.05683874 | 0.65602691 |
| MYNN | protein_coding | 0.05680892 | 0.70254031 |
| TMCC1 | protein_coding | 0.05678716 | 0.52245292 |
| AKIRIN1 | protein_coding | 0.05675765 | 1.1782594 |
| TRAPPC4 | protein_coding | 0.05672112 | 0.85622872 |
| ZDHHC15 | protein_coding | 0.05669378 | 0.59704257 |
| ANKRD36C | protein_coding | 0.05667257 | 0.42040935 |
| BCAT2 | protein_coding | 0.05642545 | 0.79866092 |
| ACOX1 | protein_coding | 0.05642194 | 0.8944686 |
| AUNIP | protein_coding | 0.0564059 | 0.3993551 |
| DMXL2 | protein_coding | 0.05632 | 1.65773998 |
| LETM2 | protein_coding | 0.05624341 | 0.25756242 |
| ZNF35 | protein_coding | 0.05624317 | 0.27116442 |
| NAA10 | protein_coding | 0.05623345 | 0.52602166 |
| SNAPC1 | protein_coding | 0.05615949 | 0.23693607 |
| NUP37 | protein_coding | 0.05613209 | 0.70419011 |
| DMXL1 | protein_coding | 0.0560675 | 1.04159367 |
| C1orf35 | protein_coding | 0.05605281 | 0.26798032 |
| RAF1 | protein_coding | 0.05603775 | 1.24590453 |
| BABAM2 | protein_coding | 0.05596603 | 0.46271365 |
| ADNP2 | protein_coding | 0.05587116 | 0.86310005 |
| MVB12B | protein_coding | 0.05585044 | 0.39167292 |
| MRPL13 | protein_coding | 0.05584666 | 0.65057376 |
| DHX38 | protein_coding | 0.05580345 | 1.06576436 |
| RPS6KA4 | protein_coding | 0.05578259 | 0.37222007 |
| ZNF197 | protein_coding | 0.05570341 | 0.49802553 |
| EXOC1 | protein_coding | 0.05562259 | 0.5162391 |
| SNX4 | protein_coding | 0.05551246 | 0.77048413 |
| XPA | protein_coding | 0.05548816 | 0.35011895 |
| RP4-622L5.7 | lncRNA | 0.05547438 | 0.04677664 |
| UFSP1 | protein_coding | 0.05529446 | 0.0759232 |
| TASP1 | protein_coding | 0.05529436 | 0.34692308 |
| RP3-508I15.9 | lncRNA | 0.05528842 | 0.21389693 |

| | | | |
|---|---|---|---|
| R3HDM1 | protein_coding | 0.05525099 | 1.1851649 |
| PANK2 | protein_coding | 0.05524431 | 1.02278678 |
| RP11-415J8.: | lncRNA | 0.05513363 | 0.0482238 |
| CCDC102B | protein_coding | 0.0550436 | 0.25422413 |
| DNAJC17 | protein_coding | 0.05479403 | 0.42904607 |
| ZFPL1 | protein_coding | 0.05478904 | 0.43556402 |
| GLE1 | protein_coding | 0.05470675 | 1.07255162 |
| WDR17 | protein_coding | 0.05465969 | 0.57454517 |
| ATG16L1 | protein_coding | 0.05454273 | 0.64493676 |
| LGALS7B | protein_coding | 0.05451017 | 0.55752539 |
| SDHB | protein_coding | 0.05448836 | 1.32895849 |
| CCAR1 | protein_coding | 0.05445391 | 1.10905768 |
| PNN | protein_coding | 0.05445156 | 1.58685153 |
| CEP57 | protein_coding | 0.05442085 | 0.9892497 |
| ZNF775 | protein_coding | 0.05442046 | 0.2722142 |
| RP11-435O5. | lncRNA | 0.05435017 | 0.05324233 |
| ZSWIM1 | protein_coding | 0.05431141 | 0.46564357 |
| RFX2 | protein_coding | 0.05428047 | 0.29158118 |
| SHLD2 | protein_coding | 0.05423428 | 0.52966311 |
| ZNF586 | protein_coding | 0.05419266 | 0.42530743 |
| TPD52L2 | protein_coding | 0.05418643 | 1.34887339 |
| MRPL42 | protein_coding | 0.05414071 | 0.73573759 |
| CKMT1B | protein_coding | 0.0541006 | 0.38820138 |
| EIF2S1 | protein_coding | 0.05408791 | 1.30490715 |
| PBX4 | protein_coding | 0.0540135 | 0.24921934 |
| TIAL1 | protein_coding | 0.05399888 | 0.8434672 |
| SLC35C2 | protein_coding | 0.05398202 | 0.92058539 |
| ATP1B1 | protein_coding | 0.05395318 | 1.34717085 |
| GNG7 | protein_coding | 0.05395049 | 0.16779236 |
| MZF1-AS1 | lncRNA | 0.05390443 | 0.20421301 |
| FOXK1 | protein_coding | 0.05386491 | 0.61151262 |
| AJ011931.1 | lncRNA | 0.05386226 | 0.04471213 |
| RRN3 | protein_coding | 0.05386088 | 0.99486013 |
| XXbac-BPG2! | lncRNA | 0.05369064 | 0.14427744 |
| TIGD1 | protein_coding | 0.05368939 | 0.34539875 |
| L3MBTL4 | protein_coding | 0.05352723 | 0.63129312 |
| SHARPIN | protein_coding | 0.05346665 | 0.4511218 |
| MRPL14 | protein_coding | 0.05346618 | 0.79595799 |
| GALE | protein_coding | 0.05345597 | 0.39762053 |
| ULK1 | protein_coding | 0.05343969 | 0.82215242 |
| CEP72 | protein_coding | 0.05341785 | 0.50282521 |
| REEP3 | protein_coding | 0.05320785 | 0.42901252 |
| TOB2 | protein_coding | 0.05310459 | 0.89144877 |
| WDR43 | protein_coding | 0.05308111 | 0.96019413 |
| FNIP2 | protein_coding | 0.05306643 | 0.91344152 |

| | | | |
|---|---|---|---|
| LAMTOR5-AS | lncRNA | 0.05291214 | 0.15339896 |
| ESRRA | protein_coding | 0.0529047 | 0.51098652 |
| EXOC8 | protein_coding | 0.05280994 | 0.61688609 |
| USP49 | protein_coding | 0.05260522 | 0.64761971 |
| JAGN1 | protein_coding | 0.05258904 | 0.4743693 |
| IFI27L1 | protein_coding | 0.05257273 | 0.50312378 |
| VPS39 | protein_coding | 0.05254803 | 1.14544037 |
| MT-CO1 | protein_coding | 0.05249886 | 2.2736917 |
| IFT74 | protein_coding | 0.05247837 | 0.26237698 |
| UBE2L3 | protein_coding | 0.05239846 | 1.04774204 |
| SREK1 | protein_coding | 0.05227056 | 0.86929525 |
| UQCRFS1 | protein_coding | 0.05214724 | 0.6870677 |
| PPIL4 | protein_coding | 0.05212744 | 0.43732785 |
| JRKL | protein_coding | 0.0521092 | 0.39379991 |
| KLF5 | protein_coding | 0.05205402 | 0.30978335 |
| SLC43A2 | protein_coding | 0.05200303 | 0.44088989 |
| MIER1 | protein_coding | 0.05196783 | 0.64874717 |
| ATXN2 | protein_coding | 0.05195964 | 0.66074791 |
| DTNB | protein_coding | 0.05193146 | 0.57157763 |
| ZNF431 | protein_coding | 0.05179623 | 0.68420269 |
| WDR48 | protein_coding | 0.05170327 | 0.61371544 |
| ZNF740 | protein_coding | 0.05160393 | 0.97299914 |
| HINT3 | protein_coding | 0.05155263 | 0.57676505 |
| RAB11FIP4 | protein_coding | 0.0515091 | 0.78204521 |
| MBIP | protein_coding | 0.05144149 | 0.38617305 |
| SNX1 | protein_coding | 0.05135555 | 1.12817621 |
| RP11-363N2 | lncRNA | 0.05122329 | 0.04415985 |
| MTM1 | protein_coding | 0.05118761 | 0.19126322 |
| GPATCH8 | protein_coding | 0.05118471 | 0.90140731 |
| PPP1R35 | protein_coding | 0.0511429 | 0.28394463 |
| FAF1 | protein_coding | 0.05113185 | 1.3474564 |
| TRAF3IP2-AS | lncRNA | 0.05111412 | 0.23487297 |
| GFPT2 | protein_coding | 0.05093564 | 0.10065445 |
| RP5-1021I20 | lncRNA | 0.05093076 | 0.04671406 |
| MRPL10 | protein_coding | 0.05083992 | 0.70730275 |
| AP4M1 | protein_coding | 0.05073636 | 0.62682864 |
| FBXW7 | protein_coding | 0.05062305 | 0.59239325 |
| RTL10 | protein_coding | 0.05061513 | 0.84936472 |
| NFYB | protein_coding | 0.05049331 | 0.69782757 |
| STX6 | protein_coding | 0.05045266 | 0.604915 |
| ZNF514 | protein_coding | 0.05041606 | 0.51257242 |
| TMEM50A | protein_coding | 0.05035837 | 0.95786747 |
| VPS18 | protein_coding | 0.05029877 | 0.48880165 |
| AC002310.7 | lncRNA | 0.05029545 | 0.06143892 |
| STK24 | protein_coding | 0.0502721 | 0.83231005 |

| | | | |
|---|---|---|---|
| CHASERR | lncRNA | 0.05017215 | 0.5492223 |
| SMYD2 | protein_coding | 0.05003895 | 0.67380572 |
| DHRS4L2 | protein_coding | 0.05000838 | 0.32680891 |
| NIPAL3 | protein_coding | 0.04999764 | 0.92571246 |
| TTL | protein_coding | 0.04995308 | 1.20613106 |
| DLG5 | protein_coding | 0.04992931 | 0.95989749 |
| POFUT2 | protein_coding | 0.04992221 | 0.46911685 |
| OCIAD1 | protein_coding | 0.04974156 | 1.28232074 |
| PPIG | protein_coding | 0.04962701 | 0.83300701 |
| TMEM30A | protein_coding | 0.04952391 | 1.0521354 |
| IWS1 | protein_coding | 0.04951342 | 1.10506497 |
| CAPZA2 | protein_coding | 0.04947649 | 0.72614839 |
| FBXO44 | protein_coding | 0.0494757 | 0.30830895 |
| TPM2 | protein_coding | 0.04938711 | 0.05600297 |
| CHMP6 | protein_coding | 0.04922411 | 0.16394256 |
| MCM9 | protein_coding | 0.04919957 | 0.38980726 |
| PUS7L | protein_coding | 0.04919082 | 0.66983297 |
| LYRM2 | protein_coding | 0.04918879 | 0.89052533 |
| CEP68 | protein_coding | 0.04912151 | 0.54190913 |
| RPE | protein_coding | 0.0490337 | 0.5083903 |
| GNPDA2 | protein_coding | 0.04899085 | 0.37832514 |
| ZNF483 | protein_coding | 0.04895909 | 0.24776969 |
| TMEM185B | protein_coding | 0.04894982 | 1.0930495 |
| MAP1LC3B | protein_coding | 0.04893709 | 0.79774839 |
| FAM66C | lncRNA | 0.04892227 | 0.06549227 |
| MRPL34 | protein_coding | 0.04884658 | 0.4837006 |
| ALOX12B | protein_coding | 0.04880478 | 0.05953316 |
| GOT1 | protein_coding | 0.04879731 | 0.95312104 |
| LYSMD3 | protein_coding | 0.04874155 | 0.39159019 |
| TSG101 | protein_coding | 0.04874091 | 0.76057729 |
| RBM25 | protein_coding | 0.04866676 | 1.05377422 |
| ESRP1 | protein_coding | 0.04860968 | 1.42926486 |
| RP11-363E6. | lncRNA | 0.04860614 | 0.0813174 |
| MSTO1 | protein_coding | 0.04860566 | 0.3257341 |
| BPNT1 | protein_coding | 0.04847612 | 0.62041666 |
| MED28 | protein_coding | 0.04838832 | 1.01845115 |
| DNAJA2 | protein_coding | 0.04824428 | 0.73023259 |
| PLPP2 | protein_coding | 0.04824121 | 0.44168891 |
| PTBP2 | protein_coding | 0.04820696 | 0.43156404 |
| ARL8B | protein_coding | 0.04820553 | 0.66859382 |
| MRPS30 | protein_coding | 0.04813785 | 0.58555528 |
| DCUN1D4 | protein_coding | 0.04812749 | 0.48206202 |
| HIP1R | protein_coding | 0.04805237 | 0.34567797 |
| MRPS25 | protein_coding | 0.04791375 | 0.52797178 |
| UQCC1 | protein_coding | 0.04786859 | 0.90197894 |

| | | | |
|---|---|---|---|
| KBTBD12 | protein_coding | 0.04773857 | 0.35022945 |
| MAILR | lncRNA | 0.04771182 | 0.13323252 |
| PDCL3 | protein_coding | 0.04759742 | 0.46350837 |
| RBM45 | protein_coding | 0.04756169 | 0.33784661 |
| CLDN5 | protein_coding | 0.04752039 | 0.14269972 |
| MAPKAPK2 | protein_coding | 0.04749185 | 0.86192885 |
| DUS1L | protein_coding | 0.04747402 | 0.81705755 |
| MROH8 | protein_coding | 0.04745227 | 0.05486452 |
| MED27 | protein_coding | 0.04743017 | 0.54920157 |
| TAF5 | protein_coding | 0.04739498 | 0.37234391 |
| SP2-AS1 | lncRNA | 0.04731139 | 0.13636494 |
| TUG1 | protein_coding | 0.0472876 | 1.39981078 |
| RNF19A | protein_coding | 0.04727607 | 0.7364613 |
| LIG3 | protein_coding | 0.04721511 | 1.04152578 |
| TINF2 | protein_coding | 0.04718788 | 0.79565231 |
| SUB1 | protein_coding | 0.04715916 | 1.22295578 |
| HEY1 | protein_coding | 0.04709773 | 0.1912108 |
| NHLRC2 | protein_coding | 0.0469968 | 0.42847988 |
| BOD1L1 | protein_coding | 0.04698407 | 0.88122281 |
| RP11-727F15 | lncRNA | 0.04694606 | 0.0365854 |
| LINC02113 | lncRNA | 0.04688677 | 0.14265397 |
| CCDC6 | protein_coding | 0.0467351 | 0.76275616 |
| ACBD5 | protein_coding | 0.04665491 | 0.37312277 |
| CNTNAP3 | protein_coding | 0.04652518 | 0.18274543 |
| MRPS21 | protein_coding | 0.04649258 | 0.77296512 |
| BRWD3 | protein_coding | 0.04643706 | 0.61892007 |
| GOLGA1 | protein_coding | 0.04642599 | 0.50182485 |
| C6orf136 | protein_coding | 0.04628482 | 0.40257365 |
| ATAD5 | protein_coding | 0.04614706 | 0.56320069 |
| CDK4 | protein_coding | 0.04614444 | 1.39977758 |
| WRNIP1 | protein_coding | 0.0461379 | 0.67352633 |
| AHCYL1 | protein_coding | 0.04609819 | 0.97375798 |
| DNA2 | protein_coding | 0.04608242 | 0.47159349 |
| RTKN2 | protein_coding | 0.0460161 | 0.25025365 |
| SHPRH | protein_coding | 0.04600164 | 0.45455284 |
| H3-3B | protein_coding | 0.04593086 | 1.69543533 |
| PES1 | protein_coding | 0.04590871 | 1.02555608 |
| MYO15B | protein_coding | 0.04590491 | 0.3460826 |
| PTCD3 | protein_coding | 0.0457791 | 0.87741743 |
| CTC-203F4.2 | lncRNA | 0.04577795 | 0.05241793 |
| SRSF2 | protein_coding | 0.04577536 | 1.47003845 |
| NOL10 | protein_coding | 0.04575399 | 0.71622849 |
| RP11-465L10 | lncRNA | 0.04573117 | 0.09973332 |
| PPFIA1 | protein_coding | 0.04568067 | 0.97147869 |
| TOP1MT | protein_coding | 0.04554375 | 0.37670847 |

| | | | |
|---|---|---|---|
| CFAP251 | protein_coding | 0.04551091 | 0.04914155 |
| TAS2R14 | protein_coding | 0.04546279 | 0.06986519 |
| ENTPD4 | protein_coding | 0.04539851 | 0.47640076 |
| EIF1AY | protein_coding | 0.04534379 | 0.74337568 |
| P2RY11 | protein_coding | 0.04523454 | 0.21114142 |
| ST7L | protein_coding | 0.0451721 | 0.49556137 |
| IPO9 | protein_coding | 0.04502938 | 1.30555393 |
| ARPC2 | protein_coding | 0.04498774 | 1.23192773 |
| INTU | protein_coding | 0.04490853 | 0.64709011 |
| FLNA | protein_coding | 0.04488926 | 0.7507685 |
| CSNK2A3 | protein_coding | 0.04477459 | 0.05380734 |
| ECSIT | protein_coding | 0.04473938 | 0.69375709 |
| LINC02035 | lncRNA | 0.04472181 | 0.13794403 |
| RNF6 | protein_coding | 0.04469738 | 0.80092315 |
| ZNF81 | protein_coding | 0.04463874 | 0.4860354 |
| NDOR1 | protein_coding | 0.04447572 | 0.38242727 |
| LZTR1 | protein_coding | 0.04442408 | 0.30600286 |
| SLC38A1 | protein_coding | 0.04432684 | 1.44901366 |
| NDUFA6-DT | lncRNA | 0.04430515 | 0.1247125 |
| RP11-414J4.[illegible] | lncRNA | 0.04429342 | 0.05561874 |
| GSTCD | protein_coding | 0.04423901 | 0.34640963 |
| SERPINB9 | protein_coding | 0.04422007 | 0.11111414 |
| EXOSC1 | protein_coding | 0.04421574 | 0.27265021 |
| ALKBH1 | protein_coding | 0.04418853 | 0.47063457 |
| MLH1 | protein_coding | 0.04409751 | 0.58083075 |
| MSMO1 | protein_coding | 0.04404527 | 0.63271302 |
| PER2 | protein_coding | 0.04403168 | 0.4545212 |
| GALNT11 | protein_coding | 0.0439334 | 0.64771102 |
| GPRC5D-AS1 | lncRNA | 0.04390706 | 0.19700679 |
| ARFGAP1 | protein_coding | 0.04367608 | 0.72108573 |
| HDHD2 | protein_coding | 0.04365914 | 0.24238965 |
| MRPS18B | protein_coding | 0.04365015 | 0.87283211 |
| FBXW2 | protein_coding | 0.04364134 | 0.99535289 |
| TBRG4 | protein_coding | 0.04356044 | 0.80483655 |
| ATXN7L2 | protein_coding | 0.04355569 | 0.39564141 |
| SSH1 | protein_coding | 0.04348302 | 0.83704187 |
| C14orf28 | protein_coding | 0.04348129 | 0.06994994 |
| CEP57L1 | protein_coding | 0.04345403 | 0.2977199 |
| UTP4 | protein_coding | 0.0434534 | 0.74166377 |
| DHX32 | protein_coding | 0.04333461 | 0.05145819 |
| RP5-971C3.1 | lncRNA | 0.04328239 | 0.06429807 |
| NUS1 | protein_coding | 0.04326186 | 0.40013825 |
| SPR | protein_coding | 0.04324774 | 0.32660634 |
| CIAO2B | protein_coding | 0.04322844 | 0.65328668 |
| PITX1 | protein_coding | 0.04322247 | 0.26122399 |

| | | | |
|---|---|---|---|
| DCTN5 | protein_coding | 0.0432098 | 1.07819151 |
| ZNF134 | protein_coding | 0.04319823 | 0.54366678 |
| ERV3-1 | protein_coding | 0.04301101 | 0.10120786 |
| MRPS36 | protein_coding | 0.04299157 | 0.39137729 |
| ACBD6 | protein_coding | 0.04297036 | 0.60333183 |
| NAA15 | protein_coding | 0.04290089 | 1.12212504 |
| FKBP14 | protein_coding | 0.04287314 | 0.23289011 |
| PARP11 | protein_coding | 0.04284994 | 0.30069584 |
| RNF149 | protein_coding | 0.04279555 | 0.40929639 |
| PCF11 | protein_coding | 0.04279443 | 0.80821417 |
| SENP2 | protein_coding | 0.0427698 | 0.83299783 |
| NRBP1 | protein_coding | 0.04273987 | 0.53108186 |
| SUPT6H | protein_coding | 0.04270389 | 1.16839072 |
| RALGAPB | protein_coding | 0.04261683 | 0.91197743 |
| GPC1 | protein_coding | 0.04261299 | 0.36403879 |
| CCDC61 | protein_coding | 0.04261154 | 0.15050829 |
| EIF2B1 | protein_coding | 0.04244133 | 0.74421584 |
| ZNF17 | protein_coding | 0.04241769 | 0.19927583 |
| CHRNA10 | protein_coding | 0.0422802 | 0.08188756 |
| MAPK8 | protein_coding | 0.04222562 | 0.55367301 |
| SEPTIN3 | protein_coding | 0.04214404 | 1.28979266 |
| ATG12 | protein_coding | 0.04210035 | 0.59766383 |
| PGPEP1 | protein_coding | 0.04194989 | 0.25847493 |
| LINC02166 | lncRNA | 0.04190346 | 0.02691364 |
| ZNF557 | protein_coding | 0.0418869 | 0.33304177 |
| TIMM29 | protein_coding | 0.04181491 | 0.2209501 |
| MRPS16 | protein_coding | 0.04177306 | 0.70819299 |
| PAAF1 | protein_coding | 0.04172715 | 0.45079699 |
| NCOR2 | protein_coding | 0.04161708 | 0.5684916 |
| GPAT4 | protein_coding | 0.0415898 | 0.75301217 |
| ZEB1-AS1 | lncRNA | 0.04141306 | 0.15048301 |
| SATB1-AS1 | lncRNA | 0.04140443 | 0.18002606 |
| LINC01106 | lncRNA | 0.0413588 | 0.06082242 |
| UHRF1BP1L | protein_coding | 0.0413256 | 0.55584387 |
| PGAP1 | protein_coding | 0.04126981 | 0.36448622 |
| MSL3 | protein_coding | 0.04115232 | 0.29634053 |
| FAM161A | protein_coding | 0.04106124 | 0.3581913 |
| ARID3B | protein_coding | 0.04102901 | 0.28787731 |
| PRKAA1 | protein_coding | 0.04102401 | 0.34076741 |
| BTAF1 | protein_coding | 0.04099394 | 0.62546807 |
| SIRT7 | protein_coding | 0.04095646 | 0.39745604 |
| ANKRD13C | protein_coding | 0.04091269 | 0.45381398 |
| RELA | protein_coding | 0.04076271 | 0.80266153 |
| TUBB2A | protein_coding | 0.0407086 | 0.27075012 |
| GLB1L2 | protein_coding | 0.04069245 | 0.38700151 |

| | | | |
|---|---|---|---|
| USPL1 | protein_coding | 0.04057604 | 0.53373973 |
| PDIA2 | protein_coding | 0.04055408 | 0.23022408 |
| XRCC6 | protein_coding | 0.04024854 | 1.29365143 |
| FAM136A | protein_coding | 0.0402184 | 0.68635387 |
| MFSD14A | protein_coding | 0.04014542 | 0.1134968 |
| LCT | protein_coding | 0.03998991 | 0.1176488 |
| HABP4 | protein_coding | 0.03998212 | 0.1913218 |
| NACA | protein_coding | 0.03997375 | 1.14298417 |
| AP5M1 | protein_coding | 0.03994722 | 0.62682278 |
| PXN | protein_coding | 0.03988826 | 0.45719923 |
| KIAA1586 | protein_coding | 0.0398471 | 0.37203527 |
| RP11-454H1: | lncRNA | 0.03978082 | 0.07992441 |
| FAM91A1 | protein_coding | 0.03971249 | 0.50970524 |
| AFG3L2 | protein_coding | 0.03969177 | 0.94874504 |
| ZNF664 | protein_coding | 0.03962116 | 1.30976764 |
| NCLN | protein_coding | 0.03953947 | 0.55996297 |
| ZNF865 | protein_coding | 0.03949426 | 0.16913241 |
| UNC13D | protein_coding | 0.03939179 | 0.07087255 |
| WDR61 | protein_coding | 0.0393604 | 0.54236626 |
| CASP8AP2 | protein_coding | 0.03932659 | 0.79658004 |
| TMEM183A | protein_coding | 0.03922791 | 0.51493967 |
| SNED1 | protein_coding | 0.03915235 | 0.04044266 |
| CWC15 | protein_coding | 0.03912182 | 0.46622118 |
| PPIL1 | protein_coding | 0.03906679 | 0.50070631 |
| XRRA1 | protein_coding | 0.03906017 | 0.29197627 |
| C1RL | protein_coding | 0.03903886 | 0.10661706 |
| SRSF5 | protein_coding | 0.03894609 | 1.08474743 |
| HAUS3 | protein_coding | 0.03894285 | 0.29649607 |
| PISD | protein_coding | 0.03890532 | 0.4373114 |
| PRRC2B | protein_coding | 0.03887713 | 1.32330288 |
| RNF146 | protein_coding | 0.03887041 | 0.16758861 |
| PKMYT1 | protein_coding | 0.03886942 | 0.44192564 |
| KMT2E | protein_coding | 0.03886817 | 0.92159341 |
| ZDHHC3 | protein_coding | 0.03881972 | 0.40034768 |
| SLC4A3 | protein_coding | 0.03880074 | 0.11173527 |
| MYOM2 | protein_coding | 0.03880064 | 0.13936561 |
| CCDC90B | protein_coding | 0.03874237 | 0.45005107 |
| TMEM223 | protein_coding | 0.03868581 | 0.30875791 |
| MCOLN1 | protein_coding | 0.03851053 | 0.25927362 |
| MAP7D3 | protein_coding | 0.03843102 | 0.24637352 |
| ST8SIA4 | protein_coding | 0.0383813 | 0.22148769 |
| KPNA5 | protein_coding | 0.03828628 | 0.23903556 |
| PUDP | protein_coding | 0.03821556 | 0.1871076 |
| SCML1 | protein_coding | 0.0381454 | 0.18234514 |
| TFG | protein_coding | 0.03808021 | 0.71421712 |

| | | | |
|---|---|---|---|
| CEBPD | protein_coding | 0.03807995 | 0.09675137 |
| PPP4R3A | protein_coding | 0.03800886 | 0.98951968 |
| LARS2 | protein_coding | 0.03799465 | 0.37638346 |
| RP13-401N8. | lncRNA | 0.03787799 | 0.04696325 |
| ZIK1 | protein_coding | 0.03781068 | 0.35160755 |
| C9orf78 | protein_coding | 0.03778341 | 0.66386134 |
| C12orf65 | protein_coding | 0.03776233 | 0.57724123 |
| NFRKB | protein_coding | 0.03775809 | 0.53645613 |
| MPDU1 | protein_coding | 0.03771463 | 0.59031064 |
| AC009120.6 | lncRNA | 0.03764771 | 0.13787191 |
| MARCHF8 | protein_coding | 0.03751671 | 0.20951888 |
| ATG3 | protein_coding | 0.03751215 | 0.60902415 |
| ARFRP1 | protein_coding | 0.03737343 | 0.37703189 |
| RPS6KA3 | protein_coding | 0.03728482 | 0.71664574 |
| RP3-465N24. | lncRNA | 0.03726076 | 0.05111793 |
| METTL16 | protein_coding | 0.03725383 | 0.43469676 |
| BCAS2 | protein_coding | 0.03723908 | 0.42485209 |
| PPP2R2B | protein_coding | 0.03717117 | 0.15556477 |
| FBH1 | protein_coding | 0.03701725 | 0.43981215 |
| INTS9 | protein_coding | 0.03699375 | 0.19393272 |
| LSG1 | protein_coding | 0.03688378 | 0.94044998 |
| DHX9 | protein_coding | 0.03687532 | 1.2403962 |
| DPP9 | protein_coding | 0.03685102 | 0.5762411 |
| YKT6 | protein_coding | 0.03678239 | 0.85469889 |
| MARK4 | protein_coding | 0.03675678 | 0.26283872 |
| TFB1M | protein_coding | 0.03675644 | 0.17472874 |
| SIK2 | protein_coding | 0.03662852 | 0.39359411 |
| NHLRC1 | protein_coding | 0.03660802 | 0.1004621 |
| DMAC2L | protein_coding | 0.0366052 | 0.22866486 |
| CTDNEP1 | protein_coding | 0.03648048 | 0.29184673 |
| GRM3-AS1 | lncRNA | 0.03637833 | 0.05847396 |
| RP11-715J22 | lncRNA | 0.03637091 | 0.07814677 |
| PLK4 | protein_coding | 0.03632015 | 0.40069423 |
| ZNF436-AS1 | lncRNA | 0.03620177 | 0.10071432 |
| FEZ1 | protein_coding | 0.03618503 | 0.24360297 |
| MRPL50 | protein_coding | 0.03617163 | 0.44599987 |
| CCDC81 | protein_coding | 0.03610644 | 0.05352744 |
| HEATR5B | protein_coding | 0.03609332 | 0.43837475 |
| PSMC3IP | protein_coding | 0.03605524 | 0.25583027 |
| CAMTA2 | protein_coding | 0.03602057 | 0.45136832 |
| MAP3K4-AS1 | lncRNA | 0.03595993 | 0.05954957 |
| AP1G1 | protein_coding | 0.03591067 | 0.87170776 |
| TM2D1 | protein_coding | 0.03586294 | 0.28707475 |
| ANKRD27 | protein_coding | 0.03583772 | 0.38676603 |
| JMJD4 | protein_coding | 0.03583438 | 0.33699704 |

| | | | |
|---|---|---|---|
| RPS12 | protein_coding | 0.03562078 | 0.68373901 |
| MAU2 | protein_coding | 0.03560614 | 0.73100074 |
| DNAH14 | protein_coding | 0.03554193 | 0.28549572 |
| TAF9B | protein_coding | 0.03539796 | 0.36694292 |
| PYCR2 | protein_coding | 0.03534061 | 0.40776655 |
| OXLD1 | protein_coding | 0.03528396 | 0.23969312 |
| DHRS4 | protein_coding | 0.03517513 | 0.20276851 |
| DMPK | protein_coding | 0.03516458 | 0.32128586 |
| UGCG | protein_coding | 0.03513293 | 0.42700501 |
| METTL2B | protein_coding | 0.0351231 | 0.44447545 |
| IDNK | protein_coding | 0.03511862 | 0.12364217 |
| CRIM1-DT | lncRNA | 0.03508965 | 0.04497013 |
| TLCD5 | protein_coding | 0.03495713 | 0.23742327 |
| NIPBL | protein_coding | 0.03495426 | 0.80791609 |
| PLEKHA5 | protein_coding | 0.03489628 | 0.48781505 |
| PPP6R2 | protein_coding | 0.03482318 | 0.32195373 |
| KLHL12 | protein_coding | 0.0347671 | 0.60768291 |
| E2F3 | protein_coding | 0.0347578 | 0.64653542 |
| LINC02352 | lncRNA | 0.03475344 | 0.03056365 |
| SARM1 | protein_coding | 0.03469528 | 0.43336452 |
| SEC23IP | protein_coding | 0.03455584 | 0.34423474 |
| PCNX3 | protein_coding | 0.03450504 | 0.58659671 |
| UBR1 | protein_coding | 0.03448267 | 0.52315498 |
| TSPYL2 | protein_coding | 0.03446782 | 0.37061635 |
| OPN3 | protein_coding | 0.03444703 | 0.17527947 |
| ZNF714 | protein_coding | 0.03438992 | 0.27137188 |
| PTPN9 | protein_coding | 0.03432572 | 0.34210153 |
| PCBD2 | protein_coding | 0.03429262 | 0.25519346 |
| CUL1 | protein_coding | 0.03427695 | 0.58681075 |
| RP11-378J18 | lncRNA | 0.03422783 | 0.03854256 |
| SH3GLB2 | protein_coding | 0.0342242 | 0.64599741 |
| SMPD2 | protein_coding | 0.03420959 | 0.15531519 |
| PRXL2C | protein_coding | 0.03414647 | 0.21736577 |
| TTC31 | protein_coding | 0.03410931 | 0.32120511 |
| LINC00900 | lncRNA | 0.03403666 | 0.06008144 |
| HADHA | protein_coding | 0.0339632 | 0.75749094 |
| RDX | protein_coding | 0.03391812 | 0.54036413 |
| ACAP3 | protein_coding | 0.03378316 | 0.35479999 |
| NEPRO | protein_coding | 0.03371124 | 0.37721204 |
| BTBD10 | protein_coding | 0.0336297 | 0.39635181 |
| RNF169 | protein_coding | 0.03362067 | 0.3701315 |
| HEMK1 | protein_coding | 0.03357298 | 0.27231461 |
| PRDM8 | protein_coding | 0.0334923 | 0.12200495 |
| ENPP5 | protein_coding | 0.03337984 | 0.11105198 |
| LINC01725 | lncRNA | 0.03332681 | 0.04028583 |

| | | | |
|---|---|---|---|
| CHD1 | protein_coding | 0.03332253 | 0.51266783 |
| STIM2 | protein_coding | 0.03332054 | 0.42254059 |
| ASCC1 | protein_coding | 0.03323522 | 0.23095561 |
| DTD2 | protein_coding | 0.03304546 | 0.17565803 |
| RP11-242D8. | lncRNA | 0.03289065 | 0.14171791 |
| VSTM2B | protein_coding | 0.032808 | 0.0648675 |
| FBXL3 | protein_coding | 0.03280539 | 0.31960165 |
| DDX52 | protein_coding | 0.03279552 | 0.41626386 |
| RIC8B | protein_coding | 0.03277322 | 0.3820813 |
| FARSA | protein_coding | 0.03276737 | 0.66519143 |
| ATP23 | protein_coding | 0.03274075 | 0.21376981 |
| KIF26B | protein_coding | 0.03270436 | 0.1109965 |
| MTA2 | protein_coding | 0.03268866 | 0.94931208 |
| RP11-286N2 | protein_coding | 0.03261327 | 0.02990135 |
| ACOX3 | protein_coding | 0.03260727 | 0.18769467 |
| CHD8 | protein_coding | 0.03255752 | 0.77022391 |
| PI4KA | protein_coding | 0.03253597 | 0.79325127 |
| SGMS1-AS1 | lncRNA | 0.03252659 | 0.06547018 |
| ZNF652 | protein_coding | 0.03238009 | 0.34209931 |
| TPRA1 | protein_coding | 0.0323583 | 0.31725713 |
| P4HA2 | protein_coding | 0.03231177 | 0.2476983 |
| FSIP1 | protein_coding | 0.03225773 | 0.05182305 |
| MAFK | protein_coding | 0.03225756 | 0.3068566 |
| ZNF532 | protein_coding | 0.0322165 | 0.73939407 |
| ZNF2 | protein_coding | 0.03220475 | 0.20168943 |
| MIRLET7A1H | lncRNA | 0.03219576 | 0.10124808 |
| DDX55 | protein_coding | 0.03219277 | 0.40739488 |
| CGGBP1 | protein_coding | 0.03212495 | 0.37246321 |
| KLHDC10 | protein_coding | 0.03211318 | 0.51057124 |
| UBA6-AS1 | lncRNA | 0.03210319 | 0.23757215 |
| RP1-309F20. | lncRNA | 0.03201156 | 0.04443979 |
| DUSP18 | protein_coding | 0.03200629 | 0.07221692 |
| GEMIN7 | protein_coding | 0.03189424 | 0.21883334 |
| ZNF561 | protein_coding | 0.03188241 | 0.41529306 |
| DCPS | protein_coding | 0.03172157 | 0.3318926 |
| KNOP1 | protein_coding | 0.03171333 | 0.49496831 |
| TUBGCP5 | protein_coding | 0.03169255 | 0.34667506 |
| BRAP | protein_coding | 0.03169123 | 0.3362916 |
| COP1 | protein_coding | 0.03166363 | 0.30260401 |
| RPL26 | protein_coding | 0.03165721 | 0.28387058 |
| TMEM79 | protein_coding | 0.03163827 | 0.2662678 |
| RP11-228B1 | lncRNA | 0.03161085 | 0.05889032 |
| PRPF3 | protein_coding | 0.03157392 | 0.60053936 |
| SIL1 | protein_coding | 0.03156768 | 0.28978528 |
| RNF123 | protein_coding | 0.03147203 | 0.26016154 |

| | | | |
|---|---|---|---|
| PCBP2 | protein_coding | 0.03135445 | 0.40754693 |
| ODAD1 | protein_coding | 0.03130805 | 0.10424058 |
| LRRC10 | protein_coding | 0.031244 | 0.04131813 |
| EFNA4 | protein_coding | 0.03124298 | 0.04497961 |
| BNIP2 | protein_coding | 0.0311566 | 0.38998305 |
| SYPL1 | protein_coding | 0.03115612 | 0.28639107 |
| SAR1B | protein_coding | 0.03114838 | 0.4568972 |
| PFN2 | protein_coding | 0.03114718 | 0.35916848 |
| NDUFB3 | protein_coding | 0.03113191 | 0.4087235 |
| PTPN4 | protein_coding | 0.0310994 | 0.31390551 |
| ZNF48 | protein_coding | 0.03092979 | 0.27986873 |
| DYRK1A | protein_coding | 0.03092685 | 0.52885511 |
| RAD50 | protein_coding | 0.0309013 | 0.14095145 |
| SSB | protein_coding | 0.03087651 | 0.60637472 |
| MAP3K4 | protein_coding | 0.03085432 | 0.23550584 |
| AMN1 | protein_coding | 0.03084498 | 0.26180942 |
| RP5-912I13.2 | lncRNA | 0.03082045 | 0.14228683 |
| MYO15A | protein_coding | 0.03071775 | 0.61543992 |
| STAG3 | protein_coding | 0.03064902 | 0.10417241 |
| ITGB3BP | protein_coding | 0.0305123 | 0.34432189 |
| GNAZ | protein_coding | 0.03051198 | 0.24826334 |
| CACNA2D1 | protein_coding | 0.03041337 | 0.45184192 |
| CCDC117 | protein_coding | 0.03040353 | 0.35155188 |
| CCNY-AS1 | lncRNA | 0.03040322 | 0.02538089 |
| SF3B1 | protein_coding | 0.03038466 | 0.81613688 |
| ZNF559 | protein_coding | 0.0303395 | 0.20029146 |
| TPP2 | protein_coding | 0.03032018 | 0.5303951 |
| LUZP1 | protein_coding | 0.030316 | 0.29729001 |
| MKNK1 | protein_coding | 0.03029124 | 0.20921502 |
| TERF2 | protein_coding | 0.03024622 | 0.44993794 |
| SLC25A33 | protein_coding | 0.03011216 | 0.20961041 |
| LINC00174 | lncRNA | 0.03006142 | 0.13322872 |
| ALG1L9P | lncRNA | 0.03000797 | 0.05246514 |
| NELFCD | protein_coding | 0.02994258 | 0.53328607 |
| UCKL1 | protein_coding | 0.02987734 | 0.32127067 |
| CAMKMT | protein_coding | 0.02982404 | 0.16642304 |
| C11orf80 | protein_coding | 0.02981008 | 0.21224269 |
| CAAP1 | protein_coding | 0.02976526 | 0.28062515 |
| OSBPL6 | protein_coding | 0.02976315 | 0.24441511 |
| VPS33B | protein_coding | 0.02974541 | 0.06788105 |
| CNNM1 | protein_coding | 0.02974326 | 0.26324844 |
| NPIPB5 | protein_coding | 0.02963232 | 0.03647071 |
| MTHFD2L | protein_coding | 0.02958886 | 0.21073709 |
| SMU1 | protein_coding | 0.02956866 | 0.51388392 |
| FBXO7 | protein_coding | 0.02955822 | 0.46751694 |

| | | | |
|---|---|---|---|
| GRID2IP | protein_coding | 0.02952186 | 0.09113454 |
| GIT2 | protein_coding | 0.02951572 | 0.25879976 |
| CLIC4 | protein_coding | 0.02949778 | 0.46634995 |
| GPR150 | protein_coding | 0.02947908 | 0.03778387 |
| KIAA0232 | protein_coding | 0.02941874 | 0.52446985 |
| JOSD2 | protein_coding | 0.029366 | 0.05619396 |
| MOSMO | protein_coding | 0.0293568 | 0.15842847 |
| NAPEPLD | protein_coding | 0.02924727 | 0.22005932 |
| UBE4A | protein_coding | 0.02918424 | 0.41883206 |
| PDSS1 | protein_coding | 0.02915357 | 0.1287497 |
| NAAA | protein_coding | 0.02913362 | 0.10229421 |
| GADD45G | protein_coding | 0.02908648 | 0.15642822 |
| LEPROTL1 | protein_coding | 0.02905574 | 0.29108134 |
| CDAN1 | protein_coding | 0.02900618 | 0.27060214 |
| ICE1 | protein_coding | 0.02893688 | 0.58276017 |
| FANCM | protein_coding | 0.02883507 | 0.24635884 |
| SDR39U1 | protein_coding | 0.02877342 | 0.32287879 |
| MSL2 | protein_coding | 0.02862924 | 0.40147799 |
| HOMEZ | protein_coding | 0.0286156 | 0.19834378 |
| RBM23 | protein_coding | 0.02860398 | 0.47315591 |
| CAPN10-DT | lncRNA | 0.02860185 | 0.08616018 |
| BCOR | protein_coding | 0.02857772 | 0.60252133 |
| TRAF6 | protein_coding | 0.02855399 | 0.23330417 |
| AP3D1 | protein_coding | 0.02853573 | 0.53676832 |
| NCAPG | protein_coding | 0.02835943 | 0.58787562 |
| CLTCL1 | protein_coding | 0.02831174 | 0.23807233 |
| ADPGK | protein_coding | 0.02821587 | 0.27974642 |
| ELL | protein_coding | 0.02818465 | 0.26936785 |
| SH3D19 | protein_coding | 0.02818125 | 0.06291797 |
| NFIL3 | protein_coding | 0.02813254 | 0.1486045 |
| ANKRD18B | protein_coding | 0.02799361 | 0.17933256 |
| ERRFI1 | protein_coding | 0.02791588 | 0.0521803 |
| TMCO6 | protein_coding | 0.02777134 | 0.19230966 |
| GABPB1-IT1 | lncRNA | 0.02759783 | 0.20216204 |
| ZNF829 | protein_coding | 0.02752441 | 0.1751376 |
| HSD17B4 | protein_coding | 0.02750575 | 0.46267799 |
| NPRL3 | protein_coding | 0.02750239 | 0.35939019 |
| STAG1 | protein_coding | 0.02748037 | 0.39539113 |
| PYGB | protein_coding | 0.02745888 | 0.35673033 |
| MDH1B | protein_coding | 0.02737674 | 0.04850944 |
| SVOP | protein_coding | 0.02735299 | 0.20066846 |
| UHRF2 | protein_coding | 0.02732967 | 0.4575232 |
| FAM120AOS | protein_coding | 0.02726071 | 0.38403418 |
| ELOA | protein_coding | 0.0272531 | 0.54245588 |
| MTFR2 | protein_coding | 0.02724258 | 0.08773003 |

| | | | |
|---|---|---|---|
| ALX1 | protein_coding | 0.02720875 | 0.43363543 |
| PSMG4 | protein_coding | 0.02719333 | 0.21280567 |
| SLC12A8 | protein_coding | 0.02716754 | 0.06174225 |
| LMTK3 | protein_coding | 0.02716333 | 0.07988213 |
| MORC4 | protein_coding | 0.02714256 | 0.28964236 |
| ANKRD26 | protein_coding | 0.02713223 | 0.16998239 |
| LNX2 | protein_coding | 0.02711874 | 0.29325178 |
| CCNH | protein_coding | 0.02703176 | 0.22666318 |
| VPS52 | protein_coding | 0.02691737 | 0.37851188 |
| CTD-3065J16 | lncRNA | 0.0269082 | 0.03220681 |
| ZNF263 | protein_coding | 0.02686383 | 0.29529497 |
| RAB5B | protein_coding | 0.02682675 | 0.47385775 |
| ERGIC2 | protein_coding | 0.02682356 | 0.24468598 |
| ZNF836 | protein_coding | 0.02682195 | 0.13254404 |
| PSMC3 | protein_coding | 0.02678013 | 0.54940484 |
| SDF4 | protein_coding | 0.0267653 | 0.19023756 |
| PRDM2 | protein_coding | 0.02675748 | 0.50208971 |
| EBAG9 | protein_coding | 0.02673049 | 0.27835006 |
| TXLNA | protein_coding | 0.02667682 | 0.61127213 |
| DESI1 | protein_coding | 0.02659336 | 0.43599715 |
| APP | protein_coding | 0.02637794 | 0.74732323 |
| ANGEL2 | protein_coding | 0.02634623 | 0.2560285 |
| HSD17B1-AS | lncRNA | 0.02632347 | 0.16975851 |
| TEX30 | protein_coding | 0.02626354 | 0.27949905 |
| RANBP2 | protein_coding | 0.02615295 | 0.68489822 |
| ATP6AP1 | protein_coding | 0.02601475 | 0.50574675 |
| GLIS2 | protein_coding | 0.0259927 | 0.11978444 |
| EPS15L1 | protein_coding | 0.02598501 | 0.32635824 |
| KCTD13 | protein_coding | 0.02593943 | 0.2141506 |
| STK25 | protein_coding | 0.02583183 | 0.36090117 |
| FEM1C | protein_coding | 0.02574067 | 0.22695691 |
| SEC23A | protein_coding | 0.02569727 | 0.26680992 |
| HES4 | protein_coding | 0.02569425 | 0.05309564 |
| SRP72 | protein_coding | 0.02563017 | 0.5178212 |
| DHX37 | protein_coding | 0.02561855 | 0.29980791 |
| SENP5 | protein_coding | 0.02552636 | 0.41419085 |
| TMED2-DT | lncRNA | 0.02551387 | 0.04440152 |
| TMED1 | protein_coding | 0.02549364 | 0.20332384 |
| RP11-679B19 | lncRNA | 0.02540849 | 0.06255378 |
| DHRS1 | protein_coding | 0.02529877 | 0.14205554 |
| TSPYL4 | protein_coding | 0.02525102 | 0.2017561 |
| POLR3B | protein_coding | 0.02523913 | 0.26354233 |
| PAQR7 | protein_coding | 0.02523514 | 0.16895347 |
| SMYD3 | protein_coding | 0.02522747 | 0.24150578 |
| CTD-2026K11 | lncRNA | 0.02517052 | 0.02741972 |

| | | | |
|---|---|---|---|
| RP11-1072A3 | lncRNA | 0.02509097 | 0.02351134 |
| MAP7D2 | protein_coding | 0.0250139 | 0.47705858 |
| FAM220A | protein_coding | 0.02497357 | 0.18780968 |
| TTC28 | protein_coding | 0.02480516 | 0.35741918 |
| DDX27 | protein_coding | 0.02480314 | 0.32412161 |
| ZNF805 | protein_coding | 0.02475812 | 0.13923701 |
| ZNF490 | protein_coding | 0.02468315 | 0.0693888 |
| NDST1 | protein_coding | 0.02459171 | 0.39859592 |
| ZNF808 | protein_coding | 0.0245266 | 0.16317844 |
| ACAT2 | protein_coding | 0.02440975 | 0.25562462 |
| MRS2 | protein_coding | 0.0243411 | 0.32451012 |
| ATP2A3 | protein_coding | 0.02428275 | 0.08697895 |
| EPM2A | protein_coding | 0.02417702 | 0.1094091 |
| ABL1 | protein_coding | 0.02413415 | 0.40204241 |
| NIPSNAP3A | protein_coding | 0.0240382 | 0.25901233 |
| ZNF169 | protein_coding | 0.02402382 | 0.06891583 |
| TADA2B | protein_coding | 0.02356147 | 0.31649257 |
| SLC17A5 | protein_coding | 0.02351628 | 0.24080239 |
| C16orf95 | protein_coding | 0.02348286 | 0.05153282 |
| RP5-827C21. | lncRNA | 0.02327909 | 0.02164731 |
| TRIT1 | protein_coding | 0.02326679 | 0.23454893 |
| LRR1 | protein_coding | 0.0232629 | 0.23191875 |
| SLC31A1 | protein_coding | 0.02324986 | 0.36238731 |
| PDS5A | protein_coding | 0.02324708 | 0.48519264 |
| DHX29 | protein_coding | 0.02322388 | 0.33643256 |
| WIPF2 | protein_coding | 0.02321737 | 0.31128175 |
| TSPAN9 | protein_coding | 0.02319683 | 0.17826345 |
| MTX1 | protein_coding | 0.02317922 | 0.14812971 |
| DCAF13 | protein_coding | 0.0231648 | 0.36157196 |
| SETBP1 | protein_coding | 0.02315222 | 0.32921421 |
| NDUFV2 | protein_coding | 0.02315172 | 0.03859256 |
| RP11-1246C1 | lncRNA | 0.02312822 | 0.13966301 |
| CCDC124 | protein_coding | 0.02309206 | 0.16632657 |
| PRKAG2-AS1 | lncRNA | 0.02306103 | 0.07007204 |
| KIF16B | protein_coding | 0.02304163 | 0.11091548 |
| CBWD5 | protein_coding | 0.02302113 | 0.11208611 |
| DCTN6 | protein_coding | 0.02301521 | 0.0973533 |
| LARP1B | protein_coding | 0.02301292 | 0.26720616 |
| NFE2L1-DT | lncRNA | 0.02297975 | 0.04534599 |
| TCOF1 | protein_coding | 0.02293691 | 0.43297929 |
| AK3 | protein_coding | 0.02278707 | 0.28517395 |
| GFPT1 | protein_coding | 0.02274985 | 0.30713626 |
| UCHL5 | protein_coding | 0.02264293 | 0.33199338 |
| SPHK2 | protein_coding | 0.02264104 | 0.17197602 |
| REXO4 | protein_coding | 0.02259013 | 0.23077042 |

| | | | |
|---|---|---|---|
| ABHD11 | protein_coding | 0.0225683 | 0.14758012 |
| ASXL1 | protein_coding | 0.02242096 | 0.38329824 |
| NME2 | protein_coding | 0.0223685 | 0.03464562 |
| MALSU1 | protein_coding | 0.02235742 | 0.14178816 |
| SLC43A3 | protein_coding | 0.02233501 | 0.39664179 |
| NDUFAF5 | protein_coding | 0.02231431 | 0.17042644 |
| DUSP28 | protein_coding | 0.02228905 | 0.10034462 |
| EMC8 | protein_coding | 0.02208018 | 0.27747726 |
| NR0B1 | protein_coding | 0.02198112 | 0.05099728 |
| NME7 | protein_coding | 0.02190947 | 0.1559978 |
| PKP4 | protein_coding | 0.02189507 | 0.41030263 |
| RIOK2 | protein_coding | 0.02185148 | 0.21739278 |
| DCLK3 | protein_coding | 0.02174695 | 0.06880619 |
| DCAF11 | protein_coding | 0.02172487 | 0.30816895 |
| CSNK1G2 | protein_coding | 0.0216102 | 0.16356246 |
| LINC02295 | lncRNA | 0.02153159 | 0.03474164 |
| PSMD9 | protein_coding | 0.02152373 | 0.04953954 |
| STARD5 | protein_coding | 0.02150136 | 0.03227615 |
| RP11-152N1 | lncRNA | 0.02142285 | 0.10325612 |
| CHMP3 | protein_coding | 0.02132217 | 0.26697371 |
| KRIT1 | protein_coding | 0.02131813 | 0.09573615 |
| PCSK1N | protein_coding | 0.021301 | 0.0399626 |
| NSMCE1 | protein_coding | 0.02128883 | 0.20108529 |
| EIF3B | protein_coding | 0.02122288 | 0.39717721 |
| D2HGDH | protein_coding | 0.02113654 | 0.12370729 |
| EIF2B5 | protein_coding | 0.02109675 | 0.4075399 |
| MAP2K2 | protein_coding | 0.02106783 | 0.17984528 |
| BBS7 | protein_coding | 0.02080599 | 0.13928474 |
| EBPL | protein_coding | 0.02077829 | 0.14518724 |
| TMPPE | protein_coding | 0.02071671 | 0.03351607 |
| WDR46 | protein_coding | 0.02069955 | 0.19729165 |
| EIF2B2 | protein_coding | 0.02069432 | 0.26957729 |
| NAIF1 | protein_coding | 0.02067203 | 0.1148406 |
| CCDC12 | protein_coding | 0.02055849 | 0.12133759 |
| FABP5 | protein_coding | 0.02051152 | 0.19464597 |
| DNAJC19 | protein_coding | 0.02049675 | 0.22063898 |
| MLLT1 | protein_coding | 0.02042928 | 0.2093502 |
| TINCR | protein_coding | 0.02031465 | 0.03703207 |
| UBXN6 | protein_coding | 0.02024922 | 0.19459454 |
| MELK | protein_coding | 0.02010235 | 0.27441489 |
| ZNF696 | protein_coding | 0.02007519 | 0.07619852 |
| HENMT1 | protein_coding | 0.02004165 | 0.11767505 |
| THRA | protein_coding | 0.02004145 | 0.38698042 |
| RAB5A | protein_coding | 0.02003252 | 0.17110051 |
| ATP5F1C | protein_coding | 0.01999182 | 0.28462177 |

| | | | |
|---|---|---|---|
| DENND6B | protein_coding | 0.01996852 | 0.04049938 |
| NRXN1 | protein_coding | 0.01991056 | 0.25623311 |
| ZDHHC21 | protein_coding | 0.0199101 | 0.11843926 |
| CT66 | lncRNA | 0.01990107 | 0.02015104 |
| REXO2 | protein_coding | 0.01988483 | 0.12880128 |
| FAM30A | lncRNA | 0.01981463 | 0.14284254 |
| CPSF4 | protein_coding | 0.01975858 | 0.25598877 |
| TBP | protein_coding | 0.01953001 | 0.12690096 |
| CTD-2335O3. | lncRNA | 0.01948775 | 0.07185982 |
| YJU2 | protein_coding | 0.01947601 | 0.13517965 |
| METTL2A | protein_coding | 0.01933165 | 0.15019444 |
| CSNK1E | protein_coding | 0.01927022 | 0.27960971 |
| GGA1 | protein_coding | 0.01922855 | 0.28488492 |
| RANBP17 | protein_coding | 0.01919456 | 0.12234032 |
| CCDC186 | protein_coding | 0.01913643 | 0.149964 |
| SSR2 | protein_coding | 0.01910708 | 0.30093363 |
| CHKB-DT | lncRNA | 0.01906287 | 0.05538729 |
| SPINT1-AS1 | lncRNA | 0.01904605 | 0.0335612 |
| RAD51-AS1 | lncRNA | 0.01903826 | 0.06212155 |
| DENND10 | protein_coding | 0.01903386 | 0.0438034 |
| PSEN1 | protein_coding | 0.01899536 | 0.30611598 |
| CCHCR1 | protein_coding | 0.01898077 | 0.24351646 |
| TIMM17A | protein_coding | 0.01897856 | 0.25421216 |
| LZIC | protein_coding | 0.0189534 | 0.14348519 |
| PBX3 | protein_coding | 0.0189419 | 0.19732468 |
| TRA2A | protein_coding | 0.01888496 | 0.21990324 |
| FAM189B | protein_coding | 0.01869981 | 0.1699502 |
| TUBB3 | protein_coding | 0.01868174 | 0.04652705 |
| RPRD1A | protein_coding | 0.01864555 | 0.21942412 |
| FIS1 | protein_coding | 0.0186176 | 0.18221968 |
| ALDH1A1 | protein_coding | 0.01855876 | 0.04180448 |
| UPF2 | protein_coding | 0.01848537 | 0.16436904 |
| INTS14 | protein_coding | 0.01845823 | 0.21540485 |
| UQCRB | protein_coding | 0.01844211 | 0.34237635 |
| PGLS | protein_coding | 0.01832609 | 0.14544598 |
| SUSD1 | protein_coding | 0.01832409 | 0.04529564 |
| PIF1 | protein_coding | 0.01830686 | 0.12472374 |
| RRS1 | protein_coding | 0.01827558 | 0.07062969 |
| RP11-552M1 | lncRNA | 0.01816815 | 0.01785847 |
| SMIM12 | protein_coding | 0.01813544 | 0.18855858 |
| IREB2 | protein_coding | 0.01812358 | 0.28041061 |
| TAF1A-AS1 | lncRNA | 0.01812021 | 0.03741376 |
| MAP2K7 | protein_coding | 0.01805318 | 0.23894323 |
| AEBP2 | protein_coding | 0.01804574 | 0.14192388 |
| GLOD4 | protein_coding | 0.01803418 | 0.22213744 |

| | | | |
|---|---|---|---|
| DNAJB9 | protein_coding | 0.01785226 | 0.13570586 |
| FAM83H | protein_coding | 0.01771615 | 0.08712088 |
| RP11-196G1 | lncRNA | 0.01768282 | 0.07505736 |
| ANO8 | protein_coding | 0.01761268 | 0.1441027 |
| DCUN1D5 | protein_coding | 0.01758206 | 0.13584521 |
| ARC | protein_coding | 0.0175124 | 0.05905173 |
| TARS3 | protein_coding | 0.01750741 | 0.12316219 |
| TMEM147-A | lncRNA | 0.01748405 | 0.09028342 |
| ABHD10 | protein_coding | 0.01742372 | 0.2311939 |
| SCYL2 | protein_coding | 0.01740522 | 0.20151424 |
| YIPF1 | protein_coding | 0.01736415 | 0.11017904 |
| PPAN | protein_coding | 0.01732525 | 0.08120668 |
| IPO8 | protein_coding | 0.01731035 | 0.22601024 |
| TANGO6 | protein_coding | 0.01727632 | 0.11961633 |
| ITGB1BP2 | protein_coding | 0.01725627 | 0.02159128 |
| HEBP2 | protein_coding | 0.01722204 | 0.1241054 |
| RP11-802E1 | lncRNA | 0.01715352 | 0.03805774 |
| ARID4B | protein_coding | 0.01714362 | 0.19730396 |
| CDV3 | protein_coding | 0.01713373 | 0.33709357 |
| PRDM15 | protein_coding | 0.0170663 | 0.13954768 |
| TMEM254 | protein_coding | 0.0169753 | 0.07899017 |
| PDE3B | protein_coding | 0.0168543 | 0.01869169 |
| MYT1L | protein_coding | 0.01666111 | 0.15225085 |
| SLC37A4 | protein_coding | 0.01655877 | 0.21644952 |
| PFDN5 | protein_coding | 0.01643992 | 0.04640129 |
| VPS26C | protein_coding | 0.01642352 | 0.19207787 |
| MTHFD1 | protein_coding | 0.01641473 | 0.38880867 |
| TMEM182 | protein_coding | 0.01640124 | 0.05743935 |
| MAP3K7CL | protein_coding | 0.01639467 | 0.02618926 |
| UBE2R2 | protein_coding | 0.01631528 | 0.23517432 |
| MCU | protein_coding | 0.01623574 | 0.09431808 |
| RBL2 | protein_coding | 0.01617088 | 0.20425916 |
| CUTC | protein_coding | 0.01616258 | 0.07370279 |
| ZNF561-AS1 | lncRNA | 0.01613732 | 0.05344889 |
| SOS1-IT1 | lncRNA | 0.0160068 | 0.03790951 |
| SLC50A1 | protein_coding | 0.01597235 | 0.17942841 |
| CGAS | protein_coding | 0.01583232 | 0.0793264 |
| MAPK1IP1L | protein_coding | 0.01570395 | 0.17656163 |
| SPATS2 | protein_coding | 0.01560442 | 0.23888591 |
| CEP128 | protein_coding | 0.01558894 | 0.10698793 |
| TOP3B | protein_coding | 0.01555732 | 0.11764411 |
| H1-10 | protein_coding | 0.01553847 | 0.07420474 |
| KCTD1 | protein_coding | 0.01541147 | 0.1033191 |
| RP4-751H13. | lncRNA | 0.01539728 | 0.01927329 |
| CTD-3092A1 | lncRNA | 0.01536445 | 0.04088291 |

| | | | |
|---|---|---|---|
| PPIF | protein_coding | 0.01526106 | 0.19512436 |
| AP1M1 | protein_coding | 0.0152487 | 0.16577265 |
| TMX2 | protein_coding | 0.01524816 | 0.26564963 |
| RP11-210M1 | lncRNA | 0.01524737 | 0.0228541 |
| ZNF329 | protein_coding | 0.0151836 | 0.10849472 |
| PLD3 | protein_coding | 0.01513071 | 0.1855181 |
| MED22 | protein_coding | 0.01512941 | 0.13864525 |
| RP11-650L12 | lncRNA | 0.01509021 | 0.03809356 |
| PEX12 | protein_coding | 0.01500848 | 0.093598 |
| NDUFS1 | protein_coding | 0.01496311 | 0.32174269 |
| NDUFS7 | protein_coding | 0.01482731 | 0.0983718 |
| EMC4 | protein_coding | 0.01480678 | 0.22385038 |
| TCAIM | protein_coding | 0.01474539 | 0.08006801 |
| NDUFS3 | protein_coding | 0.01465225 | 0.16150466 |
| TAF1 | protein_coding | 0.01458343 | 0.19199896 |
| DDAH2 | protein_coding | 0.01454728 | 0.05553019 |
| GNPAT | protein_coding | 0.01452594 | 0.15870268 |
| TIFA | protein_coding | 0.01446996 | 0.15868457 |
| RP11-166O4. | lncRNA | 0.01440617 | 0.0133977 |
| SLC33A1 | protein_coding | 0.01439465 | 0.07428488 |
| TATDN3 | protein_coding | 0.01432838 | 0.11350704 |
| GNL3L | protein_coding | 0.01426646 | 0.15034569 |
| SLC5A6 | protein_coding | 0.01424264 | 0.17653247 |
| TRPC4AP | protein_coding | 0.01423578 | 0.19064699 |
| NOC4L | protein_coding | 0.01414215 | 0.09267053 |
| BRD1 | protein_coding | 0.01406607 | 0.16389268 |
| JAZF1 | protein_coding | 0.01406009 | 0.09521981 |
| AARS2 | protein_coding | 0.01402296 | 0.16496448 |
| RNF212 | protein_coding | 0.01397102 | 0.16632853 |
| FAM118A | protein_coding | 0.01390045 | 0.13870453 |
| TMEM67 | protein_coding | 0.0138997 | 0.04959007 |
| SEPTIN2 | protein_coding | 0.01387985 | 0.32096604 |
| LINC00294 | lncRNA | 0.01374616 | 0.09975386 |
| EIF5AL1 | protein_coding | 0.01357907 | 0.0327954 |
| HDAC6 | protein_coding | 0.01348 | 0.18172344 |
| DDX5 | protein_coding | 0.01338676 | 0.33540845 |
| MPHOSPH8 | protein_coding | 0.0133777 | 0.12536131 |
| LINC02325 | lncRNA | 0.01334741 | 0.02133234 |
| CFAP20 | protein_coding | 0.01333625 | 0.14323156 |
| KIF5B | protein_coding | 0.01330786 | 0.24108517 |
| B3GALT5 | protein_coding | 0.01320041 | 0.03148608 |
| MED19 | protein_coding | 0.01318077 | 0.07739634 |
| FGF12-AS2 | lncRNA | 0.01310542 | 0.02661327 |
| PSME3 | protein_coding | 0.01306637 | 0.23270428 |
| MLF2 | protein_coding | 0.01305796 | 0.21206673 |

| | | | |
|---|---|---|---|
| ANO5 | protein_coding | 0.01304566 | 0.24390371 |
| PTPN11 | protein_coding | 0.01304056 | 0.30040106 |
| LY6G5B | protein_coding | 0.01301063 | 0.03680721 |
| ZNF8 | protein_coding | 0.01289146 | 0.09200311 |
| TRNAU1AP | protein_coding | 0.01276983 | 0.10862424 |
| IFNAR1 | protein_coding | 0.01276977 | 0.17426691 |
| KLK4 | protein_coding | 0.01264823 | 0.02010688 |
| DDX20 | protein_coding | 0.01260572 | 0.10371147 |
| LRRC47 | protein_coding | 0.01260055 | 0.10728617 |
| DNAJC25 | protein_coding | 0.01255236 | 0.06299323 |
| RGL1 | protein_coding | 0.01241257 | 0.09187031 |
| LSS | protein_coding | 0.01241021 | 0.2143297 |
| ASB8 | protein_coding | 0.01237392 | 0.13167771 |
| OSBPL11 | protein_coding | 0.01224057 | 0.12486464 |
| HBS1L | protein_coding | 0.01223698 | 0.10511589 |
| RNF113A | protein_coding | 0.0122063 | 0.07470252 |
| ZFR | protein_coding | 0.01220469 | 0.194177 |
| TERF2IP | protein_coding | 0.01210163 | 0.15026393 |
| WIZ | protein_coding | 0.01204316 | 0.16626853 |
| NRL | protein_coding | 0.01199302 | 0.03263612 |
| RBM6 | protein_coding | 0.01198935 | 0.12895092 |
| NDUFB5 | protein_coding | 0.0119071 | 0.11340471 |
| VCPIP1 | protein_coding | 0.01175251 | 0.06360834 |
| AP3S1 | protein_coding | 0.01152268 | 0.10002449 |
| ZNF132 | protein_coding | 0.01144825 | 0.04745702 |
| GOLIM4 | protein_coding | 0.01143343 | 0.15874109 |
| RP5-981L23.1 | lncRNA | 0.01143006 | 0.01062533 |
| HMOX2 | protein_coding | 0.01132329 | 0.12244044 |
| EARS2 | protein_coding | 0.01130407 | 0.14284639 |
| PTP4A1 | protein_coding | 0.01130402 | 0.01176712 |
| RPS27A | protein_coding | 0.01125971 | 0.20871892 |
| IST1 | protein_coding | 0.0112546 | 0.17954299 |
| MKKS | protein_coding | 0.01119106 | 0.10765599 |
| SLAIN2 | protein_coding | 0.01109365 | 0.10286534 |
| PNRC2 | protein_coding | 0.01106766 | 0.15390935 |
| MBTPS1 | protein_coding | 0.01103961 | 0.1812245 |
| ZNF468 | protein_coding | 0.01090235 | 0.07294746 |
| RP11-230C9. | lncRNA | 0.01088324 | 0.01078589 |
| PEPD | protein_coding | 0.01082778 | 0.11470053 |
| RP11-33N14. | lncRNA | 0.01077706 | 0.00966478 |
| CAMLG | protein_coding | 0.01072809 | 0.10772554 |
| TOP3A | protein_coding | 0.01060605 | 0.13341127 |
| MRTFB | protein_coding | 0.01044527 | 0.12489224 |
| LOH12CR2 | lncRNA | 0.01041016 | 0.02089537 |
| CDK9 | protein_coding | 0.01033156 | 0.10195201 |

| | | | |
|---|---|---|---|
| MAFA | protein_coding | 0.01024002 | 0.01291694 |
| RNF141 | protein_coding | 0.0102197 | 0.07729383 |
| UCK2 | protein_coding | 0.01019851 | 0.12783869 |
| MCTS1 | protein_coding | 0.01008101 | 0.11245474 |
| USP36 | protein_coding | 0.00997481 | 0.12136913 |
| TRAPPC12 | protein_coding | 0.00997254 | 0.07435265 |
| FBXO42 | protein_coding | 0.00994872 | 0.12411268 |
| DNMT1 | protein_coding | 0.00986901 | 0.22547229 |
| KIFAP3 | protein_coding | 0.0098536 | 0.10425648 |
| MGAT2 | protein_coding | 0.00981605 | 0.09107371 |
| NCL | protein_coding | 0.00979734 | 0.23942462 |
| RAD51 | protein_coding | 0.00979201 | 0.05688334 |
| TUSC3 | protein_coding | 0.00973824 | 0.09512978 |
| KATNB1 | protein_coding | 0.00971258 | 0.10039185 |
| MRPS14 | protein_coding | 0.00964424 | 0.07097288 |
| SYMPK | protein_coding | 0.00960584 | 0.17076984 |
| SETMAR | protein_coding | 0.00959575 | 0.05498982 |
| KIF24 | protein_coding | 0.00957373 | 0.06807806 |
| TRAPPC3 | protein_coding | 0.00955486 | 0.10327788 |
| ZNF256 | protein_coding | 0.00952982 | 0.03898982 |
| RHOG | protein_coding | 0.00951329 | 0.06084513 |
| CNP | protein_coding | 0.00950596 | 0.12278352 |
| LAMP1 | protein_coding | 0.00943378 | 0.15634917 |
| MVD | protein_coding | 0.00935566 | 0.10043537 |
| SEC22B | protein_coding | 0.00935001 | 0.08886638 |
| IMMT | protein_coding | 0.00933215 | 0.13016433 |
| SMYD5 | protein_coding | 0.00923583 | 0.06953232 |
| PDCD2 | protein_coding | 0.00912829 | 0.07655599 |
| CTD-2035E11 | lncRNA | 0.00906994 | 0.00918471 |
| ANAPC4 | protein_coding | 0.00906437 | 0.08999342 |
| TMEM135 | protein_coding | 0.00905552 | 0.06320565 |
| ASAP1 | protein_coding | 0.00899945 | 0.07214585 |
| AASDH | protein_coding | 0.00897251 | 0.04778953 |
| TMED5 | protein_coding | 0.00888857 | 0.0647173 |
| CEP126 | protein_coding | 0.00883033 | 0.05239112 |
| STARD3 | protein_coding | 0.00871369 | 0.07445475 |
| DNAJB1 | protein_coding | 0.00867587 | 0.11079263 |
| RRAGB | protein_coding | 0.00864463 | 0.0481051 |
| PROSER3 | protein_coding | 0.00836747 | 0.05152703 |
| ADCK5 | protein_coding | 0.0083598 | 0.03930764 |
| UMPS | protein_coding | 0.00827617 | 0.06671899 |
| SMPD1 | protein_coding | 0.00824474 | 0.07578701 |
| ARFGAP2 | protein_coding | 0.00823429 | 0.12079681 |
| CALU | protein_coding | 0.00818832 | 0.15631235 |
| RHEB | protein_coding | 0.0081596 | 0.10880879 |

| | | | |
|---|---|---|---|
| SPECC1L | protein_coding | 0.00807256 | 0.08759722 |
| VIRMA | protein_coding | 0.00804754 | 0.10096702 |
| ATP6V1G1 | protein_coding | 0.00802801 | 0.08045901 |
| DLG1 | protein_coding | 0.00791195 | 0.13269913 |
| LINS1 | protein_coding | 0.00781957 | 0.05593372 |
| ZNF470 | protein_coding | 0.00772506 | 0.03416653 |
| TMEM219 | protein_coding | 0.00771282 | 0.08165457 |
| AZIN1 | protein_coding | 0.00762005 | 0.13447977 |
| UBAP2 | protein_coding | 0.00760367 | 0.08710353 |
| RPUSD2 | protein_coding | 0.00757981 | 0.04293786 |
| RFC2 | protein_coding | 0.00751665 | 0.10148589 |
| TRIM33 | protein_coding | 0.00739661 | 0.08750733 |
| ANKHD1 | protein_coding | 0.00737275 | 0.01668626 |
| RASSF1 | protein_coding | 0.00730346 | 0.04220439 |
| CCDC113 | protein_coding | 0.00726463 | 0.03333633 |
| PRELID3B | protein_coding | 0.00720672 | 0.05972343 |
| RP11-61J19.! | lncRNA | 0.00719579 | 0.02381962 |
| CDC73 | protein_coding | 0.00711179 | 0.05921982 |
| HCFC1 | protein_coding | 0.00711118 | 0.10593648 |
| TSSC4 | protein_coding | 0.00708124 | 0.05929518 |
| RBM4B | protein_coding | 0.00702363 | 0.06458221 |
| ZDHHC13 | protein_coding | 0.00697181 | 0.06312787 |
| ZNF566 | protein_coding | 0.00694308 | 0.04525325 |
| APOOL | protein_coding | 0.00689933 | 0.0408409 |
| KATNIP | protein_coding | 0.00689132 | 0.05068389 |
| CNTRL | protein_coding | 0.00685892 | 0.06276167 |
| ZMYM4 | protein_coding | 0.00669042 | 0.08714881 |
| IMPDH2 | protein_coding | 0.00665318 | 0.08672297 |
| TMEM175 | protein_coding | 0.00661275 | 0.05429752 |
| SNTB2 | protein_coding | 0.00658863 | 0.04599412 |
| USP31 | protein_coding | 0.00650764 | 0.06029416 |
| RP4-660H19. | lncRNA | 0.00644899 | 0.00901171 |
| ZNF282 | protein_coding | 0.00644274 | 0.05576177 |
| PPP1R10 | protein_coding | 0.0062887 | 0.11135635 |
| SIK1 | protein_coding | 0.00628584 | 0.01150596 |
| RUSC2 | protein_coding | 0.00625755 | 0.03905286 |
| SF1 | protein_coding | 0.00620756 | 0.13029358 |
| ZFC3H1 | protein_coding | 0.00612484 | 0.05929994 |
| USP7 | protein_coding | 0.00599773 | 0.11096661 |
| MMAB | protein_coding | 0.00594593 | 0.05472568 |
| LARP4 | protein_coding | 0.00591684 | 0.05301018 |
| CPPED1 | protein_coding | 0.00568381 | 0.04569861 |
| HNRNPDL | protein_coding | 0.00568243 | 0.11446205 |
| RPP14 | protein_coding | 0.00568185 | 0.02151493 |
| CACUL1 | protein_coding | 0.00564918 | 0.05763042 |

| | | | |
|---|---|---|---|
| VCPKMT | protein_coding | 0.00560169 | 0.02757548 |
| C11orf49 | protein_coding | 0.00556215 | 0.03792402 |
| ZNF251 | protein_coding | 0.00555953 | 0.03153576 |
| QPRT | protein_coding | 0.00553456 | 0.01188297 |
| DGKZ | protein_coding | 0.00553359 | 0.06358713 |
| CSAD | protein_coding | 0.00548713 | 0.02352276 |
| FER | protein_coding | 0.00539949 | 0.04226904 |
| TIMM22 | protein_coding | 0.00531972 | 0.03176313 |
| EMC6 | protein_coding | 0.00521138 | 0.02507179 |
| PTCD1 | protein_coding | 0.00515808 | 0.01931259 |
| MACROH2A2 | protein_coding | 0.00515503 | 0.05495421 |
| CREB1 | protein_coding | 0.00512662 | 0.05058375 |
| FAM229B | protein_coding | 0.00509415 | 0.01925227 |
| YPEL5 | protein_coding | 0.00501958 | 0.0397534 |
| TIPRL | protein_coding | 0.00500162 | 0.05929934 |
| PNPLA5 | protein_coding | 0.00494525 | 0.01861674 |
| PARN | protein_coding | 0.00483353 | 0.04411117 |
| DLX6-AS1 | lncRNA | 0.00482627 | 0.01120719 |
| CLEC18B | protein_coding | 0.00476646 | 0.00972223 |
| ZNF852 | protein_coding | 0.00468885 | 0.01736204 |
| TPR | protein_coding | 0.00468477 | 0.08728264 |
| BRIX1 | protein_coding | 0.00448262 | 0.05585968 |
| BAZ2B | protein_coding | 0.0044567 | 0.04583225 |
| WDR77 | protein_coding | 0.00440748 | 0.04254253 |
| SCO1 | protein_coding | 0.00438682 | 0.03203463 |
| ZSCAN25 | protein_coding | 0.00436702 | 0.02756499 |
| RP11-506H2 | lncRNA | 0.00434043 | 0.00791368 |
| NLE1 | protein_coding | 0.00430807 | 0.03308973 |
| ARHGEF12 | protein_coding | 0.00424437 | 0.06621621 |
| MCPH1 | protein_coding | 0.00422613 | 0.03278106 |
| INTS4 | protein_coding | 0.00414818 | 0.03821568 |
| POLR2C | protein_coding | 0.00410078 | 0.04919366 |
| ZNF638 | protein_coding | 0.0039891 | 0.06725589 |
| DDX51 | protein_coding | 0.00398113 | 0.0311011 |
| RTN4 | protein_coding | 0.00395994 | 0.06235418 |
| EIF2D | protein_coding | 0.0039353 | 0.03708579 |
| KRI1 | protein_coding | 0.00382741 | 0.03349561 |
| COMT | protein_coding | 0.00380391 | 0.03663104 |
| EHMT1 | protein_coding | 0.00377438 | 0.05017437 |
| ATP5PD | protein_coding | 0.00368484 | 0.0356355 |
| CSE1L | protein_coding | 0.00366477 | 0.05998754 |
| ZNF230 | protein_coding | 0.00361176 | 0.01412445 |
| GTPBP8 | protein_coding | 0.00360599 | 0.01917138 |
| TMED4 | protein_coding | 0.00352517 | 0.04933136 |
| NSD1 | protein_coding | 0.00346213 | 0.04664431 |

| | | | |
|---|---|---|---|
| ATL1 | protein_coding | 0.003408 | 0.01714918 |
| SMC4 | protein_coding | 0.00337807 | 0.04256565 |
| INTS6 | protein_coding | 0.00336038 | 0.0329343 |
| PICK1 | protein_coding | 0.00332006 | 0.03243825 |
| SNX3 | protein_coding | 0.00331816 | 0.03095849 |
| GALNT13 | protein_coding | 0.00331569 | 0.03642561 |
| C9orf85 | protein_coding | 0.00324132 | 0.01708597 |
| SIRT1 | protein_coding | 0.00323152 | 0.02529721 |
| ZMPSTE24 | protein_coding | 0.00321174 | 0.03003577 |
| RER1 | protein_coding | 0.00320941 | 0.02930427 |
| KCTD16 | protein_coding | 0.00314937 | 0.00723111 |
| DENR | protein_coding | 0.00309575 | 0.04358913 |
| TEX2 | protein_coding | 0.00297308 | 0.02954802 |
| LINC01116 | lncRNA | 0.00289781 | 0.01755038 |
| NOL3 | protein_coding | 0.00283066 | 0.0113254 |
| ITPA | protein_coding | 0.00274535 | 0.02764846 |
| POLR3F | protein_coding | 0.00274062 | 0.02065441 |
| SMUG1 | protein_coding | 0.00273821 | 0.02526509 |
| THUMPD1 | protein_coding | 0.00269991 | 0.03039974 |
| TMEM184C | protein_coding | 0.00268475 | 0.02528845 |
| AP001372.2 | lncRNA | 0.0026562 | 0.00657467 |
| CCDC85B | protein_coding | 0.00265481 | 0.00453802 |
| DHFR2 | protein_coding | 0.00255864 | 0.0153269 |
| TRMT1L | protein_coding | 0.00255479 | 0.02125418 |
| KLHL29 | protein_coding | 0.00253345 | 0.01346177 |
| CUL4B | protein_coding | 0.00249007 | 0.02885036 |
| DGKE | protein_coding | 0.00244964 | 0.02502074 |
| TMEM109 | protein_coding | 0.00244876 | 0.02700847 |
| RNF216P1 | lncRNA | 0.00243992 | 0.0093841 |
| CNIH3 | protein_coding | 0.00235919 | 0.00640313 |
| TTC17 | protein_coding | 0.00231044 | 0.02863897 |
| RP11-147L13 | lncRNA | 0.0023052 | 0.02182146 |
| CALML4 | protein_coding | 0.00227882 | 0.0021859 |
| LDAH | protein_coding | 0.00221967 | 0.0201073 |
| NIFK | protein_coding | 0.00218483 | 0.02698048 |
| RBM15 | protein_coding | 0.00200532 | 0.0145436 |
| STRN3 | protein_coding | 0.00200225 | 0.01916825 |
| TAMM41 | protein_coding | 0.0019807 | 0.00789135 |
| CHD3 | protein_coding | 0.00191439 | 0.02490696 |
| HSDL2 | protein_coding | 0.00187541 | 0.01816401 |
| PIMREG | protein_coding | 0.00183538 | 0.01960456 |
| ZNF718 | protein_coding | 0.00182784 | 0.01067782 |
| HNRNPC | protein_coding | 0.00177384 | 0.03263843 |
| MARCHF5 | protein_coding | 0.00163082 | 0.01420215 |
| NTNG2 | protein_coding | 0.00154331 | 0.00443718 |

| | | | |
|---|---|---|---|
| TMEM94 | protein_coding | 0.00148656 | 0.01539126 |
| CIAO1 | protein_coding | 0.00146262 | 0.01793488 |
| EDRF1 | protein_coding | 0.00141297 | 0.0129471 |
| ARL17A | protein_coding | 0.0013948 | 0.0043576 |
| CEMIP2 | protein_coding | 0.00135957 | 0.00605649 |
| NTPCR | protein_coding | 0.00132088 | 0.01168762 |
| LMBR1 | protein_coding | 0.00126657 | 0.0138877 |
| NFATC2IP | protein_coding | 0.00123838 | 0.01675747 |
| AATF | protein_coding | 0.00117996 | 0.01538654 |
| HSF1 | protein_coding | 0.00104782 | 0.01148452 |
| EMC7 | protein_coding | 0.00103474 | 0.0117625 |
| CSNK2B | protein_coding | 0.00101506 | 0.00518358 |
| PTK2 | protein_coding | 0.00096694 | 0.0144037 |
| BEND3 | protein_coding | 0.00096554 | 0.00536309 |
| RAP1GDS1 | protein_coding | 0.00091364 | 0.00935433 |
| HNRNPU | protein_coding | 0.00090583 | 0.01499633 |
| TRIM8 | protein_coding | 0.00076511 | 0.00506473 |
| MACROD2 | protein_coding | 0.00074163 | 0.00457487 |
| TMEM240 | protein_coding | 0.00065008 | 0.00052701 |
| MARS2 | protein_coding | 0.00064733 | 0.0034883 |
| LINC02449 | lncRNA | 0.00056476 | 0.00014825 |
| RP11-278A2 | lncRNA | 0.00047006 | 0.00093467 |
| KIAA1191 | protein_coding | 0.0004599 | 0.00410226 |
| ATP6AP2 | protein_coding | 0.00043397 | 0.00462841 |
| PRPF4B | protein_coding | 0.00030285 | 0.00381613 |
| UVSSA | protein_coding | 0.00017184 | 0.001568 |
| SAP30L | protein_coding | 0.00015872 | 0.00152573 |
| RTL6 | protein_coding | 0.00012208 | 0.00173824 |
| SOD1 | protein_coding | 0.00011513 | 0.00175621 |
| PHB2 | protein_coding | 2.76E-05 | 0.00045886 |
| VHL | protein_coding | 1.87E-05 | 0.00016896 |
| ZSWIM3 | protein_coding | -6.51E-05 | 0.00032817 |
| ATG4C | protein_coding | -6.92E-05 | 0.00031635 |
| ZDHHC17 | protein_coding | -0.0001003 | 0.0008543 |
| NDST4 | protein_coding | -0.0002962 | 0.00073776 |
| CRTC3 | protein_coding | -0.000316 | 0.00311419 |
| MED7 | protein_coding | -0.0003633 | 0.0018041 |
| CTD-2302E2 | lncRNA | -0.0004065 | 0.0004558 |
| CDK16 | protein_coding | -0.0004206 | 0.00534732 |
| CASP6 | protein_coding | -0.0004492 | 0.00050733 |
| USP1 | protein_coding | -0.0005571 | 0.00907398 |
| KNSTRN | protein_coding | -0.0005655 | 0.00582498 |
| ZNF83 | protein_coding | -0.000619 | 0.00469592 |
| ARMC2 | protein_coding | -0.000698 | 0.00160369 |
| HULC | lncRNA | -0.0007308 | 0.00099831 |

| | | | |
|---|---|---|---|
| HOXA2 | protein_coding | -0.0007849 | 0.0014987 |
| GTF3A | protein_coding | -0.0009179 | 0.01237855 |
| MICA | protein_coding | -0.0009726 | 0.00102503 |
| RP11-550I24 | lncRNA | -0.000974 | 0.00147752 |
| NTAN1 | protein_coding | -0.0010395 | 0.00660734 |
| SYF2 | protein_coding | -0.0010484 | 0.00752159 |
| ERN1 | protein_coding | -0.0011151 | 0.00627475 |
| FYTTD1 | protein_coding | -0.0011553 | 0.01584607 |
| SCAMP4 | protein_coding | -0.0011829 | 0.01200643 |
| RAB35 | protein_coding | -0.0012051 | 0.01437579 |
| RARS2 | protein_coding | -0.0012781 | 0.01303944 |
| BIN3 | protein_coding | -0.0013209 | 0.00551495 |
| RAD54B | protein_coding | -0.0013257 | 0.00535666 |
| LINC01833 | lncRNA | -0.0013414 | 0.00420427 |
| SMYD4 | protein_coding | -0.0014401 | 0.00966271 |
| CTD-2017D1: | lncRNA | -0.0014762 | 0.00499171 |
| CKMT1A | protein_coding | -0.0015595 | 0.00795999 |
| RHOC | protein_coding | -0.00159 | 0.00056967 |
| FDX1 | protein_coding | -0.0016069 | 0.00712932 |
| TWF2 | protein_coding | -0.0016876 | 0.00830601 |
| RBFOX2 | protein_coding | -0.0016929 | 0.0297323 |
| RPRD2 | protein_coding | -0.0017522 | 0.02334031 |
| ZKSCAN2 | protein_coding | -0.0018448 | 0.01979436 |
| MPLKIP | protein_coding | -0.0018786 | 0.01320666 |
| IFT122 | protein_coding | -0.0018956 | 0.01474974 |
| ZNF417 | protein_coding | -0.0019139 | 0.01266161 |
| POMT2 | protein_coding | -0.0019547 | 0.02106498 |
| H2AC20 | protein_coding | -0.0020341 | 0.0018091 |
| NDUFA12 | protein_coding | -0.0021394 | 0.02009774 |
| RP11-547D1: | lncRNA | -0.0022011 | 0.0020628 |
| IMP3 | protein_coding | -0.0022166 | 0.01920203 |
| ZRANB3 | protein_coding | -0.0022641 | 0.01225844 |
| PPP2R3C | protein_coding | -0.0023003 | 0.0171182 |
| ZNF765 | protein_coding | -0.0023046 | 0.01217308 |
| MAP4K3 | protein_coding | -0.0025012 | 0.0218604 |
| RP11-640F2: | lncRNA | -0.0026498 | 0.00526956 |
| ZNF761 | protein_coding | -0.0026527 | 0.02209505 |
| GZF1 | protein_coding | -0.0026594 | 0.02677495 |
| NSDHL | protein_coding | -0.0027266 | 0.02172297 |
| TLNRD1 | protein_coding | -0.0027583 | 0.00893793 |
| KLHL28 | protein_coding | -0.0027629 | 0.01868337 |
| SLC38A9 | protein_coding | -0.002781 | 0.01529682 |
| TGS1 | protein_coding | -0.0027953 | 0.01778618 |
| CCDC77 | protein_coding | -0.0027962 | 0.01831862 |
| LOXL3 | protein_coding | -0.0028485 | 0.01990617 |

| | | | |
|---|---|---|---|
| SYNGAP1 | protein_coding | -0.002871 | 0.01271139 |
| MTR | protein_coding | -0.0028983 | 0.04000446 |
| LRRC8A | protein_coding | -0.002981 | 0.03533907 |
| NSD3 | protein_coding | -0.0030242 | 0.03612879 |
| CEP295 | protein_coding | -0.0030357 | 0.03543399 |
| RNASEH1 | protein_coding | -0.0030582 | 0.03789197 |
| INPP5A | protein_coding | -0.0031249 | 0.0149309 |
| AKR1B1 | protein_coding | -0.0031263 | 0.03307191 |
| RPL19 | protein_coding | -0.0031948 | 0.0680627 |
| SDE2 | protein_coding | -0.0032772 | 0.02915879 |
| PTGR2 | protein_coding | -0.0032984 | 0.01154308 |
| YEATS2 | protein_coding | -0.003317 | 0.06194673 |
| ZNF430 | protein_coding | -0.0033302 | 0.02664937 |
| XPC | protein_coding | -0.0033486 | 0.02370261 |
| TARS2 | protein_coding | -0.0033683 | 0.02924534 |
| SYDE2 | protein_coding | -0.003383 | 0.03194698 |
| TMEM259 | protein_coding | -0.0036216 | 0.02776945 |
| CISD2 | protein_coding | -0.0036587 | 0.04017346 |
| DBF4 | protein_coding | -0.0038503 | 0.03656114 |
| WBP2 | protein_coding | -0.0038562 | 0.04551032 |
| UNK | protein_coding | -0.003862 | 0.03586374 |
| KPTN | protein_coding | -0.0041748 | 0.03055424 |
| DNAJC14 | protein_coding | -0.0043439 | 0.04604964 |
| ZNF304 | protein_coding | -0.004344 | 0.03790528 |
| HSPA12A | protein_coding | -0.0043464 | 0.04060593 |
| SNX30 | protein_coding | -0.0045054 | 0.03414477 |
| RPN1 | protein_coding | -0.0045655 | 0.08968903 |
| PIGG | protein_coding | -0.0045719 | 0.0502231 |
| CNBP | protein_coding | -0.0046119 | 0.08318844 |
| TMSB10 | protein_coding | -0.0046342 | 0.0708543 |
| RNF183 | protein_coding | -0.0046399 | 0.00490746 |
| USP3 | protein_coding | -0.0046674 | 0.03806724 |
| FRMD6 | protein_coding | -0.0046747 | 0.02285005 |
| PSMG3 | protein_coding | -0.0047076 | 0.0518944 |
| TMEM179B | protein_coding | -0.0049092 | 0.03493646 |
| RANBP3 | protein_coding | -0.0049923 | 0.05028684 |
| RP5-908M14 | lncRNA | -0.0050114 | 0.01209805 |
| NEGR1 | protein_coding | -0.0050181 | 0.04233446 |
| CNOT3 | protein_coding | -0.0050902 | 0.06503651 |
| ASXL2 | protein_coding | -0.0051847 | 0.06215471 |
| RP11-400F1[illegible] | lncRNA | -0.0051885 | 0.00740932 |
| EPG5 | protein_coding | -0.0052492 | 0.0478719 |
| MTBP | protein_coding | -0.0052523 | 0.03793913 |
| FOXP2 | protein_coding | -0.0052772 | 0.03034321 |
| CENPN | protein_coding | -0.0053003 | 0.04826303 |

| | | | |
|---|---|---|---|
| SIPA1L2 | protein_coding | -0.0053425 | 0.04614762 |
| SCYL1 | protein_coding | -0.0053463 | 0.05361681 |
| DRAIC | lncRNA | -0.0053626 | 0.00590259 |
| CRADD | protein_coding | -0.0053699 | 0.02213921 |
| R3HDM4 | protein_coding | -0.0054249 | 0.05586301 |
| TNFSF13 | protein_coding | -0.0054419 | 0.00462723 |
| HIBADH | protein_coding | -0.0054443 | 0.04367611 |
| LGSN | protein_coding | -0.0054585 | 0.02140421 |
| OSTF1 | protein_coding | -0.0055053 | 0.03607348 |
| PPIP5K2 | protein_coding | -0.0055649 | 0.06529415 |
| LINGO1 | protein_coding | -0.005582 | 0.04426498 |
| MOB1A | protein_coding | -0.0057224 | 0.07625161 |
| PPP2R2D | protein_coding | -0.005763 | 0.0386051 |
| ABI1 | protein_coding | -0.0058167 | 0.04616027 |
| CCDC126 | protein_coding | -0.0058289 | 0.02011104 |
| ZNF830 | protein_coding | -0.0059355 | 0.03639148 |
| NADSYN1 | protein_coding | -0.0059408 | 0.04123569 |
| MTSS2 | protein_coding | -0.0059746 | 0.05565492 |
| PRMT2 | protein_coding | -0.0060394 | 0.05622242 |
| NAF1 | protein_coding | -0.0061769 | 0.04089856 |
| TMED7 | protein_coding | -0.0062613 | 0.02080546 |
| NUDT3 | protein_coding | -0.0063841 | 0.07679347 |
| RP11-468E2. | protein_coding | -0.0063863 | 0.01289081 |
| USP39 | protein_coding | -0.0064507 | 0.10286952 |
| ZIM2 | protein_coding | -0.0064707 | 0.00802728 |
| RUSC1 | protein_coding | -0.0065128 | 0.0705917 |
| MBTPS2 | protein_coding | -0.0065175 | 0.042736 |
| ZNF844 | protein_coding | -0.0066274 | 0.01059281 |
| SYNE4 | protein_coding | -0.0066315 | 0.00713939 |
| NUBPL | protein_coding | -0.0066494 | 0.0355551 |
| TMEM158 | protein_coding | -0.0066695 | 0.00677101 |
| SPOP | protein_coding | -0.006694 | 0.07530935 |
| ZNF567 | protein_coding | -0.0067242 | 0.04888195 |
| RPP21 | protein_coding | -0.0069289 | 0.01266898 |
| TOP2B | protein_coding | -0.0069559 | 0.11631557 |
| CYP2R1 | protein_coding | -0.0069571 | 0.04138228 |
| RAB22A | protein_coding | -0.0070536 | 0.0643968 |
| C2orf49-DT | lncRNA | -0.0071076 | 0.02914141 |
| SOAT1 | protein_coding | -0.007127 | 0.03932285 |
| ELP1 | protein_coding | -0.0072075 | 0.10659269 |
| TMEM151A | protein_coding | -0.0072172 | 0.02100102 |
| PRCC | protein_coding | -0.0072546 | 0.09318583 |
| SAFB2 | protein_coding | -0.0073003 | 0.06235659 |
| CEBPZ | protein_coding | -0.0075466 | 0.12385387 |
| SOX6 | protein_coding | -0.00757 | 0.09777849 |

| | | | |
|---|---|---|---|
| C5orf24 | protein_coding | -0.0076008 | 0.05770272 |
| AP1B1 | protein_coding | -0.0076097 | 0.11707558 |
| MDM2 | protein_coding | -0.0076103 | 0.08623391 |
| TAOK1 | protein_coding | -0.0077142 | 0.10888949 |
| NSL1 | protein_coding | -0.0077532 | 0.07638942 |
| TENT4A | protein_coding | -0.0078441 | 0.09285767 |
| GRK2 | protein_coding | -0.0079221 | 0.10966051 |
| CREBZF | protein_coding | -0.0080064 | 0.10532505 |
| FUNDC1 | protein_coding | -0.0080272 | 0.03980144 |
| SLX4 | protein_coding | -0.0080768 | 0.06911923 |
| SEC63 | protein_coding | -0.0081428 | 0.09350909 |
| RBM19 | protein_coding | -0.0081884 | 0.06699309 |
| ARFIP2 | protein_coding | -0.0081973 | 0.09096621 |
| RBM38 | protein_coding | -0.0082057 | 0.07279595 |
| AP1G2 | protein_coding | -0.0082654 | 0.08010643 |
| ZNF814 | protein_coding | -0.0083723 | 0.05409235 |
| ZNF526 | protein_coding | -0.0084908 | 0.05307709 |
| GOLGA4 | protein_coding | -0.0085231 | 0.12264469 |
| CDK13 | protein_coding | -0.0085365 | 0.09470018 |
| UBQLN2 | protein_coding | -0.0085689 | 0.08160346 |
| ARFGEF3 | protein_coding | -0.0085719 | 0.06264715 |
| C1orf112 | protein_coding | -0.0086268 | 0.05403276 |
| DNAJC18 | protein_coding | -0.0086934 | 0.05706175 |
| SLC35A4 | protein_coding | -0.0087984 | 0.12984201 |
| CCT5 | protein_coding | -0.0088022 | 0.19069279 |
| SLC39A10 | protein_coding | -0.0088307 | 0.05714242 |
| ZFAND6 | protein_coding | -0.0088489 | 0.06800464 |
| ALG13 | protein_coding | -0.0089575 | 0.07692575 |
| KLF7 | protein_coding | -0.0089767 | 0.10397794 |
| SNRPF | protein_coding | -0.0090303 | 0.12125675 |
| FASTKD5 | protein_coding | -0.0091052 | 0.07830953 |
| RP11-647K16 | lncRNA | -0.0091151 | 0.01036749 |
| FZD3 | protein_coding | -0.0092458 | 0.06131076 |
| CDIP1 | protein_coding | -0.0092596 | 0.08601267 |
| SREBF2 | protein_coding | -0.0093844 | 0.18828969 |
| ALDH9A1 | protein_coding | -0.0094601 | 0.10299026 |
| CREBL2 | protein_coding | -0.009515 | 0.1020563 |
| RAMAC | protein_coding | -0.0095274 | 0.0542791 |
| RP11-567P17 | lncRNA | -0.0095618 | 0.01404772 |
| BZW2 | protein_coding | -0.0095726 | 0.12500686 |
| MTRNR2L8 | protein_coding | -0.0095782 | 0.00714724 |
| CSK | protein_coding | -0.0095956 | 0.11502722 |
| PNPLA8 | protein_coding | -0.0096575 | 0.11537168 |
| CNST | protein_coding | -0.0096669 | 0.12055874 |
| ZNF491 | protein_coding | -0.0097765 | 0.03865317 |

| | | | |
|---|---|---|---|
| C6orf52 | protein_coding | -0.0098396 | 0.00899771 |
| NAGA | protein_coding | -0.0098701 | 0.09495213 |
| SATB1 | protein_coding | -0.0100014 | 0.1323065 |
| SYCE1L | protein_coding | -0.0101055 | 0.01537002 |
| TSR3 | protein_coding | -0.010119 | 0.06197609 |
| SBK1 | protein_coding | -0.0101221 | 0.13068504 |
| SPTLC1 | protein_coding | -0.0101484 | 0.11492818 |
| KNTC1 | protein_coding | -0.010166 | 0.15039087 |
| ELP3 | protein_coding | -0.0101741 | 0.09342674 |
| VARS2 | protein_coding | -0.0101991 | 0.01374154 |
| CEP70 | protein_coding | -0.0102123 | 0.08721956 |
| GINM1 | protein_coding | -0.0102282 | 0.06307404 |
| SCRN2 | protein_coding | -0.0102523 | 0.05910183 |
| RB1CC1 | protein_coding | -0.0102549 | 0.06191989 |
| RBMX2 | protein_coding | -0.0104175 | 0.07759317 |
| TRAIP | protein_coding | -0.0104999 | 0.05713043 |
| SLC39A9 | protein_coding | -0.0105333 | 0.18520521 |
| IGFLR1 | protein_coding | -0.0105374 | 0.01673737 |
| CTD-2553L13 | lncRNA | -0.0105826 | 0.01231487 |
| MARCHF2 | protein_coding | -0.0106146 | 0.04137447 |
| CRNKL1 | protein_coding | -0.0107558 | 0.10828348 |
| PGM2 | protein_coding | -0.0107772 | 0.10440001 |
| GSS | protein_coding | -0.0107825 | 0.082144 |
| PAXBP1 | protein_coding | -0.0108598 | 0.10733389 |
| NDUFA10 | protein_coding | -0.0108766 | 0.14481532 |
| CAPN5 | protein_coding | -0.0108897 | 0.04973129 |
| POP5 | protein_coding | -0.0110404 | 0.08964578 |
| MVB12A | protein_coding | -0.0110538 | 0.09751801 |
| FANCL | protein_coding | -0.0110791 | 0.13251244 |
| FOCAD | protein_coding | -0.011101 | 0.0953401 |
| NLK | protein_coding | -0.0111162 | 0.10239099 |
| GLYCTK | protein_coding | -0.0111983 | 0.02433974 |
| ZNF573 | protein_coding | -0.0112034 | 0.0335109 |
| CAPRIN1 | protein_coding | -0.0112351 | 0.26168568 |
| SQLE | protein_coding | -0.0113167 | 0.14903841 |
| SRRT | protein_coding | -0.0113807 | 0.18243881 |
| KIF14 | protein_coding | -0.0114678 | 0.1159054 |
| KCNH6 | protein_coding | -0.0115449 | 0.22330997 |
| CTD-2313J17 | lncRNA | -0.0115526 | 0.01210635 |
| LRRC57 | protein_coding | -0.0116175 | 0.08012108 |
| FADD | protein_coding | -0.0116205 | 0.07672317 |
| MOSPD1 | protein_coding | -0.0117349 | 0.06095547 |
| DKC1 | protein_coding | -0.0117459 | 0.20582481 |
| ELP4 | protein_coding | -0.0117572 | 0.10351116 |
| NOL4L | protein_coding | -0.0117905 | 0.12557368 |

| | | | |
|---|---|---|---|
| S100A4 | protein_coding | -0.0118954 | 0.04885474 |
| DHX35 | protein_coding | -0.0119787 | 0.10564296 |
| RNF13 | protein_coding | -0.0120546 | 0.08558546 |
| RP11-328C17 | lncRNA | -0.0120853 | 0.01588643 |
| TOR4A | protein_coding | -0.0120909 | 0.06144508 |
| SRSF11 | protein_coding | -0.0120953 | 0.14814876 |
| WAC-AS1 | lncRNA | -0.012112 | 0.10167145 |
| SLC25A51 | protein_coding | -0.0122445 | 0.07913405 |
| DYNC1LI1 | protein_coding | -0.012275 | 0.145433 |
| CDIN1 | protein_coding | -0.0123486 | 0.0849645 |
| NEMP2 | protein_coding | -0.0123802 | 0.1022197 |
| RP11-390P2. | lncRNA | -0.0124327 | 0.01277133 |
| NXF1 | protein_coding | -0.0124502 | 0.18714614 |
| YOD1 | protein_coding | -0.0127146 | 0.08265831 |
| TXLNG | protein_coding | -0.0127198 | 0.16103169 |
| CDH3 | protein_coding | -0.0129148 | 0.09295048 |
| HAUS2 | protein_coding | -0.0130902 | 0.1782499 |
| SEMA3G | protein_coding | -0.0131612 | 0.01733094 |
| RP11-111N2( | lncRNA | -0.0131937 | 0.01690288 |
| GOPC | protein_coding | -0.013232 | 0.13521286 |
| ZNF766 | protein_coding | -0.013325 | 0.15464359 |
| PARTICL | lncRNA | -0.0133368 | 0.04363106 |
| L3MBTL1 | protein_coding | -0.0133444 | 0.04605248 |
| SIRT2 | protein_coding | -0.0134457 | 0.17766594 |
| TSHZ3 | protein_coding | -0.0134841 | 0.05740494 |
| PEX11B | protein_coding | -0.0134936 | 0.09368472 |
| NPC2 | protein_coding | -0.013516 | 0.09999777 |
| SPTLC2 | protein_coding | -0.0135193 | 0.27400494 |
| CDR2 | protein_coding | -0.0135237 | 0.11134087 |
| PRPF38B | protein_coding | -0.0135335 | 0.18775621 |
| CDC26 | protein_coding | -0.0136542 | 0.04751397 |
| RAB7A | protein_coding | -0.0136758 | 0.25251588 |
| GUCY1B1 | protein_coding | -0.0138756 | 0.10913033 |
| BPGM | protein_coding | -0.0138903 | 0.11427273 |
| WBP1 | protein_coding | -0.014019 | 0.05549106 |
| CRYZL1 | protein_coding | -0.0140227 | 0.09888031 |
| POP4 | protein_coding | -0.0140494 | 0.15029115 |
| ATRIP | protein_coding | -0.0140622 | 0.06895075 |
| RP11-252I13 | lncRNA | -0.0140923 | 0.01680741 |
| RNF40 | protein_coding | -0.014126 | 0.21777477 |
| MBTD1 | protein_coding | -0.0143032 | 0.1825573 |
| ALG8 | protein_coding | -0.0143132 | 0.12347498 |
| FAM50B | protein_coding | -0.0143308 | 0.06007563 |
| NPEPPS | protein_coding | -0.0144651 | 0.23593072 |
| HOXC9 | protein_coding | -0.0145491 | 0.02062743 |

| | | | |
|---|---|---|---|
| CTSK | protein_coding | -0.0145635 | 0.04301341 |
| ORAI2 | protein_coding | -0.014565 | 0.18294824 |
| TNPO1 | protein_coding | -0.0146141 | 0.16030816 |
| AAR2 | protein_coding | -0.0147133 | 0.15245648 |
| ANAPC16 | protein_coding | -0.0147464 | 0.15175669 |
| TMCO3 | protein_coding | -0.0147843 | 0.1316226 |
| DGKH | protein_coding | -0.0148745 | 0.13441135 |
| SASS6 | protein_coding | -0.0148904 | 0.10387764 |
| TRIM65 | protein_coding | -0.0149159 | 0.10945013 |
| MAST3 | protein_coding | -0.0149732 | 0.10884215 |
| GS1-24F4.2 | lncRNA | -0.0150009 | 0.01985716 |
| OGDH | protein_coding | -0.0150054 | 0.26310405 |
| RPL36 | protein_coding | -0.0150304 | 0.22960754 |
| NMNAT3 | protein_coding | -0.0150342 | 0.08115868 |
| KMT2B | protein_coding | -0.0151386 | 0.21158939 |
| NPM1 | protein_coding | -0.0152584 | 0.39383272 |
| DIAPH1 | protein_coding | -0.0152655 | 0.29183472 |
| MYEF2 | protein_coding | -0.0152804 | 0.23775461 |
| DENND1B | protein_coding | -0.0153403 | 0.09478591 |
| SRP54 | protein_coding | -0.0154494 | 0.18602109 |
| CTSA | protein_coding | -0.0155366 | 0.14617807 |
| VAMP7 | protein_coding | -0.0155519 | 0.11941358 |
| AC092835.2 | protein_coding | -0.0156519 | 0.05987284 |
| COIL | protein_coding | -0.0156902 | 0.14900675 |
| SVIP | protein_coding | -0.015745 | 0.18073869 |
| TOMM40L | protein_coding | -0.0157694 | 0.16788026 |
| RP11-284E5. | lncRNA | -0.0158468 | 0.04558477 |
| CLDND1 | protein_coding | -0.0158728 | 0.11430953 |
| RP11-74E22. | lncRNA | -0.0158935 | 0.07757865 |
| ACTR5 | protein_coding | -0.0159085 | 0.1325335 |
| YWHAG | protein_coding | -0.0159567 | 0.37227119 |
| DCAF12 | protein_coding | -0.0161608 | 0.23654336 |
| SLC49A4 | protein_coding | -0.0161701 | 0.07438665 |
| PIK3R4 | protein_coding | -0.0162128 | 0.17391382 |
| TDRD10 | protein_coding | -0.0163077 | 0.07593187 |
| ZNF687 | protein_coding | -0.0164114 | 0.17948633 |
| B3GALT1 | protein_coding | -0.0164256 | 0.03410592 |
| ZC3H13 | protein_coding | -0.0165898 | 0.21851818 |
| SULT4A1 | protein_coding | -0.016605 | 0.17463436 |
| SCRT2 | protein_coding | -0.0167584 | 0.09700601 |
| EFHD2 | protein_coding | -0.0167672 | 0.17111299 |
| RNF41 | protein_coding | -0.016901 | 0.22047501 |
| RFXANK | protein_coding | -0.0169239 | 0.14660903 |
| ZNF738 | protein_coding | -0.0169287 | 0.06330522 |
| RNF144A | protein_coding | -0.0170063 | 0.14662056 |

| | | | |
|---|---|---|---|
| SAV1 | protein_coding | -0.0170418 | 0.14698881 |
| DNAJC5 | protein_coding | -0.0172149 | 0.29389846 |
| ZNF576 | protein_coding | -0.0172394 | 0.11811096 |
| ZNF341 | protein_coding | -0.0173211 | 0.06724915 |
| SMTN | protein_coding | -0.0173461 | 0.08672521 |
| OPA1 | protein_coding | -0.0174281 | 0.37234201 |
| MFF-DT | lncRNA | -0.0174614 | 0.02477736 |
| OIP5-AS1 | lncRNA | -0.0174959 | 0.34117608 |
| ANKRD16 | protein_coding | -0.0175601 | 0.06686064 |
| RP11-57H14. | lncRNA | -0.0175671 | 0.064181 |
| AC115522.3 | lncRNA | -0.0175732 | 0.01691544 |
| ZNF781 | protein_coding | -0.0175746 | 0.02790459 |
| FAM95C | protein_coding | -0.0175806 | 0.07709466 |
| ZNF518A | protein_coding | -0.0176558 | 0.19672451 |
| NSA2 | protein_coding | -0.01767 | 0.20107978 |
| BLOC1S4 | protein_coding | -0.0176748 | 0.09212965 |
| EBF3 | protein_coding | -0.0177967 | 0.17001102 |
| UROS | protein_coding | -0.0178556 | 0.13618416 |
| RP11-338N1( | lncRNA | -0.0178801 | 0.0180799 |
| PUS3 | protein_coding | -0.0178803 | 0.14413788 |
| PLIN4 | protein_coding | -0.0178952 | 0.03427484 |
| SUV39H2 | protein_coding | -0.017926 | 0.10667166 |
| SLC25A37 | protein_coding | -0.017974 | 0.135063 |
| SLC25A24 | protein_coding | -0.0180151 | 0.08341153 |
| SFSWAP | protein_coding | -0.018025 | 0.22777505 |
| TOMM22 | protein_coding | -0.018139 | 0.28835094 |
| DOLPP1 | protein_coding | -0.0182285 | 0.22919541 |
| TMEM251 | protein_coding | -0.0182503 | 0.1083684 |
| UMAD1 | protein_coding | -0.0182541 | 0.06829466 |
| DYNLL1 | protein_coding | -0.0183065 | 0.34472677 |
| PPP1R7 | protein_coding | -0.018406 | 0.24167183 |
| PSME3IP1 | protein_coding | -0.0184091 | 0.27404174 |
| HINFP | protein_coding | -0.0185229 | 0.18211396 |
| IFT46 | protein_coding | -0.0186522 | 0.14732178 |
| RP11-110G2 | lncRNA | -0.0187446 | 0.04326792 |
| SCAMP5 | protein_coding | -0.0188054 | 0.25726366 |
| ASPM | protein_coding | -0.0188104 | 0.37236161 |
| RP11-354K4. | lncRNA | -0.0189138 | 0.02083803 |
| RAD9A | protein_coding | -0.0189759 | 0.16995275 |
| KPNB1 | protein_coding | -0.0190553 | 0.54962108 |
| BCL2L12 | protein_coding | -0.0190644 | 0.04391319 |
| HSPD1 | protein_coding | -0.0191294 | 0.52742474 |
| LRP5 | protein_coding | -0.0191329 | 0.19349233 |
| ATP5PB | protein_coding | -0.0191363 | 0.37745017 |
| GSK3A | protein_coding | -0.0191839 | 0.22570577 |

| | | | |
|---|---|---|---|
| DDX3Y | protein_coding | -0.0191871 | 0.21903627 |
| SMIM26 | protein_coding | -0.0192018 | 0.08547584 |
| HS6ST1 | protein_coding | -0.0192828 | 0.20744034 |
| AC142528.1 | lncRNA | -0.0193356 | 0.04033004 |
| CDH7 | protein_coding | -0.0193768 | 0.08390201 |
| MRAP2 | protein_coding | -0.0194519 | 0.08291029 |
| LMF2 | protein_coding | -0.0195267 | 0.29135985 |
| RETREG3 | protein_coding | -0.0196201 | 0.2278127 |
| PGF | protein_coding | -0.0196289 | 0.05262865 |
| LENG1 | protein_coding | -0.0196424 | 0.07885315 |
| NDUFS2 | protein_coding | -0.0196572 | 0.3753151 |
| TCF20 | protein_coding | -0.0197747 | 0.31400473 |
| XKR7 | protein_coding | -0.0198071 | 0.28620634 |
| CNOT6L | protein_coding | -0.0198714 | 0.20917545 |
| RECQL4 | protein_coding | -0.0199237 | 0.2564942 |
| GNAQ | protein_coding | -0.0199332 | 0.23596189 |
| RP11-279O9. | lncRNA | -0.0199352 | 0.02890696 |
| CAPN10 | protein_coding | -0.0200312 | 0.1539589 |
| PRSS53 | protein_coding | -0.0200337 | 0.07549184 |
| TMEM263 | protein_coding | -0.0200428 | 0.12815508 |
| RP11-474G2 | lncRNA | -0.0200988 | 0.02060907 |
| RP11-983P16 | lncRNA | -0.0201669 | 0.07414061 |
| FBXO9 | protein_coding | -0.0201851 | 0.27538815 |
| SNRPE | protein_coding | -0.0201871 | 0.26204746 |
| IRF3 | protein_coding | -0.020206 | 0.21052166 |
| CST3 | protein_coding | -0.0202538 | 0.13463192 |
| PARP16 | protein_coding | -0.0202993 | 0.19195426 |
| FAM13A-AS1 | lncRNA | -0.0203575 | 0.0329673 |
| TMEM154 | protein_coding | -0.0204343 | 0.02122118 |
| AP002954.3 | lncRNA | -0.0204453 | 0.0213589 |
| SRR | protein_coding | -0.0205099 | 0.16115535 |
| VAMP2 | protein_coding | -0.0205243 | 0.22984034 |
| ZNF654 | protein_coding | -0.020622 | 0.13363563 |
| KDM4C | protein_coding | -0.0206372 | 0.20499911 |
| REV1 | protein_coding | -0.0206709 | 0.275051 |
| ASB7 | protein_coding | -0.0207274 | 0.16800044 |
| RP11-488L18 | lncRNA | -0.0207555 | 0.02806748 |
| DBR1 | protein_coding | -0.0207718 | 0.21368518 |
| SNRPB2 | protein_coding | -0.0207954 | 0.23835821 |
| IPPK | protein_coding | -0.0207954 | 0.16179929 |
| PCAT7 | lncRNA | -0.0208789 | 0.02647834 |
| ZBTB4 | protein_coding | -0.0208832 | 0.11259352 |
| ESPL1 | protein_coding | -0.0209407 | 0.30210067 |
| PTPRM | protein_coding | -0.0209478 | 0.04769528 |
| RP11-195F19 | protein_coding | -0.0210208 | 0.02023871 |

| | | | |
|---|---|---|---|
| GUF1 | protein_coding | -0.0210454 | 0.26664682 |
| PIGQ | protein_coding | -0.0210707 | 0.13805342 |
| CSTF2T | protein_coding | -0.0210874 | 0.21115314 |
| NCBP3 | protein_coding | -0.0212889 | 0.25497447 |
| RP11-109N2 | lncRNA | -0.0213642 | 0.02458535 |
| AC008982.2 | lncRNA | -0.0214042 | 0.03050845 |
| RBM10 | protein_coding | -0.0214094 | 0.29295861 |
| PHF5A | protein_coding | -0.0214427 | 0.24802479 |
| CCDC137 | protein_coding | -0.0214923 | 0.28370835 |
| ZRANB1 | protein_coding | -0.0215312 | 0.21633814 |
| WSB2 | protein_coding | -0.0215384 | 0.36037388 |
| EFNA3 | protein_coding | -0.0216608 | 0.15908175 |
| METTL18 | protein_coding | -0.021699 | 0.08012505 |
| C6orf47 | protein_coding | -0.0217059 | 0.17370436 |
| STK4 | protein_coding | -0.0217135 | 0.30414938 |
| COX18 | protein_coding | -0.0217913 | 0.16904094 |
| ZSCAN32 | protein_coding | -0.0218296 | 0.1726323 |
| RP11-156P1. | lncRNA | -0.0218806 | 0.1996917 |
| ECE1 | protein_coding | -0.0218863 | 0.26247216 |
| MTMR12 | protein_coding | -0.0219024 | 0.26236538 |
| MSRB2 | protein_coding | -0.0219187 | 0.07177261 |
| ANKRD13A | protein_coding | -0.0219461 | 0.25345045 |
| MC1R | protein_coding | -0.0219478 | 0.07007805 |
| KXD1 | protein_coding | -0.0219578 | 0.27771952 |
| MAP4K5 | protein_coding | -0.0221675 | 0.24269174 |
| SS18 | protein_coding | -0.0222144 | 0.24975585 |
| RPF1 | protein_coding | -0.0222184 | 0.26967402 |
| PYM1 | protein_coding | -0.0222289 | 0.15101008 |
| MRPL48 | protein_coding | -0.0223363 | 0.20948312 |
| PCMT1 | protein_coding | -0.0223524 | 0.28304899 |
| HJURP | protein_coding | -0.0223668 | 0.31524524 |
| AP2A1 | protein_coding | -0.0224087 | 0.24916329 |
| UBE4B | protein_coding | -0.0224387 | 0.22279713 |
| COL4A4 | protein_coding | -0.0225273 | 0.03800131 |
| MTERF3 | protein_coding | -0.0225586 | 0.20826425 |
| UPF3B | protein_coding | -0.0225964 | 0.32917248 |
| FGF12-AS3 | lncRNA | -0.0225989 | 0.06370802 |
| LRTM2 | protein_coding | -0.0225993 | 0.15035767 |
| CDC37 | protein_coding | -0.0226056 | 0.35660961 |
| BLOC1S5 | protein_coding | -0.0226224 | 0.17585528 |
| PLEKHG4B | protein_coding | -0.0226252 | 0.35918201 |
| MT-ND6 | protein_coding | -0.0226346 | 0.19573598 |
| FTSJ3 | protein_coding | -0.0226625 | 0.32604625 |
| SEC11A | protein_coding | -0.0227121 | 0.34148199 |
| ARL17B | protein_coding | -0.0227252 | 0.02952653 |

| | | | |
|---|---|---|---|
| LIPE | protein_coding | -0.0227896 | 0.12270912 |
| LINC01775 | lncRNA | -0.0228058 | 0.020695 |
| RP5-967N21. | lncRNA | -0.0228242 | 0.04728879 |
| DDX56 | protein_coding | -0.0228496 | 0.31530902 |
| YWHAH | protein_coding | -0.022963 | 0.39582453 |
| SLC22A5 | protein_coding | -0.0229944 | 0.16390969 |
| VPS4A | protein_coding | -0.0229961 | 0.28930033 |
| SS18L2 | protein_coding | -0.0230243 | 0.11407291 |
| XPO6 | protein_coding | -0.023037 | 0.44007601 |
| GTF3C2 | protein_coding | -0.0231057 | 0.38124923 |
| NEK7 | protein_coding | -0.0232034 | 0.23314462 |
| VIPAS39 | protein_coding | -0.0232967 | 0.27145681 |
| KRTCAP3 | protein_coding | -0.0233378 | 0.1070918 |
| TRIM24 | protein_coding | -0.0233439 | 0.45758428 |
| PRIM2 | protein_coding | -0.0233863 | 0.20056611 |
| CCNB2 | protein_coding | -0.0233877 | 0.36415071 |
| SLC25A17 | protein_coding | -0.0234231 | 0.26827018 |
| FEZ2 | protein_coding | -0.0234518 | 0.17804663 |
| ZWILCH | protein_coding | -0.0234839 | 0.27108961 |
| CASP2 | protein_coding | -0.0235003 | 0.4087954 |
| RP11-464F9. | lncRNA | -0.0235129 | 0.02353491 |
| KHDRBS1 | protein_coding | -0.0235244 | 0.57228959 |
| SREK1IP1 | protein_coding | -0.0235266 | 0.14840245 |
| YTHDF1 | protein_coding | -0.0235463 | 0.36491671 |
| TOX2 | protein_coding | -0.0235649 | 0.15589774 |
| CDC42BPG | protein_coding | -0.0235862 | 0.23327645 |
| CCNDBP1 | protein_coding | -0.0236222 | 0.18589706 |
| ITFG2 | protein_coding | -0.0236283 | 0.24860408 |
| REPS1 | protein_coding | -0.0236309 | 0.17723833 |
| RP11-398K2 | lncRNA | -0.0237041 | 0.17791208 |
| UBE2W | protein_coding | -0.0237429 | 0.11848835 |
| LINC01184 | lncRNA | -0.0237478 | 0.17694864 |
| NEMP1 | protein_coding | -0.0238838 | 0.44289933 |
| CARD19 | protein_coding | -0.0239214 | 0.13850629 |
| CHST7 | protein_coding | -0.0239712 | 0.11663771 |
| FARSB | protein_coding | -0.0239901 | 0.49216749 |
| EIF3I | protein_coding | -0.0241704 | 0.5446236 |
| CFAP36 | protein_coding | -0.0242445 | 0.1544189 |
| PLCE1 | protein_coding | -0.0243395 | 0.11069014 |
| AKT1S1 | protein_coding | -0.024369 | 0.35715015 |
| POLR1B | protein_coding | -0.0243772 | 0.30076319 |
| DNAH5 | protein_coding | -0.0245138 | 0.0247603 |
| MDH1 | protein_coding | -0.0246149 | 0.47452924 |
| ZNF10 | protein_coding | -0.0247119 | 0.18311327 |
| TRIM3 | protein_coding | -0.0247175 | 0.12873347 |

| | | | |
|---|---|---|---|
| BLOC1S6 | protein_coding | -0.0247287 | 0.25862861 |
| LYSMD4 | protein_coding | -0.0247411 | 0.26205829 |
| ZNF689 | protein_coding | -0.024751 | 0.22955397 |
| ZNF658 | protein_coding | -0.0248456 | 0.07477875 |
| PRUNE1 | protein_coding | -0.0248634 | 0.31415105 |
| PTS | protein_coding | -0.0248987 | 0.15342686 |
| SPRYD4 | protein_coding | -0.0249626 | 0.21003245 |
| AQR | protein_coding | -0.0250914 | 0.44816274 |
| ATP5F1A | protein_coding | -0.0251471 | 0.74352913 |
| HNRNPH3 | protein_coding | -0.02526 | 0.40586553 |
| RDH13 | protein_coding | -0.0252734 | 0.15129904 |
| MCPH1-AS1 | lncRNA | -0.0253323 | 0.08060128 |
| ELOVL5 | protein_coding | -0.0253745 | 0.30476206 |
| ATE1 | protein_coding | -0.0254261 | 0.2119693 |
| SH2B1 | protein_coding | -0.0254974 | 0.40280786 |
| ZBTB7A | protein_coding | -0.0255169 | 0.203027 |
| GATC | protein_coding | -0.0256176 | 0.31174263 |
| L3MBTL2 | protein_coding | -0.0256377 | 0.39551577 |
| RP11-121C2. | lncRNA | -0.0256853 | 0.12698394 |
| RHOBTB2 | protein_coding | -0.0257742 | 0.13797551 |
| ZNF684 | protein_coding | -0.0258603 | 0.08690723 |
| TRIM27 | protein_coding | -0.0258665 | 0.28424917 |
| COPB1 | protein_coding | -0.0258796 | 0.4012889 |
| ZNF7 | protein_coding | -0.0259555 | 0.28636867 |
| DLEU2 | lncRNA | -0.026067 | 0.10381199 |
| RAVER1 | protein_coding | -0.0260839 | 0.16690973 |
| TRMU | protein_coding | -0.0261134 | 0.32968476 |
| CHAF1B | protein_coding | -0.026187 | 0.34867328 |
| CCNQ | protein_coding | -0.0262234 | 0.15004074 |
| EP400 | protein_coding | -0.0262297 | 0.41066209 |
| ZSWIM9 | protein_coding | -0.0262352 | 0.16037976 |
| MRPL54 | protein_coding | -0.0264065 | 0.25478306 |
| GCH1 | protein_coding | -0.0264332 | 0.24935364 |
| GAK | protein_coding | -0.0264806 | 0.28701189 |
| KIAA2013 | protein_coding | -0.0264958 | 0.22598245 |
| PRKAR1A | protein_coding | -0.026499 | 0.28926847 |
| EPAS1 | protein_coding | -0.0266445 | 0.11172997 |
| RNF8 | protein_coding | -0.0267766 | 0.23823045 |
| ANKRD12 | protein_coding | -0.0268682 | 0.34780191 |
| CHAC2 | protein_coding | -0.0268848 | 0.16623152 |
| SNHG5 | lncRNA | -0.0269642 | 0.08945728 |
| RP5-965G21. | lncRNA | -0.0270532 | 0.02829307 |
| UTP23 | protein_coding | -0.0271084 | 0.28564186 |
| RP11-1396O | lncRNA | -0.0271342 | 0.06162984 |
| C14orf119 | protein_coding | -0.027217 | 0.31451048 |

| | | | |
|---|---|---|---|
| COPS4 | protein_coding | -0.0272207 | 0.35063875 |
| ULK2 | protein_coding | -0.0272258 | 0.22386662 |
| RNF32 | protein_coding | -0.027265 | 0.03435998 |
| C16orf87 | protein_coding | -0.0272783 | 0.26322418 |
| IDH3B | protein_coding | -0.027288 | 0.416503 |
| ACAD9 | protein_coding | -0.0274013 | 0.37744083 |
| GTF2F1 | protein_coding | -0.0274724 | 0.47919842 |
| ORMDL1 | protein_coding | -0.0274725 | 0.30818499 |
| C2orf49 | protein_coding | -0.0275461 | 0.2477242 |
| NFIX | protein_coding | -0.0275774 | 0.62872282 |
| CRK | protein_coding | -0.0276035 | 0.33475858 |
| FAM160A2 | protein_coding | -0.0276178 | 0.3090684 |
| PMM2 | protein_coding | -0.0278659 | 0.29066589 |
| TMEM141 | protein_coding | -0.0279377 | 0.20425124 |
| CEP83-DT | lncRNA | -0.0280576 | 0.0583631 |
| RP11-344B2: | lncRNA | -0.0280657 | 0.10747551 |
| ATP5MF | protein_coding | -0.0281106 | 0.27064101 |
| C19orf73 | protein_coding | -0.0281385 | 0.05780508 |
| SH3GL1 | protein_coding | -0.0282266 | 0.38150057 |
| DYNLT1 | protein_coding | -0.028318 | 0.33361498 |
| ZNF530 | protein_coding | -0.0283874 | 0.17022441 |
| ACD | protein_coding | -0.0283995 | 0.26295791 |
| RBM42 | protein_coding | -0.028431 | 0.22778119 |
| RP11-726G1. | lncRNA | -0.0284926 | 0.04666536 |
| SCRN1 | protein_coding | -0.0285147 | 0.79668787 |
| PHLDB1 | protein_coding | -0.0285549 | 0.48321279 |
| C2orf68 | protein_coding | -0.0285704 | 0.37107989 |
| CHRNA3 | protein_coding | -0.0286237 | 0.58916112 |
| DHPS | protein_coding | -0.0286587 | 0.09354506 |
| ENTPD1-AS1 | lncRNA | -0.028683 | 0.34277422 |
| RPL7L1 | protein_coding | -0.0287395 | 0.73650404 |
| CFAP53 | protein_coding | -0.028807 | 0.05242702 |
| NAA16 | protein_coding | -0.0288902 | 0.23878319 |
| ZNF77 | protein_coding | -0.0289948 | 0.22610274 |
| PDAP1 | protein_coding | -0.0290642 | 0.57926629 |
| DNAL4 | protein_coding | -0.0290647 | 0.20754778 |
| GGNBP2 | protein_coding | -0.0290884 | 0.49196657 |
| TIMM17B | protein_coding | -0.0291607 | 0.34760699 |
| MBLAC1 | protein_coding | -0.0292401 | 0.05245496 |
| SESN1 | protein_coding | -0.0292956 | 0.16852557 |
| FANCC | protein_coding | -0.0293769 | 0.43754557 |
| STXBP4 | protein_coding | -0.029407 | 0.21707533 |
| ASTE1 | protein_coding | -0.0294101 | 0.19936984 |
| RPN2 | protein_coding | -0.0294318 | 0.81071137 |
| RP5-1142A6. | lncRNA | -0.0294497 | 0.03126405 |

| | | | |
|---|---|---|---|
| VMP1 | protein_coding | -0.0295308 | 0.40051218 |
| MUC1 | protein_coding | -0.0297713 | 0.43253107 |
| POLR3A | protein_coding | -0.0297852 | 0.36763592 |
| CLK2 | protein_coding | -0.029807 | 0.42748113 |
| ALG11 | protein_coding | -0.0298374 | 0.13965276 |
| KATNA1 | protein_coding | -0.0298502 | 0.22970059 |
| TRAF3IP1 | protein_coding | -0.029951 | 0.17830846 |
| DNAJC3 | protein_coding | -0.0299588 | 0.31440774 |
| TUFT1 | protein_coding | -0.0299641 | 0.53935027 |
| ITGA7 | protein_coding | -0.0299785 | 0.08329672 |
| POLR2A | protein_coding | -0.030015 | 0.6461803 |
| SORT1 | protein_coding | -0.0300518 | 0.46517361 |
| ZNF593 | protein_coding | -0.0300651 | 0.17246687 |
| COX10-AS1 | lncRNA | -0.0300989 | 0.18453525 |
| ACBD3 | protein_coding | -0.0303642 | 0.31845653 |
| UBE2K | protein_coding | -0.0303909 | 0.48810284 |
| AC084809.2 | lncRNA | -0.0304856 | 0.02824949 |
| S100PBP | protein_coding | -0.0305189 | 0.41018679 |
| RNF38 | protein_coding | -0.030529 | 0.41081467 |
| AKAP8 | protein_coding | -0.0305316 | 0.36447341 |
| LARP7 | protein_coding | -0.030543 | 0.39123359 |
| LSM12 | protein_coding | -0.0305986 | 0.13370535 |
| NUCB2 | protein_coding | -0.0306038 | 0.33669817 |
| POLR1E | protein_coding | -0.0307615 | 0.47071097 |
| ZNF16 | protein_coding | -0.0308604 | 0.18410572 |
| BBS9 | protein_coding | -0.0311635 | 0.20271861 |
| CMB9-55A18 | lncRNA | -0.0311978 | 0.03312687 |
| SPRED3 | protein_coding | -0.031229 | 0.04102848 |
| PRKDC | protein_coding | -0.0313701 | 0.86245015 |
| RP11-1134I1 | lncRNA | -0.0314241 | 0.07460794 |
| TUT4 | protein_coding | -0.0314871 | 0.49298206 |
| NDUFV1 | protein_coding | -0.0316329 | 0.59726536 |
| INSM2 | protein_coding | -0.0316789 | 0.04687763 |
| NONO | protein_coding | -0.0317246 | 1.04170872 |
| WDR5 | protein_coding | -0.0317375 | 0.62663327 |
| SLC25A38 | protein_coding | -0.0317489 | 0.31447866 |
| TLK1 | protein_coding | -0.0318379 | 0.53720664 |
| LRFN5 | protein_coding | -0.0318582 | 0.31822127 |
| TMEM234 | protein_coding | -0.0319247 | 0.0931711 |
| MBNL3 | protein_coding | -0.0320051 | 0.4337206 |
| MUL1 | protein_coding | -0.0320353 | 0.3380892 |
| REEP5 | protein_coding | -0.0320879 | 0.45837186 |
| RTEL1 | protein_coding | -0.0321829 | 0.06207117 |
| FARS2 | protein_coding | -0.0322054 | 0.19964757 |
| COQ4 | protein_coding | -0.032276 | 0.35826028 |

| | | | |
|---|---|---|---|
| HSD17B10 | protein_coding | -0.0322843 | 0.62213927 |
| RP11-461A8. | lncRNA | -0.0322994 | 0.07077356 |
| CMPK1 | protein_coding | -0.032308 | 0.41791588 |
| ALKBH2 | protein_coding | -0.0323374 | 0.32887917 |
| PLEKHB2 | protein_coding | -0.0323463 | 0.61671607 |
| NDUFA11 | protein_coding | -0.0323576 | 0.21194578 |
| KLHDC2 | protein_coding | -0.0323688 | 0.28894841 |
| GDE1 | protein_coding | -0.0323831 | 0.55054025 |
| ZHX3 | protein_coding | -0.0323965 | 0.55644385 |
| GANC | protein_coding | -0.03241 | 0.2363587 |
| GATD3B | protein_coding | -0.0325189 | 0.05084783 |
| AC004156.3 | lncRNA | -0.0326584 | 0.03578743 |
| LINC01001 | lncRNA | -0.0327658 | 0.03743581 |
| LCOR | protein_coding | -0.0327812 | 0.31458618 |
| FLII | protein_coding | -0.0327846 | 0.353929 |
| TNPO2 | protein_coding | -0.0327858 | 0.93294632 |
| BZW1 | protein_coding | -0.0327875 | 0.63878034 |
| TCAP | protein_coding | -0.0328121 | 0.04791336 |
| GSPT1 | protein_coding | -0.0328214 | 0.81060604 |
| ABI2 | protein_coding | -0.0328239 | 0.61871075 |
| ATF7IP | protein_coding | -0.0328384 | 0.67975714 |
| SLC27A4 | protein_coding | -0.0330763 | 0.3468314 |
| ASCC2 | protein_coding | -0.033157 | 0.82188802 |
| RPL37 | protein_coding | -0.03324 | 1.01375576 |
| RNF34 | protein_coding | -0.0332772 | 0.44797512 |
| WWC1 | protein_coding | -0.0333947 | 0.38873862 |
| TAOK2 | protein_coding | -0.033449 | 0.64176798 |
| ZNF281 | protein_coding | -0.0334991 | 0.4375102 |
| SLC39A6 | protein_coding | -0.0335971 | 0.64094724 |
| CLCN3 | protein_coding | -0.0335975 | 0.44212941 |
| ZGPAT | protein_coding | -0.0336283 | 0.11503586 |
| PLEKHO1 | protein_coding | -0.0337042 | 0.37347473 |
| POLRMT | protein_coding | -0.0337637 | 0.2437459 |
| ATP5F1B | protein_coding | -0.0338155 | 1.29854342 |
| ATAD1 | protein_coding | -0.0339119 | 0.37385977 |
| XRN2 | protein_coding | -0.0339151 | 0.79350828 |
| SDHC | protein_coding | -0.0339546 | 0.47182557 |
| PIGC | protein_coding | -0.0340607 | 0.31527321 |
| NSUN5 | protein_coding | -0.0341047 | 0.38652205 |
| IMP4 | protein_coding | -0.0341452 | 0.58117614 |
| PEX19 | protein_coding | -0.0342048 | 0.41034032 |
| PRMT9 | protein_coding | -0.0343488 | 0.26314669 |
| WDR55 | protein_coding | -0.0343505 | 0.35286806 |
| NAA80 | protein_coding | -0.0343643 | 0.11571717 |
| EIF3E | protein_coding | -0.0343833 | 0.65605125 |

| | | | |
|---|---|---|---|
| DDHD1 | protein_coding | -0.0344123 | 0.39108689 |
| VTI1A | protein_coding | -0.0344475 | 0.21705439 |
| ZNF287 | protein_coding | -0.0344476 | 0.11139767 |
| ECI2 | protein_coding | -0.0344771 | 0.51025023 |
| SUCLA2 | protein_coding | -0.0344907 | 0.35890133 |
| TSPYL1 | protein_coding | -0.0344963 | 0.44158778 |
| SUZ12 | protein_coding | -0.0345419 | 0.56822733 |
| RO60 | protein_coding | -0.0345781 | 0.29698791 |
| SENP3 | protein_coding | -0.0346141 | 0.25786728 |
| RP6-109B7.4 | lncRNA | -0.0346212 | 0.0331757 |
| SEC14L2 | protein_coding | -0.0346395 | 0.0495064 |
| WDR12 | protein_coding | -0.0346783 | 0.5778246 |
| ZNF552 | protein_coding | -0.0347493 | 0.19194729 |
| KRR1 | protein_coding | -0.0347777 | 0.6136245 |
| RBBP8 | protein_coding | -0.0348244 | 0.76401071 |
| BAZ1B | protein_coding | -0.034898 | 0.9568415 |
| DCAF1 | protein_coding | -0.0349166 | 0.36595542 |
| LSM2 | protein_coding | -0.0349619 | 0.56822083 |
| C6orf120 | protein_coding | -0.0350644 | 0.2469868 |
| UBE2J2 | protein_coding | -0.0351223 | 0.26927532 |
| NDUFA4 | protein_coding | -0.0351896 | 0.82209051 |
| CDC25B | protein_coding | -0.0352 | 0.94890856 |
| FLOT1 | protein_coding | -0.0352056 | 0.48998555 |
| DYRK3-AS1 | lncRNA | -0.0352802 | 0.02859569 |
| SDK2 | protein_coding | -0.0353518 | 0.24245568 |
| PHYHIPL | protein_coding | -0.0353533 | 0.33432983 |
| CDKL5 | protein_coding | -0.0353602 | 0.31703583 |
| RC3H2 | protein_coding | -0.0354495 | 0.96513763 |
| ITSN1 | protein_coding | -0.0354937 | 0.40462635 |
| GRWD1 | protein_coding | -0.0355812 | 0.51456773 |
| TMEM41B | protein_coding | -0.0356886 | 0.24847897 |
| MTRES1 | protein_coding | -0.0357359 | 0.17166239 |
| RNPEPL1 | protein_coding | -0.0357798 | 0.34058399 |
| TMEM165 | protein_coding | -0.0358393 | 0.34329614 |
| KLHDC4 | protein_coding | -0.035847 | 0.50486106 |
| KDM8 | protein_coding | -0.0359639 | 0.14976893 |
| SRBD1 | protein_coding | -0.0360041 | 0.42417721 |
| RP1-266L20.4 | lncRNA | -0.0360378 | 0.03579375 |
| PFN1 | protein_coding | -0.0361006 | 0.7022284 |
| CNOT11 | protein_coding | -0.0361913 | 0.48896581 |
| RPL30 | protein_coding | -0.0362623 | 1.02528642 |
| OXCT1-AS1 | lncRNA | -0.0363186 | 0.04815965 |
| HAUS5 | protein_coding | -0.0364051 | 0.63240315 |
| PCGF3 | protein_coding | -0.0365795 | 0.63526035 |
| MPPE1 | protein_coding | -0.0366889 | 0.26387704 |

| | | | |
|---|---|---|---|
| PRKCE | protein_coding | -0.03673 | 0.22505462 |
| TMEM130 | protein_coding | -0.0367887 | 0.09820657 |
| POMZP3 | protein_coding | -0.036819 | 0.23745022 |
| FAM214B | protein_coding | -0.0368499 | 0.21169806 |
| MORF4L1 | protein_coding | -0.0368936 | 0.91037369 |
| TMBIM4 | protein_coding | -0.0369475 | 0.12698688 |
| TAF7L | protein_coding | -0.0370021 | 0.03210492 |
| NDUFB4 | protein_coding | -0.0370031 | 0.38074343 |
| ZNF79 | protein_coding | -0.0370746 | 0.20702821 |
| AGTPBP1 | protein_coding | -0.0370918 | 0.61392444 |
| GPAA1 | protein_coding | -0.0371233 | 0.73008253 |
| CCT6A | protein_coding | -0.0371715 | 1.24665064 |
| DZIP1L | protein_coding | -0.037175 | 0.07351306 |
| HAGLR | lncRNA | -0.0373229 | 0.56875667 |
| EPS15 | protein_coding | -0.0373311 | 0.43628817 |
| TP53RK | protein_coding | -0.0374576 | 0.33254386 |
| ZDHHC9 | protein_coding | -0.0376493 | 0.30881644 |
| ZNF317 | protein_coding | -0.0376618 | 0.59213508 |
| RGS2 | protein_coding | -0.0376637 | 0.28099636 |
| ARHGAP17 | protein_coding | -0.0376687 | 0.39282926 |
| RNF215 | protein_coding | -0.0377039 | 0.10126687 |
| CD151 | protein_coding | -0.0377971 | 0.53220091 |
| BUD13 | protein_coding | -0.0378386 | 0.40860946 |
| NIPA2 | protein_coding | -0.0378515 | 0.81236863 |
| LRRC58 | protein_coding | -0.0378759 | 0.40760827 |
| PTTG1IP | protein_coding | -0.0379578 | 0.76208125 |
| SRPK2 | protein_coding | -0.0379815 | 0.5524761 |
| UBE2D2 | protein_coding | -0.0380617 | 0.77384522 |
| AL133243.2 | lncRNA | -0.0381092 | 0.04035311 |
| ZNF232 | protein_coding | -0.0381775 | 0.21873812 |
| ZBTB11-AS1 | lncRNA | -0.0381884 | 0.14662477 |
| DAZAP2 | protein_coding | -0.038191 | 0.83983443 |
| NR1H2 | protein_coding | -0.0382345 | 0.33487765 |
| KCTD18 | protein_coding | -0.0382438 | 0.21662455 |
| NTMT1 | protein_coding | -0.0382693 | 0.46491204 |
| NUDT15 | protein_coding | -0.0382733 | 0.44481563 |
| ST3GAL5 | protein_coding | -0.0383958 | 0.85558007 |
| CSNK1G3 | protein_coding | -0.0384096 | 0.41134844 |
| DNM1 | protein_coding | -0.0384465 | 0.62973948 |
| WRAP73 | protein_coding | -0.0385465 | 0.4509577 |
| RP11-171I2.2 | lncRNA | -0.0385506 | 0.0372625 |
| EMC9 | protein_coding | -0.0385937 | 0.39946962 |
| PPP1R12C | protein_coding | -0.0386723 | 0.38593474 |
| ESD | protein_coding | -0.0386772 | 0.62592092 |
| WBP1L | protein_coding | -0.0387443 | 0.25680289 |

| | | | |
|---|---|---|---|
| MRM1 | protein_coding | -0.0387463 | 0.23216391 |
| ANP32B | protein_coding | -0.0388033 | 0.9173837 |
| ODF2 | protein_coding | -0.0388459 | 0.79372288 |
| RIPOR2 | protein_coding | -0.038901 | 0.60101639 |
| CCNB1IP1 | protein_coding | -0.0389042 | 0.5578045 |
| CDYL2 | protein_coding | -0.0389167 | 0.43768692 |
| SUCLG1 | protein_coding | -0.0389285 | 0.74122133 |
| CARM1 | protein_coding | -0.0389901 | 0.72787883 |
| SSR1 | protein_coding | -0.0390938 | 0.68657948 |
| SEMA4B | protein_coding | -0.0392421 | 0.21867169 |
| IDS | protein_coding | -0.0392588 | 0.11641985 |
| SLC25A25 | protein_coding | -0.0392825 | 0.60508538 |
| DNAJC16 | protein_coding | -0.0393564 | 0.33840456 |
| R3HCC1L | protein_coding | -0.0393574 | 0.25921488 |
| CDC23 | protein_coding | -0.0393876 | 0.49101621 |
| MEPCE | protein_coding | -0.0394254 | 0.86422237 |
| RP11-1275H. | lncRNA | -0.039436 | 0.15486217 |
| RUNDC1 | protein_coding | -0.039437 | 0.40309315 |
| BRCC3 | protein_coding | -0.0395456 | 0.54309706 |
| FOXC1 | protein_coding | -0.0395625 | 0.29417549 |
| RANBP6 | protein_coding | -0.0395761 | 0.6328717 |
| MT-ND5 | protein_coding | -0.0396077 | 0.77680521 |
| EXOC5 | protein_coding | -0.0396117 | 0.60210376 |
| TMEM70 | protein_coding | -0.0396383 | 0.27546464 |
| GAB2 | protein_coding | -0.0396808 | 0.77512745 |
| EXOSC6 | protein_coding | -0.0396839 | 0.54964258 |
| AGAP2-AS1 | lncRNA | -0.0397272 | 0.28412902 |
| ISG20L2 | protein_coding | -0.0397575 | 0.75345788 |
| SYNE3 | protein_coding | -0.039768 | 0.43135434 |
| INA | protein_coding | -0.0398534 | 0.72299987 |
| NARF-IT1 | lncRNA | -0.0399397 | 0.03312255 |
| SAP130 | protein_coding | -0.040004 | 0.65962124 |
| PDCD4-AS1 | lncRNA | -0.0400161 | 0.1200747 |
| ZNF19 | protein_coding | -0.0401009 | 0.08297364 |
| ZNF571 | protein_coding | -0.0401441 | 0.20892382 |
| TLE6 | protein_coding | -0.0401541 | 0.24274991 |
| TICRR | protein_coding | -0.0402152 | 0.37810924 |
| FAM98B | protein_coding | -0.0402598 | 0.58228579 |
| SAYSD1 | protein_coding | -0.0402964 | 0.33795298 |
| ZNF839 | protein_coding | -0.0403393 | 0.34596632 |
| DBNL | protein_coding | -0.0403832 | 0.64155559 |
| IKBIP | protein_coding | -0.0407082 | 0.25351894 |
| RP5-994D16. | lncRNA | -0.0408319 | 0.05500316 |
| SSH2 | protein_coding | -0.0408455 | 0.66866971 |
| TMEM115 | protein_coding | -0.0408612 | 0.26381235 |

| | | | |
|---|---|---|---|
| RPL34 | protein_coding | -0.0409126 | 0.98095356 |
| HLF | protein_coding | -0.0409556 | 0.44661952 |
| CCDC120 | protein_coding | -0.0410279 | 0.26462571 |
| NCAM1 | protein_coding | -0.041087 | 0.71836305 |
| ZNF180 | protein_coding | -0.0412736 | 0.38719974 |
| SNX17 | protein_coding | -0.0413008 | 0.63990332 |
| SWT1 | protein_coding | -0.0413241 | 0.17084971 |
| ENPP4 | protein_coding | -0.0413543 | 0.35430704 |
| HNRNPLL | protein_coding | -0.0413769 | 0.51719862 |
| ZNF782 | protein_coding | -0.0414619 | 0.25235974 |
| NRARP | protein_coding | -0.0414802 | 0.3241015 |
| PHYKPL | protein_coding | -0.041494 | 0.13349414 |
| KNL1 | protein_coding | -0.0415101 | 0.73448582 |
| CRIM1 | protein_coding | -0.0415312 | 0.30875248 |
| SNRNP27 | protein_coding | -0.0416867 | 0.3417597 |
| GOLGA5 | protein_coding | -0.0416961 | 0.6822224 |
| RCC1L | protein_coding | -0.0416984 | 0.44463343 |
| RP11-287D1. | lncRNA | -0.0417071 | 0.04184594 |
| CD2BP2 | protein_coding | -0.0417785 | 0.82488806 |
| GNA12 | protein_coding | -0.0417987 | 0.64459054 |
| C12orf60 | protein_coding | -0.0418614 | 0.10602834 |
| AL163953.3 | lncRNA | -0.0418653 | 0.04945574 |
| LONP2 | protein_coding | -0.0419665 | 0.71656512 |
| ZDBF2 | protein_coding | -0.0420953 | 0.62806798 |
| ANKRD44 | protein_coding | -0.0420954 | 0.16375075 |
| SH3BGRL | protein_coding | -0.042173 | 0.23316408 |
| CTD-2555O1( | lncRNA | -0.0421761 | 0.07350905 |
| RP11-561N1. | lncRNA | -0.0421983 | 0.08915569 |
| TEDC2 | protein_coding | -0.042453 | 0.34310228 |
| LINC00662 | lncRNA | -0.0425286 | 0.27589308 |
| PGM2L1 | protein_coding | -0.0425367 | 0.52991039 |
| ATP2B1 | protein_coding | -0.0426311 | 0.68024822 |
| PARK7 | protein_coding | -0.0426763 | 0.83254707 |
| ZRANB2-AS2 | lncRNA | -0.0426914 | 0.0426067 |
| ZNF688 | protein_coding | -0.042699 | 0.2023829 |
| TRAM1L1 | protein_coding | -0.0427272 | 0.16722973 |
| CD58 | protein_coding | -0.0427494 | 0.09211111 |
| ZNF443 | protein_coding | -0.0428363 | 0.15589739 |
| CDK11B | protein_coding | -0.0430138 | 0.39100947 |
| KAT14 | protein_coding | -0.0430409 | 0.51426797 |
| MRPL18 | protein_coding | -0.0430798 | 0.50606516 |
| VPS26A | protein_coding | -0.0431078 | 0.56095292 |
| CEP104 | protein_coding | -0.0431651 | 0.53577933 |
| PDPK1 | protein_coding | -0.0431882 | 0.73217928 |
| PHKB | protein_coding | -0.0433175 | 0.89416267 |

| | | | |
|---|---|---|---|
| ENY2 | protein_coding | -0.0433542 | 0.8203426 |
| KMT2E-AS1 | lncRNA | -0.0433776 | 0.14868674 |
| KLHL18 | protein_coding | -0.0434608 | 0.40029836 |
| PTDSS2 | protein_coding | -0.0434726 | 0.46149698 |
| CLUAP1 | protein_coding | -0.0435266 | 0.23629429 |
| STARD10 | protein_coding | -0.0435761 | 0.54027186 |
| PEX13 | protein_coding | -0.0436986 | 0.41865988 |
| FOXM1 | protein_coding | -0.0437354 | 1.14641608 |
| FAM120B | protein_coding | -0.0437684 | 0.54434466 |
| RP11-97C16. | lncRNA | -0.0437715 | 0.08744261 |
| ZNF398 | protein_coding | -0.0437796 | 0.59781719 |
| SPTY2D1 | protein_coding | -0.0437815 | 0.7011237 |
| HNRNPH1 | protein_coding | -0.0438286 | 1.30104855 |
| ILF2 | protein_coding | -0.0439187 | 1.36845207 |
| SCRIB | protein_coding | -0.0439944 | 0.48257664 |
| COG8 | protein_coding | -0.0440318 | 0.40109198 |
| SMN1 | protein_coding | -0.0441447 | 0.19768656 |
| RP11-849H4. | protein_coding | -0.0441553 | 0.24609189 |
| MFF | protein_coding | -0.0444267 | 0.7470686 |
| DERA | protein_coding | -0.0445055 | 0.54420041 |
| CTD-2368P2ː | lncRNA | -0.0445586 | 0.34777456 |
| KLHL2 | protein_coding | -0.0447185 | 0.60486165 |
| POLR2F | protein_coding | -0.0447582 | 0.69538018 |
| BPNT2 | protein_coding | -0.044786 | 0.50324354 |
| DEF8 | protein_coding | -0.0448019 | 0.63943567 |
| EIF3A | protein_coding | -0.0448384 | 1.39679596 |
| CNNM2 | protein_coding | -0.0448389 | 0.42879011 |
| MTREX | protein_coding | -0.0448448 | 1.18733869 |
| DPY30 | protein_coding | -0.0448595 | 0.5766693 |
| ZBTB8OS | protein_coding | -0.0449343 | 0.65214658 |
| NDUFC1 | protein_coding | -0.0449946 | 0.41185489 |
| GLRB | protein_coding | -0.0450379 | 0.40444876 |
| KIF17 | protein_coding | -0.0451801 | 0.10519618 |
| COPB2 | protein_coding | -0.045247 | 1.16281679 |
| SSU72 | protein_coding | -0.0452474 | 0.82634371 |
| MAFG-DT | lncRNA | -0.0453119 | 0.19469659 |
| MBD1 | protein_coding | -0.0453275 | 0.97174633 |
| ZNF570 | protein_coding | -0.0453881 | 0.3501737 |
| ACTL10 | protein_coding | -0.0454013 | 0.05416732 |
| WNT7A | protein_coding | -0.045409 | 0.11094771 |
| ANO6 | protein_coding | -0.0454263 | 0.79852594 |
| DHTKD1 | protein_coding | -0.0454683 | 1.06224078 |
| XAB2 | protein_coding | -0.0455371 | 0.73190942 |
| NUP188 | protein_coding | -0.0455404 | 1.39173268 |
| MINDY2 | protein_coding | -0.0455548 | 0.48210605 |

| | | | |
|---|---|---|---|
| PI4K2A | protein_coding | -0.0455682 | 0.28415001 |
| TDG | protein_coding | -0.0456009 | 0.90327567 |
| CEBPZOS | protein_coding | -0.0456915 | 0.42937207 |
| TMEM208 | protein_coding | -0.0457727 | 0.64125446 |
| ZNF721 | protein_coding | -0.0458317 | 0.63151113 |
| RP5-1184F4. | lncRNA | -0.045882 | 0.04201242 |
| RP11-973D8. | lncRNA | -0.0459691 | 0.04083681 |
| SGCG | protein_coding | -0.045976 | 0.6325864 |
| TMEM147 | protein_coding | -0.0460449 | 0.66747074 |
| DESI2 | protein_coding | -0.0461045 | 0.68132185 |
| KPNA3 | protein_coding | -0.046203 | 0.52972262 |
| SF3B2 | protein_coding | -0.0462995 | 1.557299 |
| USB1 | protein_coding | -0.0464177 | 0.66563127 |
| TSNAX | protein_coding | -0.0464387 | 0.40692391 |
| CRELD1 | protein_coding | -0.0464966 | 0.10232623 |
| HDAC3 | protein_coding | -0.0464975 | 0.7963972 |
| DNAAF5 | protein_coding | -0.0465151 | 0.55749992 |
| ZNF66 | protein_coding | -0.046526 | 0.16618399 |
| TRIP11 | protein_coding | -0.046549 | 0.75563267 |
| AKT3 | protein_coding | -0.0467269 | 1.06321968 |
| AQP4 | protein_coding | -0.0468078 | 0.20843328 |
| CTD-2600O9. | lncRNA | -0.0468254 | 0.05371937 |
| MFSD2B | protein_coding | -0.0468543 | 0.04414452 |
| BYSL | protein_coding | -0.0469236 | 0.58040444 |
| RP11-131L23 | lncRNA | -0.0469648 | 0.05229188 |
| GABARAPL2 | protein_coding | -0.0470018 | 0.69807753 |
| TEX10 | protein_coding | -0.0470213 | 0.46266741 |
| ADO | protein_coding | -0.0470316 | 0.40151506 |
| ERF | protein_coding | -0.0470538 | 0.76928427 |
| BRD7 | protein_coding | -0.0470938 | 1.12410265 |
| CEP20 | protein_coding | -0.0470946 | 0.56129038 |
| STAMBP | protein_coding | -0.0471187 | 0.87216868 |
| RBBP4 | protein_coding | -0.0471847 | 1.87000566 |
| GPBP1L1 | protein_coding | -0.0472149 | 0.86567392 |
| SCAF1 | protein_coding | -0.0472748 | 0.31125437 |
| ASRGL1 | protein_coding | -0.0473029 | 0.78010389 |
| RP11-95J9.3 | lncRNA | -0.0473531 | 0.04499055 |
| RBBP5 | protein_coding | -0.0474289 | 0.79383527 |
| CAMSAP3 | protein_coding | -0.0474366 | 0.38185627 |
| PITPNM3 | protein_coding | -0.0474779 | 0.06205281 |
| CPE | protein_coding | -0.0475189 | 1.62603162 |
| M6PR | protein_coding | -0.0475533 | 0.86933149 |
| WBP11 | protein_coding | -0.0475568 | 1.22530369 |
| NMT2 | protein_coding | -0.0475691 | 0.22372896 |
| IPMK | protein_coding | -0.0475899 | 0.20122625 |

| | | | |
|---|---|---|---|
| RP5-965G21. | lncRNA | -0.0476978 | 0.07275434 |
| RPS15A | protein_coding | -0.0477539 | 0.67287789 |
| MED17 | protein_coding | -0.0478474 | 0.24127982 |
| XPO1 | protein_coding | -0.0478711 | 1.18898513 |
| CMC2 | protein_coding | -0.047888 | 0.33956924 |
| AC005786.7 | lncRNA | -0.0479282 | 0.09086748 |
| MFAP3 | protein_coding | -0.0479722 | 0.75174866 |
| PLCXD1 | protein_coding | -0.0479817 | 0.40546274 |
| AP1AR | protein_coding | -0.0480084 | 0.67027786 |
| TMEM81 | protein_coding | -0.0480765 | 0.15855012 |
| MRPS7 | protein_coding | -0.0481287 | 0.69868019 |
| TMEM198 | protein_coding | -0.0481576 | 0.52230751 |
| PTGES3 | protein_coding | -0.0482318 | 1.24175388 |
| POMT1 | protein_coding | -0.0482628 | 0.5702249 |
| DRG1 | protein_coding | -0.0483246 | 0.78289464 |
| ADAMTS5 | protein_coding | -0.0483607 | 0.17166178 |
| URGCP | protein_coding | -0.0484819 | 0.57062074 |
| SUGT1 | protein_coding | -0.0485106 | 0.50072102 |
| LINC01550 | lncRNA | -0.0485469 | 0.28552973 |
| SUFU | protein_coding | -0.048558 | 0.47048752 |
| FOXK2 | protein_coding | -0.0486053 | 1.1886774 |
| MCF2L-AS1 | lncRNA | -0.048607 | 0.25958111 |
| RP3-402G11. | lncRNA | -0.048644 | 0.06527097 |
| TMEM102 | protein_coding | -0.048676 | 0.11263967 |
| ZNF674-AS1 | lncRNA | -0.0487066 | 0.2281342 |
| RNFT1 | protein_coding | -0.0487451 | 0.15344062 |
| HIBCH | protein_coding | -0.0489259 | 0.477595 |
| SNRNP40 | protein_coding | -0.0489347 | 1.08546495 |
| TMUB1 | protein_coding | -0.0489713 | 0.4610745 |
| COG4 | protein_coding | -0.049032 | 0.90211689 |
| PPP2R5A | protein_coding | -0.0490584 | 0.67349338 |
| C3orf38 | protein_coding | -0.0491976 | 0.33465119 |
| AK9 | protein_coding | -0.0492328 | 0.27850795 |
| MUC20-OT1 | lncRNA | -0.0493269 | 0.69054671 |
| PSPH | protein_coding | -0.0493323 | 0.61415319 |
| TAF5L | protein_coding | -0.049375 | 0.67271674 |
| ZFYVE1 | protein_coding | -0.0494136 | 0.54756734 |
| PET100 | protein_coding | -0.0494367 | 0.12058545 |
| NCOA2 | protein_coding | -0.0494397 | 0.54363684 |
| FAM151B | protein_coding | -0.0494574 | 0.05445892 |
| TFAP2A-AS1 | lncRNA | -0.0495024 | 0.16690102 |
| ZNF407-AS1 | lncRNA | -0.0495139 | 0.31997633 |
| STK16 | protein_coding | -0.0495624 | 0.49723771 |
| NUP214 | protein_coding | -0.0496728 | 1.09984527 |
| RPS19BP1 | protein_coding | -0.0496835 | 0.43712831 |

| | | | |
|---|---|---|---|
| AP000350.5 | lncRNA | -0.0497165 | 0.16094689 |
| RORB | protein_coding | -0.0497713 | 1.31997252 |
| AP2M1 | protein_coding | -0.0497831 | 1.92923799 |
| CENPP | protein_coding | -0.0497911 | 0.70650441 |
| GDF9 | protein_coding | -0.0498611 | 0.11304545 |
| COPS6 | protein_coding | -0.0498759 | 1.22548902 |
| CLDND2 | protein_coding | -0.0499829 | 0.05812738 |
| TMEM120A | protein_coding | -0.0501095 | 0.21108399 |
| STX12 | protein_coding | -0.0501344 | 0.70898139 |
| ZNF200 | protein_coding | -0.0502109 | 0.46726504 |
| MKRN2 | protein_coding | -0.0502134 | 0.5918907 |
| ZNF639 | protein_coding | -0.0502887 | 0.79136036 |
| PTRH1 | protein_coding | -0.0503005 | 0.16635779 |
| TBC1D14 | protein_coding | -0.050368 | 1.23795621 |
| ZNF566-AS1 | lncRNA | -0.0503913 | 0.15892906 |
| ERCC2 | protein_coding | -0.0503983 | 0.75039357 |
| RPS10 | protein_coding | -0.0504372 | 0.13516055 |
| AMZ2 | protein_coding | -0.0504766 | 0.88980931 |
| RPS26 | protein_coding | -0.0506843 | 0.98900283 |
| TTLL4 | protein_coding | -0.0506953 | 0.6268837 |
| SMIM20 | protein_coding | -0.0506972 | 0.45483441 |
| CNOT9 | protein_coding | -0.0507311 | 1.26053191 |
| RP11-809H1( | lncRNA | -0.0507486 | 0.08688448 |
| ZNF252P-AS | lncRNA | -0.0507526 | 0.04779124 |
| ABCA7 | protein_coding | -0.0507691 | 0.7523184 |
| CPLANE2 | protein_coding | -0.05078 | 0.25930184 |
| RP11-485G7. | lncRNA | -0.0508395 | 0.06980281 |
| UFD1 | protein_coding | -0.050892 | 0.77344827 |
| GBF1 | protein_coding | -0.0509156 | 1.03611264 |
| EFCAB5 | protein_coding | -0.0509339 | 0.12580727 |
| TTC39C | protein_coding | -0.0510307 | 0.39068347 |
| ARHGAP32 | protein_coding | -0.0511606 | 0.95676967 |
| RNF111 | protein_coding | -0.0511695 | 0.66741741 |
| PELP1 | protein_coding | -0.0512002 | 0.77106579 |
| MSC-AS1 | lncRNA | -0.0512314 | 0.48841672 |
| STRADA | protein_coding | -0.0513684 | 0.17646196 |
| MRPL22 | protein_coding | -0.0514203 | 0.54324297 |
| ZC3H7A | protein_coding | -0.0516306 | 0.83607782 |
| LSM14A | protein_coding | -0.0517136 | 1.4400928 |
| ZMYM1 | protein_coding | -0.0517257 | 0.37504217 |
| CNPY4 | protein_coding | -0.0518079 | 0.32869624 |
| ENO3 | protein_coding | -0.0518541 | 0.4284451 |
| ZSCAN22 | protein_coding | -0.0519084 | 0.24360897 |
| SMARCA5 | protein_coding | -0.0519774 | 1.59399219 |
| ZC3H3 | protein_coding | -0.0519903 | 0.33805722 |

| | | | |
|---|---|---|---|
| COPS3 | protein_coding | -0.0520121 | 0.8740013 |
| NKRF | protein_coding | -0.0520734 | 0.59642803 |
| THUMPD3 | protein_coding | -0.0520852 | 0.59876986 |
| TRAP1 | protein_coding | -0.0521607 | 1.30173593 |
| MIX23 | protein_coding | -0.0522027 | 0.25222694 |
| LIMD2 | protein_coding | -0.0522071 | 0.87204192 |
| OSBPL9 | protein_coding | -0.052209 | 1.01009157 |
| COX7A2 | protein_coding | -0.0523335 | 1.20742428 |
| PPP5C | protein_coding | -0.052349 | 1.42761828 |
| BTBD1 | protein_coding | -0.0524398 | 1.02445842 |
| FIBP | protein_coding | -0.0524624 | 0.89311814 |
| CLASRP | protein_coding | -0.0524982 | 0.64282988 |
| TIMM23 | protein_coding | -0.0526143 | 0.79740181 |
| RAC1 | protein_coding | -0.0526452 | 1.70343956 |
| ATXN10 | protein_coding | -0.0527061 | 1.32928314 |
| FASTKD1 | protein_coding | -0.0527364 | 0.76394423 |
| CEP55 | protein_coding | -0.0528306 | 0.69441196 |
| CHST10 | protein_coding | -0.0528629 | 0.51434419 |
| PRDX5 | protein_coding | -0.0530433 | 1.54439606 |
| TSR2 | protein_coding | -0.0530445 | 0.87483245 |
| ZNF219 | protein_coding | -0.0532238 | 0.27714909 |
| RPS6KB1 | protein_coding | -0.0532663 | 0.79843578 |
| SBDS | protein_coding | -0.0532788 | 0.86629448 |
| RP11-758H9. | lncRNA | -0.0533199 | 0.13902956 |
| GLI2 | protein_coding | -0.0533442 | 0.10870215 |
| RSRP1 | protein_coding | -0.0533615 | 0.56297482 |
| CLPTM1 | protein_coding | -0.0534454 | 0.80495778 |
| LIN7C | protein_coding | -0.0535818 | 0.52500264 |
| SEC13 | protein_coding | -0.0536524 | 0.9812253 |
| TMEM68 | protein_coding | -0.0536857 | 0.37116036 |
| HECA | protein_coding | -0.0537737 | 0.4363227 |
| LPCAT3 | protein_coding | -0.0537763 | 0.80048897 |
| PAPOLA | protein_coding | -0.0538074 | 1.47278323 |
| RP11-2E11.5 | lncRNA | -0.0540503 | 0.06024272 |
| CPSF3 | protein_coding | -0.0540886 | 1.17290124 |
| TPRKB | protein_coding | -0.0540889 | 0.54481479 |
| CTU1 | protein_coding | -0.0540957 | 0.17082631 |
| GPATCH3 | protein_coding | -0.0540962 | 0.45941848 |
| ETFRF1 | protein_coding | -0.0541265 | 0.24105799 |
| TAZ | protein_coding | -0.0541307 | 0.65015987 |
| YWHAE | protein_coding | -0.0541819 | 1.68385051 |
| MEN1 | protein_coding | -0.0542688 | 1.015821 |
| FXR2 | protein_coding | -0.0543557 | 0.8427108 |
| ABCG5 | protein_coding | -0.0543591 | 0.38415253 |
| POLG | protein_coding | -0.0544164 | 0.89641369 |

| | | | |
|---|---|---|---|
| RP13-753N3. | lncRNA | -0.0544588 | 0.10922983 |
| COPS8 | protein_coding | -0.0545094 | 0.96826067 |
| POLE4 | protein_coding | -0.054538 | 0.56017865 |
| BBS10 | protein_coding | -0.0545791 | 0.4247366 |
| THAP12 | protein_coding | -0.0545892 | 1.09098019 |
| ALPP | protein_coding | -0.054607 | 0.05865739 |
| ATG4A | protein_coding | -0.054664 | 0.58916517 |
| FNBP1L | protein_coding | -0.0547713 | 0.75503333 |
| C12orf76 | protein_coding | -0.0548036 | 0.35081489 |
| RP11-96D1.1 | lncRNA | -0.0548516 | 0.20595326 |
| GTF2E1 | protein_coding | -0.0548744 | 0.36065706 |
| ARCN1 | protein_coding | -0.0548941 | 1.58676336 |
| ZNF148 | protein_coding | -0.0549393 | 0.50071771 |
| EOLA2 | protein_coding | -0.05499 | 0.48367849 |
| GINS4 | protein_coding | -0.0551201 | 0.29441816 |
| TEDC1 | protein_coding | -0.0551272 | 0.40093892 |
| SNAI3-AS1 | lncRNA | -0.0552228 | 0.10993902 |
| GNPTG | protein_coding | -0.0552294 | 0.55176416 |
| RPL29 | protein_coding | -0.0552341 | 1.31896264 |
| PCIF1 | protein_coding | -0.0552632 | 0.82439256 |
| SYT3 | protein_coding | -0.0553575 | 0.15628642 |
| EIF3G | protein_coding | -0.055514 | 1.08082853 |
| IRAK4 | protein_coding | -0.0555141 | 0.6996852 |
| SART3 | protein_coding | -0.0555249 | 1.57470608 |
| SNHG9 | lncRNA | -0.0555405 | 0.15355141 |
| ARMC6 | protein_coding | -0.0557773 | 0.95926874 |
| PDHA1 | protein_coding | -0.0558041 | 1.54135025 |
| ARSK | protein_coding | -0.0558646 | 0.30747667 |
| FGFR1OP2 | protein_coding | -0.05587 | 0.55431336 |
| MYBL2 | protein_coding | -0.0558752 | 1.62006987 |
| HSPH1 | protein_coding | -0.0558842 | 1.85182083 |
| DEPTOR | protein_coding | -0.0559924 | 0.14793978 |
| CYB5R4 | protein_coding | -0.0560748 | 0.73135899 |
| USP12 | protein_coding | -0.0561399 | 0.99165437 |
| ADAL | protein_coding | -0.0561865 | 0.70773997 |
| PPP4C | protein_coding | -0.0562175 | 1.18002207 |
| SFI1 | protein_coding | -0.0562292 | 0.72965782 |
| LMBRD1 | protein_coding | -0.0562645 | 0.55167547 |
| CCNT1 | protein_coding | -0.0562725 | 1.16147301 |
| MIR762HG | lncRNA | -0.0563507 | 0.2593531 |
| ARL6IP6 | protein_coding | -0.056372 | 0.58968526 |
| MEAF6 | protein_coding | -0.0563751 | 1.19714513 |
| NDUFA3 | protein_coding | -0.056436 | 0.76440802 |
| CEP97 | protein_coding | -0.0564371 | 0.7770237 |
| SEC61B | protein_coding | -0.0564489 | 0.79119155 |

| | | | |
|---|---|---|---|
| USO1 | protein_coding | -0.0564574 | 1.05088822 |
| KLHL9 | protein_coding | -0.0564758 | 0.72237276 |
| ZCRB1 | protein_coding | -0.0565765 | 0.69120898 |
| STK17B | protein_coding | -0.056626 | 1.46332807 |
| ACAD8 | protein_coding | -0.0566413 | 0.45153165 |
| CSGALNACT2 | protein_coding | -0.0566429 | 0.57794266 |
| NCBP1 | protein_coding | -0.0566442 | 1.81469432 |
| FTSJ1 | protein_coding | -0.0566765 | 0.71412525 |
| SLC20A1 | protein_coding | -0.0568354 | 1.75084793 |
| SARAF | protein_coding | -0.0568377 | 1.18422701 |
| CARS2 | protein_coding | -0.0568675 | 0.94714645 |
| ZNF133 | protein_coding | -0.0569545 | 0.42101987 |
| TAX1BP1 | protein_coding | -0.0570041 | 1.26288348 |
| MED6 | protein_coding | -0.0570565 | 0.59443011 |
| ZNF212 | protein_coding | -0.0570899 | 0.53289016 |
| LETM1 | protein_coding | -0.0571597 | 1.12043063 |
| WBP2NL | protein_coding | -0.0572248 | 0.12464446 |
| HLCS | protein_coding | -0.057339 | 0.8390242 |
| MAVS | protein_coding | -0.0573688 | 1.2462552 |
| EEF1AKNMT | protein_coding | -0.0574069 | 1.04700781 |
| LINC02175 | lncRNA | -0.0574468 | 0.1353716 |
| NIPSNAP1 | protein_coding | -0.0574966 | 1.23103517 |
| WDR83 | protein_coding | -0.0575521 | 0.39554903 |
| MTDH | protein_coding | -0.0576158 | 1.50124574 |
| TMEM245 | protein_coding | -0.057637 | 1.61125098 |
| CTC-338M12 | lncRNA | -0.0576594 | 0.2043334 |
| RP11-359N1 | lncRNA | -0.0576686 | 0.14363747 |
| PUM1 | protein_coding | -0.0576725 | 1.28010168 |
| IFT52 | protein_coding | -0.0577075 | 0.67152156 |
| TALDO1 | protein_coding | -0.0577465 | 1.232031 |
| NAALADL1 | protein_coding | -0.0577534 | 0.4427905 |
| SCRN3 | protein_coding | -0.0577749 | 0.58334128 |
| UBE2G2 | protein_coding | -0.0577801 | 1.90406823 |
| TAB1 | protein_coding | -0.0577953 | 1.0056202 |
| RP11-399K21 | lncRNA | -0.0578876 | 0.11584306 |
| ZNF141 | protein_coding | -0.0578973 | 0.75195904 |
| DISC1 | protein_coding | -0.0579177 | 0.56154268 |
| RFESD | protein_coding | -0.0579286 | 0.19183053 |
| CTD-2192J16 | protein_coding | -0.0579681 | 0.23669119 |
| ZNF646 | protein_coding | -0.0580442 | 1.00022505 |
| NPHP4 | protein_coding | -0.0580511 | 0.46843192 |
| GCC2-AS1 | lncRNA | -0.0580727 | 0.08539055 |
| HEXIM1 | protein_coding | -0.0582233 | 0.71440482 |
| CMTM8 | protein_coding | -0.0582378 | 0.09216886 |
| C4orf46 | protein_coding | -0.058368 | 0.70324246 |

| | | | |
|---|---|---|---|
| SMIM27 | protein_coding | -0.0583937 | 0.19196178 |
| EIF2AK1 | protein_coding | -0.0584243 | 2.08806584 |
| C1GALT1C1 | protein_coding | -0.0585267 | 0.34943154 |
| THAP1 | protein_coding | -0.0585372 | 0.28746867 |
| STRN4 | protein_coding | -0.0585685 | 1.63217796 |
| SMDT1 | protein_coding | -0.0585953 | 0.86001959 |
| SLF2 | protein_coding | -0.058747 | 0.65475608 |
| STYX | protein_coding | -0.0588074 | 0.42029686 |
| HTRA2 | protein_coding | -0.0588222 | 0.77484047 |
| EEF1AKMT4 | protein_coding | -0.0588906 | 0.49337796 |
| PRPF8 | protein_coding | -0.0592035 | 1.87799524 |
| DMAP1 | protein_coding | -0.0592207 | 0.8155217 |
| MICU3 | protein_coding | -0.0592922 | 0.17557982 |
| PRDX1 | protein_coding | -0.0593524 | 1.54852264 |
| POLR1D | protein_coding | -0.059369 | 1.69706442 |
| BAALC-AS1 | lncRNA | -0.0593741 | 0.4867353 |
| DIP2B | protein_coding | -0.0593786 | 1.21789126 |
| POGLUT3 | protein_coding | -0.0593863 | 0.65380918 |
| ZNF568 | protein_coding | -0.05939 | 0.45371342 |
| GLI4 | protein_coding | -0.0594973 | 0.33183389 |
| ZNF284 | protein_coding | -0.0595219 | 0.12416806 |
| TOR1AIP2 | protein_coding | -0.059603 | 0.79493123 |
| CDC123 | protein_coding | -0.0596166 | 0.9394019 |
| GDPD1 | protein_coding | -0.0596336 | 0.51038766 |
| ZNF34 | protein_coding | -0.0596585 | 0.35879809 |
| ATXN7L3 | protein_coding | -0.0598873 | 1.27542091 |
| ADCY9 | protein_coding | -0.0599576 | 1.00773292 |
| RP11-16N11. | lncRNA | -0.0599696 | 0.08872618 |
| TMEM186 | protein_coding | -0.0599959 | 0.56338159 |
| ZNF384 | protein_coding | -0.0600373 | 1.27697693 |
| TOR1AIP1 | protein_coding | -0.0600782 | 0.95633259 |
| EDF1 | protein_coding | -0.060305 | 1.42772789 |
| RNF144B | protein_coding | -0.0604153 | 0.10853502 |
| STAU1 | protein_coding | -0.0604281 | 1.78669349 |
| FOXN4 | protein_coding | -0.0606728 | 0.63005179 |
| CCT3 | protein_coding | -0.0606808 | 2.37752644 |
| CHPT1 | protein_coding | -0.0606868 | 0.82470994 |
| AIFM1 | protein_coding | -0.0607672 | 0.82638257 |
| LINC00235 | lncRNA | -0.060778 | 0.14022263 |
| TFPT | protein_coding | -0.0609101 | 0.44306646 |
| DCTN2 | protein_coding | -0.0609216 | 1.7080861 |
| IDI1 | protein_coding | -0.0609242 | 0.63750381 |
| RAD51AP1 | protein_coding | -0.0609691 | 0.55043876 |
| SLC35E1 | protein_coding | -0.0609799 | 1.08565275 |
| CLTA | protein_coding | -0.0610875 | 0.74623464 |

| | | | |
|---|---|---|---|
| DMTF1 | protein_coding | -0.0611065 | 1.08838515 |
| CKMT2-AS1 | lncRNA | -0.0611329 | 0.52557949 |
| HOXA3 | protein_coding | -0.0612476 | 0.38782444 |
| CDCA5 | protein_coding | -0.061325 | 1.76065187 |
| SNX27 | protein_coding | -0.0613365 | 0.61115332 |
| AC016747.3 | lncRNA | -0.0614099 | 0.40297921 |
| SRP14 | protein_coding | -0.0614148 | 1.84965209 |
| FBXL15 | protein_coding | -0.0614404 | 0.3081609 |
| SLC35B4 | protein_coding | -0.0615966 | 0.65176788 |
| AP3M1 | protein_coding | -0.0616669 | 1.1543217 |
| PDPR | protein_coding | -0.0617244 | 1.39135254 |
| SNF8 | protein_coding | -0.0617591 | 1.05625594 |
| CCDC88A | protein_coding | -0.0618075 | 1.46908844 |
| EIF4ENIF1 | protein_coding | -0.0618615 | 1.05107597 |
| PCCB | protein_coding | -0.0618662 | 0.99619924 |
| C12orf45 | protein_coding | -0.0619711 | 0.68486605 |
| RP11-552M1 | lncRNA | -0.0619887 | 0.12035854 |
| RPS27 | protein_coding | -0.062045 | 2.13869936 |
| ATP5PF | protein_coding | -0.0620585 | 1.32409239 |
| YIPF5 | protein_coding | -0.0621351 | 0.7296694 |
| NDUFB1 | protein_coding | -0.0621838 | 0.7295427 |
| SPATA20 | protein_coding | -0.0622228 | 0.2235968 |
| YIPF6 | protein_coding | -0.0622251 | 1.2047571 |
| MIER3 | protein_coding | -0.0622907 | 0.78791716 |
| CSTF3-DT | lncRNA | -0.0623379 | 0.06694672 |
| ABRAXAS1 | protein_coding | -0.0624047 | 0.56814251 |
| CTD-3220F14 | lncRNA | -0.0624691 | 0.32215965 |
| SPTSSA | protein_coding | -0.0625475 | 0.57133115 |
| MRPS27 | protein_coding | -0.062645 | 1.56187278 |
| NDUFS6 | protein_coding | -0.0626488 | 0.89212542 |
| EPB41L1 | protein_coding | -0.0627051 | 0.4500834 |
| SFT2D2 | protein_coding | -0.0627097 | 1.04034439 |
| ZNF33A | protein_coding | -0.0627329 | 1.07791944 |
| GPRASP2 | protein_coding | -0.0627725 | 0.34739151 |
| TSN | protein_coding | -0.0629415 | 1.6282732 |
| DOC2A | protein_coding | -0.0629909 | 0.53742612 |
| SRP68 | protein_coding | -0.0630167 | 1.38304763 |
| TOE1 | protein_coding | -0.0630935 | 0.74992283 |
| AC002310.12 | lncRNA | -0.0631809 | 0.32737293 |
| YTHDC1 | protein_coding | -0.0631847 | 1.63546259 |
| TRMO | protein_coding | -0.0632418 | 0.33683092 |
| HGS | protein_coding | -0.0632492 | 1.29832946 |
| ARIH2OS | lncRNA | -0.0632503 | 0.10089361 |
| SUMO2 | protein_coding | -0.0632997 | 1.74536983 |
| NOA1 | protein_coding | -0.0634599 | 0.85978976 |

| | | | |
|---|---|---|---|
| NMRAL1 | protein_coding | -0.063499 | 1.10047223 |
| CERS4 | protein_coding | -0.0635182 | 1.00532726 |
| TRAM1 | protein_coding | -0.0635576 | 0.69457396 |
| C1orf174 | protein_coding | -0.0636124 | 0.50652733 |
| TIGAR | protein_coding | -0.0636513 | 0.20053854 |
| TMEM106B | protein_coding | -0.0636523 | 0.74019618 |
| CITF22-92A6 | lncRNA | -0.0636541 | 0.17716973 |
| PAQR3 | protein_coding | -0.0637104 | 0.71792867 |
| RP11-186B7. | lncRNA | -0.0637377 | 0.07848191 |
| APBB2 | protein_coding | -0.0637718 | 0.95214341 |
| DDX25 | protein_coding | -0.0637993 | 0.80755817 |
| FCHSD1 | protein_coding | -0.0638509 | 0.10251179 |
| FBXL17 | protein_coding | -0.0639005 | 0.65411676 |
| SERAC1 | protein_coding | -0.0639051 | 0.29771399 |
| PSMG1 | protein_coding | -0.0639368 | 1.31973841 |
| FAM3A | protein_coding | -0.063943 | 0.85322059 |
| CBY1 | protein_coding | -0.0639575 | 0.57014781 |
| NUMB | protein_coding | -0.0639845 | 1.23534022 |
| GOLGA2 | protein_coding | -0.0640168 | 1.29220499 |
| APP-DT | lncRNA | -0.064042 | 0.08699163 |
| IK | protein_coding | -0.0640558 | 1.3981625 |
| AGPS | protein_coding | -0.064081 | 1.32050108 |
| STUB1 | protein_coding | -0.0640925 | 1.19106402 |
| ITPRID2 | protein_coding | -0.0641333 | 0.66882799 |
| EBNA1BP2 | protein_coding | -0.0642443 | 1.0645847 |
| DDHD2 | protein_coding | -0.0642607 | 1.22059901 |
| MICAL1 | protein_coding | -0.0643295 | 0.61042486 |
| CENPA | protein_coding | -0.0645728 | 0.72199041 |
| SLC35E4 | protein_coding | -0.0645758 | 0.25324216 |
| GMEB2 | protein_coding | -0.0646529 | 0.80090258 |
| SP3 | protein_coding | -0.0647871 | 1.60535099 |
| FOXJ2 | protein_coding | -0.0648131 | 0.96298283 |
| BORCS7 | protein_coding | -0.0648505 | 0.28165633 |
| AC074117.1C | lncRNA | -0.0649825 | 0.32368771 |
| FRS3 | protein_coding | -0.0650072 | 0.49052562 |
| LAMTOR1 | protein_coding | -0.06507 | 1.05084642 |
| BORCS8 | protein_coding | -0.0651581 | 0.36983797 |
| SAR1A | protein_coding | -0.0651868 | 1.12841618 |
| SLC35G2 | protein_coding | -0.0652437 | 0.71403718 |
| NDUFAB1 | protein_coding | -0.0652999 | 0.97057226 |
| RTRAF | protein_coding | -0.065353 | 1.70055291 |
| AGFG1 | protein_coding | -0.0653574 | 0.73211856 |
| NFIB | protein_coding | -0.0654003 | 1.272908 |
| PTPRJ | protein_coding | -0.0654402 | 0.58304608 |
| NOP56 | protein_coding | -0.0655079 | 2.41415096 |

| | | | |
|---|---|---|---|
| CKS1B | protein_coding | -0.065588 | 1.06400344 |
| GPM6A | protein_coding | -0.065593 | 0.64963019 |
| KLF11 | protein_coding | -0.065617 | 0.55583192 |
| TONSL | protein_coding | -0.0656414 | 0.63919478 |
| RP11-758P17 | lncRNA | -0.0656619 | 0.08146651 |
| JRK | protein_coding | -0.065831 | 1.26399257 |
| ROCK1 | protein_coding | -0.0658794 | 1.61516882 |
| CTD-2089N3. | lncRNA | -0.0659212 | 0.30547101 |
| JTB | protein_coding | -0.065938 | 0.47372072 |
| SRSF4 | protein_coding | -0.0659667 | 1.68709542 |
| GMPPB | protein_coding | -0.0660058 | 0.69093457 |
| ETFDH | protein_coding | -0.0660204 | 0.8161305 |
| NUP58 | protein_coding | -0.0660573 | 1.8287915 |
| VRK3 | protein_coding | -0.066133 | 0.75159441 |
| C1orf159 | protein_coding | -0.0661514 | 0.39671921 |
| FAM120A | protein_coding | -0.0661633 | 2.08218574 |
| FBXO38 | protein_coding | -0.0661695 | 1.03698694 |
| OGG1 | protein_coding | -0.0662297 | 0.51259851 |
| RRP1B | protein_coding | -0.066273 | 1.62947922 |
| MALT1 | protein_coding | -0.0664043 | 0.89195745 |
| PMF1 | protein_coding | -0.066559 | 0.40555726 |
| HDGF | protein_coding | -0.0666498 | 2.75291674 |
| STX5 | protein_coding | -0.0666811 | 0.85251104 |
| ZNF512 | protein_coding | -0.0667005 | 0.88490784 |
| SLITRK1 | protein_coding | -0.0667205 | 0.65526893 |
| TMEM30A-D | lncRNA | -0.0667682 | 0.13719834 |
| EME1 | protein_coding | -0.066774 | 0.42540137 |
| FBXW9 | protein_coding | -0.0667913 | 0.31125595 |
| RP11-325I22 | lncRNA | -0.0668215 | 0.54322431 |
| PFAS | protein_coding | -0.0668559 | 1.01981735 |
| NAT9 | protein_coding | -0.0669806 | 0.75176372 |
| SF1-DT | lncRNA | -0.0670119 | 0.07785222 |
| CCDC134 | protein_coding | -0.0670398 | 0.92258083 |
| NAPRT | protein_coding | -0.0671211 | 0.35562838 |
| CEP78 | protein_coding | -0.0671346 | 1.29533965 |
| LA16c-431H6 | protein_coding | -0.067153 | 0.05803469 |
| EDC4 | protein_coding | -0.0672017 | 1.20256507 |
| EIF4A2 | protein_coding | -0.0672759 | 2.56304117 |
| MAST2 | protein_coding | -0.067328 | 1.40518783 |
| CRACD | protein_coding | -0.0673562 | 0.1925822 |
| PRXL2B | protein_coding | -0.0673818 | 1.21708266 |
| RBM3 | protein_coding | -0.067433 | 2.28477955 |
| NRP2 | protein_coding | -0.06747 | 0.77426373 |
| KBTBD3 | protein_coding | -0.0674717 | 0.19966957 |
| ACRBP | protein_coding | -0.0675816 | 0.08258759 |

| | | | |
|---|---|---|---|
| MANEAL | protein_coding | -0.0676215 | 1.63214943 |
| CBX3 | protein_coding | -0.0676645 | 1.34007325 |
| AMOTL1 | protein_coding | -0.067698 | 2.11082506 |
| RANGAP1 | protein_coding | -0.0678063 | 1.62455358 |
| CNDP2 | protein_coding | -0.0678636 | 1.24239866 |
| TUFM | protein_coding | -0.0678782 | 1.79605523 |
| RP11-571M6 | lncRNA | -0.0678854 | 0.079966 |
| AXIN1 | protein_coding | -0.0679011 | 0.81679808 |
| MGME1 | protein_coding | -0.0679159 | 1.05931525 |
| PTP4A2 | protein_coding | -0.0679886 | 2.02755311 |
| PEG13 | lncRNA | -0.0680097 | 0.10754568 |
| PANK1 | protein_coding | -0.0680836 | 0.55325142 |
| RP11-109E12 | lncRNA | -0.0682109 | 0.07321544 |
| POC1A | protein_coding | -0.0682329 | 0.54277168 |
| CEP44 | protein_coding | -0.0682765 | 0.93100103 |
| STXBP2 | protein_coding | -0.0683306 | 0.62700128 |
| FRMD5 | protein_coding | -0.0684357 | 0.66484937 |
| HAT1 | protein_coding | -0.0684711 | 1.18300629 |
| L3MBTL3 | protein_coding | -0.0685626 | 0.40194142 |
| PRKACB-DT | lncRNA | -0.0686171 | 0.16262335 |
| REXO1 | protein_coding | -0.0686879 | 0.67324032 |
| PAK1 | protein_coding | -0.068828 | 1.3637854 |
| B9D2 | protein_coding | -0.0688629 | 0.32674157 |
| EIPR1 | protein_coding | -0.068991 | 1.01941445 |
| PARP2 | protein_coding | -0.0690295 | 1.26133118 |
| LLPH-DT | lncRNA | -0.0690593 | 0.08546639 |
| FDFT1 | protein_coding | -0.069098 | 1.67537133 |
| UBE2Q1 | protein_coding | -0.0691403 | 2.06523062 |
| DCAF15 | protein_coding | -0.0691707 | 0.74047913 |
| MAN1B1 | protein_coding | -0.0692115 | 1.46103102 |
| H2AZ1 | protein_coding | -0.0692205 | 2.09755623 |
| MMS22L | protein_coding | -0.0692619 | 0.96869248 |
| RNF168 | protein_coding | -0.0692871 | 1.4359041 |
| KATNBL1 | protein_coding | -0.0693097 | 0.53469373 |
| VAC14 | protein_coding | -0.0694812 | 1.33006882 |
| CNOT8 | protein_coding | -0.0695055 | 1.60829175 |
| ZNF571-AS1 | lncRNA | -0.0697007 | 0.3068413 |
| RECQL5 | protein_coding | -0.0697625 | 0.84820175 |
| RABGAP1 | protein_coding | -0.0697662 | 1.53543856 |
| RP11-441K8. | lncRNA | -0.0698037 | 0.2321466 |
| AKR1A1 | protein_coding | -0.0698593 | 1.60313364 |
| RPL27A | protein_coding | -0.069928 | 2.68507844 |
| MFSD10 | protein_coding | -0.069994 | 0.50010205 |
| SRP14-AS1 | lncRNA | -0.0700164 | 0.29739548 |
| API5 | protein_coding | -0.0700867 | 2.08860398 |

| | | | |
|---|---|---|---|
| MCM8 | protein_coding | -0.0700978 | 0.74734839 |
| RPL17 | protein_coding | -0.0701075 | 0.39569262 |
| XRCC4 | protein_coding | -0.0701244 | 0.28003895 |
| CDKN2AIP | protein_coding | -0.0701328 | 1.06624277 |
| HS2ST1 | protein_coding | -0.0702078 | 0.88869654 |
| YES1 | protein_coding | -0.0702144 | 0.93248707 |
| SLC30A9 | protein_coding | -0.070308 | 1.37838811 |
| DHX30 | protein_coding | -0.0703299 | 1.54808722 |
| PAN3-AS1 | lncRNA | -0.0703804 | 0.15391618 |
| ARFGAP3 | protein_coding | -0.0704053 | 1.2890923 |
| RNF103 | protein_coding | -0.0704701 | 0.90859591 |
| RP11-266L9.8 | lncRNA | -0.0706164 | 0.09960109 |
| PPP6C | protein_coding | -0.0706459 | 1.24230072 |
| DCBLD2 | protein_coding | -0.0707435 | 0.72606528 |
| DGUOK | protein_coding | -0.0707622 | 1.333455 |
| HSF2 | protein_coding | -0.0707805 | 0.71455222 |
| CHTF18 | protein_coding | -0.0707941 | 0.71321558 |
| EHMT2 | protein_coding | -0.0709066 | 2.01256066 |
| TMEM86A | protein_coding | -0.0709464 | 0.3315826 |
| DCAF16 | protein_coding | -0.0709477 | 1.80646488 |
| SEC31A | protein_coding | -0.0709817 | 2.20768912 |
| TUBB8B | protein_coding | -0.0710122 | 0.15172818 |
| EXO1 | protein_coding | -0.0710723 | 1.14615305 |
| SLC35F5 | protein_coding | -0.0711969 | 0.75038773 |
| CTB-25B13.1 | lncRNA | -0.0713042 | 0.23861569 |
| PABIR1 | protein_coding | -0.0713601 | 0.86368466 |
| NANP | protein_coding | -0.0713758 | 1.05128308 |
| BICD2 | protein_coding | -0.0714267 | 1.28748334 |
| MMP24OS | protein_coding | -0.071467 | 0.5361663 |
| RP11-495L19 | lncRNA | -0.0714795 | 0.09512234 |
| EIF4E2 | protein_coding | -0.0715537 | 1.47549756 |
| AP006621.5 | lncRNA | -0.0715597 | 0.35868003 |
| TFIP11 | protein_coding | -0.0715723 | 1.22218313 |
| TBC1D31 | protein_coding | -0.071621 | 0.83433051 |
| NAGPA | protein_coding | -0.0716244 | 0.33863553 |
| KDM3B | protein_coding | -0.071635 | 1.92672397 |
| PIK3R3 | protein_coding | -0.0716415 | 1.82258579 |
| RABIF | protein_coding | -0.0716694 | 0.81306456 |
| LINC02256 | lncRNA | -0.0717351 | 0.11095763 |
| CCT4 | protein_coding | -0.0717579 | 2.45578559 |
| EFTUD2 | protein_coding | -0.0718078 | 2.06527273 |
| ZNF74 | protein_coding | -0.0718405 | 0.97087982 |
| CSTF2 | protein_coding | -0.0718442 | 0.95695236 |
| DAPK3 | protein_coding | -0.0718998 | 0.67075336 |
| PODN | protein_coding | -0.0720444 | 0.08436485 |

| | | | |
|---|---|---|---|
| RBM26-AS1 | lncRNA | -0.0720924 | 0.19802314 |
| FCF1 | protein_coding | -0.0721418 | 1.41288514 |
| PLEKHG2 | protein_coding | -0.0721482 | 0.95484741 |
| PKN1 | protein_coding | -0.0722236 | 1.07519695 |
| POLDIP3 | protein_coding | -0.0722625 | 2.13186807 |
| SLC6A15 | protein_coding | -0.0724801 | 1.01983363 |
| PRELID2 | protein_coding | -0.0725464 | 1.14481792 |
| ATF7 | protein_coding | -0.0725885 | 1.16373456 |
| HSDL1 | protein_coding | -0.0725897 | 1.01170615 |
| FANCI | protein_coding | -0.0726404 | 2.44068058 |
| RANBP9 | protein_coding | -0.072672 | 1.81678181 |
| BUB1 | protein_coding | -0.0726754 | 1.7133791 |
| SSRP1 | protein_coding | -0.0727103 | 3.05220944 |
| TBC1D4 | protein_coding | -0.0727335 | 0.86128876 |
| SURF6 | protein_coding | -0.0727907 | 0.7656372 |
| CTNNA1 | protein_coding | -0.0728495 | 2.33653741 |
| LINC02870 | lncRNA | -0.0728539 | 0.15582076 |
| CDK10 | protein_coding | -0.0730078 | 1.351934 |
| RTF2 | protein_coding | -0.0730252 | 1.28808179 |
| MRPS17 | protein_coding | -0.0730369 | 0.45479452 |
| FAM149B1 | protein_coding | -0.0730461 | 0.3062448 |
| TMTC3 | protein_coding | -0.0730681 | 0.96959755 |
| PCDHA13 | protein_coding | -0.0731764 | 0.22193827 |
| MT-ND4L | protein_coding | -0.073184 | 1.63266816 |
| RAB5IF | protein_coding | -0.0731913 | 0.08561886 |
| CTNNAL1 | protein_coding | -0.0732139 | 1.06480418 |
| EPCAM | protein_coding | -0.0732199 | 2.34093087 |
| MICOS13 | protein_coding | -0.0732892 | 0.96405819 |
| ELOVL3 | protein_coding | -0.0733482 | 0.07694612 |
| GATAD2B | protein_coding | -0.0733521 | 1.61775635 |
| CYB561D2 | protein_coding | -0.0733629 | 0.45006596 |
| SNRPD1 | protein_coding | -0.0734173 | 2.40600385 |
| CACNA1D | protein_coding | -0.0734391 | 0.59841709 |
| ZBTB9 | protein_coding | -0.073452 | 0.60858812 |
| LIN7B | protein_coding | -0.0734676 | 0.4515862 |
| SET | protein_coding | -0.0734713 | 2.94487816 |
| TTC32 | protein_coding | -0.0734801 | 0.34481951 |
| KTN1 | protein_coding | -0.0735652 | 2.32058375 |
| NOP16 | protein_coding | -0.0736054 | 0.72714204 |
| HAUS1 | protein_coding | -0.0737911 | 1.39031879 |
| SMC2-DT | lncRNA | -0.0738526 | 0.11555559 |
| PPP1CC | protein_coding | -0.0738883 | 2.18546906 |
| VPS35 | protein_coding | -0.0738916 | 2.06101409 |
| SMARCC2 | protein_coding | -0.0739013 | 2.2084951 |
| N4BP2 | protein_coding | -0.0739951 | 1.69703193 |

| | | | |
|---|---|---|---|
| CKAP2L | protein_coding | -0.0740133 | 1.27012132 |
| CRKL | protein_coding | -0.0740662 | 2.09925554 |
| TFAM | protein_coding | -0.074085 | 0.79001502 |
| ARHGAP10 | protein_coding | -0.0740852 | 0.70259941 |
| RPS27L | protein_coding | -0.0741572 | 0.6561624 |
| PAFAH1B1 | protein_coding | -0.0742081 | 2.11978485 |
| LINC00926 | lncRNA | -0.0743137 | 0.16165255 |
| HDAC10 | protein_coding | -0.074328 | 0.14587194 |
| NR6A1 | protein_coding | -0.0744151 | 0.27497039 |
| ALG3 | protein_coding | -0.0744398 | 1.54795713 |
| SBF2-AS1 | lncRNA | -0.0744697 | 0.4664942 |
| MRGBP | protein_coding | -0.0745492 | 0.88275475 |
| LYRM7 | protein_coding | -0.0747913 | 0.76472598 |
| ZCCHC10 | protein_coding | -0.0748117 | 0.82485972 |
| TBC1D25 | protein_coding | -0.0748907 | 0.80478357 |
| TECTA | protein_coding | -0.0749223 | 0.15162583 |
| ZNF589 | protein_coding | -0.0749353 | 1.02876146 |
| RRP36 | protein_coding | -0.0750216 | 1.37728824 |
| EEFSEC | protein_coding | -0.0750276 | 0.7852359 |
| TMEM127 | protein_coding | -0.075068 | 1.22317525 |
| NDUFA1 | protein_coding | -0.0752612 | 1.58516112 |
| KAT5 | protein_coding | -0.0752931 | 1.33013914 |
| SAAL1 | protein_coding | -0.0753315 | 1.10878566 |
| HSPB1 | protein_coding | -0.0753707 | 1.00764265 |
| FKBP3 | protein_coding | -0.0754455 | 1.91478463 |
| AVIL | protein_coding | -0.0754603 | 2.73801006 |
| ANKRD39 | protein_coding | -0.0755821 | 0.32055143 |
| ANAPC5 | protein_coding | -0.0756009 | 2.55199651 |
| UFL1 | protein_coding | -0.0757135 | 0.63364129 |
| CTXN1 | protein_coding | -0.0757901 | 0.38581555 |
| ITGB1 | protein_coding | -0.0758145 | 1.79832522 |
| MTG1 | protein_coding | -0.0759251 | 0.48200239 |
| LINC00884 | lncRNA | -0.0759955 | 0.338627 |
| RP11-372B4. | lncRNA | -0.076016 | 0.0831327 |
| G2E3 | protein_coding | -0.076018 | 1.14087447 |
| TXNRD3 | protein_coding | -0.0760718 | 0.61745033 |
| MNX1-AS1 | lncRNA | -0.0760774 | 0.5622415 |
| BHLHE40-AS | lncRNA | -0.0761173 | 0.0860043 |
| RTL8A | protein_coding | -0.076118 | 1.27275275 |
| NFS1 | protein_coding | -0.0761346 | 0.66583272 |
| RP11-44F14. | lncRNA | -0.0761969 | 0.11336367 |
| CAPN7 | protein_coding | -0.0761992 | 1.23779541 |
| LLNLF-96A1. | lncRNA | -0.0762382 | 0.14841789 |
| RP11-304L19 | lncRNA | -0.0762733 | 0.0830484 |
| USP5 | protein_coding | -0.0763109 | 2.25662644 |

| | | | |
|---|---|---|---|
| COA4 | protein_coding | -0.0763709 | 1.1152839 |
| SIGMAR1 | protein_coding | -0.0763858 | 2.02144453 |
| PLCG1-AS1 | lncRNA | -0.0763868 | 0.10283639 |
| RAB14 | protein_coding | -0.0764122 | 1.5243844 |
| AQP1 | protein_coding | -0.0764199 | 0.10934187 |
| PROK2 | protein_coding | -0.0766737 | 0.65398143 |
| HNRNPK | protein_coding | -0.0767121 | 3.69509611 |
| CHTF8 | protein_coding | -0.0767252 | 0.13271911 |
| ZMYM6 | protein_coding | -0.0767365 | 0.48732292 |
| ZNF266 | protein_coding | -0.0767854 | 1.13945911 |
| RNF121 | protein_coding | -0.0767931 | 1.26677392 |
| SH3YL1 | protein_coding | -0.0768035 | 0.94590008 |
| AK2 | protein_coding | -0.076998 | 2.31563959 |
| APEX1 | protein_coding | -0.077126 | 2.27057257 |
| IVD | protein_coding | -0.0771282 | 1.40959803 |
| MAD2L2 | protein_coding | -0.0771954 | 1.37891438 |
| RHOT2 | protein_coding | -0.0772139 | 1.54305487 |
| SP2 | protein_coding | -0.0772192 | 0.95898642 |
| HDGFL3 | protein_coding | -0.0772662 | 1.4988844 |
| TCP11L1 | protein_coding | -0.0772946 | 1.09225386 |
| NBAS | protein_coding | -0.0773098 | 1.27553658 |
| C20orf194 | protein_coding | -0.0773615 | 0.51887981 |
| ZNF426 | protein_coding | -0.0774028 | 0.62138418 |
| RNASEH1-AS | lncRNA | -0.0774193 | 0.44603283 |
| SOX12 | protein_coding | -0.0774903 | 1.17068192 |
| CELF1 | protein_coding | -0.0775033 | 2.5046827 |
| STAG2 | protein_coding | -0.0775066 | 2.1693248 |
| CTD-2555O1 | lncRNA | -0.0775557 | 0.11706391 |
| HES6 | protein_coding | -0.0775905 | 0.61452316 |
| SETD1A | protein_coding | -0.0777019 | 1.59995488 |
| GGCT | protein_coding | -0.0777535 | 1.36138708 |
| PTPN1 | protein_coding | -0.0778078 | 2.25224829 |
| EPB41L5 | protein_coding | -0.0778315 | 0.66766636 |
| YTHDF2 | protein_coding | -0.0778718 | 1.82415443 |
| BORA | protein_coding | -0.0778873 | 0.55479382 |
| TBC1D10B | protein_coding | -0.0779113 | 1.49585674 |
| TMSB15B | protein_coding | -0.0781363 | 0.10281354 |
| STEEP1 | protein_coding | -0.0781588 | 0.82857497 |
| HNRNPD | protein_coding | -0.0782491 | 3.08866761 |
| LEPROT | protein_coding | -0.0782651 | 0.99014188 |
| SRI | protein_coding | -0.078266 | 0.7241721 |
| SYT13 | protein_coding | -0.078289 | 2.12000422 |
| SWAP70 | protein_coding | -0.0783126 | 1.12802132 |
| GNL2 | protein_coding | -0.078317 | 1.85394385 |
| PRR14L | protein_coding | -0.0783627 | 2.08146226 |

| | | | |
|---|---|---|---|
| H1-10-AS1 | lncRNA | -0.0783673 | 0.14161444 |
| UGDH-AS1 | lncRNA | -0.0784986 | 0.16234408 |
| WASL-DT | lncRNA | -0.0785125 | 0.31435562 |
| MAPK14 | protein_coding | -0.0785707 | 2.27051241 |
| SF3B6 | protein_coding | -0.0785813 | 1.25753457 |
| HRH3 | protein_coding | -0.0786352 | 0.73175392 |
| ARL8A | protein_coding | -0.0787249 | 0.95584192 |
| HSPA4 | protein_coding | -0.0787712 | 2.74619414 |
| SURF1 | protein_coding | -0.0788273 | 1.1282197 |
| LINC01359 | lncRNA | -0.0788343 | 0.07933421 |
| AC006538.1 | lncRNA | -0.0789013 | 0.23811625 |
| S1PR3 | protein_coding | -0.0789073 | 0.29254248 |
| ENTPD7 | protein_coding | -0.0789286 | 0.43100517 |
| AC097724.3 | lncRNA | -0.0789318 | 0.09311945 |
| ZNF223 | protein_coding | -0.0790821 | 0.19163835 |
| NME3 | protein_coding | -0.0791248 | 0.41723336 |
| SCAF8 | protein_coding | -0.0792463 | 1.3076735 |
| OGFOD1 | protein_coding | -0.0792703 | 1.85439523 |
| DEPDC4 | protein_coding | -0.0792873 | 0.27942572 |
| CERS5 | protein_coding | -0.0793202 | 1.15500296 |
| ELK1 | protein_coding | -0.0793449 | 1.4098166 |
| COPG1 | protein_coding | -0.0793545 | 2.60414998 |
| SWSAP1 | protein_coding | -0.079422 | 0.26825249 |
| RP11-15A1.3 | lncRNA | -0.0794244 | 0.10303171 |
| WASHC5 | protein_coding | -0.0794283 | 1.56998243 |
| RP11-533E19 | lncRNA | -0.0795308 | 0.14289419 |
| AMDHD2 | protein_coding | -0.0795673 | 0.65038348 |
| SELENOH | protein_coding | -0.0795675 | 0.36265688 |
| TARBP2 | protein_coding | -0.079575 | 1.18507635 |
| ZNHIT2 | protein_coding | -0.079612 | 0.34746008 |
| RP11-7O11.9 | lncRNA | -0.0796204 | 0.21117417 |
| FANCB | protein_coding | -0.0797111 | 0.45803863 |
| N6AMT1 | protein_coding | -0.0800315 | 0.93846598 |
| CAMTA1 | protein_coding | -0.0801142 | 1.17150588 |
| CHMP2A | protein_coding | -0.0801948 | 1.45540819 |
| RNPS1 | protein_coding | -0.0802136 | 2.70101553 |
| IQGAP1 | protein_coding | -0.0802688 | 3.00669077 |
| TARDBP | protein_coding | -0.0802738 | 2.54559989 |
| MFN1 | protein_coding | -0.0803234 | 1.83076862 |
| COX11 | protein_coding | -0.0804122 | 1.07176617 |
| DDA1 | protein_coding | -0.0804607 | 1.64809221 |
| ZNF276 | protein_coding | -0.0804679 | 0.92120303 |
| PHF3 | protein_coding | -0.0805367 | 2.30539033 |
| EMSY | protein_coding | -0.0805725 | 1.44837535 |
| LA16c-380H5 | lncRNA | -0.0806152 | 0.52647665 |

| | | | |
|---|---|---|---|
| SMARCAL1 | protein_coding | -0.0806488 | 0.66492031 |
| RNH1 | protein_coding | -0.0807648 | 1.67332535 |
| PDIA6 | protein_coding | -0.0808212 | 2.80719619 |
| SPRYD3 | protein_coding | -0.0809593 | 0.91880122 |
| EIF5B | protein_coding | -0.0809644 | 2.83651662 |
| CTD-3222D19 | lncRNA | -0.0809667 | 0.20720695 |
| BCDIN3D | protein_coding | -0.0810527 | 0.63358477 |
| SEPTIN7 | protein_coding | -0.081066 | 1.77051432 |
| NPDC1 | protein_coding | -0.0811599 | 0.80570187 |
| ARRDC1-AS1 | lncRNA | -0.0811882 | 0.85164599 |
| NEK6 | protein_coding | -0.0811935 | 1.55691917 |
| SOX4 | protein_coding | -0.0812359 | 2.3701131 |
| NUP133 | protein_coding | -0.0812402 | 2.08216678 |
| UBE2G1 | protein_coding | -0.0813078 | 1.13986361 |
| TRMT2A | protein_coding | -0.0814193 | 1.61945116 |
| PLS3 | protein_coding | -0.0814632 | 1.37945298 |
| ZNF207 | protein_coding | -0.0814845 | 1.51239975 |
| CYP51A1 | protein_coding | -0.0815164 | 0.38093787 |
| PPM1L | protein_coding | -0.0815181 | 1.2323189 |
| ZDHHC8 | protein_coding | -0.0815488 | 0.75323184 |
| KAT7 | protein_coding | -0.0815704 | 2.20595662 |
| NAXE | protein_coding | -0.0815997 | 1.74980744 |
| NCOA4 | protein_coding | -0.0818982 | 2.09143009 |
| ATP6V1E1 | protein_coding | -0.0819762 | 2.48595514 |
| DIS3 | protein_coding | -0.0820385 | 2.13388967 |
| HMGN1 | protein_coding | -0.0820557 | 2.13181205 |
| RSF1 | protein_coding | -0.0821316 | 1.78147485 |
| TBC1D30 | protein_coding | -0.0821445 | 0.68162961 |
| SPINT2 | protein_coding | -0.0822295 | 1.46659109 |
| PSD | protein_coding | -0.0822311 | 0.91467895 |
| DAZAP1 | protein_coding | -0.082279 | 1.98211917 |
| PCDHGB7 | protein_coding | -0.0822909 | 0.12502874 |
| SLC16A1 | protein_coding | -0.0823002 | 2.15648102 |
| SF3B3 | protein_coding | -0.082395 | 3.3600945 |
| ATG4D | protein_coding | -0.0824845 | 1.13899683 |
| TLN1 | protein_coding | -0.0825978 | 2.79960736 |
| SLC12A2 | protein_coding | -0.0826036 | 1.11770666 |
| CTNNBL1 | protein_coding | -0.082693 | 1.47811428 |
| VPS37C | protein_coding | -0.0827588 | 1.1023145 |
| UFC1 | protein_coding | -0.0828084 | 1.60182077 |
| RUFY2 | protein_coding | -0.0828538 | 0.69771415 |
| KPNA1 | protein_coding | -0.0828579 | 1.10134995 |
| TSTD3 | protein_coding | -0.0829154 | 0.39415564 |
| WDHD1 | protein_coding | -0.082917 | 1.74674063 |
| SEPHS1 | protein_coding | -0.0830844 | 1.51418663 |

| | | | |
|---|---|---|---|
| SLC38A2 | protein_coding | -0.0832773 | 2.55766365 |
| UBAP2L | protein_coding | -0.0833086 | 2.96744019 |
| BCCIP | protein_coding | -0.0834544 | 1.69225568 |
| DGKD | protein_coding | -0.083529 | 1.6029241 |
| TP53I3 | protein_coding | -0.0835418 | 0.33428323 |
| TMEM217 | protein_coding | -0.0835491 | 0.12224871 |
| NR2C2 | protein_coding | -0.083554 | 1.74222777 |
| NKIRAS1 | protein_coding | -0.083616 | 0.48865127 |
| LGALS7 | protein_coding | -0.083653 | 0.21411732 |
| BCR | protein_coding | -0.0837653 | 1.59825659 |
| IPO9-AS1 | lncRNA | -0.0837893 | 0.14445488 |
| HOXA1 | protein_coding | -0.0838027 | 0.14765131 |
| GGH | protein_coding | -0.0838238 | 1.11511754 |
| C19orf71 | protein_coding | -0.0838464 | 0.29479397 |
| ALG5 | protein_coding | -0.0838631 | 1.20138255 |
| HIPK3 | protein_coding | -0.0838863 | 1.56441463 |
| SRSF10 | protein_coding | -0.0839337 | 2.13323947 |
| EDEM3 | protein_coding | -0.0839382 | 1.97220811 |
| PIGK | protein_coding | -0.0840626 | 1.20373044 |
| TTLL5 | protein_coding | -0.0840773 | 2.08010867 |
| PPHLN1 | protein_coding | -0.084132 | 1.7922385 |
| CIAPIN1 | protein_coding | -0.0841516 | 1.89541114 |
| ACTN3 | protein_coding | -0.0841774 | 0.15309756 |
| TMEM33 | protein_coding | -0.0842041 | 1.63636396 |
| ISCA2 | protein_coding | -0.0842197 | 1.30033967 |
| CBR3-AS1 | lncRNA | -0.0842461 | 0.12112375 |
| ODAD3 | protein_coding | -0.084266 | 0.61666519 |
| GBE1 | protein_coding | -0.0843305 | 0.5822792 |
| SUMO1 | protein_coding | -0.084435 | 1.62940211 |
| APEH | protein_coding | -0.0845227 | 1.74270336 |
| HDAC4 | protein_coding | -0.0845705 | 1.28239311 |
| CLNS1A | protein_coding | -0.0847448 | 2.32183855 |
| RP3-395C13. | lncRNA | -0.0847989 | 0.08783261 |
| ARMC7 | protein_coding | -0.0848056 | 0.84829318 |
| TPCN2 | protein_coding | -0.0848105 | 0.39373063 |
| TCEAL1 | protein_coding | -0.0848519 | 0.47910714 |
| AP001469.7 | lncRNA | -0.0848635 | 0.15355337 |
| F11R | protein_coding | -0.084942 | 1.69120457 |
| PLPBP | protein_coding | -0.0850089 | 1.06643059 |
| SH3BGRL2 | protein_coding | -0.0850423 | 1.53990583 |
| RP11-1114A! | lncRNA | -0.0850691 | 0.21047247 |
| IFT27 | protein_coding | -0.0851191 | 0.9741998 |
| ZBTB5 | protein_coding | -0.0852341 | 1.77061577 |
| RAP1B | protein_coding | -0.0852392 | 1.11701954 |
| CENPK | protein_coding | -0.0853281 | 1.03850912 |

| | | | |
|---|---|---|---|
| ZSCAN18 | protein_coding | -0.0853365 | 0.87061092 |
| SON | protein_coding | -0.0853952 | 3.59182512 |
| USP37 | protein_coding | -0.0854273 | 2.39768058 |
| FAAP20 | protein_coding | -0.0854402 | 0.65874878 |
| TIMM8B | protein_coding | -0.0855079 | 1.04797964 |
| SRSF7 | protein_coding | -0.085515 | 3.18786216 |
| SLC35A5 | protein_coding | -0.0856629 | 1.20628033 |
| WWC2 | protein_coding | -0.0856634 | 1.52108536 |
| ZNF25 | protein_coding | -0.0857013 | 0.33160414 |
| YWHAB | protein_coding | -0.0857963 | 3.19862298 |
| MED10 | protein_coding | -0.0858121 | 0.89946997 |
| APPL1 | protein_coding | -0.0858437 | 1.36903462 |
| MPG | protein_coding | -0.0859076 | 0.57697647 |
| SCCPDH | protein_coding | -0.0859466 | 1.80900743 |
| NPTN | protein_coding | -0.0860473 | 1.82506646 |
| RP11-196G1 | lncRNA | -0.0861408 | 0.10765707 |
| NOC2L | protein_coding | -0.0861454 | 1.90053616 |
| CREB3L2 | protein_coding | -0.0861809 | 2.38210789 |
| APMAP | protein_coding | -0.0862559 | 2.07707198 |
| PRR5L | protein_coding | -0.0862756 | 0.22181318 |
| ZBED4 | protein_coding | -0.0862947 | 1.61424905 |
| RXRB | protein_coding | -0.0863396 | 1.73471758 |
| MIR3936HG | lncRNA | -0.0863745 | 0.32128706 |
| NUDCD3 | protein_coding | -0.0864285 | 2.24470405 |
| KPNA6 | protein_coding | -0.0864881 | 2.13262682 |
| CARNMT1 | protein_coding | -0.0865022 | 0.98410451 |
| BANP | protein_coding | -0.0865341 | 0.71728841 |
| ENSA | protein_coding | -0.0865562 | 3.0598428 |
| GID4 | protein_coding | -0.0866824 | 1.20910764 |
| SH3BP5L | protein_coding | -0.0868005 | 1.6391289 |
| PRSS27 | protein_coding | -0.0868861 | 0.42650464 |
| CTD-2515O1 | protein_coding | -0.0869234 | 0.27062795 |
| TMEM91 | protein_coding | -0.0869448 | 0.10155136 |
| TAF12 | protein_coding | -0.0869497 | 1.05661928 |
| CS | protein_coding | -0.0870182 | 3.10952985 |
| FTCD | protein_coding | -0.0871042 | 0.110041 |
| TOM1L2 | protein_coding | -0.0871368 | 1.63432873 |
| RNF126 | protein_coding | -0.0871516 | 0.58396642 |
| PCYOX1L | protein_coding | -0.0871802 | 1.59466638 |
| TAF4 | protein_coding | -0.0872093 | 1.56961966 |
| CYB5B | protein_coding | -0.0872114 | 2.12918821 |
| THRAP3 | protein_coding | -0.0872948 | 3.10192511 |
| DNAJC4 | protein_coding | -0.0873346 | 0.76231887 |
| PRPSAP2 | protein_coding | -0.0873593 | 1.39734148 |
| GAN | protein_coding | -0.0873705 | 0.8648279 |

| | | | |
|---|---|---|---|
| EOLA1 | protein_coding | -0.0874391 | 0.8039792 |
| TIMM8A | protein_coding | -0.0875589 | 0.85657717 |
| MRPL49 | protein_coding | -0.0876532 | 2.54118001 |
| SIMC1 | protein_coding | -0.087696 | 0.86806474 |
| DDX28 | protein_coding | -0.0877741 | 0.87699254 |
| CTD-2517M2 | lncRNA | -0.0878411 | 0.12281776 |
| KIF3A | protein_coding | -0.0878499 | 1.0523831 |
| KRT8 | protein_coding | -0.0879172 | 3.45739494 |
| ADK | protein_coding | -0.0879491 | 1.71979994 |
| GTF2IRD1 | protein_coding | -0.0879949 | 1.16339084 |
| LTN1 | protein_coding | -0.0880397 | 1.9573951 |
| LHFPL4 | protein_coding | -0.0882208 | 0.89150546 |
| ZNF394 | protein_coding | -0.0882899 | 0.79707659 |
| MADD | protein_coding | -0.0883823 | 2.58771888 |
| SAMM50 | protein_coding | -0.0884719 | 1.21966629 |
| SMS | protein_coding | -0.0885339 | 2.48229321 |
| PGM3 | protein_coding | -0.0885597 | 1.45126911 |
| TRAF7 | protein_coding | -0.0885842 | 2.7514211 |
| DZANK1 | protein_coding | -0.0886145 | 0.25864657 |
| ING2 | protein_coding | -0.0887279 | 0.97760265 |
| MICU2 | protein_coding | -0.0887427 | 1.04670281 |
| GFOD2 | protein_coding | -0.0887553 | 1.28560741 |
| SACM1L | protein_coding | -0.0887848 | 0.93398435 |
| CHTOP | protein_coding | -0.0888083 | 2.18440486 |
| AP3B1 | protein_coding | -0.0888242 | 1.57202768 |
| POLE3 | protein_coding | -0.0888264 | 2.46485446 |
| TMEM242 | protein_coding | -0.0889393 | 0.77021699 |
| GPALPP1 | protein_coding | -0.0890088 | 0.90390694 |
| PHF12 | protein_coding | -0.0890716 | 2.33268465 |
| LIN9 | protein_coding | -0.0891267 | 1.20843585 |
| KCNK3 | protein_coding | -0.0891663 | 0.71767841 |
| FRA10AC1 | protein_coding | -0.0891791 | 1.29574525 |
| ZNF114 | protein_coding | -0.0891825 | 0.57086912 |
| SF3A1 | protein_coding | -0.0892025 | 3.4233529 |
| PPP1R3F | protein_coding | -0.0892452 | 0.50990846 |
| RNF43 | protein_coding | -0.0892461 | 0.0875887 |
| SDF2 | protein_coding | -0.0892476 | 1.44832755 |
| MMP11 | protein_coding | -0.0893441 | 0.1971843 |
| SUPT16H | protein_coding | -0.0893493 | 3.62090197 |
| FUT9 | protein_coding | -0.0893558 | 0.78165894 |
| MYO19 | protein_coding | -0.0893564 | 2.76591008 |
| SKOR1 | protein_coding | -0.0893779 | 0.54882014 |
| IQCN | protein_coding | -0.0894797 | 0.42661252 |
| CCT6B | protein_coding | -0.0894935 | 0.09024797 |
| REPIN1-AS1 | lncRNA | -0.0895077 | 0.10293263 |

| | | | |
|---|---|---|---|
| TOP2A | protein_coding | -0.0895286 | 4.0900441 |
| PRELID3A | protein_coding | -0.0896457 | 0.46983986 |
| TTI1 | protein_coding | -0.0896964 | 1.8502127 |
| DLAT | protein_coding | -0.0897652 | 1.56299422 |
| BCAR1 | protein_coding | -0.0899027 | 1.44167038 |
| ALKBH6 | protein_coding | -0.0899531 | 0.28864967 |
| TMEM199 | protein_coding | -0.0899779 | 1.40791216 |
| CSNK2A1 | protein_coding | -0.0900413 | 2.3378888 |
| TRAPPC9 | protein_coding | -0.0901036 | 1.60623878 |
| SCMH1 | protein_coding | -0.0901461 | 1.73172532 |
| UBXN7 | protein_coding | -0.0902738 | 2.50219825 |
| FAM78A | protein_coding | -0.0903243 | 0.72627019 |
| SAFB | protein_coding | -0.0903605 | 2.20523125 |
| ZKSCAN5 | protein_coding | -0.0903836 | 1.42000912 |
| MRPL27 | protein_coding | -0.0905345 | 1.38822675 |
| EIF4E | protein_coding | -0.0905378 | 1.61615019 |
| AC156455.1 | lncRNA | -0.0908363 | 0.15798525 |
| SMIM15 | protein_coding | -0.0910439 | 1.34235264 |
| UTP18 | protein_coding | -0.0911201 | 1.44853546 |
| PYGO2 | protein_coding | -0.0911755 | 2.56965123 |
| ZNF596 | protein_coding | -0.0912514 | 0.22322978 |
| PAF1 | protein_coding | -0.0913078 | 1.68382336 |
| RP11-613M1 | lncRNA | -0.0913526 | 0.54017736 |
| CCDC121 | protein_coding | -0.0914175 | 0.42194639 |
| NLRX1 | protein_coding | -0.0914921 | 0.68221661 |
| DIP2C | protein_coding | -0.0916006 | 0.79808823 |
| PIP4P1 | protein_coding | -0.0916432 | 1.93551412 |
| TIMM10B | protein_coding | -0.0916666 | 1.67005969 |
| KCTD3 | protein_coding | -0.0916858 | 2.98454768 |
| CTD-2256P15 | lncRNA | -0.0916977 | 0.2473343 |
| MT-ND4 | protein_coding | -0.0917119 | 2.72959744 |
| RNPC3 | protein_coding | -0.0917774 | 0.74374372 |
| NAIP | protein_coding | -0.0917844 | 0.11156238 |
| PRKCZ-AS1 | lncRNA | -0.0918309 | 0.19897114 |
| ZC3H8 | protein_coding | -0.0918365 | 1.19520853 |
| SLC30A5 | protein_coding | -0.0918692 | 2.13562389 |
| HEATR6 | protein_coding | -0.0918955 | 1.1925334 |
| SHC1 | protein_coding | -0.0919209 | 2.01801711 |
| MSH5-SAPCD | protein_coding | -0.0919424 | 0.14934027 |
| PER3 | protein_coding | -0.0919611 | 2.32939273 |
| SELENBP1 | protein_coding | -0.0919622 | 0.57785392 |
| NDUFA8 | protein_coding | -0.091992 | 1.74718439 |
| SUV39H1 | protein_coding | -0.0920367 | 1.12678731 |
| ENTPD5 | protein_coding | -0.0920377 | 3.02419191 |
| SEC61A1 | protein_coding | -0.0920785 | 4.01641367 |

| | | | |
|---|---|---|---|
| SGSM2 | protein_coding | -0.0921267 | 1.35748732 |
| LAGE3 | protein_coding | -0.0921447 | 0.79064239 |
| PNMA1 | protein_coding | -0.0922116 | 1.61085673 |
| C1orf74 | protein_coding | -0.0922792 | 0.75438913 |
| ZNF135 | protein_coding | -0.0922812 | 0.36685502 |
| C9orf40 | protein_coding | -0.0923456 | 1.13166593 |
| MCF2L2 | protein_coding | -0.0923739 | 0.7972871 |
| RP1-90K10.4 | lncRNA | -0.0923984 | 0.1882468 |
| PPIB | protein_coding | -0.0924026 | 3.01471279 |
| CUEDC1 | protein_coding | -0.092435 | 1.22055097 |
| RP11-142E9. | lncRNA | -0.09252 | 0.26978868 |
| NSMCE2 | protein_coding | -0.0925474 | 1.01334643 |
| RPGRIP1L | protein_coding | -0.0926142 | 1.54704831 |
| COMMD7 | protein_coding | -0.0926294 | 0.58411568 |
| GFM2 | protein_coding | -0.0926529 | 1.68140977 |
| SLC35E3 | protein_coding | -0.0926846 | 1.01159793 |
| ARPC5L | protein_coding | -0.0927257 | 1.53217285 |
| CWF19L1 | protein_coding | -0.0928033 | 1.0177035 |
| MYDGF | protein_coding | -0.0928365 | 1.33157835 |
| RNGTT | protein_coding | -0.092915 | 1.78618603 |
| PRPF18 | protein_coding | -0.0929316 | 0.37982537 |
| UBALD2 | protein_coding | -0.0929402 | 1.33357683 |
| BAG1 | protein_coding | -0.09302 | 1.1832 |
| CLK3 | protein_coding | -0.0930397 | 1.63234773 |
| ANKLE2 | protein_coding | -0.0930969 | 2.77299245 |
| GALNT12 | protein_coding | -0.0931224 | 0.66121506 |
| CYB5D1 | protein_coding | -0.0931296 | 0.2981829 |
| TMEM192 | protein_coding | -0.0932166 | 1.46245986 |
| ZNF710 | protein_coding | -0.0932194 | 1.12371702 |
| MDM1 | protein_coding | -0.0932912 | 1.28841504 |
| RNF138 | protein_coding | -0.093405 | 1.33047316 |
| NGRN | protein_coding | -0.09341 | 0.56869727 |
| MIGA1 | protein_coding | -0.0935287 | 1.22318381 |
| SCAPER | protein_coding | -0.0935574 | 1.50340146 |
| NIF3L1 | protein_coding | -0.0936081 | 1.4272961 |
| LACTB2 | protein_coding | -0.0936753 | 0.52671969 |
| COQ7 | protein_coding | -0.0937035 | 1.39895126 |
| DPF3 | protein_coding | -0.0938385 | 1.02243449 |
| PAXBP1-AS1 | lncRNA | -0.0938513 | 0.44321703 |
| RP11-2B2.1 | lncRNA | -0.0938763 | 0.10891591 |
| LINC01621 | lncRNA | -0.0938933 | 0.14128347 |
| HNRNPL | protein_coding | -0.0938999 | 3.18358039 |
| MORF4L2 | protein_coding | -0.0939362 | 3.3017602 |
| PTBP3 | protein_coding | -0.0939527 | 1.60614841 |
| KLHL8 | protein_coding | -0.0940502 | 1.70242278 |

| | | | |
|---|---|---|---|
| TBCB | protein_coding | -0.0940634 | 2.61415357 |
| TECPR2 | protein_coding | -0.0940833 | 1.34036428 |
| RALBP1 | protein_coding | -0.0940919 | 1.43322869 |
| PSPC1 | protein_coding | -0.0941534 | 1.83852949 |
| EFCAB6-DT | lncRNA | -0.0941571 | 0.13204794 |
| CPSF1 | protein_coding | -0.0942042 | 2.03118976 |
| PLA2G6 | protein_coding | -0.0943345 | 1.22835889 |
| BTRC | protein_coding | -0.0943741 | 1.21452431 |
| PPP2CA | protein_coding | -0.0944556 | 2.19003349 |
| SLC52A2 | protein_coding | -0.0944624 | 1.24016671 |
| KLHL21 | protein_coding | -0.0944708 | 1.07419628 |
| POLE2 | protein_coding | -0.0944979 | 1.52220523 |
| DCST1-AS1 | lncRNA | -0.0945349 | 0.18808293 |
| SPTAN1 | protein_coding | -0.0945588 | 3.29433537 |
| EXOSC3 | protein_coding | -0.0945826 | 0.77506758 |
| FOXRED1 | protein_coding | -0.0946138 | 1.93143218 |
| ACSL4 | protein_coding | -0.0946302 | 1.66830314 |
| SERINC3 | protein_coding | -0.0946357 | 2.00559465 |
| HSPA14 | protein_coding | -0.0947564 | 0.80225182 |
| FZR1 | protein_coding | -0.0947652 | 1.72380898 |
| MBLAC2 | protein_coding | -0.0947911 | 0.60859049 |
| PAX8 | protein_coding | -0.0947986 | 0.30299969 |
| SLC35G1 | protein_coding | -0.0948675 | 0.82917044 |
| PITPNA | protein_coding | -0.0949028 | 1.4347242 |
| GIPC1 | protein_coding | -0.0949155 | 1.14967948 |
| C16orf91 | protein_coding | -0.0949961 | 1.01067732 |
| KAT8 | protein_coding | -0.0950718 | 1.50883417 |
| MHENCR | lncRNA | -0.095147 | 0.43213221 |
| NCK2 | protein_coding | -0.095161 | 1.82206335 |
| CADM1 | protein_coding | -0.0952026 | 3.11834995 |
| ATP5MC3 | protein_coding | -0.095245 | 2.58158674 |
| FKBP1A | protein_coding | -0.0952541 | 2.12921716 |
| SYVN1 | protein_coding | -0.0953616 | 1.8785483 |
| C1orf21-DT | lncRNA | -0.0954014 | 0.12138521 |
| FOXN2 | protein_coding | -0.0954831 | 1.22410059 |
| U2AF1 | protein_coding | -0.0954833 | 0.10829465 |
| PRRC1 | protein_coding | -0.0954906 | 1.19776407 |
| IFT172 | protein_coding | -0.095563 | 0.39772533 |
| MCF2L | protein_coding | -0.0955648 | 1.78542805 |
| GART | protein_coding | -0.0955668 | 3.48043821 |
| KLHL23 | protein_coding | -0.0955833 | 2.03735906 |
| EPOP | protein_coding | -0.0955864 | 0.64534261 |
| HELLS | protein_coding | -0.0956232 | 1.59782767 |
| KANSL1-AS1 | lncRNA | -0.0956433 | 0.25515237 |
| SLC39A3 | protein_coding | -0.095653 | 0.49855639 |

| | | | |
|---|---|---|---|
| ZNF354B | protein_coding | -0.095713 | 0.57093387 |
| GPX1 | protein_coding | -0.0957222 | 1.1550234 |
| DDX39A | protein_coding | -0.0957426 | 3.40987193 |
| KDM5A | protein_coding | -0.0957527 | 2.3351863 |
| EEF1AKMT2 | protein_coding | -0.0957592 | 0.34001953 |
| PSMG2 | protein_coding | -0.0958144 | 1.73886151 |
| ACTN4 | protein_coding | -0.0958184 | 3.56915159 |
| SPIN1 | protein_coding | -0.0959384 | 2.39963163 |
| NKAIN1 | protein_coding | -0.096006 | 0.68630252 |
| LEO1 | protein_coding | -0.0960186 | 1.90637121 |
| HCG24 | lncRNA | -0.0960269 | 0.10064466 |
| ATP5MC1 | protein_coding | -0.0960494 | 2.27848057 |
| INTS1 | protein_coding | -0.0960507 | 1.49709815 |
| HGH1 | protein_coding | -0.096092 | 1.33164117 |
| BAP1 | protein_coding | -0.0961163 | 2.37113053 |
| SRGAP2B | protein_coding | -0.0961628 | 0.21465899 |
| MRPL35 | protein_coding | -0.0962159 | 1.72294881 |
| BTF3L4 | protein_coding | -0.0962162 | 2.04322315 |
| BRPF1 | protein_coding | -0.0962978 | 1.21809014 |
| ZNF540 | protein_coding | -0.0963143 | 0.38879568 |
| CROCC | protein_coding | -0.0963254 | 0.62238807 |
| ARF5 | protein_coding | -0.0963407 | 2.35119363 |
| TRA2B | protein_coding | -0.0964026 | 3.22938105 |
| MLEC | protein_coding | -0.0964369 | 4.05588156 |
| ATXN3 | protein_coding | -0.0965856 | 1.38978712 |
| VAPA | protein_coding | -0.0966939 | 2.97357666 |
| NOLC1 | protein_coding | -0.0966997 | 3.11493241 |
| STARD7 | protein_coding | -0.0967728 | 4.01806473 |
| RPL18 | protein_coding | -0.0968597 | 3.9114473 |
| PURA | protein_coding | -0.0969721 | 1.37355889 |
| DDX11 | protein_coding | -0.0970137 | 2.25820157 |
| NUDCD1 | protein_coding | -0.0970232 | 1.59201572 |
| DPH6 | protein_coding | -0.0970832 | 0.77246466 |
| PIGM | protein_coding | -0.0971602 | 1.47558887 |
| EXTL2 | protein_coding | -0.097256 | 0.90398579 |
| UXT | protein_coding | -0.0974255 | 1.33407925 |
| PEX16 | protein_coding | -0.0974881 | 1.08760057 |
| UNC13B | protein_coding | -0.0975752 | 2.19151202 |
| PDZD2 | protein_coding | -0.097578 | 1.30230444 |
| DYSF | protein_coding | -0.0976015 | 0.67392283 |
| KCNH2 | protein_coding | -0.0976266 | 1.65791779 |
| UQCRQ | protein_coding | -0.0977201 | 2.16148543 |
| ABHD8 | protein_coding | -0.0977288 | 0.78082671 |
| DEAF1 | protein_coding | -0.0977673 | 1.92492225 |
| EXOSC9 | protein_coding | -0.0977866 | 2.3729722 |

| | | | |
|---|---|---|---|
| PRPF31 | protein_coding | -0.0977964 | 2.18768148 |
| LINC00339 | lncRNA | -0.097867 | 0.58803586 |
| SHCBP1 | protein_coding | -0.0978817 | 1.0002876 |
| NUDCD2 | protein_coding | -0.0979198 | 1.81818556 |
| PAK2 | protein_coding | -0.0979212 | 3.28236848 |
| C20orf96 | protein_coding | -0.0979582 | 0.9822408 |
| ZNF512B | protein_coding | -0.0981698 | 2.87909382 |
| MFSD3 | protein_coding | -0.0981969 | 0.73206867 |
| CIC | protein_coding | -0.0982312 | 2.11584691 |
| ADH5 | protein_coding | -0.0983032 | 2.47480209 |
| MAGED2 | protein_coding | -0.0983119 | 1.94633636 |
| PINK1 | protein_coding | -0.0983903 | 1.66119887 |
| TXNDC5 | protein_coding | -0.0984346 | 0.15619825 |
| GHET1 | lncRNA | -0.0984948 | 0.19366406 |
| THAP10 | protein_coding | -0.0985483 | 0.24556869 |
| EEF2KMT | protein_coding | -0.0986556 | 1.26404393 |
| DNASE1L2 | protein_coding | -0.0988248 | 0.3170007 |
| FPGT | protein_coding | -0.0988577 | 0.72565964 |
| FAF2 | protein_coding | -0.0988793 | 2.17774125 |
| ZXDC | protein_coding | -0.0988875 | 1.5896219 |
| SCAMP1 | protein_coding | -0.0988904 | 1.20823582 |
| RP11-708J19 | lncRNA | -0.0989271 | 0.17165525 |
| PHPT1 | protein_coding | -0.0990108 | 1.65576917 |
| PDZD11 | protein_coding | -0.0992054 | 1.50197614 |
| HMGCL | protein_coding | -0.09923 | 0.85996481 |
| GTF2B | protein_coding | -0.0992705 | 1.3394905 |
| SRD5A3 | protein_coding | -0.0994602 | 0.61081133 |
| TRPT1 | protein_coding | -0.0994876 | 1.15238553 |
| LDHB | protein_coding | -0.0996507 | 0.72180539 |
| RP5-1021I20 | lncRNA | -0.0997352 | 0.29485651 |
| RP11-448G1! | lncRNA | -0.0997556 | 0.17201083 |
| RPS13 | protein_coding | -0.0997849 | 4.21248321 |
| PHF6 | protein_coding | -0.0998094 | 1.66940632 |
| TBC1D8B | protein_coding | -0.0998579 | 0.34251117 |
| SHE | protein_coding | -0.0999524 | 2.83747897 |
| CKAP5 | protein_coding | -0.1001912 | 3.96613591 |
| GMFB | protein_coding | -0.1002766 | 1.14660807 |
| GAPVD1 | protein_coding | -0.1003022 | 2.02681421 |
| AC114271.2 | lncRNA | -0.1003336 | 0.65160723 |
| BAG6 | protein_coding | -0.1003673 | 3.72386289 |
| THAP5 | protein_coding | -0.1003833 | 0.97719057 |
| TMEM218 | protein_coding | -0.1003903 | 0.79622095 |
| FRYL | protein_coding | -0.1003912 | 2.94669861 |
| PSMD8 | protein_coding | -0.1005001 | 3.30633678 |
| CTD-2287O1( | lncRNA | -0.1005325 | 0.22362788 |

| | | | |
|---|---|---|---|
| DNAL1 | protein_coding | -0.1005583 | 0.82501892 |
| SYNM | protein_coding | -0.1005814 | 2.25993228 |
| ABHD14A | protein_coding | -0.1005841 | 0.29327764 |
| NDUFB10 | protein_coding | -0.1005856 | 2.20033234 |
| BRK1 | protein_coding | -0.1007169 | 2.2184596 |
| FAM234B | protein_coding | -0.1007322 | 2.20444977 |
| SLC25A3 | protein_coding | -0.1007757 | 3.83904106 |
| LRRC59 | protein_coding | -0.100785 | 3.34291261 |
| DLST | protein_coding | -0.1008504 | 3.62152061 |
| HIF1AN | protein_coding | -0.1008574 | 1.88433633 |
| NME4 | protein_coding | -0.1008866 | 1.82358848 |
| FBXL4 | protein_coding | -0.1009745 | 0.71690274 |
| UBXN2A | protein_coding | -0.101007 | 1.60751493 |
| SRSF3 | protein_coding | -0.1010127 | 3.10689611 |
| AJM1 | protein_coding | -0.1010545 | 0.48356133 |
| TIMM50 | protein_coding | -0.1010995 | 2.46665114 |
| RP11-982M1 | lncRNA | -0.101102 | 0.10523581 |
| CSMD2 | protein_coding | -0.1011749 | 0.23931298 |
| ZFYVE9 | protein_coding | -0.1011829 | 1.08467392 |
| RP1-257I20.1 | lncRNA | -0.1012099 | 0.19917416 |
| CHST11 | protein_coding | -0.1012467 | 1.17297103 |
| TYW1B | protein_coding | -0.1012601 | 0.40327678 |
| DCUN1D1 | protein_coding | -0.1012624 | 1.4674388 |
| FDPS | protein_coding | -0.1012739 | 2.98877144 |
| ASPHD1 | protein_coding | -0.1013114 | 1.29713103 |
| ZDHHC18 | protein_coding | -0.1013372 | 1.70287839 |
| BRSK1 | protein_coding | -0.1013627 | 1.47846782 |
| ABCB10 | protein_coding | -0.1013827 | 0.94213222 |
| BBS5 | protein_coding | -0.1013965 | 0.27922703 |
| TAS2R20 | protein_coding | -0.1014172 | 0.1661814 |
| NDUFV3 | protein_coding | -0.1014265 | 1.71093748 |
| C22orf39 | protein_coding | -0.1014778 | 1.57848971 |
| WDR82 | protein_coding | -0.1014795 | 3.11520753 |
| SUPT5H | protein_coding | -0.1014802 | 3.30090572 |
| SPDL1 | protein_coding | -0.1014988 | 1.71895592 |
| ZBTB14 | protein_coding | -0.1015413 | 1.43349929 |
| INTS11 | protein_coding | -0.1015782 | 1.78887844 |
| CAND1 | protein_coding | -0.1016232 | 3.79362126 |
| SRSF9 | protein_coding | -0.1016435 | 2.95988221 |
| IDH3G | protein_coding | -0.1016864 | 1.93924466 |
| COQ3 | protein_coding | -0.1017288 | 0.76035375 |
| MON1B | protein_coding | -0.1017991 | 2.57113908 |
| MBOAT7 | protein_coding | -0.1018332 | 1.94118354 |
| KAT6B | protein_coding | -0.1018513 | 1.5611629 |
| RP3-475N16. | lncRNA | -0.1019059 | 0.14726511 |

| | | | |
|---|---|---|---|
| DDX54 | protein_coding | -0.1020353 | 1.83038129 |
| RIOK3 | protein_coding | -0.1021365 | 1.73411308 |
| EAPP | protein_coding | -0.102196 | 1.33655756 |
| MEMO1 | protein_coding | -0.1022877 | 0.25887499 |
| MYL6 | protein_coding | -0.1023011 | 1.69534557 |
| ZNF510 | protein_coding | -0.1023017 | 1.50654504 |
| FAM221A | protein_coding | -0.1023383 | 1.17204263 |
| GCDH | protein_coding | -0.102373 | 1.43789448 |
| SLC15A2 | protein_coding | -0.1024846 | 0.14055077 |
| RAB5C | protein_coding | -0.1025976 | 1.12704248 |
| RP1-223E5.4 | lncRNA | -0.1026484 | 0.29097006 |
| WDR89 | protein_coding | -0.1027289 | 1.32657306 |
| CCDC91 | protein_coding | -0.1028116 | 1.20299853 |
| TPM3 | protein_coding | -0.102858 | 3.8659315 |
| UBL3 | protein_coding | -0.102866 | 1.57511275 |
| SPAG7 | protein_coding | -0.1029176 | 1.69946299 |
| ARRDC2 | protein_coding | -0.1029443 | 0.74615585 |
| PDIK1L | protein_coding | -0.1029589 | 2.07342561 |
| MCMBP | protein_coding | -0.1030241 | 2.32789811 |
| CCDC183 | protein_coding | -0.1031811 | 0.18377527 |
| CASC2 | lncRNA | -0.1031815 | 0.26185998 |
| FUBP1 | protein_coding | -0.1032161 | 2.91450166 |
| CYSLTR2 | protein_coding | -0.1032517 | 0.5029429 |
| GPR137 | protein_coding | -0.1032568 | 2.63966574 |
| RP11-783K16 | lncRNA | -0.1033079 | 0.23267916 |
| CNKSR2 | protein_coding | -0.1033168 | 1.06305479 |
| SMN2 | protein_coding | -0.1034874 | 0.45844631 |
| SUGP1 | protein_coding | -0.1035391 | 1.47626665 |
| RNPEP | protein_coding | -0.1035638 | 1.27233237 |
| RNMT | protein_coding | -0.1036281 | 2.48198875 |
| ZNF790 | protein_coding | -0.1036379 | 0.6118427 |
| TRIM68 | protein_coding | -0.1036412 | 0.67419194 |
| NUP35 | protein_coding | -0.1037121 | 1.61848133 |
| RNF2 | protein_coding | -0.1037662 | 1.04639879 |
| WDR76 | protein_coding | -0.1040294 | 3.01804451 |
| DPY19L3 | protein_coding | -0.1040526 | 1.58736472 |
| ENOX2 | protein_coding | -0.1040632 | 1.82686032 |
| ZNF622 | protein_coding | -0.1041065 | 1.41730057 |
| CHST6 | protein_coding | -0.1041191 | 0.18377527 |
| SELENOF | protein_coding | -0.1041373 | 1.94718124 |
| NAA50 | protein_coding | -0.1041408 | 2.59053609 |
| USP22 | protein_coding | -0.1041437 | 3.82031029 |
| PTGFRN | protein_coding | -0.1041473 | 0.4894197 |
| SIRT6 | protein_coding | -0.1041497 | 1.23147445 |
| RNF139 | protein_coding | -0.1041559 | 1.3842077 |

| | | | |
|---|---|---|---|
| SCARF2 | protein_coding | -0.1042361 | 0.17375347 |
| QSER1 | protein_coding | -0.104288 | 2.57937291 |
| ERICH1 | protein_coding | -0.1043744 | 0.71186684 |
| COMMD3 | protein_coding | -0.1048071 | 0.47004269 |
| RIBC1 | protein_coding | -0.1049625 | 0.17828769 |
| CRMP1 | protein_coding | -0.104972 | 4.02992095 |
| WDR97 | protein_coding | -0.1050069 | 0.173888 |
| ZNF70 | protein_coding | -0.1050401 | 0.58094787 |
| C11orf58 | protein_coding | -0.1050995 | 3.05444377 |
| TOLLIP | protein_coding | -0.1052405 | 1.8415404 |
| PDHB | protein_coding | -0.1052824 | 1.66707825 |
| KIF13B | protein_coding | -0.1053139 | 1.10347641 |
| POLR2M | protein_coding | -0.1053357 | 0.98332297 |
| ACYP2 | protein_coding | -0.1053692 | 0.48250962 |
| MGA | protein_coding | -0.1053765 | 3.23357606 |
| MINDY4 | protein_coding | -0.1053864 | 0.12380969 |
| RHOB | protein_coding | -0.1053972 | 1.25207684 |
| CYCS | protein_coding | -0.10541 | 1.37131746 |
| LINC01968 | lncRNA | -0.1054555 | 0.15047498 |
| SAP30L-AS1 | lncRNA | -0.1054577 | 0.19502837 |
| MED21 | protein_coding | -0.105501 | 1.36254969 |
| ERC1 | protein_coding | -0.1055532 | 2.93047594 |
| SLC24A1 | protein_coding | -0.105566 | 0.6493216 |
| CCSER2 | protein_coding | -0.1055674 | 1.77646943 |
| MLXIPL | protein_coding | -0.1055683 | 1.43686399 |
| CASKIN2 | protein_coding | -0.1055807 | 1.82739936 |
| ZMYND10 | protein_coding | -0.1055938 | 0.21989054 |
| WDR4 | protein_coding | -0.1056452 | 1.52304737 |
| RAB11A | protein_coding | -0.1056515 | 2.85939133 |
| ZNF705E | protein_coding | -0.1056989 | 0.1259622 |
| FUCA2 | protein_coding | -0.1058328 | 0.42119327 |
| XRCC3 | protein_coding | -0.1059289 | 2.06229378 |
| TEX15 | protein_coding | -0.1064237 | 1.25771051 |
| COX6C | protein_coding | -0.1064377 | 2.43681353 |
| SLC22A18 | protein_coding | -0.106444 | 0.47725459 |
| DPYSL2 | protein_coding | -0.1064524 | 2.26440863 |
| DYNC1I2 | protein_coding | -0.1066208 | 2.11608883 |
| MIS12 | protein_coding | -0.1067378 | 1.47704158 |
| EXOSC2 | protein_coding | -0.1067529 | 2.32128214 |
| MAPK8IP3 | protein_coding | -0.1067967 | 2.34431051 |
| VWCE | protein_coding | -0.1068042 | 0.52059067 |
| ITPRIP | protein_coding | -0.1068417 | 1.21224056 |
| MIB2 | protein_coding | -0.1069862 | 0.52847096 |
| MIDN | protein_coding | -0.1070262 | 1.21530298 |
| SCAMP2 | protein_coding | -0.1070363 | 2.33359699 |

| | | | |
|---|---|---|---|
| SEC24C | protein_coding | -0.107127 | 2.56477668 |
| RUVBL1 | protein_coding | -0.1071471 | 2.84934336 |
| EIF3K | protein_coding | -0.1071797 | 3.97773153 |
| KHDRBS2 | protein_coding | -0.107202 | 1.16362654 |
| HCK | protein_coding | -0.1072476 | 0.11918503 |
| SERBP1 | protein_coding | -0.1072807 | 4.1916182 |
| SKP2 | protein_coding | -0.1076681 | 2.25057849 |
| PAPSS1 | protein_coding | -0.1079017 | 2.7922981 |
| SPINT1 | protein_coding | -0.1079045 | 1.66152354 |
| RAD51B | protein_coding | -0.1080028 | 0.42808588 |
| CNTN3 | protein_coding | -0.1080437 | 0.51492448 |
| PLCL2 | protein_coding | -0.1080672 | 1.51934259 |
| OSBPL10 | protein_coding | -0.1081126 | 1.52975313 |
| ACTR3 | protein_coding | -0.1081288 | 3.23955119 |
| ATXN7L3B | protein_coding | -0.1082547 | 4.28931784 |
| MAPRE1 | protein_coding | -0.1082686 | 3.93957933 |
| PATZ1 | protein_coding | -0.1083152 | 2.2712478 |
| CTR9 | protein_coding | -0.1083454 | 3.09818378 |
| JAK3 | protein_coding | -0.108427 | 0.12945442 |
| CACNB2 | protein_coding | -0.1084517 | 1.13796663 |
| SULT1A1 | protein_coding | -0.1085492 | 0.47834012 |
| SLC12A9 | protein_coding | -0.1086147 | 1.46079648 |
| ADIPOR2 | protein_coding | -0.1086486 | 3.22281246 |
| MYO5A | protein_coding | -0.1086643 | 2.73268093 |
| RAVER2 | protein_coding | -0.1086885 | 2.70902848 |
| AGPAT2 | protein_coding | -0.1087517 | 1.41844792 |
| POLR3GL | protein_coding | -0.1087708 | 0.75664342 |
| MANBAL | protein_coding | -0.1087708 | 1.86888266 |
| DNAJB12 | protein_coding | -0.1089854 | 1.05793023 |
| TNRC6B | protein_coding | -0.1090258 | 2.36896015 |
| RPL11 | protein_coding | -0.1090275 | 4.78104035 |
| SLC25A23 | protein_coding | -0.1091503 | 1.93150284 |
| ARRDC3-AS1 | lncRNA | -0.109338 | 0.2563424 |
| RP11-284F21 | lncRNA | -0.1094635 | 0.28257241 |
| SPATA5 | protein_coding | -0.1095182 | 0.87767086 |
| ZNF501 | protein_coding | -0.1095213 | 0.3885105 |
| ATP9A | protein_coding | -0.1096345 | 3.42817286 |
| COL5A2 | protein_coding | -0.1096838 | 0.25925003 |
| YWHAQ | protein_coding | -0.1097398 | 4.23741658 |
| PIAS1 | protein_coding | -0.1097978 | 2.09165393 |
| KMT5B | protein_coding | -0.1099355 | 1.76248198 |
| OTUD6B-AS1 | lncRNA | -0.1099494 | 1.2469556 |
| ALDH18A1 | protein_coding | -0.1100822 | 2.01419102 |
| GUCA1B | protein_coding | -0.1100984 | 0.29186467 |
| THAP7 | protein_coding | -0.1101606 | 1.49067911 |

| | | | |
|---|---|---|---|
| SGPP1 | protein_coding | -0.1101727 | 0.8409295 |
| OTUD7A | protein_coding | -0.110198 | 1.78981504 |
| CEP85 | protein_coding | -0.110231 | 3.51485436 |
| BDNF-AS | lncRNA | -0.1103097 | 0.29461306 |
| RAD51C | protein_coding | -0.110391 | 1.75398283 |
| CPSF6 | protein_coding | -0.1104222 | 2.89404254 |
| LL22NC03-2H | lncRNA | -0.1104232 | 0.18330055 |
| CCDC15-DT | lncRNA | -0.1104368 | 0.19046426 |
| FBXO46 | protein_coding | -0.1104535 | 1.07787101 |
| PDCD10 | protein_coding | -0.1104893 | 2.39177065 |
| DPH5 | protein_coding | -0.1105575 | 1.73852779 |
| NCOA5 | protein_coding | -0.1106257 | 2.67031833 |
| ILF3 | protein_coding | -0.1106984 | 4.83289504 |
| RP11-71L14.4 | lncRNA | -0.1108535 | 0.1000752 |
| CYRIB | protein_coding | -0.110869 | 3.07950983 |
| SKA3 | protein_coding | -0.1109441 | 1.3549337 |
| SMIM4 | protein_coding | -0.1109614 | 0.50899039 |
| KCTD5 | protein_coding | -0.1112587 | 2.15161957 |
| SHKBP1 | protein_coding | -0.1114039 | 2.54913967 |
| IFT80 | protein_coding | -0.1115522 | 0.69917941 |
| RP13-714J12 | lncRNA | -0.1115594 | 0.19643386 |
| QARS1 | protein_coding | -0.1115705 | 2.78816578 |
| NEK2 | protein_coding | -0.1115872 | 2.11000608 |
| RPUSD3 | protein_coding | -0.1116542 | 0.79939195 |
| PHF8 | protein_coding | -0.1117343 | 2.15696432 |
| MRPL51 | protein_coding | -0.1117746 | 3.11721828 |
| OTUD6B | protein_coding | -0.1119561 | 0.77242461 |
| ZNF326 | protein_coding | -0.1119583 | 2.85620262 |
| WARS2 | protein_coding | -0.111969 | 1.12550913 |
| CDC5L | protein_coding | -0.1120452 | 3.48359524 |
| RP11-395N3. | lncRNA | -0.112061 | 0.09047396 |
| EXD3 | protein_coding | -0.112071 | 0.77873043 |
| KCNA4 | protein_coding | -0.1120715 | 1.55540275 |
| TADA2A | protein_coding | -0.1120838 | 1.80213754 |
| RP11-342L8. | lncRNA | -0.112155 | 0.18282719 |
| DIPK1A | protein_coding | -0.1122823 | 0.18391896 |
| TMEM231 | protein_coding | -0.1122893 | 0.15468747 |
| DFFB | protein_coding | -0.1123281 | 0.76975478 |
| TAF6 | protein_coding | -0.1123842 | 2.81864843 |
| VAPB | protein_coding | -0.1124655 | 4.13724816 |
| EVA1C | protein_coding | -0.1124929 | 0.93634135 |
| TPRN | protein_coding | -0.1124995 | 1.20776236 |
| COL9A2 | protein_coding | -0.1125975 | 2.14838529 |
| SART1 | protein_coding | -0.1126332 | 1.430911 |
| PPCS | protein_coding | -0.1126366 | 1.2524359 |

| | | | |
|---|---|---|---|
| C19orf25 | protein_coding | -0.1127377 | 1.81260208 |
| IRX3 | protein_coding | -0.1127615 | 0.28268093 |
| RP11-54C4.3 | lncRNA | -0.1128922 | 0.27875415 |
| GOLPH3L | protein_coding | -0.112933 | 1.63278558 |
| SLC9B1 | protein_coding | -0.1129738 | 0.30155476 |
| DHX57 | protein_coding | -0.1130328 | 1.87175495 |
| GPSM1 | protein_coding | -0.1131001 | 1.43011879 |
| AF127936.9 | lncRNA | -0.113288 | 0.09937575 |
| BROX | protein_coding | -0.1133282 | 1.32811786 |
| PITHD1 | protein_coding | -0.113526 | 2.88887951 |
| CARMIL2 | protein_coding | -0.1135715 | 2.77371428 |
| RPL31 | protein_coding | -0.1135957 | 4.84881633 |
| COG3 | protein_coding | -0.1136041 | 1.51359703 |
| SEMA6A | protein_coding | -0.1136192 | 3.36582341 |
| RETSAT | protein_coding | -0.1136299 | 0.94394922 |
| RAD9B | protein_coding | -0.1137018 | 0.18033097 |
| OSBP2 | protein_coding | -0.1137111 | 1.24470531 |
| FPGS | protein_coding | -0.1137258 | 1.45217314 |
| LLPH | protein_coding | -0.1139789 | 1.26042779 |
| ZNF124 | protein_coding | -0.1140553 | 1.38021813 |
| ZNF319 | protein_coding | -0.1141598 | 0.73259335 |
| TFDP1 | protein_coding | -0.114226 | 4.68033243 |
| PPP1R2 | protein_coding | -0.1142495 | 1.77886168 |
| VAX2 | protein_coding | -0.1143052 | 0.5295792 |
| SPPL2B | protein_coding | -0.1144239 | 1.96506678 |
| ITPK1 | protein_coding | -0.1144713 | 2.50728599 |
| WDR18 | protein_coding | -0.1144784 | 1.43871504 |
| ATPAF2 | protein_coding | -0.114545 | 0.91711959 |
| WDR53 | protein_coding | -0.1145875 | 2.00613875 |
| RIMKLB | protein_coding | -0.1147051 | 0.99096724 |
| RPL8 | protein_coding | -0.1147772 | 3.86402422 |
| CCDC171 | protein_coding | -0.1148325 | 1.08751958 |
| MRPL36 | protein_coding | -0.114834 | 0.96071243 |
| C4orf19 | protein_coding | -0.1148549 | 1.78699761 |
| CABIN1 | protein_coding | -0.1148932 | 3.32244847 |
| NCOA3 | protein_coding | -0.1149104 | 2.27175496 |
| PEX6 | protein_coding | -0.114929 | 2.49766394 |
| RNF157 | protein_coding | -0.1149561 | 3.77209644 |
| CCDC154 | protein_coding | -0.1149573 | 0.30194985 |
| AP000442.1 | lncRNA | -0.1149689 | 0.1717398 |
| DHRS7B | protein_coding | -0.1150787 | 0.7241291 |
| HDLBP | protein_coding | -0.1151078 | 4.80032682 |
| MEA1 | protein_coding | -0.1151964 | 3.006101 |
| FAAP100 | protein_coding | -0.1152813 | 1.81241418 |
| CERCAM | protein_coding | -0.1152895 | 0.74683858 |

| | | | |
|---|---|---|---|
| KCTD2 | protein_coding | -0.1153167 | 2.65816257 |
| DRAXIN | protein_coding | -0.1153523 | 0.71514605 |
| PIGO | protein_coding | -0.1153565 | 1.66459395 |
| TAF11 | protein_coding | -0.1155576 | 2.36286848 |
| PARPBP | protein_coding | -0.1156357 | 1.3726852 |
| VEZF1 | protein_coding | -0.1156402 | 3.85950097 |
| ATRAID | protein_coding | -0.115656 | 2.50792538 |
| ZNF616 | protein_coding | -0.1156588 | 1.33652476 |
| IPO7 | protein_coding | -0.1156723 | 3.92800753 |
| DAP | protein_coding | -0.1158144 | 3.72003865 |
| ALDH3A1 | protein_coding | -0.1158346 | 0.24443243 |
| PRPF6 | protein_coding | -0.1159258 | 3.83319725 |
| COX5A | protein_coding | -0.1160443 | 2.26561808 |
| EFNB1 | protein_coding | -0.1161161 | 1.62148578 |
| CUL3 | protein_coding | -0.1161548 | 3.02270653 |
| GGCX | protein_coding | -0.1161801 | 1.8319521 |
| SFPQ | protein_coding | -0.1162105 | 3.70218769 |
| COX15 | protein_coding | -0.1162203 | 1.33300334 |
| SNRNP200 | protein_coding | -0.1162478 | 4.97097469 |
| ZC3H18 | protein_coding | -0.1162589 | 3.17159229 |
| SLC39A13 | protein_coding | -0.1162654 | 1.4624401 |
| LSM3 | protein_coding | -0.1162859 | 1.71724516 |
| ATP8A2 | protein_coding | -0.1163037 | 3.55394208 |
| RP11-161I6.2 | lncRNA | -0.1163516 | 0.49660077 |
| MRFAP1L1 | protein_coding | -0.116411 | 3.1088968 |
| GBA | protein_coding | -0.1164383 | 1.47537929 |
| NAPG | protein_coding | -0.116622 | 1.14935733 |
| GNB1 | protein_coding | -0.1166359 | 4.94376772 |
| RP11-685N1( | lncRNA | -0.1166441 | 0.14269649 |
| HSPBP1 | protein_coding | -0.116676 | 1.35316114 |
| SMAD4 | protein_coding | -0.1166987 | 2.09657961 |
| KLC4 | protein_coding | -0.1167315 | 1.47686952 |
| AP1S2 | protein_coding | -0.1168372 | 1.2842174 |
| EIF3D | protein_coding | -0.1169506 | 4.39292931 |
| CZIB | protein_coding | -0.1170336 | 2.03226602 |
| SOCS6 | protein_coding | -0.117039 | 0.83320123 |
| RMDN1 | protein_coding | -0.1171095 | 2.13483403 |
| ACTL6A | protein_coding | -0.1171546 | 4.31000731 |
| CBFA2T2 | protein_coding | -0.1171729 | 4.37878111 |
| MOB4 | protein_coding | -0.117387 | 0.88555304 |
| TGFBRAP1 | protein_coding | -0.1174702 | 2.61452812 |
| PPFIBP1 | protein_coding | -0.1175174 | 2.20096764 |
| ATP2B1-AS1 | lncRNA | -0.1175935 | 0.21658187 |
| LPCAT4 | protein_coding | -0.1176575 | 1.5242457 |
| IGBP1 | protein_coding | -0.1177006 | 2.01155276 |

| | | | |
|---|---|---|---|
| CTA-292E10. | lncRNA | -0.1177071 | 0.41509759 |
| LONRF1 | protein_coding | -0.1179475 | 1.18934985 |
| MRPL20 | protein_coding | -0.1179523 | 2.28579716 |
| MED9 | protein_coding | -0.1179563 | 1.42978102 |
| PEX14 | protein_coding | -0.1179689 | 1.22608105 |
| VPS51 | protein_coding | -0.1180126 | 1.39211704 |
| TMEM222 | protein_coding | -0.1180779 | 1.92330614 |
| ZFP30 | protein_coding | -0.1182095 | 2.03247291 |
| GLT8D1 | protein_coding | -0.1182352 | 1.67830533 |
| DEPDC1B | protein_coding | -0.1182929 | 1.58849525 |
| AGAP3 | protein_coding | -0.1183272 | 2.85443922 |
| TMED8 | protein_coding | -0.1183689 | 3.51551997 |
| LMNB2 | protein_coding | -0.1183808 | 2.78880271 |
| BCL2 | protein_coding | -0.1184044 | 4.0341815 |
| PIGH | protein_coding | -0.1187733 | 1.62258063 |
| CTD-2012K1 | lncRNA | -0.1187925 | 0.09973853 |
| MAP2K5-DT | lncRNA | -0.1188501 | 0.31033196 |
| DYM | protein_coding | -0.1190302 | 3.35137771 |
| CEP152 | protein_coding | -0.1190387 | 2.01069878 |
| RP11-394I13 | lncRNA | -0.1193403 | 0.13266124 |
| PHB | protein_coding | -0.1193996 | 3.66216512 |
| MAML1 | protein_coding | -0.119404 | 2.62251676 |
| DAXX | protein_coding | -0.1194056 | 2.99331191 |
| RPP30 | protein_coding | -0.1194306 | 1.32022638 |
| LIN28B-AS1 | lncRNA | -0.1194392 | 0.3854695 |
| MYH10 | protein_coding | -0.1194699 | 1.75274668 |
| CDR2L | protein_coding | -0.1194725 | 0.91108377 |
| GRM8 | protein_coding | -0.1195789 | 2.00698243 |
| COX19 | protein_coding | -0.1196304 | 1.56071141 |
| TBC1D7 | protein_coding | -0.1198225 | 1.96442714 |
| OTUD5 | protein_coding | -0.1198458 | 2.29830783 |
| CLPP | protein_coding | -0.1199277 | 1.51783539 |
| AC092535.3 | lncRNA | -0.1199547 | 0.11883315 |
| MCRS1 | protein_coding | -0.1199805 | 2.84406284 |
| KPNA2 | protein_coding | -0.1200264 | 4.38327136 |
| JADE1 | protein_coding | -0.1200271 | 2.99125242 |
| CDC42 | protein_coding | -0.1200814 | 3.40509361 |
| RPL39 | protein_coding | -0.1201765 | 2.64449723 |
| TNK2 | protein_coding | -0.1201885 | 1.43944529 |
| RP11-660L1 | lncRNA | -0.1202434 | 0.48031334 |
| ZNF473 | protein_coding | -0.1203738 | 2.22339943 |
| RP11-356B1 | lncRNA | -0.1203933 | 0.13012796 |
| SMC5-DT | lncRNA | -0.1204337 | 0.42627797 |
| EZR | protein_coding | -0.1204467 | 3.7559759 |
| DOCK7 | protein_coding | -0.1204704 | 1.52990146 |

| | | | |
|---|---|---|---|
| PRPF4 | protein_coding | -0.1206209 | 3.24439704 |
| PEX5 | protein_coding | -0.1206214 | 2.35047683 |
| STRAP | protein_coding | -0.1206383 | 3.61194227 |
| DYNLT2B | protein_coding | -0.1206465 | 0.38965116 |
| PTPRA | protein_coding | -0.1207209 | 3.63767603 |
| ZER1 | protein_coding | -0.1207528 | 2.03949109 |
| DNLZ | protein_coding | -0.1207824 | 0.11480893 |
| MGMT | protein_coding | -0.1207865 | 1.13426851 |
| ORMDL2 | protein_coding | -0.1207889 | 1.68175367 |
| SRSF8 | protein_coding | -0.1209181 | 1.87937352 |
| MIR17HG | lncRNA | -0.1209943 | 0.39217973 |
| ZNF785 | protein_coding | -0.1210002 | 1.59054513 |
| ZBTB3 | protein_coding | -0.1210034 | 0.8964217 |
| KCTD6 | protein_coding | -0.1210079 | 0.86042765 |
| VDAC1 | protein_coding | -0.1210681 | 4.33322158 |
| C9orf72 | protein_coding | -0.1211015 | 1.64822165 |
| TCFL5 | protein_coding | -0.1211228 | 1.66023414 |
| RAP2A | protein_coding | -0.1212536 | 1.7486617 |
| CCSAP | protein_coding | -0.1213091 | 2.19161133 |
| IFT140 | protein_coding | -0.1213376 | 1.41014803 |
| MACO1 | protein_coding | -0.1213831 | 3.01064861 |
| CIRBP | protein_coding | -0.1216002 | 4.45542614 |
| TEPSIN | protein_coding | -0.1216338 | 1.49124904 |
| HSD17B7 | protein_coding | -0.1217253 | 1.21857642 |
| ANAPC1P2 | lncRNA | -0.1217487 | 0.12129454 |
| PPM1B | protein_coding | -0.1218863 | 1.17938643 |
| SELENOS | protein_coding | -0.1219132 | 2.4242749 |
| STX2 | protein_coding | -0.1219871 | 2.52085488 |
| ZFAND2B | protein_coding | -0.1220152 | 1.26292128 |
| PANO1 | protein_coding | -0.122071 | 0.35734505 |
| NUTM2B-AS | lncRNA | -0.1221889 | 0.8276923 |
| CACYBP | protein_coding | -0.1221952 | 3.51515958 |
| B3GNTL1 | protein_coding | -0.1222102 | 0.30633509 |
| TMEM209 | protein_coding | -0.1222684 | 3.07364905 |
| DDIT4 | protein_coding | -0.1222877 | 2.8866517 |
| PRKCZ | protein_coding | -0.1223468 | 1.07401691 |
| ARV1 | protein_coding | -0.1223542 | 1.92878446 |
| OSBPL2 | protein_coding | -0.1225125 | 2.23669207 |
| CCDC86 | protein_coding | -0.1225154 | 2.54183554 |
| SCARB1 | protein_coding | -0.1225258 | 2.30580163 |
| SDHAF2 | protein_coding | -0.1225342 | 0.81294393 |
| FBXO28 | protein_coding | -0.1225727 | 1.97894111 |
| CTD-2201E18 | lncRNA | -0.12261 | 1.34906201 |
| CEP192 | protein_coding | -0.1226323 | 2.86480427 |
| LAPTM4A | protein_coding | -0.1226825 | 3.25365348 |

| | | | |
|---|---|---|---|
| UCK1 | protein_coding | -0.1227095 | 2.88231178 |
| UBA1 | protein_coding | -0.1227124 | 5.37938466 |
| ZNF445 | protein_coding | -0.1227278 | 2.48726425 |
| MAP3K14-AS | lncRNA | -0.1228054 | 0.6013171 |
| NBEAL2 | protein_coding | -0.1228279 | 2.3351849 |
| DPF1 | protein_coding | -0.122869 | 1.05034759 |
| PDE7A | protein_coding | -0.1229226 | 1.95174607 |
| LAS1L | protein_coding | -0.1229282 | 3.15798268 |
| AURKB | protein_coding | -0.1230573 | 1.59095502 |
| MRPL57 | protein_coding | -0.1231494 | 1.14171533 |
| WDCP | protein_coding | -0.1231499 | 2.29838451 |
| HSPA8 | protein_coding | -0.1231751 | 6.02714213 |
| LIPT2 | protein_coding | -0.1232763 | 0.47300099 |
| ANKRD13D | protein_coding | -0.1232966 | 1.71238361 |
| VPS13A | protein_coding | -0.1233124 | 4.26943961 |
| RP11-1277A3 | lncRNA | -0.1233852 | 0.31058753 |
| RASA1 | protein_coding | -0.1234109 | 1.46316342 |
| EMD | protein_coding | -0.1236007 | 2.0254951 |
| CEP170B | protein_coding | -0.1236839 | 1.63403688 |
| VPS13B-DT | lncRNA | -0.1237487 | 0.35066809 |
| RBM8A | protein_coding | -0.1237805 | 4.14984536 |
| MESD | protein_coding | -0.1238495 | 3.30836939 |
| U2AF2 | protein_coding | -0.1239158 | 4.00793946 |
| FAM174C | protein_coding | -0.1239911 | 1.49293968 |
| AC084219.4 | lncRNA | -0.1243786 | 1.0948298 |
| KANK3 | protein_coding | -0.1244167 | 0.20072372 |
| PRSS22 | protein_coding | -0.1244237 | 0.14276625 |
| ETAA1 | protein_coding | -0.1244923 | 1.59810149 |
| ZNRD2 | protein_coding | -0.1244947 | 1.30903025 |
| MBNL1 | protein_coding | -0.1245127 | 2.0193779 |
| SLC8A2 | protein_coding | -0.124513 | 1.53417224 |
| RRAGA | protein_coding | -0.1246318 | 2.88914162 |
| EIF2B3 | protein_coding | -0.1246748 | 2.35635528 |
| SH2D2A | protein_coding | -0.1246796 | 0.90533456 |
| CCN1 | protein_coding | -0.1247095 | 0.3587033 |
| AGO2 | protein_coding | -0.1247187 | 4.43856742 |
| SCAI | protein_coding | -0.1247579 | 1.62767604 |
| POLR2G | protein_coding | -0.1248223 | 3.18567621 |
| TTC21A | protein_coding | -0.124984 | 0.67291463 |
| LARP4B | protein_coding | -0.1250039 | 2.67329022 |
| RAD21 | protein_coding | -0.1251477 | 4.89566188 |
| PPP6R3 | protein_coding | -0.1251832 | 3.30188005 |
| ZNF428 | protein_coding | -0.1251849 | 1.24949975 |
| MFAP1 | protein_coding | -0.1251895 | 2.81760856 |
| KLF7-IT1 | lncRNA | -0.125273 | 0.15365586 |

| | | | |
|---|---|---|---|
| TMEM250 | protein_coding | -0.1253212 | 1.59030398 |
| EEF1AKMT1 | protein_coding | -0.1253684 | 0.35540098 |
| ACLY | protein_coding | -0.1254415 | 5.24375628 |
| CAMSAP1 | protein_coding | -0.1255261 | 3.79248776 |
| ALMS1 | protein_coding | -0.125551 | 3.41126653 |
| TENM3-AS1 | lncRNA | -0.1255864 | 1.11410749 |
| RBX1 | protein_coding | -0.1257069 | 2.89409952 |
| SLC46A3 | protein_coding | -0.1257077 | 1.77060708 |
| SMARCC1 | protein_coding | -0.1259692 | 3.95979666 |
| BCAS3 | protein_coding | -0.1259865 | 1.25467524 |
| PCYT2 | protein_coding | -0.1260628 | 1.9012156 |
| NELFA | protein_coding | -0.1261027 | 1.5465659 |
| ARL16 | protein_coding | -0.1261061 | 2.17947895 |
| URB1 | protein_coding | -0.1261996 | 3.63302021 |
| GSKIP | protein_coding | -0.1262877 | 1.77279206 |
| MT-ND2 | protein_coding | -0.126405 | 3.4121257 |
| ACTR2 | protein_coding | -0.1264127 | 3.82535624 |
| ZNF841 | protein_coding | -0.1266959 | 0.48291026 |
| BICRA | protein_coding | -0.126769 | 1.11915072 |
| ZNF550 | protein_coding | -0.1268367 | 1.99718313 |
| CACTIN | protein_coding | -0.1269356 | 1.26456559 |
| BOP1 | protein_coding | -0.1271175 | 2.43836301 |
| CBX1 | protein_coding | -0.1272181 | 5.0632904 |
| HACE1 | protein_coding | -0.1272399 | 1.7926623 |
| ECM1 | protein_coding | -0.1272603 | 0.60400737 |
| EIF3J-DT | lncRNA | -0.1274662 | 0.75933919 |
| SEC14L1 | protein_coding | -0.1275404 | 3.82745035 |
| XPO7 | protein_coding | -0.1275443 | 3.71434585 |
| METTL8 | protein_coding | -0.1275477 | 1.85651378 |
| PIGT | protein_coding | -0.1275732 | 2.80073198 |
| SLC6A3 | protein_coding | -0.1275892 | 1.14023522 |
| WDR6 | protein_coding | -0.1276226 | 4.21063907 |
| SIRT3 | protein_coding | -0.1276288 | 1.270546 |
| MRPL17 | protein_coding | -0.1276717 | 3.0468742 |
| GEMIN5 | protein_coding | -0.1277155 | 3.02500546 |
| ZNF525 | protein_coding | -0.1277942 | 1.96692992 |
| YDJC | protein_coding | -0.1278154 | 1.00395048 |
| RRAS2 | protein_coding | -0.1278822 | 0.93378171 |
| FBXO21 | protein_coding | -0.1279601 | 3.41919621 |
| ALKBH4 | protein_coding | -0.1279879 | 1.99485478 |
| NUDT5 | protein_coding | -0.1280472 | 2.96260076 |
| SNX22 | protein_coding | -0.1281423 | 1.40647064 |
| CKAP2 | protein_coding | -0.1281995 | 2.52669327 |
| CDK14 | protein_coding | -0.1282146 | 2.03133309 |
| MAP2K4 | protein_coding | -0.1283613 | 1.6161997 |

| | | | |
|---|---|---|---|
| SENP8 | protein_coding | -0.1284053 | 0.71893034 |
| RPS15 | protein_coding | -0.1284127 | 3.25289701 |
| FBXW11 | protein_coding | -0.1285104 | 2.8111169 |
| DHRS12 | protein_coding | -0.1285262 | 0.34088954 |
| CNOT7 | protein_coding | -0.1285765 | 2.39054025 |
| SNHG3 | lncRNA | -0.1286092 | 1.26190078 |
| LCORL | protein_coding | -0.128623 | 2.60194877 |
| RNF112 | protein_coding | -0.1287632 | 0.18945761 |
| SZRD1 | protein_coding | -0.1288441 | 4.12076795 |
| TSPAN31 | protein_coding | -0.1288678 | 1.04973505 |
| TRIAP1 | protein_coding | -0.1288954 | 1.80788108 |
| EIF6 | protein_coding | -0.1289267 | 2.71039775 |
| SPOCK1 | protein_coding | -0.129036 | 2.0112501 |
| ESYT3 | protein_coding | -0.1290469 | 0.39106656 |
| RP11-73M18 | lncRNA | -0.1290477 | 0.82437239 |
| NOVA1 | protein_coding | -0.129051 | 2.33591773 |
| ZNF157 | protein_coding | -0.1291148 | 0.54830802 |
| PIGW | protein_coding | -0.1293946 | 2.26966838 |
| RAN | protein_coding | -0.1295553 | 5.64116705 |
| SNX18 | protein_coding | -0.1295875 | 3.13163454 |
| RP11-545E17 | lncRNA | -0.1295938 | 0.49946391 |
| ZNF793-AS1 | lncRNA | -0.1296169 | 0.21469818 |
| CCM2 | protein_coding | -0.1297456 | 2.69460619 |
| NCBP2 | protein_coding | -0.1297978 | 4.45665031 |
| RELCH | protein_coding | -0.1298426 | 3.30105803 |
| METTL7A | protein_coding | -0.1298611 | 2.38452203 |
| INTS5 | protein_coding | -0.1299368 | 2.92616487 |
| MGAT3 | protein_coding | -0.1299401 | 1.88741915 |
| ERMP1 | protein_coding | -0.1299667 | 2.23710833 |
| FH | protein_coding | -0.12997 | 3.24949233 |
| MRPS15 | protein_coding | -0.1300366 | 3.25782571 |
| ABHD5 | protein_coding | -0.1300963 | 1.61318754 |
| ADPRS | protein_coding | -0.1301202 | 2.01358083 |
| ARHGAP19-S | protein_coding | -0.1301208 | 0.19849926 |
| FAM111A | protein_coding | -0.1301367 | 2.41053561 |
| RP11-544I20 | lncRNA | -0.1302001 | 0.24678837 |
| C16orf71 | protein_coding | -0.1302032 | 0.25303273 |
| FAM193B | protein_coding | -0.1302255 | 1.72586098 |
| NRM | protein_coding | -0.1302497 | 2.71829653 |
| YWHAZ | protein_coding | -0.1303278 | 3.9391138 |
| ARPC3 | protein_coding | -0.1303661 | 3.65685155 |
| PLEKHA6 | protein_coding | -0.1303742 | 5.28099784 |
| GPD1L | protein_coding | -0.1305812 | 2.0523045 |
| RP11-365P13 | lncRNA | -0.1306782 | 0.1630948 |
| KLHL11 | protein_coding | -0.1307303 | 1.3052739 |

| | | | |
|---|---|---|---|
| HSD11B1L | protein_coding | -0.1307685 | 0.78873542 |
| TRAPPC2B | protein_coding | -0.1308117 | 0.59842061 |
| MARVELD2 | protein_coding | -0.1308509 | 0.85867096 |
| BTBD7 | protein_coding | -0.1309039 | 3.29397365 |
| OSBP | protein_coding | -0.1309207 | 4.45870879 |
| LMO7 | protein_coding | -0.1309709 | 1.16342508 |
| RP11-74E22. | lncRNA | -0.1309963 | 0.42092868 |
| ZNF562 | protein_coding | -0.1310118 | 1.56174696 |
| NRF1 | protein_coding | -0.1311956 | 1.76818851 |
| ERCC1 | protein_coding | -0.1312643 | 1.80316412 |
| RABL2A | protein_coding | -0.1312745 | 1.05839092 |
| OSTC | protein_coding | -0.1313053 | 2.33458938 |
| ALYREF | protein_coding | -0.1314031 | 3.4752777 |
| TYW1 | protein_coding | -0.1314094 | 2.13009162 |
| SF3B4 | protein_coding | -0.1314964 | 3.5010705 |
| RP13-1032I1 | lncRNA | -0.1315127 | 0.25641412 |
| RBAK | protein_coding | -0.1315377 | 1.86864403 |
| SEPTIN7-DT | lncRNA | -0.1315377 | 0.43210861 |
| ZYG11B | protein_coding | -0.1315437 | 2.56202301 |
| TTF1 | protein_coding | -0.1315653 | 2.68505073 |
| AC113189.5 | lncRNA | -0.1316093 | 1.38751141 |
| LCMT1 | protein_coding | -0.1316274 | 2.54611933 |
| CCDC168 | protein_coding | -0.1316709 | 0.35843314 |
| YY1 | protein_coding | -0.1316965 | 4.48371192 |
| ASB16-AS1 | lncRNA | -0.1317658 | 1.11914989 |
| UQCC3 | protein_coding | -0.1318066 | 1.35150698 |
| NPAT | protein_coding | -0.1318914 | 1.82879865 |
| KDM1A | protein_coding | -0.1319868 | 4.96779971 |
| PRKACB | protein_coding | -0.132012 | 1.48110467 |
| IRS2 | protein_coding | -0.1320183 | 0.97749853 |
| HNRNPA2B1 | protein_coding | -0.1320334 | 6.57640804 |
| UQCRC1 | protein_coding | -0.1321886 | 3.35810004 |
| KANSL3 | protein_coding | -0.132257 | 3.78710987 |
| YEATS4 | protein_coding | -0.1322635 | 2.19880284 |
| NOP14-AS1 | lncRNA | -0.1322796 | 0.82880049 |
| RWDD4 | protein_coding | -0.1323668 | 1.50828965 |
| RCE1 | protein_coding | -0.1324036 | 2.18668433 |
| URB2 | protein_coding | -0.1324708 | 3.26391398 |
| ABT1 | protein_coding | -0.1325173 | 1.71756199 |
| SSC5D | protein_coding | -0.1325254 | 0.38480603 |
| CALHM2 | protein_coding | -0.1325382 | 0.48418904 |
| MYBL1 | protein_coding | -0.1326097 | 0.53791612 |
| PIGA | protein_coding | -0.1326277 | 1.60784575 |
| WTIP | protein_coding | -0.1326414 | 0.27995959 |
| PCBP1 | protein_coding | -0.1326804 | 4.73244904 |

| | | | |
|---|---|---|---|
| ZBTB44 | protein_coding | -0.1327267 | 3.35251082 |
| ACER3 | protein_coding | -0.1327413 | 1.53724204 |
| DHODH | protein_coding | -0.1327607 | 1.58180562 |
| RP4-668E10. | lncRNA | -0.1328689 | 4.17742442 |
| ZNF629 | protein_coding | -0.1328792 | 2.51171552 |
| DHX8 | protein_coding | -0.1329947 | 3.36333246 |
| NCAPH2 | protein_coding | -0.1330148 | 4.38988786 |
| PPM1G | protein_coding | -0.1330525 | 5.06878185 |
| BEX2 | protein_coding | -0.1332249 | 3.05676707 |
| FOXRED2 | protein_coding | -0.1334785 | 3.53792564 |
| RFX1 | protein_coding | -0.1335005 | 0.81285908 |
| LBR | protein_coding | -0.1337712 | 2.81761505 |
| NRDE2 | protein_coding | -0.133788 | 2.10826025 |
| SLU7 | protein_coding | -0.1338429 | 2.81588474 |
| GNA11 | protein_coding | -0.1338907 | 2.73203708 |
| CHD1L | protein_coding | -0.1338943 | 0.17824584 |
| POLD2 | protein_coding | -0.1339291 | 4.64356959 |
| ACP1 | protein_coding | -0.1339976 | 4.47928574 |
| CACNB1 | protein_coding | -0.1340001 | 1.01224155 |
| MAPK9 | protein_coding | -0.1340127 | 2.22660091 |
| TAFA4 | protein_coding | -0.1340343 | 0.19062923 |
| PARVB | protein_coding | -0.1342558 | 0.2125825 |
| PAQR4 | protein_coding | -0.1342737 | 3.94856216 |
| LRRC36 | protein_coding | -0.1343032 | 0.38399237 |
| PTPA | protein_coding | -0.1343139 | 4.06119925 |
| RAB1A | protein_coding | -0.1343141 | 3.45899646 |
| ATP13A3 | protein_coding | -0.1344306 | 3.83534175 |
| CTD-2206N4. | lncRNA | -0.1344844 | 0.12235365 |
| ITCH | protein_coding | -0.1347411 | 3.32789817 |
| ZNF248 | protein_coding | -0.1347434 | 1.85057106 |
| FMNL1 | protein_coding | -0.1347979 | 0.9534107 |
| SLC25A29 | protein_coding | -0.1348393 | 3.49180292 |
| ZNF611 | protein_coding | -0.1348615 | 1.69970483 |
| MTA3 | protein_coding | -0.1348903 | 2.86798239 |
| HNRNPF | protein_coding | -0.1349739 | 4.76329765 |
| TRIM11 | protein_coding | -0.1350877 | 1.49015002 |
| RP11-683L23 | lncRNA | -0.1350889 | 0.1724831 |
| RPS23 | protein_coding | -0.1350993 | 5.82046329 |
| KCNQ5 | protein_coding | -0.135214 | 2.19536047 |
| DDI2 | protein_coding | -0.1352716 | 3.39605085 |
| SCAF4 | protein_coding | -0.1353091 | 3.58440751 |
| ZNF701 | protein_coding | -0.135421 | 1.40046831 |
| SAMD8 | protein_coding | -0.1355278 | 1.57468845 |
| MRPS18A | protein_coding | -0.1358013 | 1.73653429 |
| RPS17 | protein_coding | -0.1358855 | 1.00851945 |

| | | | |
|---|---|---|---|
| HIPK1 | protein_coding | -0.135886 | 3.6273198 |
| GK5 | protein_coding | -0.135907 | 2.41948109 |
| GPHN | protein_coding | -0.1359513 | 2.78953861 |
| INTS2 | protein_coding | -0.1359697 | 2.42301506 |
| HDHD5 | protein_coding | -0.1360628 | 3.29049541 |
| ZDHHC5 | protein_coding | -0.1360672 | 4.2637625 |
| RP11-471B2: | lncRNA | -0.1360928 | 0.20105001 |
| UBE2I | protein_coding | -0.1361203 | 4.54066761 |
| RPL36AL | protein_coding | -0.1361262 | 3.69540527 |
| GPR153 | protein_coding | -0.1361862 | 2.14485247 |
| VPS25 | protein_coding | -0.1362672 | 3.41813592 |
| RAB8A | protein_coding | -0.136286 | 2.99845858 |
| MROH1 | protein_coding | -0.1364304 | 1.83939401 |
| SLC35B2 | protein_coding | -0.1364554 | 2.87843472 |
| JMJD8 | protein_coding | -0.1364931 | 2.29696408 |
| MRPS23 | protein_coding | -0.1365775 | 3.22263138 |
| ZBTB33 | protein_coding | -0.1366384 | 2.24607359 |
| FAM20B | protein_coding | -0.1366409 | 4.04496043 |
| MPHOSPH9 | protein_coding | -0.1366705 | 3.44974538 |
| DVL2 | protein_coding | -0.1367019 | 2.65053177 |
| QRICH1 | protein_coding | -0.1367365 | 3.26142369 |
| RARA | protein_coding | -0.1367967 | 2.51046175 |
| MRPL2 | protein_coding | -0.1368053 | 0.59911046 |
| ZNHIT1 | protein_coding | -0.1368415 | 2.86476281 |
| SAE1 | protein_coding | -0.1369472 | 4.93693692 |
| TMPO | protein_coding | -0.1370558 | 4.57334242 |
| TUBGCP4 | protein_coding | -0.1370639 | 3.57108148 |
| C19orf54 | protein_coding | -0.137113 | 2.91380446 |
| ZNF213-AS1 | lncRNA | -0.1371191 | 1.51404769 |
| AKAP10 | protein_coding | -0.1371544 | 1.72331399 |
| RPL24 | protein_coding | -0.1372048 | 5.66593781 |
| DECR1 | protein_coding | -0.1373039 | 0.50496031 |
| SMIM13 | protein_coding | -0.1373051 | 2.80101162 |
| SEC62 | protein_coding | -0.1373186 | 3.37228908 |
| FBXO45 | protein_coding | -0.1373627 | 2.58165089 |
| AASDHPPT | protein_coding | -0.1373979 | 2.53887048 |
| LINC00667 | lncRNA | -0.137508 | 2.84581817 |
| TXNDC17 | protein_coding | -0.1375243 | 1.51475547 |
| SLX4IP | protein_coding | -0.1376132 | 1.88716648 |
| LAMTOR2 | protein_coding | -0.1376877 | 1.59342169 |
| GUCD1 | protein_coding | -0.137771 | 3.87814986 |
| RP5-940J5.6 | lncRNA | -0.1378084 | 0.1316251 |
| ANKRD54 | protein_coding | -0.1378298 | 2.50057878 |
| GOLGA8N | protein_coding | -0.1382395 | 0.32029246 |
| NEUROD2 | protein_coding | -0.1383995 | 1.89745698 |

| | | | |
|---|---|---|---|
| TUSC2 | protein_coding | -0.1384353 | 1.38007082 |
| CYFIP2 | protein_coding | -0.1387047 | 5.40848405 |
| ABR | protein_coding | -0.1387312 | 2.9232078 |
| EMC3 | protein_coding | -0.138742 | 1.98938813 |
| MST1 | protein_coding | -0.1387798 | 0.27124751 |
| SMC3 | protein_coding | -0.1388623 | 4.2042867 |
| CHMP4B | protein_coding | -0.1388681 | 2.6006658 |
| TCEAL4 | protein_coding | -0.1389554 | 2.46986507 |
| NAALADL2 | protein_coding | -0.1390304 | 0.27258901 |
| PUF60 | protein_coding | -0.1390374 | 3.97273913 |
| CCDC107 | protein_coding | -0.1392148 | 0.9185142 |
| GOLM2 | protein_coding | -0.139246 | 2.51043264 |
| SPAG5 | protein_coding | -0.1392657 | 3.84296336 |
| SLC4A2 | protein_coding | -0.1393863 | 3.1109232 |
| DRD4 | protein_coding | -0.1395183 | 0.39970944 |
| RP11-295P9. | lncRNA | -0.1395434 | 0.85650146 |
| RAB26 | protein_coding | -0.1396003 | 1.60767139 |
| FAM166A | protein_coding | -0.1397177 | 0.25525188 |
| SIK1B | protein_coding | -0.1397421 | 0.39972322 |
| RP11-934B9. | lncRNA | -0.1398001 | 0.35811714 |
| BANF1 | protein_coding | -0.1398956 | 4.26355883 |
| NUTF2 | protein_coding | -0.1400349 | 3.44750728 |
| CD2AP | protein_coding | -0.1400466 | 2.62842948 |
| CLCC1 | protein_coding | -0.140084 | 3.34173609 |
| ADD1 | protein_coding | -0.140121 | 4.42981489 |
| RP11-426D1 | lncRNA | -0.1401871 | 0.26680177 |
| MKRN1 | protein_coding | -0.1402825 | 3.57208352 |
| VPS26B | protein_coding | -0.1403119 | 3.54212508 |
| PRORP | protein_coding | -0.1403699 | 0.79312324 |
| IARS2 | protein_coding | -0.1403817 | 4.25537143 |
| FBXL19-AS1 | lncRNA | -0.1404651 | 0.507183 |
| EYA3 | protein_coding | -0.1405347 | 3.58264117 |
| DLL1 | protein_coding | -0.1406653 | 0.24775162 |
| PINLYP | protein_coding | -0.1407458 | 1.75735094 |
| POLR3H | protein_coding | -0.1407654 | 3.43844144 |
| SLC4A7 | protein_coding | -0.1407719 | 2.10756851 |
| ZNF480 | protein_coding | -0.1408667 | 2.92477258 |
| DYNLL2 | protein_coding | -0.1410071 | 5.66864393 |
| SHQ1 | protein_coding | -0.1410729 | 1.55858344 |
| RPUSD1 | protein_coding | -0.1410785 | 1.65701036 |
| RP11-57A19. | lncRNA | -0.1411209 | 1.0776609 |
| SERF2 | protein_coding | -0.1411358 | 5.29337771 |
| SOCS5 | protein_coding | -0.1411478 | 1.81087555 |
| RPL27 | protein_coding | -0.1411488 | 5.68164145 |
| MACROH2A1 | protein_coding | -0.1412104 | 6.20042353 |

| | | | |
|---|---|---|---|
| LSM7 | protein_coding | -0.141212 | 2.03345087 |
| ZNF345 | protein_coding | -0.1412266 | 1.16552514 |
| CTC-329D1.2 | lncRNA | -0.1412832 | 0.36017476 |
| RITA1 | protein_coding | -0.1415092 | 2.49010663 |
| CENPJ | protein_coding | -0.1416189 | 3.1362753 |
| RP3-475N16. | lncRNA | -0.141631 | 0.28656388 |
| MSL1 | protein_coding | -0.1416896 | 4.5882115 |
| AP000662.4 | lncRNA | -0.1416936 | 0.24156213 |
| PTPN12 | protein_coding | -0.1417153 | 2.41154276 |
| PIP4K2B | protein_coding | -0.1417591 | 4.79669149 |
| LINC00997 | lncRNA | -0.1417667 | 0.86984297 |
| EML5 | protein_coding | -0.141895 | 4.06775973 |
| ELAVL1 | protein_coding | -0.1420267 | 5.10656233 |
| WDR33 | protein_coding | -0.1421087 | 3.93498817 |
| PSD3 | protein_coding | -0.1422383 | 2.91381831 |
| AK1 | protein_coding | -0.1422678 | 0.75525306 |
| ZNF275 | protein_coding | -0.1422705 | 2.44496561 |
| ZNF555 | protein_coding | -0.1422716 | 1.19228937 |
| TMTC4 | protein_coding | -0.1424286 | 2.45595591 |
| ADIRF-AS1 | lncRNA | -0.1425092 | 1.40317731 |
| RPS19 | protein_coding | -0.1425784 | 5.75822622 |
| TMEM214 | protein_coding | -0.1425952 | 3.77598478 |
| LRRC74B | protein_coding | -0.1426283 | 0.14935812 |
| POLR2J | protein_coding | -0.1427453 | 1.79619407 |
| CLEC4F | protein_coding | -0.1428105 | 0.17677591 |
| SEPSECS | protein_coding | -0.142901 | 1.95231554 |
| CEP76 | protein_coding | -0.1429919 | 2.29571731 |
| SIN3A | protein_coding | -0.1430968 | 4.96973981 |
| ZMYND8 | protein_coding | -0.1431512 | 3.84526424 |
| PRKCB | protein_coding | -0.143165 | 0.92567329 |
| PEX1 | protein_coding | -0.143226 | 1.58728862 |
| OXCT1 | protein_coding | -0.1433111 | 5.53667801 |
| CAT | protein_coding | -0.1433243 | 1.95958621 |
| GEMIN6 | protein_coding | -0.1433635 | 1.42924066 |
| RP13-467H1 | lncRNA | -0.1433803 | 0.19028972 |
| KIAA0895L | protein_coding | -0.1433816 | 2.81684247 |
| ELMO2 | protein_coding | -0.1433932 | 3.80009007 |
| ELP5 | protein_coding | -0.1433997 | 2.43915038 |
| RP4-545L17. | lncRNA | -0.1435013 | 0.1313441 |
| POMGNT1 | protein_coding | -0.1436077 | 3.8983125 |
| SNAPIN | protein_coding | -0.1437452 | 2.12821031 |
| ANKMY2 | protein_coding | -0.1437747 | 1.39140962 |
| CHUK | protein_coding | -0.1438642 | 2.70381538 |
| TEX45 | protein_coding | -0.1439026 | 0.28677337 |
| PYGM | protein_coding | -0.1439322 | 0.21203478 |

| | | | |
|---|---|---|---|
| POP1 | protein_coding | -0.1439505 | 2.3308885 |
| DNAJC8 | protein_coding | -0.1439761 | 4.1689207 |
| SLC16A14 | protein_coding | -0.1440918 | 0.55238168 |
| SMARCA4 | protein_coding | -0.1441821 | 4.65832126 |
| NT5C | protein_coding | -0.1442812 | 1.64318937 |
| EXO5 | protein_coding | -0.1443003 | 1.86135114 |
| AP4S1 | protein_coding | -0.1443358 | 0.77040559 |
| RP1-101A2.1 | lncRNA | -0.1443407 | 0.24335575 |
| DYNLRB1 | protein_coding | -0.1443448 | 0.89444925 |
| COX6B1 | protein_coding | -0.1443782 | 4.54018236 |
| MUC5AC | protein_coding | -0.1444899 | 0.30061666 |
| HOXD9 | protein_coding | -0.144794 | 1.91215634 |
| CLP1 | protein_coding | -0.1448311 | 1.45556467 |
| CEP250 | protein_coding | -0.144948 | 4.39498203 |
| OSBPL8 | protein_coding | -0.1449756 | 3.03690716 |
| ELOC | protein_coding | -0.1450194 | 2.41008185 |
| NPR2 | protein_coding | -0.1450502 | 0.77456146 |
| RP1-118J21.[illegible] | lncRNA | -0.1450704 | 0.18489961 |
| MIR4453HG | lncRNA | -0.1450891 | 1.03694601 |
| PRKCI | protein_coding | -0.1450894 | 3.09303951 |
| ZNF300 | protein_coding | -0.1450897 | 1.25206401 |
| PRPS1 | protein_coding | -0.1451961 | 4.76540451 |
| ATP11B | protein_coding | -0.1452076 | 3.59177427 |
| PPOX | protein_coding | -0.1452498 | 1.46044425 |
| C11orf54 | protein_coding | -0.14534 | 1.50958025 |
| GMNN | protein_coding | -0.1454425 | 3.11769539 |
| NELFB | protein_coding | -0.1455369 | 3.58389716 |
| CAMK1D | protein_coding | -0.1456044 | 2.09670069 |
| RPL10A | protein_coding | -0.1457392 | 5.77365006 |
| ASTN2 | protein_coding | -0.1457963 | 2.64441882 |
| BPHL | protein_coding | -0.145941 | 2.20791508 |
| USP25 | protein_coding | -0.1460735 | 2.38919658 |
| ZNF695 | protein_coding | -0.1460831 | 0.3406606 |
| LINC01413 | lncRNA | -0.1460945 | 0.82774125 |
| PFDN1 | protein_coding | -0.1462098 | 2.50934071 |
| SNX12 | protein_coding | -0.1462515 | 2.69906686 |
| FLVCR1 | protein_coding | -0.1462531 | 2.99682847 |
| XPNPEP3 | protein_coding | -0.1462614 | 2.31040447 |
| ALG12 | protein_coding | -0.1463296 | 2.45322275 |
| ZNF574 | protein_coding | -0.146427 | 2.13484073 |
| GRB2 | protein_coding | -0.1464547 | 4.46978066 |
| KDSR | protein_coding | -0.1465282 | 2.79955812 |
| FUBP3 | protein_coding | -0.1466124 | 4.82181206 |
| XYLB | protein_coding | -0.1466201 | 2.06258546 |
| CCDC138 | protein_coding | -0.1466629 | 1.68588167 |

| | | | |
|---|---|---|---|
| LRP11 | protein_coding | -0.1466646 | 0.25707881 |
| TMEM38A | protein_coding | -0.1468339 | 1.22257161 |
| CLYBL | protein_coding | -0.1469781 | 0.74118971 |
| FAM168B | protein_coding | -0.1470547 | 6.20659752 |
| DLG4 | protein_coding | -0.1471066 | 1.16617933 |
| HOXD-AS2 | lncRNA | -0.1472706 | 0.51714166 |
| TMEM50B | protein_coding | -0.1473133 | 2.11934774 |
| FAM184A | protein_coding | -0.1473226 | 2.02835175 |
| AC068831.18 | protein_coding | -0.1473478 | 0.87287051 |
| SERINC1 | protein_coding | -0.1473914 | 3.12954564 |
| PC | protein_coding | -0.1474612 | 2.34629683 |
| COX7B | protein_coding | -0.1475569 | 4.20261675 |
| GOLGA7 | protein_coding | -0.1475824 | 2.82743767 |
| CDKAL1 | protein_coding | -0.1477036 | 5.62693482 |
| NUP155 | protein_coding | -0.1477663 | 4.6153515 |
| TIMM10 | protein_coding | -0.1478789 | 2.24621204 |
| UBAC1 | protein_coding | -0.1479564 | 3.67480558 |
| EIF4H | protein_coding | -0.1479672 | 5.45786812 |
| BCL7B | protein_coding | -0.1480624 | 3.0798994 |
| ALKBH5 | protein_coding | -0.1481326 | 2.28986714 |
| PABPC4L | protein_coding | -0.1481415 | 1.91796744 |
| VPS33A | protein_coding | -0.1481588 | 1.59492253 |
| DOHH | protein_coding | -0.1481858 | 0.86774528 |
| SF3A3 | protein_coding | -0.1482028 | 5.08263084 |
| NNT-AS1 | lncRNA | -0.1482159 | 3.11410854 |
| LRPPRC | protein_coding | -0.1482515 | 5.92043896 |
| SEC16A | protein_coding | -0.1484057 | 5.19925532 |
| NBPF3 | protein_coding | -0.1484098 | 1.54010547 |
| PPP3CA | protein_coding | -0.1484249 | 3.10353424 |
| OTUB1 | protein_coding | -0.1484255 | 3.76190885 |
| DUT | protein_coding | -0.1484746 | 3.20168593 |
| FBXO34 | protein_coding | -0.1485039 | 2.29237682 |
| UTP11 | protein_coding | -0.1485304 | 3.27200456 |
| SLC2A4RG | protein_coding | -0.1486077 | 1.71172784 |
| RP11-347I19 | lncRNA | -0.148633 | 0.2368696 |
| RP11-250B2. | lncRNA | -0.1487808 | 0.20713899 |
| PDLIM5 | protein_coding | -0.1488892 | 3.55960009 |
| RP11-51J9.7 | lncRNA | -0.148913 | 0.22379994 |
| EEF2 | protein_coding | -0.1490803 | 5.32034035 |
| FGD3 | protein_coding | -0.1491186 | 1.32985635 |
| CTD-2350C19 | lncRNA | -0.1491507 | 0.16249476 |
| GRSF1 | protein_coding | -0.1492933 | 4.21193997 |
| AMER2 | protein_coding | -0.1493176 | 0.5256225 |
| KCNQ1OT1 | lncRNA | -0.1493939 | 1.31728151 |
| CD2BP2-DT | lncRNA | -0.1494629 | 0.28436511 |

| | | | |
|---|---|---|---|
| NAP1L3 | protein_coding | -0.1494724 | 2.37532971 |
| B3GAT3 | protein_coding | -0.1494924 | 4.72547635 |
| PHF10 | protein_coding | -0.14951 | 3.7036621 |
| IGSF8 | protein_coding | -0.1496016 | 2.18800484 |
| RTN4RL2 | protein_coding | -0.149618 | 1.91013792 |
| MIR924HG | lncRNA | -0.1496475 | 1.70256946 |
| SYAP1 | protein_coding | -0.1496861 | 2.62813806 |
| RP11-196B3. | lncRNA | -0.1497066 | 0.32048381 |
| SPPL3 | protein_coding | -0.1497928 | 3.667218 |
| KLK3 | protein_coding | -0.1498277 | 0.15568467 |
| TJP3 | protein_coding | -0.1499361 | 1.89863945 |
| CRCP | protein_coding | -0.1499413 | 3.13253413 |
| AP000704.5 | lncRNA | -0.1500206 | 0.60302869 |
| VPS9D1-AS1 | lncRNA | -0.1500561 | 1.50486997 |
| MB21D2 | protein_coding | -0.1501263 | 2.05449161 |
| PA2G4 | protein_coding | -0.1501362 | 5.32164614 |
| CCDC83 | protein_coding | -0.1501574 | 0.15271678 |
| USP13 | protein_coding | -0.1501804 | 3.20162382 |
| SRSF1 | protein_coding | -0.1501952 | 1.75180239 |
| CENPQ | protein_coding | -0.1502747 | 1.08503868 |
| FAM160B2 | protein_coding | -0.1503262 | 1.31885355 |
| ERH | protein_coding | -0.1503268 | 3.94353599 |
| AAAS | protein_coding | -0.1503599 | 3.57537029 |
| EXD2 | protein_coding | -0.1503807 | 3.79156878 |
| ICMT | protein_coding | -0.150387 | 4.44661575 |
| VRK1 | protein_coding | -0.150409 | 3.43623837 |
| NPTXR | protein_coding | -0.1504321 | 4.83750417 |
| SPRED2 | protein_coding | -0.1504552 | 1.09978349 |
| VAV3 | protein_coding | -0.1509113 | 3.12393374 |
| COX7A2L | protein_coding | -0.1509286 | 3.73896975 |
| PJA2 | protein_coding | -0.1509581 | 3.70663898 |
| NDUFB11 | protein_coding | -0.1509582 | 4.02589946 |
| EXOSC4 | protein_coding | -0.1509647 | 1.70263813 |
| RNF4 | protein_coding | -0.151003 | 4.35078568 |
| CEP19 | protein_coding | -0.1510508 | 1.11271562 |
| PCDHB1 | protein_coding | -0.1510835 | 0.15014929 |
| BMI1 | protein_coding | -0.1511114 | 1.199204 |
| SNRNP70 | protein_coding | -0.1511198 | 2.81257954 |
| PRR15L | protein_coding | -0.1513086 | 2.59695518 |
| POLA1 | protein_coding | -0.15135 | 4.88125601 |
| CBX8 | protein_coding | -0.1513558 | 2.15163923 |
| CENPE | protein_coding | -0.151374 | 4.35313015 |
| LINC01088 | lncRNA | -0.1514533 | 0.38243687 |
| RTTN | protein_coding | -0.1515933 | 2.85072953 |
| FBXL18 | protein_coding | -0.151656 | 2.20337807 |

| | | | |
|---|---|---|---|
| TRMT10C | protein_coding | -0.1516641 | 2.16608975 |
| NEIL3 | protein_coding | -0.1516826 | 2.50887751 |
| METTL9 | protein_coding | -0.1518675 | 4.80069075 |
| FBRSL1 | protein_coding | -0.1520074 | 1.50983903 |
| PDXDC1 | protein_coding | -0.152014 | 5.85425199 |
| LMAN2L | protein_coding | -0.1520249 | 2.61811269 |
| ARRB2 | protein_coding | -0.1520438 | 3.44015945 |
| WASF1 | protein_coding | -0.1520927 | 3.46195751 |
| LYRM1 | protein_coding | -0.1521182 | 1.62245103 |
| AHSA1 | protein_coding | -0.1521337 | 5.7079809 |
| TMEM63B | protein_coding | -0.1521382 | 4.30376282 |
| PPAT | protein_coding | -0.1523511 | 4.54237435 |
| NDUFAF3 | protein_coding | -0.1523557 | 1.50814985 |
| GOLT1B | protein_coding | -0.1523838 | 1.9039259 |
| SCAP | protein_coding | -0.1524316 | 3.45824808 |
| CELSR2 | protein_coding | -0.1524865 | 4.77735964 |
| NOTUM | protein_coding | -0.1525386 | 2.21905925 |
| RP11-390F4. | lncRNA | -0.1525739 | 0.60826293 |
| FAM8A1 | protein_coding | -0.1526141 | 3.01662066 |
| GALNS | protein_coding | -0.1526363 | 1.96009104 |
| ATG13 | protein_coding | -0.1526539 | 4.14621545 |
| NRBP2 | protein_coding | -0.1526626 | 2.00356425 |
| RPL13 | protein_coding | -0.1526892 | 1.55471288 |
| PEBP1 | protein_coding | -0.1526996 | 5.2696431 |
| C8orf33 | protein_coding | -0.1527432 | 4.08826153 |
| NAP1L4 | protein_coding | -0.1527947 | 5.64077379 |
| GRPEL1 | protein_coding | -0.152863 | 3.26981453 |
| MPP1 | protein_coding | -0.1528875 | 1.90027521 |
| RP3-337H4.9 | lncRNA | -0.1529467 | 0.36027437 |
| ZNF624 | protein_coding | -0.1529743 | 0.57284353 |
| DICER1 | protein_coding | -0.1530509 | 4.74991508 |
| NOP9 | protein_coding | -0.1530514 | 4.39456085 |
| SPINDOC | protein_coding | -0.1531729 | 3.05888845 |
| NUP50 | protein_coding | -0.153294 | 5.45977277 |
| RPL23 | protein_coding | -0.1533635 | 6.08607115 |
| CYHR1 | protein_coding | -0.1533692 | 2.860683 |
| ENAH | protein_coding | -0.1533757 | 3.67746196 |
| TUT1 | protein_coding | -0.1534802 | 2.91733933 |
| THOC6 | protein_coding | -0.153554 | 3.77327469 |
| CRBN | protein_coding | -0.1536482 | 1.77350155 |
| INSIG1-DT | lncRNA | -0.1537372 | 1.20032733 |
| NUDT19 | protein_coding | -0.1537493 | 0.95867295 |
| UCP3 | protein_coding | -0.1538846 | 0.40280869 |
| KIF4A | protein_coding | -0.1539136 | 3.67513885 |
| WDR37 | protein_coding | -0.1539518 | 1.80334671 |

| | | | |
|---|---|---|---|
| B4GALNT4 | protein_coding | -0.1539588 | 1.25896584 |
| DUSP19 | protein_coding | -0.1539958 | 0.71934108 |
| CYB5R2 | protein_coding | -0.154079 | 1.26105753 |
| RP11-88E10. | lncRNA | -0.1543448 | 0.53498722 |
| CDT1 | protein_coding | -0.1543998 | 2.01027117 |
| SCAND1 | protein_coding | -0.1544012 | 0.94562318 |
| MED29 | protein_coding | -0.1544746 | 4.49326507 |
| RPS9 | protein_coding | -0.1546196 | 5.0638268 |
| MMUT | protein_coding | -0.1547024 | 3.57210433 |
| RP5-940J5.9 | lncRNA | -0.1547734 | 0.4388729 |
| RP11-4O1.2 | lncRNA | -0.1548315 | 0.21684853 |
| GPLD1 | protein_coding | -0.1548721 | 0.4778278 |
| SVBP | protein_coding | -0.1548839 | 2.79334758 |
| SPATA2L | protein_coding | -0.1552377 | 1.22755623 |
| NRCAM | protein_coding | -0.1552409 | 3.17445864 |
| DRP2 | protein_coding | -0.1553695 | 0.21119139 |
| MTF2 | protein_coding | -0.1554079 | 3.05754285 |
| APOO | protein_coding | -0.1554139 | 2.09628235 |
| CHML | protein_coding | -0.1556808 | 3.60129417 |
| CLCN7 | protein_coding | -0.1558183 | 2.49867375 |
| IL17RE | protein_coding | -0.155839 | 0.87874564 |
| MICOS10 | protein_coding | -0.1559764 | 1.23929103 |
| TNKS2 | protein_coding | -0.1560593 | 2.82613701 |
| PPP1R12A | protein_coding | -0.1561389 | 3.5197561 |
| SNAP29 | protein_coding | -0.1561845 | 3.88719936 |
| POLE | protein_coding | -0.156256 | 5.27511758 |
| GARRE1 | protein_coding | -0.1564509 | 2.32998869 |
| MOCS2 | protein_coding | -0.156474 | 2.70649169 |
| ODC1 | protein_coding | -0.1564784 | 5.30890172 |
| ACE | protein_coding | -0.1565706 | 3.03724254 |
| PGAM5 | protein_coding | -0.1566409 | 4.4946781 |
| AOC2 | protein_coding | -0.1566461 | 0.25166843 |
| SPCS1 | protein_coding | -0.1566877 | 3.50472723 |
| PIGX | protein_coding | -0.156755 | 3.18415555 |
| FIGNL1 | protein_coding | -0.1569195 | 3.03996529 |
| ERCC3 | protein_coding | -0.1569713 | 4.07094969 |
| TRNT1 | protein_coding | -0.1570061 | 2.0693701 |
| SYNGR4 | protein_coding | -0.1570595 | 0.37971736 |
| TAC3 | protein_coding | -0.157122 | 1.2305256 |
| POLA2 | protein_coding | -0.1571222 | 1.6325461 |
| MYH7B | protein_coding | -0.1572763 | 0.38140562 |
| CANT1 | protein_coding | -0.1573491 | 2.75468222 |
| KIAA1143 | protein_coding | -0.1573887 | 2.56901155 |
| GPR37 | protein_coding | -0.157389 | 1.12276946 |
| VDAC2 | protein_coding | -0.157443 | 3.70053523 |

| | | | |
|---|---|---|---|
| C1orf198 | protein_coding | -0.1575331 | 4.60530943 |
| BNIP1 | protein_coding | -0.1575907 | 1.21038097 |
| DR1 | protein_coding | -0.1576118 | 3.60757839 |
| MRPL11 | protein_coding | -0.1576248 | 3.22856233 |
| ETFB | protein_coding | -0.1576285 | 2.20096794 |
| AC106786.1 | lncRNA | -0.1576666 | 0.48874878 |
| PSRC1 | protein_coding | -0.1576846 | 2.18386879 |
| SMIM2-AS1 | lncRNA | -0.1577179 | 0.1146854 |
| OCEL1 | protein_coding | -0.1577808 | 0.38552765 |
| CTCF | protein_coding | -0.1577954 | 4.67739261 |
| HP1BP3 | protein_coding | -0.1578426 | 6.61366461 |
| KHSRP | protein_coding | -0.1579385 | 5.51274749 |
| MRPL55 | protein_coding | -0.1580471 | 2.28413442 |
| MYB | protein_coding | -0.1581207 | 1.78787842 |
| CTBP1 | protein_coding | -0.15826 | 3.6812805 |
| COPS9 | protein_coding | -0.1584356 | 2.58083607 |
| RP11-723O4. | lncRNA | -0.1584606 | 0.85675087 |
| CHID1 | protein_coding | -0.1584936 | 3.5267151 |
| MYO9B | protein_coding | -0.1584993 | 3.07266267 |
| RAB1B | protein_coding | -0.1587339 | 5.50412828 |
| CENPM | protein_coding | -0.1587467 | 2.78301357 |
| TCEANC2 | protein_coding | -0.1588998 | 3.38668721 |
| AGPAT1 | protein_coding | -0.1589454 | 5.56376569 |
| KLHL42 | protein_coding | -0.1590231 | 4.91033863 |
| ATOX1 | protein_coding | -0.1590674 | 2.14313333 |
| RNF167 | protein_coding | -0.1591208 | 3.012416 |
| ZFP1 | protein_coding | -0.1591667 | 2.83871279 |
| GMEB1 | protein_coding | -0.1593213 | 1.54964898 |
| AC093627.9 | lncRNA | -0.1593682 | 1.23235635 |
| REEP4 | protein_coding | -0.159408 | 2.42708229 |
| ZP3 | protein_coding | -0.1594217 | 1.82869877 |
| POLR1G | protein_coding | -0.1594285 | 2.58393244 |
| TMEM139-A! | lncRNA | -0.1594874 | 0.15957983 |
| H2AC15 | protein_coding | -0.1595917 | 0.18718186 |
| RANBP1 | protein_coding | -0.1596215 | 4.64315947 |
| RP11-700J17 | lncRNA | -0.1597092 | 0.27879832 |
| DVL3 | protein_coding | -0.159732 | 6.26324758 |
| DLL3 | protein_coding | -0.1597805 | 0.95341893 |
| CHAMP1 | protein_coding | -0.1599409 | 4.00362219 |
| CEP162 | protein_coding | -0.1600111 | 1.18553351 |
| FAM200A | protein_coding | -0.1600998 | 1.74045804 |
| ZNF407 | protein_coding | -0.1601473 | 1.79529977 |
| PRRT3 | protein_coding | -0.1601871 | 0.61185735 |
| CERS2 | protein_coding | -0.1602326 | 4.94825346 |
| ARID1A | protein_coding | -0.1603693 | 6.08592335 |

| | | | |
|---|---|---|---|
| RP4-646N3.2 | lncRNA | -0.160377 | 0.36059962 |
| MOAP1 | protein_coding | -0.1605022 | 3.09219028 |
| ARHGEF18 | protein_coding | -0.1606227 | 2.35140982 |
| UPK3A | protein_coding | -0.1606304 | 0.28858632 |
| NOP53 | protein_coding | -0.1606687 | 1.79821118 |
| AP006621.6 | lncRNA | -0.1607909 | 0.32523378 |
| LRRC24 | protein_coding | -0.1607929 | 0.73042897 |
| SELENOT | protein_coding | -0.1607981 | 3.15902299 |
| GNPNAT1 | protein_coding | -0.1608688 | 1.96190134 |
| CTD-2267D1 | lncRNA | -0.1609275 | 0.3503399 |
| LYPLA1 | protein_coding | -0.1609373 | 1.97675421 |
| ZNF354A | protein_coding | -0.160986 | 1.59523314 |
| CHRNE | protein_coding | -0.1610197 | 0.2036693 |
| RPS11 | protein_coding | -0.1611909 | 7.05325971 |
| RND3 | protein_coding | -0.1612188 | 3.81540867 |
| PTK6 | protein_coding | -0.1612556 | 0.38410128 |
| GSK3B | protein_coding | -0.1612975 | 3.73130444 |
| ZNF92 | protein_coding | -0.1613076 | 1.76238295 |
| MINK1 | protein_coding | -0.1614313 | 3.11673541 |
| FOXP3 | protein_coding | -0.1614783 | 0.39548225 |
| MFSD6 | protein_coding | -0.1614911 | 3.62781143 |
| AC079807.2 | lncRNA | -0.1615087 | 0.21513939 |
| ERP29 | protein_coding | -0.1616435 | 5.57832955 |
| MMAA | protein_coding | -0.1616952 | 1.15988914 |
| SCOC | protein_coding | -0.1617161 | 2.21091436 |
| DIDO1 | protein_coding | -0.1618404 | 4.2839249 |
| OGDHL | protein_coding | -0.161948 | 1.45863739 |
| TXN2 | protein_coding | -0.1620131 | 3.84666482 |
| RP11-817O1 | lncRNA | -0.1621593 | 0.60175971 |
| CTPS1 | protein_coding | -0.1623143 | 6.39915937 |
| RAB15 | protein_coding | -0.1623385 | 5.08180389 |
| ILVBL | protein_coding | -0.1623588 | 3.74484114 |
| NOSIP | protein_coding | -0.1625253 | 2.52131291 |
| RPL28 | protein_coding | -0.1625272 | 5.16621012 |
| RP11-225B1 | lncRNA | -0.1625337 | 0.34319923 |
| NDUFS8 | protein_coding | -0.1625511 | 3.13867909 |
| SUMF2 | protein_coding | -0.1626351 | 5.47204306 |
| CAND2 | protein_coding | -0.1626798 | 1.49771716 |
| MTA1 | protein_coding | -0.1626974 | 2.94043971 |
| RAD1 | protein_coding | -0.1627713 | 3.82540555 |
| ZMAT3 | protein_coding | -0.1627848 | 2.0027049 |
| B3GNT9 | protein_coding | -0.1628024 | 1.63640009 |
| WRAP53 | protein_coding | -0.1628319 | 1.76387312 |
| GNAI3 | protein_coding | -0.1628621 | 4.43968111 |
| HAUS6 | protein_coding | -0.1629016 | 3.44051764 |

| | | | |
|---|---|---|---|
| FBXO11 | protein_coding | -0.1629534 | 4.60652615 |
| C2orf69 | protein_coding | -0.1629591 | 2.18176164 |
| GIHCG | lncRNA | -0.1630056 | 1.07240699 |
| RCCD1 | protein_coding | -0.1630214 | 2.79732939 |
| TRIP13 | protein_coding | -0.1631634 | 3.28727498 |
| GADD45GIP1 | protein_coding | -0.1631918 | 1.53539661 |
| STK39 | protein_coding | -0.1631926 | 4.32108503 |
| GPR173 | protein_coding | -0.163246 | 0.78771358 |
| NUP85 | protein_coding | -0.1636339 | 4.03339413 |
| ERBB2 | protein_coding | -0.1637512 | 1.30939882 |
| DNMT3B | protein_coding | -0.163902 | 1.31623604 |
| MRFAP1 | protein_coding | -0.1639282 | 6.59958455 |
| XRCC5 | protein_coding | -0.1639617 | 7.21552248 |
| SPAG16 | protein_coding | -0.1640299 | 1.34440362 |
| RSU1 | protein_coding | -0.1641125 | 1.31688511 |
| PPP1R3D | protein_coding | -0.1641358 | 0.3271737 |
| WWOX | protein_coding | -0.1641466 | 2.84484721 |
| ERI2 | protein_coding | -0.1642111 | 3.13602991 |
| DHRS4-AS1 | lncRNA | -0.1642586 | 1.90007732 |
| PACC1 | protein_coding | -0.1643532 | 2.34539079 |
| MAP3K9 | protein_coding | -0.1643666 | 4.26402395 |
| DTNBP1 | protein_coding | -0.1644382 | 3.35205676 |
| CTD-2265O2 | lncRNA | -0.1645036 | 0.20284213 |
| ZNF774 | protein_coding | -0.1645875 | 1.77882545 |
| ERG28 | protein_coding | -0.1646312 | 5.44013763 |
| CTD-3074O7 | lncRNA | -0.1646826 | 0.47358651 |
| DCP2 | protein_coding | -0.1647142 | 4.14270598 |
| TTI2 | protein_coding | -0.1647192 | 1.50137667 |
| SNIP1 | protein_coding | -0.1647463 | 2.79523894 |
| PRPF38A | protein_coding | -0.1648227 | 5.06073143 |
| EMC10 | protein_coding | -0.1648615 | 4.32406946 |
| TCERG1 | protein_coding | -0.1648841 | 5.76895041 |
| METTL26 | protein_coding | -0.1649659 | 2.18947069 |
| CRTC1 | protein_coding | -0.1650805 | 2.43447837 |
| LSM8 | protein_coding | -0.1650824 | 4.23527737 |
| LIMK2 | protein_coding | -0.1650879 | 3.03833098 |
| LRRIQ3 | protein_coding | -0.1652718 | 0.51471994 |
| SLC25A26 | protein_coding | -0.1654417 | 1.24669458 |
| MARCHF1 | protein_coding | -0.1655278 | 0.50416132 |
| NSD2 | protein_coding | -0.1655962 | 6.48650552 |
| CCNB1 | protein_coding | -0.1656136 | 5.40572597 |
| TRIR | protein_coding | -0.1656619 | 1.5703207 |
| FKBPL | protein_coding | -0.1657299 | 1.54869748 |
| TIMELESS | protein_coding | -0.1657406 | 6.15553086 |
| RP11-120K24 | lncRNA | -0.1657624 | 0.84405311 |

| | | | |
|---|---|---|---|
| ADCK1 | protein_coding | -0.1657705 | 2.18757898 |
| GPS1 | protein_coding | -0.1657741 | 4.26388478 |
| EEF1D | protein_coding | -0.1657862 | 3.00746069 |
| DOLK | protein_coding | -0.1658151 | 1.90073011 |
| MTAP | protein_coding | -0.1658736 | 2.49487005 |
| LINC00476 | lncRNA | -0.1658891 | 0.75948855 |
| AURKAIP1 | protein_coding | -0.1658963 | 1.34586681 |
| YRDC | protein_coding | -0.1659191 | 3.2183163 |
| BTBD2 | protein_coding | -0.1659652 | 3.0801582 |
| NOP10 | protein_coding | -0.1660097 | 4.01203572 |
| MRPS34 | protein_coding | -0.1660551 | 1.95378114 |
| BRWD1 | protein_coding | -0.1660966 | 5.07087752 |
| PPM1D | protein_coding | -0.1661393 | 3.33177579 |
| DTYMK | protein_coding | -0.1664919 | 3.85648077 |
| CENPF | protein_coding | -0.1665019 | 6.7276681 |
| EID2 | protein_coding | -0.1666413 | 2.55322299 |
| ZNF614 | protein_coding | -0.1667244 | 1.48052314 |
| CLTRN | protein_coding | -0.1667811 | 0.22920489 |
| PRR12 | protein_coding | -0.1668002 | 2.84570657 |
| FBXL19 | protein_coding | -0.1668369 | 1.57482344 |
| CIB1 | protein_coding | -0.1669596 | 2.01864958 |
| KLLN | protein_coding | -0.1669672 | 0.25198955 |
| SMPD4 | protein_coding | -0.1669928 | 4.66903377 |
| DDX46 | protein_coding | -0.1671203 | 5.75258524 |
| USP27X-DT | lncRNA | -0.1671341 | 0.23632669 |
| RPL32 | protein_coding | -0.1673441 | 5.44195221 |
| NDUFAF6 | protein_coding | -0.1674082 | 3.21138925 |
| MYL12B | protein_coding | -0.1675675 | 5.01828796 |
| MAN1A1 | protein_coding | -0.1675701 | 1.96414767 |
| ABCG4 | protein_coding | -0.167665 | 0.17773723 |
| ESS2 | protein_coding | -0.1676916 | 2.51061274 |
| RP11-305O2( | lncRNA | -0.1677078 | 0.53966902 |
| RPS2 | protein_coding | -0.167806 | 3.58736372 |
| TPCN1 | protein_coding | -0.1680095 | 4.17832488 |
| PLEKHG5 | protein_coding | -0.1680183 | 2.01624925 |
| PLA2G12A | protein_coding | -0.1681046 | 3.79515925 |
| ZW10 | protein_coding | -0.1681062 | 1.82324449 |
| PRKAR2A | protein_coding | -0.1681071 | 2.37644529 |
| SAP18 | protein_coding | -0.1682002 | 4.22200951 |
| SRGAP2 | protein_coding | -0.1682013 | 3.55689955 |
| VLDLR | protein_coding | -0.1682057 | 2.18830894 |
| POLD3 | protein_coding | -0.168336 | 4.09159622 |
| SEC24B | protein_coding | -0.1683725 | 2.50441869 |
| BCKDHB | protein_coding | -0.1683825 | 2.68670777 |
| CERT1 | protein_coding | -0.168478 | 3.29799117 |

| | | | |
|---|---|---|---|
| RP11-156L14 | lncRNA | -0.1685112 | 0.57350904 |
| WEE1 | protein_coding | -0.1685147 | 4.39833903 |
| SIAH3 | protein_coding | -0.1685193 | 0.71250521 |
| SMNDC1 | protein_coding | -0.1686067 | 2.96012225 |
| SMIM19 | protein_coding | -0.1687949 | 1.44262939 |
| ADCK2 | protein_coding | -0.168795 | 3.05714558 |
| FAM204A | protein_coding | -0.1688035 | 2.62195405 |
| TRAK2 | protein_coding | -0.1688525 | 3.63687894 |
| CLN6 | protein_coding | -0.1688763 | 3.45968644 |
| RHNO1 | protein_coding | -0.1690455 | 3.19648325 |
| ZFP69 | protein_coding | -0.1690542 | 1.84561269 |
| MARVELD1 | protein_coding | -0.1691009 | 2.01470071 |
| ARSB | protein_coding | -0.16913 | 2.72313699 |
| UBE2T | protein_coding | -0.1691843 | 4.25475652 |
| MRPL28 | protein_coding | -0.1691956 | 4.30437327 |
| RAPGEF2 | protein_coding | -0.1692429 | 2.20054538 |
| KCTD10 | protein_coding | -0.169243 | 3.61931894 |
| BBIP1 | protein_coding | -0.1692593 | 1.20827646 |
| UBA2 | protein_coding | -0.1692884 | 6.45124872 |
| NABP2 | protein_coding | -0.1693025 | 4.66137357 |
| FAM174A | protein_coding | -0.1693056 | 2.24120261 |
| RP11-127B2( | lncRNA | -0.1693269 | 0.25279054 |
| TGM1 | protein_coding | -0.1694818 | 0.80604191 |
| DROSHA | protein_coding | -0.1694833 | 5.73448038 |
| SERP1 | protein_coding | -0.1695175 | 4.54710832 |
| HSPE1 | protein_coding | -0.1700302 | 2.25210146 |
| COX8A | protein_coding | -0.1700357 | 4.20324763 |
| HNRNPM | protein_coding | -0.170039 | 4.0997189 |
| RP4-809E13. | lncRNA | -0.1700926 | 0.29802915 |
| MPP6 | protein_coding | -0.1701226 | 2.43576014 |
| FRMD8 | protein_coding | -0.1701541 | 3.13433288 |
| UIMC1 | protein_coding | -0.1701862 | 2.56934754 |
| COPZ1 | protein_coding | -0.1701928 | 5.51551948 |
| ARMH3 | protein_coding | -0.1702627 | 2.28328605 |
| RNF130 | protein_coding | -0.1704332 | 3.81698893 |
| ATP6V1B2 | protein_coding | -0.1704958 | 4.6874937 |
| AC079305.8 | lncRNA | -0.1706563 | 0.26320717 |
| EID2B | protein_coding | -0.1707037 | 0.83811869 |
| RFWD3 | protein_coding | -0.1707652 | 5.75565991 |
| DYRK1B | protein_coding | -0.1708044 | 1.09765249 |
| HSPA13 | protein_coding | -0.1708439 | 2.70301025 |
| STAU2 | protein_coding | -0.1709365 | 2.11383925 |
| TBL1XR1 | protein_coding | -0.1710079 | 2.76337156 |
| GMCL1 | protein_coding | -0.171257 | 3.05401863 |
| AOPEP | protein_coding | -0.1712704 | 2.01740931 |

| | | | |
|---|---|---|---|
| RP11-319G6. | lncRNA | -0.1712708 | 0.39845227 |
| TESK1 | protein_coding | -0.1714239 | 2.87023189 |
| KRAS | protein_coding | -0.1715787 | 3.6548413 |
| ELOVL4 | protein_coding | -0.1716984 | 2.27383601 |
| SRPRB | protein_coding | -0.1717029 | 5.37294554 |
| EIF1AX | protein_coding | -0.1717045 | 3.69879812 |
| ELF4 | protein_coding | -0.1717858 | 3.99938442 |
| RP11-38L15. | lncRNA | -0.1717867 | 0.24061751 |
| ZC3H10 | protein_coding | -0.1718471 | 1.89334518 |
| FRG1 | protein_coding | -0.1719494 | 2.89591952 |
| JPT2 | protein_coding | -0.1719606 | 6.58080119 |
| CDCA7 | protein_coding | -0.1720684 | 5.33716874 |
| BRMS1 | protein_coding | -0.172133 | 3.84589457 |
| ATP6V1D | protein_coding | -0.1721727 | 3.56629335 |
| COX20 | protein_coding | -0.1722195 | 2.36035795 |
| RP11-98I9.4 | lncRNA | -0.1722319 | 0.61993778 |
| TAGLN3 | protein_coding | -0.1722939 | 4.205383 |
| RP13-512J5. | lncRNA | -0.1723004 | 0.72785876 |
| TRIM35 | protein_coding | -0.1723207 | 2.33451916 |
| MAPK1 | protein_coding | -0.1723566 | 6.13486515 |
| FAM201A | lncRNA | -0.1724359 | 1.97847915 |
| HMGB2 | protein_coding | -0.1724467 | 5.80828758 |
| MPRIP | protein_coding | -0.1724781 | 5.46569694 |
| ZBTB11 | protein_coding | -0.1726457 | 4.28598614 |
| TRAPPC1 | protein_coding | -0.1727158 | 2.13828575 |
| ZNF816 | protein_coding | -0.1727297 | 1.12601071 |
| TCHP | protein_coding | -0.1728049 | 3.09370435 |
| ZNF845 | protein_coding | -0.1729593 | 2.64409574 |
| ACTR1A | protein_coding | -0.1729829 | 5.62244573 |
| GINS1 | protein_coding | -0.1731013 | 2.83688353 |
| RP5-832C2.5 | lncRNA | -0.1731075 | 0.30184556 |
| TP53INP1 | protein_coding | -0.1731467 | 2.10170484 |
| MARCHF9 | protein_coding | -0.1731842 | 2.79109302 |
| IER3IP1 | protein_coding | -0.173353 | 1.68072317 |
| PLS1 | protein_coding | -0.1733541 | 2.95131445 |
| RP1-101K10. | lncRNA | -0.1734021 | 0.45703064 |
| ZMYM3 | protein_coding | -0.1735295 | 5.19852239 |
| ING1 | protein_coding | -0.1735817 | 2.49355483 |
| PHF20 | protein_coding | -0.1735951 | 4.55803644 |
| UGP2 | protein_coding | -0.1737388 | 3.60125697 |
| MADCAM1 | protein_coding | -0.1738562 | 0.40125073 |
| COX14 | protein_coding | -0.1739231 | 2.19152777 |
| CSTF3 | protein_coding | -0.1739646 | 4.89763378 |
| NACC1 | protein_coding | -0.1740962 | 3.20125226 |
| PREB | protein_coding | -0.1741357 | 4.62595259 |

| | | | |
|---|---|---|---|
| CTD-2017C7. | lncRNA | -0.1741581 | 0.48194576 |
| HSPA1L | protein_coding | -0.1741693 | 0.52839483 |
| PACS2 | protein_coding | -0.174206 | 4.73448038 |
| DOK3 | protein_coding | -0.1742699 | 0.9787247 |
| TMED3 | protein_coding | -0.1742983 | 4.22905708 |
| NPIPB15 | protein_coding | -0.1743301 | 0.55251761 |
| DSN1 | protein_coding | -0.174337 | 3.54290659 |
| ABHD17C | protein_coding | -0.174372 | 3.25216508 |
| RAD54L | protein_coding | -0.1744157 | 3.01930962 |
| RALGPS1 | protein_coding | -0.1745023 | 4.12508108 |
| SLC25A46 | protein_coding | -0.1745319 | 2.62158008 |
| ZNF891 | protein_coding | -0.1746303 | 1.81786975 |
| PPDPF | protein_coding | -0.1747172 | 1.74084992 |
| FBXO5 | protein_coding | -0.174761 | 2.48441667 |
| BTF3 | protein_coding | -0.1747936 | 6.26064892 |
| CARD9 | protein_coding | -0.1748273 | 0.42715694 |
| PPP2R5C | protein_coding | -0.1748552 | 5.82898324 |
| QSOX2 | protein_coding | -0.1748889 | 6.40288459 |
| TUT7 | protein_coding | -0.1749552 | 3.54644281 |
| ARHGAP1 | protein_coding | -0.1750171 | 4.73591234 |
| INO80E | protein_coding | -0.1750515 | 3.47545919 |
| TCF19 | protein_coding | -0.1751104 | 6.21908773 |
| PTRHD1 | protein_coding | -0.1751302 | 2.81873886 |
| OARD1 | protein_coding | -0.1752135 | 2.85505899 |
| RPL10 | protein_coding | -0.1753333 | 6.98205925 |
| NDC80 | protein_coding | -0.1753989 | 3.23147785 |
| ARHGEF26 | protein_coding | -0.1754408 | 2.3412851 |
| LINC00205 | lncRNA | -0.175517 | 3.68115435 |
| TADA1 | protein_coding | -0.1755186 | 2.67018618 |
| RP11-96H17. | lncRNA | -0.1757304 | 0.25900874 |
| APAF1 | protein_coding | -0.1757444 | 3.4333639 |
| ZMAT2 | protein_coding | -0.1757456 | 6.12541369 |
| CHCHD1 | protein_coding | -0.1757467 | 2.17216022 |
| KLHL24 | protein_coding | -0.1757486 | 4.99562567 |
| FAM32A | protein_coding | -0.1758904 | 4.97039436 |
| PIH1D1 | protein_coding | -0.1758937 | 3.65191707 |
| FAM89A | protein_coding | -0.1759449 | 0.32198247 |
| VPS45 | protein_coding | -0.1759603 | 4.45968778 |
| CCNK | protein_coding | -0.175965 | 5.4030666 |
| TMEM101 | protein_coding | -0.1759944 | 2.40966522 |
| HAUS8 | protein_coding | -0.1760642 | 2.69034158 |
| BORCS5 | protein_coding | -0.1760825 | 2.29777123 |
| HDAC2 | protein_coding | -0.1761384 | 5.69011432 |
| PPIL2 | protein_coding | -0.1762859 | 4.56127076 |
| TMED2 | protein_coding | -0.1763203 | 5.72825641 |

| | | | |
|---|---|---|---|
| METAP1 | protein_coding | -0.1764705 | 4.92554985 |
| HOMER3 | protein_coding | -0.1764937 | 0.71639463 |
| HEPACAM2 | protein_coding | -0.1765289 | 6.49974305 |
| GKAP1 | protein_coding | -0.1765986 | 2.00457289 |
| DEPDC5 | protein_coding | -0.1766634 | 2.51674846 |
| GTDC1 | protein_coding | -0.1767149 | 2.45698583 |
| CENPU | protein_coding | -0.1767302 | 5.41217234 |
| RAB13 | protein_coding | -0.1768593 | 0.56003858 |
| SLC25A19 | protein_coding | -0.1769148 | 3.38480365 |
| DDX24 | protein_coding | -0.1769451 | 6.72029509 |
| CTB-147N14. | lncRNA | -0.177053 | 0.26725374 |
| POLD1 | protein_coding | -0.1771517 | 3.59967263 |
| SORD | protein_coding | -0.1771582 | 3.96158506 |
| HSP90AA1 | protein_coding | -0.1771846 | 8.89664233 |
| PHF2 | protein_coding | -0.1772154 | 6.14856695 |
| NFYA | protein_coding | -0.1773208 | 4.04072801 |
| RP11-1C8.4 | lncRNA | -0.1773954 | 0.34417032 |
| GRK4 | protein_coding | -0.1774901 | 1.63577431 |
| ZNF595 | protein_coding | -0.1777927 | 2.14785311 |
| SPATA24 | protein_coding | -0.1778504 | 0.67076303 |
| BRICD5 | protein_coding | -0.1778776 | 0.715121 |
| VMA21 | protein_coding | -0.1779219 | 3.45649539 |
| ANP32A | protein_coding | -0.1780017 | 6.45025508 |
| IQCC | protein_coding | -0.1781823 | 2.42249823 |
| C9orf16 | protein_coding | -0.178472 | 3.03342079 |
| SWI5 | protein_coding | -0.1784767 | 2.80076469 |
| RFC5 | protein_coding | -0.1785073 | 4.18244573 |
| BRD3 | protein_coding | -0.1785264 | 4.47697274 |
| RP11-352M1 | lncRNA | -0.1785332 | 2.79757243 |
| ATP6V0B | protein_coding | -0.1785922 | 6.92332429 |
| CRYBG1 | protein_coding | -0.1787293 | 2.21457961 |
| ELF2 | protein_coding | -0.1787556 | 5.07313191 |
| CHEK1 | protein_coding | -0.1787582 | 3.56902922 |
| TTC28-AS1 | lncRNA | -0.1788587 | 2.39435766 |
| CRYBG2 | protein_coding | -0.1790595 | 0.86770195 |
| ORC6 | protein_coding | -0.1790727 | 3.51613565 |
| METTL14-DT | lncRNA | -0.1792058 | 0.41857562 |
| STX1A | protein_coding | -0.1796193 | 4.96613416 |
| BRAT1 | protein_coding | -0.1796323 | 3.71920968 |
| CHMP1B | protein_coding | -0.1796676 | 3.73640854 |
| BLOC1S1 | protein_coding | -0.1796727 | 0.67393768 |
| VTI1B | protein_coding | -0.179698 | 4.03416422 |
| KANTR | protein_coding | -0.1797004 | 1.50328367 |
| TRIM32 | protein_coding | -0.1798969 | 3.45076958 |
| TANC2 | protein_coding | -0.1800862 | 5.99622135 |

| | | | |
|---|---|---|---|
| LRIG1 | protein_coding | -0.180312 | 4.14560504 |
| RAB33B-AS1 | lncRNA | -0.1803234 | 1.28625345 |
| YIPF3 | protein_coding | -0.1803522 | 3.83594241 |
| MCAT | protein_coding | -0.1803906 | 2.51303723 |
| CASZ1 | protein_coding | -0.1805427 | 3.5499493 |
| AGBL5 | protein_coding | -0.1805481 | 3.42192508 |
| MAPT-IT1 | lncRNA | -0.1805779 | 1.64623295 |
| GPR146 | protein_coding | -0.180583 | 0.37470427 |
| RPL38 | protein_coding | -0.1805892 | 5.65660794 |
| LINC02062 | lncRNA | -0.1806352 | 0.27442722 |
| RP11-249O6. | lncRNA | -0.1808429 | 0.47592497 |
| MRTFA | protein_coding | -0.1808908 | 3.18147964 |
| USP44 | protein_coding | -0.1809821 | 0.63222387 |
| KCNC1 | protein_coding | -0.1809919 | 1.96698939 |
| FICD | protein_coding | -0.1810752 | 1.62397334 |
| RPS25 | protein_coding | -0.1811001 | 6.71977537 |
| MOK | protein_coding | -0.1811652 | 0.65364703 |
| ACSF3 | protein_coding | -0.1815708 | 2.64469455 |
| C19orf47 | protein_coding | -0.181656 | 3.18237624 |
| ZNF486 | protein_coding | -0.1816837 | 1.98536738 |
| RBMX | protein_coding | -0.1816863 | 7.06772489 |
| C1orf122 | protein_coding | -0.1817128 | 1.79621725 |
| C4orf3 | protein_coding | -0.1817796 | 2.58211689 |
| RP11-44N22. | lncRNA | -0.1817918 | 0.22008153 |
| PEX11A | protein_coding | -0.1818747 | 0.99898943 |
| BRME1 | protein_coding | -0.1819605 | 3.05480339 |
| MAGIX | protein_coding | -0.1820793 | 0.54550465 |
| PCSK4 | protein_coding | -0.1820828 | 0.44785788 |
| RORB-AS1 | lncRNA | -0.1821111 | 0.26928037 |
| RP11-723O4. | lncRNA | -0.182133 | 0.51786521 |
| STRN | protein_coding | -0.1822509 | 3.78835671 |
| ATP6V0D1-D | lncRNA | -0.1824022 | 0.83748133 |
| SLC10A3 | protein_coding | -0.1824729 | 1.99011624 |
| SRGAP2C | protein_coding | -0.1825227 | 1.03443868 |
| FKBP8 | protein_coding | -0.1825442 | 5.20501549 |
| CDHR2 | protein_coding | -0.1826507 | 0.31074263 |
| HOXD1 | protein_coding | -0.1826854 | 1.12978945 |
| GTF2E2 | protein_coding | -0.1826952 | 2.12262898 |
| ZC3H14 | protein_coding | -0.1828185 | 5.11575659 |
| ZNF236 | protein_coding | -0.1828288 | 2.91774242 |
| NAB1 | protein_coding | -0.1828919 | 4.88649386 |
| RPL35 | protein_coding | -0.182997 | 7.05618672 |
| ALG2 | protein_coding | -0.1830615 | 3.39205377 |
| GTF3C5 | protein_coding | -0.1832164 | 4.93586896 |
| PKIB | protein_coding | -0.1834192 | 4.43554868 |

| | | | |
|---|---|---|---|
| SEMA3A | protein_coding | -0.1835105 | 2.64413513 |
| DYNLT2 | protein_coding | -0.1835548 | 0.32334494 |
| HERPUD2-AS | lncRNA | -0.1836395 | 2.52724607 |
| THADA | protein_coding | -0.1836758 | 3.19843917 |
| SDHA | protein_coding | -0.1837185 | 1.45262739 |
| RPL14 | protein_coding | -0.1838537 | 6.17618671 |
| RNF139-AS1 | lncRNA | -0.1838648 | 0.49863884 |
| LYPD6 | protein_coding | -0.1838939 | 2.8640545 |
| ZBTB6 | protein_coding | -0.1839871 | 1.98891478 |
| E4F1 | protein_coding | -0.1839898 | 2.13900279 |
| ARL5A | protein_coding | -0.1839919 | 1.79056175 |
| LCLAT1 | protein_coding | -0.1840111 | 3.39140772 |
| IER5L | protein_coding | -0.1840846 | 0.66924798 |
| SETD3 | protein_coding | -0.1841495 | 6.36541905 |
| HDDC2 | protein_coding | -0.1842112 | 2.25820724 |
| HOXA5 | protein_coding | -0.1843163 | 1.76895309 |
| TTK | protein_coding | -0.1843626 | 3.89984627 |
| CARD10 | protein_coding | -0.1843827 | 2.21454582 |
| PSKH1 | protein_coding | -0.1844048 | 3.16965091 |
| PPP1CA | protein_coding | -0.1845807 | 6.36840579 |
| POLR2B | protein_coding | -0.1846265 | 6.94753366 |
| LHFPL2 | protein_coding | -0.1846787 | 1.40566329 |
| BSG | protein_coding | -0.1846954 | 6.33445762 |
| SRRM1 | protein_coding | -0.1847802 | 5.60916365 |
| TMED10 | protein_coding | -0.1848231 | 5.3388815 |
| MOB2 | protein_coding | -0.1848506 | 3.13718832 |
| CYBC1 | protein_coding | -0.184962 | 2.59860195 |
| ENOPH1 | protein_coding | -0.1850715 | 5.99158049 |
| RP5-1142A6. | lncRNA | -0.1850838 | 0.50896399 |
| ACVR1B | protein_coding | -0.1850947 | 4.86156918 |
| NEO1 | protein_coding | -0.1854257 | 2.48913226 |
| EED | protein_coding | -0.1854302 | 3.8492523 |
| ISCA1 | protein_coding | -0.185501 | 3.35401746 |
| SELENOP | protein_coding | -0.1858588 | 0.17597721 |
| RPLP1 | protein_coding | -0.1859108 | 5.74482554 |
| AP1S1 | protein_coding | -0.185957 | 3.70030826 |
| PPIH | protein_coding | -0.1861075 | 3.33475079 |
| MPHOSPH6 | protein_coding | -0.1861459 | 3.42612478 |
| SCFD2 | protein_coding | -0.1861775 | 2.91593324 |
| CCDC115 | protein_coding | -0.1862174 | 3.80564552 |
| COX6A1 | protein_coding | -0.1862436 | 4.25497524 |
| SPTB | protein_coding | -0.1863184 | 3.97592734 |
| ECH1 | protein_coding | -0.1865697 | 3.65332148 |
| LINC00551 | lncRNA | -0.18657 | 0.20376881 |
| NDRG3 | protein_coding | -0.1866539 | 2.90903038 |

| | | | |
|---|---|---|---|
| EFHC1 | protein_coding | -0.1866589 | 2.36419288 |
| NEBL | protein_coding | -0.1867088 | 3.69025825 |
| MORC2 | protein_coding | -0.1867393 | 5.49338568 |
| MROH6 | protein_coding | -0.1867547 | 1.11701503 |
| IAH1 | protein_coding | -0.1867616 | 3.96454542 |
| LNPK | protein_coding | -0.1867752 | 4.00904915 |
| PGRMC1 | protein_coding | -0.1868596 | 4.33902701 |
| KLC1 | protein_coding | -0.1868625 | 2.97169273 |
| TTC39B | protein_coding | -0.1869098 | 2.16353834 |
| BNIPL | protein_coding | -0.1869378 | 0.34148503 |
| PLCXD2 | protein_coding | -0.1869382 | 2.76982578 |
| AKT2 | protein_coding | -0.1871687 | 6.42251296 |
| PLOD2 | protein_coding | -0.1872238 | 4.44465891 |
| ACOT8 | protein_coding | -0.1873241 | 3.15234192 |
| CTD-2371O3. | lncRNA | -0.1873336 | 0.64659331 |
| RP11-290D2. | lncRNA | -0.1874733 | 0.29545768 |
| RFC1 | protein_coding | -0.1876205 | 5.87215321 |
| ZBTB47 | protein_coding | -0.1876418 | 1.57600799 |
| WDR25 | protein_coding | -0.1876431 | 1.88972402 |
| RAB11B | protein_coding | -0.1876538 | 3.48513373 |
| TULP3 | protein_coding | -0.1876718 | 2.82050039 |
| MTIF3 | protein_coding | -0.1877271 | 2.89418863 |
| RP4-733M16 | lncRNA | -0.1877943 | 2.22297406 |
| NKPD1 | protein_coding | -0.1879017 | 0.22100483 |
| CCDC153 | protein_coding | -0.1880158 | 0.47002695 |
| WDR47 | protein_coding | -0.1881223 | 3.35675864 |
| LINC00526 | lncRNA | -0.1882513 | 1.49140126 |
| MOCS3 | protein_coding | -0.1883061 | 1.76611431 |
| FARP2 | protein_coding | -0.1883355 | 4.24157984 |
| COA3 | protein_coding | -0.188507 | 2.88591375 |
| ZNF584 | protein_coding | -0.1885242 | 2.89410401 |
| THAP7-AS1 | lncRNA | -0.1886551 | 0.99829926 |
| MTERF1 | protein_coding | -0.1886833 | 2.07695303 |
| THAP11 | protein_coding | -0.1886945 | 3.58425917 |
| STX7 | protein_coding | -0.1887182 | 1.85859145 |
| TTC7B | protein_coding | -0.1888929 | 3.76922684 |
| HYKK | protein_coding | -0.1889296 | 0.64012335 |
| PPP2R2C | protein_coding | -0.1892682 | 5.7740542 |
| PPP1R8 | protein_coding | -0.1892734 | 4.83067496 |
| SYTL2 | protein_coding | -0.1894057 | 5.07940667 |
| FAM133B | protein_coding | -0.1895118 | 1.92848647 |
| BSCL2 | protein_coding | -0.189641 | 1.51502057 |
| SMC1B | protein_coding | -0.1897448 | 0.25820853 |
| DCTPP1 | protein_coding | -0.1897889 | 5.57002646 |
| TRUB2 | protein_coding | -0.1898135 | 3.56186966 |

| | | | |
|---|---|---|---|
| KASH5 | protein_coding | -0.1900471 | 0.30862496 |
| CFAP20DC | protein_coding | -0.190138 | 1.75675083 |
| ADSS2 | protein_coding | -0.1902716 | 2.87082071 |
| PPP1R14B-A! | lncRNA | -0.190295 | 0.47242304 |
| PLPP5 | protein_coding | -0.1903933 | 2.73244604 |
| PDXP | protein_coding | -0.1904171 | 0.2410974 |
| GDI2 | protein_coding | -0.1905136 | 5.95406271 |
| SMIM7 | protein_coding | -0.19095 | 3.73068813 |
| ELOF1 | protein_coding | -0.1909972 | 4.12794302 |
| GCAT | protein_coding | -0.1910075 | 2.46562922 |
| AKAP5 | protein_coding | -0.1911205 | 3.51133983 |
| MCM10 | protein_coding | -0.1911256 | 3.01631105 |
| NFYC | protein_coding | -0.1911829 | 5.16163568 |
| ATP6V1H | protein_coding | -0.1911941 | 2.71774972 |
| KIDINS220 | protein_coding | -0.1913691 | 6.60197886 |
| NDUFB6 | protein_coding | -0.1914925 | 4.17349797 |
| SFT2D3 | protein_coding | -0.1915371 | 2.10986127 |
| HNRNPA3 | protein_coding | -0.1916547 | 7.24211779 |
| PPIE | protein_coding | -0.191693 | 4.98762115 |
| H2BC9 | protein_coding | -0.1917136 | 0.2536758 |
| NASP | protein_coding | -0.1918002 | 7.95558767 |
| IL10RB | protein_coding | -0.1918671 | 0.32905812 |
| MAF1 | protein_coding | -0.1918947 | 3.99337104 |
| SKAP2 | protein_coding | -0.1922211 | 3.65112198 |
| RXYLT1 | protein_coding | -0.1923363 | 3.51915816 |
| KDELR2 | protein_coding | -0.1924648 | 5.16714939 |
| OGFOD2 | protein_coding | -0.192513 | 0.99245378 |
| RP11-77P6.2 | lncRNA | -0.1927602 | 0.46455794 |
| OCIAD2 | protein_coding | -0.1927855 | 4.67426237 |
| DBI | protein_coding | -0.1927932 | 4.73801866 |
| C2orf88 | protein_coding | -0.1928789 | 0.40432977 |
| RP11-679B1! | lncRNA | -0.1928816 | 0.29509328 |
| LAMTOR4 | protein_coding | -0.1929635 | 4.08579903 |
| STOX2 | protein_coding | -0.193011 | 3.77505874 |
| BEX1 | protein_coding | -0.1930578 | 5.46805485 |
| CEP89 | protein_coding | -0.1933455 | 2.97600732 |
| SGCB | protein_coding | -0.193434 | 4.48055575 |
| LINC01311 | lncRNA | -0.1934523 | 0.4837837 |
| CACNA1I | protein_coding | -0.1934537 | 1.62489278 |
| DPCD | protein_coding | -0.1934899 | 0.76776789 |
| DDX23 | protein_coding | -0.193561 | 6.83449295 |
| WDFY3-AS2 | lncRNA | -0.1937005 | 0.62464738 |
| DCAF7 | protein_coding | -0.193771 | 7.67030739 |
| CYP26B1 | protein_coding | -0.1937916 | 2.70773388 |
| ANKRD42 | protein_coding | -0.1938481 | 0.63677395 |

| | | | |
|---|---|---|---|
| SORCS2 | protein_coding | -0.1938777 | 1.1121345 |
| ADAT3 | protein_coding | -0.1940313 | 0.29492153 |
| KIFBP | protein_coding | -0.1940763 | 2.06974553 |
| PRIM1 | protein_coding | -0.1942183 | 3.55170392 |
| USP3-AS1 | lncRNA | -0.1942353 | 0.96304293 |
| OGA | protein_coding | -0.1942393 | 5.48875364 |
| RP11-147L13 | lncRNA | -0.1943119 | 0.78519081 |
| DPM2 | protein_coding | -0.194405 | 4.53111789 |
| RFC4 | protein_coding | -0.1944552 | 6.26698277 |
| ZFYVE27 | protein_coding | -0.1945804 | 3.11583505 |
| RP11-455F5. | lncRNA | -0.1946751 | 0.90627776 |
| QDPR | protein_coding | -0.1947911 | 4.56793751 |
| TIGD2 | protein_coding | -0.1948187 | 2.08387168 |
| CCT7 | protein_coding | -0.1948689 | 7.43296339 |
| GDI1 | protein_coding | -0.1948769 | 7.2189491 |
| MRM3 | protein_coding | -0.1948956 | 1.99110323 |
| NYAP1 | protein_coding | -0.1949479 | 0.90121058 |
| GLCE | protein_coding | -0.1950503 | 5.48741209 |
| ABHD17A | protein_coding | -0.1950742 | 2.07357789 |
| DNAJC30 | protein_coding | -0.1951294 | 0.99015571 |
| ZNF324 | protein_coding | -0.195217 | 3.003445 |
| TKFC | protein_coding | -0.195281 | 4.51027192 |
| EXOC6 | protein_coding | -0.1953274 | 2.3014199 |
| SERPINI1 | protein_coding | -0.1953567 | 4.09533802 |
| DDX49 | protein_coding | -0.195361 | 4.04915614 |
| ENO2 | protein_coding | -0.1953797 | 7.85632741 |
| GARNL3 | protein_coding | -0.1954237 | 1.08641189 |
| CES4A | protein_coding | -0.1954357 | 0.89591657 |
| SUMO3 | protein_coding | -0.1954402 | 6.07478984 |
| SLC12A5 | protein_coding | -0.1955957 | 1.44551901 |
| CTD-2083E4. | lncRNA | -0.1956112 | 0.16003349 |
| U91328.19 | lncRNA | -0.1956893 | 1.07955668 |
| AC006277.2 | lncRNA | -0.1957426 | 0.58011558 |
| FBXO41 | protein_coding | -0.1957527 | 2.62506751 |
| CABYR | protein_coding | -0.1958622 | 0.43738431 |
| BICDL1 | protein_coding | -0.1961014 | 4.68588521 |
| ADIPOR1 | protein_coding | -0.1961185 | 6.37186392 |
| PLCG1 | protein_coding | -0.1962692 | 6.79728906 |
| KIFC2 | protein_coding | -0.1963492 | 4.36215186 |
| EIF3F | protein_coding | -0.196398 | 5.94077916 |
| QPCTL | protein_coding | -0.1964662 | 1.76525217 |
| WDR20 | protein_coding | -0.1964868 | 4.21708012 |
| ZNF707 | protein_coding | -0.1965554 | 1.50349762 |
| WDR92 | protein_coding | -0.1965804 | 1.09917968 |
| HNRNPA1 | protein_coding | -0.196748 | 8.49333958 |

| | | | |
|---|---|---|---|
| URB1-AS1 | lncRNA | -0.1968337 | 1.02400124 |
| ANTXR2 | protein_coding | -0.196834 | 1.7421873 |
| LLGL2 | protein_coding | -0.1970367 | 3.55552536 |
| SMC5 | protein_coding | -0.1970851 | 5.65012206 |
| SLC25A14 | protein_coding | -0.1971212 | 1.12548937 |
| ZNF610 | protein_coding | -0.197239 | 0.71273736 |
| BRD3OS | protein_coding | -0.1972413 | 2.56236221 |
| RAB4A | protein_coding | -0.1972438 | 2.73032721 |
| DCAF6 | protein_coding | -0.1972945 | 4.31127164 |
| TPPP | protein_coding | -0.1975076 | 0.98235758 |
| PAFAH2 | protein_coding | -0.1975255 | 1.92769093 |
| TACC3 | protein_coding | -0.1976019 | 5.84800182 |
| ELAC2 | protein_coding | -0.1976087 | 4.33442648 |
| RTL8B | protein_coding | -0.1977655 | 2.06793753 |
| NF2 | protein_coding | -0.1977828 | 3.4883601 |
| RCN2 | protein_coding | -0.1978254 | 5.19305651 |
| B3GALT6 | protein_coding | -0.1979177 | 1.99270873 |
| DAGLB | protein_coding | -0.1979364 | 3.24919985 |
| RP11-546B8. | lncRNA | -0.1980633 | 3.16256447 |
| PARAL1 | lncRNA | -0.1981004 | 0.44420492 |
| ZMYND11 | protein_coding | -0.1981235 | 3.78052909 |
| MFSD13A | protein_coding | -0.1981942 | 3.16268583 |
| RPS24 | protein_coding | -0.1983067 | 7.40616152 |
| HPS6 | protein_coding | -0.1983744 | 1.72675716 |
| MRPS12 | protein_coding | -0.1984054 | 3.5802824 |
| RNF145 | protein_coding | -0.1984762 | 5.95156007 |
| HNRNPA0 | protein_coding | -0.1989393 | 6.17594483 |
| IFITM10 | protein_coding | -0.1990236 | 0.29979269 |
| PARVG | protein_coding | -0.1991367 | 0.92617061 |
| CASP3 | protein_coding | -0.1991747 | 3.91917916 |
| RPL37A | protein_coding | -0.1992866 | 8.09752556 |
| RBM4 | protein_coding | -0.1994109 | 2.77058889 |
| L2HGDH | protein_coding | -0.1994436 | 4.06247414 |
| CPT2 | protein_coding | -0.199548 | 3.41480014 |
| FSD1 | protein_coding | -0.1995702 | 4.65653139 |
| GLUD1 | protein_coding | -0.199617 | 4.90125106 |
| CMBL | protein_coding | -0.1996189 | 3.15520561 |
| ATP6V1C1 | protein_coding | -0.1996264 | 3.7908657 |
| ZBED8 | protein_coding | -0.1996293 | 2.11974435 |
| MPV17L2 | protein_coding | -0.1998125 | 4.03901017 |
| PEMT | protein_coding | -0.199857 | 1.82615659 |
| ANAPC13 | protein_coding | -0.1999358 | 4.8009939 |
| MIR9-3HG | lncRNA | -0.2000337 | 2.30119683 |
| NR2F1-AS1 | lncRNA | -0.200097 | 0.29872072 |
| ATP11A | protein_coding | -0.2001192 | 6.06701069 |

| | | | |
|---|---|---|---|
| DHX15 | protein_coding | -0.2001988 | 7.47959453 |
| RASA3 | protein_coding | -0.2002448 | 0.35076889 |
| MCM5 | protein_coding | -0.2002584 | 5.74080872 |
| RALB | protein_coding | -0.2002858 | 1.49511573 |
| LRFN1 | protein_coding | -0.2002899 | 0.44818994 |
| CTNNA3 | protein_coding | -0.2004537 | 1.04225385 |
| FXR1 | protein_coding | -0.2005307 | 6.96131275 |
| MRTO4 | protein_coding | -0.2005608 | 4.81899255 |
| ZNF346 | protein_coding | -0.2006172 | 3.33635132 |
| RP11-522I20 | lncRNA | -0.2006379 | 0.77277116 |
| RPL35A | protein_coding | -0.2006843 | 7.48661114 |
| TRMT61A | protein_coding | -0.2007368 | 3.63172715 |
| GAS2L3 | protein_coding | -0.2008673 | 2.97158484 |
| ELK4 | protein_coding | -0.2008869 | 3.82520858 |
| LRP3 | protein_coding | -0.2009353 | 2.44103042 |
| ZNF670 | protein_coding | -0.2010034 | 3.01028325 |
| PIN4 | protein_coding | -0.201008 | 3.55184163 |
| RPS18 | protein_coding | -0.2011575 | 8.57762445 |
| SUV39H2-DT | lncRNA | -0.2011869 | 0.4220854 |
| GPN3 | protein_coding | -0.2012631 | 3.92083574 |
| MMS19 | protein_coding | -0.2013037 | 4.47897612 |
| FYN | protein_coding | -0.2013586 | 6.27207365 |
| FBLN7 | protein_coding | -0.2013871 | 3.71904478 |
| TMX1 | protein_coding | -0.2013972 | 6.29516175 |
| SPATA7 | protein_coding | -0.2015435 | 0.79927162 |
| EVL | protein_coding | -0.2016208 | 6.21973316 |
| RP11-355O1 | lncRNA | -0.2016455 | 0.61809023 |
| BOLA2B | protein_coding | -0.2016665 | 0.2923762 |
| RIC8A | protein_coding | -0.2017199 | 6.32647982 |
| SPC24 | protein_coding | -0.2017208 | 2.90696637 |
| MRPL30 | protein_coding | -0.2017322 | 4.63631328 |
| MORN1 | protein_coding | -0.2019838 | 0.88397664 |
| RYR1 | protein_coding | -0.2020813 | 2.267973 |
| KBTBD6 | protein_coding | -0.2021892 | 2.84226392 |
| CALCOCO1 | protein_coding | -0.2021936 | 3.50345541 |
| CCDC87 | protein_coding | -0.2023102 | 0.26215798 |
| CCAR2 | protein_coding | -0.2023126 | 4.55412758 |
| SF3A2 | protein_coding | -0.2023358 | 3.87094141 |
| MFSD1 | protein_coding | -0.2023814 | 3.0924242 |
| UQCC2 | protein_coding | -0.2024403 | 5.47761374 |
| ALKBH7 | protein_coding | -0.2024776 | 1.34467718 |
| TOR2A | protein_coding | -0.2024804 | 2.5745734 |
| CNNM3 | protein_coding | -0.202493 | 4.59732985 |
| LINC01023 | lncRNA | -0.2025312 | 0.31013961 |
| NUP43 | protein_coding | -0.2027381 | 2.93251366 |

| | | | |
|---|---|---|---|
| PGRMC2 | protein_coding | -0.202774 | 4.29437674 |
| ARPP19 | protein_coding | -0.2027982 | 5.78402441 |
| POLR2D | protein_coding | -0.2028075 | 5.10127226 |
| HECTD3 | protein_coding | -0.2029307 | 4.98014388 |
| TUBGCP2 | protein_coding | -0.203011 | 3.50570288 |
| SRMS | protein_coding | -0.2030539 | 0.65437652 |
| RPS14 | protein_coding | -0.2030691 | 7.62476668 |
| DNM2 | protein_coding | -0.2031064 | 5.79574443 |
| FASN | protein_coding | -0.2031132 | 4.23883297 |
| NTAQ1 | protein_coding | -0.2032679 | 3.38231853 |
| CACNA1B | protein_coding | -0.203301 | 3.98272018 |
| RP11-477D19 | lncRNA | -0.2033339 | 0.61517322 |
| BMT2 | protein_coding | -0.2034211 | 2.24470754 |
| ZDHHC16 | protein_coding | -0.203914 | 2.07469295 |
| ANKZF1 | protein_coding | -0.2039457 | 3.50778387 |
| CBLB | protein_coding | -0.2041021 | 4.54188694 |
| ALDH16A1 | protein_coding | -0.2041428 | 1.74649552 |
| SLC38A10 | protein_coding | -0.204278 | 3.22813261 |
| PDP1 | protein_coding | -0.2043484 | 2.81922588 |
| KLHL25 | protein_coding | -0.2044745 | 2.57956791 |
| ZBTB34 | protein_coding | -0.2045264 | 2.88533774 |
| DVL1 | protein_coding | -0.2045925 | 3.32953569 |
| TMEM14C | protein_coding | -0.2046144 | 3.74626799 |
| SLC25A16 | protein_coding | -0.2046293 | 1.93819595 |
| CTDSPL2 | protein_coding | -0.204658 | 5.15494303 |
| SERTAD3 | protein_coding | -0.2046905 | 2.19111652 |
| TMCO1 | protein_coding | -0.2047404 | 5.70355227 |
| SDHAF3 | protein_coding | -0.2047952 | 2.52359879 |
| ARMC1 | protein_coding | -0.2048073 | 2.88094174 |
| SPOPL | protein_coding | -0.2048666 | 3.04077487 |
| RALGPS2 | protein_coding | -0.2049321 | 0.88118868 |
| SLC7A6 | protein_coding | -0.204937 | 6.4042072 |
| KIF22 | protein_coding | -0.20503 | 6.1150735 |
| RP11-46H11. | lncRNA | -0.2050855 | 0.25398002 |
| RPS3 | protein_coding | -0.2053109 | 8.68758741 |
| PKIA | protein_coding | -0.2053809 | 3.29303288 |
| RPL9 | protein_coding | -0.2054003 | 6.84603948 |
| NPAS4 | protein_coding | -0.2055544 | 0.32282949 |
| NCK1-DT | lncRNA | -0.2055686 | 1.1782644 |
| RSPH4A | protein_coding | -0.2055747 | 0.21995708 |
| AP000487.5 | lncRNA | -0.2056141 | 0.38633444 |
| UNC119B | protein_coding | -0.2056365 | 5.81281014 |
| SLC9A3-AS1 | lncRNA | -0.2056585 | 3.46788582 |
| HMCES | protein_coding | -0.2056852 | 5.0993696 |
| PRDX6 | protein_coding | -0.2057401 | 4.52999406 |

| | | | |
|---|---|---|---|
| MTFR1L | protein_coding | -0.2057767 | 4.94766746 |
| RP5-1142A6. | lncRNA | -0.2057793 | 0.22753143 |
| LAMTOR3 | protein_coding | -0.2059679 | 3.37982193 |
| EIF1B | protein_coding | -0.2061508 | 3.25670608 |
| CSTF1 | protein_coding | -0.206151 | 3.88650045 |
| TPT1 | protein_coding | -0.2062404 | 8.04635037 |
| TMEM248 | protein_coding | -0.2064647 | 5.64723867 |
| MYT1 | protein_coding | -0.2065099 | 4.28966419 |
| MCM3 | protein_coding | -0.2065149 | 8.25843327 |
| ARNTL | protein_coding | -0.2067014 | 4.0180032 |
| AC002116.7 | lncRNA | -0.2067461 | 0.61371473 |
| SOX30 | protein_coding | -0.2068223 | 0.19461977 |
| UBE2V2 | protein_coding | -0.2069377 | 3.35783583 |
| YBEY | protein_coding | -0.2069554 | 1.53220436 |
| CCNA2 | protein_coding | -0.2070634 | 5.08091346 |
| DDN | protein_coding | -0.2071015 | 3.77387767 |
| FAU | protein_coding | -0.2071666 | 6.29330336 |
| ARL6IP1 | protein_coding | -0.2072144 | 5.25393049 |
| AC137932.4 | lncRNA | -0.2072303 | 0.46545636 |
| PABPC1 | protein_coding | -0.207243 | 7.43556932 |
| FBXL16 | protein_coding | -0.2073829 | 3.43851583 |
| PRDX2 | protein_coding | -0.2074256 | 8.80123635 |
| MAPK12 | protein_coding | -0.2075271 | 1.69968048 |
| RP11-284M1 | lncRNA | -0.2077182 | 0.23110867 |
| CTD-3185P2. | lncRNA | -0.207719 | 0.60658811 |
| RMC1 | protein_coding | -0.2077532 | 2.8989253 |
| RBM12 | protein_coding | -0.207854 | 6.01990484 |
| TMEM237 | protein_coding | -0.2079066 | 3.71636767 |
| PDCL | protein_coding | -0.208056 | 4.1390366 |
| COX5B | protein_coding | -0.2080575 | 4.51153164 |
| RP11-438B2: | lncRNA | -0.2081254 | 0.72252484 |
| VEGFB | protein_coding | -0.2081776 | 3.81869857 |
| DSCC1 | protein_coding | -0.2082237 | 3.12660982 |
| CENPI | protein_coding | -0.2082277 | 2.96021492 |
| C17orf100 | protein_coding | -0.2082637 | 0.92906037 |
| METTL4 | protein_coding | -0.2082927 | 3.21515895 |
| CDCA2 | protein_coding | -0.2083025 | 2.64478656 |
| COPS7A | protein_coding | -0.2083768 | 3.89178404 |
| TOMM40 | protein_coding | -0.2085445 | 6.43915035 |
| PPCDC | protein_coding | -0.2086199 | 2.74457425 |
| THYN1 | protein_coding | -0.2086669 | 3.16164611 |
| SV2A | protein_coding | -0.2088643 | 4.69207949 |
| TNFRSF25 | protein_coding | -0.2088693 | 1.57340445 |
| RP11-212P7. | lncRNA | -0.2090479 | 2.2407024 |
| PGAP2 | protein_coding | -0.2091568 | 3.44116162 |

| | | | |
|---|---|---|---|
| KIAA1614-AS | lncRNA | -0.2091955 | 0.4697952 |
| B3GALT4 | protein_coding | -0.2092061 | 0.36706945 |
| CACHD1 | protein_coding | -0.2093296 | 4.71452301 |
| SSR3 | protein_coding | -0.2094584 | 6.45531358 |
| CCDC71 | protein_coding | -0.2094838 | 1.80425704 |
| MANSC1 | protein_coding | -0.2095357 | 3.59609272 |
| POM121 | protein_coding | -0.2096016 | 4.93596434 |
| SMG9 | protein_coding | -0.2096547 | 4.9751314 |
| WDR7 | protein_coding | -0.209787 | 4.75454372 |
| C16orf95-DT | lncRNA | -0.2098028 | 0.29325509 |
| PAFAH1B2 | protein_coding | -0.2098389 | 6.39566117 |
| DIRAS1 | protein_coding | -0.2098534 | 5.98541482 |
| TMEM243 | protein_coding | -0.2099658 | 1.14647721 |
| NDUFB7 | protein_coding | -0.2100397 | 4.2074207 |
| RP11-295D4. | lncRNA | -0.2100581 | 0.53357087 |
| PCGF2 | protein_coding | -0.210446 | 4.98796946 |
| PABIR2 | protein_coding | -0.2104626 | 4.07676825 |
| PTBP1 | protein_coding | -0.2104859 | 5.62894576 |
| ACER2 | protein_coding | -0.2104868 | 1.43987026 |
| SLC29A1 | protein_coding | -0.2105476 | 6.94731967 |
| EIF1AD | protein_coding | -0.2105731 | 3.58957445 |
| STK11IP | protein_coding | -0.2109188 | 2.88712109 |
| PPP1R14B | protein_coding | -0.2109505 | 5.53192271 |
| TMEM203 | protein_coding | -0.2109846 | 3.39368205 |
| NCKIPSD | protein_coding | -0.211046 | 3.4673433 |
| POLR2K | protein_coding | -0.211087 | 4.03302555 |
| CHST9 | protein_coding | -0.211108 | 4.25467697 |
| GTSE1 | protein_coding | -0.2111747 | 4.5959168 |
| ALG1 | protein_coding | -0.2112451 | 3.2782966 |
| FAM120C | protein_coding | -0.2113222 | 2.46296105 |
| EIF3H | protein_coding | -0.2113366 | 7.78501151 |
| TPX2 | protein_coding | -0.2114248 | 7.3083849 |
| ZNF101 | protein_coding | -0.2114518 | 2.31868865 |
| SKA2 | protein_coding | -0.2114905 | 4.69389671 |
| NCAPD3 | protein_coding | -0.211497 | 6.2066655 |
| CD99L2 | protein_coding | -0.2114983 | 3.87727018 |
| PADI3 | protein_coding | -0.2115106 | 0.28548321 |
| RP11-368I7.6 | lncRNA | -0.2115322 | 0.75722208 |
| PRDM13 | protein_coding | -0.2116056 | 2.46781155 |
| KIF23 | protein_coding | -0.2116933 | 5.34050435 |
| CUEDC2 | protein_coding | -0.2117248 | 3.8740338 |
| TCTN2 | protein_coding | -0.2117254 | 0.90505477 |
| RP11-363H1? | lncRNA | -0.2118472 | 0.25652402 |
| ZC3H4 | protein_coding | -0.2118603 | 4.42908933 |
| SMARCA2 | protein_coding | -0.2118869 | 5.56704408 |

| | | | |
|---|---|---|---|
| RNF227 | protein_coding | -0.211911 | 0.94889871 |
| ENKUR | protein_coding | -0.2119188 | 0.30061272 |
| CEL | protein_coding | -0.2119264 | 0.69684104 |
| KIF5C | protein_coding | -0.2119477 | 6.77219865 |
| SUN1 | protein_coding | -0.2121248 | 7.87131972 |
| ZBTB8A | protein_coding | -0.2121411 | 1.69576227 |
| PTER | protein_coding | -0.2122024 | 2.74471435 |
| RP11-1148L6 | lncRNA | -0.2122163 | 0.31921766 |
| ACAD10 | protein_coding | -0.2122364 | 3.92697663 |
| PPP1R3E | protein_coding | -0.2124638 | 4.86646332 |
| ATP6V1E2 | protein_coding | -0.2127629 | 2.45526489 |
| SKP1 | protein_coding | -0.2128321 | 6.10293425 |
| SMARCAD1 | protein_coding | -0.2130915 | 5.40104742 |
| SMURF2 | protein_coding | -0.2131145 | 4.15126566 |
| ATL3 | protein_coding | -0.2132259 | 5.08593922 |
| DBNDD1 | protein_coding | -0.2132616 | 4.05905205 |
| CDK8 | protein_coding | -0.2133318 | 4.35390536 |
| UBL7-AS1 | lncRNA | -0.2133409 | 1.01578246 |
| PAN3 | protein_coding | -0.2137114 | 3.79529628 |
| HSD17B12 | protein_coding | -0.2137939 | 2.07920842 |
| PLEK2 | protein_coding | -0.2138061 | 0.85397785 |
| CHRNA5 | protein_coding | -0.2138087 | 3.32076137 |
| CDC7 | protein_coding | -0.2139556 | 4.9175033 |
| PREPL | protein_coding | -0.2139981 | 6.02425909 |
| WIPI2 | protein_coding | -0.2140838 | 4.98885206 |
| TMF1 | protein_coding | -0.2140874 | 2.64317578 |
| TOPBP1 | protein_coding | -0.214091 | 6.25557723 |
| PDLIM7 | protein_coding | -0.2141093 | 1.35196627 |
| LDB1 | protein_coding | -0.2141404 | 4.9849811 |
| RALGAPA2 | protein_coding | -0.2145693 | 4.08166946 |
| CD79A | protein_coding | -0.214769 | 0.41669919 |
| KDM2B-DT | lncRNA | -0.2147774 | 0.36987365 |
| DNMBP | protein_coding | -0.2150805 | 2.91882289 |
| ORC1 | protein_coding | -0.2150925 | 3.74767628 |
| SCP2 | protein_coding | -0.215133 | 5.21520287 |
| FUNDC2 | protein_coding | -0.2151622 | 4.82407096 |
| MAP3K21 | protein_coding | -0.2152737 | 2.92649598 |
| CTBP1-DT | lncRNA | -0.2152816 | 2.48884585 |
| ITPKA | protein_coding | -0.2152904 | 1.22053405 |
| DDX6 | protein_coding | -0.2154131 | 7.1901992 |
| URM1 | protein_coding | -0.2156123 | 3.75783024 |
| NUCKS1 | protein_coding | -0.2156339 | 7.20790579 |
| TBCCD1 | protein_coding | -0.2156562 | 3.72774771 |
| RP11-46A10. | lncRNA | -0.2157451 | 0.45306497 |
| KITLG | protein_coding | -0.2157678 | 2.48769635 |

| | | | |
|---|---|---|---|
| RPLP2 | protein_coding | -0.2158523 | 5.14577929 |
| LHPP | protein_coding | -0.215918 | 1.87926441 |
| BCL2L1 | protein_coding | -0.2161746 | 7.60223979 |
| TMEM151B | protein_coding | -0.2162192 | 2.93049319 |
| KCNA5 | protein_coding | -0.2163119 | 0.75873791 |
| BUD23 | protein_coding | -0.2163897 | 5.95497662 |
| NDFIP2 | protein_coding | -0.2166359 | 4.83815737 |
| LPAR2 | protein_coding | -0.2166729 | 2.4283776 |
| RIOX1 | protein_coding | -0.2166943 | 3.93838395 |
| PABPC4 | protein_coding | -0.2167283 | 5.67984934 |
| INO80B | protein_coding | -0.2167438 | 0.24262926 |
| ARL15 | protein_coding | -0.2167444 | 1.60939042 |
| PDXDC2P-NP | lncRNA | -0.2170467 | 0.60169356 |
| SLC16A13 | protein_coding | -0.2170831 | 0.66778672 |
| WFS1 | protein_coding | -0.2171004 | 2.32672042 |
| CHERP | protein_coding | -0.2171572 | 4.98792171 |
| EPB41L4B | protein_coding | -0.2172601 | 1.40627699 |
| CTD-2005H7. | lncRNA | -0.2172693 | 0.22233601 |
| ZNF672 | protein_coding | -0.2173891 | 3.30978754 |
| DNAJC27-AS | lncRNA | -0.2174172 | 0.42950097 |
| PRDX3 | protein_coding | -0.217477 | 5.83077134 |
| FAM131A | protein_coding | -0.2176585 | 3.85362809 |
| OSGEPL1 | protein_coding | -0.2176865 | 1.55537589 |
| OSER1-DT | lncRNA | -0.2177629 | 1.15517902 |
| SARS2 | protein_coding | -0.2177859 | 1.43312183 |
| GSE1 | protein_coding | -0.2179211 | 8.23162296 |
| RPS7 | protein_coding | -0.2180417 | 7.75448928 |
| SLC25A30 | protein_coding | -0.2181312 | 1.38981227 |
| MRPL23 | protein_coding | -0.218175 | 2.15320228 |
| DCUN1D2 | protein_coding | -0.2182732 | 3.29553818 |
| CYB561A3 | protein_coding | -0.2183576 | 3.67170604 |
| TSEN54 | protein_coding | -0.2183933 | 3.65843797 |
| SPEN | protein_coding | -0.2184487 | 7.2246885 |
| PFKP | protein_coding | -0.2184538 | 5.778675 |
| STRA6LP | lncRNA | -0.2185451 | 0.32231047 |
| ZFP28 | protein_coding | -0.2185777 | 1.80887226 |
| RP11-594N1! | lncRNA | -0.2185894 | 0.29704603 |
| INAFM1 | protein_coding | -0.2185934 | 1.10960107 |
| BACE1 | protein_coding | -0.2186083 | 6.50823916 |
| RP11-372K1⁄ | lncRNA | -0.2187226 | 0.44540415 |
| CDC34 | protein_coding | -0.2188006 | 4.13611314 |
| DDOST | protein_coding | -0.2188796 | 7.99148849 |
| TMEM14B-D | lncRNA | -0.2189979 | 0.46991343 |
| SND1 | protein_coding | -0.2190427 | 7.71843793 |
| MED4 | protein_coding | -0.2190573 | 3.22489187 |

| | | | |
|---|---|---|---|
| CAPN15 | protein_coding | -0.2191844 | 2.12126263 |
| TMEM25 | protein_coding | -0.2192042 | 3.09624765 |
| AFMID | protein_coding | -0.2192336 | 4.24743998 |
| NKIRAS2 | protein_coding | -0.2195052 | 5.89626498 |
| RTL9 | protein_coding | -0.2196627 | 0.27205464 |
| ZNF446 | protein_coding | -0.219713 | 3.08863842 |
| LMTK2 | protein_coding | -0.2197195 | 4.76542272 |
| POM121C | protein_coding | -0.2197295 | 4.89496756 |
| RADIL | protein_coding | -0.2198318 | 2.69431611 |
| ZNF324B | protein_coding | -0.2200233 | 2.39926119 |
| SPCS2 | protein_coding | -0.2202175 | 5.06692567 |
| APH1B | protein_coding | -0.2202737 | 2.57964202 |
| RAPGEF6 | protein_coding | -0.2202872 | 1.7246959 |
| NNT | protein_coding | -0.2202942 | 5.85691148 |
| BUB3 | protein_coding | -0.2203456 | 5.92772971 |
| FHDC1 | protein_coding | -0.2206486 | 4.3849165 |
| CENPB | protein_coding | -0.220978 | 4.92248135 |
| HACD3 | protein_coding | -0.2211072 | 7.02471426 |
| ABHD14B | protein_coding | -0.2211794 | 2.98364777 |
| TTYH3 | protein_coding | -0.2212531 | 4.87030502 |
| HIRIP3 | protein_coding | -0.2213499 | 4.22971245 |
| RTN2 | protein_coding | -0.2214492 | 3.01457064 |
| TRABD | protein_coding | -0.2217541 | 2.35001509 |
| SS18L1 | protein_coding | -0.2218062 | 5.68354505 |
| LHFPL5 | protein_coding | -0.2218278 | 4.15378737 |
| PGBD1 | protein_coding | -0.2218497 | 1.54197348 |
| DDB1 | protein_coding | -0.2219248 | 8.7070859 |
| HOXA7 | protein_coding | -0.2220221 | 5.20036246 |
| RP11-34F20. | lncRNA | -0.2220651 | 0.7703596 |
| RPL6 | protein_coding | -0.2221111 | 8.37485568 |
| AP000251.3 | lncRNA | -0.2221374 | 1.05366214 |
| TRIM28 | protein_coding | -0.2222687 | 7.48303066 |
| RP11-454E5. | lncRNA | -0.2224717 | 0.7264672 |
| RPS21 | protein_coding | -0.2224897 | 8.72680072 |
| FAM110A | protein_coding | -0.2226278 | 1.9975167 |
| OBSCN-AS1 | lncRNA | -0.2228062 | 0.93034537 |
| MYLK-AS1 | lncRNA | -0.222859 | 0.54947753 |
| NT5DC3 | protein_coding | -0.2229771 | 5.01448018 |
| PMM1 | protein_coding | -0.223005 | 4.02811311 |
| HMGB1 | protein_coding | -0.2230432 | 7.40127503 |
| MED31 | protein_coding | -0.2230474 | 1.12391288 |
| RPL3 | protein_coding | -0.2230596 | 9.57405912 |
| VEGFA | protein_coding | -0.2231195 | 6.58405836 |
| VCAN | protein_coding | -0.2231574 | 1.1885665 |
| MFSD9 | protein_coding | -0.2232975 | 4.02903438 |

| | | | |
|---|---|---|---|
| ARG2 | protein_coding | -0.223317 | 3.06135995 |
| BAX | protein_coding | -0.2233741 | 5.42808494 |
| PARS2 | protein_coding | -0.2233997 | 1.89451783 |
| KTI12 | protein_coding | -0.223475 | 1.4744144 |
| ERLEC1 | protein_coding | -0.2234788 | 3.87403495 |
| FTO | protein_coding | -0.2236285 | 5.37069397 |
| RP1-267D11. | lncRNA | -0.2237041 | 1.81740546 |
| FITM2 | protein_coding | -0.2238353 | 1.4736481 |
| MKI67 | protein_coding | -0.2238628 | 8.11572527 |
| MCM3AP-AS | lncRNA | -0.2238823 | 2.99312933 |
| MSH6 | protein_coding | -0.2240135 | 8.08084919 |
| KLHL36 | protein_coding | -0.2240721 | 5.50992628 |
| MMGT1 | protein_coding | -0.2241092 | 4.63119237 |
| SNHG14 | lncRNA | -0.2241134 | 7.20006381 |
| EBF4 | protein_coding | -0.2242557 | 0.37839634 |
| TBC1D12 | protein_coding | -0.2242559 | 2.30264753 |
| PUM2 | protein_coding | -0.2242804 | 6.04035703 |
| GNAS | protein_coding | -0.2245004 | 9.76550566 |
| LINC-PINT | lncRNA | -0.2245931 | 0.65947721 |
| SNRNP48 | protein_coding | -0.2247327 | 3.77868361 |
| FAAP24 | protein_coding | -0.2247971 | 1.66504288 |
| SNRPA | protein_coding | -0.2248037 | 5.85237674 |
| ARAP1 | protein_coding | -0.2248473 | 4.93072948 |
| LIPA | protein_coding | -0.2250175 | 4.80417923 |
| ANKRD18A | protein_coding | -0.2250371 | 3.02816734 |
| NSF | protein_coding | -0.2250676 | 5.4938475 |
| WAS | protein_coding | -0.2250756 | 0.36792829 |
| GNG5 | protein_coding | -0.2251001 | 0.79925657 |
| DHCR7 | protein_coding | -0.2251628 | 6.32320833 |
| ZMIZ1 | protein_coding | -0.2251899 | 6.63983705 |
| CYB561D1 | protein_coding | -0.2252307 | 3.83358501 |
| CORO6 | protein_coding | -0.2253782 | 0.46989432 |
| DPP3 | protein_coding | -0.2254626 | 6.01649431 |
| IPO5 | protein_coding | -0.2255158 | 8.50638909 |
| PIDD1 | protein_coding | -0.2256176 | 4.93025939 |
| CHM | protein_coding | -0.2258387 | 4.13858053 |
| EI24 | protein_coding | -0.2259277 | 7.79459588 |
| RTF1 | protein_coding | -0.2259434 | 7.11428504 |
| IRF7 | protein_coding | -0.2260805 | 0.35115103 |
| RP11-440L14 | lncRNA | -0.2261339 | 2.43669232 |
| ECHDC3 | protein_coding | -0.2262016 | 1.3499551 |
| LINC00337 | lncRNA | -0.2262698 | 0.3361262 |
| RRP1 | protein_coding | -0.2263164 | 5.40772819 |
| ARHGAP21 | protein_coding | -0.2263357 | 5.98220431 |
| KBTBD7 | protein_coding | -0.226367 | 2.49220574 |

| | | | |
|---|---|---|---|
| PCYOX1 | protein_coding | -0.2263869 | 4.34891496 |
| CDC25A | protein_coding | -0.2265166 | 4.48692273 |
| RP11-439E19 | lncRNA | -0.2266205 | 0.65211659 |
| GDPD3 | protein_coding | -0.2266392 | 0.86299542 |
| FAM160A1-D | lncRNA | -0.2266861 | 0.27582854 |
| RP11-661A12 | lncRNA | -0.2267022 | 0.22627284 |
| ZNF337-AS1 | lncRNA | -0.2267412 | 0.72747711 |
| MRPL24 | protein_coding | -0.2268713 | 3.67945022 |
| DIO2 | protein_coding | -0.2269267 | 3.80554121 |
| TAF1B | protein_coding | -0.2269871 | 2.34451882 |
| PHKG2 | protein_coding | -0.2270215 | 3.00257374 |
| TREML1 | protein_coding | -0.227037 | 1.59488522 |
| NAA40 | protein_coding | -0.2270401 | 5.73538592 |
| CCDC15 | protein_coding | -0.2270923 | 3.14446243 |
| LINC02245 | lncRNA | -0.2271184 | 0.94935115 |
| LINC02447 | lncRNA | -0.2272913 | 0.29512339 |
| HNRNPUL2 | protein_coding | -0.2273167 | 8.0179976 |
| ANLN | protein_coding | -0.2274638 | 6.68986728 |
| MAMSTR | protein_coding | -0.2275476 | 2.52874697 |
| EXTL1 | protein_coding | -0.2275708 | 0.24669753 |
| FAM76A | protein_coding | -0.2277588 | 3.88185443 |
| RP11-229P13 | lncRNA | -0.2279416 | 0.23616159 |
| CDK5 | protein_coding | -0.2279559 | 4.24920939 |
| EDN2 | protein_coding | -0.2280197 | 1.98554983 |
| CDCA8 | protein_coding | -0.2280366 | 5.62576651 |
| RCAN1 | protein_coding | -0.22809 | 5.8341059 |
| FUS | protein_coding | -0.2281228 | 8.8156731 |
| PRDM10 | protein_coding | -0.2281812 | 3.58246231 |
| PPP1R16A | protein_coding | -0.2282221 | 1.64280586 |
| BDH2 | protein_coding | -0.2283389 | 0.63793797 |
| TP53INP2 | protein_coding | -0.2284055 | 5.47145587 |
| MNS1 | protein_coding | -0.228424 | 2.23993367 |
| BCKDHA | protein_coding | -0.2284836 | 0.24639722 |
| UNC45A | protein_coding | -0.2285335 | 6.17834851 |
| CHCHD5 | protein_coding | -0.2285458 | 2.32504841 |
| LINC00271 | lncRNA | -0.2286387 | 0.27319694 |
| TTLL1-AS1 | lncRNA | -0.2286491 | 0.25504318 |
| CDC45 | protein_coding | -0.2286997 | 5.09420785 |
| BLCAP | protein_coding | -0.2287847 | 5.95341217 |
| IL11 | protein_coding | -0.229199 | 0.33437441 |
| KIAA1841 | protein_coding | -0.2292225 | 2.64802245 |
| PASK | protein_coding | -0.2292741 | 5.48911179 |
| CNGA2 | protein_coding | -0.2292827 | 0.71950525 |
| FLNB | protein_coding | -0.2293307 | 7.83032536 |
| ZNF668 | protein_coding | -0.229425 | 1.95252671 |

| | | | |
|---|---|---|---|
| C1QTNF3 | protein_coding | -0.2294796 | 0.9281701 |
| TM7SF2 | protein_coding | -0.2295729 | 4.59112278 |
| WASF2 | protein_coding | -0.2295834 | 8.08894389 |
| CHAF1A | protein_coding | -0.229619 | 6.373113 |
| FNTB | protein_coding | -0.2297174 | 2.41939847 |
| POMK | protein_coding | -0.2297308 | 3.05629622 |
| TUBG1 | protein_coding | -0.2298272 | 5.86976779 |
| UBL7 | protein_coding | -0.2299954 | 4.83674465 |
| ZBTB18 | protein_coding | -0.2300183 | 5.66845289 |
| DTWD2 | protein_coding | -0.2301529 | 0.75611229 |
| XPNPEP1 | protein_coding | -0.2302277 | 4.93209775 |
| GATAD2A | protein_coding | -0.2302931 | 6.54752264 |
| LTA4H | protein_coding | -0.2303556 | 5.31272814 |
| MED11 | protein_coding | -0.2303981 | 1.304959 |
| CPNE3 | protein_coding | -0.2304006 | 5.35457449 |
| MAD2L1 | protein_coding | -0.2304558 | 5.37956727 |
| YIF1B | protein_coding | -0.230616 | 3.98576762 |
| TIGD5 | protein_coding | -0.2307838 | 1.36277494 |
| MAP3K3 | protein_coding | -0.2309179 | 3.0306775 |
| CENPC | protein_coding | -0.2310183 | 3.98350211 |
| NCDN | protein_coding | -0.2311942 | 4.74550997 |
| PLK3 | protein_coding | -0.2312975 | 0.98895308 |
| SLC25A32 | protein_coding | -0.2313077 | 0.50709649 |
| ZNF556 | protein_coding | -0.2313194 | 1.69892997 |
| ALMS1-IT1 | lncRNA | -0.2314124 | 0.51795777 |
| TPST2 | protein_coding | -0.2314302 | 2.29708432 |
| NCBP2AS2 | protein_coding | -0.2314845 | 2.78317135 |
| COPE | protein_coding | -0.2315049 | 5.74743755 |
| NPNT | protein_coding | -0.2315198 | 6.11727529 |
| AKAP1 | protein_coding | -0.2315665 | 6.30295837 |
| PTCH1 | protein_coding | -0.2316165 | 4.74343463 |
| USP11 | protein_coding | -0.2316334 | 6.33381558 |
| RP11-554A1[illegible] | protein_coding | -0.2318306 | 0.31440499 |
| CCDC177 | protein_coding | -0.2318452 | 0.43412707 |
| AC007879.5 | lncRNA | -0.2318673 | 0.81223267 |
| RNF220 | protein_coding | -0.2318954 | 6.72565173 |
| GNG4 | protein_coding | -0.2318985 | 8.15213168 |
| VPS4B | protein_coding | -0.2319228 | 5.25186089 |
| TACC1 | protein_coding | -0.2319871 | 3.51853651 |
| IMPACT | protein_coding | -0.2321623 | 3.3507984 |
| GNB5 | protein_coding | -0.2321722 | 4.49458307 |
| NEURL1B | protein_coding | -0.2322269 | 4.52283324 |
| KLHL7 | protein_coding | -0.2322319 | 4.72807402 |
| WSB1 | protein_coding | -0.2322827 | 5.86790531 |
| ZNF835 | protein_coding | -0.232372 | 1.0647405 |

| | | | |
|---|---|---|---|
| STIL | protein_coding | -0.2323993 | 5.14301522 |
| DEK | protein_coding | -0.232408 | 6.92900438 |
| NCAPH | protein_coding | -0.2324327 | 4.61143298 |
| LAPTM4B | protein_coding | -0.2325253 | 7.73658802 |
| RP13-1032I1 | lncRNA | -0.2326449 | 2.04535229 |
| ISLR2 | protein_coding | -0.2326517 | 4.47929646 |
| GGA2 | protein_coding | -0.2327219 | 7.80725804 |
| MSH4 | protein_coding | -0.2328241 | 0.50081193 |
| AC007383.3 | lncRNA | -0.2328305 | 1.07620178 |
| GPN2 | protein_coding | -0.2329308 | 4.44593349 |
| EFNA5 | protein_coding | -0.232977 | 4.87806022 |
| SDHAF1 | protein_coding | -0.233148 | 2.23521071 |
| MTARC1 | protein_coding | -0.2331761 | 1.96359059 |
| LANCL1 | protein_coding | -0.2334386 | 4.16956377 |
| GTPBP3 | protein_coding | -0.2336199 | 4.33563838 |
| IMMP1L | protein_coding | -0.2336602 | 3.04027298 |
| HDGFL2 | protein_coding | -0.233661 | 3.6052697 |
| RP11-864I4.1 | protein_coding | -0.2337193 | 0.42380675 |
| GLCCI1 | protein_coding | -0.2340375 | 4.63521257 |
| RTL5 | protein_coding | -0.2340634 | 3.30896793 |
| FUT10 | protein_coding | -0.2340863 | 1.55488802 |
| FOXP1 | protein_coding | -0.2340911 | 2.07827837 |
| PDRG1 | protein_coding | -0.2341765 | 4.191752 |
| SLC66A2 | protein_coding | -0.2341884 | 3.58172045 |
| MIEN1 | protein_coding | -0.234202 | 5.10290077 |
| TM9SF3 | protein_coding | -0.2343179 | 5.64617729 |
| ACSL3 | protein_coding | -0.2343655 | 4.38230468 |
| NCKAP5L | protein_coding | -0.2344806 | 4.40928034 |
| PPP1R37 | protein_coding | -0.2344906 | 2.66658492 |
| RPL26L1 | protein_coding | -0.2344972 | 2.60909308 |
| HINT2 | protein_coding | -0.2345517 | 1.3369518 |
| RGS12 | protein_coding | -0.234557 | 3.53102959 |
| SLC25A6 | protein_coding | -0.2345882 | 6.1822202 |
| ZNF132-DT | lncRNA | -0.234807 | 0.45840593 |
| RPL4 | protein_coding | -0.2349537 | 9.48647225 |
| LINC02012 | lncRNA | -0.2352069 | 0.66021834 |
| MRPL41 | protein_coding | -0.2355438 | 2.18325593 |
| PPM1F | protein_coding | -0.2359557 | 4.48273186 |
| SYP | protein_coding | -0.2360877 | 6.35566284 |
| PPP2R5D | protein_coding | -0.2361274 | 6.51296733 |
| NUF2 | protein_coding | -0.2362181 | 5.38821208 |
| ARHGDIG | protein_coding | -0.2364549 | 2.93457767 |
| COX7C | protein_coding | -0.2365456 | 7.64194415 |
| FAXC | protein_coding | -0.236574 | 2.53590021 |
| FAM182B | lncRNA | -0.237016 | 0.50117261 |

| | | | |
|---|---|---|---|
| SNRK | protein_coding | -0.2370406 | 3.93696162 |
| RTL8C | protein_coding | -0.2371164 | 4.2062494 |
| RNF170 | protein_coding | -0.2371591 | 2.43976226 |
| LL22NC03-N1 | lncRNA | -0.2373032 | 0.47424638 |
| DCLRE1B | protein_coding | -0.2373254 | 4.4122629 |
| ACTR3B | protein_coding | -0.2374288 | 3.81980983 |
| MTMR6 | protein_coding | -0.2374835 | 4.58211415 |
| PREP | protein_coding | -0.2374959 | 4.76014584 |
| SRF | protein_coding | -0.2375407 | 5.2093246 |
| MRPL20-AS1 | lncRNA | -0.2375674 | 2.89028308 |
| VPS53 | protein_coding | -0.2376202 | 5.2307339 |
| CALM2 | protein_coding | -0.2377297 | 6.13567873 |
| TCTA | protein_coding | -0.2378177 | 5.30655975 |
| MFSD12 | protein_coding | -0.2379266 | 4.44428814 |
| RTN1 | protein_coding | -0.2379503 | 6.36332862 |
| EEF1A1 | protein_coding | -0.2379802 | 10.7124921 |
| CHMP7 | protein_coding | -0.2379804 | 4.4440182 |
| RCC1 | protein_coding | -0.2381384 | 6.70697098 |
| KDSR-DT | lncRNA | -0.238197 | 0.89619878 |
| ERO1A | protein_coding | -0.2382854 | 5.67978719 |
| IL17RA | protein_coding | -0.2383484 | 4.86177196 |
| CATSPERB | protein_coding | -0.2386405 | 0.32266844 |
| GPC6 | protein_coding | -0.2387334 | 2.55590906 |
| GLO1 | protein_coding | -0.2387922 | 6.96815007 |
| PPP3CB | protein_coding | -0.238938 | 5.22285268 |
| RP11-11M20 | lncRNA | -0.2390039 | 0.96979959 |
| EVI5 | protein_coding | -0.2390794 | 1.51402567 |
| MAEA | protein_coding | -0.2391055 | 6.24621433 |
| PIP5KL1 | protein_coding | -0.2391601 | 0.69876962 |
| ISOC2 | protein_coding | -0.2392287 | 4.1901933 |
| MEX3B | protein_coding | -0.2393019 | 3.86145794 |
| RGMB | protein_coding | -0.2393282 | 6.35972966 |
| GNB4 | protein_coding | -0.2393937 | 3.86092992 |
| RP11-259K21 | lncRNA | -0.2394101 | 0.48923282 |
| VDAC3 | protein_coding | -0.2396094 | 6.61559089 |
| IRX5 | protein_coding | -0.2396158 | 1.93472445 |
| ORMDL3 | protein_coding | -0.23968 | 5.45690481 |
| RPA3 | protein_coding | -0.239823 | 3.95200598 |
| APEX2 | protein_coding | -0.2399365 | 5.3346605 |
| TSPAN3 | protein_coding | -0.2400632 | 7.41681995 |
| MT-CYB | protein_coding | -0.240185 | 7.76943588 |
| PLEKHJ1 | protein_coding | -0.2402311 | 4.77679974 |
| SNPH | protein_coding | -0.2404612 | 0.18917782 |
| IER5 | protein_coding | -0.2405222 | 3.04238613 |
| ZNF420 | protein_coding | -0.2411258 | 3.87437243 |

| | | | |
|---|---|---|---|
| PDHX | protein_coding | -0.2412144 | 4.23481643 |
| RALY | protein_coding | -0.241247 | 7.11169918 |
| YIPF4 | protein_coding | -0.2413044 | 5.6456812 |
| ZYG11A | protein_coding | -0.241374 | 3.02082053 |
| CTA-276F8.1 | lncRNA | -0.2413947 | 0.24795749 |
| HSF4 | protein_coding | -0.2415164 | 3.06835792 |
| RAE1 | protein_coding | -0.2416544 | 6.52644955 |
| NUDT22 | protein_coding | -0.2416895 | 3.63291909 |
| E2F2 | protein_coding | -0.241844 | 7.41619636 |
| TBC1D22A | protein_coding | -0.2419351 | 4.20983377 |
| RHPN1 | protein_coding | -0.242013 | 1.7602017 |
| CTB-58E17.1 | lncRNA | -0.2420355 | 1.78252052 |
| LYRM4 | protein_coding | -0.2420894 | 3.58805076 |
| MARCKSL1 | protein_coding | -0.2420965 | 7.49467888 |
| DOT1L | protein_coding | -0.2420992 | 4.34800859 |
| SLC25A5 | protein_coding | -0.242144 | 8.87310985 |
| IGFBPL1 | protein_coding | -0.2421722 | 8.41755542 |
| CH17-55D9.1 | lncRNA | -0.2421826 | 0.77703311 |
| ARF1 | protein_coding | -0.2422379 | 9.29466651 |
| SNRPB | protein_coding | -0.2422953 | 8.2340941 |
| HTATSF1 | protein_coding | -0.2423339 | 9.01714663 |
| PHF19 | protein_coding | -0.2423958 | 6.76206248 |
| METTL23 | protein_coding | -0.2424137 | 2.41033293 |
| MRNIP | protein_coding | -0.2424967 | 2.93147574 |
| CXXC1 | protein_coding | -0.2425549 | 6.27948754 |
| H2AX | protein_coding | -0.2425962 | 2.43882608 |
| PREX1 | protein_coding | -0.2426293 | 2.40065595 |
| NSMCE3 | protein_coding | -0.2427257 | 2.37096387 |
| RP5-1050D4. | lncRNA | -0.242752 | 0.64583029 |
| EFHC2 | protein_coding | -0.2427549 | 0.68065433 |
| BTG3 | protein_coding | -0.2428438 | 5.13801208 |
| RP11-930P14 | lncRNA | -0.2428819 | 0.23459893 |
| RP11-550A5. | lncRNA | -0.2429545 | 0.28280013 |
| LMNTD2 | protein_coding | -0.2429681 | 1.54279684 |
| MTMR7 | protein_coding | -0.2430207 | 3.12675248 |
| TMEM51-AS | lncRNA | -0.2430725 | 1.1255196 |
| TNRC18 | protein_coding | -0.2431426 | 2.79320573 |
| NPEPL1 | protein_coding | -0.243146 | 0.67483753 |
| PHKA1 | protein_coding | -0.2432614 | 5.58225031 |
| CKS2 | protein_coding | -0.2433427 | 6.46700181 |
| UBA52 | protein_coding | -0.2435493 | 8.02140291 |
| RP11-386G1 | lncRNA | -0.2436005 | 0.65496331 |
| HMGB3 | protein_coding | -0.2436269 | 7.41682684 |
| TEKT2 | protein_coding | -0.2436426 | 1.03995689 |
| HEY2 | protein_coding | -0.2436427 | 1.61204472 |

| | | | |
|---|---|---|---|
| LMBRD2 | protein_coding | -0.2436661 | 3.99290845 |
| PUSL1 | protein_coding | -0.2436726 | 3.11507368 |
| ATPSCKMT | protein_coding | -0.2437493 | 2.11303695 |
| MNT | protein_coding | -0.2438307 | 4.39077241 |
| POLR2H | protein_coding | -0.2440054 | 6.50933968 |
| ZNF888 | protein_coding | -0.2440825 | 1.79315385 |
| EFCAB14 | protein_coding | -0.2441992 | 6.07677051 |
| GAS5-AS1 | lncRNA | -0.244247 | 0.64303758 |
| TNFRSF21 | protein_coding | -0.2443642 | 6.3338648 |
| LIN52 | protein_coding | -0.2444298 | 4.90959874 |
| ASB3 | protein_coding | -0.2444887 | 3.15704874 |
| TPGS2 | protein_coding | -0.2445366 | 6.54480412 |
| TTLL12 | protein_coding | -0.2449305 | 7.83204582 |
| SMAP2 | protein_coding | -0.2451442 | 5.9583481 |
| MPZL1 | protein_coding | -0.2452006 | 6.42194085 |
| CENPV | protein_coding | -0.2452658 | 5.43746821 |
| AURKA | protein_coding | -0.2452794 | 6.57772431 |
| RP11-646I6.5 | lncRNA | -0.24557 | 0.46828753 |
| RCOR1 | protein_coding | -0.2456617 | 7.55357323 |
| ISOC1 | protein_coding | -0.245722 | 6.09769669 |
| MRPS26 | protein_coding | -0.2457467 | 2.19014203 |
| CTNNBIP1 | protein_coding | -0.2457628 | 3.51102915 |
| MIER2 | protein_coding | -0.2458576 | 4.88113014 |
| ATP1A1-AS1 | lncRNA | -0.2459933 | 0.5564324 |
| TCERG1L | protein_coding | -0.2460668 | 4.50860824 |
| ABHD12 | protein_coding | -0.2463161 | 5.17454962 |
| PFKFB3 | protein_coding | -0.2464029 | 4.20553851 |
| PROB1 | protein_coding | -0.246667 | 1.01571253 |
| CCDC9B | protein_coding | -0.2467297 | 0.75459282 |
| BET1L | protein_coding | -0.246798 | 5.76134078 |
| NUDT7 | protein_coding | -0.2469524 | 1.22685616 |
| RP11-582J16 | lncRNA | -0.2469967 | 0.93250323 |
| LINC01424 | lncRNA | -0.2470595 | 0.37233941 |
| ZMAT5 | protein_coding | -0.2472309 | 4.27426908 |
| MRPS28 | protein_coding | -0.2472415 | 1.23633692 |
| NLGN2 | protein_coding | -0.2472679 | 2.69560193 |
| CDK5RAP2 | protein_coding | -0.2472751 | 7.56064838 |
| HOMER1 | protein_coding | -0.247423 | 3.22981148 |
| NUDT4 | protein_coding | -0.2475395 | 2.87907581 |
| ARHGAP20 | protein_coding | -0.2476025 | 0.75211397 |
| ADGRL1 | protein_coding | -0.2476521 | 6.06648349 |
| RP11-323N1 | lncRNA | -0.247719 | 0.46931826 |
| RHOA | protein_coding | -0.247791 | 7.70726111 |
| CFAP410 | protein_coding | -0.2478649 | 2.79217428 |
| SLCO4A1-AS | lncRNA | -0.2479303 | 1.10531635 |

| | | | |
|---|---|---|---|
| BUB1B | protein_coding | -0.2480487 | 7.29352049 |
| SGMS1 | protein_coding | -0.2481123 | 3.4401056 |
| PROC | protein_coding | -0.2481504 | 0.48331742 |
| EIF4A3 | protein_coding | -0.2481519 | 5.69560901 |
| STMP1 | protein_coding | -0.2481843 | 4.66036221 |
| APPL2 | protein_coding | -0.2482677 | 3.78023275 |
| CCDC127 | protein_coding | -0.2483276 | 3.94134455 |
| RMND5B | protein_coding | -0.248328 | 5.04773529 |
| PIGU | protein_coding | -0.2483653 | 5.19894984 |
| C1D | protein_coding | -0.2483715 | 2.79383328 |
| LINC00682 | lncRNA | -0.2483897 | 0.2316838 |
| MYOZ1 | protein_coding | -0.2487755 | 0.27094784 |
| CHPF | protein_coding | -0.2488744 | 3.61355469 |
| LZTS1 | protein_coding | -0.2489032 | 5.57042631 |
| RP5-884G6.2 | lncRNA | -0.2489429 | 0.26427105 |
| UBL4A | protein_coding | -0.2489826 | 5.92671646 |
| RHOBTB1 | protein_coding | -0.2490096 | 1.84926056 |
| ZNF789 | protein_coding | -0.2490546 | 3.15218324 |
| RP11-22N19. | lncRNA | -0.2491729 | 3.10272628 |
| POLR2I | protein_coding | -0.2492099 | 4.77066529 |
| KLHL15 | protein_coding | -0.2492583 | 4.59413271 |
| TUBB | protein_coding | -0.2492779 | 10.7866154 |
| TBATA | protein_coding | -0.2495433 | 0.42574212 |
| ATG9A | protein_coding | -0.2496438 | 5.21683931 |
| SNRPG | protein_coding | -0.2497165 | 6.92799688 |
| TMEM18 | protein_coding | -0.2497483 | 5.31521291 |
| CDKN2D | protein_coding | -0.2500027 | 4.05948408 |
| RP5-1014D1: | lncRNA | -0.2500502 | 0.645659 |
| RABL2B | protein_coding | -0.2501169 | 3.518399 |
| MDH2 | protein_coding | -0.2501212 | 8.61788872 |
| RABEP2 | protein_coding | -0.2501688 | 2.8692815 |
| FAHD1 | protein_coding | -0.2502613 | 3.12893716 |
| SH3GL2 | protein_coding | -0.250399 | 4.48785595 |
| ANKDD1B | protein_coding | -0.2504209 | 0.51365442 |
| ACTB | protein_coding | -0.250587 | 9.98888728 |
| NR2C2AP | protein_coding | -0.2508149 | 4.12800073 |
| AC002467.7 | lncRNA | -0.2509874 | 0.77191741 |
| PRAME | protein_coding | -0.2510507 | 4.64346856 |
| PCDHGA12 | protein_coding | -0.2512096 | 0.29502294 |
| CLHC1 | protein_coding | -0.251235 | 1.44820915 |
| ZNF813 | protein_coding | -0.2513724 | 1.63446354 |
| TBC1D10A | protein_coding | -0.251425 | 1.98953617 |
| STK35 | protein_coding | -0.2514957 | 5.5656709 |
| PPM1E | protein_coding | -0.2515847 | 5.6153347 |
| GPSM2 | protein_coding | -0.2516311 | 6.3865266 |

| | | | |
|---|---|---|---|
| UBE2C | protein_coding | -0.2516581 | 7.13739706 |
| RPS4X | protein_coding | -0.2517549 | 9.66046698 |
| PTPMT1 | protein_coding | -0.2517835 | 3.1384951 |
| KLHL17 | protein_coding | -0.2518988 | 1.35962986 |
| R3HCC1 | protein_coding | -0.2521908 | 2.84601508 |
| NDUFS5 | protein_coding | -0.2524286 | 6.66955659 |
| TMEM161A | protein_coding | -0.2525603 | 4.31844257 |
| TRO | protein_coding | -0.2526766 | 4.14925861 |
| DARS2 | protein_coding | -0.252899 | 6.24745506 |
| DNAJC9 | protein_coding | -0.2530009 | 5.82957213 |
| RPP25 | protein_coding | -0.2530931 | 3.72433808 |
| PCDHGC5 | protein_coding | -0.2531166 | 0.5347464 |
| DCXR | protein_coding | -0.2531183 | 4.56279156 |
| ZBTB42 | protein_coding | -0.2531429 | 3.08523328 |
| SUPT3H | protein_coding | -0.2531588 | 2.00479548 |
| PPM1N | protein_coding | -0.2532872 | 1.54738408 |
| ST6GALNAC4 | protein_coding | -0.253302 | 6.48845394 |
| PPP2R3B | protein_coding | -0.2533603 | 2.14737562 |
| STRADB | protein_coding | -0.2534597 | 2.96725116 |
| RP11-463C8. | lncRNA | -0.2534786 | 0.38254221 |
| BLMH | protein_coding | -0.2535008 | 4.75504762 |
| ITGB1BP1 | protein_coding | -0.253511 | 6.56682419 |
| AC093323.3 | protein_coding | -0.2535304 | 4.18385444 |
| NAA60 | protein_coding | -0.2536715 | 6.08367247 |
| RAB3IP | protein_coding | -0.2536851 | 9.01799977 |
| ATP6V1F | protein_coding | -0.2537601 | 6.56828686 |
| PRKX | protein_coding | -0.2538478 | 5.64065777 |
| RP11-1000B6 | lncRNA | -0.2539952 | 0.48425517 |
| RBM14 | protein_coding | -0.254109 | 7.65269679 |
| PHRF1 | protein_coding | -0.2541857 | 5.1357254 |
| RPL23A | protein_coding | -0.2542059 | 8.69839588 |
| PTPRG | protein_coding | -0.2542436 | 6.2861524 |
| RBM27 | protein_coding | -0.2543106 | 3.47800028 |
| SPECC1-DT | lncRNA | -0.2545234 | 0.33411809 |
| DNAJB11 | protein_coding | -0.2546387 | 6.69816577 |
| HEBP1 | protein_coding | -0.2546424 | 1.96486623 |
| NDUFB2-AS1 | lncRNA | -0.2547981 | 0.24016075 |
| TCP1 | protein_coding | -0.2549358 | 8.20498319 |
| ANO10 | protein_coding | -0.2553992 | 1.72703488 |
| GTF2I | protein_coding | -0.2554439 | 5.45690294 |
| C15orf61 | protein_coding | -0.2555474 | 3.94933898 |
| RP11-108K14 | protein_coding | -0.255693 | 0.37048353 |
| CHCHD7 | protein_coding | -0.2560499 | 2.58746481 |
| BRI3BP | protein_coding | -0.2560802 | 8.16997372 |
| TMX4 | protein_coding | -0.2563628 | 7.87041672 |

| | | | |
|---|---|---|---|
| KBTBD4 | protein_coding | -0.2563707 | 4.13546331 |
| DOK1 | protein_coding | -0.2563785 | 2.19638305 |
| AC002511.3 | lncRNA | -0.2564346 | 0.33277836 |
| FAM210A | protein_coding | -0.2565099 | 3.16009858 |
| ZNF143 | protein_coding | -0.2565377 | 3.72534384 |
| WWP2 | protein_coding | -0.2565835 | 6.01060653 |
| MANF | protein_coding | -0.2566274 | 5.59267279 |
| CCT8 | protein_coding | -0.2566469 | 8.97979486 |
| NUDT1 | protein_coding | -0.256682 | 5.77459759 |
| TMEM104 | protein_coding | -0.2567535 | 5.90409415 |
| RTN3 | protein_coding | -0.2568216 | 10.0609556 |
| FAM83D | protein_coding | -0.2568552 | 5.54629031 |
| CLGN | protein_coding | -0.2570487 | 2.05473548 |
| NUDC | protein_coding | -0.2570908 | 7.37629975 |
| RP11-96K19. | lncRNA | -0.2571116 | 0.25353078 |
| TCF4 | protein_coding | -0.257193 | 8.5457006 |
| CTA-250D10. | lncRNA | -0.2572918 | 0.29272175 |
| MCM7 | protein_coding | -0.2573889 | 9.6311005 |
| AC142472.6 | lncRNA | -0.2574252 | 0.42615284 |
| CORO1B | protein_coding | -0.2574456 | 5.00348939 |
| CEP63 | protein_coding | -0.2574878 | 2.46646946 |
| RP11-575F12 | lncRNA | -0.2575302 | 0.50572467 |
| CLTB | protein_coding | -0.2575748 | 4.9612207 |
| ADIRF | protein_coding | -0.2577084 | 4.08933803 |
| AMFR | protein_coding | -0.2577827 | 5.13355186 |
| ELP6 | protein_coding | -0.2582606 | 4.52474612 |
| KLHL22 | protein_coding | -0.2583712 | 4.88501796 |
| ANKRD7 | protein_coding | -0.2586265 | 0.316941 |
| KDM5B | protein_coding | -0.2586307 | 5.77951093 |
| LMCD1-AS1 | lncRNA | -0.2587196 | 0.24730947 |
| COA8 | protein_coding | -0.258735 | 3.23485292 |
| PLA2G4B | protein_coding | -0.2590506 | 0.37640427 |
| YBX2 | protein_coding | -0.2590589 | 1.78289493 |
| ANKRD46 | protein_coding | -0.2591812 | 3.8913378 |
| LRPAP1 | protein_coding | -0.2592227 | 5.60459525 |
| CHMP1A | protein_coding | -0.2592604 | 6.58137895 |
| ZNF273 | protein_coding | -0.2593105 | 2.46788756 |
| LRCH4 | protein_coding | -0.259311 | 2.82066958 |
| FAM72C | protein_coding | -0.2593171 | 0.85710911 |
| LINC01560 | lncRNA | -0.2594745 | 2.05904448 |
| EPHA10 | protein_coding | -0.2595761 | 3.29211962 |
| ATP6V0E2 | protein_coding | -0.2596886 | 7.29898381 |
| SEL1L3 | protein_coding | -0.2598199 | 8.48470295 |
| LSM14B | protein_coding | -0.2599571 | 8.08716482 |
| RP11-19F9.2 | lncRNA | -0.2601209 | 1.35575203 |

| | | | |
|---|---|---|---|
| RPP25L | protein_coding | -0.2601237 | 3.10891716 |
| DNASE1 | protein_coding | -0.2604147 | 5.23458939 |
| CD248 | protein_coding | -0.2604328 | 3.33524562 |
| SH3PXD2A | protein_coding | -0.2604732 | 7.78011506 |
| FBL | protein_coding | -0.2605597 | 6.22214473 |
| MAP3K20 | protein_coding | -0.2606354 | 5.61597849 |
| BCLAF1 | protein_coding | -0.2612691 | 6.856583 |
| PPME1 | protein_coding | -0.2614228 | 7.06820838 |
| DDB2 | protein_coding | -0.2614409 | 4.81046784 |
| MXI1 | protein_coding | -0.2615486 | 3.8848322 |
| RMND5A | protein_coding | -0.2616384 | 7.92103187 |
| KLF16 | protein_coding | -0.2616655 | 4.15172501 |
| LRRC63 | protein_coding | -0.2618487 | 0.4299755 |
| TMEM258 | protein_coding | -0.2620123 | 4.59004884 |
| RETREG2 | protein_coding | -0.2620282 | 6.8929423 |
| FBXO3 | protein_coding | -0.2620953 | 5.57127914 |
| RSBN1 | protein_coding | -0.2621302 | 5.16672539 |
| BCAP31 | protein_coding | -0.2621647 | 7.23196212 |
| EPM2AIP1 | protein_coding | -0.2621767 | 5.09969148 |
| PIK3C3 | protein_coding | -0.2622715 | 5.74860729 |
| SNRPC | protein_coding | -0.2622727 | 7.26958011 |
| IRF2BP1 | protein_coding | -0.2622891 | 1.91347557 |
| ATP9B | protein_coding | -0.2623496 | 3.31358177 |
| FMNL2 | protein_coding | -0.262397 | 4.97711327 |
| POLR2L | protein_coding | -0.262499 | 6.7904656 |
| RFC3 | protein_coding | -0.2626005 | 3.24876195 |
| BOLA3-AS1 | lncRNA | -0.2626504 | 2.08671084 |
| RP11-7O11.3 | lncRNA | -0.2630618 | 0.63719457 |
| RPL5 | protein_coding | -0.2630818 | 9.86481051 |
| INAFM2 | protein_coding | -0.2631181 | 4.80694626 |
| RP11-705C15 | lncRNA | -0.2631908 | 0.77392988 |
| ELMO3 | protein_coding | -0.2632184 | 2.69070007 |
| C11orf65 | protein_coding | -0.2632737 | 0.49224172 |
| TNFAIP8 | protein_coding | -0.263371 | 2.55136757 |
| GTF2H5 | protein_coding | -0.2634005 | 3.74682252 |
| SPAG9 | protein_coding | -0.2634418 | 8.69269426 |
| PTPRT | protein_coding | -0.2634498 | 3.48683268 |
| DIAPH3 | protein_coding | -0.2636752 | 6.40027402 |
| LINC01002 | lncRNA | -0.2638446 | 0.82025552 |
| STX10 | protein_coding | -0.2638583 | 5.20087931 |
| PIAS3 | protein_coding | -0.2638744 | 4.9610902 |
| NUDT21 | protein_coding | -0.2639403 | 2.6464023 |
| PPP2R1A | protein_coding | -0.2640481 | 9.3100909 |
| PWWP3A | protein_coding | -0.2641433 | 5.66989295 |
| SC5D | protein_coding | -0.2641656 | 5.47149571 |

| | | | |
|---|---|---|---|
| RPL15 | protein_coding | -0.264227 | 8.36785283 |
| ZNF292 | protein_coding | -0.2642363 | 7.19399185 |
| C6orf226 | protein_coding | -0.2646464 | 0.95144111 |
| GNAO1-AS1 | lncRNA | -0.2647385 | 0.25971172 |
| STIP1 | protein_coding | -0.2648026 | 8.39485874 |
| ROGDI | protein_coding | -0.2648076 | 1.98651493 |
| SBNO1 | protein_coding | -0.2648175 | 7.55781751 |
| BTBD6 | protein_coding | -0.2648322 | 6.42845592 |
| RPL7A | protein_coding | -0.26486 | 10.858658 |
| KCNJ11 | protein_coding | -0.2649634 | 1.44410383 |
| GNAI1 | protein_coding | -0.2649727 | 4.93592369 |
| ARHGEF25 | protein_coding | -0.2651315 | 3.82312803 |
| GSTA4 | protein_coding | -0.2652272 | 4.60625595 |
| C8orf88 | protein_coding | -0.2653268 | 2.75678561 |
| ZNF367 | protein_coding | -0.2653703 | 5.10603735 |
| ARL2 | protein_coding | -0.2659831 | 4.50128462 |
| TMEM59 | protein_coding | -0.266098 | 8.54476546 |
| ENDOD1 | protein_coding | -0.266269 | 2.62012038 |
| HDAC1 | protein_coding | -0.2663614 | 7.77473218 |
| DIPK2A | protein_coding | -0.2664854 | 3.891426 |
| MZT1 | protein_coding | -0.2667316 | 4.27373192 |
| CAMKV | protein_coding | -0.2667464 | 4.06369069 |
| MRPL12 | protein_coding | -0.2667852 | 1.34585387 |
| CYB5R3 | protein_coding | -0.2667892 | 8.59463956 |
| UQCRH | protein_coding | -0.2668343 | 8.06594736 |
| PET117 | protein_coding | -0.2668362 | 0.33890301 |
| PAIP2 | protein_coding | -0.2668509 | 7.37093513 |
| CDK6 | protein_coding | -0.266863 | 1.08929543 |
| UBL5 | protein_coding | -0.2672942 | 7.05919077 |
| SP4 | protein_coding | -0.2674326 | 3.84684252 |
| CNPPD1 | protein_coding | -0.267621 | 6.51186557 |
| UBTD2 | protein_coding | -0.2676347 | 3.12500025 |
| STIM1 | protein_coding | -0.2676565 | 5.59108132 |
| AGFG2 | protein_coding | -0.2677421 | 4.19283977 |
| ZNF414 | protein_coding | -0.2678828 | 2.64624787 |
| PTPN2 | protein_coding | -0.2679077 | 5.49880199 |
| EIF4B | protein_coding | -0.2680154 | 9.59937396 |
| ERICH3 | protein_coding | -0.2682987 | 3.1259421 |
| PCNA | protein_coding | -0.268303 | 9.92191296 |
| MAPRE2 | protein_coding | -0.2683717 | 7.10737137 |
| RPS6 | protein_coding | -0.268411 | 9.6997887 |
| USP51 | protein_coding | -0.2684352 | 0.87479524 |
| SMIM22 | protein_coding | -0.2684728 | 0.49991592 |
| ZDHHC20 | protein_coding | -0.2685355 | 4.32302664 |
| SLC26A2 | protein_coding | -0.2689121 | 2.55625734 |

| | | | |
|---|---|---|---|
| BCDIN3D-AS | lncRNA | -0.2690041 | 0.95611444 |
| LPIN2 | protein_coding | -0.2690336 | 7.34302269 |
| SLC25A36 | protein_coding | -0.2695694 | 6.11004773 |
| PPP3R1 | protein_coding | -0.2697895 | 4.74598142 |
| SNU13 | protein_coding | -0.2699758 | 8.23523888 |
| SPIN4 | protein_coding | -0.2699968 | 3.81105767 |
| CCP110 | protein_coding | -0.2700506 | 8.6111065 |
| SLC7A7 | protein_coding | -0.2701658 | 0.2619882 |
| ZNF295-AS1 | lncRNA | -0.270171 | 0.31876662 |
| TFAP4 | protein_coding | -0.2702548 | 4.0229114 |
| CDK3 | protein_coding | -0.2702583 | 0.38782378 |
| RP11-326K13 | lncRNA | -0.2703581 | 0.30068296 |
| LYPLA2 | protein_coding | -0.2704458 | 6.62034128 |
| ZNF432 | protein_coding | -0.2705761 | 1.24534041 |
| ANKRD34A | protein_coding | -0.2705953 | 0.39734597 |
| PTTG1 | protein_coding | -0.2707786 | 7.01158283 |
| SNHG29 | lncRNA | -0.2708465 | 7.94548562 |
| NR4A1 | protein_coding | -0.2708831 | 4.50862955 |
| AKTIP | protein_coding | -0.2709996 | 2.87032838 |
| ZNF565 | protein_coding | -0.2711093 | 1.11460017 |
| WDFY2 | protein_coding | -0.2712115 | 4.41978775 |
| PKM | protein_coding | -0.271221 | 10.4711918 |
| MED16 | protein_coding | -0.2713712 | 3.84810153 |
| RHBDD2 | protein_coding | -0.2713834 | 5.94580606 |
| AC097468.4 | lncRNA | -0.2713983 | 1.93418161 |
| INSIG2 | protein_coding | -0.271624 | 3.19745522 |
| ZNF519 | protein_coding | -0.2716714 | 2.01429424 |
| RAP1A | protein_coding | -0.2717495 | 4.92699676 |
| CAPZB | protein_coding | -0.2717959 | 7.89140097 |
| MID1IP1 | protein_coding | -0.2719817 | 5.19575904 |
| AMN | protein_coding | -0.2720694 | 1.6984268 |
| RP11-588G2 | lncRNA | -0.272102 | 0.63135903 |
| SYN3 | protein_coding | -0.2722176 | 4.4307254 |
| ANKS1B | protein_coding | -0.2722582 | 3.5279388 |
| MCRIP1 | protein_coding | -0.2727024 | 4.23164191 |
| HMGA1 | protein_coding | -0.2727453 | 9.29383728 |
| BIRC5 | protein_coding | -0.2729771 | 8.07247561 |
| PIGV | protein_coding | -0.2729799 | 3.54101823 |
| CBX5 | protein_coding | -0.2729859 | 10.1157456 |
| CTA-796E4.4 | lncRNA | -0.273267 | 1.19588718 |
| C19orf53 | protein_coding | -0.2733241 | 5.90983723 |
| PIK3AP1 | protein_coding | -0.2734154 | 0.57214459 |
| TEX261 | protein_coding | -0.2734611 | 7.5995278 |
| TSEN34 | protein_coding | -0.273565 | 6.9881553 |
| HILPDA | protein_coding | -0.2736423 | 5.51771924 |

| | | | |
|---|---|---|---|
| RP11-770J1.3 | lncRNA | -0.2736853 | 0.68661061 |
| CTD-2562G15 | lncRNA | -0.2737057 | 0.36923135 |
| PARD6A | protein_coding | -0.2737288 | 3.53054861 |
| CACNB3 | protein_coding | -0.2738453 | 6.69909386 |
| RP5-892C22. | lncRNA | -0.2739366 | 0.39870366 |
| E2F1 | protein_coding | -0.2739785 | 7.54985513 |
| CDS2 | protein_coding | -0.2740515 | 7.47968994 |
| CUTA | protein_coding | -0.2740739 | 7.12343216 |
| UQCRHL | protein_coding | -0.2740748 | 2.23240169 |
| ATG9B | protein_coding | -0.2741546 | 1.4272553 |
| RP11-775C24 | lncRNA | -0.274258 | 0.42827989 |
| HNRNPR | protein_coding | -0.2744182 | 8.24879723 |
| NDFIP1 | protein_coding | -0.2745218 | 7.3255411 |
| ZFPM1 | protein_coding | -0.2745372 | 1.44014612 |
| MACF1 | protein_coding | -0.2745604 | 7.67428766 |
| ZNF496 | protein_coding | -0.2746907 | 4.40761932 |
| PSIP1 | protein_coding | -0.2752877 | 7.88004656 |
| ESRP2 | protein_coding | -0.275348 | 4.11897977 |
| RPL21 | protein_coding | -0.2756233 | 9.5899175 |
| EPHX1 | protein_coding | -0.2757846 | 6.90781379 |
| TLE5 | protein_coding | -0.275916 | 7.85937778 |
| TGFBR1 | protein_coding | -0.2760988 | 6.1208242 |
| RPL41 | protein_coding | -0.2761364 | 7.95158718 |
| PPP1R21 | protein_coding | -0.2761531 | 4.93449376 |
| CREB3L4 | protein_coding | -0.2763782 | 5.39296548 |
| MZT2A | protein_coding | -0.2764673 | 2.90898328 |
| RUNDC3A | protein_coding | -0.2765183 | 6.41377155 |
| RP11-111A2 | lncRNA | -0.2765428 | 0.35290856 |
| DEPDC1 | protein_coding | -0.2765513 | 5.73045784 |
| NUMA1 | protein_coding | -0.2765785 | 8.75703003 |
| ATAD3A | protein_coding | -0.2766941 | 5.06233623 |
| TIMM13 | protein_coding | -0.2767176 | 7.38239394 |
| PSMA3-AS1 | lncRNA | -0.2768247 | 4.63472709 |
| RP11-634H2 | lncRNA | -0.2769714 | 1.54331831 |
| PPL | protein_coding | -0.2770699 | 1.07577928 |
| ECT2 | protein_coding | -0.2773635 | 7.68764802 |
| GIGYF1 | protein_coding | -0.2773728 | 6.61342702 |
| STARD13 | protein_coding | -0.2774536 | 0.54364306 |
| TTC21B | protein_coding | -0.2775282 | 6.07108723 |
| QRFPR | protein_coding | -0.2776472 | 3.43151977 |
| RP11-731C17 | lncRNA | -0.2777555 | 0.79406817 |
| ZC3HAV1L | protein_coding | -0.2778196 | 2.4971405 |
| TMOD2 | protein_coding | -0.2779428 | 6.94926704 |
| GATD1 | protein_coding | -0.2780483 | 6.0734088 |
| STT3A | protein_coding | -0.2781431 | 8.26373814 |

| | | | |
|---|---|---|---|
| STXBP1 | protein_coding | -0.2783258 | 8.71825551 |
| SPTBN4 | protein_coding | -0.2786968 | 4.18177602 |
| ZNF575 | protein_coding | -0.2787386 | 1.36279115 |
| CADM4 | protein_coding | -0.2788289 | 5.27043054 |
| MIS18BP1 | protein_coding | -0.2789367 | 5.38437676 |
| ARHGDIA | protein_coding | -0.2789942 | 7.12195092 |
| PPP6R1 | protein_coding | -0.2791734 | 7.53689563 |
| AC007292.6 | lncRNA | -0.2792289 | 1.34971179 |
| CDPF1 | protein_coding | -0.2793028 | 2.54863283 |
| ATP2A2 | protein_coding | -0.2796692 | 9.98202296 |
| MRI1 | protein_coding | -0.2797658 | 4.64493052 |
| PCDHA8 | protein_coding | -0.2799567 | 0.51631916 |
| LMLN | protein_coding | -0.2800965 | 1.985863 |
| CYP2U1-AS1 | lncRNA | -0.2801813 | 1.56052166 |
| GID8 | protein_coding | -0.2802056 | 7.80633836 |
| MYLIP | protein_coding | -0.280478 | 2.67067002 |
| ARPC5 | protein_coding | -0.2805223 | 7.55618909 |
| UBTF | protein_coding | -0.2805968 | 8.80017035 |
| TMEM177 | protein_coding | -0.2806563 | 3.16126541 |
| USF2 | protein_coding | -0.2809861 | 5.03769225 |
| RP11-967K21 | lncRNA | -0.2811974 | 0.99387837 |
| ARL9 | protein_coding | -0.2813524 | 2.12084171 |
| STXBP5L | protein_coding | -0.2814397 | 5.48416053 |
| FAM13B | protein_coding | -0.2814772 | 6.70977191 |
| ATOH1 | protein_coding | -0.2815745 | 8.95320518 |
| MAP9 | protein_coding | -0.2817154 | 1.82929043 |
| SNRPD2 | protein_coding | -0.2818032 | 8.90721108 |
| DLL4 | protein_coding | -0.2818767 | 3.90262996 |
| DCK | protein_coding | -0.2819061 | 5.05419452 |
| FBXO33 | protein_coding | -0.2819939 | 4.09647243 |
| UCN | protein_coding | -0.2819962 | 0.35104586 |
| TANGO2 | protein_coding | -0.2820907 | 3.75068593 |
| AHCTF1 | protein_coding | -0.2821368 | 6.71217092 |
| CTC-575N7.1 | lncRNA | -0.2822953 | 0.46432729 |
| RP11-155O18 | lncRNA | -0.2824544 | 1.04477316 |
| MBOAT2 | protein_coding | -0.2827681 | 7.55633207 |
| BIK | protein_coding | -0.2828847 | 2.00094308 |
| EVPL | protein_coding | -0.2828951 | 0.55731362 |
| RP11-95D17. | lncRNA | -0.2830026 | 1.48792884 |
| STT3B | protein_coding | -0.2830099 | 8.3446195 |
| LINC01128 | lncRNA | -0.2830314 | 3.70911805 |
| TIAM1 | protein_coding | -0.28311 | 7.24653044 |
| RLF | protein_coding | -0.283202 | 5.91322096 |
| TMEM164 | protein_coding | -0.2834397 | 7.86314767 |
| PAM | protein_coding | -0.2834836 | 4.65778633 |

| | | | |
|---|---|---|---|
| PAXX | protein_coding | -0.2835317 | 4.18847901 |
| LIMS1 | protein_coding | -0.2837761 | 5.03579689 |
| EBP | protein_coding | -0.2840604 | 5.26908608 |
| HOOK3 | protein_coding | -0.2841945 | 4.10553103 |
| CLPB | protein_coding | -0.2845725 | 5.95243171 |
| NGB | protein_coding | -0.2845743 | 6.36894556 |
| SURF4 | protein_coding | -0.284744 | 7.97742422 |
| SMC1A | protein_coding | -0.2851088 | 9.57471368 |
| PPP3CC | protein_coding | -0.2851935 | 2.23824021 |
| PRPF19 | protein_coding | -0.285363 | 10.0363243 |
| SOGA3 | protein_coding | -0.2854803 | 0.49283383 |
| C1orf116 | protein_coding | -0.2857041 | 1.15295094 |
| RRM1 | protein_coding | -0.2858108 | 9.93274958 |
| ATPAF1 | protein_coding | -0.2858657 | 6.8723507 |
| TBCD | protein_coding | -0.2859123 | 8.59772375 |
| CTC-429P9.3 | lncRNA | -0.28596 | 1.47464025 |
| PYROXD2 | protein_coding | -0.2859894 | 0.8998027 |
| SEH1L | protein_coding | -0.2860318 | 8.25965531 |
| JKAMP | protein_coding | -0.2860941 | 5.90317642 |
| MAPK7 | protein_coding | -0.2861908 | 4.79695604 |
| C16orf70 | protein_coding | -0.2862608 | 5.40211148 |
| ERGIC1 | protein_coding | -0.2865457 | 5.47562866 |
| RP11-791G1! | lncRNA | -0.2868739 | 0.35847785 |
| GMPS | protein_coding | -0.2869431 | 8.68079281 |
| MCM2 | protein_coding | -0.2871716 | 10.0081939 |
| OTUD3 | protein_coding | -0.2872363 | 5.88794256 |
| RP1-317E23. | lncRNA | -0.2876913 | 0.38327613 |
| PTAR1 | protein_coding | -0.2877971 | 5.22130936 |
| UQCR10 | protein_coding | -0.2878978 | 7.85330375 |
| DNAJC12 | protein_coding | -0.2880123 | 1.49240539 |
| RP13-516M1 | lncRNA | -0.2880497 | 0.85220886 |
| LINGO2 | protein_coding | -0.2881111 | 2.15236358 |
| DFFA | protein_coding | -0.2882604 | 5.74505443 |
| TMEM8B | protein_coding | -0.2883331 | 3.23729319 |
| C8orf82 | protein_coding | -0.2883834 | 4.25580343 |
| NFYC-AS1 | lncRNA | -0.2884302 | 1.63595675 |
| CTC-353G13. | lncRNA | -0.2884819 | 0.53253875 |
| TECPR1 | protein_coding | -0.2884916 | 6.36627416 |
| SLC44A2 | protein_coding | -0.2885154 | 1.68906847 |
| G3BP2 | protein_coding | -0.2890921 | 8.93162942 |
| RP11-379L12 | lncRNA | -0.2891922 | 4.07814599 |
| CYTH3 | protein_coding | -0.2892532 | 5.50276144 |
| DGCR2 | protein_coding | -0.2892855 | 7.64748181 |
| ERI3 | protein_coding | -0.2895538 | 6.63653592 |
| PGP | protein_coding | -0.2897543 | 4.97170532 |

| | | | |
|---|---|---|---|
| MXD4 | protein_coding | -0.2898026 | 5.67745494 |
| DLGAP5 | protein_coding | -0.2899751 | 6.96617143 |
| ZFYVE21 | protein_coding | -0.2900714 | 4.37217445 |
| CCNE2 | protein_coding | -0.2902756 | 8.35794556 |
| ANAPC11 | protein_coding | -0.2902801 | 5.12996398 |
| RP11-358L22 | lncRNA | -0.2904095 | 1.30418943 |
| UBE2Q2 | protein_coding | -0.2904135 | 3.81834274 |
| GORASP2 | protein_coding | -0.290491 | 8.29646365 |
| KIF20A | protein_coding | -0.290517 | 6.56484167 |
| STAM | protein_coding | -0.2906234 | 5.22548755 |
| VGLL3 | protein_coding | -0.2906698 | 0.53453686 |
| DCAKD | protein_coding | -0.2906711 | 6.3030172 |
| TMEM168 | protein_coding | -0.2907071 | 5.05578884 |
| GPRIN1 | protein_coding | -0.290751 | 6.42006862 |
| SYNJ1 | protein_coding | -0.2907906 | 7.44045252 |
| CNOT6 | protein_coding | -0.2908711 | 6.65223475 |
| MAGEF1 | protein_coding | -0.2909013 | 5.12917342 |
| SHMT1 | protein_coding | -0.2909695 | 4.94375439 |
| DPY19L4 | protein_coding | -0.2916445 | 5.18258422 |
| NT5C3B | protein_coding | -0.291781 | 5.72326904 |
| ABCD3 | protein_coding | -0.2919987 | 5.8920103 |
| RPGR | protein_coding | -0.2920133 | 2.14428416 |
| ZFP14 | protein_coding | -0.2920415 | 4.75352899 |
| MFSD14B | protein_coding | -0.2923793 | 7.00893539 |
| ZNF687-AS1 | lncRNA | -0.2926547 | 0.40328243 |
| GABPA | protein_coding | -0.2926844 | 4.4276967 |
| PANK4 | protein_coding | -0.2927432 | 4.24003439 |
| SSR4 | protein_coding | -0.2928761 | 7.69598827 |
| FKBP7 | protein_coding | -0.2928884 | 3.89769326 |
| PMPCA | protein_coding | -0.2929705 | 7.09279826 |
| UPF1 | protein_coding | -0.2930698 | 7.62246485 |
| KLF13 | protein_coding | -0.2931351 | 9.40761878 |
| TBX6 | protein_coding | -0.2931521 | 1.35895096 |
| INCENP | protein_coding | -0.2932943 | 6.69829854 |
| GSTZ1 | protein_coding | -0.2934012 | 5.46975331 |
| CTA-204B4.2 | lncRNA | -0.2935495 | 1.7442875 |
| LMNB1 | protein_coding | -0.2936798 | 9.75337741 |
| AGO1 | protein_coding | -0.2937003 | 8.25450501 |
| PPM1F-AS1 | lncRNA | -0.2937314 | 0.77984391 |
| IGSF3 | protein_coding | -0.2937389 | 1.3675347 |
| STARD9 | protein_coding | -0.293858 | 6.0636544 |
| ANKEF1 | protein_coding | -0.2939305 | 6.7508973 |
| SLC11A1 | protein_coding | -0.2939441 | 0.4219421 |
| ELOVL1 | protein_coding | -0.2940438 | 4.75888385 |
| TELO2 | protein_coding | -0.2941631 | 3.46857685 |

| | | | |
|---|---|---|---|
| RASD2 | protein_coding | -0.2943212 | 8.30967761 |
| GPD2 | protein_coding | -0.2943598 | 8.07341876 |
| SLC39A7 | protein_coding | -0.2944082 | 8.7634263 |
| SIGIRR | protein_coding | -0.2945217 | 3.75938106 |
| RP11-61K9.3 | lncRNA | -0.2945775 | 0.95390553 |
| SLC2A1 | protein_coding | -0.2946537 | 9.30000759 |
| PTMA | protein_coding | -0.2947155 | 10.1986556 |
| TLX3 | protein_coding | -0.2947693 | 1.98279105 |
| PTPN3 | protein_coding | -0.2948502 | 6.17812408 |
| AK4 | protein_coding | -0.2948607 | 7.33570417 |
| MAP6D1 | protein_coding | -0.2950203 | 0.27399998 |
| MRE11 | protein_coding | -0.2950956 | 5.6204221 |
| ZNF76 | protein_coding | -0.2952754 | 5.92187974 |
| PLK1 | protein_coding | -0.2953367 | 7.5343426 |
| GUCY1A2 | protein_coding | -0.2955701 | 3.38868415 |
| KAZALD1 | protein_coding | -0.2956597 | 0.81162157 |
| CD320 | protein_coding | -0.2958688 | 5.57346314 |
| ZNF230-DT | lncRNA | -0.2960106 | 0.45131768 |
| ZNF837 | protein_coding | -0.2962336 | 1.17641544 |
| TMEM107 | protein_coding | -0.2965533 | 1.17009898 |
| RP11-656D1( | lncRNA | -0.2966191 | 0.57747453 |
| BTG3-AS1 | lncRNA | -0.296738 | 0.73839843 |
| PTOV1-AS1 | lncRNA | -0.2967414 | 1.21025856 |
| C1QL2 | protein_coding | -0.2970998 | 1.0696284 |
| SSNA1 | protein_coding | -0.2971166 | 5.27814477 |
| MCM6 | protein_coding | -0.2971615 | 9.52790846 |
| NEDD8 | protein_coding | -0.2972334 | 4.44622106 |
| LHX1 | protein_coding | -0.2973473 | 2.77542837 |
| KIF2C | protein_coding | -0.2974435 | 7.50385779 |
| RPS20 | protein_coding | -0.2977069 | 8.5188283 |
| RP11-13K12. | lncRNA | -0.2980115 | 1.18365365 |
| ANXA5 | protein_coding | -0.2980396 | 9.43701823 |
| OLMALINC | lncRNA | -0.2980539 | 3.41050468 |
| ASAP2 | protein_coding | -0.2981171 | 5.90710284 |
| UBAC2-AS1 | lncRNA | -0.298231 | 1.03405378 |
| PHKA2 | protein_coding | -0.2984228 | 7.38001958 |
| ACYP1 | protein_coding | -0.2984548 | 5.75344341 |
| CORO7 | protein_coding | -0.2985619 | 3.34298366 |
| LINC00638 | lncRNA | -0.298578 | 2.52301584 |
| AC090587.4 | lncRNA | -0.2986796 | 0.97960607 |
| RPL13A | protein_coding | -0.29876 | 10.0521561 |
| DSTN | protein_coding | -0.2987725 | 8.89317763 |
| CDS1 | protein_coding | -0.2989931 | 2.47733482 |
| CETN3 | protein_coding | -0.2990725 | 5.44874505 |
| RNF208 | protein_coding | -0.2991392 | 2.59228668 |

| | | | |
|---|---|---|---|
| PSMD10 | protein_coding | -0.299392 | 4.9400249 |
| RABL6 | protein_coding | -0.2994052 | 7.22124764 |
| MSH2 | protein_coding | -0.2994183 | 9.56542482 |
| ZNF487 | protein_coding | -0.2994399 | 0.45044133 |
| JPH1 | protein_coding | -0.2994826 | 3.63023596 |
| KCNN4 | protein_coding | -0.299582 | 3.79405355 |
| LRWD1 | protein_coding | -0.2996327 | 4.16514693 |
| ARPC4 | protein_coding | -0.2996763 | 5.68224753 |
| PAGR1 | protein_coding | -0.299772 | 4.45314794 |
| RAB11FIP3 | protein_coding | -0.2999523 | 5.83211869 |
| GPR26 | protein_coding | -0.3002422 | 1.35372136 |
| RP11-384O8. | protein_coding | -0.3004074 | 0.54332203 |
| DENND6A | protein_coding | -0.3004086 | 5.49788846 |
| TSNARE1 | protein_coding | -0.3009327 | 3.5401057 |
| UBXN11 | protein_coding | -0.3010461 | 4.00333063 |
| MEGF9 | protein_coding | -0.3010722 | 8.49256924 |
| LYRM9 | protein_coding | -0.3011713 | 2.10835273 |
| C2orf74 | protein_coding | -0.3013376 | 0.41363434 |
| SMIM10L2B | protein_coding | -0.301368 | 1.11301254 |
| ZMYND19 | protein_coding | -0.3014627 | 6.18934448 |
| IQANK1 | protein_coding | -0.3015434 | 0.66454065 |
| TEX264 | protein_coding | -0.3015645 | 4.99901462 |
| MBD3 | protein_coding | -0.3015675 | 5.23703395 |
| STK26 | protein_coding | -0.3016934 | 5.15157363 |
| XPR1 | protein_coding | -0.3017877 | 8.09058593 |
| PIGP | protein_coding | -0.3022086 | 4.59649879 |
| CTD-2203K17 | lncRNA | -0.3024547 | 1.41784216 |
| C2orf15 | protein_coding | -0.3028534 | 1.82818494 |
| SATL1 | protein_coding | -0.3028566 | 2.10005138 |
| ROMO1 | protein_coding | -0.3029751 | 6.44020062 |
| TEAD4 | protein_coding | -0.3029886 | 4.3087859 |
| SEM1 | protein_coding | -0.3029913 | 7.40089603 |
| FBXO36 | protein_coding | -0.3031134 | 2.55813139 |
| TC2N | protein_coding | -0.3032007 | 5.67065063 |
| CDHR1 | protein_coding | -0.3032134 | 3.28040353 |
| LCMT2 | protein_coding | -0.3033724 | 6.17876833 |
| HIC2 | protein_coding | -0.3034235 | 4.23302876 |
| TBC1D2 | protein_coding | -0.3034927 | 2.23487166 |
| POLH | protein_coding | -0.303512 | 6.65349802 |
| RDM1 | protein_coding | -0.3035307 | 1.25797469 |
| EIF3L | protein_coding | -0.303789 | 9.23582144 |
| BAG4 | protein_coding | -0.3038546 | 3.59525164 |
| FSBP | protein_coding | -0.3041135 | 2.00752158 |
| TMEM181 | protein_coding | -0.3041656 | 6.62925481 |
| RP11-467D6. | lncRNA | -0.3041758 | 0.58527047 |

| | | | |
|---|---|---|---|
| RP11-216L13 | lncRNA | -0.304205 | 1.139401 |
| SLC25A35 | protein_coding | -0.3044807 | 2.28314923 |
| PPP1R12A-AS | lncRNA | -0.3045287 | 0.751552 |
| CTB-129O4.1 | lncRNA | -0.3047195 | 0.54314142 |
| C9orf43 | protein_coding | -0.3049012 | 0.79580722 |
| NDUFC2 | protein_coding | -0.3050584 | 3.18378124 |
| CYP46A1 | protein_coding | -0.305135 | 3.43314017 |
| TEX29 | protein_coding | -0.3051937 | 0.50927347 |
| MMACHC | protein_coding | -0.3052516 | 4.98203098 |
| CDKN2AIPNL | protein_coding | -0.3053337 | 5.3891992 |
| NETO2 | protein_coding | -0.3053344 | 8.56745121 |
| AC069513.4 | lncRNA | -0.3053389 | 0.5841918 |
| IQGAP3 | protein_coding | -0.3054296 | 4.5515013 |
| FGD5-AS1 | lncRNA | -0.3056783 | 8.60423922 |
| RNF5 | protein_coding | -0.3057343 | 6.77079373 |
| MITF | protein_coding | -0.3057679 | 1.1599086 |
| TNFRSF13C | protein_coding | -0.305794 | 2.53856358 |
| CANX | protein_coding | -0.3058119 | 9.92122881 |
| CRLS1 | protein_coding | -0.3058742 | 0.67415974 |
| ZNF23 | protein_coding | -0.3059041 | 0.82685113 |
| CTDSP1 | protein_coding | -0.3059416 | 6.73279735 |
| GVQW3 | protein_coding | -0.3059801 | 5.3431888 |
| PFDN6 | protein_coding | -0.3060155 | 5.65869263 |
| RPS8 | protein_coding | -0.3061112 | 10.5748256 |
| CIRBP-AS1 | lncRNA | -0.3063996 | 1.33233064 |
| BCL2L11 | protein_coding | -0.306426 | 7.14375314 |
| MTCH1 | protein_coding | -0.3066206 | 10.190092 |
| GAS2L1 | protein_coding | -0.3068023 | 2.5635245 |
| CCDC85C | protein_coding | -0.30706 | 8.79141631 |
| CASP9 | protein_coding | -0.3070643 | 3.06346062 |
| RUVBL2 | protein_coding | -0.3072158 | 7.16962612 |
| LRP2BP | protein_coding | -0.3072521 | 1.01674736 |
| BHLHA15 | protein_coding | -0.3072913 | 2.1657233 |
| NAA38 | protein_coding | -0.3076476 | 3.74421154 |
| SINHCAF | protein_coding | -0.3077566 | 9.55836793 |
| C10orf95-AS | lncRNA | -0.3078533 | 6.15090852 |
| F8A1 | protein_coding | -0.3078955 | 3.52015501 |
| RP11-412D9. | lncRNA | -0.3079553 | 2.18692265 |
| RP4-591C20. | lncRNA | -0.308053 | 2.02679757 |
| MAPKAPK5-A | lncRNA | -0.3080536 | 4.33097167 |
| TYSND1 | protein_coding | -0.3080934 | 2.65240767 |
| LRFN3 | protein_coding | -0.3082073 | 3.51511609 |
| PIAS2 | protein_coding | -0.3084207 | 6.48031904 |
| SIX5 | protein_coding | -0.3085275 | 2.4116846 |
| RACK1 | protein_coding | -0.3086554 | 11.1980268 |

| | | | |
|---|---|---|---|
| ELMOD2 | protein_coding | -0.3086993 | 5.4269194 |
| NEXMIF | protein_coding | -0.3086995 | 5.32798039 |
| UPRT | protein_coding | -0.3087328 | 3.4029098 |
| HSD17B6 | protein_coding | -0.3088583 | 1.6291796 |
| SLC16A10 | protein_coding | -0.3090823 | 3.44256358 |
| EIF4EBP2 | protein_coding | -0.3092556 | 9.47095591 |
| ELAVL2 | protein_coding | -0.3093613 | 8.03877045 |
| STRIP2 | protein_coding | -0.3096412 | 4.61447651 |
| POLR2E | protein_coding | -0.309756 | 8.56442157 |
| RP11-92K15. | lncRNA | -0.3097936 | 0.40124258 |
| CPT1C | protein_coding | -0.3098144 | 3.4794558 |
| FKBP1C | protein_coding | -0.3099323 | 0.38493023 |
| SNAPC2 | protein_coding | -0.309954 | 4.47093663 |
| AC093388.3 | lncRNA | -0.3100937 | 1.3530728 |
| PDLIM1 | protein_coding | -0.3101225 | 5.11318015 |
| HRAT92 | lncRNA | -0.3102135 | 0.43653059 |
| PGAP4 | protein_coding | -0.3102315 | 7.40078661 |
| DARS1-AS1 | lncRNA | -0.3102558 | 0.83527471 |
| CCND3 | protein_coding | -0.3103877 | 7.50078783 |
| CTD-3051D2 | lncRNA | -0.3106803 | 0.45378166 |
| CBLN1 | protein_coding | -0.3108451 | 5.40546128 |
| CDC14B | protein_coding | -0.3108886 | 1.516538 |
| RPA2 | protein_coding | -0.3108889 | 9.40897725 |
| RP1-257A7.5 | lncRNA | -0.3109119 | 0.57002127 |
| UBE2N | protein_coding | -0.31093 | 8.15184333 |
| IGSF21 | protein_coding | -0.3110144 | 2.35131251 |
| SNTA1 | protein_coding | -0.3110602 | 5.37225012 |
| MRPL33 | protein_coding | -0.3112234 | 5.7329054 |
| UBE2S | protein_coding | -0.3114486 | 2.77471292 |
| SP5 | protein_coding | -0.3115543 | 0.50008053 |
| SHPK | protein_coding | -0.3116305 | 0.98172 |
| HDAC7 | protein_coding | -0.311631 | 4.41340859 |
| RP11-147L1 | lncRNA | -0.3116462 | 0.63301601 |
| LL21NC02-1 | lncRNA | -0.3117914 | 0.43734801 |
| GOLGA8R | protein_coding | -0.3118135 | 0.57151617 |
| ZNF32 | protein_coding | -0.3120928 | 3.19010823 |
| CPEB3 | protein_coding | -0.3121652 | 2.79950217 |
| BEX4 | protein_coding | -0.3121976 | 5.63293319 |
| FAM111B | protein_coding | -0.3122427 | 9.13702461 |
| FRG1-DT | lncRNA | -0.3122706 | 0.50585031 |
| CELF4 | protein_coding | -0.3123366 | 6.32991484 |
| PHETA2 | protein_coding | -0.3123605 | 3.77616641 |
| ARHGAP33 | protein_coding | -0.3123955 | 7.45376758 |
| REP15 | protein_coding | -0.3125333 | 0.51151403 |
| KHK | protein_coding | -0.3125745 | 5.62803851 |

| | | | |
|---|---|---|---|
| A4GALT | protein_coding | -0.312701 | 1.85595828 |
| RP11-179B2. | lncRNA | -0.312781 | 0.58814207 |
| BCL2L15 | protein_coding | -0.3127861 | 0.64258241 |
| RP11-304L19 | lncRNA | -0.3128608 | 0.37793682 |
| HNRNPH2 | protein_coding | -0.312906 | 6.79079182 |
| CEP41 | protein_coding | -0.3129126 | 5.86194886 |
| TRIM36 | protein_coding | -0.3129223 | 3.48020426 |
| NUDT2 | protein_coding | -0.3130388 | 2.9078345 |
| FAM207A | protein_coding | -0.313292 | 3.15775491 |
| CRIP1 | protein_coding | -0.3136183 | 1.28306513 |
| RMI1 | protein_coding | -0.3136189 | 6.28392551 |
| ZNF385A | protein_coding | -0.3136608 | 10.3495129 |
| KIFC1 | protein_coding | -0.3138175 | 8.70346195 |
| HEXIM2 | protein_coding | -0.3139304 | 1.84667329 |
| MRPL43 | protein_coding | -0.3140172 | 5.14854839 |
| FAM72D | protein_coding | -0.314248 | 2.00071472 |
| ZNF680 | protein_coding | -0.3142856 | 3.96784011 |
| SLC25A15 | protein_coding | -0.3143849 | 3.07240826 |
| PCLAF | protein_coding | -0.3146694 | 1.4684261 |
| CCDC40 | protein_coding | -0.3147832 | 4.45104134 |
| ATP5F1D | protein_coding | -0.3147947 | 3.08352778 |
| RPA1 | protein_coding | -0.3148935 | 9.23309593 |
| DMAC2 | protein_coding | -0.3153812 | 7.08608902 |
| SPCS3 | protein_coding | -0.3156066 | 6.30046051 |
| SPIRE1 | protein_coding | -0.3156076 | 1.57353023 |
| MLF1 | protein_coding | -0.3156272 | 4.52316587 |
| MAGED1 | protein_coding | -0.3156352 | 10.225065 |
| RPS16 | protein_coding | -0.3156723 | 10.1400928 |
| RP11-452L6. | lncRNA | -0.3156974 | 2.62776449 |
| RP11-797A18 | lncRNA | -0.3159762 | 0.95253297 |
| KCND2 | protein_coding | -0.3160038 | 7.02636997 |
| KIF11 | protein_coding | -0.3161577 | 8.16222511 |
| GEMIN4 | protein_coding | -0.3162918 | 6.79155956 |
| BCL11B | protein_coding | -0.3163612 | 8.1522982 |
| ACAA2 | protein_coding | -0.3163711 | 7.63718805 |
| RP11-395A13 | lncRNA | -0.3163905 | 0.61644058 |
| LRRC34 | protein_coding | -0.3164049 | 1.5064194 |
| ATN1 | protein_coding | -0.316424 | 8.13223109 |
| RILP | protein_coding | -0.3166055 | 0.99219094 |
| ARMCX5-GPI | lncRNA | -0.3166306 | 2.36315439 |
| SPSB1 | protein_coding | -0.3167678 | 2.89256249 |
| LGR6 | protein_coding | -0.3168811 | 2.58723477 |
| LIG1 | protein_coding | -0.3169616 | 7.50339564 |
| DEDD2 | protein_coding | -0.3170294 | 5.00309209 |
| ADAMTS8 | protein_coding | -0.3171407 | 1.16698171 |

| | | | |
|---|---|---|---|
| FTCDNL1 | protein_coding | -0.3172759 | 0.87509772 |
| MAZ | protein_coding | -0.3173932 | 9.25299082 |
| TCEAL3 | protein_coding | -0.317464 | 3.57199165 |
| IFT88 | protein_coding | -0.3175116 | 1.55056137 |
| TMEM167A | protein_coding | -0.3175517 | 5.77311988 |
| NUBP2 | protein_coding | -0.3177929 | 7.26028767 |
| MGAT1 | protein_coding | -0.3178168 | 6.45129561 |
| ERCC6L | protein_coding | -0.3178467 | 3.38284983 |
| MPI | protein_coding | -0.3178879 | 6.37477789 |
| SUPT4H1 | protein_coding | -0.3179374 | 0.54223973 |
| DLX4 | protein_coding | -0.3179504 | 9.13022267 |
| CDK2 | protein_coding | -0.3184041 | 9.24854726 |
| MIAT | lncRNA | -0.3185264 | 10.8280888 |
| SLC36A4 | protein_coding | -0.3185885 | 7.24873836 |
| ANKRD50 | protein_coding | -0.3188915 | 8.23785929 |
| C11orf1 | protein_coding | -0.3190412 | 2.30015531 |
| ZBTB20 | protein_coding | -0.319084 | 0.35462304 |
| SLC23A2 | protein_coding | -0.3192651 | 7.79332871 |
| MRPL52 | protein_coding | -0.3193992 | 6.05619633 |
| ATP6V1FNB | protein_coding | -0.3194199 | 1.78322482 |
| LAMTOR5 | protein_coding | -0.3194236 | 5.8190748 |
| PLA2G15 | protein_coding | -0.3196183 | 4.06534447 |
| ASMTL | protein_coding | -0.31997 | 3.62377512 |
| AURKC | protein_coding | -0.3200076 | 0.47260339 |
| HAUS4 | protein_coding | -0.3200613 | 3.28666849 |
| GOLGA8A | protein_coding | -0.3200762 | 6.14998008 |
| C8orf44 | lncRNA | -0.3201775 | 0.91203857 |
| BCL6B | protein_coding | -0.3203115 | 7.23581924 |
| EEPD1 | protein_coding | -0.3203762 | 4.22453534 |
| PDS5B | protein_coding | -0.3204303 | 6.96322216 |
| EGFL7 | protein_coding | -0.3204925 | 2.11418105 |
| DYNC2I2 | protein_coding | -0.3204986 | 6.56109622 |
| TUBA1B | protein_coding | -0.3205974 | 12.1253578 |
| ATP8A1 | protein_coding | -0.3206393 | 8.32517519 |
| TMEFF1 | protein_coding | -0.3206555 | 0.43528775 |
| VASP | protein_coding | -0.3208085 | 6.60840487 |
| RP11-323F24 | lncRNA | -0.3209974 | 1.35517917 |
| CRYBA2 | protein_coding | -0.3210654 | 2.37615566 |
| DCTN3 | protein_coding | -0.3210834 | 7.68260716 |
| CNBD2 | protein_coding | -0.3211637 | 1.39540058 |
| GS1-72M22.1 | lncRNA | -0.3212391 | 2.99725098 |
| DYDC2 | protein_coding | -0.3212456 | 0.66224795 |
| DET1 | protein_coding | -0.3215553 | 0.47133021 |
| GSDMD | protein_coding | -0.3216474 | 1.57421024 |
| LIPE-AS1 | lncRNA | -0.3217082 | 1.05348395 |

| | | | |
|---|---|---|---|
| HSD17B8 | protein_coding | -0.3217624 | 2.81713548 |
| RP11-458J1.1 | lncRNA | -0.3217844 | 2.2850505 |
| RP6-65G23.3 | lncRNA | -0.321844 | 2.95424077 |
| RP11-146F11 | lncRNA | -0.3218605 | 0.41783424 |
| UQCR11 | protein_coding | -0.3219807 | 4.54324636 |
| CTD-2001C12 | lncRNA | -0.322082 | 0.40837601 |
| PLAUR | protein_coding | -0.3221303 | 3.55419338 |
| PSENEN | protein_coding | -0.3222291 | 2.73692918 |
| FGFBP3 | protein_coding | -0.3222578 | 1.25497626 |
| TMCO1-AS1 | lncRNA | -0.3222979 | 0.7936402 |
| CNTNAP3B | protein_coding | -0.3223032 | 1.63231744 |
| INKA2 | protein_coding | -0.3224054 | 6.21554829 |
| RPS28 | protein_coding | -0.3224242 | 5.67983358 |
| AFF3 | protein_coding | -0.3224877 | 0.36252409 |
| ANKRD65 | protein_coding | -0.3226765 | 2.80142707 |
| VTA1 | protein_coding | -0.3227838 | 6.7818513 |
| TBC1D8-AS1 | lncRNA | -0.3229827 | 1.19960766 |
| GMPPA | protein_coding | -0.3230203 | 7.52758431 |
| AC092198.1 | lncRNA | -0.323027 | 0.40095978 |
| SMIM14 | protein_coding | -0.3231336 | 5.2220216 |
| ZBTB12 | protein_coding | -0.3232104 | 3.62389482 |
| BAALC | protein_coding | -0.3233198 | 1.01155429 |
| ATCAY | protein_coding | -0.32336 | 0.37949295 |
| RPL7 | protein_coding | -0.3233969 | 9.04162802 |
| ARMCX4 | protein_coding | -0.3235058 | 1.19431743 |
| ENDOG | protein_coding | -0.3235594 | 3.71944415 |
| PLEKHB1 | protein_coding | -0.3235968 | 3.87899759 |
| RP11-46C24. | lncRNA | -0.3238246 | 1.65176391 |
| CDIPT | protein_coding | -0.3239601 | 7.97985705 |
| SCX | protein_coding | -0.3239719 | 0.53928472 |
| SGO2 | protein_coding | -0.3239856 | 6.02888929 |
| NISCH | protein_coding | -0.3240978 | 8.47736992 |
| GS1-279B7.2 | lncRNA | -0.3241148 | 0.82482632 |
| NUDT16L1 | protein_coding | -0.3245465 | 4.94450713 |
| HAGH | protein_coding | -0.3245803 | 6.4118217 |
| RP3-402G11. | lncRNA | -0.3246262 | 1.41284428 |
| RP6-201G10. | lncRNA | -0.3249765 | 0.46205034 |
| PIEZO2 | protein_coding | -0.3250335 | 8.56572589 |
| MRPL20-DT | lncRNA | -0.3252691 | 2.14145832 |
| CLN8 | protein_coding | -0.3255043 | 2.45105121 |
| FGF12 | protein_coding | -0.3255103 | 5.7823023 |
| IQCG | protein_coding | -0.325561 | 3.28445734 |
| CACFD1 | protein_coding | -0.325574 | 6.55273507 |
| CAP1 | protein_coding | -0.3256216 | 9.72757338 |
| CERS1 | protein_coding | -0.325704 | 1.78855256 |

| | | | |
|---|---|---|---|
| CDKN2C | protein_coding | -0.325704 | 7.82768686 |
| UBE2D3-AS1 | lncRNA | -0.3257118 | 0.64138847 |
| GLMP | protein_coding | -0.3257318 | 3.04870146 |
| PTDSS1 | protein_coding | -0.3258297 | 9.2915314 |
| PODXL2 | protein_coding | -0.3258759 | 7.69585341 |
| TMEM44-AS | lncRNA | -0.3259959 | 1.9838549 |
| PRR34-AS1 | lncRNA | -0.3261505 | 1.41859752 |
| LMAN2 | protein_coding | -0.326266 | 8.3600507 |
| NAB2 | protein_coding | -0.3264756 | 4.38316652 |
| RP5-940J5.3 | lncRNA | -0.3267772 | 0.62213181 |
| PGGHG | protein_coding | -0.3267886 | 5.18715475 |
| LINC00659 | lncRNA | -0.3269599 | 1.32643198 |
| CFDP1 | protein_coding | -0.3276844 | 7.01685123 |
| SSX2IP | protein_coding | -0.327862 | 9.19924913 |
| ARMH1 | protein_coding | -0.3282002 | 0.74407589 |
| NUDT12 | protein_coding | -0.3282464 | 3.82388355 |
| NDC1 | protein_coding | -0.3284599 | 8.27085765 |
| KIF18B | protein_coding | -0.3285337 | 7.98986404 |
| ATP5F1E | protein_coding | -0.3286233 | 9.09100846 |
| SLC41A3 | protein_coding | -0.3286438 | 6.40616812 |
| GPR143 | protein_coding | -0.3286552 | 0.78523642 |
| MGST3 | protein_coding | -0.3286698 | 6.71898334 |
| LINC02693 | lncRNA | -0.3287663 | 2.58506811 |
| DIPK1B | protein_coding | -0.3289197 | 5.03686657 |
| MED18 | protein_coding | -0.3289762 | 4.12867151 |
| MFNG | protein_coding | -0.3295912 | 7.02649051 |
| ZNF449 | protein_coding | -0.3297572 | 1.64065631 |
| SMAD5 | protein_coding | -0.3300442 | 6.55696171 |
| MAP10 | protein_coding | -0.3301359 | 1.30335025 |
| ZNHIT3 | protein_coding | -0.3302852 | 6.29455812 |
| SPAST | protein_coding | -0.3303929 | 7.08350577 |
| KDELR1 | protein_coding | -0.3304747 | 9.6125672 |
| SHMT2 | protein_coding | -0.3304879 | 8.56952986 |
| AE000661.37 | lncRNA | -0.3305585 | 0.81328368 |
| GOT2 | protein_coding | -0.330662 | 9.83207048 |
| DLGAP3 | protein_coding | -0.3307402 | 4.49814673 |
| HRAS | protein_coding | -0.3309403 | 7.65907953 |
| FRMPD1 | protein_coding | -0.3309494 | 1.16346035 |
| SLC15A4 | protein_coding | -0.3313563 | 6.4340795 |
| APRT | protein_coding | -0.3318399 | 6.50040417 |
| DEPDC7 | protein_coding | -0.3318472 | 4.23467432 |
| MAP2K3 | protein_coding | -0.3318739 | 5.79225089 |
| ATP6V0D1 | protein_coding | -0.332116 | 7.49842725 |
| HYAL3 | protein_coding | -0.332367 | 2.26162268 |
| PHF7 | protein_coding | -0.332523 | 2.34246883 |

| | | | |
|---|---|---|---|
| FURIN | protein_coding | -0.332567 | 8.0296654 |
| MYRFL | protein_coding | -0.3326399 | 0.76920064 |
| ZFP62 | protein_coding | -0.3329832 | 6.66491475 |
| LL09NC01-13 | lncRNA | -0.3330487 | 0.41333827 |
| PPARG | protein_coding | -0.3335047 | 1.14104673 |
| BRIP1 | protein_coding | -0.3337205 | 5.9520105 |
| NPPA | protein_coding | -0.33389 | 5.34555979 |
| HOXC6 | protein_coding | -0.3339888 | 2.06339101 |
| GPR6 | protein_coding | -0.3340203 | 2.00302519 |
| CTD-2366F13 | lncRNA | -0.3340865 | 0.9151413 |
| SKI | protein_coding | -0.3341426 | 5.68647476 |
| GYS1 | protein_coding | -0.3341731 | 6.31069513 |
| CETN2 | protein_coding | -0.3341929 | 5.10066304 |
| RP3-438D16. | lncRNA | -0.3342549 | 0.41948785 |
| ALDH6A1 | protein_coding | -0.3342561 | 6.65424606 |
| ZNF511 | protein_coding | -0.3343152 | 2.47393885 |
| FAM104B | protein_coding | -0.3343625 | 4.6585354 |
| CDK1 | protein_coding | -0.3345822 | 6.62417462 |
| RP5-1198O2( | lncRNA | -0.3349136 | 0.66894917 |
| B4GAT1 | protein_coding | -0.3349579 | 4.83033471 |
| TMEM160 | protein_coding | -0.3349782 | 0.74787361 |
| NDUFAF8 | protein_coding | -0.3350558 | 4.0268241 |
| TBC1D9B | protein_coding | -0.3351847 | 6.91051481 |
| SP2-DT | lncRNA | -0.335246 | 1.8008842 |
| ARHGEF19 | protein_coding | -0.3353678 | 6.78544149 |
| SLC16A7 | protein_coding | -0.335567 | 3.52300837 |
| DYNC2LI1 | protein_coding | -0.3359463 | 3.49060423 |
| RP1-179N16. | lncRNA | -0.3360446 | 0.60704523 |
| FAM167B | protein_coding | -0.3360986 | 0.63038062 |
| ZCCHC17 | protein_coding | -0.3361816 | 6.64154234 |
| HSBP1L1 | protein_coding | -0.3362739 | 0.56139374 |
| CPEB4 | protein_coding | -0.3368499 | 5.78534822 |
| DOP1B | protein_coding | -0.3369295 | 8.16130039 |
| FAM107B | protein_coding | -0.3373974 | 1.94924598 |
| MTPN | protein_coding | -0.3374723 | 7.88770465 |
| LGALS4 | protein_coding | -0.3374943 | 0.40347687 |
| CYP2U1 | protein_coding | -0.337738 | 3.17480333 |
| HSBP1 | protein_coding | -0.3379704 | 9.98009316 |
| DNPH1 | protein_coding | -0.3380035 | 4.32816308 |
| NHLH2 | protein_coding | -0.3380103 | 3.05146329 |
| PPP1R35-AS1 | lncRNA | -0.3380203 | 0.37324889 |
| GTF2A1 | protein_coding | -0.3380829 | 6.76088842 |
| BCL2L10 | protein_coding | -0.3383188 | 0.75276318 |
| CAVIN4 | protein_coding | -0.3383995 | 2.7600325 |
| RP11-345P4. | lncRNA | -0.3386863 | 2.07283747 |

| | | | |
|---|---|---|---|
| TCAF1 | protein_coding | -0.3390229 | 6.64248267 |
| GRIN2D | protein_coding | -0.3391322 | 2.13680686 |
| TRMT112 | protein_coding | -0.339356 | 8.56751064 |
| PITPNM2 | protein_coding | -0.3398438 | 7.41230507 |
| RP1-198K11. | lncRNA | -0.3398546 | 0.87819786 |
| CTD-2020K17 | lncRNA | -0.3399095 | 1.56595174 |
| RUNX1T1 | protein_coding | -0.339962 | 6.81632933 |
| GLB1L | protein_coding | -0.340094 | 1.44192411 |
| CAPS | protein_coding | -0.3401339 | 1.00535306 |
| TXNDC12 | protein_coding | -0.3403433 | 2.15006451 |
| POT1-AS1 | lncRNA | -0.3405038 | 0.44895816 |
| MIR600HG | lncRNA | -0.3405264 | 4.51571786 |
| PAICS | protein_coding | -0.3405959 | 10.9463473 |
| INPP5K | protein_coding | -0.3406875 | 5.65134517 |
| TRAPPC2L | protein_coding | -0.3407201 | 5.59082219 |
| CTD-3064H18 | lncRNA | -0.3411895 | 0.82304352 |
| SLCO4A1 | protein_coding | -0.3412029 | 4.26257893 |
| PNP | protein_coding | -0.3413931 | 8.36235589 |
| UROD | protein_coding | -0.341397 | 6.85815765 |
| MGAT5 | protein_coding | -0.3414536 | 8.25885623 |
| PKNOX1 | protein_coding | -0.3415899 | 6.83770889 |
| TLE1 | protein_coding | -0.3417214 | 7.49259695 |
| DPYSL5 | protein_coding | -0.3418094 | 10.5457679 |
| SLC35C1 | protein_coding | -0.34189 | 4.67601269 |
| C21orf62-AS | lncRNA | -0.3419008 | 1.33412394 |
| SCUBE2 | protein_coding | -0.3419498 | 0.49061573 |
| CEP131 | protein_coding | -0.3419936 | 3.9068242 |
| ASPH | protein_coding | -0.3421727 | 6.75386472 |
| RNF122 | protein_coding | -0.3422139 | 2.21403402 |
| TLCD3A | protein_coding | -0.3422238 | 6.64685748 |
| DBP | protein_coding | -0.3426743 | 4.31196028 |
| LNP1 | protein_coding | -0.3427363 | 2.3965097 |
| SACS | protein_coding | -0.3429397 | 7.69227262 |
| NAPSA | protein_coding | -0.343167 | 0.53302717 |
| HOXA11-AS | lncRNA | -0.3432602 | 0.51784395 |
| VKORC1L1 | protein_coding | -0.3433255 | 7.06436998 |
| OSBPL3 | protein_coding | -0.3434859 | 8.4139126 |
| CTC-429P9.5 | lncRNA | -0.343607 | 1.08550503 |
| MPC2 | protein_coding | -0.3436452 | 8.07508451 |
| LYPD3 | protein_coding | -0.3436886 | 3.94772236 |
| TIMP2 | protein_coding | -0.3439163 | 7.37238027 |
| SHC2 | protein_coding | -0.3441202 | 4.89924641 |
| SOWAHA | protein_coding | -0.3444572 | 5.62606175 |
| AP3B2 | protein_coding | -0.3446105 | 6.99713785 |
| TXNL4A | protein_coding | -0.3447504 | 7.61653475 |

| | | | |
|---|---|---|---|
| ARRDC1 | protein_coding | -0.3447813 | 1.88004258 |
| RP11-203J24 | protein_coding | -0.3448692 | 0.5916014 |
| RP11-307L14 | lncRNA | -0.3449504 | 0.47382014 |
| RPLP0 | protein_coding | -0.3450992 | 12.10898 |
| PTPRN | protein_coding | -0.3454313 | 4.22568641 |
| KATNAL1 | protein_coding | -0.3455221 | 7.43530338 |
| LINC01144 | lncRNA | -0.3456086 | 1.72837856 |
| LINC01136 | lncRNA | -0.3456351 | 0.50779889 |
| HNRNPUL1 | protein_coding | -0.3461606 | 9.83615369 |
| RSRC1 | protein_coding | -0.3462546 | 6.65532994 |
| SMC2 | protein_coding | -0.3463412 | 10.2716769 |
| HOXA10 | protein_coding | -0.3467844 | 5.50984732 |
| RP11-216B9. | lncRNA | -0.3470548 | 0.80551587 |
| AGGF1 | protein_coding | -0.3470733 | 3.73320026 |
| AK7 | protein_coding | -0.3472253 | 4.50298258 |
| G6PC3 | protein_coding | -0.3472668 | 5.52553533 |
| IPP | protein_coding | -0.3473206 | 6.22163855 |
| RGS5 | lncRNA | -0.347335 | 1.21697225 |
| CDNF | protein_coding | -0.3473528 | 0.53613916 |
| GMDS | protein_coding | -0.3475244 | 6.69968095 |
| RPL22 | protein_coding | -0.3475918 | 7.05433051 |
| TBL2 | protein_coding | -0.3476144 | 6.69654896 |
| RP11-727F15 | lncRNA | -0.3476289 | 1.72671986 |
| TTC30A | protein_coding | -0.3476364 | 2.69161283 |
| ACTA1 | protein_coding | -0.3477814 | 0.64549248 |
| ATG10 | protein_coding | -0.3480031 | 3.82916843 |
| CHCHD6 | protein_coding | -0.3482176 | 3.93210591 |
| MAPK8IP1 | protein_coding | -0.3484608 | 6.42879668 |
| SMIM8 | protein_coding | -0.3484994 | 5.3036672 |
| INPP5F | protein_coding | -0.348776 | 5.45329123 |
| PDZD8 | protein_coding | -0.3489205 | 7.57236167 |
| TTC8 | protein_coding | -0.3490746 | 4.29270783 |
| ATP5IF1 | protein_coding | -0.3491912 | 9.33030437 |
| RP1-63M2.7 | lncRNA | -0.3493182 | 0.5305507 |
| TEF | protein_coding | -0.3495309 | 6.87221877 |
| PELI3 | protein_coding | -0.3497495 | 5.33493961 |
| MSI2 | protein_coding | -0.3498745 | 9.29280366 |
| MYCL | protein_coding | -0.3499274 | 6.9433662 |
| CABLES1 | protein_coding | -0.3500133 | 6.92913367 |
| CNIH1 | protein_coding | -0.3501635 | 7.55353184 |
| STARD3NL | protein_coding | -0.350224 | 4.67926915 |
| SCNN1D | protein_coding | -0.3502338 | 2.00209744 |
| EPN1 | protein_coding | -0.3502594 | 5.07273948 |
| INKA2-AS1 | lncRNA | -0.3503548 | 1.74476675 |
| CCDC9 | protein_coding | -0.350607 | 4.78942402 |

| | | | |
|---|---|---|---|
| ZADH2 | protein_coding | -0.3508677 | 5.82917083 |
| RP11-52L5.6 | lncRNA | -0.3511279 | 0.51345683 |
| DONSON | protein_coding | -0.3513682 | 7.55950155 |
| DDX41 | protein_coding | -0.3515721 | 7.63245664 |
| CTC-241N9.1 | lncRNA | -0.3516541 | 0.82869295 |
| EVI5L | protein_coding | -0.351885 | 5.05989293 |
| CTC-260E6.6 | lncRNA | -0.3520115 | 1.94044251 |
| SAC3D1 | protein_coding | -0.3521101 | 4.43009605 |
| LINC01346 | lncRNA | -0.3521449 | 2.33535177 |
| MCM4 | protein_coding | -0.3523342 | 10.0744595 |
| DSEL | protein_coding | -0.3523377 | 7.05422743 |
| ARPC1A | protein_coding | -0.3527764 | 5.29203927 |
| SPESP1 | protein_coding | -0.352848 | 2.63786129 |
| PITRM1-AS1 | lncRNA | -0.3532548 | 0.70147315 |
| TRIM13 | protein_coding | -0.3534074 | 5.0466182 |
| LURAP1 | protein_coding | -0.3534799 | 0.7963379 |
| CDCA3 | protein_coding | -0.353539 | 6.67363451 |
| ZNF653 | protein_coding | -0.3535639 | 1.98769291 |
| RP11-73K9.2 | lncRNA | -0.3536441 | 0.83580892 |
| NCAPD2 | protein_coding | -0.3540634 | 11.1254984 |
| SLC25A39 | protein_coding | -0.3541722 | 11.2326946 |
| RP11-438L19 | lncRNA | -0.3542035 | 0.67219027 |
| SLC36A1 | protein_coding | -0.3542513 | 7.09645853 |
| RET | protein_coding | -0.354348 | 3.56792345 |
| PRDM11 | protein_coding | -0.3545525 | 5.09956771 |
| TCF7 | protein_coding | -0.3549265 | 3.87226672 |
| MAPK3 | protein_coding | -0.355025 | 7.8194042 |
| PLIN3 | protein_coding | -0.3551908 | 1.54225274 |
| LRFN4 | protein_coding | -0.3556669 | 4.51240367 |
| KATNAL2 | protein_coding | -0.3558159 | 2.34040296 |
| SNRNP35 | protein_coding | -0.3559121 | 4.99766565 |
| HOXA4 | protein_coding | -0.3561567 | 2.49319055 |
| UXT-AS1 | lncRNA | -0.3562126 | 1.36504697 |
| CINP | protein_coding | -0.3562163 | 7.77305532 |
| ZSWIM4 | protein_coding | -0.3562278 | 4.56398229 |
| CDC6 | protein_coding | -0.3569414 | 8.19955028 |
| MT-ND3 | protein_coding | -0.3570348 | 6.60883639 |
| TP53I13 | protein_coding | -0.3572624 | 3.14741499 |
| FAM117B | protein_coding | -0.3573592 | 7.94114814 |
| BBOF1 | protein_coding | -0.3574206 | 4.12877979 |
| PPP1R32 | protein_coding | -0.3576103 | 1.47679101 |
| MPND | protein_coding | -0.3576618 | 5.59302514 |
| RP11-301O19 | lncRNA | -0.3577181 | 1.15818735 |
| CRTAP | protein_coding | -0.3579541 | 6.10157591 |
| CTD-2201E18 | lncRNA | -0.3581108 | 0.61353464 |

| | | | |
|---|---|---|---|
| PCNT | protein_coding | -0.3582691 | 10.0972997 |
| RNF182 | protein_coding | -0.3584983 | 2.26795311 |
| LRP4 | protein_coding | -0.3586388 | 4.18500863 |
| GNB2 | protein_coding | -0.3587327 | 8.87252773 |
| MXRA7 | protein_coding | -0.3587497 | 4.26799215 |
| ENDOV | protein_coding | -0.3588126 | 5.19795251 |
| STMN1 | protein_coding | -0.3590338 | 12.9929296 |
| CDK2AP2 | protein_coding | -0.3591039 | 6.3480679 |
| KANK4 | protein_coding | -0.3598745 | 0.89221857 |
| RPS29 | protein_coding | -0.3598963 | 8.19512293 |
| RP11-517P14 | lncRNA | -0.3603 | 1.59197439 |
| HPRT1 | protein_coding | -0.3603923 | 7.53880334 |
| RP1-39G22.7 | lncRNA | -0.3605214 | 4.3151144 |
| SCML2 | protein_coding | -0.3607525 | 7.0513005 |
| PLP2 | protein_coding | -0.3607592 | 10.0597436 |
| RANBP3-DT | lncRNA | -0.3608613 | 0.70367723 |
| RP11-715J22 | lncRNA | -0.3608637 | 7.19847657 |
| VPS37B | protein_coding | -0.3609107 | 6.34730475 |
| PRICKLE2 | protein_coding | -0.3613785 | 1.72748935 |
| FANCG | protein_coding | -0.3615597 | 8.12561317 |
| FUZ | protein_coding | -0.3616116 | 5.66417902 |
| SPATA33 | protein_coding | -0.3617253 | 5.23116859 |
| MAGEE1 | protein_coding | -0.3618886 | 0.56338811 |
| ZNF572 | protein_coding | -0.3618942 | 0.85047161 |
| NDUFA2 | protein_coding | -0.3621799 | 7.60761738 |
| COQ10A | protein_coding | -0.3623075 | 5.03047172 |
| TFIP11-DT | lncRNA | -0.3624019 | 2.75680505 |
| RBBP7 | protein_coding | -0.3624316 | 11.3604189 |
| TOMM20 | protein_coding | -0.3625215 | 9.63588483 |
| KLRG2 | protein_coding | -0.3627245 | 1.01923415 |
| TYRO3 | protein_coding | -0.3628948 | 7.34014145 |
| RP11-498P14 | lncRNA | -0.3629158 | 0.77490332 |
| NCAN | protein_coding | -0.3630282 | 3.28548029 |
| PRKACA | protein_coding | -0.3630754 | 9.11961592 |
| TBCE | protein_coding | -0.363371 | 2.58502846 |
| DMAC1 | protein_coding | -0.3634597 | 5.7353448 |
| HMX2 | protein_coding | -0.3637127 | 0.64374905 |
| INF2 | protein_coding | -0.3637142 | 7.37714032 |
| EXOSC5 | protein_coding | -0.3638235 | 7.09765088 |
| RUBCN | protein_coding | -0.3640304 | 9.09774558 |
| PPIA | protein_coding | -0.3640738 | 11.8385722 |
| HNRNPAB | protein_coding | -0.3641955 | 9.36279339 |
| AC083843.1 | lncRNA | -0.3641975 | 1.71156048 |
| HPSE | protein_coding | -0.3642556 | 6.53032741 |
| NEFL | protein_coding | -0.3643475 | 10.6874932 |

| | | | |
|---|---|---|---|
| SCN2A | protein_coding | -0.3644438 | 2.64230483 |
| CLSPN | protein_coding | -0.3645555 | 8.89769591 |
| RP11-299G2( | lncRNA | -0.3646196 | 0.72249417 |
| BACH2 | protein_coding | -0.364856 | 1.49008889 |
| RP11-362K14 | lncRNA | -0.3649891 | 0.59533941 |
| MEX3D | protein_coding | -0.3650028 | 2.15246996 |
| EXOC3-AS1 | lncRNA | -0.365333 | 2.27476833 |
| E2F7 | protein_coding | -0.3654449 | 7.86384632 |
| SLC25A34 | protein_coding | -0.3655258 | 1.09645284 |
| BAIAP2L1 | protein_coding | -0.3655787 | 8.24197755 |
| ROR1 | protein_coding | -0.3655953 | 0.79444365 |
| RP11-18H7.1 | lncRNA | -0.3656539 | 1.15065368 |
| ADD3 | protein_coding | -0.3663093 | 5.55162443 |
| TMEM143 | protein_coding | -0.3663303 | 3.30949483 |
| SPTBN2 | protein_coding | -0.3664918 | 8.88691936 |
| LDHC | protein_coding | -0.3665754 | 2.15323983 |
| ARPIN | protein_coding | -0.3669831 | 0.75080287 |
| GM2A | protein_coding | -0.3669862 | 4.78808587 |
| RP11-455O6. | lncRNA | -0.366995 | 0.55880622 |
| THOP1 | protein_coding | -0.3670294 | 6.89981766 |
| LRRC43 | protein_coding | -0.3676584 | 2.1591642 |
| ADAMTS6 | protein_coding | -0.367694 | 2.01496318 |
| RP11-384P7. | lncRNA | -0.3679464 | 5.76596749 |
| TGFB1 | protein_coding | -0.3680062 | 0.65016418 |
| MYLK2 | protein_coding | -0.3680164 | 0.69691129 |
| AQP3 | protein_coding | -0.3680725 | 5.47602369 |
| ATP1B3 | protein_coding | -0.368191 | 9.34749134 |
| XYLT1 | protein_coding | -0.3682996 | 6.37917122 |
| TRIM52 | protein_coding | -0.3684198 | 4.78233543 |
| RAB2A | protein_coding | -0.3684414 | 7.13610148 |
| CNNM3-DT | lncRNA | -0.36846 | 1.04995843 |
| NIPBL-DT | lncRNA | -0.3686795 | 6.79735299 |
| RP5-968P14. | lncRNA | -0.3686994 | 0.82551242 |
| P2RX3 | protein_coding | -0.3687806 | 0.70318833 |
| RABGEF1 | protein_coding | -0.3688444 | 1.88660468 |
| TTC26 | protein_coding | -0.3691176 | 3.73850869 |
| MEF2C | protein_coding | -0.3691715 | 0.70684491 |
| RAP2B | protein_coding | -0.369607 | 4.86236819 |
| GAREM2 | protein_coding | -0.3697365 | 5.63684042 |
| HIGD1A | protein_coding | -0.369776 | 3.83653913 |
| TRIM41 | protein_coding | -0.3698721 | 7.28969355 |
| DTL | protein_coding | -0.3698779 | 9.91395196 |
| HAGHL | protein_coding | -0.3700347 | 2.01822037 |
| PSPN | protein_coding | -0.3700473 | 2.2562186 |
| BORCS6 | protein_coding | -0.3700559 | 2.30316947 |

| | | | |
|---|---|---|---|
| ARHGAP19 | protein_coding | -0.370234 | 6.40497964 |
| NHP2 | protein_coding | -0.3702869 | 8.46209102 |
| FAM216A | protein_coding | -0.3702907 | 6.95188036 |
| AC079922.3 | lncRNA | -0.3704023 | 2.06275793 |
| TTC33 | protein_coding | -0.3705133 | 5.26646406 |
| ATAD2 | protein_coding | -0.3705765 | 10.6687259 |
| SYNGR1 | protein_coding | -0.3705819 | 8.93077879 |
| TUBB4B | protein_coding | -0.3706142 | 11.7476021 |
| ILF3-DT | lncRNA | -0.3707553 | 6.80757449 |
| ITM2C | protein_coding | -0.371083 | 7.26907106 |
| SPDEF | protein_coding | -0.3711728 | 3.33874533 |
| SAPCD2 | protein_coding | -0.3713939 | 4.916246 |
| HOMER2 | protein_coding | -0.3714336 | 6.14382684 |
| NR2F6 | protein_coding | -0.3716083 | 3.31487757 |
| TMPRSS6 | protein_coding | -0.3716247 | 0.7789462 |
| ACVR2B | protein_coding | -0.3717411 | 5.89237233 |
| HSPB11 | protein_coding | -0.3720923 | 6.70882746 |
| SLC25A4 | protein_coding | -0.3720968 | 7.73853782 |
| ZBTB45 | protein_coding | -0.3723595 | 4.40805316 |
| GS1-238N13 | lncRNA | -0.3725537 | 0.67845868 |
| RP11-422P24 | lncRNA | -0.372715 | 1.28844195 |
| BLM | protein_coding | -0.3728279 | 7.79891401 |
| CCDC30 | protein_coding | -0.3728473 | 1.71836701 |
| RP11-742B18 | lncRNA | -0.3730737 | 0.39612062 |
| IFT22 | protein_coding | -0.3731227 | 3.1499295 |
| TSSK6 | protein_coding | -0.3732489 | 1.71575098 |
| KLRG1 | protein_coding | -0.3732543 | 0.99756802 |
| CHCHD2 | protein_coding | -0.3733056 | 8.778355 |
| BDH1 | protein_coding | -0.3733172 | 6.62399065 |
| SMARCD2 | protein_coding | -0.3733185 | 10.2670939 |
| PRR36 | protein_coding | -0.3735114 | 7.22410967 |
| SLBP | protein_coding | -0.3735972 | 10.4038329 |
| RP11-367N14 | lncRNA | -0.3737661 | 0.47688524 |
| CTD-2292M1 | lncRNA | -0.3738204 | 1.36726583 |
| AAMDC | protein_coding | -0.3738929 | 3.29050567 |
| ATP5MC2 | protein_coding | -0.3741004 | 9.79299125 |
| PTOV1 | protein_coding | -0.3743594 | 8.70832066 |
| MOSPD3 | protein_coding | -0.3744635 | 4.21382246 |
| RELA-DT | lncRNA | -0.3745064 | 2.53902452 |
| RBPJ | protein_coding | -0.3747066 | 8.27577105 |
| PINK1-AS | lncRNA | -0.3747707 | 3.72239828 |
| CDK5R1 | protein_coding | -0.3748447 | 9.1195259 |
| PIP5K1C | protein_coding | -0.3748695 | 5.80945299 |
| CACNA1A | protein_coding | -0.3753083 | 8.81700399 |
| ATP5ME | protein_coding | -0.3753325 | 8.61531546 |

| | | | |
|---|---|---|---|
| ID4 | protein_coding | -0.3753935 | 7.49813752 |
| HAGLROS | lncRNA | -0.3755779 | 2.04701999 |
| HYAL2 | protein_coding | -0.375826 | 4.9114669 |
| BAG5 | protein_coding | -0.3759029 | 10.6903637 |
| ARL3 | protein_coding | -0.376411 | 5.9448735 |
| SLC25A11 | protein_coding | -0.3764484 | 7.86896175 |
| PXN-AS1 | lncRNA | -0.3764991 | 2.35236102 |
| SAMD1 | protein_coding | -0.3765686 | 6.27927505 |
| RSBN1L | protein_coding | -0.3772931 | 7.7294936 |
| SUMF1 | protein_coding | -0.3776902 | 2.75547533 |
| POP7 | protein_coding | -0.3777143 | 6.40870782 |
| UBOX5 | protein_coding | -0.3778312 | 3.72111997 |
| RAB33A | protein_coding | -0.3779234 | 3.83076522 |
| ZNF497 | protein_coding | -0.377957 | 1.87426927 |
| TMEM167B-I | lncRNA | -0.378079 | 0.89157575 |
| LA16c-313D1 | lncRNA | -0.378124 | 3.5999891 |
| RP11-124N2. | lncRNA | -0.3781965 | 2.2512706 |
| CHRNB2 | protein_coding | -0.3784641 | 8.77655183 |
| INTS6-AS1 | lncRNA | -0.378537 | 0.49251508 |
| NRG3 | protein_coding | -0.3785846 | 3.20513889 |
| INSR | protein_coding | -0.3786561 | 7.77533917 |
| XKR8 | protein_coding | -0.3787194 | 3.75971835 |
| RP11-977G1 | lncRNA | -0.3787204 | 1.6393757 |
| DLX5 | protein_coding | -0.378762 | 3.27170609 |
| CDKN3 | protein_coding | -0.3788063 | 7.4730326 |
| SOD2-OT1 | lncRNA | -0.3791176 | 0.48024304 |
| ZNF57 | protein_coding | -0.379168 | 4.75522604 |
| IDH2 | protein_coding | -0.3792081 | 10.0005726 |
| C18orf32 | protein_coding | -0.3793164 | 3.19596395 |
| SCD | protein_coding | -0.3793435 | 11.3277775 |
| B4GALT2 | protein_coding | -0.379354 | 8.77023008 |
| LRRC38 | protein_coding | -0.3795907 | 6.86202185 |
| SREBF1 | protein_coding | -0.3797586 | 8.05020024 |
| FRMD6-AS1 | lncRNA | -0.380138 | 0.56314172 |
| NOX5 | protein_coding | -0.380224 | 0.75122556 |
| RAB11B-AS1 | lncRNA | -0.3804721 | 1.6303136 |
| CPOX | protein_coding | -0.3806355 | 4.44080093 |
| DNM1P35 | lncRNA | -0.3808076 | 1.23435775 |
| GLIDR | lncRNA | -0.380934 | 0.78441227 |
| ALG14 | protein_coding | -0.3809881 | 2.41528307 |
| TERF1 | protein_coding | -0.3812733 | 6.16722047 |
| RP11-9E17.1 | lncRNA | -0.3812854 | 2.65863465 |
| TMED9 | protein_coding | -0.3814308 | 8.98346103 |
| ZNRD2-AS1 | lncRNA | -0.3818043 | 1.22529131 |
| CTD-2298J14 | lncRNA | -0.3818709 | 2.57177016 |

| | | | |
|---|---|---|---|
| PRTFDC1 | protein_coding | -0.3826833 | 5.77029497 |
| GABARAP | protein_coding | -0.3826982 | 2.10550679 |
| TMEM14B | protein_coding | -0.3827122 | 6.12407995 |
| RP11-159D12 | lncRNA | -0.3831133 | 0.89471819 |
| SH3BGRL3 | protein_coding | -0.3831273 | 7.31519008 |
| RP11-432J22 | lncRNA | -0.3838181 | 1.41563138 |
| CDCP1 | protein_coding | -0.3841528 | 5.05941815 |
| AMIGO2 | protein_coding | -0.3842929 | 9.02496696 |
| ACOT4 | protein_coding | -0.3842931 | 3.74556807 |
| FAM72B | protein_coding | -0.3843264 | 4.96162364 |
| GTF2A2 | protein_coding | -0.3845286 | 6.39906062 |
| CAPN9 | protein_coding | -0.3848019 | 1.09839885 |
| GPR19 | protein_coding | -0.3852962 | 3.61984839 |
| SCART1 | protein_coding | -0.3853925 | 0.68818758 |
| TMEM241 | protein_coding | -0.3855927 | 5.70267316 |
| RP13-516M1 | lncRNA | -0.3855974 | 0.8352306 |
| PRDX4 | protein_coding | -0.3856402 | 9.80797831 |
| SYBU | protein_coding | -0.385972 | 7.69862515 |
| MIR9-1HG | lncRNA | -0.3861854 | 0.49876461 |
| RP11-364B14 | protein_coding | -0.3862366 | 2.60262475 |
| SPACA9 | protein_coding | -0.386258 | 1.73557231 |
| RP11-230F18 | lncRNA | -0.3863489 | 0.68159053 |
| FAM183A | protein_coding | -0.3864754 | 0.51199809 |
| RP11-253M7 | lncRNA | -0.3867169 | 0.50500945 |
| AC010894.3 | lncRNA | -0.386853 | 0.44459312 |
| XXYLT1 | protein_coding | -0.3869324 | 7.9398795 |
| FKBP4 | protein_coding | -0.3870956 | 11.0170052 |
| PLEKHA7 | protein_coding | -0.3873575 | 6.30697058 |
| CLDN9 | protein_coding | -0.3873762 | 5.24588793 |
| AP2S1 | protein_coding | -0.3875754 | 8.63308983 |
| APTR | lncRNA | -0.387584 | 5.12147322 |
| TTC3 | protein_coding | -0.3876822 | 11.8769163 |
| UBR7 | protein_coding | -0.3877113 | 11.7916175 |
| FBXL20 | protein_coding | -0.3878152 | 5.98283961 |
| TRAPPC5 | protein_coding | -0.3878475 | 1.52922213 |
| PCMTD2 | protein_coding | -0.3879299 | 9.12359413 |
| RASL11B | protein_coding | -0.3879922 | 1.12616238 |
| DGAT1 | protein_coding | -0.3880601 | 6.03409547 |
| ZNF346-IT1 | lncRNA | -0.3881383 | 0.48323538 |
| KCNMB2 | protein_coding | -0.388355 | 2.51148867 |
| RP11-350E12 | lncRNA | -0.3885079 | 1.55565981 |
| AQP11 | protein_coding | -0.3885425 | 1.62172973 |
| INSM1 | protein_coding | -0.3885697 | 6.88617996 |
| MIEF2 | protein_coding | -0.3887592 | 2.52683391 |
| SRP9 | protein_coding | -0.3892837 | 5.79726905 |

| | | | |
|---|---|---|---|
| NAT14 | protein_coding | -0.3893158 | 5.69712648 |
| DMC1 | protein_coding | -0.3894946 | 5.27653702 |
| MAPK4 | protein_coding | -0.389593 | 2.39768428 |
| SLC25A53 | protein_coding | -0.3897581 | 1.42634771 |
| ANKRD13B | protein_coding | -0.389873 | 5.79339274 |
| HNRNPA1P4: | protein_coding | -0.3901354 | 1.4079725 |
| ATP2A1-AS1 | lncRNA | -0.3901447 | 2.12229037 |
| PTMS | protein_coding | -0.3904015 | 10.2534277 |
| FBXO43 | protein_coding | -0.3904588 | 3.18704887 |
| BEX3 | protein_coding | -0.3905452 | 9.29684059 |
| TSPYL5 | protein_coding | -0.3906445 | 7.33982764 |
| SORBS1 | protein_coding | -0.3908597 | 5.83445614 |
| LHX9 | protein_coding | -0.3908806 | 7.23666843 |
| CCNF | protein_coding | -0.3909675 | 9.90124598 |
| UBASH3B | protein_coding | -0.3910544 | 2.30348763 |
| PDZD7 | protein_coding | -0.391084 | 2.72014663 |
| TRIQK | protein_coding | -0.3914658 | 5.4636849 |
| TMEM11 | protein_coding | -0.3921538 | 4.9876882 |
| SCHIP1 | protein_coding | -0.3923226 | 3.96681143 |
| EEF1B2 | protein_coding | -0.392586 | 11.5440033 |
| CCNY | protein_coding | -0.3929901 | 7.71246409 |
| KIAA0825 | protein_coding | -0.3929919 | 0.71323797 |
| C8orf58 | protein_coding | -0.3931178 | 1.59686561 |
| PPP4R1 | protein_coding | -0.3931809 | 9.39041742 |
| KLHL32 | protein_coding | -0.3934274 | 1.8597355 |
| AKAIN1 | protein_coding | -0.3935005 | 3.60339813 |
| DAD1 | protein_coding | -0.3939904 | 8.76819989 |
| MAPKAP1 | protein_coding | -0.3943182 | 8.62471267 |
| RP11-932O9. | lncRNA | -0.3946305 | 0.68719739 |
| ARHGAP44 | protein_coding | -0.3948765 | 4.98247326 |
| RP11-977B1( | lncRNA | -0.3950396 | 0.78611333 |
| PRR19 | protein_coding | -0.3953035 | 2.37214228 |
| RP11-435O5. | lncRNA | -0.3954953 | 0.75989476 |
| PIGF | protein_coding | -0.3959076 | 5.96954681 |
| ADRA2B | protein_coding | -0.3959779 | 9.58725547 |
| RP11-696N1· | lncRNA | -0.3960712 | 0.62958848 |
| RNF150 | protein_coding | -0.3960716 | 4.2228168 |
| RP11-722E2: | lncRNA | -0.3962377 | 0.48278398 |
| KCNIP2-AS1 | lncRNA | -0.3962661 | 0.45825283 |
| GCSH | protein_coding | -0.3963355 | 0.9632184 |
| MIR4458HG | lncRNA | -0.3964563 | 6.36053331 |
| COMMD10 | protein_coding | -0.3965307 | 5.66320954 |
| PRPS2 | protein_coding | -0.3965825 | 6.61645936 |
| RP11-119F7. | lncRNA | -0.3966411 | 0.93006867 |
| ZWINT | protein_coding | -0.3966594 | 8.94586026 |

| | | | |
|---|---|---|---|
| RP11-158H18 | lncRNA | -0.3966612 | 2.49651454 |
| RP11-435O5. | lncRNA | -0.3967224 | 1.37403055 |
| KDM4B | protein_coding | -0.3968419 | 7.26447585 |
| SLC9A6 | protein_coding | -0.3968757 | 6.57229145 |
| AGPAT5 | protein_coding | -0.3970132 | 8.34184417 |
| PBXIP1 | protein_coding | -0.3971745 | 4.16946577 |
| NBDY | protein_coding | -0.3973322 | 6.25226907 |
| POLD4 | protein_coding | -0.3974472 | 0.76662397 |
| CALM3 | protein_coding | -0.3975282 | 12.3679836 |
| DUSP2 | protein_coding | -0.3976169 | 4.41990356 |
| LDLRAP1 | protein_coding | -0.3978844 | 9.91514671 |
| LINC01963 | lncRNA | -0.3979595 | 4.65761179 |
| LHX1-DT | lncRNA | -0.3981555 | 1.94902938 |
| RYK | protein_coding | -0.3982428 | 7.83419042 |
| HS3ST5 | protein_coding | -0.3983304 | 4.32921613 |
| LRRC45 | protein_coding | -0.3983911 | 5.6580969 |
| FN3KRP | protein_coding | -0.3984564 | 7.56394926 |
| PAFAH1B3 | protein_coding | -0.3990192 | 8.23262006 |
| MBOAT1 | protein_coding | -0.3990415 | 3.93255352 |
| KLHDC3 | protein_coding | -0.3997219 | 10.6021543 |
| SMIM1 | protein_coding | -0.4001519 | 1.11560119 |
| LINC02878 | lncRNA | -0.4002706 | 0.61863042 |
| RP11-363E7. | lncRNA | -0.4003333 | 2.01572069 |
| LPIN3 | protein_coding | -0.4003702 | 3.97886458 |
| ZNF804A | protein_coding | -0.4005298 | 4.18187079 |
| C1orf21 | protein_coding | -0.4006524 | 8.34303625 |
| FDXACB1 | protein_coding | -0.4008629 | 1.13346601 |
| H3-3A | protein_coding | -0.4012165 | 7.20995968 |
| CACNA1G | protein_coding | -0.4015867 | 2.85120915 |
| EFCAB11 | protein_coding | -0.4026721 | 2.58380557 |
| GCNT1 | protein_coding | -0.4026828 | 4.50911085 |
| NRXN2 | protein_coding | -0.4028109 | 4.99239027 |
| HMG20B | protein_coding | -0.4028347 | 5.32553241 |
| LSM4 | protein_coding | -0.4028791 | 9.02263947 |
| GLT1D1 | protein_coding | -0.4029818 | 8.11870873 |
| RP11-305K5. | lncRNA | -0.4030071 | 0.82524125 |
| COMMD9 | protein_coding | -0.4030569 | 6.97796921 |
| FEN1 | protein_coding | -0.4030846 | 11.3991178 |
| DCTN6-DT | lncRNA | -0.403155 | 0.65997797 |
| PPP1R3G | protein_coding | -0.4033008 | 1.15249846 |
| FLJ46284 | lncRNA | -0.4033201 | 0.69639811 |
| AVPI1 | protein_coding | -0.4039588 | 1.38728109 |
| RP11-254F7. | lncRNA | -0.4040078 | 0.82092802 |
| DEGS1 | protein_coding | -0.4041289 | 9.24830928 |
| YBX1 | protein_coding | -0.4042756 | 11.2834264 |

| | | | |
|---|---|---|---|
| SPNS2 | protein_coding | -0.4045603 | 1.45427039 |
| CTDP1 | protein_coding | -0.4047877 | 4.21871375 |
| OXR1 | protein_coding | -0.4048439 | 1.46354333 |
| RP11-96H19. | lncRNA | -0.4049354 | 1.38734861 |
| ALX3 | protein_coding | -0.4049642 | 5.53663151 |
| FBXO25 | protein_coding | -0.4050843 | 6.07530401 |
| KCTD12 | protein_coding | -0.4050863 | 5.88693015 |
| GET3 | protein_coding | -0.4051451 | 7.61979663 |
| ACTL6B | protein_coding | -0.4055716 | 8.95406764 |
| PHLDA3 | protein_coding | -0.4056144 | 4.02059669 |
| GPANK1 | protein_coding | -0.4057804 | 8.42569707 |
| HSD11B2 | protein_coding | -0.4059198 | 3.62162616 |
| CTDSPL | protein_coding | -0.4060541 | 7.89917399 |
| BOC | protein_coding | -0.4061275 | 3.03071514 |
| CTB-104F4.3 | lncRNA | -0.4062258 | 0.45555722 |
| KLHL26 | protein_coding | -0.4062804 | 3.65898206 |
| RB1 | protein_coding | -0.4063147 | 5.74729176 |
| C11orf95 | protein_coding | -0.4063532 | 6.1581029 |
| WDR1 | protein_coding | -0.4064198 | 9.79971832 |
| SPC25 | protein_coding | -0.4067499 | 6.24180019 |
| DGKA | protein_coding | -0.4067659 | 1.56348013 |
| TAFA1 | protein_coding | -0.4067772 | 4.57972054 |
| HOOK1 | protein_coding | -0.4070703 | 7.03760629 |
| P2RY1 | protein_coding | -0.4071853 | 7.66581266 |
| GAPDH | protein_coding | -0.4073037 | 14.1470235 |
| GNB1L | protein_coding | -0.407616 | 4.26050363 |
| POLR3K | protein_coding | -0.4077177 | 7.67553957 |
| HADH | protein_coding | -0.4077723 | 10.8912958 |
| TMEM106C | protein_coding | -0.4078405 | 10.441638 |
| PIP4P2 | protein_coding | -0.4082792 | 3.02539876 |
| RP11-382A2( | lncRNA | -0.4083549 | 0.57172325 |
| LASP1 | protein_coding | -0.4084817 | 6.97939403 |
| NR4A3 | protein_coding | -0.4086468 | 0.61427149 |
| OIP5 | protein_coding | -0.4092453 | 6.473157 |
| HOXA6 | protein_coding | -0.4093711 | 3.89599994 |
| FAM241B | protein_coding | -0.4094214 | 4.54685833 |
| HOXB3 | protein_coding | -0.4096826 | 0.67026836 |
| FGFR4 | protein_coding | -0.4098404 | 2.19777469 |
| DKK3 | protein_coding | -0.4100775 | 0.77297442 |
| RP11-152N1: | lncRNA | -0.4100968 | 0.74395832 |
| KB-1440D3.1 | lncRNA | -0.410127 | 0.78772005 |
| DLK2 | protein_coding | -0.4104105 | 3.29751387 |
| NHS | protein_coding | -0.4107875 | 6.17476344 |
| SLF1 | protein_coding | -0.4109249 | 6.58397361 |
| AIP | protein_coding | -0.4110464 | 7.44577043 |

| | | | |
|---|---|---|---|
| RP11-421K1. | lncRNA | -0.411054 | 0.78661012 |
| DNM3 | protein_coding | -0.4112174 | 4.1989612 |
| IQCE | protein_coding | -0.411633 | 6.90248974 |
| HMMR | protein_coding | -0.4122943 | 7.09978471 |
| ZNF138 | protein_coding | -0.4123441 | 6.39180831 |
| RGL2 | protein_coding | -0.4124157 | 5.82654254 |
| SERINC5 | protein_coding | -0.4127633 | 5.82684269 |
| CFAP43 | protein_coding | -0.4133217 | 0.5312691 |
| UNG | protein_coding | -0.4134315 | 11.253297 |
| ARID3A | protein_coding | -0.4134332 | 6.08701214 |
| GABRE | protein_coding | -0.4137668 | 2.29035442 |
| FAM89B | protein_coding | -0.4137693 | 3.58982438 |
| RP11-253E3. | lncRNA | -0.4139611 | 3.3083484 |
| CRIPT | protein_coding | -0.4143382 | 6.65198828 |
| LINC01356 | lncRNA | -0.4146275 | 1.27424956 |
| SLC27A5 | protein_coding | -0.4146554 | 2.56058938 |
| IQSEC3 | protein_coding | -0.414836 | 4.12528936 |
| ERMAP | protein_coding | -0.4152745 | 6.06956152 |
| LINC01534 | lncRNA | -0.4155899 | 2.89531144 |
| ARHGEF1 | protein_coding | -0.4157606 | 7.10302614 |
| TMEM60 | protein_coding | -0.4157833 | 4.5898241 |
| PWP2 | protein_coding | -0.4158526 | 4.30044899 |
| SEMA4F | protein_coding | -0.4159123 | 5.20122358 |
| CAMK1 | protein_coding | -0.4159379 | 0.82633778 |
| RAPSN | protein_coding | -0.4163944 | 6.50242972 |
| FBXL8 | protein_coding | -0.416542 | 3.77338203 |
| RPS6KA5 | protein_coding | -0.4169536 | 8.8045781 |
| ZFYVE28 | protein_coding | -0.4170087 | 4.68608552 |
| RP11-574F11 | lncRNA | -0.4170217 | 0.91204198 |
| CCNB3 | protein_coding | -0.417027 | 7.48210688 |
| TMEM44 | protein_coding | -0.4170709 | 3.98280491 |
| C7orf50 | protein_coding | -0.417201 | 6.35293508 |
| MT-ATP6 | protein_coding | -0.417247 | 8.92551494 |
| MIF4GD | protein_coding | -0.4174362 | 5.62512442 |
| RUSF1 | protein_coding | -0.4176992 | 7.45893567 |
| CTD-2342J14 | lncRNA | -0.4178265 | 1.07856243 |
| FAM72A | protein_coding | -0.4179914 | 4.1720354 |
| TMEM184A | protein_coding | -0.4181327 | 4.82257224 |
| LINC01574 | lncRNA | -0.4184758 | 0.50966358 |
| RP4-583P15. | lncRNA | -0.4189554 | 0.86136273 |
| HDAC11 | protein_coding | -0.4190076 | 2.847736 |
| PXMP2 | protein_coding | -0.4190243 | 2.72922559 |
| RP11-390F4. | lncRNA | -0.4190346 | 1.09577763 |
| PROCA1 | protein_coding | -0.4192893 | 2.21963327 |
| COBL | protein_coding | -0.4195719 | 6.10440137 |

| | | | |
|---|---|---|---|
| PPFIA2 | protein_coding | -0.4196302 | 3.64155653 |
| CKLF | protein_coding | -0.4197837 | 1.00639844 |
| UTP3 | protein_coding | -0.4200039 | 6.74069576 |
| KLHDC8A | protein_coding | -0.4201242 | 2.78046024 |
| WDTC1 | protein_coding | -0.4201433 | 10.7708082 |
| TMEM132D | protein_coding | -0.4206439 | 3.73327313 |
| TIGD3 | protein_coding | -0.4207228 | 3.19740645 |
| DLK1 | protein_coding | -0.4208647 | 2.12892455 |
| PRR7 | protein_coding | -0.4208843 | 1.16251736 |
| FGF9 | protein_coding | -0.4210668 | 3.4808988 |
| ING4 | protein_coding | -0.4210723 | 6.4985446 |
| PIEZO1 | protein_coding | -0.4215207 | 2.53025729 |
| RBFOX1 | protein_coding | -0.4215623 | 10.2820945 |
| NRSN1 | protein_coding | -0.4215781 | 10.411268 |
| RP11-575F12 | lncRNA | -0.4217591 | 1.49565861 |
| RCOR2 | protein_coding | -0.4218192 | 10.5460388 |
| TAFA2 | protein_coding | -0.4218753 | 0.66481125 |
| UBALD1 | protein_coding | -0.4228898 | 4.86549312 |
| ST6GALNAC6 | protein_coding | -0.4229226 | 1.65405087 |
| SLC25A5-AS1 | lncRNA | -0.4232548 | 2.19492515 |
| TFR2 | protein_coding | -0.4233029 | 5.36076096 |
| COX16 | protein_coding | -0.4234004 | 2.06865882 |
| RP11-677M1 | lncRNA | -0.4237734 | 0.49688116 |
| CNIH2 | protein_coding | -0.4242266 | 9.10194485 |
| TM4SF19-AS | lncRNA | -0.424299 | 0.57058888 |
| HLTF | protein_coding | -0.4243067 | 7.97459293 |
| LINC02894 | lncRNA | -0.4244118 | 5.05223873 |
| MZT2B | protein_coding | -0.424717 | 3.1707495 |
| TMC7 | protein_coding | -0.4248734 | 2.80964842 |
| CCDC62 | protein_coding | -0.4248994 | 1.3667491 |
| CDK2AP1 | protein_coding | -0.4249314 | 10.301678 |
| AP001412.1 | lncRNA | -0.424998 | 0.67004983 |
| SERGEF | protein_coding | -0.4254537 | 4.87518975 |
| GRAMD4 | protein_coding | -0.4259758 | 9.06406234 |
| ACVR2B-AS1 | lncRNA | -0.4259885 | 2.04477777 |
| RP11-767N6. | lncRNA | -0.426136 | 1.46817078 |
| EGLN1 | protein_coding | -0.4264032 | 1.59132954 |
| GRK6 | protein_coding | -0.4264259 | 8.07871339 |
| ZNF485 | protein_coding | -0.4264376 | 1.29781574 |
| SLC2A8 | protein_coding | -0.4264461 | 6.12989983 |
| KCTD7 | protein_coding | -0.4265503 | 8.28872678 |
| LZTS2 | protein_coding | -0.4266541 | 5.49893829 |
| LAMB2 | protein_coding | -0.4267219 | 2.01895667 |
| DHX40 | protein_coding | -0.4268775 | 9.54998432 |
| ALDOA | protein_coding | -0.4272336 | 3.3743677 |

| | | | |
|---|---|---|---|
| RP11-190A12 | lncRNA | -0.427409 | 2.42966091 |
| SEC61G | protein_coding | -0.4277921 | 7.09273915 |
| HID1 | protein_coding | -0.4280676 | 7.86100263 |
| RP11-15A1.8 | lncRNA | -0.4283504 | 0.65940301 |
| TMEM97 | protein_coding | -0.4286928 | 11.378105 |
| BBS12 | protein_coding | -0.4289786 | 0.62345431 |
| SGTA | protein_coding | -0.4290104 | 11.0365293 |
| OBI1 | protein_coding | -0.4292255 | 9.65539753 |
| RAI1 | protein_coding | -0.4292595 | 8.37421833 |
| AKR7A2 | protein_coding | -0.429323 | 7.85733771 |
| PCLO | protein_coding | -0.4295938 | 11.528541 |
| FOXO4 | protein_coding | -0.4298052 | 3.69417336 |
| RP11-393I2.4 | lncRNA | -0.4298879 | 1.05339159 |
| RP11-468E2. | lncRNA | -0.4299991 | 0.81292808 |
| RP11-395A13 | lncRNA | -0.4301294 | 0.98444137 |
| TMEM201 | protein_coding | -0.4301891 | 6.03369 |
| GLRX5 | protein_coding | -0.4302128 | 11.1470725 |
| PADI2 | protein_coding | -0.4303416 | 11.7747839 |
| RP11-326C3. | lncRNA | -0.4303664 | 1.99598469 |
| RRBP1 | protein_coding | -0.4305566 | 6.69064866 |
| POR | protein_coding | -0.4305743 | 7.90368326 |
| MYO1H | protein_coding | -0.4305849 | 1.18107628 |
| PMEL | protein_coding | -0.4307149 | 0.95178408 |
| AC000068.9 | lncRNA | -0.4308087 | 0.75465377 |
| NFATC4 | protein_coding | -0.4308647 | 3.37906908 |
| RP11-783K16 | lncRNA | -0.4309062 | 1.10709576 |
| CRACR2B | protein_coding | -0.4309158 | 4.27049698 |
| MLYCD | protein_coding | -0.4310096 | 4.59703276 |
| HEG1 | protein_coding | -0.4311804 | 1.3778066 |
| P4HA3 | protein_coding | -0.4312359 | 0.61093452 |
| NACAD | protein_coding | -0.4315851 | 0.91211981 |
| GIT1 | protein_coding | -0.4317624 | 10.3951561 |
| HDAC2-AS2 | lncRNA | -0.4318015 | 1.49563321 |
| RP11-115C21 | lncRNA | -0.4318717 | 2.68830638 |
| TESK2 | protein_coding | -0.4319567 | 4.42023904 |
| LINGO3 | protein_coding | -0.4321319 | 1.31738794 |
| FAM210B | protein_coding | -0.4322661 | 12.0149338 |
| TNFRSF1A | protein_coding | -0.4323115 | 4.12003835 |
| FAM126A | protein_coding | -0.4329579 | 10.3368684 |
| SPRING1 | protein_coding | -0.4329605 | 9.73876488 |
| DICER1-AS1 | lncRNA | -0.4330963 | 2.9534199 |
| CTD-2168K21 | lncRNA | -0.4331694 | 1.27353005 |
| CDC20 | protein_coding | -0.4331992 | 9.90753587 |
| TYMS | protein_coding | -0.4332233 | 10.7293217 |
| ARHGAP39 | protein_coding | -0.4332423 | 4.50703363 |

| | | | |
|---|---|---|---|
| PRKAR1B | protein_coding | -0.4335298 | 8.8428331 |
| TMEM170B | protein_coding | -0.4335611 | 6.11802139 |
| COMMD4 | protein_coding | -0.4338212 | 9.15457508 |
| LSM11 | protein_coding | -0.4338689 | 7.22988884 |
| ATP5MG | protein_coding | -0.4341798 | 7.1642465 |
| E2F8 | protein_coding | -0.4342274 | 4.92576186 |
| SLC30A6-DT | lncRNA | -0.4343 | 0.68410268 |
| FAM241A | protein_coding | -0.4345228 | 3.00682293 |
| MXD3 | protein_coding | -0.4346009 | 5.58814541 |
| SNHG30 | lncRNA | -0.4348241 | 4.98832329 |
| EID1 | protein_coding | -0.4350885 | 7.51803298 |
| RP11-644F5. | protein_coding | -0.4354043 | 1.04420875 |
| SPON2 | protein_coding | -0.4356535 | 0.85567525 |
| HDDC3 | protein_coding | -0.4357341 | 6.62840302 |
| BATF3 | protein_coding | -0.435742 | 2.03720658 |
| KB-318B8.7 | lncRNA | -0.4357805 | 0.51318586 |
| TP73 | protein_coding | -0.4360307 | 7.87703226 |
| NUDT14 | protein_coding | -0.4360708 | 8.09166395 |
| LINC01816 | lncRNA | -0.4361499 | 1.81132828 |
| ZNF615 | protein_coding | -0.4361674 | 2.93486455 |
| DPY19L3-DT | lncRNA | -0.4364636 | 0.79902675 |
| APBB1 | protein_coding | -0.4364659 | 7.68603129 |
| ARL6 | protein_coding | -0.4369022 | 4.763859 |
| CFAP298 | protein_coding | -0.4369435 | 4.21948319 |
| ELAVL3 | protein_coding | -0.4370868 | 10.9567473 |
| C12orf57 | protein_coding | -0.437165 | 10.8860001 |
| MARCHF3 | protein_coding | -0.4372751 | 7.40054644 |
| PGBD5 | protein_coding | -0.4373698 | 7.29126533 |
| OTUD1 | protein_coding | -0.4383002 | 1.20281542 |
| TECR | protein_coding | -0.4390738 | 10.6794193 |
| TOB1-AS1 | lncRNA | -0.4391547 | 2.92993822 |
| TNS4 | protein_coding | -0.4392518 | 1.22349705 |
| MIS18A | protein_coding | -0.4395537 | 8.24430884 |
| RP11-677M1 | lncRNA | -0.4400788 | 0.95323468 |
| CCER2 | protein_coding | -0.4401385 | 11.7243525 |
| ZNF45-AS1 | lncRNA | -0.4402056 | 0.8079283 |
| PDE4C | protein_coding | -0.4404798 | 0.65601495 |
| VWA5B2 | protein_coding | -0.440537 | 8.61554754 |
| SEC24B-AS1 | lncRNA | -0.4407717 | 0.93611079 |
| TTC23L-AS1 | lncRNA | -0.4407755 | 0.49412429 |
| BFSP1 | protein_coding | -0.4408134 | 1.10059778 |
| SLC39A4 | protein_coding | -0.4411325 | 4.5887886 |
| RP11-295H2 | lncRNA | -0.4411998 | 0.90681618 |
| TRAK1 | protein_coding | -0.4412356 | 8.9114113 |
| RIN1 | protein_coding | -0.4415088 | 1.14604058 |

| | | | |
|---|---|---|---|
| FAM172A | protein_coding | -0.4415351 | 6.37269196 |
| ZNF160 | protein_coding | -0.4419537 | 8.62917394 |
| FUCA1 | protein_coding | -0.4420435 | 3.80354285 |
| BRSK2 | protein_coding | -0.4421657 | 6.98998644 |
| GABPB2 | protein_coding | -0.4423588 | 6.21487697 |
| CACNA2D2 | protein_coding | -0.442752 | 8.64354684 |
| SPA17 | protein_coding | -0.4430173 | 2.59330168 |
| PKN3 | protein_coding | -0.4431275 | 5.47160779 |
| RRM2 | protein_coding | -0.4431657 | 12.5108569 |
| KCTD21-AS1 | lncRNA | -0.4431904 | 2.20530392 |
| OAZ2 | protein_coding | -0.4432338 | 7.2754127 |
| CHRFAM7A | protein_coding | -0.4438036 | 0.68943573 |
| TMEM54 | protein_coding | -0.4438915 | 2.88112403 |
| FAM98C | protein_coding | -0.4440333 | 5.06654231 |
| TNIK | protein_coding | -0.4440873 | 11.7553903 |
| DLEU1 | lncRNA | -0.4441471 | 3.36879707 |
| TMEM53 | protein_coding | -0.4442189 | 4.39276189 |
| FGFRL1 | protein_coding | -0.4444581 | 4.14863235 |
| PIN1 | protein_coding | -0.4444839 | 8.74209138 |
| CPTP | protein_coding | -0.4449656 | 5.14099667 |
| SUN2 | protein_coding | -0.4453184 | 10.9506759 |
| CAMK2N2 | protein_coding | -0.4456214 | 6.28460935 |
| PACSIN2 | protein_coding | -0.4458068 | 10.2272135 |
| NR2F2 | protein_coding | -0.4460139 | 5.14457797 |
| DENND5B | protein_coding | -0.4462199 | 5.10692505 |
| TDRD5 | protein_coding | -0.4468675 | 3.89733804 |
| RAPGEF5 | protein_coding | -0.4471372 | 5.51969154 |
| IRS1 | protein_coding | -0.4472006 | 10.8978911 |
| SF3B5 | protein_coding | -0.4473054 | 8.10175641 |
| FADS1 | protein_coding | -0.4475277 | 12.0693555 |
| GRB14 | protein_coding | -0.4478552 | 1.63376969 |
| GON7 | protein_coding | -0.4483976 | 7.79763725 |
| GALM | protein_coding | -0.4485 | 0.62420195 |
| LINC00471 | lncRNA | -0.4485258 | 1.1965563 |
| TTC5 | protein_coding | -0.4486433 | 7.53589638 |
| SGSM1 | protein_coding | -0.4487091 | 3.84758818 |
| PYCR3 | protein_coding | -0.4487137 | 5.50331849 |
| PALD1 | protein_coding | -0.4492121 | 2.30687869 |
| ARRDC4 | protein_coding | -0.4493932 | 6.32408428 |
| PRRT1 | protein_coding | -0.4496462 | 0.99417986 |
| DACT3 | protein_coding | -0.4496527 | 5.19969999 |
| ZP1 | protein_coding | -0.4498595 | 0.68120942 |
| CCDC88C | protein_coding | -0.4498808 | 10.1076947 |
| CENPX | protein_coding | -0.4499363 | 10.0876812 |
| PPT2 | protein_coding | -0.4501708 | 5.38983811 |

| | | | |
|---|---|---|---|
| LEMD3 | protein_coding | -0.4502834 | 9.94851538 |
| RNF217 | protein_coding | -0.4505406 | 2.19253837 |
| RP11-403P17 | lncRNA | -0.4508066 | 0.81636724 |
| C1QL4 | protein_coding | -0.4508876 | 0.87705717 |
| GUK1 | protein_coding | -0.4509059 | 9.14955214 |
| ZSWIM6 | protein_coding | -0.451733 | 4.91545903 |
| LLGL1 | protein_coding | -0.4519856 | 6.64194515 |
| ACSL6 | protein_coding | -0.4523618 | 6.19680863 |
| CFL1 | protein_coding | -0.4535231 | 13.6236445 |
| PACSIN1 | protein_coding | -0.4536159 | 5.67817692 |
| DNAAF3 | protein_coding | -0.4536566 | 5.20811658 |
| SIRT5 | protein_coding | -0.4540714 | 7.22607775 |
| RPSA | protein_coding | -0.4541357 | 11.6040381 |
| HCFC1R1 | protein_coding | -0.4542722 | 8.84544295 |
| RP11-73M18 | lncRNA | -0.4545998 | 1.04643863 |
| NTN3 | protein_coding | -0.4556006 | 2.44344842 |
| CNPY3 | protein_coding | -0.4556689 | 8.9100118 |
| AC129492.6 | lncRNA | -0.4558616 | 2.01816249 |
| ACVR2A | protein_coding | -0.456082 | 7.57905506 |
| SLC6A8 | protein_coding | -0.4563674 | 4.63930686 |
| TMEM249 | protein_coding | -0.4564934 | 0.52510485 |
| SNX25 | protein_coding | -0.4566047 | 5.72523878 |
| SPSB4 | protein_coding | -0.4570707 | 8.21339177 |
| RP11-10K17. | lncRNA | -0.4571177 | 0.51646623 |
| ST8SIA2 | protein_coding | -0.4571578 | 0.50899428 |
| RECQL | protein_coding | -0.4572477 | 8.96061882 |
| DHFR | protein_coding | -0.4573654 | 9.55936566 |
| ISL1 | protein_coding | -0.4577691 | 12.2778346 |
| ARPC1B | protein_coding | -0.4579059 | 1.36213554 |
| DPM3 | protein_coding | -0.4579413 | 3.79881075 |
| ELAVL4 | protein_coding | -0.457952 | 5.9243928 |
| MRPS24 | protein_coding | -0.4579521 | 0.41471976 |
| CIDEB | protein_coding | -0.4579856 | 3.25683137 |
| RALYL | protein_coding | -0.4591694 | 0.60935496 |
| CLCN2 | protein_coding | -0.4591821 | 4.64244347 |
| RP11-6O2.4 | lncRNA | -0.4594489 | 0.57781594 |
| RP4-753M9.1 | lncRNA | -0.4594571 | 1.08931136 |
| RAPH1 | protein_coding | -0.4596504 | 4.26934178 |
| FADS2 | protein_coding | -0.4597489 | 13.1472274 |
| INKA1 | protein_coding | -0.4597935 | 1.0240609 |
| CD8B | protein_coding | -0.4598583 | 1.76486076 |
| GPATCH11 | protein_coding | -0.4598994 | 9.19261102 |
| GJC2 | protein_coding | -0.4599329 | 1.87612074 |
| NUP93-DT | lncRNA | -0.4600676 | 0.54064819 |
| NRG2 | protein_coding | -0.4601768 | 0.63790734 |

| | | | |
|---|---|---|---|
| DOK7 | protein_coding | -0.4603494 | 0.88202822 |
| TUB | protein_coding | -0.4604619 | 9.25764364 |
| JAM3 | protein_coding | -0.4604944 | 7.36993579 |
| SLC20A2 | protein_coding | -0.4606635 | 5.2970528 |
| ADAMTS2 | protein_coding | -0.4609247 | 1.0042364 |
| RP11-637A1. | lncRNA | -0.4609292 | 0.96778527 |
| SEPSECS-AS1 | lncRNA | -0.4609975 | 1.83765493 |
| DPYSL3 | protein_coding | -0.4610641 | 13.5069318 |
| SDHAF4 | protein_coding | -0.4614783 | 5.19805376 |
| AC079145.4 | lncRNA | -0.4615266 | 0.62413233 |
| MIF | protein_coding | -0.4616204 | 7.63569453 |
| CCDC187 | protein_coding | -0.4624875 | 7.04101214 |
| FOXE1 | protein_coding | -0.4626259 | 5.21323745 |
| UCP2 | protein_coding | -0.4626744 | 10.5545058 |
| CTD-2574D2. | lncRNA | -0.4628594 | 1.39826731 |
| RPL34-DT | lncRNA | -0.46306 | 0.58926787 |
| RPS5 | protein_coding | -0.4633009 | 13.3907081 |
| FRMD3 | protein_coding | -0.4637431 | 8.80439363 |
| SFXN3 | protein_coding | -0.4637636 | 5.78292281 |
| SLC17A9 | protein_coding | -0.4639521 | 0.73764612 |
| ENTPD3-AS1 | lncRNA | -0.4641796 | 0.67101283 |
| MPZL3 | protein_coding | -0.4644266 | 3.11692316 |
| MYO1B | protein_coding | -0.464968 | 11.005323 |
| MCRIP2 | protein_coding | -0.4650793 | 5.52694529 |
| RND2 | protein_coding | -0.4651425 | 6.79966776 |
| RP11-154J22 | lncRNA | -0.4653156 | 2.0423032 |
| RP11-640I2.2 | lncRNA | -0.4655157 | 3.62029839 |
| HOXA-AS3 | lncRNA | -0.4656569 | 1.51630806 |
| PGAM1 | protein_coding | -0.4661775 | 9.35781336 |
| RP11-80A15. | lncRNA | -0.4665383 | 3.2949441 |
| APOM | protein_coding | -0.4666178 | 1.81054288 |
| SSBP2 | protein_coding | -0.4666866 | 8.49504684 |
| DUSP26 | protein_coding | -0.4668368 | 9.14887855 |
| C6orf99 | lncRNA | -0.467061 | 1.34108436 |
| KSR2 | protein_coding | -0.4673005 | 9.12327258 |
| ZNF362 | protein_coding | -0.4674246 | 9.53948293 |
| SLC29A4 | protein_coding | -0.4676078 | 3.64414259 |
| ELOVL2 | protein_coding | -0.4676511 | 9.3719333 |
| LINC00867 | lncRNA | -0.4678635 | 4.8484601 |
| H2AZ1-DT | lncRNA | -0.4683867 | 1.02545133 |
| RP1-170O19. | lncRNA | -0.4687086 | 0.81687019 |
| LINC00652 | lncRNA | -0.4690323 | 0.64797244 |
| CDKL3 | protein_coding | -0.4690586 | 1.09234938 |
| SLC25A10 | protein_coding | -0.4691032 | 3.33801151 |
| MRPS30-DT | lncRNA | -0.4692852 | 0.74173827 |

| | | | |
|---|---|---|---|
| GREM1 | protein_coding | -0.4693268 | 2.74516458 |
| BOLA1 | protein_coding | -0.4694787 | 6.23539761 |
| COX17 | protein_coding | -0.4695436 | 5.76659509 |
| KIAA0895 | protein_coding | -0.4696487 | 6.67079505 |
| CBFB | protein_coding | -0.4696977 | 9.21421678 |
| LINC00921 | lncRNA | -0.470084 | 1.38393676 |
| PRR11 | protein_coding | -0.4700987 | 11.057139 |
| RORC | protein_coding | -0.4701412 | 1.21057427 |
| LRRC20 | protein_coding | -0.4703656 | 6.84860589 |
| MACROD1 | protein_coding | -0.4706077 | 4.69196918 |
| LPL | protein_coding | -0.4709017 | 9.15907625 |
| ADAM11 | protein_coding | -0.4713916 | 10.5017834 |
| TWSG1 | protein_coding | -0.4716243 | 5.62179728 |
| TPI1 | protein_coding | -0.4716686 | 13.9753655 |
| RACGAP1 | protein_coding | -0.4717606 | 11.2939645 |
| STPG3-AS1 | lncRNA | -0.4719363 | 2.85835528 |
| PLPPR2 | protein_coding | -0.4726405 | 7.76801193 |
| ATP1A3 | protein_coding | -0.4727266 | 2.50291625 |
| ACADS | protein_coding | -0.472746 | 3.99712232 |
| SLAIN1 | protein_coding | -0.4734128 | 9.12768043 |
| RTBDN | protein_coding | -0.4736384 | 7.40852136 |
| RP11-632L2.: | lncRNA | -0.4740722 | 1.36618094 |
| SNHG21 | lncRNA | -0.4741293 | 1.20774487 |
| RP11-172F4. | lncRNA | -0.4744532 | 1.36854781 |
| MT-ATP8 | protein_coding | -0.4748626 | 11.1377649 |
| PRAG1 | protein_coding | -0.4748737 | 6.01328048 |
| IMPDH1 | protein_coding | -0.4748825 | 8.78789405 |
| MAGI1 | protein_coding | -0.4755368 | 7.35021299 |
| MISP3 | protein_coding | -0.4768141 | 4.52651374 |
| SMAP1 | protein_coding | -0.4768582 | 8.5531644 |
| RP11-1191J2 | lncRNA | -0.4769452 | 0.66142751 |
| RP11-214N9. | lncRNA | -0.4770448 | 1.62501325 |
| DACH1 | protein_coding | -0.4770963 | 10.1566001 |
| RP4-669P10. | lncRNA | -0.4771276 | 1.66115185 |
| TAS1R3 | protein_coding | -0.477262 | 1.76586468 |
| RORA | protein_coding | -0.4776793 | 4.93617185 |
| BMP1 | protein_coding | -0.4777728 | 2.68854316 |
| RP11-65L3.2 | lncRNA | -0.4778748 | 0.74086844 |
| CCDC163 | protein_coding | -0.4781757 | 2.63746206 |
| RNASEH2C | protein_coding | -0.4782907 | 8.56895205 |
| APBA1 | protein_coding | -0.478777 | 2.74127593 |
| LINC01063 | lncRNA | -0.4789299 | 0.83068164 |
| TLCD3B | protein_coding | -0.4793156 | 7.6115935 |
| CTB-50L17.7 | lncRNA | -0.4793807 | 0.64870826 |
| MAP2 | protein_coding | -0.4796507 | 10.2003772 |

| | | | |
|---|---|---|---|
| RP11-77K12. | lncRNA | -0.479684 | 1.99266738 |
| RGS14 | protein_coding | -0.4797675 | 8.61509423 |
| ESR2 | protein_coding | -0.4798023 | 1.01970321 |
| RP1-206D15. | lncRNA | -0.4798624 | 0.82946947 |
| CSKMT | protein_coding | -0.4798957 | 2.88770196 |
| ZNF737 | protein_coding | -0.4799096 | 7.7082377 |
| CCNE1 | protein_coding | -0.4802186 | 9.42641632 |
| TK1 | protein_coding | -0.48036 | 8.16988965 |
| COA6-AS1 | lncRNA | -0.4807073 | 0.82253675 |
| RP11-70P17. | lncRNA | -0.4808798 | 0.70269368 |
| MTCP1 | protein_coding | -0.4808829 | 0.94238087 |
| FUT8 | protein_coding | -0.4809393 | 9.13303826 |
| COQ8A | protein_coding | -0.4809923 | 1.24542883 |
| OTX1 | protein_coding | -0.4813629 | 3.2251167 |
| DNAJC1 | protein_coding | -0.4814037 | 6.62065034 |
| RP11-318A1! | lncRNA | -0.4814493 | 1.45173441 |
| LTBP3 | protein_coding | -0.4815499 | 5.61628616 |
| RBM15-AS1 | lncRNA | -0.4818113 | 1.34497848 |
| PSTPIP2 | protein_coding | -0.4819501 | 0.884393 |
| LINC01623 | lncRNA | -0.4820161 | 1.12051173 |
| CXADR | protein_coding | -0.4821938 | 7.10443368 |
| ECHS1 | protein_coding | -0.4822924 | 11.6979767 |
| LINC02500 | lncRNA | -0.4823255 | 2.75767841 |
| RP11-61L14.' | lncRNA | -0.4823355 | 1.8520452 |
| PAQR5 | protein_coding | -0.4824584 | 4.33649837 |
| RP11-304L1! | lncRNA | -0.4833 | 0.88550751 |
| MORN2 | protein_coding | -0.483819 | 3.26058701 |
| TCEAL8 | protein_coding | -0.4839777 | 6.84507828 |
| DLX6 | protein_coding | -0.4841108 | 5.04696918 |
| EPPK1 | protein_coding | -0.4844559 | 1.33393268 |
| TMEM74 | protein_coding | -0.4845133 | 11.6468057 |
| PEDS1 | protein_coding | -0.4846651 | 8.99989208 |
| TSPAN6 | protein_coding | -0.4851063 | 5.57995052 |
| KMT5C | protein_coding | -0.485259 | 7.70433267 |
| NELL2 | protein_coding | -0.4852786 | 11.7582599 |
| FAT1 | protein_coding | -0.4852951 | 9.14878415 |
| RPS3A | protein_coding | -0.4854899 | 13.3621301 |
| CDCA4 | protein_coding | -0.4856025 | 11.092833 |
| COL6A2 | protein_coding | -0.4860792 | 3.04997156 |
| ZNF860 | protein_coding | -0.4865325 | 1.4666018 |
| RHCE | protein_coding | -0.4867909 | 2.80327233 |
| GET1 | protein_coding | -0.4870494 | 4.00252385 |
| HMSD | protein_coding | -0.4873689 | 1.55798127 |
| ENO1 | protein_coding | -0.4877247 | 14.4914492 |
| MAP6 | protein_coding | -0.488109 | 0.85637223 |

| | | | |
|---|---|---|---|
| PRDX6-AS1 | lncRNA | -0.4888501 | 1.27929261 |
| LINC00847 | lncRNA | -0.4892124 | 3.82578393 |
| C2orf81 | protein_coding | -0.4896395 | 0.84492869 |
| ATP5MJ | protein_coding | -0.4896879 | 11.3374041 |
| PRC1 | protein_coding | -0.4898858 | 2.64943278 |
| WDR90 | protein_coding | -0.4903274 | 8.09013398 |
| FOXO6 | protein_coding | -0.4903375 | 6.95603875 |
| PECR | protein_coding | -0.4903853 | 6.32408246 |
| ELFN2 | protein_coding | -0.4904634 | 10.7068324 |
| LHX6 | protein_coding | -0.4913361 | 2.80770988 |
| MRC2 | protein_coding | -0.4917935 | 4.43505137 |
| ARHGEF7-IT1 | lncRNA | -0.4919864 | 0.83774748 |
| RNFT2 | protein_coding | -0.4924651 | 2.16275492 |
| CTBP2 | protein_coding | -0.4930219 | 6.04286961 |
| AHCY | protein_coding | -0.4931123 | 12.8258662 |
| RP3-402G11. | lncRNA | -0.4933153 | 0.78691465 |
| GSTM3 | protein_coding | -0.4933714 | 5.85862532 |
| CDC37L1-DT | lncRNA | -0.4935811 | 0.69163732 |
| ACOT7 | protein_coding | -0.4935993 | 9.40650577 |
| OLFML2B | protein_coding | -0.4936327 | 5.43662931 |
| LIMA1 | protein_coding | -0.4936961 | 10.9615074 |
| HDAC5 | protein_coding | -0.4940069 | 9.6657856 |
| SPSB3 | protein_coding | -0.4941064 | 7.61957533 |
| CLPSL2 | protein_coding | -0.4947815 | 0.65526404 |
| GPSM3 | protein_coding | -0.4948233 | 2.38274227 |
| DKFZP434A0 | lncRNA | -0.4952305 | 0.60159243 |
| RP11-685M7 | lncRNA | -0.4953931 | 0.71939516 |
| CLCN5 | protein_coding | -0.4957618 | 12.8088439 |
| STEAP1B | protein_coding | -0.4961995 | 1.41941584 |
| DNAJC9-AS1 | lncRNA | -0.4962814 | 2.50517388 |
| HLX | protein_coding | -0.4964557 | 0.68382342 |
| FOXJ1 | protein_coding | -0.4966506 | 7.11096161 |
| ADAMTS12 | protein_coding | -0.4968041 | 2.78046075 |
| RP11-317B1 | lncRNA | -0.4970023 | 2.10386745 |
| TCF15 | protein_coding | -0.4970142 | 4.33079369 |
| TXNIP | protein_coding | -0.4973967 | 10.4886361 |
| ULK4 | protein_coding | -0.4974315 | 2.26937064 |
| SGK3 | protein_coding | -0.4976687 | 2.60078752 |
| CHRNA7 | protein_coding | -0.4979203 | 2.95180063 |
| RP5-901A4.1 | lncRNA | -0.4981724 | 1.48240716 |
| PRX | protein_coding | -0.4985008 | 3.11208365 |
| NUP62CL | protein_coding | -0.4985071 | 4.02420267 |
| ADGRA3 | protein_coding | -0.4987769 | 8.58797395 |
| ZBED5-AS1 | lncRNA | -0.4991639 | 6.499817 |
| TRIM52-AS1 | lncRNA | -0.4994613 | 2.93464598 |

| | | | |
|---|---|---|---|
| DISP2 | protein_coding | -0.500068 | 7.06236377 |
| PHF21B | protein_coding | -0.5001512 | 8.71656577 |
| FLICR | lncRNA | -0.5004196 | 0.8926723 |
| KRT85 | protein_coding | -0.500564 | 6.54365747 |
| SPAG1 | protein_coding | -0.5006431 | 4.35643172 |
| SLC6A17 | protein_coding | -0.500753 | 5.12610611 |
| CAMTA1-DT | lncRNA | -0.5010602 | 0.78537657 |
| RAC3 | protein_coding | -0.5011728 | 8.52869128 |
| RP1-310O13. | lncRNA | -0.5013353 | 2.74218867 |
| ANK2 | protein_coding | -0.5018537 | 7.8086657 |
| AC002480.4 | lncRNA | -0.5019005 | 1.77116164 |
| HK1 | protein_coding | -0.5021718 | 9.65499106 |
| GTSE1-DT | lncRNA | -0.5024369 | 2.87615166 |
| TSTD1 | protein_coding | -0.5026519 | 5.97851566 |
| NEFM | protein_coding | -0.5027143 | 11.9575713 |
| FUOM | protein_coding | -0.5030255 | 3.43858326 |
| OXA1L | protein_coding | -0.5032314 | 11.5185701 |
| RIPPLY2 | protein_coding | -0.5034581 | 1.67547981 |
| SEPTIN9 | protein_coding | -0.5035234 | 11.3767132 |
| POMGNT2 | protein_coding | -0.5036191 | 6.51541915 |
| PDK2 | protein_coding | -0.504119 | 7.02696602 |
| C1orf216 | protein_coding | -0.504161 | 6.01040126 |
| WNT8B | protein_coding | -0.5046977 | 3.05503523 |
| AKT1 | protein_coding | -0.5048252 | 13.1364441 |
| ZNF236-DT | lncRNA | -0.5051654 | 0.90768081 |
| LINC01134 | lncRNA | -0.5058519 | 2.83491542 |
| NT5M | protein_coding | -0.505922 | 5.89975503 |
| KCNMA1 | protein_coding | -0.5061713 | 9.14325808 |
| DGCR6L | protein_coding | -0.5061857 | 8.82449836 |
| C20orf204 | protein_coding | -0.5062247 | 3.14617573 |
| CTD-2501B8. | protein_coding | -0.5062953 | 0.65966001 |
| GPAM | protein_coding | -0.506424 | 8.57640301 |
| CORO1C | protein_coding | -0.5065972 | 13.2185755 |
| SFRP5 | protein_coding | -0.506617 | 1.22414341 |
| FOXJ3 | protein_coding | -0.5068299 | 10.6432229 |
| PCDH17 | protein_coding | -0.5070733 | 6.41419037 |
| TOMM7 | protein_coding | -0.5071177 | 10.5463151 |
| EDA | protein_coding | -0.5073567 | 3.70419901 |
| GEMIN8 | protein_coding | -0.5076586 | 4.87996668 |
| CREG2 | protein_coding | -0.5081741 | 8.53180421 |
| PRMT6 | protein_coding | -0.5081948 | 8.28383653 |
| JARID2-DT | lncRNA | -0.5082806 | 1.2863733 |
| SMAGP | protein_coding | -0.5086551 | 0.89455563 |
| RABAC1 | protein_coding | -0.5089524 | 2.84831116 |
| ZNF704 | protein_coding | -0.5090241 | 10.4360463 |

| | | | |
|---|---|---|---|
| VSIG10 | protein_coding | -0.5090422 | 8.54371469 |
| RP11-395I6.3 | lncRNA | -0.5094361 | 2.12800657 |
| GFUS | protein_coding | -0.5095393 | 9.32752993 |
| NUSAP1 | protein_coding | -0.5097031 | 1.40012165 |
| PRANCR | lncRNA | -0.5110126 | 4.13828636 |
| SYN2 | protein_coding | -0.5110375 | 11.3447394 |
| RINL | protein_coding | -0.5112028 | 6.81336441 |
| MAG | protein_coding | -0.5113186 | 1.77139878 |
| ITPRIPL1 | protein_coding | -0.5113443 | 9.13656806 |
| PIP4K2A | protein_coding | -0.5115604 | 6.27371675 |
| PMVK | protein_coding | -0.5121015 | 6.71935285 |
| RP11-739L10 | lncRNA | -0.5121351 | 0.71033319 |
| FLVCR1-DT | lncRNA | -0.5132937 | 2.26631065 |
| FAM157C | lncRNA | -0.5133054 | 0.80191792 |
| TBXAS1 | protein_coding | -0.5136117 | 2.50277739 |
| CH17-340M2 | lncRNA | -0.5139361 | 1.56358804 |
| TMEM14A | protein_coding | -0.5147248 | 5.51118902 |
| PHF13 | protein_coding | -0.5148544 | 7.02895733 |
| RP11-505K9. | lncRNA | -0.5150676 | 0.9023205 |
| SNRNP25 | protein_coding | -0.5152258 | 10.3303832 |
| ZNF215 | protein_coding | -0.5155178 | 7.32989582 |
| TMEM184B | protein_coding | -0.5158647 | 10.9077941 |
| DNAJC28 | protein_coding | -0.5163758 | 2.19023513 |
| RP11-314A2( | lncRNA | -0.5170571 | 0.71938757 |
| STK32C | protein_coding | -0.5171323 | 5.96745834 |
| RP11-663P9. | lncRNA | -0.5173322 | 4.53343761 |
| TNS3 | protein_coding | -0.5175012 | 3.20354487 |
| CCDC189 | protein_coding | -0.5175829 | 1.97389088 |
| MED25 | protein_coding | -0.5177437 | 11.0165875 |
| AOC3 | protein_coding | -0.5177979 | 0.97666309 |
| FAH | protein_coding | -0.5178652 | 4.63963015 |
| LHFPL3 | protein_coding | -0.5178854 | 5.57961881 |
| NARF | protein_coding | -0.5179846 | 10.7711916 |
| NIPA1 | protein_coding | -0.5180535 | 9.23759368 |
| ZDHHC24 | protein_coding | -0.5184398 | 6.6440425 |
| PARVA | protein_coding | -0.5193361 | 1.18159074 |
| RP11-792A8. | lncRNA | -0.5203536 | 2.43535338 |
| AQP4-AS1 | lncRNA | -0.5204569 | 1.73044891 |
| EPHB4 | protein_coding | -0.520646 | 8.61651319 |
| RNASET2 | protein_coding | -0.5210843 | 1.51151461 |
| TCEA2 | protein_coding | -0.5213214 | 5.38587922 |
| ERBB3 | protein_coding | -0.5213597 | 8.02251187 |
| BEST3 | protein_coding | -0.5214994 | 13.4579545 |
| PTPN13 | protein_coding | -0.5216543 | 1.24822104 |
| NUDT6 | protein_coding | -0.5220548 | 1.69804011 |

| | | | |
|---|---|---|---|
| PTPN18 | protein_coding | -0.5222801 | 6.60925759 |
| B4GALNT3 | protein_coding | -0.52239 | 14.1037684 |
| CPNE2 | protein_coding | -0.5231048 | 10.8547323 |
| DHCR24 | protein_coding | -0.5232368 | 14.0839453 |
| RP11-339B2: | lncRNA | -0.5233651 | 1.68592276 |
| SIVA1 | protein_coding | -0.5234456 | 8.65478355 |
| ID2-AS1 | lncRNA | -0.5241582 | 1.20331193 |
| CR2 | protein_coding | -0.5244072 | 2.32119783 |
| WDR5B-DT | lncRNA | -0.5249683 | 1.35475875 |
| PHC2-AS1 | lncRNA | -0.5257527 | 1.22757352 |
| TUBA4A | protein_coding | -0.5258415 | 12.2450548 |
| SLC35A2 | protein_coding | -0.5259284 | 8.87762727 |
| PLEKHD1 | protein_coding | -0.5261718 | 0.81684644 |
| NPHP3 | protein_coding | -0.5263401 | 2.79204741 |
| PARD6G | protein_coding | -0.5269618 | 4.56879175 |
| TDP1 | protein_coding | -0.526974 | 7.86858107 |
| CCDC34 | protein_coding | -0.5269767 | 5.40316871 |
| ELOB | protein_coding | -0.5272611 | 12.1652792 |
| NEDD4L | protein_coding | -0.5280391 | 7.74964296 |
| SUGT1-DT | lncRNA | -0.5285405 | 0.94972013 |
| ALG10 | protein_coding | -0.5286352 | 0.95937819 |
| LINC02206 | lncRNA | -0.5286487 | 1.21094906 |
| RP11-407N1 | lncRNA | -0.528912 | 0.6252633 |
| AC012354.6 | lncRNA | -0.5291513 | 4.26565381 |
| AP001505.1( | lncRNA | -0.5298731 | 4.26132045 |
| CFAP73 | protein_coding | -0.5301728 | 0.85669746 |
| PPP1R13L | protein_coding | -0.5306371 | 4.40674902 |
| GRID1 | protein_coding | -0.5309133 | 3.70255102 |
| ANK3 | protein_coding | -0.5311572 | 8.93650759 |
| PPT1 | protein_coding | -0.5314871 | 12.9025874 |
| AHNAK2 | protein_coding | -0.5317787 | 11.5179665 |
| ANXA2R | protein_coding | -0.5327093 | 3.81394346 |
| NPR3 | protein_coding | -0.5327122 | 1.27024637 |
| CABLES2 | protein_coding | -0.5329188 | 9.24187128 |
| SLC26A6 | protein_coding | -0.5332797 | 6.16572419 |
| TMEM19 | protein_coding | -0.5333055 | 8.99635857 |
| C21orf58 | protein_coding | -0.5334015 | 8.95447511 |
| ORAI3 | protein_coding | -0.53345 | 0.71049032 |
| C2orf76 | protein_coding | -0.5334561 | 4.17789991 |
| CASKIN1 | protein_coding | -0.5334722 | 4.1681639 |
| EXPH5 | protein_coding | -0.5334987 | 1.39316666 |
| SLC29A3 | protein_coding | -0.5336085 | 3.45631883 |
| RP4-644F6.4 | lncRNA | -0.533638 | 1.11566054 |
| MEF2D | protein_coding | -0.5339113 | 10.0484527 |
| INHBB | protein_coding | -0.5339656 | 1.49855607 |

| | | | |
|---|---|---|---|
| HIC1 | protein_coding | -0.5340225 | 1.64423635 |
| LINC02861 | lncRNA | -0.5342528 | 0.79922589 |
| UBL4B | protein_coding | -0.5346956 | 0.87106194 |
| ASPG | protein_coding | -0.5347082 | 1.13122616 |
| CENPS | protein_coding | -0.5353476 | 0.94141076 |
| ARHGEF39 | protein_coding | -0.5353679 | 7.46305666 |
| TSHZ1 | protein_coding | -0.5361605 | 9.68360555 |
| RP4-549L20.: | lncRNA | -0.5366765 | 0.71706011 |
| TMA7 | protein_coding | -0.5370965 | 8.94108065 |
| ATP1A1 | protein_coding | -0.5371199 | 16.030717 |
| RIBC2 | protein_coding | -0.5371227 | 6.59654186 |
| N4BP3 | protein_coding | -0.5371528 | 5.66314259 |
| MSC | protein_coding | -0.5371671 | 8.00463212 |
| SESTD1 | protein_coding | -0.5371686 | 9.55646922 |
| GSTP1 | protein_coding | -0.5373227 | 11.9575745 |
| HOXA13 | protein_coding | -0.537414 | 4.80269246 |
| H2AZ2 | protein_coding | -0.5377034 | 14.2022556 |
| RP11-600F24 | lncRNA | -0.5380572 | 3.95099998 |
| OPHN1 | protein_coding | -0.5382442 | 10.4217317 |
| FER1L5 | protein_coding | -0.5382909 | 0.72149803 |
| ART5 | protein_coding | -0.5385545 | 2.19556315 |
| MPP7 | protein_coding | -0.5387288 | 1.15891352 |
| KCTD19 | protein_coding | -0.5388767 | 3.17376239 |
| TOX | protein_coding | -0.5395703 | 4.29089077 |
| PRKCA | protein_coding | -0.54012 | 12.7192362 |
| B4GALT7 | protein_coding | -0.5407083 | 5.88232557 |
| DCHS1 | protein_coding | -0.5408119 | 9.76925327 |
| TLE2 | protein_coding | -0.5410141 | 10.0083821 |
| RP11-388C12 | lncRNA | -0.5410208 | 0.77666736 |
| CTD-2314G24 | lncRNA | -0.5412463 | 10.1584652 |
| RP11-31H5.3 | lncRNA | -0.5414729 | 1.01953677 |
| RP11-778D9. | lncRNA | -0.5415534 | 0.9255314 |
| BEND5 | protein_coding | -0.541979 | 7.59012886 |
| SORBS2 | protein_coding | -0.5420984 | 7.59676276 |
| SLC19A2 | protein_coding | -0.542186 | 5.17045232 |
| MT-CO3 | protein_coding | -0.5433412 | 13.9928273 |
| AC009133.15 | lncRNA | -0.543483 | 1.07442968 |
| GS1-124K5.4 | lncRNA | -0.5435033 | 2.30388785 |
| KLHDC8B | protein_coding | -0.5435126 | 2.09976022 |
| PYGL | protein_coding | -0.5440759 | 0.79474803 |
| PRRG2 | protein_coding | -0.5443871 | 2.00311482 |
| MALINC1 | lncRNA | -0.5446045 | 1.13238014 |
| PLCB3 | protein_coding | -0.5447026 | 8.80296208 |
| DNAJC3-DT | lncRNA | -0.5448418 | 4.2640836 |
| WNK3 | protein_coding | -0.5448466 | 3.68989726 |

| | | | |
|---|---|---|---|
| MIMT1 | lncRNA | -0.5448782 | 3.09246246 |
| RP11-247C2. | lncRNA | -0.5449779 | 8.777988 |
| PAQR9 | protein_coding | -0.5450703 | 4.86250295 |
| ARHGAP5-AS | lncRNA | -0.5453502 | 4.24965684 |
| TMEM145 | protein_coding | -0.545357 | 6.69064265 |
| TPBGL | protein_coding | -0.5453771 | 4.01946351 |
| LRRN3 | protein_coding | -0.5456498 | 4.61809738 |
| ABHD15 | protein_coding | -0.5461493 | 5.89619 |
| C11orf96 | protein_coding | -0.5466583 | 1.98439742 |
| TYMSOS | lncRNA | -0.5470226 | 3.65482847 |
| ANKRD9 | protein_coding | -0.5471684 | 7.91148989 |
| SYT1 | protein_coding | -0.5474148 | 11.874794 |
| SIX3 | protein_coding | -0.5474298 | 7.08237119 |
| SLC27A3 | protein_coding | -0.5476865 | 6.38152781 |
| ATP2B2 | protein_coding | -0.5478206 | 8.84791769 |
| SNHG10 | lncRNA | -0.548167 | 7.04005175 |
| LY6K | protein_coding | -0.5481744 | 9.40947278 |
| DDX11-AS1 | lncRNA | -0.5482339 | 4.06266991 |
| PNPLA2 | protein_coding | -0.548428 | 8.17069253 |
| GSTO2 | protein_coding | -0.5484613 | 2.90520364 |
| CHMP4C | protein_coding | -0.548804 | 2.56832636 |
| DACH2 | protein_coding | -0.548978 | 10.8579264 |
| RP11-585P4. | lncRNA | -0.5494833 | 1.01802635 |
| HBQ1 | protein_coding | -0.5496399 | 1.54047872 |
| RP11-327P2. | lncRNA | -0.5497281 | 2.1990694 |
| ST8SIA5 | protein_coding | -0.549874 | 9.64673528 |
| RP11-158I9.8 | lncRNA | -0.5500406 | 1.8377118 |
| TMEM139 | protein_coding | -0.5500426 | 1.48701276 |
| BAIAP2 | protein_coding | -0.5504132 | 6.48845475 |
| VAT1 | protein_coding | -0.5508501 | 12.1881514 |
| FAM166C | protein_coding | -0.5512274 | 1.30221949 |
| RP1-92O14.3 | lncRNA | -0.551628 | 0.91448644 |
| GLDC | protein_coding | -0.5520609 | 7.77313323 |
| GORASP1 | protein_coding | -0.5522166 | 9.67886728 |
| TPD52 | protein_coding | -0.5526376 | 6.32185727 |
| HNRNPD-DT | lncRNA | -0.5526528 | 1.72702068 |
| ASF1B | protein_coding | -0.5526896 | 12.5334683 |
| BCKDK | protein_coding | -0.5528115 | 8.76899636 |
| DMRTA2 | protein_coding | -0.5528254 | 1.14145251 |
| LLNLR-245B6 | lncRNA | -0.5528973 | 4.05447774 |
| LRRC42 | protein_coding | -0.5530933 | 10.2769847 |
| WSCD1 | protein_coding | -0.5531295 | 1.19898348 |
| IFTAP | protein_coding | -0.553258 | 2.03040248 |
| ARHGEF7 | protein_coding | -0.5535701 | 14.2584833 |
| INSYN1 | protein_coding | -0.5536552 | 8.64599622 |

| | | | |
|---|---|---|---|
| CCM2L | protein_coding | -0.5537981 | 5.252292 |
| ISL2 | protein_coding | -0.5538461 | 5.47174773 |
| TNS2 | protein_coding | -0.5542133 | 5.21445626 |
| ITGA9-AS1 | lncRNA | -0.5543184 | 2.12233104 |
| GALNT17 | protein_coding | -0.5544883 | 4.496461 |
| RNF20 | protein_coding | -0.5545883 | 12.3488466 |
| SLC12A6 | protein_coding | -0.5548048 | 11.4773909 |
| KCNQ2 | protein_coding | -0.5556816 | 6.33535931 |
| AC006115.7 | lncRNA | -0.5565565 | 1.68267798 |
| GLUD2 | protein_coding | -0.5574323 | 0.73893336 |
| TFAP2A-AS2 | lncRNA | -0.5575892 | 1.9302503 |
| FAM135B | protein_coding | -0.557844 | 3.54310087 |
| MEIS2 | protein_coding | -0.5578518 | 9.27116715 |
| LRRC28 | protein_coding | -0.5581476 | 10.2351824 |
| CD81 | protein_coding | -0.5581626 | 11.3168838 |
| TET1 | protein_coding | -0.5582009 | 7.72070563 |
| BEX5 | protein_coding | -0.5586218 | 2.37526169 |
| RP11-10K16. | lncRNA | -0.5593444 | 1.19661827 |
| RP5-1136G1 | lncRNA | -0.5593659 | 4.43326555 |
| SEMA4G | protein_coding | -0.5595245 | 3.92938134 |
| MPST | protein_coding | -0.5597049 | 5.60305293 |
| NPAS2 | protein_coding | -0.56034 | 0.73870764 |
| NPW | protein_coding | -0.5604772 | 1.6693246 |
| RTN4RL1 | protein_coding | -0.5606163 | 6.43790134 |
| MYLK3 | protein_coding | -0.5606504 | 4.7049057 |
| PLCD3 | protein_coding | -0.5611583 | 3.70036313 |
| CYB5D2 | protein_coding | -0.5614929 | 4.09795299 |
| ABHD17B | protein_coding | -0.5615759 | 8.53563408 |
| MELTF-AS1 | lncRNA | -0.5617454 | 3.80143279 |
| RP11-129M1 | lncRNA | -0.5618826 | 1.6468861 |
| RP11-674E16 | lncRNA | -0.5620835 | 1.0234396 |
| AC007879.2 | lncRNA | -0.5624579 | 1.8468104 |
| PCDH10 | protein_coding | -0.562492 | 7.64564972 |
| AC093627.10 | lncRNA | -0.5628139 | 2.90672006 |
| UHRF1 | protein_coding | -0.5631149 | 11.0161258 |
| PAPOLA-DT | lncRNA | -0.5631179 | 3.78199878 |
| LINC02688 | lncRNA | -0.5637288 | 0.92591236 |
| LPAR5 | protein_coding | -0.5639555 | 0.81816628 |
| YTHDF3-DT | lncRNA | -0.5641628 | 0.84540323 |
| GLS2 | protein_coding | -0.5654394 | 4.16307667 |
| SNAI1 | protein_coding | -0.5655646 | 3.12562206 |
| PHLPP1 | protein_coding | -0.5661291 | 8.25203272 |
| RP11-666O2. | lncRNA | -0.5665942 | 1.15633726 |
| RP11-762B2 | lncRNA | -0.5668795 | 0.89342925 |
| DUSP23 | protein_coding | -0.5668848 | 6.21713145 |

| | | | |
|---|---|---|---|
| AC105760.2 | lncRNA | -0.5673705 | 1.8578241 |
| C5orf38 | lncRNA | -0.5674417 | 1.07331858 |
| STMN3 | protein_coding | -0.5674917 | 13.4891143 |
| DBN1 | protein_coding | -0.5681125 | 10.6415418 |
| RP11-307C12 | lncRNA | -0.5688449 | 2.29546138 |
| SELPLG | protein_coding | -0.5690851 | 1.29873837 |
| PXMP4 | protein_coding | -0.5696089 | 10.1441523 |
| NOTCH3 | protein_coding | -0.5697315 | 1.75023358 |
| MFSD4A | protein_coding | -0.5701328 | 4.04038114 |
| GPI | protein_coding | -0.5703388 | 13.7380646 |
| SYNE1 | protein_coding | -0.57063 | 11.4385141 |
| RP11-15L13.4 | lncRNA | -0.5706831 | 0.74445963 |
| HYI | protein_coding | -0.5714499 | 5.89210262 |
| RASGRP1 | protein_coding | -0.5720873 | 10.9164875 |
| RP11-712L6.5 | lncRNA | -0.572344 | 1.08255091 |
| FAM171A2 | protein_coding | -0.5726142 | 6.88637955 |
| RNF157-AS1 | lncRNA | -0.573122 | 2.39288385 |
| RASL10B | protein_coding | -0.5731981 | 4.41007641 |
| RP11-88E10. | lncRNA | -0.5732528 | 3.13716766 |
| GRIN2A | protein_coding | -0.5738013 | 6.82645701 |
| RP1-40E16.1 | lncRNA | -0.5741093 | 1.91027871 |
| TBC1D8 | protein_coding | -0.5741365 | 9.27587647 |
| MAGI2 | protein_coding | -0.5741379 | 2.50533962 |
| LRATD1 | protein_coding | -0.5743446 | 4.55313909 |
| VASH1 | protein_coding | -0.5749958 | 9.03197059 |
| TMEM229A | protein_coding | -0.5756717 | 1.84064559 |
| ONECUT1 | protein_coding | -0.575722 | 1.78275723 |
| AGXT | protein_coding | -0.5760424 | 0.67840782 |
| UBE2V1 | protein_coding | -0.5764494 | 2.40316614 |
| RP11-498C9. | lncRNA | -0.5767572 | 2.90111173 |
| TIMM21 | protein_coding | -0.5770975 | 8.69978654 |
| YPEL1 | protein_coding | -0.5772033 | 9.27392079 |
| LINC01547 | lncRNA | -0.5778458 | 4.86779897 |
| TAFA5 | protein_coding | -0.5784444 | 7.51750845 |
| KAT2B | protein_coding | -0.5785457 | 7.9226774 |
| TMEM59L | protein_coding | -0.5796744 | 4.00745695 |
| FOXP4-AS1 | lncRNA | -0.5799283 | 2.93893731 |
| TMEM225B | protein_coding | -0.5800208 | 1.13449402 |
| TSPAN17 | protein_coding | -0.5808649 | 8.10020499 |
| RP11-120D5. | lncRNA | -0.5813493 | 1.75277605 |
| RP5-882C2.2 | lncRNA | -0.5823691 | 2.40758163 |
| OSCP1 | protein_coding | -0.5828961 | 2.99244432 |
| CTD-2619J13 | lncRNA | -0.582933 | 6.34057771 |
| SLC19A1 | protein_coding | -0.5834221 | 7.12705416 |
| ECE2 | protein_coding | -0.5840494 | 1.71271419 |

| | | | |
|---|---|---|---|
| RENO1 | lncRNA | -0.5841417 | 3.75995444 |
| SLC6A11 | protein_coding | -0.5847147 | 6.18749793 |
| AP000320.6 | lncRNA | -0.5849659 | 1.20807493 |
| HMGN2 | protein_coding | -0.5850962 | 12.0216365 |
| ZNF423 | protein_coding | -0.585152 | 2.56420917 |
| RP11-347C18 | lncRNA | -0.5853655 | 2.13229844 |
| NIFK-AS1 | lncRNA | -0.585559 | 5.45297656 |
| KIAA1614 | protein_coding | -0.5863562 | 9.92545511 |
| FLJ20021 | lncRNA | -0.586872 | 4.20062691 |
| KRT86 | protein_coding | -0.5872606 | 2.4290574 |
| MIF4GD-DT | lncRNA | -0.5874556 | 3.103895 |
| LRTOMT | protein_coding | -0.5874948 | 2.04207452 |
| NPM2 | protein_coding | -0.5875919 | 2.46863983 |
| BRF1 | protein_coding | -0.5876749 | 11.6955646 |
| SEC11C | protein_coding | -0.5878753 | 12.5385056 |
| RASSF3 | protein_coding | -0.5881386 | 9.10661158 |
| ISYNA1 | protein_coding | -0.588473 | 7.64591708 |
| FUT11 | protein_coding | -0.5885208 | 5.72771954 |
| TPGS1 | protein_coding | -0.5887028 | 2.46380137 |
| MIF-AS1 | lncRNA | -0.5891687 | 3.20061111 |
| CCNP | protein_coding | -0.5894381 | 1.26099234 |
| LPCAT2 | protein_coding | -0.5894516 | 3.25499097 |
| NTF3 | protein_coding | -0.5900182 | 2.52404435 |
| ADAMTS20 | protein_coding | -0.5901555 | 2.60446165 |
| UBC | protein_coding | -0.590315 | 11.2858914 |
| TMEM169 | protein_coding | -0.5907941 | 5.68185497 |
| LTK | protein_coding | -0.5908271 | 7.79358626 |
| EIF2AK3-DT | lncRNA | -0.5916159 | 2.09303257 |
| SRGAP3 | protein_coding | -0.591705 | 4.94725276 |
| NKX3-1 | protein_coding | -0.5917711 | 1.10438214 |
| HS6ST2 | protein_coding | -0.5925727 | 3.57695261 |
| NSMF | protein_coding | -0.5926793 | 10.4073886 |
| TST | protein_coding | -0.5932428 | 2.86604675 |
| NR4A2 | protein_coding | -0.5939323 | 6.04470286 |
| RAPGEF4 | protein_coding | -0.5942752 | 5.40690141 |
| BAHCC1 | protein_coding | -0.5943236 | 10.5759889 |
| CHD7 | protein_coding | -0.596268 | 10.1584862 |
| ID3 | protein_coding | -0.5962954 | 6.00954954 |
| TMEM191C | protein_coding | -0.5967815 | 0.92735864 |
| TMEM179 | protein_coding | -0.5970316 | 12.1035595 |
| STARD8 | protein_coding | -0.5971902 | 7.49673895 |
| ITPR3 | protein_coding | -0.5978725 | 14.0168115 |
| GPC3 | protein_coding | -0.5987024 | 11.7180966 |
| PRSS36 | protein_coding | -0.5989324 | 3.01293762 |
| TMEM9 | protein_coding | -0.5999316 | 9.30816044 |

| | | | |
|---|---|---|---|
| FKBP14-AS1 | lncRNA | -0.600377 | 1.28193344 |
| BEST4 | protein_coding | -0.6003791 | 0.84221417 |
| TCEAL9 | protein_coding | -0.6005847 | 7.9135063 |
| AC022182.3 | lncRNA | -0.6006699 | 2.63858618 |
| RP11-146F11 | lncRNA | -0.6013226 | 3.40683754 |
| EPN2 | protein_coding | -0.6017481 | 8.17925168 |
| DHRS2 | protein_coding | -0.6021079 | 4.02785339 |
| LEPR | protein_coding | -0.6022112 | 4.08005953 |
| RBMS2 | protein_coding | -0.6022193 | 2.21804022 |
| RP11-93B14. | lncRNA | -0.6023447 | 1.11189856 |
| VKORC1 | protein_coding | -0.6023704 | 7.64241424 |
| SMIM10L2A | protein_coding | -0.6028083 | 1.41320635 |
| DDAH1 | protein_coding | -0.6033413 | 10.6618867 |
| CTB-25B13.5 | lncRNA | -0.6034295 | 2.94248397 |
| RXFP3 | protein_coding | -0.6041867 | 6.26284148 |
| RAB3C | protein_coding | -0.6046622 | 6.7972331 |
| RIN2 | protein_coding | -0.6051675 | 1.49775434 |
| RP11-269F19 | lncRNA | -0.6052419 | 1.68180517 |
| CDKN2B | protein_coding | -0.6062739 | 9.85776442 |
| TRMT9B | protein_coding | -0.6062934 | 8.01576897 |
| XXbac-B135H | lncRNA | -0.6063844 | 1.41874011 |
| MYRF | protein_coding | -0.606903 | 4.45917305 |
| PEG3 | protein_coding | -0.6071017 | 14.5548963 |
| MYCBPAP | protein_coding | -0.6081237 | 2.01185054 |
| ADGRL3 | protein_coding | -0.6088995 | 8.14206712 |
| SPATC1L | protein_coding | -0.6089677 | 5.45349202 |
| MESP1 | protein_coding | -0.6097905 | 4.63833411 |
| NOTCH1 | protein_coding | -0.6101361 | 1.58278174 |
| RP11-1017G1 | lncRNA | -0.6101768 | 2.8948756 |
| AC009961.3 | lncRNA | -0.6101802 | 2.05587259 |
| CDC25C | protein_coding | -0.6111087 | 5.86051115 |
| NETO1 | protein_coding | -0.6115243 | 3.55705615 |
| SEMA3F | protein_coding | -0.6123418 | 5.20044872 |
| SREBF2-AS1 | lncRNA | -0.6123649 | 1.49699522 |
| FLYWCH2 | protein_coding | -0.6130056 | 9.0368909 |
| SDF2L1 | protein_coding | -0.6131388 | 5.72125003 |
| LPAR3 | protein_coding | -0.6136571 | 1.62847603 |
| PCDHGC4 | protein_coding | -0.6137697 | 5.87460551 |
| ZDHHC12 | protein_coding | -0.6138102 | 11.3509622 |
| EPS8L1 | protein_coding | -0.6139211 | 6.30131578 |
| LINC00957 | lncRNA | -0.6140666 | 8.16262764 |
| SFXN1 | protein_coding | -0.61427 | 9.87956634 |
| SDC1 | protein_coding | -0.6146198 | 9.16714596 |
| HIGD2A | protein_coding | -0.6146204 | 9.94415838 |
| SLC18A2 | protein_coding | -0.6156602 | 3.32684243 |

| | | | |
|---|---|---|---|
| XKRX | protein_coding | -0.6175884 | 3.14831219 |
| RP11-474G2: | lncRNA | -0.61772 | 1.18392999 |
| RP11-669I1.2 | lncRNA | -0.6183305 | 2.20297928 |
| GPER1 | protein_coding | -0.6186764 | 2.18046764 |
| ARID5B | protein_coding | -0.6189586 | 5.3824754 |
| IQCD | protein_coding | -0.6196047 | 3.1267809 |
| RP13-270P17 | lncRNA | -0.619986 | 0.99648268 |
| GMIP | protein_coding | -0.6203024 | 8.71059108 |
| RP11-394O2. | lncRNA | -0.62045 | 0.77446552 |
| RP11-527J8.1 | lncRNA | -0.6206796 | 1.63852678 |
| NENF | protein_coding | -0.6207853 | 11.1493802 |
| NAPA-AS1 | lncRNA | -0.6210596 | 1.70450104 |
| ANXA11 | protein_coding | -0.6214769 | 10.2100787 |
| STOML3 | protein_coding | -0.6216754 | 1.02699046 |
| ZNF710-AS1 | lncRNA | -0.6218122 | 6.11778737 |
| RP11-378A13 | lncRNA | -0.6218139 | 1.64705737 |
| GRB7 | protein_coding | -0.6223239 | 7.38408495 |
| PHACTR2 | protein_coding | -0.6225617 | 2.93909211 |
| RP11-172H24 | lncRNA | -0.6226022 | 2.07673275 |
| ZNF706 | protein_coding | -0.6236526 | 11.6552018 |
| RP11-521B24 | lncRNA | -0.6238124 | 8.3725908 |
| NSG1 | protein_coding | -0.6238174 | 11.2436505 |
| AC004540.4 | lncRNA | -0.6239312 | 2.33577332 |
| MID1 | protein_coding | -0.6245218 | 6.61360448 |
| ZDHHC2 | protein_coding | -0.6248092 | 4.83134225 |
| ZSCAN1 | protein_coding | -0.6249998 | 1.40182065 |
| PRXL2A | protein_coding | -0.6255845 | 11.8843988 |
| FHOD1 | protein_coding | -0.6266029 | 9.22481341 |
| RP11-485M7 | lncRNA | -0.62681 | 1.47721108 |
| PHLDB3 | protein_coding | -0.627171 | 11.7767401 |
| RAB40B | protein_coding | -0.6273314 | 7.72665729 |
| SLC2A1-AS1 | lncRNA | -0.6277727 | 1.79260081 |
| TTC14 | protein_coding | -0.6279465 | 10.0854854 |
| FAM20C | protein_coding | -0.6280984 | 7.37887675 |
| VWA2 | protein_coding | -0.6281548 | 2.8231975 |
| CH17-343M1 | lncRNA | -0.6288247 | 0.97697565 |
| KDM7A | protein_coding | -0.6289213 | 9.40342456 |
| NSG2 | protein_coding | -0.6294222 | 8.96929776 |
| RP11-363J20 | lncRNA | -0.629969 | 1.71108557 |
| ELK3 | protein_coding | -0.6321403 | 2.08144318 |
| TRERF1 | protein_coding | -0.6325366 | 8.7445114 |
| SLC9A1 | protein_coding | -0.6331199 | 12.0293627 |
| SH3PXD2B | protein_coding | -0.6331729 | 8.67425551 |
| STX1B | protein_coding | -0.6332131 | 2.67008099 |
| ULBP2 | protein_coding | -0.6341294 | 1.15136764 |

| | | | |
|---|---|---|---|
| BICC1 | protein_coding | -0.6344867 | 3.93168048 |
| DYRK4 | protein_coding | -0.6349563 | 2.29624297 |
| GAB1 | protein_coding | -0.6351052 | 5.99964247 |
| FAM167A-AS | lncRNA | -0.635644 | 4.36881358 |
| CCDC78 | protein_coding | -0.637829 | 2.31270326 |
| LINC00115 | lncRNA | -0.6380528 | 0.96731525 |
| DISP1 | protein_coding | -0.638083 | 8.48153039 |
| JAKMIP3 | protein_coding | -0.638097 | 4.44811794 |
| YEATS2-AS1 | lncRNA | -0.6381084 | 1.54503765 |
| PHTF2 | protein_coding | -0.6381386 | 12.0673147 |
| ZNNT1 | lncRNA | -0.6389386 | 3.77338146 |
| CTB-31O20.2 | lncRNA | -0.6392911 | 5.36486897 |
| RP11-9G1.3 | lncRNA | -0.6393333 | 2.04143797 |
| LAMC3 | protein_coding | -0.6396951 | 3.50201726 |
| AC002480.3 | lncRNA | -0.639779 | 3.48445018 |
| CPLX2 | protein_coding | -0.6401499 | 14.56168 |
| RUNDC3A-AS | lncRNA | -0.640484 | 6.77725916 |
| KPNB1-DT | lncRNA | -0.6411746 | 0.72496197 |
| PNPLA7 | protein_coding | -0.6417361 | 6.29500235 |
| RP11-867G2 | lncRNA | -0.6418188 | 4.33653563 |
| LZTFL1 | protein_coding | -0.6428989 | 4.98889964 |
| FOXO1 | protein_coding | -0.6429859 | 1.52349346 |
| TMEM256 | protein_coding | -0.6436333 | 1.54431487 |
| PLEKHA2 | protein_coding | -0.6438278 | 7.95308659 |
| ITGAE | protein_coding | -0.6439283 | 7.17282927 |
| RP11-245D1 | lncRNA | -0.6439331 | 2.48864573 |
| SLC4A8 | protein_coding | -0.644089 | 3.24916896 |
| TMEM185A | protein_coding | -0.6446182 | 6.45470953 |
| PCDH11X | protein_coding | -0.6447839 | 7.16478507 |
| RASSF7 | protein_coding | -0.6453669 | 9.22231039 |
| KRT10 | protein_coding | -0.6454716 | 6.68203898 |
| NUP62 | protein_coding | -0.6466257 | 13.779423 |
| RP11-575F1 | lncRNA | -0.6468636 | 4.86244518 |
| CDK19 | protein_coding | -0.6481442 | 10.1298709 |
| THEG | protein_coding | -0.648421 | 6.36869646 |
| SNAP25 | protein_coding | -0.6488206 | 13.1933829 |
| FRRS1L | protein_coding | -0.6489035 | 2.02798384 |
| LA16c-OS12. | lncRNA | -0.6489618 | 1.07616477 |
| LPCAT1 | protein_coding | -0.6490816 | 13.8277559 |
| TSPAN5 | protein_coding | -0.649224 | 4.99234669 |
| KDM3A | protein_coding | -0.6497156 | 13.0552164 |
| RIMBP2 | protein_coding | -0.6498555 | 5.81099865 |
| ZNF497-AS1 | lncRNA | -0.6500461 | 1.56983884 |
| KDM4A-AS1 | lncRNA | -0.6503513 | 3.73361044 |
| RP11-449P1 | lncRNA | -0.6517161 | 1.00275617 |

| | | | |
|---|---|---|---|
| RP11-247I13 | lncRNA | -0.6522546 | 2.3416436 |
| UACA | protein_coding | -0.6523425 | 3.92043191 |
| TOX3 | protein_coding | -0.6526927 | 8.16182022 |
| CHADL | protein_coding | -0.6537823 | 0.89522104 |
| ARHGEF7-AS | lncRNA | -0.653784 | 1.24730363 |
| DISP3 | protein_coding | -0.6544472 | 2.65432508 |
| KCNAB3 | protein_coding | -0.6545674 | 7.01038291 |
| IKZF2 | protein_coding | -0.6547193 | 0.90557827 |
| KRT81 | protein_coding | -0.6548048 | 2.25964989 |
| RP3-467L1.6 | lncRNA | -0.6548558 | 0.85929378 |
| PCOLCE2 | protein_coding | -0.6551881 | 10.8304153 |
| ENTPD8 | protein_coding | -0.6564659 | 3.65986622 |
| SEPTIN5 | protein_coding | -0.6567835 | 1.65512652 |
| MYORG | protein_coding | -0.6570011 | 6.86258412 |
| ZFHX3 | protein_coding | -0.6570647 | 4.91907741 |
| MID1IP1-AS1 | lncRNA | -0.6571191 | 1.80553336 |
| RP11-135M8 | lncRNA | -0.6578206 | 1.15161168 |
| RP11-124N1 | lncRNA | -0.6588742 | 3.84107384 |
| RP11-128N1 | lncRNA | -0.6592117 | 1.67951267 |
| RP5-1024G6. | lncRNA | -0.65944 | 1.17155198 |
| TEK | protein_coding | -0.6595779 | 1.81176249 |
| NCALD | protein_coding | -0.6600824 | 6.38846003 |
| PDK3 | protein_coding | -0.6601706 | 10.7744212 |
| INHBC | protein_coding | -0.6601782 | 1.2188425 |
| FHL3 | protein_coding | -0.660512 | 5.80267415 |
| EIF2B5-DT | lncRNA | -0.6605923 | 2.43463365 |
| PDE1C | protein_coding | -0.6609113 | 8.63466778 |
| THOC3 | protein_coding | -0.6614642 | 5.97946824 |
| FRAT2 | protein_coding | -0.6614759 | 5.43677812 |
| AMD1 | protein_coding | -0.6618514 | 12.5322055 |
| RTN4R | protein_coding | -0.6619105 | 6.67645776 |
| BRWD1-AS2 | lncRNA | -0.6621614 | 2.22137423 |
| SCNN1A | protein_coding | -0.6637511 | 15.8878748 |
| LINC01521 | lncRNA | -0.6651304 | 5.80608871 |
| REPS2 | protein_coding | -0.6652758 | 7.702869 |
| IGSF22 | protein_coding | -0.6657972 | 1.90105406 |
| ZCCHC24 | protein_coding | -0.6659335 | 4.06822986 |
| RP11-79N23. | lncRNA | -0.6659516 | 1.61420623 |
| TMEM134 | protein_coding | -0.6670888 | 9.17401662 |
| FOXD2-AS1 | lncRNA | -0.6679426 | 8.06565898 |
| DBH-AS1 | lncRNA | -0.6680641 | 5.98215563 |
| RAB8B | protein_coding | -0.6682518 | 8.80196555 |
| SHROOM2 | protein_coding | -0.6688913 | 11.0493485 |
| PCSK1 | protein_coding | -0.6697036 | 12.211124 |
| CTSL | protein_coding | -0.6697512 | 1.0727126 |

| | | | |
|---|---|---|---|
| CALM1 | protein_coding | -0.6698627 | 16.2885579 |
| ARHGAP29 | protein_coding | -0.6706961 | 2.43053094 |
| CCNI2 | protein_coding | -0.6724876 | 1.56295008 |
| S100A1 | protein_coding | -0.6725277 | 4.91457286 |
| LINC01315 | lncRNA | -0.6726007 | 3.22288039 |
| TMEM255B | protein_coding | -0.6726994 | 1.40877187 |
| SPAG5-AS1 | lncRNA | -0.6727361 | 0.93597787 |
| BHLHE22 | protein_coding | -0.6736024 | 8.58001671 |
| LINC01006 | lncRNA | -0.6740961 | 2.6717845 |
| KCNK15-AS1 | lncRNA | -0.6749933 | 1.34074116 |
| ZNF395 | protein_coding | -0.675115 | 9.51667727 |
| RP11-91G21. | lncRNA | -0.6756691 | 2.93280199 |
| PGK1 | protein_coding | -0.6763318 | 15.6736215 |
| EP300-AS1 | lncRNA | -0.6764325 | 3.14311494 |
| RP4-605O3.4 | lncRNA | -0.676502 | 1.30908555 |
| SEPT5-GP1B | protein_coding | -0.6768557 | 1.76667655 |
| ASB1 | protein_coding | -0.6775168 | 12.8158201 |
| PLCL1 | protein_coding | -0.6776629 | 5.0442851 |
| ASPDH | protein_coding | -0.6787422 | 7.50252218 |
| NAV2 | protein_coding | -0.6797288 | 14.5453504 |
| SSBP4 | protein_coding | -0.6797643 | 7.22163003 |
| SALL4 | protein_coding | -0.6800499 | 3.56142578 |
| METTL27 | protein_coding | -0.6803007 | 2.23646347 |
| BTF3-DT | lncRNA | -0.6806052 | 2.09353378 |
| LINC02607 | lncRNA | -0.68068 | 5.58020742 |
| BEGAIN | protein_coding | -0.6810297 | 7.28951678 |
| FNDC11 | protein_coding | -0.6811458 | 3.9851898 |
| TRIB1 | protein_coding | -0.6815628 | 9.29544494 |
| WIPI1 | protein_coding | -0.6822925 | 1.41916353 |
| CTD-2555A7. | lncRNA | -0.682693 | 4.86650238 |
| CTD-2506P8. | lncRNA | -0.6832162 | 3.19343037 |
| GINS2 | protein_coding | -0.6832705 | 13.4047252 |
| RP11-575F12 | lncRNA | -0.6843559 | 0.74636938 |
| RP11-149I2.4 | lncRNA | -0.6848165 | 0.80884594 |
| NAV1 | protein_coding | -0.6851865 | 13.0057482 |
| RP11-15H20. | lncRNA | -0.6866292 | 3.91747859 |
| KB-1410C5.5 | lncRNA | -0.6872517 | 0.74650243 |
| ADGRE2 | protein_coding | -0.6876275 | 0.98059409 |
| SLC25A1 | protein_coding | -0.6885757 | 8.70159833 |
| PXDN | protein_coding | -0.6886131 | 8.54715762 |
| CRYL1 | protein_coding | -0.6888257 | 3.92353004 |
| C4orf54 | protein_coding | -0.6889203 | 1.93878099 |
| APC2 | protein_coding | -0.6889218 | 2.1840218 |
| TMPO-AS1 | lncRNA | -0.6902371 | 8.63989513 |
| GPR68 | protein_coding | -0.6903244 | 1.82292954 |

| | | | |
|---|---|---|---|
| GEMIN7-AS1 | lncRNA | -0.690593 | 1.05107645 |
| MARCHF4 | protein_coding | -0.6907034 | 1.05582451 |
| C11orf94 | protein_coding | -0.6917587 | 1.07812469 |
| BAIAP2-DT | lncRNA | -0.6920151 | 8.77726988 |
| PITPNA-AS1 | lncRNA | -0.6936261 | 5.50341213 |
| CTD-2270P14 | lncRNA | -0.6941893 | 3.15088975 |
| TMEM64 | protein_coding | -0.6949537 | 4.21425428 |
| FOXN3 | protein_coding | -0.6957131 | 15.2036042 |
| NKD1 | protein_coding | -0.695822 | 5.99262659 |
| CITED2 | protein_coding | -0.6967767 | 8.64795516 |
| INPP5B | protein_coding | -0.6970841 | 9.76736381 |
| MAP4K1 | protein_coding | -0.6976172 | 8.06386981 |
| SNAI3 | protein_coding | -0.6987587 | 1.1277533 |
| PRRG1 | protein_coding | -0.6993081 | 8.00169673 |
| TLR4 | protein_coding | -0.6995643 | 1.28357966 |
| RGS11 | protein_coding | -0.6995792 | 3.20976329 |
| SVIL | protein_coding | -0.6998622 | 8.16555058 |
| RP1-8B1.4 | lncRNA | -0.6999507 | 1.33944677 |
| FLNC | protein_coding | -0.7001163 | 2.5622246 |
| C3orf70 | protein_coding | -0.7002424 | 4.99191158 |
| RP11-118J22 | lncRNA | -0.7005556 | 1.66887278 |
| ATRN | protein_coding | -0.7009673 | 8.38941056 |
| RP4-768P8.2 | lncRNA | -0.7012632 | 1.75591607 |
| CDH23 | protein_coding | -0.7019536 | 7.35404467 |
| CCDC136 | protein_coding | -0.7023571 | 8.9143863 |
| THRB | protein_coding | -0.7032439 | 4.74298561 |
| BRINP1 | protein_coding | -0.7032877 | 11.4588163 |
| ASAP3 | protein_coding | -0.7036125 | 7.01107583 |
| KRT19 | protein_coding | -0.704429 | 8.71059941 |
| RP11-45P15. | lncRNA | -0.7044494 | 1.7790023 |
| CTD-3220F14 | lncRNA | -0.7050324 | 0.58530082 |
| PCOLCE | protein_coding | -0.7050458 | 2.83025409 |
| LLNLR-225F1 | protein_coding | -0.7060207 | 1.39643839 |
| RP11-318E3. | lncRNA | -0.7062264 | 1.23486743 |
| CFD | protein_coding | -0.7065439 | 5.3325858 |
| RP13-131K19 | lncRNA | -0.7065963 | 1.47405104 |
| RP1-315G1.3 | lncRNA | -0.7067652 | 1.60753693 |
| RAI14 | protein_coding | -0.7072601 | 9.07135221 |
| FOXN3-AS1 | lncRNA | -0.7074459 | 3.91490042 |
| AC005301.9 | lncRNA | -0.7079415 | 8.10783099 |
| CTSV | protein_coding | -0.7081543 | 6.82490401 |
| FAAH | protein_coding | -0.708338 | 9.06979128 |
| YPEL2 | protein_coding | -0.7085506 | 8.94993394 |
| BCAM | protein_coding | -0.7086537 | 1.99390938 |
| WDR54 | protein_coding | -0.7088465 | 10.3822704 |

| | | | |
|---|---|---|---|
| SAP30-DT | lncRNA | -0.7092719 | 5.48368748 |
| CPNE7 | protein_coding | -0.709465 | 5.66448084 |
| AP006621.8 | lncRNA | -0.7095294 | 4.0089794 |
| ZNF778 | protein_coding | -0.7101845 | 13.2139895 |
| RASAL3 | protein_coding | -0.7104916 | 1.13292099 |
| LINC01659 | lncRNA | -0.7106736 | 1.66017876 |
| TFAP2B | protein_coding | -0.7107374 | 11.2842187 |
| RP11-1072C1 | lncRNA | -0.7108803 | 1.91688404 |
| SOBP | protein_coding | -0.7108911 | 6.03544276 |
| RHPN1-AS1 | lncRNA | -0.7110117 | 1.64270944 |
| VAMP1 | protein_coding | -0.7112855 | 6.1522477 |
| CARMIL1 | protein_coding | -0.7116982 | 11.2702712 |
| HHAT | protein_coding | -0.7118989 | 1.34821428 |
| P3H4 | protein_coding | -0.7127042 | 6.63924285 |
| UGT8 | protein_coding | -0.7131123 | 12.5185039 |
| LINC01305 | lncRNA | -0.713335 | 2.02687468 |
| DLG5-AS1 | lncRNA | -0.713375 | 2.23275119 |
| SIAE | protein_coding | -0.7135756 | 2.74253884 |
| WASIR2 | lncRNA | -0.7137375 | 3.2904075 |
| POU2F2 | protein_coding | -0.7151674 | 2.90715133 |
| SNX32 | protein_coding | -0.7152156 | 3.04846889 |
| DTD1 | protein_coding | -0.7152838 | 3.58418711 |
| MORF4L2-AS | lncRNA | -0.7163685 | 1.51056963 |
| MAST4 | protein_coding | -0.717048 | 6.24089857 |
| LFNG | protein_coding | -0.7171318 | 8.06915856 |
| OST4 | protein_coding | -0.7176157 | 10.6650715 |
| LINC02106 | lncRNA | -0.7177706 | 4.11694528 |
| SLAMF8 | protein_coding | -0.7178151 | 1.5003443 |
| CDK15 | protein_coding | -0.7179592 | 0.93161536 |
| TNNT1 | protein_coding | -0.718147 | 11.1904755 |
| RIMKLA | protein_coding | -0.7188919 | 10.8469848 |
| ITPRIP-AS1 | lncRNA | -0.7201664 | 1.29947504 |
| CNKSR3 | protein_coding | -0.7203029 | 2.33087478 |
| INSYN1-AS1 | lncRNA | -0.7204747 | 3.4512003 |
| CDCA7L | protein_coding | -0.7210519 | 14.7767471 |
| GLDN | protein_coding | -0.7214742 | 1.54818335 |
| FAM102B | protein_coding | -0.7223482 | 12.5905924 |
| RP11-316O1 | lncRNA | -0.7225439 | 2.83041805 |
| F3 | protein_coding | -0.7232729 | 2.03359186 |
| TRAPPC6A | protein_coding | -0.7241614 | 7.85408226 |
| SDCBP2 | protein_coding | -0.7243806 | 1.59161073 |
| ELOA-AS1 | lncRNA | -0.7250256 | 3.89782201 |
| TMEM99 | lncRNA | -0.725068 | 5.75530592 |
| B3GNT4 | protein_coding | -0.7255682 | 10.1420668 |
| PAK6 | protein_coding | -0.7257833 | 9.80610271 |

| | | | |
|---|---|---|---|
| SYNGAP1-AS | lncRNA | -0.7275821 | 1.07697918 |
| GRIN2C | protein_coding | -0.7277551 | 9.49721449 |
| RDH10 | protein_coding | -0.7281597 | 9.85880923 |
| FZD1 | protein_coding | -0.7282081 | 6.99920112 |
| POF1B | protein_coding | -0.7295208 | 9.79921373 |
| NPTX2 | protein_coding | -0.7296263 | 9.59789556 |
| PTGR1 | protein_coding | -0.7296547 | 10.0686433 |
| FN3K | protein_coding | -0.7298412 | 4.5552734 |
| ACVR1C | protein_coding | -0.7303284 | 1.09745905 |
| DENND2A | protein_coding | -0.7308695 | 6.19078367 |
| RP11-182L21 | lncRNA | -0.731648 | 1.73332176 |
| PABPC5 | protein_coding | -0.7322821 | 5.78979635 |
| BCORL1 | protein_coding | -0.732554 | 8.06666247 |
| RP11-111M2 | lncRNA | -0.7327491 | 6.68040521 |
| SNTG2-AS1 | lncRNA | -0.733322 | 4.40074515 |
| HEIH | lncRNA | -0.734025 | 8.16951119 |
| GAS2L2 | protein_coding | -0.7341987 | 1.44346624 |
| FBXL22 | protein_coding | -0.7351286 | 2.5294627 |
| WASF3 | protein_coding | -0.7357872 | 11.1955208 |
| CFAP61-AS1 | lncRNA | -0.7362276 | 6.85317242 |
| RUNX3 | protein_coding | -0.7363014 | 6.93784093 |
| LTBR | protein_coding | -0.7366782 | 7.00213422 |
| EMC1-AS1 | lncRNA | -0.7368253 | 1.60509816 |
| CCL28 | protein_coding | -0.7368337 | 10.4783611 |
| RP11-477E3. | lncRNA | -0.7371113 | 3.47378877 |
| RGR | protein_coding | -0.7385537 | 1.2067819 |
| ST8SIA5-DT | lncRNA | -0.7387386 | 7.4575324 |
| NXPH3 | protein_coding | -0.7388088 | 7.50315023 |
| CTB-131K11. | lncRNA | -0.738929 | 2.25544522 |
| RP11-331F9. | lncRNA | -0.7391947 | 1.264795 |
| ASIC4 | protein_coding | -0.7408309 | 10.4078845 |
| C6orf223 | lncRNA | -0.7408648 | 8.28009162 |
| CKB | protein_coding | -0.7410241 | 10.8800229 |
| AC000068.5 | lncRNA | -0.7412453 | 3.95917138 |
| TRIM62 | protein_coding | -0.7414151 | 11.8324079 |
| DRC3 | protein_coding | -0.7414899 | 1.70895289 |
| SLC45A3 | protein_coding | -0.7417724 | 2.4031669 |
| SLC2A4 | protein_coding | -0.7431218 | 0.78669759 |
| VPS37D | protein_coding | -0.7431501 | 5.51716477 |
| ZSCAN16-AS | lncRNA | -0.7432076 | 6.81508722 |
| FAM163B | protein_coding | -0.7432732 | 3.51942543 |
| EFHD1 | protein_coding | -0.7438444 | 2.53585939 |
| CATSPERE | protein_coding | -0.7446561 | 1.11669754 |
| NR3C2 | protein_coding | -0.7449376 | 4.84070403 |
| RP5-1148A2: | lncRNA | -0.7449435 | 1.39220735 |

| | | | |
|---|---|---|---|
| REEP6 | protein_coding | -0.7457489 | 4.46216983 |
| KCNG3 | protein_coding | -0.746051 | 5.59997994 |
| FHL1 | protein_coding | -0.7461926 | 10.3283672 |
| KRT15 | protein_coding | -0.7466642 | 6.05384501 |
| SERINC2 | protein_coding | -0.7466806 | 13.8684171 |
| CNMD | protein_coding | -0.7468969 | 1.76768618 |
| FERMT3 | protein_coding | -0.7471737 | 6.28985304 |
| MAML3 | protein_coding | -0.7474589 | 12.6360859 |
| SHISAL1 | protein_coding | -0.7484335 | 5.10566559 |
| LINC01353 | lncRNA | -0.7490837 | 2.11993658 |
| LINC01882 | lncRNA | -0.7493497 | 3.20914484 |
| POU4F1 | protein_coding | -0.7496929 | 14.6666955 |
| KDF1 | protein_coding | -0.7504128 | 8.83081327 |
| DNAJC6 | protein_coding | -0.7512441 | 12.2665658 |
| TTLL9 | protein_coding | -0.7516375 | 0.76968139 |
| FGGY-DT | lncRNA | -0.7518546 | 2.17029575 |
| CTD-2314B2: | lncRNA | -0.7527681 | 1.75119819 |
| KCTD8 | protein_coding | -0.7531397 | 4.94781149 |
| SNRPD3 | protein_coding | -0.75439 | 14.4805679 |
| ZNF846 | protein_coding | -0.7553903 | 1.22387151 |
| ALDOC | protein_coding | -0.7554427 | 12.0080696 |
| AC009506.1 | lncRNA | -0.7556592 | 5.71300971 |
| CTNND2 | protein_coding | -0.7559389 | 3.44081448 |
| CCDC3 | protein_coding | -0.7559596 | 11.4963323 |
| PPP1R21-DT | lncRNA | -0.7560885 | 1.96695085 |
| PAK5 | protein_coding | -0.7567103 | 5.75839919 |
| TTC9 | protein_coding | -0.7570297 | 3.63796913 |
| FLJ37453 | lncRNA | -0.7571023 | 7.39550985 |
| ADAP1 | protein_coding | -0.7575827 | 1.3225221 |
| LINC01786 | lncRNA | -0.7578945 | 0.96283547 |
| CTC-563A5.5 | lncRNA | -0.7579931 | 5.67848031 |
| AC034243.1 | lncRNA | -0.7580064 | 2.2081404 |
| CABP7 | protein_coding | -0.7581577 | 14.3915717 |
| RP3-508I15.2 | lncRNA | -0.7582841 | 1.33062168 |
| CNFN | protein_coding | -0.7590304 | 2.93815039 |
| GRIN1 | protein_coding | -0.759847 | 8.89556674 |
| CFAP99 | protein_coding | -0.7598591 | 1.00533385 |
| CTC-425O23. | lncRNA | -0.7602051 | 2.51840584 |
| MAP3K13 | protein_coding | -0.7602101 | 9.3827549 |
| SYCE2 | protein_coding | -0.7606322 | 5.07343538 |
| BEND4 | protein_coding | -0.7611805 | 9.57876141 |
| CSPG5 | protein_coding | -0.7613063 | 6.49779204 |
| TSPAN7 | protein_coding | -0.7623198 | 8.72680162 |
| RP11-398J5.: | lncRNA | -0.7625308 | 1.77686754 |
| PGM1 | protein_coding | -0.7625982 | 9.18071101 |

| | | | |
|---|---|---|---|
| SCG2 | protein_coding | -0.7630738 | 2.78545848 |
| RP11-798M1 | lncRNA | -0.7631309 | 2.95287676 |
| RP11-486I11 | lncRNA | -0.7637451 | 1.65637576 |
| HBEGF | protein_coding | -0.7638719 | 4.29391792 |
| DHRS3 | protein_coding | -0.7639619 | 9.33894347 |
| CTD-2286N8. | lncRNA | -0.7642366 | 1.8958239 |
| CYBA | protein_coding | -0.7650171 | 9.4500439 |
| OTULINL | protein_coding | -0.765296 | 12.8202702 |
| C1orf115 | protein_coding | -0.7653305 | 11.5954225 |
| TMEM132E | protein_coding | -0.7653343 | 7.8650846 |
| LINC02874 | lncRNA | -0.7655881 | 1.55428574 |
| SNHG19 | lncRNA | -0.7663487 | 5.65281254 |
| YPEL3 | protein_coding | -0.7663867 | 11.4292906 |
| ZNF8-DT | lncRNA | -0.7672731 | 5.33908775 |
| SRRM2-AS1 | lncRNA | -0.7677047 | 2.67516216 |
| TM6SF1 | protein_coding | -0.7680206 | 2.01292068 |
| THORLNC | lncRNA | -0.7680946 | 1.09637278 |
| RP11-256E16 | lncRNA | -0.7686431 | 0.73664702 |
| RETREG1 | protein_coding | -0.76995 | 9.01459419 |
| KIAA1549 | protein_coding | -0.7704782 | 5.43830936 |
| CYB5A | protein_coding | -0.7707457 | 12.0045498 |
| FES | protein_coding | -0.7711882 | 3.43458575 |
| FIGNL2 | protein_coding | -0.7717012 | 3.61868561 |
| ZFPM2 | protein_coding | -0.7717238 | 1.37184287 |
| KIF21B | protein_coding | -0.7723677 | 4.33740943 |
| PLD5 | protein_coding | -0.772473 | 12.6757365 |
| RIMS4 | protein_coding | -0.7735833 | 11.8678061 |
| C9orf163 | lncRNA | -0.7738069 | 2.43783974 |
| CD14 | protein_coding | -0.7741877 | 1.72502116 |
| MAFF | protein_coding | -0.7748077 | 5.49392028 |
| ANAPC15 | protein_coding | -0.7748431 | 11.6156347 |
| NANOS1 | protein_coding | -0.7760707 | 4.42131022 |
| P4HA1 | protein_coding | -0.7769847 | 11.0413673 |
| KCNH1 | protein_coding | -0.7770385 | 2.61886568 |
| SEPTIN1 | protein_coding | -0.777178 | 2.19957309 |
| STKLD1 | protein_coding | -0.7785051 | 3.09030948 |
| LINC02696 | lncRNA | -0.7785492 | 1.7471536 |
| HPDL | protein_coding | -0.7785924 | 8.93997956 |
| NPR1 | protein_coding | -0.7792806 | 1.24388532 |
| PDZRN3 | protein_coding | -0.7797218 | 11.7516788 |
| DACT1 | protein_coding | -0.7798892 | 13.3751217 |
| PRIMA1 | protein_coding | -0.7800781 | 6.7791868 |
| SIPA1 | protein_coding | -0.7810382 | 9.78736772 |
| PLA2G4F | protein_coding | -0.7812995 | 8.88201206 |
| SOX15 | protein_coding | -0.7816343 | 6.56093832 |

| | | | |
|---|---|---|---|
| RP11-755F1( | lncRNA | -0.781777 | 1.21844817 |
| H3-3A-DT | lncRNA | -0.781821 | 1.10243787 |
| TP53I11 | protein_coding | -0.7819331 | 13.4531685 |
| RP11-281A2( | lncRNA | -0.7821559 | 2.08682023 |
| SHD | protein_coding | -0.7825285 | 10.1907484 |
| CHST1 | protein_coding | -0.7840835 | 4.88351675 |
| DAPL1 | protein_coding | -0.7860739 | 4.73297099 |
| XXbac-B476( | lncRNA | -0.7862706 | 1.22862687 |
| ASTL | protein_coding | -0.786633 | 3.97903226 |
| DUSP15 | protein_coding | -0.7870275 | 4.84074237 |
| CFAP298-TCI | protein_coding | -0.7879703 | 2.10389075 |
| NDUFV2-AS1 | lncRNA | -0.7896186 | 1.95855548 |
| DLC1 | protein_coding | -0.7904659 | 2.81836992 |
| CCDC96 | protein_coding | -0.7908369 | 2.90760313 |
| C1QL1 | protein_coding | -0.7913298 | 8.15599373 |
| ANO7 | protein_coding | -0.7914415 | 1.07639961 |
| CHD5 | protein_coding | -0.791661 | 2.89148182 |
| RP11-73M18 | lncRNA | -0.7916949 | 1.41868584 |
| LLNLR-268E1 | lncRNA | -0.7917408 | 1.80647008 |
| CBLN4 | protein_coding | -0.791857 | 2.37037515 |
| SCIRT | lncRNA | -0.7923243 | 6.51003703 |
| RP11-83N9.6 | lncRNA | -0.7923796 | 1.55716084 |
| MESP2 | protein_coding | -0.7926981 | 1.84706199 |
| RP11-382N1 | lncRNA | -0.7947399 | 2.21913471 |
| SEC24D | protein_coding | -0.7954839 | 9.6340962 |
| CTC-786C10. | lncRNA | -0.795856 | 2.03698079 |
| ZFP92 | protein_coding | -0.7960492 | 3.08545691 |
| CTSH | protein_coding | -0.7961819 | 6.0907475 |
| PPP4R3B-DT | lncRNA | -0.7969684 | 2.56411822 |
| RUFY3 | protein_coding | -0.7971516 | 7.94821004 |
| ZNF521 | protein_coding | -0.797202 | 5.791803 |
| SLC66A3 | protein_coding | -0.7980483 | 4.92482805 |
| PROX1 | protein_coding | -0.7988039 | 13.0529811 |
| SEMA6A-AS2 | lncRNA | -0.7997108 | 2.04194052 |
| RP11-416N2. | lncRNA | -0.7997661 | 1.7589208 |
| ADORA2A | protein_coding | -0.7997834 | 1.53954942 |
| CNGB1 | protein_coding | -0.799816 | 1.51021913 |
| F12 | protein_coding | -0.8014578 | 6.59682729 |
| TUBB8 | protein_coding | -0.8028721 | 7.59122366 |
| SELENON | protein_coding | -0.803358 | 12.3494006 |
| RP11-482D24 | lncRNA | -0.8046804 | 1.04565845 |
| PAQR8 | protein_coding | -0.8064407 | 6.93000888 |
| STAT6 | protein_coding | -0.8066986 | 3.79734071 |
| CFAP100 | protein_coding | -0.8080976 | 3.00626482 |
| NACC2 | protein_coding | -0.8087258 | 4.0133633 |

| | | | |
|---|---|---|---|
| RP11-118F19 | lncRNA | -0.8091405 | 2.2543554 |
| ODF2L | protein_coding | -0.8104232 | 3.62717664 |
| CARHSP1 | protein_coding | -0.8105594 | 15.7142905 |
| FZD8 | protein_coding | -0.8106006 | 5.16925775 |
| HRH1 | protein_coding | -0.8112766 | 6.24837185 |
| LRRTM1 | protein_coding | -0.8122662 | 10.5251782 |
| MPP2 | protein_coding | -0.8124209 | 9.16859498 |
| AGMAT | protein_coding | -0.8136305 | 2.68451137 |
| MGAT5B | protein_coding | -0.815559 | 6.50619367 |
| AKAP1-DT | lncRNA | -0.8157773 | 2.14175812 |
| RP11-378A13 | lncRNA | -0.8188254 | 1.14546297 |
| SLC8A1 | protein_coding | -0.8190246 | 13.2427721 |
| AMOT | protein_coding | -0.820031 | 11.7310398 |
| TMEM121 | protein_coding | -0.8208356 | 4.23467948 |
| H19 | lncRNA | -0.8212991 | 2.81970645 |
| CCSER1 | protein_coding | -0.8216244 | 7.23453326 |
| RP11-167N4. | lncRNA | -0.8217152 | 2.66230148 |
| CTD-2162K18 | lncRNA | -0.8239848 | 7.53017207 |
| HCN2 | protein_coding | -0.8241755 | 10.4779419 |
| RP4-647C14. | lncRNA | -0.8247495 | 2.10340957 |
| CHRNA4 | protein_coding | -0.8251388 | 4.52836721 |
| PALM | protein_coding | -0.8254714 | 12.1884853 |
| ZBTB7C | protein_coding | -0.8257286 | 5.56464387 |
| TUBA4B | protein_coding | -0.8268621 | 6.82259832 |
| EHBP1L1 | protein_coding | -0.8273952 | 9.3609768 |
| ZNF853 | protein_coding | -0.8275564 | 1.98692573 |
| CTB-113I20.2 | lncRNA | -0.8280859 | 1.31597387 |
| KANK2 | protein_coding | -0.8285181 | 10.7236893 |
| RP11-108L7. | lncRNA | -0.8300874 | 2.73974282 |
| CLIC6 | protein_coding | -0.8320665 | 10.6683043 |
| ABHD4 | protein_coding | -0.8322561 | 4.14520948 |
| RP11-802D6. | lncRNA | -0.8326318 | 1.68617649 |
| DDX19A-DT | lncRNA | -0.8329262 | 5.20103801 |
| RP11-112J3. | lncRNA | -0.833231 | 2.06831963 |
| PITPNM1 | protein_coding | -0.8342575 | 3.98441385 |
| SAMD15 | protein_coding | -0.8362853 | 1.42238658 |
| BMP8A | protein_coding | -0.8364845 | 4.9556214 |
| C10orf82 | protein_coding | -0.8375029 | 3.09766579 |
| ETV1 | protein_coding | -0.8376181 | 10.8174919 |
| FAM162A | protein_coding | -0.8377471 | 11.9651683 |
| RBP4 | protein_coding | -0.8378673 | 1.60572742 |
| PIGZ | protein_coding | -0.8400018 | 1.91045597 |
| RP11-475I24 | lncRNA | -0.8413631 | 1.43485713 |
| CTA-29F11.1 | lncRNA | -0.8419852 | 7.93848995 |
| PRADC1 | protein_coding | -0.8424635 | 11.2255919 |

| | | | |
|---|---|---|---|
| FBLIM1 | protein_coding | -0.8425929 | 1.90560541 |
| VWA1 | protein_coding | -0.8435552 | 5.71536848 |
| MYO1A | protein_coding | -0.8445785 | 4.17476452 |
| AC079807.4 | lncRNA | -0.8446334 | 2.16845707 |
| KIF7 | protein_coding | -0.8447598 | 2.893109 |
| CALN1 | protein_coding | -0.8450097 | 12.5204201 |
| KLF2 | protein_coding | -0.8463708 | 2.8213119 |
| DMRTA1 | protein_coding | -0.8465414 | 14.8308777 |
| RP11-384P7. | lncRNA | -0.8466459 | 3.73210251 |
| SUSD2 | protein_coding | -0.8467096 | 2.29963144 |
| LRRC4 | protein_coding | -0.8496763 | 5.98696893 |
| METTL24 | protein_coding | -0.8497648 | 3.1304669 |
| BMP3 | protein_coding | -0.8501561 | 1.30152475 |
| MIR4733HG | lncRNA | -0.8531744 | 1.26940824 |
| CTB-178M22 | lncRNA | -0.8533029 | 2.05786167 |
| RP11-481J2.3 | lncRNA | -0.8539491 | 3.01951242 |
| ADGRA2 | protein_coding | -0.8544416 | 7.85768354 |
| ADARB2 | protein_coding | -0.8544712 | 5.79104224 |
| PDK1 | protein_coding | -0.8547588 | 12.7435011 |
| SLC7A14 | protein_coding | -0.8552212 | 13.1686495 |
| CADM2 | protein_coding | -0.8569353 | 4.87509466 |
| REEP2 | protein_coding | -0.8571983 | 2.25957569 |
| C17orf97 | protein_coding | -0.8580656 | 2.77998834 |
| CTD-2260A1 | lncRNA | -0.8581689 | 7.09436272 |
| CACNA1H | protein_coding | -0.8588899 | 6.8896193 |
| TPM4 | protein_coding | -0.8593706 | 13.6567265 |
| JAKMIP1 | protein_coding | -0.859549 | 1.72227849 |
| RECK | protein_coding | -0.8599119 | 2.56235967 |
| AC005329.7 | lncRNA | -0.8602326 | 6.05189243 |
| SLIT2 | protein_coding | -0.8603802 | 12.9344457 |
| SCN9A | protein_coding | -0.8604914 | 4.43591529 |
| ETV2 | protein_coding | -0.8606192 | 2.15370209 |
| ARL4C | protein_coding | -0.8613401 | 5.98887566 |
| BMP7 | protein_coding | -0.8615192 | 15.500064 |
| RCCD1-AS1 | lncRNA | -0.8616812 | 1.6175065 |
| GPR156 | protein_coding | -0.8619443 | 2.38926061 |
| MAPK13 | protein_coding | -0.8624171 | 12.6064614 |
| SARDH | protein_coding | -0.8628193 | 8.14525253 |
| CTD-2015H6. | lncRNA | -0.8628966 | 5.81609188 |
| CTB-58E17.5 | lncRNA | -0.8632474 | 5.80066228 |
| CAMK2B | protein_coding | -0.8647979 | 9.2069805 |
| BICDL2 | protein_coding | -0.865899 | 2.85797607 |
| RP11-1055B8 | lncRNA | -0.8660514 | 3.827033 |
| ENHO | protein_coding | -0.8671733 | 8.69812602 |
| IRX2 | protein_coding | -0.8684969 | 6.69975149 |

| | | | |
|---|---|---|---|
| RP11-314B1. | lncRNA | -0.8701002 | 9.51150932 |
| RNASEH2A | protein_coding | -0.8707172 | 2.66115414 |
| BMX | protein_coding | -0.8716542 | 3.10886619 |
| FBXO15 | protein_coding | -0.8721794 | 2.25462901 |
| TEX9 | protein_coding | -0.8733098 | 6.82053108 |
| PTPRE | protein_coding | -0.873781 | 3.55860861 |
| RGS9BP | protein_coding | -0.8738441 | 1.97285261 |
| ZNF471 | protein_coding | -0.8753686 | 6.79995469 |
| TNFRSF10A | protein_coding | -0.8756087 | 4.94190136 |
| THEM6 | protein_coding | -0.8760574 | 13.2938546 |
| KCNJ3 | protein_coding | -0.8762319 | 2.43841569 |
| ADPRHL1 | protein_coding | -0.8765765 | 6.96404355 |
| DOK4 | protein_coding | -0.8814584 | 3.8309036 |
| GRIN3A | protein_coding | -0.8818474 | 8.33933698 |
| LMCD1 | protein_coding | -0.8820578 | 4.48174698 |
| NRGN | protein_coding | -0.8821862 | 9.28481225 |
| AGBL4 | protein_coding | -0.8823031 | 6.55573683 |
| KAZN | protein_coding | -0.882517 | 3.74449796 |
| CENATAC-DT | lncRNA | -0.8828071 | 8.20448232 |
| CASTOR2 | protein_coding | -0.8832336 | 8.61462998 |
| EFNB3 | protein_coding | -0.8845071 | 5.33512898 |
| PDGFC | protein_coding | -0.8846353 | 5.26190095 |
| INPPL1 | protein_coding | -0.8854943 | 17.1610354 |
| HECW2 | protein_coding | -0.8855303 | 6.12471269 |
| SHOX2 | protein_coding | -0.886335 | 15.5635256 |
| ELOVL2-AS1 | lncRNA | -0.8876241 | 1.61956254 |
| RP11-685G9. | lncRNA | -0.8877687 | 1.62794082 |
| CFAP65 | protein_coding | -0.8879338 | 3.66436995 |
| GHRL | protein_coding | -0.8882576 | 1.64635714 |
| ILDR2 | protein_coding | -0.888399 | 13.1067745 |
| STAMBPL1 | protein_coding | -0.8888555 | 4.22174235 |
| FBXO24 | protein_coding | -0.8895482 | 1.81844048 |
| RP11-296I10 | lncRNA | -0.8895839 | 3.98989943 |
| FBN3 | protein_coding | -0.8909595 | 9.48822707 |
| C4orf48 | protein_coding | -0.8921086 | 1.75882853 |
| RAMP2-AS1 | lncRNA | -0.892141 | 2.19572578 |
| MYADML2 | protein_coding | -0.8923253 | 1.84507956 |
| HOXA11 | protein_coding | -0.8930498 | 5.65562431 |
| BAIAP3 | protein_coding | -0.8931707 | 8.3390444 |
| RAB3B | protein_coding | -0.8957884 | 11.2779745 |
| LRRC8D-DT | lncRNA | -0.8960585 | 4.0326824 |
| NYAP2 | protein_coding | -0.8966897 | 11.3275811 |
| SIT1 | protein_coding | -0.9013605 | 4.50854178 |
| CAVIN1 | protein_coding | -0.9014751 | 8.10522707 |
| PFKFB4 | protein_coding | -0.9021444 | 9.33662503 |

| | | | |
|---|---|---|---|
| FGD4 | protein_coding | -0.9031051 | 4.27683959 |
| GPAT2 | protein_coding | -0.9034969 | 7.33518857 |
| ADAM28 | protein_coding | -0.9039389 | 3.89337158 |
| BMP8B | protein_coding | -0.9049461 | 9.31947817 |
| MINCR | lncRNA | -0.9057122 | 4.8845272 |
| NUDT18 | protein_coding | -0.9068085 | 2.78088848 |
| LINC01143 | lncRNA | -0.9070069 | 5.34160511 |
| PAIP2B | protein_coding | -0.9072003 | 13.1878822 |
| ANKRD37 | protein_coding | -0.9087949 | 7.09833638 |
| C2CD4C | protein_coding | -0.9097772 | 6.89850684 |
| DPYSL4 | protein_coding | -0.9099536 | 12.0764491 |
| ILRUN-AS1 | lncRNA | -0.9117162 | 4.70568644 |
| TFDP2 | protein_coding | -0.9123191 | 6.66005399 |
| KCTD17 | protein_coding | -0.9124522 | 10.6518518 |
| NSMAF | protein_coding | -0.9133895 | 1.95556499 |
| LLfos-22E10. | lncRNA | -0.9136542 | 7.43799052 |
| ZNF850 | protein_coding | -0.9137902 | 9.74859249 |
| SESN3 | protein_coding | -0.9140876 | 15.4724221 |
| PLEKHF1 | protein_coding | -0.9145371 | 3.33471556 |
| AC022154.7 | lncRNA | -0.9163663 | 1.79290702 |
| LCMT1-AS1 | lncRNA | -0.916534 | 1.9557328 |
| DCXR-DT | lncRNA | -0.9170205 | 1.49126314 |
| RP11-18O11. | lncRNA | -0.9171608 | 5.58510405 |
| WI2-1959D1! | lncRNA | -0.9175327 | 1.82324663 |
| RP1-159A19. | lncRNA | -0.9179574 | 2.47357628 |
| TNNI1 | protein_coding | -0.9190879 | 1.33738469 |
| RP11-531A2 | lncRNA | -0.9206779 | 1.39179536 |
| MELTF | protein_coding | -0.9211308 | 8.4478575 |
| FOXD2 | protein_coding | -0.9215046 | 6.41725845 |
| FRAT1 | protein_coding | -0.9246551 | 3.29470644 |
| AC137932.5 | lncRNA | -0.924971 | 3.3823568 |
| MPP3 | protein_coding | -0.9250351 | 3.18678937 |
| PTGES2-AS1 | lncRNA | -0.9256017 | 1.22279982 |
| KLF3 | protein_coding | -0.9259314 | 8.95746737 |
| RP11-568J23 | lncRNA | -0.9260186 | 1.52490758 |
| SMIM3 | protein_coding | -0.9262461 | 2.33536643 |
| MMP28 | protein_coding | -0.9264427 | 2.28713609 |
| AMPD3 | protein_coding | -0.9271552 | 4.51583624 |
| ANO3 | protein_coding | -0.9272913 | 4.32478353 |
| VIPR1 | protein_coding | -0.9277106 | 5.30367415 |
| RP11-295G2 | lncRNA | -0.928084 | 2.92479371 |
| JAG2 | protein_coding | -0.9287736 | 11.1937192 |
| FOXD1 | protein_coding | -0.9295453 | 6.51813963 |
| NOS1 | protein_coding | -0.9340548 | 2.32850027 |
| AC006026.13 | lncRNA | -0.9369468 | 2.61245764 |

| | | | |
|---|---|---|---|
| RP11-25K19. | lncRNA | -0.9371983 | 1.98223803 |
| LINC01971 | lncRNA | -0.9400203 | 4.83324319 |
| PLAAT1 | protein_coding | -0.9406652 | 5.81304578 |
| MEP1B | protein_coding | -0.9426771 | 2.06322609 |
| NIBAN2 | protein_coding | -0.943007 | 11.8480485 |
| LINC01566 | lncRNA | -0.9435203 | 7.09970908 |
| CLDN7 | protein_coding | -0.9456112 | 8.2314306 |
| KRT24 | protein_coding | -0.9457501 | 3.76626448 |
| AMH | protein_coding | -0.9469205 | 4.76721306 |
| RP4-561L24.3 | lncRNA | -0.9476851 | 1.69287372 |
| ARL10 | protein_coding | -0.9480321 | 9.77996245 |
| CCDC28A-AS1 | lncRNA | -0.9485849 | 1.84583836 |
| RP11-15E18. | lncRNA | -0.949743 | 1.69454468 |
| LINC01239 | lncRNA | -0.9499792 | 4.87234325 |
| MTSS1 | protein_coding | -0.9502082 | 6.97335643 |
| ST | protein_coding | -0.950971 | 2.32953598 |
| CBLN2 | protein_coding | -0.9520553 | 4.06631363 |
| AJAP1 | protein_coding | -0.9524447 | 4.37192346 |
| ITGA2B | protein_coding | -0.9547365 | 10.1004081 |
| ZNF185 | protein_coding | -0.9550819 | 6.31509367 |
| RP11-793H13 | lncRNA | -0.9552243 | 3.14013814 |
| LINC02765 | lncRNA | -0.9564526 | 2.22561012 |
| LINC01431 | lncRNA | -0.9571717 | 2.4349845 |
| AFF2 | protein_coding | -0.9573361 | 1.22528562 |
| RAB3A | protein_coding | -0.9582406 | 13.1953123 |
| IGFBP2 | protein_coding | -0.959421 | 15.3701687 |
| NR0B2 | protein_coding | -0.9601692 | 3.89407148 |
| ADSS1 | protein_coding | -0.9606303 | 3.49394571 |
| CNTN4 | protein_coding | -0.961058 | 5.80134312 |
| ZMAT4 | protein_coding | -0.9611248 | 9.03878363 |
| TMEM191B | protein_coding | -0.9652562 | 2.54117094 |
| PTPN21 | protein_coding | -0.9670505 | 8.85675787 |
| CAMK4 | protein_coding | -0.9682178 | 15.0230879 |
| CERNA1 | lncRNA | -0.9698453 | 2.92600759 |
| RP11-373A6. | lncRNA | -0.9699457 | 3.71943729 |
| RP11-567M1 | lncRNA | -0.9704062 | 7.54453862 |
| SORL1 | protein_coding | -0.9708502 | 9.91047877 |
| SOX2 | protein_coding | -0.9725018 | 15.1408865 |
| REEP1 | protein_coding | -0.9728267 | 8.16020019 |
| CD207 | protein_coding | -0.9728825 | 3.84338438 |
| DNAI4 | protein_coding | -0.97321 | 2.67805122 |
| SCN4B | protein_coding | -0.974977 | 5.12747902 |
| NAT8L | protein_coding | -0.9751397 | 8.59474884 |
| GAMT | protein_coding | -0.9754406 | 11.0044413 |
| HOXB4 | protein_coding | -0.9797339 | 3.12701588 |

| | | | |
|---|---|---|---|
| GRK1 | protein_coding | -0.9816556 | 2.38808213 |
| NEUROG2-A | lncRNA | -0.9845393 | 1.89162269 |
| LINC01719 | lncRNA | -0.9859435 | 1.5523782 |
| RP11-286N2: | lncRNA | -0.9861833 | 2.58818606 |
| C19orf81 | protein_coding | -0.9869449 | 2.26119081 |
| F8 | protein_coding | -0.9870563 | 4.36588781 |
| LINC01484 | lncRNA | -0.9883616 | 3.85888578 |
| RP11-152P23 | lncRNA | -0.9885383 | 2.24592865 |
| HCG25 | lncRNA | -0.9886009 | 1.95062301 |
| LINC01679 | lncRNA | -0.9892092 | 10.5336029 |
| RP11-672L1( | lncRNA | -0.9898986 | 1.70661978 |
| LINC01994 | lncRNA | -0.9904148 | 1.6680976 |
| MGAT4C | protein_coding | -0.9908368 | 9.0596402 |
| BSG-AS1 | lncRNA | -0.9914929 | 2.59286206 |
| MINAR1 | protein_coding | -0.993226 | 9.44501805 |
| DPP10-AS1 | lncRNA | -0.9943764 | 4.71242861 |
| RP11-382A2( | lncRNA | -0.9954335 | 1.33293771 |
| TRIM29 | protein_coding | -0.9954599 | 5.88742354 |
| CTD-2336O2. | lncRNA | -0.9970809 | 2.02780185 |
| GPX7 | protein_coding | -0.9999788 | 7.81805356 |
| B3GLCT | protein_coding | -1.0000111 | 12.5149237 |
| KIF5A | protein_coding | -1.0002023 | 9.5619599 |
| RP5-908M14 | lncRNA | -1.0009214 | 5.35762148 |
| HTR1D | protein_coding | -1.001732 | 4.19001414 |
| GRTP1 | protein_coding | -1.0022685 | 1.99851288 |
| AP003068.12 | lncRNA | -1.0050865 | 1.7192747 |
| COL8A1 | protein_coding | -1.0062643 | 14.3016268 |
| SCOC-AS1 | lncRNA | -1.006667 | 3.9288448 |
| TIMP3 | protein_coding | -1.006757 | 13.9618593 |
| EIF4E1B | protein_coding | -1.0069052 | 2.12302604 |
| PTENP1-AS | lncRNA | -1.0073299 | 9.77814511 |
| DAGLA | protein_coding | -1.0081489 | 14.2049218 |
| ADCYAP1R1 | protein_coding | -1.0101316 | 2.17299057 |
| RP1-76B20.1 | lncRNA | -1.0110212 | 1.9935334 |
| SEMA6B | protein_coding | -1.0113748 | 5.2647675 |
| BTBD8 | protein_coding | -1.013808 | 3.09548968 |
| KANK1 | protein_coding | -1.0149365 | 15.2104909 |
| XKR5 | protein_coding | -1.0163997 | 5.51302686 |
| PHKA2-AS1 | lncRNA | -1.01657 | 2.29900049 |
| BTBD11 | protein_coding | -1.016725 | 2.58874991 |
| ADORA2B | protein_coding | -1.0176066 | 2.70958699 |
| AMOTL2 | protein_coding | -1.0220687 | 10.7168112 |
| DANCR | lncRNA | -1.0229235 | 13.2909886 |
| PEG10 | protein_coding | -1.0241746 | 14.4409972 |
| AC004540.5 | lncRNA | -1.0244194 | 2.07691681 |

| | | | |
|---|---|---|---|
| RP11-657O9. | lncRNA | -1.0260243 | 3.09642531 |
| RP11-480D4. | lncRNA | -1.026463 | 2.31345634 |
| AC013463.2 | lncRNA | -1.0273477 | 6.94335857 |
| KCNT2 | protein_coding | -1.0284318 | 1.82532565 |
| ACTG1 | protein_coding | -1.0284436 | 18.586501 |
| MFSD2A | protein_coding | -1.0325693 | 12.4097053 |
| CBARP | protein_coding | -1.0327118 | 4.30039971 |
| AC012314.8 | lncRNA | -1.0347293 | 2.19941752 |
| PLOD1 | protein_coding | -1.0353315 | 10.5134498 |
| SOAT2 | protein_coding | -1.0354718 | 1.453581 |
| C1QL3 | protein_coding | -1.0359074 | 2.26395293 |
| CORO1A | protein_coding | -1.0359691 | 12.3686776 |
| RP11-139K4. | lncRNA | -1.0371608 | 2.06697849 |
| TNNI3 | protein_coding | -1.0388664 | 7.82792912 |
| HK2-DT | lncRNA | -1.0390845 | 4.96923765 |
| LRRC10B | protein_coding | -1.0408894 | 7.80736674 |
| CCNT2-AS1 | lncRNA | -1.0414436 | 4.98513212 |
| CECR2 | protein_coding | -1.0428259 | 10.4151053 |
| CYP51A1-AS1 | lncRNA | -1.0428341 | 1.65051354 |
| SERTAD4 | protein_coding | -1.0436575 | 7.71541471 |
| NT5DC2 | protein_coding | -1.045802 | 16.1879321 |
| FAM13A | protein_coding | -1.0464631 | 10.0805038 |
| LIMD1 | protein_coding | -1.0474929 | 9.96520154 |
| ASIC1 | protein_coding | -1.0477613 | 8.63800225 |
| KB-1440D3.1 | lncRNA | -1.0487402 | 2.68995217 |
| RNF213-AS1 | lncRNA | -1.0491298 | 2.46547154 |
| TRPA1 | protein_coding | -1.0497307 | 5.99437743 |
| PNMA2 | protein_coding | -1.0500873 | 5.9482396 |
| MYCBP | protein_coding | -1.0503659 | 11.7790377 |
| TOLLIP-AS1 | lncRNA | -1.0507487 | 3.24955957 |
| AC083864.4 | lncRNA | -1.0508216 | 2.08967873 |
| ZSWIM5 | protein_coding | -1.0515985 | 8.88501778 |
| TFAP2E | protein_coding | -1.0535482 | 2.55183297 |
| ACO1 | protein_coding | -1.0568538 | 13.5547864 |
| ADAMTS15 | protein_coding | -1.0571053 | 8.96211659 |
| SBNO1-AS1 | lncRNA | -1.0586675 | 3.53999918 |
| AMPD2 | protein_coding | -1.0587374 | 15.724354 |
| IMPA2 | protein_coding | -1.0603825 | 9.00187709 |
| RN7SL832P | lncRNA | -1.0623707 | 1.79815392 |
| CPLX1 | protein_coding | -1.0626947 | 9.25423636 |
| PERP | protein_coding | -1.0633768 | 10.6176108 |
| KCNQ1 | protein_coding | -1.0643285 | 10.8440732 |
| RP11-571L19 | lncRNA | -1.0648461 | 1.76143415 |
| BNIP3L | protein_coding | -1.0655162 | 14.6688534 |
| GAL3ST1 | protein_coding | -1.0671094 | 11.4785437 |

| | | | |
|---|---|---|---|
| DYNC1I1 | protein_coding | -1.068315 | 5.39583694 |
| STK32A-AS1 | lncRNA | -1.0729929 | 2.57991924 |
| PTPRD | protein_coding | -1.0757085 | 12.4300609 |
| CLEC11A | protein_coding | -1.0765953 | 4.7356085 |
| CHRM4 | protein_coding | -1.079097 | 10.0531987 |
| PSLNR | lncRNA | -1.0798183 | 5.89509199 |
| RP4-657D16. | lncRNA | -1.0804316 | 1.46252805 |
| CTSF | protein_coding | -1.0808388 | 11.7506792 |
| U47924.31 | lncRNA | -1.0824614 | 6.01145701 |
| OVOL2 | protein_coding | -1.0830171 | 6.02441351 |
| RP4-564M11 | lncRNA | -1.083524 | 2.32974155 |
| RP11-712B9. | lncRNA | -1.0852695 | 4.71015236 |
| PCDH19 | protein_coding | -1.0862851 | 7.49457975 |
| MTNR1B | protein_coding | -1.0874046 | 1.75468586 |
| NAV3 | protein_coding | -1.0897311 | 5.16267097 |
| KLHL35 | protein_coding | -1.0904701 | 2.96711058 |
| OPCML | protein_coding | -1.0911759 | 2.22923614 |
| RP11-197P3. | lncRNA | -1.092034 | 1.3197323 |
| DEGS2 | protein_coding | -1.0927467 | 8.7377165 |
| RFLNB | protein_coding | -1.0946297 | 11.0508635 |
| MT3 | protein_coding | -1.0959024 | 7.54390157 |
| ADGRD2 | protein_coding | -1.0983816 | 1.91308877 |
| LINC02846 | lncRNA | -1.1002924 | 6.21011037 |
| CLMN | protein_coding | -1.1012679 | 11.3626667 |
| DHDDS-AS1 | lncRNA | -1.1017387 | 2.84780581 |
| MEF2C-AS2 | lncRNA | -1.1050373 | 2.08925992 |
| ZNF232-AS1 | lncRNA | -1.105504 | 5.5504205 |
| UPP1 | protein_coding | -1.1065018 | 5.7503672 |
| GAL3ST3 | protein_coding | -1.1066671 | 4.73021074 |
| RGS7BP | protein_coding | -1.1067773 | 2.72986419 |
| LEF1 | protein_coding | -1.1069118 | 3.75781984 |
| MECOM | protein_coding | -1.1071271 | 10.8035479 |
| RP11-351C21 | lncRNA | -1.1079606 | 2.19426713 |
| GNB3 | protein_coding | -1.1090296 | 8.52764332 |
| CTC-338M12 | lncRNA | -1.1098197 | 2.62116734 |
| RP11-467K18 | lncRNA | -1.1106847 | 4.74072261 |
| MTMR2 | protein_coding | -1.1163805 | 7.83639061 |
| ADRA2C | protein_coding | -1.1174995 | 11.2152872 |
| FLRT1 | protein_coding | -1.1177269 | 1.87337466 |
| PROX1-AS1 | lncRNA | -1.1196208 | 7.7934983 |
| MAGEH1 | protein_coding | -1.1196871 | 2.32121254 |
| NKX2-4 | protein_coding | -1.1236864 | 5.97579536 |
| LARGE2 | protein_coding | -1.1238292 | 10.7525544 |
| KLHL14 | protein_coding | -1.1239759 | 5.58159042 |
| PLEKHH1 | protein_coding | -1.1240809 | 6.31693136 |

| | | | |
|---|---|---|---|
| SLITRK4 | protein_coding | -1.1248286 | 6.82047054 |
| CERK | protein_coding | -1.1261942 | 17.8290844 |
| RP11-80F22. | lncRNA | -1.1264852 | 1.9750084 |
| RP11-389G6. | lncRNA | -1.1270413 | 2.23444465 |
| IER3-AS1 | lncRNA | -1.1289197 | 1.62683335 |
| FMN2 | protein_coding | -1.1289597 | 12.0663572 |
| RP1-212P9.3 | lncRNA | -1.1289983 | 1.98521147 |
| DPP10 | protein_coding | -1.1303493 | 11.0075532 |
| NDST3 | protein_coding | -1.1310851 | 5.06895931 |
| MYOCD | protein_coding | -1.1323122 | 11.1795511 |
| MTURN | protein_coding | -1.1326497 | 16.6143618 |
| MPPED1 | protein_coding | -1.1327067 | 7.59739159 |
| SMIM5 | protein_coding | -1.1358398 | 2.76132759 |
| MTFP1 | protein_coding | -1.1368272 | 4.24111706 |
| WNT16 | protein_coding | -1.1375265 | 5.06727966 |
| KCNJ12 | protein_coding | -1.1378682 | 5.12802784 |
| P3H3 | protein_coding | -1.1379109 | 10.4238615 |
| ADGRG5 | protein_coding | -1.1388256 | 6.05316863 |
| GALK1 | protein_coding | -1.1393329 | 10.2443281 |
| RP11-686O6. | lncRNA | -1.1406326 | 4.88500616 |
| PHYHIP | protein_coding | -1.1411066 | 2.37632285 |
| PCCA | protein_coding | -1.1425065 | 5.05305397 |
| AC005076.5 | lncRNA | -1.1425116 | 4.04373621 |
| RP11-326F6. | lncRNA | -1.1434655 | 5.85443425 |
| EYA1 | protein_coding | -1.1436999 | 5.70697218 |
| XXbac-BPG1! | lncRNA | -1.144185 | 2.32848406 |
| NEURL1 | protein_coding | -1.1443218 | 11.4750255 |
| CPM | protein_coding | -1.1451557 | 9.81634358 |
| MARVELD3 | protein_coding | -1.1465567 | 8.93732753 |
| LINC00592 | lncRNA | -1.1467863 | 3.05654006 |
| RP1-118J21.: | lncRNA | -1.1469906 | 2.08808489 |
| PRRT4 | protein_coding | -1.1506773 | 5.80219332 |
| ARHGEF11 | protein_coding | -1.1509557 | 9.10462943 |
| PRKD3 | protein_coding | -1.1515228 | 11.4063925 |
| STK32A | protein_coding | -1.1516453 | 15.5894879 |
| MOCOS | protein_coding | -1.1519401 | 4.96950579 |
| TMEM159 | protein_coding | -1.1521034 | 3.46381399 |
| KHDC1 | protein_coding | -1.1528626 | 5.24398347 |
| KCNT1 | protein_coding | -1.1544013 | 7.82862596 |
| GLB1L3 | protein_coding | -1.1551462 | 6.64357165 |
| SEPTIN10 | protein_coding | -1.1554905 | 4.67240287 |
| PHF24 | protein_coding | -1.1557468 | 5.17520589 |
| TMIGD2 | protein_coding | -1.1572292 | 7.49496638 |
| SLC13A3 | protein_coding | -1.1577556 | 9.04321316 |
| Tcommon | protein_coding | -1.1605055 | 8.45906528 |

| | | | |
|---|---|---|---|
| KCNG1 | protein_coding | -1.1635922 | 10.0946875 |
| EPHA5 | protein_coding | -1.1703492 | 3.00220644 |
| HHEX | protein_coding | -1.1733532 | 6.57834073 |
| CA9 | protein_coding | -1.1743873 | 17.9556028 |
| LINC02691 | lncRNA | -1.1746709 | 8.48875192 |
| CTSB | protein_coding | -1.1777526 | 6.4645194 |
| HEYL | protein_coding | -1.1788605 | 6.52471807 |
| TGFB1I1 | protein_coding | -1.1814014 | 3.59828033 |
| ESAM | protein_coding | -1.1836073 | 3.95176962 |
| KCNIP2 | protein_coding | -1.1880814 | 9.80635333 |
| RAB9B | protein_coding | -1.1881989 | 4.35422158 |
| RASGRP2 | protein_coding | -1.1955928 | 2.63316486 |
| C1GALT1C1L | protein_coding | -1.1956728 | 3.8860549 |
| DOCK5 | protein_coding | -1.1985241 | 3.94219012 |
| PDE3A | protein_coding | -1.2004241 | 10.9584631 |
| UBE2QL1 | protein_coding | -1.2011787 | 8.06969657 |
| RP3-454M7.: | lncRNA | -1.2018738 | 3.6241848 |
| RP11-738E2: | lncRNA | -1.2027064 | 2.12170649 |
| GPR161 | protein_coding | -1.2032299 | 8.89993643 |
| RGS19 | protein_coding | -1.2033092 | 7.44756601 |
| PCDH7 | protein_coding | -1.2039047 | 4.62837439 |
| CNTNAP2 | protein_coding | -1.2043654 | 10.0133933 |
| NUDT8 | protein_coding | -1.2046931 | 4.98344862 |
| TUBB6 | protein_coding | -1.210482 | 2.88149598 |
| ITGB2 | protein_coding | -1.2106396 | 2.85355032 |
| TMEM98 | protein_coding | -1.2123969 | 11.9578147 |
| RUNDC3B | protein_coding | -1.2137129 | 3.82613353 |
| PRDM6 | protein_coding | -1.2160248 | 8.536799 |
| UNC5A | protein_coding | -1.2198448 | 13.2853652 |
| ODC1-DT | lncRNA | -1.2214279 | 3.35885308 |
| KIF26A | protein_coding | -1.2217524 | 10.6744409 |
| CD6 | protein_coding | -1.2218105 | 2.35112646 |
| NDRG1 | protein_coding | -1.2264539 | 17.0235567 |
| RP11-316M2 | lncRNA | -1.2266393 | 3.68979589 |
| GPR62 | protein_coding | -1.2289049 | 2.19884186 |
| RP11-6N13.1 | lncRNA | -1.2293191 | 2.25196549 |
| RP5-921G16. | lncRNA | -1.2296794 | 3.18991051 |
| GABBR2 | protein_coding | -1.2304826 | 8.08571143 |
| HMGCLL1 | protein_coding | -1.2324015 | 2.83871405 |
| POU4F2 | protein_coding | -1.2337322 | 4.29125468 |
| RP4-813D12. | lncRNA | -1.2354326 | 2.32263387 |
| SLC22A17 | protein_coding | -1.2356373 | 8.74759191 |
| TMEM117 | protein_coding | -1.2364068 | 4.68668506 |
| ANO4 | protein_coding | -1.2412279 | 7.28637114 |
| NPTX1 | protein_coding | -1.2412973 | 7.26956708 |

| | | | |
|---|---|---|---|
| NATD1 | protein_coding | -1.2428499 | 6.06244895 |
| FUT8-AS1 | lncRNA | -1.2466039 | 2.33876548 |
| RP11-867G2: | lncRNA | -1.2499118 | 6.09043772 |
| OLFM2 | protein_coding | -1.2500101 | 6.57249858 |
| AP002528.1 | lncRNA | -1.2507537 | 4.36922683 |
| SRC | protein_coding | -1.2511935 | 7.19093993 |
| ZNF488 | protein_coding | -1.2530154 | 9.72310864 |
| CHSY1 | protein_coding | -1.2538826 | 11.1374451 |
| KB-1410C5.6 | lncRNA | -1.2542443 | 6.11253342 |
| NELL1 | protein_coding | -1.255977 | 13.6351063 |
| C1orf127 | protein_coding | -1.2568912 | 2.25467665 |
| MEIOC | protein_coding | -1.2579458 | 12.0704839 |
| TSPAN10 | protein_coding | -1.258288 | 3.65750363 |
| RP11-206F17 | lncRNA | -1.2584797 | 5.44144519 |
| LINC02844 | lncRNA | -1.2587317 | 5.45730837 |
| C16orf74 | protein_coding | -1.2604245 | 6.66496723 |
| FAT4 | protein_coding | -1.2618662 | 13.4563253 |
| ZNF618 | protein_coding | -1.2627467 | 6.47946304 |
| PHLDA2 | protein_coding | -1.2667434 | 9.56213164 |
| ACOXL-AS1 | lncRNA | -1.2668027 | 3.35664808 |
| SNTG2 | protein_coding | -1.2715459 | 7.72641258 |
| ISM2 | protein_coding | -1.2716773 | 8.95527378 |
| DSG2 | protein_coding | -1.2742373 | 5.13849862 |
| PPP1R1B | protein_coding | -1.2751925 | 5.12348745 |
| GRK5 | protein_coding | -1.2753034 | 14.8497666 |
| ABLIM1 | protein_coding | -1.2762817 | 8.31730088 |
| PNMA8A | protein_coding | -1.278857 | 3.6066797 |
| LYNX1 | protein_coding | -1.2789819 | 7.82170288 |
| CTD-3001H1: | lncRNA | -1.2797529 | 2.52179015 |
| STMND1 | protein_coding | -1.2813235 | 3.08823826 |
| SCD5 | protein_coding | -1.2860238 | 14.7938914 |
| RIMS1 | protein_coding | -1.2874432 | 15.3992332 |
| LRRN2 | protein_coding | -1.2902724 | 3.77972132 |
| CNRIP1 | protein_coding | -1.2903518 | 2.58979915 |
| CELF5 | protein_coding | -1.290822 | 10.1537653 |
| BNIP3 | protein_coding | -1.2922552 | 16.9841057 |
| CDH15 | protein_coding | -1.2972996 | 4.98919023 |
| AIF1L | protein_coding | -1.2988732 | 13.2493222 |
| LINC01625 | lncRNA | -1.3012848 | 3.10491674 |
| DIXDC1 | protein_coding | -1.3030963 | 5.99024919 |
| GPM6B | protein_coding | -1.3065599 | 11.5463698 |
| CTD-3094K11 | lncRNA | -1.307681 | 3.81515537 |
| PTGDR2 | protein_coding | -1.3077967 | 4.05981333 |
| MAB21L4 | protein_coding | -1.3091538 | 8.27863011 |
| SLC44A3 | protein_coding | -1.3097286 | 7.21191882 |

| | | | |
|---|---|---|---|
| KRT20 | protein_coding | -1.3147297 | 18.4248677 |
| PPP1R16B | protein_coding | -1.3163017 | 9.80837927 |
| PRSS3 | protein_coding | -1.3164982 | 2.28264445 |
| SHF | protein_coding | -1.3177783 | 10.5564436 |
| NYNRIN | protein_coding | -1.3188328 | 13.068558 |
| SIX1 | protein_coding | -1.3197968 | 6.39008105 |
| MMP15 | protein_coding | -1.3250697 | 7.18600045 |
| HOXC10 | protein_coding | -1.3260225 | 8.85197583 |
| SNCB | protein_coding | -1.3295123 | 10.9353018 |
| SULF1 | protein_coding | -1.3308954 | 11.3442534 |
| RIPPLY3 | protein_coding | -1.334242 | 11.2324701 |
| FNDC10 | protein_coding | -1.3345673 | 10.3096335 |
| RILPL1 | protein_coding | -1.3357143 | 5.95994903 |
| B3GAT1 | protein_coding | -1.3364159 | 7.27470212 |
| OLFML2A | protein_coding | -1.3379302 | 6.30822655 |
| SHISA7 | protein_coding | -1.3413803 | 5.07685602 |
| LINC00870 | lncRNA | -1.3416943 | 7.15006304 |
| RP11-2E11.9 | lncRNA | -1.3418994 | 3.1300559 |
| CCNO | protein_coding | -1.3422786 | 5.22694341 |
| FAM153A | protein_coding | -1.3424451 | 4.60071966 |
| RP11-380N8. | lncRNA | -1.3426651 | 2.38208107 |
| LMNTD2-AS1 | lncRNA | -1.3486845 | 6.95703386 |
| DBH | protein_coding | -1.3508898 | 8.75840639 |
| SCN5A | protein_coding | -1.3519629 | 8.04765993 |
| PLCH2 | protein_coding | -1.3533334 | 6.92030396 |
| CNTFR | protein_coding | -1.3557216 | 9.67681081 |
| LDHA | protein_coding | -1.358738 | 19.9771973 |
| BACE2 | protein_coding | -1.3617438 | 6.73093387 |
| BTNL9 | protein_coding | -1.3631999 | 4.83424444 |
| KRT12 | protein_coding | -1.3650453 | 2.68684199 |
| LAMA5 | protein_coding | -1.3713039 | 10.9272699 |
| RP11-677M1 | lncRNA | -1.3714848 | 3.42961696 |
| RP11-139E24 | lncRNA | -1.3721989 | 3.53864412 |
| NRN1 | protein_coding | -1.3727157 | 10.8055749 |
| TCP11L2 | protein_coding | -1.3763301 | 4.65342978 |
| RP4-564M11 | lncRNA | -1.3779788 | 4.21240277 |
| GSTT2B | protein_coding | -1.3802529 | 3.67550073 |
| TSHZ2 | protein_coding | -1.3817181 | 4.89645116 |
| NRIP3 | protein_coding | -1.3822977 | 13.4189156 |
| RP1-102E24. | lncRNA | -1.383551 | 2.99224154 |
| OPRD1 | protein_coding | -1.3874391 | 6.58936119 |
| SERPINH1 | protein_coding | -1.3875457 | 7.9696487 |
| RAMP2 | protein_coding | -1.3908742 | 3.10861607 |
| TBKBP1 | protein_coding | -1.3937187 | 6.8866687 |
| PTPRD-AS1 | lncRNA | -1.3946261 | 3.65097801 |

| | | | |
|---|---|---|---|
| B3GAT2 | protein_coding | -1.3962179 | 10.4125221 |
| PHETA1 | protein_coding | -1.4034072 | 4.54375794 |
| HAP1 | protein_coding | -1.4046466 | 3.93400715 |
| LINC01016 | lncRNA | -1.4073733 | 2.48776409 |
| CLSTN2 | protein_coding | -1.4078211 | 13.1700501 |
| TLCD4 | protein_coding | -1.4122707 | 8.6166191 |
| KCNJ6 | protein_coding | -1.4123766 | 12.10707 |
| RP11-480D4. | lncRNA | -1.4148608 | 4.22215369 |
| EDN3 | protein_coding | -1.4166192 | 2.88929 |
| FCHO2 | protein_coding | -1.4188447 | 7.9686045 |
| LINC01018 | lncRNA | -1.4197955 | 7.58847205 |
| RP11-416O1 | lncRNA | -1.4236723 | 5.10192386 |
| SLC30A3 | protein_coding | -1.4236813 | 5.0485782 |
| MFAP3L | protein_coding | -1.4237776 | 3.1891727 |
| ELN | protein_coding | -1.4256081 | 11.0428411 |
| LGR4 | protein_coding | -1.4271622 | 9.66049889 |
| CCDC178 | protein_coding | -1.4280927 | 13.9998906 |
| MCUB | protein_coding | -1.4284481 | 8.74043934 |
| ARSG | protein_coding | -1.4292652 | 4.96388547 |
| OCLN | protein_coding | -1.4304703 | 4.88946363 |
| TMEM125 | protein_coding | -1.4336106 | 3.74218696 |
| KISS1R | protein_coding | -1.4354863 | 6.36435219 |
| KCNB1 | protein_coding | -1.443373 | 5.84416861 |
| CDH4 | protein_coding | -1.4471023 | 8.94293291 |
| PLIN2 | protein_coding | -1.4472696 | 11.2397507 |
| SHISA9 | protein_coding | -1.4473734 | 7.38448717 |
| TBX20 | protein_coding | -1.4524985 | 3.27429059 |
| CDKN1C | protein_coding | -1.4594933 | 13.0422591 |
| RP11-535M1 | lncRNA | -1.4601213 | 6.19575285 |
| FOLR1 | protein_coding | -1.4610152 | 2.07221228 |
| RP11-557N2 | lncRNA | -1.4612953 | 2.4786484 |
| MUC5B | protein_coding | -1.4623907 | 2.77727486 |
| MAN1C1 | protein_coding | -1.4639112 | 8.17692374 |
| NEUROD1 | protein_coding | -1.4656772 | 7.38578346 |
| JAKMIP2 | protein_coding | -1.4690428 | 5.35346199 |
| HMX1 | protein_coding | -1.4702508 | 4.06969949 |
| ASGR1 | protein_coding | -1.4706814 | 3.49763264 |
| RGS4 | protein_coding | -1.4713407 | 7.57016297 |
| FLI1 | protein_coding | -1.4722337 | 10.4440981 |
| LANCL3 | protein_coding | -1.4742082 | 4.62847015 |
| PLCD4 | protein_coding | -1.4748049 | 8.30888474 |
| FGD1 | protein_coding | -1.4767872 | 9.4274698 |
| IGFBP5 | protein_coding | -1.4834746 | 19.9957579 |
| ATXN8OS | lncRNA | -1.4851197 | 6.51745508 |
| RP11-21C4.1 | lncRNA | -1.4879669 | 4.17872889 |

| | | | |
|---|---|---|---|
| DNAH6 | protein_coding | -1.4889095 | 4.09584523 |
| ST6GALNAC5 | protein_coding | -1.4901532 | 15.7761143 |
| AC012442.5 | lncRNA | -1.4911584 | 2.63806129 |
| LINC00051 | lncRNA | -1.4925084 | 2.23625667 |
| RP5-983L19.: | lncRNA | -1.4933403 | 2.56520351 |
| RP11-1149O: | lncRNA | -1.4943403 | 5.68266668 |
| TRIM7 | protein_coding | -1.4969644 | 4.38220811 |
| MMP17 | protein_coding | -1.5001353 | 8.74683246 |
| USH1C | protein_coding | -1.504879 | 5.85581533 |
| GPR157 | protein_coding | -1.5078367 | 2.65271403 |
| EHHADH | protein_coding | -1.5087878 | 9.36388962 |
| MYO5B | protein_coding | -1.5096629 | 14.9705734 |
| GPR12 | protein_coding | -1.5107068 | 14.0699037 |
| RNF39 | protein_coding | -1.5111481 | 5.72178094 |
| GALNT1 | protein_coding | -1.5125392 | 12.3532039 |
| GRAMD2A | protein_coding | -1.5146776 | 2.73662729 |
| ABC10-4437: | lncRNA | -1.5151702 | 3.30848875 |
| PLPPR1 | protein_coding | -1.5163554 | 9.36052254 |
| THBD | protein_coding | -1.5171357 | 5.05639884 |
| ELL2 | protein_coding | -1.5188836 | 13.2135163 |
| SHROOM3 | protein_coding | -1.5235521 | 8.95983909 |
| RAB6B | protein_coding | -1.5265127 | 8.29844325 |
| RP11-478J18 | lncRNA | -1.5278684 | 3.64640865 |
| RP11-433J8.: | lncRNA | -1.5283868 | 5.39999353 |
| NXPH4 | protein_coding | -1.5284478 | 15.8242469 |
| TMCO4 | protein_coding | -1.52916 | 3.45125016 |
| ATP10B | protein_coding | -1.5305611 | 2.43270591 |
| ANKRD35 | protein_coding | -1.5323451 | 5.60051358 |
| RP11-227D2. | lncRNA | -1.5390387 | 2.51725939 |
| KDM7A-DT | lncRNA | -1.5390746 | 5.12779322 |
| RP11-507K2. | lncRNA | -1.5427272 | 4.51483343 |
| ANKRD33 | protein_coding | -1.5529209 | 2.7564305 |
| REM2 | protein_coding | -1.5585644 | 5.55136941 |
| OBSL1 | protein_coding | -1.5620085 | 6.56364288 |
| RGCC | protein_coding | -1.5629029 | 5.77164002 |
| SLC22A15 | protein_coding | -1.5636184 | 4.38207633 |
| MAGI3 | protein_coding | -1.5658357 | 18.1229914 |
| LINC00641 | lncRNA | -1.5662083 | 13.9116389 |
| MIR210HG | lncRNA | -1.5744342 | 10.6581742 |
| KLHL1 | protein_coding | -1.5745051 | 12.2709678 |
| EFCAB6 | protein_coding | -1.5784075 | 3.17655759 |
| GRIP2 | protein_coding | -1.5813912 | 5.87689983 |
| MYO1G | protein_coding | -1.5819033 | 3.39238121 |
| MYBPHL | protein_coding | -1.5824313 | 2.94894166 |
| LCK | protein_coding | -1.5831989 | 4.928351 |

| | | | |
|---|---|---|---|
| RP11-23J9.8 | lncRNA | -1.5851331 | 8.73739307 |
| PDE11A | protein_coding | -1.5861272 | 3.0583448 |
| ZNF599 | protein_coding | -1.5862692 | 5.2003013 |
| LINC00467 | lncRNA | -1.5924237 | 5.05966043 |
| CROCC2 | protein_coding | -1.5987281 | 2.63364791 |
| NID2 | protein_coding | -1.601663 | 6.2242851 |
| ZNF667-AS1 | lncRNA | -1.6068333 | 11.5756029 |
| SOWAHC | protein_coding | -1.6102706 | 13.4419619 |
| FSCN2 | protein_coding | -1.6171546 | 3.57401856 |
| KL | protein_coding | -1.6173565 | 3.68429992 |
| DGCR5 | lncRNA | -1.6210143 | 11.6538055 |
| ETS1 | protein_coding | -1.6294803 | 14.1554696 |
| CTD-2223O1 | lncRNA | -1.630568 | 8.48121454 |
| FEV | protein_coding | -1.636769 | 4.87017118 |
| CCN2 | protein_coding | -1.6401941 | 3.76152992 |
| DHRS7 | protein_coding | -1.6446076 | 13.3795874 |
| CYYR1 | protein_coding | -1.6549361 | 11.3105407 |
| SMCO4 | protein_coding | -1.6553155 | 7.01902394 |
| SP9 | protein_coding | -1.655523 | 5.88783289 |
| RP11-1108I1 | lncRNA | -1.6599381 | 8.62930558 |
| CD44 | protein_coding | -1.6627623 | 20.5662475 |
| LRRK1 | protein_coding | -1.6629388 | 4.8214738 |
| RASD1 | protein_coding | -1.6643213 | 5.41838728 |
| SLC8A3 | protein_coding | -1.6646422 | 10.7142714 |
| C6orf132 | protein_coding | -1.6669978 | 5.53951521 |
| PLCB2 | protein_coding | -1.6693345 | 4.04700793 |
| TRHDE | protein_coding | -1.676067 | 3.76149332 |
| BARX1 | protein_coding | -1.6774632 | 4.91123031 |
| SP8 | protein_coding | -1.6791946 | 7.55824397 |
| GATM | protein_coding | -1.6793588 | 10.3732251 |
| HAND2 | protein_coding | -1.6856886 | 8.51808491 |
| LINC02600 | lncRNA | -1.686087 | 16.0300643 |
| RLBP1 | protein_coding | -1.6877548 | 3.20823665 |
| RP11-54O7.1 | lncRNA | -1.6884679 | 3.60015259 |
| MOB1B | protein_coding | -1.6887183 | 8.19409103 |
| SHANK1 | protein_coding | -1.6887511 | 4.75986237 |
| FXYD6 | protein_coding | -1.6902343 | 8.004573 |
| CGREF1 | protein_coding | -1.6904166 | 6.88079111 |
| FERMT2 | protein_coding | -1.6989595 | 13.2177376 |
| NEUROG2 | protein_coding | -1.700583 | 4.34581651 |
| HUNK | protein_coding | -1.7106777 | 11.5817164 |
| KCNK5 | protein_coding | -1.7117044 | 5.67889771 |
| ARHGAP28 | protein_coding | -1.7182482 | 4.49828854 |
| PPP1R3C | protein_coding | -1.7200153 | 10.7293077 |
| NBL1 | protein_coding | -1.7262399 | 11.9792857 |

| | | | |
|---|---|---|---|
| PSD2 | protein_coding | -1.7330234 | 4.197412 |
| RP11-753H1( | lncRNA | -1.7382427 | 2.91465196 |
| SASH1 | protein_coding | -1.7420453 | 8.10382398 |
| GSTO1 | protein_coding | -1.7429669 | 6.40656266 |
| BOK | protein_coding | -1.7429788 | 6.55591672 |
| MCIDAS | protein_coding | -1.7430697 | 5.18805347 |
| PDE5A | protein_coding | -1.7440987 | 4.59332129 |
| NXPH2 | protein_coding | -1.759301 | 4.30420094 |
| RP11-753H1( | lncRNA | -1.7600415 | 3.81286828 |
| SLC10A4 | protein_coding | -1.7608279 | 4.25047277 |
| LINC01829 | lncRNA | -1.7609133 | 6.30569528 |
| SYK | protein_coding | -1.7632899 | 12.7992185 |
| ANTXR1 | protein_coding | -1.7645703 | 6.28529122 |
| ATOH7 | protein_coding | -1.767899 | 8.74409656 |
| CRB1 | protein_coding | -1.772998 | 5.95219339 |
| FUT3 | protein_coding | -1.7730898 | 6.2271516 |
| PCCA-DT | lncRNA | -1.7733845 | 4.92200985 |
| GPR160 | protein_coding | -1.7841129 | 10.1202387 |
| MAPK10 | protein_coding | -1.7866093 | 6.06072766 |
| TRHDE-AS1 | lncRNA | -1.7888641 | 4.88606318 |
| JPH3 | protein_coding | -1.7963092 | 11.4528433 |
| LBH | protein_coding | -1.7972129 | 15.6047883 |
| FGFR3 | protein_coding | -1.7982549 | 7.6704131 |
| ATP4A | protein_coding | -1.8005184 | 7.15274043 |
| TMC3-AS1 | lncRNA | -1.8054152 | 2.27739108 |
| STON1 | protein_coding | -1.8061731 | 10.1995624 |
| EPHA6 | protein_coding | -1.8063555 | 12.339733 |
| PDZD4 | protein_coding | -1.8065441 | 11.0421138 |
| C4orf47 | protein_coding | -1.8108302 | 4.60286981 |
| DZIP1 | protein_coding | -1.8130713 | 8.17309855 |
| COPG2IT1 | lncRNA | -1.8143574 | 12.93853 |
| RP3-395M20 | lncRNA | -1.8197521 | 3.34267768 |
| CNN2 | protein_coding | -1.8238619 | 6.36452708 |
| STC1 | protein_coding | -1.827708 | 17.4214094 |
| AATK | protein_coding | -1.8301396 | 9.50749569 |
| RCAN3 | protein_coding | -1.8315366 | 12.6235806 |
| PIP5K1B | protein_coding | -1.8326869 | 8.34898574 |
| CDH20 | protein_coding | -1.8373984 | 12.205101 |
| PCBP4 | protein_coding | -1.8395806 | 7.66028508 |
| ABCG1 | protein_coding | -1.8422428 | 12.0551594 |
| LINC02743 | lncRNA | -1.8437285 | 4.49059699 |
| WIF1 | protein_coding | -1.8555298 | 12.1129489 |
| LINC02180 | lncRNA | -1.8607076 | 2.40117776 |
| NOS2 | protein_coding | -1.8652792 | 5.4631753 |
| PIANP | protein_coding | -1.8662645 | 3.3226326 |

| | | | |
|---|---|---|---|
| UTRN | protein_coding | -1.8820095 | 6.74271898 |
| CACNA2D4 | protein_coding | -1.8849739 | 8.96286979 |
| COL28A1 | protein_coding | -1.889554 | 12.0093369 |
| SCN1A | protein_coding | -1.8909033 | 6.78275237 |
| TPTE2 | protein_coding | -1.8970736 | 3.73200061 |
| PDE1B | protein_coding | -1.8974075 | 4.0000179 |
| AK5 | protein_coding | -1.8986718 | 13.2058517 |
| ZNF667 | protein_coding | -1.9018597 | 11.272741 |
| TNFRSF19 | protein_coding | -1.9025998 | 14.8171203 |
| AC011288.2 | lncRNA | -1.9123109 | 5.05517875 |
| AC007405.6 | lncRNA | -1.9192774 | 3.91103404 |
| SLC30A4 | protein_coding | -1.9221363 | 8.75375544 |
| RBMS1 | protein_coding | -1.924178 | 11.7582856 |
| RP1-237I15.2 | lncRNA | -1.9253432 | 3.83483097 |
| HSD17B13 | protein_coding | -1.9333673 | 10.8912759 |
| PLEKHH2 | protein_coding | -1.9344317 | 10.0084021 |
| S1PR5 | protein_coding | -1.935356 | 5.53742543 |
| RP13-766D2( | lncRNA | -1.94572 | 4.06539218 |
| PCDH12 | protein_coding | -1.9494606 | 8.5238629 |
| TSPAN19 | protein_coding | -1.9501239 | 3.7854454 |
| PXDC1 | protein_coding | -1.9544792 | 3.81455152 |
| TMSB15A | protein_coding | -1.9640046 | 10.053769 |
| METRN | protein_coding | -1.9647501 | 8.43950932 |
| KIRREL3 | protein_coding | -1.9715091 | 3.10249641 |
| ADGRL2 | protein_coding | -1.9764604 | 6.50242191 |
| VWA5B1 | protein_coding | -1.9839239 | 18.0358514 |
| SHISAL2B | protein_coding | -1.9843581 | 16.2209974 |
| HK2 | protein_coding | -1.9869471 | 17.9870357 |
| LRRC4C | protein_coding | -1.9874802 | 5.61146311 |
| LINC00958 | lncRNA | -1.9940586 | 5.18939479 |
| RP11-343J3.2 | lncRNA | -1.9972743 | 3.10369798 |
| RNF152 | protein_coding | -2.0001121 | 6.47937616 |
| RP11-675F6. | lncRNA | -2.0084881 | 2.62697663 |
| HRH2 | protein_coding | -2.011842 | 4.76522275 |
| PPP1R3B | protein_coding | -2.0180826 | 3.29303009 |
| RP11-323C15 | lncRNA | -2.0230729 | 4.61862604 |
| GPR162 | protein_coding | -2.0340102 | 9.09234133 |
| PDGFB | protein_coding | -2.0462758 | 11.8130868 |
| LRRC53 | protein_coding | -2.0508512 | 5.66513734 |
| NCKAP1L | protein_coding | -2.0627844 | 8.11998611 |
| FARP1 | protein_coding | -2.0726991 | 11.1701485 |
| MAB21L1 | protein_coding | -2.0769085 | 13.2225075 |
| RP11-157P1. | lncRNA | -2.0774714 | 3.7218721 |
| RIMBP3 | protein_coding | -2.0845368 | 3.50580796 |
| FGFR2 | protein_coding | -2.0872935 | 5.09595262 |

| | | | |
|---|---|---|---|
| LINC02694 | lncRNA | -2.103282 | 11.0736491 |
| GRIK3 | protein_coding | -2.1091439 | 14.6629606 |
| DSP | protein_coding | -2.1105418 | 16.9804911 |
| LINC02299 | lncRNA | -2.1170632 | 4.68057205 |
| AP1M2 | protein_coding | -2.1194897 | 8.14294816 |
| SYNDIG1L | protein_coding | -2.1230972 | 15.1414782 |
| ITGA6 | protein_coding | -2.127892 | 8.07029966 |
| SOX5 | protein_coding | -2.1286771 | 12.1912623 |
| HAND1 | protein_coding | -2.1347189 | 3.60366336 |
| HGF | protein_coding | -2.1365832 | 14.293452 |
| HIF1A-AS3 | lncRNA | -2.1430835 | 3.68220952 |
| NDUFA4L2 | protein_coding | -2.1446168 | 17.2522154 |
| MED12L | protein_coding | -2.15404 | 6.36481923 |
| PPM1H | protein_coding | -2.1544967 | 13.2157249 |
| RP11-62C7.2 | lncRNA | -2.1557378 | 3.42299076 |
| MMP24 | protein_coding | -2.1585031 | 8.39250956 |
| CD9 | protein_coding | -2.1607017 | 5.83385812 |
| OPRL1 | protein_coding | -2.17517 | 5.86383307 |
| SLC16A11 | protein_coding | -2.1777008 | 4.0422392 |
| HSD17B11 | protein_coding | -2.1802387 | 3.13352024 |
| ADARB2-AS1 | lncRNA | -2.2038421 | 2.90624742 |
| RP11-737O2[illegible] | lncRNA | -2.2117189 | 4.39313305 |
| CUX2 | protein_coding | -2.2141502 | 5.83430084 |
| DCBLD1 | protein_coding | -2.2145184 | 5.6558955 |
| BLK | protein_coding | -2.225975 | 6.96654202 |
| RP11-21L23.[illegible] | lncRNA | -2.2352298 | 9.38670241 |
| HTRA1 | protein_coding | -2.2406835 | 3.77034035 |
| TSKU | protein_coding | -2.2549786 | 14.4340773 |
| MCTP1 | protein_coding | -2.2816627 | 9.08948098 |
| ZIC2 | protein_coding | -2.2894476 | 9.0908652 |
| LINC01234 | lncRNA | -2.2910594 | 5.72900538 |
| TMEM45A | protein_coding | -2.3006525 | 10.1194921 |
| CBFA2T3 | protein_coding | -2.3051382 | 11.1727684 |
| KCNH5 | protein_coding | -2.3197463 | 9.40472133 |
| SNCA | protein_coding | -2.3222556 | 8.12511689 |
| ABCC8 | protein_coding | -2.334984 | 5.46961589 |
| RP11-1263C[illegible] | lncRNA | -2.33842 | 8.00272868 |
| SNX31 | protein_coding | -2.349788 | 5.3732471 |
| HOXC12 | protein_coding | -2.3511255 | 6.77916865 |
| TCEA3 | protein_coding | -2.356872 | 16.6724344 |
| PCP4L1 | protein_coding | -2.3661987 | 5.04436313 |
| ACTL8 | protein_coding | -2.3743396 | 17.9580415 |
| ANOS1 | protein_coding | -2.3827652 | 10.5503693 |
| FOXA2 | protein_coding | -2.3880978 | 5.96847843 |
| SLC22A16 | protein_coding | -2.3896068 | 3.48385311 |

| | | | |
|---|---|---|---|
| APLN | protein_coding | -2.3960411 | 9.68110895 |
| DCAF12L2 | protein_coding | -2.3969596 | 4.50749476 |
| SIRPA | protein_coding | -2.3973188 | 11.5256385 |
| MMP14 | protein_coding | -2.400432 | 16.742084 |
| SALL1 | protein_coding | -2.4044324 | 6.80721819 |
| RP11-430C7. | lncRNA | -2.4076927 | 4.70025823 |
| DOCK3 | protein_coding | -2.4104638 | 5.24799147 |
| FGFR1 | protein_coding | -2.4507213 | 10.1526831 |
| LAMA2 | protein_coding | -2.4518397 | 5.3579738 |
| KIAA1549L | protein_coding | -2.4578168 | 4.10790045 |
| CLEC4E | protein_coding | -2.4634359 | 6.90361261 |
| ARHGEF33 | protein_coding | -2.466338 | 14.1285946 |
| SPAG6 | protein_coding | -2.4861383 | 6.75693245 |
| SMIM18 | protein_coding | -2.4921348 | 3.85500454 |
| PELI2 | protein_coding | -2.496907 | 11.9987165 |
| LGALSL | protein_coding | -2.4987472 | 8.17035013 |
| IGFBP3 | protein_coding | -2.500105 | 9.23976723 |
| ST6GALNAC3 | protein_coding | -2.5088557 | 4.75127303 |
| FHL2 | protein_coding | -2.5358069 | 9.57445203 |
| TMEM271 | protein_coding | -2.547743 | 8.31551361 |
| EFS | protein_coding | -2.5687878 | 16.1065344 |
| PRR35 | protein_coding | -2.600051 | 4.85746751 |
| AC004490.1 | lncRNA | -2.614879 | 4.28428382 |
| EPHB1 | protein_coding | -2.6191235 | 12.4793804 |
| RP11-346M1 | lncRNA | -2.6344429 | 4.21702287 |
| ANKRD63 | protein_coding | -2.6381625 | 5.92208175 |
| TFAP2C | protein_coding | -2.64067 | 9.85059526 |
| CCDC39 | protein_coding | -2.6453633 | 8.85963224 |
| SHISA8 | protein_coding | -2.658353 | 3.77835998 |
| NEUROG3 | protein_coding | -2.661061 | 8.10799896 |
| RP4-607I7.1 | lncRNA | -2.6660483 | 8.85866113 |
| KCNK13 | protein_coding | -2.6987725 | 16.4557567 |
| FAM43B | protein_coding | -2.7021791 | 10.4419327 |
| RP11-245J24 | lncRNA | -2.7297436 | 6.59932812 |
| FST | protein_coding | -2.7510082 | 18.0585684 |
| NMBR | protein_coding | -2.7615451 | 5.00034482 |
| SYT17 | protein_coding | -2.7896399 | 7.21348653 |
| MXRA8 | protein_coding | -2.7912916 | 11.0000616 |
| ANKRD30A | protein_coding | -2.8451565 | 5.31493924 |
| GABRD | protein_coding | -2.8481082 | 11.1712494 |
| HEMGN | protein_coding | -2.8575142 | 5.93099855 |
| RP11-482M8 | lncRNA | -2.8843759 | 4.92312831 |
| SLC1A1 | protein_coding | -2.888864 | 6.05510061 |
| SOX2-OT | lncRNA | -2.8912205 | 16.2181283 |
| USH2A | protein_coding | -2.907346 | 9.13846266 |

| | | | |
|---|---|---|---|
| GRM5 | protein_coding | -2.9216307 | 7.93464739 |
| SMOC1 | protein_coding | -2.9388138 | 10.0670789 |
| CDH13 | protein_coding | -2.953193 | 5.57405677 |
| ZIC5 | protein_coding | -2.9585021 | 7.77299884 |
| TSPAN2 | protein_coding | -2.9974465 | 6.23750894 |
| DIAPH2 | protein_coding | -3.0032252 | 8.80488584 |
| AOC1 | protein_coding | -3.011217 | 4.05081792 |
| EYA2 | protein_coding | -3.046996 | 15.0615027 |
| BARHL1 | protein_coding | -3.0807622 | 14.5527332 |
| ADAMTS19 | protein_coding | -3.1209524 | 8.06802809 |
| SLC16A3 | protein_coding | -3.1228377 | 13.9881689 |
| AADAT | protein_coding | -3.1545788 | 11.0479368 |
| AC124789.1 | lncRNA | -3.1546437 | 11.4138245 |
| ACTG2 | protein_coding | -3.1571339 | 13.1518722 |
| RP4-647J21.1 | lncRNA | -3.1593962 | 5.16243606 |
| FLT4 | protein_coding | -3.1817186 | 10.7078355 |
| SLC38A11 | protein_coding | -3.2138954 | 12.8577739 |
| KCNK6 | protein_coding | -3.24405 | 10.8762415 |
| MEIOB | protein_coding | -3.2757603 | 10.854331 |
| MOCS1 | protein_coding | -3.292207 | 7.92843541 |
| EMILIN3 | protein_coding | -3.3023887 | 4.89416621 |
| GPRIN3 | protein_coding | -3.3388163 | 11.228158 |
| UTS2R | protein_coding | -3.4281022 | 4.65203544 |
| EPHB6 | protein_coding | -3.4442606 | 9.29197112 |
| EHF | protein_coding | -3.4580776 | 7.89396203 |
| RP5-1120P11 | lncRNA | -3.4738265 | 10.5434543 |
| CFAP77 | protein_coding | -3.4944836 | 9.81376492 |
| SLC38A3 | protein_coding | -3.5081562 | 5.67710756 |
| MAP2K6 | protein_coding | -3.5560016 | 9.91264266 |
| SORBS3 | protein_coding | -3.5805312 | 11.1191451 |
| TFCP2L1 | protein_coding | -3.6057723 | 10.7442366 |
| SYTL4 | protein_coding | -3.6377506 | 7.97082621 |
| HPCAL4 | protein_coding | -3.6598179 | 7.40101288 |
| TCEAL2 | protein_coding | -3.6870156 | 6.73844789 |
| TRIM71 | protein_coding | -3.8829699 | 4.29990835 |
| HTR6 | protein_coding | -3.9516882 | 6.78725414 |
| SLC4A4 | protein_coding | -4.0253121 | 16.9564516 |
| CYYR1-AS1 | lncRNA | -4.0618832 | 10.6507117 |
| LINC02031 | lncRNA | -4.1419204 | 9.12295476 |
| GSC2 | protein_coding | -4.1725335 | 7.32528946 |
| WLS | protein_coding | -4.2323933 | 11.2870105 |
| PRSS2 | protein_coding | -4.4532877 | 6.7892854 |
| CHRDL1 | protein_coding | -4.4560381 | 8.54879758 |
| NWD2 | protein_coding | -4.5568752 | 11.1109572 |
| AGR2 | protein_coding | -4.5601386 | 13.1893368 |

| | | | |
|---|---|---|---|
| LINC01754 | lncRNA | -4.6319814 | 7.02332046 |
| SYT6 | protein_coding | -4.8339069 | 12.1148567 |
| DIPK1C | protein_coding | -4.8966987 | 8.56512086 |
| CRLF1 | protein_coding | -4.9160997 | 10.6644598 |
| RBP7 | protein_coding | -4.9771164 | 9.2227236 |
| GPR139 | protein_coding | -5.0502587 | 5.71294593 |
| GJD2 | protein_coding | -5.2619555 | 8.7370324 |
| PKD1L3 | protein_coding | -5.6102059 | 6.467044 |
| SKAP1 | protein_coding | -5.6888091 | 7.10299004 |
| RP11-304L19 | lncRNA | -5.7710439 | 9.09998924 |
| RP11-304L19 | lncRNA | -5.9110247 | 8.94090763 |
| SPON1 | protein_coding | -6.5501749 | 5.0450934 |
| SLC52A3 | protein_coding | -6.9314644 | 5.82430212 |
| MOBP | protein_coding | -8.4783639 | 8.27823536 |
| ERVK9-11 | lncRNA | -8.6902265 | 8.95798747 |
| LINC02510 | lncRNA | -8.7869225 | 9.26294381 |
| MIR133A1HG | lncRNA | -9.2862242 | 10.2907813 |
