## Supplementary dataset 2 for "Polyomavirus ALTOs, but not MTs, downregulate viral early gene expression by activating the NF-κB pathway"

Table: Gene sets enriched in MKL-2 ALTO vs. RFP

| | GS<br>follow link to MSigDB | SIZE | ES | NES | NOM p-val | FDR q-val |
|---|---|---|---|---|---|---|
| 1 | REACTOME_INTERFERON_SIGNALING | 172 | 0.84 | 1.98 | 0 | 0 |
| 2 | BLANCO_MELO_COVID19_SARS_COV_2_INFECTIOI | 254 | 0.81 | 1.96 | 0 | 0 |
| 3 | BLANCO_MELO_HUMAN_PARAINFLUENZA_VIRUS_ | 169 | 0.83 | 1.94 | 0 | 0 |
| 4 | BLANCO_MELO_RESPIRATORY_SYNCYTIAL_VIRUS_ | 223 | 0.81 | 1.92 | 0 | 0 |
| 5 | SANA_RESPONSE_TO_IFNG_UP | 67 | 0.88 | 1.91 | 0 | 0 |
| 6 | BLANCO_MELO_COVID19_BRONCHIAL_EPITHELIAL | 100 | 0.84 | 1.9 | 0 | 0 |
| 7 | SEITZ_NEOPLASTIC_TRANSFORMATION_BY_8P_DE | 59 | 0.89 | 1.89 | 0 | 0 |
| 8 | MCLACHLAN_DENTAL_CARIES_UP | 151 | 0.8 | 1.89 | 0 | 0 |
| 9 | DER_IFN_ALPHA_RESPONSE_UP | 72 | 0.86 | 1.88 | 0 | 0 |
| 10 | SANA_TNF_SIGNALING_UP | 70 | 0.87 | 1.88 | 0 | 0 |
| 11 | KEGG_CYTOKINE_CYTOKINE_RECEPTOR_INTERACT | 120 | 0.81 | 1.88 | 0 | 0 |
| 12 | BROWNE_INTERFERON_RESPONSIVE_GENES | 63 | 0.87 | 1.88 | 0 | 0 |
| 13 | DER_IFN_BETA_RESPONSE_UP | 100 | 0.82 | 1.88 | 0 | 0 |
| 14 | WP_ALLOGRAFT_REJECTION | 51 | 0.88 | 1.87 | 0 | 0 |
| 15 | REACTOME_INTERFERON_ALPHA_BETA_SIGNALIN | 56 | 0.88 | 1.87 | 0 | 0 |
| 16 | GAURNIER_PSMD4_TARGETS | 46 | 0.9 | 1.86 | 0 | 0 |
| 17 | HELLER_SILENCED_BY_METHYLATION_UP | 216 | 0.78 | 1.86 | 0 | 0 |
| 18 | REACTOME_INTERFERON_GAMMA_SIGNALING | 76 | 0.84 | 1.85 | 0 | 0 |
| 19 | TAKEDA_TARGETS_OF_NUP98_HOXA9_FUSION_3I | 143 | 0.8 | 1.85 | 0 | 0 |
| 20 | XIE_ST_HSC_S1PR3_OE_UP | 130 | 0.81 | 1.85 | 0 | 0 |
| 21 | WIELAND_UP_BY_HBV_INFECTION | 78 | 0.83 | 1.85 | 0 | 0 |
| 22 | HECKER_IFNB1_TARGETS | 65 | 0.85 | 1.85 | 0 | 0 |
| 23 | ZHANG_RESPONSE_TO_IKK_INHIBITOR_AND_TNF_ | 185 | 0.78 | 1.84 | 0 | 0 |
| 24 | WP_OVERVIEW_OF_PROINFLAMMATORY_AND_PI | 40 | 0.9 | 1.84 | 0 | 0 |
| 25 | ICHIBA_GRAFT_VERSUS_HOST_DISEASE_D7_UP | 80 | 0.83 | 1.84 | 0 | 0 |
| 26 | SEKI_INFLAMMATORY_RESPONSE_LPS_UP | 60 | 0.84 | 1.84 | 0 | 0 |
| 27 | KEGG_SYSTEMIC_LUPUS_ERYTHEMATOSUS | 67 | 0.84 | 1.83 | 0 | 0 |
| 28 | BLANCO_MELO_BETA_INTERFERON_TREATED_BR | 297 | 0.76 | 1.83 | 0 | 0 |
| 29 | DER_IFN_GAMMA_RESPONSE_UP | 68 | 0.84 | 1.83 | 0 | 0 |
| 30 | KEGG_PORPHYRIN_AND_CHLOROPHYLL_METABOL | 29 | 0.93 | 1.83 | 0 | 0 |

| | | | | | | |
|---|---|---|---|---|---|---|
| 31 | BOSCO_TH1_CYTOTOXIC_MODULE | 55 | 0.85 | 1.82 | 0 | 0 |
| 32 | MOSERLE_IFNA_RESPONSE | 30 | 0.92 | 1.81 | 0 | 0 |
| 33 | WP_SARSCOV2_INNATE_IMMUNITY_EVASION_AN | 56 | 0.84 | 1.81 | 0 | 0 |
| 34 | BLANCO_MELO_BRONCHIAL_EPITHELIAL_CELLS_IN | 117 | 0.79 | 1.8 | 0 | 0 |
| 35 | KEGG_ANTIGEN_PROCESSING_AND_PRESENTATIC | 53 | 0.84 | 1.8 | 0 | 0 |
| 36 | WALLACE_PROSTATE_CANCER_RACE_UP | 184 | 0.76 | 1.8 | 0 | 0 |
| 37 | RADAEVA_RESPONSE_TO_IFNA1_UP | 45 | 0.87 | 1.8 | 0 | 0 |
| 38 | KEGG_STARCH_AND_SUCROSE_METABOLISM | 28 | 0.91 | 1.79 | 0 | 0 |
| 39 | DEBIASI_APOPTOSIS_BY_REOVIRUS_INFECTION_U | 282 | 0.74 | 1.79 | 0 | 0 |
| 40 | DAUER_STAT3_TARGETS_DN | 46 | 0.87 | 1.79 | 0 | 0 |
| 41 | BOSCO_INTERFERON_INDUCED_ANTIVIRAL_MODU | 65 | 0.83 | 1.78 | 0 | 0 |
| 42 | BLANCO_MELO_COVID19_SARS_COV_2_POS_PATI | 72 | 0.81 | 1.78 | 0 | 0 |
| 43 | ZHU_CMV_ALL_UP | 99 | 0.78 | 1.78 | 0 | 0 |
| 44 | WP_NETWORK_MAP_OF_SARSCOV2_SIGNALING_ | 141 | 0.77 | 1.77 | 0 | 0 |
| 45 | REACTOME_COSTIMULATION_BY_THE_CD28_FAM | 52 | 0.84 | 1.77 | 0 | 0 |
| 46 | BASSO_CD40_SIGNALING_UP | 90 | 0.78 | 1.77 | 0 | 0 |
| 47 | BERTUCCI_MEDULLARY_VS_DUCTAL_BREAST_CAN | 187 | 0.75 | 1.77 | 0 | 0 |
| 48 | REACTOME_ANTIVIRAL_MECHANISM_BY_IFN_STIM | 81 | 0.8 | 1.77 | 0 | 0 |
| 49 | JI_RESPONSE_TO_FSH_UP | 61 | 0.81 | 1.77 | 0 | 0 |
| 50 | KEGG_JAK_STAT_SIGNALING_PATHWAY | 88 | 0.78 | 1.76 | 0 | 0 |
| 51 | RODWELL_AGING_KIDNEY_UP | 358 | 0.73 | 1.76 | 0 | 0 |
| 52 | KIM_GLIS2_TARGETS_UP | 55 | 0.82 | 1.76 | 0 | 0 |
| 53 | REACTOME_TNFR2_NON_CANONICAL_NF_KB_PAT | 84 | 0.79 | 1.76 | 0 | 0 |
| 54 | KEGG_INTESTINAL_IMMUNE_NETWORK_FOR_IGA | 29 | 0.9 | 1.76 | 0 | 0 |
| 55 | WP_EBOLA_VIRUS_INFECTION_IN_HOST | 114 | 0.77 | 1.75 | 0 | 0 |
| 56 | BLANCO_MELO_COVID19_SARS_COV_2_LOW_MO | 136 | 0.75 | 1.75 | 0 | 0 |
| 57 | CROMER_TUMORIGENESIS_UP | 36 | 0.87 | 1.75 | 0 | 0 |
| 58 | KEGG_TYPE_I_DIABETES_MELLITUS | 27 | 0.89 | 1.75 | 0 | 0 |
| 59 | PHONG_TNF_RESPONSE_NOT_VIA_P38 | 309 | 0.72 | 1.75 | 0 | 0 |
| 60 | JISON_SICKLE_CELL_DISEASE_UP | 150 | 0.74 | 1.75 | 0 | 0 |
| 61 | HINATA_NFKB_TARGETS_KERATINOCYTE_UP | 68 | 0.8 | 1.75 | 0 | 0 |
| 62 | ZHU_CMV_8_HR_UP | 37 | 0.86 | 1.74 | 0 | 0 |
| 63 | LU_IL4_SIGNALING | 71 | 0.79 | 1.74 | 0 | 0 |

| | | | | | | |
|---|---|---|---|---|---|---|
| 64 | REACTOME_DDX58_IFIH1_MEDIATED_INDUCTION_ | 65 | 0.8 | 1.74 | 0 | 0 |
| 65 | KEGG_PENTOSE_AND_GLUCURONATE_INTERCON' | 18 | 0.94 | 1.74 | 0 | 0 |
| 66 | GRAHAM_CML_DIVIDING_VS_NORMAL_QUIESCEN | 68 | 0.78 | 1.74 | 0 | 0 |
| 67 | WP_SELENIUM_MICRONUTRIENT_NETWORK | 57 | 0.81 | 1.73 | 0 | 0 |
| 68 | JINESH_BLEBBISHIELD_VS_LIVE_CONTROL_DN | 239 | 0.73 | 1.73 | 0 | 0 |
| 69 | REACTOME_CHEMOKINE_RECEPTORS_BIND_CHEN | 23 | 0.92 | 1.73 | 0 | 0 |
| 70 | KEGG_RETINOL_METABOLISM | 26 | 0.88 | 1.73 | 0 | 0 |
| 71 | LINDSTEDT_DENDRITIC_CELL_MATURATION_A | 41 | 0.85 | 1.73 | 0 | 0 |
| 72 | WP_CYTOSOLIC_DNASENSING_PATHWAY | 52 | 0.82 | 1.73 | 0 | 0 |
| 73 | KEGG_DRUG_METABOLISM_OTHER_ENZYMES | 28 | 0.9 | 1.72 | 0 | 0 |
| 74 | KEGG_AUTOIMMUNE_THYROID_DISEASE | 21 | 0.91 | 1.72 | 0 | 0 |
| 75 | BLANCO_MELO_COVID19_SARS_COV_2_INFECTIOI | 449 | 0.71 | 1.72 | 0 | 0 |
| 76 | KEGG_CHEMOKINE_SIGNALING_PATHWAY | 130 | 0.74 | 1.72 | 0 | 0 |
| 77 | ALTEMEIER_RESPONSE_TO_LPS_WITH_MECHANIC | 73 | 0.78 | 1.72 | 0 | 0.001 |
| 78 | MEL18_DN.V1_UP | 110 | 0.75 | 1.72 | 0 | 0.001 |
| 79 | KUROZUMI_RESPONSE_TO_ONCOCYTIC_VIRUS | 18 | 0.93 | 1.72 | 0 | 0.001 |
| 80 | POOLA_INVASIVE_BREAST_CANCER_UP | 172 | 0.73 | 1.71 | 0 | 0.001 |
| 81 | STTTCRNTTT_IRF_Q6 | 158 | 0.73 | 1.71 | 0 | 0.001 |
| 82 | SENGUPTA_NASOPHARYNGEAL_CARCINOMA_UP | 271 | 0.71 | 1.71 | 0 | 0.001 |
| 83 | PHONG_TNF_TARGETS_UP | 55 | 0.8 | 1.71 | 0 | 0.001 |
| 84 | BENNETT_SYSTEMIC_LUPUS_ERYTHEMATOSUS | 23 | 0.89 | 1.71 | 0 | 0.001 |
| 85 | FULCHER_INFLAMMATORY_RESPONSE_LECTIN_VS | 350 | 0.7 | 1.71 | 0 | 0.001 |
| 86 | STAMBOLSKY_TARGETS_OF_MUTATED_TP53_DN | 45 | 0.81 | 1.71 | 0 | 0.001 |
| 87 | ZHANG_INTERFERON_RESPONSE | 23 | 0.9 | 1.71 | 0 | 0.001 |
| 88 | KIM_LRRC3B_TARGETS | 27 | 0.89 | 1.71 | 0 | 0.001 |
| 89 | KEGG_VIRAL_MYOCARDITIS | 52 | 0.8 | 1.7 | 0 | 0.001 |
| 90 | KRASNOSELSKAYA_ILF3_TARGETS_UP | 29 | 0.86 | 1.7 | 0 | 0.001 |
| 91 | KEGG_CYTOSOLIC_DNA_SENSING_PATHWAY | 35 | 0.85 | 1.7 | 0 | 0.001 |
| 92 | DOUGLAS_BMI1_TARGETS_DN | 288 | 0.7 | 1.7 | 0 | 0.001 |
| 93 | KEGG_GRAFT_VERSUS_HOST_DISEASE | 21 | 0.91 | 1.7 | 0 | 0.001 |
| 94 | HUMMERICH_SKIN_CANCER_PROGRESSION_UP | 71 | 0.78 | 1.7 | 0 | 0.001 |
| 95 | GRAESSMANN_RESPONSE_TO_MC_AND_SERUM_ | 162 | 0.72 | 1.7 | 0 | 0.001 |
| 96 | WP_VITAMIN_B12_METABOLISM | 30 | 0.85 | 1.69 | 0 | 0.001 |

| | | | | | | |
|---|---|---|---|---|---|---|
| 97 | DIRMEIER_LMP1_RESPONSE_EARLY | 50 | 0.81 | 1.69 | 0 | 0.001 |
| 98 | REACTOME_INTERLEUKIN_10_SIGNALING | 20 | 0.93 | 1.69 | 0 | 0.001 |
| 99 | WP_CHEMOKINE_SIGNALING_PATHWAY | 121 | 0.73 | 1.69 | 0 | 0.001 |
| 100 | SAKAI_CHRONIC_HEPATITIS_VS_LIVER_CANCER_U | 76 | 0.77 | 1.69 | 0 | 0.001 |
| 101 | KEGG_CELL_ADHESION_MOLECULES_CAMS | 92 | 0.75 | 1.69 | 0 | 0.001 |
| 102 | WP_GLUCURONIDATION | 16 | 0.97 | 1.69 | 0 | 0.001 |
| 103 | BMI1_DN_MEL18_DN.V1_UP | 110 | 0.74 | 1.69 | 0 | 0.001 |
| 104 | WUNDER_INFLAMMATORY_RESPONSE_AND_CHOI | 44 | 0.81 | 1.69 | 0 | 0.001 |
| 105 | MORI_LARGE_PRE_BII_LYMPHOCYTE_DN | 49 | 0.8 | 1.69 | 0 | 0.002 |
| 106 | JOHNSTONE_PARVB_TARGETS_2_UP | 130 | 0.72 | 1.68 | 0 | 0.002 |
| 107 | KEGG_LEISHMANIA_INFECTION | 51 | 0.79 | 1.68 | 0 | 0.002 |
| 108 | KEGG_STEROID_HORMONE_BIOSYNTHESIS | 23 | 0.9 | 1.68 | 0 | 0.002 |
| 109 | GHANDHI_DIRECT_IRRADIATION_UP | 68 | 0.78 | 1.68 | 0 | 0.002 |
| 110 | KEGG_ALLOGRAFT_REJECTION | 21 | 0.91 | 1.68 | 0 | 0.002 |
| 111 | YORDY_RECIPROCAL_REGULATION_BY_ETS1_AN | 55 | 0.78 | 1.68 | 0 | 0.002 |
| 112 | HOFFMANN_SMALL_PRE_BII_TO_IMMATURE_B_LY | 49 | 0.8 | 1.68 | 0 | 0.002 |
| 113 | REACTOME_MHC_CLASS_II_ANTIGEN_PRESENTA | 113 | 0.72 | 1.68 | 0 | 0.002 |
| 114 | WP_NONALCOHOLIC_FATTY_LIVER_DISEASE | 141 | 0.72 | 1.68 | 0 | 0.002 |
| 115 | PID_ATF2_PATHWAY | 50 | 0.79 | 1.68 | 0.001 | 0.002 |
| 116 | ZHU_CMV_24_HR_UP | 78 | 0.75 | 1.68 | 0 | 0.002 |
| 117 | EINAV_INTERFERON_SIGNATURE_IN_CANCER | 26 | 0.87 | 1.68 | 0 | 0.002 |
| 118 | LIU_VAV3_PROSTATE_CARCINOGENESIS_UP | 42 | 0.82 | 1.67 | 0 | 0.002 |
| 119 | TAKEDA_TARGETS_OF_NUP98_HOXA9_FUSION_1 | 131 | 0.72 | 1.67 | 0 | 0.002 |
| 120 | SARRIO_EPITHELIAL_MESENCHYMAL_TRANSITIOI | 111 | 0.73 | 1.67 | 0 | 0.002 |
| 121 | MIR1287_3P | 29 | 0.86 | 1.67 | 0 | 0.002 |
| 122 | SMID_BREAST_CANCER_NORMAL_LIKE_UP | 265 | 0.69 | 1.67 | 0 | 0.002 |
| 123 | REACTOME_PEPTIDE_LIGAND_BINDING_RECEPTC | 68 | 0.76 | 1.67 | 0 | 0.002 |
| 124 | WP_PHOTODYNAMIC_THERAPYINDUCED_NFKB_SI | 25 | 0.88 | 1.67 | 0 | 0.002 |
| 125 | NABA_SECRETED_FACTORS | 148 | 0.72 | 1.67 | 0 | 0.002 |
| 126 | KEGG_ASTHMA | 15 | 0.95 | 1.67 | 0 | 0.002 |
| 127 | TAKEDA_TARGETS_OF_NUP98_HOXA9_FUSION_8 | 101 | 0.73 | 1.67 | 0 | 0.002 |
| 128 | RODWELL_AGING_KIDNEY_NO_BLOOD_UP | 155 | 0.71 | 1.67 | 0 | 0.002 |
| 129 | BEIER_GLIOMA_STEM_CELL_DN | 62 | 0.78 | 1.67 | 0 | 0.002 |

| | | | | | | |
|---|---|---|---|---|---|---|
| 130 | WP_MEASLES_VIRUS_INFECTION | 104 | 0.73 | 1.67 | 0 | 0.002 |
| 131 | FONTAINE_THYROID_TUMOR_UNCERTAIN_MALIGN | 16 | 0.93 | 1.67 | 0 | 0.002 |
| 132 | MEK_UP.V1_DN | 162 | 0.71 | 1.67 | 0 | 0.002 |
| 133 | KRAS.DF.V1_UP | 156 | 0.7 | 1.67 | 0 | 0.002 |
| 134 | HORIUCHI_WTAP_TARGETS_UP | 253 | 0.69 | 1.66 | 0 | 0.003 |
| 135 | REACTOME_TCR_SIGNALING | 98 | 0.73 | 1.66 | 0 | 0.003 |
| 136 | KRIEG_KDM3A_TARGETS_NOT_HYPOXIA | 185 | 0.7 | 1.66 | 0 | 0.003 |
| 137 | GGGNNTTTCC_NFKB_Q6_01 | 113 | 0.72 | 1.66 | 0.001 | 0.003 |
| 138 | CHIN_BREAST_CANCER_COPY_NUMBER_UP | 23 | 0.88 | 1.66 | 0 | 0.003 |
| 139 | WP_TNFRELATED_WEAK_INDUCER_OF_APOPTOS | 40 | 0.81 | 1.66 | 0 | 0.003 |
| 140 | BURTON_ADIPOGENESIS_8 | 62 | 0.77 | 1.66 | 0.001 | 0.003 |
| 141 | ZHONG_SECRETOME_OF_LUNG_CANCER_AND_EI | 55 | 0.77 | 1.66 | 0 | 0.003 |
| 142 | CHANG_IMMORTALIZED_BY_HPV31_DN | 41 | 0.81 | 1.66 | 0 | 0.003 |
| 143 | ISRE_01 | 200 | 0.7 | 1.66 | 0 | 0.003 |
| 144 | HOSHIDA_LIVER_CANCER_SUBCLASS_S1 | 185 | 0.7 | 1.65 | 0 | 0.003 |
| 145 | ICHIBA_GRAFT_VERSUS_HOST_DISEASE_35D_UP | 87 | 0.73 | 1.65 | 0 | 0.003 |
| 146 | BOSCO_ALLERGEN_INDUCED_TH2_ASSOCIATED_ | 116 | 0.72 | 1.65 | 0 | 0.004 |
| 147 | PETROVA_PROX1_TARGETS_DN | 43 | 0.8 | 1.65 | 0 | 0.004 |
| 148 | DEMAGALHAES_AGING_UP | 36 | 0.82 | 1.65 | 0 | 0.004 |
| 149 | REACTOME_ARACHIDONIC_ACID_METABOLISM | 28 | 0.84 | 1.65 | 0 | 0.004 |
| 150 | NAKAYAMA_SOFT_TISSUE_TUMORS_PCA1_UP | 37 | 0.82 | 1.65 | 0 | 0.004 |
| 151 | YANG_BCL3_TARGETS_UP | 254 | 0.69 | 1.65 | 0 | 0.004 |
| 152 | MIR571 | 48 | 0.78 | 1.65 | 0 | 0.004 |
| 153 | LEE_LIVER_CANCER_MYC_E2F1_UP | 39 | 0.81 | 1.65 | 0.001 | 0.004 |
| 154 | SALVADOR_MARTIN_PEDIATRIC_TBD_ANTI_TNF_T | 19 | 0.89 | 1.65 | 0.001 | 0.004 |
| 155 | LENAOUR_DENDRITIC_CELL_MATURATION_UP | 83 | 0.74 | 1.65 | 0 | 0.004 |
| 156 | WP_PROSTAGLANDIN_SIGNALING | 18 | 0.92 | 1.65 | 0 | 0.004 |
| 157 | GRAHAM_CML_QUIESCENT_VS_NORMAL_DIVIDING | 37 | 0.81 | 1.64 | 0 | 0.004 |
| 158 | PICCALUGA_ANGIOIMMUNOBLASTIC_LYMPHOMA_ | 128 | 0.71 | 1.64 | 0 | 0.004 |
| 159 | TSUNODA_CISPLATIN_RESISTANCE_DN | 38 | 0.8 | 1.64 | 0 | 0.004 |
| 160 | LIANG_SILENCED_BY_METHYLATION_2 | 37 | 0.81 | 1.64 | 0 | 0.005 |
| 161 | WP_NONGENOMIC_ACTIONS_OF_125_DIHYDROXY | 60 | 0.77 | 1.64 | 0 | 0.005 |
| 162 | MARZEC_IL2_SIGNALING_UP | 93 | 0.73 | 1.64 | 0 | 0.005 |

| | | | | | | |
|---|---|---|---|---|---|---|
| 163 | WP_ARYL_HYDROCARBON_RECEPTOR_PATHWAY | 36 | 0.81 | 1.64 | 0 | 0.005 |
| 164 | KEGG_ASCORBATE_AND_ALDARATE_METABOLIS | 16 | 0.92 | 1.64 | 0 | 0.005 |
| 165 | WANG_ESOPHAGUS_CANCER_VS_NORMAL_UP | 96 | 0.72 | 1.64 | 0 | 0.005 |
| 166 | WP_OVERVIEW_OF_NANOPARTICLE_EFFECTS | 15 | 0.92 | 1.64 | 0 | 0.005 |
| 167 | RPS14_DN.V1_UP | 108 | 0.72 | 1.64 | 0 | 0.005 |
| 168 | LINDGREN_BLADDER_CANCER_CLUSTER_2B | 280 | 0.67 | 1.64 | 0 | 0.006 |
| 169 | PID_CD8_TCR_DOWNSTREAM_PATHWAY | 37 | 0.8 | 1.64 | 0 | 0.006 |
| 170 | WP_APOPTOSISRELATED_NETWORK_DUE_TO_AL | 47 | 0.78 | 1.64 | 0.001 | 0.006 |
| 171 | LIN_SILENCED_BY_TUMOR_MICROENVIRONMENT | 61 | 0.75 | 1.63 | 0 | 0.006 |
| 172 | LINDSTEDT_DENDRITIC_CELL_MATURATION_B | 44 | 0.79 | 1.63 | 0 | 0.006 |
| 173 | KEGG_LEUKOCYTE_TRANSENDOTHELIAL_MIGRAT | 90 | 0.72 | 1.63 | 0 | 0.006 |
| 174 | WP_P53_TRANSCRIPTIONAL_GENE_NETWORK | 63 | 0.75 | 1.63 | 0 | 0.006 |
| 175 | REACTOME_NEGATIVE_REGULATORS_OF_DDX58_ | 34 | 0.82 | 1.63 | 0.001 | 0.006 |
| 176 | TSAI_RESPONSE_TO_IONIZING_RADIATION | 126 | 0.7 | 1.63 | 0 | 0.006 |
| 177 | GRAESSMANN_APOPTOSIS_BY_SERUM_DEPRIVA | 449 | 0.67 | 1.63 | 0 | 0.006 |
| 178 | FOSTER_TOLERANT_MACROPHAGE_DN | 354 | 0.67 | 1.63 | 0 | 0.007 |
| 179 | WIERENGA_STAT5A_TARGETS_UP | 153 | 0.69 | 1.62 | 0 | 0.007 |
| 180 | REACTOME_ESTROGEN_DEPENDENT_GENE_EXP | 112 | 0.71 | 1.62 | 0 | 0.008 |
| 181 | RASHI_NFKB1_TARGETS | 18 | 0.89 | 1.62 | 0 | 0.008 |
| 182 | REACTOME_SENESCENCE_ASSOCIATED_SECRET | 78 | 0.74 | 1.62 | 0 | 0.007 |
| 183 | MARTIN_NFKB_TARGETS_UP | 28 | 0.84 | 1.62 | 0 | 0.008 |
| 184 | TURASHVILI_BREAST_LOBULAR_CARCINOMA_VS_ | 45 | 0.77 | 1.62 | 0 | 0.008 |
| 185 | ZWANG_CLASS_3_TRANSIENTLY_INDUCED_BY_E | 183 | 0.68 | 1.62 | 0 | 0.008 |
| 186 | PHONG_TNF_RESPONSE_VIA_P38_PARTIAL | 142 | 0.68 | 1.62 | 0 | 0.008 |
| 187 | WP_TCELL_ACTIVATION_SARSCOV2 | 64 | 0.74 | 1.62 | 0 | 0.008 |
| 188 | REACTOME_CONDENSATION_OF_PROPHASE_CHI | 42 | 0.77 | 1.62 | 0.003 | 0.008 |
| 189 | WP_SPINAL_CORD_INJURY | 79 | 0.73 | 1.62 | 0 | 0.008 |
| 190 | SHIN_B_CELL_LYMPHOMA_CLUSTER_8 | 29 | 0.82 | 1.62 | 0 | 0.009 |
| 191 | CENPT_TARGET_GENES | 59 | 0.75 | 1.62 | 0.001 | 0.009 |
| 192 | REACTOME_INTEGRIN_CELL_SURFACE_INTERACT | 49 | 0.77 | 1.62 | 0 | 0.009 |
| 193 | WEST_ADRENOCORTICAL_CARCINOMA_VS_ADEN | 15 | 0.91 | 1.62 | 0 | 0.009 |
| 194 | BMI1_DN.V1_UP | 109 | 0.71 | 1.62 | 0 | 0.009 |
| 195 | NOJIMA_SFRP2_TARGETS_DN | 22 | 0.86 | 1.62 | 0 | 0.009 |

| | | | | | | |
|---|---|---|---|---|---|---|
| 196 | WP_FOLATE_METABOLISM | 43 | 0.77 | 1.62 | 0 | 0.009 |
| 197 | HINATA_NFKB_TARGETS_FIBROBLAST_UP | 70 | 0.73 | 1.61 | 0 | 0.009 |
| 198 | WIERENGA_STAT5A_TARGETS_GROUP2 | 37 | 0.8 | 1.61 | 0.001 | 0.009 |
| 199 | JINESH_BLEBBISHIELD_TO_IMMUNE_CELL_FUSIO | 301 | 0.67 | 1.61 | 0 | 0.009 |
| 200 | DIRMEIER_LMP1_RESPONSE_LATE_UP | 50 | 0.77 | 1.61 | 0.001 | 0.009 |
| 201 | SMIRNOV_RESPONSE_TO_IR_2HR_UP | 49 | 0.77 | 1.61 | 0.001 | 0.009 |
| 202 | UROSEVIC_RESPONSE_TO_IMIQUIMOD | 17 | 0.9 | 1.61 | 0 | 0.009 |
| 203 | BROWNE_HCMV_INFECTION_8HR_UP | 88 | 0.72 | 1.61 | 0 | 0.009 |
| 204 | WP_HOSTPATHOGEN_INTERACTION_OF_HUMAN_( | 32 | 0.81 | 1.61 | 0.001 | 0.009 |
| 205 | CHIARADONNA_NEOPLASTIC_TRANSFORMATION_ | 93 | 0.72 | 1.61 | 0 | 0.009 |
| 206 | GEISS_RESPONSE_TO_DSRNA_UP | 34 | 0.81 | 1.61 | 0 | 0.009 |
| 207 | NABA_MATRISOME_ASSOCIATED | 345 | 0.66 | 1.61 | 0 | 0.01 |
| 208 | WP_EBSTEINBARR_VIRUS_LMP1_SIGNALING | 22 | 0.87 | 1.61 | 0 | 0.01 |
| 209 | REACTOME_INTERLEUKIN_4_AND_INTERLEUKIN_ | 70 | 0.73 | 1.61 | 0 | 0.01 |
| 210 | REACTOME_INFLAMMASOMES | 17 | 0.89 | 1.61 | 0 | 0.01 |
| 211 | GAZDA_DIAMOND_BLACKFAN_ANEMIA_ERYTHROI | 24 | 0.84 | 1.61 | 0 | 0.01 |
| 212 | MORI_MATURE_B_LYMPHOCYTE_UP | 81 | 0.73 | 1.61 | 0 | 0.01 |
| 213 | REACTOME_ERCC6_CSB_AND_EHMT2_G9A_POSI | 45 | 0.78 | 1.61 | 0 | 0.01 |
| 214 | WP_TYPE_II_INTERFERON_SIGNALING_IFNG | 32 | 0.81 | 1.61 | 0.001 | 0.011 |
| 215 | CHEN_HOXA5_TARGETS_9HR_UP | 212 | 0.67 | 1.61 | 0 | 0.011 |
| 216 | WP_MIRNAS_INVOLVEMENT_IN_THE_IMMUNE_RES | 28 | 0.82 | 1.61 | 0 | 0.011 |
| 217 | MIR6813_3P | 31 | 0.81 | 1.61 | 0 | 0.011 |
| 218 | BIOCARTA_ERK_PATHWAY | 25 | 0.84 | 1.61 | 0 | 0.011 |
| 219 | BIOCARTA_IL2RB_PATHWAY | 34 | 0.81 | 1.6 | 0 | 0.011 |
| 220 | ZHANG_ANTIVIRAL_RESPONSE_TO_RIBAVIRIN_DN | 32 | 0.82 | 1.6 | 0 | 0.011 |
| 221 | CUI_TCF21_TARGETS_UP | 27 | 0.84 | 1.6 | 0.004 | 0.011 |
| 222 | SCHUETZ_BREAST_CANCER_DUCTAL_INVASIVE_ | 227 | 0.67 | 1.6 | 0 | 0.011 |
| 223 | REACTOME_CYTOSOLIC_SENSORS_OF_PATHOGE | 60 | 0.75 | 1.6 | 0 | 0.011 |
| 224 | REACTOME_ASSEMBLY_OF_THE_ORC_COMPLEX_ | 38 | 0.79 | 1.6 | 0 | 0.012 |
| 225 | IRF_Q6 | 199 | 0.67 | 1.6 | 0 | 0.012 |
| 226 | GRAHAM_CML_QUIESCENT_VS_NORMAL_QUIESC | 30 | 0.81 | 1.6 | 0.001 | 0.012 |
| 227 | REACTOME_TNFS_BIND_THEIR_PHYSIOLOGICAL_ | 16 | 0.89 | 1.6 | 0 | 0.012 |
| 228 | LU_TUMOR_VASCULATURE_UP | 18 | 0.87 | 1.6 | 0 | 0.012 |

| | | | | | | |
|---|---|---|---|---|---|---|
| 229 | WP_IMMUNE_RESPONSE_TO_TUBERCULOSIS | 23 | 0.84 | 1.6 | 0 | 0.012 |
| 230 | MARKEY_RB1_ACUTE_LOF_DN | 164 | 0.68 | 1.6 | 0 | 0.013 |
| 231 | BECKER_TAMOXIFEN_RESISTANCE_UP | 41 | 0.78 | 1.6 | 0.004 | 0.013 |
| 232 | WP_ANTIVIRAL_AND_ANTIINFLAMMATORY_EFFEC | 26 | 0.81 | 1.6 | 0.003 | 0.013 |
| 233 | GRAHAM_NORMAL_QUIESCENT_VS_NORMAL_DIV | 42 | 0.78 | 1.6 | 0 | 0.013 |
| 234 | REACTOME_DNA_METHYLATION | 33 | 0.79 | 1.6 | 0.001 | 0.013 |
| 235 | RUTELLA_RESPONSE_TO_CSF2RB_AND_IL4_UP | 291 | 0.66 | 1.6 | 0 | 0.013 |
| 236 | GALE_APL_WITH_FLT3_MUTATED_DN | 16 | 0.88 | 1.6 | 0.002 | 0.013 |
| 237 | SYED_ESTRADIOL_RESPONSE | 17 | 0.88 | 1.6 | 0 | 0.013 |
| 238 | MARTINEZ_RESPONSE_TO_TRABECTEDIN_UP | 66 | 0.74 | 1.59 | 0.001 | 0.014 |
| 239 | REACTOME_TRANSPORT_OF_VITAMINS_NUCLEOS | 33 | 0.78 | 1.59 | 0.001 | 0.014 |
| 240 | MANNE_COVID19_NONICU_VS_HEALTHY_DONOR_ | 127 | 0.69 | 1.59 | 0 | 0.014 |
| 241 | BOWIE_RESPONSE_TO_EXTRACELLULAR_MATRIX | 17 | 0.88 | 1.59 | 0 | 0.014 |
| 242 | SMID_BREAST_CANCER_LUMINAL_B_DN | 335 | 0.65 | 1.59 | 0 | 0.014 |
| 243 | MIR6839_3P | 133 | 0.68 | 1.59 | 0 | 0.014 |
| 244 | REACTOME_TAK1_ACTIVATES_NFKB_BY_PHOSPH | 29 | 0.81 | 1.59 | 0 | 0.014 |
| 245 | REACTOME_PRC2_METHYLATES_HISTONES_AND_ | 42 | 0.78 | 1.59 | 0 | 0.014 |
| 246 | MUELLER_COMMON_TARGETS_OF_AML_FUSIONS | 28 | 0.82 | 1.59 | 0 | 0.014 |
| 247 | MARTIN_VIRAL_GPCR_SIGNALING_UP | 56 | 0.76 | 1.59 | 0 | 0.014 |
| 248 | KEGG_NOD_LIKE_RECEPTOR_SIGNALING_PATHW | 47 | 0.77 | 1.59 | 0.001 | 0.015 |
| 249 | BOWIE_RESPONSE_TO_TAMOXIFEN | 18 | 0.89 | 1.59 | 0 | 0.015 |
| 250 | LTE2_UP.V1_DN | 170 | 0.67 | 1.59 | 0 | 0.015 |
| 251 | OSWALD_HEMATOPOIETIC_STEM_CELL_IN_COLL/ | 177 | 0.66 | 1.59 | 0 | 0.015 |
| 252 | STEIN_ESRRA_TARGETS_RESPONSIVE_TO_ESTR | 25 | 0.82 | 1.59 | 0 | 0.015 |
| 253 | EIF4E_DN | 90 | 0.71 | 1.59 | 0 | 0.015 |
| 254 | XU_AKT1_TARGETS_6HR | 26 | 0.83 | 1.59 | 0.003 | 0.015 |
| 255 | REACTOME_HDACS_DEACETYLATE_HISTONES | 59 | 0.74 | 1.59 | 0 | 0.015 |
| 256 | NFKAPPAB_01 | 218 | 0.66 | 1.59 | 0 | 0.015 |
| 257 | REACTOME_BINDING_AND_UPTAKE_OF_LIGANDS_ | 19 | 0.86 | 1.59 | 0.003 | 0.015 |
| 258 | CHIBA_RESPONSE_TO_TSA_DN | 18 | 0.86 | 1.59 | 0.001 | 0.016 |
| 259 | BLANCO_MELO_COVID19_SARS_COV_2_INFECTIC | 400 | 0.65 | 1.59 | 0 | 0.016 |
| 260 | VALK_AML_CLUSTER_6 | 28 | 0.81 | 1.59 | 0.004 | 0.016 |
| 261 | ZHANG_ANTIVIRAL_RESPONSE_TO_RIBAVIRIN_UF | 22 | 0.84 | 1.59 | 0.003 | 0.016 |

| | | | | | | |
|---|---|---|---|---|---|---|
| 262 | SUNG_METASTASIS_STROMA_DN | 46 | 0.76 | 1.59 | 0.001 | 0.016 |
| 263 | GHANDHI_BYSTANDER_IRRADIATION_UP | 56 | 0.74 | 1.58 | 0 | 0.016 |
| 264 | BROCKE_APOPTOSIS_REVERSED_BY_IL6 | 132 | 0.68 | 1.58 | 0 | 0.016 |
| 265 | QI_PLASMACYTOMA_UP | 188 | 0.67 | 1.58 | 0 | 0.016 |
| 266 | HOXA9_DN.V1_UP | 151 | 0.67 | 1.58 | 0 | 0.016 |
| 267 | REACTOME_RECOGNITION_AND_ASSOCIATION_O | 32 | 0.79 | 1.58 | 0.001 | 0.017 |
| 268 | REACTOME_ANTIGEN_PRESENTATION_FOLDING_ | 25 | 0.83 | 1.58 | 0 | 0.017 |
| 269 | NABA_MATRISOME | 495 | 0.64 | 1.58 | 0 | 0.018 |
| 270 | MIR6736_3P | 106 | 0.69 | 1.58 | 0.001 | 0.018 |
| 271 | REACTOME_ACTIVATED_PKN1_STIMULATES_TRAI | 34 | 0.8 | 1.58 | 0 | 0.018 |
| 272 | REACTOME_IMMUNOREGULATORY_INTERACTIONS | 44 | 0.75 | 1.58 | 0.005 | 0.018 |
| 273 | WP_IL18_SIGNALING_PATHWAY | 210 | 0.67 | 1.58 | 0 | 0.018 |
| 274 | TURASHVILI_BREAST_LOBULAR_CARCINOMA_VS_ | 38 | 0.78 | 1.58 | 0.001 | 0.018 |
| 275 | WP_TOLLLIKE_RECEPTOR_SIGNALING_PATHWAY | 73 | 0.72 | 1.58 | 0 | 0.018 |
| 276 | KERLEY_RESPONSE_TO_CISPLATIN_UP | 34 | 0.79 | 1.58 | 0.001 | 0.018 |
| 277 | REACTOME_DNA_DAMAGE_TELOMERE_STRESS_I | 51 | 0.74 | 1.58 | 0.001 | 0.018 |
| 278 | REACTOME_CLASS_A_1_RHODOPSIN_LIKE_RECE | 116 | 0.68 | 1.58 | 0 | 0.018 |
| 279 | ZHAN_MULTIPLE_MYELOMA_LB_DN | 36 | 0.78 | 1.58 | 0 | 0.018 |
| 280 | WP_INTERACTIONS_BETWEEN_IMMUNE_CELLS_A | 18 | 0.88 | 1.58 | 0 | 0.018 |
| 281 | WIEDERSCHAIN_TARGETS_OF_BMI1_AND_PCGF2 | 42 | 0.76 | 1.58 | 0.001 | 0.019 |
| 282 | NUTT_GBM_VS_AO_GLIOMA_UP | 44 | 0.76 | 1.58 | 0.001 | 0.019 |
| 283 | SNF5_DN.V1_UP | 123 | 0.69 | 1.58 | 0 | 0.019 |
| 284 | MIR6875_5P | 55 | 0.75 | 1.58 | 0.004 | 0.019 |
| 285 | PID_AP1_PATHWAY | 51 | 0.74 | 1.57 | 0.003 | 0.019 |
| 286 | PETROVA_ENDOTHELIUM_LYMPHATIC_VS_BLOOI | 125 | 0.68 | 1.57 | 0 | 0.02 |
| 287 | REACTOME_INTERLEUKIN_3_INTERLEUKIN_5_ANI | 36 | 0.79 | 1.57 | 0.003 | 0.02 |
| 288 | NAGASHIMA_NRG1_SIGNALING_UP | 155 | 0.67 | 1.57 | 0 | 0.02 |
| 289 | SHIN_B_CELL_LYMPHOMA_CLUSTER_2 | 25 | 0.82 | 1.57 | 0.003 | 0.02 |
| 290 | JOHANSSON_GLIOMAGENESIS_BY_PDGFB_UP | 54 | 0.74 | 1.57 | 0.001 | 0.02 |
| 291 | WP_VITAMIN_D_RECEPTOR_PATHWAY | 104 | 0.69 | 1.57 | 0.001 | 0.02 |
| 292 | AMIT_SERUM_RESPONSE_120_MCF10A | 61 | 0.73 | 1.57 | 0.001 | 0.02 |
| 293 | FARMER_BREAST_CANCER_CLUSTER_1 | 21 | 0.84 | 1.57 | 0.001 | 0.02 |
| 294 | REACTOME_SIRT1_NEGATIVELY_REGULATES_RR | 37 | 0.78 | 1.57 | 0.001 | 0.02 |

| | | | | | | |
|---|---|---|---|---|---|---|
| 295 | REACTOME_GENE_SILENCING_BY_RNA | 98 | 0.7 | 1.57 | 0 | 0.02 |
| 296 | KEGG_ARACHIDONIC_ACID_METABOLISM | 26 | 0.81 | 1.57 | 0.003 | 0.022 |
| 297 | WP_IL7_SIGNALING_PATHWAY | 23 | 0.82 | 1.57 | 0.003 | 0.022 |
| 298 | CAHOY_ASTROGLIAL | 68 | 0.72 | 1.57 | 0 | 0.022 |
| 299 | TARTE_PLASMA_CELL_VS_PLASMABLAST_UP | 261 | 0.65 | 1.57 | 0 | 0.022 |
| 300 | WP_MITOCHONDRIAL_IMMUNE_RESPONSE_TO_S/ | 28 | 0.82 | 1.57 | 0 | 0.023 |
| 301 | WU_HBX_TARGETS_2_UP | 20 | 0.85 | 1.56 | 0.001 | 0.023 |
| 302 | GAL_LEUKEMIC_STEM_CELL_DN | 146 | 0.66 | 1.56 | 0.001 | 0.023 |
| 303 | BORLAK_LIVER_CANCER_EGF_UP | 41 | 0.77 | 1.56 | 0 | 0.023 |
| 304 | GRAHAM_CML_QUIESCENT_VS_CML_DIVIDING_UI | 15 | 0.89 | 1.56 | 0 | 0.024 |
| 305 | REACTOME_INTERLEUKIN_RECEPTOR_SHC_SIGN | 18 | 0.86 | 1.56 | 0.003 | 0.024 |
| 306 | NFKB_Q6_01 | 201 | 0.66 | 1.56 | 0 | 0.024 |
| 307 | WORSCHECH_TUMOR_REJECTION_UP | 27 | 0.82 | 1.56 | 0.004 | 0.024 |
| 308 | JEON_SMAD6_TARGETS_UP | 21 | 0.84 | 1.56 | 0 | 0.025 |
| 309 | PAPASPYRIDONOS_UNSTABLE_ATEROSCLEROTI( | 30 | 0.79 | 1.56 | 0 | 0.025 |
| 310 | REACTOME_SIGNALING_BY_INTERLEUKINS | 328 | 0.65 | 1.56 | 0 | 0.025 |
| 311 | REACTOME_METABOLISM_OF_PORPHYRINS | 20 | 0.85 | 1.56 | 0.003 | 0.025 |
| 312 | SALVADOR_MARTIN_PEDIATRIC_TBD_ANTI_TNF_T | 20 | 0.86 | 1.56 | 0.001 | 0.025 |
| 313 | CHAUHAN_RESPONSE_TO_METHOXYESTRADIOL_ | 96 | 0.69 | 1.56 | 0.002 | 0.025 |
| 314 | LIM_MAMMARY_LUMINAL_PROGENITOR_UP | 37 | 0.77 | 1.56 | 0.001 | 0.025 |
| 315 | DAUER_STAT3_TARGETS_UP | 37 | 0.76 | 1.56 | 0.004 | 0.025 |
| 316 | RYAN_MANTLE_CELL_LYMPHOMA_NOTCH_DIREC | 115 | 0.68 | 1.56 | 0.001 | 0.026 |
| 317 | RASHI_RESPONSE_TO_IONIZING_RADIATION_1 | 32 | 0.78 | 1.56 | 0.003 | 0.026 |
| 318 | WP_TYPE_I_INTERFERON_INDUCTION_AND_SIGN/ | 26 | 0.81 | 1.56 | 0.004 | 0.026 |
| 319 | GENTILE_UV_RESPONSE_CLUSTER_D6 | 32 | 0.78 | 1.56 | 0.004 | 0.026 |
| 320 | REACTOME_BASE_EXCISION_REPAIR_AP_SITE_F | 38 | 0.77 | 1.56 | 0.003 | 0.026 |
| 321 | WP_NONSMALL_CELL_LUNG_CANCER | 71 | 0.71 | 1.56 | 0.002 | 0.026 |
| 322 | KEGG_APOPTOSIS | 69 | 0.71 | 1.56 | 0.001 | 0.026 |
| 323 | IRF2_01 | 106 | 0.68 | 1.56 | 0 | 0.026 |
| 324 | RUTELLA_RESPONSE_TO_HGF_VS_CSF2RB_AND_ | 218 | 0.65 | 1.56 | 0 | 0.026 |
| 325 | ZHOU_TNF_SIGNALING_30MIN | 49 | 0.74 | 1.56 | 0.003 | 0.026 |
| 326 | REACTOME_DEPOSITION_OF_NEW_CENPA_CONT | 50 | 0.73 | 1.56 | 0.002 | 0.026 |
| 327 | TAKEDA_TARGETS_OF_NUP98_HOXA9_FUSION_1 | 106 | 0.68 | 1.56 | 0.001 | 0.026 |

| | | | | | | |
|---|---|---|---|---|---|---|
| 328 | ATM_DN.V1_UP | 87 | 0.69 | 1.56 | 0 | 0.027 |
| 329 | DUTTA_APOPTOSIS_VIA_NFKB | 28 | 0.8 | 1.55 | 0.004 | 0.027 |
| 330 | WEIGEL_OXIDATIVE_STRESS_RESPONSE | 30 | 0.8 | 1.55 | 0.004 | 0.027 |
| 331 | KEGG_TOLL_LIKE_RECEPTOR_SIGNALING_PATHW | 72 | 0.72 | 1.55 | 0 | 0.027 |
| 332 | DAZARD_RESPONSE_TO_UV_NHEK_UP | 181 | 0.66 | 1.55 | 0 | 0.027 |
| 333 | ZHOU_INFLAMMATORY_RESPONSE_LPS_UP | 269 | 0.65 | 1.55 | 0 | 0.027 |
| 334 | GREGORY_SYNTHETIC_LETHAL_WITH_IMATINIB | 122 | 0.67 | 1.55 | 0 | 0.027 |
| 335 | KRCTCNNNNMANAGC_UNKNOWN | 38 | 0.76 | 1.55 | 0 | 0.027 |
| 336 | GRUETZMANN_PANCREATIC_CANCER_UP | 310 | 0.64 | 1.55 | 0 | 0.028 |
| 337 | ZWANG_EGF_PERSISTENTLY_UP | 26 | 0.81 | 1.55 | 0.001 | 0.028 |
| 338 | VEGF_A_UP.V1_UP | 139 | 0.66 | 1.55 | 0.001 | 0.028 |
| 339 | GALINDO_IMMUNE_RESPONSE_TO_ENTEROTOXIN | 67 | 0.71 | 1.55 | 0 | 0.028 |
| 340 | OUELLET_CULTURED_OVARIAN_CANCER_INVASIV | 62 | 0.72 | 1.55 | 0.004 | 0.027 |
| 341 | MARKEY_RB1_CHRONIC_LOF_DN | 78 | 0.7 | 1.55 | 0.002 | 0.028 |
| 342 | RELA_DN.V1_DN | 64 | 0.72 | 1.55 | 0.001 | 0.028 |
| 343 | HAN_JNK_SINGALING_UP | 29 | 0.8 | 1.55 | 0.001 | 0.028 |
| 344 | FURUKAWA_DUSP6_TARGETS_PCI35_UP | 54 | 0.73 | 1.55 | 0.001 | 0.028 |
| 345 | WANG_CISPLATIN_RESPONSE_AND_XPC_DN | 192 | 0.65 | 1.55 | 0 | 0.028 |
| 346 | REACTOME_INHIBITION_OF_DNA_RECOMBINATIO | 44 | 0.75 | 1.55 | 0.004 | 0.028 |
| 347 | LEI_MYB_TARGETS | 269 | 0.65 | 1.55 | 0 | 0.028 |
| 348 | HOWLIN_PUBERTAL_MAMMARY_GLAND | 36 | 0.78 | 1.55 | 0.003 | 0.028 |
| 349 | REACTOME_PURINERGIC_SIGNALING_IN_LEISHM | 17 | 0.87 | 1.55 | 0 | 0.028 |
| 350 | KEGG_CHRONIC_MYELOID_LEUKEMIA | 68 | 0.71 | 1.55 | 0 | 0.028 |
| 351 | ZHOU_TNF_SIGNALING_4HR | 49 | 0.73 | 1.55 | 0.003 | 0.028 |
| 352 | MAHAJAN_RESPONSE_TO_IL1A_UP | 53 | 0.73 | 1.55 | 0.004 | 0.028 |
| 353 | VERHAAK_AML_WITH_NPM1_MUTATED_UP | 118 | 0.67 | 1.55 | 0 | 0.028 |
| 354 | SIRNA_EIF4GI_DN | 77 | 0.7 | 1.55 | 0.002 | 0.028 |
| 355 | LAU_APOPTOSIS_CDKN2A_UP | 53 | 0.74 | 1.55 | 0.001 | 0.029 |
| 356 | WP_NUCLEOTIDEBINDING_OLIGOMERIZATION_DO | 25 | 0.81 | 1.55 | 0.004 | 0.029 |
| 357 | BAE_BRCA1_TARGETS_DN | 29 | 0.79 | 1.55 | 0.004 | 0.029 |
| 358 | IRF7_01 | 205 | 0.65 | 1.55 | 0 | 0.029 |
| 359 | DOANE_BREAST_CANCER_CLASSES_DN | 30 | 0.78 | 1.55 | 0.004 | 0.029 |
| 360 | BOQUEST_STEM_CELL_DN | 154 | 0.65 | 1.55 | 0 | 0.029 |

| | | | | | | |
|---|---|---|---|---|---|---|
| 361 | BOQUEST_STEM_CELL_CULTURED_VS_FRESH_UI | 314 | 0.64 | 1.55 | 0 | 0.029 |
| 362 | REACTOME_RHO_GTPASES_ACTIVATE_PKNS | 58 | 0.73 | 1.55 | 0.003 | 0.029 |
| 363 | REACTOME_DISEASES_OF_PROGRAMMED_CELL_ | 69 | 0.71 | 1.55 | 0.001 | 0.029 |
| 364 | SERVITJA_ISLET_HNF1A_TARGETS_UP | 119 | 0.67 | 1.55 | 0 | 0.03 |
| 365 | PLASARI_TGFB1_TARGETS_1HR_UP | 29 | 0.78 | 1.55 | 0.003 | 0.03 |
| 366 | WP_PROTEASOME_DEGRADATION | 62 | 0.72 | 1.55 | 0.001 | 0.03 |
| 367 | ODONNELL_TARGETS_OF_MYC_AND_TFRC_UP | 61 | 0.72 | 1.55 | 0.001 | 0.03 |
| 368 | ONO_FOXP3_TARGETS_DN | 27 | 0.79 | 1.55 | 0.003 | 0.03 |
| 369 | BRACHAT_RESPONSE_TO_METHOTREXATE_DN | 25 | 0.8 | 1.55 | 0.008 | 0.03 |
| 370 | TURASHVILI_BREAST_DUCTAL_CARCINOMA_VS_L | 45 | 0.74 | 1.54 | 0.004 | 0.031 |
| 371 | KEGG_METABOLISM_OF_XENOBIOTICS_BY_CYTC | 27 | 0.8 | 1.54 | 0.001 | 0.031 |
| 372 | TIAN_TNF_SIGNALING_VIA_NFKB | 23 | 0.81 | 1.54 | 0.001 | 0.031 |
| 373 | KEGG_DRUG_METABOLISM_CYTOCHROME_P450 | 23 | 0.82 | 1.54 | 0.003 | 0.032 |
| 374 | PID_CERAMIDE_PATHWAY | 44 | 0.73 | 1.54 | 0.003 | 0.033 |
| 375 | REACTOME_GENERATION_OF_SECOND_MESSENG | 21 | 0.82 | 1.54 | 0.004 | 0.033 |
| 376 | LEE_LIVER_CANCER_CIPROFIBRATE_DN | 28 | 0.79 | 1.54 | 0.004 | 0.033 |
| 377 | WORSCHECH_TUMOR_EVASION_AND_TOLEROGE | 17 | 0.85 | 1.54 | 0.004 | 0.034 |
| 378 | P53_DN.V2_UP | 73 | 0.69 | 1.54 | 0.004 | 0.034 |
| 379 | RUTELLA_RESPONSE_TO_HGF_UP | 339 | 0.63 | 1.54 | 0 | 0.034 |
| 380 | TONKS_TARGETS_OF_RUNX1_RUNX1T1_FUSION_ | 16 | 0.87 | 1.54 | 0.002 | 0.034 |
| 381 | MELLMAN_TUT1_TARGETS_DN | 42 | 0.74 | 1.54 | 0.005 | 0.034 |
| 382 | KEGG_NON_SMALL_CELL_LUNG_CANCER | 52 | 0.73 | 1.54 | 0.003 | 0.034 |
| 383 | MEDINA_SMARCA4_TARGETS | 32 | 0.77 | 1.54 | 0.001 | 0.035 |
| 384 | REACTOME_GPCR_LIGAND_BINDING | 176 | 0.65 | 1.54 | 0 | 0.035 |
| 385 | GOTZMANN_EPITHELIAL_TO_MESENCHYMAL_TRA | 53 | 0.72 | 1.54 | 0.005 | 0.035 |
| 386 | WP_GASTRIN_SIGNALING_PATHWAY | 101 | 0.67 | 1.54 | 0.002 | 0.035 |
| 387 | LIU_NASOPHARYNGEAL_CARCINOMA | 63 | 0.71 | 1.54 | 0.004 | 0.035 |
| 388 | WU_HBX_TARGETS_3_UP | 18 | 0.84 | 1.54 | 0.003 | 0.035 |
| 389 | REACTOME_OTHER_INTERLEUKIN_SIGNALING | 18 | 0.84 | 1.54 | 0.006 | 0.035 |
| 390 | WP_CONSTITUTIVE_ANDROSTANE_RECEPTOR_PA | 20 | 0.84 | 1.54 | 0.007 | 0.036 |
| 391 | REACTOME_TRANSCRIPTIONAL_REGULATION_BY_ | 76 | 0.7 | 1.54 | 0.001 | 0.036 |
| 392 | PID_MYC_REPRESS_PATHWAY | 57 | 0.72 | 1.53 | 0.003 | 0.036 |
| 393 | STK33_UP | 220 | 0.65 | 1.53 | 0 | 0.036 |

| | | | | | | |
|---|---|---|---|---|---|---|
| 394 | TIEN_INTESTINE_PROBIOTICS_2HR_UP | 29 | 0.79 | 1.53 | 0.007 | 0.037 |
| 395 | TIMOFEEVA_GROWTH_STRESS_VIA_STAT1_DN | 16 | 0.86 | 1.53 | 0.001 | 0.037 |
| 396 | DIAZ_CHRONIC_MYELOGENOUS_LEUKEMIA_DN | 72 | 0.7 | 1.53 | 0 | 0.037 |
| 397 | HOLLERN_SQUAMOUS_BREAST_TUMOR | 86 | 0.68 | 1.53 | 0.001 | 0.038 |
| 398 | PTEN_DN.V2_UP | 88 | 0.69 | 1.53 | 0.001 | 0.038 |
| 399 | RHEIN_ALL_GLUCOCORTICOID_THERAPY_UP | 49 | 0.73 | 1.53 | 0.001 | 0.038 |
| 400 | ANASTASSIOU_MULTICANCER_INVASIVENESS_SI | 35 | 0.75 | 1.53 | 0.009 | 0.038 |
| 401 | REACTOME_SIGNALING_BY_ALK | 27 | 0.8 | 1.53 | 0 | 0.038 |
| 402 | TURASHVILI_BREAST_DUCTAL_CARCINOMA_VS_D | 142 | 0.66 | 1.53 | 0 | 0.038 |
| 403 | LINDGREN_BLADDER_CANCER_HIGH_RECURRENC | 39 | 0.75 | 1.53 | 0.007 | 0.038 |
| 404 | DANG_REGULATED_BY_MYC_DN | 207 | 0.64 | 1.53 | 0 | 0.039 |
| 405 | MIR3126_5P | 54 | 0.71 | 1.53 | 0.005 | 0.039 |
| 406 | JINESH_BLEBBISHIELD_TRANSFORMED_STEM_CE | 131 | 0.66 | 1.53 | 0 | 0.04 |
| 407 | MIKKELSEN_MCV6_LCP_WITH_H3K4ME3 | 110 | 0.67 | 1.53 | 0.001 | 0.04 |
| 408 | HOFMANN_CELL_LYMPHOMA_DN | 29 | 0.77 | 1.53 | 0.011 | 0.04 |
| 409 | YAGI_AML_WITH_T_8_21_TRANSLOCATION | 315 | 0.63 | 1.53 | 0 | 0.04 |
| 410 | WP_PI3KAKTMTOR_VITD3_SIGNALING | 17 | 0.85 | 1.53 | 0.006 | 0.04 |
| 411 | WP_NOVEL_INTRACELLULAR_COMPONENTS_OF_ | 55 | 0.71 | 1.53 | 0.004 | 0.041 |
| 412 | PEDERSEN_TARGETS_OF_611CTF_ISOFORM_OF_ | 59 | 0.71 | 1.53 | 0.001 | 0.041 |
| 413 | TONKS_TARGETS_OF_RUNX1_RUNX1T1_FUSION_ | 135 | 0.66 | 1.53 | 0 | 0.041 |
| 414 | CERIBELLI_PROMOTERS_INACTIVE_AND_BOUND_ | 20 | 0.82 | 1.53 | 0.003 | 0.041 |
| 415 | HOLLERN_SOLID_NODULAR_BREAST_TUMOR_DN | 23 | 0.8 | 1.53 | 0.019 | 0.041 |
| 416 | GILDEA_METASTASIS | 17 | 0.85 | 1.53 | 0 | 0.041 |
| 417 | WP_METAPATHWAY_BIOTRANSFORMATION_PHAS | 92 | 0.68 | 1.53 | 0 | 0.042 |
| 418 | MIR1255A_MIR1255B_5P | 54 | 0.72 | 1.52 | 0.005 | 0.042 |
| 419 | REACTOME_PHASE_II_CONJUGATION_OF_COMPC | 65 | 0.69 | 1.52 | 0.001 | 0.042 |
| 420 | REACTOME_POSITIVE_EPIGENETIC_REGULATION | 75 | 0.69 | 1.52 | 0.004 | 0.042 |
| 421 | SENESE_HDAC3_TARGETS_UP | 418 | 0.62 | 1.52 | 0 | 0.042 |
| 422 | SERVITJA_LIVER_HNF1A_TARGETS_DN | 87 | 0.68 | 1.52 | 0.002 | 0.042 |
| 423 | REACTOME_B_WICH_COMPLEX_POSITIVELY_REG | 60 | 0.71 | 1.52 | 0.002 | 0.042 |
| 424 | REACTOME_RNA_POLYMERASE_I_PROMOTER_ES | 60 | 0.71 | 1.52 | 0.001 | 0.042 |
| 425 | BIOCARTA_HIVNEF_PATHWAY | 54 | 0.71 | 1.52 | 0.001 | 0.043 |
| 426 | DEURIG_T_CELL_PROLYMPHOCYTIC_LEUKEMIA_I | 258 | 0.64 | 1.52 | 0 | 0.043 |

| | | | | | | |
|---|---|---|---|---|---|---|
| 427 | AMIT_EGF_RESPONSE_60_MCF10A | 33 | 0.77 | 1.52 | 0.005 | 0.043 |
| 428 | REACTOME_MEIOTIC_SYNAPSIS | 48 | 0.72 | 1.52 | 0.001 | 0.043 |
| 429 | BIOCARTA_SPRY_PATHWAY | 16 | 0.86 | 1.52 | 0.003 | 0.043 |
| 430 | ALK_DN.V1_UP | 77 | 0.68 | 1.52 | 0.001 | 0.043 |
| 431 | DIRMEIER_LMP1_RESPONSE_LATE_DN | 26 | 0.79 | 1.52 | 0.006 | 0.043 |
| 432 | HUANG_DASATINIB_SENSITIVITY_UP | 64 | 0.7 | 1.52 | 0.007 | 0.044 |
| 433 | HOSHIDA_LIVER_CANCER_SURVIVAL_UP | 58 | 0.71 | 1.52 | 0.001 | 0.044 |
| 434 | WP_COMPLEMENT_SYSTEM | 50 | 0.73 | 1.52 | 0.005 | 0.044 |
| 435 | REACTOME_ZBP1_DAI_MEDIATED_INDUCTION_OF | 19 | 0.83 | 1.52 | 0.009 | 0.045 |
| 436 | PID_FRA_PATHWAY | 32 | 0.77 | 1.52 | 0.006 | 0.045 |
| 437 | CHIARADONNA_NEOPLASTIC_TRANSFORMATION_ | 31 | 0.77 | 1.52 | 0.001 | 0.046 |
| 438 | WP_NRF2_PATHWAY | 99 | 0.67 | 1.52 | 0.001 | 0.046 |
| 439 | REACTOME_CD209_DC_SIGN_SIGNALING | 18 | 0.83 | 1.52 | 0.005 | 0.046 |
| 440 | WP_DNA_DAMAGE_RESPONSE | 66 | 0.7 | 1.52 | 0.001 | 0.046 |
| 441 | REACTOME_HATS_ACETYLATE_HISTONES | 106 | 0.67 | 1.52 | 0.001 | 0.047 |
| 442 | VERHAAK_GLIOBLASTOMA_MESENCHYMAL | 152 | 0.65 | 1.52 | 0 | 0.047 |
| 443 | WP_TLR4_SIGNALING_AND_TOLERANCE | 27 | 0.79 | 1.52 | 0.01 | 0.048 |
| 444 | REACTOME_REGULATED_NECROSIS | 47 | 0.73 | 1.52 | 0.003 | 0.048 |
| 445 | LEE_LIVER_CANCER_MYC_TGFA_DN | 35 | 0.76 | 1.51 | 0.005 | 0.049 |
| 446 | IRF1_01 | 196 | 0.64 | 1.51 | 0 | 0.049 |
| 447 | SCHRAETS_MLL_TARGETS_UP | 23 | 0.8 | 1.51 | 0.01 | 0.049 |
| 448 | NOJIMA_SFRP2_TARGETS_UP | 26 | 0.79 | 1.51 | 0.011 | 0.049 |
| 449 | PID_FOXM1_PATHWAY | 37 | 0.75 | 1.51 | 0.007 | 0.049 |
| 450 | CHARAFE_BREAST_CANCER_LUMINAL_VS_BASAL | 358 | 0.63 | 1.51 | 0 | 0.049 |
| 451 | MIR218_2_3P | 39 | 0.74 | 1.51 | 0.011 | 0.05 |
| 452 | CAIRO_LIVER_DEVELOPMENT_DN | 131 | 0.65 | 1.51 | 0 | 0.05 |
| 453 | ZHAN_EARLY_DIFFERENTIATION_GENES_DN | 36 | 0.74 | 1.51 | 0.004 | 0.051 |
| 454 | PID_P53_DOWNSTREAM_PATHWAY | 123 | 0.65 | 1.51 | 0.001 | 0.051 |
| 455 | WP_MIRNA_REGULATION_OF_DNA_DAMAGE_RES | 69 | 0.69 | 1.51 | 0.001 | 0.051 |
| 456 | REACTOME_ESR_MEDIATED_SIGNALING | 170 | 0.64 | 1.51 | 0 | 0.052 |
| 457 | WP_FERROPTOSIS | 55 | 0.72 | 1.51 | 0.007 | 0.052 |
| 458 | ALCALAY_AML_BY_NPM1_LOCALIZATION_DN | 153 | 0.65 | 1.51 | 0.001 | 0.052 |
| 459 | MISSIAGLIA_REGULATED_BY_METHYLATION_UP | 102 | 0.66 | 1.51 | 0.001 | 0.053 |

| | | | | | | |
|---|---|---|---|---|---|---|
| 460 | PID_REG_GR_PATHWAY | 63 | 0.69 | 1.51 | 0.007 | 0.053 |
| 461 | GARGALOVIC_RESPONSE_TO_OXIDIZED_PHOSPH | 22 | 0.8 | 1.51 | 0.018 | 0.053 |
| 462 | MYLLYKANGAS_AMPLIFICATION_HOT_SPOT_23 | 17 | 0.84 | 1.51 | 0.005 | 0.053 |
| 463 | REACTOME_RMTS_METHYLATE_HISTONE_ARGINI | 51 | 0.71 | 1.51 | 0.009 | 0.053 |
| 464 | WP_PREGNANE_X_RECEPTOR_PATHWAY | 17 | 0.83 | 1.51 | 0.011 | 0.054 |
| 465 | LENAOUR_DENDRITIC_CELL_MATURATION_DN | 85 | 0.67 | 1.51 | 0.001 | 0.054 |
| 466 | NUNODA_RESPONSE_TO_DASATINIB_IMATINIB_UI | 29 | 0.76 | 1.51 | 0.012 | 0.054 |
| 467 | REACTOME_PRE_NOTCH_EXPRESSION_AND_PRC | 76 | 0.68 | 1.51 | 0.002 | 0.054 |
| 468 | NFKAPPAB65_01 | 200 | 0.63 | 1.51 | 0 | 0.054 |
| 469 | GOLDRATH_ANTIGEN_RESPONSE | 292 | 0.62 | 1.51 | 0 | 0.054 |
| 470 | WP_SARS_CORONAVIRUS_AND_INNATE_IMMUNIT | 18 | 0.83 | 1.51 | 0.012 | 0.054 |
| 471 | RIZKI_TUMOR_INVASIVENESS_3D_DN | 207 | 0.63 | 1.51 | 0 | 0.054 |
| 472 | REACTOME_UB_SPECIFIC_PROCESSING_PROTEA | 173 | 0.64 | 1.51 | 0 | 0.055 |
| 473 | REACTOME_NEGATIVE_EPIGENETIC_REGULATIOI | 78 | 0.68 | 1.51 | 0.004 | 0.055 |
| 474 | REACTOME_TRAF6_MEDIATED_NF_KB_ACTIVATIC | 23 | 0.8 | 1.5 | 0.016 | 0.056 |
| 475 | REACTOME_NUCLEOTIDE_BINDING_DOMAIN_LEU( | 51 | 0.71 | 1.5 | 0.003 | 0.056 |
| 476 | WP_PROSTAGLANDIN_SYNTHESIS_AND_REGULAT | 23 | 0.79 | 1.5 | 0.017 | 0.056 |
| 477 | KEGG_ACUTE_MYELOID_LEUKEMIA | 53 | 0.7 | 1.5 | 0.005 | 0.056 |
| 478 | MORI_IMMATURE_B_LYMPHOCYTE_UP | 41 | 0.72 | 1.5 | 0.009 | 0.056 |
| 479 | VANASSE_BCL2_TARGETS_DN | 47 | 0.71 | 1.5 | 0.012 | 0.056 |
| 480 | GRABARCZYK_BCL11B_TARGETS_UP | 67 | 0.69 | 1.5 | 0.002 | 0.057 |
| 481 | MACAEVA_PBMC_RESPONSE_TO_IR | 82 | 0.68 | 1.5 | 0.006 | 0.056 |
| 482 | BURTON_ADIPOGENESIS_4 | 41 | 0.73 | 1.5 | 0.012 | 0.057 |
| 483 | NFE2L2.V2 | 309 | 0.62 | 1.5 | 0 | 0.057 |
| 484 | KIM_RESPONSE_TO_TSA_AND_DECITABINE_UP | 84 | 0.67 | 1.5 | 0.002 | 0.056 |
| 485 | WP_NETRINUNC5B_SIGNALING_PATHWAY | 42 | 0.72 | 1.5 | 0.012 | 0.057 |
| 486 | PID_HIV_NEF_PATHWAY | 33 | 0.75 | 1.5 | 0.007 | 0.057 |
| 487 | APPIERTO_RESPONSE_TO_FENRETINIDE_DN | 46 | 0.72 | 1.5 | 0.008 | 0.057 |
| 488 | PODAR_RESPONSE_TO_ADAPHOSTIN_UP | 137 | 0.65 | 1.5 | 0 | 0.058 |
| 489 | WINNEPENNINCKX_MELANOMA_METASTASIS_DN | 30 | 0.76 | 1.5 | 0.007 | 0.059 |
| 490 | PID_TELOMERASE_PATHWAY | 63 | 0.69 | 1.5 | 0.004 | 0.06 |
| 491 | WP_TOLLLIKE_RECEPTOR_SIGNALING_RELATED_ | 25 | 0.78 | 1.5 | 0.012 | 0.06 |
| 492 | PID_NFKAPPAB_CANONICAL_PATHWAY | 22 | 0.8 | 1.5 | 0.011 | 0.06 |

| | | | | | | |
|---|---|---|---|---|---|---|
| 493 | SENESE_HDAC1_AND_HDAC2_TARGETS_UP | 177 | 0.63 | 1.5 | 0 | 0.061 |
| 494 | CROMER_TUMORIGENESIS_DN | 24 | 0.79 | 1.5 | 0.013 | 0.061 |
| 495 | LINDSTEDT_DENDRITIC_CELL_MATURATION_C | 50 | 0.71 | 1.5 | 0.006 | 0.061 |
| 496 | REACTOME_RUNX1_REGULATES_GENES_INVOLV | 60 | 0.7 | 1.5 | 0.005 | 0.062 |
| 497 | MCDOWELL_ACUTE_LUNG_INJURY_UP | 30 | 0.76 | 1.5 | 0.018 | 0.063 |
| 498 | ONDER_CDH1_TARGETS_1_UP | 119 | 0.65 | 1.5 | 0.002 | 0.063 |
| 499 | PID_SYNDECAN_4_PATHWAY | 26 | 0.78 | 1.49 | 0.01 | 0.065 |
| 500 | DELYS_THYROID_CANCER_UP | 331 | 0.62 | 1.49 | 0 | 0.065 |
| 501 | KEGG_GLIOMA | 59 | 0.69 | 1.49 | 0.008 | 0.065 |
| 502 | ALCALA_APOPTOSIS | 78 | 0.68 | 1.49 | 0.004 | 0.066 |
| 503 | PID_NFAT_TFPATHWAY | 27 | 0.76 | 1.49 | 0.012 | 0.067 |
| 504 | BROWNE_HCMV_INFECTION_6HR_UP | 42 | 0.72 | 1.49 | 0.01 | 0.067 |
| 505 | WP_COMPLEMENT_SYSTEM_IN_NEURONAL_DEVE | 67 | 0.68 | 1.49 | 0.006 | 0.067 |
| 506 | TTGCWCAAY_CEBPB_02 | 53 | 0.71 | 1.49 | 0.005 | 0.068 |
| 507 | REACTOME_OVARIAN_TUMOR_DOMAIN_PROTEAS | 36 | 0.74 | 1.49 | 0.011 | 0.068 |
| 508 | NABA_COLLAGENS | 21 | 0.8 | 1.49 | 0.016 | 0.068 |
| 509 | KIM_BIPOLAR_DISORDER_OLIGODENDROCYTE_D | 38 | 0.73 | 1.49 | 0.008 | 0.068 |
| 510 | ZNF318_TARGET_GENES | 390 | 0.61 | 1.49 | 0 | 0.068 |
| 511 | LIU_SMARCA4_TARGETS | 39 | 0.73 | 1.49 | 0.009 | 0.068 |
| 512 | REACTOME_TNFR1_INDUCED_NFKAPPAB_SIGNAL | 30 | 0.75 | 1.49 | 0.009 | 0.068 |
| 513 | BIOCARTA_HER2_PATHWAY | 21 | 0.8 | 1.49 | 0.016 | 0.068 |
| 514 | WP_CORTICOTROPINRELEASING_HORMONE_SIGN | 74 | 0.67 | 1.49 | 0.004 | 0.069 |
| 515 | PID_TNF_PATHWAY | 45 | 0.72 | 1.49 | 0.013 | 0.071 |
| 516 | AKL_HTLV1_INFECTION_UP | 27 | 0.77 | 1.49 | 0.011 | 0.07 |
| 517 | PROVENZANI_METASTASIS_DN | 120 | 0.64 | 1.49 | 0.009 | 0.072 |
| 518 | WP_TCELL_ANTIGEN_RECEPTOR_TCR_PATHWAY_ | 43 | 0.71 | 1.49 | 0.013 | 0.072 |
| 519 | YAN_ESCAPE_FROM_ANOIKIS | 22 | 0.8 | 1.49 | 0.016 | 0.072 |
| 520 | WP_MAMMARY_GLAND_DEVELOPMENT_PATHWAY | 24 | 0.78 | 1.49 | 0.013 | 0.072 |
| 521 | DUAN_PRDM5_TARGETS | 65 | 0.69 | 1.49 | 0.011 | 0.072 |
| 522 | REACTOME_SIGNALING_BY_NUCLEAR_RECEPTOI | 226 | 0.62 | 1.49 | 0 | 0.073 |
| 523 | PEART_HDAC_PROLIFERATION_CLUSTER_DN | 65 | 0.69 | 1.49 | 0.005 | 0.073 |
| 524 | RYTAAWNNNTGAY_UNKNOWN | 41 | 0.71 | 1.49 | 0.015 | 0.073 |
| 525 | WP_RENINANGIOTENSINALDOSTERONE_SYSTEM_ | 30 | 0.76 | 1.49 | 0.008 | 0.073 |

| | | | | | | |
|---|---|---|---|---|---|---|
| 526 | WP_HEPATITIS_B_INFECTION | 131 | 0.63 | 1.48 | 0.005 | 0.073 |
| 527 | DALESSIO_TSA_RESPONSE | 16 | 0.82 | 1.48 | 0.011 | 0.074 |
| 528 | REACTOME_MAP3K8_TPL2_DEPENDENT_MAPK1_ | 16 | 0.82 | 1.48 | 0.014 | 0.075 |
| 529 | CAMPS_COLON_CANCER_COPY_NUMBER_DN | 33 | 0.75 | 1.48 | 0.016 | 0.075 |
| 530 | HU_GENOTOXIN_ACTION_DIRECT_VS_INDIRECT_4 | 29 | 0.76 | 1.48 | 0.016 | 0.075 |
| 531 | LIANG_SILENCED_BY_METHYLATION_UP | 24 | 0.79 | 1.48 | 0.007 | 0.075 |
| 532 | CASTELLANO_NRAS_TARGETS_UP | 55 | 0.69 | 1.48 | 0.002 | 0.076 |
| 533 | KYNG_DNA_DAMAGE_DN | 149 | 0.64 | 1.48 | 0 | 0.077 |
| 534 | HESS_TARGETS_OF_HOXA9_AND_MEIS1_DN | 54 | 0.7 | 1.48 | 0.009 | 0.077 |
| 535 | DOANE_BREAST_CANCER_ESR1_DN | 29 | 0.76 | 1.48 | 0.018 | 0.077 |
| 536 | REACTOME_AMYLOID_FIBER_FORMATION | 63 | 0.68 | 1.48 | 0.005 | 0.077 |
| 537 | KEGG_HEMATOPOIETIC_CELL_LINEAGE | 33 | 0.74 | 1.48 | 0.022 | 0.077 |
| 538 | WP_RAS_AND_BRADYKININ_PATHWAYS_IN_COVII | 20 | 0.81 | 1.48 | 0.015 | 0.077 |
| 539 | REACTOME_MEIOTIC_RECOMBINATION | 54 | 0.7 | 1.48 | 0.009 | 0.077 |
| 540 | BIOCARTA_DEATH_PATHWAY | 28 | 0.76 | 1.48 | 0.013 | 0.078 |
| 541 | KOKKINAKIS_METHIONINE_DEPRIVATION_48HR_U | 111 | 0.65 | 1.48 | 0.005 | 0.078 |
| 542 | BIOCARTA_RAC1_PATHWAY | 18 | 0.81 | 1.48 | 0.015 | 0.078 |
| 543 | NABA_ECM_AFFILIATED | 90 | 0.66 | 1.48 | 0.002 | 0.078 |
| 544 | STK33_SKM_UP | 212 | 0.62 | 1.48 | 0 | 0.08 |
| 545 | REACTOME_RUNX1_REGULATES_TRANSCRIPTION | 92 | 0.66 | 1.48 | 0.004 | 0.08 |
| 546 | BURTON_ADIPOGENESIS_PEAK_AT_2HR | 41 | 0.7 | 1.48 | 0.016 | 0.081 |
| 547 | SARTIPY_BLUNTED_BY_INSULIN_RESISTANCE_UI | 15 | 0.84 | 1.48 | 0.016 | 0.081 |
| 548 | MIKKELSEN_MEF_LCP_WITH_H3K4ME3 | 86 | 0.66 | 1.48 | 0.004 | 0.081 |
| 549 | FOURNIER_ACINAR_DEVELOPMENT_LATE_DN | 20 | 0.8 | 1.48 | 0.008 | 0.081 |
| 550 | GENTILE_UV_LOW_DOSE_UP | 24 | 0.78 | 1.48 | 0.017 | 0.081 |
| 551 | ZNF777_TARGET_GENES | 39 | 0.72 | 1.48 | 0.019 | 0.081 |
| 552 | TSAI_RESPONSE_TO_RADIATION_THERAPY | 21 | 0.8 | 1.48 | 0.01 | 0.081 |
| 553 | BIOCARTA_CELLCYCLE_PATHWAY | 23 | 0.79 | 1.48 | 0.012 | 0.081 |
| 554 | ESC_J1_UP_EARLY.V1_DN | 140 | 0.64 | 1.48 | 0.004 | 0.081 |
| 555 | RICKMAN_HEAD_AND_NECK_CANCER_C | 44 | 0.71 | 1.48 | 0.006 | 0.081 |
| 556 | WP_FAS_LIGAND_PATHWAY_AND_STRESS_INDUC | 38 | 0.73 | 1.48 | 0.01 | 0.081 |
| 557 | JIANG_AGING_HYPOTHALAMUS_DN | 31 | 0.75 | 1.48 | 0.013 | 0.081 |
| 558 | PID_INTEGRIN2_PATHWAY | 15 | 0.83 | 1.48 | 0.011 | 0.081 |

| | | | | | | |
|---|---|---|---|---|---|---|
| 559 | MANN_RESPONSE_TO_AMIFOSTINE_UP | 19 | 0.81 | 1.47 | 0.007 | 0.082 |
| 560 | NFKB_C | 221 | 0.62 | 1.47 | 0 | 0.082 |
| 561 | KEGG_PRION_DISEASES | 23 | 0.77 | 1.47 | 0.019 | 0.082 |
| 562 | FONTAINE_FOLLICULAR_THYROID_ADENOMA_DN | 55 | 0.69 | 1.47 | 0.011 | 0.083 |
| 563 | PID_CASPASE_PATHWAY | 46 | 0.71 | 1.47 | 0.013 | 0.084 |
| 564 | HASLINGER_B_CLL_WITH_6Q21_DELETION | 17 | 0.82 | 1.47 | 0.02 | 0.084 |
| 565 | AMIT_EGF_RESPONSE_40_HELA | 39 | 0.72 | 1.47 | 0.017 | 0.084 |
| 566 | PTEN_DN.V1_DN | 95 | 0.65 | 1.47 | 0.003 | 0.084 |
| 567 | SEIDEN_MET_SIGNALING | 18 | 0.83 | 1.47 | 0.014 | 0.084 |
| 568 | BRACHAT_RESPONSE_TO_CAMPTOTHECIN_DN | 41 | 0.72 | 1.47 | 0.013 | 0.084 |
| 569 | WP_GASTRIC_CANCER_NETWORK_2 | 29 | 0.75 | 1.47 | 0.016 | 0.085 |
| 570 | REACTOME_PKMTS_METHYLATE_HISTONE_LYSIN | 52 | 0.69 | 1.47 | 0.005 | 0.084 |
| 571 | WP_FATTY_ACID_BIOSYNTHESIS | 22 | 0.78 | 1.47 | 0.017 | 0.084 |
| 572 | MIR4771 | 39 | 0.72 | 1.47 | 0.011 | 0.084 |
| 573 | HELLER_HDAC_TARGETS_SILENCED_BY_METHYL | 236 | 0.61 | 1.47 | 0 | 0.084 |
| 574 | SCHEIDEREIT_IKK_TARGETS | 16 | 0.83 | 1.47 | 0.012 | 0.085 |
| 575 | MCBRYAN_PUBERTAL_TGFB1_TARGETS_DN | 47 | 0.69 | 1.47 | 0.015 | 0.085 |
| 576 | ODONNELL_TFRC_TARGETS_UP | 279 | 0.61 | 1.47 | 0 | 0.085 |
| 577 | KRIGE_AMINO_ACID_DEPRIVATION | 28 | 0.75 | 1.47 | 0.021 | 0.085 |
| 578 | REACTOME_REGULATION_OF_TNFR1_SIGNALING | 35 | 0.72 | 1.47 | 0.026 | 0.085 |
| 579 | MUNSHI_MULTIPLE_MYELOMA_UP | 65 | 0.68 | 1.47 | 0.01 | 0.085 |
| 580 | JACKSON_DNMT1_TARGETS_UP | 67 | 0.67 | 1.47 | 0.009 | 0.085 |
| 581 | REACTOME_BILE_ACID_AND_BILE_SALT_METABC | 23 | 0.78 | 1.47 | 0.024 | 0.085 |
| 582 | ZHAN_MULTIPLE_MYELOMA_UP | 61 | 0.67 | 1.47 | 0.015 | 0.085 |
| 583 | MIR6732_5P | 24 | 0.77 | 1.47 | 0.016 | 0.085 |
| 584 | SESTO_RESPONSE_TO_UV_C0 | 99 | 0.65 | 1.47 | 0.006 | 0.085 |
| 585 | ZHANG_BREAST_CANCER_PROGENITORS_DN | 124 | 0.63 | 1.47 | 0.002 | 0.085 |
| 586 | TAKAO_RESPONSE_TO_UVB_RADIATION_UP | 80 | 0.66 | 1.47 | 0.009 | 0.086 |
| 587 | LIEN_BREAST_CARCINOMA_METAPLASTIC_VS_DU | 55 | 0.67 | 1.47 | 0.015 | 0.086 |
| 588 | REACTOME_COLLAGEN_CHAIN_TRIMERIZATION | 21 | 0.8 | 1.47 | 0.021 | 0.086 |
| 589 | NAM_FXYD5_TARGETS_DN | 16 | 0.82 | 1.47 | 0.017 | 0.086 |
| 590 | LEE_AGING_CEREBELLUM_UP | 57 | 0.68 | 1.47 | 0.006 | 0.086 |
| 591 | KEGG_P53_SIGNALING_PATHWAY | 66 | 0.67 | 1.47 | 0.011 | 0.086 |

| | | | | | |
|---|---|---|---|---|---|
| 592 WP_BURN_WOUND_HEALING | 68 | 0.67 | 1.47 | 0.009 | 0.087 |
| 593 REACTOME_SMAD2_SMAD3_SMAD4_HETEROTRIM | 31 | 0.74 | 1.47 | 0.018 | 0.087 |
| 594 NFKB_Q6 | 218 | 0.61 | 1.47 | 0 | 0.087 |
| 595 REACTOME_RIP_MEDIATED_NFKB_ACTIVATION_V | 16 | 0.82 | 1.47 | 0.027 | 0.088 |
| 596 TAKEDA_TARGETS_OF_NUP98_HOXA9_FUSION_6 | 34 | 0.73 | 1.47 | 0.019 | 0.088 |
| 597 CREL_01 | 220 | 0.61 | 1.47 | 0.001 | 0.088 |
| 598 DARWICHE_SQUAMOUS_CELL_CARCINOMA_UP | 110 | 0.64 | 1.47 | 0.001 | 0.088 |
| 599 RUTELLA_RESPONSE_TO_HGF_VS_CSF2RB_AND_ | 317 | 0.61 | 1.47 | 0 | 0.088 |
| 600 BROWN_MYELOID_CELL_DEVELOPMENT_UP | 102 | 0.64 | 1.47 | 0.003 | 0.088 |
| 601 PID_IL4_2PATHWAY | 39 | 0.72 | 1.47 | 0.016 | 0.088 |
| 602 RODRIGUES_THYROID_CARCINOMA_DN | 68 | 0.68 | 1.47 | 0.008 | 0.088 |
| 603 KRIEG_HYPOXIA_VIA_KDM3A | 44 | 0.71 | 1.47 | 0.008 | 0.088 |
| 604 WP_OREXIN_RECEPTOR_PATHWAY | 104 | 0.65 | 1.47 | 0.001 | 0.089 |
| 605 VANTVEER_BREAST_CANCER_ESR1_DN | 196 | 0.62 | 1.47 | 0.001 | 0.089 |
| 606 TOOKER_GEMCITABINE_RESISTANCE_UP | 71 | 0.67 | 1.47 | 0.006 | 0.089 |
| 607 PETRETTO_HEART_MASS_QTL_CIS_DN | 22 | 0.79 | 1.47 | 0.018 | 0.089 |
| 608 HELLEBREKERS_SILENCED_DURING_TUMOR_ANG | 57 | 0.69 | 1.46 | 0.009 | 0.089 |
| 609 ZHU_SKIL_TARGETS_UP | 16 | 0.82 | 1.46 | 0.025 | 0.089 |
| 610 REACTOME_INTERLEUKIN_2_FAMILY_SIGNALING | 32 | 0.73 | 1.46 | 0.026 | 0.09 |
| 611 REACTOME_G_ALPHA_I_SIGNALLING_EVENTS | 169 | 0.62 | 1.46 | 0.001 | 0.09 |
| 612 NAGASHIMA_EGF_SIGNALING_UP | 50 | 0.7 | 1.46 | 0.01 | 0.09 |
| 613 KYNG_RESPONSE_TO_H2O2_VIA_ERCC6_DN | 42 | 0.71 | 1.46 | 0.017 | 0.09 |
| 614 KOBAYASHI_RESPONSE_TO_ROMIDEPSIN | 19 | 0.8 | 1.46 | 0.016 | 0.09 |
| 615 REACTOME_FORMATION_OF_THE_BETA_CATENIN | 59 | 0.68 | 1.46 | 0.016 | 0.09 |
| 616 WP_PATHWAYS_OF_NUCLEIC_ACID_METABOLISM | 16 | 0.81 | 1.46 | 0.02 | 0.091 |
| 617 MAML1_TARGET_GENES | 192 | 0.62 | 1.46 | 0 | 0.091 |
| 618 SARTIPY_BLUNTED_BY_INSULIN_RESISTANCE_DN | 17 | 0.81 | 1.46 | 0.031 | 0.091 |
| 619 KLEIN_PRIMARY_EFFUSION_LYMPHOMA_DN | 47 | 0.7 | 1.46 | 0.005 | 0.091 |
| 620 CCAGGGG_MIR331 | 87 | 0.65 | 1.46 | 0.012 | 0.092 |
| 621 GUENTHER_GROWTH_SPHERICAL_VS_ADHERENT | 23 | 0.77 | 1.46 | 0.026 | 0.092 |
| 622 AP1_Q4 | 219 | 0.61 | 1.46 | 0 | 0.092 |
| 623 PID_IFNG_PATHWAY | 38 | 0.71 | 1.46 | 0.014 | 0.092 |
| 624 PID_IL2_STAT5_PATHWAY | 25 | 0.77 | 1.46 | 0.012 | 0.092 |

| | | | | | | |
|---|---|---|---|---|---|---|
| 625 | ENK_UV_RESPONSE_EPIDERMIS_UP | 254 | 0.6 | 1.46 | 0 | 0.093 |
| 626 | REACTOME_CLEC7A_DECTIN_1_SIGNALING | 95 | 0.64 | 1.46 | 0.005 | 0.093 |
| 627 | LEIN_ASTROCYTE_MARKERS | 26 | 0.76 | 1.46 | 0.012 | 0.094 |
| 628 | VANTVEER_BREAST_CANCER_BRCA1_UP | 29 | 0.75 | 1.46 | 0.025 | 0.094 |
| 629 | EBAUER_TARGETS_OF_PAX3_FOXO1_FUSION_UI | 137 | 0.63 | 1.46 | 0.001 | 0.095 |
| 630 | PID_IL12_STAT4_PATHWAY | 17 | 0.81 | 1.46 | 0.016 | 0.094 |
| 631 | CAMP_UP.V1_UP | 190 | 0.61 | 1.46 | 0 | 0.094 |
| 632 | HAN_JNK_SINGALING_DN | 22 | 0.77 | 1.46 | 0.016 | 0.096 |
| 633 | RASHI_RESPONSE_TO_IONIZING_RADIATION_2 | 112 | 0.64 | 1.46 | 0.007 | 0.096 |
| 634 | KEGG_RIG_I_LIKE_RECEPTOR_SIGNALING_PATH | 51 | 0.69 | 1.46 | 0.022 | 0.097 |
| 635 | FULCHER_INFLAMMATORY_RESPONSE_LECTIN_V | 490 | 0.6 | 1.46 | 0 | 0.096 |
| 636 | REACTOME_P75NTR_SIGNALS_VIA_NF_KB | 15 | 0.83 | 1.46 | 0.008 | 0.097 |
| 637 | PID_IL3_PATHWAY | 22 | 0.78 | 1.46 | 0.024 | 0.099 |
| 638 | SAGIV_CD24_TARGETS_DN | 41 | 0.71 | 1.46 | 0.014 | 0.099 |
| 639 | SMITH_TERT_TARGETS_UP | 139 | 0.62 | 1.46 | 0.001 | 0.099 |
| 640 | ONO_AML1_TARGETS_DN | 22 | 0.78 | 1.46 | 0.015 | 0.099 |
| 641 | CAVARD_LIVER_CANCER_MALIGNANT_VS_BENIGI | 21 | 0.78 | 1.46 | 0.02 | 0.099 |
| 642 | REACTOME_NONHOMOLOGOUS_END_JOINING_NI | 50 | 0.68 | 1.46 | 0.008 | 0.099 |
| 643 | SIMBULAN_PARP1_TARGETS_UP | 23 | 0.77 | 1.46 | 0.028 | 0.099 |
| 644 | WP_ONCOSTATIN_M_SIGNALING_PATHWAY | 58 | 0.67 | 1.46 | 0.01 | 0.099 |
| 645 | PID_IL27_PATHWAY | 15 | 0.82 | 1.45 | 0.023 | 0.1 |
| 646 | AP1FJ_Q2 | 218 | 0.61 | 1.45 | 0 | 0.1 |
| 647 | ROZANOV_MMP14_TARGETS_UP | 207 | 0.61 | 1.45 | 0 | 0.1 |
| 648 | BIOCARTA_CASPASE_PATHWAY | 18 | 0.8 | 1.45 | 0.02 | 0.1 |
| 649 | REACTOME_TELOMERE_MAINTENANCE | 89 | 0.65 | 1.45 | 0.005 | 0.1 |
| 650 | BAKER_HEMATOPOIESIS_STAT3_TARGETS | 15 | 0.83 | 1.45 | 0.022 | 0.1 |
| 651 | MIR1322 | 62 | 0.67 | 1.45 | 0.018 | 0.1 |
| 652 | LEE_LIVER_CANCER_E2F1_UP | 46 | 0.7 | 1.45 | 0.01 | 0.1 |
| 653 | BOYAULT_LIVER_CANCER_SUBCLASS_G5_DN | 21 | 0.78 | 1.45 | 0.014 | 0.1 |
| 654 | REACTOME_BASE_EXCISION_REPAIR | 66 | 0.67 | 1.45 | 0.007 | 0.1 |
| 655 | SHAFFER_IRF4_TARGETS_IN_PLASMA_CELL_VS_ | 61 | 0.67 | 1.45 | 0.009 | 0.1 |
| 656 | LY_AGING_PREMATURE_DN | 25 | 0.77 | 1.45 | 0.026 | 0.1 |
| 657 | BASSO_B_LYMPHOCYTE_NETWORK | 140 | 0.62 | 1.45 | 0.001 | 0.1 |

| | | | | | | |
|---|---|---|---|---|---|---|
| 658 | DELACROIX_RAR_TARGETS_UP | 37 | 0.71 | 1.45 | 0.02 | 0.101 |
| 659 | XU_HGF_TARGETS_INDUCED_BY_AKT1_6HR | 18 | 0.79 | 1.45 | 0.028 | 0.101 |
| 660 | REACTOME_ELASTIC_FIBRE_FORMATION | 31 | 0.73 | 1.45 | 0.023 | 0.102 |
| 661 | TAVOR_CEBPA_TARGETS_DN | 23 | 0.77 | 1.45 | 0.024 | 0.102 |
| 662 | WANG_SMARCE1_TARGETS_DN | 329 | 0.6 | 1.45 | 0 | 0.102 |
| 663 | MARKS_HDAC_TARGETS_UP | 20 | 0.79 | 1.45 | 0.015 | 0.102 |
| 664 | REACTOME_TICAM1_RIP1_MEDIATED_IKK_COMPL | 18 | 0.8 | 1.45 | 0.027 | 0.102 |
| 665 | SA_TRKA_RECEPTOR | 16 | 0.83 | 1.45 | 0.016 | 0.102 |
| 666 | AZARE_STAT3_TARGETS | 20 | 0.78 | 1.45 | 0.034 | 0.102 |
| 667 | MARSON_FOXP3_TARGETS_DN | 40 | 0.7 | 1.45 | 0.018 | 0.103 |
| 668 | VANASSE_BCL2_TARGETS_UP | 22 | 0.78 | 1.45 | 0.03 | 0.103 |
| 669 | ANDERSEN_CHOLANGIOCARCINOMA_CLASS2 | 112 | 0.63 | 1.45 | 0.006 | 0.103 |
| 670 | PEDERSEN_METASTASIS_BY_ERBB2_ISOFORM_3 | 15 | 0.82 | 1.45 | 0.022 | 0.104 |
| 671 | PELLICCIOTTA_HDAC_IN_ANTIGEN_PRESENTATIC | 49 | 0.69 | 1.45 | 0.013 | 0.104 |
| 672 | REACTOME_COMPLEMENT_CASCADE | 20 | 0.78 | 1.45 | 0.032 | 0.104 |
| 673 | REACTOME_VEGFR2_MEDIATED_VASCULAR_PER | 27 | 0.74 | 1.45 | 0.024 | 0.105 |
| 674 | SABATES_COLORECTAL_ADENOMA_UP | 71 | 0.66 | 1.45 | 0.017 | 0.105 |
| 675 | REACTOME_BIOLOGICAL_OXIDATIONS | 120 | 0.63 | 1.45 | 0.006 | 0.106 |
| 676 | MIR4520_2_3P | 101 | 0.63 | 1.45 | 0.005 | 0.106 |
| 677 | YU_MYC_TARGETS_DN | 47 | 0.69 | 1.45 | 0.018 | 0.106 |
| 678 | KOBAYASHI_EGFR_SIGNALING_24HR_UP | 82 | 0.65 | 1.45 | 0.011 | 0.106 |
| 679 | DAZARD_RESPONSE_TO_UV_SCC_UP | 92 | 0.64 | 1.45 | 0.009 | 0.107 |
| 680 | SHAFFER_IRF4_TARGETS_IN_ACTIVATED_DENDRI | 59 | 0.68 | 1.45 | 0.008 | 0.107 |
| 681 | MIR3174 | 37 | 0.7 | 1.45 | 0.023 | 0.107 |
| 682 | REACTOME_FLT3_SIGNALING_IN_DISEASE | 27 | 0.75 | 1.45 | 0.014 | 0.107 |
| 683 | WP_IL6_SIGNALING_PATHWAY | 39 | 0.71 | 1.45 | 0.011 | 0.107 |
| 684 | REACTOME_INSERTION_OF_TAIL_ANCHORED_PR | 22 | 0.76 | 1.45 | 0.025 | 0.107 |
| 685 | REACTOME_EXTRACELLULAR_MATRIX_ORGANIZA | 192 | 0.61 | 1.45 | 0.004 | 0.107 |
| 686 | GHO_ATF5_TARGETS_DN | 16 | 0.81 | 1.45 | 0.03 | 0.108 |
| 687 | PID_PI3K_PLC_TRK_PATHWAY | 34 | 0.72 | 1.45 | 0.02 | 0.107 |
| 688 | GOTTWEIN_TARGETS_OF_KSHV_MIR_K12_11 | 57 | 0.67 | 1.45 | 0.011 | 0.107 |
| 689 | SATO_SILENCED_BY_METHYLATION_IN_PANCREA | 287 | 0.6 | 1.44 | 0 | 0.108 |
| 690 | MIR6834_3P | 39 | 0.71 | 1.44 | 0.027 | 0.108 |

| | | | | | | |
|---|---|---|---|---|---|---|
| 691 | KIM_ALL_DISORDERS_OLIGODENDROCYTE_NUMB | 20 | 0.78 | 1.44 | 0.013 | 0.109 |
| 692 | KEGG_NATURAL_KILLER_CELL_MEDIATED_CYTOT | 79 | 0.66 | 1.44 | 0.008 | 0.109 |
| 693 | REACTOME_TRAF6_MEDIATED_IRF7_ACTIVATION | 17 | 0.79 | 1.44 | 0.025 | 0.109 |
| 694 | STEARMAN_TUMOR_FIELD_EFFECT_UP | 24 | 0.76 | 1.44 | 0.027 | 0.109 |
| 695 | MIR7158_3P | 48 | 0.68 | 1.44 | 0.023 | 0.109 |
| 696 | PID_P73PATHWAY | 71 | 0.66 | 1.44 | 0.011 | 0.109 |
| 697 | BLANCO_MELO_INFLUENZA_A_INFECTION_A594_ | 34 | 0.72 | 1.44 | 0.024 | 0.109 |
| 698 | KRAS.KIDNEY_UP.V1_DN | 70 | 0.66 | 1.44 | 0.012 | 0.109 |
| 699 | WP_TYROBP_CAUSAL_NETWORK_IN_MICROGLIA | 35 | 0.72 | 1.44 | 0.012 | 0.109 |
| 700 | KYNG_DNA_DAMAGE_BY_GAMMA_RADIATION | 61 | 0.66 | 1.44 | 0.027 | 0.109 |
| 701 | ERBB2_UP.V1_DN | 178 | 0.61 | 1.44 | 0 | 0.109 |
| 702 | SENGUPTA_NASOPHARYNGEAL_CARCINOMA_WIT | 321 | 0.6 | 1.44 | 0.001 | 0.11 |
| 703 | ZHANG_TARGETS_OF_EWSR1_FLI1_FUSION | 74 | 0.65 | 1.44 | 0.015 | 0.11 |
| 704 | ATF2_UP.V1_UP | 119 | 0.63 | 1.44 | 0.006 | 0.11 |
| 705 | MIR193A_5P | 33 | 0.72 | 1.44 | 0.025 | 0.11 |
| 706 | VILIMAS_NOTCH1_TARGETS_UP | 29 | 0.74 | 1.44 | 0.015 | 0.11 |
| 707 | AZARE_NEOPLASTIC_TRANSFORMATION_BY_STA | 95 | 0.63 | 1.44 | 0.008 | 0.111 |
| 708 | VERHAAK_AML_WITH_NPM1_MUTATED_DN | 177 | 0.61 | 1.44 | 0 | 0.111 |
| 709 | WP_IMATINIB_AND_CHRONIC_MYELOID_LEUKEMI | 15 | 0.82 | 1.44 | 0.019 | 0.111 |
| 710 | DE_YY1_TARGETS_DN | 82 | 0.65 | 1.44 | 0.008 | 0.111 |
| 711 | SMIRNOV_RESPONSE_TO_IR_6HR_UP | 152 | 0.61 | 1.44 | 0.003 | 0.111 |
| 712 | WP_G1_TO_S_CELL_CYCLE_CONTROL | 64 | 0.66 | 1.44 | 0.012 | 0.112 |
| 713 | ONDER_CDH1_TARGETS_2_DN | 339 | 0.59 | 1.44 | 0.001 | 0.112 |
| 714 | KAAB_FAILED_HEART_VENTRICLE_DN | 37 | 0.71 | 1.44 | 0.033 | 0.114 |
| 715 | STAT3_01 | 19 | 0.77 | 1.44 | 0.035 | 0.115 |
| 716 | REACTOME_SIGNALING_BY_PTK6 | 51 | 0.68 | 1.44 | 0.025 | 0.115 |
| 717 | REACTOME_SIGNALING_BY_NOTCH | 194 | 0.61 | 1.44 | 0.001 | 0.115 |
| 718 | VART_KSHV_INFECTION_ANGIOGENIC_MARKERS_ | 106 | 0.63 | 1.44 | 0.006 | 0.115 |
| 719 | PAL_PRMT5_TARGETS_DN | 21 | 0.77 | 1.44 | 0.028 | 0.115 |
| 720 | WESTON_VEGFA_TARGETS | 63 | 0.67 | 1.44 | 0.015 | 0.115 |
| 721 | CEBPE_TARGET_GENES | 21 | 0.77 | 1.44 | 0.036 | 0.115 |
| 722 | WIERENGA_STAT5A_TARGETS_GROUP1 | 102 | 0.63 | 1.44 | 0.007 | 0.115 |
| 723 | ALONSO_METASTASIS_DN | 16 | 0.81 | 1.44 | 0.022 | 0.115 |

| | | | | | | |
|---|---|---|---|---|---|---|
| 724 | DESERT_EXTRACELLULAR_MATRIX_HEPATOCELL | 15 | 0.8 | 1.44 | 0.032 | 0.115 |
| 725 | KANNAN_TP53_TARGETS_DN | 20 | 0.79 | 1.44 | 0.023 | 0.115 |
| 726 | KORKOLA_EMBRYONAL_CARCINOMA_UP | 35 | 0.73 | 1.44 | 0.023 | 0.115 |
| 727 | WP_ACUTE_VIRAL_MYOCARDITIS | 72 | 0.65 | 1.44 | 0.012 | 0.116 |
| 728 | KRAS.LUNG.BREAST_UP.V1_UP | 70 | 0.66 | 1.44 | 0.023 | 0.116 |
| 729 | WP_IL4_SIGNALING_PATHWAY | 50 | 0.67 | 1.44 | 0.021 | 0.116 |
| 730 | MIR3085_5P | 33 | 0.73 | 1.44 | 0.025 | 0.116 |
| 731 | TONKS_TARGETS_OF_RUNX1_RUNX1T1_FUSION_ | 175 | 0.61 | 1.44 | 0.002 | 0.116 |
| 732 | BIOCARTA_NFKB_PATHWAY | 19 | 0.77 | 1.44 | 0.037 | 0.116 |
| 733 | WP_MELATONIN_METABOLISM_AND_EFFECTS | 28 | 0.74 | 1.44 | 0.025 | 0.117 |
| 734 | IZADPANAH_STEM_CELL_ADIPOSE_VS_BONE_DN | 72 | 0.66 | 1.44 | 0.01 | 0.117 |
| 735 | MIR4633_3P_MIR6500_5P | 34 | 0.72 | 1.44 | 0.021 | 0.117 |
| 736 | BAELDE_DIABETIC_NEPHROPATHY_DN | 378 | 0.59 | 1.44 | 0 | 0.116 |
| 737 | WP_EXTRACELLULAR_VESICLEMEDIATED_SIGNAI | 25 | 0.76 | 1.44 | 0.022 | 0.116 |
| 738 | ICSBP_Q6 | 199 | 0.61 | 1.44 | 0.002 | 0.117 |
| 739 | GROSS_HYPOXIA_VIA_ELK3_DN | 130 | 0.62 | 1.43 | 0.002 | 0.117 |
| 740 | MIR6742_3P | 85 | 0.65 | 1.43 | 0.013 | 0.117 |
| 741 | KYNG_RESPONSE_TO_H2O2 | 59 | 0.66 | 1.43 | 0.012 | 0.117 |
| 742 | LEE_RECENT_THYMIC_EMIGRANT | 202 | 0.6 | 1.43 | 0.004 | 0.117 |
| 743 | REACTOME_DNA_REPLICATION_PRE_INITIATION | 114 | 0.62 | 1.43 | 0.009 | 0.117 |
| 744 | MIR1251_3P | 139 | 0.61 | 1.43 | 0.008 | 0.117 |
| 745 | JOHNSTONE_PARVB_TARGETS_3_UP | 396 | 0.59 | 1.43 | 0 | 0.118 |
| 746 | REACTOME_RNA_POLYMERASE_I_TRANSCRIPTIO | 80 | 0.65 | 1.43 | 0.011 | 0.118 |
| 747 | RASHI_RESPONSE_TO_IONIZING_RADIATION_6 | 57 | 0.67 | 1.43 | 0.022 | 0.117 |
| 748 | AP1_C | 213 | 0.6 | 1.43 | 0.001 | 0.118 |
| 749 | TAKEDA_TARGETS_OF_NUP98_HOXA9_FUSION_1 | 84 | 0.65 | 1.43 | 0.013 | 0.118 |
| 750 | GOLUB_ALL_VS_AML_DN | 17 | 0.8 | 1.43 | 0.021 | 0.118 |
| 751 | BOGNI_TREATMENT_RELATED_MYELOID_LEUKEM | 31 | 0.73 | 1.43 | 0.029 | 0.119 |
| 752 | SIG_PIP3_SIGNALING_IN_CARDIAC_MYOCTES | 63 | 0.66 | 1.43 | 0.019 | 0.119 |
| 753 | MIR645 | 37 | 0.71 | 1.43 | 0.02 | 0.12 |
| 754 | YU_BAP1_TARGETS | 25 | 0.76 | 1.43 | 0.022 | 0.121 |
| 755 | MIR3606_5P | 117 | 0.62 | 1.43 | 0.009 | 0.121 |
| 756 | THEILGAARD_NEUTROPHIL_AT_SKIN_WOUND_UP | 63 | 0.66 | 1.43 | 0.02 | 0.122 |

| | | | | | | |
|---|---|---|---|---|---|---|
| 757 | PID_IL23_PATHWAY | 21 | 0.78 | 1.43 | 0.029 | 0.122 |
| 758 | FLECHNER_PBL_KIDNEY_TRANSPLANT_REJECTEI | 49 | 0.68 | 1.43 | 0.02 | 0.122 |
| 759 | WP_NUCLEAR_RECEPTORS_METAPATHWAY | 214 | 0.6 | 1.43 | 0.003 | 0.122 |
| 760 | KIM_MYCN_AMPLIFICATION_TARGETS_DN | 96 | 0.63 | 1.43 | 0.013 | 0.122 |
| 761 | FLECHNER_PBL_KIDNEY_TRANSPLANT_OK_VS_D | 33 | 0.73 | 1.43 | 0.017 | 0.122 |
| 762 | DARWICHE_SKIN_TUMOR_PROMOTER_UP | 109 | 0.63 | 1.43 | 0.007 | 0.122 |
| 763 | STK33_NOMO_UP | 222 | 0.6 | 1.43 | 0 | 0.122 |
| 764 | AMIT_SERUM_RESPONSE_40_MCF10A | 26 | 0.74 | 1.43 | 0.034 | 0.122 |
| 765 | PEDRIOLI_MIR31_TARGETS_DN | 307 | 0.59 | 1.43 | 0 | 0.123 |
| 766 | MTOR_UP.V1_UP | 129 | 0.61 | 1.43 | 0.003 | 0.123 |
| 767 | REACTOME_TRANSCRIPTIONAL_REGULATION_OF | 55 | 0.67 | 1.43 | 0.022 | 0.123 |
| 768 | PID_DNA_PK_PATHWAY | 15 | 0.79 | 1.43 | 0.032 | 0.123 |
| 769 | DASU_IL6_SIGNALING_UP | 55 | 0.67 | 1.43 | 0.023 | 0.123 |
| 770 | KAYO_CALORIE_RESTRICTION_MUSCLE_DN | 69 | 0.66 | 1.43 | 0.014 | 0.123 |
| 771 | KORKOLA_YOLK_SAC_TUMOR_UP | 20 | 0.77 | 1.43 | 0.033 | 0.124 |
| 772 | RUTELLA_RESPONSE_TO_HGF_DN | 196 | 0.6 | 1.43 | 0.001 | 0.124 |
| 773 | HELLER_HDAC_TARGETS_DN | 236 | 0.59 | 1.43 | 0.003 | 0.124 |
| 774 | ONDER_CDH1_SIGNALING_VIA_CTNNB1 | 66 | 0.66 | 1.43 | 0.018 | 0.125 |
| 775 | FLECHNER_BIOPSY_KIDNEY_TRANSPLANT_REJE( | 54 | 0.66 | 1.43 | 0.028 | 0.125 |
| 776 | COULOUARN_TEMPORAL_TGFB1_SIGNATURE_DN | 114 | 0.62 | 1.43 | 0.015 | 0.125 |
| 777 | MANNE_COVID19_COMBINED_COHORT_VS_HEAL | 146 | 0.61 | 1.43 | 0.004 | 0.125 |
| 778 | WILENSKY_RESPONSE_TO_DARAPLADIB | 16 | 0.82 | 1.43 | 0.02 | 0.125 |
| 779 | FIGUEROA_AML_METHYLATION_CLUSTER_3_DN | 30 | 0.72 | 1.43 | 0.033 | 0.125 |
| 780 | BIOCARTA_41BB_PATHWAY | 15 | 0.84 | 1.43 | 0.013 | 0.125 |
| 781 | SENESE_HDAC2_TARGETS_UP | 89 | 0.64 | 1.43 | 0.009 | 0.126 |
| 782 | REACTOME_GROWTH_HORMONE_RECEPTOR_SIG | 19 | 0.77 | 1.43 | 0.023 | 0.126 |
| 783 | KAYO_AGING_MUSCLE_DN | 108 | 0.62 | 1.43 | 0.008 | 0.126 |
| 784 | REACTOME_MEIOSIS | 79 | 0.65 | 1.43 | 0.013 | 0.127 |
| 785 | REACTOME_ESTROGEN_DEPENDENT_NUCLEAR_I | 21 | 0.75 | 1.43 | 0.036 | 0.127 |
| 786 | WANG_CISPLATIN_RESPONSE_AND_XPC_UP | 139 | 0.62 | 1.43 | 0.005 | 0.127 |
| 787 | MISHRA_CARCINOMA_ASSOCIATED_FIBROBLAST_ | 17 | 0.8 | 1.43 | 0.037 | 0.127 |
| 788 | REACTOME_ATF4_ACTIVATES_GENES_IN_RESPO | 26 | 0.74 | 1.43 | 0.029 | 0.126 |
| 789 | TERF1_TARGET_GENES | 205 | 0.6 | 1.43 | 0.002 | 0.127 |

| | | | | | | |
|---|---|---|---|---|---|---|
| 790 | REACTOME_HCMV_EARLY_EVENTS | 98 | 0.63 | 1.43 | 0.012 | 0.126 |
| 791 | MIR11181_5P | 61 | 0.65 | 1.42 | 0.016 | 0.127 |
| 792 | WP_CAMKK2_PATHWAY | 35 | 0.71 | 1.42 | 0.027 | 0.127 |
| 793 | MIR146B_5P | 91 | 0.63 | 1.42 | 0.015 | 0.127 |
| 794 | BIOCARTA_CDMAC_PATHWAY | 16 | 0.81 | 1.42 | 0.036 | 0.127 |
| 795 | REACTOME_ANTIGEN_PROCESSING_CROSS_PRE | 88 | 0.63 | 1.42 | 0.022 | 0.128 |
| 796 | GALE_APL_WITH_FLT3_MUTATED_UP | 56 | 0.67 | 1.42 | 0.028 | 0.128 |
| 797 | BIOCARTA_EGF_PATHWAY | 27 | 0.74 | 1.42 | 0.028 | 0.128 |
| 798 | YAGI_AML_WITH_T_9_11_TRANSLOCATION | 111 | 0.62 | 1.42 | 0.01 | 0.129 |
| 799 | PID_DELTA_NP63_PATHWAY | 43 | 0.68 | 1.42 | 0.032 | 0.129 |
| 800 | LINDVALL_IMMORTALIZED_BY_TERT_DN | 55 | 0.66 | 1.42 | 0.034 | 0.129 |
| 801 | WP_PI3KAKT_SIGNALING_PATHWAY | 230 | 0.59 | 1.42 | 0 | 0.129 |
| 802 | MIR4727_3P | 28 | 0.73 | 1.42 | 0.038 | 0.129 |
| 803 | FOSTER_KDM1A_TARGETS_UP | 141 | 0.61 | 1.42 | 0.008 | 0.13 |
| 804 | PID_LYSOPHOSPHOLIPID_PATHWAY | 56 | 0.66 | 1.42 | 0.023 | 0.13 |
| 805 | JAZAG_TGFB1_SIGNALING_VIA_SMAD4_UP | 81 | 0.64 | 1.42 | 0.012 | 0.131 |
| 806 | PID_INTEGRIN1_PATHWAY | 42 | 0.69 | 1.42 | 0.033 | 0.131 |
| 807 | WOO_LIVER_CANCER_RECURRENCE_DN | 47 | 0.69 | 1.42 | 0.028 | 0.131 |
| 808 | WP_IL2_SIGNALING_PATHWAY | 40 | 0.69 | 1.42 | 0.028 | 0.131 |
| 809 | REACTOME_DEUBIQUITINATION | 244 | 0.59 | 1.42 | 0 | 0.131 |
| 810 | GROSS_HYPOXIA_VIA_HIF1A_UP | 71 | 0.65 | 1.42 | 0.013 | 0.132 |
| 811 | PID_IL2_1PATHWAY | 51 | 0.67 | 1.42 | 0.033 | 0.132 |
| 812 | WINZEN_DEGRADED_VIA_KHSRP | 77 | 0.64 | 1.42 | 0.02 | 0.132 |
| 813 | REACTOME_HCMV_LATE_EVENTS | 82 | 0.64 | 1.42 | 0.012 | 0.133 |
| 814 | REACTOME_DETOXIFICATION_OF_REACTIVE_OXY | 30 | 0.72 | 1.42 | 0.039 | 0.133 |
| 815 | CHEBOTAEV_GR_TARGETS_UP | 56 | 0.67 | 1.42 | 0.026 | 0.133 |
| 816 | VERNELL_RETINOBLASTOMA_PATHWAY_DN | 22 | 0.77 | 1.42 | 0.028 | 0.134 |
| 817 | REACTOME_RAF_INDEPENDENT_MAPK1_3_ACTIV | 22 | 0.76 | 1.42 | 0.036 | 0.134 |
| 818 | REACTOME_PROCESSING_OF_DNA_DOUBLE_STF | 79 | 0.64 | 1.42 | 0.012 | 0.134 |
| 819 | AMIT_SERUM_RESPONSE_60_MCF10A | 48 | 0.67 | 1.42 | 0.029 | 0.135 |
| 820 | TTCYNRGAA_STAT5B_01 | 257 | 0.59 | 1.42 | 0 | 0.135 |
| 821 | WANG_RESPONSE_TO_GSK3_INHIBITOR_SB2167 | 305 | 0.59 | 1.42 | 0.001 | 0.136 |
| 822 | HOLLERN_EMT_BREAST_TUMOR_UP | 102 | 0.62 | 1.42 | 0.01 | 0.136 |

| | | | | | | |
|---|---|---|---|---|---|---|
| 823 | NEMETH_INFLAMMATORY_RESPONSE_LPS_UP | 68 | 0.65 | 1.42 | 0.018 | 0.136 |
| 824 | CROONQUIST_NRAS_VS_STROMAL_STIMULATION | 77 | 0.64 | 1.42 | 0.014 | 0.136 |
| 825 | REACTOME_WNT5A_DEPENDENT_INTERNALIZATIC | 15 | 0.81 | 1.42 | 0.027 | 0.136 |
| 826 | FERRARI_RESPONSE_TO_FENRETINIDE_UP | 19 | 0.78 | 1.42 | 0.036 | 0.136 |
| 827 | CUI_GLUCOSE_DEPRIVATION | 52 | 0.68 | 1.42 | 0.02 | 0.136 |
| 828 | LI_INDUCED_T_TO_NATURAL_KILLER_UP | 230 | 0.59 | 1.42 | 0.004 | 0.137 |
| 829 | WESTON_VEGFA_TARGETS_12HR | 17 | 0.79 | 1.42 | 0.04 | 0.137 |
| 830 | PID_ALK1_PATHWAY | 20 | 0.76 | 1.42 | 0.037 | 0.137 |
| 831 | CAMP_UP.V1_DN | 176 | 0.6 | 1.42 | 0.003 | 0.137 |
| 832 | REACTOME_SIGNALING_BY_GPCR | 363 | 0.58 | 1.42 | 0.001 | 0.137 |
| 833 | NABA_PROTEOGLYCANS | 15 | 0.8 | 1.42 | 0.039 | 0.137 |
| 834 | THUM_SYSTOLIC_HEART_FAILURE_UP | 305 | 0.59 | 1.42 | 0 | 0.137 |
| 835 | BIOCARTA_ARF_PATHWAY | 15 | 0.79 | 1.42 | 0.048 | 0.137 |
| 836 | WP_TCELL_RECEPTOR_TCR_SIGNALING_PATHWA | 69 | 0.64 | 1.42 | 0.019 | 0.137 |
| 837 | MIKKELSEN_ES_ICP_WITH_H3K4ME3_AND_H3K27 | 76 | 0.64 | 1.41 | 0.014 | 0.14 |
| 838 | YAMASHITA_LIVER_CANCER_STEM_CELL_DN | 32 | 0.72 | 1.41 | 0.036 | 0.14 |
| 839 | BIOCARTA_VIP_PATHWAY | 20 | 0.76 | 1.41 | 0.039 | 0.141 |
| 840 | KEGG_BLADDER_CANCER | 40 | 0.68 | 1.41 | 0.031 | 0.141 |
| 841 | HSIAO_HOUSEKEEPING_GENES | 382 | 0.58 | 1.41 | 0 | 0.141 |
| 842 | CHIANG_LIVER_CANCER_SUBCLASS_CTNNB1_DN | 101 | 0.62 | 1.41 | 0.008 | 0.141 |
| 843 | BILD_MYC_ONCOGENIC_SIGNATURE | 191 | 0.6 | 1.41 | 0 | 0.144 |
| 844 | PID_NFAT_3PATHWAY | 50 | 0.67 | 1.41 | 0.028 | 0.144 |
| 845 | WP_DNA_DAMAGE_RESPONSE_ONLY_ATM_DEPE | 100 | 0.63 | 1.41 | 0.007 | 0.144 |
| 846 | TIAN_TNF_SIGNALING_NOT_VIA_NFKB | 19 | 0.77 | 1.41 | 0.051 | 0.144 |
| 847 | HUPER_BREAST_BASAL_VS_LUMINAL_UP | 27 | 0.73 | 1.41 | 0.04 | 0.145 |
| 848 | HONMA_DOCETAXEL_RESISTANCE | 35 | 0.7 | 1.41 | 0.029 | 0.146 |
| 849 | BORCZUK_MALIGNANT_MESOTHELIOMA_DN | 60 | 0.65 | 1.41 | 0.024 | 0.146 |
| 850 | PLASARI_TGFB1_SIGNALING_VIA_NFIC_1HR_UP | 21 | 0.76 | 1.41 | 0.03 | 0.146 |
| 851 | KYNG_DNA_DAMAGE_BY_GAMMA_AND_UV_RADIA | 58 | 0.66 | 1.41 | 0.032 | 0.147 |
| 852 | BIOCARTA_CCR5_PATHWAY | 15 | 0.8 | 1.41 | 0.033 | 0.147 |
| 853 | REACTOME_CHROMATIN_MODIFYING_ENZYMES | 232 | 0.58 | 1.41 | 0.002 | 0.147 |
| 854 | BOQUEST_STEM_CELL_UP | 149 | 0.61 | 1.41 | 0.004 | 0.147 |
| 855 | SMITH_LIVER_CANCER | 43 | 0.68 | 1.41 | 0.032 | 0.149 |

| | | | | | | |
|---|---|---|---|---|---|---|
| 856 | REACTOME_RAC3_GTPASE_CYCLE | 85 | 0.63 | 1.41 | 0.01 | 0.149 |
| 857 | GAUSSMANN_MLL_AF4_FUSION_TARGETS_F_UP | 134 | 0.61 | 1.41 | 0.011 | 0.15 |
| 858 | MARTENS_BOUND_BY_PML_RARA_FUSION | 383 | 0.58 | 1.41 | 0 | 0.15 |
| 859 | MIR7153_5P | 93 | 0.62 | 1.41 | 0.021 | 0.15 |
| 860 | YIH_RESPONSE_TO_ARSENITE_C1 | 20 | 0.77 | 1.41 | 0.052 | 0.15 |
| 861 | AMIT_EGF_RESPONSE_60_HELA | 44 | 0.68 | 1.41 | 0.027 | 0.15 |
| 862 | REACTOME_RIPK1_MEDIATED_REGULATED_NECF | 26 | 0.73 | 1.41 | 0.038 | 0.15 |
| 863 | REACTOME_PERK_REGULATES_GENE_EXPRESSI | 31 | 0.71 | 1.41 | 0.043 | 0.152 |
| 864 | REACTOME_C_TYPE_LECTIN_RECEPTORS_CLRS | 118 | 0.61 | 1.41 | 0.012 | 0.151 |
| 865 | COLINA_TARGETS_OF_4EBP1_AND_4EBP2 | 291 | 0.58 | 1.41 | 0.001 | 0.152 |
| 866 | MAGRANGEAS_MULTIPLE_MYELOMA_IGG_VS_IG/ | 19 | 0.77 | 1.41 | 0.034 | 0.152 |
| 867 | REACTOME_SURFACTANT_METABOLISM | 18 | 0.77 | 1.4 | 0.049 | 0.152 |
| 868 | BIOCARTA_FAS_PATHWAY | 27 | 0.72 | 1.4 | 0.046 | 0.152 |
| 869 | CONCANNON_APOPTOSIS_BY_EPOXOMICIN_UP | 209 | 0.59 | 1.4 | 0.004 | 0.152 |
| 870 | ESC_V6.5_UP_LATE.V1_DN | 147 | 0.6 | 1.4 | 0.01 | 0.152 |
| 871 | MOHANKUMAR_HOXA1_TARGETS_DN | 110 | 0.61 | 1.4 | 0.015 | 0.152 |
| 872 | CHICAS_RB1_TARGETS_SENESCENT | 465 | 0.57 | 1.4 | 0 | 0.152 |
| 873 | REACTOME_DOWNREGULATION_OF_SMAD2_3_SM | 26 | 0.73 | 1.4 | 0.04 | 0.152 |
| 874 | SCHEIDEREIT_IKK_INTERACTING_PROTEINS | 47 | 0.67 | 1.4 | 0.035 | 0.152 |
| 875 | DORN_ADENOVIRUS_INFECTION_12HR_UP | 23 | 0.73 | 1.4 | 0.05 | 0.152 |
| 876 | BOSCO_EPITHELIAL_DIFFERENTIATION_MODULE | 30 | 0.71 | 1.4 | 0.034 | 0.152 |
| 877 | PLASARI_TGFB1_SIGNALING_VIA_NFIC_10HR_UP | 38 | 0.7 | 1.4 | 0.032 | 0.152 |
| 878 | YIH_RESPONSE_TO_ARSENITE_C4 | 17 | 0.78 | 1.4 | 0.04 | 0.152 |
| 879 | ZHAN_MULTIPLE_MYELOMA_MS_DN | 40 | 0.68 | 1.4 | 0.039 | 0.152 |
| 880 | WP_REGULATORY_CIRCUITS_OF_THE_STAT3_SIG | 59 | 0.65 | 1.4 | 0.034 | 0.152 |
| 881 | TONKS_TARGETS_OF_RUNX1_RUNX1T1_FUSION_ | 30 | 0.71 | 1.4 | 0.04 | 0.152 |
| 882 | RUTELLA_RESPONSE_TO_CSF2RB_AND_IL4_DN | 244 | 0.58 | 1.4 | 0.005 | 0.153 |
| 883 | REACTOME_TRAF3_DEPENDENT_IRF_ACTIVATION | 15 | 0.8 | 1.4 | 0.044 | 0.153 |
| 884 | REACTOME_ECM_PROTEOGLYCANS | 42 | 0.68 | 1.4 | 0.033 | 0.154 |
| 885 | TONKS_TARGETS_OF_RUNX1_RUNX1T1_FUSION_ | 125 | 0.61 | 1.4 | 0.011 | 0.154 |
| 886 | AGGAAGC_MIR5163P | 107 | 0.61 | 1.4 | 0.011 | 0.154 |
| 887 | VANHARANTA_UTERINE_FIBROID_DN | 54 | 0.66 | 1.4 | 0.029 | 0.154 |
| 888 | REACTOME_ASSEMBLY_OF_COLLAGEN_FIBRILS_ | 36 | 0.7 | 1.4 | 0.03 | 0.154 |

| | | | | | | |
|---|---|---|---|---|---|---|
| 889 | WP_LUNG_FIBROSIS | 40 | 0.68 | 1.4 | 0.042 | 0.154 |
| 890 | PECE_MAMMARY_STEM_CELL_UP | 130 | 0.6 | 1.4 | 0.014 | 0.154 |
| 891 | LINDSTEDT_DENDRITIC_CELL_MATURATION_D | 43 | 0.69 | 1.4 | 0.034 | 0.155 |
| 892 | WP_MIRNA_REGULATION_OF_PROSTATE_CANCEI | 37 | 0.69 | 1.4 | 0.036 | 0.155 |
| 893 | WP_TNFALPHA_SIGNALING_PATHWAY | 91 | 0.62 | 1.4 | 0.017 | 0.154 |
| 894 | WP_APOPTOSIS | 77 | 0.63 | 1.4 | 0.02 | 0.155 |
| 895 | REACTOME_MATURATION_OF_SARS_COV_2_NUC | 15 | 0.79 | 1.4 | 0.058 | 0.155 |
| 896 | SMID_BREAST_CANCER_RELAPSE_IN_BONE_DN | 206 | 0.58 | 1.4 | 0.001 | 0.155 |
| 897 | SESTO_RESPONSE_TO_UV_C1 | 63 | 0.64 | 1.4 | 0.02 | 0.155 |
| 898 | AMUNDSON_DNA_DAMAGE_RESPONSE_TP53 | 16 | 0.77 | 1.4 | 0.052 | 0.155 |
| 899 | SCHAVOLT_TARGETS_OF_TP53_AND_TP63 | 15 | 0.8 | 1.4 | 0.039 | 0.156 |
| 900 | WP_ARYL_HYDROCARBON_RECEPTOR_PATHWAY | 40 | 0.69 | 1.4 | 0.027 | 0.156 |
| 901 | WANG_TNF_TARGETS | 18 | 0.78 | 1.4 | 0.038 | 0.157 |
| 902 | GUILLAUMOND_KLF10_TARGETS_UP | 34 | 0.7 | 1.4 | 0.045 | 0.157 |
| 903 | BLANCO_MELO_COVID19_SARS_COV_2_INFECTIC | 61 | 0.64 | 1.4 | 0.028 | 0.157 |
| 904 | TSENG_IRS1_TARGETS_UP | 97 | 0.62 | 1.4 | 0.011 | 0.157 |
| 905 | WESTON_VEGFA_TARGETS_3HR | 45 | 0.67 | 1.4 | 0.044 | 0.157 |
| 906 | IGLESIAS_E2F_TARGETS_UP | 106 | 0.62 | 1.4 | 0.015 | 0.157 |
| 907 | TOMIDA_METASTASIS_UP | 22 | 0.74 | 1.4 | 0.049 | 0.157 |
| 908 | GROSS_HYPOXIA_VIA_ELK3_AND_HIF1A_UP | 128 | 0.6 | 1.4 | 0.012 | 0.157 |
| 909 | ZHANG_PROLIFERATING_VS_QUIESCENT | 44 | 0.67 | 1.4 | 0.028 | 0.157 |
| 910 | RICKMAN_HEAD_AND_NECK_CANCER_A | 75 | 0.64 | 1.4 | 0.017 | 0.159 |
| 911 | PID_NOTCH_PATHWAY | 52 | 0.65 | 1.4 | 0.023 | 0.158 |
| 912 | BROWNE_HCMV_INFECTION_14HR_UP | 140 | 0.6 | 1.4 | 0.007 | 0.159 |
| 913 | MIR4256 | 65 | 0.64 | 1.4 | 0.022 | 0.159 |
| 914 | MIR101_5P | 31 | 0.71 | 1.4 | 0.065 | 0.159 |
| 915 | GOTZMANN_EPITHELIAL_TO_MESENCHYMAL_TRA | 170 | 0.6 | 1.4 | 0.004 | 0.159 |
| 916 | KLEIN_TARGETS_OF_BCR_ABL1_FUSION | 24 | 0.74 | 1.4 | 0.047 | 0.159 |
| 917 | WP_PHOTODYNAMIC_THERAPYINDUCED_AP1_SUI | 46 | 0.67 | 1.4 | 0.04 | 0.159 |
| 918 | ONDER_CDH1_TARGETS_1_DN | 132 | 0.6 | 1.4 | 0.007 | 0.159 |
| 919 | REACTOME_SIGNALING_BY_FLT3_FUSION_PROTE | 19 | 0.76 | 1.4 | 0.06 | 0.159 |
| 920 | GAJATE_RESPONSE_TO_TRABECTEDIN_UP | 56 | 0.66 | 1.4 | 0.029 | 0.159 |
| 921 | CERIBELLI_GENES_INACTIVE_AND_BOUND_BY_N | 25 | 0.74 | 1.4 | 0.043 | 0.159 |

| | | | | | | |
|---|---|---|---|---|---|---|
| 922 | GAUSSMANN_MLL_AF4_FUSION_TARGETS_A_DN | 65 | 0.65 | 1.4 | 0.028 | 0.159 |
| 923 | TTCCGTT_MIR191 | 29 | 0.72 | 1.4 | 0.052 | 0.16 |
| 924 | REACTOME_PIWI_INTERACTING_RNA_PIRNA_BIOG | 20 | 0.74 | 1.4 | 0.049 | 0.161 |
| 925 | MIR6804_5P | 58 | 0.65 | 1.4 | 0.043 | 0.161 |
| 926 | TARTE_PLASMA_CELL_VS_PLASMABLAST_DN | 290 | 0.58 | 1.39 | 0 | 0.162 |
| 927 | KEGG_REGULATION_OF_AUTOPHAGY | 19 | 0.76 | 1.39 | 0.043 | 0.163 |
| 928 | SMID_BREAST_CANCER_BASAL_UP | 483 | 0.57 | 1.39 | 0 | 0.163 |
| 929 | ZNF490_TARGET_GENES | 258 | 0.58 | 1.39 | 0.001 | 0.163 |
| 930 | NICK_RESPONSE_TO_PROC_TREATMENT_DN | 26 | 0.73 | 1.39 | 0.043 | 0.163 |
| 931 | GARGALOVIC_RESPONSE_TO_OXIDIZED_PHOSPH | 76 | 0.63 | 1.39 | 0.03 | 0.164 |
| 932 | BURTON_ADIPOGENESIS_9 | 67 | 0.64 | 1.39 | 0.031 | 0.164 |
| 933 | WP_OVERVIEW_OF_INTERFERONSMEDIATED_SIG | 17 | 0.77 | 1.39 | 0.06 | 0.164 |
| 934 | BIOCARTA_IL6_PATHWAY | 21 | 0.75 | 1.39 | 0.048 | 0.164 |
| 935 | MIR3674 | 102 | 0.61 | 1.39 | 0.015 | 0.165 |
| 936 | REACTOME_CHONDROITIN_SULFATE_BIOSYNTHE | 17 | 0.77 | 1.39 | 0.064 | 0.165 |
| 937 | CYCLIN_D1_UP.V1_DN | 141 | 0.59 | 1.39 | 0.012 | 0.166 |
| 938 | MIR203A_5P | 54 | 0.66 | 1.39 | 0.029 | 0.166 |
| 939 | WNT_UP.V1_DN | 114 | 0.6 | 1.39 | 0.013 | 0.167 |
| 940 | BROWNE_HCMV_INFECTION_4HR_UP | 39 | 0.69 | 1.39 | 0.043 | 0.167 |
| 941 | LEE_LIVER_CANCER_MYC_E2F1_DN | 28 | 0.72 | 1.39 | 0.04 | 0.167 |
| 942 | LIU_SOX4_TARGETS_UP | 126 | 0.6 | 1.39 | 0.016 | 0.167 |
| 943 | LANDIS_ERBB2_BREAST_TUMORS_324_DN | 112 | 0.6 | 1.39 | 0.011 | 0.168 |
| 944 | REACTOME_MITOTIC_PROPHASE | 111 | 0.6 | 1.39 | 0.01 | 0.168 |
| 945 | WANG_TARGETS_OF_MLL_CBP_FUSION_UP | 35 | 0.69 | 1.39 | 0.046 | 0.168 |
| 946 | TAVOR_CEBPA_TARGETS_UP | 37 | 0.69 | 1.39 | 0.048 | 0.169 |
| 947 | DANG_MYC_TARGETS_DN | 30 | 0.72 | 1.39 | 0.047 | 0.168 |
| 948 | ATF2_UP.V1_DN | 127 | 0.6 | 1.39 | 0.017 | 0.169 |
| 949 | SHAFFER_IRF4_MULTIPLE_MYELOMA_PROGRAM | 31 | 0.7 | 1.39 | 0.056 | 0.169 |
| 950 | PID_INTEGRIN_A9B1_PATHWAY | 17 | 0.76 | 1.39 | 0.062 | 0.169 |
| 951 | WP_ADIPOGENESIS | 100 | 0.61 | 1.39 | 0.016 | 0.169 |
| 952 | REACTOME_INTERLEUKIN_20_FAMILY_SIGNALING | 17 | 0.78 | 1.39 | 0.051 | 0.17 |
| 953 | REACTOME_SIGNALING_BY_ALK_IN_CANCER | 54 | 0.66 | 1.39 | 0.03 | 0.17 |
| 954 | REACTOME_BMAL1_CLOCK_NPAS2_ACTIVATES_C | 25 | 0.72 | 1.39 | 0.057 | 0.17 |

| | | | | | | |
|---|---|---|---|---|---|---|
| 955 | IIZUKA_LIVER_CANCER_PROGRESSION_L0_L1_DI | 16 | 0.77 | 1.39 | 0.068 | 0.17 |
| 956 | REACTOME_SEMAPHORIN_INTERACTIONS | 57 | 0.65 | 1.39 | 0.035 | 0.17 |
| 957 | NADLER_OBESITY_UP | 43 | 0.66 | 1.39 | 0.039 | 0.171 |
| 958 | MIR513B_3P | 136 | 0.6 | 1.39 | 0.012 | 0.172 |
| 959 | REACTOME_DECTIN_1_MEDIATED_NONCANONICA | 59 | 0.65 | 1.39 | 0.032 | 0.172 |
| 960 | MIR4443 | 138 | 0.59 | 1.39 | 0.008 | 0.172 |
| 961 | HIRSCH_CELLULAR_TRANSFORMATION_SIGNATUI | 211 | 0.58 | 1.39 | 0.005 | 0.172 |
| 962 | SENESE_HDAC1_TARGETS_UP | 373 | 0.57 | 1.39 | 0 | 0.172 |
| 963 | MIR6507_5P | 283 | 0.57 | 1.39 | 0.001 | 0.172 |
| 964 | REACTOME_PYROPTOSIS | 21 | 0.74 | 1.39 | 0.052 | 0.172 |
| 965 | STAT5A_01 | 196 | 0.58 | 1.39 | 0.006 | 0.172 |
| 966 | REACTOME_METABOLIC_DISORDERS_OF_BIOLOC | 22 | 0.74 | 1.39 | 0.048 | 0.172 |
| 967 | WP_7Q1123_COPY_NUMBER_VARIATION_SYNDRC | 97 | 0.61 | 1.39 | 0.018 | 0.174 |
| 968 | REACTOME_TP53_REGULATES_TRANSCRIPTION_ | 20 | 0.75 | 1.39 | 0.046 | 0.174 |
| 969 | BARIS_THYROID_CANCER_DN | 52 | 0.66 | 1.39 | 0.027 | 0.175 |
| 970 | BACH1_01 | 209 | 0.58 | 1.38 | 0.001 | 0.175 |
| 971 | MIR146A_5P | 92 | 0.62 | 1.38 | 0.019 | 0.176 |
| 972 | WU_CELL_MIGRATION | 130 | 0.6 | 1.38 | 0.013 | 0.176 |
| 973 | MIR92A_1_5P | 39 | 0.68 | 1.38 | 0.035 | 0.176 |
| 974 | SCHOEN_NFKB_SIGNALING | 27 | 0.72 | 1.38 | 0.045 | 0.176 |
| 975 | AACATTC_MIR4093P | 124 | 0.6 | 1.38 | 0.009 | 0.177 |
| 976 | PID_CD40_PATHWAY | 28 | 0.71 | 1.38 | 0.052 | 0.177 |
| 977 | KAMMINGA_EZH2_TARGETS | 41 | 0.66 | 1.38 | 0.046 | 0.177 |
| 978 | HOLLMANN_APOPTOSIS_VIA_CD40_DN | 233 | 0.58 | 1.38 | 0.004 | 0.177 |
| 979 | KIM_PTEN_TARGETS_UP | 15 | 0.8 | 1.38 | 0.039 | 0.178 |
| 980 | MILI_PSEUDOPODIA | 46 | 0.67 | 1.38 | 0.044 | 0.18 |
| 981 | AMIT_EGF_RESPONSE_120_HELA | 57 | 0.64 | 1.38 | 0.029 | 0.181 |
| 982 | SA_CASPASE_CASCADE | 15 | 0.78 | 1.38 | 0.07 | 0.181 |
| 983 | GROSS_ELK3_TARGETS_DN | 28 | 0.72 | 1.38 | 0.059 | 0.181 |
| 984 | SWEET_KRAS_TARGETS_UP | 69 | 0.63 | 1.38 | 0.03 | 0.181 |
| 985 | WP_GASTRIC_CANCER_NETWORK_1 | 27 | 0.71 | 1.38 | 0.056 | 0.181 |
| 986 | HOFFMANN_PRE_BI_TO_LARGE_PRE_BII_LYMPHC | 22 | 0.73 | 1.38 | 0.061 | 0.182 |
| 987 | AP1_Q2 | 210 | 0.58 | 1.38 | 0.002 | 0.182 |

| | | | | | | |
|---|---|---|---|---|---|---|
| 988 | PID_IL8_CXCR2_PATHWAY | 28 | 0.7 | 1.38 | 0.057 | 0.182 |
| 989 | DARWICHE_PAPILLOMA_RISK_HIGH_UP | 109 | 0.61 | 1.38 | 0.011 | 0.183 |
| 990 | NIKOLSKY_BREAST_CANCER_22Q13_AMPLICON | 16 | 0.76 | 1.38 | 0.063 | 0.184 |
| 991 | WP_ENDOTHELIN_PATHWAYS | 22 | 0.74 | 1.38 | 0.059 | 0.184 |
| 992 | REACTOME_TRANSCRIPTIONAL_ACTIVITY_OF_SM | 44 | 0.67 | 1.38 | 0.036 | 0.185 |
| 993 | TARTE_PLASMA_CELL_VS_B_LYMPHOCYTE_DN | 30 | 0.69 | 1.38 | 0.065 | 0.185 |
| 994 | BIOCARTA_GLEEVEC_PATHWAY | 22 | 0.75 | 1.38 | 0.055 | 0.185 |
| 995 | KEGG_ALDOSTERONE_REGULATED_SODIUM_REA | 31 | 0.69 | 1.38 | 0.055 | 0.185 |
| 996 | MIR4473 | 80 | 0.62 | 1.38 | 0.017 | 0.187 |
| 997 | PID_S1P_S1P2_PATHWAY | 23 | 0.72 | 1.38 | 0.056 | 0.188 |
| 998 | BRUECKNER_TARGETS_OF_MIRLET7A3_DN | 58 | 0.64 | 1.38 | 0.035 | 0.188 |
| 999 | BIOCARTA_PDGF_PATHWAY | 27 | 0.71 | 1.38 | 0.056 | 0.188 |
| 1000 | NABA_CORE_MATRISOME | 150 | 0.59 | 1.38 | 0.008 | 0.188 |
| 1001 | PID_INTEGRIN3_PATHWAY | 27 | 0.7 | 1.38 | 0.06 | 0.188 |
| 1002 | P53_DN.V1_DN | 133 | 0.59 | 1.38 | 0.018 | 0.188 |
| 1003 | REACTOME_NICOTINATE_METABOLISM | 21 | 0.74 | 1.38 | 0.056 | 0.188 |
| 1004 | RIZKI_TUMOR_INVASIVENESS_3D_UP | 164 | 0.58 | 1.38 | 0.011 | 0.188 |
| 1005 | REACTOME_METABOLISM_OF_STEROIDS | 107 | 0.6 | 1.38 | 0.023 | 0.188 |
| 1006 | TAKEDA_TARGETS_OF_NUP98_HOXA9_FUSION_8 | 114 | 0.6 | 1.38 | 0.018 | 0.188 |
| 1007 | LEE_CALORIE_RESTRICTION_MUSCLE_DN | 40 | 0.67 | 1.38 | 0.053 | 0.188 |
| 1008 | FOSTER_TOLERANT_MACROPHAGE_UP | 121 | 0.59 | 1.38 | 0.018 | 0.188 |
| 1009 | HUMMERICH_SKIN_CANCER_PROGRESSION_DN | 83 | 0.61 | 1.38 | 0.041 | 0.188 |
| 1010 | CHIARETTI_T_ALL_RELAPSE_PROGNOSIS | 17 | 0.77 | 1.38 | 0.06 | 0.189 |
| 1011 | WP_NRP1TRIGGERED_SIGNALING_PATHWAYS_IN | 48 | 0.65 | 1.38 | 0.05 | 0.189 |
| 1012 | ADDYA_ERYTHROID_DIFFERENTIATION_BY_HEMIN | 62 | 0.64 | 1.38 | 0.041 | 0.189 |
| 1013 | CHEN_LVAD_SUPPORT_OF_FAILING_HEART_DN | 26 | 0.72 | 1.37 | 0.049 | 0.19 |
| 1014 | MIR3960_MIR8072 | 18 | 0.75 | 1.37 | 0.074 | 0.19 |
| 1015 | MIR4764_3P | 62 | 0.63 | 1.37 | 0.038 | 0.19 |
| 1016 | KAN_RESPONSE_TO_ARSENIC_TRIOXIDE | 100 | 0.61 | 1.37 | 0.027 | 0.19 |
| 1017 | LI_WILMS_TUMOR_VS_FETAL_KIDNEY_1_UP | 156 | 0.58 | 1.37 | 0.008 | 0.19 |
| 1018 | REACTOME_CHROMOSOME_MAINTENANCE | 116 | 0.6 | 1.37 | 0.011 | 0.19 |
| 1019 | JECHLINGER_EPITHELIAL_TO_MESENCHYMAL_TF | 51 | 0.65 | 1.37 | 0.034 | 0.19 |
| 1020 | CGCTGCT_MIR503 | 21 | 0.74 | 1.37 | 0.056 | 0.19 |

| | | | | | | |
|---|---|---|---|---|---|---|
| 1021 | ATF2_S_UP.V1_DN | 120 | 0.6 | 1.37 | 0.02 | 0.19 |
| 1022 | PID_ECADHERIN_STABILIZATION_PATHWAY | 38 | 0.68 | 1.37 | 0.054 | 0.19 |
| 1023 | SARTIPY_NORMAL_AT_INSULIN_RESISTANCE_UP | 27 | 0.71 | 1.37 | 0.066 | 0.191 |
| 1024 | KORKOLA_EMBRYONIC_CARCINOMA_VS_SEMINO | 15 | 0.79 | 1.37 | 0.066 | 0.191 |
| 1025 | LEE_EARLY_T_LYMPHOCYTE_DN | 43 | 0.66 | 1.37 | 0.062 | 0.192 |
| 1026 | KIM_WT1_TARGETS_12HR_DN | 190 | 0.58 | 1.37 | 0.002 | 0.192 |
| 1027 | ONDER_CDH1_TARGETS_3_DN | 29 | 0.7 | 1.37 | 0.062 | 0.192 |
| 1028 | BILD_HRAS_ONCOGENIC_SIGNATURE | 202 | 0.58 | 1.37 | 0.008 | 0.192 |
| 1029 | WP_IL9_SIGNALING_PATHWAY | 15 | 0.78 | 1.37 | 0.06 | 0.194 |
| 1030 | LUI_THYROID_CANCER_PAX8_PPARG_DN | 45 | 0.66 | 1.37 | 0.052 | 0.194 |
| 1031 | DORN_ADENOVIRUS_INFECTION_32HR_DN | 36 | 0.67 | 1.37 | 0.053 | 0.194 |
| 1032 | RUIZ_TNC_TARGETS_UP | 128 | 0.59 | 1.37 | 0.013 | 0.194 |
| 1033 | REACTOME_RND1_GTPASE_CYCLE | 40 | 0.67 | 1.37 | 0.051 | 0.194 |
| 1034 | RIGGINS_TAMOXIFEN_RESISTANCE_DN | 190 | 0.57 | 1.37 | 0.01 | 0.194 |
| 1035 | REACTOME_NEUTROPHIL_DEGRANULATION | 352 | 0.57 | 1.37 | 0.001 | 0.195 |
| 1036 | KEGG_B_CELL_RECEPTOR_SIGNALING_PATHWA' | 62 | 0.64 | 1.37 | 0.032 | 0.195 |
| 1037 | BALDWIN_PRKCI_TARGETS_UP | 34 | 0.69 | 1.37 | 0.052 | 0.195 |
| 1038 | MIR432_3P | 49 | 0.65 | 1.37 | 0.051 | 0.195 |
| 1039 | BAELDE_DIABETIC_NEPHROPATHY_UP | 65 | 0.63 | 1.37 | 0.035 | 0.194 |
| 1040 | HAHTOLA_MYCOSIS_FUNGOIDES_SKIN_DN | 26 | 0.72 | 1.37 | 0.051 | 0.196 |
| 1041 | ANDERSEN_LIVER_CANCER_KRT19_DN | 44 | 0.66 | 1.37 | 0.051 | 0.196 |
| 1042 | HENDRICKS_SMARCA4_TARGETS_DN | 32 | 0.68 | 1.37 | 0.055 | 0.197 |
| 1043 | XU_CREBBP_TARGETS_DN | 31 | 0.69 | 1.37 | 0.061 | 0.197 |
| 1044 | BIOCARTA_TNFR2_PATHWAY | 17 | 0.76 | 1.37 | 0.075 | 0.198 |
| 1045 | KRAS.600.LUNG.BREAST_UP.V1_UP | 144 | 0.59 | 1.37 | 0.008 | 0.198 |
| 1046 | REACTOME_REPRODUCTION | 90 | 0.6 | 1.37 | 0.035 | 0.198 |
| 1047 | CHOW_RASSF1_TARGETS_DN | 30 | 0.69 | 1.37 | 0.07 | 0.198 |
| 1048 | SESTO_RESPONSE_TO_UV_C8 | 64 | 0.63 | 1.37 | 0.048 | 0.198 |
| 1049 | RICKMAN_METASTASIS_DN | 225 | 0.57 | 1.37 | 0.005 | 0.199 |
| 1050 | EGFR_UP.V1_UP | 153 | 0.59 | 1.37 | 0.008 | 0.2 |
| 1051 | ABRAHAM_ALPC_VS_MULTIPLE_MYELOMA_UP | 22 | 0.73 | 1.37 | 0.061 | 0.2 |
| 1052 | REACTOME_TNF_SIGNALING | 44 | 0.65 | 1.37 | 0.057 | 0.2 |
| 1053 | LEE_LIVER_CANCER_E2F1_DN | 29 | 0.69 | 1.37 | 0.067 | 0.2 |

| | | | | | | |
|---|---|---|---|---|---|---|
| 1054 | ENK_UV_RESPONSE_KERATINOCYTE_UP | 457 | 0.56 | 1.37 | 0 | 0.2 |
| 1055 | BOHN_PRIMARY_IMMUNODEFICIENCY_SYNDROM_ | 30 | 0.69 | 1.37 | 0.075 | 0.2 |
| 1056 | CHICAS_RB1_TARGETS_GROWING | 207 | 0.57 | 1.37 | 0.008 | 0.201 |
| 1057 | BERENJENO_ROCK_SIGNALING_NOT_VIA_RHOA_ | 18 | 0.75 | 1.37 | 0.066 | 0.202 |
| 1058 | BEGUM_TARGETS_OF_PAX3_FOXO1_FUSION_DN | 38 | 0.67 | 1.37 | 0.058 | 0.202 |
| 1059 | ROSS_AML_WITH_CBFB_MYH11_FUSION | 32 | 0.69 | 1.37 | 0.051 | 0.202 |
| 1060 | AP1_Q6 | 209 | 0.57 | 1.36 | 0.004 | 0.204 |
| 1061 | MIR4638_3P | 56 | 0.63 | 1.36 | 0.043 | 0.204 |
| 1062 | PID_SMAD2_3NUCLEAR_PATHWAY | 67 | 0.62 | 1.36 | 0.042 | 0.205 |
| 1063 | WP_NEPHROTIC_SYNDROME | 35 | 0.68 | 1.36 | 0.051 | 0.204 |
| 1064 | PID_WNT_SIGNALING_PATHWAY | 20 | 0.74 | 1.36 | 0.067 | 0.204 |
| 1065 | REACTOME_NOD1_2_SIGNALING_PATHWAY | 35 | 0.68 | 1.36 | 0.062 | 0.205 |
| 1066 | CSHL1_TARGET_GENES | 204 | 0.57 | 1.36 | 0.006 | 0.206 |
| 1067 | MEBARKI_HCC_PROGENITOR_FZD8CRD_DN | 258 | 0.56 | 1.36 | 0.002 | 0.206 |
| 1068 | SUH_COEXPRESSED_WITH_ID1_AND_ID2_UP | 16 | 0.77 | 1.36 | 0.069 | 0.206 |
| 1069 | KEGG_CELL_CYCLE | 121 | 0.59 | 1.36 | 0.013 | 0.206 |
| 1070 | TERF2_TARGET_GENES | 98 | 0.6 | 1.36 | 0.025 | 0.206 |
| 1071 | KAUFFMANN_DNA_REPLICATION_GENES | 133 | 0.59 | 1.36 | 0.02 | 0.207 |
| 1072 | DORN_ADENOVIRUS_INFECTION_48HR_DN | 36 | 0.68 | 1.36 | 0.052 | 0.208 |
| 1073 | PID_UPA_UPAR_PATHWAY | 26 | 0.71 | 1.36 | 0.055 | 0.208 |
| 1074 | AR_02 | 34 | 0.68 | 1.36 | 0.065 | 0.209 |
| 1075 | BARRIER_CANCER_RELAPSE_NORMAL_SAMPLE_ | 23 | 0.72 | 1.36 | 0.064 | 0.209 |
| 1076 | ZHOU_INFLAMMATORY_RESPONSE_LIVE_UP | 324 | 0.56 | 1.36 | 0.001 | 0.209 |
| 1077 | MODY_HIPPOCAMPUS_POSTNATAL | 50 | 0.64 | 1.36 | 0.058 | 0.209 |
| 1078 | REACTOME_SIGNALING_BY_FLT3_ITD_AND_TKD_ | 15 | 0.78 | 1.36 | 0.068 | 0.209 |
| 1079 | BLUM_RESPONSE_TO_SALIRASIB_UP | 233 | 0.56 | 1.36 | 0.006 | 0.21 |
| 1080 | LI_WILMS_TUMOR_VS_FETAL_KIDNEY_2_DN | 33 | 0.68 | 1.36 | 0.07 | 0.21 |
| 1081 | MARSON_FOXP3_TARGETS_UP | 60 | 0.64 | 1.36 | 0.056 | 0.21 |
| 1082 | RELA_DN.V1_UP | 104 | 0.6 | 1.36 | 0.03 | 0.21 |
| 1083 | KEGG_PANCREATIC_CANCER | 65 | 0.63 | 1.36 | 0.04 | 0.21 |
| 1084 | JAK2_DN.V1_DN | 124 | 0.59 | 1.36 | 0.022 | 0.211 |
| 1085 | WP_FOCAL_ADHESION_PI3KAKTMTORSIGNALING_ | 219 | 0.57 | 1.36 | 0.007 | 0.211 |
| 1086 | REACTOME_REGULATION_OF_TP53_EXPRESSION | 37 | 0.68 | 1.36 | 0.047 | 0.212 |

| | | | | | | |
|---|---|---|---|---|---|---|
| 1087 | TONG_INTERACT_WITH_PTTG1 | 54 | 0.64 | 1.36 | 0.047 | 0.212 |
| 1088 | MIR6728_5P | 57 | 0.64 | 1.36 | 0.046 | 0.212 |
| 1089 | MIR450A_1_3P | 155 | 0.58 | 1.36 | 0.009 | 0.213 |
| 1090 | SASAKI_ADULT_T_CELL_LEUKEMIA | 149 | 0.58 | 1.36 | 0.014 | 0.213 |
| 1091 | REACTOME_STRIATED_MUSCLE_CONTRACTION | 21 | 0.72 | 1.36 | 0.078 | 0.213 |
| 1092 | SMIRNOV_CIRCULATING_ENDOTHELIOCYTES_IN_( | 117 | 0.6 | 1.36 | 0.031 | 0.215 |
| 1093 | MIR2681_3P | 31 | 0.69 | 1.36 | 0.055 | 0.215 |
| 1094 | KEGG_FOCAL_ADHESION | 157 | 0.58 | 1.36 | 0.008 | 0.215 |
| 1095 | MITSIADES_RESPONSE_TO_APLIDIN_UP | 388 | 0.55 | 1.36 | 0 | 0.215 |
| 1096 | TERAMOTO_OPN_TARGETS_CLUSTER_6 | 21 | 0.72 | 1.36 | 0.071 | 0.215 |
| 1097 | HBZ_TARGET_GENES | 499 | 0.55 | 1.36 | 0 | 0.215 |
| 1098 | SMID_BREAST_CANCER_LUMINAL_A_DN | 17 | 0.76 | 1.36 | 0.072 | 0.215 |
| 1099 | WP_PLURIPOTENT_STEM_CELL_DIFFERENTIATIOI | 22 | 0.72 | 1.36 | 0.071 | 0.215 |
| 1100 | SCHAEFFER_PROSTATE_DEVELOPMENT_6HR_UP | 125 | 0.59 | 1.36 | 0.017 | 0.215 |
| 1101 | MIR3929 | 76 | 0.62 | 1.36 | 0.04 | 0.216 |
| 1102 | KEGG_UBIQUITIN_MEDIATED_PROTEOLYSIS | 129 | 0.58 | 1.36 | 0.02 | 0.216 |
| 1103 | WP_OVARIAN_INFERTILITY | 20 | 0.74 | 1.36 | 0.08 | 0.216 |
| 1104 | REACTOME_PLASMA_LIPOPROTEIN_REMODELINC | 16 | 0.77 | 1.36 | 0.079 | 0.216 |
| 1105 | WALLACE_PROSTATE_CANCER_UP | 19 | 0.73 | 1.36 | 0.083 | 0.216 |
| 1106 | SMID_BREAST_CANCER_RELAPSE_IN_PLEURA_D | 15 | 0.76 | 1.36 | 0.092 | 0.217 |
| 1107 | WP_IL3_SIGNALING_PATHWAY | 40 | 0.65 | 1.36 | 0.076 | 0.217 |
| 1108 | LINDGREN_BLADDER_CANCER_CLUSTER_2A_DN | 116 | 0.59 | 1.35 | 0.022 | 0.219 |
| 1109 | PID_IL8_CXCR1_PATHWAY | 21 | 0.73 | 1.35 | 0.068 | 0.219 |
| 1110 | REACTOME_TOLL_LIKE_RECEPTOR_CASCADES | 129 | 0.58 | 1.35 | 0.029 | 0.219 |
| 1111 | REACTOME_DNA_REPLICATION | 157 | 0.58 | 1.35 | 0.009 | 0.219 |
| 1112 | NOTCH_DN.V1_DN | 107 | 0.59 | 1.35 | 0.024 | 0.219 |
| 1113 | LINDGREN_BLADDER_CANCER_CLUSTER_1_DN | 334 | 0.55 | 1.35 | 0.003 | 0.22 |
| 1114 | PID_GMCSF_PATHWAY | 31 | 0.69 | 1.35 | 0.074 | 0.22 |
| 1115 | TAKAO_RESPONSE_TO_UVB_RADIATION_DN | 93 | 0.59 | 1.35 | 0.03 | 0.22 |
| 1116 | PTTG1_TARGET_GENES | 59 | 0.63 | 1.35 | 0.059 | 0.22 |
| 1117 | WP_FIBRIN_COMPLEMENT_RECEPTOR_3_SIGNAL | 34 | 0.68 | 1.35 | 0.075 | 0.221 |
| 1118 | HTF_01 | 60 | 0.63 | 1.35 | 0.051 | 0.221 |
| 1119 | MIR3188 | 70 | 0.62 | 1.35 | 0.023 | 0.222 |

| | | | | | | |
|---|---|---|---|---|---|---|
| 1120 | KEGG_HEDGEHOG_SIGNALING_PATHWAY | 35 | 0.66 | 1.35 | 0.069 | 0.222 |
| 1121 | MIR7702 | 49 | 0.64 | 1.35 | 0.058 | 0.222 |
| 1122 | AML_Q6 | 196 | 0.57 | 1.35 | 0.005 | 0.223 |
| 1123 | MAF_Q6 | 218 | 0.56 | 1.35 | 0.008 | 0.223 |
| 1124 | REACTOME_RHO_GTPASES_ACTIVATE_NADPH_O | 19 | 0.75 | 1.35 | 0.075 | 0.223 |
| 1125 | REACTOME_EPIGENETIC_REGULATION_OF_GENE | 116 | 0.59 | 1.35 | 0.021 | 0.223 |
| 1126 | BANDRES_RESPONSE_TO_CARMUSTIN_MGMT_48 | 15 | 0.77 | 1.35 | 0.084 | 0.223 |
| 1127 | SASAI_RESISTANCE_TO_NEOPLASTIC_TRANSFRC | 43 | 0.65 | 1.35 | 0.082 | 0.223 |
| 1128 | VALK_AML_CLUSTER_5 | 19 | 0.74 | 1.35 | 0.073 | 0.223 |
| 1129 | ROSS_ACUTE_MYELOID_LEUKEMIA_CBF | 65 | 0.62 | 1.35 | 0.033 | 0.224 |
| 1130 | HERNANDEZ_MITOTIC_ARREST_BY_DOCETAXEL_ | 43 | 0.66 | 1.35 | 0.077 | 0.224 |
| 1131 | WP_OXIDATION_BY_CYTOCHROME_P450 | 23 | 0.72 | 1.35 | 0.076 | 0.224 |
| 1132 | TBX1_TARGET_GENES | 147 | 0.58 | 1.35 | 0.015 | 0.225 |
| 1133 | SCIAN_INVERSED_TARGETS_OF_TP53_AND_TP73 | 28 | 0.69 | 1.35 | 0.057 | 0.225 |
| 1134 | WEST_ADRENOCORTICAL_TUMOR_DN | 424 | 0.55 | 1.35 | 0.001 | 0.225 |
| 1135 | REACTOME_RND2_GTPASE_CYCLE | 41 | 0.66 | 1.35 | 0.059 | 0.225 |
| 1136 | MAGRANGEAS_MULTIPLE_MYELOMA_IGLL_VS_IG | 19 | 0.73 | 1.35 | 0.08 | 0.225 |
| 1137 | AMIT_EGF_RESPONSE_40_MCF10A | 16 | 0.77 | 1.35 | 0.077 | 0.225 |
| 1138 | MIR921 | 50 | 0.65 | 1.35 | 0.071 | 0.225 |
| 1139 | STAT5B_01 | 192 | 0.56 | 1.35 | 0.006 | 0.226 |
| 1140 | PID_ERA_GENOMIC_PATHWAY | 58 | 0.63 | 1.35 | 0.061 | 0.226 |
| 1141 | CSR_LATE_UP.V1_DN | 115 | 0.59 | 1.35 | 0.02 | 0.226 |
| 1142 | LI_WILMS_TUMOR_ANAPLASTIC_UP | 17 | 0.75 | 1.35 | 0.082 | 0.226 |
| 1143 | WP_COMPLEMENT_AND_COAGULATION_CASCADI | 25 | 0.7 | 1.35 | 0.081 | 0.227 |
| 1144 | MILI_PSEUDOPODIA_CHEMOTAXIS_UP | 80 | 0.6 | 1.35 | 0.045 | 0.227 |
| 1145 | FISCHER_DIRECT_P53_TARGETS_META_ANALYSI | 279 | 0.56 | 1.35 | 0.004 | 0.226 |
| 1146 | BURTON_ADIPOGENESIS_PEAK_AT_0HR | 46 | 0.65 | 1.35 | 0.07 | 0.228 |
| 1147 | ALCALAY_AML_BY_NPM1_LOCALIZATION_UP | 114 | 0.58 | 1.35 | 0.029 | 0.228 |
| 1148 | MIR4761_5P | 51 | 0.64 | 1.35 | 0.066 | 0.228 |
| 1149 | MIKKELSEN_IPS_LCP_WITH_H3K4ME3 | 117 | 0.58 | 1.35 | 0.025 | 0.229 |
| 1150 | GAVIN_FOXP3_TARGETS_CLUSTER_P2 | 60 | 0.63 | 1.35 | 0.053 | 0.23 |
| 1151 | CHICAS_RB1_TARGETS_CONFLUENT | 442 | 0.55 | 1.35 | 0 | 0.23 |
| 1152 | DELPUECH_FOXO3_TARGETS_UP | 59 | 0.62 | 1.35 | 0.06 | 0.23 |

| | | | | | | |
|---|---|---|---|---|---|---|
| 1153 | WANG_LSD1_TARGETS_DN | 31 | 0.68 | 1.35 | 0.083 | 0.23 |
| 1154 | LEE_AGING_MUSCLE_DN | 32 | 0.68 | 1.35 | 0.08 | 0.229 |
| 1155 | REACTOME_ACTIVATION_OF_ANTERIOR_HOX_GE | 88 | 0.6 | 1.35 | 0.047 | 0.232 |
| 1156 | KEGG_COMPLEMENT_AND_COAGULATION_CASCA | 26 | 0.69 | 1.35 | 0.088 | 0.232 |
| 1157 | DELASERNA_MYOD_TARGETS_UP | 72 | 0.61 | 1.35 | 0.048 | 0.232 |
| 1158 | TGFB_UP.V1_DN | 156 | 0.57 | 1.35 | 0.014 | 0.232 |
| 1159 | WP_FBXL10_ENHANCEMENT_OF_MAPERK_SIGNA | 25 | 0.7 | 1.35 | 0.091 | 0.232 |
| 1160 | MIR6515_5P | 71 | 0.62 | 1.35 | 0.046 | 0.232 |
| 1161 | BIOCARTA_TEL_PATHWAY | 17 | 0.74 | 1.35 | 0.078 | 0.232 |
| 1162 | ATOH8_TARGET_GENES | 28 | 0.69 | 1.35 | 0.079 | 0.232 |
| 1163 | WANG_CLIM2_TARGETS_DN | 169 | 0.57 | 1.34 | 0.014 | 0.232 |
| 1164 | MIR4712_3P | 44 | 0.65 | 1.34 | 0.076 | 0.234 |
| 1165 | CAFFAREL_RESPONSE_TO_THC_UP | 31 | 0.68 | 1.34 | 0.062 | 0.235 |
| 1166 | HOFMANN_MYELODYSPLASTIC_SYNDROM_LOW_I | 16 | 0.76 | 1.34 | 0.079 | 0.235 |
| 1167 | MIR15A_3P | 28 | 0.68 | 1.34 | 0.089 | 0.236 |
| 1168 | CHUANG_OXIDATIVE_STRESS_RESPONSE_UP | 24 | 0.7 | 1.34 | 0.087 | 0.236 |
| 1169 | KEGG_T_CELL_RECEPTOR_SIGNALING_PATHWAY | 81 | 0.6 | 1.34 | 0.049 | 0.236 |
| 1170 | VALK_AML_CLUSTER_15 | 24 | 0.71 | 1.34 | 0.082 | 0.236 |
| 1171 | NRF2_Q4 | 215 | 0.56 | 1.34 | 0.01 | 0.236 |
| 1172 | KEGG_PATHWAYS_IN_CANCER | 271 | 0.56 | 1.34 | 0.006 | 0.236 |
| 1173 | XU_HGF_TARGETS_REPRESSED_BY_AKT1_DN | 50 | 0.63 | 1.34 | 0.064 | 0.236 |
| 1174 | BHATI_G2M_ARREST_BY_2METHOXYESTRADIOL_ | 99 | 0.59 | 1.34 | 0.036 | 0.236 |
| 1175 | COULOUARN_TEMPORAL_TGFB1_SIGNATURE_UP | 93 | 0.6 | 1.34 | 0.04 | 0.237 |
| 1176 | BRCA1_DN.V1_UP | 57 | 0.63 | 1.34 | 0.055 | 0.237 |
| 1177 | BOYLAN_MULTIPLE_MYELOMA_D_DN | 65 | 0.62 | 1.34 | 0.056 | 0.238 |
| 1178 | KEGG_ENDOMETRIAL_CANCER | 51 | 0.63 | 1.34 | 0.05 | 0.238 |
| 1179 | MIR6822_5P | 55 | 0.63 | 1.34 | 0.048 | 0.238 |
| 1180 | MULLIGHAN_MLL_SIGNATURE_1_DN | 183 | 0.56 | 1.34 | 0.013 | 0.238 |
| 1181 | KOKKINAKIS_METHIONINE_DEPRIVATION_96HR_U | 98 | 0.59 | 1.34 | 0.031 | 0.239 |
| 1182 | WP_RELATIONSHIP_BETWEEN_INFLAMMATION_C( | 16 | 0.74 | 1.34 | 0.097 | 0.239 |
| 1183 | WANG_BARRETTS_ESOPHAGUS_UP | 35 | 0.66 | 1.34 | 0.077 | 0.239 |
| 1184 | CHIBA_RESPONSE_TO_TSA | 43 | 0.65 | 1.34 | 0.039 | 0.239 |
| 1185 | AP3_Q6 | 181 | 0.57 | 1.34 | 0.011 | 0.24 |

| | | | | | | |
|---|---|---|---|---|---|---|
| 1186 | WANG_LMO4_TARGETS_DN | 334 | 0.55 | 1.34 | 0.003 | 0.24 |
| 1187 | WP_IL5_SIGNALING_PATHWAY | 35 | 0.68 | 1.34 | 0.062 | 0.24 |
| 1188 | WP_CELL_CYCLE | 119 | 0.58 | 1.34 | 0.03 | 0.24 |
| 1189 | CROONQUIST_NRAS_SIGNALING_UP | 23 | 0.7 | 1.34 | 0.113 | 0.24 |
| 1190 | REACTOME_PLASMA_LIPOPROTEIN_ASSEMBLY_I | 47 | 0.65 | 1.34 | 0.065 | 0.24 |
| 1191 | MUELLER_METHYLATED_IN_GLIOBLASTOMA | 21 | 0.72 | 1.34 | 0.085 | 0.241 |
| 1192 | MIR6817_3P | 183 | 0.57 | 1.34 | 0.016 | 0.241 |
| 1193 | MCBRYAN_PUBERTAL_BREAST_3_4WK_UP | 156 | 0.57 | 1.34 | 0.015 | 0.241 |
| 1194 | CROMER_METASTASIS_DN | 67 | 0.61 | 1.34 | 0.048 | 0.241 |
| 1195 | REACTOME_TERMINATION_OF_TRANSLESION_DN | 31 | 0.67 | 1.34 | 0.065 | 0.241 |
| 1196 | ESC_V6.5_UP_EARLY.V1_DN | 113 | 0.58 | 1.34 | 0.027 | 0.242 |
| 1197 | PID_AVB3_INTEGRIN_PATHWAY | 51 | 0.63 | 1.34 | 0.048 | 0.243 |
| 1198 | BIOCARTA_CXCR4_PATHWAY | 18 | 0.73 | 1.34 | 0.123 | 0.243 |
| 1199 | CADWELL_ATG16L1_TARGETS_UP | 49 | 0.63 | 1.34 | 0.07 | 0.243 |
| 1200 | DORN_ADENOVIRUS_INFECTION_24HR_DN | 38 | 0.65 | 1.34 | 0.086 | 0.243 |
| 1201 | VALK_AML_CLUSTER_9 | 23 | 0.71 | 1.34 | 0.083 | 0.244 |
| 1202 | MIR532_3P | 66 | 0.62 | 1.34 | 0.058 | 0.244 |
| 1203 | REACTOME_G2_M_DNA_DAMAGE_CHECKPOINT | 76 | 0.6 | 1.34 | 0.061 | 0.244 |
| 1204 | STOSSI_RESPONSE_TO_ESTRADIOL | 20 | 0.71 | 1.34 | 0.107 | 0.245 |
| 1205 | PARK_APL_PATHOGENESIS_DN | 37 | 0.67 | 1.34 | 0.063 | 0.247 |
| 1206 | BILD_SRC_ONCOGENIC_SIGNATURE | 54 | 0.63 | 1.34 | 0.065 | 0.247 |
| 1207 | GAVIN_IL2_RESPONSIVE_FOXP3_TARGETS_UP | 16 | 0.75 | 1.34 | 0.086 | 0.247 |
| 1208 | MACLACHLAN_BRCA1_TARGETS_UP | 20 | 0.72 | 1.34 | 0.08 | 0.247 |
| 1209 | PID_HES_HEY_PATHWAY | 38 | 0.66 | 1.34 | 0.077 | 0.247 |
| 1210 | PID_EPO_PATHWAY | 28 | 0.69 | 1.34 | 0.098 | 0.248 |
| 1211 | FARMER_BREAST_CANCER_BASAL_VS_LULMINAL | 275 | 0.55 | 1.33 | 0.005 | 0.248 |
| 1212 | WP_16P112_DISTAL_DELETION_SYNDROME | 28 | 0.69 | 1.33 | 0.08 | 0.248 |
| 1213 | LEE_LIVER_CANCER_ACOX1_UP | 49 | 0.63 | 1.33 | 0.074 | 0.249 |
| 1214 | REACTOME_SIGNALING_BY_CSF3_G_CSF | 27 | 0.69 | 1.33 | 0.087 | 0.249 |
| 1215 | AMIT_EGF_RESPONSE_120_MCF10A | 34 | 0.68 | 1.33 | 0.065 | 0.25 |
| 1216 | YAO_TEMPORAL_RESPONSE_TO_PROGESTERON | 24 | 0.69 | 1.33 | 0.099 | 0.25 |
| 1217 | TURASHVILI_BREAST_LOBULAR_CARCINOMA_VS_ | 68 | 0.61 | 1.33 | 0.056 | 0.25 |
| 1218 | BUCKANOVICH_T_LYMPHOCYTE_HOMING_ON_TUI | 16 | 0.75 | 1.33 | 0.088 | 0.25 |

| | | | | | | |
|---|---|---|---|---|---|---|
| 1219 | PLASARI_TGFB1_TARGETS_10HR_UP | 145 | 0.57 | 1.33 | 0.018 | 0.25 |
| 1220 | REACTOME_TRANSCRIPTIONAL_REGULATION_BY_ | 101 | 0.58 | 1.33 | 0.047 | 0.25 |
| 1221 | WILCOX_RESPONSE_TO_PROGESTERONE_DN | 44 | 0.64 | 1.33 | 0.069 | 0.25 |
| 1222 | COLLER_MYC_TARGETS_UP | 23 | 0.7 | 1.33 | 0.094 | 0.25 |
| 1223 | DLX2_TARGET_GENES | 206 | 0.56 | 1.33 | 0.008 | 0.251 |
| 1224 | SERVITJA_ISLET_HNF1A_TARGETS_DN | 59 | 0.63 | 1.33 | 0.058 | 0.251 |
| 1225 | TONKS_TARGETS_OF_RUNX1_RUNX1T1_FUSION_ | 105 | 0.59 | 1.33 | 0.034 | 0.251 |
| 1226 | KEGG_MELANOGENESIS | 79 | 0.6 | 1.33 | 0.044 | 0.251 |
| 1227 | REACTOME_SIGNALING_BY_ERYTHROPOIETIN | 22 | 0.7 | 1.33 | 0.104 | 0.251 |
| 1228 | JECHLINGER_EPITHELIAL_TO_MESENCHYMAL_TF | 45 | 0.65 | 1.33 | 0.078 | 0.251 |
| 1229 | CHYLA_CBFA2T3_TARGETS_UP | 252 | 0.55 | 1.33 | 0.012 | 0.251 |
| 1230 | PANGAS_TUMOR_SUPPRESSION_BY_SMAD1_ANE | 106 | 0.59 | 1.33 | 0.036 | 0.251 |
| 1231 | REACTOME_NUCLEAR_EVENTS_STIMULATED_BY_ | 18 | 0.73 | 1.33 | 0.084 | 0.252 |
| 1232 | MIR10397_5P | 66 | 0.61 | 1.33 | 0.047 | 0.253 |
| 1233 | ZHOU_INFLAMMATORY_RESPONSE_FIMA_UP | 346 | 0.55 | 1.33 | 0.001 | 0.253 |
| 1234 | PHONG_TNF_RESPONSE_VIA_P38_COMPLETE | 212 | 0.56 | 1.33 | 0.015 | 0.253 |
| 1235 | MIR6871_3P | 101 | 0.58 | 1.33 | 0.029 | 0.253 |
| 1236 | HOLLEMAN_ASPARAGINASE_RESISTANCE_ALL_U | 22 | 0.71 | 1.33 | 0.085 | 0.254 |
| 1237 | LANG_MYB_FAMILY_TARGETS | 27 | 0.68 | 1.33 | 0.086 | 0.253 |
| 1238 | LIAN_LIPA_TARGETS_6M | 32 | 0.67 | 1.33 | 0.067 | 0.253 |
| 1239 | MIR4661_3P | 27 | 0.69 | 1.33 | 0.091 | 0.255 |
| 1240 | SHEDDEN_LUNG_CANCER_GOOD_SURVIVAL_A4 | 149 | 0.57 | 1.33 | 0.029 | 0.256 |
| 1241 | CEBPDELTA_Q6 | 196 | 0.56 | 1.33 | 0.016 | 0.256 |
| 1242 | WESTON_VEGFA_TARGETS_6HR | 29 | 0.68 | 1.33 | 0.105 | 0.257 |
| 1243 | PHESSE_TARGETS_OF_APC_AND_MBD2_UP | 15 | 0.77 | 1.33 | 0.095 | 0.258 |
| 1244 | KHETCHOUMIAN_TRIM24_TARGETS_UP | 32 | 0.67 | 1.33 | 0.076 | 0.258 |
| 1245 | MIR6748_3P | 67 | 0.6 | 1.33 | 0.053 | 0.258 |
| 1246 | PID_SYNDECAN_1_PATHWAY | 29 | 0.67 | 1.33 | 0.097 | 0.258 |
| 1247 | JOHANSSON_BRAIN_CANCER_EARLY_VS_LATE_D | 33 | 0.66 | 1.33 | 0.093 | 0.258 |
| 1248 | ALK_DN.V1_DN | 82 | 0.6 | 1.33 | 0.043 | 0.258 |
| 1249 | WANG_TARGETS_OF_MLL_CBP_FUSION_DN | 42 | 0.65 | 1.33 | 0.089 | 0.259 |
| 1250 | SCHLINGEMANN_SKIN_CARCINOGENESIS_TPA_UI | 26 | 0.68 | 1.33 | 0.098 | 0.259 |
| 1251 | YAGI_AML_RELAPSE_PROGNOSIS | 32 | 0.67 | 1.33 | 0.085 | 0.259 |

| | | | | | | |
|---|---|---|---|---|---|---|
| 1252 | MIR4711_3P | 63 | 0.61 | 1.33 | 0.07 | 0.259 |
| 1253 | OISHI_CHOLANGIOMA_STEM_CELL_LIKE_DN | 203 | 0.56 | 1.33 | 0.014 | 0.259 |
| 1254 | MIR6839_5P | 91 | 0.59 | 1.33 | 0.045 | 0.26 |
| 1255 | MANNE_COVID19_COMBINED_COHORT_VS_HEAL | 139 | 0.57 | 1.33 | 0.032 | 0.26 |
| 1256 | MIR1343_3P | 134 | 0.57 | 1.33 | 0.022 | 0.262 |
| 1257 | DASU_IL6_SIGNALING_SCAR_UP | 27 | 0.69 | 1.33 | 0.093 | 0.263 |
| 1258 | PDGF_UP.V1_DN | 73 | 0.6 | 1.33 | 0.051 | 0.264 |
| 1259 | DARWICHE_PAPILLOMA_RISK_LOW_UP | 116 | 0.58 | 1.33 | 0.034 | 0.264 |
| 1260 | FUJII_YBX1_TARGETS_UP | 34 | 0.66 | 1.33 | 0.089 | 0.264 |
| 1261 | WP_HEPATITIS_C_AND_HEPATOCELLULAR_CARC | 41 | 0.64 | 1.33 | 0.08 | 0.264 |
| 1262 | MIR6716_5P | 111 | 0.57 | 1.32 | 0.037 | 0.265 |
| 1263 | MANNE_COVID19_NONICU_VS_HEALTHY_DONOR_ | 133 | 0.57 | 1.32 | 0.029 | 0.265 |
| 1264 | REACTOME_COLLAGEN_FORMATION | 58 | 0.62 | 1.32 | 0.06 | 0.265 |
| 1265 | BIOCARTA_G1_PATHWAY | 27 | 0.69 | 1.32 | 0.096 | 0.265 |
| 1266 | PID_IL12_2PATHWAY | 36 | 0.66 | 1.32 | 0.08 | 0.265 |
| 1267 | REACTOME_TRANSCRIPTIONAL_REGULATION_BY_ | 183 | 0.56 | 1.32 | 0.014 | 0.265 |
| 1268 | WARTERS_IR_RESPONSE_5GY | 41 | 0.64 | 1.32 | 0.09 | 0.267 |
| 1269 | REACTOME_CELLULAR_RESPONSE_TO_CHEMICA | 145 | 0.57 | 1.32 | 0.032 | 0.268 |
| 1270 | FERNANDEZ_BOUND_BY_MYC | 156 | 0.56 | 1.32 | 0.021 | 0.268 |
| 1271 | DAZARD_UV_RESPONSE_CLUSTER_G2 | 25 | 0.69 | 1.32 | 0.09 | 0.268 |
| 1272 | RGTTAMWNATT_HNF1_01 | 45 | 0.64 | 1.32 | 0.071 | 0.268 |
| 1273 | CHARAFE_BREAST_CANCER_LUMINAL_VS_MESEI | 375 | 0.54 | 1.32 | 0.003 | 0.268 |
| 1274 | WP_RESISTIN_AS_A_REGULATOR_OF_INFLAMMA | 29 | 0.68 | 1.32 | 0.071 | 0.269 |
| 1275 | SKP2_TARGET_GENES | 46 | 0.63 | 1.32 | 0.074 | 0.269 |
| 1276 | ROVERSI_GLIOMA_COPY_NUMBER_DN | 41 | 0.65 | 1.32 | 0.083 | 0.269 |
| 1277 | RIGGINS_TAMOXIFEN_RESISTANCE_UP | 51 | 0.62 | 1.32 | 0.072 | 0.27 |
| 1278 | REACTOME_DEATH_RECEPTOR_SIGNALLING | 129 | 0.58 | 1.32 | 0.031 | 0.27 |
| 1279 | REACTOME_DISEASES_OF_IMMUNE_SYSTEM | 19 | 0.72 | 1.32 | 0.125 | 0.27 |
| 1280 | WP_MODULATORS_OF_TCR_SIGNALING_AND_T_( | 48 | 0.63 | 1.32 | 0.078 | 0.271 |
| 1281 | BLALOCK_ALZHEIMERS_DISEASE_INCIPIENT_UP | 331 | 0.55 | 1.32 | 0.002 | 0.271 |
| 1282 | KEGG_TGF_BETA_SIGNALING_PATHWAY | 65 | 0.61 | 1.32 | 0.066 | 0.27 |
| 1283 | WP_PANCREATIC_ADENOCARCINOMA_PATHWAY | 86 | 0.6 | 1.32 | 0.041 | 0.27 |
| 1284 | PRC2_SUZ12_UP.V1_DN | 112 | 0.57 | 1.32 | 0.04 | 0.271 |

| | | | | | | |
|---|---|---|---|---|---|---|
| 1285 | REACTOME_RND3_GTPASE_CYCLE | 39 | 0.64 | 1.32 | 0.079 | 0.271 |
| 1286 | IL2_UP.V1_DN | 122 | 0.57 | 1.32 | 0.039 | 0.271 |
| 1287 | PKCA_DN.V1_DN | 108 | 0.58 | 1.32 | 0.041 | 0.271 |
| 1288 | WP_UNFOLDED_PROTEIN_RESPONSE | 22 | 0.7 | 1.32 | 0.122 | 0.272 |
| 1289 | E2F3_UP.V1_DN | 115 | 0.57 | 1.32 | 0.042 | 0.272 |
| 1290 | REACTOME_CA2_PATHWAY | 52 | 0.62 | 1.32 | 0.054 | 0.272 |
| 1291 | WP_HYPOTHESIZED_PATHWAYS_IN_PATHOGENE | 18 | 0.73 | 1.32 | 0.096 | 0.273 |
| 1292 | SA_PTEN_PATHWAY | 16 | 0.75 | 1.32 | 0.11 | 0.273 |
| 1293 | E2F1_UP.V1_DN | 152 | 0.56 | 1.32 | 0.025 | 0.276 |
| 1294 | CIZ_01 | 198 | 0.55 | 1.32 | 0.023 | 0.277 |
| 1295 | GU_PDEF_TARGETS_UP | 57 | 0.62 | 1.32 | 0.075 | 0.278 |
| 1296 | LEE_AGING_CEREBELLUM_DN | 54 | 0.62 | 1.32 | 0.062 | 0.278 |
| 1297 | WP_HAIR_FOLLICLE_DEVELOPMENT_ORGANOGE | 22 | 0.7 | 1.32 | 0.101 | 0.278 |
| 1298 | KEGG_ERBB_SIGNALING_PATHWAY | 81 | 0.59 | 1.32 | 0.067 | 0.278 |
| 1299 | REACTOME_HCMV_INFECTION | 122 | 0.57 | 1.32 | 0.039 | 0.278 |
| 1300 | WP_NANOMATERIAL_INDUCED_APOPTOSIS | 18 | 0.73 | 1.32 | 0.094 | 0.278 |
| 1301 | BOGNI_TREATMENT_RELATED_MYELOID_LEUKEM | 22 | 0.7 | 1.32 | 0.105 | 0.279 |
| 1302 | HOFFMANN_IMMATURE_TO_MATURE_B_LYMPHOC | 37 | 0.65 | 1.32 | 0.077 | 0.279 |
| 1303 | WP_STING_PATHWAY_IN_KAWASAKILIKE_DISEAS | 18 | 0.73 | 1.32 | 0.107 | 0.28 |
| 1304 | CAIRO_HEPATOBLASTOMA_CLASSES_DN | 141 | 0.57 | 1.32 | 0.026 | 0.28 |
| 1305 | WP_FOCAL_ADHESION | 160 | 0.56 | 1.32 | 0.03 | 0.28 |
| 1306 | DURAND_STROMA_S_UP | 220 | 0.55 | 1.32 | 0.019 | 0.281 |
| 1307 | MIR4687_3P | 70 | 0.6 | 1.32 | 0.065 | 0.281 |
| 1308 | BIOCARTA_BIOPEPTIDES_PATHWAY | 25 | 0.69 | 1.32 | 0.104 | 0.281 |
| 1309 | WP_APOPTOSIS_MODULATION_AND_SIGNALING | 78 | 0.59 | 1.32 | 0.063 | 0.281 |
| 1310 | CEBALLOS_TARGETS_OF_TP53_AND_MYC_UP | 17 | 0.72 | 1.32 | 0.111 | 0.281 |
| 1311 | ZWANG_CLASS_1_TRANSIENTLY_INDUCED_BY_E | 418 | 0.54 | 1.31 | 0.003 | 0.282 |
| 1312 | MIR372_5P | 210 | 0.55 | 1.31 | 0.02 | 0.282 |
| 1313 | IVANOVA_HEMATOPOIESIS_STEM_CELL | 144 | 0.56 | 1.31 | 0.026 | 0.282 |
| 1314 | OHGUCHI_LIVER_HNF4A_TARGETS_UP | 29 | 0.67 | 1.31 | 0.113 | 0.282 |
| 1315 | ZHENG_IL22_SIGNALING_UP | 30 | 0.67 | 1.31 | 0.107 | 0.282 |
| 1316 | REACTOME_DOWNSTREAM_SIGNALING_EVENTS_ | 78 | 0.59 | 1.31 | 0.049 | 0.282 |
| 1317 | GERHOLD_ADIPOGENESIS_DN | 51 | 0.61 | 1.31 | 0.081 | 0.282 |

| | | | | | | |
|---|---|---|---|---|---|---|
| 1318 | HOUSTIS_ROS | 28 | 0.68 | 1.31 | 0.088 | 0.283 |
| 1319 | MIR6130 | 164 | 0.55 | 1.31 | 0.033 | 0.283 |
| 1320 | REACTOME_INTERLEUKIN_1_FAMILY_SIGNALING | 109 | 0.57 | 1.31 | 0.04 | 0.283 |
| 1321 | ZHENG_FOXP3_TARGETS_UP | 21 | 0.71 | 1.31 | 0.098 | 0.283 |
| 1322 | REACTOME_ASSOCIATION_OF_TRIC_CCT_WITH_T | 37 | 0.65 | 1.31 | 0.097 | 0.283 |
| 1323 | MIR4510 | 165 | 0.56 | 1.31 | 0.035 | 0.283 |
| 1324 | KEGG_ECM_RECEPTOR_INTERACTION | 50 | 0.62 | 1.31 | 0.091 | 0.283 |
| 1325 | FARMER_BREAST_CANCER_APOCRINE_VS_BASA | 272 | 0.55 | 1.31 | 0.009 | 0.283 |
| 1326 | MYC_UP.V1_DN | 121 | 0.58 | 1.31 | 0.036 | 0.284 |
| 1327 | KRAS.LUNG_UP.V1_UP | 73 | 0.6 | 1.31 | 0.048 | 0.284 |
| 1328 | BIOCARTA_GH_PATHWAY | 23 | 0.69 | 1.31 | 0.101 | 0.284 |
| 1329 | NAKAJIMA_MAST_CELL | 27 | 0.68 | 1.31 | 0.113 | 0.284 |
| 1330 | MIYAGAWA_TARGETS_OF_EWSR1_ETS_FUSIONS_ | 165 | 0.56 | 1.31 | 0.017 | 0.286 |
| 1331 | ROSS_LEUKEMIA_WITH_MLL_FUSIONS | 65 | 0.59 | 1.31 | 0.073 | 0.286 |
| 1332 | REACTOME_CLASS_I_MHC_MEDIATED_ANTIGEN_I | 332 | 0.54 | 1.31 | 0.007 | 0.287 |
| 1333 | HELLER_HDAC_TARGETS_SILENCED_BY_METHYL | 380 | 0.53 | 1.31 | 0.005 | 0.287 |
| 1334 | NAKAMURA_METASTASIS_MODEL_DN | 32 | 0.66 | 1.31 | 0.102 | 0.287 |
| 1335 | CROONQUIST_STROMAL_STIMULATION_UP | 34 | 0.65 | 1.31 | 0.104 | 0.287 |
| 1336 | GNATENKO_PLATELET_SIGNATURE | 37 | 0.65 | 1.31 | 0.085 | 0.287 |
| 1337 | FALVELLA_SMOKERS_WITH_LUNG_CANCER | 62 | 0.61 | 1.31 | 0.067 | 0.287 |
| 1338 | WP_HEDGEHOG_SIGNALING_PATHWAY_WP4249 | 33 | 0.65 | 1.31 | 0.107 | 0.287 |
| 1339 | WINTER_HYPOXIA_DN | 32 | 0.66 | 1.31 | 0.091 | 0.288 |
| 1340 | MIR6757_5P | 47 | 0.62 | 1.31 | 0.091 | 0.289 |
| 1341 | LIANG_HEMATOPOIESIS_STEM_CELL_NUMBER_L/ | 32 | 0.66 | 1.31 | 0.103 | 0.289 |
| 1342 | REACTOME_INACTIVATION_OF_CSF3_G_CSF_SIG | 22 | 0.7 | 1.31 | 0.091 | 0.29 |
| 1343 | SANSOM_APC_TARGETS_DN | 251 | 0.54 | 1.31 | 0.01 | 0.29 |
| 1344 | GARGALOVIC_RESPONSE_TO_OXIDIZED_PHOSPH | 67 | 0.6 | 1.31 | 0.07 | 0.29 |
| 1345 | HEIDENBLAD_AMPLIFIED_IN_PANCREATIC_CANCE | 50 | 0.63 | 1.31 | 0.081 | 0.29 |
| 1346 | HARRIS_BRAIN_CANCER_PROGENITORS | 26 | 0.69 | 1.31 | 0.103 | 0.29 |
| 1347 | MIR6827_5P | 65 | 0.61 | 1.31 | 0.076 | 0.29 |
| 1348 | REACTOME_MOLECULES_ASSOCIATED_WITH_EL/ | 25 | 0.68 | 1.31 | 0.105 | 0.29 |
| 1349 | WP_ERBB_SIGNALING_PATHWAY | 87 | 0.58 | 1.31 | 0.068 | 0.291 |
| 1350 | REACTOME_CONSTITUTIVE_SIGNALING_BY_EGFF | 15 | 0.73 | 1.31 | 0.107 | 0.291 |

| | | | | | | |
|---|---|---|---|---|---|---|
| 1351 | MIKKELSEN_ES_HCP_WITH_H3_UNMETHYLATED | 28 | 0.67 | 1.31 | 0.115 | 0.291 |
| 1352 | BOYLAN_MULTIPLE_MYELOMA_PCA1_UP | 59 | 0.61 | 1.31 | 0.067 | 0.291 |
| 1353 | MIR339_5P | 59 | 0.61 | 1.31 | 0.068 | 0.291 |
| 1354 | YOKOE_CANCER_TESTIS_ANTIGENS | 27 | 0.69 | 1.31 | 0.091 | 0.291 |
| 1355 | URS_ADIPOCYTE_DIFFERENTIATION_DN | 21 | 0.71 | 1.31 | 0.099 | 0.291 |
| 1356 | ZNF418_TARGET_GENES | 27 | 0.67 | 1.31 | 0.108 | 0.293 |
| 1357 | BIOCARTA_RELA_PATHWAY | 15 | 0.75 | 1.31 | 0.119 | 0.293 |
| 1358 | WP_BREAST_CANCER_PATHWAY | 126 | 0.57 | 1.31 | 0.041 | 0.293 |
| 1359 | OKUMURA_INFLAMMATORY_RESPONSE_LPS | 149 | 0.56 | 1.31 | 0.032 | 0.293 |
| 1360 | AMIT_SERUM_RESPONSE_480_MCF10A | 29 | 0.68 | 1.31 | 0.101 | 0.293 |
| 1361 | LAIHO_COLORECTAL_CANCER_SERRATED_UP | 104 | 0.57 | 1.31 | 0.054 | 0.293 |
| 1362 | DAZARD_UV_RESPONSE_CLUSTER_G28 | 15 | 0.75 | 1.31 | 0.11 | 0.293 |
| 1363 | GAUSSMANN_MLL_AF4_FUSION_TARGETS_E_UP | 64 | 0.6 | 1.31 | 0.08 | 0.293 |
| 1364 | ABBUD_LIF_SIGNALING_1_UP | 28 | 0.67 | 1.31 | 0.11 | 0.294 |
| 1365 | REACTOME_ACTIVATION_OF_BH3_ONLY_PROTEII | 30 | 0.66 | 1.31 | 0.116 | 0.294 |
| 1366 | MIR6866_5P | 64 | 0.61 | 1.31 | 0.072 | 0.296 |
| 1367 | OUELLET_OVARIAN_CANCER_INVASIVE_VS_LMP_ | 113 | 0.56 | 1.31 | 0.047 | 0.296 |
| 1368 | BOYAULT_LIVER_CANCER_SUBCLASS_G1_DN | 38 | 0.64 | 1.31 | 0.108 | 0.296 |
| 1369 | GTCAACC_MIR3805P | 18 | 0.73 | 1.31 | 0.099 | 0.297 |
| 1370 | WANG_IMMORTALIZED_BY_HOXA9_AND_MEIS1_U | 18 | 0.72 | 1.31 | 0.099 | 0.296 |
| 1371 | REACTOME_DEGRADATION_OF_THE_EXTRACELLU | 81 | 0.59 | 1.31 | 0.076 | 0.297 |
| 1372 | FERRANDO_T_ALL_WITH_MLL_ENL_FUSION_DN | 77 | 0.6 | 1.3 | 0.064 | 0.297 |
| 1373 | KIM_WT1_TARGETS_UP | 181 | 0.55 | 1.3 | 0.039 | 0.297 |
| 1374 | MIR6740_5P | 52 | 0.61 | 1.3 | 0.085 | 0.297 |
| 1375 | REACTOME_LEISHMANIA_INFECTION | 149 | 0.56 | 1.3 | 0.047 | 0.297 |
| 1376 | PID_BETA_CATENIN_NUC_PATHWAY | 65 | 0.59 | 1.3 | 0.088 | 0.298 |
| 1377 | BIOCARTA_BAD_PATHWAY | 19 | 0.71 | 1.3 | 0.119 | 0.299 |
| 1378 | MIR7850_5P | 104 | 0.57 | 1.3 | 0.057 | 0.299 |
| 1379 | SMID_BREAST_CANCER_RELAPSE_IN_BRAIN_UP | 27 | 0.68 | 1.3 | 0.105 | 0.299 |
| 1380 | HOFFMANN_PRE_BI_TO_LARGE_PRE_BII_LYMPHO | 56 | 0.61 | 1.3 | 0.081 | 0.3 |
| 1381 | MIR4506 | 33 | 0.66 | 1.3 | 0.095 | 0.299 |
| 1382 | FRIDMAN_SENESCENCE_UP | 60 | 0.6 | 1.3 | 0.069 | 0.299 |
| 1383 | PSMB5_TARGET_GENES | 259 | 0.54 | 1.3 | 0.018 | 0.3 |

| | | | | | | |
|---|---|---|---|---|---|---|
| 1384 | BILBAN_B_CLL_LPL_UP | 54 | 0.61 | 1.3 | 0.095 | 0.301 |
| 1385 | KRAS.BREAST_UP.V1_UP | 69 | 0.6 | 1.3 | 0.069 | 0.301 |
| 1386 | KYNG_NORMAL_AGING_UP | 15 | 0.74 | 1.3 | 0.12 | 0.301 |
| 1387 | REACTOME_TP53_REGULATES_TRANSCRIPTION_ | 39 | 0.64 | 1.3 | 0.099 | 0.301 |
| 1388 | WARTERS_RESPONSE_TO_IR_SKIN | 69 | 0.59 | 1.3 | 0.077 | 0.301 |
| 1389 | BERENJENO_TRANSFORMED_BY_RHOA_FOREVEI | 19 | 0.72 | 1.3 | 0.109 | 0.301 |
| 1390 | BIOCARTA_BARRESTIN_SRC_PATHWAY | 15 | 0.74 | 1.3 | 0.113 | 0.302 |
| 1391 | BIOCARTA_TFF_PATHWAY | 17 | 0.72 | 1.3 | 0.141 | 0.302 |
| 1392 | PID_IL2_PI3K_PATHWAY | 32 | 0.66 | 1.3 | 0.11 | 0.302 |
| 1393 | MARSON_FOXP3_TARGETS_STIMULATED_UP | 17 | 0.72 | 1.3 | 0.121 | 0.303 |
| 1394 | MIR532_5P | 68 | 0.6 | 1.3 | 0.067 | 0.303 |
| 1395 | TAKEDA_TARGETS_OF_NUP98_HOXA9_FUSION_6 | 57 | 0.61 | 1.3 | 0.086 | 0.303 |
| 1396 | CRX_DN.V1_DN | 94 | 0.58 | 1.3 | 0.052 | 0.303 |
| 1397 | REACTOME_DISEASES_ASSOCIATED_WITH_GLYC | 31 | 0.66 | 1.3 | 0.106 | 0.303 |
| 1398 | RIGGI_EWING_SARCOMA_PROGENITOR_DN | 129 | 0.56 | 1.3 | 0.04 | 0.304 |
| 1399 | RUIZ_TNC_TARGETS_DN | 131 | 0.56 | 1.3 | 0.046 | 0.304 |
| 1400 | NAKAMURA_TUMOR_ZONE_PERIPHERAL_VS_CEN | 234 | 0.54 | 1.3 | 0.015 | 0.305 |
| 1401 | TCGA_GLIOBLASTOMA_COPY_NUMBER_UP | 68 | 0.6 | 1.3 | 0.081 | 0.304 |
| 1402 | WP_MIRNA_REGULATION_OF_P53_PATHWAY_IN_I | 22 | 0.7 | 1.3 | 0.104 | 0.304 |
| 1403 | REACTOME_INTERLEUKIN_7_SIGNALING | 23 | 0.69 | 1.3 | 0.117 | 0.304 |
| 1404 | KEGG_PROSTATE_CANCER | 84 | 0.58 | 1.3 | 0.073 | 0.304 |
| 1405 | MIR3161 | 61 | 0.6 | 1.3 | 0.094 | 0.304 |
| 1406 | ZHAN_V1_LATE_DIFFERENTIATION_GENES_UP | 30 | 0.65 | 1.3 | 0.119 | 0.304 |
| 1407 | CERVERA_SDHB_TARGETS_2 | 75 | 0.59 | 1.3 | 0.069 | 0.304 |
| 1408 | SWEET_KRAS_ONCOGENIC_SIGNATURE | 77 | 0.59 | 1.3 | 0.08 | 0.305 |
| 1409 | REACTOME_BETA_CATENIN_INDEPENDENT_WNT_ | 132 | 0.56 | 1.3 | 0.033 | 0.306 |
| 1410 | REACTOME_SCF_SKP2_MEDIATED_DEGRADATION | 60 | 0.59 | 1.3 | 0.083 | 0.306 |
| 1411 | MIR6769B_5P | 63 | 0.59 | 1.3 | 0.088 | 0.306 |
| 1412 | REACTOME_DISEASES_OF_METABOLISM | 186 | 0.55 | 1.3 | 0.028 | 0.307 |
| 1413 | KEGG_SELENOAMINO_ACID_METABOLISM | 25 | 0.67 | 1.3 | 0.113 | 0.307 |
| 1414 | MIR6129 | 166 | 0.55 | 1.3 | 0.026 | 0.307 |
| 1415 | COATES_MACROPHAGE_M1_VS_M2_UP | 67 | 0.6 | 1.3 | 0.074 | 0.307 |
| 1416 | WP_HISTONE_MODIFICATIONS | 49 | 0.63 | 1.3 | 0.093 | 0.307 |

| | | | | | | |
|---|---|---|---|---|---|---|
| 1417 | ROZANOV_MMP14_TARGETS_SUBSET | 22 | 0.69 | 1.3 | 0.116 | 0.308 |
| 1418 | MISHRA_CARCINOMA_ASSOCIATED_FIBROBLAST_ | 22 | 0.69 | 1.3 | 0.127 | 0.308 |
| 1419 | REACTOME_CONSTITUTIVE_SIGNALING_BY_LIGAI | 19 | 0.7 | 1.3 | 0.134 | 0.308 |
| 1420 | FERRANDO_T_ALL_WITH_MLL_ENL_FUSION_UP | 72 | 0.59 | 1.3 | 0.077 | 0.308 |
| 1421 | MIKKELSEN_IPS_ICP_WITH_H3K4ME3_AND_H327I | 70 | 0.59 | 1.3 | 0.092 | 0.308 |
| 1422 | FERRANDO_HOX11_NEIGHBORS | 16 | 0.73 | 1.3 | 0.135 | 0.308 |
| 1423 | KONDO_COLON_CANCER_HCP_WITH_H3K27ME1 | 20 | 0.71 | 1.3 | 0.113 | 0.308 |
| 1424 | BIOCARTA_ATM_PATHWAY | 20 | 0.71 | 1.3 | 0.117 | 0.308 |
| 1425 | MIR6133 | 169 | 0.55 | 1.3 | 0.025 | 0.308 |
| 1426 | MARTENS_TRETINOIN_RESPONSE_UP | 426 | 0.53 | 1.3 | 0.001 | 0.308 |
| 1427 | REACTOME_SIGNALING_BY_NOTCH1 | 70 | 0.59 | 1.3 | 0.078 | 0.309 |
| 1428 | MIR6505_3P | 77 | 0.59 | 1.3 | 0.077 | 0.309 |
| 1429 | HERNANDEZ_ABERRANT_MITOSIS_BY_DOCETACE | 52 | 0.61 | 1.3 | 0.081 | 0.309 |
| 1430 | REACTOME_RAC2_GTPASE_CYCLE | 83 | 0.58 | 1.3 | 0.073 | 0.309 |
| 1431 | WP_TP53_NETWORK | 19 | 0.7 | 1.3 | 0.129 | 0.309 |
| 1432 | KEGG_BASAL_CELL_CARCINOMA | 39 | 0.63 | 1.3 | 0.119 | 0.309 |
| 1433 | WP_PROSTAGLANDIN_AND_LEUKOTRIENE_METAI | 18 | 0.71 | 1.3 | 0.144 | 0.309 |
| 1434 | BROWNE_HCMV_INFECTION_16HR_DN | 64 | 0.6 | 1.3 | 0.094 | 0.309 |
| 1435 | RBL1_TARGET_GENES | 38 | 0.63 | 1.3 | 0.111 | 0.309 |
| 1436 | GALLUZZI_PERMEABILIZE_MITOCHONDRIA | 38 | 0.63 | 1.3 | 0.107 | 0.311 |
| 1437 | KINSEY_TARGETS_OF_EWSR1_FLII_FUSION_DN | 281 | 0.54 | 1.29 | 0.019 | 0.313 |
| 1438 | ZHAN_MULTIPLE_MYELOMA_CD1_DN | 42 | 0.63 | 1.29 | 0.105 | 0.313 |
| 1439 | LMO2COM_02 | 181 | 0.55 | 1.29 | 0.026 | 0.313 |
| 1440 | WP_STRIATED_MUSCLE_CONTRACTION_PATHWAY | 20 | 0.7 | 1.29 | 0.116 | 0.313 |
| 1441 | WP_MIRNAS_INVOLVED_IN_DNA_DAMAGE_RESPC | 17 | 0.72 | 1.29 | 0.122 | 0.314 |
| 1442 | PEA3_Q6 | 214 | 0.54 | 1.29 | 0.026 | 0.314 |
| 1443 | BOYLAN_MULTIPLE_MYELOMA_PCA3_UP | 66 | 0.59 | 1.29 | 0.083 | 0.314 |
| 1444 | WP_WNT_SIGNALING | 94 | 0.57 | 1.29 | 0.069 | 0.314 |
| 1445 | FLORIO_NEOCORTEX_BASAL_RADIAL_GLIA_DN | 171 | 0.55 | 1.29 | 0.033 | 0.314 |
| 1446 | STAT_01 | 206 | 0.54 | 1.29 | 0.025 | 0.314 |
| 1447 | JIANG_VHL_TARGETS | 84 | 0.58 | 1.29 | 0.069 | 0.315 |
| 1448 | WIKMAN_ASBESTOS_LUNG_CANCER_DN | 21 | 0.68 | 1.29 | 0.138 | 0.315 |
| 1449 | MCCABE_HOXC6_TARGETS_DN | 18 | 0.72 | 1.29 | 0.117 | 0.316 |

| | | | | | | |
|---|---|---|---|---|---|---|
| 1450 | INGRAM_SHH_TARGETS_DN | 46 | 0.62 | 1.29 | 0.099 | 0.316 |
| 1451 | KRAS.LUNG_UP.V1_DN | 75 | 0.58 | 1.29 | 0.072 | 0.316 |
| 1452 | FLECHNER_PBL_KIDNEY_TRANSPLANT_OK_VS_D | 144 | 0.55 | 1.29 | 0.04 | 0.316 |
| 1453 | RB_DN.V1_DN | 105 | 0.57 | 1.29 | 0.062 | 0.316 |
| 1454 | MIR10524_5P | 73 | 0.59 | 1.29 | 0.091 | 0.317 |
| 1455 | HOSHIDA_LIVER_CANCER_SUBCLASS_S3 | 166 | 0.55 | 1.29 | 0.039 | 0.317 |
| 1456 | BIOCARTA_EPO_PATHWAY | 18 | 0.72 | 1.29 | 0.135 | 0.318 |
| 1457 | KIM_GERMINAL_CENTER_T_HELPER_UP | 55 | 0.61 | 1.29 | 0.103 | 0.318 |
| 1458 | IL15_UP.V1_UP | 132 | 0.55 | 1.29 | 0.054 | 0.318 |
| 1459 | PGF_UP.V1_DN | 127 | 0.56 | 1.29 | 0.05 | 0.318 |
| 1460 | ZHONG_SECRETOME_OF_LUNG_CANCER_AND_FI | 110 | 0.56 | 1.29 | 0.054 | 0.319 |
| 1461 | SHETH_LIVER_CANCER_VS_TXNIP_LOSS_PAM3 | 56 | 0.6 | 1.29 | 0.08 | 0.319 |
| 1462 | LOPES_METHYLATED_IN_COLON_CANCER_UP | 16 | 0.72 | 1.29 | 0.127 | 0.319 |
| 1463 | BURTON_ADIPOGENESIS_7 | 37 | 0.64 | 1.29 | 0.101 | 0.319 |
| 1464 | AMIT_EGF_RESPONSE_240_MCF10A | 18 | 0.71 | 1.29 | 0.136 | 0.319 |
| 1465 | VECCHI_GASTRIC_CANCER_ADVANCED_VS_EARL | 115 | 0.56 | 1.29 | 0.054 | 0.319 |
| 1466 | JIANG_AGING_HYPOTHALAMUS_UP | 43 | 0.63 | 1.29 | 0.102 | 0.319 |
| 1467 | MIR6127 | 179 | 0.54 | 1.29 | 0.027 | 0.319 |
| 1468 | CAVIN1_TARGET_GENES | 69 | 0.59 | 1.29 | 0.082 | 0.319 |
| 1469 | MIR5706 | 136 | 0.55 | 1.29 | 0.047 | 0.319 |
| 1470 | GOBERT_CORE_OLIGODENDROCYTE_DIFFERENTI | 40 | 0.63 | 1.29 | 0.109 | 0.319 |
| 1471 | WP_TGFBETA_RECEPTOR_SIGNALING_IN_SKELET | 44 | 0.62 | 1.29 | 0.114 | 0.32 |
| 1472 | GARGALOVIC_RESPONSE_TO_OXIDIZED_PHOSPH | 132 | 0.56 | 1.29 | 0.047 | 0.32 |
| 1473 | DORN_ADENOVIRUS_INFECTION_12HR_DN | 30 | 0.65 | 1.29 | 0.118 | 0.321 |
| 1474 | MATZUK_SPERMATOCYTE | 59 | 0.6 | 1.29 | 0.07 | 0.32 |
| 1475 | HASLINGER_B_CLL_WITH_11Q23_DELETION | 22 | 0.69 | 1.29 | 0.131 | 0.32 |
| 1476 | LEE_LIVER_CANCER_MYC_TGFA_UP | 46 | 0.62 | 1.29 | 0.091 | 0.321 |
| 1477 | MIR7106_3P | 39 | 0.64 | 1.29 | 0.117 | 0.322 |
| 1478 | SHAFFER_IRF4_TARGETS_IN_MYELOMA_VS_MATU | 95 | 0.57 | 1.29 | 0.067 | 0.322 |
| 1479 | ZHONG_SECRETOME_OF_LUNG_CANCER_AND_MA | 69 | 0.59 | 1.29 | 0.08 | 0.324 |
| 1480 | WP_THYMIC_STROMAL_LYMPHOPOIETIN_TSLP_S | 40 | 0.63 | 1.29 | 0.104 | 0.324 |
| 1481 | STEARMAN_LUNG_CANCER_EARLY_VS_LATE_DN | 47 | 0.6 | 1.29 | 0.115 | 0.324 |
| 1482 | ZIRN_TRETINOIN_RESPONSE_UP | 18 | 0.71 | 1.29 | 0.129 | 0.325 |

| | | | | | | |
|---|---|---|---|---|---|---|
| 1483 | REACTOME_NOTCH1_INTRACELLULAR_DOMAIN_F | 46 | 0.61 | 1.29 | 0.101 | 0.325 |
| 1484 | TCF11_01 | 204 | 0.54 | 1.29 | 0.038 | 0.326 |
| 1485 | MIR6737_5P | 62 | 0.59 | 1.29 | 0.086 | 0.326 |
| 1486 | GAZDA_DIAMOND_BLACKFAN_ANEMIA_MYELOID_ | 33 | 0.65 | 1.29 | 0.1 | 0.326 |
| 1487 | MIR4781_3P | 31 | 0.65 | 1.29 | 0.111 | 0.326 |
| 1488 | WP_PPARALPHA_PATHWAY | 18 | 0.71 | 1.29 | 0.115 | 0.326 |
| 1489 | WP_INFLAMMATORY_RESPONSE_PATHWAY | 17 | 0.71 | 1.29 | 0.142 | 0.327 |
| 1490 | PID_CXCR3_PATHWAY | 38 | 0.63 | 1.29 | 0.128 | 0.326 |
| 1491 | JAEGER_METASTASIS_DN | 158 | 0.54 | 1.29 | 0.046 | 0.326 |
| 1492 | RICKMAN_HEAD_AND_NECK_CANCER_F | 27 | 0.66 | 1.29 | 0.121 | 0.328 |
| 1493 | MULLIGHAN_MLL_SIGNATURE_2_DN | 220 | 0.53 | 1.29 | 0.021 | 0.329 |
| 1494 | MIR5001_3P | 130 | 0.56 | 1.28 | 0.047 | 0.33 |
| 1495 | PEDERSEN_METASTASIS_BY_ERBB2_ISOFORM_7 | 345 | 0.53 | 1.28 | 0.007 | 0.331 |
| 1496 | AMIT_SERUM_RESPONSE_240_MCF10A | 42 | 0.62 | 1.28 | 0.101 | 0.331 |
| 1497 | KEGG_NOTCH_SIGNALING_PATHWAY | 44 | 0.62 | 1.28 | 0.124 | 0.331 |
| 1498 | REACTOME_FATTY_ACID_METABOLISM | 129 | 0.56 | 1.28 | 0.05 | 0.331 |
| 1499 | OLSSON_E2F3_TARGETS_UP | 19 | 0.7 | 1.28 | 0.148 | 0.331 |
| 1500 | MIR4433A_3P | 52 | 0.61 | 1.28 | 0.086 | 0.331 |
| 1501 | MORI_MATURE_B_LYMPHOCYTE_DN | 65 | 0.59 | 1.28 | 0.099 | 0.331 |
| 1502 | WP_CCL18_SIGNALING_PATHWAY | 38 | 0.64 | 1.28 | 0.102 | 0.332 |
| 1503 | MIR6816_3P | 29 | 0.65 | 1.28 | 0.12 | 0.332 |
| 1504 | OSADA_ASCL1_TARGETS_UP | 38 | 0.63 | 1.28 | 0.13 | 0.332 |
| 1505 | MIR605_3P | 230 | 0.54 | 1.28 | 0.026 | 0.332 |
| 1506 | MIR185_3P | 133 | 0.55 | 1.28 | 0.044 | 0.332 |
| 1507 | LEE_AGING_NEOCORTEX_DN | 46 | 0.62 | 1.28 | 0.103 | 0.334 |
| 1508 | REACTOME_NRIF_SIGNALS_CELL_DEATH_FROM_ | 15 | 0.73 | 1.28 | 0.148 | 0.334 |
| 1509 | LEE_LIVER_CANCER_DENA_DN | 35 | 0.64 | 1.28 | 0.113 | 0.334 |
| 1510 | LUND_SILENCED_BY_METHYLATION | 15 | 0.71 | 1.28 | 0.149 | 0.334 |
| 1511 | CYCLIN_D1_KE_.V1_DN | 123 | 0.55 | 1.28 | 0.054 | 0.336 |
| 1512 | PID_TOLL_ENDOGENOUS_PATHWAY | 18 | 0.7 | 1.28 | 0.145 | 0.335 |
| 1513 | WU_ALZHEIMER_DISEASE_DN | 17 | 0.72 | 1.28 | 0.136 | 0.336 |
| 1514 | REACTOME_SYNTHESIS_OF_VERY_LONG_CHAIN_ | 18 | 0.7 | 1.28 | 0.141 | 0.337 |
| 1515 | MIR6741_5P | 18 | 0.69 | 1.28 | 0.138 | 0.337 |

| | | | | | | |
|---|---|---|---|---|---|---|
| 1516 | MIR8079 | 28 | 0.65 | 1.28 | 0.144 | 0.337 |
| 1517 | ONDER_CDH1_TARGETS_2_UP | 202 | 0.54 | 1.28 | 0.036 | 0.337 |
| 1518 | MIR4999_5P | 80 | 0.57 | 1.28 | 0.082 | 0.338 |
| 1519 | PID_SYNDECAN_2_PATHWAY | 27 | 0.67 | 1.28 | 0.13 | 0.338 |
| 1520 | IVANOVSKA_MIR106B_TARGETS | 85 | 0.57 | 1.28 | 0.072 | 0.339 |
| 1521 | WEIGEL_OXIDATIVE_STRESS_BY_TBH_AND_H2O2 | 29 | 0.66 | 1.28 | 0.135 | 0.339 |
| 1522 | ROY_WOUND_BLOOD_VESSEL_UP | 36 | 0.63 | 1.28 | 0.123 | 0.339 |
| 1523 | MIR1301_5P | 44 | 0.62 | 1.28 | 0.111 | 0.339 |
| 1524 | BACH2_01 | 218 | 0.53 | 1.28 | 0.035 | 0.338 |
| 1525 | PID_TCR_CALCIUM_PATHWAY | 18 | 0.7 | 1.28 | 0.145 | 0.339 |
| 1526 | REACTOME_G0_AND_EARLY_G1 | 27 | 0.66 | 1.28 | 0.154 | 0.339 |
| 1527 | BIOCARTA_PTEN_PATHWAY | 17 | 0.73 | 1.28 | 0.135 | 0.34 |
| 1528 | KEGG_MELANOMA | 54 | 0.6 | 1.28 | 0.096 | 0.34 |
| 1529 | MIR601 | 34 | 0.65 | 1.28 | 0.121 | 0.34 |
| 1530 | WP_NCRNAS_INVOLVED_IN_WNT_SIGNALING_IN_ | 68 | 0.59 | 1.28 | 0.085 | 0.34 |
| 1531 | STAT5A_02 | 112 | 0.57 | 1.28 | 0.063 | 0.34 |
| 1532 | MIR5701 | 164 | 0.54 | 1.28 | 0.041 | 0.34 |
| 1533 | CHIANG_LIVER_CANCER_SUBCLASS_POLYSOMY7 | 20 | 0.69 | 1.28 | 0.128 | 0.341 |
| 1534 | GUILLAUMOND_KLF10_TARGETS_DN | 21 | 0.69 | 1.28 | 0.137 | 0.341 |
| 1535 | MIR3064_3P | 140 | 0.55 | 1.28 | 0.053 | 0.343 |
| 1536 | OHM_METHYLATED_IN_ADULT_CANCERS | 21 | 0.68 | 1.28 | 0.145 | 0.343 |
| 1537 | HAN_SATB1_TARGETS_DN | 311 | 0.52 | 1.28 | 0.025 | 0.343 |
| 1538 | HUNSBERGER_EXERCISE_REGULATED_GENES | 23 | 0.68 | 1.28 | 0.153 | 0.343 |
| 1539 | MARTINEZ_TP53_TARGETS_DN | 498 | 0.52 | 1.28 | 0.002 | 0.343 |
| 1540 | REACTOME_UNFOLDED_PROTEIN_RESPONSE_UF | 88 | 0.57 | 1.28 | 0.076 | 0.344 |
| 1541 | HOFMANN_MYELODYSPLASTIC_SYNDROM_LOW_I | 23 | 0.67 | 1.28 | 0.16 | 0.344 |
| 1542 | FLORIO_NEOCORTEX_BASAL_RADIAL_GLIA_UP | 172 | 0.54 | 1.28 | 0.044 | 0.344 |
| 1543 | MORI_EMU_MYC_LYMPHOMA_BY_ONSET_TIME_D | 15 | 0.73 | 1.28 | 0.139 | 0.344 |
| 1544 | SWEET_LUNG_CANCER_KRAS_DN | 262 | 0.53 | 1.28 | 0.028 | 0.346 |
| 1545 | BERTUCCI_INVASIVE_CARCINOMA_DUCTAL_VS_L | 25 | 0.67 | 1.28 | 0.142 | 0.346 |
| 1546 | REACTOME_CIRCADIAN_CLOCK | 67 | 0.58 | 1.28 | 0.095 | 0.346 |
| 1547 | ANDERSEN_LIVER_CANCER_KRT19_UP | 34 | 0.64 | 1.28 | 0.135 | 0.347 |
| 1548 | PAPASPYRIDONOS_UNSTABLE_ATEROSCLEROTIC | 41 | 0.63 | 1.28 | 0.116 | 0.347 |

| | | | | | | |
|---|---|---|---|---|---|---|
| 1549 | MIR4804_3P | 74 | 0.58 | 1.28 | 0.079 | 0.347 |
| 1550 | UZONYI_RESPONSE_TO_LEUKOTRIENE_AND_THF | 31 | 0.65 | 1.27 | 0.141 | 0.348 |
| 1551 | WP_PHOTODYNAMIC_THERAPYINDUCED_NFE2L2_ | 19 | 0.7 | 1.27 | 0.133 | 0.348 |
| 1552 | NOUSHMEHR_GBM_SILENCED_BY_METHYLATION | 30 | 0.64 | 1.27 | 0.148 | 0.35 |
| 1553 | PICCALUGA_ANGIOIMMUNOBLASTIC_LYMPHOMA_ | 128 | 0.55 | 1.27 | 0.073 | 0.351 |
| 1554 | FOROUTAN_PRODRANK_TGFB_EMT_UP | 145 | 0.55 | 1.27 | 0.053 | 0.351 |
| 1555 | KAAB_FAILED_HEART_ATRIUM_DN | 135 | 0.54 | 1.27 | 0.063 | 0.352 |
| 1556 | GTTAAAG_MIR302B | 62 | 0.59 | 1.27 | 0.103 | 0.351 |
| 1557 | ZHAN_V2_LATE_DIFFERENTIATION_GENES | 43 | 0.61 | 1.27 | 0.118 | 0.351 |
| 1558 | NEWMAN_ERCC6_TARGETS_UP | 18 | 0.7 | 1.27 | 0.168 | 0.351 |
| 1559 | BENPORATH_MYC_TARGETS_WITH_EBOX | 208 | 0.53 | 1.27 | 0.035 | 0.352 |
| 1560 | GAVIN_FOXP3_TARGETS_CLUSTER_P6 | 79 | 0.58 | 1.27 | 0.081 | 0.352 |
| 1561 | HADDAD_T_LYMPHOCYTE_AND_NK_PROGENITOR | 33 | 0.64 | 1.27 | 0.139 | 0.352 |
| 1562 | MARIADASON_RESPONSE_TO_CURCUMIN_SULINE | 22 | 0.68 | 1.27 | 0.155 | 0.352 |
| 1563 | WILLIAMS_ESR1_TARGETS_UP | 21 | 0.69 | 1.27 | 0.142 | 0.352 |
| 1564 | KEGG_PRIMARY_IMMUNODEFICIENCY | 20 | 0.69 | 1.27 | 0.126 | 0.353 |
| 1565 | CGGAARNGGCNG_UNKNOWN | 54 | 0.6 | 1.27 | 0.119 | 0.353 |
| 1566 | CDH4_TARGET_GENES | 63 | 0.59 | 1.27 | 0.09 | 0.355 |
| 1567 | MIR4694_3P | 253 | 0.52 | 1.27 | 0.036 | 0.354 |
| 1568 | BROWNE_HCMV_INFECTION_14HR_DN | 242 | 0.53 | 1.27 | 0.025 | 0.355 |
| 1569 | MIR9898 | 51 | 0.6 | 1.27 | 0.093 | 0.357 |
| 1570 | REACTOME_TELOMERE_EXTENSION_BY_TELOME | 23 | 0.68 | 1.27 | 0.147 | 0.358 |
| 1571 | KENNY_CTNNB1_TARGETS_UP | 44 | 0.6 | 1.27 | 0.135 | 0.36 |
| 1572 | MIR8055 | 64 | 0.59 | 1.27 | 0.096 | 0.36 |
| 1573 | MIR6811_5P | 62 | 0.58 | 1.27 | 0.111 | 0.361 |
| 1574 | BASSO_CD40_SIGNALING_DN | 51 | 0.61 | 1.27 | 0.097 | 0.361 |
| 1575 | KEGG_VEGF_SIGNALING_PATHWAY | 60 | 0.58 | 1.27 | 0.096 | 0.36 |
| 1576 | GINESTIER_BREAST_CANCER_20Q13_AMPLIFICAT | 179 | 0.53 | 1.27 | 0.044 | 0.36 |
| 1577 | WP_MICROGLIA_PATHOGEN_PHAGOCYTOSIS_PA | 29 | 0.65 | 1.27 | 0.144 | 0.361 |
| 1578 | REACTOME_SIGNALING_BY_ERBB2_ECD_MUTANT | 16 | 0.71 | 1.27 | 0.164 | 0.361 |
| 1579 | LUI_THYROID_CANCER_PAX8_PPARG_UP | 41 | 0.62 | 1.27 | 0.137 | 0.361 |
| 1580 | ROSS_AML_WITH_PML_RARA_FUSION | 64 | 0.58 | 1.27 | 0.109 | 0.361 |
| 1581 | MIR5704 | 45 | 0.6 | 1.27 | 0.114 | 0.361 |

| | | | | | | |
|---|---|---|---|---|---|---|
| 1582 | REACTOME_SIGNALING_BY_CYTOSOLIC_FGFR1_I | 18 | 0.69 | 1.27 | 0.184 | 0.361 |
| 1583 | VERRECCHIA_DELAYED_RESPONSE_TO_TGFB1 | 31 | 0.64 | 1.27 | 0.141 | 0.361 |
| 1584 | WP_CELL_MIGRATION_AND_INVASION_THROUGH_ | 27 | 0.66 | 1.27 | 0.152 | 0.361 |
| 1585 | GERHOLD_ADIPOGENESIS_UP | 39 | 0.61 | 1.27 | 0.138 | 0.361 |
| 1586 | IL21_UP.V1_DN | 108 | 0.56 | 1.27 | 0.083 | 0.361 |
| 1587 | MIR2052 | 188 | 0.53 | 1.27 | 0.036 | 0.361 |
| 1588 | GUO_TARGETS_OF_IRS1_AND_IRS2 | 82 | 0.58 | 1.27 | 0.076 | 0.362 |
| 1589 | WP_EICOSANOID_METABOLISM_VIA_LIPOOXYGEI | 17 | 0.7 | 1.27 | 0.153 | 0.363 |
| 1590 | AHRARNT_01 | 128 | 0.55 | 1.27 | 0.066 | 0.363 |
| 1591 | WP_THERMOGENESIS | 95 | 0.56 | 1.27 | 0.086 | 0.363 |
| 1592 | HOELZEL_NF1_TARGETS_DN | 86 | 0.57 | 1.27 | 0.095 | 0.363 |
| 1593 | RIGGI_EWING_SARCOMA_PROGENITOR_UP | 311 | 0.53 | 1.27 | 0.021 | 0.364 |
| 1594 | GENTILE_UV_RESPONSE_CLUSTER_D2 | 35 | 0.62 | 1.27 | 0.138 | 0.364 |
| 1595 | REACTOME_EXTRA_NUCLEAR_ESTROGEN_SIGNA | 61 | 0.59 | 1.27 | 0.114 | 0.364 |
| 1596 | REACTOME_CYCLIN_A_CDK2_ASSOCIATED_EVEN | 85 | 0.56 | 1.27 | 0.082 | 0.364 |
| 1597 | REACTOME_TRANSLESION_SYNTHESIS_BY_Y_FA | 38 | 0.63 | 1.27 | 0.132 | 0.364 |
| 1598 | WP_AGERAGE_PATHWAY | 60 | 0.59 | 1.27 | 0.107 | 0.364 |
| 1599 | LI_AMPLIFIED_IN_LUNG_CANCER | 156 | 0.54 | 1.27 | 0.047 | 0.364 |
| 1600 | AP1_Q6_01 | 202 | 0.53 | 1.27 | 0.036 | 0.364 |
| 1601 | MIR3667_5P | 49 | 0.6 | 1.27 | 0.117 | 0.364 |
| 1602 | NR1H4_TARGET_GENES | 226 | 0.52 | 1.27 | 0.034 | 0.364 |
| 1603 | BREDEMEYER_RAG_SIGNALING_NOT_VIA_ATM_U | 50 | 0.6 | 1.27 | 0.114 | 0.364 |
| 1604 | SERVITJA_LIVER_HNF1A_TARGETS_UP | 95 | 0.56 | 1.27 | 0.084 | 0.363 |
| 1605 | GGCCAGT_MIR193A_MIR193B | 83 | 0.56 | 1.27 | 0.08 | 0.363 |
| 1606 | REACTOME_G_ALPHA_Q_SIGNALLING_EVENTS | 111 | 0.55 | 1.27 | 0.072 | 0.364 |
| 1607 | PID_ARF6_TRAFFICKING_PATHWAY | 37 | 0.63 | 1.27 | 0.121 | 0.364 |
| 1608 | ONGUSAHA_TP53_TARGETS | 31 | 0.65 | 1.27 | 0.153 | 0.364 |
| 1609 | PIONTEK_PKD1_TARGETS_UP | 27 | 0.66 | 1.27 | 0.144 | 0.364 |
| 1610 | REACTOME_FOXO_MEDIATED_TRANSCRIPTION | 51 | 0.6 | 1.27 | 0.12 | 0.365 |
| 1611 | WP_EPITHELIAL_TO_MESENCHYMAL_TRANSITION | 124 | 0.55 | 1.26 | 0.07 | 0.365 |
| 1612 | TBK1.DF_UP | 259 | 0.53 | 1.26 | 0.023 | 0.366 |
| 1613 | ROVERSI_GLIOMA_COPY_NUMBER_UP | 76 | 0.58 | 1.26 | 0.092 | 0.367 |
| 1614 | BROWNE_HCMV_INFECTION_24HR_UP | 128 | 0.54 | 1.26 | 0.073 | 0.368 |

| | | | | | | |
|---|---|---|---|---|---|---|
| 1615 | MAGRANGEAS_MULTIPLE_MYELOMA_IGLL_VS_IG | 31 | 0.65 | 1.26 | 0.131 | 0.369 |
| 1616 | REACTOME_NR1H2_AND_NR1H3_MEDIATED_SIGN | 39 | 0.62 | 1.26 | 0.126 | 0.369 |
| 1617 | UDAYAKUMAR_MED1_TARGETS_DN | 225 | 0.53 | 1.26 | 0.031 | 0.369 |
| 1618 | DACOSTA_UV_RESPONSE_VIA_ERCC3_XPCS_UP | 23 | 0.67 | 1.26 | 0.17 | 0.369 |
| 1619 | KEGG_VASOPRESSIN_REGULATED_WATER_REAE | 39 | 0.62 | 1.26 | 0.122 | 0.369 |
| 1620 | LE_EGR2_TARGETS_UP | 103 | 0.55 | 1.26 | 0.069 | 0.369 |
| 1621 | YAGI_AML_SURVIVAL | 107 | 0.55 | 1.26 | 0.077 | 0.369 |
| 1622 | PID_HNF3A_PATHWAY | 31 | 0.63 | 1.26 | 0.158 | 0.369 |
| 1623 | MIR4507 | 27 | 0.64 | 1.26 | 0.156 | 0.369 |
| 1624 | WILSON_PROTEASES_AT_TUMOR_BONE_INTERF/ | 16 | 0.73 | 1.26 | 0.173 | 0.369 |
| 1625 | PRAMOONJAGO_SOX4_TARGETS_UP | 48 | 0.6 | 1.26 | 0.129 | 0.37 |
| 1626 | REACTOME_COLLAGEN_BIOSYNTHESIS_AND_MO | 40 | 0.61 | 1.26 | 0.137 | 0.369 |
| 1627 | MEL18_DN.V1_DN | 113 | 0.55 | 1.26 | 0.077 | 0.37 |
| 1628 | MCBRYAN_PUBERTAL_BREAST_4_5WK_UP | 205 | 0.53 | 1.26 | 0.03 | 0.37 |
| 1629 | IKEDA_MIR1_TARGETS_UP | 52 | 0.59 | 1.26 | 0.108 | 0.37 |
| 1630 | MIR122B_5P | 179 | 0.53 | 1.26 | 0.053 | 0.372 |
| 1631 | REACTOME_CLASS_B_2_SECRETIN_FAMILY_RECI | 48 | 0.61 | 1.26 | 0.13 | 0.373 |
| 1632 | REACTOME_SIGNALING_BY_PDGFR_IN_DISEASE | 18 | 0.69 | 1.26 | 0.165 | 0.373 |
| 1633 | TOMIDA_METASTASIS_DN | 18 | 0.7 | 1.26 | 0.139 | 0.373 |
| 1634 | MIR6771_3P | 86 | 0.56 | 1.26 | 0.102 | 0.373 |
| 1635 | MIKKELSEN_ES_ICP_WITH_H3K27ME3 | 19 | 0.68 | 1.26 | 0.164 | 0.372 |
| 1636 | MED25_TARGET_GENES | 70 | 0.58 | 1.26 | 0.101 | 0.373 |
| 1637 | BONOME_OVARIAN_CANCER_SURVIVAL_OPTIMAL | 196 | 0.53 | 1.26 | 0.039 | 0.374 |
| 1638 | MIR3135A | 28 | 0.64 | 1.26 | 0.149 | 0.374 |
| 1639 | BROWNE_HCMV_INFECTION_8HR_DN | 35 | 0.63 | 1.26 | 0.15 | 0.374 |
| 1640 | BCL6B_TARGET_GENES | 139 | 0.54 | 1.26 | 0.06 | 0.375 |
| 1641 | MIR200A_3P | 355 | 0.52 | 1.26 | 0.024 | 0.375 |
| 1642 | NATSUME_RESPONSE_TO_INTERFERON_BETA_UI | 59 | 0.59 | 1.26 | 0.101 | 0.376 |
| 1643 | WP_VASOPRESSINREGULATED_WATER_REABSOI | 39 | 0.62 | 1.26 | 0.129 | 0.376 |
| 1644 | MIR6824_5P | 62 | 0.58 | 1.26 | 0.121 | 0.376 |
| 1645 | BROWNE_HCMV_INFECTION_12HR_UP | 103 | 0.55 | 1.26 | 0.084 | 0.376 |
| 1646 | HU_GENOTOXIC_DAMAGE_24HR | 35 | 0.63 | 1.26 | 0.138 | 0.377 |
| 1647 | MIKKELSEN_ES_HCP_WITH_H3K27ME3 | 25 | 0.65 | 1.26 | 0.167 | 0.377 |

| | | | | | | |
|---|---|---|---|---|---|---|
| 1648 | WP_WNT_SIGNALING_PATHWAY_AND_PLURIPOTE | 85 | 0.56 | 1.26 | 0.102 | 0.377 |
| 1649 | REACTOME_SIGNALING_BY_ERBB4 | 45 | 0.61 | 1.26 | 0.145 | 0.378 |
| 1650 | BENPORATH_ES_CORE_NINE_CORRELATED | 90 | 0.56 | 1.26 | 0.098 | 0.377 |
| 1651 | WP_MALE_INFERTILITY | 98 | 0.56 | 1.26 | 0.075 | 0.377 |
| 1652 | KEGG_O_GLYCAN_BIOSYNTHESIS | 22 | 0.68 | 1.26 | 0.153 | 0.379 |
| 1653 | MIR630 | 72 | 0.57 | 1.26 | 0.11 | 0.379 |
| 1654 | MCMURRAY_TP53_HRAS_COOPERATION_RESPON | 53 | 0.6 | 1.26 | 0.122 | 0.379 |
| 1655 | FONTAINE_PAPILLARY_THYROID_CARCINOMA_DN | 54 | 0.59 | 1.26 | 0.116 | 0.379 |
| 1656 | MIR12116 | 45 | 0.61 | 1.26 | 0.136 | 0.38 |
| 1657 | WP_INTERACTIONS_BETWEEN_LOXL4_AND_OXID | 17 | 0.71 | 1.26 | 0.172 | 0.38 |
| 1658 | PAX_Q6 | 197 | 0.53 | 1.26 | 0.037 | 0.381 |
| 1659 | CHEOK_RESPONSE_TO_MERCAPTOPURINE_DN | 16 | 0.71 | 1.26 | 0.173 | 0.381 |
| 1660 | TOMLINS_PROSTATE_CANCER_DN | 33 | 0.62 | 1.26 | 0.145 | 0.381 |
| 1661 | REACTOME_INTERLEUKIN_1_SIGNALING | 91 | 0.56 | 1.26 | 0.096 | 0.382 |
| 1662 | MIR3529_5P | 30 | 0.64 | 1.26 | 0.163 | 0.382 |
| 1663 | GAVIN_FOXP3_TARGETS_CLUSTER_T4 | 86 | 0.56 | 1.26 | 0.093 | 0.383 |
| 1664 | KATSANOU_ELAVL1_TARGETS_UP | 115 | 0.55 | 1.26 | 0.087 | 0.384 |
| 1665 | KATSANOU_ELAVL1_TARGETS_DN | 110 | 0.55 | 1.26 | 0.09 | 0.384 |
| 1666 | RSRFC4_Q2 | 152 | 0.54 | 1.25 | 0.058 | 0.385 |
| 1667 | LU_EZH2_TARGETS_UP | 261 | 0.52 | 1.25 | 0.038 | 0.385 |
| 1668 | MIR1265 | 38 | 0.61 | 1.25 | 0.144 | 0.385 |
| 1669 | DAVICIONI_TARGETS_OF_PAX_FOXO1_FUSIONS_ | 50 | 0.59 | 1.25 | 0.147 | 0.386 |
| 1670 | MIR4324 | 232 | 0.52 | 1.25 | 0.047 | 0.386 |
| 1671 | MCRS1_TARGET_GENES | 35 | 0.61 | 1.25 | 0.155 | 0.387 |
| 1672 | GU_PDEF_TARGETS_DN | 28 | 0.65 | 1.25 | 0.149 | 0.388 |
| 1673 | VANDESLUIS_COMMD1_TARGETS_GROUP_3_DN | 26 | 0.65 | 1.25 | 0.168 | 0.388 |
| 1674 | REACTOME_MAPK_FAMILY_SIGNALING_CASCADE | 268 | 0.52 | 1.25 | 0.039 | 0.388 |
| 1675 | ESC_J1_UP_LATE.V1_DN | 145 | 0.54 | 1.25 | 0.073 | 0.388 |
| 1676 | REACTOME_EGFR_DOWNREGULATION | 26 | 0.66 | 1.25 | 0.168 | 0.388 |
| 1677 | REACTOME_CELLULAR_SENESCENCE | 156 | 0.53 | 1.25 | 0.056 | 0.388 |
| 1678 | BREDEMEYER_RAG_SIGNALING_VIA_ATM_NOT_V | 35 | 0.62 | 1.25 | 0.137 | 0.388 |
| 1679 | MIR7978 | 76 | 0.56 | 1.25 | 0.109 | 0.389 |
| 1680 | TEL2_Q6 | 218 | 0.52 | 1.25 | 0.031 | 0.389 |

| | | | | | | |
|---|---|---|---|---|---|---|
| 1681 | HWANG_PROSTATE_CANCER_MARKERS | 24 | 0.67 | 1.25 | 0.175 | 0.389 |
| 1682 | MIR4663 | 55 | 0.58 | 1.25 | 0.129 | 0.389 |
| 1683 | SHETH_LIVER_CANCER_VS_TXNIP_LOSS_PAM5 | 77 | 0.56 | 1.25 | 0.109 | 0.39 |
| 1684 | BROWNE_HCMV_INFECTION_2HR_UP | 27 | 0.64 | 1.25 | 0.16 | 0.39 |
| 1685 | WP_INHIBITION_OF_EXOSOME_BIOGENESIS_AND | 17 | 0.69 | 1.25 | 0.173 | 0.391 |
| 1686 | BIOCARTA_TPO_PATHWAY | 21 | 0.68 | 1.25 | 0.165 | 0.39 |
| 1687 | DUNNE_TARGETS_OF_AML1_MTG8_FUSION_UP | 31 | 0.64 | 1.25 | 0.157 | 0.39 |
| 1688 | MIKKELSEN_ES_LCP_WITH_H3K4ME3 | 87 | 0.56 | 1.25 | 0.105 | 0.391 |
| 1689 | MIR4793_3P | 66 | 0.56 | 1.25 | 0.124 | 0.391 |
| 1690 | REACTOME_HOMOLOGY_DIRECTED_REPAIR | 118 | 0.54 | 1.25 | 0.075 | 0.391 |
| 1691 | BIOCARTA_AT1R_PATHWAY | 25 | 0.65 | 1.25 | 0.162 | 0.391 |
| 1692 | HNF1_01 | 147 | 0.53 | 1.25 | 0.068 | 0.391 |
| 1693 | REACTOME_VISUAL_PHOTOTRANSDUCTION | 58 | 0.58 | 1.25 | 0.126 | 0.392 |
| 1694 | GARY_CD5_TARGETS_UP | 447 | 0.51 | 1.25 | 0.011 | 0.392 |
| 1695 | PID_SHP2_PATHWAY | 48 | 0.59 | 1.25 | 0.132 | 0.392 |
| 1696 | MIR6762_5P | 51 | 0.59 | 1.25 | 0.134 | 0.392 |
| 1697 | DELLA_RESPONSE_TO_TSA_AND_BUTYRATE | 19 | 0.69 | 1.25 | 0.181 | 0.393 |
| 1698 | WP_MALIGNANT_PLEURAL_MESOTHELIOMA | 352 | 0.52 | 1.25 | 0.018 | 0.394 |
| 1699 | FOROUTAN_TGFB_EMT_UP | 148 | 0.54 | 1.25 | 0.076 | 0.394 |
| 1700 | WP_HOSTPATHOGEN_INTERACTION_OF_HUMAN_( | 20 | 0.67 | 1.25 | 0.167 | 0.394 |
| 1701 | KASLER_HDAC7_TARGETS_1_UP | 168 | 0.53 | 1.25 | 0.068 | 0.396 |
| 1702 | WP_INITIATION_OF_TRANSCRIPTION_AND_TRANS | 33 | 0.62 | 1.25 | 0.15 | 0.396 |
| 1703 | PID_ERBB1_INTERNALIZATION_PATHWAY | 40 | 0.6 | 1.25 | 0.15 | 0.397 |
| 1704 | REACTOME_SIGNALING_BY_TGF_BETA_RECEPTC | 71 | 0.57 | 1.25 | 0.125 | 0.396 |
| 1705 | MIR141_3P | 357 | 0.51 | 1.25 | 0.017 | 0.397 |
| 1706 | WP_TRANSCRIPTION_FACTOR_REGULATION_IN_A | 19 | 0.69 | 1.25 | 0.153 | 0.397 |
| 1707 | WP_MICRORNAS_IN_CARDIOMYOCYTE_HYPERTR( | 75 | 0.57 | 1.25 | 0.12 | 0.397 |
| 1708 | REACTOME_SIGNALING_BY_WNT | 262 | 0.52 | 1.25 | 0.043 | 0.397 |
| 1709 | PTEN_DN.V1_UP | 103 | 0.55 | 1.25 | 0.09 | 0.397 |
| 1710 | STAEGE_EWING_FAMILY_TUMOR | 19 | 0.67 | 1.25 | 0.194 | 0.397 |
| 1711 | PID_TAP63_PATHWAY | 50 | 0.59 | 1.25 | 0.153 | 0.399 |
| 1712 | MIR6845_5P | 54 | 0.58 | 1.25 | 0.125 | 0.399 |
| 1713 | BIOCARTA_CARM_ER_PATHWAY | 23 | 0.65 | 1.25 | 0.189 | 0.4 |

| | | | | | | |
|---|---|---|---|---|---|---|
| 1714 | MIR1238_5P_MIR4758_5P | 38 | 0.61 | 1.25 | 0.136 | 0.401 |
| 1715 | MIR711 | 35 | 0.61 | 1.25 | 0.158 | 0.402 |
| 1716 | REACTOME_NGF_STIMULATED_TRANSCRIPTION | 30 | 0.65 | 1.25 | 0.139 | 0.402 |
| 1717 | BROWNE_HCMV_INFECTION_30MIN_UP | 37 | 0.61 | 1.25 | 0.158 | 0.403 |
| 1718 | MIR4747_5P | 234 | 0.52 | 1.25 | 0.034 | 0.403 |
| 1719 | KEGG_GLYCEROPHOSPHOLIPID_METABOLISM | 60 | 0.57 | 1.25 | 0.118 | 0.402 |
| 1720 | MIR665 | 110 | 0.54 | 1.25 | 0.104 | 0.403 |
| 1721 | MIR143_3P_MIR4770 | 174 | 0.53 | 1.25 | 0.065 | 0.404 |
| 1722 | WP_SMALL_CELL_LUNG_CANCER | 85 | 0.56 | 1.25 | 0.108 | 0.404 |
| 1723 | VECCHI_GASTRIC_CANCER_EARLY_UP | 393 | 0.51 | 1.25 | 0.023 | 0.404 |
| 1724 | MIR6088 | 144 | 0.54 | 1.24 | 0.08 | 0.404 |
| 1725 | KEGG_MAPK_SIGNALING_PATHWAY | 212 | 0.52 | 1.24 | 0.058 | 0.404 |
| 1726 | NFY_Q6 | 212 | 0.52 | 1.24 | 0.048 | 0.404 |
| 1727 | AHR_01 | 61 | 0.58 | 1.24 | 0.129 | 0.405 |
| 1728 | KEGG_HYPERTROPHIC_CARDIOMYOPATHY_HCM | 54 | 0.58 | 1.24 | 0.147 | 0.406 |
| 1729 | VIETOR_IFRD1_TARGETS | 18 | 0.69 | 1.24 | 0.175 | 0.406 |
| 1730 | BIOCARTA_NGF_PATHWAY | 17 | 0.69 | 1.24 | 0.203 | 0.406 |
| 1731 | IVANOVA_HEMATOPOIESIS_STEM_CELL_SHORT_ | 16 | 0.69 | 1.24 | 0.188 | 0.406 |
| 1732 | E2F_01 | 65 | 0.57 | 1.24 | 0.124 | 0.406 |
| 1733 | BRUINS_UVC_RESPONSE_VIA_TP53_GROUP_B | 432 | 0.51 | 1.24 | 0.018 | 0.406 |
| 1734 | MIR219B_5P | 42 | 0.61 | 1.24 | 0.163 | 0.407 |
| 1735 | CHARAFE_BREAST_CANCER_BASAL_VS_MESENC | 93 | 0.56 | 1.24 | 0.104 | 0.407 |
| 1736 | WP_CANCER_IMMUNOTHERAPY_BY_PD1_BLOCKA | 16 | 0.7 | 1.24 | 0.177 | 0.407 |
| 1737 | WONG_ENDMETRIUM_CANCER_UP | 15 | 0.7 | 1.24 | 0.182 | 0.407 |
| 1738 | WINTER_HYPOXIA_METAGENE | 200 | 0.52 | 1.24 | 0.047 | 0.407 |
| 1739 | REACTOME_GLYOXYLATE_METABOLISM_AND_GL | 26 | 0.64 | 1.24 | 0.152 | 0.407 |
| 1740 | YAGI_AML_FAB_MARKERS | 159 | 0.53 | 1.24 | 0.075 | 0.407 |
| 1741 | MIR3913_5P | 70 | 0.56 | 1.24 | 0.129 | 0.407 |
| 1742 | YANG_BREAST_CANCER_ESR1_UP | 25 | 0.65 | 1.24 | 0.172 | 0.408 |
| 1743 | YAMASHITA_METHYLATED_IN_PROSTATE_CANCE | 25 | 0.66 | 1.24 | 0.165 | 0.408 |
| 1744 | BIOCARTA_P53_PATHWAY | 16 | 0.7 | 1.24 | 0.194 | 0.408 |
| 1745 | HIRSCH_CELLULAR_TRANSFORMATION_SIGNATUI | 92 | 0.55 | 1.24 | 0.118 | 0.409 |
| 1746 | MIR943 | 42 | 0.6 | 1.24 | 0.153 | 0.408 |

| | | | | | | |
|---|---|---|---|---|---|---|
| 1747 | KLEIN_PRIMARY_EFFUSION_LYMPHOMA_UP | 41 | 0.61 | 1.24 | 0.149 | 0.408 |
| 1748 | DOANE_RESPONSE_TO_ANDROGEN_DN | 218 | 0.52 | 1.24 | 0.047 | 0.409 |
| 1749 | MIR4645_5P | 103 | 0.55 | 1.24 | 0.108 | 0.409 |
| 1750 | REACTOME_TRANSCRIPTIONAL_REGULATION_BY_ | 32 | 0.63 | 1.24 | 0.163 | 0.409 |
| 1751 | HAHTOLA_MYCOSIS_FUNGOIDES_SKIN_UP | 158 | 0.53 | 1.24 | 0.084 | 0.409 |
| 1752 | PID_CMYB_PATHWAY | 62 | 0.58 | 1.24 | 0.13 | 0.41 |
| 1753 | REACTOME_FORMATION_OF_THE_CORNIFIED_EN | 40 | 0.61 | 1.24 | 0.149 | 0.41 |
| 1754 | PID_HIF2PATHWAY | 29 | 0.64 | 1.24 | 0.188 | 0.41 |
| 1755 | MA_MYELOID_DIFFERENTIATION_UP | 33 | 0.62 | 1.24 | 0.173 | 0.41 |
| 1756 | MIR4433B_3P | 27 | 0.64 | 1.24 | 0.156 | 0.41 |
| 1757 | REN_ALVEOLAR_RHABDOMYOSARCOMA_DN | 356 | 0.51 | 1.24 | 0.022 | 0.41 |
| 1758 | WP_MELANOMA | 63 | 0.58 | 1.24 | 0.141 | 0.41 |
| 1759 | ELVIDGE_HYPOXIA_BY_DMOG_UP | 118 | 0.54 | 1.24 | 0.095 | 0.41 |
| 1760 | REACTOME_KERATINIZATION | 40 | 0.61 | 1.24 | 0.145 | 0.41 |
| 1761 | REACTOME_DOWNREGULATION_OF_ERBB2_SIGN | 26 | 0.64 | 1.24 | 0.183 | 0.41 |
| 1762 | REACTOME_CYTOCHROME_P450_ARRANGED_BY_ | 25 | 0.65 | 1.24 | 0.166 | 0.41 |
| 1763 | MIR3940_5P | 26 | 0.65 | 1.24 | 0.176 | 0.41 |
| 1764 | DUTERTRE_ESTRADIOL_RESPONSE_24HR_DN | 444 | 0.51 | 1.24 | 0.012 | 0.41 |
| 1765 | DELACROIX_RARG_BOUND_MEF | 310 | 0.51 | 1.24 | 0.035 | 0.41 |
| 1766 | HAHTOLA_MYCOSIS_FUNGOIDES_CD4_UP | 43 | 0.6 | 1.24 | 0.168 | 0.412 |
| 1767 | SCHUETZ_BREAST_CANCER_DUCTAL_INVASIVE_[illegible] | 67 | 0.57 | 1.24 | 0.143 | 0.412 |
| 1768 | WP_NEUROINFLAMMATION_AND_GLUTAMATERGI[illegible] | 107 | 0.54 | 1.24 | 0.095 | 0.412 |
| 1769 | PID_PDGFRA_PATHWAY | 21 | 0.67 | 1.24 | 0.171 | 0.412 |
| 1770 | ROSS_AML_WITH_AML1_ETO_FUSION | 62 | 0.57 | 1.24 | 0.109 | 0.412 |
| 1771 | CEBPB_01 | 211 | 0.52 | 1.24 | 0.056 | 0.412 |
| 1772 | GENTILE_UV_LOW_DOSE_DN | 60 | 0.57 | 1.24 | 0.15 | 0.412 |
| 1773 | BOCHKIS_FOXA2_TARGETS | 334 | 0.51 | 1.24 | 0.023 | 0.413 |
| 1774 | ASTIER_INTEGRIN_SIGNALING | 48 | 0.59 | 1.24 | 0.14 | 0.413 |
| 1775 | MIR4305 | 52 | 0.58 | 1.24 | 0.129 | 0.413 |
| 1776 | JAK2_DN.V1_UP | 86 | 0.55 | 1.24 | 0.126 | 0.413 |
| 1777 | MIR6819_5P | 65 | 0.58 | 1.24 | 0.129 | 0.414 |
| 1778 | MULLIGAN_NTF3_SIGNALING_VIA_INSR_AND_IGF[illegible] | 19 | 0.68 | 1.24 | 0.192 | 0.414 |
| 1779 | KOKKINAKIS_METHIONINE_DEPRIVATION_96HR_[illegible] | 68 | 0.57 | 1.24 | 0.129 | 0.414 |

| | | | | | | |
|---|---|---|---|---|---|---|
| 1780 | MIR8062 | 93 | 0.55 | 1.24 | 0.102 | 0.414 |
| 1781 | YAO_TEMPORAL_RESPONSE_TO_PROGESTERON | 61 | 0.57 | 1.24 | 0.145 | 0.414 |
| 1782 | ZHAN_MULTIPLE_MYELOMA_CD1_AND_CD2_DN | 53 | 0.58 | 1.24 | 0.147 | 0.414 |
| 1783 | PR_02 | 105 | 0.55 | 1.24 | 0.103 | 0.414 |
| 1784 | WNT_SIGNALING | 70 | 0.56 | 1.24 | 0.129 | 0.414 |
| 1785 | MIR4423_5P | 23 | 0.66 | 1.24 | 0.191 | 0.414 |
| 1786 | MIR548B_3P | 95 | 0.55 | 1.24 | 0.107 | 0.415 |
| 1787 | LIU_TARGETS_OF_VMYB_VS_CMYB_DN | 27 | 0.64 | 1.24 | 0.175 | 0.414 |
| 1788 | MIR670_5P | 216 | 0.52 | 1.24 | 0.048 | 0.414 |
| 1789 | REACTOME_REGULATION_OF_HMOX1_EXPRESSI | 63 | 0.57 | 1.24 | 0.131 | 0.415 |
| 1790 | BRACHAT_RESPONSE_TO_CAMPTOTHECIN_UP | 23 | 0.66 | 1.24 | 0.173 | 0.415 |
| 1791 | REACTOME_O_LINKED_GLYCOSYLATION_OF_MU | 39 | 0.6 | 1.24 | 0.158 | 0.415 |
| 1792 | REACTOME_SENSORY_PERCEPTION | 139 | 0.53 | 1.24 | 0.094 | 0.416 |
| 1793 | BAKKER_FOXO3_TARGETS_DN | 149 | 0.53 | 1.24 | 0.072 | 0.416 |
| 1794 | JAZAG_TGFB1_SIGNALING_UP | 88 | 0.55 | 1.24 | 0.117 | 0.416 |
| 1795 | MIR6886_3P | 79 | 0.56 | 1.24 | 0.117 | 0.416 |
| 1796 | MIR761 | 207 | 0.52 | 1.24 | 0.068 | 0.416 |
| 1797 | MIR4496 | 249 | 0.52 | 1.24 | 0.046 | 0.416 |
| 1798 | MIKKELSEN_MEF_ICP_WITH_H3K27ME3 | 109 | 0.53 | 1.24 | 0.104 | 0.417 |
| 1799 | REACTOME_RHOJ_GTPASE_CYCLE | 50 | 0.58 | 1.24 | 0.146 | 0.417 |
| 1800 | REACTOME_TRIGLYCERIDE_METABOLISM | 22 | 0.67 | 1.24 | 0.195 | 0.417 |
| 1801 | MIR3679_5P | 58 | 0.57 | 1.24 | 0.149 | 0.418 |
| 1802 | DELPUECH_FOXO3_TARGETS_DN | 38 | 0.61 | 1.24 | 0.156 | 0.418 |
| 1803 | GYORFFY_DOXORUBICIN_RESISTANCE | 26 | 0.65 | 1.24 | 0.17 | 0.418 |
| 1804 | MIR3201 | 71 | 0.56 | 1.23 | 0.137 | 0.419 |
| 1805 | SHIN_B_CELL_LYMPHOMA_CLUSTER_3 | 22 | 0.65 | 1.23 | 0.189 | 0.419 |
| 1806 | MIR4720_3P | 91 | 0.55 | 1.23 | 0.101 | 0.419 |
| 1807 | REACTOME_DNA_DAMAGE_BYPASS | 47 | 0.6 | 1.23 | 0.134 | 0.419 |
| 1808 | MIR4782_5P | 137 | 0.53 | 1.23 | 0.083 | 0.419 |
| 1809 | MIR6511B_5P | 63 | 0.57 | 1.23 | 0.167 | 0.42 |
| 1810 | MIR4662A_3P | 54 | 0.58 | 1.23 | 0.155 | 0.419 |
| 1811 | CASORELLI_APL_SECONDARY_VS_DE_NOVO_UP | 26 | 0.64 | 1.23 | 0.17 | 0.421 |
| 1812 | DAZARD_UV_RESPONSE_CLUSTER_G1 | 52 | 0.59 | 1.23 | 0.134 | 0.422 |

| | | | | | | |
|---|---|---|---|---|---|---|
| 1813 | HOXA7_TARGET_GENES | 102 | 0.54 | 1.23 | 0.108 | 0.422 |
| 1814 | WP_LNCRNA_IN_CANONICAL_WNT_SIGNALING_AN | 77 | 0.56 | 1.23 | 0.134 | 0.424 |
| 1815 | DASU_IL6_SIGNALING_SCAR_DN | 15 | 0.69 | 1.23 | 0.206 | 0.424 |
| 1816 | SCIBETTA_KDM5B_TARGETS_DN | 77 | 0.56 | 1.23 | 0.122 | 0.424 |
| 1817 | HAN_SATB1_TARGETS_UP | 350 | 0.5 | 1.23 | 0.022 | 0.424 |
| 1818 | BURTON_ADIPOGENESIS_PEAK_AT_16HR | 37 | 0.61 | 1.23 | 0.18 | 0.424 |
| 1819 | REACTOME_CYCLIN_A_B1_B2_ASSOCIATED_EVE | 25 | 0.64 | 1.23 | 0.173 | 0.424 |
| 1820 | REACTOME_NUCLEAR_SIGNALING_BY_ERBB4 | 23 | 0.65 | 1.23 | 0.198 | 0.424 |
| 1821 | BOYLAN_MULTIPLE_MYELOMA_C_CLUSTER_UP | 34 | 0.62 | 1.23 | 0.183 | 0.424 |
| 1822 | PASINI_SUZ12_TARGETS_UP | 83 | 0.55 | 1.23 | 0.126 | 0.425 |
| 1823 | WP_EGFR_TYROSINE_KINASE_INHIBITOR_RESIST | 78 | 0.56 | 1.23 | 0.137 | 0.426 |
| 1824 | PID_VEGFR1_PATHWAY | 25 | 0.64 | 1.23 | 0.18 | 0.427 |
| 1825 | REACTOME_SENSORY_PROCESSING_OF_SOUND | 61 | 0.57 | 1.23 | 0.142 | 0.426 |
| 1826 | MIR4524A_3P | 194 | 0.52 | 1.23 | 0.074 | 0.426 |
| 1827 | ZHAN_MULTIPLE_MYELOMA_DN | 23 | 0.65 | 1.23 | 0.173 | 0.426 |
| 1828 | GAUSSMANN_MLL_AF4_FUSION_TARGETS_C_DN | 16 | 0.69 | 1.23 | 0.202 | 0.427 |
| 1829 | SHETH_LIVER_CANCER_VS_TXNIP_LOSS_PAM4 | 199 | 0.52 | 1.23 | 0.076 | 0.427 |
| 1830 | GRADE_COLON_VS_RECTAL_CANCER_UP | 31 | 0.62 | 1.23 | 0.185 | 0.427 |
| 1831 | BAUS_TFF2_TARGETS_UP | 16 | 0.69 | 1.23 | 0.199 | 0.429 |
| 1832 | FARMER_BREAST_CANCER_APOCRINE_VS_LUMIN | 256 | 0.51 | 1.23 | 0.047 | 0.428 |
| 1833 | HNF1_Q6 | 145 | 0.53 | 1.23 | 0.093 | 0.429 |
| 1834 | MIR3689A_3P_MIR3689B_3P_MIR3689C | 137 | 0.53 | 1.23 | 0.094 | 0.429 |
| 1835 | BIOCARTA_RACCYCD_PATHWAY | 26 | 0.63 | 1.23 | 0.196 | 0.429 |
| 1836 | JAZAERI_BREAST_CANCER_BRCA1_VS_BRCA2_U | 44 | 0.59 | 1.23 | 0.164 | 0.429 |
| 1837 | MIR3616_5P | 70 | 0.56 | 1.23 | 0.131 | 0.429 |
| 1838 | MIR379_3P_MIR411_3P | 150 | 0.52 | 1.23 | 0.085 | 0.429 |
| 1839 | KAYO_CALORIE_RESTRICTION_MUSCLE_UP | 67 | 0.57 | 1.23 | 0.149 | 0.429 |
| 1840 | WP_MAPK_PATHWAY_IN_CONGENITAL_THYROID_ | 16 | 0.67 | 1.23 | 0.231 | 0.43 |
| 1841 | JAIN_NFKB_SIGNALING | 69 | 0.56 | 1.23 | 0.153 | 0.43 |
| 1842 | MIR518A_5P_MIR527 | 228 | 0.51 | 1.23 | 0.063 | 0.43 |
| 1843 | MIR6877_3P | 30 | 0.63 | 1.23 | 0.189 | 0.43 |
| 1844 | MIR197_5P | 58 | 0.57 | 1.23 | 0.134 | 0.43 |
| 1845 | GCNF_01 | 59 | 0.57 | 1.23 | 0.144 | 0.431 |

| | | | | | | |
|---|---|---|---|---|---|---|
| 1846 | MIR5192 | 41 | 0.61 | 1.23 | 0.166 | 0.431 |
| 1847 | MIR6851_3P | 53 | 0.58 | 1.23 | 0.17 | 0.432 |
| 1848 | IRF1_Q6 | 205 | 0.51 | 1.23 | 0.07 | 0.432 |
| 1849 | REACTOME_TIGHT_JUNCTION_INTERACTIONS | 19 | 0.67 | 1.23 | 0.191 | 0.432 |
| 1850 | REACTOME_MYD88_INDEPENDENT_TLR4_CASCAD | 91 | 0.55 | 1.23 | 0.104 | 0.432 |
| 1851 | REACTOME_REGULATION_OF_RUNX2_EXPRESSIO | 66 | 0.56 | 1.23 | 0.132 | 0.432 |
| 1852 | WP_BLADDER_CANCER | 39 | 0.6 | 1.23 | 0.174 | 0.433 |
| 1853 | SASSON_RESPONSE_TO_FORSKOLIN_UP | 77 | 0.55 | 1.23 | 0.126 | 0.433 |
| 1854 | SABATES_COLORECTAL_ADENOMA_SIZE_UP | 22 | 0.65 | 1.23 | 0.206 | 0.434 |
| 1855 | ILK_TARGET_GENES | 17 | 0.68 | 1.23 | 0.191 | 0.434 |
| 1856 | MANALO_HYPOXIA_UP | 175 | 0.52 | 1.23 | 0.066 | 0.434 |
| 1857 | BYSTROEM_CORRELATED_WITH_IL5_UP | 29 | 0.63 | 1.23 | 0.199 | 0.434 |
| 1858 | WP_EXTRACELLULAR_VESICLES_IN_THE_CROSS | 16 | 0.68 | 1.23 | 0.216 | 0.434 |
| 1859 | REACTOME_REGULATION_OF_RUNX1_EXPRESSIO | 17 | 0.68 | 1.23 | 0.199 | 0.436 |
| 1860 | LEF1_UP.V1_DN | 146 | 0.53 | 1.23 | 0.085 | 0.436 |
| 1861 | MIR4673 | 125 | 0.53 | 1.23 | 0.109 | 0.436 |
| 1862 | SEIDEN_ONCOGENESIS_BY_MET | 86 | 0.54 | 1.23 | 0.147 | 0.436 |
| 1863 | WP_EGFEGFR_SIGNALING_PATHWAY | 153 | 0.52 | 1.23 | 0.097 | 0.438 |
| 1864 | REACTOME_TOLL_LIKE_RECEPTOR_TLR1_TLR2_( | 87 | 0.54 | 1.23 | 0.12 | 0.438 |
| 1865 | TGTGTGA_MIR377 | 179 | 0.52 | 1.23 | 0.088 | 0.438 |
| 1866 | MIR4469 | 95 | 0.54 | 1.23 | 0.13 | 0.438 |
| 1867 | WP_INFLUENCE_OF_LAMINOPATHIES_ON_WNT_S | 32 | 0.61 | 1.23 | 0.177 | 0.438 |
| 1868 | TURASHVILI_BREAST_DUCTAL_CARCINOMA_VS_L | 65 | 0.57 | 1.23 | 0.162 | 0.438 |
| 1869 | SASSON_RESPONSE_TO_GONADOTROPHINS_UP | 75 | 0.56 | 1.23 | 0.138 | 0.438 |
| 1870 | PID_RAS_PATHWAY | 28 | 0.63 | 1.23 | 0.184 | 0.437 |
| 1871 | KIM_MYC_AMPLIFICATION_TARGETS_DN | 87 | 0.54 | 1.22 | 0.128 | 0.438 |
| 1872 | MIR4639_5P | 78 | 0.56 | 1.22 | 0.146 | 0.438 |
| 1873 | HSIAO_LIVER_SPECIFIC_GENES | 103 | 0.54 | 1.22 | 0.115 | 0.439 |
| 1874 | RAY_TUMORIGENESIS_BY_ERBB2_CDC25A_DN | 230 | 0.51 | 1.22 | 0.068 | 0.44 |
| 1875 | MIR18A_3P | 77 | 0.56 | 1.22 | 0.122 | 0.439 |
| 1876 | LEIN_MEDULLA_MARKERS | 64 | 0.57 | 1.22 | 0.155 | 0.439 |
| 1877 | MIR3189_5P | 43 | 0.59 | 1.22 | 0.176 | 0.44 |
| 1878 | HMCES_TARGET_GENES | 20 | 0.66 | 1.22 | 0.201 | 0.44 |

| | | | | | | |
|---|---|---|---|---|---|---|
| 1879 | HAHTOLA_SEZARY_SYNDROM_UP | 71 | 0.55 | 1.22 | 0.13 | 0.441 |
| 1880 | GCNP_SHH_UP_LATE.V1_UP | 163 | 0.52 | 1.22 | 0.087 | 0.441 |
| 1881 | BROWNE_HCMV_INFECTION_18HR_DN | 137 | 0.53 | 1.22 | 0.11 | 0.441 |
| 1882 | PID_HDAC_CLASSII_PATHWAY | 31 | 0.63 | 1.22 | 0.182 | 0.441 |
| 1883 | SESTO_RESPONSE_TO_UV_C5 | 43 | 0.59 | 1.22 | 0.172 | 0.443 |
| 1884 | ZHANG_GATA6_TARGETS_DN | 54 | 0.58 | 1.22 | 0.146 | 0.443 |
| 1885 | MIR12121 | 18 | 0.67 | 1.22 | 0.179 | 0.443 |
| 1886 | BIOCARTA_IL1R_PATHWAY | 24 | 0.65 | 1.22 | 0.206 | 0.443 |
| 1887 | FLECHNER_PBL_KIDNEY_TRANSPLANT_REJECTEI | 61 | 0.57 | 1.22 | 0.164 | 0.443 |
| 1888 | SHEDDEN_LUNG_CANCER_GOOD_SURVIVAL_A12 | 205 | 0.51 | 1.22 | 0.077 | 0.444 |
| 1889 | CCCNNNNNNAAGWT_UNKNOWN | 93 | 0.54 | 1.22 | 0.122 | 0.444 |
| 1890 | HELLER_SILENCED_BY_METHYLATION_DN | 87 | 0.53 | 1.22 | 0.122 | 0.446 |
| 1891 | WOO_LIVER_CANCER_RECURRENCE_UP | 84 | 0.55 | 1.22 | 0.15 | 0.446 |
| 1892 | BLANCO_MELO_MERS_COV_INFECTION_MCR5_CE | 220 | 0.51 | 1.22 | 0.072 | 0.446 |
| 1893 | PEDERSEN_METASTASIS_BY_ERBB2_ISOFORM_1 | 34 | 0.62 | 1.22 | 0.165 | 0.446 |
| 1894 | WP_SIGNAL_TRANSDUCTION_THROUGH_IL1R | 26 | 0.65 | 1.22 | 0.211 | 0.445 |
| 1895 | BYSTRYKH_HEMATOPOIESIS_STEM_CELL_SCP2_ | 16 | 0.69 | 1.22 | 0.219 | 0.446 |
| 1896 | MIR5196_5P | 245 | 0.51 | 1.22 | 0.063 | 0.447 |
| 1897 | REACTOME_SIGNAL_TRANSDUCTION_BY_L1 | 20 | 0.66 | 1.22 | 0.205 | 0.448 |
| 1898 | KEGG_EPITHELIAL_CELL_SIGNALING_IN_HELICOI | 60 | 0.57 | 1.22 | 0.149 | 0.448 |
| 1899 | PID_LKB1_PATHWAY | 44 | 0.59 | 1.22 | 0.165 | 0.448 |
| 1900 | REACTOME_CARGO_RECOGNITION_FOR_CLATHR | 83 | 0.54 | 1.22 | 0.144 | 0.448 |
| 1901 | EBAUER_MYOGENIC_TARGETS_OF_PAX3_FOXO1 | 31 | 0.62 | 1.22 | 0.192 | 0.448 |
| 1902 | MIR8060 | 83 | 0.54 | 1.22 | 0.141 | 0.448 |
| 1903 | TAANNYSGCG_UNKNOWN | 77 | 0.55 | 1.22 | 0.142 | 0.448 |
| 1904 | MIR6841_3P | 42 | 0.59 | 1.22 | 0.197 | 0.448 |
| 1905 | PECE_MAMMARY_STEM_CELL_DN | 130 | 0.53 | 1.22 | 0.1 | 0.448 |
| 1906 | KEGG_WNT_SIGNALING_PATHWAY | 125 | 0.53 | 1.22 | 0.119 | 0.449 |
| 1907 | LU_TUMOR_ANGIOGENESIS_UP | 24 | 0.65 | 1.22 | 0.203 | 0.449 |
| 1908 | WATANABE_RECTAL_CANCER_RADIOTHERAPY_R | 81 | 0.54 | 1.22 | 0.149 | 0.45 |
| 1909 | VARELA_ZMPSTE24_TARGETS_DN | 23 | 0.63 | 1.22 | 0.211 | 0.45 |
| 1910 | REACTOME_RHOV_GTPASE_CYCLE | 37 | 0.6 | 1.22 | 0.182 | 0.449 |
| 1911 | SNIJDERS_AMPLIFIED_IN_HEAD_AND_NECK_TUM( | 32 | 0.61 | 1.22 | 0.189 | 0.45 |

| | | | | | | |
|---|---|---|---|---|---|---|
| 1912 | CCATCCA_MIR432 | 51 | 0.58 | 1.22 | 0.155 | 0.45 |
| 1913 | BIOCARTA_MITOCHONDRIA_PATHWAY | 19 | 0.67 | 1.22 | 0.213 | 0.45 |
| 1914 | CTCTATG_MIR368 | 34 | 0.61 | 1.22 | 0.186 | 0.45 |
| 1915 | ENK_UV_RESPONSE_KERATINOCYTE_DN | 459 | 0.5 | 1.22 | 0.022 | 0.452 |
| 1916 | MANNE_COVID19_ICU_VS_HEALTHY_DONOR_PLA | 102 | 0.53 | 1.22 | 0.133 | 0.452 |
| 1917 | HU_ANGIOGENESIS_UP | 19 | 0.67 | 1.22 | 0.196 | 0.453 |
| 1918 | PID_ERBB1_RECEPTOR_PROXIMAL_PATHWAY | 35 | 0.6 | 1.22 | 0.187 | 0.453 |
| 1919 | REACTOME_PCP_CE_PATHWAY | 86 | 0.55 | 1.22 | 0.135 | 0.454 |
| 1920 | AKT_UP.V1_UP | 127 | 0.52 | 1.22 | 0.113 | 0.454 |
| 1921 | ICHIBA_GRAFT_VERSUS_HOST_DISEASE_D7_DN | 22 | 0.65 | 1.22 | 0.19 | 0.454 |
| 1922 | SESTO_RESPONSE_TO_UV_C6 | 35 | 0.6 | 1.22 | 0.173 | 0.454 |
| 1923 | MIR6782_5P | 29 | 0.61 | 1.22 | 0.213 | 0.454 |
| 1924 | BOYLAN_MULTIPLE_MYELOMA_C_D_DN | 158 | 0.52 | 1.22 | 0.097 | 0.454 |
| 1925 | REACTOME_PROGRAMMED_CELL_DEATH | 187 | 0.51 | 1.22 | 0.086 | 0.455 |
| 1926 | MIR3132 | 63 | 0.56 | 1.22 | 0.157 | 0.455 |
| 1927 | FIGUEROA_AML_METHYLATION_CLUSTER_6_DN | 20 | 0.65 | 1.22 | 0.216 | 0.455 |
| 1928 | KONDO_PROSTATE_CANCER_HCP_WITH_H3K27M | 67 | 0.55 | 1.22 | 0.153 | 0.455 |
| 1929 | MIR4463 | 67 | 0.56 | 1.22 | 0.133 | 0.455 |
| 1930 | MIR4289 | 59 | 0.57 | 1.22 | 0.162 | 0.456 |
| 1931 | POOLA_INVASIVE_BREAST_CANCER_DN | 92 | 0.53 | 1.22 | 0.14 | 0.456 |
| 1932 | SEITZ_NEOPLASTIC_TRANSFORMATION_BY_8P_D | 26 | 0.63 | 1.22 | 0.207 | 0.456 |
| 1933 | KEGG_SMALL_CELL_LUNG_CANCER | 76 | 0.55 | 1.22 | 0.146 | 0.457 |
| 1934 | MIR619_3P | 47 | 0.57 | 1.22 | 0.18 | 0.457 |
| 1935 | KEGG_ETHER_LIPID_METABOLISM | 18 | 0.67 | 1.21 | 0.234 | 0.457 |
| 1936 | GERY_CEBP_TARGETS | 99 | 0.53 | 1.21 | 0.128 | 0.457 |
| 1937 | ATF2_S_UP.V1_UP | 125 | 0.53 | 1.21 | 0.121 | 0.458 |
| 1938 | HOQUE_METHYLATED_IN_CANCER | 40 | 0.6 | 1.21 | 0.184 | 0.458 |
| 1939 | RGAANNTTC_HSF1_01 | 360 | 0.5 | 1.21 | 0.038 | 0.458 |
| 1940 | REACTOME_ABERRANT_REGULATION_OF_MITOTI | 17 | 0.67 | 1.21 | 0.224 | 0.459 |
| 1941 | WP_TGFBETA_RECEPTOR_SIGNALING | 41 | 0.6 | 1.21 | 0.2 | 0.459 |
| 1942 | ACEVEDO_LIVER_CANCER_WITH_H3K27ME3_DN | 136 | 0.52 | 1.21 | 0.106 | 0.459 |
| 1943 | MORI_SMALL_PRE_BII_LYMPHOCYTE_DN | 63 | 0.56 | 1.21 | 0.163 | 0.46 |
| 1944 | GRESHOCK_CANCER_COPY_NUMBER_UP | 281 | 0.5 | 1.21 | 0.062 | 0.46 |

| | | | | | | |
|---|---|---|---|---|---|---|
| 1945 | PID_AURORA_A_PATHWAY | 31 | 0.61 | 1.21 | 0.193 | 0.461 |
| 1946 | WP_EMBRYONIC_STEM_CELL_PLURIPOTENCY_PA | 87 | 0.54 | 1.21 | 0.151 | 0.461 |
| 1947 | NABA_ECM_REGULATORS | 107 | 0.53 | 1.21 | 0.124 | 0.461 |
| 1948 | GAUSSMANN_MLL_AF4_FUSION_TARGETS_C_UP | 151 | 0.52 | 1.21 | 0.102 | 0.462 |
| 1949 | MIR7851_3P | 51 | 0.58 | 1.21 | 0.164 | 0.462 |
| 1950 | XU_CREBBP_TARGETS_UP | 20 | 0.66 | 1.21 | 0.23 | 0.462 |
| 1951 | DURAND_STROMA_NS_UP | 134 | 0.52 | 1.21 | 0.121 | 0.461 |
| 1952 | SCHAEFFER_PROSTATE_DEVELOPMENT_48HR_U | 331 | 0.5 | 1.21 | 0.044 | 0.461 |
| 1953 | WP_IL17_SIGNALING_PATHWAY | 24 | 0.65 | 1.21 | 0.219 | 0.462 |
| 1954 | MIR378J | 70 | 0.55 | 1.21 | 0.158 | 0.462 |
| 1955 | WP_TYPE_II_DIABETES_MELLITUS | 18 | 0.66 | 1.21 | 0.215 | 0.462 |
| 1956 | WP_NAD_METABOLISM_IN_ONCOGENEINDUCED_S | 23 | 0.64 | 1.21 | 0.195 | 0.462 |
| 1957 | SIMBULAN_UV_RESPONSE_NORMAL_DN | 28 | 0.63 | 1.21 | 0.195 | 0.464 |
| 1958 | CHOW_RASSF1_TARGETS_UP | 26 | 0.63 | 1.21 | 0.206 | 0.465 |
| 1959 | PURBEY_TARGETS_OF_CTBP1_AND_SATB1_DN | 135 | 0.52 | 1.21 | 0.115 | 0.465 |
| 1960 | TRAYNOR_RETT_SYNDROM_UP | 25 | 0.64 | 1.21 | 0.2 | 0.465 |
| 1961 | JINESH_BLEBBISHIELD_TRANSFORMED_STEM_CE | 282 | 0.5 | 1.21 | 0.052 | 0.466 |
| 1962 | GAVIN_FOXP3_TARGETS_CLUSTER_P4 | 76 | 0.54 | 1.21 | 0.136 | 0.466 |
| 1963 | REACTOME_SLC_MEDIATED_TRANSMEMBRANE_T | 163 | 0.51 | 1.21 | 0.101 | 0.466 |
| 1964 | PRC2_SUZ12_UP.V1_UP | 127 | 0.52 | 1.21 | 0.126 | 0.467 |
| 1965 | WP_KISSPEPTINKISSPEPTIN_RECEPTOR_SYSTEM | 32 | 0.61 | 1.21 | 0.206 | 0.467 |
| 1966 | ZNF512_TARGET_GENES | 299 | 0.5 | 1.21 | 0.049 | 0.466 |
| 1967 | BIOCARTA_KERATINOCYTE_PATHWAY | 43 | 0.58 | 1.21 | 0.201 | 0.466 |
| 1968 | REACTOME_HDMS_DEMETHYLATE_HISTONES | 32 | 0.62 | 1.21 | 0.21 | 0.466 |
| 1969 | GLI1_UP.V1_UP | 18 | 0.66 | 1.21 | 0.21 | 0.466 |
| 1970 | IZADPANAH_STEM_CELL_ADIPOSE_VS_BONE_UP | 107 | 0.53 | 1.21 | 0.116 | 0.467 |
| 1971 | SWEET_LUNG_CANCER_KRAS_UP | 390 | 0.5 | 1.21 | 0.037 | 0.467 |
| 1972 | HOXC13_TARGET_GENES | 104 | 0.53 | 1.21 | 0.131 | 0.468 |
| 1973 | CHIARADONNA_NEOPLASTIC_TRANSFORMATION_ | 107 | 0.53 | 1.21 | 0.126 | 0.468 |
| 1974 | MIR6502_5P | 45 | 0.58 | 1.21 | 0.183 | 0.468 |
| 1975 | MIR1249_5P | 168 | 0.51 | 1.21 | 0.107 | 0.468 |
| 1976 | WILLIAMS_ESR2_TARGETS_UP | 24 | 0.64 | 1.21 | 0.201 | 0.468 |
| 1977 | FISCHER_G2_M_CELL_CYCLE | 230 | 0.5 | 1.21 | 0.082 | 0.468 |

| | | | | | | |
|---|---|---|---|---|---|---|
| 1978 | TBK1.DN.48HRS_DN | 44 | 0.58 | 1.21 | 0.184 | 0.468 |
| 1979 | SUBTIL_PROGESTIN_TARGETS | 35 | 0.6 | 1.21 | 0.195 | 0.469 |
| 1980 | MIR6819_3P | 31 | 0.61 | 1.21 | 0.189 | 0.468 |
| 1981 | MIR3927_5P | 37 | 0.6 | 1.21 | 0.18 | 0.468 |
| 1982 | MIR4723_5P | 172 | 0.51 | 1.21 | 0.09 | 0.469 |
| 1983 | KANG_IMMORTALIZED_BY_TERT_UP | 67 | 0.56 | 1.21 | 0.172 | 0.469 |
| 1984 | CEBP_C | 161 | 0.51 | 1.21 | 0.087 | 0.47 |
| 1985 | MIR1245A | 32 | 0.61 | 1.21 | 0.216 | 0.47 |
| 1986 | VERRECCHIA_RESPONSE_TO_TGFB1_C5 | 17 | 0.67 | 1.21 | 0.232 | 0.472 |
| 1987 | KYNG_DNA_DAMAGE_BY_4NQO | 27 | 0.62 | 1.21 | 0.193 | 0.472 |
| 1988 | REACTOME_SHC1_EVENTS_IN_ERBB2_SIGNALIN( | 20 | 0.66 | 1.21 | 0.22 | 0.472 |
| 1989 | MATSUDA_NATURAL_KILLER_DIFFERENTIATION | 374 | 0.49 | 1.21 | 0.034 | 0.472 |
| 1990 | SIG_IL4RECEPTOR_IN_B_LYPHOCYTES | 26 | 0.62 | 1.21 | 0.223 | 0.472 |
| 1991 | PARK_HSC_MARKERS | 38 | 0.59 | 1.21 | 0.193 | 0.473 |
| 1992 | KEGG_PROTEASOME | 41 | 0.6 | 1.21 | 0.186 | 0.473 |
| 1993 | WP_OLIGODENDROCYTE_SPECIFICATION_AND_DI | 15 | 0.69 | 1.21 | 0.259 | 0.473 |
| 1994 | MARTORIATI_MDM4_TARGETS_NEUROEPITHELIUM | 146 | 0.51 | 1.21 | 0.106 | 0.473 |
| 1995 | MIR648 | 42 | 0.58 | 1.21 | 0.192 | 0.473 |
| 1996 | MIR4664_5P | 62 | 0.56 | 1.21 | 0.175 | 0.473 |
| 1997 | BROWNE_HCMV_INFECTION_24HR_DN | 115 | 0.53 | 1.21 | 0.135 | 0.473 |
| 1998 | MIR7110_3P | 339 | 0.5 | 1.21 | 0.056 | 0.474 |
| 1999 | REACTOME_SIGNALING_BY_RECEPTOR_TYROSIN | 429 | 0.49 | 1.21 | 0.046 | 0.474 |
| 2000 | GTF2A2_TARGET_GENES | 424 | 0.49 | 1.21 | 0.04 | 0.475 |
| 2001 | PKCA_DN.V1_UP | 96 | 0.53 | 1.21 | 0.133 | 0.475 |
| 2002 | MIR320A_5P | 89 | 0.53 | 1.21 | 0.149 | 0.475 |
| 2003 | REACTOME_CELL_JUNCTION_ORGANIZATION | 72 | 0.55 | 1.21 | 0.158 | 0.475 |
| 2004 | TARTE_PLASMA_CELL_VS_B_LYMPHOCYTE_UP | 69 | 0.56 | 1.21 | 0.153 | 0.475 |
| 2005 | ZNF746_TARGET_GENES | 63 | 0.55 | 1.21 | 0.177 | 0.475 |
| 2006 | REACTOME_DNA_DOUBLE_STRAND_BREAK_RESI | 59 | 0.57 | 1.21 | 0.17 | 0.475 |
| 2007 | KEGG_NEUROTROPHIN_SIGNALING_PATHWAY | 115 | 0.53 | 1.2 | 0.134 | 0.477 |
| 2008 | PRC2_EED_DN.V1_UP | 128 | 0.52 | 1.2 | 0.138 | 0.477 |
| 2009 | CORRE_MULTIPLE_MYELOMA_UP | 49 | 0.57 | 1.2 | 0.185 | 0.477 |
| 2010 | CCANNAGRKGGC_UNKNOWN | 93 | 0.54 | 1.2 | 0.147 | 0.477 |

| | | | | | | |
|---|---|---|---|---|---|---|
| 2011 | REACTOME_SEMA4D_IN_SEMAPHORIN_SIGNALIN( | 22 | 0.64 | 1.2 | 0.217 | 0.477 |
| 2012 | REACTOME_SYNDECAN_INTERACTIONS | 20 | 0.66 | 1.2 | 0.218 | 0.478 |
| 2013 | MIR6779_5P | 143 | 0.52 | 1.2 | 0.132 | 0.479 |
| 2014 | WP_SUDDEN_INFANT_DEATH_SYNDROME_SIDS_S | 113 | 0.52 | 1.2 | 0.135 | 0.479 |
| 2015 | LIM_MAMMARY_STEM_CELL_DN | 373 | 0.5 | 1.2 | 0.047 | 0.479 |
| 2016 | MIR6750_5P | 54 | 0.57 | 1.2 | 0.182 | 0.479 |
| 2017 | MIR8068 | 316 | 0.49 | 1.2 | 0.054 | 0.479 |
| 2018 | TAKEDA_TARGETS_OF_NUP98_HOXA9_FUSION_1 | 70 | 0.55 | 1.2 | 0.164 | 0.48 |
| 2019 | LIU_PROSTATE_CANCER_DN | 360 | 0.49 | 1.2 | 0.05 | 0.48 |
| 2020 | DESERT_STEM_CELL_HEPATOCELLULAR_CARCIN | 218 | 0.5 | 1.2 | 0.096 | 0.48 |
| 2021 | MIR4786_3P | 141 | 0.51 | 1.2 | 0.113 | 0.48 |
| 2022 | WGGAATGY_TEF1_Q6 | 284 | 0.5 | 1.2 | 0.064 | 0.481 |
| 2023 | VANTVEER_BREAST_CANCER_METASTASIS_UP | 34 | 0.6 | 1.2 | 0.2 | 0.481 |
| 2024 | LEE_AGING_NEOCORTEX_UP | 60 | 0.55 | 1.2 | 0.191 | 0.481 |
| 2025 | MIR7153_3P | 47 | 0.57 | 1.2 | 0.192 | 0.481 |
| 2026 | ZHENG_GLIOBLASTOMA_PLASTICITY_UP | 204 | 0.5 | 1.2 | 0.089 | 0.481 |
| 2027 | KUMAR_TARGETS_OF_MLL_AF9_FUSION | 301 | 0.5 | 1.2 | 0.07 | 0.481 |
| 2028 | PID_ERBB4_PATHWAY | 33 | 0.6 | 1.2 | 0.221 | 0.481 |
| 2029 | REACTOME_STABILIZATION_OF_P53 | 55 | 0.57 | 1.2 | 0.175 | 0.482 |
| 2030 | CDPCR1_01 | 103 | 0.53 | 1.2 | 0.152 | 0.481 |
| 2031 | GGCNNMSMYNTTG_UNKNOWN | 72 | 0.55 | 1.2 | 0.169 | 0.482 |
| 2032 | MIR140_3P | 125 | 0.52 | 1.2 | 0.13 | 0.482 |
| 2033 | MIR214_3P | 222 | 0.51 | 1.2 | 0.087 | 0.481 |
| 2034 | REACTOME_DISEASES_OF_MITOTIC_CELL_CYCLE | 38 | 0.6 | 1.2 | 0.186 | 0.481 |
| 2035 | BROWNE_HCMV_INFECTION_18HR_UP | 155 | 0.52 | 1.2 | 0.118 | 0.481 |
| 2036 | JAATINEN_HEMATOPOIETIC_STEM_CELL_DN | 115 | 0.52 | 1.2 | 0.155 | 0.482 |
| 2037 | RAO_BOUND_BY_SALL4_ISOFORM_B | 415 | 0.49 | 1.2 | 0.047 | 0.482 |
| 2038 | MIR375_3P | 32 | 0.6 | 1.2 | 0.211 | 0.482 |
| 2039 | OSADA_ASCL1_TARGETS_DN | 15 | 0.68 | 1.2 | 0.26 | 0.482 |
| 2040 | BIOCARTA_TGFB_PATHWAY | 17 | 0.68 | 1.2 | 0.23 | 0.482 |
| 2041 | GENTILE_UV_HIGH_DOSE_DN | 271 | 0.5 | 1.2 | 0.06 | 0.483 |
| 2042 | SABATES_COLORECTAL_ADENOMA_DN | 143 | 0.52 | 1.2 | 0.112 | 0.484 |
| 2043 | CHANDRAN_METASTASIS_DN | 246 | 0.5 | 1.2 | 0.087 | 0.484 |

| | | | | | | |
|---|---|---|---|---|---|---|
| 2044 | STAT5A_03 | 212 | 0.5 | 1.2 | 0.091 | 0.484 |
| 2045 | RICKMAN_TUMOR_DIFFERENTIATED_WELL_VS_M( | 80 | 0.54 | 1.2 | 0.158 | 0.485 |
| 2046 | PID_CXCR4_PATHWAY | 85 | 0.54 | 1.2 | 0.152 | 0.485 |
| 2047 | VALK_AML_CLUSTER_7 | 19 | 0.66 | 1.2 | 0.236 | 0.487 |
| 2048 | NFXL1_TARGET_GENES | 294 | 0.49 | 1.2 | 0.07 | 0.487 |
| 2049 | DACOSTA_UV_RESPONSE_VIA_ERCC3_TTD_DN | 77 | 0.55 | 1.2 | 0.17 | 0.487 |
| 2050 | SPIRA_SMOKERS_LUNG_CANCER_UP | 30 | 0.62 | 1.2 | 0.221 | 0.487 |
| 2051 | YANG_BREAST_CANCER_ESR1_DN | 22 | 0.65 | 1.2 | 0.23 | 0.488 |
| 2052 | MIR4662A_5P | 73 | 0.54 | 1.2 | 0.178 | 0.489 |
| 2053 | MIR6731_3P | 57 | 0.56 | 1.2 | 0.185 | 0.489 |
| 2054 | REACTOME_TRANSCRIPTIONAL_REGULATION_BY_ | 37 | 0.58 | 1.2 | 0.211 | 0.489 |
| 2055 | HNF6_Q6 | 186 | 0.51 | 1.2 | 0.104 | 0.489 |
| 2056 | PID_TCPTP_PATHWAY | 38 | 0.58 | 1.2 | 0.221 | 0.49 |
| 2057 | MIR6747_5P | 32 | 0.6 | 1.2 | 0.22 | 0.49 |
| 2058 | RATTENBACHER_BOUND_BY_CELF1 | 345 | 0.49 | 1.2 | 0.052 | 0.492 |
| 2059 | MARTINEZ_RB1_AND_TP53_TARGETS_DN | 500 | 0.49 | 1.2 | 0.03 | 0.492 |
| 2060 | REACTOME_G_BETA_GAMMA_SIGNALLING_THRO | 18 | 0.67 | 1.2 | 0.225 | 0.492 |
| 2061 | REACTOME_SIGNALING_BY_ERBB2_IN_CANCER | 24 | 0.62 | 1.2 | 0.235 | 0.493 |
| 2062 | LEE_CALORIE_RESTRICTION_NEOCORTEX_DN | 51 | 0.56 | 1.2 | 0.172 | 0.494 |
| 2063 | MANNE_COVID19_ICU_VS_HEALTHY_DONOR_PLA | 103 | 0.53 | 1.2 | 0.155 | 0.494 |
| 2064 | SIRNA_EIF4GI_UP | 84 | 0.54 | 1.2 | 0.174 | 0.495 |
| 2065 | WNT_UP.V1_UP | 121 | 0.52 | 1.2 | 0.142 | 0.495 |
| 2066 | ZHU_CMV_24_HR_DN | 62 | 0.55 | 1.2 | 0.181 | 0.495 |
| 2067 | MIYAGAWA_TARGETS_OF_EWSR1_ETS_FUSIONS_ | 216 | 0.5 | 1.2 | 0.11 | 0.495 |
| 2068 | MIR4797_3P | 82 | 0.54 | 1.2 | 0.161 | 0.495 |
| 2069 | GAVIN_FOXP3_TARGETS_CLUSTER_P3 | 126 | 0.52 | 1.2 | 0.138 | 0.495 |
| 2070 | REACTOME_TRANSCRIPTIONAL_REGULATION_OF_ | 75 | 0.54 | 1.2 | 0.177 | 0.496 |
| 2071 | PIAS4_TARGET_GENES | 59 | 0.55 | 1.2 | 0.18 | 0.496 |
| 2072 | MARKEY_RB1_ACUTE_LOF_UP | 219 | 0.5 | 1.19 | 0.085 | 0.497 |
| 2073 | BIOCARTA_EDG1_PATHWAY | 18 | 0.65 | 1.19 | 0.268 | 0.497 |
| 2074 | KEGG_MTOR_SIGNALING_PATHWAY | 47 | 0.57 | 1.19 | 0.199 | 0.497 |
| 2075 | GESERICK_TERT_TARGETS_DN | 19 | 0.66 | 1.19 | 0.225 | 0.498 |
| 2076 | PID_AVB3_OPN_PATHWAY | 29 | 0.61 | 1.19 | 0.217 | 0.499 |

| | | | | | | |
|---|---|---|---|---|---|---|
| 2077 | FRASOR_RESPONSE_TO_SERM_OR_FULVESTRAN | 49 | 0.57 | 1.19 | 0.189 | 0.499 |
| 2078 | PID_TXA2PATHWAY | 45 | 0.57 | 1.19 | 0.22 | 0.499 |
| 2079 | IL2_UP.V1_UP | 130 | 0.51 | 1.19 | 0.138 | 0.5 |
| 2080 | MARIADASON_REGULATED_BY_HISTONE_ACETYL | 56 | 0.56 | 1.19 | 0.182 | 0.501 |
| 2081 | LIM_MAMMARY_STEM_CELL_UP | 359 | 0.49 | 1.19 | 0.066 | 0.501 |
| 2082 | HADDAD_B_LYMPHOCYTE_PROGENITOR | 239 | 0.49 | 1.19 | 0.084 | 0.501 |
| 2083 | WIERENGA_STAT5A_TARGETS_DN | 139 | 0.51 | 1.19 | 0.125 | 0.505 |
| 2084 | KUUSELO_PANCREATIC_CANCER_19Q13_AMPLIFI | 26 | 0.62 | 1.19 | 0.216 | 0.504 |
| 2085 | METTL14_TARGET_GENES | 99 | 0.52 | 1.19 | 0.163 | 0.505 |
| 2086 | MEBARKI_HCC_PROGENITOR_WNT_UP | 123 | 0.51 | 1.19 | 0.135 | 0.506 |
| 2087 | BROWNE_HCMV_INFECTION_1HR_DN | 169 | 0.51 | 1.19 | 0.125 | 0.506 |
| 2088 | NFE2_01 | 225 | 0.5 | 1.19 | 0.094 | 0.507 |
| 2089 | BIOCARTA_INTEGRIN_PATHWAY | 33 | 0.61 | 1.19 | 0.218 | 0.508 |
| 2090 | BYSTRYKH_HEMATOPOIESIS_STEM_CELL_QTL_C | 113 | 0.52 | 1.19 | 0.162 | 0.509 |
| 2091 | MIR6729_3P | 58 | 0.56 | 1.19 | 0.192 | 0.509 |
| 2092 | LEE_LIVER_CANCER_ACOX1_DN | 31 | 0.6 | 1.19 | 0.223 | 0.509 |
| 2093 | ACEVEDO_LIVER_CANCER_WITH_H3K27ME3_UP | 156 | 0.51 | 1.19 | 0.13 | 0.508 |
| 2094 | PASQUALUCCI_LYMPHOMA_BY_GC_STAGE_DN | 150 | 0.51 | 1.19 | 0.125 | 0.508 |
| 2095 | BOYLAN_MULTIPLE_MYELOMA_C_DN | 49 | 0.57 | 1.19 | 0.196 | 0.508 |
| 2096 | BMI1_DN.V1_DN | 105 | 0.52 | 1.19 | 0.146 | 0.508 |
| 2097 | LEE_LIVER_CANCER_DENA_UP | 37 | 0.59 | 1.19 | 0.226 | 0.509 |
| 2098 | KARLSSON_TGFB1_TARGETS_UP | 115 | 0.52 | 1.19 | 0.162 | 0.509 |
| 2099 | KRAS.PROSTATE_UP.V1_UP | 80 | 0.53 | 1.19 | 0.175 | 0.508 |
| 2100 | TIEN_INTESTINE_PROBIOTICS_6HR_UP | 52 | 0.57 | 1.19 | 0.174 | 0.508 |
| 2101 | REACTOME_COLLAGEN_DEGRADATION | 34 | 0.59 | 1.19 | 0.24 | 0.511 |
| 2102 | TGCTGAY_UNKNOWN | 417 | 0.49 | 1.19 | 0.039 | 0.511 |
| 2103 | QI_HYPOXIA_TARGETS_OF_HIF1A_AND_FOXA2 | 31 | 0.6 | 1.19 | 0.247 | 0.511 |
| 2104 | JEPSEN_SMRT_TARGETS | 25 | 0.63 | 1.19 | 0.239 | 0.511 |
| 2105 | TCTAGAG_MIR517 | 43 | 0.58 | 1.19 | 0.199 | 0.511 |
| 2106 | AP1_01 | 206 | 0.5 | 1.19 | 0.099 | 0.511 |
| 2107 | DORMOY_ELAVL1_TARGETS | 15 | 0.68 | 1.19 | 0.27 | 0.511 |
| 2108 | REACTOME_AUF1_HNRNP_D0_BINDS_AND_DESTA | 53 | 0.56 | 1.19 | 0.199 | 0.511 |
| 2109 | GATA1_04 | 167 | 0.5 | 1.19 | 0.137 | 0.51 |

| | | | | | | |
|---|---|---|---|---|---|---|
| 2110 | MIF1_01 | 161 | 0.51 | 1.19 | 0.133 | 0.51 |
| 2111 | ROYLANCE_BREAST_CANCER_16Q_COPY_NUMBE | 19 | 0.64 | 1.19 | 0.253 | 0.51 |
| 2112 | REACTOME_TCF_DEPENDENT_SIGNALING_IN_RE: | 178 | 0.5 | 1.19 | 0.13 | 0.51 |
| 2113 | MIR4683 | 39 | 0.58 | 1.19 | 0.211 | 0.51 |
| 2114 | MIR6769A_5P | 40 | 0.59 | 1.19 | 0.206 | 0.511 |
| 2115 | REACTOME_SIGNALING_BY_VEGF | 99 | 0.53 | 1.19 | 0.163 | 0.511 |
| 2116 | MIR558 | 96 | 0.52 | 1.19 | 0.186 | 0.511 |
| 2117 | MIR603 | 147 | 0.51 | 1.19 | 0.147 | 0.511 |
| 2118 | CREB_Q2_01 | 198 | 0.49 | 1.19 | 0.126 | 0.511 |
| 2119 | MIR4450 | 64 | 0.54 | 1.19 | 0.186 | 0.511 |
| 2120 | MIR526B_5P | 87 | 0.53 | 1.19 | 0.175 | 0.511 |
| 2121 | REACTOME_NEGATIVE_REGULATION_OF_NOTCH4 | 54 | 0.56 | 1.19 | 0.202 | 0.513 |
| 2122 | REACTOME_REGULATION_OF_INSULIN_LIKE_GRC | 72 | 0.54 | 1.19 | 0.195 | 0.512 |
| 2123 | MIR3972 | 56 | 0.55 | 1.19 | 0.218 | 0.513 |
| 2124 | MIR5693 | 69 | 0.54 | 1.19 | 0.173 | 0.513 |
| 2125 | DEURIG_T_CELL_PROLYMPHOCYTIC_LEUKEMIA_U | 303 | 0.49 | 1.19 | 0.08 | 0.514 |
| 2126 | WP_HIPPOMERLIN_SIGNALING_DYSREGULATION | 93 | 0.52 | 1.19 | 0.17 | 0.514 |
| 2127 | WP_MAPK_SIGNALING_PATHWAY | 203 | 0.5 | 1.19 | 0.112 | 0.514 |
| 2128 | RSRFC4_01 | 183 | 0.51 | 1.19 | 0.115 | 0.514 |
| 2129 | MIR548AO_3P | 17 | 0.67 | 1.19 | 0.221 | 0.515 |
| 2130 | REACTOME_SIGNALING_BY_EGFR | 45 | 0.56 | 1.19 | 0.217 | 0.515 |
| 2131 | MIR504_3P | 124 | 0.51 | 1.19 | 0.153 | 0.517 |
| 2132 | MIDORIKAWA_AMPLIFIED_IN_LIVER_CANCER | 42 | 0.58 | 1.19 | 0.224 | 0.516 |
| 2133 | AP1_Q2_01 | 215 | 0.5 | 1.19 | 0.121 | 0.516 |
| 2134 | HAMAI_APOPTOSIS_VIA_TRAIL_DN | 162 | 0.51 | 1.19 | 0.128 | 0.517 |
| 2135 | AR_03 | 49 | 0.57 | 1.19 | 0.205 | 0.517 |
| 2136 | RAPA_EARLY_UP.V1_UP | 118 | 0.52 | 1.18 | 0.16 | 0.518 |
| 2137 | CUI_TCF21_TARGETS_2_UP | 366 | 0.48 | 1.18 | 0.08 | 0.518 |
| 2138 | CAAGGAT_MIR362 | 59 | 0.55 | 1.18 | 0.208 | 0.519 |
| 2139 | WANG_METHYLATED_IN_BREAST_CANCER | 30 | 0.61 | 1.18 | 0.244 | 0.519 |
| 2140 | WP_COPPER_HOMEOSTASIS | 43 | 0.57 | 1.18 | 0.205 | 0.52 |
| 2141 | BRUECKNER_TARGETS_OF_MIRLET7A3_UP | 92 | 0.53 | 1.18 | 0.175 | 0.52 |
| 2142 | REACTOME_CELL_CELL_COMMUNICATION | 103 | 0.52 | 1.18 | 0.159 | 0.52 |

| | | | | | | |
|---|---|---|---|---|---|---|
| 2143 | PURBEY_TARGETS_OF_CTBP1_NOT_SATB1_UP | 237 | 0.49 | 1.18 | 0.108 | 0.52 |
| 2144 | MARTINEZ_RB1_AND_TP53_TARGETS_UP | 468 | 0.49 | 1.18 | 0.051 | 0.52 |
| 2145 | MIR2467_5P | 96 | 0.52 | 1.18 | 0.185 | 0.52 |
| 2146 | MIR3156_5P | 60 | 0.55 | 1.18 | 0.197 | 0.521 |
| 2147 | REACTOME_REGULATION_OF_PTEN_STABILITY_A | 66 | 0.53 | 1.18 | 0.204 | 0.521 |
| 2148 | WP_INTEGRATED_CANCER_PATHWAY | 43 | 0.58 | 1.18 | 0.218 | 0.521 |
| 2149 | MIR4653_5P | 117 | 0.51 | 1.18 | 0.165 | 0.521 |
| 2150 | AKT_UP_MTOR_DN.V1_UP | 142 | 0.51 | 1.18 | 0.154 | 0.522 |
| 2151 | MIR4487 | 60 | 0.54 | 1.18 | 0.195 | 0.523 |
| 2152 | JAZAERI_BREAST_CANCER_BRCA1_VS_BRCA2_D | 34 | 0.59 | 1.18 | 0.227 | 0.522 |
| 2153 | XU_HGF_SIGNALING_NOT_VIA_AKT1_48HR_UP | 30 | 0.6 | 1.18 | 0.234 | 0.522 |
| 2154 | MIR20B_3P | 52 | 0.55 | 1.18 | 0.198 | 0.522 |
| 2155 | MIR125A_5P | 278 | 0.49 | 1.18 | 0.11 | 0.522 |
| 2156 | SIG_CHEMOTAXIS | 41 | 0.58 | 1.18 | 0.229 | 0.522 |
| 2157 | GGTAACC_MIR4095P | 27 | 0.61 | 1.18 | 0.256 | 0.522 |
| 2158 | STEIN_ESR1_TARGETS | 72 | 0.54 | 1.18 | 0.185 | 0.522 |
| 2159 | WANG_RESPONSE_TO_BEXAROTENE_UP | 26 | 0.61 | 1.18 | 0.225 | 0.522 |
| 2160 | MIR4648 | 31 | 0.59 | 1.18 | 0.242 | 0.523 |
| 2161 | BIOCARTA_ARAP_PATHWAY | 17 | 0.66 | 1.18 | 0.265 | 0.524 |
| 2162 | PID_KIT_PATHWAY | 44 | 0.57 | 1.18 | 0.222 | 0.524 |
| 2163 | MIR502_5P | 61 | 0.54 | 1.18 | 0.193 | 0.525 |
| 2164 | ACCAATC_MIR509 | 43 | 0.57 | 1.18 | 0.238 | 0.525 |
| 2165 | SUNG_METASTASIS_STROMA_UP | 91 | 0.53 | 1.18 | 0.18 | 0.525 |
| 2166 | BROWNE_HCMV_INFECTION_48HR_DN | 366 | 0.48 | 1.18 | 0.071 | 0.527 |
| 2167 | MIR7114_5P | 124 | 0.51 | 1.18 | 0.154 | 0.527 |
| 2168 | CEBPB_02 | 205 | 0.49 | 1.18 | 0.131 | 0.527 |
| 2169 | KEGG_DILATED_CARDIOMYOPATHY | 57 | 0.55 | 1.18 | 0.216 | 0.527 |
| 2170 | SOTIRIOU_BREAST_CANCER_GRADE_1_VS_3_DN | 44 | 0.57 | 1.18 | 0.231 | 0.527 |
| 2171 | PID_ECADHERIN_KERATINOCYTE_PATHWAY | 21 | 0.63 | 1.18 | 0.237 | 0.527 |
| 2172 | MIR3912_5P | 75 | 0.54 | 1.18 | 0.192 | 0.527 |
| 2173 | HSF2_TARGET_GENES | 200 | 0.5 | 1.18 | 0.117 | 0.527 |
| 2174 | REACTOME_G2_M_CHECKPOINTS | 147 | 0.5 | 1.18 | 0.136 | 0.528 |
| 2175 | MIR5572 | 72 | 0.54 | 1.18 | 0.201 | 0.528 |

| | | | | | | |
|---|---|---|---|---|---|---|
| 2176 | ENGELMANN_CANCER_PROGENITORS_DN | 50 | 0.55 | 1.18 | 0.208 | 0.528 |
| 2177 | MIR449C_3P | 47 | 0.56 | 1.18 | 0.232 | 0.528 |
| 2178 | STREICHER_LSM1_TARGETS_UP | 39 | 0.58 | 1.18 | 0.233 | 0.528 |
| 2179 | MIR874_5P | 53 | 0.56 | 1.18 | 0.217 | 0.528 |
| 2180 | ZHENG_BOUND_BY_FOXP3 | 380 | 0.48 | 1.18 | 0.072 | 0.529 |
| 2181 | LI_PROSTATE_CANCER_EPIGENETIC | 24 | 0.63 | 1.18 | 0.261 | 0.529 |
| 2182 | WOOD_EBV_EBNA1_TARGETS_DN | 34 | 0.59 | 1.18 | 0.225 | 0.53 |
| 2183 | MIR522_3P | 161 | 0.5 | 1.18 | 0.146 | 0.534 |
| 2184 | CHIARETTI_ACUTE_LYMPHOBLASTIC_LEUKEMIA_ | 57 | 0.55 | 1.18 | 0.218 | 0.534 |
| 2185 | TTTNNANAGCYR_UNKNOWN | 85 | 0.53 | 1.18 | 0.18 | 0.533 |
| 2186 | COLIN_PILOCYTIC_ASTROCYTOMA_VS_GLIOBLAS | 21 | 0.63 | 1.18 | 0.254 | 0.533 |
| 2187 | WP_OXIDATIVE_DAMAGE_RESPONSE | 35 | 0.58 | 1.18 | 0.232 | 0.533 |
| 2188 | HOELZEL_NF1_TARGETS_UP | 112 | 0.52 | 1.18 | 0.158 | 0.533 |
| 2189 | VANTVEER_BREAST_CANCER_POOR_PROGNOSIS | 40 | 0.58 | 1.18 | 0.236 | 0.533 |
| 2190 | FOROUTAN_INTEGRATED_TGFB_EMT_UP | 87 | 0.52 | 1.18 | 0.184 | 0.533 |
| 2191 | SCIBETTA_KDM5B_TARGETS_UP | 18 | 0.65 | 1.18 | 0.259 | 0.533 |
| 2192 | MIR3200_3P | 25 | 0.61 | 1.18 | 0.24 | 0.533 |
| 2193 | DELASERNA_MYOD_TARGETS_DN | 47 | 0.56 | 1.18 | 0.228 | 0.533 |
| 2194 | CSR_EARLY_UP.V1_DN | 113 | 0.51 | 1.18 | 0.162 | 0.532 |
| 2195 | MIR374A_5P | 355 | 0.48 | 1.18 | 0.084 | 0.533 |
| 2196 | REACTOME_WNT_LIGAND_BIOGENESIS_AND_TRA | 18 | 0.65 | 1.18 | 0.279 | 0.533 |
| 2197 | MIR6797_5P | 171 | 0.5 | 1.18 | 0.155 | 0.533 |
| 2198 | YAO_HOXA10_TARGETS_VIA_PROGESTERONE_U | 57 | 0.55 | 1.18 | 0.212 | 0.533 |
| 2199 | WP_STATIN_INHIBITION_OF_CHOLESTEROL_PROI | 19 | 0.64 | 1.18 | 0.273 | 0.534 |
| 2200 | MIR4678 | 97 | 0.52 | 1.18 | 0.197 | 0.534 |
| 2201 | LY_AGING_OLD_DN | 50 | 0.56 | 1.18 | 0.207 | 0.534 |
| 2202 | KANG_FLUOROURACIL_RESISTANCE_UP | 21 | 0.63 | 1.18 | 0.252 | 0.534 |
| 2203 | REACTOME_METABOLISM_OF_FAT_SOLUBLE_VIT | 28 | 0.61 | 1.18 | 0.248 | 0.535 |
| 2204 | SAKAI_CHRONIC_HEPATITIS_VS_LIVER_CANCER_ | 23 | 0.63 | 1.18 | 0.265 | 0.535 |
| 2205 | WP_NEURAL_CREST_CELL_MIGRATION_DURING_I | 35 | 0.58 | 1.18 | 0.248 | 0.534 |
| 2206 | GARGALOVIC_RESPONSE_TO_OXIDIZED_PHOSPH | 15 | 0.68 | 1.18 | 0.295 | 0.534 |
| 2207 | EGR2_01 | 167 | 0.5 | 1.18 | 0.15 | 0.534 |
| 2208 | VALK_AML_CLUSTER_3 | 29 | 0.6 | 1.18 | 0.255 | 0.534 |

| | | | | | | |
|---|---|---|---|---|---|---|
| 2209 | REACTOME_DEFECTIVE_CFTR_CAUSES_CYSTIC_I | 57 | 0.54 | 1.18 | 0.21 | 0.534 |
| 2210 | WP_PDGFRBETA_PATHWAY | 29 | 0.6 | 1.18 | 0.252 | 0.534 |
| 2211 | KEGG_REGULATION_OF_ACTIN_CYTOSKELETON | 167 | 0.5 | 1.18 | 0.156 | 0.534 |
| 2212 | PARENT_MTOR_SIGNALING_DN | 32 | 0.6 | 1.18 | 0.242 | 0.535 |
| 2213 | CEBP_Q2_01 | 214 | 0.49 | 1.18 | 0.121 | 0.535 |
| 2214 | MIR125B_5P | 276 | 0.49 | 1.18 | 0.116 | 0.535 |
| 2215 | KEGG_ADIPOCYTOKINE_SIGNALING_PATHWAY | 53 | 0.56 | 1.17 | 0.2 | 0.535 |
| 2216 | ERBB2_UP.V1_UP | 154 | 0.5 | 1.17 | 0.154 | 0.535 |
| 2217 | MIR4735_3P | 100 | 0.52 | 1.17 | 0.176 | 0.535 |
| 2218 | BOYAULT_LIVER_CANCER_SUBCLASS_G6_DN | 18 | 0.64 | 1.17 | 0.269 | 0.535 |
| 2219 | BCAT_BILD_ET_AL_DN | 40 | 0.57 | 1.17 | 0.215 | 0.535 |
| 2220 | GR_01 | 154 | 0.5 | 1.17 | 0.156 | 0.535 |
| 2221 | MCDOWELL_ACUTE_LUNG_INJURY_DN | 35 | 0.58 | 1.17 | 0.242 | 0.536 |
| 2222 | ROY_WOUND_BLOOD_VESSEL_DN | 15 | 0.66 | 1.17 | 0.295 | 0.536 |
| 2223 | REACTOME_RHO_GTPASES_ACTIVATE_PAKS | 19 | 0.64 | 1.17 | 0.265 | 0.536 |
| 2224 | MIR3921 | 115 | 0.51 | 1.17 | 0.179 | 0.536 |
| 2225 | MCCLUNG_COCAINE_REWARD_5D | 65 | 0.54 | 1.17 | 0.2 | 0.537 |
| 2226 | BRUINS_UVC_RESPONSE_MIDDLE | 91 | 0.53 | 1.17 | 0.176 | 0.537 |
| 2227 | ELVIDGE_HIF1A_AND_HIF2A_TARGETS_DN | 92 | 0.52 | 1.17 | 0.205 | 0.537 |
| 2228 | REACTOME_SIGNALING_BY_EGFR_IN_CANCER | 22 | 0.62 | 1.17 | 0.257 | 0.537 |
| 2229 | BASAKI_YBX1_TARGETS_DN | 338 | 0.48 | 1.17 | 0.075 | 0.537 |
| 2230 | MIR6732_3P | 83 | 0.52 | 1.17 | 0.19 | 0.537 |
| 2231 | BRUINS_UVC_RESPONSE_VIA_TP53_GROUP_D | 207 | 0.49 | 1.17 | 0.134 | 0.539 |
| 2232 | MEF2_03 | 171 | 0.5 | 1.17 | 0.148 | 0.539 |
| 2233 | MIR6514_5P | 35 | 0.59 | 1.17 | 0.256 | 0.541 |
| 2234 | MIR5698 | 166 | 0.5 | 1.17 | 0.161 | 0.542 |
| 2235 | ELVIDGE_HYPOXIA_UP | 153 | 0.5 | 1.17 | 0.163 | 0.542 |
| 2236 | LIAN_LIPA_TARGETS_3M | 24 | 0.62 | 1.17 | 0.264 | 0.542 |
| 2237 | REACTOME_NUCLEAR_EVENTS_KINASE_AND_TR/ | 51 | 0.55 | 1.17 | 0.232 | 0.542 |
| 2238 | WOOD_EBV_EBNA1_TARGETS_UP | 102 | 0.52 | 1.17 | 0.194 | 0.543 |
| 2239 | HELLER_HDAC_TARGETS_UP | 286 | 0.49 | 1.17 | 0.101 | 0.543 |
| 2240 | MUELLER_PLURINET | 287 | 0.49 | 1.17 | 0.111 | 0.544 |
| 2241 | WP_NOTCH_SIGNALING | 42 | 0.57 | 1.17 | 0.214 | 0.544 |

| | | | | | | |
|---|---|---|---|---|---|---|
| 2242 | WP_OXIDATIVE_STRESS_RESPONSE | 28 | 0.61 | 1.17 | 0.252 | 0.544 |
| 2243 | BIDUS_METASTASIS_DN | 125 | 0.51 | 1.17 | 0.164 | 0.544 |
| 2244 | NAKAMURA_METASTASIS_MODEL_UP | 34 | 0.59 | 1.17 | 0.236 | 0.544 |
| 2245 | NABA_ECM_GLYCOPROTEINS | 114 | 0.51 | 1.17 | 0.18 | 0.543 |
| 2246 | NGUYEN_NOTCH1_TARGETS_UP | 20 | 0.64 | 1.17 | 0.287 | 0.544 |
| 2247 | XU_GH1_AUTOCRINE_TARGETS_UP | 140 | 0.5 | 1.17 | 0.164 | 0.544 |
| 2248 | SHIN_B_CELL_LYMPHOMA_CLUSTER_7 | 26 | 0.6 | 1.17 | 0.258 | 0.546 |
| 2249 | RGAGGAARY_PU1_Q6 | 392 | 0.48 | 1.17 | 0.078 | 0.546 |
| 2250 | MTOR_UP.V1_DN | 143 | 0.5 | 1.17 | 0.158 | 0.546 |
| 2251 | WANG_BARRETTS_ESOPHAGUS_AND_ESOPHAGU | 24 | 0.61 | 1.17 | 0.269 | 0.546 |
| 2252 | CHNG_MULTIPLE_MYELOMA_HYPERPLOID_DN | 29 | 0.58 | 1.17 | 0.263 | 0.546 |
| 2253 | MIR4317 | 78 | 0.52 | 1.17 | 0.189 | 0.546 |
| 2254 | PTEN_DN.V2_DN | 111 | 0.5 | 1.17 | 0.172 | 0.545 |
| 2255 | RYAAAKNNNNNNTTGW_UNKNOWN | 61 | 0.54 | 1.17 | 0.224 | 0.545 |
| 2256 | CAFFAREL_RESPONSE_TO_THC_24HR_5_UP | 32 | 0.59 | 1.17 | 0.253 | 0.546 |
| 2257 | MIR342_5P | 49 | 0.56 | 1.17 | 0.22 | 0.546 |
| 2258 | KYNG_DNA_DAMAGE_UP | 160 | 0.5 | 1.17 | 0.149 | 0.546 |
| 2259 | MIR3156_3P | 78 | 0.53 | 1.17 | 0.2 | 0.547 |
| 2260 | VALK_AML_CLUSTER_2 | 23 | 0.62 | 1.17 | 0.265 | 0.547 |
| 2261 | AKT_UP.V1_DN | 136 | 0.5 | 1.17 | 0.176 | 0.548 |
| 2262 | NCX_01 | 136 | 0.51 | 1.17 | 0.174 | 0.548 |
| 2263 | REACTOME_ASYMMETRIC_LOCALIZATION_OF_PC | 61 | 0.54 | 1.17 | 0.216 | 0.548 |
| 2264 | HOSHIDA_LIVER_CANCER_LATE_RECURRENCE_U | 47 | 0.56 | 1.17 | 0.227 | 0.549 |
| 2265 | REACTOME_ANTIGEN_PROCESSING_UBIQUITINAT | 279 | 0.48 | 1.17 | 0.102 | 0.549 |
| 2266 | ZHAN_MULTIPLE_MYELOMA_CD1_UP | 37 | 0.58 | 1.17 | 0.256 | 0.55 |
| 2267 | HOXB7_TARGET_GENES | 24 | 0.61 | 1.17 | 0.257 | 0.55 |
| 2268 | MIR6762_3P | 106 | 0.51 | 1.17 | 0.18 | 0.55 |
| 2269 | MIR224_3P | 162 | 0.5 | 1.17 | 0.137 | 0.551 |
| 2270 | WP_PHOTODYNAMIC_THERAPYINDUCED_UNFOLD | 26 | 0.61 | 1.17 | 0.26 | 0.551 |
| 2271 | REACTOME_CD28_DEPENDENT_PI3K_AKT_SIGNA | 20 | 0.63 | 1.17 | 0.284 | 0.552 |
| 2272 | GAZDA_DIAMOND_BLACKFAN_ANEMIA_PROGENIT | 36 | 0.57 | 1.17 | 0.234 | 0.553 |
| 2273 | REACTOME_SIGNALING_BY_NOTCH2 | 28 | 0.6 | 1.17 | 0.264 | 0.553 |
| 2274 | MIR6812_5P | 53 | 0.55 | 1.17 | 0.229 | 0.553 |

| | | | | | | |
|---|---|---|---|---|---|---|
| 2275 | VALK_AML_CLUSTER_11 | 30 | 0.59 | 1.17 | 0.255 | 0.553 |
| 2276 | BURTON_ADIPOGENESIS_11 | 46 | 0.56 | 1.17 | 0.234 | 0.555 |
| 2277 | REACTOME_SENSORY_PROCESSING_OF_SOUND_ | 42 | 0.56 | 1.17 | 0.231 | 0.555 |
| 2278 | WP_NEURAL_CREST_CELL_MIGRATION_IN_CANCI | 37 | 0.58 | 1.17 | 0.252 | 0.555 |
| 2279 | MIR7111_5P | 167 | 0.5 | 1.17 | 0.156 | 0.554 |
| 2280 | MIR379_5P | 39 | 0.57 | 1.17 | 0.244 | 0.555 |
| 2281 | WP_NOTCH_SIGNALING_PATHWAY | 58 | 0.54 | 1.17 | 0.204 | 0.555 |
| 2282 | GRAESSMANN_RESPONSE_TO_MC_AND_SERUM_ | 78 | 0.53 | 1.16 | 0.21 | 0.556 |
| 2283 | PRC2_EZH2_UP.V1_UP | 135 | 0.5 | 1.16 | 0.164 | 0.556 |
| 2284 | RNCTGNYNRNCTGNY_UNKNOWN | 61 | 0.54 | 1.16 | 0.228 | 0.556 |
| 2285 | REACTOME_SIGNALING_BY_THE_B_CELL_RECEP | 104 | 0.51 | 1.16 | 0.181 | 0.557 |
| 2286 | BOYLAN_MULTIPLE_MYELOMA_C_CLUSTER_DN | 22 | 0.62 | 1.16 | 0.269 | 0.557 |
| 2287 | MIR3920 | 122 | 0.5 | 1.16 | 0.171 | 0.557 |
| 2288 | MIR1_5P | 74 | 0.54 | 1.16 | 0.201 | 0.558 |
| 2289 | FOXJ2_01 | 142 | 0.5 | 1.16 | 0.169 | 0.558 |
| 2290 | MIR4264 | 86 | 0.52 | 1.16 | 0.216 | 0.558 |
| 2291 | MYLLYKANGAS_AMPLIFICATION_HOT_SPOT_17 | 15 | 0.66 | 1.16 | 0.276 | 0.558 |
| 2292 | DCA_UP.V1_UP | 115 | 0.51 | 1.16 | 0.191 | 0.558 |
| 2293 | SCHLINGEMANN_SKIN_CARCINOGENESIS_TPA_DI | 20 | 0.63 | 1.16 | 0.277 | 0.558 |
| 2294 | REACTOME_DOWNSTREAM_SIGNALING_OF_ACTIV | 17 | 0.65 | 1.16 | 0.285 | 0.558 |
| 2295 | MIR569 | 67 | 0.53 | 1.16 | 0.228 | 0.559 |
| 2296 | MIR6505_5P | 169 | 0.49 | 1.16 | 0.16 | 0.56 |
| 2297 | MIR30B_3P | 138 | 0.5 | 1.16 | 0.176 | 0.56 |
| 2298 | HOFMANN_MYELODYSPLASTIC_SYNDROM_RISK_I | 18 | 0.64 | 1.16 | 0.265 | 0.56 |
| 2299 | NAKAMURA_ADIPOGENESIS_LATE_DN | 24 | 0.62 | 1.16 | 0.274 | 0.563 |
| 2300 | SATO_SILENCED_BY_METHYLATION_IN_PANCREA | 36 | 0.57 | 1.16 | 0.239 | 0.563 |
| 2301 | GARGALOVIC_RESPONSE_TO_OXIDIZED_PHOSPI | 21 | 0.63 | 1.16 | 0.279 | 0.564 |
| 2302 | MIR4752 | 69 | 0.53 | 1.16 | 0.219 | 0.564 |
| 2303 | REACTOME_RUNX1_INTERACTS_WITH_CO_FACTO | 37 | 0.57 | 1.16 | 0.266 | 0.565 |
| 2304 | REACTOME_CYTOPROTECTION_BY_HMOX1 | 116 | 0.51 | 1.16 | 0.187 | 0.564 |
| 2305 | GCNP_SHH_UP_EARLY.V1_DN | 120 | 0.5 | 1.16 | 0.185 | 0.565 |
| 2306 | MIR580_3P | 184 | 0.49 | 1.16 | 0.167 | 0.566 |
| 2307 | GATA3_01 | 172 | 0.49 | 1.16 | 0.18 | 0.566 |

| | | | | | | |
|---|---|---|---|---|---|---|
| 2308 | MIR12115 | 56 | 0.54 | 1.16 | 0.212 | 0.567 |
| 2309 | TGANNYRGCA_TCF11MAFG_01 | 248 | 0.48 | 1.16 | 0.134 | 0.567 |
| 2310 | WP_AMPACTIVATED_PROTEIN_KINASE_AMPK_SIG | 58 | 0.54 | 1.16 | 0.213 | 0.568 |
| 2311 | EGFR_UP.V1_DN | 167 | 0.49 | 1.16 | 0.178 | 0.568 |
| 2312 | TEF1_Q6 | 176 | 0.49 | 1.16 | 0.175 | 0.569 |
| 2313 | PARK_TRETINOIN_RESPONSE_AND_PML_RARA_F | 23 | 0.61 | 1.16 | 0.265 | 0.57 |
| 2314 | MORI_PRE_BI_LYMPHOCYTE_DN | 63 | 0.54 | 1.16 | 0.227 | 0.57 |
| 2315 | MIR6776_5P | 73 | 0.53 | 1.16 | 0.228 | 0.569 |
| 2316 | FAELT_B_CLL_WITH_VH_REARRANGEMENTS_UP | 43 | 0.56 | 1.16 | 0.248 | 0.569 |
| 2317 | MIR106A_3P | 84 | 0.52 | 1.16 | 0.217 | 0.569 |
| 2318 | MIR6870_5P | 177 | 0.49 | 1.16 | 0.168 | 0.569 |
| 2319 | MIR9986 | 141 | 0.5 | 1.16 | 0.17 | 0.569 |
| 2320 | JINESH_BLEBBISHIELD_VS_LIVE_CONTROL_UP | 356 | 0.48 | 1.16 | 0.102 | 0.57 |
| 2321 | REACTOME_PHASE_I_FUNCTIONALIZATION_OF_C | 51 | 0.54 | 1.16 | 0.245 | 0.571 |
| 2322 | REACTOME_SIGNALING_BY_PDGF | 44 | 0.55 | 1.16 | 0.245 | 0.571 |
| 2323 | MIR331_3P | 57 | 0.55 | 1.16 | 0.231 | 0.571 |
| 2324 | MIR4761_3P | 41 | 0.57 | 1.16 | 0.265 | 0.571 |
| 2325 | WP_EPO_RECEPTOR_SIGNALING | 23 | 0.61 | 1.16 | 0.287 | 0.571 |
| 2326 | COWLING_MYCN_TARGETS | 26 | 0.6 | 1.16 | 0.27 | 0.571 |
| 2327 | XU_RESPONSE_TO_TRETINOIN_UP | 15 | 0.66 | 1.16 | 0.301 | 0.574 |
| 2328 | REACTOME_O_LINKED_GLYCOSYLATION | 77 | 0.53 | 1.16 | 0.245 | 0.574 |
| 2329 | MIR1294 | 142 | 0.49 | 1.16 | 0.171 | 0.573 |
| 2330 | TGASTMAGC_NFE2_01 | 155 | 0.49 | 1.16 | 0.173 | 0.575 |
| 2331 | COLLIS_PRKDC_SUBSTRATES | 17 | 0.63 | 1.16 | 0.298 | 0.575 |
| 2332 | REACTOME_PLASMA_LIPOPROTEIN_CLEARANCE | 26 | 0.6 | 1.16 | 0.284 | 0.575 |
| 2333 | REACTOME_NCAM1_INTERACTIONS | 29 | 0.59 | 1.16 | 0.258 | 0.576 |
| 2334 | REACTOME_DISEASES_OF_GLYCOSYLATION | 113 | 0.51 | 1.16 | 0.205 | 0.576 |
| 2335 | ZIC1_01 | 226 | 0.49 | 1.16 | 0.149 | 0.576 |
| 2336 | MIR6802_3P | 36 | 0.58 | 1.16 | 0.24 | 0.576 |
| 2337 | NCOA2_TARGET_GENES | 391 | 0.47 | 1.16 | 0.093 | 0.577 |
| 2338 | IL21_UP.V1_UP | 124 | 0.5 | 1.16 | 0.189 | 0.578 |
| 2339 | BIOCARTA_FMLP_PATHWAY | 31 | 0.58 | 1.16 | 0.286 | 0.579 |
| 2340 | REACTOME_DEGRADATION_OF_AXIN | 53 | 0.55 | 1.16 | 0.258 | 0.579 |

| | | | | | | |
|---|---|---|---|---|---|---|
| 2341 | MARTINEZ_RB1_TARGETS_DN | 434 | 0.47 | 1.15 | 0.08 | 0.58 |
| 2342 | SMID_BREAST_CANCER_LUMINAL_A_UP | 40 | 0.57 | 1.15 | 0.26 | 0.58 |
| 2343 | PID_RETINOIC_ACID_PATHWAY | 28 | 0.59 | 1.15 | 0.263 | 0.582 |
| 2344 | MIR4528 | 176 | 0.49 | 1.15 | 0.188 | 0.582 |
| 2345 | AREB6_04 | 200 | 0.48 | 1.15 | 0.175 | 0.583 |
| 2346 | BIOCARTA_IL2_PATHWAY | 20 | 0.63 | 1.15 | 0.294 | 0.583 |
| 2347 | MIR6796_3P | 56 | 0.54 | 1.15 | 0.256 | 0.583 |
| 2348 | ALFANO_MYC_TARGETS | 208 | 0.48 | 1.15 | 0.156 | 0.583 |
| 2349 | AP1_Q4_01 | 210 | 0.48 | 1.15 | 0.167 | 0.583 |
| 2350 | GAUSSMANN_MLL_AF4_FUSION_TARGETS_G_UP | 176 | 0.49 | 1.15 | 0.178 | 0.583 |
| 2351 | RAAGNYNNCTTY_UNKNOWN | 114 | 0.5 | 1.15 | 0.201 | 0.583 |
| 2352 | CYCLIN_D1_UP.V1_UP | 136 | 0.5 | 1.15 | 0.189 | 0.583 |
| 2353 | MIR3659 | 62 | 0.53 | 1.15 | 0.251 | 0.585 |
| 2354 | MIR192_3P | 84 | 0.51 | 1.15 | 0.221 | 0.586 |
| 2355 | SPIELMAN_LYMPHOBLAST_EUROPEAN_VS_ASIAN | 466 | 0.47 | 1.15 | 0.095 | 0.587 |
| 2356 | FOXD3_TARGET_GENES | 30 | 0.58 | 1.15 | 0.255 | 0.588 |
| 2357 | NFAT_Q6 | 193 | 0.49 | 1.15 | 0.155 | 0.588 |
| 2358 | HOFMANN_CELL_LYMPHOMA_UP | 40 | 0.56 | 1.15 | 0.252 | 0.588 |
| 2359 | KYNG_WERNER_SYNDROM_AND_NORMAL_AGING | 68 | 0.53 | 1.15 | 0.23 | 0.588 |
| 2360 | REACTOME_UCH_PROTEINASES | 90 | 0.51 | 1.15 | 0.226 | 0.588 |
| 2361 | MIR1185_5P | 48 | 0.55 | 1.15 | 0.259 | 0.588 |
| 2362 | BROWN_MYELOID_CELL_DEVELOPMENT_DN | 108 | 0.51 | 1.15 | 0.218 | 0.588 |
| 2363 | NIKOLSKY_BREAST_CANCER_21Q22_AMPLICON | 15 | 0.65 | 1.15 | 0.3 | 0.589 |
| 2364 | LANDIS_BREAST_CANCER_PROGRESSION_DN | 49 | 0.54 | 1.15 | 0.241 | 0.589 |
| 2365 | ARGGGTTAA_UNKNOWN | 92 | 0.51 | 1.15 | 0.225 | 0.59 |
| 2366 | MIR18A_5P_MIR18B_5P | 110 | 0.51 | 1.15 | 0.201 | 0.59 |
| 2367 | BERENJENO_ROCK_SIGNALING_NOT_VIA_RHOA_ | 41 | 0.56 | 1.15 | 0.279 | 0.591 |
| 2368 | DACOSTA_ERCC3_ALLELE_XPCS_VS_TTD_UP | 21 | 0.62 | 1.15 | 0.297 | 0.59 |
| 2369 | MIR519A_2_5P_MIR520B_5P | 170 | 0.49 | 1.15 | 0.171 | 0.59 |
| 2370 | ZHAN_MULTIPLE_MYELOMA_CD1_VS_CD2_UP | 52 | 0.55 | 1.15 | 0.261 | 0.59 |
| 2371 | BROWNE_HCMV_INFECTION_12HR_DN | 79 | 0.52 | 1.15 | 0.22 | 0.59 |
| 2372 | FREAC4_01 | 107 | 0.51 | 1.15 | 0.223 | 0.59 |
| 2373 | NAKAYAMA_FRA2_TARGETS | 36 | 0.58 | 1.15 | 0.272 | 0.591 |

| | | | | | | |
|---|---|---|---|---|---|---|
| 2374 | WHITFIELD_CELL_CYCLE_G2_M | 182 | 0.49 | 1.15 | 0.188 | 0.592 |
| 2375 | LI_INDUCED_T_TO_NATURAL_KILLER_DN | 107 | 0.5 | 1.15 | 0.214 | 0.592 |
| 2376 | CAIRO_PML_TARGETS_BOUND_BY_MYC_UP | 21 | 0.62 | 1.15 | 0.287 | 0.592 |
| 2377 | CREIGHTON_ENDOCRINE_THERAPY_RESISTANCE | 459 | 0.47 | 1.15 | 0.086 | 0.592 |
| 2378 | MIR655_5P | 66 | 0.53 | 1.15 | 0.24 | 0.592 |
| 2379 | TORCHIA_TARGETS_OF_EWSR1_FLI1_FUSION_UF | 220 | 0.48 | 1.15 | 0.162 | 0.595 |
| 2380 | SIRT3_TARGET_GENES | 39 | 0.57 | 1.15 | 0.278 | 0.595 |
| 2381 | BURTON_ADIPOGENESIS_PEAK_AT_8HR | 33 | 0.57 | 1.15 | 0.265 | 0.595 |
| 2382 | ACAACCT_MIR453 | 39 | 0.56 | 1.15 | 0.272 | 0.596 |
| 2383 | MIR6883_5P | 348 | 0.47 | 1.15 | 0.115 | 0.596 |
| 2384 | MIR653_5P | 104 | 0.5 | 1.15 | 0.217 | 0.597 |
| 2385 | PID_THROMBIN_PAR1_PATHWAY | 36 | 0.57 | 1.15 | 0.286 | 0.598 |
| 2386 | PEREZ_TP63_TARGETS | 301 | 0.48 | 1.15 | 0.133 | 0.598 |
| 2387 | HALMOS_CEBPA_TARGETS_DN | 36 | 0.57 | 1.15 | 0.277 | 0.598 |
| 2388 | PODAR_RESPONSE_TO_ADAPHOSTIN_DN | 16 | 0.64 | 1.15 | 0.299 | 0.599 |
| 2389 | ROVERSI_GLIOMA_LOH_REGIONS | 33 | 0.57 | 1.15 | 0.277 | 0.599 |
| 2390 | HOOI_ST7_TARGETS_UP | 78 | 0.52 | 1.15 | 0.229 | 0.599 |
| 2391 | MIR573 | 71 | 0.52 | 1.15 | 0.243 | 0.599 |
| 2392 | BERENJENO_TRANSFORMED_BY_RHOA_DN | 345 | 0.47 | 1.15 | 0.13 | 0.6 |
| 2393 | WP_SEROTONIN_RECEPTOR_2_AND_ELKSRFGAT | 15 | 0.65 | 1.15 | 0.32 | 0.599 |
| 2394 | DAIRKEE_CANCER_PRONE_RESPONSE_BPA_E2 | 103 | 0.5 | 1.15 | 0.23 | 0.6 |
| 2395 | LIU_CMYB_TARGETS_UP | 149 | 0.49 | 1.15 | 0.193 | 0.6 |
| 2396 | LINDGREN_BLADDER_CANCER_CLUSTER_3_DN | 201 | 0.48 | 1.15 | 0.176 | 0.6 |
| 2397 | PDGF_ERK_DN.V1_DN | 133 | 0.5 | 1.15 | 0.212 | 0.6 |
| 2398 | MIR4733_5P | 78 | 0.52 | 1.15 | 0.244 | 0.6 |
| 2399 | MIR4272 | 175 | 0.49 | 1.15 | 0.181 | 0.6 |
| 2400 | YAUCH_HEDGEHOG_SIGNALING_PARACRINE_UP | 89 | 0.51 | 1.15 | 0.219 | 0.6 |
| 2401 | OSF2_Q6 | 199 | 0.48 | 1.15 | 0.179 | 0.6 |
| 2402 | ATGTCAC_MIR489 | 85 | 0.51 | 1.15 | 0.216 | 0.6 |
| 2403 | PILON_KLF1_TARGETS_UP | 361 | 0.47 | 1.15 | 0.126 | 0.6 |
| 2404 | BLANCO_MELO_BETA_INTERFERON_TREATED_BI | 143 | 0.49 | 1.15 | 0.189 | 0.601 |
| 2405 | HMGB2_TARGET_GENES | 314 | 0.47 | 1.14 | 0.156 | 0.602 |
| 2406 | HALMOS_CEBPA_TARGETS_UP | 38 | 0.56 | 1.14 | 0.265 | 0.602 |

| | | | | | | |
|---|---|---|---|---|---|---|
| 2407 | KRAS.600.LUNG.BREAST_UP.V1_DN | 167 | 0.49 | 1.14 | 0.187 | 0.602 |
| 2408 | REACTOME_ORC1_REMOVAL_FROM_CHROMATIN | 69 | 0.52 | 1.14 | 0.26 | 0.602 |
| 2409 | RODWELL_AGING_KIDNEY_NO_BLOOD_DN | 133 | 0.49 | 1.14 | 0.24 | 0.601 |
| 2410 | PID_INTEGRIN_CS_PATHWAY | 15 | 0.65 | 1.14 | 0.313 | 0.603 |
| 2411 | HILLION_HMGA1B_TARGETS | 72 | 0.52 | 1.14 | 0.242 | 0.604 |
| 2412 | LE_NEURONAL_DIFFERENTIATION_DN | 19 | 0.62 | 1.14 | 0.323 | 0.604 |
| 2413 | HOEBEKE_LYMPHOID_STEM_CELL_UP | 81 | 0.51 | 1.14 | 0.252 | 0.604 |
| 2414 | GCNP_SHH_UP_EARLY.V1_UP | 155 | 0.49 | 1.14 | 0.201 | 0.604 |
| 2415 | EPPERT_LSC_R | 36 | 0.57 | 1.14 | 0.284 | 0.605 |
| 2416 | WP_GPR143_IN_MELANOCYTES_AND_RETINAL_P | 22 | 0.6 | 1.14 | 0.297 | 0.605 |
| 2417 | KIM_MYCN_AMPLIFICATION_TARGETS_UP | 77 | 0.52 | 1.14 | 0.23 | 0.606 |
| 2418 | LIN_MELANOMA_COPY_NUMBER_UP | 62 | 0.53 | 1.14 | 0.248 | 0.607 |
| 2419 | GROSS_HYPOXIA_VIA_ELK3_AND_HIF1A_DN | 90 | 0.5 | 1.14 | 0.241 | 0.606 |
| 2420 | BILBAN_B_CLL_LPL_DN | 35 | 0.57 | 1.14 | 0.283 | 0.606 |
| 2421 | MULLIGHAN_NPM1_SIGNATURE_3_DN | 134 | 0.49 | 1.14 | 0.218 | 0.606 |
| 2422 | PID_E2F_PATHWAY | 70 | 0.52 | 1.14 | 0.253 | 0.606 |
| 2423 | ACTGTGA_MIR27A_MIR27B | 433 | 0.47 | 1.14 | 0.119 | 0.606 |
| 2424 | SENESE_HDAC2_TARGETS_DN | 92 | 0.5 | 1.14 | 0.235 | 0.607 |
| 2425 | WP_HIPPOYAP_SIGNALING_PATHWAY | 21 | 0.6 | 1.14 | 0.291 | 0.607 |
| 2426 | CAATGCA_MIR33 | 81 | 0.52 | 1.14 | 0.228 | 0.607 |
| 2427 | NR0B1_TARGET_GENES | 199 | 0.48 | 1.14 | 0.182 | 0.608 |
| 2428 | WP_ONECARBON_METABOLISM | 26 | 0.59 | 1.14 | 0.29 | 0.608 |
| 2429 | WP_VEGFAVEGFR2_SIGNALING_PATHWAY | 387 | 0.47 | 1.14 | 0.136 | 0.608 |
| 2430 | LUI_THYROID_CANCER_CLUSTER_1 | 47 | 0.55 | 1.14 | 0.261 | 0.608 |
| 2431 | MATTIOLI_MGUS_VS_PCL | 90 | 0.5 | 1.14 | 0.245 | 0.609 |
| 2432 | PID_MYC_PATHWAY | 25 | 0.6 | 1.14 | 0.287 | 0.609 |
| 2433 | DACOSTA_UV_RESPONSE_VIA_ERCC3_COMMON_ | 446 | 0.46 | 1.14 | 0.121 | 0.609 |
| 2434 | REACTOME_CHONDROITIN_SULFATE_DERMATAN_ | 44 | 0.55 | 1.14 | 0.265 | 0.609 |
| 2435 | OHGUCHI_LIVER_HNF4A_TARGETS_DN | 75 | 0.51 | 1.14 | 0.257 | 0.609 |
| 2436 | ABRAHAM_ALPC_VS_MULTIPLE_MYELOMA_DN | 20 | 0.62 | 1.14 | 0.303 | 0.61 |
| 2437 | ZFP36L1_TARGET_GENES | 15 | 0.65 | 1.14 | 0.325 | 0.611 |
| 2438 | NIKOLSKY_BREAST_CANCER_8Q23_Q24_AMPLIC | 123 | 0.49 | 1.14 | 0.22 | 0.611 |
| 2439 | TENEDINI_MEGAKARYOCYTE_MARKERS | 49 | 0.54 | 1.14 | 0.27 | 0.612 |

| | | | | | | |
|---|---|---|---|---|---|---|
| 2440 | ARP1_01 | 126 | 0.49 | 1.14 | 0.216 | 0.612 |
| 2441 | PID_TRAIL_PATHWAY | 24 | 0.61 | 1.14 | 0.33 | 0.613 |
| 2442 | MIR3124_3P | 182 | 0.48 | 1.14 | 0.201 | 0.613 |
| 2443 | NAKAYAMA_SOFT_TISSUE_TUMORS_PCA2_DN | 35 | 0.57 | 1.14 | 0.283 | 0.613 |
| 2444 | ACOSTA_PROLIFERATION_INDEPENDENT_MYC_T/ | 91 | 0.5 | 1.14 | 0.232 | 0.613 |
| 2445 | MIR4478 | 159 | 0.48 | 1.14 | 0.206 | 0.614 |
| 2446 | KEGG_AMINO_SUGAR_AND_NUCLEOTIDE_SUGAR | 41 | 0.55 | 1.14 | 0.259 | 0.613 |
| 2447 | REACTOME_NOTCH4_INTRACELLULAR_DOMAIN_F | 17 | 0.62 | 1.14 | 0.299 | 0.613 |
| 2448 | PID_ERBB1_DOWNSTREAM_PATHWAY | 101 | 0.5 | 1.14 | 0.252 | 0.614 |
| 2449 | GARY_CD5_TARGETS_DN | 420 | 0.47 | 1.14 | 0.126 | 0.615 |
| 2450 | HOXA4_Q2 | 197 | 0.48 | 1.14 | 0.186 | 0.615 |
| 2451 | MIR3976 | 88 | 0.51 | 1.14 | 0.237 | 0.615 |
| 2452 | GOLDRATH_IMMUNE_MEMORY | 55 | 0.54 | 1.14 | 0.259 | 0.614 |
| 2453 | ZNF41_TARGET_GENES | 74 | 0.52 | 1.14 | 0.238 | 0.614 |
| 2454 | MIR4652_3P | 282 | 0.47 | 1.14 | 0.177 | 0.615 |
| 2455 | WP_PROLACTIN_SIGNALING_PATHWAY | 73 | 0.51 | 1.14 | 0.229 | 0.615 |
| 2456 | WP_HEMATOPOIETIC_STEM_CELL_DIFFERENTIAT | 34 | 0.57 | 1.14 | 0.283 | 0.615 |
| 2457 | BOYLAN_MULTIPLE_MYELOMA_PCA3_DN | 47 | 0.54 | 1.14 | 0.292 | 0.616 |
| 2458 | PID_PTP1B_PATHWAY | 42 | 0.55 | 1.14 | 0.271 | 0.617 |
| 2459 | ROPERO_HDAC2_TARGETS | 77 | 0.51 | 1.14 | 0.24 | 0.617 |
| 2460 | POU3F2_01 | 66 | 0.52 | 1.14 | 0.269 | 0.617 |
| 2461 | AMIT_EGF_RESPONSE_480_HELA | 145 | 0.49 | 1.14 | 0.211 | 0.618 |
| 2462 | REACTOME_REGULATION_OF_MRNA_STABILITY_I | 85 | 0.51 | 1.14 | 0.254 | 0.618 |
| 2463 | ANDERSEN_CHOLANGIOCARCINOMA_CLASS1 | 42 | 0.55 | 1.14 | 0.29 | 0.619 |
| 2464 | WHITEHURST_PACLITAXEL_SENSITIVITY | 33 | 0.58 | 1.14 | 0.291 | 0.619 |
| 2465 | REACTOME_GLYCOSAMINOGLYCAN_METABOLISM | 97 | 0.5 | 1.14 | 0.244 | 0.618 |
| 2466 | GAUSSMANN_MLL_AF4_FUSION_TARGETS_G_DN | 32 | 0.57 | 1.14 | 0.289 | 0.618 |
| 2467 | MIR10397_3P | 39 | 0.55 | 1.14 | 0.291 | 0.618 |
| 2468 | CHEN_LVAD_SUPPORT_OF_FAILING_HEART_UP | 81 | 0.51 | 1.14 | 0.253 | 0.619 |
| 2469 | BILD_CTNNB1_ONCOGENIC_SIGNATURE | 66 | 0.52 | 1.14 | 0.247 | 0.619 |
| 2470 | MIR578 | 143 | 0.48 | 1.14 | 0.225 | 0.619 |
| 2471 | LEF1_UP.V1_UP | 128 | 0.49 | 1.14 | 0.241 | 0.62 |
| 2472 | MIR6823_3P | 46 | 0.55 | 1.14 | 0.281 | 0.62 |

| | | | | | | |
|---|---|---|---|---|---|---|
| 2473 | VETTER_TARGETS_OF_PRKCA_AND_ETS1_UP | 15 | 0.65 | 1.14 | 0.315 | 0.621 |
| 2474 | MIR874_3P | 48 | 0.54 | 1.14 | 0.273 | 0.621 |
| 2475 | MIR758_5P | 28 | 0.59 | 1.14 | 0.302 | 0.621 |
| 2476 | LIM_MAMMARY_LUMINAL_MATURE_UP | 87 | 0.5 | 1.14 | 0.263 | 0.621 |
| 2477 | MIR151A_3P | 27 | 0.58 | 1.14 | 0.305 | 0.621 |
| 2478 | JINESH_BLEBBISHIELD_TO_IMMUNE_CELL_FUSIO | 327 | 0.47 | 1.13 | 0.148 | 0.623 |
| 2479 | WP_NUCLEAR_RECEPTORS_IN_LIPID_METABOLIS | 18 | 0.61 | 1.13 | 0.312 | 0.624 |
| 2480 | KIM_LIVER_CANCER_POOR_SURVIVAL_DN | 17 | 0.63 | 1.13 | 0.319 | 0.624 |
| 2481 | PID_LYMPH_ANGIOGENESIS_PATHWAY | 21 | 0.61 | 1.13 | 0.307 | 0.624 |
| 2482 | HJURP_TARGET_GENES | 24 | 0.6 | 1.13 | 0.28 | 0.625 |
| 2483 | CEBP_Q3 | 190 | 0.48 | 1.13 | 0.199 | 0.626 |
| 2484 | EFC_Q6 | 225 | 0.47 | 1.13 | 0.189 | 0.626 |
| 2485 | CHEOK_RESPONSE_TO_MERCAPTOPURINE_AND_ | 15 | 0.63 | 1.13 | 0.337 | 0.628 |
| 2486 | KYNG_DNA_DAMAGE_BY_UV | 42 | 0.55 | 1.13 | 0.287 | 0.629 |
| 2487 | TGFB_UP.V1_UP | 150 | 0.48 | 1.13 | 0.237 | 0.629 |
| 2488 | WP_REGULATION_OF_ACTIN_CYTOSKELETON | 120 | 0.5 | 1.13 | 0.223 | 0.629 |
| 2489 | WP_TRANSCRIPTION_COFACTORS_SKI_AND_SKIL | 16 | 0.64 | 1.13 | 0.33 | 0.629 |
| 2490 | REACTOME_SYNTHESIS_OF_BILE_ACIDS_AND_BI | 20 | 0.61 | 1.13 | 0.313 | 0.629 |
| 2491 | GACAGGG_MIR339 | 61 | 0.52 | 1.13 | 0.272 | 0.629 |
| 2492 | GAGCTGG_MIR337 | 142 | 0.49 | 1.13 | 0.222 | 0.63 |
| 2493 | KEGG_GLYCEROLIPID_METABOLISM | 39 | 0.55 | 1.13 | 0.274 | 0.629 |
| 2494 | PID_RXR_VDR_PATHWAY | 24 | 0.6 | 1.13 | 0.322 | 0.629 |
| 2495 | MIR4772_3P | 76 | 0.51 | 1.13 | 0.277 | 0.63 |
| 2496 | MULLIGHAN_MLL_SIGNATURE_2_UP | 357 | 0.46 | 1.13 | 0.138 | 0.63 |
| 2497 | REACTOME_DEFECTIVE_B4GALT7_CAUSES_EDS_ | 17 | 0.63 | 1.13 | 0.316 | 0.63 |
| 2498 | MIR1914_3P | 22 | 0.6 | 1.13 | 0.32 | 0.63 |
| 2499 | SESTO_RESPONSE_TO_UV_C3 | 18 | 0.62 | 1.13 | 0.322 | 0.63 |
| 2500 | MIR6512_5P | 102 | 0.49 | 1.13 | 0.257 | 0.63 |
| 2501 | MIR6849_3P | 70 | 0.52 | 1.13 | 0.266 | 0.63 |
| 2502 | KORKOLA_SEMINOMA_UP | 39 | 0.55 | 1.13 | 0.293 | 0.63 |
| 2503 | REACTOME_SIGNALING_BY_RETINOIC_ACID | 31 | 0.57 | 1.13 | 0.3 | 0.63 |
| 2504 | MIR6847_5P | 130 | 0.49 | 1.13 | 0.245 | 0.631 |
| 2505 | YAMASHITA_LIVER_CANCER_STEM_CELL_UP | 37 | 0.56 | 1.13 | 0.29 | 0.63 |

| | | | | | | |
|---|---|---|---|---|---|---|
| 2506 | GCGCCTT_MIR525_MIR524 | 15 | 0.64 | 1.13 | 0.328 | 0.631 |
| 2507 | MIR4745_5P | 68 | 0.52 | 1.13 | 0.271 | 0.631 |
| 2508 | REACTOME_CLATHRIN_MEDIATED_ENDOCYTOSIS | 121 | 0.49 | 1.13 | 0.232 | 0.631 |
| 2509 | WP_G13_SIGNALING_PATHWAY | 37 | 0.56 | 1.13 | 0.282 | 0.631 |
| 2510 | MIR1202 | 61 | 0.53 | 1.13 | 0.263 | 0.631 |
| 2511 | YAP1_UP | 41 | 0.55 | 1.13 | 0.297 | 0.631 |
| 2512 | ZA_UNIPROT_Q9UM89_UNREVIEWED_TARGET_GE | 51 | 0.53 | 1.13 | 0.27 | 0.632 |
| 2513 | LEF1_Q2 | 183 | 0.48 | 1.13 | 0.205 | 0.632 |
| 2514 | HENDRICKS_SMARCA4_TARGETS_UP | 41 | 0.55 | 1.13 | 0.287 | 0.632 |
| 2515 | TURASHVILI_BREAST_DUCTAL_CARCINOMA_VS_D | 34 | 0.58 | 1.13 | 0.31 | 0.632 |
| 2516 | REACTOME_ACYL_CHAIN_REMODELLING_OF_PE | 15 | 0.65 | 1.13 | 0.343 | 0.632 |
| 2517 | MIR4531 | 284 | 0.46 | 1.13 | 0.178 | 0.632 |
| 2518 | QUINTENS_EMBRYONIC_BRAIN_RESPONSE_TO_II | 75 | 0.51 | 1.13 | 0.273 | 0.632 |
| 2519 | BREDEMEYER_RAG_SIGNALING_VIA_ATM_NOT_V | 47 | 0.54 | 1.13 | 0.28 | 0.632 |
| 2520 | MIR4766_3P | 127 | 0.49 | 1.13 | 0.236 | 0.633 |
| 2521 | CATTTCA_MIR203 | 260 | 0.47 | 1.13 | 0.193 | 0.633 |
| 2522 | SIMBULAN_UV_RESPONSE_IMMORTALIZED_DN | 28 | 0.58 | 1.13 | 0.317 | 0.633 |
| 2523 | RICKMAN_TUMOR_DIFFERENTIATED_WELL_VS_PC | 232 | 0.47 | 1.13 | 0.187 | 0.633 |
| 2524 | RAGHAVACHARI_PLATELET_SPECIFIC_GENES | 57 | 0.53 | 1.13 | 0.281 | 0.634 |
| 2525 | REACTOME_CELLULAR_RESPONSE_TO_HEAT_ST | 93 | 0.51 | 1.13 | 0.255 | 0.634 |
| 2526 | LUI_TARGETS_OF_PAX8_PPARG_FUSION | 34 | 0.56 | 1.13 | 0.299 | 0.634 |
| 2527 | REACTOME_TP53_REGULATES_METABOLIC_GENI | 85 | 0.5 | 1.13 | 0.261 | 0.635 |
| 2528 | ZWANG_EGF_INTERVAL_DN | 173 | 0.48 | 1.13 | 0.211 | 0.635 |
| 2529 | MIR4704_3P | 130 | 0.49 | 1.13 | 0.238 | 0.635 |
| 2530 | SRC_UP.V1_UP | 110 | 0.49 | 1.13 | 0.25 | 0.635 |
| 2531 | HLF_01 | 214 | 0.47 | 1.13 | 0.198 | 0.635 |
| 2532 | MIR377_5P | 40 | 0.55 | 1.13 | 0.316 | 0.635 |
| 2533 | CHUNG_BLISTER_CYTOTOXICITY_DN | 35 | 0.56 | 1.13 | 0.306 | 0.636 |
| 2534 | FOXQ1_TARGET_GENES | 37 | 0.56 | 1.13 | 0.295 | 0.635 |
| 2535 | CLIMENT_BREAST_CANCER_COPY_NUMBER_UP | 18 | 0.63 | 1.13 | 0.333 | 0.636 |
| 2536 | ATAGGAA_MIR202 | 96 | 0.5 | 1.13 | 0.26 | 0.636 |
| 2537 | MIR4329 | 63 | 0.53 | 1.13 | 0.269 | 0.636 |
| 2538 | ASTON_MAJOR_DEPRESSIVE_DISORDER_DN | 130 | 0.48 | 1.13 | 0.233 | 0.636 |

| | | | | | | |
|---|---|---|---|---|---|---|
| 2539 | TURASHVILI_BREAST_NORMAL_DUCTAL_VS_LOBI | 61 | 0.52 | 1.13 | 0.278 | 0.637 |
| 2540 | MIR6822_3P | 47 | 0.55 | 1.13 | 0.293 | 0.637 |
| 2541 | TAVAZOIE_METASTASIS | 63 | 0.52 | 1.13 | 0.278 | 0.637 |
| 2542 | BOUDOUKHA_BOUND_BY_IGF2BP2 | 99 | 0.5 | 1.13 | 0.262 | 0.639 |
| 2543 | REACTOME_TRANSCRIPTIONAL_REGULATION_BY_ | 89 | 0.5 | 1.13 | 0.254 | 0.639 |
| 2544 | REACTOME_NS1_MEDIATED_EFFECTS_ON_HOST_ | 40 | 0.55 | 1.13 | 0.296 | 0.639 |
| 2545 | ZHU_CMV_ALL_DN | 89 | 0.5 | 1.13 | 0.263 | 0.639 |
| 2546 | MIR4521 | 62 | 0.51 | 1.13 | 0.287 | 0.64 |
| 2547 | HUANG_FOXA2_TARGETS_DN | 20 | 0.62 | 1.13 | 0.323 | 0.64 |
| 2548 | RP58_01 | 155 | 0.48 | 1.13 | 0.227 | 0.64 |
| 2549 | MIR3122 | 53 | 0.53 | 1.13 | 0.298 | 0.64 |
| 2550 | WP_OVERLAP_BETWEEN_SIGNAL_TRANSDUCTIOI | 47 | 0.54 | 1.13 | 0.278 | 0.64 |
| 2551 | WANG_TUMOR_INVASIVENESS_UP | 359 | 0.46 | 1.13 | 0.169 | 0.64 |
| 2552 | KEGG_TYPE_II_DIABETES_MELLITUS | 38 | 0.56 | 1.13 | 0.289 | 0.64 |
| 2553 | CHNG_MULTIPLE_MYELOMA_HYPERPLOID_UP | 52 | 0.53 | 1.13 | 0.277 | 0.64 |
| 2554 | SP1_01 | 208 | 0.47 | 1.13 | 0.217 | 0.641 |
| 2555 | MIR27B_5P | 40 | 0.56 | 1.13 | 0.291 | 0.641 |
| 2556 | DITTMER_PTHLH_TARGETS_DN | 64 | 0.52 | 1.13 | 0.273 | 0.641 |
| 2557 | LEE_NEURAL_CREST_STEM_CELL_UP | 103 | 0.49 | 1.13 | 0.257 | 0.641 |
| 2558 | MIR4803 | 379 | 0.46 | 1.13 | 0.147 | 0.641 |
| 2559 | HUANG_GATA2_TARGETS_UP | 110 | 0.49 | 1.12 | 0.248 | 0.641 |
| 2560 | REACTOME_TRANSPORT_OF_INORGANIC_CATION | 73 | 0.51 | 1.12 | 0.253 | 0.642 |
| 2561 | AGCTCCT_MIR28 | 77 | 0.5 | 1.12 | 0.291 | 0.642 |
| 2562 | MIR4642 | 86 | 0.5 | 1.12 | 0.258 | 0.643 |
| 2563 | ROSS_AML_WITH_MLL_FUSIONS | 64 | 0.52 | 1.12 | 0.287 | 0.643 |
| 2564 | REACTOME_CDT1_ASSOCIATION_WITH_THE_CDC( | 57 | 0.52 | 1.12 | 0.299 | 0.644 |
| 2565 | WP_THYROID_HORMONES_PRODUCTION_AND_PE | 70 | 0.51 | 1.12 | 0.264 | 0.646 |
| 2566 | WATANABE_ULCERATIVE_COLITIS_WITH_CANCEF | 17 | 0.61 | 1.12 | 0.334 | 0.646 |
| 2567 | KEGG_DORSO_VENTRAL_AXIS_FORMATION | 22 | 0.59 | 1.12 | 0.316 | 0.646 |
| 2568 | JNK_DN.V1_UP | 119 | 0.49 | 1.12 | 0.246 | 0.646 |
| 2569 | MIR204_3P | 123 | 0.49 | 1.12 | 0.253 | 0.647 |
| 2570 | REACTOME_DEGRADATION_OF_GLI1_BY_THE_PF | 56 | 0.53 | 1.12 | 0.276 | 0.647 |
| 2571 | SU_LIVER | 16 | 0.63 | 1.12 | 0.336 | 0.648 |

| | | | | | | |
|---|---|---|---|---|---|---|
| 2572 | ZNF585B_TARGET_GENES | 107 | 0.49 | 1.12 | 0.259 | 0.648 |
| 2573 | MIR1273H_5P | 160 | 0.47 | 1.12 | 0.242 | 0.648 |
| 2574 | WAMUNYOKOLI_OVARIAN_CANCER_LMP_UP | 249 | 0.46 | 1.12 | 0.216 | 0.647 |
| 2575 | YOSHIMURA_MAPK8_TARGETS_DN | 312 | 0.46 | 1.12 | 0.184 | 0.648 |
| 2576 | MULLIGHAN_NPM1_MUTATED_SIGNATURE_1_DN | 106 | 0.49 | 1.12 | 0.267 | 0.651 |
| 2577 | BECKER_TAMOXIFEN_RESISTANCE_DN | 37 | 0.55 | 1.12 | 0.288 | 0.651 |
| 2578 | MIR3140_5P | 107 | 0.49 | 1.12 | 0.273 | 0.651 |
| 2579 | MIR6738_5P | 24 | 0.59 | 1.12 | 0.325 | 0.652 |
| 2580 | KEGG_THYROID_CANCER | 28 | 0.57 | 1.12 | 0.29 | 0.651 |
| 2581 | ELF1_Q6 | 202 | 0.47 | 1.12 | 0.223 | 0.651 |
| 2582 | YKACATTT_UNKNOWN | 234 | 0.47 | 1.12 | 0.216 | 0.653 |
| 2583 | SAFFORD_T_LYMPHOCYTE_ANERGY | 70 | 0.51 | 1.12 | 0.277 | 0.653 |
| 2584 | RICKMAN_HEAD_AND_NECK_CANCER_E | 24 | 0.58 | 1.12 | 0.344 | 0.654 |
| 2585 | CHYLA_CBFA2T3_TARGETS_DN | 180 | 0.47 | 1.12 | 0.234 | 0.654 |
| 2586 | REACTOME_E3_UBIQUITIN_LIGASES_UBIQUITINA[illegible] | 52 | 0.53 | 1.12 | 0.299 | 0.654 |
| 2587 | POMEROY_MEDULLOBLASTOMA_DESMOPLASIC_[illegible] | 57 | 0.52 | 1.12 | 0.287 | 0.655 |
| 2588 | WATANABE_RECTAL_CANCER_RADIOTHERAPY_R | 104 | 0.49 | 1.12 | 0.262 | 0.655 |
| 2589 | VALK_AML_CLUSTER_16 | 22 | 0.6 | 1.12 | 0.334 | 0.654 |
| 2590 | RACCACAR_AML_Q6 | 196 | 0.47 | 1.12 | 0.21 | 0.654 |
| 2591 | MIR4314 | 108 | 0.49 | 1.12 | 0.263 | 0.654 |
| 2592 | VALK_AML_CLUSTER_1 | 23 | 0.6 | 1.12 | 0.336 | 0.654 |
| 2593 | MIR12122 | 184 | 0.47 | 1.12 | 0.23 | 0.654 |
| 2594 | OZEN_MIR125B1_TARGETS | 21 | 0.61 | 1.12 | 0.336 | 0.655 |
| 2595 | MIR5089_5P | 89 | 0.5 | 1.12 | 0.252 | 0.656 |
| 2596 | YANG_BREAST_CANCER_ESR1_LASER_DN | 41 | 0.54 | 1.12 | 0.306 | 0.657 |
| 2597 | ESC_J1_UP_EARLY.V1_UP | 143 | 0.48 | 1.12 | 0.243 | 0.657 |
| 2598 | MIR8076 | 100 | 0.49 | 1.12 | 0.261 | 0.66 |
| 2599 | DAVICIONI_TARGETS_OF_PAX_FOXO1_FUSIONS_ | 211 | 0.47 | 1.12 | 0.225 | 0.66 |
| 2600 | NR1D1_TARGET_GENES | 106 | 0.49 | 1.12 | 0.253 | 0.661 |
| 2601 | RODRIGUES_DCC_TARGETS_DN | 105 | 0.49 | 1.12 | 0.258 | 0.661 |
| 2602 | HEIDENBLAD_AMPLICON_8Q24_DN | 34 | 0.56 | 1.12 | 0.308 | 0.661 |
| 2603 | MIR650 | 95 | 0.49 | 1.12 | 0.28 | 0.661 |
| 2604 | CRX_DN.V1_UP | 99 | 0.49 | 1.12 | 0.269 | 0.661 |

| | | | | | | |
|---|---|---|---|---|---|---|
| 2605 | MIR10393_5P | 122 | 0.49 | 1.12 | 0.254 | 0.663 |
| 2606 | MIR2116_5P | 120 | 0.49 | 1.12 | 0.244 | 0.664 |
| 2607 | GRUETZMANN_PANCREATIC_CANCER_DN | 156 | 0.48 | 1.12 | 0.249 | 0.664 |
| 2608 | YAO_TEMPORAL_RESPONSE_TO_PROGESTERON | 59 | 0.52 | 1.12 | 0.31 | 0.664 |
| 2609 | MIR4686 | 72 | 0.51 | 1.12 | 0.282 | 0.665 |
| 2610 | MIR6881_3P | 154 | 0.48 | 1.12 | 0.224 | 0.665 |
| 2611 | MIR6780A_5P | 181 | 0.47 | 1.12 | 0.246 | 0.665 |
| 2612 | GRADE_COLON_AND_RECTAL_CANCER_UP | 272 | 0.46 | 1.12 | 0.218 | 0.666 |
| 2613 | AML1_01 | 211 | 0.46 | 1.12 | 0.234 | 0.666 |
| 2614 | WANG_SMARCE1_TARGETS_UP | 207 | 0.46 | 1.11 | 0.239 | 0.666 |
| 2615 | KRAS.600_UP.V1_DN | 144 | 0.48 | 1.11 | 0.247 | 0.666 |
| 2616 | KOYAMA_SEMA3B_TARGETS_UP | 218 | 0.46 | 1.11 | 0.225 | 0.665 |
| 2617 | BONOME_OVARIAN_CANCER_SURVIVAL_SUBOPT | 425 | 0.46 | 1.11 | 0.173 | 0.665 |
| 2618 | MIR7845_5P | 49 | 0.53 | 1.11 | 0.309 | 0.665 |
| 2619 | WP_HEAD_AND_NECK_SQUAMOUS_CELL_CARCIN | 71 | 0.5 | 1.11 | 0.284 | 0.665 |
| 2620 | MIR5194 | 32 | 0.56 | 1.11 | 0.317 | 0.665 |
| 2621 | GATA_Q6 | 134 | 0.48 | 1.11 | 0.257 | 0.665 |
| 2622 | WP_TRANSSULFURATION_AND_ONECARBON_MET | 29 | 0.57 | 1.11 | 0.341 | 0.666 |
| 2623 | MIR6894_3P | 55 | 0.53 | 1.11 | 0.295 | 0.666 |
| 2624 | MIR6801_5P | 97 | 0.49 | 1.11 | 0.283 | 0.666 |
| 2625 | MIR2115_3P | 198 | 0.47 | 1.11 | 0.222 | 0.668 |
| 2626 | BROWNE_HCMV_INFECTION_48HR_UP | 153 | 0.47 | 1.11 | 0.244 | 0.668 |
| 2627 | GRE_C | 104 | 0.49 | 1.11 | 0.288 | 0.668 |
| 2628 | VALK_AML_CLUSTER_13 | 25 | 0.57 | 1.11 | 0.344 | 0.668 |
| 2629 | REACTOME_IRON_UPTAKE_AND_TRANSPORT | 45 | 0.53 | 1.11 | 0.305 | 0.668 |
| 2630 | MIR4529_5P | 46 | 0.53 | 1.11 | 0.31 | 0.668 |
| 2631 | RFX1_01 | 219 | 0.47 | 1.11 | 0.218 | 0.668 |
| 2632 | REACTOME_METABOLISM_OF_AMINO_ACIDS_ANI | 304 | 0.46 | 1.11 | 0.191 | 0.668 |
| 2633 | REACTOME_DEGRADATION_OF_DVL | 54 | 0.52 | 1.11 | 0.305 | 0.668 |
| 2634 | MIR5094 | 78 | 0.5 | 1.11 | 0.292 | 0.668 |
| 2635 | MIR377_3P | 208 | 0.47 | 1.11 | 0.237 | 0.669 |
| 2636 | BIOCARTA_ETS_PATHWAY | 16 | 0.63 | 1.11 | 0.331 | 0.67 |
| 2637 | DURCHDEWALD_SKIN_CARCINOGENESIS_UP | 68 | 0.52 | 1.11 | 0.297 | 0.67 |

| | | | | | | |
|---|---|---|---|---|---|---|
| 2638 | MIR4763_5P | 46 | 0.53 | 1.11 | 0.312 | 0.67 |
| 2639 | MIR12128 | 116 | 0.48 | 1.11 | 0.273 | 0.67 |
| 2640 | STAT4_01 | 189 | 0.47 | 1.11 | 0.234 | 0.672 |
| 2641 | MIR4433B_5P | 49 | 0.53 | 1.11 | 0.29 | 0.672 |
| 2642 | DUNNE_TARGETS_OF_AML1_MTG8_FUSION_DN | 17 | 0.61 | 1.11 | 0.363 | 0.673 |
| 2643 | YAO_TEMPORAL_RESPONSE_TO_PROGESTERON | 175 | 0.47 | 1.11 | 0.239 | 0.673 |
| 2644 | CTAGGAA_MIR384 | 58 | 0.51 | 1.11 | 0.302 | 0.673 |
| 2645 | REACTOME_PI_METABOLISM | 77 | 0.5 | 1.11 | 0.281 | 0.672 |
| 2646 | GROSS_HYPOXIA_VIA_HIF1A_DN | 92 | 0.49 | 1.11 | 0.306 | 0.672 |
| 2647 | MIR146A_3P | 176 | 0.47 | 1.11 | 0.258 | 0.673 |
| 2648 | PID_TCR_PATHWAY | 48 | 0.53 | 1.11 | 0.306 | 0.672 |
| 2649 | HNF3ALPHA_Q6 | 163 | 0.47 | 1.11 | 0.252 | 0.673 |
| 2650 | CSR_EARLY_UP.V1_UP | 141 | 0.47 | 1.11 | 0.257 | 0.673 |
| 2651 | MCBRYAN_PUBERTAL_TGFB1_TARGETS_UP | 141 | 0.48 | 1.11 | 0.256 | 0.672 |
| 2652 | MIR3934_5P | 55 | 0.52 | 1.11 | 0.303 | 0.672 |
| 2653 | REACTOME_ION_CHANNEL_TRANSPORT | 129 | 0.47 | 1.11 | 0.276 | 0.673 |
| 2654 | REACTOME_STIMULI_SENSING_CHANNELS | 71 | 0.51 | 1.11 | 0.293 | 0.674 |
| 2655 | BLANCO_MELO_COVID19_SARS_COV_2_INFECTIC | 49 | 0.52 | 1.11 | 0.308 | 0.674 |
| 2656 | REACTOME_RAC1_GTPASE_CYCLE | 163 | 0.47 | 1.11 | 0.238 | 0.674 |
| 2657 | MIR6769B_3P | 65 | 0.51 | 1.11 | 0.301 | 0.674 |
| 2658 | MIR1298_3P | 31 | 0.56 | 1.11 | 0.341 | 0.675 |
| 2659 | KRAS.600_UP.V1_UP | 169 | 0.47 | 1.11 | 0.237 | 0.675 |
| 2660 | RODRIGUES_NTN1_TARGETS_DN | 128 | 0.48 | 1.11 | 0.259 | 0.675 |
| 2661 | MCGARVEY_SILENCED_BY_METHYLATION_IN_CO | 23 | 0.58 | 1.11 | 0.361 | 0.674 |
| 2662 | PID_TRKR_PATHWAY | 55 | 0.52 | 1.11 | 0.311 | 0.676 |
| 2663 | MORI_LARGE_PRE_BII_LYMPHOCYTE_UP | 83 | 0.5 | 1.11 | 0.29 | 0.675 |
| 2664 | RAPA_EARLY_UP.V1_DN | 134 | 0.48 | 1.11 | 0.273 | 0.675 |
| 2665 | MIR616_3P | 33 | 0.56 | 1.11 | 0.317 | 0.676 |
| 2666 | MIR5002_5P | 224 | 0.47 | 1.11 | 0.242 | 0.677 |
| 2667 | MIR194_3P | 74 | 0.5 | 1.11 | 0.296 | 0.678 |
| 2668 | MIR4276 | 223 | 0.46 | 1.11 | 0.248 | 0.678 |
| 2669 | HUPER_BREAST_BASAL_VS_LUMINAL_DN | 48 | 0.53 | 1.11 | 0.306 | 0.678 |
| 2670 | RREB1_01 | 173 | 0.47 | 1.11 | 0.249 | 0.678 |

| | | | | | | |
|---|---|---|---|---|---|---|
| 2671 | MIR2110 | 149 | 0.47 | 1.11 | 0.277 | 0.678 |
| 2672 | KONDO_EZH2_TARGETS | 159 | 0.47 | 1.11 | 0.275 | 0.678 |
| 2673 | MIR6086 | 38 | 0.54 | 1.11 | 0.322 | 0.679 |
| 2674 | MIR4653_3P | 45 | 0.53 | 1.11 | 0.319 | 0.679 |
| 2675 | TACAATC_MIR508 | 59 | 0.52 | 1.11 | 0.301 | 0.679 |
| 2676 | MIR4743_3P | 173 | 0.47 | 1.11 | 0.248 | 0.679 |
| 2677 | REACTOME_SIGNALING_BY_NOTCH4 | 79 | 0.49 | 1.11 | 0.291 | 0.679 |
| 2678 | TERAMOTO_OPN_TARGETS_CLUSTER_7 | 15 | 0.63 | 1.11 | 0.363 | 0.679 |
| 2679 | MIR6873_3P | 316 | 0.45 | 1.11 | 0.208 | 0.679 |
| 2680 | REACTOME_RORA_ACTIVATES_GENE_EXPRESSIO | 18 | 0.61 | 1.11 | 0.35 | 0.68 |
| 2681 | MIR4723_3P | 65 | 0.51 | 1.11 | 0.302 | 0.682 |
| 2682 | SCHAEFFER_PROSTATE_DEVELOPMENT_AND_CA | 32 | 0.55 | 1.11 | 0.318 | 0.682 |
| 2683 | MIR149_3P | 374 | 0.45 | 1.11 | 0.204 | 0.683 |
| 2684 | REACTOME_ROS_AND_RNS_PRODUCTION_IN_PHA | 29 | 0.57 | 1.11 | 0.345 | 0.683 |
| 2685 | CHEN_LUNG_CANCER_SURVIVAL | 23 | 0.59 | 1.11 | 0.329 | 0.683 |
| 2686 | REACTOME_ACTIVATED_NOTCH1_TRANSMITS_SIG | 29 | 0.56 | 1.11 | 0.337 | 0.684 |
| 2687 | SMIRNOV_RESPONSE_TO_IR_2HR_DN | 51 | 0.52 | 1.11 | 0.333 | 0.684 |
| 2688 | REACTOME_INTERLEUKIN_12_FAMILY_SIGNALING | 47 | 0.52 | 1.11 | 0.327 | 0.683 |
| 2689 | SENESE_HDAC1_AND_HDAC2_TARGETS_DN | 177 | 0.47 | 1.11 | 0.268 | 0.683 |
| 2690 | FOROUTAN_INTEGRATED_TGFB_EMT_DN | 55 | 0.52 | 1.11 | 0.301 | 0.683 |
| 2691 | BOYLAN_MULTIPLE_MYELOMA_C_D_UP | 133 | 0.48 | 1.11 | 0.28 | 0.683 |
| 2692 | GOZGIT_ESR1_TARGETS_UP | 114 | 0.48 | 1.11 | 0.286 | 0.685 |
| 2693 | REACTOME_OXIDATIVE_STRESS_INDUCED_SENE | 93 | 0.49 | 1.11 | 0.301 | 0.685 |
| 2694 | PLASARI_TGFB1_TARGETS_10HR_DN | 190 | 0.47 | 1.1 | 0.264 | 0.686 |
| 2695 | MIR7156_3P | 81 | 0.5 | 1.1 | 0.287 | 0.686 |
| 2696 | PID_AMB2_NEUTROPHILS_PATHWAY | 32 | 0.56 | 1.1 | 0.318 | 0.686 |
| 2697 | TGGNNNNNNKCCAR_UNKNOWN | 340 | 0.45 | 1.1 | 0.193 | 0.686 |
| 2698 | CYCLIN_D1_KE_.V1_UP | 154 | 0.47 | 1.1 | 0.284 | 0.687 |
| 2699 | WP_FACTORS_AND_PATHWAYS_AFFECTING_INSU | 30 | 0.56 | 1.1 | 0.331 | 0.687 |
| 2700 | ZSCAN5C_TARGET_GENES | 117 | 0.48 | 1.1 | 0.293 | 0.687 |
| 2701 | ICHIBA_GRAFT_VERSUS_HOST_DISEASE_35D_DN | 28 | 0.57 | 1.1 | 0.339 | 0.687 |
| 2702 | MIR7108_5P | 39 | 0.54 | 1.1 | 0.338 | 0.687 |
| 2703 | WANG_NEOPLASTIC_TRANSFORMATION_BY_CCNI | 15 | 0.62 | 1.1 | 0.373 | 0.687 |

| | | | | | | |
|---|---|---|---|---|---|---|
| 2704 | STAT6_01 | 218 | 0.46 | 1.1 | 0.23 | 0.687 |
| 2705 | MIR374B_5P | 305 | 0.45 | 1.1 | 0.222 | 0.687 |
| 2706 | MIR1304_5P | 109 | 0.48 | 1.1 | 0.285 | 0.689 |
| 2707 | MIR6759_3P | 91 | 0.49 | 1.1 | 0.291 | 0.689 |
| 2708 | GAUSSMANN_MLL_AF4_FUSION_TARGETS_D_UP | 25 | 0.57 | 1.1 | 0.332 | 0.689 |
| 2709 | MYOGNF1_01 | 37 | 0.54 | 1.1 | 0.34 | 0.69 |
| 2710 | FRASOR_RESPONSE_TO_ESTRADIOL_DN | 61 | 0.51 | 1.1 | 0.315 | 0.69 |
| 2711 | NGUYEN_NOTCH1_TARGETS_DN | 74 | 0.49 | 1.1 | 0.307 | 0.69 |
| 2712 | MIR3612 | 96 | 0.49 | 1.1 | 0.3 | 0.691 |
| 2713 | MIR8073 | 69 | 0.5 | 1.1 | 0.306 | 0.693 |
| 2714 | ZNF133_TARGET_GENES | 130 | 0.48 | 1.1 | 0.285 | 0.693 |
| 2715 | HUTTMANN_B_CLL_POOR_SURVIVAL_UP | 226 | 0.46 | 1.1 | 0.247 | 0.694 |
| 2716 | MIR6797_3P | 55 | 0.52 | 1.1 | 0.304 | 0.694 |
| 2717 | WP_HAIR_FOLLICLE_DEVELOPMENT_CYTODIFFEI | 54 | 0.51 | 1.1 | 0.316 | 0.694 |
| 2718 | WP_LEPTIN_SIGNALING_PATHWAY | 71 | 0.5 | 1.1 | 0.302 | 0.694 |
| 2719 | MIR6830_5P | 159 | 0.47 | 1.1 | 0.283 | 0.694 |
| 2720 | MIR4700_5P | 185 | 0.47 | 1.1 | 0.266 | 0.695 |
| 2721 | MIR3622B_5P | 45 | 0.53 | 1.1 | 0.339 | 0.695 |
| 2722 | LET_7G_3P | 245 | 0.46 | 1.1 | 0.254 | 0.696 |
| 2723 | HEIDENBLAD_AMPLICON_12P11_12_DN | 22 | 0.59 | 1.1 | 0.35 | 0.695 |
| 2724 | REACTOME_FATTY_ACYL_COA_BIOSYNTHESIS | 30 | 0.56 | 1.1 | 0.362 | 0.695 |
| 2725 | MARTINEZ_TP53_TARGETS_UP | 475 | 0.45 | 1.1 | 0.196 | 0.695 |
| 2726 | CORONEL_RFX7_DIRECT_TARGETS_UP | 44 | 0.53 | 1.1 | 0.35 | 0.695 |
| 2727 | MIR7113_3P | 79 | 0.49 | 1.1 | 0.295 | 0.695 |
| 2728 | ATXN7L3_TARGET_GENES | 253 | 0.46 | 1.1 | 0.246 | 0.696 |
| 2729 | LET_7A_2_3P | 245 | 0.46 | 1.1 | 0.241 | 0.696 |
| 2730 | WP_HOSTPATHOGEN_INTERACTION_OF_HUMAN_( | 36 | 0.55 | 1.1 | 0.339 | 0.696 |
| 2731 | CHIANG_LIVER_CANCER_SUBCLASS_PROLIFERAT | 91 | 0.49 | 1.1 | 0.295 | 0.696 |
| 2732 | REACTOME_HEMOSTASIS | 430 | 0.45 | 1.1 | 0.193 | 0.697 |
| 2733 | RTTTNNNYTGGM_UNKNOWN | 116 | 0.48 | 1.1 | 0.283 | 0.697 |
| 2734 | MIR6829_5P | 80 | 0.49 | 1.1 | 0.292 | 0.697 |
| 2735 | HOFFMANN_LARGE_TO_SMALL_PRE_BII_LYMPHC | 49 | 0.53 | 1.1 | 0.315 | 0.697 |
| 2736 | ZNF704_TARGET_GENES | 64 | 0.51 | 1.1 | 0.323 | 0.697 |

| | | | | | | |
|---|---|---|---|---|---|---|
| 2737 | REACTOME_FOXO_MEDIATED_TRANSCRIPTION_C | 15 | 0.62 | 1.1 | 0.363 | 0.697 |
| 2738 | TURASHVILI_BREAST_CARCINOMA_DUCTAL_VS_L | 18 | 0.62 | 1.1 | 0.382 | 0.697 |
| 2739 | GAUSSMANN_MLL_AF4_FUSION_TARGETS_F_DN | 20 | 0.6 | 1.1 | 0.377 | 0.697 |
| 2740 | PID_MYC_ACTIV_PATHWAY | 75 | 0.5 | 1.1 | 0.307 | 0.698 |
| 2741 | FRASOR_TAMOXIFEN_RESPONSE_UP | 43 | 0.53 | 1.1 | 0.331 | 0.698 |
| 2742 | REACTOME_TRANSLATION_OF_SARS_COV_1_STI | 28 | 0.56 | 1.1 | 0.345 | 0.699 |
| 2743 | CCAWWNAAGG_SRF_Q4 | 69 | 0.5 | 1.1 | 0.313 | 0.699 |
| 2744 | PID_ILK_PATHWAY | 41 | 0.54 | 1.1 | 0.321 | 0.7 |
| 2745 | ENK_UV_RESPONSE_EPIDERMIS_DN | 437 | 0.45 | 1.1 | 0.203 | 0.7 |
| 2746 | AML1_Q6 | 211 | 0.46 | 1.1 | 0.265 | 0.7 |
| 2747 | WP_MAMMARY_GLAND_DEVELOPMENT_PATHWAY | 15 | 0.62 | 1.1 | 0.387 | 0.7 |
| 2748 | BLANCO_MELO_COVID19_SARS_COV_2_POS_PA | 170 | 0.47 | 1.1 | 0.285 | 0.7 |
| 2749 | MIR6815_3P | 184 | 0.46 | 1.1 | 0.271 | 0.701 |
| 2750 | BIOCARTA_MEF2D_PATHWAY | 16 | 0.61 | 1.1 | 0.372 | 0.701 |
| 2751 | REACTOME_DISEASES_OF_SIGNAL_TRANSDUCTI | 383 | 0.45 | 1.1 | 0.219 | 0.701 |
| 2752 | NKX22_01 | 145 | 0.47 | 1.1 | 0.29 | 0.701 |
| 2753 | WU_SILENCED_BY_METHYLATION_IN_BLADDER_ | 35 | 0.55 | 1.1 | 0.356 | 0.702 |
| 2754 | TERT_TARGET_GENES | 81 | 0.5 | 1.1 | 0.299 | 0.702 |
| 2755 | WP_GPCRS_CLASS_A_RHODOPSINLIKE | 62 | 0.51 | 1.1 | 0.344 | 0.702 |
| 2756 | CERVERA_SDHB_TARGETS_1_DN | 18 | 0.61 | 1.1 | 0.379 | 0.702 |
| 2757 | ZNF34_TARGET_GENES | 59 | 0.51 | 1.1 | 0.314 | 0.702 |
| 2758 | MIR3663_3P | 132 | 0.48 | 1.1 | 0.282 | 0.703 |
| 2759 | REACTOME_INTERLEUKIN_6_FAMILY_SIGNALING | 18 | 0.61 | 1.1 | 0.379 | 0.703 |
| 2760 | MIR3660 | 88 | 0.49 | 1.1 | 0.297 | 0.704 |
| 2761 | MIR3657 | 46 | 0.53 | 1.1 | 0.317 | 0.704 |
| 2762 | STAT6_02 | 201 | 0.46 | 1.1 | 0.265 | 0.705 |
| 2763 | ADNP_TARGET_GENES | 73 | 0.51 | 1.1 | 0.315 | 0.705 |
| 2764 | WP_PEPTIDE_GPCRS | 15 | 0.62 | 1.1 | 0.405 | 0.705 |
| 2765 | MIKKELSEN_MEF_HCP_WITH_H3_UNMETHYLATED | 129 | 0.47 | 1.1 | 0.286 | 0.705 |
| 2766 | WP_CHROMOSOMAL_AND_MICROSATELLITE_INST | 68 | 0.5 | 1.1 | 0.343 | 0.705 |
| 2767 | CHICAS_RB1_TARGETS_LOW_SERUM | 89 | 0.49 | 1.09 | 0.318 | 0.706 |
| 2768 | GENTLES_LEUKEMIC_STEM_CELL_UP | 19 | 0.6 | 1.09 | 0.386 | 0.706 |
| 2769 | ZHENG_GLIOBLASTOMA_PLASTICITY_DN | 44 | 0.53 | 1.09 | 0.324 | 0.706 |

| | | | | | | |
|---|---|---|---|---|---|---|
| 2770 | WP_PRADERWILLI_AND_ANGELMAN_SYNDROME | 40 | 0.54 | 1.09 | 0.335 | 0.706 |
| 2771 | WP_EXERCISEINDUCED_CIRCADIAN_REGULATION | 44 | 0.52 | 1.09 | 0.348 | 0.706 |
| 2772 | ZNF436_TARGET_GENES | 385 | 0.45 | 1.09 | 0.215 | 0.706 |
| 2773 | BIOCARTA_CALCINEURIN_PATHWAY | 18 | 0.6 | 1.09 | 0.359 | 0.707 |
| 2774 | REACTOME_CASPASE_ACTIVATION_VIA_EXTRINS | 23 | 0.58 | 1.09 | 0.376 | 0.707 |
| 2775 | FONTAINE_PAPILLARY_THYROID_CARCINOMA_UF | 49 | 0.51 | 1.09 | 0.335 | 0.708 |
| 2776 | MYB_Q5_01 | 217 | 0.46 | 1.09 | 0.262 | 0.707 |
| 2777 | WP_CARDIAC_HYPERTROPHIC_RESPONSE | 50 | 0.52 | 1.09 | 0.319 | 0.709 |
| 2778 | WP_BRAINDERIVED_NEUROTROPHIC_FACTOR_BD | 130 | 0.47 | 1.09 | 0.299 | 0.709 |
| 2779 | MIR449B_3P | 111 | 0.48 | 1.09 | 0.303 | 0.708 |
| 2780 | WP_OSTEOBLAST_DIFFERENTIATION_AND_RELAT | 87 | 0.48 | 1.09 | 0.316 | 0.708 |
| 2781 | KUNINGER_IGF1_VS_PDGFB_TARGETS_DN | 32 | 0.55 | 1.09 | 0.36 | 0.71 |
| 2782 | MARKEY_RB1_CHRONIC_LOF_UP | 96 | 0.48 | 1.09 | 0.325 | 0.711 |
| 2783 | MIR5002_3P | 40 | 0.53 | 1.09 | 0.34 | 0.711 |
| 2784 | SRF_Q6 | 196 | 0.46 | 1.09 | 0.276 | 0.711 |
| 2785 | PID_ANGIOPOIETIN_RECEPTOR_PATHWAY | 41 | 0.53 | 1.09 | 0.351 | 0.711 |
| 2786 | ZNF595_TARGET_GENES | 78 | 0.5 | 1.09 | 0.311 | 0.711 |
| 2787 | DACOSTA_UV_RESPONSE_VIA_ERCC3_UP | 305 | 0.45 | 1.09 | 0.239 | 0.711 |
| 2788 | KANG_FLUOROURACIL_RESISTANCE_DN | 15 | 0.61 | 1.09 | 0.385 | 0.711 |
| 2789 | MIR6758_3P | 91 | 0.49 | 1.09 | 0.307 | 0.71 |
| 2790 | REACTOME_PLATELET_ACTIVATION_SIGNALING_ | 184 | 0.46 | 1.09 | 0.273 | 0.71 |
| 2791 | DAVIES_MULTIPLE_MYELOMA_VS_MGUS_DN | 17 | 0.61 | 1.09 | 0.385 | 0.711 |
| 2792 | CLASPER_LYMPHATIC_VESSELS_DURING_METAS | 25 | 0.57 | 1.09 | 0.353 | 0.711 |
| 2793 | BIOCARTA_ARENRF2_PATHWAY | 18 | 0.59 | 1.09 | 0.374 | 0.711 |
| 2794 | MIR1200 | 288 | 0.45 | 1.09 | 0.256 | 0.712 |
| 2795 | PID_TGFBR_PATHWAY | 48 | 0.52 | 1.09 | 0.343 | 0.712 |
| 2796 | IWANAGA_CARCINOGENESIS_BY_KRAS_UP | 134 | 0.48 | 1.09 | 0.295 | 0.712 |
| 2797 | MIR384 | 95 | 0.48 | 1.09 | 0.314 | 0.713 |
| 2798 | MIR10400_3P_MIR4674 | 27 | 0.56 | 1.09 | 0.358 | 0.714 |
| 2799 | AMUNDSON_RESPONSE_TO_ARSENITE | 196 | 0.46 | 1.09 | 0.3 | 0.714 |
| 2800 | MIR7113_5P | 123 | 0.47 | 1.09 | 0.312 | 0.714 |
| 2801 | BRUINS_UVC_RESPONSE_VIA_TP53_GROUP_C | 66 | 0.5 | 1.09 | 0.33 | 0.714 |
| 2802 | GENTILE_UV_HIGH_DOSE_UP | 19 | 0.6 | 1.09 | 0.385 | 0.714 |

| | | | | | | |
|---|---|---|---|---|---|---|
| 2803 | PID_INTEGRIN_A4B1_PATHWAY | 28 | 0.56 | 1.09 | 0.363 | 0.714 |
| 2804 | ZNF454_TARGET_GENES | 16 | 0.61 | 1.09 | 0.396 | 0.714 |
| 2805 | GRAHAM_CML_QUIESCENT_VS_NORMAL_QUIESC | 70 | 0.49 | 1.09 | 0.339 | 0.714 |
| 2806 | LEE_CALORIE_RESTRICTION_MUSCLE_UP | 29 | 0.56 | 1.09 | 0.335 | 0.713 |
| 2807 | DORSEY_GAB2_TARGETS | 16 | 0.62 | 1.09 | 0.39 | 0.714 |
| 2808 | WP_PHYSICOCHEMICAL_FEATURES_AND_TOXICIT | 60 | 0.5 | 1.09 | 0.325 | 0.714 |
| 2809 | GTAGGCA_MIR189 | 27 | 0.56 | 1.09 | 0.381 | 0.714 |
| 2810 | WP_CANONICAL_AND_NONCANONICAL_TGFB_SIG | 15 | 0.62 | 1.09 | 0.397 | 0.715 |
| 2811 | FLECHNER_BIOPSY_KIDNEY_TRANSPLANT_OK_V | 21 | 0.59 | 1.09 | 0.382 | 0.716 |
| 2812 | MIR3190_5P | 104 | 0.48 | 1.09 | 0.334 | 0.716 |
| 2813 | WANG_RESPONSE_TO_GSK3_INHIBITOR_SB2167 | 326 | 0.45 | 1.09 | 0.266 | 0.716 |
| 2814 | REACTOME_ABC_FAMILY_PROTEINS_MEDIATED_ | 86 | 0.48 | 1.09 | 0.31 | 0.716 |
| 2815 | MORI_PLASMA_CELL_DN | 32 | 0.55 | 1.09 | 0.358 | 0.716 |
| 2816 | MIR552_3P | 52 | 0.52 | 1.09 | 0.349 | 0.716 |
| 2817 | MONNIER_POSTRADIATION_TUMOR_ESCAPE_DN | 334 | 0.45 | 1.09 | 0.254 | 0.717 |
| 2818 | WP_NEOVASCULARISATION_PROCESSES | 33 | 0.54 | 1.09 | 0.353 | 0.717 |
| 2819 | BHAT_ESR1_TARGETS_VIA_AKT1_UP | 245 | 0.45 | 1.09 | 0.267 | 0.718 |
| 2820 | SMAD3_Q6 | 186 | 0.46 | 1.09 | 0.294 | 0.718 |
| 2821 | MIR4791 | 55 | 0.51 | 1.09 | 0.323 | 0.718 |
| 2822 | ZNF165_TARGET_GENES | 71 | 0.49 | 1.09 | 0.315 | 0.718 |
| 2823 | HMGIY_Q6 | 185 | 0.46 | 1.09 | 0.29 | 0.718 |
| 2824 | MIR6783_3P | 174 | 0.46 | 1.09 | 0.298 | 0.718 |
| 2825 | MIR4766_5P | 275 | 0.45 | 1.09 | 0.278 | 0.718 |
| 2826 | MIR3619_5P | 251 | 0.45 | 1.09 | 0.276 | 0.718 |
| 2827 | NIKOLSKY_BREAST_CANCER_8P12_P11_AMPLIC | 41 | 0.52 | 1.09 | 0.35 | 0.719 |
| 2828 | MIR6859_5P | 175 | 0.46 | 1.09 | 0.29 | 0.719 |
| 2829 | RORIE_TARGETS_OF_EWSR1_FLI1_FUSION_UP | 23 | 0.59 | 1.09 | 0.364 | 0.718 |
| 2830 | WP_PROTEOGLYCAN_BIOSYNTHESIS | 18 | 0.59 | 1.09 | 0.389 | 0.719 |
| 2831 | MIR3654 | 42 | 0.53 | 1.09 | 0.349 | 0.719 |
| 2832 | STAT5A_04 | 161 | 0.46 | 1.09 | 0.308 | 0.721 |
| 2833 | MIR623 | 104 | 0.47 | 1.09 | 0.328 | 0.721 |
| 2834 | AFFAR_YY1_TARGETS_UP | 138 | 0.46 | 1.09 | 0.305 | 0.72 |
| 2835 | SIG_CD40PATHWAYMAP | 34 | 0.55 | 1.09 | 0.352 | 0.721 |

| | | | | | | |
|---|---|---|---|---|---|---|
| 2836 | MIR501_5P | 128 | 0.47 | 1.09 | 0.309 | 0.721 |
| 2837 | CONRAD_STEM_CELL | 27 | 0.57 | 1.09 | 0.372 | 0.721 |
| 2838 | MIR6742_5P | 37 | 0.54 | 1.09 | 0.36 | 0.721 |
| 2839 | MIR759 | 58 | 0.51 | 1.09 | 0.342 | 0.721 |
| 2840 | DACOSTA_UV_RESPONSE_VIA_ERCC3_XPCS_DN | 80 | 0.5 | 1.09 | 0.332 | 0.721 |
| 2841 | LIEN_BREAST_CARCINOMA_METAPLASTIC | 21 | 0.58 | 1.09 | 0.373 | 0.721 |
| 2842 | REACTOME_P75_NTR_RECEPTOR_MEDIATED_SIG | 89 | 0.49 | 1.09 | 0.325 | 0.721 |
| 2843 | MIR561_5P | 126 | 0.47 | 1.09 | 0.317 | 0.721 |
| 2844 | PXR_Q2 | 200 | 0.45 | 1.09 | 0.309 | 0.721 |
| 2845 | MIR499A_5P | 154 | 0.47 | 1.09 | 0.309 | 0.721 |
| 2846 | MIR6813_5P | 38 | 0.53 | 1.09 | 0.345 | 0.722 |
| 2847 | REACTOME_INTERLEUKIN_37_SIGNALING | 15 | 0.62 | 1.09 | 0.387 | 0.721 |
| 2848 | WP_T_CELL_RECEPTOR_AND_COSTIMULATORY_S | 24 | 0.57 | 1.09 | 0.357 | 0.722 |
| 2849 | MIR4526 | 86 | 0.49 | 1.09 | 0.338 | 0.722 |
| 2850 | PBX1_01 | 186 | 0.46 | 1.09 | 0.29 | 0.722 |
| 2851 | LEE_CALORIE_RESTRICTION_NEOCORTEX_UP | 55 | 0.51 | 1.09 | 0.342 | 0.722 |
| 2852 | KOBAYASHI_EGFR_SIGNALING_24HR_DN | 237 | 0.45 | 1.09 | 0.261 | 0.722 |
| 2853 | MIR5584_3P | 123 | 0.47 | 1.09 | 0.32 | 0.722 |
| 2854 | ODONNELL_METASTASIS_DN | 18 | 0.59 | 1.09 | 0.38 | 0.722 |
| 2855 | SHEPARD_BMYB_MORPHOLINO_UP | 184 | 0.46 | 1.08 | 0.29 | 0.722 |
| 2856 | HOEBEKE_LYMPHOID_STEM_CELL_DN | 80 | 0.49 | 1.08 | 0.323 | 0.722 |
| 2857 | MIR2276_3P | 104 | 0.47 | 1.08 | 0.321 | 0.722 |
| 2858 | NFY_01 | 227 | 0.45 | 1.08 | 0.291 | 0.722 |
| 2859 | MIR583 | 149 | 0.47 | 1.08 | 0.316 | 0.723 |
| 2860 | KEGG_LONG_TERM_DEPRESSION | 51 | 0.51 | 1.08 | 0.345 | 0.723 |
| 2861 | MIR3123 | 193 | 0.46 | 1.08 | 0.281 | 0.723 |
| 2862 | HAHTOLA_SEZARY_SYNDROM_DN | 22 | 0.58 | 1.08 | 0.345 | 0.723 |
| 2863 | SENGUPTA_NASOPHARYNGEAL_CARCINOMA_WIT | 70 | 0.49 | 1.08 | 0.341 | 0.724 |
| 2864 | SCHAEFFER_PROSTATE_DEVELOPMENT_48HR_D | 293 | 0.45 | 1.08 | 0.28 | 0.724 |
| 2865 | SHIPP_DLBCL_VS_FOLLICULAR_LYMPHOMA_DN | 24 | 0.57 | 1.08 | 0.385 | 0.724 |
| 2866 | GGAMTNNNNNTCCY_UNKNOWN | 112 | 0.47 | 1.08 | 0.329 | 0.724 |
| 2867 | MARTIN_INTERACT_WITH_HDAC | 36 | 0.54 | 1.08 | 0.35 | 0.724 |
| 2868 | MIR6750_3P | 60 | 0.5 | 1.08 | 0.345 | 0.724 |

| | | | | | | |
|---|---|---|---|---|---|---|
| 2869 | MIR6719_3P | 106 | 0.47 | 1.08 | 0.319 | 0.724 |
| 2870 | SHEPARD_BMYB_MORPHOLINO_DN | 161 | 0.46 | 1.08 | 0.306 | 0.724 |
| 2871 | MIR629_3P | 155 | 0.46 | 1.08 | 0.302 | 0.724 |
| 2872 | SIX1_TARGET_GENES | 273 | 0.45 | 1.08 | 0.276 | 0.724 |
| 2873 | KARLSSON_TGFB1_TARGETS_DN | 178 | 0.46 | 1.08 | 0.292 | 0.725 |
| 2874 | BROWNE_HCMV_INFECTION_30MIN_DN | 99 | 0.47 | 1.08 | 0.315 | 0.726 |
| 2875 | CHO_NR4A1_TARGETS | 24 | 0.58 | 1.08 | 0.378 | 0.726 |
| 2876 | WP_PROXIMAL_TUBULE_TRANSPORT | 36 | 0.53 | 1.08 | 0.368 | 0.726 |
| 2877 | NAKAMURA_ADIPOGENESIS_EARLY_DN | 23 | 0.57 | 1.08 | 0.376 | 0.727 |
| 2878 | RODRIGUES_NTN1_AND_DCC_TARGETS | 32 | 0.55 | 1.08 | 0.372 | 0.727 |
| 2879 | RAF_UP.V1_DN | 153 | 0.46 | 1.08 | 0.298 | 0.728 |
| 2880 | WP_PHYSIOLOGICAL_AND_PATHOLOGICAL_HYPE | 22 | 0.58 | 1.08 | 0.375 | 0.728 |
| 2881 | RB_P130_DN.V1_DN | 117 | 0.47 | 1.08 | 0.331 | 0.728 |
| 2882 | REACTOME_MITOTIC_G1_PHASE_AND_G1_S_TRA | 147 | 0.47 | 1.08 | 0.306 | 0.728 |
| 2883 | MIKKELSEN_MCV6_ICP_WITH_H3K4ME3_AND_H3K | 18 | 0.6 | 1.08 | 0.389 | 0.728 |
| 2884 | ZNF596_TARGET_GENES | 488 | 0.44 | 1.08 | 0.223 | 0.728 |
| 2885 | ATF4_Q2 | 229 | 0.45 | 1.08 | 0.282 | 0.728 |
| 2886 | WP_RANKLRANK_SIGNALING_PATHWAY | 48 | 0.52 | 1.08 | 0.349 | 0.728 |
| 2887 | GGCAGAC_MIR346 | 35 | 0.53 | 1.08 | 0.366 | 0.728 |
| 2888 | TONKS_TARGETS_OF_RUNX1_RUNX1T1_FUSION_ | 34 | 0.54 | 1.08 | 0.348 | 0.728 |
| 2889 | REACTOME_LDL_CLEARANCE | 17 | 0.6 | 1.08 | 0.415 | 0.728 |
| 2890 | ACATTCC_MIR1_MIR206 | 279 | 0.45 | 1.08 | 0.278 | 0.729 |
| 2891 | MIR301A_5P | 74 | 0.49 | 1.08 | 0.344 | 0.729 |
| 2892 | ZNF586_TARGET_GENES | 29 | 0.55 | 1.08 | 0.373 | 0.73 |
| 2893 | MIR6785_5P | 438 | 0.44 | 1.08 | 0.246 | 0.73 |
| 2894 | KOYAMA_SEMA3B_TARGETS_DN | 313 | 0.44 | 1.08 | 0.282 | 0.73 |
| 2895 | MIR6792_3P | 102 | 0.47 | 1.08 | 0.335 | 0.731 |
| 2896 | SNAPC4_TARGET_GENES | 73 | 0.49 | 1.08 | 0.331 | 0.732 |
| 2897 | SEKI_INFLAMMATORY_RESPONSE_LPS_DN | 15 | 0.61 | 1.08 | 0.389 | 0.732 |
| 2898 | WP_WNT_SIGNALING_IN_KIDNEY_DISEASE | 29 | 0.55 | 1.08 | 0.363 | 0.732 |
| 2899 | MILI_PSEUDOPODIA_CHEMOTAXIS_DN | 398 | 0.44 | 1.08 | 0.264 | 0.732 |
| 2900 | OUYANG_PROSTATE_CANCER_MARKERS | 18 | 0.59 | 1.08 | 0.405 | 0.733 |
| 2901 | RB_P107_DN.V1_DN | 113 | 0.48 | 1.08 | 0.34 | 0.734 |

| | | | | | | |
|---|---|---|---|---|---|---|
| 2902 | MIR6501_3P | 128 | 0.47 | 1.08 | 0.313 | 0.734 |
| 2903 | PEDERSEN_METASTASIS_BY_ERBB2_ISOFORM_4 | 78 | 0.49 | 1.08 | 0.332 | 0.734 |
| 2904 | MIR6755_5P | 26 | 0.56 | 1.08 | 0.377 | 0.734 |
| 2905 | MCCABE_BOUND_BY_HOXC6 | 318 | 0.45 | 1.08 | 0.258 | 0.734 |
| 2906 | MIR203A_3P | 303 | 0.45 | 1.08 | 0.286 | 0.734 |
| 2907 | MIR3916 | 185 | 0.46 | 1.08 | 0.304 | 0.734 |
| 2908 | JIANG_HYPOXIA_CANCER | 56 | 0.51 | 1.08 | 0.356 | 0.734 |
| 2909 | KYNG_ENVIRONMENTAL_STRESS_RESPONSE_NC | 36 | 0.54 | 1.08 | 0.376 | 0.734 |
| 2910 | TATCTGG_MIR488 | 56 | 0.5 | 1.08 | 0.37 | 0.735 |
| 2911 | SPIELMAN_LYMPHOBLAST_EUROPEAN_VS_ASIAN | 15 | 0.62 | 1.08 | 0.403 | 0.735 |
| 2912 | DELACROIX_RAR_TARGETS_DN | 16 | 0.6 | 1.08 | 0.416 | 0.735 |
| 2913 | REACTOME_DNA_DOUBLE_STRAND_BREAK_REPA | 147 | 0.46 | 1.08 | 0.338 | 0.734 |
| 2914 | KEGG_COLORECTAL_CANCER | 59 | 0.51 | 1.08 | 0.361 | 0.734 |
| 2915 | MEK_UP.V1_UP | 159 | 0.46 | 1.08 | 0.341 | 0.736 |
| 2916 | AGCATTA_MIR155 | 116 | 0.47 | 1.08 | 0.325 | 0.736 |
| 2917 | YWATTWNNRGCT_UNKNOWN | 54 | 0.5 | 1.08 | 0.344 | 0.736 |
| 2918 | MIR636 | 196 | 0.45 | 1.08 | 0.318 | 0.736 |
| 2919 | WIERENGA_PML_INTERACTOME | 38 | 0.53 | 1.08 | 0.362 | 0.737 |
| 2920 | PU1_Q6 | 183 | 0.45 | 1.08 | 0.313 | 0.738 |
| 2921 | BAHD1_TARGET_GENES | 18 | 0.59 | 1.08 | 0.373 | 0.738 |
| 2922 | REACTOME_HSF1_DEPENDENT_TRANSACTIVATIO | 31 | 0.54 | 1.08 | 0.389 | 0.738 |
| 2923 | GGARNTKYCCA_UNKNOWN | 61 | 0.5 | 1.08 | 0.361 | 0.738 |
| 2924 | ZNF257_TARGET_GENES | 481 | 0.44 | 1.08 | 0.229 | 0.738 |
| 2925 | REACTOME_KERATAN_SULFATE_BIOSYNTHESIS | 18 | 0.6 | 1.08 | 0.394 | 0.738 |
| 2926 | MIKKELSEN_NPC_ICP_WITH_H3K4ME3 | 367 | 0.44 | 1.08 | 0.294 | 0.738 |
| 2927 | KEGG_OLFACTORY_TRANSDUCTION | 19 | 0.59 | 1.08 | 0.377 | 0.74 |
| 2928 | TGGTGCT_MIR29A_MIR29B_MIR29C | 468 | 0.44 | 1.08 | 0.253 | 0.741 |
| 2929 | STK33_SKM_DN | 201 | 0.45 | 1.08 | 0.314 | 0.741 |
| 2930 | TATTATA_MIR374 | 256 | 0.45 | 1.08 | 0.297 | 0.741 |
| 2931 | MIR301B_5P | 74 | 0.49 | 1.08 | 0.354 | 0.741 |
| 2932 | CTCAAGA_MIR526B | 60 | 0.5 | 1.08 | 0.339 | 0.741 |
| 2933 | BRUNO_HEMATOPOIESIS | 46 | 0.51 | 1.08 | 0.35 | 0.741 |
| 2934 | BIOCARTA_ALK_PATHWAY | 23 | 0.57 | 1.07 | 0.395 | 0.742 |

| | | | | | | |
|---|---|---|---|---|---|---|
| 2935 | MIR6830_3P | 162 | 0.46 | 1.07 | 0.348 | 0.742 |
| 2936 | MIR450A_2_3P | 110 | 0.47 | 1.07 | 0.348 | 0.742 |
| 2937 | ROESSLER_LIVER_CANCER_METASTASIS_DN | 44 | 0.51 | 1.07 | 0.382 | 0.743 |
| 2938 | PR_01 | 116 | 0.47 | 1.07 | 0.355 | 0.743 |
| 2939 | REACTOME_RHOC_GTPASE_CYCLE | 73 | 0.49 | 1.07 | 0.347 | 0.743 |
| 2940 | DANG_REGULATED_BY_MYC_UP | 65 | 0.5 | 1.07 | 0.351 | 0.743 |
| 2941 | MASSARWEH_TAMOXIFEN_RESISTANCE_UP | 497 | 0.44 | 1.07 | 0.258 | 0.743 |
| 2942 | GATA_C | 189 | 0.45 | 1.07 | 0.334 | 0.743 |
| 2943 | WP_CIRCADIAN_RHYTHM_GENES | 160 | 0.46 | 1.07 | 0.339 | 0.743 |
| 2944 | PID_HDAC_CLASSI_PATHWAY | 64 | 0.49 | 1.07 | 0.358 | 0.744 |
| 2945 | FARMER_BREAST_CANCER_CLUSTER_7 | 16 | 0.6 | 1.07 | 0.42 | 0.745 |
| 2946 | MIR4316 | 121 | 0.47 | 1.07 | 0.335 | 0.744 |
| 2947 | CAIRO_HEPATOBLASTOMA_DN | 157 | 0.46 | 1.07 | 0.327 | 0.745 |
| 2948 | MIR4455 | 87 | 0.47 | 1.07 | 0.344 | 0.745 |
| 2949 | MIR2054 | 134 | 0.46 | 1.07 | 0.334 | 0.745 |
| 2950 | MIR3121_5P | 87 | 0.47 | 1.07 | 0.342 | 0.746 |
| 2951 | REACTOME_INTRINSIC_PATHWAY_FOR_APOPTOS | 54 | 0.51 | 1.07 | 0.382 | 0.746 |
| 2952 | HOOI_ST7_TARGETS_DN | 94 | 0.48 | 1.07 | 0.335 | 0.746 |
| 2953 | PID_INSULIN_PATHWAY | 41 | 0.53 | 1.07 | 0.361 | 0.746 |
| 2954 | CRX_NRL_DN.V1_UP | 106 | 0.47 | 1.07 | 0.349 | 0.746 |
| 2955 | REACTOME_GASTRIN_CREB_SIGNALLING_PATHW | 15 | 0.61 | 1.07 | 0.414 | 0.747 |
| 2956 | PURBEY_TARGETS_OF_CTBP1_NOT_SATB1_DN | 308 | 0.44 | 1.07 | 0.316 | 0.746 |
| 2957 | WP_TGFBETA_SIGNALING_PATHWAY | 123 | 0.46 | 1.07 | 0.335 | 0.747 |
| 2958 | MYAATNNNNNNNGGC_UNKNOWN | 95 | 0.48 | 1.07 | 0.358 | 0.747 |
| 2959 | MIR6760_5P | 94 | 0.47 | 1.07 | 0.347 | 0.747 |
| 2960 | GTF2E2_TARGET_GENES | 326 | 0.44 | 1.07 | 0.306 | 0.746 |
| 2961 | JOHNSTONE_PARVB_TARGETS_1_DN | 57 | 0.5 | 1.07 | 0.357 | 0.747 |
| 2962 | MIR1296_5P | 29 | 0.56 | 1.07 | 0.397 | 0.747 |
| 2963 | TSENG_ADIPOGENIC_POTENTIAL_UP | 27 | 0.56 | 1.07 | 0.404 | 0.747 |
| 2964 | NATSUME_RESPONSE_TO_INTERFERON_BETA_DI | 46 | 0.51 | 1.07 | 0.373 | 0.747 |
| 2965 | MIR4744 | 143 | 0.46 | 1.07 | 0.347 | 0.747 |
| 2966 | IWANAGA_CARCINOGENESIS_BY_KRAS_PTEN_UI | 144 | 0.46 | 1.07 | 0.342 | 0.747 |
| 2967 | SCHLOSSER_SERUM_RESPONSE_UP | 111 | 0.47 | 1.07 | 0.345 | 0.748 |

| | | | | | | |
|---|---|---|---|---|---|---|
| 2968 | WP_PREIMPLANTATION_EMBRYO | 37 | 0.53 | 1.07 | 0.381 | 0.748 |
| 2969 | WP_PARKINSONS_DISEASE_PATHWAY | 37 | 0.53 | 1.07 | 0.351 | 0.748 |
| 2970 | MIR4693_3P | 48 | 0.51 | 1.07 | 0.382 | 0.748 |
| 2971 | PID_WNT_NONCANONICAL_PATHWAY | 30 | 0.55 | 1.07 | 0.386 | 0.75 |
| 2972 | GRAHAM_NORMAL_QUIESCENT_VS_NORMAL_DIV | 83 | 0.49 | 1.07 | 0.367 | 0.75 |
| 2973 | IWANAGA_CARCINOGENESIS_BY_KRAS_PTEN_DN | 256 | 0.44 | 1.07 | 0.327 | 0.75 |
| 2974 | ABE_VEGFA_TARGETS_30MIN | 20 | 0.58 | 1.07 | 0.402 | 0.751 |
| 2975 | GCACCTT_MIR18A_MIR18B | 110 | 0.46 | 1.07 | 0.337 | 0.751 |
| 2976 | OSWALD_HEMATOPOIETIC_STEM_CELL_IN_COLL | 232 | 0.45 | 1.07 | 0.33 | 0.751 |
| 2977 | REACTOME_NEGATIVE_REGULATION_OF_MAPK_I | 41 | 0.52 | 1.07 | 0.392 | 0.751 |
| 2978 | YAGI_AML_WITH_11Q23_REARRANGED | 292 | 0.45 | 1.07 | 0.298 | 0.752 |
| 2979 | MIR5695 | 79 | 0.48 | 1.07 | 0.372 | 0.752 |
| 2980 | SIPA1_TARGET_GENES | 113 | 0.46 | 1.07 | 0.361 | 0.752 |
| 2981 | BHATI_G2M_ARREST_BY_2METHOXYESTRADIOL_ | 100 | 0.47 | 1.07 | 0.348 | 0.752 |
| 2982 | SHEPARD_CRASH_AND_BURN_MUTANT_UP | 185 | 0.45 | 1.07 | 0.328 | 0.755 |
| 2983 | MIR5009_3P | 145 | 0.45 | 1.07 | 0.328 | 0.754 |
| 2984 | MIR4468 | 92 | 0.48 | 1.07 | 0.37 | 0.755 |
| 2985 | BILANGES_RAPAMYCIN_SENSITIVE_VIA_TSC1_AN | 65 | 0.49 | 1.07 | 0.36 | 0.755 |
| 2986 | MIR1237_3P | 108 | 0.47 | 1.07 | 0.333 | 0.755 |
| 2987 | SNF5_DN.V1_DN | 95 | 0.47 | 1.07 | 0.359 | 0.755 |
| 2988 | MIR627_3P | 394 | 0.44 | 1.07 | 0.284 | 0.754 |
| 2989 | VRTN_TARGET_GENES | 18 | 0.59 | 1.07 | 0.403 | 0.754 |
| 2990 | MIR4330 | 69 | 0.48 | 1.07 | 0.351 | 0.754 |
| 2991 | REACTOME_NON_INTEGRIN_MEMBRANE_ECM_INT | 41 | 0.52 | 1.07 | 0.385 | 0.754 |
| 2992 | ACEVEDO_METHYLATED_IN_LIVER_CANCER_DN | 479 | 0.44 | 1.07 | 0.28 | 0.754 |
| 2993 | CCCACAT_MIR2993P | 49 | 0.5 | 1.07 | 0.381 | 0.754 |
| 2994 | EVI1_06 | 15 | 0.59 | 1.07 | 0.415 | 0.754 |
| 2995 | MIR944 | 335 | 0.44 | 1.07 | 0.285 | 0.754 |
| 2996 | JOSEPH_RESPONSE_TO_SODIUM_BUTYRATE_UP | 27 | 0.56 | 1.07 | 0.4 | 0.754 |
| 2997 | YAATNRNNNYNATT_UNKNOWN | 79 | 0.48 | 1.07 | 0.368 | 0.754 |
| 2998 | MULLIGHAN_MLL_SIGNATURE_1_UP | 322 | 0.44 | 1.07 | 0.298 | 0.754 |
| 2999 | JAEGER_METASTASIS_UP | 37 | 0.52 | 1.07 | 0.384 | 0.755 |
| 3000 | GATA6_01 | 185 | 0.45 | 1.07 | 0.346 | 0.755 |

| | | | | | | |
|---|---|---|---|---|---|---|
| 3001 | OCT1_01 | 205 | 0.44 | 1.07 | 0.346 | 0.755 |
| 3002 | MATZUK_CENTRAL_FOR_FEMALE_FERTILITY | 17 | 0.59 | 1.07 | 0.428 | 0.756 |
| 3003 | MIR337_3P | 108 | 0.47 | 1.07 | 0.351 | 0.756 |
| 3004 | BIOCARTA_NTHI_PATHWAY | 21 | 0.57 | 1.07 | 0.417 | 0.757 |
| 3005 | MIR4263 | 227 | 0.45 | 1.07 | 0.342 | 0.759 |
| 3006 | MIR29B_3P_MIR29C_3P | 380 | 0.44 | 1.07 | 0.302 | 0.758 |
| 3007 | VART_KSHV_INFECTION_ANGIOGENIC_MARKERS_ | 94 | 0.47 | 1.07 | 0.373 | 0.758 |
| 3008 | PID_ARF6_PATHWAY | 25 | 0.56 | 1.07 | 0.405 | 0.76 |
| 3009 | CERVERA_SDHB_TARGETS_1_UP | 85 | 0.48 | 1.07 | 0.357 | 0.76 |
| 3010 | IL15_UP.V1_DN | 106 | 0.46 | 1.06 | 0.367 | 0.761 |
| 3011 | MIR4687_5P | 42 | 0.52 | 1.06 | 0.392 | 0.761 |
| 3012 | HDGF_TARGET_GENES | 247 | 0.44 | 1.06 | 0.333 | 0.761 |
| 3013 | GARGALOVIC_RESPONSE_TO_OXIDIZED_PHOSPH | 31 | 0.54 | 1.06 | 0.391 | 0.761 |
| 3014 | YYCATTCAWW_UNKNOWN | 158 | 0.45 | 1.06 | 0.34 | 0.761 |
| 3015 | GRAESSMANN_APOPTOSIS_BY_SERUM_DEPRIVA | 211 | 0.45 | 1.06 | 0.342 | 0.762 |
| 3016 | MIR634 | 75 | 0.48 | 1.06 | 0.352 | 0.762 |
| 3017 | KEGG_CALCIUM_SIGNALING_PATHWAY | 119 | 0.46 | 1.06 | 0.349 | 0.762 |
| 3018 | MIR4691_5P | 103 | 0.47 | 1.06 | 0.369 | 0.762 |
| 3019 | WP_SLEEP_REGULATION | 19 | 0.58 | 1.06 | 0.407 | 0.762 |
| 3020 | MIR4288 | 126 | 0.46 | 1.06 | 0.365 | 0.762 |
| 3021 | MIR24_3P | 245 | 0.44 | 1.06 | 0.327 | 0.763 |
| 3022 | MIR877_3P | 167 | 0.45 | 1.06 | 0.337 | 0.763 |
| 3023 | MIR29A_3P | 380 | 0.44 | 1.06 | 0.315 | 0.762 |
| 3024 | MIR4259 | 22 | 0.56 | 1.06 | 0.382 | 0.762 |
| 3025 | MIR6804_3P | 79 | 0.48 | 1.06 | 0.366 | 0.762 |
| 3026 | LEIN_NEURON_MARKERS | 63 | 0.49 | 1.06 | 0.386 | 0.762 |
| 3027 | CHIBA_RESPONSE_TO_TSA_UP | 36 | 0.53 | 1.06 | 0.387 | 0.762 |
| 3028 | MIR765 | 238 | 0.44 | 1.06 | 0.323 | 0.762 |
| 3029 | ODONNELL_TFRC_TARGETS_DN | 115 | 0.46 | 1.06 | 0.349 | 0.762 |
| 3030 | SA_B_CELL_RECEPTOR_COMPLEXES | 22 | 0.57 | 1.06 | 0.411 | 0.762 |
| 3031 | WP_ENDOMETRIAL_CANCER | 61 | 0.49 | 1.06 | 0.371 | 0.762 |
| 3032 | REACTOME_RHOU_GTPASE_CYCLE | 39 | 0.53 | 1.06 | 0.394 | 0.762 |
| 3033 | REACTOME_CELL_SURFACE_INTERACTIONS_AT_ | 80 | 0.48 | 1.06 | 0.355 | 0.763 |

| | | | | | | |
|---|---|---|---|---|---|---|
| 3034 | MIR6765_3P | 82 | 0.48 | 1.06 | 0.364 | 0.763 |
| 3035 | ROYLANCE_BREAST_CANCER_16Q_COPY_NUMBE | 51 | 0.51 | 1.06 | 0.372 | 0.763 |
| 3036 | MIKKELSEN_NPC_HCP_WITH_H3K4ME3_AND_H3K | 149 | 0.45 | 1.06 | 0.352 | 0.764 |
| 3037 | EN1_01 | 92 | 0.46 | 1.06 | 0.365 | 0.765 |
| 3038 | TIEN_INTESTINE_PROBIOTICS_24HR_DN | 205 | 0.45 | 1.06 | 0.345 | 0.766 |
| 3039 | SWEET_KRAS_TARGETS_DN | 48 | 0.51 | 1.06 | 0.383 | 0.767 |
| 3040 | SIG_PIP3_SIGNALING_IN_B_LYMPHOCYTES | 30 | 0.54 | 1.06 | 0.395 | 0.767 |
| 3041 | TTF1_Q6 | 217 | 0.44 | 1.06 | 0.336 | 0.767 |
| 3042 | BOYLAN_MULTIPLE_MYELOMA_D_CLUSTER_UP | 26 | 0.55 | 1.06 | 0.409 | 0.767 |
| 3043 | CDPCR3_01 | 38 | 0.53 | 1.06 | 0.405 | 0.767 |
| 3044 | MIR4474_3P | 71 | 0.48 | 1.06 | 0.385 | 0.769 |
| 3045 | LIU_VMYB_TARGETS_UP | 115 | 0.46 | 1.06 | 0.366 | 0.769 |
| 3046 | MODY_HIPPOCAMPUS_NEONATAL | 30 | 0.53 | 1.06 | 0.417 | 0.768 |
| 3047 | MIR885_5P | 102 | 0.47 | 1.06 | 0.386 | 0.768 |
| 3048 | CTGRYYYNATT_UNKNOWN | 53 | 0.5 | 1.06 | 0.379 | 0.769 |
| 3049 | MIKKELSEN_NPC_LCP_WITH_H3K4ME3 | 40 | 0.52 | 1.06 | 0.38 | 0.77 |
| 3050 | MIR5186 | 28 | 0.55 | 1.06 | 0.417 | 0.769 |
| 3051 | SATO_SILENCED_BY_DEACETYLATION_IN_PANCF | 31 | 0.54 | 1.06 | 0.389 | 0.77 |
| 3052 | MIR496 | 83 | 0.48 | 1.06 | 0.371 | 0.77 |
| 3053 | ODONNELL_METASTASIS_UP | 48 | 0.51 | 1.06 | 0.396 | 0.769 |
| 3054 | DAVICIONI_PAX_FOXO1_SIGNATURE_IN_ARMS_UI | 49 | 0.5 | 1.06 | 0.381 | 0.769 |
| 3055 | MIR6752_3P | 128 | 0.45 | 1.06 | 0.336 | 0.769 |
| 3056 | IK3_01 | 177 | 0.45 | 1.06 | 0.361 | 0.77 |
| 3057 | MIR2114_5P | 32 | 0.53 | 1.06 | 0.382 | 0.77 |
| 3058 | WP_INTERFERON_TYPE_I_SIGNALING_PATHWAYS | 54 | 0.5 | 1.06 | 0.388 | 0.77 |
| 3059 | HOLLERN_MICROACINAR_BREAST_TUMOR_UP | 34 | 0.52 | 1.06 | 0.39 | 0.77 |
| 3060 | MIR1286 | 73 | 0.48 | 1.06 | 0.38 | 0.77 |
| 3061 | WP_GLUCOCORTICOID_RECEPTOR_PATHWAY | 51 | 0.5 | 1.06 | 0.39 | 0.77 |
| 3062 | RB_DN.V1_UP | 106 | 0.46 | 1.06 | 0.373 | 0.772 |
| 3063 | WP_GLIOBLASTOMA_SIGNALING_PATHWAYS | 79 | 0.48 | 1.06 | 0.384 | 0.772 |
| 3064 | SASSON_RESPONSE_TO_GONADOTROPHINS_DN | 78 | 0.47 | 1.06 | 0.387 | 0.773 |
| 3065 | MIR589_5P | 76 | 0.48 | 1.06 | 0.376 | 0.773 |
| 3066 | KEGG_TIGHT_JUNCTION | 100 | 0.47 | 1.06 | 0.376 | 0.774 |

| | | | | | | |
|---|---|---|---|---|---|---|
| 3067 | REACTOME_RHO_GTPASE_EFFECTORS | 271 | 0.44 | 1.06 | 0.35 | 0.774 |
| 3068 | IK1_01 | 214 | 0.45 | 1.06 | 0.356 | 0.775 |
| 3069 | CTIP_DN.V1_DN | 82 | 0.47 | 1.06 | 0.388 | 0.775 |
| 3070 | MIR2053 | 139 | 0.45 | 1.06 | 0.377 | 0.774 |
| 3071 | MIR5692B_MIR5692C | 336 | 0.43 | 1.06 | 0.326 | 0.775 |
| 3072 | PID_RAC1_REG_PATHWAY | 34 | 0.52 | 1.06 | 0.397 | 0.775 |
| 3073 | GATA1_05 | 204 | 0.45 | 1.06 | 0.354 | 0.774 |
| 3074 | LEE_TARGETS_OF_PTCH1_AND_SUFU_UP | 46 | 0.5 | 1.06 | 0.378 | 0.775 |
| 3075 | MIR5682 | 186 | 0.45 | 1.06 | 0.357 | 0.775 |
| 3076 | MONNIER_POSTRADIATION_TUMOR_ESCAPE_UP | 380 | 0.43 | 1.06 | 0.334 | 0.775 |
| 3077 | MIR3679_3P | 144 | 0.45 | 1.06 | 0.365 | 0.775 |
| 3078 | MIR574_5P | 63 | 0.49 | 1.06 | 0.386 | 0.774 |
| 3079 | REACTOME_DNA_REPAIR | 305 | 0.43 | 1.06 | 0.333 | 0.775 |
| 3080 | BLANCO_MELO_COVID19_SARS_COV_2_INFECTIC | 68 | 0.48 | 1.06 | 0.385 | 0.775 |
| 3081 | FRASOR_RESPONSE_TO_SERM_OR_FULVESTRAN | 22 | 0.56 | 1.06 | 0.416 | 0.775 |
| 3082 | CTIP_DN.V1_UP | 71 | 0.48 | 1.06 | 0.39 | 0.775 |
| 3083 | MIR4308 | 124 | 0.46 | 1.06 | 0.363 | 0.774 |
| 3084 | MIR624_3P | 86 | 0.48 | 1.06 | 0.377 | 0.774 |
| 3085 | MIR6884_3P | 51 | 0.49 | 1.06 | 0.392 | 0.774 |
| 3086 | REACTOME_RHOQ_GTPASE_CYCLE | 54 | 0.5 | 1.06 | 0.394 | 0.776 |
| 3087 | MATTIOLI_MULTIPLE_MYELOMA_WITH_14Q32_TR/ | 29 | 0.54 | 1.06 | 0.41 | 0.776 |
| 3088 | ETS2_B | 220 | 0.44 | 1.06 | 0.357 | 0.777 |
| 3089 | PUJANA_BREAST_CANCER_LIT_INT_NETWORK | 98 | 0.47 | 1.06 | 0.358 | 0.777 |
| 3090 | PID_RB_1PATHWAY | 55 | 0.5 | 1.06 | 0.382 | 0.776 |
| 3091 | ZHAN_MULTIPLE_MYELOMA_LB_UP | 39 | 0.51 | 1.06 | 0.396 | 0.777 |
| 3092 | MIR133A_3P_MIR133B | 217 | 0.44 | 1.06 | 0.34 | 0.777 |
| 3093 | MIR637 | 76 | 0.48 | 1.06 | 0.4 | 0.777 |
| 3094 | MIR8078 | 20 | 0.58 | 1.06 | 0.442 | 0.777 |
| 3095 | REACTOME_SARS_COV_1_INFECTION | 49 | 0.5 | 1.06 | 0.386 | 0.777 |
| 3096 | KRAS.300_UP.V1_DN | 78 | 0.47 | 1.06 | 0.375 | 0.777 |
| 3097 | TCF11MAFG_01 | 171 | 0.45 | 1.05 | 0.348 | 0.777 |
| 3098 | ISSAEVA_MLL2_TARGETS | 40 | 0.52 | 1.05 | 0.399 | 0.777 |
| 3099 | MIR544B | 195 | 0.45 | 1.05 | 0.378 | 0.777 |

| | | | | | | |
|---|---|---|---|---|---|---|
| 3100 | MIR4727_5P | 101 | 0.46 | 1.05 | 0.378 | 0.777 |
| 3101 | REACTOME_PHOSPHOLIPID_METABOLISM | 173 | 0.44 | 1.05 | 0.351 | 0.777 |
| 3102 | MIR219A_2_3P | 52 | 0.5 | 1.05 | 0.398 | 0.777 |
| 3103 | LANDIS_ERBB2_BREAST_TUMORS_65_DN | 27 | 0.55 | 1.05 | 0.401 | 0.777 |
| 3104 | MIR4703_3P | 29 | 0.54 | 1.05 | 0.414 | 0.777 |
| 3105 | REACTOME_APC_C_MEDIATED_DEGRADATION_OI | 86 | 0.47 | 1.05 | 0.383 | 0.777 |
| 3106 | RAF_UP.V1_UP | 158 | 0.45 | 1.05 | 0.365 | 0.778 |
| 3107 | YAMAZAKI_TCEB3_TARGETS_UP | 136 | 0.45 | 1.05 | 0.376 | 0.778 |
| 3108 | CORRE_MULTIPLE_MYELOMA_DN | 38 | 0.51 | 1.05 | 0.381 | 0.778 |
| 3109 | MIR4749_3P | 45 | 0.51 | 1.05 | 0.403 | 0.779 |
| 3110 | WANG_HCP_PROSTATE_CANCER | 90 | 0.47 | 1.05 | 0.381 | 0.779 |
| 3111 | KRAS.300_UP.V1_UP | 86 | 0.47 | 1.05 | 0.371 | 0.779 |
| 3112 | HOXC11_TARGET_GENES | 44 | 0.51 | 1.05 | 0.402 | 0.779 |
| 3113 | MIR498_3P | 187 | 0.44 | 1.05 | 0.352 | 0.779 |
| 3114 | HMX1_01 | 38 | 0.52 | 1.05 | 0.407 | 0.78 |
| 3115 | TGAYRTCA_ATF3_Q6 | 421 | 0.43 | 1.05 | 0.338 | 0.78 |
| 3116 | MIR5691_MIR6805_3P | 132 | 0.45 | 1.05 | 0.381 | 0.78 |
| 3117 | WP_EICOSANOID_METABOLISM_VIA_CYCLOOXYG | 17 | 0.58 | 1.05 | 0.436 | 0.78 |
| 3118 | IGLV5_37_TARGET_GENES | 436 | 0.43 | 1.05 | 0.331 | 0.78 |
| 3119 | AGCACTT_MIR93_MIR302A_MIR302B_MIR302C_MII | 314 | 0.43 | 1.05 | 0.347 | 0.78 |
| 3120 | BENITEZ_GBM_PROTEASOME_INHIBITION_RESPC | 47 | 0.5 | 1.05 | 0.404 | 0.78 |
| 3121 | CCTGAGT_MIR510 | 40 | 0.51 | 1.05 | 0.39 | 0.78 |
| 3122 | DOANE_BREAST_CANCER_ESR1_UP | 77 | 0.47 | 1.05 | 0.389 | 0.781 |
| 3123 | KEGG_PEROXISOME | 70 | 0.48 | 1.05 | 0.382 | 0.781 |
| 3124 | XU_GH1_EXOGENOUS_TARGETS_DN | 71 | 0.48 | 1.05 | 0.393 | 0.78 |
| 3125 | FOXO1_01 | 198 | 0.44 | 1.05 | 0.368 | 0.782 |
| 3126 | WP_CLEAR_CELL_RENAL_CELL_CARCINOMA_PAT | 71 | 0.48 | 1.05 | 0.378 | 0.783 |
| 3127 | MOREAUX_MULTIPLE_MYELOMA_BY_TACI_UP | 256 | 0.44 | 1.05 | 0.372 | 0.783 |
| 3128 | KEGG_AXON_GUIDANCE | 116 | 0.46 | 1.05 | 0.395 | 0.783 |
| 3129 | MULLIGHAN_NPM1_SIGNATURE_3_UP | 274 | 0.44 | 1.05 | 0.369 | 0.783 |
| 3130 | STAT3_02 | 128 | 0.46 | 1.05 | 0.394 | 0.784 |
| 3131 | CAGCACT_MIR5123P | 140 | 0.45 | 1.05 | 0.395 | 0.784 |
| 3132 | PID_P75_NTR_PATHWAY | 61 | 0.49 | 1.05 | 0.403 | 0.786 |

| | | | | | | |
|---|---|---|---|---|---|---|
| 3133 | CHEN_LIVER_METABOLISM_QTL_CIS | 74 | 0.47 | 1.05 | 0.392 | 0.787 |
| 3134 | TTANWNANTGGM_UNKNOWN | 45 | 0.51 | 1.05 | 0.427 | 0.788 |
| 3135 | CCAWYNNGAAR_UNKNOWN | 126 | 0.46 | 1.05 | 0.368 | 0.788 |
| 3136 | MIR10522_5P | 195 | 0.44 | 1.05 | 0.373 | 0.789 |
| 3137 | F10_TARGET_GENES | 203 | 0.44 | 1.05 | 0.388 | 0.788 |
| 3138 | REACTOME_CD28_CO_STIMULATION | 29 | 0.53 | 1.05 | 0.416 | 0.789 |
| 3139 | MIR942_5P | 184 | 0.44 | 1.05 | 0.38 | 0.789 |
| 3140 | ESC_J1_UP_LATE.V1_UP | 133 | 0.45 | 1.05 | 0.391 | 0.789 |
| 3141 | MTOR_UP.N4.V1_DN | 163 | 0.45 | 1.05 | 0.382 | 0.79 |
| 3142 | EBAUER_TARGETS_OF_PAX3_FOXO1_FUSION_DN | 39 | 0.5 | 1.05 | 0.409 | 0.791 |
| 3143 | RB_P107_DN.V1_UP | 114 | 0.45 | 1.05 | 0.381 | 0.791 |
| 3144 | REACTOME_RHOG_GTPASE_CYCLE | 69 | 0.48 | 1.05 | 0.402 | 0.791 |
| 3145 | REACTOME_TRANSCRIPTION_OF_E2F_TARGETS_ | 19 | 0.57 | 1.05 | 0.425 | 0.791 |
| 3146 | MIR4291 | 328 | 0.43 | 1.05 | 0.363 | 0.791 |
| 3147 | TST1_01 | 205 | 0.44 | 1.05 | 0.377 | 0.791 |
| 3148 | ER_Q6 | 222 | 0.44 | 1.05 | 0.383 | 0.792 |
| 3149 | MIR23A_5P | 23 | 0.56 | 1.05 | 0.433 | 0.792 |
| 3150 | MIR567 | 89 | 0.46 | 1.05 | 0.39 | 0.793 |
| 3151 | REACTOME_MAPK6_MAPK4_SIGNALING | 81 | 0.47 | 1.05 | 0.385 | 0.792 |
| 3152 | REACTOME_DISEASES_ASSOCIATED_WITH_O_GL | 48 | 0.5 | 1.05 | 0.411 | 0.793 |
| 3153 | ACEVEDO_LIVER_TUMOR_VS_NORMAL_ADJACEN | 197 | 0.44 | 1.05 | 0.371 | 0.794 |
| 3154 | MIR584_5P | 65 | 0.48 | 1.05 | 0.397 | 0.794 |
| 3155 | BOYAULT_LIVER_CANCER_SUBCLASS_G123_DN | 35 | 0.52 | 1.05 | 0.426 | 0.794 |
| 3156 | BRCA1_DN.V1_DN | 95 | 0.47 | 1.05 | 0.378 | 0.794 |
| 3157 | BYSTRYKH_HEMATOPOIESIS_STEM_CELL_AND_B | 64 | 0.48 | 1.05 | 0.401 | 0.795 |
| 3158 | OSMAN_BLADDER_CANCER_DN | 394 | 0.43 | 1.05 | 0.361 | 0.795 |
| 3159 | CAGCAGG_MIR370 | 136 | 0.45 | 1.05 | 0.392 | 0.795 |
| 3160 | DROSHA_TARGET_GENES | 60 | 0.49 | 1.05 | 0.399 | 0.795 |
| 3161 | TRIP13_TARGET_GENES | 74 | 0.48 | 1.05 | 0.407 | 0.795 |
| 3162 | TTTGTAG_MIR520D | 288 | 0.43 | 1.05 | 0.375 | 0.795 |
| 3163 | MIR664B_5P | 46 | 0.51 | 1.05 | 0.408 | 0.795 |
| 3164 | MIR6889_3P | 96 | 0.46 | 1.05 | 0.37 | 0.796 |
| 3165 | YANAGIHARA_ESX1_TARGETS | 23 | 0.55 | 1.05 | 0.43 | 0.797 |

| | | | | | | |
|---|---|---|---|---|---|---|
| 3166 | PID_P38_MK2_PATHWAY | 20 | 0.57 | 1.04 | 0.435 | 0.797 |
| 3167 | RASHI_RESPONSE_TO_IONIZING_RADIATION_4 | 39 | 0.52 | 1.04 | 0.413 | 0.797 |
| 3168 | KIM_WT1_TARGETS_12HR_UP | 135 | 0.45 | 1.04 | 0.392 | 0.798 |
| 3169 | WANG_MLL_TARGETS | 177 | 0.44 | 1.04 | 0.404 | 0.798 |
| 3170 | WP_GPCRS_OTHER | 30 | 0.53 | 1.04 | 0.419 | 0.799 |
| 3171 | WP_GLYCEROPHOSPHOLIPID_BIOSYNTHETIC_PA | 27 | 0.54 | 1.04 | 0.428 | 0.799 |
| 3172 | MEISSNER_NPC_HCP_WITH_H3_UNMETHYLATED | 359 | 0.43 | 1.04 | 0.361 | 0.799 |
| 3173 | AKT_UP_MTOR_DN.V1_DN | 143 | 0.44 | 1.04 | 0.397 | 0.799 |
| 3174 | WANG_TUMOR_INVASIVENESS_DN | 198 | 0.44 | 1.04 | 0.389 | 0.799 |
| 3175 | MIR5585_5P | 101 | 0.46 | 1.04 | 0.407 | 0.8 |
| 3176 | WP_ONECARBON_METABOLISM_AND_RELATED_F | 41 | 0.51 | 1.04 | 0.417 | 0.8 |
| 3177 | MIR2116_3P | 57 | 0.49 | 1.04 | 0.417 | 0.801 |
| 3178 | GATA1_02 | 184 | 0.44 | 1.04 | 0.394 | 0.801 |
| 3179 | HEIDENBLAD_AMPLICON_12P11_12_UP | 33 | 0.52 | 1.04 | 0.411 | 0.801 |
| 3180 | REACTOME_METABOLISM_OF_STEROID_HORMON | 16 | 0.59 | 1.04 | 0.458 | 0.801 |
| 3181 | OCT1_B | 192 | 0.44 | 1.04 | 0.377 | 0.801 |
| 3182 | CTR9_TARGET_GENES | 56 | 0.49 | 1.04 | 0.432 | 0.802 |
| 3183 | WP_ALZHEIMERS_DISEASE | 227 | 0.44 | 1.04 | 0.38 | 0.803 |
| 3184 | RIZ_ERYTHROID_DIFFERENTIATION_APOBEC2 | 17 | 0.58 | 1.04 | 0.451 | 0.802 |
| 3185 | CORDENONSI_YAP_CONSERVED_SIGNATURE | 43 | 0.5 | 1.04 | 0.41 | 0.802 |
| 3186 | REACTOME_METABOLISM_OF_POLYAMINES | 56 | 0.49 | 1.04 | 0.397 | 0.802 |
| 3187 | BIOCARTA_TID_PATHWAY | 17 | 0.59 | 1.04 | 0.466 | 0.803 |
| 3188 | CHEOK_RESPONSE_TO_MERCAPTOPURINE_AND_ | 22 | 0.55 | 1.04 | 0.436 | 0.803 |
| 3189 | LEE_DIFFERENTIATING_T_LYMPHOCYTE | 138 | 0.45 | 1.04 | 0.393 | 0.803 |
| 3190 | KESHELAVA_MULTIPLE_DRUG_RESISTANCE | 72 | 0.47 | 1.04 | 0.406 | 0.803 |
| 3191 | HEDENFALK_BREAST_CANCER_BRACX_UP | 17 | 0.58 | 1.04 | 0.447 | 0.804 |
| 3192 | MIR6509_3P | 116 | 0.45 | 1.04 | 0.412 | 0.804 |
| 3193 | ZHENG_IL22_SIGNALING_DN | 23 | 0.55 | 1.04 | 0.454 | 0.804 |
| 3194 | BIOCARTA_GPCR_PATHWAY | 28 | 0.53 | 1.04 | 0.419 | 0.804 |
| 3195 | NKX2_3_TARGET_GENES | 469 | 0.43 | 1.04 | 0.375 | 0.804 |
| 3196 | MIR3117_3P | 30 | 0.53 | 1.04 | 0.444 | 0.804 |
| 3197 | WOTTON_RUNX_TARGETS_DN | 25 | 0.55 | 1.04 | 0.442 | 0.805 |
| 3198 | KYNG_ENVIRONMENTAL_STRESS_RESPONSE_NC | 29 | 0.53 | 1.04 | 0.434 | 0.805 |

| | | | | | | |
|---|---|---|---|---|---|---|
| 3199 | WAGNER_APO2_SENSITIVITY | 21 | 0.55 | 1.04 | 0.438 | 0.804 |
| 3200 | WP_AMYOTROPHIC_LATERAL_SCLEROSIS_ALS | 36 | 0.51 | 1.04 | 0.421 | 0.805 |
| 3201 | MIR3169 | 51 | 0.49 | 1.04 | 0.411 | 0.805 |
| 3202 | NIKOLSKY_BREAST_CANCER_7Q21_Q22_AMPLIC | 54 | 0.49 | 1.04 | 0.408 | 0.805 |
| 3203 | NFY_Q6_01 | 229 | 0.43 | 1.04 | 0.411 | 0.806 |
| 3204 | WANG_PROSTATE_CANCER_ANDROGEN_INDEPEI | 55 | 0.48 | 1.04 | 0.429 | 0.806 |
| 3205 | IKEDA_MIR30_TARGETS_UP | 114 | 0.45 | 1.04 | 0.406 | 0.806 |
| 3206 | REACTOME_PROTEIN_UBIQUITINATION | 72 | 0.47 | 1.04 | 0.409 | 0.807 |
| 3207 | PID_NETRIN_PATHWAY | 31 | 0.53 | 1.04 | 0.444 | 0.807 |
| 3208 | REACTOME_SWITCHING_OF_ORIGINS_TO_A_POS | 89 | 0.46 | 1.04 | 0.394 | 0.808 |
| 3209 | KIM_MYCL1_AMPLIFICATION_TARGETS_DN | 18 | 0.57 | 1.04 | 0.44 | 0.808 |
| 3210 | TCGA_GLIOBLASTOMA_COPY_NUMBER_DN | 24 | 0.54 | 1.04 | 0.441 | 0.808 |
| 3211 | WP_FLUOROPYRIMIDINE_ACTIVITY | 25 | 0.54 | 1.04 | 0.43 | 0.808 |
| 3212 | BOYAULT_LIVER_CANCER_SUBCLASS_G3_DN | 32 | 0.53 | 1.04 | 0.437 | 0.808 |
| 3213 | GAL_LEUKEMIC_STEM_CELL_UP | 99 | 0.46 | 1.04 | 0.427 | 0.808 |
| 3214 | MAGRANGEAS_MULTIPLE_MYELOMA_IGG_VS_IG/ | 17 | 0.58 | 1.04 | 0.446 | 0.808 |
| 3215 | MIR10393_3P | 95 | 0.46 | 1.04 | 0.399 | 0.808 |
| 3216 | MIR4666B | 105 | 0.45 | 1.04 | 0.41 | 0.808 |
| 3217 | WP_INTEGRINMEDIATED_CELL_ADHESION | 85 | 0.46 | 1.04 | 0.411 | 0.809 |
| 3218 | SHI_SPARC_TARGETS_UP | 19 | 0.57 | 1.04 | 0.471 | 0.808 |
| 3219 | WP_MYOMETRIAL_RELAXATION_AND_CONTRACTI | 119 | 0.45 | 1.04 | 0.404 | 0.808 |
| 3220 | TURASHVILI_BREAST_LOBULAR_CARCINOMA_VS_ | 63 | 0.47 | 1.04 | 0.426 | 0.808 |
| 3221 | REACTOME_RAP1_SIGNALLING | 15 | 0.58 | 1.04 | 0.48 | 0.808 |
| 3222 | REACTOME_MET_ACTIVATES_PTK2_SIGNALING | 24 | 0.54 | 1.04 | 0.445 | 0.809 |
| 3223 | TATA_01 | 157 | 0.44 | 1.04 | 0.396 | 0.809 |
| 3224 | TFCP2_TARGET_GENES | 61 | 0.48 | 1.04 | 0.425 | 0.809 |
| 3225 | BIOCARTA_TNFR1_PATHWAY | 28 | 0.53 | 1.04 | 0.433 | 0.81 |
| 3226 | MIR4742_5P | 152 | 0.44 | 1.04 | 0.409 | 0.81 |
| 3227 | IGARASHI_ATF4_TARGETS_DN | 86 | 0.46 | 1.04 | 0.412 | 0.81 |
| 3228 | MIR4755_5P | 313 | 0.42 | 1.04 | 0.389 | 0.81 |
| 3229 | MIR1258 | 50 | 0.48 | 1.04 | 0.406 | 0.81 |
| 3230 | MIR34A_3P | 61 | 0.48 | 1.04 | 0.422 | 0.809 |
| 3231 | MIR4301 | 33 | 0.51 | 1.04 | 0.425 | 0.809 |

| | | | | | | |
|---|---|---|---|---|---|---|
| 3232 | MIR346 | 55 | 0.48 | 1.04 | 0.433 | 0.809 |
| 3233 | MIR4695_3P | 59 | 0.48 | 1.04 | 0.418 | 0.81 |
| 3234 | JIANG_AGING_CEREBRAL_CORTEX_DN | 47 | 0.5 | 1.04 | 0.436 | 0.81 |
| 3235 | NGFIC_01 | 231 | 0.43 | 1.04 | 0.394 | 0.81 |
| 3236 | SCHRAETS_MLL_TARGETS_DN | 16 | 0.59 | 1.04 | 0.47 | 0.81 |
| 3237 | SAMOLS_TARGETS_OF_KHSV_MIRNAS_DN | 46 | 0.5 | 1.04 | 0.434 | 0.809 |
| 3238 | VECCHI_GASTRIC_CANCER_EARLY_DN | 197 | 0.44 | 1.04 | 0.407 | 0.809 |
| 3239 | ZAMORA_NOS2_TARGETS_DN | 63 | 0.47 | 1.04 | 0.417 | 0.809 |
| 3240 | MIR4777_5P | 85 | 0.46 | 1.04 | 0.417 | 0.809 |
| 3241 | CDP_01 | 61 | 0.48 | 1.04 | 0.411 | 0.809 |
| 3242 | MIR24_1_5P_MIR24_2_5P | 42 | 0.5 | 1.04 | 0.414 | 0.809 |
| 3243 | SMID_BREAST_CANCER_BASAL_DN | 495 | 0.43 | 1.04 | 0.381 | 0.809 |
| 3244 | CEBP_Q2 | 185 | 0.44 | 1.04 | 0.402 | 0.81 |
| 3245 | MIR1228_3P | 66 | 0.46 | 1.04 | 0.414 | 0.81 |
| 3246 | ZSCAN5B_TARGET_GENES | 56 | 0.49 | 1.04 | 0.418 | 0.81 |
| 3247 | MIR4728_5P | 482 | 0.42 | 1.04 | 0.373 | 0.81 |
| 3248 | MIR1263 | 48 | 0.49 | 1.04 | 0.413 | 0.809 |
| 3249 | LHX3_01 | 160 | 0.44 | 1.04 | 0.406 | 0.809 |
| 3250 | BOYLAN_MULTIPLE_MYELOMA_C_UP | 44 | 0.5 | 1.04 | 0.42 | 0.809 |
| 3251 | KANNAN_TP53_TARGETS_UP | 51 | 0.48 | 1.04 | 0.408 | 0.81 |
| 3252 | SOX10_TARGET_GENES | 186 | 0.43 | 1.04 | 0.409 | 0.809 |
| 3253 | WONG_ENDMETRIUM_CANCER_DN | 39 | 0.5 | 1.04 | 0.421 | 0.81 |
| 3254 | PRC1_BMI_UP.V1_DN | 119 | 0.45 | 1.04 | 0.412 | 0.81 |
| 3255 | BRACHAT_RESPONSE_TO_METHOTREXATE_UP | 21 | 0.54 | 1.04 | 0.437 | 0.81 |
| 3256 | KRASNOSELSKAYA_ILF3_TARGETS_DN | 37 | 0.51 | 1.04 | 0.431 | 0.812 |
| 3257 | MIR6810_3P | 66 | 0.47 | 1.04 | 0.434 | 0.812 |
| 3258 | LU_EZH2_TARGETS_DN | 346 | 0.42 | 1.03 | 0.386 | 0.813 |
| 3259 | MIR4652_5P | 31 | 0.52 | 1.03 | 0.448 | 0.813 |
| 3260 | FRASOR_RESPONSE_TO_ESTRADIOL_UP | 23 | 0.55 | 1.03 | 0.456 | 0.813 |
| 3261 | AAGGGAT_MIR188 | 68 | 0.47 | 1.03 | 0.413 | 0.814 |
| 3262 | REACTOME_KERATAN_SULFATE_KERATIN_METAE | 24 | 0.54 | 1.03 | 0.443 | 0.813 |
| 3263 | WP_KIT_RECEPTOR_SIGNALING_PATHWAY | 52 | 0.49 | 1.03 | 0.406 | 0.814 |
| 3264 | MORI_SMALL_PRE_BII_LYMPHOCYTE_UP | 62 | 0.48 | 1.03 | 0.421 | 0.815 |

| | | | | | | |
|---|---|---|---|---|---|---|
| 3265 | PID_HIF1_TFPATHWAY | 55 | 0.48 | 1.03 | 0.457 | 0.816 |
| 3266 | MIR6733_5P | 237 | 0.43 | 1.03 | 0.395 | 0.816 |
| 3267 | SANSOM_APC_TARGETS_UP | 112 | 0.45 | 1.03 | 0.427 | 0.817 |
| 3268 | MIR664A_3P | 330 | 0.43 | 1.03 | 0.413 | 0.817 |
| 3269 | AGCGCTT_MIR518F_MIR518E_MIR518A | 15 | 0.58 | 1.03 | 0.465 | 0.817 |
| 3270 | REACTOME_CYTOSOLIC_TRNA_AMINOACYLATION | 23 | 0.55 | 1.03 | 0.443 | 0.817 |
| 3271 | MIR4660 | 118 | 0.45 | 1.03 | 0.424 | 0.817 |
| 3272 | MIR5187_5P | 83 | 0.46 | 1.03 | 0.414 | 0.818 |
| 3273 | KRAS.50_UP.V1_UP | 33 | 0.52 | 1.03 | 0.438 | 0.818 |
| 3274 | SESTO_RESPONSE_TO_UV_C2 | 51 | 0.49 | 1.03 | 0.424 | 0.819 |
| 3275 | BLALOCK_ALZHEIMERS_DISEASE_INCIPIENT_DN | 149 | 0.44 | 1.03 | 0.43 | 0.819 |
| 3276 | REACTOME_SIGNALING_BY_MET | 70 | 0.47 | 1.03 | 0.433 | 0.819 |
| 3277 | MIR205_3P | 343 | 0.43 | 1.03 | 0.403 | 0.82 |
| 3278 | PTF1BETA_Q6 | 202 | 0.43 | 1.03 | 0.408 | 0.821 |
| 3279 | ZIC2_01 | 218 | 0.43 | 1.03 | 0.427 | 0.821 |
| 3280 | BROWNE_HCMV_INFECTION_10HR_UP | 84 | 0.46 | 1.03 | 0.444 | 0.821 |
| 3281 | AP4_Q6_01 | 210 | 0.43 | 1.03 | 0.409 | 0.821 |
| 3282 | CHANG_IMMORTALIZED_BY_HPV31_UP | 52 | 0.49 | 1.03 | 0.435 | 0.821 |
| 3283 | BCAT_BILD_ET_AL_UP | 38 | 0.51 | 1.03 | 0.437 | 0.822 |
| 3284 | BIOCARTA_PROTEASOME_PATHWAY | 19 | 0.57 | 1.03 | 0.488 | 0.822 |
| 3285 | MIR539_5P | 321 | 0.42 | 1.03 | 0.4 | 0.822 |
| 3286 | MIR934 | 60 | 0.48 | 1.03 | 0.434 | 0.822 |
| 3287 | SPZ1_01 | 199 | 0.43 | 1.03 | 0.435 | 0.822 |
| 3288 | MIR1236_3P | 159 | 0.44 | 1.03 | 0.438 | 0.822 |
| 3289 | JI_METASTASIS_REPRESSED_BY_STK11 | 16 | 0.57 | 1.03 | 0.473 | 0.822 |
| 3290 | BIOCARTA_WNT_PATHWAY | 22 | 0.55 | 1.03 | 0.473 | 0.823 |
| 3291 | MIR3617_3P | 74 | 0.46 | 1.03 | 0.428 | 0.823 |
| 3292 | WP_CANONICAL_AND_NONCANONICAL_NOTCH_SI | 25 | 0.54 | 1.03 | 0.463 | 0.823 |
| 3293 | VERHAAK_GLIOBLASTOMA_NEURAL | 99 | 0.45 | 1.03 | 0.427 | 0.824 |
| 3294 | ZNF354B_TARGET_GENES | 179 | 0.43 | 1.03 | 0.438 | 0.825 |
| 3295 | MIR7977 | 197 | 0.43 | 1.03 | 0.45 | 0.825 |
| 3296 | REACTOME_CELL_CELL_JUNCTION_ORGANIZATIC | 46 | 0.49 | 1.03 | 0.432 | 0.825 |
| 3297 | MIR6739_5P | 238 | 0.43 | 1.03 | 0.427 | 0.825 |

| | | | | | | |
|---|---|---|---|---|---|---|
| 3298 | MIR143_5P | 117 | 0.45 | 1.03 | 0.445 | 0.825 |
| 3299 | BROWNE_HCMV_INFECTION_16HR_UP | 205 | 0.43 | 1.03 | 0.425 | 0.825 |
| 3300 | KEGG_ARRHYTHMOGENIC_RIGHT_VENTRICULAR_ | 56 | 0.48 | 1.03 | 0.416 | 0.826 |
| 3301 | KASLER_HDAC7_TARGETS_2_DN | 31 | 0.52 | 1.03 | 0.461 | 0.826 |
| 3302 | CREB_Q3 | 227 | 0.43 | 1.03 | 0.426 | 0.826 |
| 3303 | E12_Q6 | 214 | 0.43 | 1.03 | 0.422 | 0.826 |
| 3304 | RRCCGTTA_UNKNOWN | 80 | 0.46 | 1.03 | 0.434 | 0.827 |
| 3305 | HOFFMANN_IMMATURE_TO_MATURE_B_LYMPHOC | 31 | 0.52 | 1.03 | 0.432 | 0.827 |
| 3306 | MSX1_TARGET_GENES | 168 | 0.44 | 1.03 | 0.426 | 0.828 |
| 3307 | MIR642B_5P | 68 | 0.47 | 1.03 | 0.442 | 0.83 |
| 3308 | NIKOLSKY_BREAST_CANCER_17Q11_Q21_AMPLI( | 96 | 0.45 | 1.03 | 0.414 | 0.83 |
| 3309 | REACTOME_G1_S_DNA_DAMAGE_CHECKPOINTS | 66 | 0.47 | 1.03 | 0.443 | 0.83 |
| 3310 | REACTOME_ANTI_INFLAMMATORY_RESPONSE_F/ | 76 | 0.46 | 1.03 | 0.43 | 0.83 |
| 3311 | MIR7160_5P | 163 | 0.44 | 1.03 | 0.435 | 0.83 |
| 3312 | MORI_PRE_BI_LYMPHOCYTE_UP | 69 | 0.47 | 1.03 | 0.445 | 0.83 |
| 3313 | NELSON_RESPONSE_TO_ANDROGEN_UP | 73 | 0.47 | 1.03 | 0.447 | 0.83 |
| 3314 | MIR4726_3P | 63 | 0.47 | 1.03 | 0.435 | 0.829 |
| 3315 | MEISSNER_BRAIN_HCP_WITH_H3K4ME2_AND_H3I | 36 | 0.51 | 1.03 | 0.45 | 0.829 |
| 3316 | MIR4518 | 81 | 0.46 | 1.03 | 0.445 | 0.829 |
| 3317 | SCIAN_CELL_CYCLE_TARGETS_OF_TP53_AND_TI | 21 | 0.55 | 1.03 | 0.456 | 0.829 |
| 3318 | MIR499B_5P | 87 | 0.46 | 1.03 | 0.444 | 0.829 |
| 3319 | SASSON_RESPONSE_TO_FORSKOLIN_DN | 77 | 0.47 | 1.03 | 0.446 | 0.829 |
| 3320 | CHAUHAN_RESPONSE_TO_METHOXYESTRADIOL_ | 45 | 0.49 | 1.03 | 0.438 | 0.829 |
| 3321 | MIR555 | 40 | 0.5 | 1.03 | 0.454 | 0.829 |
| 3322 | MIR6529_3P | 75 | 0.46 | 1.03 | 0.431 | 0.829 |
| 3323 | BIOCARTA_CTCF_PATHWAY | 21 | 0.55 | 1.03 | 0.44 | 0.83 |
| 3324 | CHANDRAN_METASTASIS_UP | 182 | 0.43 | 1.03 | 0.43 | 0.83 |
| 3325 | VERRECCHIA_EARLY_RESPONSE_TO_TGFB1 | 42 | 0.5 | 1.03 | 0.444 | 0.83 |
| 3326 | MIR3120_5P | 71 | 0.47 | 1.03 | 0.456 | 0.83 |
| 3327 | MIR4311 | 300 | 0.42 | 1.03 | 0.446 | 0.83 |
| 3328 | MCCLUNG_COCAIN_REWARD_4WK | 57 | 0.48 | 1.02 | 0.437 | 0.831 |
| 3329 | INGRAM_SHH_TARGETS_UP | 87 | 0.46 | 1.02 | 0.429 | 0.831 |
| 3330 | MIR4633_5P | 43 | 0.5 | 1.02 | 0.448 | 0.831 |

| | | | | | | |
|---|---|---|---|---|---|---|
| 3331 | WP_FATTY_ACID_BETAOXIDATION | 30 | 0.52 | 1.02 | 0.436 | 0.831 |
| 3332 | CLASPER_LYMPHATIC_VESSELS_DURING_METAS | 15 | 0.58 | 1.02 | 0.478 | 0.831 |
| 3333 | MIR511_5P | 254 | 0.43 | 1.02 | 0.425 | 0.831 |
| 3334 | VANLOO_SP3_TARGETS_DN | 66 | 0.47 | 1.02 | 0.454 | 0.831 |
| 3335 | MIR4525 | 30 | 0.52 | 1.02 | 0.457 | 0.831 |
| 3336 | JIANG_AGING_CEREBRAL_CORTEX_UP | 27 | 0.53 | 1.02 | 0.448 | 0.831 |
| 3337 | PID_PS1_PATHWAY | 42 | 0.5 | 1.02 | 0.441 | 0.832 |
| 3338 | MIR7152_3P | 85 | 0.46 | 1.02 | 0.445 | 0.832 |
| 3339 | HUANG_GATA2_TARGETS_DN | 67 | 0.47 | 1.02 | 0.438 | 0.832 |
| 3340 | MIR4802_3P | 147 | 0.44 | 1.02 | 0.446 | 0.832 |
| 3341 | NAKAMURA_CANCER_MICROENVIRONMENT_DN | 44 | 0.5 | 1.02 | 0.455 | 0.832 |
| 3342 | MIR6502_3P | 52 | 0.49 | 1.02 | 0.473 | 0.833 |
| 3343 | GCAAGAC_MIR431 | 37 | 0.5 | 1.02 | 0.429 | 0.833 |
| 3344 | MIR3611 | 131 | 0.44 | 1.02 | 0.429 | 0.834 |
| 3345 | MIR4319 | 377 | 0.42 | 1.02 | 0.432 | 0.835 |
| 3346 | MIR6728_3P | 122 | 0.44 | 1.02 | 0.424 | 0.835 |
| 3347 | ESC_V6.5_UP_LATE.V1_UP | 135 | 0.44 | 1.02 | 0.455 | 0.835 |
| 3348 | NF1_Q6_01 | 217 | 0.43 | 1.02 | 0.436 | 0.836 |
| 3349 | AACYNNNNTTCCS_UNKNOWN | 95 | 0.46 | 1.02 | 0.444 | 0.836 |
| 3350 | MIR4462 | 15 | 0.57 | 1.02 | 0.493 | 0.836 |
| 3351 | VISALA_AGING_LYMPHOCYTE_DN | 16 | 0.57 | 1.02 | 0.479 | 0.836 |
| 3352 | REACTOME_SIGNALING_BY_NOTCH3 | 44 | 0.49 | 1.02 | 0.454 | 0.836 |
| 3353 | MIR1976 | 69 | 0.46 | 1.02 | 0.46 | 0.837 |
| 3354 | AMIT_DELAYED_EARLY_GENES | 17 | 0.57 | 1.02 | 0.463 | 0.837 |
| 3355 | ZWANG_EGF_INTERVAL_UP | 64 | 0.46 | 1.02 | 0.442 | 0.838 |
| 3356 | MIR654_5P | 81 | 0.45 | 1.02 | 0.453 | 0.839 |
| 3357 | MIR3180_5P | 260 | 0.42 | 1.02 | 0.451 | 0.841 |
| 3358 | GTGCAAA_MIR507 | 113 | 0.45 | 1.02 | 0.442 | 0.841 |
| 3359 | REACTOME_SIGNALING_BY_NOTCH1_PEST_DOM/ | 56 | 0.48 | 1.02 | 0.471 | 0.841 |
| 3360 | SHETH_LIVER_CANCER_VS_TXNIP_LOSS_PAM2 | 138 | 0.44 | 1.02 | 0.447 | 0.841 |
| 3361 | KONDO_PROSTATE_CANCER_WITH_H3K27ME3 | 52 | 0.47 | 1.02 | 0.446 | 0.841 |
| 3362 | LANDEMAINE_LUNG_METASTASIS | 18 | 0.56 | 1.02 | 0.464 | 0.842 |
| 3363 | E4F1_Q6 | 256 | 0.43 | 1.02 | 0.452 | 0.842 |

| | | | | | | |
|---|---|---|---|---|---|---|
| 3364 | MIR4646_5P | 127 | 0.44 | 1.02 | 0.44 | 0.843 |
| 3365 | WP_ARRHYTHMOGENIC_RIGHT_VENTRICULAR_CA | 56 | 0.48 | 1.02 | 0.475 | 0.843 |
| 3366 | MIR12120 | 160 | 0.43 | 1.02 | 0.47 | 0.843 |
| 3367 | MIR888_3P | 104 | 0.44 | 1.02 | 0.452 | 0.844 |
| 3368 | WTTGKCTG_UNKNOWN | 426 | 0.42 | 1.02 | 0.451 | 0.844 |
| 3369 | WP_ALZHEIMERS_DISEASE_AND_MIRNA_EFFECTS | 235 | 0.42 | 1.02 | 0.443 | 0.844 |
| 3370 | NFAT_Q4_01 | 211 | 0.43 | 1.02 | 0.446 | 0.844 |
| 3371 | CGTSACG_PAX3_B | 136 | 0.44 | 1.02 | 0.419 | 0.844 |
| 3372 | CREB_Q2 | 236 | 0.43 | 1.02 | 0.459 | 0.844 |
| 3373 | MIR582_5P | 318 | 0.42 | 1.02 | 0.438 | 0.844 |
| 3374 | KEGG_RENAL_CELL_CARCINOMA | 67 | 0.47 | 1.02 | 0.466 | 0.844 |
| 3375 | MIR6853_3P | 175 | 0.43 | 1.02 | 0.445 | 0.844 |
| 3376 | AAAYWAACM_HFH4_01 | 201 | 0.43 | 1.02 | 0.455 | 0.844 |
| 3377 | MIR3688_3P | 343 | 0.42 | 1.02 | 0.437 | 0.845 |
| 3378 | BURTON_ADIPOGENESIS_1 | 30 | 0.51 | 1.02 | 0.464 | 0.845 |
| 3379 | WP_SYNAPTIC_SIGNALING_PATHWAYS_ASSOCIA | 44 | 0.49 | 1.02 | 0.468 | 0.845 |
| 3380 | MIR767_5P | 164 | 0.43 | 1.02 | 0.456 | 0.845 |
| 3381 | MIR221_5P | 67 | 0.47 | 1.02 | 0.452 | 0.845 |
| 3382 | MIR4281 | 17 | 0.56 | 1.02 | 0.486 | 0.845 |
| 3383 | MIKKELSEN_IPS_HCP_WITH_H3_UNMETHYLATED | 42 | 0.5 | 1.02 | 0.463 | 0.845 |
| 3384 | CYTAGCAAY_UNKNOWN | 127 | 0.44 | 1.02 | 0.454 | 0.845 |
| 3385 | ZNF33A_TARGET_GENES | 186 | 0.43 | 1.02 | 0.451 | 0.845 |
| 3386 | MARTORIATI_MDM4_TARGETS_NEUROEPITHELIUM | 127 | 0.44 | 1.02 | 0.429 | 0.845 |
| 3387 | HFH4_01 | 159 | 0.43 | 1.02 | 0.469 | 0.845 |
| 3388 | GENTILE_RESPONSE_CLUSTER_D3 | 51 | 0.48 | 1.02 | 0.453 | 0.845 |
| 3389 | SHIPP_DLBCL_VS_FOLLICULAR_LYMPHOMA_UP | 44 | 0.5 | 1.02 | 0.462 | 0.845 |
| 3390 | WP_G_PROTEIN_SIGNALING_PATHWAYS | 74 | 0.46 | 1.02 | 0.447 | 0.845 |
| 3391 | CACCCBINDINGFACTOR_Q6 | 229 | 0.43 | 1.02 | 0.446 | 0.845 |
| 3392 | MIR6126 | 99 | 0.45 | 1.02 | 0.449 | 0.845 |
| 3393 | MIR152_3P | 309 | 0.42 | 1.02 | 0.458 | 0.846 |
| 3394 | WP_INTERLEUKIN11_SIGNALING_PATHWAY | 42 | 0.49 | 1.02 | 0.458 | 0.845 |
| 3395 | BIOCARTA_NO1_PATHWAY | 23 | 0.55 | 1.02 | 0.485 | 0.846 |
| 3396 | KEGG_HUNTINGTONS_DISEASE | 163 | 0.43 | 1.02 | 0.46 | 0.846 |

| | | | | | | |
|---|---|---|---|---|---|---|
| 3397 | NKX62_Q2 | 185 | 0.43 | 1.02 | 0.459 | 0.846 |
| 3398 | KEGG_ALANINE_ASPARTATE_AND_GLUTAMATE_I | 27 | 0.53 | 1.02 | 0.454 | 0.846 |
| 3399 | PID_PRL_SIGNALING_EVENTS_PATHWAY | 21 | 0.55 | 1.02 | 0.472 | 0.845 |
| 3400 | MIR9851_3P | 162 | 0.43 | 1.02 | 0.474 | 0.845 |
| 3401 | WAKABAYASHI_ADIPOGENESIS_PPARG_RXRA_B( | 137 | 0.44 | 1.02 | 0.464 | 0.846 |
| 3402 | RADMACHER_AML_PROGNOSIS | 64 | 0.47 | 1.02 | 0.461 | 0.846 |
| 3403 | HFH3_01 | 154 | 0.43 | 1.02 | 0.462 | 0.846 |
| 3404 | BLANCO_MELO_MERS_COV_INFECTION_MCR5_CE | 21 | 0.55 | 1.02 | 0.471 | 0.846 |
| 3405 | YAUCH_HEDGEHOG_SIGNALING_PARACRINE_DN | 141 | 0.43 | 1.02 | 0.479 | 0.846 |
| 3406 | HNF1_C | 159 | 0.43 | 1.02 | 0.473 | 0.847 |
| 3407 | YNTTTNNNANGCARM_UNKNOWN | 46 | 0.48 | 1.02 | 0.46 | 0.846 |
| 3408 | MIR148B_3P | 308 | 0.42 | 1.02 | 0.473 | 0.847 |
| 3409 | PID_FCER1_PATHWAY | 47 | 0.48 | 1.01 | 0.454 | 0.847 |
| 3410 | MIR148A_3P | 309 | 0.42 | 1.01 | 0.455 | 0.847 |
| 3411 | WP_RAS_SIGNALING | 147 | 0.43 | 1.01 | 0.457 | 0.847 |
| 3412 | EBNA1BP2_TARGET_GENES | 285 | 0.42 | 1.01 | 0.462 | 0.848 |
| 3413 | REACTOME_S_PHASE | 160 | 0.43 | 1.01 | 0.451 | 0.848 |
| 3414 | PAX8_TARGET_GENES | 76 | 0.46 | 1.01 | 0.469 | 0.849 |
| 3415 | YAO_TEMPORAL_RESPONSE_TO_PROGESTERON | 60 | 0.47 | 1.01 | 0.459 | 0.849 |
| 3416 | RAY_TUMORIGENESIS_BY_ERBB2_CDC25A_UP | 140 | 0.43 | 1.01 | 0.475 | 0.849 |
| 3417 | MIR199B_5P | 162 | 0.43 | 1.01 | 0.469 | 0.85 |
| 3418 | MIR660_3P | 78 | 0.45 | 1.01 | 0.462 | 0.85 |
| 3419 | ATGCTGG_MIR338 | 103 | 0.44 | 1.01 | 0.468 | 0.851 |
| 3420 | TOOKER_GEMCITABINE_RESISTANCE_DN | 114 | 0.44 | 1.01 | 0.468 | 0.851 |
| 3421 | DAZARD_RESPONSE_TO_UV_NHEK_DN | 287 | 0.42 | 1.01 | 0.446 | 0.852 |
| 3422 | DING_LUNG_CANCER_BY_MUTATION_RATE | 17 | 0.57 | 1.01 | 0.486 | 0.851 |
| 3423 | PAX4_04 | 168 | 0.43 | 1.01 | 0.451 | 0.852 |
| 3424 | PASINI_SUZ12_TARGETS_DN | 265 | 0.42 | 1.01 | 0.46 | 0.853 |
| 3425 | WONG_PROTEASOME_GENE_MODULE | 50 | 0.48 | 1.01 | 0.468 | 0.852 |
| 3426 | MIR4646_3P | 50 | 0.48 | 1.01 | 0.45 | 0.852 |
| 3427 | MIR296_3P | 28 | 0.52 | 1.01 | 0.458 | 0.852 |
| 3428 | GARGALOVIC_RESPONSE_TO_OXIDIZED_PHOSPI | 31 | 0.51 | 1.01 | 0.444 | 0.852 |
| 3429 | MIR1266_5P | 85 | 0.45 | 1.01 | 0.465 | 0.852 |

| | | | | | | |
|---|---|---|---|---|---|---|
| 3430 | MIR9851_5P | 99 | 0.45 | 1.01 | 0.467 | 0.852 |
| 3431 | MIR517_5P | 75 | 0.45 | 1.01 | 0.459 | 0.852 |
| 3432 | MIR3682_3P | 61 | 0.46 | 1.01 | 0.45 | 0.852 |
| 3433 | ELLWOOD_MYC_TARGETS_DN | 39 | 0.5 | 1.01 | 0.478 | 0.852 |
| 3434 | MIR545_3P | 330 | 0.42 | 1.01 | 0.471 | 0.852 |
| 3435 | MIR4522 | 44 | 0.49 | 1.01 | 0.454 | 0.852 |
| 3436 | H1_6_TARGET_GENES | 417 | 0.41 | 1.01 | 0.464 | 0.852 |
| 3437 | BURTON_ADIPOGENESIS_3 | 95 | 0.45 | 1.01 | 0.471 | 0.852 |
| 3438 | MIR3154 | 95 | 0.45 | 1.01 | 0.46 | 0.853 |
| 3439 | KRAS.AMP.LUNG_UP.V1_UP | 62 | 0.46 | 1.01 | 0.479 | 0.853 |
| 3440 | SEAVEY_EPITHELIOID_HEMANGIOENDOTHELIOMA | 50 | 0.48 | 1.01 | 0.454 | 0.853 |
| 3441 | WP_TYPE_I_COLLAGEN_SYNTHESIS_IN_THE_CON | 24 | 0.53 | 1.01 | 0.476 | 0.853 |
| 3442 | REACTOME_ACTIVATION_OF_GENE_EXPRESSION | 41 | 0.49 | 1.01 | 0.467 | 0.852 |
| 3443 | CTCTGGA_MIR520A_MIR525 | 142 | 0.43 | 1.01 | 0.471 | 0.853 |
| 3444 | GROSS_HYPOXIA_VIA_ELK3_UP | 190 | 0.43 | 1.01 | 0.468 | 0.853 |
| 3445 | MIR6758_5P | 89 | 0.45 | 1.01 | 0.46 | 0.854 |
| 3446 | WP_INSULIN_SIGNALING | 145 | 0.43 | 1.01 | 0.456 | 0.854 |
| 3447 | NAKAMURA_ADIPOGENESIS_EARLY_UP | 43 | 0.49 | 1.01 | 0.488 | 0.854 |
| 3448 | ZHAN_MULTIPLE_MYELOMA_MF_UP | 32 | 0.51 | 1.01 | 0.477 | 0.854 |
| 3449 | MIR329_5P | 123 | 0.44 | 1.01 | 0.466 | 0.854 |
| 3450 | MIR5683 | 211 | 0.42 | 1.01 | 0.464 | 0.854 |
| 3451 | MIR9718 | 89 | 0.45 | 1.01 | 0.468 | 0.855 |
| 3452 | MIR4294 | 60 | 0.47 | 1.01 | 0.47 | 0.855 |
| 3453 | MIR671_5P | 107 | 0.44 | 1.01 | 0.47 | 0.856 |
| 3454 | AFFAR_YY1_TARGETS_DN | 193 | 0.42 | 1.01 | 0.479 | 0.856 |
| 3455 | MEF2_02 | 164 | 0.43 | 1.01 | 0.489 | 0.856 |
| 3456 | CHIARADONNA_NEOPLASTIC_TRANSFORMATION_ | 45 | 0.48 | 1.01 | 0.473 | 0.856 |
| 3457 | SHIPP_DLBCL_CURED_VS_FATAL_DN | 42 | 0.49 | 1.01 | 0.48 | 0.856 |
| 3458 | NMYC_01 | 245 | 0.42 | 1.01 | 0.46 | 0.857 |
| 3459 | MIR541_3P | 80 | 0.46 | 1.01 | 0.469 | 0.857 |
| 3460 | MIR5010_5P | 31 | 0.51 | 1.01 | 0.487 | 0.858 |
| 3461 | WP_RAC1PAK1P38MMP2_PATHWAY | 60 | 0.46 | 1.01 | 0.467 | 0.857 |
| 3462 | REACTOME_SULFUR_AMINO_ACID_METABOLISM | 24 | 0.54 | 1.01 | 0.491 | 0.857 |

| | | | | | | |
|---|---|---|---|---|---|---|
| 3463 | MIR4799_5P | 343 | 0.41 | 1.01 | 0.497 | 0.857 |
| 3464 | PGF_UP.V1_UP | 166 | 0.43 | 1.01 | 0.464 | 0.857 |
| 3465 | KEGG_GLYCOSAMINOGLYCAN_BIOSYNTHESIS_CI | 20 | 0.55 | 1.01 | 0.515 | 0.857 |
| 3466 | REACTOME_SIGNALING_BY_ERBB2 | 47 | 0.48 | 1.01 | 0.489 | 0.857 |
| 3467 | IK2_01 | 209 | 0.42 | 1.01 | 0.465 | 0.858 |
| 3468 | BONCI_TARGETS_OF_MIR15A_AND_MIR16_1 | 77 | 0.46 | 1.01 | 0.469 | 0.858 |
| 3469 | WP_NANOPARTICLEMEDIATED_ACTIVATION_OF_F | 27 | 0.52 | 1.01 | 0.478 | 0.859 |
| 3470 | MIR4756_3P | 96 | 0.45 | 1.01 | 0.483 | 0.859 |
| 3471 | MIR6795_5P | 112 | 0.44 | 1.01 | 0.454 | 0.858 |
| 3472 | GFI1_01 | 197 | 0.42 | 1.01 | 0.47 | 0.858 |
| 3473 | REACTOME_CONSTITUTIVE_SIGNALING_BY_ABEF | 54 | 0.47 | 1.01 | 0.485 | 0.858 |
| 3474 | COREBINDINGFACTOR_Q6 | 189 | 0.42 | 1.01 | 0.493 | 0.858 |
| 3475 | MIR3922_3P | 83 | 0.45 | 1.01 | 0.479 | 0.858 |
| 3476 | MIR1207_3P | 114 | 0.44 | 1.01 | 0.48 | 0.858 |
| 3477 | BCAT.100_UP.V1_UP | 29 | 0.51 | 1.01 | 0.464 | 0.858 |
| 3478 | MIR216A_3P | 405 | 0.41 | 1.01 | 0.477 | 0.859 |
| 3479 | REACTOME_FGFR1_MUTANT_RECEPTOR_ACTIVAT | 24 | 0.53 | 1.01 | 0.475 | 0.859 |
| 3480 | HARRIS_HYPOXIA | 64 | 0.46 | 1.01 | 0.486 | 0.859 |
| 3481 | REACTOME_G_PROTEIN_MEDIATED_EVENTS | 45 | 0.48 | 1.01 | 0.465 | 0.858 |
| 3482 | MIR891B | 105 | 0.44 | 1.01 | 0.476 | 0.859 |
| 3483 | MIR1233_3P | 75 | 0.46 | 1.01 | 0.479 | 0.859 |
| 3484 | MCLACHLAN_DENTAL_CARIES_DN | 67 | 0.45 | 1.01 | 0.473 | 0.859 |
| 3485 | PPARA_TARGET_GENES | 59 | 0.47 | 1.01 | 0.468 | 0.861 |
| 3486 | TTGGGAG_MIR150 | 82 | 0.46 | 1.01 | 0.484 | 0.861 |
| 3487 | MIR505_5P | 45 | 0.48 | 1.01 | 0.473 | 0.861 |
| 3488 | RHODES_UNDIFFERENTIATED_CANCER | 67 | 0.46 | 1.01 | 0.445 | 0.861 |
| 3489 | KEGG_ENDOCYTOSIS | 158 | 0.43 | 1.01 | 0.471 | 0.861 |
| 3490 | DNMT1_TARGET_GENES | 151 | 0.43 | 1.01 | 0.488 | 0.861 |
| 3491 | FIRESTEIN_PROLIFERATION | 136 | 0.43 | 1.01 | 0.461 | 0.861 |
| 3492 | SILIGAN_BOUND_BY_EWS_FLT1_FUSION | 33 | 0.5 | 1.01 | 0.465 | 0.861 |
| 3493 | MIR11399 | 66 | 0.46 | 1 | 0.463 | 0.862 |
| 3494 | REACTOME_ONCOGENIC_MAPK_SIGNALING | 73 | 0.45 | 1 | 0.472 | 0.862 |
| 3495 | PID_ERB_GENOMIC_PATHWAY | 15 | 0.57 | 1 | 0.513 | 0.862 |

| | | | | | | |
|---|---|---|---|---|---|---|
| 3496 | REACTOME_INTERLEUKIN_17_SIGNALING | 65 | 0.46 | 1 | 0.47 | 0.862 |
| 3497 | MIR6827_3P | 67 | 0.46 | 1 | 0.49 | 0.862 |
| 3498 | WILLERT_WNT_SIGNALING | 17 | 0.56 | 1 | 0.499 | 0.862 |
| 3499 | PEREZ_TP53_AND_TP63_TARGETS | 180 | 0.42 | 1 | 0.49 | 0.863 |
| 3500 | REACTOME_APOPTOSIS | 164 | 0.43 | 1 | 0.473 | 0.863 |
| 3501 | MIR4425 | 77 | 0.46 | 1 | 0.477 | 0.863 |
| 3502 | BAE_BRCA1_TARGETS_UP | 65 | 0.45 | 1 | 0.478 | 0.863 |
| 3503 | PID_CD8_TCR_PATHWAY | 40 | 0.49 | 1 | 0.462 | 0.864 |
| 3504 | JAZAG_TGFB1_SIGNALING_VIA_SMAD4_DN | 50 | 0.48 | 1 | 0.483 | 0.865 |
| 3505 | MIR8057 | 78 | 0.45 | 1 | 0.472 | 0.865 |
| 3506 | REACTOME_ADORA2B_MEDIATED_ANTI_INFLAMM | 55 | 0.47 | 1 | 0.466 | 0.865 |
| 3507 | DELYS_THYROID_CANCER_DN | 156 | 0.43 | 1 | 0.486 | 0.865 |
| 3508 | BMI1_DN_MEL18_DN.V1_DN | 113 | 0.43 | 1 | 0.483 | 0.865 |
| 3509 | AREB6_02 | 211 | 0.42 | 1 | 0.484 | 0.865 |
| 3510 | LA_MEN1_TARGETS | 19 | 0.55 | 1 | 0.495 | 0.865 |
| 3511 | WP_KENNEDY_PATHWAY_FROM_SPHINGOLIPIDS | 15 | 0.57 | 1 | 0.526 | 0.865 |
| 3512 | ETS1_B | 224 | 0.42 | 1 | 0.495 | 0.865 |
| 3513 | MIR2355_3P | 91 | 0.45 | 1 | 0.484 | 0.865 |
| 3514 | GCNP_SHH_UP_LATE.V1_DN | 135 | 0.43 | 1 | 0.504 | 0.865 |
| 3515 | MIR199A_5P | 163 | 0.42 | 1 | 0.494 | 0.865 |
| 3516 | LCORL_TARGET_GENES | 355 | 0.41 | 1 | 0.489 | 0.865 |
| 3517 | MIR12135 | 161 | 0.43 | 1 | 0.487 | 0.865 |
| 3518 | MIR613 | 296 | 0.41 | 1 | 0.496 | 0.866 |
| 3519 | MIR6083 | 223 | 0.42 | 1 | 0.496 | 0.866 |
| 3520 | FOXO4_01 | 198 | 0.42 | 1 | 0.494 | 0.866 |
| 3521 | MIR769_5P | 39 | 0.49 | 1 | 0.492 | 0.866 |
| 3522 | NF1_Q6 | 219 | 0.42 | 1 | 0.485 | 0.866 |
| 3523 | WP_DIFFERENTIATION_OF_WHITE_AND_BROWN_[illegible] | 15 | 0.57 | 1 | 0.517 | 0.866 |
| 3524 | NKX25_01 | 94 | 0.44 | 1 | 0.49 | 0.867 |
| 3525 | MIR3183 | 66 | 0.46 | 1 | 0.492 | 0.867 |
| 3526 | MIR632 | 53 | 0.46 | 1 | 0.488 | 0.867 |
| 3527 | MIR6124 | 375 | 0.41 | 1 | 0.484 | 0.867 |
| 3528 | MIR182_3P | 57 | 0.47 | 1 | 0.492 | 0.867 |

| | | | | | | |
|---|---|---|---|---|---|---|
| 3529 | GTGCCAA_MIR96 | 273 | 0.42 | 1 | 0.494 | 0.867 |
| 3530 | MIR6733_3P | 121 | 0.43 | 1 | 0.507 | 0.867 |
| 3531 | MIR6801_3P | 48 | 0.48 | 1 | 0.479 | 0.867 |
| 3532 | MIR5587_5P | 98 | 0.44 | 1 | 0.49 | 0.867 |
| 3533 | MIR3934_3P | 86 | 0.44 | 1 | 0.467 | 0.868 |
| 3534 | MCBRYAN_PUBERTAL_BREAST_6_7WK_UP | 166 | 0.42 | 1 | 0.485 | 0.868 |
| 3535 | MIR514A_3P_MIR514B_3P | 89 | 0.45 | 1 | 0.495 | 0.868 |
| 3536 | MIR5006_3P | 314 | 0.41 | 1 | 0.501 | 0.868 |
| 3537 | REACTOME_REGULATION_OF_RAS_BY_GAPS | 66 | 0.46 | 1 | 0.488 | 0.868 |
| 3538 | SMITH_TERT_TARGETS_DN | 73 | 0.45 | 1 | 0.484 | 0.868 |
| 3539 | LEE_LIVER_CANCER_MYC_DN | 31 | 0.51 | 1 | 0.499 | 0.869 |
| 3540 | SU_PANCREAS | 23 | 0.53 | 1 | 0.493 | 0.869 |
| 3541 | MIR520G_3P_MIR520H | 186 | 0.42 | 1 | 0.488 | 0.869 |
| 3542 | TBK1.DF_DN | 268 | 0.41 | 1 | 0.488 | 0.869 |
| 3543 | YCATTAA_UNKNOWN | 409 | 0.41 | 1 | 0.514 | 0.869 |
| 3544 | REACTOME_DOWNSTREAM_SIGNAL_TRANSDUCTI | 29 | 0.51 | 1 | 0.502 | 0.87 |
| 3545 | CHIANG_LIVER_CANCER_SUBCLASS_CTNNB1_UP | 111 | 0.44 | 1 | 0.481 | 0.87 |
| 3546 | MIR34C_3P | 57 | 0.46 | 1 | 0.492 | 0.871 |
| 3547 | HOFFMANN_SMALL_PRE_BII_TO_IMMATURE_B_LY | 30 | 0.5 | 1 | 0.489 | 0.871 |
| 3548 | MIR6802_5P | 23 | 0.53 | 1 | 0.496 | 0.871 |
| 3549 | REACTOME_CDC42_GTPASE_CYCLE | 135 | 0.43 | 1 | 0.491 | 0.871 |
| 3550 | MIR556_3P | 105 | 0.44 | 1 | 0.498 | 0.871 |
| 3551 | WP_PROGERIAASSOCIATED_LIPODYSTROPHY | 20 | 0.53 | 1 | 0.496 | 0.871 |
| 3552 | MIR6718_5P | 62 | 0.45 | 1 | 0.491 | 0.871 |
| 3553 | ZNF510_TARGET_GENES | 36 | 0.5 | 1 | 0.487 | 0.871 |
| 3554 | DCA_UP.V1_DN | 144 | 0.43 | 1 | 0.492 | 0.871 |
| 3555 | MIR19B_1_5P_MIR19B_2_5P | 141 | 0.43 | 1 | 0.501 | 0.871 |
| 3556 | MIR3125 | 198 | 0.42 | 1 | 0.493 | 0.871 |
| 3557 | MIR7154_5P | 212 | 0.41 | 1 | 0.513 | 0.872 |
| 3558 | MIR4437 | 85 | 0.45 | 1 | 0.492 | 0.872 |
| 3559 | GCTNWTTGK_UNKNOWN | 245 | 0.41 | 1 | 0.505 | 0.873 |
| 3560 | REACTOME_TRANSLATION_OF_SARS_COV_2_STF | 48 | 0.48 | 1 | 0.484 | 0.873 |
| 3561 | SHETH_LIVER_CANCER_VS_TXNIP_LOSS_PAM1 | 212 | 0.42 | 1 | 0.526 | 0.873 |

| | | | | | | |
|---|---|---|---|---|---|---|
| 3562 | DURCHDEWALD_SKIN_CARCINOGENESIS_DN | 231 | 0.42 | 1 | 0.5 | 0.873 |
| 3563 | KRAS.AMP.LUNG_UP.V1_DN | 81 | 0.45 | 1 | 0.494 | 0.873 |
| 3564 | MIR3688_5P | 120 | 0.43 | 1 | 0.498 | 0.873 |
| 3565 | MIR203B_5P | 62 | 0.45 | 1 | 0.48 | 0.874 |
| 3566 | SP1_Q2_01 | 221 | 0.42 | 1 | 0.507 | 0.874 |
| 3567 | MIR6823_5P | 55 | 0.47 | 1 | 0.494 | 0.875 |
| 3568 | REACTOME_FGFR2_MUTANT_RECEPTOR_ACTIVAT | 21 | 0.54 | 1 | 0.512 | 0.875 |
| 3569 | ATF3_Q6 | 203 | 0.42 | 1 | 0.511 | 0.875 |
| 3570 | PID_NEPHRIN_NEPH1_PATHWAY | 27 | 0.51 | 1 | 0.511 | 0.876 |
| 3571 | PAX6_01 | 80 | 0.44 | 1 | 0.498 | 0.876 |
| 3572 | MIR579_3P | 413 | 0.41 | 1 | 0.528 | 0.876 |
| 3573 | PID_A6B1_A6B4_INTEGRIN_PATHWAY | 42 | 0.48 | 1 | 0.508 | 0.876 |
| 3574 | MIR4708_5P | 80 | 0.45 | 1 | 0.499 | 0.876 |
| 3575 | HOWLIN_CITED1_TARGETS_1_UP | 28 | 0.51 | 1 | 0.482 | 0.876 |
| 3576 | PID_IL6_7_PATHWAY | 41 | 0.49 | 0.99 | 0.477 | 0.876 |
| 3577 | MTOR_UP.N4.V1_UP | 176 | 0.42 | 0.99 | 0.517 | 0.876 |
| 3578 | MIR1909_5P | 32 | 0.5 | 0.99 | 0.492 | 0.877 |
| 3579 | PAX4_01 | 238 | 0.42 | 0.99 | 0.503 | 0.876 |
| 3580 | MIR3978 | 220 | 0.42 | 0.99 | 0.5 | 0.877 |
| 3581 | REACTOME_FCERI_MEDIATED_CA_2_MOBILIZATIC | 22 | 0.54 | 0.99 | 0.504 | 0.877 |
| 3582 | BANDRES_RESPONSE_TO_CARMUSTIN_WITHOUT_ | 26 | 0.51 | 0.99 | 0.503 | 0.877 |
| 3583 | ZNF654_TARGET_GENES | 448 | 0.41 | 0.99 | 0.518 | 0.877 |
| 3584 | CHIANG_LIVER_CANCER_SUBCLASS_UNANNOTAT | 63 | 0.46 | 0.99 | 0.502 | 0.878 |
| 3585 | BROWNE_HCMV_INFECTION_2HR_DN | 43 | 0.48 | 0.99 | 0.492 | 0.879 |
| 3586 | MIR4705 | 106 | 0.43 | 0.99 | 0.506 | 0.879 |
| 3587 | MIR766_5P | 176 | 0.42 | 0.99 | 0.51 | 0.878 |
| 3588 | GROSS_ELK3_TARGETS_UP | 24 | 0.53 | 0.99 | 0.509 | 0.878 |
| 3589 | MIR664B_3P | 410 | 0.4 | 0.99 | 0.519 | 0.879 |
| 3590 | MIR1225_5P | 29 | 0.5 | 0.99 | 0.492 | 0.879 |
| 3591 | REACTOME_RESPONSE_TO_ELEVATED_PLATELE | 90 | 0.44 | 0.99 | 0.511 | 0.879 |
| 3592 | NIKOLSKY_OVERCONNECTED_IN_BREAST_CANCE | 18 | 0.54 | 0.99 | 0.493 | 0.88 |
| 3593 | FOX_Q2 | 167 | 0.42 | 0.99 | 0.505 | 0.881 |
| 3594 | EPPERT_HSC_R | 97 | 0.44 | 0.99 | 0.511 | 0.882 |

| | | | | | | |
|---|---|---|---|---|---|---|
| 3595 | MIR128_3P | 404 | 0.41 | 0.99 | 0.538 | 0.882 |
| 3596 | MIR5702 | 58 | 0.46 | 0.99 | 0.496 | 0.881 |
| 3597 | RICKMAN_HEAD_AND_NECK_CANCER_B | 23 | 0.53 | 0.99 | 0.49 | 0.881 |
| 3598 | STAT1_03 | 210 | 0.42 | 0.99 | 0.522 | 0.881 |
| 3599 | MIR6761_5P | 97 | 0.44 | 0.99 | 0.51 | 0.881 |
| 3600 | KYNG_DNA_DAMAGE_BY_4NQO_OR_UV | 49 | 0.47 | 0.99 | 0.513 | 0.881 |
| 3601 | REACTOME_MET_PROMOTES_CELL_MOTILITY | 35 | 0.5 | 0.99 | 0.481 | 0.881 |
| 3602 | KORKOLA_YOLK_SAC_TUMOR | 42 | 0.49 | 0.99 | 0.525 | 0.881 |
| 3603 | MIR518C_5P | 158 | 0.42 | 0.99 | 0.512 | 0.881 |
| 3604 | MIR4307 | 347 | 0.41 | 0.99 | 0.53 | 0.881 |
| 3605 | MIR6833_5P | 103 | 0.44 | 0.99 | 0.504 | 0.881 |
| 3606 | CTACTGT_MIR199A | 167 | 0.42 | 0.99 | 0.527 | 0.881 |
| 3607 | DORSAM_HOXA9_TARGETS_UP | 32 | 0.5 | 0.99 | 0.493 | 0.881 |
| 3608 | KEGG_CARDIAC_MUSCLE_CONTRACTION | 54 | 0.46 | 0.99 | 0.503 | 0.881 |
| 3609 | BIOCARTA_NUCLEARRS_PATHWAY | 19 | 0.54 | 0.99 | 0.5 | 0.881 |
| 3610 | ZHAN_LATE_DIFFERENTIATION_GENES_UP | 26 | 0.51 | 0.99 | 0.486 | 0.882 |
| 3611 | GARCIA_TARGETS_OF_FLI1_AND_DAX1_UP | 44 | 0.48 | 0.99 | 0.51 | 0.882 |
| 3612 | REACTOME_TOLL_LIKE_RECEPTOR_9_TLR9_CAS | 87 | 0.45 | 0.99 | 0.484 | 0.883 |
| 3613 | MIR224_5P | 155 | 0.43 | 0.99 | 0.523 | 0.883 |
| 3614 | KIM_WT1_TARGETS_8HR_UP | 139 | 0.42 | 0.99 | 0.513 | 0.883 |
| 3615 | MIR6868_5P | 97 | 0.44 | 0.99 | 0.509 | 0.883 |
| 3616 | MIR19A_5P | 142 | 0.43 | 0.99 | 0.523 | 0.883 |
| 3617 | MIR760 | 115 | 0.43 | 0.99 | 0.514 | 0.883 |
| 3618 | MIR4790_3P | 67 | 0.45 | 0.99 | 0.481 | 0.883 |
| 3619 | SHAFFER_IRF4_TARGETS_IN_ACTIVATED_B_LYMI | 77 | 0.45 | 0.99 | 0.493 | 0.883 |
| 3620 | MIR656_3P | 337 | 0.41 | 0.99 | 0.543 | 0.884 |
| 3621 | MIR3622A_5P | 38 | 0.49 | 0.99 | 0.501 | 0.884 |
| 3622 | KEGG_PARKINSONS_DISEASE | 116 | 0.43 | 0.99 | 0.508 | 0.884 |
| 3623 | MIR23B_5P | 36 | 0.5 | 0.99 | 0.516 | 0.885 |
| 3624 | REACTOME_AMINO_ACID_TRANSPORT_ACROSS_ | 24 | 0.53 | 0.99 | 0.52 | 0.885 |
| 3625 | FOROUTAN_TGFB_EMT_DN | 84 | 0.45 | 0.99 | 0.508 | 0.885 |
| 3626 | MIR486_5P | 78 | 0.45 | 0.99 | 0.484 | 0.885 |
| 3627 | MIR4738_3P | 148 | 0.42 | 0.99 | 0.522 | 0.885 |

| | | | | | | |
|---|---|---|---|---|---|---|
| 3628 | CROMER_METASTASIS_UP | 68 | 0.45 | 0.99 | 0.525 | 0.885 |
| 3629 | WAMUNYOKOLI_OVARIAN_CANCER_GRADES_1_2 | 52 | 0.46 | 0.99 | 0.497 | 0.885 |
| 3630 | MIR183_3P | 159 | 0.42 | 0.99 | 0.51 | 0.885 |
| 3631 | ATF6_01 | 101 | 0.43 | 0.99 | 0.522 | 0.885 |
| 3632 | HNF3B_01 | 152 | 0.42 | 0.99 | 0.523 | 0.886 |
| 3633 | ZHU_CMV_8_HR_DN | 40 | 0.49 | 0.99 | 0.503 | 0.886 |
| 3634 | REACTOME_JNK_C_JUN_KINASES_PHOSPHORYL/ | 21 | 0.53 | 0.99 | 0.505 | 0.886 |
| 3635 | GTF3A_TARGET_GENES | 16 | 0.55 | 0.99 | 0.512 | 0.886 |
| 3636 | MIR30D_3P | 340 | 0.41 | 0.99 | 0.545 | 0.886 |
| 3637 | MIR378G | 70 | 0.45 | 0.99 | 0.505 | 0.886 |
| 3638 | MIR142_3P | 159 | 0.42 | 0.99 | 0.506 | 0.886 |
| 3639 | MIR4667_3P | 107 | 0.44 | 0.99 | 0.521 | 0.886 |
| 3640 | MIR5583_3P | 67 | 0.45 | 0.99 | 0.511 | 0.886 |
| 3641 | GACTGTT_MIR212_MIR132 | 151 | 0.43 | 0.99 | 0.522 | 0.888 |
| 3642 | MIR4753_3P | 454 | 0.4 | 0.99 | 0.547 | 0.889 |
| 3643 | AAGCACT_MIR520F | 216 | 0.41 | 0.99 | 0.517 | 0.889 |
| 3644 | HAHTOLA_MYCOSIS_FUNGOIDES_CD4_DN | 112 | 0.43 | 0.99 | 0.525 | 0.889 |
| 3645 | MIR939_3P | 126 | 0.43 | 0.99 | 0.512 | 0.889 |
| 3646 | MIR877_5P | 62 | 0.46 | 0.99 | 0.497 | 0.889 |
| 3647 | WP_VITAMIN_D_IN_INFLAMMATORY_DISEASES | 22 | 0.53 | 0.99 | 0.52 | 0.889 |
| 3648 | REACTOME_REGULATION_OF_SIGNALING_BY_CB | 20 | 0.54 | 0.99 | 0.518 | 0.889 |
| 3649 | JIANG_TIP30_TARGETS_UP | 40 | 0.48 | 0.99 | 0.512 | 0.89 |
| 3650 | OCT1_02 | 154 | 0.42 | 0.99 | 0.519 | 0.89 |
| 3651 | REACTOME_CYCLIN_D_ASSOCIATED_EVENTS_IN_ | 46 | 0.47 | 0.99 | 0.512 | 0.889 |
| 3652 | MIR1283 | 404 | 0.4 | 0.99 | 0.549 | 0.889 |
| 3653 | WILCOX_RESPONSE_TO_PROGESTERONE_UP | 128 | 0.43 | 0.99 | 0.508 | 0.889 |
| 3654 | PAL_PRMT5_TARGETS_UP | 188 | 0.42 | 0.99 | 0.538 | 0.889 |
| 3655 | MIR3152_3P | 62 | 0.45 | 0.99 | 0.526 | 0.889 |
| 3656 | ZNF781_TARGET_GENES | 116 | 0.43 | 0.99 | 0.519 | 0.889 |
| 3657 | MOOTHA_GLUCONEOGENESIS | 24 | 0.52 | 0.99 | 0.519 | 0.889 |
| 3658 | IVANOVA_HEMATOPOIESIS_LATE_PROGENITOR | 425 | 0.4 | 0.98 | 0.553 | 0.89 |
| 3659 | FLOTHO_PEDIATRIC_ALL_THERAPY_RESPONSE_I | 26 | 0.52 | 0.98 | 0.516 | 0.89 |
| 3660 | BREDEMEYER_RAG_SIGNALING_NOT_VIA_ATM_D | 48 | 0.47 | 0.98 | 0.499 | 0.89 |

| | | | | | | |
|---|---|---|---|---|---|---|
| 3661 | MIR10398_5P | 66 | 0.45 | 0.98 | 0.509 | 0.89 |
| 3662 | P53_DN.V2_DN | 93 | 0.44 | 0.98 | 0.519 | 0.891 |
| 3663 | MIR6798_3P | 32 | 0.5 | 0.98 | 0.509 | 0.891 |
| 3664 | GGGGCCC_MIR296 | 63 | 0.45 | 0.98 | 0.514 | 0.891 |
| 3665 | POS_HISTAMINE_RESPONSE_NETWORK | 28 | 0.51 | 0.98 | 0.52 | 0.892 |
| 3666 | GUTIERREZ_CHRONIC_LYMPHOCYTIC_LEUKEMIA_ | 46 | 0.47 | 0.98 | 0.518 | 0.892 |
| 3667 | TAZ_TARGET_GENES | 377 | 0.41 | 0.98 | 0.565 | 0.892 |
| 3668 | WP_PPAR_SIGNALING_PATHWAY | 37 | 0.49 | 0.98 | 0.495 | 0.893 |
| 3669 | MIR6734_3P | 124 | 0.43 | 0.98 | 0.531 | 0.893 |
| 3670 | MIR4267 | 190 | 0.41 | 0.98 | 0.544 | 0.893 |
| 3671 | REACTOME_SARS_COV_INFECTIONS | 140 | 0.42 | 0.98 | 0.523 | 0.893 |
| 3672 | CAGCTTT_MIR320 | 234 | 0.41 | 0.98 | 0.542 | 0.893 |
| 3673 | ATCTTGC_MIR31 | 71 | 0.45 | 0.98 | 0.522 | 0.894 |
| 3674 | CREBP1CJUN_01 | 227 | 0.41 | 0.98 | 0.531 | 0.894 |
| 3675 | KRAS.PROSTATE_UP.V1_DN | 71 | 0.45 | 0.98 | 0.52 | 0.894 |
| 3676 | MIR1253 | 130 | 0.43 | 0.98 | 0.55 | 0.894 |
| 3677 | ZNF558_TARGET_GENES | 231 | 0.41 | 0.98 | 0.539 | 0.894 |
| 3678 | NIKOLSKY_BREAST_CANCER_17Q21_Q25_AMPLI( | 257 | 0.41 | 0.98 | 0.542 | 0.894 |
| 3679 | MIR4290 | 57 | 0.46 | 0.98 | 0.511 | 0.894 |
| 3680 | REACTOME_G_PROTEIN_BETA_GAMMA_SIGNALLI | 24 | 0.51 | 0.98 | 0.521 | 0.894 |
| 3681 | RAX2_TARGET_GENES | 64 | 0.45 | 0.98 | 0.509 | 0.895 |
| 3682 | MADAN_DPPA4_TARGETS | 32 | 0.49 | 0.98 | 0.523 | 0.895 |
| 3683 | HSF_Q6 | 167 | 0.42 | 0.98 | 0.528 | 0.895 |
| 3684 | YATGNWAAT_OCT_C | 269 | 0.41 | 0.98 | 0.547 | 0.895 |
| 3685 | REACTOME_SIGNALING_BY_MODERATE_KINASE_ | 39 | 0.48 | 0.98 | 0.525 | 0.895 |
| 3686 | ZNF623_TARGET_GENES | 84 | 0.43 | 0.98 | 0.503 | 0.895 |
| 3687 | DAVICIONI_MOLECULAR_ARMS_VS_ERMS_DN | 142 | 0.42 | 0.98 | 0.544 | 0.894 |
| 3688 | MIR490_5P | 36 | 0.49 | 0.98 | 0.527 | 0.895 |
| 3689 | HESS_TARGETS_OF_HOXA9_AND_MEIS1_UP | 61 | 0.45 | 0.98 | 0.518 | 0.895 |
| 3690 | CHOP_01 | 199 | 0.41 | 0.98 | 0.533 | 0.895 |
| 3691 | MIR3923 | 42 | 0.48 | 0.98 | 0.511 | 0.897 |
| 3692 | MIR548T_5P | 368 | 0.4 | 0.98 | 0.585 | 0.897 |
| 3693 | MIR4481 | 107 | 0.43 | 0.98 | 0.538 | 0.897 |

| | | | | | | |
|---|---|---|---|---|---|---|
| 3694 | ABE_VEGFA_TARGETS_2HR | 25 | 0.51 | 0.98 | 0.515 | 0.897 |
| 3695 | REACTOME_G1_S_SPECIFIC_TRANSCRIPTION | 29 | 0.5 | 0.98 | 0.533 | 0.897 |
| 3696 | NIKOLSKY_BREAST_CANCER_11Q12_Q14_AMPLI( | 121 | 0.42 | 0.98 | 0.528 | 0.899 |
| 3697 | REACTOME_NETRIN_1_SIGNALING | 40 | 0.48 | 0.98 | 0.532 | 0.899 |
| 3698 | MIR3126_3P | 37 | 0.47 | 0.98 | 0.524 | 0.9 |
| 3699 | WANG_METASTASIS_OF_BREAST_CANCER_ESR1_ | 23 | 0.52 | 0.98 | 0.521 | 0.899 |
| 3700 | MIR8081 | 47 | 0.47 | 0.98 | 0.506 | 0.899 |
| 3701 | TGCACTG_MIR148A_MIR152_MIR148B | 276 | 0.41 | 0.98 | 0.555 | 0.899 |
| 3702 | MIR484 | 79 | 0.44 | 0.98 | 0.526 | 0.899 |
| 3703 | MIR4680_5P | 65 | 0.45 | 0.98 | 0.508 | 0.899 |
| 3704 | SRCAP_TARGET_GENES | 373 | 0.4 | 0.98 | 0.575 | 0.9 |
| 3705 | MIR6876_5P | 194 | 0.41 | 0.98 | 0.538 | 0.9 |
| 3706 | ACEVEDO_NORMAL_TISSUE_ADJACENT_TO_LIVE | 139 | 0.42 | 0.98 | 0.55 | 0.9 |
| 3707 | MARIADASON_RESPONSE_TO_BUTYRATE_SULINI | 45 | 0.47 | 0.98 | 0.513 | 0.9 |
| 3708 | REACTOME_PI3K_AKT_SIGNALING_IN_CANCER | 81 | 0.44 | 0.98 | 0.52 | 0.9 |
| 3709 | JOSEPH_RESPONSE_TO_SODIUM_BUTYRATE_DN | 47 | 0.47 | 0.98 | 0.51 | 0.9 |
| 3710 | MIR6778_3P | 194 | 0.41 | 0.98 | 0.551 | 0.9 |
| 3711 | AREB6_01 | 207 | 0.41 | 0.98 | 0.546 | 0.9 |
| 3712 | REACTOME_NEDDYLATION | 208 | 0.41 | 0.98 | 0.549 | 0.9 |
| 3713 | MIR581 | 65 | 0.45 | 0.98 | 0.525 | 0.9 |
| 3714 | LAIHO_COLORECTAL_CANCER_SERRATED_DN | 83 | 0.44 | 0.98 | 0.531 | 0.9 |
| 3715 | MIR6165 | 105 | 0.43 | 0.98 | 0.534 | 0.901 |
| 3716 | MOREIRA_RESPONSE_TO_TSA_UP | 27 | 0.5 | 0.98 | 0.523 | 0.9 |
| 3717 | FIRESTEIN_CTNNB1_PATHWAY | 31 | 0.5 | 0.98 | 0.527 | 0.9 |
| 3718 | BARRIER_CANCER_RELAPSE_NORMAL_SAMPLE_ | 31 | 0.5 | 0.98 | 0.51 | 0.9 |
| 3719 | PLASARI_TGFB1_SIGNALING_VIA_NFIC_1HR_DN | 92 | 0.44 | 0.98 | 0.532 | 0.9 |
| 3720 | UEDA_PERIFERAL_CLOCK | 143 | 0.42 | 0.98 | 0.553 | 0.9 |
| 3721 | WP_MECP2_AND_ASSOCIATED_RETT_SYNDROME | 60 | 0.45 | 0.98 | 0.513 | 0.9 |
| 3722 | WELCSH_BRCA1_TARGETS_UP | 182 | 0.41 | 0.98 | 0.548 | 0.901 |
| 3723 | THRAP3_TARGET_GENES | 19 | 0.53 | 0.98 | 0.532 | 0.9 |
| 3724 | LASTOWSKA_COAMPLIFIED_WITH_MYCN | 33 | 0.49 | 0.98 | 0.527 | 0.902 |
| 3725 | REACTOME_DAG_AND_IP3_SIGNALING | 35 | 0.49 | 0.98 | 0.507 | 0.903 |
| 3726 | MZF1_01 | 201 | 0.41 | 0.98 | 0.562 | 0.903 |

| | | | | | | |
|---|---|---|---|---|---|---|
| 3727 | IIZUKA_LIVER_CANCER_PROGRESSION_G2_G3_U | 22 | 0.52 | 0.98 | 0.528 | 0.903 |
| 3728 | CRX_Q4 | 193 | 0.41 | 0.98 | 0.551 | 0.903 |
| 3729 | MIR4423_3P | 112 | 0.43 | 0.98 | 0.531 | 0.903 |
| 3730 | MIR6740_3P | 128 | 0.42 | 0.98 | 0.55 | 0.903 |
| 3731 | SREBP_Q3 | 198 | 0.41 | 0.98 | 0.55 | 0.903 |
| 3732 | MIR4721 | 59 | 0.45 | 0.98 | 0.513 | 0.903 |
| 3733 | GTTATAT_MIR410 | 80 | 0.44 | 0.98 | 0.522 | 0.903 |
| 3734 | MIR6780A_3P | 137 | 0.42 | 0.98 | 0.56 | 0.903 |
| 3735 | WP_CALCIUM_REGULATION_IN_CARDIAC_CELLS | 109 | 0.42 | 0.98 | 0.549 | 0.904 |
| 3736 | MIR4668_5P | 407 | 0.4 | 0.97 | 0.593 | 0.904 |
| 3737 | PID_ENDOTHELIN_PATHWAY | 49 | 0.46 | 0.97 | 0.518 | 0.904 |
| 3738 | LMO2COM_01 | 224 | 0.4 | 0.97 | 0.556 | 0.905 |
| 3739 | GARGALOVIC_RESPONSE_TO_OXIDIZED_PHOSPH | 16 | 0.55 | 0.97 | 0.542 | 0.905 |
| 3740 | REACTOME_GPVI_MEDIATED_ACTIVATION_CASCA | 28 | 0.5 | 0.97 | 0.514 | 0.906 |
| 3741 | STAMBOLSKY_RESPONSE_TO_VITAMIN_D3_DN | 19 | 0.53 | 0.97 | 0.525 | 0.906 |
| 3742 | GCCNNNWTAAR_UNKNOWN | 125 | 0.42 | 0.97 | 0.554 | 0.907 |
| 3743 | REACTOME_IKK_COMPLEX_RECRUITMENT_MEDIA | 21 | 0.52 | 0.97 | 0.533 | 0.907 |
| 3744 | REACTOME_INTRACELLULAR_SIGNALING_BY_SE( | 267 | 0.4 | 0.97 | 0.578 | 0.907 |
| 3745 | MIR103A_1_5P | 126 | 0.42 | 0.97 | 0.544 | 0.907 |
| 3746 | MIR8089 | 150 | 0.41 | 0.97 | 0.561 | 0.906 |
| 3747 | BROWNE_HCMV_INFECTION_6HR_DN | 145 | 0.41 | 0.97 | 0.547 | 0.906 |
| 3748 | MIR3187_5P | 94 | 0.43 | 0.97 | 0.526 | 0.906 |
| 3749 | CACTGCC_MIR34A_MIR34C_MIR449 | 258 | 0.4 | 0.97 | 0.572 | 0.906 |
| 3750 | MEISSNER_BRAIN_HCP_WITH_H3K27ME3 | 148 | 0.42 | 0.97 | 0.536 | 0.906 |
| 3751 | MARSHALL_VIRAL_INFECTION_RESPONSE_DN | 22 | 0.53 | 0.97 | 0.554 | 0.906 |
| 3752 | MXD1_TARGET_GENES | 134 | 0.42 | 0.97 | 0.562 | 0.906 |
| 3753 | LINDGREN_BLADDER_CANCER_CLUSTER_3_UP | 307 | 0.4 | 0.97 | 0.589 | 0.906 |
| 3754 | LEIN_CEREBELLUM_MARKERS | 65 | 0.45 | 0.97 | 0.519 | 0.905 |
| 3755 | RODWELL_AGING_KIDNEY_DN | 130 | 0.42 | 0.97 | 0.551 | 0.905 |
| 3756 | ACACTGG_MIR199A_MIR199B | 146 | 0.42 | 0.97 | 0.543 | 0.906 |
| 3757 | YAO_TEMPORAL_RESPONSE_TO_PROGESTERON | 27 | 0.5 | 0.97 | 0.537 | 0.906 |
| 3758 | HOXD1_TARGET_GENES | 53 | 0.45 | 0.97 | 0.533 | 0.905 |
| 3759 | GARGALOVIC_RESPONSE_TO_OXIDIZED_PHOSPH | 16 | 0.55 | 0.97 | 0.544 | 0.905 |

| | | | | | | |
|---|---|---|---|---|---|---|
| 3760 | MIR3678_3P | 99 | 0.43 | 0.97 | 0.548 | 0.906 |
| 3761 | ETS_Q4 | 213 | 0.41 | 0.97 | 0.563 | 0.906 |
| 3762 | MIR3661 | 53 | 0.46 | 0.97 | 0.537 | 0.907 |
| 3763 | MIR6888_3P | 73 | 0.44 | 0.97 | 0.538 | 0.907 |
| 3764 | MIR6738_3P | 129 | 0.42 | 0.97 | 0.552 | 0.906 |
| 3765 | REACTOME_NEGATIVE_REGULATION_OF_FGFR4_ | 21 | 0.53 | 0.97 | 0.553 | 0.906 |
| 3766 | MIR622 | 73 | 0.44 | 0.97 | 0.528 | 0.906 |
| 3767 | P300_01 | 215 | 0.4 | 0.97 | 0.551 | 0.907 |
| 3768 | MIR12123 | 393 | 0.4 | 0.97 | 0.596 | 0.907 |
| 3769 | MIR4682 | 112 | 0.42 | 0.97 | 0.55 | 0.907 |
| 3770 | MIR548AT_5P | 313 | 0.4 | 0.97 | 0.579 | 0.907 |
| 3771 | CHESLER_BRAIN_QTL_CIS | 65 | 0.44 | 0.97 | 0.535 | 0.907 |
| 3772 | WHITFIELD_CELL_CYCLE_S | 139 | 0.42 | 0.97 | 0.54 | 0.909 |
| 3773 | GAUSSMANN_MLL_AF4_FUSION_TARGETS_A_UP | 163 | 0.41 | 0.97 | 0.56 | 0.909 |
| 3774 | MIR675_3P | 54 | 0.46 | 0.97 | 0.555 | 0.91 |
| 3775 | WP_CONGENITAL_GENERALIZED_LIPODYSTROPH | 18 | 0.54 | 0.97 | 0.53 | 0.91 |
| 3776 | BERENJENO_TRANSFORMED_BY_RHOA_REVERSI | 26 | 0.5 | 0.97 | 0.54 | 0.912 |
| 3777 | MIR6806_5P | 55 | 0.46 | 0.97 | 0.546 | 0.912 |
| 3778 | PID_HIF1A_PATHWAY | 19 | 0.53 | 0.97 | 0.546 | 0.912 |
| 3779 | MIR4666A_5P | 211 | 0.4 | 0.97 | 0.586 | 0.912 |
| 3780 | REACTOME_NEGATIVE_REGULATION_OF_THE_PI3 | 85 | 0.43 | 0.97 | 0.556 | 0.914 |
| 3781 | DING_LUNG_CANCER_MUTATED_SIGNIFICANTLY | 24 | 0.51 | 0.97 | 0.544 | 0.914 |
| 3782 | KTGGYRSGAA_UNKNOWN | 73 | 0.44 | 0.97 | 0.546 | 0.914 |
| 3783 | WANG_LSD1_TARGETS_UP | 19 | 0.53 | 0.97 | 0.538 | 0.913 |
| 3784 | MIR635 | 39 | 0.47 | 0.97 | 0.52 | 0.913 |
| 3785 | IPF1_Q4 | 186 | 0.41 | 0.97 | 0.58 | 0.913 |
| 3786 | ZWANG_CLASS_2_TRANSIENTLY_INDUCED_BY_E( | 32 | 0.49 | 0.97 | 0.549 | 0.913 |
| 3787 | MIR6888_5P | 39 | 0.47 | 0.97 | 0.528 | 0.913 |
| 3788 | PPARGC1A_TARGET_GENES | 84 | 0.43 | 0.97 | 0.534 | 0.913 |
| 3789 | GAZDA_DIAMOND_BLACKFAN_ANEMIA_ERYTHROI | 396 | 0.4 | 0.97 | 0.614 | 0.913 |
| 3790 | STAMBOLSKY_RESPONSE_TO_VITAMIN_D3_UP | 64 | 0.45 | 0.97 | 0.533 | 0.914 |
| 3791 | NABA_BASEMENT_MEMBRANES | 25 | 0.5 | 0.97 | 0.541 | 0.914 |
| 3792 | BIOCARTA_ATRBRCA_PATHWAY | 22 | 0.52 | 0.97 | 0.543 | 0.914 |

| | | | | | | |
|---|---|---|---|---|---|---|
| 3793 | MIR150_5P | 148 | 0.41 | 0.97 | 0.574 | 0.914 |
| 3794 | MIR5586_3P | 38 | 0.47 | 0.97 | 0.548 | 0.914 |
| 3795 | MIR3187_3P | 67 | 0.45 | 0.97 | 0.549 | 0.914 |
| 3796 | P53_DN.V1_UP | 153 | 0.41 | 0.97 | 0.561 | 0.913 |
| 3797 | KYNG_ENVIRONMENTAL_STRESS_RESPONSE_DN | 16 | 0.55 | 0.97 | 0.545 | 0.913 |
| 3798 | CAHOY_NEURONAL | 67 | 0.45 | 0.97 | 0.557 | 0.914 |
| 3799 | PETROVA_ENDOTHELIUM_LYMPHATIC_VS_BLOOD | 105 | 0.42 | 0.97 | 0.565 | 0.913 |
| 3800 | PITX2_Q2 | 193 | 0.4 | 0.97 | 0.563 | 0.913 |
| 3801 | HMEF2_Q6 | 92 | 0.43 | 0.97 | 0.55 | 0.915 |
| 3802 | REACTOME_SLC_TRANSPORTER_DISORDERS | 63 | 0.45 | 0.97 | 0.537 | 0.915 |
| 3803 | PPARA_01 | 28 | 0.5 | 0.97 | 0.525 | 0.915 |
| 3804 | YTTCCNNNGGAMR_UNKNOWN | 48 | 0.47 | 0.97 | 0.534 | 0.915 |
| 3805 | CROONQUIST_IL6_DEPRIVATION_DN | 90 | 0.43 | 0.97 | 0.53 | 0.915 |
| 3806 | REACTOME_METABOLISM_OF_FOLATE_AND_PTE | 15 | 0.55 | 0.97 | 0.586 | 0.916 |
| 3807 | MIR4271 | 179 | 0.41 | 0.97 | 0.579 | 0.916 |
| 3808 | MIR30E_3P | 338 | 0.4 | 0.97 | 0.597 | 0.916 |
| 3809 | MIR3670 | 49 | 0.45 | 0.97 | 0.536 | 0.916 |
| 3810 | VERNELL_RETINOBLASTOMA_PATHWAY_UP | 69 | 0.44 | 0.97 | 0.554 | 0.916 |
| 3811 | WP_SPHINGOLIPID_PATHWAY | 26 | 0.5 | 0.97 | 0.567 | 0.916 |
| 3812 | MIR587 | 298 | 0.4 | 0.97 | 0.584 | 0.916 |
| 3813 | CTAWWWATA_RSRFC4_Q2 | 258 | 0.4 | 0.97 | 0.594 | 0.916 |
| 3814 | HNF3_Q6 | 139 | 0.41 | 0.97 | 0.559 | 0.916 |
| 3815 | TANAKA_METHYLATED_IN_ESOPHAGEAL_CARCIN | 73 | 0.44 | 0.97 | 0.555 | 0.917 |
| 3816 | CP2_02 | 208 | 0.4 | 0.97 | 0.578 | 0.916 |
| 3817 | REACTOME_APC_C_CDH1_MEDIATED_DEGRADAT | 72 | 0.44 | 0.97 | 0.528 | 0.916 |
| 3818 | MIR503_5P | 92 | 0.43 | 0.96 | 0.557 | 0.916 |
| 3819 | MIR152_5P | 79 | 0.43 | 0.96 | 0.525 | 0.916 |
| 3820 | MIR12118 | 85 | 0.43 | 0.96 | 0.554 | 0.916 |
| 3821 | ZNF394_TARGET_GENES | 335 | 0.4 | 0.96 | 0.614 | 0.918 |
| 3822 | REACTOME_PEPTIDE_HORMONE_METABOLISM | 49 | 0.45 | 0.96 | 0.546 | 0.919 |
| 3823 | SESTO_RESPONSE_TO_UV_C7 | 60 | 0.44 | 0.96 | 0.547 | 0.919 |
| 3824 | ZHAN_MULTIPLE_MYELOMA_HP_UP | 38 | 0.47 | 0.96 | 0.535 | 0.919 |
| 3825 | REACTOME_SARS_COV_2_INFECTION | 70 | 0.44 | 0.96 | 0.553 | 0.919 |

| | | | | | | |
|---|---|---|---|---|---|---|
| 3826 | AP2REP_01 | 146 | 0.42 | 0.96 | 0.591 | 0.919 |
| 3827 | MIR134_3P | 73 | 0.44 | 0.96 | 0.553 | 0.919 |
| 3828 | MIR7976 | 89 | 0.42 | 0.96 | 0.545 | 0.919 |
| 3829 | MIR3191_5P | 133 | 0.41 | 0.96 | 0.576 | 0.92 |
| 3830 | MIR3151_3P | 32 | 0.5 | 0.96 | 0.562 | 0.919 |
| 3831 | TFIIA_Q6 | 207 | 0.4 | 0.96 | 0.57 | 0.919 |
| 3832 | REACTOME_RHOA_GTPASE_CYCLE | 134 | 0.42 | 0.96 | 0.576 | 0.919 |
| 3833 | WP_MONOAMINE_TRANSPORT | 19 | 0.53 | 0.96 | 0.576 | 0.919 |
| 3834 | MIR5190 | 114 | 0.42 | 0.96 | 0.572 | 0.919 |
| 3835 | REACTOME_SIGNALING_BY_FGFR1_IN_DISEASE | 31 | 0.48 | 0.96 | 0.539 | 0.92 |
| 3836 | MIR3173_3P | 87 | 0.43 | 0.96 | 0.557 | 0.92 |
| 3837 | HOEGERKORP_CD44_TARGETS_DIRECT_UP | 19 | 0.53 | 0.96 | 0.57 | 0.92 |
| 3838 | MIR369_3P | 257 | 0.4 | 0.96 | 0.605 | 0.92 |
| 3839 | MIR4427 | 197 | 0.4 | 0.96 | 0.578 | 0.921 |
| 3840 | MIR103A_2_5P | 129 | 0.42 | 0.96 | 0.572 | 0.92 |
| 3841 | HOXA10_TARGET_GENES | 245 | 0.4 | 0.96 | 0.601 | 0.921 |
| 3842 | WGTTNNNNNAAA_UNKNOWN | 438 | 0.39 | 0.96 | 0.621 | 0.921 |
| 3843 | MIR6799_5P | 120 | 0.42 | 0.96 | 0.549 | 0.921 |
| 3844 | MEBARKI_HCC_PROGENITOR_WNT_UP_CTNNB1_[illegible] | 55 | 0.45 | 0.96 | 0.56 | 0.921 |
| 3845 | MIR3129_5P | 130 | 0.41 | 0.96 | 0.567 | 0.921 |
| 3846 | MIR30A_3P | 337 | 0.4 | 0.96 | 0.617 | 0.921 |
| 3847 | MIR4326 | 66 | 0.45 | 0.96 | 0.562 | 0.921 |
| 3848 | KRAS.50_UP.V1_DN | 23 | 0.51 | 0.96 | 0.55 | 0.921 |
| 3849 | RICKMAN_TUMOR_DIFFERENTIATED_WELL_VS_P[illegible] | 317 | 0.39 | 0.96 | 0.627 | 0.921 |
| 3850 | REACTOME_L1CAM_INTERACTIONS | 103 | 0.42 | 0.96 | 0.553 | 0.921 |
| 3851 | MIR192_5P | 33 | 0.48 | 0.96 | 0.538 | 0.921 |
| 3852 | REACTOME_ACTIVATION_OF_IRF3_IRF7_MEDIATE | 16 | 0.54 | 0.96 | 0.567 | 0.921 |
| 3853 | MIR6848_3P | 77 | 0.44 | 0.96 | 0.57 | 0.921 |
| 3854 | GATA1_03 | 178 | 0.41 | 0.96 | 0.58 | 0.922 |
| 3855 | KAAB_HEART_ATRIUM_VS_VENTRICLE_UP | 195 | 0.4 | 0.96 | 0.602 | 0.921 |
| 3856 | MIR1301_3P_MIR5047 | 98 | 0.42 | 0.96 | 0.564 | 0.921 |
| 3857 | SENGUPTA_NASOPHARYNGEAL_CARCINOMA_DN | 209 | 0.41 | 0.96 | 0.595 | 0.922 |
| 3858 | REACTOME_NOTCH_HLH_TRANSCRIPTION_PATHW | 27 | 0.5 | 0.96 | 0.557 | 0.922 |

| | | | | | | |
|---|---|---|---|---|---|---|
| 3859 | BIOCARTA_TCR_PATHWAY | 38 | 0.48 | 0.96 | 0.555 | 0.922 |
| 3860 | HUTTMANN_B_CLL_POOR_SURVIVAL_DN | 49 | 0.45 | 0.96 | 0.539 | 0.922 |
| 3861 | MIR548AV_5P_MIR548K | 257 | 0.4 | 0.96 | 0.613 | 0.922 |
| 3862 | MIR3127_5P | 55 | 0.45 | 0.96 | 0.556 | 0.922 |
| 3863 | MIR888_5P | 203 | 0.41 | 0.96 | 0.592 | 0.922 |
| 3864 | KORKOLA_EMBRYONIC_CARCINOMA_VS_SEMINO | 16 | 0.54 | 0.96 | 0.575 | 0.923 |
| 3865 | MIR141_5P | 78 | 0.43 | 0.96 | 0.572 | 0.923 |
| 3866 | GROSS_HIF1A_TARGETS_DN | 21 | 0.52 | 0.96 | 0.553 | 0.923 |
| 3867 | HUANG_FOXA2_TARGETS_UP | 43 | 0.46 | 0.96 | 0.565 | 0.923 |
| 3868 | CETS1P54_01 | 240 | 0.4 | 0.96 | 0.607 | 0.924 |
| 3869 | MIR4679 | 121 | 0.41 | 0.96 | 0.555 | 0.924 |
| 3870 | MIR6838_3P | 52 | 0.45 | 0.96 | 0.539 | 0.924 |
| 3871 | PID_FAK_PATHWAY | 56 | 0.45 | 0.96 | 0.576 | 0.924 |
| 3872 | TGAATGT_MIR181A_MIR181B_MIR181C_MIR181D | 445 | 0.39 | 0.96 | 0.644 | 0.924 |
| 3873 | CHIARETTI_T_ALL_REFRACTORY_TO_THERAPY | 26 | 0.5 | 0.96 | 0.557 | 0.925 |
| 3874 | BYSTRYKH_HEMATOPOIESIS_STEM_CELL_AND_E | 134 | 0.41 | 0.96 | 0.572 | 0.925 |
| 3875 | NKX2_8_TARGET_GENES | 30 | 0.48 | 0.96 | 0.557 | 0.925 |
| 3876 | GTGACTT_MIR224 | 141 | 0.41 | 0.96 | 0.576 | 0.925 |
| 3877 | MIR4667_5P | 151 | 0.41 | 0.96 | 0.581 | 0.925 |
| 3878 | REACTOME_O_GLYCOSYLATION_OF_TSR_DOMAI | 28 | 0.49 | 0.96 | 0.558 | 0.925 |
| 3879 | MIR6895_3P | 69 | 0.43 | 0.96 | 0.571 | 0.925 |
| 3880 | WP_WHITE_FAT_CELL_DIFFERENTIATION | 29 | 0.49 | 0.96 | 0.539 | 0.925 |
| 3881 | PASQUALUCCI_LYMPHOMA_BY_GC_STAGE_UP | 252 | 0.4 | 0.96 | 0.63 | 0.925 |
| 3882 | KEGG_GAP_JUNCTION | 71 | 0.44 | 0.96 | 0.574 | 0.925 |
| 3883 | HOFFMANN_LARGE_TO_SMALL_PRE_BII_LYMPHC | 146 | 0.41 | 0.96 | 0.581 | 0.925 |
| 3884 | MIR3176 | 66 | 0.44 | 0.96 | 0.546 | 0.925 |
| 3885 | MIR4446_5P | 256 | 0.4 | 0.96 | 0.613 | 0.925 |
| 3886 | CIC_TARGET_GENES | 349 | 0.4 | 0.96 | 0.636 | 0.926 |
| 3887 | MIR6843_3P | 77 | 0.44 | 0.96 | 0.589 | 0.927 |
| 3888 | MIR4279 | 188 | 0.4 | 0.96 | 0.606 | 0.928 |
| 3889 | MIR196A_1_3P | 337 | 0.4 | 0.96 | 0.646 | 0.928 |
| 3890 | MIR3133 | 436 | 0.39 | 0.96 | 0.647 | 0.928 |
| 3891 | MIR4795_5P | 57 | 0.45 | 0.96 | 0.55 | 0.928 |

| | | | | | | |
|---|---|---|---|---|---|---|
| 3892 | GCTGAGT_MIR5125P | 46 | 0.46 | 0.96 | 0.546 | 0.928 |
| 3893 | LI_CYTIDINE_ANALOGS_CYCTOTOXICITY | 15 | 0.54 | 0.96 | 0.563 | 0.928 |
| 3894 | BASAKI_YBX1_TARGETS_UP | 268 | 0.39 | 0.96 | 0.622 | 0.928 |
| 3895 | GATA4_Q3 | 195 | 0.4 | 0.96 | 0.591 | 0.928 |
| 3896 | MIR586 | 235 | 0.39 | 0.95 | 0.6 | 0.929 |
| 3897 | EPPERT_CE_HSC_LSC | 27 | 0.49 | 0.95 | 0.543 | 0.93 |
| 3898 | ZNF697_TARGET_GENES | 47 | 0.46 | 0.95 | 0.549 | 0.93 |
| 3899 | RAMASWAMY_METASTASIS_DN | 41 | 0.46 | 0.95 | 0.562 | 0.93 |
| 3900 | OCT1_Q6 | 209 | 0.4 | 0.95 | 0.596 | 0.93 |
| 3901 | ACEVEDO_LIVER_CANCER_DN | 420 | 0.39 | 0.95 | 0.657 | 0.93 |
| 3902 | CHEBOTAEV_GR_TARGETS_DN | 87 | 0.43 | 0.95 | 0.575 | 0.93 |
| 3903 | MYOD_Q6 | 197 | 0.4 | 0.95 | 0.616 | 0.93 |
| 3904 | REACTOME_GLYCEROPHOSPHOLIPID_BIOSYNTHI | 97 | 0.42 | 0.95 | 0.567 | 0.93 |
| 3905 | MIR146B_3P | 60 | 0.44 | 0.95 | 0.576 | 0.931 |
| 3906 | MIR520F_5P | 161 | 0.41 | 0.95 | 0.583 | 0.931 |
| 3907 | NRL_DN.V1_UP | 107 | 0.42 | 0.95 | 0.587 | 0.931 |
| 3908 | MIR4657 | 57 | 0.44 | 0.95 | 0.558 | 0.931 |
| 3909 | MIR4524B_5P | 208 | 0.4 | 0.95 | 0.617 | 0.931 |
| 3910 | MIR8054 | 257 | 0.4 | 0.95 | 0.634 | 0.931 |
| 3911 | MIR1276 | 148 | 0.41 | 0.95 | 0.585 | 0.931 |
| 3912 | WP_NUCLEOTIDE_EXCISION_REPAIR_IN_XERODE | 65 | 0.44 | 0.95 | 0.569 | 0.932 |
| 3913 | FOXD3_01 | 162 | 0.4 | 0.95 | 0.584 | 0.932 |
| 3914 | MIR421 | 151 | 0.41 | 0.95 | 0.595 | 0.932 |
| 3915 | MIR155_5P | 197 | 0.4 | 0.95 | 0.608 | 0.932 |
| 3916 | MIR510_5P | 60 | 0.44 | 0.95 | 0.567 | 0.932 |
| 3917 | CREBP1_Q2 | 223 | 0.4 | 0.95 | 0.625 | 0.932 |
| 3918 | MIR6757_3P | 97 | 0.42 | 0.95 | 0.578 | 0.932 |
| 3919 | CASORELLI_ACUTE_PROMYELOCYTIC_LEUKEMIA | 136 | 0.41 | 0.95 | 0.588 | 0.932 |
| 3920 | BHATTACHARYA_EMBRYONIC_STEM_CELL | 70 | 0.43 | 0.95 | 0.561 | 0.932 |
| 3921 | MIR6871_5P | 52 | 0.45 | 0.95 | 0.561 | 0.933 |
| 3922 | MIR3085_3P | 133 | 0.41 | 0.95 | 0.6 | 0.933 |
| 3923 | MIR3145_5P | 86 | 0.42 | 0.95 | 0.589 | 0.933 |
| 3924 | MIR652_5P | 32 | 0.48 | 0.95 | 0.554 | 0.933 |

| | | | | | | |
|---|---|---|---|---|---|---|
| 3925 | MIR3189_3P | 65 | 0.44 | 0.95 | 0.572 | 0.933 |
| 3926 | REACTOME_NRAGE_SIGNALS_DEATH_THROUGH_ | 52 | 0.45 | 0.95 | 0.583 | 0.933 |
| 3927 | MASSARWEH_RESPONSE_TO_ESTRADIOL | 52 | 0.45 | 0.95 | 0.569 | 0.933 |
| 3928 | MIR211_3P | 72 | 0.43 | 0.95 | 0.549 | 0.933 |
| 3929 | MIR326 | 110 | 0.42 | 0.95 | 0.591 | 0.933 |
| 3930 | TGACATY_UNKNOWN | 495 | 0.39 | 0.95 | 0.675 | 0.933 |
| 3931 | MIR4524A_5P | 208 | 0.4 | 0.95 | 0.61 | 0.933 |
| 3932 | MIR670_3P | 254 | 0.4 | 0.95 | 0.639 | 0.933 |
| 3933 | MIR3919 | 255 | 0.4 | 0.95 | 0.637 | 0.933 |
| 3934 | PIT1_Q6 | 180 | 0.4 | 0.95 | 0.611 | 0.933 |
| 3935 | LUCAS_HNF4A_TARGETS_UP | 49 | 0.45 | 0.95 | 0.573 | 0.934 |
| 3936 | WP_PHOTODYNAMIC_THERAPYINDUCED_HIF1_SU | 31 | 0.49 | 0.95 | 0.584 | 0.934 |
| 3937 | DBP_Q6 | 199 | 0.4 | 0.95 | 0.61 | 0.934 |
| 3938 | MIR5189_5P | 118 | 0.41 | 0.95 | 0.599 | 0.935 |
| 3939 | ACEVEDO_LIVER_CANCER_WITH_H3K9ME3_DN | 78 | 0.43 | 0.95 | 0.575 | 0.935 |
| 3940 | MIR186_3P | 128 | 0.41 | 0.95 | 0.601 | 0.935 |
| 3941 | KIM_WT1_TARGETS_DN | 435 | 0.39 | 0.95 | 0.671 | 0.935 |
| 3942 | MIR610 | 70 | 0.43 | 0.95 | 0.57 | 0.935 |
| 3943 | AR_Q6 | 205 | 0.4 | 0.95 | 0.611 | 0.936 |
| 3944 | MIR383_3P | 111 | 0.41 | 0.95 | 0.592 | 0.936 |
| 3945 | REACTOME_SUMOYLATION | 167 | 0.4 | 0.95 | 0.617 | 0.936 |
| 3946 | KYNG_ENVIRONMENTAL_STRESS_RESPONSE_NC | 28 | 0.49 | 0.95 | 0.579 | 0.935 |
| 3947 | GYORFFY_MITOXANTRONE_RESISTANCE | 27 | 0.49 | 0.95 | 0.576 | 0.935 |
| 3948 | MIR4286 | 80 | 0.42 | 0.95 | 0.57 | 0.935 |
| 3949 | CEBPA_01 | 194 | 0.4 | 0.95 | 0.628 | 0.935 |
| 3950 | MIR215_3P | 109 | 0.41 | 0.95 | 0.582 | 0.936 |
| 3951 | GARGALOVIC_RESPONSE_TO_OXIDIZED_PHOSPH | 53 | 0.45 | 0.95 | 0.595 | 0.935 |
| 3952 | MIR4325 | 233 | 0.4 | 0.95 | 0.62 | 0.935 |
| 3953 | WAMUNYOKOLI_OVARIAN_CANCER_GRADES_1_2 | 124 | 0.41 | 0.95 | 0.593 | 0.935 |
| 3954 | MIR3681_3P | 353 | 0.39 | 0.95 | 0.668 | 0.935 |
| 3955 | BIOCARTA_GSK3_PATHWAY | 17 | 0.52 | 0.95 | 0.58 | 0.936 |
| 3956 | KRAS.LUNG.BREAST_UP.V1_DN | 83 | 0.43 | 0.95 | 0.585 | 0.936 |
| 3957 | MIR221_3P | 153 | 0.4 | 0.95 | 0.595 | 0.936 |

| | | | | | | |
|---|---|---|---|---|---|---|
| 3958 | MIR6881_5P | 47 | 0.45 | 0.95 | 0.576 | 0.936 |
| 3959 | IBRAHIM_NRF1_UP | 403 | 0.39 | 0.95 | 0.655 | 0.937 |
| 3960 | MIR222_3P | 150 | 0.41 | 0.95 | 0.6 | 0.937 |
| 3961 | AAANWWTGC_UNKNOWN | 150 | 0.41 | 0.95 | 0.608 | 0.937 |
| 3962 | LEIN_LOCALIZED_TO_DISTAL_AND_PROXIMAL_DE | 18 | 0.52 | 0.95 | 0.58 | 0.938 |
| 3963 | MIR4789_3P | 291 | 0.39 | 0.95 | 0.652 | 0.938 |
| 3964 | SCHLOSSER_MYC_TARGETS_AND_SERUM_RESPO | 48 | 0.45 | 0.95 | 0.582 | 0.938 |
| 3965 | MIR1_3P | 344 | 0.39 | 0.95 | 0.65 | 0.938 |
| 3966 | ZHAN_MULTIPLE_MYELOMA_PR_DN | 43 | 0.46 | 0.95 | 0.576 | 0.938 |
| 3967 | PAX8_B | 82 | 0.43 | 0.95 | 0.577 | 0.939 |
| 3968 | MIR5008_3P | 147 | 0.4 | 0.95 | 0.625 | 0.939 |
| 3969 | CTTTGCA_MIR527 | 220 | 0.39 | 0.95 | 0.635 | 0.939 |
| 3970 | MIR561_3P | 433 | 0.39 | 0.95 | 0.698 | 0.939 |
| 3971 | MIR6737_3P | 169 | 0.4 | 0.95 | 0.626 | 0.939 |
| 3972 | MIR873_5P | 97 | 0.41 | 0.95 | 0.582 | 0.941 |
| 3973 | MIR3618 | 85 | 0.42 | 0.95 | 0.597 | 0.941 |
| 3974 | MIR4670_3P | 119 | 0.41 | 0.95 | 0.608 | 0.94 |
| 3975 | MIR1278 | 54 | 0.45 | 0.94 | 0.587 | 0.941 |
| 3976 | MIR5588_5P | 72 | 0.43 | 0.94 | 0.593 | 0.941 |
| 3977 | PBXIP1_TARGET_GENES | 178 | 0.4 | 0.94 | 0.631 | 0.941 |
| 3978 | MIR199A_3P_MIR199B_3P | 131 | 0.41 | 0.94 | 0.615 | 0.941 |
| 3979 | LINDGREN_BLADDER_CANCER_CLUSTER_1_UP | 112 | 0.41 | 0.94 | 0.613 | 0.941 |
| 3980 | SIX4_TARGET_GENES | 37 | 0.46 | 0.94 | 0.57 | 0.941 |
| 3981 | NFY_C | 209 | 0.39 | 0.94 | 0.64 | 0.941 |
| 3982 | COLDREN_GEFITINIB_RESISTANCE_DN | 185 | 0.4 | 0.94 | 0.639 | 0.941 |
| 3983 | AP2ALPHA_01 | 224 | 0.4 | 0.94 | 0.656 | 0.941 |
| 3984 | KEGG_LONG_TERM_POTENTIATION | 58 | 0.44 | 0.94 | 0.586 | 0.941 |
| 3985 | DELACROIX_RAR_BOUND_ES | 369 | 0.39 | 0.94 | 0.664 | 0.941 |
| 3986 | MIR215_5P | 35 | 0.47 | 0.94 | 0.565 | 0.941 |
| 3987 | STAT1_02 | 225 | 0.4 | 0.94 | 0.628 | 0.941 |
| 3988 | PID_FANCONI_PATHWAY | 47 | 0.45 | 0.94 | 0.593 | 0.941 |
| 3989 | MIR6895_5P | 43 | 0.46 | 0.94 | 0.563 | 0.942 |
| 3990 | MIR3152_5P | 51 | 0.44 | 0.94 | 0.578 | 0.942 |

| | | | | | | |
|---|---|---|---|---|---|---|
| 3991 | DUTERTRE_ESTRADIOL_RESPONSE_6HR_UP | 205 | 0.4 | 0.94 | 0.631 | 0.942 |
| 3992 | MIR6715B_3P | 114 | 0.41 | 0.94 | 0.593 | 0.942 |
| 3993 | MIR5571_3P | 36 | 0.47 | 0.94 | 0.57 | 0.942 |
| 3994 | CDPCR3HD_01 | 199 | 0.4 | 0.94 | 0.637 | 0.942 |
| 3995 | KEGG_PPAR_SIGNALING_PATHWAY | 40 | 0.45 | 0.94 | 0.569 | 0.942 |
| 3996 | FXR_Q3 | 84 | 0.42 | 0.94 | 0.585 | 0.942 |
| 3997 | MARTORIATI_MDM4_TARGETS_FETAL_LIVER_DN | 451 | 0.39 | 0.94 | 0.704 | 0.943 |
| 3998 | MIR4284 | 95 | 0.42 | 0.94 | 0.618 | 0.944 |
| 3999 | BLANCO_MELO_HUMAN_PARAINFLUENZA_VIRUS_ | 86 | 0.42 | 0.94 | 0.571 | 0.944 |
| 4000 | MIR148A_5P | 98 | 0.41 | 0.94 | 0.594 | 0.944 |
| 4001 | MIR548AZ_5P | 370 | 0.39 | 0.94 | 0.68 | 0.944 |
| 4002 | MIR4640_5P | 121 | 0.41 | 0.94 | 0.614 | 0.944 |
| 4003 | MIR6832_5P | 61 | 0.44 | 0.94 | 0.606 | 0.945 |
| 4004 | KORKOLA_TERATOMA | 32 | 0.47 | 0.94 | 0.581 | 0.944 |
| 4005 | MIR7151_5P | 40 | 0.45 | 0.94 | 0.573 | 0.945 |
| 4006 | MIR1910_3P | 160 | 0.4 | 0.94 | 0.635 | 0.945 |
| 4007 | GR_Q6 | 198 | 0.39 | 0.94 | 0.645 | 0.945 |
| 4008 | MIR222_5P | 155 | 0.4 | 0.94 | 0.616 | 0.945 |
| 4009 | STARK_HYPPOCAMPUS_22Q11_DELETION_UP | 49 | 0.45 | 0.94 | 0.567 | 0.945 |
| 4010 | HADDAD_T_LYMPHOCYTE_AND_NK_PROGENITOR | 65 | 0.43 | 0.94 | 0.588 | 0.944 |
| 4011 | MIR7157_3P | 170 | 0.4 | 0.94 | 0.656 | 0.944 |
| 4012 | WP_NRF2ARE_REGULATION | 22 | 0.5 | 0.94 | 0.571 | 0.945 |
| 4013 | BOHN_PRIMARY_IMMUNODEFICIENCY_SYNDROM_ | 45 | 0.45 | 0.94 | 0.59 | 0.945 |
| 4014 | MIR1913 | 127 | 0.4 | 0.94 | 0.629 | 0.945 |
| 4015 | WP_INTERLEUKIN1_IL1_STRUCTURAL_PATHWAY | 45 | 0.46 | 0.94 | 0.586 | 0.945 |
| 4016 | GTGGGTGK_UNKNOWN | 228 | 0.39 | 0.94 | 0.656 | 0.945 |
| 4017 | MIR7159_3P | 186 | 0.4 | 0.94 | 0.633 | 0.945 |
| 4018 | MIR4799_3P | 70 | 0.43 | 0.94 | 0.585 | 0.945 |
| 4019 | MIR4794 | 90 | 0.42 | 0.94 | 0.613 | 0.945 |
| 4020 | BUCKANOVICH_T_LYMPHOCYTE_HOMING_ON_TUI | 17 | 0.52 | 0.94 | 0.599 | 0.945 |
| 4021 | REACTOME_FCERI_MEDIATED_NF_KB_ACTIVATIO | 73 | 0.43 | 0.94 | 0.614 | 0.945 |
| 4022 | FOXC1_TARGET_GENES | 32 | 0.47 | 0.94 | 0.581 | 0.945 |
| 4023 | HNF4_01_B | 206 | 0.4 | 0.94 | 0.662 | 0.945 |

| | | | | | | |
|---|---|---|---|---|---|---|
| 4024 | PAX2_02 | 200 | 0.39 | 0.94 | 0.643 | 0.945 |
| 4025 | RODRIGUES_THYROID_CARCINOMA_ANAPLASTIC_ | 460 | 0.39 | 0.94 | 0.714 | 0.945 |
| 4026 | SHEN_SMARCA2_TARGETS_DN | 153 | 0.4 | 0.94 | 0.627 | 0.945 |
| 4027 | SHETH_LIVER_CANCER_VS_TXNIP_LOSS_PAM6 | 46 | 0.45 | 0.94 | 0.576 | 0.945 |
| 4028 | MIR382_5P | 49 | 0.45 | 0.94 | 0.593 | 0.945 |
| 4029 | CAIRO_LIVER_DEVELOPMENT_UP | 126 | 0.4 | 0.94 | 0.614 | 0.945 |
| 4030 | CDX2_Q5 | 201 | 0.39 | 0.94 | 0.647 | 0.945 |
| 4031 | SANSOM_APC_TARGETS | 168 | 0.39 | 0.94 | 0.634 | 0.945 |
| 4032 | FRIDMAN_IMMORTALIZATION_DN | 25 | 0.49 | 0.94 | 0.588 | 0.945 |
| 4033 | BENPORATH_PROLIFERATION | 144 | 0.4 | 0.94 | 0.629 | 0.946 |
| 4034 | MIR548AO_5P_MIR548AX | 145 | 0.4 | 0.94 | 0.638 | 0.946 |
| 4035 | QI_PLASMACYTOMA_DN | 80 | 0.42 | 0.94 | 0.587 | 0.946 |
| 4036 | NKX3A_01 | 176 | 0.4 | 0.94 | 0.614 | 0.945 |
| 4037 | BIOCARTA_PTDINS_PATHWAY | 21 | 0.5 | 0.94 | 0.59 | 0.947 |
| 4038 | MIR153_3P | 295 | 0.39 | 0.94 | 0.67 | 0.946 |
| 4039 | MIR8064 | 74 | 0.43 | 0.94 | 0.601 | 0.946 |
| 4040 | MIR4429 | 282 | 0.39 | 0.94 | 0.687 | 0.947 |
| 4041 | MIR298 | 151 | 0.4 | 0.94 | 0.632 | 0.946 |
| 4042 | GTGCCAT_MIR183 | 163 | 0.4 | 0.94 | 0.632 | 0.947 |
| 4043 | COBLL1_TARGET_GENES | 122 | 0.41 | 0.94 | 0.596 | 0.947 |
| 4044 | YTAATTAA_LHX3_01 | 148 | 0.4 | 0.94 | 0.63 | 0.947 |
| 4045 | MIR4774_5P | 79 | 0.42 | 0.94 | 0.612 | 0.948 |
| 4046 | REACTOME_SYNTHESIS_OF_PIPS_AT_THE_PLASI | 48 | 0.45 | 0.94 | 0.587 | 0.948 |
| 4047 | REACTOME_ION_TRANSPORT_BY_P_TYPE_ATPAS | 40 | 0.46 | 0.94 | 0.589 | 0.948 |
| 4048 | MIR5088_3P | 107 | 0.41 | 0.94 | 0.622 | 0.949 |
| 4049 | MIR6774_5P | 41 | 0.46 | 0.94 | 0.591 | 0.948 |
| 4050 | CTCNANGTGNY_UNKNOWN | 77 | 0.43 | 0.94 | 0.602 | 0.949 |
| 4051 | MIR3160_5P | 175 | 0.4 | 0.94 | 0.635 | 0.949 |
| 4052 | MIR3116 | 79 | 0.42 | 0.94 | 0.598 | 0.949 |
| 4053 | REACTOME_EXTENSION_OF_TELOMERES | 51 | 0.44 | 0.94 | 0.601 | 0.949 |
| 4054 | MIR6504_3P | 323 | 0.39 | 0.94 | 0.682 | 0.949 |
| 4055 | PETRETTO_CARDIAC_HYPERTROPHY | 25 | 0.49 | 0.94 | 0.573 | 0.949 |
| 4056 | CDC73_TARGET_GENES | 443 | 0.39 | 0.94 | 0.701 | 0.95 |

| | | | | | | |
|---|---|---|---|---|---|---|
| 4057 | MIR1827 | 321 | 0.38 | 0.94 | 0.688 | 0.95 |
| 4058 | MYB_Q6 | 203 | 0.39 | 0.94 | 0.636 | 0.949 |
| 4059 | AATGTGA_MIR23A_MIR23B | 364 | 0.38 | 0.94 | 0.699 | 0.95 |
| 4060 | MIR373_3P | 288 | 0.39 | 0.94 | 0.691 | 0.95 |
| 4061 | MIR664A_5P | 69 | 0.42 | 0.93 | 0.594 | 0.95 |
| 4062 | ZNF555_TARGET_GENES | 35 | 0.46 | 0.93 | 0.586 | 0.95 |
| 4063 | GRADE_COLON_AND_RECTAL_CANCER_DN | 79 | 0.42 | 0.93 | 0.62 | 0.95 |
| 4064 | REACTOME_HEME_SIGNALING | 40 | 0.46 | 0.93 | 0.598 | 0.95 |
| 4065 | MCBRYAN_PUBERTAL_BREAST_3_4WK_DN | 28 | 0.49 | 0.93 | 0.57 | 0.95 |
| 4066 | RIZKI_TUMOR_INVASIVENESS_2D_DN | 51 | 0.44 | 0.93 | 0.598 | 0.95 |
| 4067 | GTGCAAT_MIR25_MIR32_MIR92_MIR363_MIR367 | 270 | 0.39 | 0.93 | 0.671 | 0.95 |
| 4068 | MIR4491 | 58 | 0.43 | 0.93 | 0.594 | 0.95 |
| 4069 | DACOSTA_UV_RESPONSE_VIA_ERCC3_TTD_UP | 57 | 0.43 | 0.93 | 0.597 | 0.951 |
| 4070 | REACTOME_FOXO_MEDIATED_TRANSCRIPTION_C | 15 | 0.53 | 0.93 | 0.586 | 0.951 |
| 4071 | MEISSNER_NPC_HCP_WITH_H3K4ME2 | 383 | 0.38 | 0.93 | 0.716 | 0.951 |
| 4072 | MIR6878_5P | 181 | 0.4 | 0.93 | 0.626 | 0.951 |
| 4073 | EGR_Q6 | 247 | 0.39 | 0.93 | 0.661 | 0.95 |
| 4074 | BENPORATH_PRC2_TARGETS | 372 | 0.38 | 0.93 | 0.702 | 0.951 |
| 4075 | MIR320A_3P_MIR320B_MIR320C_MIR320D | 283 | 0.39 | 0.93 | 0.689 | 0.951 |
| 4076 | MIR5193 | 135 | 0.4 | 0.93 | 0.626 | 0.951 |
| 4077 | WONG_EMBRYONIC_STEM_CELL_CORE | 328 | 0.38 | 0.93 | 0.703 | 0.953 |
| 4078 | WP_ATM_SIGNALING_PATHWAY | 40 | 0.45 | 0.93 | 0.596 | 0.953 |
| 4079 | MIR4768_5P | 274 | 0.39 | 0.93 | 0.69 | 0.953 |
| 4080 | MIR661 | 134 | 0.4 | 0.93 | 0.655 | 0.954 |
| 4081 | T3R_Q6 | 196 | 0.39 | 0.93 | 0.662 | 0.955 |
| 4082 | ZNF669_TARGET_GENES | 111 | 0.4 | 0.93 | 0.61 | 0.955 |
| 4083 | KRAS.DF.V1_DN | 137 | 0.4 | 0.93 | 0.633 | 0.956 |
| 4084 | MIR659_3P | 189 | 0.39 | 0.93 | 0.647 | 0.956 |
| 4085 | MIR6787_5P | 22 | 0.5 | 0.93 | 0.586 | 0.956 |
| 4086 | FISCHER_G1_S_CELL_CYCLE | 180 | 0.39 | 0.93 | 0.645 | 0.956 |
| 4087 | RODRIGUES_THYROID_CARCINOMA_UP | 15 | 0.53 | 0.93 | 0.616 | 0.956 |
| 4088 | MIR206 | 345 | 0.38 | 0.93 | 0.695 | 0.956 |
| 4089 | MIR3190_3P | 95 | 0.41 | 0.93 | 0.617 | 0.956 |

| | | | | | | |
|---|---|---|---|---|---|---|
| 4090 | MEISSNER_NPC_HCP_WITH_H3K4ME3_AND_H3K2 | 114 | 0.4 | 0.93 | 0.619 | 0.956 |
| 4091 | ARID3B_TARGET_GENES | 70 | 0.42 | 0.93 | 0.599 | 0.956 |
| 4092 | REACTOME_SIGNALING_BY_NTRKS | 118 | 0.4 | 0.93 | 0.641 | 0.956 |
| 4093 | ALPHACP1_01 | 223 | 0.39 | 0.93 | 0.656 | 0.956 |
| 4094 | OCT_Q6 | 188 | 0.4 | 0.93 | 0.681 | 0.955 |
| 4095 | MIR6756_3P | 86 | 0.41 | 0.93 | 0.623 | 0.955 |
| 4096 | TFAM_TARGET_GENES | 21 | 0.5 | 0.93 | 0.594 | 0.955 |
| 4097 | WP_HIPPO_SIGNALING_REGULATION_PATHWAYS | 79 | 0.42 | 0.93 | 0.626 | 0.956 |
| 4098 | MIR1269A_MIR1269B | 42 | 0.45 | 0.93 | 0.581 | 0.956 |
| 4099 | MURAKAMI_UV_RESPONSE_6HR_UP | 29 | 0.47 | 0.93 | 0.614 | 0.956 |
| 4100 | AAGCACA_MIR218 | 360 | 0.38 | 0.93 | 0.712 | 0.956 |
| 4101 | MIR9_5P | 421 | 0.38 | 0.93 | 0.719 | 0.956 |
| 4102 | ATAACCT_MIR154 | 56 | 0.44 | 0.93 | 0.61 | 0.956 |
| 4103 | AGTCAGC_MIR345 | 56 | 0.43 | 0.93 | 0.576 | 0.956 |
| 4104 | MIR7155_5P | 88 | 0.41 | 0.93 | 0.624 | 0.956 |
| 4105 | MIR6764_5P | 192 | 0.39 | 0.93 | 0.658 | 0.956 |
| 4106 | WTGAAAT_UNKNOWN | 450 | 0.38 | 0.93 | 0.735 | 0.956 |
| 4107 | MIR6883_3P | 52 | 0.44 | 0.93 | 0.608 | 0.957 |
| 4108 | MIR590_5P | 106 | 0.41 | 0.93 | 0.639 | 0.956 |
| 4109 | MIR3977 | 135 | 0.4 | 0.93 | 0.639 | 0.956 |
| 4110 | LEIN_PONS_MARKERS | 68 | 0.42 | 0.93 | 0.598 | 0.956 |
| 4111 | MIR5589_3P | 144 | 0.39 | 0.93 | 0.642 | 0.957 |
| 4112 | KIM_MYC_AMPLIFICATION_TARGETS_UP | 182 | 0.39 | 0.93 | 0.652 | 0.957 |
| 4113 | FREAC7_01 | 144 | 0.39 | 0.93 | 0.646 | 0.957 |
| 4114 | BOYAULT_LIVER_CANCER_SUBCLASS_G2 | 25 | 0.48 | 0.93 | 0.583 | 0.958 |
| 4115 | MEIS1_01 | 201 | 0.39 | 0.93 | 0.663 | 0.957 |
| 4116 | MIR6811_3P | 48 | 0.45 | 0.93 | 0.612 | 0.957 |
| 4117 | REACTOME_SIGNALING_BY_WNT_IN_CANCER | 30 | 0.48 | 0.93 | 0.605 | 0.957 |
| 4118 | REACTOME_PTEN_REGULATION | 133 | 0.4 | 0.93 | 0.643 | 0.958 |
| 4119 | REACTOME_SUMOYLATION_OF_DNA_DAMAGE_RE | 76 | 0.42 | 0.93 | 0.633 | 0.958 |
| 4120 | REACTOME_THE_PHOTOTRANSDUCTION_CASCAD | 21 | 0.49 | 0.93 | 0.6 | 0.958 |
| 4121 | KAMMINGA_SENESCENCE | 35 | 0.46 | 0.93 | 0.595 | 0.958 |
| 4122 | MIR3140_3P | 205 | 0.39 | 0.93 | 0.657 | 0.958 |

| | | | | | | |
|---|---|---|---|---|---|---|
| 4123 | MIR3184_3P | 67 | 0.43 | 0.93 | 0.619 | 0.958 |
| 4124 | NIKOLSKY_BREAST_CANCER_8Q12_Q22_AMPLIC( | 106 | 0.4 | 0.93 | 0.64 | 0.958 |
| 4125 | COUP_DR1_Q6 | 211 | 0.39 | 0.93 | 0.676 | 0.958 |
| 4126 | MIR612 | 82 | 0.42 | 0.93 | 0.608 | 0.958 |
| 4127 | MIR6080 | 42 | 0.45 | 0.93 | 0.613 | 0.958 |
| 4128 | WP_PATHWAYS_AFFECTED_IN_ADENOID_CYSTIC_ | 61 | 0.42 | 0.93 | 0.621 | 0.958 |
| 4129 | MIR3975 | 49 | 0.44 | 0.93 | 0.605 | 0.958 |
| 4130 | MIR6885_3P | 334 | 0.38 | 0.93 | 0.718 | 0.958 |
| 4131 | MIR7848_3P | 24 | 0.49 | 0.93 | 0.611 | 0.959 |
| 4132 | REACTOME_TRANSCRIPTIONAL_REGULATION_BY_ | 353 | 0.38 | 0.93 | 0.717 | 0.959 |
| 4133 | MIR4492 | 366 | 0.38 | 0.93 | 0.706 | 0.959 |
| 4134 | TSHZ1_TARGET_GENES | 154 | 0.4 | 0.93 | 0.658 | 0.959 |
| 4135 | MIR3179 | 107 | 0.4 | 0.93 | 0.641 | 0.959 |
| 4136 | SIG_INSULIN_RECEPTOR_PATHWAY_IN_CARDIAC_ | 47 | 0.44 | 0.93 | 0.611 | 0.959 |
| 4137 | LIN_APC_TARGETS | 66 | 0.43 | 0.93 | 0.615 | 0.959 |
| 4138 | MIR150_3P | 107 | 0.41 | 0.93 | 0.633 | 0.96 |
| 4139 | MIR5590_5P | 98 | 0.41 | 0.92 | 0.632 | 0.96 |
| 4140 | HFH1_01 | 195 | 0.39 | 0.92 | 0.669 | 0.96 |
| 4141 | WP_RETINOBLASTOMA_GENE_IN_CANCER | 87 | 0.41 | 0.92 | 0.61 | 0.96 |
| 4142 | CAGGGTC_MIR504 | 74 | 0.42 | 0.92 | 0.63 | 0.96 |
| 4143 | VALK_AML_WITH_FLT3_ITD | 32 | 0.47 | 0.92 | 0.594 | 0.96 |
| 4144 | S8_01 | 183 | 0.39 | 0.92 | 0.662 | 0.96 |
| 4145 | REACTOME_SEMA4D_INDUCED_CELL_MIGRATION | 18 | 0.51 | 0.92 | 0.603 | 0.96 |
| 4146 | SOX3_TARGET_GENES | 421 | 0.38 | 0.92 | 0.742 | 0.96 |
| 4147 | MIR6809_3P | 459 | 0.38 | 0.92 | 0.758 | 0.96 |
| 4148 | MIR1911_3P | 39 | 0.45 | 0.92 | 0.608 | 0.96 |
| 4149 | TCCATTKW_UNKNOWN | 204 | 0.39 | 0.92 | 0.7 | 0.96 |
| 4150 | MIR3137 | 49 | 0.43 | 0.92 | 0.581 | 0.96 |
| 4151 | MIR2355_5P | 118 | 0.4 | 0.92 | 0.637 | 0.961 |
| 4152 | GENTILE_UV_RESPONSE_CLUSTER_D4 | 53 | 0.44 | 0.92 | 0.601 | 0.961 |
| 4153 | MIR520E_5P | 144 | 0.4 | 0.92 | 0.647 | 0.961 |
| 4154 | MIR4777_3P | 58 | 0.42 | 0.92 | 0.614 | 0.961 |
| 4155 | REACTOME_FOXO_MEDIATED_TRANSCRIPTION_C | 18 | 0.52 | 0.92 | 0.622 | 0.961 |

| | | | | | | |
|---|---|---|---|---|---|---|
| 4156 | REACTOME_CELL_CYCLE_CHECKPOINTS | 270 | 0.38 | 0.92 | 0.706 | 0.961 |
| 4157 | REACTOME_POTENTIAL_THERAPEUTICS_FOR_SA | 72 | 0.42 | 0.92 | 0.613 | 0.961 |
| 4158 | BROWNE_HCMV_INFECTION_20HR_DN | 82 | 0.41 | 0.92 | 0.637 | 0.961 |
| 4159 | MIR3155A_MIR3155B | 71 | 0.42 | 0.92 | 0.632 | 0.961 |
| 4160 | SRF_Q4 | 184 | 0.39 | 0.92 | 0.658 | 0.961 |
| 4161 | MIR4711_5P | 89 | 0.41 | 0.92 | 0.642 | 0.961 |
| 4162 | VERHAAK_GLIOBLASTOMA_CLASSICAL | 138 | 0.4 | 0.92 | 0.664 | 0.961 |
| 4163 | MIR11181_3P | 169 | 0.39 | 0.92 | 0.677 | 0.961 |
| 4164 | MIR378A_5P | 117 | 0.4 | 0.92 | 0.644 | 0.961 |
| 4165 | CROONQUIST_NRAS_SIGNALING_DN | 70 | 0.42 | 0.92 | 0.622 | 0.961 |
| 4166 | MIR6837_3P | 86 | 0.41 | 0.92 | 0.64 | 0.961 |
| 4167 | MIR1183 | 112 | 0.4 | 0.92 | 0.637 | 0.961 |
| 4168 | MIR6893_5P | 159 | 0.39 | 0.92 | 0.659 | 0.961 |
| 4169 | GGCAGTG_MIR3243P | 83 | 0.41 | 0.92 | 0.626 | 0.961 |
| 4170 | RIZ_ERYTHROID_DIFFERENTIATION_12HR | 30 | 0.47 | 0.92 | 0.617 | 0.961 |
| 4171 | CRX_NRL_DN.V1_DN | 105 | 0.4 | 0.92 | 0.634 | 0.962 |
| 4172 | MIR625_5P | 173 | 0.39 | 0.92 | 0.662 | 0.962 |
| 4173 | TCF1P_Q6 | 223 | 0.39 | 0.92 | 0.699 | 0.962 |
| 4174 | WP_SEROTONIN_RECEPTOR_467_AND_NR3C_SIG | 17 | 0.51 | 0.92 | 0.601 | 0.962 |
| 4175 | HERNANDEZ_MITOTIC_ARREST_BY_DOCETAXEL_ | 18 | 0.51 | 0.92 | 0.623 | 0.963 |
| 4176 | GGTGTGT_MIR329 | 103 | 0.4 | 0.92 | 0.649 | 0.963 |
| 4177 | HFH8_01 | 167 | 0.39 | 0.92 | 0.668 | 0.962 |
| 4178 | BHAT_ESR1_TARGETS_NOT_VIA_AKT1_DN | 79 | 0.41 | 0.92 | 0.632 | 0.963 |
| 4179 | MIR593_5P | 146 | 0.4 | 0.92 | 0.652 | 0.963 |
| 4180 | MIR411_5P | 42 | 0.44 | 0.92 | 0.614 | 0.963 |
| 4181 | MIR6852_5P | 168 | 0.39 | 0.92 | 0.671 | 0.963 |
| 4182 | RIEGE_DELTANP63_DIRECT_TARGETS_UP | 119 | 0.4 | 0.92 | 0.651 | 0.963 |
| 4183 | MIR500B_5P | 48 | 0.44 | 0.92 | 0.61 | 0.964 |
| 4184 | MIR140_5P | 82 | 0.41 | 0.92 | 0.648 | 0.963 |
| 4185 | SRF_C | 168 | 0.39 | 0.92 | 0.673 | 0.964 |
| 4186 | MATZUK_IMPLANTATION_AND_UTERINE | 15 | 0.53 | 0.92 | 0.622 | 0.964 |
| 4187 | MIR6727_3P | 100 | 0.41 | 0.92 | 0.636 | 0.964 |
| 4188 | MIR6861_5P | 50 | 0.43 | 0.92 | 0.639 | 0.964 |

| | | | | | | |
|---|---|---|---|---|---|---|
| 4189 | MIR29B_1_5P | 139 | 0.39 | 0.92 | 0.644 | 0.964 |
| 4190 | GRYDER_PAX3FOXO1_ENHANCERS_KO_DOWN | 394 | 0.38 | 0.92 | 0.739 | 0.964 |
| 4191 | REACTOME_SIGNALING_BY_FGFR2_IN_DISEASE | 31 | 0.46 | 0.92 | 0.601 | 0.964 |
| 4192 | DMRT1_TARGET_GENES | 359 | 0.38 | 0.92 | 0.74 | 0.964 |
| 4193 | MIR10396B_5P | 17 | 0.51 | 0.92 | 0.61 | 0.964 |
| 4194 | MIR4456 | 117 | 0.4 | 0.92 | 0.673 | 0.964 |
| 4195 | SAKAI_TUMOR_INFILTRATING_MONOCYTES_DN | 72 | 0.42 | 0.92 | 0.639 | 0.964 |
| 4196 | REACTOME_RHO_GTPASE_CYCLE | 405 | 0.38 | 0.92 | 0.771 | 0.964 |
| 4197 | TFIII_Q6 | 168 | 0.39 | 0.92 | 0.67 | 0.964 |
| 4198 | MIR4278 | 26 | 0.48 | 0.92 | 0.621 | 0.964 |
| 4199 | GATAAGR_GATA_C | 205 | 0.38 | 0.92 | 0.694 | 0.964 |
| 4200 | REACTOME_ABC_TRANSPORTER_DISORDERS | 69 | 0.42 | 0.92 | 0.628 | 0.964 |
| 4201 | BIOCARTA_CHEMICAL_PATHWAY | 18 | 0.5 | 0.92 | 0.614 | 0.964 |
| 4202 | MIR548S | 111 | 0.4 | 0.92 | 0.654 | 0.965 |
| 4203 | TTCYRGAA_UNKNOWN | 261 | 0.38 | 0.92 | 0.708 | 0.964 |
| 4204 | CHARAFE_BREAST_CANCER_LUMINAL_VS_MESEI | 401 | 0.38 | 0.92 | 0.758 | 0.964 |
| 4205 | HAND1_TARGET_GENES | 388 | 0.38 | 0.92 | 0.759 | 0.965 |
| 4206 | REACTOME_DOWNSTREAM_SIGNALING_OF_ACTIV | 18 | 0.51 | 0.92 | 0.598 | 0.965 |
| 4207 | APPIERTO_RESPONSE_TO_FENRETINIDE_UP | 33 | 0.46 | 0.92 | 0.608 | 0.965 |
| 4208 | AGGGCAG_MIR18A | 125 | 0.39 | 0.92 | 0.649 | 0.965 |
| 4209 | TGACGTCA_ATF3_Q6 | 202 | 0.39 | 0.92 | 0.701 | 0.965 |
| 4210 | MEBARKI_HCC_PROGENITOR_WNT_UP_BLOCKED | 80 | 0.41 | 0.92 | 0.64 | 0.965 |
| 4211 | BROWNE_HCMV_INFECTION_4HR_DN | 215 | 0.39 | 0.92 | 0.721 | 0.965 |
| 4212 | SMID_BREAST_CANCER_RELAPSE_IN_LUNG_DN | 16 | 0.51 | 0.92 | 0.617 | 0.965 |
| 4213 | PRDM5_TARGET_GENES | 69 | 0.42 | 0.92 | 0.639 | 0.965 |
| 4214 | MIR4280 | 40 | 0.45 | 0.92 | 0.612 | 0.965 |
| 4215 | MYOD_Q6_01 | 218 | 0.38 | 0.92 | 0.721 | 0.965 |
| 4216 | MIR885_3P | 25 | 0.48 | 0.92 | 0.61 | 0.965 |
| 4217 | PPAR_DR1_Q2 | 212 | 0.39 | 0.92 | 0.68 | 0.965 |
| 4218 | YATTNATC_UNKNOWN | 264 | 0.38 | 0.92 | 0.706 | 0.965 |
| 4219 | HIF1_Q5 | 223 | 0.38 | 0.92 | 0.681 | 0.965 |
| 4220 | MIR4439 | 62 | 0.42 | 0.92 | 0.628 | 0.965 |
| 4221 | MIR12119 | 153 | 0.39 | 0.92 | 0.68 | 0.965 |

| | | | | | | |
|---|---|---|---|---|---|---|
| 4222 | MIR455_3P | 84 | 0.41 | 0.92 | 0.66 | 0.965 |
| 4223 | MIR1825 | 145 | 0.39 | 0.92 | 0.67 | 0.965 |
| 4224 | RHODES_CANCER_META_SIGNATURE | 64 | 0.42 | 0.92 | 0.624 | 0.965 |
| 4225 | REACTOME_PLATELET_AGGREGATION_PLUG_FO | 26 | 0.47 | 0.92 | 0.608 | 0.965 |
| 4226 | REACTOME_PROTEIN_FOLDING | 81 | 0.41 | 0.92 | 0.624 | 0.965 |
| 4227 | KEGG_AMYOTROPHIC_LATERAL_SCLEROSIS_ALS | 47 | 0.44 | 0.92 | 0.619 | 0.965 |
| 4228 | BIOCARTA_TOLL_PATHWAY | 23 | 0.49 | 0.92 | 0.619 | 0.965 |
| 4229 | MIR6796_5P | 71 | 0.42 | 0.92 | 0.63 | 0.964 |
| 4230 | MIR3138 | 59 | 0.43 | 0.92 | 0.607 | 0.964 |
| 4231 | DOANE_RESPONSE_TO_ANDROGEN_UP | 144 | 0.39 | 0.92 | 0.684 | 0.964 |
| 4232 | WP_JOUBERT_SYNDROME | 71 | 0.41 | 0.92 | 0.628 | 0.964 |
| 4233 | ACAACTT_MIR382 | 65 | 0.42 | 0.92 | 0.627 | 0.965 |
| 4234 | BIOCARTA_FCER1_PATHWAY | 33 | 0.46 | 0.92 | 0.612 | 0.965 |
| 4235 | XPO1_TARGET_GENES | 16 | 0.51 | 0.91 | 0.618 | 0.965 |
| 4236 | MIR1296_3P | 27 | 0.47 | 0.91 | 0.614 | 0.965 |
| 4237 | MIR550A_3P | 59 | 0.42 | 0.91 | 0.631 | 0.965 |
| 4238 | MIR6833_3P | 337 | 0.37 | 0.91 | 0.738 | 0.965 |
| 4239 | ATTCTTT_MIR186 | 249 | 0.38 | 0.91 | 0.714 | 0.965 |
| 4240 | REACTOME_HS_GAG_BIOSYNTHESIS | 24 | 0.48 | 0.91 | 0.64 | 0.965 |
| 4241 | MIR6891_5P | 102 | 0.41 | 0.91 | 0.652 | 0.966 |
| 4242 | E4BP4_01 | 192 | 0.39 | 0.91 | 0.699 | 0.965 |
| 4243 | SENESE_HDAC1_TARGETS_DN | 220 | 0.38 | 0.91 | 0.708 | 0.966 |
| 4244 | MIKKELSEN_MCV6_ICP_WITH_H3K27ME3 | 34 | 0.46 | 0.91 | 0.621 | 0.966 |
| 4245 | DR4_Q2 | 224 | 0.38 | 0.91 | 0.706 | 0.966 |
| 4246 | MIR647 | 68 | 0.42 | 0.91 | 0.633 | 0.966 |
| 4247 | MIR145_5P | 262 | 0.38 | 0.91 | 0.739 | 0.966 |
| 4248 | NAKAMURA_METASTASIS | 39 | 0.45 | 0.91 | 0.633 | 0.966 |
| 4249 | MIR1288_3P | 39 | 0.45 | 0.91 | 0.627 | 0.966 |
| 4250 | PAX5_01 | 137 | 0.39 | 0.91 | 0.677 | 0.966 |
| 4251 | MIR663A | 15 | 0.52 | 0.91 | 0.633 | 0.966 |
| 4252 | NERF_Q2 | 214 | 0.38 | 0.91 | 0.711 | 0.967 |
| 4253 | MIR518E_5P_MIR519A_5P_MIR519B_5P_MIR519C_ | 74 | 0.42 | 0.91 | 0.636 | 0.967 |
| 4254 | MIR3689A_5P_MIR3689B_5P_MIR3689E_MIR3689F | 96 | 0.41 | 0.91 | 0.662 | 0.967 |

| | | | | | | |
|---|---|---|---|---|---|---|
| 4255 | MIR4697_3P | 99 | 0.4 | 0.91 | 0.661 | 0.967 |
| 4256 | GTACAGG_MIR486 | 55 | 0.43 | 0.91 | 0.644 | 0.966 |
| 4257 | MIR1303 | 167 | 0.39 | 0.91 | 0.696 | 0.967 |
| 4258 | MOHANKUMAR_HOXA1_TARGETS_UP | 383 | 0.37 | 0.91 | 0.759 | 0.967 |
| 4259 | CCAATNNSNNNGCG_UNKNOWN | 55 | 0.43 | 0.91 | 0.62 | 0.966 |
| 4260 | IVANOVA_HEMATOPOIESIS_STEM_CELL_AND_PR | 462 | 0.37 | 0.91 | 0.766 | 0.967 |
| 4261 | MIR331_5P | 66 | 0.42 | 0.91 | 0.638 | 0.967 |
| 4262 | MIR6826_3P | 88 | 0.41 | 0.91 | 0.664 | 0.967 |
| 4263 | MIR96_3P | 58 | 0.42 | 0.91 | 0.637 | 0.968 |
| 4264 | WP_NEURAL_CREST_DIFFERENTIATION | 69 | 0.41 | 0.91 | 0.632 | 0.967 |
| 4265 | MIR6845_3P | 118 | 0.4 | 0.91 | 0.677 | 0.968 |
| 4266 | GENTILE_UV_RESPONSE_CLUSTER_D1 | 17 | 0.5 | 0.91 | 0.614 | 0.968 |
| 4267 | MIR548M | 165 | 0.39 | 0.91 | 0.683 | 0.967 |
| 4268 | MIR4715_5P | 35 | 0.45 | 0.91 | 0.631 | 0.967 |
| 4269 | MIR875_5P | 44 | 0.43 | 0.91 | 0.642 | 0.967 |
| 4270 | MEF2_Q6_01 | 175 | 0.39 | 0.91 | 0.693 | 0.968 |
| 4271 | WAMUNYOKOLI_OVARIAN_CANCER_LMP_DN | 166 | 0.39 | 0.91 | 0.699 | 0.967 |
| 4272 | MIR4720_5P | 71 | 0.41 | 0.91 | 0.641 | 0.967 |
| 4273 | MIR505_3P | 97 | 0.4 | 0.91 | 0.68 | 0.967 |
| 4274 | MODY_HIPPOCAMPUS_PRENATAL | 43 | 0.44 | 0.91 | 0.65 | 0.967 |
| 4275 | BYSTROEM_CORRELATED_WITH_IL5_DN | 49 | 0.43 | 0.91 | 0.628 | 0.967 |
| 4276 | TGATTTRY_GFI1_01 | 205 | 0.38 | 0.91 | 0.705 | 0.967 |
| 4277 | MIR1343_5P | 104 | 0.4 | 0.91 | 0.665 | 0.967 |
| 4278 | MIR3127_3P | 89 | 0.4 | 0.91 | 0.669 | 0.967 |
| 4279 | REACTOME_SIGNALING_BY_TGFB_FAMILY_MEMB | 88 | 0.41 | 0.91 | 0.659 | 0.967 |
| 4280 | LEIN_CHOROID_PLEXUS_MARKERS | 63 | 0.42 | 0.91 | 0.648 | 0.967 |
| 4281 | REACTOME_CELL_DEATH_SIGNALLING_VIA_NRA( | 69 | 0.42 | 0.91 | 0.644 | 0.967 |
| 4282 | ZNF563_TARGET_GENES | 495 | 0.37 | 0.91 | 0.781 | 0.967 |
| 4283 | MIR8080 | 118 | 0.39 | 0.91 | 0.668 | 0.967 |
| 4284 | LMTK3_TARGET_GENES | 444 | 0.37 | 0.91 | 0.777 | 0.967 |
| 4285 | MOOTHA_PGC | 391 | 0.37 | 0.91 | 0.78 | 0.968 |
| 4286 | SCHAEFFER_PROSTATE_DEVELOPMENT_12HR_U | 76 | 0.41 | 0.91 | 0.655 | 0.968 |
| 4287 | FOURNIER_ACINAR_DEVELOPMENT_LATE_2 | 271 | 0.38 | 0.91 | 0.739 | 0.967 |

| | | | | | | |
|---|---|---|---|---|---|---|
| 4288 | MIR6749_3P | 92 | 0.4 | 0.91 | 0.67 | 0.968 |
| 4289 | REACTOME_GLUCAGON_LIKE_PEPTIDE_1_GLP1_ | 30 | 0.46 | 0.91 | 0.607 | 0.968 |
| 4290 | MULLIGHAN_NPM1_MUTATED_SIGNATURE_1_UP | 212 | 0.38 | 0.91 | 0.719 | 0.968 |
| 4291 | KEGG_FATTY_ACID_METABOLISM | 33 | 0.46 | 0.91 | 0.618 | 0.968 |
| 4292 | JIANG_HYPOXIA_VIA_VHL | 19 | 0.48 | 0.91 | 0.61 | 0.968 |
| 4293 | REACTOME_EFFECTS_OF_PIP2_HYDROLYSIS | 20 | 0.49 | 0.91 | 0.622 | 0.968 |
| 4294 | LYF1_01 | 210 | 0.38 | 0.91 | 0.709 | 0.968 |
| 4295 | MIR1285_3P | 129 | 0.39 | 0.91 | 0.696 | 0.969 |
| 4296 | WENG_POR_TARGETS_LIVER_UP | 30 | 0.47 | 0.91 | 0.636 | 0.969 |
| 4297 | MIR544A | 193 | 0.38 | 0.91 | 0.699 | 0.969 |
| 4298 | OCT1_07 | 127 | 0.39 | 0.91 | 0.669 | 0.969 |
| 4299 | YAO_TEMPORAL_RESPONSE_TO_PROGESTERON | 38 | 0.44 | 0.91 | 0.623 | 0.969 |
| 4300 | REACTOME_ACTIVATED_TAK1_MEDIATES_P38_M/ | 22 | 0.48 | 0.91 | 0.631 | 0.969 |
| 4301 | MIR549A_3P | 69 | 0.42 | 0.91 | 0.631 | 0.97 |
| 4302 | MIR362_5P | 47 | 0.44 | 0.91 | 0.636 | 0.97 |
| 4303 | MIR33A_5P_MIR33B_5P | 97 | 0.41 | 0.91 | 0.665 | 0.969 |
| 4304 | TNCATNTCCYR_UNKNOWN | 117 | 0.39 | 0.91 | 0.663 | 0.97 |
| 4305 | ATGCAGT_MIR217 | 107 | 0.4 | 0.91 | 0.662 | 0.97 |
| 4306 | MIR940 | 159 | 0.39 | 0.91 | 0.697 | 0.97 |
| 4307 | OUILLETTE_CLL_13Q14_DELETION_UP | 68 | 0.42 | 0.91 | 0.647 | 0.97 |
| 4308 | MIKKELSEN_IPS_WITH_HCP_H3K27ME3 | 59 | 0.42 | 0.91 | 0.637 | 0.97 |
| 4309 | REACTOME_SEMA3A_PAK_DEPENDENT_AXON_RI | 16 | 0.51 | 0.91 | 0.642 | 0.971 |
| 4310 | MIR3153 | 196 | 0.38 | 0.91 | 0.729 | 0.972 |
| 4311 | AAGCCAT_MIR135A_MIR135B | 302 | 0.38 | 0.91 | 0.744 | 0.973 |
| 4312 | GRADE_COLON_VS_RECTAL_CANCER_DN | 46 | 0.43 | 0.91 | 0.634 | 0.973 |
| 4313 | MIR518D_5P_MIR518F_5P_MIR520C_5P_MIR526A_ | 77 | 0.41 | 0.91 | 0.654 | 0.973 |
| 4314 | VANOEVELEN_MYOGENESIS_SIN3A_TARGETS | 208 | 0.38 | 0.91 | 0.712 | 0.973 |
| 4315 | MIR4515 | 32 | 0.46 | 0.91 | 0.638 | 0.972 |
| 4316 | MAHAJAN_RESPONSE_TO_IL1A_DN | 58 | 0.42 | 0.91 | 0.634 | 0.972 |
| 4317 | USF_C | 245 | 0.38 | 0.91 | 0.725 | 0.972 |
| 4318 | KOHOUTEK_CCNT1_TARGETS | 42 | 0.44 | 0.91 | 0.64 | 0.973 |
| 4319 | MIR6715A_3P | 142 | 0.39 | 0.9 | 0.693 | 0.973 |
| 4320 | MIR597_5P | 62 | 0.42 | 0.9 | 0.654 | 0.973 |

| | | | | | | |
|---|---|---|---|---|---|---|
| 4321 | MIR92B_3P | 424 | 0.37 | 0.9 | 0.782 | 0.973 |
| 4322 | REACTOME_SYNTHESIS_OF_PC | 23 | 0.47 | 0.9 | 0.6 | 0.973 |
| 4323 | MIR320E | 66 | 0.41 | 0.9 | 0.633 | 0.973 |
| 4324 | MIR766_3P | 125 | 0.39 | 0.9 | 0.684 | 0.974 |
| 4325 | MIR3675_5P | 30 | 0.46 | 0.9 | 0.64 | 0.974 |
| 4326 | PID_ER_NONGENOMIC_PATHWAY | 37 | 0.44 | 0.9 | 0.644 | 0.974 |
| 4327 | YTCCCRNNAGGY_UNKNOWN | 66 | 0.41 | 0.9 | 0.661 | 0.973 |
| 4328 | MIR6887_5P | 112 | 0.4 | 0.9 | 0.694 | 0.974 |
| 4329 | ZID_01 | 225 | 0.38 | 0.9 | 0.718 | 0.974 |
| 4330 | CEBP_01 | 207 | 0.38 | 0.9 | 0.715 | 0.974 |
| 4331 | ATF_B | 157 | 0.38 | 0.9 | 0.706 | 0.974 |
| 4332 | KEGG_OOCYTE_MEIOSIS | 95 | 0.4 | 0.9 | 0.665 | 0.974 |
| 4333 | IWANAGA_CARCINOGENESIS_BY_KRAS_DN | 89 | 0.4 | 0.9 | 0.663 | 0.974 |
| 4334 | MIR942_3P | 93 | 0.4 | 0.9 | 0.668 | 0.974 |
| 4335 | MSX1_01 | 131 | 0.39 | 0.9 | 0.701 | 0.974 |
| 4336 | MIR4500 | 454 | 0.37 | 0.9 | 0.785 | 0.974 |
| 4337 | RRAGTTGT_UNKNOWN | 217 | 0.38 | 0.9 | 0.73 | 0.974 |
| 4338 | HSF1_01 | 217 | 0.37 | 0.9 | 0.717 | 0.974 |
| 4339 | MARTORIATI_MDM4_TARGETS_FETAL_LIVER_UP | 209 | 0.38 | 0.9 | 0.721 | 0.975 |
| 4340 | MIR5008_5P | 66 | 0.41 | 0.9 | 0.658 | 0.975 |
| 4341 | CAHOY_ASTROCYTIC | 67 | 0.41 | 0.9 | 0.652 | 0.975 |
| 4342 | PAX4_02 | 174 | 0.38 | 0.9 | 0.718 | 0.975 |
| 4343 | ZNF426_TARGET_GENES | 48 | 0.43 | 0.9 | 0.651 | 0.975 |
| 4344 | REACTOME_RHO_GTPASES_ACTIVATE_WASPS_A | 35 | 0.45 | 0.9 | 0.614 | 0.975 |
| 4345 | PID_PI3KCI_PATHWAY | 41 | 0.44 | 0.9 | 0.62 | 0.975 |
| 4346 | MIR3924 | 282 | 0.37 | 0.9 | 0.737 | 0.975 |
| 4347 | GGGCATT_MIR365 | 101 | 0.4 | 0.9 | 0.686 | 0.976 |
| 4348 | GGCKCATGS_UNKNOWN | 62 | 0.41 | 0.9 | 0.658 | 0.977 |
| 4349 | DAVICIONI_MOLECULAR_ARMS_VS_ERMS_UP | 306 | 0.37 | 0.9 | 0.772 | 0.977 |
| 4350 | MIR6860 | 86 | 0.39 | 0.9 | 0.664 | 0.976 |
| 4351 | REACTOME_CELLULAR_RESPONSE_TO_HYPOXIA | 72 | 0.41 | 0.9 | 0.671 | 0.977 |
| 4352 | MIR486_3P | 149 | 0.39 | 0.9 | 0.693 | 0.977 |
| 4353 | CREIGHTON_ENDOCRINE_THERAPY_RESISTANCE | 263 | 0.37 | 0.9 | 0.759 | 0.977 |

| | | | | | | |
|---|---|---|---|---|---|---|
| 4354 | CREIGHTON_AKT1_SIGNALING_VIA_MTOR_UP | 34 | 0.45 | 0.9 | 0.637 | 0.977 |
| 4355 | MIR6856_5P | 84 | 0.4 | 0.9 | 0.669 | 0.977 |
| 4356 | MIR1288_5P | 45 | 0.43 | 0.9 | 0.653 | 0.977 |
| 4357 | TERAO_AOX4_TARGETS_SKIN_DN | 15 | 0.51 | 0.9 | 0.622 | 0.977 |
| 4358 | LHX3_TARGET_GENES | 29 | 0.46 | 0.9 | 0.656 | 0.977 |
| 4359 | PRC2_EED_DN.V1_DN | 177 | 0.38 | 0.9 | 0.721 | 0.976 |
| 4360 | MIR5590_3P | 301 | 0.37 | 0.9 | 0.768 | 0.976 |
| 4361 | MIR32_5P | 427 | 0.37 | 0.9 | 0.805 | 0.976 |
| 4362 | ZNF486_TARGET_GENES | 136 | 0.38 | 0.9 | 0.69 | 0.976 |
| 4363 | FOXJ2_02 | 186 | 0.38 | 0.9 | 0.725 | 0.976 |
| 4364 | YAGI_AML_WITH_INV_16_TRANSLOCATION | 323 | 0.37 | 0.9 | 0.754 | 0.976 |
| 4365 | MIR520D_3P | 297 | 0.37 | 0.9 | 0.759 | 0.976 |
| 4366 | REACTOME_ION_HOMEOSTASIS | 40 | 0.43 | 0.9 | 0.637 | 0.976 |
| 4367 | SMAD_Q6 | 211 | 0.38 | 0.9 | 0.741 | 0.976 |
| 4368 | MIR424_3P | 29 | 0.46 | 0.9 | 0.653 | 0.976 |
| 4369 | MIR4476 | 193 | 0.38 | 0.9 | 0.749 | 0.976 |
| 4370 | YAO_TEMPORAL_RESPONSE_TO_PROGESTERON | 68 | 0.41 | 0.9 | 0.656 | 0.976 |
| 4371 | CDP_02 | 78 | 0.4 | 0.9 | 0.676 | 0.976 |
| 4372 | MIR6840_3P | 145 | 0.38 | 0.9 | 0.708 | 0.976 |
| 4373 | MIR7703 | 95 | 0.4 | 0.9 | 0.657 | 0.976 |
| 4374 | MIR1231 | 43 | 0.43 | 0.9 | 0.641 | 0.976 |
| 4375 | MIR92A_3P | 427 | 0.37 | 0.9 | 0.808 | 0.976 |
| 4376 | MIR6747_3P | 107 | 0.39 | 0.9 | 0.702 | 0.976 |
| 4377 | KEGG_PROGESTERONE_MEDIATED_OOCYTE_MA | 75 | 0.41 | 0.9 | 0.656 | 0.977 |
| 4378 | CHARAFE_BREAST_CANCER_LUMINAL_VS_BASAL | 341 | 0.37 | 0.9 | 0.784 | 0.977 |
| 4379 | DAZARD_UV_RESPONSE_CLUSTER_G4 | 16 | 0.5 | 0.9 | 0.66 | 0.977 |
| 4380 | MIR550B_3P | 52 | 0.43 | 0.9 | 0.645 | 0.977 |
| 4381 | SCHAEFFER_PROSTATE_DEVELOPMENT_12HR_D | 38 | 0.44 | 0.9 | 0.655 | 0.977 |
| 4382 | REACTOME_MAP2K_AND_MAPK_ACTIVATION | 33 | 0.45 | 0.9 | 0.646 | 0.978 |
| 4383 | MYB_Q3 | 213 | 0.38 | 0.9 | 0.748 | 0.978 |
| 4384 | MIR4524B_3P | 61 | 0.42 | 0.9 | 0.66 | 0.978 |
| 4385 | APPEL_IMATINIB_RESPONSE | 23 | 0.48 | 0.9 | 0.632 | 0.978 |
| 4386 | MIR7515 | 126 | 0.39 | 0.9 | 0.717 | 0.978 |

| | | | | | | |
|---|---|---|---|---|---|---|
| 4387 | SIG_BCR_SIGNALING_PATHWAY | 39 | 0.44 | 0.9 | 0.66 | 0.978 |
| 4388 | MIR627_5P | 35 | 0.44 | 0.9 | 0.646 | 0.978 |
| 4389 | KEGG_ALZHEIMERS_DISEASE | 147 | 0.38 | 0.9 | 0.728 | 0.978 |
| 4390 | MIR6076 | 47 | 0.43 | 0.9 | 0.668 | 0.978 |
| 4391 | LIM_MAMMARY_LUMINAL_MATURE_DN | 68 | 0.42 | 0.9 | 0.663 | 0.978 |
| 4392 | MIR6754_5P | 93 | 0.4 | 0.9 | 0.677 | 0.978 |
| 4393 | GCGSCMNTTT_UNKNOWN | 68 | 0.41 | 0.9 | 0.686 | 0.977 |
| 4394 | MIR29A_5P | 166 | 0.38 | 0.9 | 0.713 | 0.977 |
| 4395 | MIR299_5P | 122 | 0.39 | 0.9 | 0.699 | 0.978 |
| 4396 | MIR6868_3P | 221 | 0.37 | 0.9 | 0.741 | 0.978 |
| 4397 | MIR6791_3P | 70 | 0.41 | 0.9 | 0.689 | 0.978 |
| 4398 | MIR12125 | 88 | 0.4 | 0.9 | 0.691 | 0.978 |
| 4399 | MIR193B_5P | 52 | 0.43 | 0.9 | 0.658 | 0.978 |
| 4400 | P53_DECAMER_Q2 | 212 | 0.37 | 0.9 | 0.735 | 0.978 |
| 4401 | MIR3681_5P | 25 | 0.47 | 0.9 | 0.645 | 0.979 |
| 4402 | CTGYNNCTYTAA_UNKNOWN | 64 | 0.41 | 0.9 | 0.666 | 0.979 |
| 4403 | MIR4691_3P | 72 | 0.41 | 0.9 | 0.665 | 0.979 |
| 4404 | WP_INTRACELLULAR_TRAFFICKING_PROTEINS_IN | 24 | 0.47 | 0.9 | 0.632 | 0.978 |
| 4405 | MIR8077 | 52 | 0.42 | 0.9 | 0.654 | 0.978 |
| 4406 | SANSOM_APC_TARGETS_REQUIRE_MYC | 211 | 0.38 | 0.89 | 0.741 | 0.979 |
| 4407 | BENPORATH_OCT4_TARGETS | 251 | 0.37 | 0.89 | 0.775 | 0.979 |
| 4408 | MIR5195_3P | 266 | 0.37 | 0.89 | 0.77 | 0.979 |
| 4409 | MIR576_3P | 48 | 0.43 | 0.89 | 0.656 | 0.979 |
| 4410 | PRC1_BMI_UP.V1_UP | 113 | 0.39 | 0.89 | 0.694 | 0.979 |
| 4411 | CDC5_01 | 196 | 0.38 | 0.89 | 0.743 | 0.979 |
| 4412 | MIR3136_5P | 66 | 0.41 | 0.89 | 0.663 | 0.979 |
| 4413 | ATACCTC_MIR202 | 158 | 0.38 | 0.89 | 0.711 | 0.979 |
| 4414 | MIR432_5P | 83 | 0.4 | 0.89 | 0.66 | 0.979 |
| 4415 | MIR5580_3P | 203 | 0.38 | 0.89 | 0.746 | 0.979 |
| 4416 | MIR144_3P | 434 | 0.37 | 0.89 | 0.827 | 0.979 |
| 4417 | REACTOME_NOTCH3_ACTIVATION_AND_TRANSMI | 25 | 0.47 | 0.89 | 0.646 | 0.979 |
| 4418 | GLI1_UP.V1_DN | 15 | 0.51 | 0.89 | 0.646 | 0.979 |
| 4419 | MIR7853_5P | 198 | 0.37 | 0.89 | 0.739 | 0.979 |

| | | | | | | |
|---|---|---|---|---|---|---|
| 4420 | MIR6809_5P | 119 | 0.38 | 0.89 | 0.693 | 0.979 |
| 4421 | MIR6893_3P | 80 | 0.4 | 0.89 | 0.677 | 0.979 |
| 4422 | KYNG_ENVIRONMENTAL_STRESS_RESPONSE_NC | 22 | 0.48 | 0.89 | 0.671 | 0.98 |
| 4423 | MIR6829_3P | 71 | 0.41 | 0.89 | 0.675 | 0.98 |
| 4424 | REACTOME_SUMOYLATION_OF_CHROMATIN_ORG | 61 | 0.41 | 0.89 | 0.671 | 0.98 |
| 4425 | TBPL1_TARGET_GENES | 96 | 0.39 | 0.89 | 0.679 | 0.981 |
| 4426 | MIR6506_5P | 82 | 0.4 | 0.89 | 0.683 | 0.981 |
| 4427 | PLASARI_TGFB1_SIGNALING_VIA_NFIC_10HR_DN | 19 | 0.49 | 0.89 | 0.637 | 0.98 |
| 4428 | MIR2681_5P | 126 | 0.39 | 0.89 | 0.702 | 0.98 |
| 4429 | MIR12131 | 148 | 0.38 | 0.89 | 0.738 | 0.98 |
| 4430 | REACTOME_SIGNALING_BY_FGFR4 | 31 | 0.45 | 0.89 | 0.66 | 0.98 |
| 4431 | FIGUEROA_AML_METHYLATION_CLUSTER_1_DN | 35 | 0.44 | 0.89 | 0.651 | 0.98 |
| 4432 | SMIRNOV_RESPONSE_TO_IR_6HR_DN | 93 | 0.39 | 0.89 | 0.682 | 0.98 |
| 4433 | MARZEC_IL2_SIGNALING_DN | 29 | 0.46 | 0.89 | 0.634 | 0.98 |
| 4434 | MIR302E | 423 | 0.36 | 0.89 | 0.8 | 0.98 |
| 4435 | MIR3692_5P | 130 | 0.38 | 0.89 | 0.713 | 0.98 |
| 4436 | PAX2_01 | 49 | 0.42 | 0.89 | 0.661 | 0.98 |
| 4437 | LFA1_Q6 | 189 | 0.38 | 0.89 | 0.75 | 0.98 |
| 4438 | TCTCTCC_MIR185 | 117 | 0.39 | 0.89 | 0.71 | 0.98 |
| 4439 | MIR10399_5P | 309 | 0.37 | 0.89 | 0.786 | 0.98 |
| 4440 | REACTOME_G_ALPHA_S_SIGNALLING_EVENTS | 65 | 0.41 | 0.89 | 0.679 | 0.98 |
| 4441 | MIR219A_1_3P | 46 | 0.43 | 0.89 | 0.662 | 0.98 |
| 4442 | GAUSSMANN_MLL_AF4_FUSION_TARGETS_E_DN | 16 | 0.5 | 0.89 | 0.667 | 0.98 |
| 4443 | THEODOROU_MAMMARY_TUMORIGENESIS | 15 | 0.51 | 0.89 | 0.649 | 0.98 |
| 4444 | MIR6808_5P | 158 | 0.38 | 0.89 | 0.72 | 0.98 |
| 4445 | MIR372_3P | 296 | 0.37 | 0.89 | 0.776 | 0.981 |
| 4446 | CREB_01 | 233 | 0.37 | 0.89 | 0.76 | 0.98 |
| 4447 | HANSON_HRAS_SIGNALING_VIA_NFKB | 17 | 0.5 | 0.89 | 0.631 | 0.981 |
| 4448 | MIR520B_3P_MIR520C_3P_MIR520E_3P | 295 | 0.37 | 0.89 | 0.772 | 0.981 |
| 4449 | REACTOME_NCAM_SIGNALING_FOR_NEURITE_OU | 49 | 0.42 | 0.89 | 0.665 | 0.981 |
| 4450 | WANG_CLIM2_TARGETS_UP | 229 | 0.37 | 0.89 | 0.773 | 0.981 |
| 4451 | MIR7847_3P | 63 | 0.41 | 0.89 | 0.699 | 0.981 |
| 4452 | REACTOME_HEDGEHOG_LIGAND_BIOGENESIS | 59 | 0.42 | 0.89 | 0.676 | 0.981 |

| | | | | | | |
|---|---|---|---|---|---|---|
| 4453 | MIR302B_5P | 171 | 0.38 | 0.89 | 0.725 | 0.981 |
| 4454 | MIR12127 | 64 | 0.41 | 0.89 | 0.671 | 0.981 |
| 4455 | MIR6884_5P | 124 | 0.39 | 0.89 | 0.71 | 0.981 |
| 4456 | MIR6508_5P | 291 | 0.37 | 0.89 | 0.783 | 0.981 |
| 4457 | MIR4800_5P | 18 | 0.49 | 0.89 | 0.645 | 0.982 |
| 4458 | MIR492 | 20 | 0.48 | 0.89 | 0.638 | 0.982 |
| 4459 | LMX1B_TARGET_GENES | 24 | 0.47 | 0.89 | 0.653 | 0.982 |
| 4460 | HEB_Q6 | 214 | 0.38 | 0.89 | 0.775 | 0.982 |
| 4461 | WP_MBDNF_AND_PROBDNF_REGULATION_OF_GA | 26 | 0.46 | 0.89 | 0.661 | 0.982 |
| 4462 | WP_VITAMIN_DSENSITIVE_CALCIUM_SIGNALING_I | 33 | 0.44 | 0.89 | 0.661 | 0.981 |
| 4463 | ROESSLER_LIVER_CANCER_METASTASIS_UP | 83 | 0.4 | 0.89 | 0.7 | 0.982 |
| 4464 | MIR3915 | 170 | 0.38 | 0.89 | 0.752 | 0.981 |
| 4465 | FOXO4_02 | 219 | 0.37 | 0.89 | 0.748 | 0.981 |
| 4466 | TAAWWATAG_RSRFC4_Q2 | 127 | 0.38 | 0.89 | 0.72 | 0.981 |
| 4467 | BIOCARTA_HDAC_PATHWAY | 21 | 0.48 | 0.89 | 0.649 | 0.981 |
| 4468 | WP_LEUKOCYTEINTRINSIC_HIPPO_PATHWAY_FUN | 28 | 0.45 | 0.89 | 0.638 | 0.981 |
| 4469 | HATADA_METHYLATED_IN_LUNG_CANCER_UP | 252 | 0.37 | 0.89 | 0.773 | 0.982 |
| 4470 | ZSCAN21_TARGET_GENES | 430 | 0.36 | 0.89 | 0.828 | 0.982 |
| 4471 | HILLION_HMGA1_TARGETS | 65 | 0.4 | 0.89 | 0.694 | 0.982 |
| 4472 | CCAWNWWNNNGGC_UNKNOWN | 79 | 0.4 | 0.89 | 0.699 | 0.982 |
| 4473 | STAT_Q6 | 202 | 0.37 | 0.89 | 0.746 | 0.981 |
| 4474 | TGTYNNNNNRGCARM_UNKNOWN | 72 | 0.41 | 0.89 | 0.688 | 0.981 |
| 4475 | FIGUEROA_AML_METHYLATION_CLUSTER_5_DN | 26 | 0.46 | 0.89 | 0.651 | 0.981 |
| 4476 | UBE2I_TARGET_GENES | 61 | 0.41 | 0.89 | 0.667 | 0.981 |
| 4477 | CP2_01 | 217 | 0.37 | 0.89 | 0.758 | 0.982 |
| 4478 | CACTTTG_MIR520G_MIR520H | 220 | 0.37 | 0.89 | 0.759 | 0.982 |
| 4479 | LIEN_BREAST_CARCINOMA_METAPLASTIC_VS_DU | 76 | 0.4 | 0.89 | 0.687 | 0.983 |
| 4480 | MIR488_3P | 146 | 0.38 | 0.89 | 0.726 | 0.982 |
| 4481 | ATACTGT_MIR144 | 178 | 0.37 | 0.89 | 0.728 | 0.982 |
| 4482 | MIR2114_3P | 32 | 0.44 | 0.89 | 0.643 | 0.982 |
| 4483 | MIR1915_5P | 37 | 0.43 | 0.89 | 0.651 | 0.982 |
| 4484 | POU1F1_Q6 | 181 | 0.38 | 0.89 | 0.751 | 0.982 |
| 4485 | MIR4696 | 83 | 0.4 | 0.89 | 0.694 | 0.982 |

| | | | | | | |
|---|---|---|---|---|---|---|
| 4486 | SATO_SILENCED_EPIGENETICALLY_IN_PANCREA | 29 | 0.46 | 0.89 | 0.65 | 0.982 |
| 4487 | SMID_BREAST_CANCER_ERBB2_UP | 100 | 0.39 | 0.89 | 0.708 | 0.982 |
| 4488 | PID_IL1_PATHWAY | 28 | 0.46 | 0.89 | 0.647 | 0.983 |
| 4489 | LAMB_CCND1_TARGETS | 17 | 0.49 | 0.89 | 0.646 | 0.983 |
| 4490 | MIR4703_5P | 203 | 0.37 | 0.89 | 0.779 | 0.983 |
| 4491 | MIR6073 | 88 | 0.39 | 0.89 | 0.697 | 0.984 |
| 4492 | PID_SMAD2_3PATHWAY | 16 | 0.49 | 0.89 | 0.647 | 0.984 |
| 4493 | MIR4452 | 135 | 0.38 | 0.88 | 0.739 | 0.984 |
| 4494 | KANG_IMMORTALIZED_BY_TERT_DN | 59 | 0.41 | 0.88 | 0.671 | 0.984 |
| 4495 | MIR5004_3P | 111 | 0.39 | 0.88 | 0.713 | 0.984 |
| 4496 | MIR3158_5P | 141 | 0.38 | 0.88 | 0.731 | 0.984 |
| 4497 | MIR628_5P | 72 | 0.4 | 0.88 | 0.7 | 0.984 |
| 4498 | MIR608 | 171 | 0.38 | 0.88 | 0.756 | 0.984 |
| 4499 | MIR651_5P | 98 | 0.39 | 0.88 | 0.702 | 0.984 |
| 4500 | REACTOME_ADRENALINE_NORADRENALINE_INHIE | 22 | 0.47 | 0.88 | 0.642 | 0.984 |
| 4501 | SCHLOSSER_MYC_TARGETS_AND_SERUM_RESP( | 48 | 0.42 | 0.88 | 0.684 | 0.985 |
| 4502 | TTGCCAA_MIR182 | 302 | 0.37 | 0.88 | 0.785 | 0.985 |
| 4503 | DACOSTA_UV_RESPONSE_VIA_ERCC3_COMMON_ | 72 | 0.4 | 0.88 | 0.682 | 0.985 |
| 4504 | REACTOME_REGULATION_OF_CHOLESTEROL_BI( | 54 | 0.41 | 0.88 | 0.671 | 0.984 |
| 4505 | GUENTHER_GROWTH_SPHERICAL_VS_ADHERENT | 17 | 0.5 | 0.88 | 0.665 | 0.985 |
| 4506 | MIR4726_5P | 89 | 0.39 | 0.88 | 0.696 | 0.985 |
| 4507 | WP_BMP_SIGNALING_IN_EYELID_DEVELOPMENT | 15 | 0.5 | 0.88 | 0.661 | 0.985 |
| 4508 | MIR7159_5P | 314 | 0.36 | 0.88 | 0.816 | 0.985 |
| 4509 | SENGUPTA_EBNA1_ANTICORRELATED | 124 | 0.38 | 0.88 | 0.729 | 0.985 |
| 4510 | RYTGCNNRGNAAC_MIF1_01 | 77 | 0.4 | 0.88 | 0.717 | 0.985 |
| 4511 | ZNF584_TARGET_GENES | 147 | 0.38 | 0.88 | 0.731 | 0.984 |
| 4512 | MIR4441 | 239 | 0.37 | 0.88 | 0.776 | 0.985 |
| 4513 | MIR1185_1_3P_MIR1185_2_3P | 351 | 0.36 | 0.88 | 0.822 | 0.985 |
| 4514 | BIOCARTA_PPARA_PATHWAY | 44 | 0.43 | 0.88 | 0.68 | 0.985 |
| 4515 | MIR142_5P | 300 | 0.37 | 0.88 | 0.81 | 0.985 |
| 4516 | REACTOME_A_TETRASACCHARIDE_LINKER_SEQU | 23 | 0.46 | 0.88 | 0.677 | 0.985 |
| 4517 | MIR520A_3P | 298 | 0.37 | 0.88 | 0.792 | 0.985 |
| 4518 | KEGG_ABC_TRANSPORTERS | 27 | 0.46 | 0.88 | 0.652 | 0.985 |

| | | | | | | |
|---|---|---|---|---|---|---|
| 4519 | AAAYRNCTG_UNKNOWN | 285 | 0.36 | 0.88 | 0.795 | 0.985 |
| 4520 | HIF1_Q3 | 212 | 0.37 | 0.88 | 0.768 | 0.985 |
| 4521 | DITTMER_PTHLH_TARGETS_UP | 106 | 0.39 | 0.88 | 0.722 | 0.985 |
| 4522 | MIR4472 | 89 | 0.4 | 0.88 | 0.721 | 0.985 |
| 4523 | BIOCARTA_G2_PATHWAY | 24 | 0.45 | 0.88 | 0.681 | 0.984 |
| 4524 | MIR4725_3P | 233 | 0.37 | 0.88 | 0.774 | 0.984 |
| 4525 | MIR370_3P | 82 | 0.39 | 0.88 | 0.697 | 0.984 |
| 4526 | MIR4712_5P | 86 | 0.39 | 0.88 | 0.708 | 0.984 |
| 4527 | MIR605_5P | 115 | 0.38 | 0.88 | 0.729 | 0.984 |
| 4528 | VERRECCHIA_RESPONSE_TO_TGFB1_C2 | 15 | 0.5 | 0.88 | 0.675 | 0.984 |
| 4529 | MIR4474_5P | 39 | 0.43 | 0.88 | 0.681 | 0.984 |
| 4530 | SRF_Q5_01 | 176 | 0.37 | 0.88 | 0.751 | 0.984 |
| 4531 | YAO_TEMPORAL_RESPONSE_TO_PROGESTERON | 137 | 0.38 | 0.88 | 0.736 | 0.984 |
| 4532 | BLANCO_MELO_BRONCHIAL_EPITHELIAL_CELLS_ | 64 | 0.41 | 0.88 | 0.683 | 0.984 |
| 4533 | TSENG_IRS1_TARGETS_DN | 97 | 0.39 | 0.88 | 0.723 | 0.984 |
| 4534 | FONTAINE_FOLLICULAR_THYROID_ADENOMA_UP | 67 | 0.4 | 0.88 | 0.7 | 0.984 |
| 4535 | PID_ECADHERIN_NASCENT_AJ_PATHWAY | 38 | 0.43 | 0.88 | 0.676 | 0.984 |
| 4536 | DESERT_PERIVENOUS_HEPATOCELLULAR_CARCI | 125 | 0.38 | 0.88 | 0.729 | 0.984 |
| 4537 | REACTOME_PROTEIN_LOCALIZATION | 155 | 0.37 | 0.88 | 0.75 | 0.984 |
| 4538 | MIR1206 | 98 | 0.39 | 0.88 | 0.715 | 0.985 |
| 4539 | MCCABE_HOXC6_TARGETS_CANCER_UP | 22 | 0.48 | 0.88 | 0.687 | 0.985 |
| 4540 | STK33_DN | 221 | 0.37 | 0.88 | 0.762 | 0.985 |
| 4541 | ATM_DN.V1_DN | 73 | 0.39 | 0.88 | 0.7 | 0.984 |
| 4542 | RUVBL1_TARGET_GENES | 150 | 0.38 | 0.88 | 0.74 | 0.985 |
| 4543 | MIR6864_5P | 68 | 0.4 | 0.88 | 0.705 | 0.985 |
| 4544 | MIR16_1_3P | 159 | 0.38 | 0.88 | 0.746 | 0.985 |
| 4545 | MIR922 | 165 | 0.37 | 0.88 | 0.765 | 0.985 |
| 4546 | ATF_01 | 224 | 0.37 | 0.88 | 0.782 | 0.985 |
| 4547 | MIR4688 | 41 | 0.43 | 0.88 | 0.679 | 0.985 |
| 4548 | BRUINS_UVC_RESPONSE_EARLY_LATE | 287 | 0.36 | 0.88 | 0.812 | 0.985 |
| 4549 | MIR10401_5P | 61 | 0.41 | 0.88 | 0.691 | 0.985 |
| 4550 | MIR2276_5P | 64 | 0.41 | 0.88 | 0.698 | 0.986 |
| 4551 | PRDM12_TARGET_GENES | 90 | 0.39 | 0.88 | 0.722 | 0.986 |

| | | | | | | |
|---|---|---|---|---|---|---|
| 4552 | MIR129_1_3P_MIR129_2_3P | 119 | 0.38 | 0.88 | 0.731 | 0.986 |
| 4553 | MIR33A_3P | 357 | 0.36 | 0.88 | 0.84 | 0.986 |
| 4554 | MIR4654 | 38 | 0.43 | 0.88 | 0.668 | 0.987 |
| 4555 | MIR4718 | 93 | 0.39 | 0.88 | 0.732 | 0.987 |
| 4556 | MIR4776_5P | 44 | 0.42 | 0.88 | 0.692 | 0.987 |
| 4557 | FERREIRA_EWINGS_SARCOMA_UNSTABLE_VS_ST | 76 | 0.39 | 0.88 | 0.72 | 0.988 |
| 4558 | MIR448 | 314 | 0.36 | 0.88 | 0.823 | 0.988 |
| 4559 | MIR2682_5P | 97 | 0.39 | 0.88 | 0.723 | 0.988 |
| 4560 | CREB_Q4 | 238 | 0.36 | 0.88 | 0.799 | 0.988 |
| 4561 | MIR9500 | 136 | 0.38 | 0.88 | 0.754 | 0.988 |
| 4562 | KEGG_ADHERENS_JUNCTION | 70 | 0.4 | 0.88 | 0.711 | 0.989 |
| 4563 | MIR6792_5P | 108 | 0.38 | 0.88 | 0.72 | 0.988 |
| 4564 | MIR363_3P | 378 | 0.36 | 0.88 | 0.825 | 0.988 |
| 4565 | MIR194_5P | 171 | 0.37 | 0.88 | 0.753 | 0.989 |
| 4566 | WANG_ESOPHAGUS_CANCER_VS_NORMAL_DN | 89 | 0.39 | 0.88 | 0.708 | 0.989 |
| 4567 | IVANOVA_HEMATOPOIESIS_STEM_CELL_LONG_T | 169 | 0.37 | 0.88 | 0.77 | 0.989 |
| 4568 | MCBRYAN_PUBERTAL_BREAST_5_6WK_UP | 93 | 0.39 | 0.88 | 0.713 | 0.989 |
| 4569 | AR_01 | 127 | 0.38 | 0.88 | 0.75 | 0.989 |
| 4570 | SUZUKI_RESPONSE_TO_TSA_AND_DECITABINE_1 | 15 | 0.5 | 0.88 | 0.675 | 0.989 |
| 4571 | MIR887_5P | 88 | 0.39 | 0.88 | 0.704 | 0.989 |
| 4572 | DAIRKEE_CANCER_PRONE_RESPONSE_BPA | 44 | 0.42 | 0.88 | 0.68 | 0.989 |
| 4573 | REACTOME_REGULATION_OF_INSULIN_SECRETIC | 60 | 0.4 | 0.87 | 0.693 | 0.99 |
| 4574 | MIR6799_3P | 107 | 0.38 | 0.87 | 0.721 | 0.99 |
| 4575 | GENTILE_UV_RESPONSE_CLUSTER_D9 | 24 | 0.46 | 0.87 | 0.655 | 0.99 |
| 4576 | MIR367_3P | 379 | 0.36 | 0.87 | 0.842 | 0.989 |
| 4577 | GAVIN_FOXP3_TARGETS_CLUSTER_T7 | 90 | 0.39 | 0.87 | 0.712 | 0.99 |
| 4578 | GTAAACC_MIR2995P | 44 | 0.42 | 0.87 | 0.685 | 0.99 |
| 4579 | RLF_TARGET_GENES | 302 | 0.36 | 0.87 | 0.82 | 0.99 |
| 4580 | REACTOME_HEPARAN_SULFATE_HEPARIN_HS_G/ | 47 | 0.42 | 0.87 | 0.678 | 0.989 |
| 4581 | MIR939_5P | 107 | 0.38 | 0.87 | 0.735 | 0.99 |
| 4582 | MIR3529_3P | 480 | 0.36 | 0.87 | 0.863 | 0.99 |
| 4583 | MIR4312 | 36 | 0.43 | 0.87 | 0.668 | 0.99 |
| 4584 | REACTOME_FACTORS_INVOLVED_IN_MEGAKARY( | 124 | 0.38 | 0.87 | 0.731 | 0.99 |

| | | | | | | |
|---|---|---|---|---|---|---|
| 4585 | LMNB2_TARGET_GENES | 31 | 0.44 | 0.87 | 0.682 | 0.99 |
| 4586 | PHB2_TARGET_GENES | 138 | 0.38 | 0.87 | 0.74 | 0.99 |
| 4587 | BASSO_HAIRY_CELL_LEUKEMIA_UP | 54 | 0.41 | 0.87 | 0.7 | 0.991 |
| 4588 | FOROUTAN_PRODRANK_TGFB_EMT_DN | 68 | 0.4 | 0.87 | 0.706 | 0.991 |
| 4589 | MIR4684_3P | 96 | 0.39 | 0.87 | 0.734 | 0.991 |
| 4590 | MIR4645_3P | 88 | 0.39 | 0.87 | 0.732 | 0.992 |
| 4591 | MIR6836_5P | 118 | 0.38 | 0.87 | 0.751 | 0.992 |
| 4592 | NOTCH_DN.V1_UP | 120 | 0.38 | 0.87 | 0.753 | 0.992 |
| 4593 | MIR6880_5P | 74 | 0.4 | 0.87 | 0.737 | 0.992 |
| 4594 | MIR219B_3P | 44 | 0.43 | 0.87 | 0.685 | 0.992 |
| 4595 | YAO_TEMPORAL_RESPONSE_TO_PROGESTERON | 44 | 0.43 | 0.87 | 0.686 | 0.991 |
| 4596 | CHIARADONNA_NEOPLASTIC_TRANSFORMATION_ | 117 | 0.38 | 0.87 | 0.739 | 0.992 |
| 4597 | MIR6511A_5P | 165 | 0.37 | 0.87 | 0.77 | 0.992 |
| 4598 | XU_HGF_TARGETS_INDUCED_BY_AKT1_48HR_DN | 21 | 0.47 | 0.87 | 0.675 | 0.992 |
| 4599 | MIR770_5P | 88 | 0.39 | 0.87 | 0.726 | 0.992 |
| 4600 | TIEN_INTESTINE_PROBIOTICS_2HR_DN | 82 | 0.39 | 0.87 | 0.717 | 0.992 |
| 4601 | VECCHI_GASTRIC_CANCER_ADVANCED_VS_EARL | 83 | 0.39 | 0.87 | 0.724 | 0.991 |
| 4602 | MIR548G_3P | 212 | 0.37 | 0.87 | 0.793 | 0.992 |
| 4603 | ATGAAGG_MIR205 | 145 | 0.37 | 0.87 | 0.744 | 0.992 |
| 4604 | MIR6734_5P | 55 | 0.41 | 0.87 | 0.697 | 0.992 |
| 4605 | MIR489_3P | 62 | 0.4 | 0.87 | 0.703 | 0.992 |
| 4606 | MIR6794_3P | 74 | 0.39 | 0.87 | 0.722 | 0.992 |
| 4607 | REACTOME_VOLTAGE_GATED_POTASSIUM_CHAN | 35 | 0.43 | 0.87 | 0.687 | 0.992 |
| 4608 | REACTOME_MUSCLE_CONTRACTION | 125 | 0.37 | 0.87 | 0.729 | 0.992 |
| 4609 | NIKOLSKY_MUTATED_AND_AMPLIFIED_IN_BREAS | 81 | 0.39 | 0.87 | 0.711 | 0.992 |
| 4610 | MIR3667_3P | 126 | 0.38 | 0.87 | 0.741 | 0.992 |
| 4611 | MIR7160_3P | 28 | 0.45 | 0.87 | 0.686 | 0.992 |
| 4612 | MIR183_5P | 216 | 0.36 | 0.87 | 0.794 | 0.992 |
| 4613 | CART1_01 | 174 | 0.37 | 0.87 | 0.771 | 0.992 |
| 4614 | MIR4428 | 106 | 0.38 | 0.87 | 0.743 | 0.992 |
| 4615 | GCGCTTT_MIR518B_MIR518C_MIR518D | 17 | 0.49 | 0.87 | 0.692 | 0.992 |
| 4616 | MIR6081 | 91 | 0.38 | 0.87 | 0.722 | 0.992 |
| 4617 | DESERT_PERIPORTAL_HEPATOCELLULAR_CARCI | 71 | 0.4 | 0.87 | 0.704 | 0.992 |

| | | | | | | |
|---|---|---|---|---|---|---|
| 4618 | REACTOME_REGULATION_OF_TP53_ACTIVITY | 157 | 0.37 | 0.87 | 0.782 | 0.992 |
| 4619 | DORSAM_HOXA9_TARGETS_DN | 29 | 0.44 | 0.87 | 0.692 | 0.993 |
| 4620 | MIR4292 | 143 | 0.37 | 0.87 | 0.761 | 0.993 |
| 4621 | ACAGGGT_MIR10A_MIR10B | 109 | 0.38 | 0.87 | 0.737 | 0.993 |
| 4622 | MIR29B_2_5P | 182 | 0.37 | 0.87 | 0.767 | 0.993 |
| 4623 | MIR513A_5P | 319 | 0.36 | 0.87 | 0.839 | 0.993 |
| 4624 | MIR6748_5P | 54 | 0.41 | 0.87 | 0.709 | 0.993 |
| 4625 | MIR455_5P | 73 | 0.4 | 0.87 | 0.727 | 0.993 |
| 4626 | MIR6798_5P | 37 | 0.43 | 0.87 | 0.683 | 0.993 |
| 4627 | LUI_THYROID_CANCER_CLUSTER_2 | 40 | 0.42 | 0.87 | 0.684 | 0.993 |
| 4628 | ROZANOV_MMP14_TARGETS_DN | 24 | 0.45 | 0.87 | 0.672 | 0.993 |
| 4629 | MIR7111_3P | 94 | 0.38 | 0.87 | 0.747 | 0.993 |
| 4630 | REACTOME_METABOLISM_OF_VITAMINS_AND_CC | 140 | 0.37 | 0.87 | 0.775 | 0.993 |
| 4631 | MIR202_3P | 209 | 0.37 | 0.87 | 0.798 | 0.993 |
| 4632 | MIR4457 | 106 | 0.38 | 0.87 | 0.737 | 0.993 |
| 4633 | MIR31_5P | 126 | 0.38 | 0.87 | 0.747 | 0.993 |
| 4634 | ZNF404_TARGET_GENES | 20 | 0.47 | 0.87 | 0.671 | 0.993 |
| 4635 | MIR4283 | 172 | 0.37 | 0.87 | 0.779 | 0.992 |
| 4636 | MIR34B_3P | 97 | 0.39 | 0.87 | 0.737 | 0.993 |
| 4637 | BCAT_GDS748_UP | 37 | 0.43 | 0.87 | 0.696 | 0.993 |
| 4638 | MIR138_2_3P | 147 | 0.37 | 0.87 | 0.771 | 0.993 |
| 4639 | HOLLERN_EMT_BREAST_TUMOR_DN | 101 | 0.38 | 0.87 | 0.752 | 0.993 |
| 4640 | MIR507 | 231 | 0.36 | 0.87 | 0.807 | 0.993 |
| 4641 | ACACTCC_MIR122A | 71 | 0.39 | 0.87 | 0.713 | 0.994 |
| 4642 | TAYLOR_METHYLATED_IN_ACUTE_LYMPHOBLAST | 72 | 0.39 | 0.87 | 0.709 | 0.994 |
| 4643 | REACTOME_DOWNREGULATION_OF_TGF_BETA_F | 24 | 0.46 | 0.87 | 0.669 | 0.994 |
| 4644 | WP_ANDROGEN_RECEPTOR_SIGNALING_PATHWA | 88 | 0.39 | 0.87 | 0.734 | 0.994 |
| 4645 | CARD_MIR302A_TARGETS | 72 | 0.39 | 0.87 | 0.725 | 0.994 |
| 4646 | VARELA_ZMPSTE24_TARGETS_UP | 42 | 0.42 | 0.87 | 0.708 | 0.994 |
| 4647 | PRC2_EZH2_UP.V1_DN | 153 | 0.37 | 0.87 | 0.79 | 0.995 |
| 4648 | MIR3135B | 90 | 0.38 | 0.87 | 0.739 | 0.994 |
| 4649 | MIR7854_3P | 88 | 0.39 | 0.87 | 0.741 | 0.994 |
| 4650 | REACTOME_RAS_ACTIVATION_UPON_CA2_INFLUX | 17 | 0.49 | 0.87 | 0.695 | 0.995 |

| | | | | | | |
|---|---|---|---|---|---|---|
| 4651 | PRDM4_TARGET_GENES | 366 | 0.35 | 0.87 | 0.852 | 0.995 |
| 4652 | MIR1208 | 120 | 0.37 | 0.87 | 0.752 | 0.995 |
| 4653 | KYNG_WERNER_SYNDROM_AND_NORMAL_AGING | 155 | 0.37 | 0.87 | 0.776 | 0.995 |
| 4654 | MIR3194_5P | 63 | 0.4 | 0.86 | 0.705 | 0.995 |
| 4655 | NADLER_OBESITY_DN | 36 | 0.43 | 0.86 | 0.687 | 0.995 |
| 4656 | MIR548L | 300 | 0.36 | 0.86 | 0.837 | 0.995 |
| 4657 | MIR137_3P | 470 | 0.35 | 0.86 | 0.883 | 0.995 |
| 4658 | MYOGENIN_Q6 | 209 | 0.36 | 0.86 | 0.803 | 0.995 |
| 4659 | CACTGTG_MIR128A_MIR128B | 306 | 0.36 | 0.86 | 0.834 | 0.995 |
| 4660 | ER_Q6_01 | 218 | 0.36 | 0.86 | 0.818 | 0.995 |
| 4661 | MIR6776_3P | 54 | 0.4 | 0.86 | 0.71 | 0.996 |
| 4662 | BIOCARTA_RAS_PATHWAY | 20 | 0.47 | 0.86 | 0.697 | 0.995 |
| 4663 | GTACTGT_MIR101 | 228 | 0.36 | 0.86 | 0.808 | 0.996 |
| 4664 | E47_01 | 210 | 0.36 | 0.86 | 0.812 | 0.995 |
| 4665 | GGGATGC_MIR3245P | 46 | 0.42 | 0.86 | 0.687 | 0.996 |
| 4666 | WP_PURINERGIC_SIGNALING | 18 | 0.47 | 0.86 | 0.69 | 0.996 |
| 4667 | MIR6132 | 121 | 0.37 | 0.86 | 0.749 | 0.996 |
| 4668 | WP_PDGF_PATHWAY | 39 | 0.42 | 0.86 | 0.706 | 0.996 |
| 4669 | MIR1843 | 44 | 0.41 | 0.86 | 0.701 | 0.996 |
| 4670 | MIR619_5P | 116 | 0.38 | 0.86 | 0.767 | 0.996 |
| 4671 | HNF4ALPHA_Q6 | 214 | 0.36 | 0.86 | 0.811 | 0.996 |
| 4672 | MIR6846_5P | 65 | 0.4 | 0.86 | 0.719 | 0.996 |
| 4673 | MIR497_3P | 210 | 0.36 | 0.86 | 0.814 | 0.996 |
| 4674 | REACTOME_LAMININ_INTERACTIONS | 23 | 0.45 | 0.86 | 0.668 | 0.996 |
| 4675 | MIR3185 | 153 | 0.37 | 0.86 | 0.784 | 0.996 |
| 4676 | BEIER_GLIOMA_STEM_CELL_UP | 34 | 0.43 | 0.86 | 0.708 | 0.995 |
| 4677 | CUI_TCF21_TARGETS_DN | 23 | 0.45 | 0.86 | 0.688 | 0.996 |
| 4678 | MIR25_3P | 373 | 0.36 | 0.86 | 0.868 | 0.996 |
| 4679 | GCM_Q2 | 204 | 0.36 | 0.86 | 0.823 | 0.996 |
| 4680 | MIR6789_5P | 18 | 0.47 | 0.86 | 0.675 | 0.995 |
| 4681 | MIR154_5P | 52 | 0.4 | 0.86 | 0.713 | 0.996 |
| 4682 | MIR3650 | 43 | 0.42 | 0.86 | 0.72 | 0.996 |
| 4683 | MIR6832_3P | 212 | 0.36 | 0.86 | 0.811 | 0.996 |

| | | | | | | |
|---|---|---|---|---|---|---|
| 4684 | MIR6769A_3P | 15 | 0.49 | 0.86 | 0.673 | 0.996 |
| 4685 | MIR1233_5P | 79 | 0.39 | 0.86 | 0.728 | 0.997 |
| 4686 | MIR5580_5P | 104 | 0.38 | 0.86 | 0.749 | 0.997 |
| 4687 | CADWELL_ATG16L1_TARGETS_DN | 55 | 0.4 | 0.86 | 0.723 | 0.997 |
| 4688 | GROSS_HYPOXIA_VIA_ELK3_ONLY_UP | 29 | 0.44 | 0.86 | 0.704 | 0.997 |
| 4689 | MIR22_3P | 123 | 0.37 | 0.86 | 0.765 | 0.998 |
| 4690 | REACTOME_SIGNALING_BY_ROBO_RECEPTORS | 207 | 0.36 | 0.86 | 0.806 | 0.998 |
| 4691 | IVANOVA_HEMATOPOIESIS_MATURE_CELL | 244 | 0.36 | 0.86 | 0.815 | 0.998 |
| 4692 | GTAAGAT_MIR200A | 48 | 0.41 | 0.86 | 0.708 | 0.998 |
| 4693 | MIR18B_3P | 43 | 0.41 | 0.86 | 0.692 | 0.998 |
| 4694 | WP_THYROID_STIMULATING_HORMONE_TSH_SIGI | 61 | 0.4 | 0.86 | 0.714 | 0.998 |
| 4695 | MIR4731_5P | 215 | 0.36 | 0.86 | 0.805 | 0.998 |
| 4696 | GGTGAAG_MIR412 | 56 | 0.4 | 0.86 | 0.735 | 0.998 |
| 4697 | NADLER_HYPERGLYCEMIA_AT_OBESITY | 49 | 0.41 | 0.86 | 0.711 | 0.999 |
| 4698 | AP4_Q5 | 216 | 0.36 | 0.86 | 0.813 | 0.998 |
| 4699 | REACTOME_DISORDERS_OF_TRANSMEMBRANE_T | 132 | 0.37 | 0.86 | 0.781 | 0.999 |
| 4700 | MIR6848_5P | 68 | 0.39 | 0.86 | 0.717 | 0.999 |
| 4701 | MIR4715_3P | 52 | 0.4 | 0.86 | 0.713 | 0.999 |
| 4702 | ELVIDGE_HIF1A_TARGETS_DN | 81 | 0.38 | 0.86 | 0.737 | 0.999 |
| 4703 | HP1SITEFACTOR_Q6 | 189 | 0.36 | 0.86 | 0.783 | 0.999 |
| 4704 | MIR557 | 198 | 0.36 | 0.86 | 0.799 | 0.999 |
| 4705 | HOEGERKORP_CD44_TARGETS_TEMPORAL_DN | 19 | 0.46 | 0.86 | 0.71 | 0.999 |
| 4706 | MIR4494 | 51 | 0.41 | 0.86 | 0.72 | 0.999 |
| 4707 | MIR6509_5P | 77 | 0.39 | 0.86 | 0.738 | 0.999 |
| 4708 | GUTIERREZ_MULTIPLE_MYELOMA_UP | 35 | 0.43 | 0.86 | 0.711 | 1 |
| 4709 | IVANOVA_HEMATOPOIESIS_INTERMEDIATE_PROG | 123 | 0.36 | 0.86 | 0.791 | 1 |
| 4710 | REACTOME_NOTCH3_INTRACELLULAR_DOMAIN_F | 20 | 0.46 | 0.86 | 0.691 | 1 |
| 4711 | MEBARKI_HCC_PROGENITOR_WNT_UP_CTNNB1_I | 29 | 0.44 | 0.86 | 0.705 | 1 |
| 4712 | REACTOME_INTEGRATION_OF_ENERGY_METABO | 84 | 0.38 | 0.86 | 0.734 | 1 |
| 4713 | MIR5694 | 47 | 0.41 | 0.86 | 0.721 | 1 |
| 4714 | HOX13_01 | 36 | 0.43 | 0.86 | 0.699 | 1 |
| 4715 | MIR654_3P | 68 | 0.39 | 0.86 | 0.726 | 1 |
| 4716 | REACTOME_DEGRADATION_OF_BETA_CATENIN_E | 83 | 0.38 | 0.86 | 0.726 | 1 |

| | | | | | | |
|---|---|---|---|---|---|---|
| 4717 | ELK1_01 | 242 | 0.35 | 0.86 | 0.821 | 1 |
| 4718 | REACTOME_TRAF6_MEDIATED_INDUCTION_OF_TA | 15 | 0.48 | 0.86 | 0.678 | 1 |
| 4719 | LET_7F_2_3P | 344 | 0.35 | 0.86 | 0.859 | 1 |
| 4720 | KANG_CISPLATIN_RESISTANCE_UP | 20 | 0.46 | 0.86 | 0.711 | 1 |
| 4721 | MIR3675_3P | 75 | 0.39 | 0.86 | 0.745 | 1 |
| 4722 | MCBRYAN_PUBERTAL_BREAST_6_7WK_DN | 56 | 0.4 | 0.86 | 0.73 | 1 |
| 4723 | WP_ANGIOTENSIN_II_RECEPTOR_TYPE_1_PATHW | 23 | 0.45 | 0.86 | 0.688 | 1 |
| 4724 | MIR4802_5P | 44 | 0.41 | 0.85 | 0.717 | 1 |
| 4725 | BENPORATH_NOS_TARGETS | 149 | 0.37 | 0.85 | 0.797 | 1 |
| 4726 | PID_FAS_PATHWAY | 35 | 0.43 | 0.85 | 0.716 | 1 |
| 4727 | BEGUM_TARGETS_OF_PAX3_FOXO1_FUSION_UP | 45 | 0.41 | 0.85 | 0.714 | 1 |
| 4728 | MIR1271_5P | 333 | 0.35 | 0.85 | 0.861 | 1 |
| 4729 | MATZUK_MEIOTIC_AND_DNA_REPAIR | 32 | 0.43 | 0.85 | 0.697 | 1 |
| 4730 | GGGACCA_MIR133A_MIR133B | 189 | 0.36 | 0.85 | 0.805 | 1 |
| 4731 | DARWICHE_PAPILLOMA_RISK_LOW_DN | 136 | 0.37 | 0.85 | 0.79 | 1 |
| 4732 | SREBP1_Q6 | 202 | 0.36 | 0.85 | 0.813 | 1 |
| 4733 | GARGALOVIC_RESPONSE_TO_OXIDIZED_PHOSPH | 23 | 0.45 | 0.85 | 0.696 | 1 |
| 4734 | MIR4684_5P | 143 | 0.37 | 0.85 | 0.777 | 1 |
| 4735 | MIR4755_3P | 129 | 0.37 | 0.85 | 0.769 | 1 |
| 4736 | CAGNYGKNAAA_UNKNOWN | 65 | 0.39 | 0.85 | 0.727 | 1 |
| 4737 | MIR205_5P | 190 | 0.36 | 0.85 | 0.791 | 1 |
| 4738 | IBRAHIM_NRF1_DOWN | 151 | 0.37 | 0.85 | 0.782 | 1 |
| 4739 | BRF1_TARGET_GENES | 41 | 0.42 | 0.85 | 0.711 | 1 |
| 4740 | MIR6856_3P | 53 | 0.4 | 0.85 | 0.738 | 1 |
| 4741 | REACTOME_RET_SIGNALING | 33 | 0.43 | 0.85 | 0.715 | 1 |
| 4742 | MIR345_3P | 184 | 0.36 | 0.85 | 0.804 | 1 |
| 4743 | MIR5579_5P | 66 | 0.39 | 0.85 | 0.736 | 1 |
| 4744 | GCAAAAA_MIR129 | 175 | 0.36 | 0.85 | 0.812 | 1 |
| 4745 | HNF4_DR1_Q3 | 214 | 0.36 | 0.85 | 0.832 | 1 |
| 4746 | MIR4784 | 120 | 0.37 | 0.85 | 0.772 | 1 |
| 4747 | COLIN_PILOCYTIC_ASTROCYTOMA_VS_GLIOBLAS | 24 | 0.44 | 0.85 | 0.711 | 1 |
| 4748 | MIR376A_2_5P | 55 | 0.4 | 0.85 | 0.701 | 1 |
| 4749 | GCTTGAA_MIR498 | 99 | 0.38 | 0.85 | 0.747 | 1 |

| | | | | | | |
|---|---|---|---|---|---|---|
| 4750 | MIR6516_3P | 115 | 0.37 | 0.85 | 0.772 | 1 |
| 4751 | PARK_TRETINOIN_RESPONSE_AND_RARA_PLZF_I | 16 | 0.48 | 0.85 | 0.697 | 1 |
| 4752 | MIR302C_5P | 484 | 0.35 | 0.85 | 0.915 | 1 |
| 4753 | YORDY_RECIPROCAL_REGULATION_BY_ETS1_AN | 20 | 0.46 | 0.85 | 0.694 | 1 |
| 4754 | MIR487A_5P | 91 | 0.38 | 0.85 | 0.746 | 1 |
| 4755 | WP_BDNFTRKB_SIGNALING | 29 | 0.44 | 0.85 | 0.716 | 1 |
| 4756 | UEDA_CENTRAL_CLOCK | 81 | 0.38 | 0.85 | 0.746 | 1 |
| 4757 | CTGTTAC_MIR194 | 95 | 0.38 | 0.85 | 0.769 | 1 |
| 4758 | OCT1_05 | 180 | 0.36 | 0.85 | 0.806 | 1 |
| 4759 | MIR6773_3P | 87 | 0.38 | 0.85 | 0.757 | 1 |
| 4760 | MIR4309 | 36 | 0.42 | 0.85 | 0.715 | 1 |
| 4761 | ULE_SPLICING_VIA_NOVA2 | 40 | 0.42 | 0.85 | 0.724 | 1 |
| 4762 | REACTOME_CROSS_PRESENTATION_OF_SOLUBL | 45 | 0.41 | 0.85 | 0.733 | 1 |
| 4763 | TACTTGA_MIR26A_MIR26B | 278 | 0.36 | 0.85 | 0.838 | 1 |
| 4764 | MIR3619_3P | 44 | 0.41 | 0.85 | 0.724 | 1 |
| 4765 | MIR7156_5P | 107 | 0.37 | 0.85 | 0.771 | 1 |
| 4766 | MIR1295B_3P | 15 | 0.49 | 0.85 | 0.688 | 1 |
| 4767 | MIR4270 | 139 | 0.37 | 0.85 | 0.783 | 1 |
| 4768 | MIR3168 | 113 | 0.37 | 0.85 | 0.773 | 1 |
| 4769 | STAT1_01 | 60 | 0.39 | 0.85 | 0.739 | 1 |
| 4770 | MIR1587 | 51 | 0.4 | 0.85 | 0.732 | 1 |
| 4771 | REACTOME_INTERLEUKIN_12_SIGNALING | 38 | 0.41 | 0.85 | 0.712 | 1 |
| 4772 | TORCHIA_TARGETS_OF_EWSR1_FLI1_FUSION_DN | 253 | 0.35 | 0.85 | 0.831 | 1 |
| 4773 | MIR633 | 122 | 0.37 | 0.85 | 0.784 | 1 |
| 4774 | MIR595 | 51 | 0.4 | 0.85 | 0.72 | 1 |
| 4775 | MIR12133 | 332 | 0.35 | 0.85 | 0.875 | 1 |
| 4776 | YNGTTNNNATT_UNKNOWN | 295 | 0.35 | 0.85 | 0.857 | 1 |
| 4777 | MIR4643 | 182 | 0.36 | 0.85 | 0.825 | 1 |
| 4778 | MIR4266 | 73 | 0.38 | 0.85 | 0.748 | 1 |
| 4779 | GAUSSMANN_MLL_AF4_FUSION_TARGETS_B_UP | 18 | 0.47 | 0.85 | 0.681 | 1 |
| 4780 | MIR487B_5P | 90 | 0.38 | 0.85 | 0.773 | 0.999 |
| 4781 | REACTOME_SIGNALING_BY_BRAF_AND_RAF1_FU | 57 | 0.4 | 0.85 | 0.741 | 0.999 |
| 4782 | REACTOME_ANTIGEN_ACTIVATES_B_CELL_RECE | 27 | 0.44 | 0.85 | 0.703 | 0.999 |

| | | | | | | |
|---|---|---|---|---|---|---|
| 4783 | PBX1_02 | 91 | 0.38 | 0.85 | 0.76 | 1 |
| 4784 | MIR3942_5P | 177 | 0.36 | 0.85 | 0.819 | 1 |
| 4785 | NOUZOVA_METHYLATED_IN_APL | 51 | 0.41 | 0.85 | 0.743 | 1 |
| 4786 | MIR323B_3P | 51 | 0.4 | 0.85 | 0.753 | 1 |
| 4787 | WP_WNT_SIGNALING_PATHWAY | 48 | 0.41 | 0.85 | 0.735 | 1 |
| 4788 | WP_DNA_IRDAMAGE_AND_CELLULAR_RESPONSE | 81 | 0.38 | 0.85 | 0.764 | 1 |
| 4789 | MIR4741 | 41 | 0.41 | 0.85 | 0.739 | 1 |
| 4790 | SANSOM_APC_MYC_TARGETS | 231 | 0.35 | 0.85 | 0.851 | 1 |
| 4791 | MIR767_3P | 112 | 0.37 | 0.85 | 0.796 | 1 |
| 4792 | NOUZOVA_TRETINOIN_AND_H4_ACETYLATION | 134 | 0.36 | 0.85 | 0.775 | 1 |
| 4793 | MIR4689 | 65 | 0.38 | 0.85 | 0.751 | 1 |
| 4794 | TAAYNRNNTCC_UNKNOWN | 134 | 0.37 | 0.85 | 0.792 | 1 |
| 4795 | WP_IL1_SIGNALING_PATHWAY | 50 | 0.4 | 0.85 | 0.741 | 1 |
| 4796 | MIR3689D | 85 | 0.38 | 0.85 | 0.738 | 1 |
| 4797 | LBP1_Q6 | 178 | 0.36 | 0.85 | 0.823 | 1 |
| 4798 | ABRAMSON_INTERACT_WITH_AIRE | 43 | 0.41 | 0.85 | 0.722 | 1 |
| 4799 | MASSARWEH_TAMOXIFEN_RESISTANCE_DN | 212 | 0.35 | 0.85 | 0.846 | 1 |
| 4800 | TONKS_TARGETS_OF_RUNX1_RUNX1T1_FUSION_ | 42 | 0.41 | 0.85 | 0.721 | 1 |
| 4801 | WP_WNTBETACATENIN_SIGNALING_PATHWAY_IN_ | 21 | 0.45 | 0.85 | 0.696 | 1 |
| 4802 | MIR511_3P | 128 | 0.36 | 0.85 | 0.785 | 1 |
| 4803 | REACTOME_METABOLISM_OF_CARBOHYDRATES | 242 | 0.35 | 0.85 | 0.845 | 1 |
| 4804 | MIR606 | 82 | 0.38 | 0.85 | 0.75 | 1 |
| 4805 | MIR190B_3P | 121 | 0.37 | 0.85 | 0.79 | 1 |
| 4806 | MYOCD_TARGET_GENES | 420 | 0.35 | 0.85 | 0.9 | 1 |
| 4807 | SANA_RESPONSE_TO_IFNG_DN | 74 | 0.39 | 0.85 | 0.752 | 1 |
| 4808 | MIR185_5P | 242 | 0.35 | 0.85 | 0.854 | 1 |
| 4809 | MIR6788_5P | 102 | 0.38 | 0.85 | 0.779 | 1 |
| 4810 | CTCTAGA_MIR526C_MIR518F_MIR526A | 51 | 0.4 | 0.85 | 0.727 | 1 |
| 4811 | OCT_C | 192 | 0.36 | 0.85 | 0.835 | 1 |
| 4812 | MIR1293 | 53 | 0.4 | 0.85 | 0.735 | 1 |
| 4813 | GARGALOVIC_RESPONSE_TO_OXIDIZED_PHOSPH | 55 | 0.39 | 0.85 | 0.751 | 1 |
| 4814 | REACTOME_TGF_BETA_RECEPTOR_SIGNALING_A | 30 | 0.42 | 0.85 | 0.723 | 1 |
| 4815 | WP_ESTROGEN_SIGNALING_PATHWAY | 21 | 0.46 | 0.85 | 0.733 | 1 |

| | | | | | | |
|---|---|---|---|---|---|---|
| 4816 | HNF4_Q6 | 209 | 0.36 | 0.85 | 0.831 | 1 |
| 4817 | GR_Q6_01 | 220 | 0.35 | 0.85 | 0.842 | 1 |
| 4818 | NIKOLSKY_BREAST_CANCER_5P15_AMPLICON | 25 | 0.44 | 0.85 | 0.711 | 1 |
| 4819 | MIR4675 | 40 | 0.42 | 0.85 | 0.726 | 1 |
| 4820 | MIR193A_3P_MIR193B_3P | 74 | 0.39 | 0.85 | 0.777 | 1 |
| 4821 | AAAGGAT_MIR501 | 116 | 0.37 | 0.84 | 0.792 | 1 |
| 4822 | AMIT_EGF_RESPONSE_240_HELA | 51 | 0.4 | 0.84 | 0.75 | 1 |
| 4823 | MIR1199_5P | 64 | 0.39 | 0.84 | 0.737 | 1 |
| 4824 | VILLANUEVA_LIVER_CANCER_KRT19_UP | 164 | 0.36 | 0.84 | 0.822 | 1 |
| 4825 | MIR3065_5P | 408 | 0.35 | 0.84 | 0.901 | 1 |
| 4826 | CHX10_01 | 158 | 0.36 | 0.84 | 0.811 | 1 |
| 4827 | MIKKELSEN_MEF_HCP_WITH_H3K27ME3 | 358 | 0.34 | 0.84 | 0.887 | 1 |
| 4828 | MIR26A_5P | 444 | 0.34 | 0.84 | 0.91 | 1 |
| 4829 | MIR4436B_5P | 128 | 0.36 | 0.84 | 0.787 | 1 |
| 4830 | TCCCCAC_MIR491 | 53 | 0.4 | 0.84 | 0.737 | 1 |
| 4831 | MIR6882_3P | 98 | 0.37 | 0.84 | 0.76 | 1 |
| 4832 | MIR6768_5P | 173 | 0.36 | 0.84 | 0.836 | 1 |
| 4833 | MIR6077 | 192 | 0.36 | 0.84 | 0.859 | 1 |
| 4834 | BIOCARTA_MAPK_PATHWAY | 80 | 0.38 | 0.84 | 0.769 | 1 |
| 4835 | GATA2_01 | 95 | 0.37 | 0.84 | 0.774 | 1 |
| 4836 | MIR2115_5P | 119 | 0.37 | 0.84 | 0.781 | 1 |
| 4837 | P53_02 | 211 | 0.35 | 0.84 | 0.845 | 1 |
| 4838 | TASOR_TARGET_GENES | 255 | 0.35 | 0.84 | 0.855 | 1 |
| 4839 | MCM2_TARGET_GENES | 56 | 0.39 | 0.84 | 0.773 | 1 |
| 4840 | MIR223_5P | 130 | 0.36 | 0.84 | 0.804 | 1 |
| 4841 | CHENG_IMPRINTED_BY_ESTRADIOL | 98 | 0.37 | 0.84 | 0.754 | 1 |
| 4842 | GATA1_01 | 205 | 0.35 | 0.84 | 0.842 | 1 |
| 4843 | BANDRES_RESPONSE_TO_CARMUSTIN_MGMT_48 | 116 | 0.37 | 0.84 | 0.805 | 1 |
| 4844 | MIR6503_5P | 50 | 0.4 | 0.84 | 0.744 | 1 |
| 4845 | MIR12130 | 24 | 0.45 | 0.84 | 0.712 | 1 |
| 4846 | REACTOME_TRANSCRIPTIONAL_ACTIVATION_OF_ | 53 | 0.39 | 0.84 | 0.736 | 1 |
| 4847 | MIR186_5P | 478 | 0.34 | 0.84 | 0.921 | 1 |
| 4848 | BURTON_ADIPOGENESIS_2 | 59 | 0.39 | 0.84 | 0.778 | 1 |

| | | | | | | |
|---|---|---|---|---|---|---|
| 4849 | MIR210_5P | 44 | 0.4 | 0.84 | 0.726 | 1 |
| 4850 | MIR3130_3P | 43 | 0.4 | 0.84 | 0.729 | 1 |
| 4851 | MIR26B_5P | 444 | 0.34 | 0.84 | 0.921 | 1 |
| 4852 | YAMAZAKI_TCEB3_TARGETS_DN | 175 | 0.35 | 0.84 | 0.834 | 1 |
| 4853 | MIR3182 | 225 | 0.35 | 0.84 | 0.855 | 1 |
| 4854 | MCBRYAN_PUBERTAL_BREAST_5_6WK_DN | 117 | 0.36 | 0.84 | 0.79 | 1 |
| 4855 | ZIC3_01 | 215 | 0.35 | 0.84 | 0.855 | 1 |
| 4856 | ALX4_TARGET_GENES | 26 | 0.44 | 0.84 | 0.72 | 1 |
| 4857 | YBX1_TARGET_GENES | 60 | 0.39 | 0.84 | 0.765 | 1 |
| 4858 | BLANCO_MELO_INFLUENZA_A_INFECTION_A594_( | 115 | 0.36 | 0.84 | 0.814 | 1 |
| 4859 | MIR3651 | 15 | 0.47 | 0.84 | 0.711 | 1 |
| 4860 | REACTOME_RUNX2_REGULATES_OSTEOBLAST_D | 17 | 0.46 | 0.84 | 0.734 | 1 |
| 4861 | MIR4644 | 251 | 0.35 | 0.84 | 0.868 | 1 |
| 4862 | CREBP1_01 | 161 | 0.35 | 0.84 | 0.814 | 1 |
| 4863 | MIR1261 | 66 | 0.39 | 0.84 | 0.747 | 1 |
| 4864 | MIR8067 | 246 | 0.35 | 0.84 | 0.854 | 1 |
| 4865 | KYNG_ENVIRONMENTAL_STRESS_RESPONSE_UP | 46 | 0.41 | 0.84 | 0.74 | 1 |
| 4866 | MIR342_3P | 108 | 0.37 | 0.84 | 0.784 | 1 |
| 4867 | BERNARD_PPAPDC1B_TARGETS_UP | 35 | 0.41 | 0.84 | 0.723 | 1 |
| 4868 | LU_AGING_BRAIN_UP | 203 | 0.35 | 0.84 | 0.853 | 1 |
| 4869 | MIR5689 | 135 | 0.36 | 0.84 | 0.815 | 1 |
| 4870 | CHUNG_BLISTER_CYTOTOXICITY_UP | 121 | 0.36 | 0.84 | 0.82 | 1 |
| 4871 | REACTOME_DAP12_INTERACTIONS | 24 | 0.44 | 0.84 | 0.728 | 1 |
| 4872 | STEIN_ESRRA_TARGETS | 494 | 0.34 | 0.84 | 0.936 | 1 |
| 4873 | REACTOME_NUCLEOTIDE_SALVAGE | 20 | 0.44 | 0.84 | 0.717 | 1 |
| 4874 | MIR1246 | 47 | 0.4 | 0.84 | 0.736 | 1 |
| 4875 | TCANNTGAY_SREBP1_01 | 408 | 0.34 | 0.84 | 0.902 | 1 |
| 4876 | PID_BCR_5PATHWAY | 53 | 0.4 | 0.84 | 0.745 | 1 |
| 4877 | MIR3145_3P | 182 | 0.36 | 0.84 | 0.839 | 1 |
| 4878 | MIR3150B_3P | 119 | 0.37 | 0.84 | 0.817 | 1 |
| 4879 | MIR4424 | 98 | 0.37 | 0.84 | 0.78 | 1 |
| 4880 | MIR302A_3P_MIR302B_3P_MIR302C_3P_MIR302D_ | 314 | 0.35 | 0.84 | 0.89 | 1 |
| 4881 | MIR4498 | 221 | 0.35 | 0.84 | 0.844 | 1 |

| | | | | | | |
|---|---|---|---|---|---|---|
| 4882 | DUTERTRE_ESTRADIOL_RESPONSE_6HR_DN | 77 | 0.38 | 0.84 | 0.766 | 1 |
| 4883 | ACTACCT_MIR196A_MIR196B | 124 | 0.36 | 0.84 | 0.809 | 1 |
| 4884 | REACTOME_REGULATION_OF_BETA_CELL_DEVEI | 26 | 0.44 | 0.84 | 0.728 | 1 |
| 4885 | AAGCAAT_MIR137 | 196 | 0.35 | 0.84 | 0.866 | 1 |
| 4886 | MIR629_5P | 82 | 0.37 | 0.84 | 0.762 | 1 |
| 4887 | MIR3200_5P | 58 | 0.39 | 0.84 | 0.733 | 1 |
| 4888 | REACTOME_CA_DEPENDENT_EVENTS | 30 | 0.42 | 0.84 | 0.711 | 1 |
| 4889 | REN_ALVEOLAR_RHABDOMYOSARCOMA_UP | 83 | 0.37 | 0.84 | 0.786 | 1 |
| 4890 | MIR585_5P | 114 | 0.36 | 0.84 | 0.796 | 1 |
| 4891 | MORI_PLASMA_CELL_UP | 49 | 0.39 | 0.84 | 0.727 | 1 |
| 4892 | MIR4490 | 84 | 0.37 | 0.84 | 0.776 | 1 |
| 4893 | FOXM1_01 | 197 | 0.35 | 0.84 | 0.849 | 1 |
| 4894 | REACTOME_POST_TRANSLATIONAL_MODIFICATIC | 54 | 0.39 | 0.84 | 0.747 | 1 |
| 4895 | JAATINEN_HEMATOPOIETIC_STEM_CELL_UP | 261 | 0.35 | 0.84 | 0.866 | 1 |
| 4896 | SQSTM1_TARGET_GENES | 38 | 0.41 | 0.84 | 0.741 | 1 |
| 4897 | MIR6499_3P | 110 | 0.37 | 0.84 | 0.804 | 1 |
| 4898 | SMTTTTGT_UNKNOWN | 352 | 0.34 | 0.84 | 0.891 | 1 |
| 4899 | REACTOME_REGULATION_OF_HSF1_MEDIATED_H | 75 | 0.38 | 0.84 | 0.781 | 1 |
| 4900 | ZNF577_TARGET_GENES | 412 | 0.34 | 0.83 | 0.923 | 1 |
| 4901 | WP_APOPTOSIS_MODULATION_BY_HSP70 | 18 | 0.46 | 0.83 | 0.721 | 1 |
| 4902 | KEGG_VASCULAR_SMOOTH_MUSCLE_CONTRACT | 76 | 0.38 | 0.83 | 0.78 | 1 |
| 4903 | MIR3925_5P | 128 | 0.36 | 0.83 | 0.822 | 1 |
| 4904 | MIR6751_3P | 64 | 0.39 | 0.83 | 0.779 | 1 |
| 4905 | WU_APOPTOSIS_BY_CDKN1A_VIA_TP53 | 50 | 0.4 | 0.83 | 0.749 | 1 |
| 4906 | MIR218_1_3P | 80 | 0.38 | 0.83 | 0.795 | 1 |
| 4907 | MIR892A | 111 | 0.37 | 0.83 | 0.806 | 1 |
| 4908 | MIR3173_5P | 93 | 0.37 | 0.83 | 0.775 | 1 |
| 4909 | MIR4699_3P | 411 | 0.34 | 0.83 | 0.921 | 1 |
| 4910 | WP_ENVELOPE_PROTEINS_AND_THEIR_POTENTI/ | 42 | 0.4 | 0.83 | 0.748 | 1 |
| 4911 | MIR450B_5P | 408 | 0.34 | 0.83 | 0.923 | 1 |
| 4912 | SHIPP_DLBCL_CURED_VS_FATAL_UP | 25 | 0.44 | 0.83 | 0.732 | 1 |
| 4913 | MIR591 | 41 | 0.4 | 0.83 | 0.747 | 1 |
| 4914 | MIR631 | 40 | 0.41 | 0.83 | 0.76 | 1 |

| | | | | | | |
|---|---|---|---|---|---|---|
| 4915 | MIR4306 | 390 | 0.34 | 0.83 | 0.924 | 1 |
| 4916 | MIR4776_3P | 130 | 0.36 | 0.83 | 0.8 | 1 |
| 4917 | ZBTB44_TARGET_GENES | 286 | 0.35 | 0.83 | 0.87 | 1 |
| 4918 | PID_PDGFRB_PATHWAY | 121 | 0.36 | 0.83 | 0.817 | 1 |
| 4919 | MIER1_TARGET_GENES | 296 | 0.34 | 0.83 | 0.882 | 1 |
| 4920 | MIR4448 | 59 | 0.39 | 0.83 | 0.794 | 1 |
| 4921 | MYOD_01 | 223 | 0.35 | 0.83 | 0.864 | 1 |
| 4922 | ZBED4_TARGET_GENES | 162 | 0.35 | 0.83 | 0.844 | 1 |
| 4923 | ZIRN_TRETINOIN_RESPONSE_WT1_UP | 17 | 0.47 | 0.83 | 0.715 | 1 |
| 4924 | GTATGAT_MIR154_MIR487 | 68 | 0.38 | 0.83 | 0.782 | 1 |
| 4925 | MIR5690 | 103 | 0.36 | 0.83 | 0.799 | 1 |
| 4926 | CEBALLOS_TARGETS_OF_TP53_AND_MYC_DN | 29 | 0.42 | 0.83 | 0.747 | 1 |
| 4927 | MIR559 | 478 | 0.34 | 0.83 | 0.934 | 1 |
| 4928 | MIR490_3P | 44 | 0.4 | 0.83 | 0.757 | 1 |
| 4929 | LEE_METASTASIS_AND_ALTERNATIVE_SPLICING_ | 56 | 0.39 | 0.83 | 0.762 | 1 |
| 4930 | MIR550B_2_5P | 40 | 0.4 | 0.83 | 0.742 | 1 |
| 4931 | AR_Q2 | 93 | 0.37 | 0.83 | 0.805 | 1 |
| 4932 | TTTTGAG_MIR373 | 208 | 0.35 | 0.83 | 0.856 | 1 |
| 4933 | REACTOME_OPIOID_SIGNALLING | 72 | 0.38 | 0.83 | 0.773 | 1 |
| 4934 | MIR4668_3P | 271 | 0.34 | 0.83 | 0.882 | 1 |
| 4935 | MIR4511 | 140 | 0.36 | 0.83 | 0.835 | 1 |
| 4936 | MIR7162_3P | 190 | 0.36 | 0.83 | 0.851 | 1 |
| 4937 | BURTON_ADIPOGENESIS_5 | 114 | 0.36 | 0.83 | 0.792 | 1 |
| 4938 | MIR6504_5P | 129 | 0.36 | 0.83 | 0.835 | 1 |
| 4939 | ZWANG_DOWN_BY_2ND_EGF_PULSE | 236 | 0.35 | 0.83 | 0.887 | 1 |
| 4940 | GRATIAS_RETINOBLASTOMA_16Q24 | 18 | 0.46 | 0.83 | 0.732 | 1 |
| 4941 | PID_FOXO_PATHWAY | 46 | 0.4 | 0.83 | 0.761 | 1 |
| 4942 | MIR3685 | 158 | 0.35 | 0.83 | 0.839 | 1 |
| 4943 | TGACCTTG_SF1_Q6 | 220 | 0.35 | 0.83 | 0.86 | 1 |
| 4944 | MIR4713_5P | 133 | 0.36 | 0.83 | 0.83 | 1 |
| 4945 | DOANE_BREAST_CANCER_CLASSES_UP | 45 | 0.4 | 0.83 | 0.754 | 1 |
| 4946 | EGR1_01 | 248 | 0.34 | 0.83 | 0.874 | 1 |
| 4947 | WP_ECTODERM_DIFFERENTIATION | 115 | 0.36 | 0.83 | 0.808 | 1 |

| | | | | | | |
|---|---|---|---|---|---|---|
| 4948 | RFX1_02 | 228 | 0.34 | 0.83 | 0.866 | 1 |
| 4949 | MIR3680_3P | 454 | 0.34 | 0.83 | 0.937 | 1 |
| 4950 | TAKADA_GASTRIC_CANCER_COPY_NUMBER_DN | 21 | 0.44 | 0.83 | 0.75 | 1 |
| 4951 | MIR491_3P | 98 | 0.37 | 0.83 | 0.803 | 1 |
| 4952 | CTCAGGG_MIR125B_MIR125A | 300 | 0.34 | 0.83 | 0.894 | 1 |
| 4953 | SMARCA1_TARGET_GENES | 109 | 0.36 | 0.83 | 0.807 | 1 |
| 4954 | OSMAN_BLADDER_CANCER_UP | 370 | 0.34 | 0.83 | 0.91 | 1 |
| 4955 | MIR3064_5P | 133 | 0.36 | 0.83 | 0.829 | 1 |
| 4956 | MIR577 | 273 | 0.34 | 0.83 | 0.886 | 1 |
| 4957 | PETRETTO_HEART_MASS_QTL_CIS_UP | 29 | 0.41 | 0.83 | 0.74 | 1 |
| 4958 | TTGCACT_MIR130A_MIR301_MIR130B | 368 | 0.34 | 0.83 | 0.913 | 1 |
| 4959 | MIR548AP_5P_MIR548J_5P | 422 | 0.34 | 0.83 | 0.92 | 1 |
| 4960 | MIR4438 | 60 | 0.38 | 0.83 | 0.77 | 1 |
| 4961 | NME2_TARGET_GENES | 36 | 0.41 | 0.83 | 0.755 | 1 |
| 4962 | POMEROY_MEDULLOBLASTOMA_PROGNOSIS_UP | 40 | 0.4 | 0.83 | 0.733 | 1 |
| 4963 | REACTOME_GENE_AND_PROTEIN_EXPRESSION_I | 33 | 0.41 | 0.83 | 0.747 | 1 |
| 4964 | MIR548AB | 425 | 0.34 | 0.83 | 0.926 | 1 |
| 4965 | REACTOME_NR1H3_NR1H2_REGULATE_GENE_EX | 32 | 0.42 | 0.83 | 0.752 | 1 |
| 4966 | IVANOVA_HEMATOPOIESIS_EARLY_PROGENITOR | 402 | 0.34 | 0.83 | 0.915 | 1 |
| 4967 | REACTOME_RHOB_GTPASE_CYCLE | 68 | 0.38 | 0.83 | 0.798 | 1 |
| 4968 | MIR7106_5P | 304 | 0.34 | 0.83 | 0.908 | 1 |
| 4969 | WP_GENES_CONTROLLING_NEPHROGENESIS | 30 | 0.43 | 0.83 | 0.756 | 1 |
| 4970 | TEF_Q6 | 199 | 0.35 | 0.83 | 0.852 | 1 |
| 4971 | ATATGCA_MIR448 | 205 | 0.35 | 0.83 | 0.892 | 1 |
| 4972 | DR1_Q3 | 216 | 0.34 | 0.83 | 0.868 | 1 |
| 4973 | PR_Q2 | 202 | 0.35 | 0.83 | 0.85 | 1 |
| 4974 | MIKKELSEN_MCV6_HCP_WITH_H3K27ME3 | 291 | 0.34 | 0.82 | 0.901 | 1 |
| 4975 | MIR548BB_5P | 427 | 0.34 | 0.82 | 0.926 | 1 |
| 4976 | CATTGTYY_SOX9_B1 | 301 | 0.34 | 0.82 | 0.902 | 1 |
| 4977 | JIANG_HYPOXIA_NORMAL | 186 | 0.35 | 0.82 | 0.856 | 1 |
| 4978 | ATF5_TARGET_GENES | 376 | 0.34 | 0.82 | 0.919 | 1 |
| 4979 | MIR548I | 423 | 0.34 | 0.82 | 0.941 | 1 |
| 4980 | MIR302D_5P | 167 | 0.35 | 0.82 | 0.848 | 1 |

| | | | | | | |
|---|---|---|---|---|---|---|
| 4981 | MIR1290 | 215 | 0.35 | 0.82 | 0.868 | 1 |
| 4982 | BHAT_ESR1_TARGETS_VIA_AKT1_DN | 71 | 0.37 | 0.82 | 0.794 | 1 |
| 4983 | MIR138_5P | 184 | 0.35 | 0.82 | 0.864 | 1 |
| 4984 | MIR548AR_5P | 430 | 0.34 | 0.82 | 0.935 | 1 |
| 4985 | MIR4725_5P | 47 | 0.39 | 0.82 | 0.755 | 1 |
| 4986 | ZNF547_TARGET_GENES | 125 | 0.36 | 0.82 | 0.836 | 1 |
| 4987 | PCGF6_TARGET_GENES | 65 | 0.38 | 0.82 | 0.766 | 1 |
| 4988 | MIR4477A | 328 | 0.34 | 0.82 | 0.914 | 1 |
| 4989 | CAGTATT_MIR200B_MIR200C_MIR429 | 426 | 0.34 | 0.82 | 0.94 | 1 |
| 4990 | FOURNIER_ACINAR_DEVELOPMENT_EARLY_UP | 18 | 0.46 | 0.82 | 0.729 | 1 |
| 4991 | E47_02 | 204 | 0.35 | 0.82 | 0.871 | 1 |
| 4992 | ZNF37A_TARGET_GENES | 122 | 0.36 | 0.82 | 0.823 | 1 |
| 4993 | E2A_Q2 | 210 | 0.34 | 0.82 | 0.863 | 1 |
| 4994 | BIOCARTA_IGF1_PATHWAY | 19 | 0.45 | 0.82 | 0.733 | 1 |
| 4995 | MIR510_3P | 320 | 0.34 | 0.82 | 0.904 | 1 |
| 4996 | TATA_C | 191 | 0.35 | 0.82 | 0.874 | 1 |
| 4997 | MIR548AQ_5P | 426 | 0.34 | 0.82 | 0.931 | 1 |
| 4998 | MIR6756_5P | 213 | 0.34 | 0.82 | 0.872 | 1 |
| 4999 | MIR8070 | 110 | 0.36 | 0.82 | 0.831 | 1 |
| 5000 | MIR6791_5P | 147 | 0.35 | 0.82 | 0.86 | 1 |
| 5001 | E2F1_Q4 | 229 | 0.34 | 0.82 | 0.883 | 1 |
| 5002 | FOXJ2_TARGET_GENES | 467 | 0.34 | 0.82 | 0.938 | 1 |
| 5003 | MIR6508_3P | 38 | 0.4 | 0.82 | 0.759 | 1 |
| 5004 | WANG_ADIPOGENIC_GENES_REPRESSED_BY_SII | 23 | 0.44 | 0.82 | 0.742 | 1 |
| 5005 | JI_RESPONSE_TO_FSH_DN | 50 | 0.39 | 0.82 | 0.788 | 1 |
| 5006 | VDR_Q6 | 219 | 0.34 | 0.82 | 0.87 | 1 |
| 5007 | COUP_01 | 206 | 0.34 | 0.82 | 0.858 | 1 |
| 5008 | MIR548O_5P_MIR548W | 424 | 0.34 | 0.82 | 0.932 | 1 |
| 5009 | MIR548AS_5P | 423 | 0.34 | 0.82 | 0.924 | 1 |
| 5010 | CTCCAAG_MIR432 | 72 | 0.37 | 0.82 | 0.81 | 1 |
| 5011 | MIR548AK_MIR548AM_5P_MIR548C_5P_MIR548H_ | 424 | 0.34 | 0.82 | 0.942 | 1 |
| 5012 | WENG_POR_TARGETS_GLOBAL_DN | 18 | 0.45 | 0.82 | 0.739 | 1 |
| 5013 | MIR548Y | 431 | 0.34 | 0.82 | 0.939 | 1 |

| | | | | | | |
|---|---|---|---|---|---|---|
| 5014 | MIR548AD_5P_MIR548AE_5P_MIR548AY_5P_MIR5 | 425 | 0.34 | 0.82 | 0.939 | 1 |
| 5015 | MIR4533 | 272 | 0.34 | 0.82 | 0.892 | 1 |
| 5016 | BENPORATH_ES_1 | 335 | 0.34 | 0.82 | 0.935 | 1 |
| 5017 | MIR198 | 79 | 0.37 | 0.82 | 0.793 | 1 |
| 5018 | SU_TESTIS | 58 | 0.37 | 0.82 | 0.775 | 1 |
| 5019 | MIR4458 | 308 | 0.34 | 0.82 | 0.912 | 1 |
| 5020 | BIOCARTA_AKT_PATHWAY | 18 | 0.45 | 0.82 | 0.74 | 1 |
| 5021 | MIR548A_5P | 424 | 0.34 | 0.82 | 0.932 | 1 |
| 5022 | MIR6739_3P | 156 | 0.35 | 0.82 | 0.858 | 1 |
| 5023 | MIR6844 | 272 | 0.34 | 0.82 | 0.892 | 1 |
| 5024 | TERAO_AOX4_TARGETS_HG_UP | 27 | 0.43 | 0.82 | 0.763 | 1 |
| 5025 | MIR200C_3P | 454 | 0.33 | 0.82 | 0.932 | 1 |
| 5026 | WP_TUMOR_SUPPRESSOR_ACTIVITY_OF_SMARCI | 28 | 0.42 | 0.82 | 0.758 | 1 |
| 5027 | WP_VITAMIN_A_AND_CAROTENOID_METABOLISM | 29 | 0.41 | 0.82 | 0.756 | 1 |
| 5028 | MIR3918 | 79 | 0.37 | 0.82 | 0.797 | 1 |
| 5029 | MIR204_5P | 319 | 0.34 | 0.82 | 0.906 | 1 |
| 5030 | LIU_LIVER_CANCER | 27 | 0.43 | 0.82 | 0.759 | 1 |
| 5031 | AMEF2_Q6 | 192 | 0.34 | 0.82 | 0.864 | 1 |
| 5032 | MARIADASON_REGULATED_BY_HISTONE_ACETYL | 37 | 0.4 | 0.82 | 0.76 | 1 |
| 5033 | MIR4251 | 333 | 0.33 | 0.82 | 0.919 | 1 |
| 5034 | MIR6783_5P | 129 | 0.35 | 0.82 | 0.832 | 1 |
| 5035 | SEMENZA_HIF1_TARGETS | 30 | 0.42 | 0.82 | 0.759 | 1 |
| 5036 | HASLINGER_B_CLL_WITH_17P13_DELETION | 22 | 0.43 | 0.82 | 0.741 | 1 |
| 5037 | SYATTGTG_UNKNOWN | 178 | 0.34 | 0.82 | 0.869 | 1 |
| 5038 | BROWNE_HCMV_INFECTION_20HR_UP | 216 | 0.34 | 0.82 | 0.894 | 1 |
| 5039 | BOYAULT_LIVER_CANCER_SUBCLASS_G1_UP | 112 | 0.36 | 0.82 | 0.829 | 1 |
| 5040 | RAHMAN_TP53_TARGETS_PHOSPHORYLATED | 20 | 0.44 | 0.82 | 0.744 | 1 |
| 5041 | ALONSO_METASTASIS_UP | 169 | 0.35 | 0.82 | 0.877 | 1 |
| 5042 | REACTOME_SYNTHESIS_OF_DNA | 118 | 0.36 | 0.82 | 0.849 | 1 |
| 5043 | MAZR_01 | 194 | 0.34 | 0.82 | 0.875 | 1 |
| 5044 | MIR6085 | 37 | 0.4 | 0.82 | 0.791 | 1 |
| 5045 | MIR330_5P | 156 | 0.35 | 0.82 | 0.854 | 1 |
| 5046 | MAPK3_TARGET_GENES | 54 | 0.38 | 0.82 | 0.777 | 1 |

| | | | | | | |
|---|---|---|---|---|---|---|
| 5047 | ZNF560_TARGET_GENES | 43 | 0.4 | 0.82 | 0.765 | 1 |
| 5048 | MIR335_5P | 92 | 0.37 | 0.81 | 0.798 | 1 |
| 5049 | ACTTTAT_MIR1425P | 264 | 0.34 | 0.81 | 0.904 | 1 |
| 5050 | TGCACTT_MIR519C_MIR519B_MIR519A | 414 | 0.33 | 0.81 | 0.934 | 1 |
| 5051 | SMCHD1_TARGET_GENES | 40 | 0.4 | 0.81 | 0.776 | 1 |
| 5052 | MIR642A_5P | 71 | 0.37 | 0.81 | 0.799 | 1 |
| 5053 | AAGWWRNYGGC_UNKNOWN | 116 | 0.35 | 0.81 | 0.842 | 1 |
| 5054 | FAELT_B_CLL_WITH_VH3_21_DN | 47 | 0.38 | 0.81 | 0.765 | 1 |
| 5055 | MIR200B_3P | 455 | 0.33 | 0.81 | 0.943 | 1 |
| 5056 | LOPEZ_TRANSLATION_VIA_FN1_SIGNALING | 31 | 0.41 | 0.81 | 0.774 | 1 |
| 5057 | MIR21_5P | 119 | 0.35 | 0.81 | 0.822 | 1 |
| 5058 | MIR7975 | 40 | 0.4 | 0.81 | 0.767 | 1 |
| 5059 | MIR1305 | 445 | 0.33 | 0.81 | 0.943 | 1 |
| 5060 | ZNF816_TARGET_GENES | 384 | 0.34 | 0.81 | 0.939 | 1 |
| 5061 | MIR1915_3P | 143 | 0.35 | 0.81 | 0.864 | 1 |
| 5062 | OCT1_06 | 194 | 0.34 | 0.81 | 0.884 | 1 |
| 5063 | MIR3117_5P | 49 | 0.39 | 0.81 | 0.772 | 1 |
| 5064 | SRY_01 | 179 | 0.34 | 0.81 | 0.859 | 1 |
| 5065 | WP_DISRUPTION_OF_POSTSYNAPTIC_SIGNALING | 28 | 0.41 | 0.81 | 0.768 | 1 |
| 5066 | RICKMAN_TUMOR_DIFFERENTIATED_MODERATEL` | 15 | 0.46 | 0.81 | 0.748 | 1 |
| 5067 | GLI1_TARGET_GENES | 401 | 0.33 | 0.81 | 0.941 | 1 |
| 5068 | ENGELMANN_CANCER_PROGENITORS_UP | 42 | 0.4 | 0.81 | 0.772 | 1 |
| 5069 | PROVENZANI_METASTASIS_UP | 184 | 0.34 | 0.81 | 0.886 | 1 |
| 5070 | MIR3194_3P | 84 | 0.36 | 0.81 | 0.816 | 1 |
| 5071 | MIR5007_5P | 48 | 0.39 | 0.81 | 0.793 | 1 |
| 5072 | E2F1_Q6_01 | 234 | 0.34 | 0.81 | 0.891 | 1 |
| 5073 | CTTGTAT_MIR381 | 189 | 0.34 | 0.81 | 0.893 | 1 |
| 5074 | GREENBAUM_E2A_TARGETS_DN | 15 | 0.46 | 0.81 | 0.782 | 1 |
| 5075 | GGGNRMNNYCAT_UNKNOWN | 74 | 0.37 | 0.81 | 0.803 | 1 |
| 5076 | LIN_NPAS4_TARGETS_DN | 55 | 0.38 | 0.81 | 0.784 | 1 |
| 5077 | FORTSCHEGGER_PHF8_TARGETS_UP | 238 | 0.34 | 0.81 | 0.895 | 1 |
| 5078 | MIR105_5P | 197 | 0.34 | 0.81 | 0.885 | 1 |
| 5079 | REACTOME_NEGATIVE_REGULATION_OF_FGFR2_ | 22 | 0.43 | 0.81 | 0.744 | 1 |

| | | | | | | |
|---|---|---|---|---|---|---|
| 5080 | MEISSNER_NPC_HCP_WITH_H3K4ME2_AND_H3K2 | 220 | 0.34 | 0.81 | 0.891 | 1 |
| 5081 | MIR3143 | 385 | 0.33 | 0.81 | 0.937 | 1 |
| 5082 | MIR371A_5P | 161 | 0.35 | 0.81 | 0.863 | 1 |
| 5083 | ACCTGTTG_UNKNOWN | 136 | 0.34 | 0.81 | 0.839 | 1 |
| 5084 | POU3F2_02 | 201 | 0.34 | 0.81 | 0.886 | 1 |
| 5085 | MIR758_3P | 103 | 0.36 | 0.81 | 0.834 | 1 |
| 5086 | MIR582_3P | 84 | 0.36 | 0.81 | 0.827 | 1 |
| 5087 | MIR1298_5P | 35 | 0.4 | 0.81 | 0.764 | 1 |
| 5088 | EVI1_02 | 90 | 0.36 | 0.81 | 0.81 | 1 |
| 5089 | MIR4328 | 492 | 0.33 | 0.81 | 0.959 | 1 |
| 5090 | CREIGHTON_ENDOCRINE_THERAPY_RESISTANCE | 423 | 0.33 | 0.81 | 0.937 | 1 |
| 5091 | MIR6873_5P | 85 | 0.36 | 0.81 | 0.818 | 1 |
| 5092 | NEWMAN_ERCC6_TARGETS_DN | 22 | 0.43 | 0.81 | 0.732 | 1 |
| 5093 | REACTOME_HS_GAG_DEGRADATION | 20 | 0.43 | 0.81 | 0.747 | 1 |
| 5094 | LET_7I_5P | 356 | 0.33 | 0.81 | 0.928 | 1 |
| 5095 | SP1_Q6_01 | 229 | 0.34 | 0.81 | 0.905 | 1 |
| 5096 | ZHANG_RESPONSE_TO_IKK_INHIBITOR_AND_TNF | 94 | 0.35 | 0.81 | 0.841 | 1 |
| 5097 | MIR1275 | 244 | 0.34 | 0.81 | 0.904 | 1 |
| 5098 | MIR6744_5P | 35 | 0.4 | 0.81 | 0.774 | 1 |
| 5099 | MIR147B_5P | 274 | 0.33 | 0.81 | 0.916 | 1 |
| 5100 | CAGTGTT_MIR141_MIR200A | 292 | 0.33 | 0.81 | 0.92 | 1 |
| 5101 | MIR1299 | 287 | 0.33 | 0.81 | 0.915 | 1 |
| 5102 | REACTOME_RUNX2_REGULATES_BONE_DEVELOI | 22 | 0.43 | 0.81 | 0.769 | 1 |
| 5103 | KEGG_GLUTATHIONE_METABOLISM | 37 | 0.4 | 0.81 | 0.782 | 1 |
| 5104 | MIR211_5P | 318 | 0.33 | 0.81 | 0.921 | 1 |
| 5105 | YTAAYNGCT_UNKNOWN | 121 | 0.35 | 0.81 | 0.86 | 1 |
| 5106 | REACTOME_TRANSCRIPTIONAL_REGULATION_BY | 51 | 0.38 | 0.81 | 0.797 | 1 |
| 5107 | MIR4677_3P | 106 | 0.35 | 0.81 | 0.85 | 1 |
| 5108 | HUANG_DASATINIB_RESISTANCE_DN | 61 | 0.38 | 0.81 | 0.815 | 1 |
| 5109 | AFP1_Q6 | 205 | 0.34 | 0.81 | 0.887 | 1 |
| 5110 | MIR568 | 223 | 0.34 | 0.81 | 0.896 | 1 |
| 5111 | MIR1178_5P | 29 | 0.42 | 0.81 | 0.775 | 1 |
| 5112 | WP_3Q29_COPY_NUMBER_VARIATION_SYNDROM | 54 | 0.38 | 0.81 | 0.793 | 1 |

| | | | | | | |
|---|---|---|---|---|---|---|
| 5113 | MIR3926 | 147 | 0.34 | 0.81 | 0.87 | 1 |
| 5114 | MIR4434 | 139 | 0.35 | 0.81 | 0.862 | 1 |
| 5115 | EHLERS_ANEUPLOIDY_UP | 37 | 0.4 | 0.81 | 0.8 | 1 |
| 5116 | MIR3944_5P | 104 | 0.36 | 0.81 | 0.838 | 1 |
| 5117 | MED16_TARGET_GENES | 28 | 0.41 | 0.8 | 0.769 | 1 |
| 5118 | MIR676_5P | 125 | 0.35 | 0.8 | 0.854 | 1 |
| 5119 | LET_7G_5P | 355 | 0.33 | 0.8 | 0.932 | 1 |
| 5120 | MCBRYAN_PUBERTAL_BREAST_4_5WK_DN | 163 | 0.34 | 0.8 | 0.892 | 1 |
| 5121 | MIR429 | 462 | 0.33 | 0.8 | 0.956 | 1 |
| 5122 | REACTOME_G_ALPHA_12_13_SIGNALLING_EVEN[illegible] | 63 | 0.37 | 0.8 | 0.812 | 1 |
| 5123 | REACTOME_CREB1_PHOSPHORYLATION_THROU[illegible] | 25 | 0.42 | 0.8 | 0.761 | 1 |
| 5124 | FOXO1_02 | 205 | 0.34 | 0.8 | 0.906 | 1 |
| 5125 | SRC_UP.V1_DN | 126 | 0.35 | 0.8 | 0.859 | 1 |
| 5126 | MIR7114_3P | 36 | 0.4 | 0.8 | 0.772 | 1 |
| 5127 | MIR4538 | 24 | 0.42 | 0.8 | 0.776 | 1 |
| 5128 | MIR508_5P | 86 | 0.36 | 0.8 | 0.846 | 1 |
| 5129 | WP_COHESIN_COMPLEX_CORNELIA_DE_LANGE_[illegible] | 35 | 0.4 | 0.8 | 0.776 | 1 |
| 5130 | PID_NFKAPPAB_ATYPICAL_PATHWAY | 17 | 0.45 | 0.8 | 0.756 | 1 |
| 5131 | MIR362_3P | 111 | 0.35 | 0.8 | 0.859 | 1 |
| 5132 | MIR3935 | 60 | 0.38 | 0.8 | 0.8 | 1 |
| 5133 | LET_7D_5P | 305 | 0.33 | 0.8 | 0.921 | 1 |
| 5134 | ZHANG_RESPONSE_TO_CANTHARIDIN_DN | 65 | 0.37 | 0.8 | 0.808 | 1 |
| 5135 | MA_MYELOID_DIFFERENTIATION_DN | 33 | 0.4 | 0.8 | 0.784 | 1 |
| 5136 | MIR374C_3P | 46 | 0.39 | 0.8 | 0.766 | 1 |
| 5137 | MIR96_5P | 386 | 0.33 | 0.8 | 0.94 | 1 |
| 5138 | MIR130A_5P | 277 | 0.33 | 0.8 | 0.924 | 1 |
| 5139 | REACTOME_FC_EPSILON_RECEPTOR_FCERI_SIG[illegible] | 116 | 0.35 | 0.8 | 0.868 | 1 |
| 5140 | REACTOME_HEDGEHOG_ON_STATE | 76 | 0.36 | 0.8 | 0.838 | 1 |
| 5141 | WP_SPLICING_FACTOR_NOVA_REGULATED_SYN[illegible] | 35 | 0.4 | 0.8 | 0.786 | 1 |
| 5142 | CCTGCTG_MIR214 | 224 | 0.33 | 0.8 | 0.905 | 1 |
| 5143 | MIR329_3P | 111 | 0.35 | 0.8 | 0.85 | 1 |
| 5144 | MIR499A_3P | 144 | 0.34 | 0.8 | 0.887 | 1 |
| 5145 | AGGTGCA_MIR500 | 82 | 0.36 | 0.8 | 0.841 | 1 |

| | | | | | | |
|---|---|---|---|---|---|---|
| 5146 | MIR3157_5P | 92 | 0.36 | 0.8 | 0.848 | 1 |
| 5147 | MIR6751_5P | 54 | 0.38 | 0.8 | 0.831 | 1 |
| 5148 | MIR499B_3P | 146 | 0.34 | 0.8 | 0.863 | 1 |
| 5149 | MIR4426 | 124 | 0.35 | 0.8 | 0.852 | 1 |
| 5150 | BHAT_ESR1_TARGETS_NOT_VIA_AKT1_UP | 179 | 0.34 | 0.8 | 0.89 | 1 |
| 5151 | MIR3649 | 17 | 0.45 | 0.8 | 0.753 | 1 |
| 5152 | AHR_Q5 | 207 | 0.34 | 0.8 | 0.894 | 1 |
| 5153 | CROONQUIST_NRAS_VS_STROMAL_STIMULATION | 30 | 0.41 | 0.8 | 0.774 | 1 |
| 5154 | DNMT3A_TARGET_GENES | 496 | 0.33 | 0.8 | 0.969 | 1 |
| 5155 | MIKKELSEN_MEF_ICP_WITH_H3K4ME3_AND_H3K2 | 27 | 0.41 | 0.8 | 0.774 | 1 |
| 5156 | MOOTHA_MITOCHONDRIA | 411 | 0.33 | 0.8 | 0.953 | 1 |
| 5157 | MIR1321 | 120 | 0.35 | 0.8 | 0.871 | 1 |
| 5158 | REACTOME_PLATELET_CALCIUM_HOMEOSTASIS | 23 | 0.42 | 0.8 | 0.775 | 1 |
| 5159 | LET_7B_5P | 355 | 0.33 | 0.8 | 0.944 | 1 |
| 5160 | ZNF140_TARGET_GENES | 462 | 0.33 | 0.8 | 0.969 | 1 |
| 5161 | FURUKAWA_DUSP6_TARGETS_PCI35_DN | 68 | 0.36 | 0.8 | 0.834 | 1 |
| 5162 | MIR4773 | 163 | 0.34 | 0.8 | 0.884 | 1 |
| 5163 | MIR4768_3P | 160 | 0.34 | 0.8 | 0.888 | 1 |
| 5164 | PACHER_TARGETS_OF_IGF1_AND_IGF2_UP | 31 | 0.4 | 0.8 | 0.759 | 1 |
| 5165 | MIR5787 | 212 | 0.33 | 0.8 | 0.909 | 1 |
| 5166 | SATOH_COLORECTAL_CANCER_MYC_DN | 57 | 0.37 | 0.8 | 0.817 | 1 |
| 5167 | TTAYRTAA_E4BP4_01 | 220 | 0.33 | 0.8 | 0.905 | 1 |
| 5168 | MIR548V | 113 | 0.35 | 0.8 | 0.873 | 1 |
| 5169 | CHIARADONNA_NEOPLASTIC_TRANSFORMATION_ | 131 | 0.34 | 0.8 | 0.886 | 1 |
| 5170 | REACTOME_REGULATION_OF_MECP2_EXPRESSI( | 30 | 0.41 | 0.8 | 0.775 | 1 |
| 5171 | MIR9900 | 39 | 0.39 | 0.8 | 0.793 | 1 |
| 5172 | COATES_MACROPHAGE_M1_VS_M2_DN | 58 | 0.37 | 0.8 | 0.848 | 1 |
| 5173 | MIR6862_5P | 40 | 0.39 | 0.8 | 0.791 | 1 |
| 5174 | MIR3914 | 49 | 0.37 | 0.8 | 0.819 | 1 |
| 5175 | MIR876_3P | 67 | 0.36 | 0.8 | 0.815 | 1 |
| 5176 | MIR513C_5P | 152 | 0.34 | 0.8 | 0.87 | 1 |
| 5177 | MIR4762_3P | 233 | 0.33 | 0.8 | 0.93 | 1 |
| 5178 | MIR6760_3P | 84 | 0.36 | 0.8 | 0.846 | 1 |

| | | | | | | |
|---|---|---|---|---|---|---|
| 5179 | MIR4477B | 222 | 0.33 | 0.8 | 0.911 | 1 |
| 5180 | AGGGCCA_MIR328 | 71 | 0.36 | 0.8 | 0.82 | 1 |
| 5181 | CHANDRAN_METASTASIS_TOP50_DN | 44 | 0.39 | 0.8 | 0.806 | 1 |
| 5182 | WP_NOCGMPPKG_MEDIATED_NEUROPROTECTIOI | 40 | 0.39 | 0.8 | 0.816 | 1 |
| 5183 | MIR449C_5P | 114 | 0.35 | 0.8 | 0.872 | 1 |
| 5184 | PID_BMP_PATHWAY | 32 | 0.41 | 0.8 | 0.796 | 1 |
| 5185 | MIR1207_5P | 285 | 0.33 | 0.8 | 0.927 | 1 |
| 5186 | MIR6874_5P | 30 | 0.4 | 0.8 | 0.788 | 1 |
| 5187 | MIR324_5P | 40 | 0.39 | 0.8 | 0.795 | 1 |
| 5188 | CEBPGAMMA_Q6 | 196 | 0.33 | 0.8 | 0.903 | 1 |
| 5189 | GENTILE_UV_RESPONSE_CLUSTER_D8 | 34 | 0.4 | 0.8 | 0.787 | 1 |
| 5190 | FOXO3_01 | 196 | 0.33 | 0.8 | 0.903 | 1 |
| 5191 | CASP8AP2_TARGET_GENES | 290 | 0.33 | 0.8 | 0.934 | 1 |
| 5192 | MIR196A_3P | 109 | 0.35 | 0.79 | 0.866 | 1 |
| 5193 | MIR485_5P | 120 | 0.34 | 0.79 | 0.867 | 1 |
| 5194 | MIR4719 | 444 | 0.33 | 0.79 | 0.969 | 1 |
| 5195 | WP_ATM_SIGNALING_IN_DEVELOPMENT_AND_DIS | 45 | 0.38 | 0.79 | 0.82 | 1 |
| 5196 | YRTCANNRCGC_UNKNOWN | 57 | 0.37 | 0.79 | 0.828 | 1 |
| 5197 | MIR328_3P | 33 | 0.4 | 0.79 | 0.805 | 1 |
| 5198 | WELCSH_BRCA1_TARGETS_DN | 137 | 0.34 | 0.79 | 0.878 | 1 |
| 5199 | LEE_METASTASIS_AND_ALTERNATIVE_SPLICING_ | 39 | 0.39 | 0.79 | 0.82 | 1 |
| 5200 | FOSTER_KDM1A_TARGETS_DN | 190 | 0.34 | 0.79 | 0.906 | 1 |
| 5201 | JNK_DN.V1_DN | 123 | 0.35 | 0.79 | 0.87 | 1 |
| 5202 | VANDESLUIS_COMMD1_TARGETS_GROUP_3_UP | 59 | 0.37 | 0.79 | 0.842 | 1 |
| 5203 | LET_7A_5P_LET_7C_5P_LET_7E_5P | 353 | 0.33 | 0.79 | 0.958 | 1 |
| 5204 | PID_EPHA2_FWD_PATHWAY | 19 | 0.43 | 0.79 | 0.776 | 1 |
| 5205 | MIR4709_3P | 127 | 0.34 | 0.79 | 0.867 | 1 |
| 5206 | MIR4756_5P | 119 | 0.35 | 0.79 | 0.86 | 1 |
| 5207 | ZHOU_CELL_CYCLE_GENES_IN_IR_RESPONSE_2 | 119 | 0.35 | 0.79 | 0.88 | 1 |
| 5208 | MEINHOLD_OVARIAN_CANCER_LOW_GRADE_DN | 19 | 0.43 | 0.79 | 0.771 | 1 |
| 5209 | LIAO_METASTASIS | 495 | 0.33 | 0.79 | 0.973 | 1 |
| 5210 | TTCNRGNNNNTTC_HSF_Q6 | 123 | 0.34 | 0.79 | 0.897 | 1 |
| 5211 | MIR34B_5P | 113 | 0.35 | 0.79 | 0.871 | 1 |

| | | | | | | |
|---|---|---|---|---|---|---|
| 5212 | AP4_01 | 219 | 0.33 | 0.79 | 0.913 | 1 |
| 5213 | GARGALOVIC_RESPONSE_TO_OXIDIZED_PHOSPH | 22 | 0.42 | 0.79 | 0.773 | 1 |
| 5214 | VALK_AML_CLUSTER_12 | 24 | 0.41 | 0.79 | 0.774 | 1 |
| 5215 | REACTOME_REGULATION_OF_LIPID_METABOLISM | 103 | 0.35 | 0.79 | 0.86 | 1 |
| 5216 | EVI1_04 | 179 | 0.34 | 0.79 | 0.898 | 1 |
| 5217 | PHF21A_TARGET_GENES | 241 | 0.33 | 0.79 | 0.91 | 1 |
| 5218 | MIR6131 | 88 | 0.36 | 0.79 | 0.874 | 1 |
| 5219 | LET_7F_5P | 354 | 0.33 | 0.79 | 0.947 | 1 |
| 5220 | MIR4517 | 167 | 0.34 | 0.79 | 0.902 | 1 |
| 5221 | REACTOME_SIGNALING_BY_SCF_KIT | 37 | 0.39 | 0.79 | 0.812 | 1 |
| 5222 | MIR4739 | 120 | 0.34 | 0.79 | 0.871 | 1 |
| 5223 | REACTOME_G_ALPHA_Z_SIGNALLING_EVENTS | 33 | 0.4 | 0.79 | 0.787 | 1 |
| 5224 | MIR7161_5P | 175 | 0.34 | 0.79 | 0.92 | 1 |
| 5225 | NRSF_01 | 74 | 0.36 | 0.79 | 0.846 | 1 |
| 5226 | MIR98_5P | 354 | 0.33 | 0.79 | 0.949 | 1 |
| 5227 | MIR10A_5P_MIR10B_5P | 89 | 0.35 | 0.79 | 0.845 | 1 |
| 5228 | MIR4460 | 75 | 0.36 | 0.79 | 0.836 | 1 |
| 5229 | CCCAGAG_MIR326 | 136 | 0.34 | 0.79 | 0.898 | 1 |
| 5230 | MIR5699_5P | 96 | 0.35 | 0.79 | 0.873 | 1 |
| 5231 | KUNINGER_IGF1_VS_PDGFB_TARGETS_UP | 52 | 0.37 | 0.79 | 0.818 | 1 |
| 5232 | KEGG_PHOSPHATIDYLINOSITOL_SIGNALING_SYS | 67 | 0.36 | 0.79 | 0.842 | 1 |
| 5233 | MIR6800_5P | 37 | 0.39 | 0.79 | 0.8 | 1 |
| 5234 | AAAGACA_MIR511 | 176 | 0.33 | 0.79 | 0.907 | 1 |
| 5235 | REACTOME_SYNTHESIS_OF_SUBSTRATES_IN_N_ | 56 | 0.37 | 0.79 | 0.816 | 1 |
| 5236 | MIR3158_3P | 37 | 0.39 | 0.79 | 0.807 | 1 |
| 5237 | HSF2_01 | 203 | 0.33 | 0.79 | 0.92 | 1 |
| 5238 | CACBINDINGPROTEIN_Q6 | 207 | 0.33 | 0.79 | 0.932 | 1 |
| 5239 | VDR_Q3 | 197 | 0.33 | 0.79 | 0.924 | 1 |
| 5240 | MIR4527 | 62 | 0.37 | 0.79 | 0.828 | 1 |
| 5241 | USF_Q6_01 | 219 | 0.33 | 0.79 | 0.922 | 1 |
| 5242 | BIOCARTA_NFAT_PATHWAY | 40 | 0.38 | 0.79 | 0.802 | 1 |
| 5243 | REACTOME_REGULATION_OF_TP53_ACTIVITY_TH | 89 | 0.35 | 0.79 | 0.857 | 1 |
| 5244 | SCHAEFFER_SOX9_TARGETS_IN_PROSTATE_DEV | 40 | 0.39 | 0.79 | 0.81 | 1 |

| | | | | | | |
|---|---|---|---|---|---|---|
| 5245 | WP_FOLLICLE_STIMULATING_HORMONE_FSH_SIG | 21 | 0.43 | 0.79 | 0.791 | 1 |
| 5246 | TSENG_ADIPOGENIC_POTENTIAL_DN | 31 | 0.4 | 0.79 | 0.793 | 1 |
| 5247 | WATANABE_COLON_CANCER_MSI_VS_MSS_UP | 24 | 0.42 | 0.79 | 0.796 | 1 |
| 5248 | MIR4716_5P | 104 | 0.35 | 0.79 | 0.878 | 1 |
| 5249 | DACOSTA_ERCC3_ALLELE_XPCS_VS_TTD_DN | 23 | 0.42 | 0.79 | 0.791 | 1 |
| 5250 | MIR4453 | 23 | 0.42 | 0.79 | 0.796 | 1 |
| 5251 | WP_PARKINUBIQUITIN_PROTEASOMAL_SYSTEM_I | 62 | 0.36 | 0.79 | 0.846 | 1 |
| 5252 | MIR4465 | 383 | 0.32 | 0.79 | 0.954 | 1 |
| 5253 | SARRIO_EPITHELIAL_MESENCHYMAL_TRANSITIOI | 153 | 0.33 | 0.79 | 0.896 | 1 |
| 5254 | MIR1252_3P | 250 | 0.33 | 0.79 | 0.934 | 1 |
| 5255 | MIR218_5P | 368 | 0.32 | 0.79 | 0.955 | 1 |
| 5256 | CHEMNITZ_RESPONSE_TO_PROSTAGLANDIN_E2_ | 321 | 0.32 | 0.79 | 0.941 | 1 |
| 5257 | MIR6818_3P | 96 | 0.35 | 0.79 | 0.857 | 1 |
| 5258 | REACTOME_M_PHASE | 375 | 0.32 | 0.79 | 0.959 | 1 |
| 5259 | BERTUCCI_INVASIVE_CARCINOMA_DUCTAL_VS_L | 24 | 0.41 | 0.79 | 0.786 | 1 |
| 5260 | MIR6730_5P | 108 | 0.34 | 0.79 | 0.893 | 1 |
| 5261 | MIR493_3P | 56 | 0.36 | 0.79 | 0.823 | 1 |
| 5262 | MIR937_5P | 47 | 0.37 | 0.78 | 0.824 | 1 |
| 5263 | ACCAAAG_MIR9 | 452 | 0.32 | 0.78 | 0.968 | 1 |
| 5264 | NKX61_01 | 179 | 0.33 | 0.78 | 0.924 | 1 |
| 5265 | MIR223_3P | 112 | 0.34 | 0.78 | 0.873 | 1 |
| 5266 | REACTOME_METALLOPROTEASE_DUBS | 27 | 0.41 | 0.78 | 0.8 | 1 |
| 5267 | MIR642A_3P_MIR642B_3P | 114 | 0.34 | 0.78 | 0.897 | 1 |
| 5268 | TAXCREB_01 | 117 | 0.34 | 0.78 | 0.876 | 1 |
| 5269 | MIR4495 | 410 | 0.32 | 0.78 | 0.961 | 1 |
| 5270 | KEGG_SPHINGOLIPID_METABOLISM | 33 | 0.39 | 0.78 | 0.803 | 1 |
| 5271 | CAFFAREL_RESPONSE_TO_THC_8HR_5_UP | 15 | 0.43 | 0.78 | 0.763 | 1 |
| 5272 | PID_AJDISS_2PATHWAY | 44 | 0.38 | 0.78 | 0.831 | 1 |
| 5273 | MIR10226 | 65 | 0.36 | 0.78 | 0.848 | 1 |
| 5274 | MIR875_3P | 256 | 0.32 | 0.78 | 0.925 | 1 |
| 5275 | KOHOUTEK_CCNT2_TARGETS | 39 | 0.39 | 0.78 | 0.819 | 1 |
| 5276 | MIR7151_3P | 233 | 0.33 | 0.78 | 0.929 | 1 |
| 5277 | ZNF433_TARGET_GENES | 45 | 0.38 | 0.78 | 0.822 | 1 |

| | | | | | | |
|---|---|---|---|---|---|---|
| 5278 | SHARMA_PILOCYTIC_ASTROCYTOMA_LOCATION_ | 20 | 0.42 | 0.78 | 0.767 | 1 |
| 5279 | ASGHARZADEH_NEUROBLASTOMA_POOR_SURVI\ | 40 | 0.39 | 0.78 | 0.817 | 1 |
| 5280 | HOXA9_DN.V1_DN | 163 | 0.33 | 0.78 | 0.905 | 1 |
| 5281 | CATRRAGC_UNKNOWN | 115 | 0.34 | 0.78 | 0.889 | 1 |
| 5282 | MIR136_5P | 90 | 0.35 | 0.78 | 0.87 | 1 |
| 5283 | PAX3_B | 81 | 0.35 | 0.78 | 0.861 | 1 |
| 5284 | MIR4484 | 32 | 0.4 | 0.78 | 0.801 | 1 |
| 5285 | MIR5703 | 140 | 0.34 | 0.78 | 0.9 | 1 |
| 5286 | MIR514B_5P | 153 | 0.34 | 0.78 | 0.902 | 1 |
| 5287 | TCATCTC_MIR143 | 135 | 0.33 | 0.78 | 0.906 | 1 |
| 5288 | MIR4269 | 186 | 0.33 | 0.78 | 0.917 | 1 |
| 5289 | MIR6780B_5P | 233 | 0.33 | 0.78 | 0.945 | 1 |
| 5290 | WP_REGULATION_OF_APOPTOSIS_BY_PARATHYI | 19 | 0.43 | 0.78 | 0.782 | 1 |
| 5291 | OCT1_03 | 184 | 0.33 | 0.78 | 0.916 | 1 |
| 5292 | MIR5197_5P | 167 | 0.33 | 0.78 | 0.91 | 1 |
| 5293 | WP_H19_ACTION_RBE2F1_SIGNALING_AND_CDKE | 15 | 0.45 | 0.78 | 0.795 | 1 |
| 5294 | ABE_INNER_EAR | 32 | 0.39 | 0.78 | 0.797 | 1 |
| 5295 | REACTOME_INTEGRIN_SIGNALING | 21 | 0.41 | 0.78 | 0.785 | 1 |
| 5296 | KAAB_HEART_ATRIUM_VS_VENTRICLE_DN | 226 | 0.32 | 0.78 | 0.914 | 1 |
| 5297 | MIR5006_5P | 103 | 0.34 | 0.78 | 0.881 | 1 |
| 5298 | HDAC8_TARGET_GENES | 109 | 0.34 | 0.78 | 0.883 | 1 |
| 5299 | SRY_02 | 207 | 0.32 | 0.78 | 0.922 | 1 |
| 5300 | BRACHAT_RESPONSE_TO_CISPLATIN | 20 | 0.42 | 0.78 | 0.814 | 1 |
| 5301 | MIR3714 | 390 | 0.32 | 0.78 | 0.972 | 1 |
| 5302 | KEGG_INSULIN_SIGNALING_PATHWAY | 113 | 0.34 | 0.78 | 0.888 | 1 |
| 5303 | KEGG_LYSOSOME | 111 | 0.34 | 0.78 | 0.897 | 1 |
| 5304 | MIR5579_3P | 134 | 0.33 | 0.78 | 0.898 | 1 |
| 5305 | MIR200C_5P | 47 | 0.37 | 0.78 | 0.826 | 1 |
| 5306 | BURTON_ADIPOGENESIS_PEAK_AT_24HR | 39 | 0.38 | 0.78 | 0.83 | 1 |
| 5307 | GTATTAT_MIR3693P | 191 | 0.33 | 0.78 | 0.933 | 1 |
| 5308 | MIR12129 | 123 | 0.34 | 0.78 | 0.895 | 1 |
| 5309 | XU_GH1_AUTOCRINE_TARGETS_DN | 112 | 0.34 | 0.78 | 0.883 | 1 |
| 5310 | GAANYNYGACNY_UNKNOWN | 61 | 0.36 | 0.78 | 0.844 | 1 |

| | | | | | | |
|---|---|---|---|---|---|---|
| 5311 | REACTOME_THROMBIN_SIGNALLING_THROUGH_F | 21 | 0.43 | 0.78 | 0.804 | 1 |
| 5312 | MIR6875_3P | 328 | 0.32 | 0.78 | 0.963 | 1 |
| 5313 | MIR4769_3P | 98 | 0.34 | 0.78 | 0.888 | 1 |
| 5314 | LEE_LIVER_CANCER_MYC_UP | 43 | 0.37 | 0.78 | 0.818 | 1 |
| 5315 | ACCATTT_MIR522 | 148 | 0.33 | 0.78 | 0.914 | 1 |
| 5316 | MIR4637 | 52 | 0.36 | 0.78 | 0.806 | 1 |
| 5317 | ZNF532_TARGET_GENES | 60 | 0.36 | 0.78 | 0.84 | 1 |
| 5318 | MIR4268 | 52 | 0.37 | 0.78 | 0.849 | 1 |
| 5319 | KMCATNNWGGA_UNKNOWN | 92 | 0.34 | 0.78 | 0.874 | 1 |
| 5320 | MIR3665 | 68 | 0.36 | 0.78 | 0.863 | 1 |
| 5321 | GGCNKCCATNK_UNKNOWN | 116 | 0.34 | 0.78 | 0.889 | 1 |
| 5322 | TITF1_Q3 | 195 | 0.32 | 0.78 | 0.927 | 1 |
| 5323 | WINTER_HYPOXIA_UP | 88 | 0.35 | 0.77 | 0.883 | 1 |
| 5324 | PID_AR_PATHWAY | 56 | 0.36 | 0.77 | 0.853 | 1 |
| 5325 | MIR122_5P | 80 | 0.35 | 0.77 | 0.873 | 1 |
| 5326 | MIR6864_3P | 36 | 0.39 | 0.77 | 0.819 | 1 |
| 5327 | MARCINIAK_ER_STRESS_RESPONSE_VIA_CHOP | 19 | 0.43 | 0.77 | 0.789 | 1 |
| 5328 | MIR3177_5P | 130 | 0.34 | 0.77 | 0.911 | 1 |
| 5329 | MIR6862_3P | 26 | 0.4 | 0.77 | 0.807 | 1 |
| 5330 | FAC1_01 | 178 | 0.33 | 0.77 | 0.922 | 1 |
| 5331 | XBP1_01 | 114 | 0.34 | 0.77 | 0.899 | 1 |
| 5332 | MIR483_3P | 48 | 0.37 | 0.77 | 0.859 | 1 |
| 5333 | GGCAGCT_MIR22 | 205 | 0.33 | 0.77 | 0.94 | 1 |
| 5334 | AACTGAC_MIR223 | 91 | 0.34 | 0.77 | 0.888 | 1 |
| 5335 | MIR3120_3P | 307 | 0.32 | 0.77 | 0.953 | 1 |
| 5336 | MIR6516_5P | 160 | 0.33 | 0.77 | 0.925 | 1 |
| 5337 | MIR599 | 69 | 0.35 | 0.77 | 0.855 | 1 |
| 5338 | MIR3664_3P | 129 | 0.33 | 0.77 | 0.905 | 1 |
| 5339 | MIR6782_3P | 77 | 0.35 | 0.77 | 0.869 | 1 |
| 5340 | MIR3162_3P | 16 | 0.44 | 0.77 | 0.801 | 1 |
| 5341 | MIR3136_3P | 55 | 0.36 | 0.77 | 0.842 | 1 |
| 5342 | KIM_ALL_DISORDERS_DURATION_CORR_DN | 133 | 0.33 | 0.77 | 0.916 | 1 |
| 5343 | TAXCREB_02 | 22 | 0.41 | 0.77 | 0.808 | 1 |

| | | | | | | |
|---|---|---|---|---|---|---|
| 5344 | LI_WILMS_TUMOR_VS_FETAL_KIDNEY_1_DN | 156 | 0.33 | 0.77 | 0.913 | 1 |
| 5345 | IIZUKA_LIVER_CANCER_PROGRESSION_L1_G1_U | 17 | 0.43 | 0.77 | 0.816 | 1 |
| 5346 | ELVIDGE_HYPOXIA_DN | 140 | 0.33 | 0.77 | 0.914 | 1 |
| 5347 | MIR525_5P | 140 | 0.33 | 0.77 | 0.899 | 1 |
| 5348 | WANG_RESPONSE_TO_BEXAROTENE_DN | 22 | 0.41 | 0.77 | 0.827 | 1 |
| 5349 | MIR6855_5P | 60 | 0.36 | 0.77 | 0.867 | 1 |
| 5350 | OUYANG_PROSTATE_CANCER_PROGRESSION_DI | 20 | 0.41 | 0.77 | 0.799 | 1 |
| 5351 | REACTOME_DISASSEMBLY_OF_THE_DESTRUCTIC | 27 | 0.4 | 0.77 | 0.819 | 1 |
| 5352 | BCAT.100_UP.V1_DN | 24 | 0.41 | 0.77 | 0.822 | 1 |
| 5353 | KEGG_FC_EPSILON_RI_SIGNALING_PATHWAY | 56 | 0.36 | 0.77 | 0.848 | 1 |
| 5354 | WP_TARGET_OF_RAPAMYCIN_TOR_SIGNALING | 35 | 0.38 | 0.77 | 0.84 | 1 |
| 5355 | TAATAAT_MIR126 | 190 | 0.33 | 0.77 | 0.932 | 1 |
| 5356 | MIR126_5P | 350 | 0.32 | 0.77 | 0.957 | 1 |
| 5357 | MIR297 | 116 | 0.33 | 0.77 | 0.893 | 1 |
| 5358 | ZNF331_TARGET_GENES | 288 | 0.32 | 0.77 | 0.952 | 1 |
| 5359 | MIR600 | 118 | 0.33 | 0.77 | 0.903 | 1 |
| 5360 | MIR4273 | 85 | 0.35 | 0.77 | 0.877 | 1 |
| 5361 | MIR519A_3P_MIR519B_3P | 358 | 0.32 | 0.77 | 0.959 | 1 |
| 5362 | MIR6887_3P | 177 | 0.32 | 0.77 | 0.921 | 1 |
| 5363 | MIR6089 | 16 | 0.43 | 0.77 | 0.804 | 1 |
| 5364 | MIR3908 | 100 | 0.34 | 0.77 | 0.895 | 1 |
| 5365 | REACTOME_FLT3_SIGNALING | 34 | 0.38 | 0.77 | 0.838 | 1 |
| 5366 | DAZARD_RESPONSE_TO_UV_SCC_DN | 109 | 0.34 | 0.77 | 0.902 | 1 |
| 5367 | REACTOME_GLYCOSPHINGOLIPID_METABOLISM | 39 | 0.37 | 0.77 | 0.843 | 1 |
| 5368 | MIR374A_3P | 115 | 0.33 | 0.77 | 0.907 | 1 |
| 5369 | MIR1250_3P | 416 | 0.32 | 0.77 | 0.977 | 1 |
| 5370 | MIR1252_5P | 265 | 0.32 | 0.77 | 0.954 | 1 |
| 5371 | MIR4651 | 163 | 0.33 | 0.77 | 0.917 | 1 |
| 5372 | MIR520A_5P | 140 | 0.33 | 0.77 | 0.908 | 1 |
| 5373 | AGTCTAG_MIR151 | 22 | 0.41 | 0.77 | 0.803 | 1 |
| 5374 | MIR4796_3P | 219 | 0.32 | 0.77 | 0.946 | 1 |
| 5375 | MIR200B_5P | 94 | 0.34 | 0.77 | 0.875 | 1 |
| 5376 | MIR519C_3P | 359 | 0.32 | 0.77 | 0.974 | 1 |

| | | | | | | |
|---|---|---|---|---|---|---|
| 5377 | STONER_ESOPHAGEAL_CARCINOGENESIS_UP | 36 | 0.38 | 0.77 | 0.834 | 1 |
| 5378 | BIOCARTA_STRESS_PATHWAY | 24 | 0.4 | 0.77 | 0.836 | 1 |
| 5379 | REACTOME_ADHERENS_JUNCTIONS_INTERACTIOI | 25 | 0.4 | 0.77 | 0.82 | 1 |
| 5380 | MIR216B_3P | 111 | 0.34 | 0.77 | 0.889 | 1 |
| 5381 | REACTOME_HOST_INTERACTIONS_OF_HIV_FACTC | 126 | 0.33 | 0.77 | 0.919 | 1 |
| 5382 | MIR4534 | 122 | 0.33 | 0.77 | 0.902 | 1 |
| 5383 | LANDIS_ERBB2_BREAST_PRENEOPLASTIC_UP | 17 | 0.42 | 0.76 | 0.809 | 1 |
| 5384 | MIR4635 | 108 | 0.34 | 0.76 | 0.893 | 1 |
| 5385 | MIR493_5P | 399 | 0.31 | 0.76 | 0.974 | 1 |
| 5386 | MIR361_3P | 90 | 0.34 | 0.76 | 0.894 | 1 |
| 5387 | HAND1E47_01 | 203 | 0.32 | 0.76 | 0.941 | 1 |
| 5388 | LANDIS_ERBB2_BREAST_PRENEOPLASTIC_DN | 45 | 0.37 | 0.76 | 0.856 | 1 |
| 5389 | DAIRKEE_CANCER_PRONE_RESPONSE_E2 | 22 | 0.41 | 0.76 | 0.809 | 1 |
| 5390 | MIR6513_3P | 124 | 0.33 | 0.76 | 0.919 | 1 |
| 5391 | REACTOME_REGULATION_OF_EXPRESSION_OF_S | 163 | 0.33 | 0.76 | 0.93 | 1 |
| 5392 | WHITFIELD_CELL_CYCLE_M_G1 | 124 | 0.33 | 0.76 | 0.908 | 1 |
| 5393 | MIR7849_3P | 103 | 0.34 | 0.76 | 0.903 | 1 |
| 5394 | ZFP3_TARGET_GENES | 291 | 0.32 | 0.76 | 0.969 | 1 |
| 5395 | REACTOME_ROLE_OF_PHOSPHOLIPIDS_IN_PHAG | 19 | 0.42 | 0.76 | 0.798 | 1 |
| 5396 | CHEOK_RESPONSE_TO_HD_MTX_DN | 23 | 0.4 | 0.76 | 0.819 | 1 |
| 5397 | MIR6857_3P | 58 | 0.35 | 0.76 | 0.872 | 1 |
| 5398 | GGCACAT_MIR455 | 53 | 0.36 | 0.76 | 0.869 | 1 |
| 5399 | ERR1_Q2 | 238 | 0.32 | 0.76 | 0.952 | 1 |
| 5400 | JOHNSTONE_PARVB_TARGETS_2_DN | 321 | 0.31 | 0.76 | 0.969 | 1 |
| 5401 | MIR3928_3P | 80 | 0.34 | 0.76 | 0.895 | 1 |
| 5402 | HASLINGER_B_CLL_WITH_13Q14_DELETION | 19 | 0.41 | 0.76 | 0.814 | 1 |
| 5403 | ZAMORA_NOS2_TARGETS_UP | 52 | 0.36 | 0.76 | 0.853 | 1 |
| 5404 | MIR4261 | 375 | 0.31 | 0.76 | 0.979 | 1 |
| 5405 | AREB6_03 | 227 | 0.32 | 0.76 | 0.954 | 1 |
| 5406 | HOLLEMAN_PREDNISOLONE_RESISTANCE_ALL_U | 20 | 0.41 | 0.76 | 0.815 | 1 |
| 5407 | PRAMOONJAGO_SOX4_TARGETS_DN | 47 | 0.36 | 0.76 | 0.87 | 1 |
| 5408 | MIR3059_5P | 250 | 0.31 | 0.76 | 0.949 | 1 |
| 5409 | MIR4676_5P | 48 | 0.36 | 0.76 | 0.836 | 1 |

| | | | | | | |
|---|---|---|---|---|---|---|
| 5410 | MIR6817_5P | 98 | 0.34 | 0.76 | 0.915 | 1 |
| 5411 | MIR1909_3P | 123 | 0.33 | 0.76 | 0.911 | 1 |
| 5412 | MIR10394_5P | 155 | 0.32 | 0.76 | 0.929 | 1 |
| 5413 | MIR4323 | 37 | 0.37 | 0.76 | 0.842 | 1 |
| 5414 | PID_BETA_CATENIN_DEG_PATHWAY | 17 | 0.42 | 0.76 | 0.795 | 1 |
| 5415 | WP_METHIONINE_DE_NOVO_AND_SALVAGE_PATI | 18 | 0.43 | 0.76 | 0.832 | 1 |
| 5416 | MIR4789_5P | 336 | 0.31 | 0.76 | 0.975 | 1 |
| 5417 | KAYO_AGING_MUSCLE_UP | 162 | 0.32 | 0.76 | 0.93 | 1 |
| 5418 | MIR4774_3P | 64 | 0.35 | 0.76 | 0.884 | 1 |
| 5419 | BILANGES_SERUM_SENSITIVE_GENES | 61 | 0.35 | 0.76 | 0.867 | 1 |
| 5420 | USF_Q6 | 235 | 0.32 | 0.76 | 0.954 | 1 |
| 5421 | HU_GENOTOXIN_ACTION_DIRECT_VS_INDIRECT_2 | 46 | 0.36 | 0.76 | 0.863 | 1 |
| 5422 | MIR7152_5P | 212 | 0.32 | 0.76 | 0.953 | 1 |
| 5423 | CCCNNGGGAR_OLF1_01 | 272 | 0.32 | 0.76 | 0.954 | 1 |
| 5424 | GAGCCAG_MIR149 | 135 | 0.33 | 0.76 | 0.927 | 1 |
| 5425 | KIM_GASTRIC_CANCER_CHEMOSENSITIVITY | 79 | 0.34 | 0.76 | 0.896 | 1 |
| 5426 | SNAI1_TARGET_GENES | 430 | 0.31 | 0.76 | 0.981 | 1 |
| 5427 | ZNF512B_TARGET_GENES | 174 | 0.32 | 0.76 | 0.946 | 1 |
| 5428 | CCAGGTT_MIR490 | 61 | 0.35 | 0.76 | 0.894 | 1 |
| 5429 | MIR7155_3P | 56 | 0.35 | 0.76 | 0.87 | 1 |
| 5430 | OCT1_Q5_01 | 189 | 0.32 | 0.76 | 0.963 | 1 |
| 5431 | USF_02 | 233 | 0.31 | 0.76 | 0.964 | 1 |
| 5432 | REACTOME_TRNA_MODIFICATION_IN_THE_NUCLE | 42 | 0.37 | 0.76 | 0.868 | 1 |
| 5433 | TING_SILENCED_BY_DICER | 19 | 0.42 | 0.76 | 0.82 | 1 |
| 5434 | BIOCARTA_CERAMIDE_PATHWAY | 21 | 0.41 | 0.76 | 0.83 | 1 |
| 5435 | MIR938 | 67 | 0.35 | 0.76 | 0.88 | 1 |
| 5436 | MIR4421 | 43 | 0.37 | 0.76 | 0.849 | 1 |
| 5437 | MIR4297 | 143 | 0.32 | 0.76 | 0.939 | 1 |
| 5438 | TCCAGAG_MIR518C | 139 | 0.32 | 0.76 | 0.935 | 1 |
| 5439 | MIR494_3P | 319 | 0.31 | 0.76 | 0.961 | 1 |
| 5440 | PID_REELIN_PATHWAY | 27 | 0.39 | 0.76 | 0.839 | 1 |
| 5441 | RAO_BOUND_BY_SALL4_ISOFORM_A | 124 | 0.33 | 0.75 | 0.921 | 1 |
| 5442 | MIR5000_5P | 92 | 0.33 | 0.75 | 0.915 | 1 |

| | | | | | | |
|---|---|---|---|---|---|---|
| 5443 | MIR6715B_5P | 199 | 0.32 | 0.75 | 0.946 | 1 |
| 5444 | ZNF157_TARGET_GENES | 42 | 0.37 | 0.75 | 0.851 | 1 |
| 5445 | TTTGCAC_MIR19A_MIR19B | 478 | 0.31 | 0.75 | 0.987 | 1 |
| 5446 | MIR4690_3P | 62 | 0.35 | 0.75 | 0.873 | 1 |
| 5447 | FOXO4_TARGET_GENES | 268 | 0.31 | 0.75 | 0.973 | 1 |
| 5448 | MOOTHA_GLYCOLYSIS | 15 | 0.43 | 0.75 | 0.821 | 1 |
| 5449 | MIR3910 | 359 | 0.31 | 0.75 | 0.98 | 1 |
| 5450 | MIR129_5P | 329 | 0.31 | 0.75 | 0.978 | 1 |
| 5451 | PID_VEGFR1_2_PATHWAY | 62 | 0.34 | 0.75 | 0.853 | 1 |
| 5452 | EVI1_05 | 119 | 0.33 | 0.75 | 0.939 | 1 |
| 5453 | TTTGCAG_MIR518A2 | 190 | 0.32 | 0.75 | 0.94 | 1 |
| 5454 | MIR6793_5P | 49 | 0.36 | 0.75 | 0.866 | 1 |
| 5455 | WP_ID_SIGNALING_PATHWAY | 15 | 0.43 | 0.75 | 0.81 | 1 |
| 5456 | WEIGEL_OXIDATIVE_STRESS_BY_HNE_AND_TBH | 54 | 0.35 | 0.75 | 0.882 | 1 |
| 5457 | ZHOU_INFLAMMATORY_RESPONSE_FIMA_DN | 217 | 0.32 | 0.75 | 0.96 | 1 |
| 5458 | MIR4275 | 133 | 0.32 | 0.75 | 0.934 | 1 |
| 5459 | WP_STEROL_REGULATORY_ELEMENTBINDING_PI | 65 | 0.34 | 0.75 | 0.895 | 1 |
| 5460 | MIR8074 | 17 | 0.42 | 0.75 | 0.82 | 1 |
| 5461 | URS_ADIPOCYTE_DIFFERENTIATION_UP | 49 | 0.36 | 0.75 | 0.874 | 1 |
| 5462 | MIR138_1_3P | 72 | 0.34 | 0.75 | 0.874 | 1 |
| 5463 | RBM15_TARGET_GENES | 114 | 0.33 | 0.75 | 0.918 | 1 |
| 5464 | ZNF708_TARGET_GENES | 69 | 0.34 | 0.75 | 0.87 | 1 |
| 5465 | MIR4422 | 397 | 0.31 | 0.75 | 0.979 | 1 |
| 5466 | XIE_LT_HSC_S1PR3_OE_UP | 17 | 0.42 | 0.75 | 0.832 | 1 |
| 5467 | MEISSNER_ES_ICP_WITH_H3K4ME3 | 27 | 0.39 | 0.75 | 0.839 | 1 |
| 5468 | MIR6879_5P | 95 | 0.33 | 0.75 | 0.908 | 1 |
| 5469 | MIR6503_3P | 25 | 0.39 | 0.75 | 0.844 | 1 |
| 5470 | YIH_RESPONSE_TO_ARSENITE_C3 | 30 | 0.38 | 0.75 | 0.831 | 1 |
| 5471 | RODRIGUES_NTN1_TARGETS_UP | 16 | 0.42 | 0.75 | 0.822 | 1 |
| 5472 | MIR6721_5P | 300 | 0.31 | 0.75 | 0.975 | 1 |
| 5473 | MIR3672 | 38 | 0.37 | 0.75 | 0.858 | 1 |
| 5474 | MIR135A_5P | 267 | 0.31 | 0.75 | 0.966 | 1 |
| 5475 | MIR1227_5P | 88 | 0.34 | 0.75 | 0.903 | 1 |

| | | | | | | |
|---|---|---|---|---|---|---|
| 5476 | MIR545_5P | 266 | 0.31 | 0.75 | 0.971 | 1 |
| 5477 | SMAD4_Q6 | 205 | 0.31 | 0.75 | 0.951 | 1 |
| 5478 | MIR200A_5P | 94 | 0.33 | 0.75 | 0.915 | 1 |
| 5479 | MIR7150 | 113 | 0.33 | 0.75 | 0.919 | 1 |
| 5480 | MIR3691_3P | 63 | 0.34 | 0.75 | 0.895 | 1 |
| 5481 | MIR8075 | 72 | 0.34 | 0.75 | 0.893 | 1 |
| 5482 | WP_GLYCOLYSIS_AND_GLUCONEOGENESIS | 34 | 0.38 | 0.75 | 0.879 | 1 |
| 5483 | SHEPARD_CRASH_AND_BURN_MUTANT_DN | 160 | 0.32 | 0.75 | 0.961 | 1 |
| 5484 | CATGTAA_MIR496 | 158 | 0.32 | 0.75 | 0.948 | 1 |
| 5485 | MIR3913_3P | 60 | 0.34 | 0.75 | 0.88 | 1 |
| 5486 | FERREIRA_EWINGS_SARCOMA_UNSTABLE_VS_ST | 156 | 0.32 | 0.75 | 0.954 | 1 |
| 5487 | ER_Q6_02 | 211 | 0.31 | 0.75 | 0.957 | 1 |
| 5488 | MULLIGHAN_NPM1_MUTATED_SIGNATURE_2_UP | 110 | 0.33 | 0.75 | 0.941 | 1 |
| 5489 | CHARAFE_BREAST_CANCER_BASAL_VS_MESENC | 44 | 0.36 | 0.75 | 0.865 | 1 |
| 5490 | BRN2_01 | 178 | 0.32 | 0.75 | 0.961 | 1 |
| 5491 | MIR1273H_3P | 78 | 0.33 | 0.75 | 0.921 | 1 |
| 5492 | MIR3620_5P | 45 | 0.36 | 0.75 | 0.876 | 1 |
| 5493 | MIR4722_3P | 76 | 0.34 | 0.75 | 0.902 | 1 |
| 5494 | LEE_LIVER_CANCER_SURVIVAL_DN | 166 | 0.32 | 0.75 | 0.947 | 1 |
| 5495 | STEIN_ESRRA_TARGETS_UP | 369 | 0.31 | 0.75 | 0.987 | 1 |
| 5496 | YGACNNYACAR_UNKNOWN | 70 | 0.34 | 0.74 | 0.906 | 1 |
| 5497 | REACTOME_REGULATION_OF_RUNX3_EXPRESSIO | 55 | 0.35 | 0.74 | 0.88 | 1 |
| 5498 | MIR135B_5P | 266 | 0.31 | 0.74 | 0.971 | 1 |
| 5499 | MIR299_3P | 41 | 0.36 | 0.74 | 0.854 | 1 |
| 5500 | TAGGTCA_MIR192_MIR215 | 44 | 0.36 | 0.74 | 0.877 | 1 |
| 5501 | REACTOME_SPHINGOLIPID_METABOLISM | 77 | 0.34 | 0.74 | 0.899 | 1 |
| 5502 | REACTOME_NUCLEOTIDE_CATABOLISM | 24 | 0.4 | 0.74 | 0.848 | 1 |
| 5503 | RPS14_DN.V1_DN | 164 | 0.31 | 0.74 | 0.95 | 1 |
| 5504 | TOMLINS_PROSTATE_CANCER_UP | 38 | 0.36 | 0.74 | 0.876 | 1 |
| 5505 | YY1_01 | 206 | 0.31 | 0.74 | 0.956 | 1 |
| 5506 | MIR187_5P | 143 | 0.32 | 0.74 | 0.951 | 1 |
| 5507 | MIR5680 | 404 | 0.31 | 0.74 | 0.986 | 1 |
| 5508 | YAO_TEMPORAL_RESPONSE_TO_PROGESTERON | 57 | 0.35 | 0.74 | 0.886 | 1 |

| | | | | | | |
|---|---|---|---|---|---|---|
| 5509 | MIR4713_3P | 37 | 0.37 | 0.74 | 0.848 | 1 |
| 5510 | MIR520G_5P | 45 | 0.36 | 0.74 | 0.89 | 1 |
| 5511 | VERRECCHIA_RESPONSE_TO_TGFB1_C1 | 15 | 0.42 | 0.74 | 0.825 | 1 |
| 5512 | MIR512_3P | 96 | 0.33 | 0.74 | 0.918 | 1 |
| 5513 | WP_PRIMARY_FOCAL_SEGMENTAL_GLOMERULOS | 60 | 0.34 | 0.74 | 0.891 | 1 |
| 5514 | BIOCARTA_BCR_PATHWAY | 31 | 0.38 | 0.74 | 0.85 | 1 |
| 5515 | CTTTGTA_MIR524 | 396 | 0.3 | 0.74 | 0.983 | 1 |
| 5516 | VANTVEER_BREAST_CANCER_BRCA1_DN | 31 | 0.37 | 0.74 | 0.848 | 1 |
| 5517 | MIR1289 | 46 | 0.36 | 0.74 | 0.876 | 1 |
| 5518 | MIR676_3P | 46 | 0.36 | 0.74 | 0.894 | 1 |
| 5519 | REACTOME_TRISTETRAPROLIN_TTP_ZFP36_BINDS | 17 | 0.41 | 0.74 | 0.849 | 1 |
| 5520 | LIAO_HAVE_SOX4_BINDING_SITES | 34 | 0.37 | 0.74 | 0.85 | 1 |
| 5521 | MIR330_3P | 292 | 0.31 | 0.74 | 0.978 | 1 |
| 5522 | MIR6808_3P | 36 | 0.37 | 0.74 | 0.857 | 1 |
| 5523 | CREB_Q4_01 | 188 | 0.31 | 0.74 | 0.964 | 1 |
| 5524 | MIR802 | 119 | 0.32 | 0.74 | 0.946 | 1 |
| 5525 | RPA1_TARGET_GENES | 47 | 0.36 | 0.74 | 0.876 | 1 |
| 5526 | MIR660_5P | 63 | 0.34 | 0.74 | 0.891 | 1 |
| 5527 | REACTOME_BRANCHED_CHAIN_AMINO_ACID_CAT | 20 | 0.41 | 0.74 | 0.827 | 1 |
| 5528 | MIR6858_5P | 46 | 0.35 | 0.74 | 0.854 | 1 |
| 5529 | TAF9B_TARGET_GENES | 472 | 0.3 | 0.74 | 0.984 | 1 |
| 5530 | ZNF548_TARGET_GENES | 42 | 0.36 | 0.74 | 0.87 | 1 |
| 5531 | MIR548BA | 142 | 0.32 | 0.74 | 0.945 | 1 |
| 5532 | MIR4252 | 80 | 0.33 | 0.74 | 0.902 | 1 |
| 5533 | MIR8082 | 86 | 0.34 | 0.74 | 0.92 | 1 |
| 5534 | MIR6515_3P | 185 | 0.31 | 0.74 | 0.962 | 1 |
| 5535 | MIR3202 | 209 | 0.31 | 0.74 | 0.963 | 1 |
| 5536 | MIR103B | 34 | 0.37 | 0.74 | 0.874 | 1 |
| 5537 | MIR449B_5P | 232 | 0.31 | 0.74 | 0.978 | 1 |
| 5538 | MIR6867_3P | 91 | 0.33 | 0.74 | 0.921 | 1 |
| 5539 | NPAT_TARGET_GENES | 238 | 0.31 | 0.74 | 0.978 | 1 |
| 5540 | PID_FGF_PATHWAY | 43 | 0.35 | 0.74 | 0.875 | 1 |
| 5541 | BIOCARTA_AGR_PATHWAY | 32 | 0.37 | 0.74 | 0.866 | 1 |

| | | | | | | |
|---|---|---|---|---|---|---|
| 5542 | MIR376A_3P_MIR376B_3P | 114 | 0.32 | 0.74 | 0.94 | 1 |
| 5543 | RHEIN_ALL_GLUCOCORTICOID_THERAPY_DN | 344 | 0.3 | 0.74 | 0.979 | 1 |
| 5544 | PID_ARF_3PATHWAY | 18 | 0.4 | 0.74 | 0.814 | 1 |
| 5545 | MIR656_5P | 30 | 0.37 | 0.74 | 0.874 | 1 |
| 5546 | GTGTGAG_MIR342 | 60 | 0.34 | 0.74 | 0.908 | 1 |
| 5547 | MIR34C_5P | 233 | 0.31 | 0.74 | 0.979 | 1 |
| 5548 | LINSLEY_MIR16_TARGETS | 198 | 0.31 | 0.74 | 0.969 | 1 |
| 5549 | ZHONG_RESPONSE_TO_AZACITIDINE_AND_TSA_U | 148 | 0.31 | 0.74 | 0.951 | 1 |
| 5550 | ZNF169_TARGET_GENES | 37 | 0.37 | 0.74 | 0.876 | 1 |
| 5551 | MIR9902 | 240 | 0.31 | 0.74 | 0.976 | 1 |
| 5552 | REACTOME_TRANSPORT_OF_BILE_SALTS_AND_C | 50 | 0.35 | 0.74 | 0.885 | 1 |
| 5553 | MIR890 | 78 | 0.33 | 0.74 | 0.909 | 1 |
| 5554 | MIR503_3P | 31 | 0.37 | 0.74 | 0.841 | 1 |
| 5555 | KUMAR_PATHOGEN_LOAD_BY_MACROPHAGES | 206 | 0.31 | 0.74 | 0.965 | 1 |
| 5556 | MIR4327 | 72 | 0.33 | 0.74 | 0.913 | 1 |
| 5557 | MIR5584_5P | 222 | 0.31 | 0.74 | 0.977 | 1 |
| 5558 | MIR4483 | 77 | 0.33 | 0.74 | 0.914 | 1 |
| 5559 | MIR423_5P | 121 | 0.32 | 0.74 | 0.935 | 1 |
| 5560 | REACTOME_METABOLISM_OF_COFACTORS | 19 | 0.4 | 0.73 | 0.834 | 1 |
| 5561 | PGM3_TARGET_GENES | 247 | 0.31 | 0.73 | 0.977 | 1 |
| 5562 | MIR3609 | 310 | 0.3 | 0.73 | 0.986 | 1 |
| 5563 | MIR4435 | 73 | 0.33 | 0.73 | 0.911 | 1 |
| 5564 | NRL_DN.V1_DN | 102 | 0.32 | 0.73 | 0.919 | 1 |
| 5565 | HEY2_TARGET_GENES | 58 | 0.34 | 0.73 | 0.902 | 1 |
| 5566 | CREIGHTON_ENDOCRINE_THERAPY_RESISTANCE | 357 | 0.3 | 0.73 | 0.987 | 1 |
| 5567 | SP3_Q3 | 219 | 0.31 | 0.73 | 0.976 | 1 |
| 5568 | REACTOME_HEDGEHOG_OFF_STATE | 99 | 0.32 | 0.73 | 0.928 | 1 |
| 5569 | MIR514A_5P | 51 | 0.35 | 0.73 | 0.894 | 1 |
| 5570 | SHEPARD_BMYB_TARGETS | 65 | 0.34 | 0.73 | 0.919 | 1 |
| 5571 | MARTINEZ_RESPONSE_TO_TRABECTEDIN_DN | 262 | 0.3 | 0.73 | 0.978 | 1 |
| 5572 | WP_PYRIMIDINE_METABOLISM | 79 | 0.33 | 0.73 | 0.907 | 1 |
| 5573 | MIR6836_3P | 44 | 0.35 | 0.73 | 0.888 | 1 |
| 5574 | SMID_BREAST_CANCER_RELAPSE_IN_BRAIN_DN | 65 | 0.34 | 0.73 | 0.898 | 1 |

| | | | | | | |
|---|---|---|---|---|---|---|
| 5575 | LEE_DOUBLE_POLAR_THYMOCYTE | 20 | 0.4 | 0.73 | 0.834 | 1 |
| 5576 | MIR501_3P_MIR502_3P | 63 | 0.34 | 0.73 | 0.904 | 1 |
| 5577 | MIKKELSEN_IPS_ICP_WITH_H3K27ME3 | 24 | 0.39 | 0.73 | 0.869 | 1 |
| 5578 | TBK1.DN.48HRS_UP | 48 | 0.35 | 0.73 | 0.908 | 1 |
| 5579 | AIYAR_COBRA1_TARGETS_UP | 29 | 0.37 | 0.73 | 0.88 | 1 |
| 5580 | MIR504_5P | 48 | 0.35 | 0.73 | 0.892 | 1 |
| 5581 | MIR7112_3P | 81 | 0.33 | 0.73 | 0.92 | 1 |
| 5582 | THUM_SYSTOLIC_HEART_FAILURE_DN | 195 | 0.31 | 0.73 | 0.975 | 1 |
| 5583 | NAKAMURA_ADIPOGENESIS_LATE_UP | 82 | 0.33 | 0.73 | 0.92 | 1 |
| 5584 | MIR498_5P | 217 | 0.31 | 0.73 | 0.977 | 1 |
| 5585 | BANDRES_RESPONSE_TO_CARMUSTIN_MGMT_24 | 24 | 0.39 | 0.73 | 0.859 | 1 |
| 5586 | MIR9983_3P | 411 | 0.3 | 0.73 | 0.988 | 1 |
| 5587 | DAZARD_UV_RESPONSE_CLUSTER_G6 | 137 | 0.31 | 0.73 | 0.968 | 1 |
| 5588 | MIR145_3P | 63 | 0.33 | 0.73 | 0.91 | 1 |
| 5589 | LEE_EARLY_T_LYMPHOCYTE_UP | 97 | 0.32 | 0.73 | 0.945 | 1 |
| 5590 | MIR1264 | 105 | 0.32 | 0.73 | 0.941 | 1 |
| 5591 | PCGF1_TARGET_GENES | 495 | 0.3 | 0.73 | 0.992 | 1 |
| 5592 | MIR892C_5P | 236 | 0.3 | 0.73 | 0.979 | 1 |
| 5593 | MIR6841_5P | 32 | 0.36 | 0.73 | 0.886 | 1 |
| 5594 | MIR3191_3P | 20 | 0.4 | 0.73 | 0.848 | 1 |
| 5595 | MIR646 | 220 | 0.3 | 0.73 | 0.974 | 1 |
| 5596 | GEORGES_CELL_CYCLE_MIR192_TARGETS | 60 | 0.34 | 0.73 | 0.912 | 1 |
| 5597 | MARTIN_VIRAL_GPCR_SIGNALING_DN | 36 | 0.36 | 0.73 | 0.877 | 1 |
| 5598 | MIR873_3P | 43 | 0.35 | 0.73 | 0.884 | 1 |
| 5599 | MIR3170 | 46 | 0.34 | 0.73 | 0.89 | 1 |
| 5600 | HEN1_02 | 174 | 0.31 | 0.73 | 0.961 | 1 |
| 5601 | RIZKI_TUMOR_INVASIVENESS_2D_UP | 55 | 0.34 | 0.73 | 0.895 | 1 |
| 5602 | MIR2278 | 92 | 0.32 | 0.73 | 0.931 | 1 |
| 5603 | BOYLAN_MULTIPLE_MYELOMA_D_UP | 83 | 0.32 | 0.73 | 0.932 | 1 |
| 5604 | REACTOME_THE_ROLE_OF_GTSE1_IN_G2_M_PR( | 71 | 0.33 | 0.73 | 0.927 | 1 |
| 5605 | MIR6878_3P | 53 | 0.34 | 0.73 | 0.91 | 1 |
| 5606 | ACTGAAA_MIR30A3P_MIR30E3P | 174 | 0.31 | 0.73 | 0.965 | 1 |
| 5607 | MIR4530 | 121 | 0.32 | 0.73 | 0.951 | 1 |

| | | | | | | |
|---|---|---|---|---|---|---|
| 5608 | MIR4779 | 116 | 0.32 | 0.73 | 0.948 | 1 |
| 5609 | LEE_LIVER_CANCER_SURVIVAL_UP | 109 | 0.32 | 0.73 | 0.946 | 1 |
| 5610 | REACTOME_REGULATION_OF_PTEN_GENE_TRAN | 58 | 0.34 | 0.73 | 0.916 | 1 |
| 5611 | MIR3198 | 38 | 0.36 | 0.73 | 0.877 | 1 |
| 5612 | MIR4516 | 262 | 0.3 | 0.73 | 0.98 | 1 |
| 5613 | KEGG_GLYCINE_SERINE_AND_THREONINE_META | 24 | 0.38 | 0.72 | 0.873 | 1 |
| 5614 | HUI_MAPK14_TARGETS_UP | 15 | 0.41 | 0.72 | 0.834 | 1 |
| 5615 | REACTOME_PEROXISOMAL_PROTEIN_IMPORT | 57 | 0.34 | 0.72 | 0.891 | 1 |
| 5616 | MYCMAX_03 | 238 | 0.3 | 0.72 | 0.98 | 1 |
| 5617 | PARK_HSC_VS_MULTIPOTENT_PROGENITORS_UF | 17 | 0.41 | 0.72 | 0.862 | 1 |
| 5618 | MIR744_3P | 52 | 0.34 | 0.72 | 0.902 | 1 |
| 5619 | MIR1270 | 63 | 0.33 | 0.72 | 0.919 | 1 |
| 5620 | MIR212_3P | 171 | 0.31 | 0.72 | 0.965 | 1 |
| 5621 | TGCTTTG_MIR330 | 295 | 0.3 | 0.72 | 0.992 | 1 |
| 5622 | NIKOLSKY_BREAST_CANCER_12Q13_Q21_AMPLI( | 39 | 0.35 | 0.72 | 0.9 | 1 |
| 5623 | LIU_CDX2_TARGETS_UP | 20 | 0.39 | 0.72 | 0.873 | 1 |
| 5624 | MIR6810_5P | 61 | 0.34 | 0.72 | 0.911 | 1 |
| 5625 | KIM_WT1_TARGETS_8HR_DN | 99 | 0.32 | 0.72 | 0.937 | 1 |
| 5626 | MIR6858_3P | 34 | 0.36 | 0.72 | 0.888 | 1 |
| 5627 | WP_NUCLEAR_RECEPTORS | 30 | 0.37 | 0.72 | 0.883 | 1 |
| 5628 | MIR466 | 349 | 0.3 | 0.72 | 0.987 | 1 |
| 5629 | MIR7107_3P | 47 | 0.34 | 0.72 | 0.892 | 1 |
| 5630 | REACTOME_BIOSYNTHESIS_OF_THE_N_GLYCAN_ | 70 | 0.33 | 0.72 | 0.936 | 1 |
| 5631 | TCTATGA_MIR376A_MIR376B | 76 | 0.33 | 0.72 | 0.936 | 1 |
| 5632 | PAX8_01 | 34 | 0.36 | 0.72 | 0.877 | 1 |
| 5633 | WP_ETHANOL_EFFECTS_ON_HISTONE_MODIFICA' | 25 | 0.38 | 0.72 | 0.876 | 1 |
| 5634 | WP_7OXOC_AND_7BETAHC_PATHWAYS | 21 | 0.39 | 0.72 | 0.85 | 1 |
| 5635 | MIR4724_3P | 69 | 0.33 | 0.72 | 0.929 | 1 |
| 5636 | MIR1226_5P | 39 | 0.35 | 0.72 | 0.903 | 1 |
| 5637 | AMBROSINI_FLAVOPIRIDOL_TREATMENT_TP53 | 97 | 0.32 | 0.72 | 0.939 | 1 |
| 5638 | MIR2861 | 82 | 0.32 | 0.72 | 0.949 | 1 |
| 5639 | REACTOME_DOWNSTREAM_SIGNALING_OF_ACTI\ | 18 | 0.4 | 0.72 | 0.854 | 1 |
| 5640 | MIR4735_5P | 149 | 0.31 | 0.72 | 0.967 | 1 |

| | | | | | | |
|---|---|---|---|---|---|---|
| 5641 | RORA1_01 | 205 | 0.3 | 0.72 | 0.976 | 1 |
| 5642 | ZHAN_MULTIPLE_MYELOMA_HP_DN | 40 | 0.35 | 0.72 | 0.89 | 1 |
| 5643 | ELVIDGE_HYPOXIA_BY_DMOG_DN | 55 | 0.34 | 0.72 | 0.911 | 1 |
| 5644 | MIR6869_5P | 34 | 0.36 | 0.72 | 0.9 | 1 |
| 5645 | MIR4310 | 251 | 0.3 | 0.72 | 0.987 | 1 |
| 5646 | MEF2_01 | 91 | 0.32 | 0.72 | 0.942 | 1 |
| 5647 | MIR3165 | 30 | 0.37 | 0.72 | 0.875 | 1 |
| 5648 | MIR6838_5P | 480 | 0.29 | 0.72 | 0.995 | 1 |
| 5649 | MIR548AH_5P | 310 | 0.3 | 0.72 | 0.988 | 1 |
| 5650 | MIR3692_3P | 260 | 0.3 | 0.72 | 0.986 | 1 |
| 5651 | MIR8069 | 49 | 0.34 | 0.72 | 0.913 | 1 |
| 5652 | METHYLCYTOSINE_DIOXYGENASE_TET_UNIPROT | 136 | 0.31 | 0.72 | 0.959 | 1 |
| 5653 | PAF1_TARGET_GENES | 62 | 0.33 | 0.72 | 0.919 | 1 |
| 5654 | MIR216B_5P | 75 | 0.32 | 0.72 | 0.945 | 1 |
| 5655 | MIR367_5P | 58 | 0.34 | 0.72 | 0.923 | 1 |
| 5656 | MIR212_5P | 84 | 0.32 | 0.72 | 0.939 | 1 |
| 5657 | SF1_Q6 | 233 | 0.3 | 0.72 | 0.979 | 1 |
| 5658 | JEON_SMAD6_TARGETS_DN | 18 | 0.39 | 0.72 | 0.862 | 1 |
| 5659 | KEGG_ONE_CARBON_POOL_BY_FOLATE | 16 | 0.4 | 0.72 | 0.839 | 1 |
| 5660 | MIR3677_5P | 22 | 0.39 | 0.72 | 0.853 | 1 |
| 5661 | MIR132_3P | 173 | 0.3 | 0.72 | 0.984 | 1 |
| 5662 | MIR548O_3P | 252 | 0.3 | 0.72 | 0.989 | 1 |
| 5663 | CHEN_PDGF_TARGETS | 17 | 0.39 | 0.72 | 0.861 | 1 |
| 5664 | MIR513B_5P | 251 | 0.3 | 0.72 | 0.988 | 1 |
| 5665 | REACTOME_PKA_MEDIATED_PHOSPHORYLATION | 15 | 0.4 | 0.72 | 0.846 | 1 |
| 5666 | MIR361_5P | 133 | 0.31 | 0.72 | 0.96 | 1 |
| 5667 | MIR1245B_5P | 54 | 0.33 | 0.72 | 0.922 | 1 |
| 5668 | MIR6852_3P | 124 | 0.31 | 0.71 | 0.959 | 1 |
| 5669 | MIR340_3P | 44 | 0.35 | 0.71 | 0.91 | 1 |
| 5670 | TAATGTG_MIR323 | 151 | 0.31 | 0.71 | 0.977 | 1 |
| 5671 | MIR4436B_3P | 94 | 0.32 | 0.71 | 0.954 | 1 |
| 5672 | MIR6837_5P | 101 | 0.31 | 0.71 | 0.955 | 1 |
| 5673 | MIR1248 | 106 | 0.32 | 0.71 | 0.953 | 1 |

| | | | | | | |
|---|---|---|---|---|---|---|
| 5674 | MIR576_5P | 299 | 0.29 | 0.71 | 0.99 | 1 |
| 5675 | MIR5093 | 203 | 0.3 | 0.71 | 0.981 | 1 |
| 5676 | REACTOME_MITOTIC_G2_G2_M_PHASES | 190 | 0.3 | 0.71 | 0.981 | 1 |
| 5677 | ZNF101_TARGET_GENES | 217 | 0.3 | 0.71 | 0.991 | 1 |
| 5678 | MIR4649_3P | 176 | 0.3 | 0.71 | 0.982 | 1 |
| 5679 | REACTOME_ER_QUALITY_CONTROL_COMPARTME | 21 | 0.38 | 0.71 | 0.865 | 1 |
| 5680 | KAMIKUBO_MYELOID_MN1_NETWORK | 17 | 0.4 | 0.71 | 0.857 | 1 |
| 5681 | MIR182_5P | 492 | 0.29 | 0.71 | 0.999 | 1 |
| 5682 | MIR5100 | 79 | 0.32 | 0.71 | 0.934 | 1 |
| 5683 | REACTOME_NEPHRIN_FAMILY_INTERACTIONS | 21 | 0.38 | 0.71 | 0.866 | 1 |
| 5684 | FIGUEROA_AML_METHYLATION_CLUSTER_6_UP | 116 | 0.31 | 0.71 | 0.963 | 1 |
| 5685 | MIR6768_3P | 125 | 0.3 | 0.71 | 0.971 | 1 |
| 5686 | HOLLEMAN_ASPARAGINASE_RESISTANCE_B_ALL | 25 | 0.37 | 0.71 | 0.895 | 1 |
| 5687 | ATGCTGC_MIR103_MIR107 | 198 | 0.3 | 0.71 | 0.977 | 1 |
| 5688 | AUTS2_TARGET_GENES | 399 | 0.29 | 0.71 | 0.998 | 1 |
| 5689 | MIR154_3P | 89 | 0.32 | 0.71 | 0.952 | 1 |
| 5690 | NRF1_Q6 | 249 | 0.29 | 0.71 | 0.984 | 1 |
| 5691 | HOLLEMAN_VINCRISTINE_RESISTANCE_B_ALL_UI | 38 | 0.35 | 0.71 | 0.912 | 1 |
| 5692 | MIR3192_3P | 68 | 0.32 | 0.71 | 0.949 | 1 |
| 5693 | PDGF_UP.V1_UP | 129 | 0.31 | 0.71 | 0.964 | 1 |
| 5694 | MIR4759 | 40 | 0.35 | 0.71 | 0.897 | 1 |
| 5695 | REACTOME_THROMBOXANE_SIGNALLING_THROU | 16 | 0.4 | 0.71 | 0.863 | 1 |
| 5696 | MORI_EMU_MYC_LYMPHOMA_BY_ONSET_TIME_U | 107 | 0.31 | 0.71 | 0.956 | 1 |
| 5697 | REACTOME_TRNA_AMINOACYLATION | 41 | 0.35 | 0.71 | 0.92 | 1 |
| 5698 | MIR6870_3P | 37 | 0.35 | 0.71 | 0.892 | 1 |
| 5699 | RAMALHO_STEMNESS_UP | 188 | 0.3 | 0.71 | 0.976 | 1 |
| 5700 | MIR597_3P | 92 | 0.32 | 0.71 | 0.965 | 1 |
| 5701 | REACTOME_N_GLYCAN_TRIMMING_IN_THE_ER_AI | 35 | 0.35 | 0.71 | 0.896 | 1 |
| 5702 | MIR4253 | 42 | 0.34 | 0.71 | 0.91 | 1 |
| 5703 | MIR513A_3P_MIR513C_3P | 383 | 0.29 | 0.71 | 0.997 | 1 |
| 5704 | AGUIRRE_PANCREATIC_CANCER_COPY_NUMBER_ | 284 | 0.29 | 0.71 | 0.993 | 1 |
| 5705 | MIR5581_3P | 112 | 0.31 | 0.71 | 0.96 | 1 |
| 5706 | ZHOU_INFLAMMATORY_RESPONSE_LIVE_DN | 308 | 0.29 | 0.71 | 0.996 | 1 |

| | | | | | | |
|---|---|---|---|---|---|---|
| 5707 | IKEDA_MIR133_TARGETS_UP | 43 | 0.34 | 0.71 | 0.933 | 1 |
| 5708 | NR1I2_TARGET_GENES | 304 | 0.29 | 0.71 | 0.996 | 1 |
| 5709 | ZFP28_TARGET_GENES | 74 | 0.32 | 0.71 | 0.944 | 1 |
| 5710 | BOYAULT_LIVER_CANCER_SUBCLASS_G123_UP | 43 | 0.34 | 0.71 | 0.914 | 1 |
| 5711 | MIR30C_1_3P | 105 | 0.31 | 0.71 | 0.955 | 1 |
| 5712 | BURTON_ADIPOGENESIS_6 | 159 | 0.3 | 0.71 | 0.972 | 1 |
| 5713 | MIR3613_5P | 64 | 0.32 | 0.71 | 0.93 | 1 |
| 5714 | MIR8087 | 128 | 0.3 | 0.71 | 0.966 | 1 |
| 5715 | VALK_AML_CLUSTER_4 | 22 | 0.38 | 0.71 | 0.896 | 1 |
| 5716 | MIR3907 | 41 | 0.34 | 0.71 | 0.919 | 1 |
| 5717 | MIR548AI_MIR570_5P | 144 | 0.3 | 0.7 | 0.975 | 1 |
| 5718 | MIR3942_3P | 284 | 0.29 | 0.7 | 0.995 | 1 |
| 5719 | SOX5_01 | 214 | 0.29 | 0.7 | 0.985 | 1 |
| 5720 | MIR6818_5P | 91 | 0.31 | 0.7 | 0.961 | 1 |
| 5721 | MIR3184_5P | 117 | 0.31 | 0.7 | 0.965 | 1 |
| 5722 | MIR4302 | 116 | 0.31 | 0.7 | 0.966 | 1 |
| 5723 | MIR1284 | 201 | 0.29 | 0.7 | 0.99 | 1 |
| 5724 | NADERI_BREAST_CANCER_PROGNOSIS_UP | 44 | 0.34 | 0.7 | 0.912 | 1 |
| 5725 | DANG_MYC_TARGETS_UP | 122 | 0.31 | 0.7 | 0.979 | 1 |
| 5726 | CAGNWMCNNNGAC_UNKNOWN | 67 | 0.32 | 0.7 | 0.944 | 1 |
| 5727 | MARTINEZ_RESPONSE_TO_TRABECTEDIN | 47 | 0.34 | 0.7 | 0.924 | 1 |
| 5728 | MIR548U | 140 | 0.3 | 0.7 | 0.977 | 1 |
| 5729 | BACOLOD_RESISTANCE_TO_ALKYLATING_AGENT | 18 | 0.38 | 0.7 | 0.869 | 1 |
| 5730 | MTF1_Q4 | 227 | 0.29 | 0.7 | 0.988 | 1 |
| 5731 | ONKEN_UVEAL_MELANOMA_DN | 462 | 0.29 | 0.7 | 0.999 | 1 |
| 5732 | KEGG_FC_GAMMA_R_MEDIATED_PHAGOCYTOSIS | 80 | 0.31 | 0.7 | 0.961 | 1 |
| 5733 | GCTCTTG_MIR335 | 81 | 0.31 | 0.7 | 0.957 | 1 |
| 5734 | REACTOME_UNBLOCKING_OF_NMDA_RECEPTORS | 16 | 0.39 | 0.7 | 0.858 | 1 |
| 5735 | MIR3192_5P | 136 | 0.3 | 0.7 | 0.973 | 1 |
| 5736 | CAGCCTC_MIR4855P | 131 | 0.3 | 0.7 | 0.975 | 1 |
| 5737 | YRCCAKNNGNCGC_UNKNOWN | 62 | 0.32 | 0.7 | 0.945 | 1 |
| 5738 | MIR6764_3P | 95 | 0.31 | 0.7 | 0.965 | 1 |
| 5739 | LOCKWOOD_AMPLIFIED_IN_LUNG_CANCER | 210 | 0.29 | 0.7 | 0.991 | 1 |

| | | | | | | |
|---|---|---|---|---|---|---|
| 5740 | MIR12124 | 281 | 0.29 | 0.7 | 0.991 | 1 |
| 5741 | TCCCRNNRTGC_UNKNOWN | 203 | 0.29 | 0.7 | 0.99 | 1 |
| 5742 | AP2GAMMA_01 | 224 | 0.29 | 0.7 | 0.99 | 1 |
| 5743 | MIR424_5P | 468 | 0.29 | 0.7 | 0.999 | 1 |
| 5744 | REACTOME_INTRA_GOLGI_TRAFFIC | 43 | 0.34 | 0.7 | 0.924 | 1 |
| 5745 | MIR6500_3P | 70 | 0.32 | 0.7 | 0.94 | 1 |
| 5746 | MIR6513_5P | 78 | 0.31 | 0.7 | 0.949 | 1 |
| 5747 | SESTO_RESPONSE_TO_UV_C4 | 17 | 0.39 | 0.7 | 0.877 | 1 |
| 5748 | MIR3142 | 53 | 0.33 | 0.7 | 0.936 | 1 |
| 5749 | SREBP1_02 | 68 | 0.32 | 0.7 | 0.949 | 1 |
| 5750 | MIR487A_3P | 90 | 0.31 | 0.7 | 0.962 | 1 |
| 5751 | REACTOME_SIGNALING_BY_NODAL | 15 | 0.39 | 0.7 | 0.881 | 1 |
| 5752 | MIR589_3P | 214 | 0.29 | 0.7 | 0.996 | 1 |
| 5753 | MIR7110_5P | 141 | 0.3 | 0.7 | 0.982 | 1 |
| 5754 | SRPK2_TARGET_GENES | 45 | 0.33 | 0.7 | 0.927 | 1 |
| 5755 | MIR5583_5P | 86 | 0.32 | 0.7 | 0.952 | 1 |
| 5756 | OUILLETTE_CLL_13Q14_DELETION_DN | 47 | 0.33 | 0.7 | 0.924 | 1 |
| 5757 | MIR3686 | 289 | 0.29 | 0.7 | 0.996 | 1 |
| 5758 | MIR6821_3P | 42 | 0.34 | 0.7 | 0.929 | 1 |
| 5759 | ZBTB5_TARGET_GENES | 159 | 0.3 | 0.7 | 0.987 | 1 |
| 5760 | REACTOME_MITOCHONDRIAL_BIOGENESIS | 90 | 0.31 | 0.7 | 0.969 | 1 |
| 5761 | MIR9_3P | 309 | 0.29 | 0.7 | 0.994 | 1 |
| 5762 | MILI_PSEUDOPODIA_HAPTOTAXIS_UP | 499 | 0.29 | 0.7 | 0.999 | 1 |
| 5763 | MIR4303 | 191 | 0.29 | 0.7 | 0.987 | 1 |
| 5764 | MIR4769_5P | 29 | 0.36 | 0.7 | 0.917 | 1 |
| 5765 | ACTGCCT_MIR34B | 202 | 0.29 | 0.7 | 0.995 | 1 |
| 5766 | HOXD11_TARGET_GENES | 129 | 0.3 | 0.7 | 0.975 | 1 |
| 5767 | MIR6894_5P | 33 | 0.35 | 0.7 | 0.912 | 1 |
| 5768 | MIR6824_3P | 95 | 0.31 | 0.7 | 0.961 | 1 |
| 5769 | REACTOME_CRMPS_IN_SEMA3A_SIGNALING | 16 | 0.39 | 0.7 | 0.874 | 1 |
| 5770 | REACTOME_BASIGIN_INTERACTIONS | 20 | 0.38 | 0.7 | 0.875 | 1 |
| 5771 | BIOCARTA_ECM_PATHWAY | 17 | 0.39 | 0.69 | 0.873 | 1 |
| 5772 | YEGNASUBRAMANIAN_PROSTATE_CANCER | 93 | 0.31 | 0.69 | 0.97 | 1 |

| | | | | | | |
|---|---|---|---|---|---|---|
| 5773 | EMX1_TARGET_GENES | 198 | 0.29 | 0.69 | 0.992 | 1 |
| 5774 | REACTOME_SIGNALING_BY_HEDGEHOG | 130 | 0.3 | 0.69 | 0.975 | 1 |
| 5775 | REACTOME_RECYCLING_PATHWAY_OF_L1 | 42 | 0.33 | 0.69 | 0.916 | 1 |
| 5776 | WP_DOPAMINERGIC_NEUROGENESIS | 19 | 0.38 | 0.69 | 0.886 | 1 |
| 5777 | MIR4632_5P | 88 | 0.31 | 0.69 | 0.97 | 1 |
| 5778 | EGR3_01 | 71 | 0.32 | 0.69 | 0.954 | 1 |
| 5779 | MIR1229_5P | 28 | 0.36 | 0.69 | 0.909 | 1 |
| 5780 | SP1_Q4_01 | 237 | 0.29 | 0.69 | 0.997 | 1 |
| 5781 | WP_HEMATOPOIETIC_STEM_CELL_GENE_REGUL/ | 20 | 0.37 | 0.69 | 0.889 | 1 |
| 5782 | REACTOME_TRAFFICKING_OF_AMPA_RECEPTORS | 25 | 0.36 | 0.69 | 0.903 | 1 |
| 5783 | ARNT_02 | 233 | 0.29 | 0.69 | 0.992 | 1 |
| 5784 | ZNF784_TARGET_GENES | 373 | 0.28 | 0.69 | 0.998 | 1 |
| 5785 | MIR4505 | 221 | 0.29 | 0.69 | 0.991 | 1 |
| 5786 | MIR6882_5P | 151 | 0.29 | 0.69 | 0.987 | 1 |
| 5787 | DARWICHE_SKIN_TUMOR_PROMOTER_DN | 143 | 0.29 | 0.69 | 0.988 | 1 |
| 5788 | PUJANA_BRCA_CENTERED_NETWORK | 118 | 0.3 | 0.69 | 0.974 | 1 |
| 5789 | LIU_COMMON_CANCER_GENES | 58 | 0.32 | 0.69 | 0.946 | 1 |
| 5790 | LEE_TARGETS_OF_PTCH1_AND_SUFU_DN | 64 | 0.32 | 0.69 | 0.94 | 1 |
| 5791 | MIR4772_5P | 42 | 0.33 | 0.69 | 0.931 | 1 |
| 5792 | MIR410_3P | 277 | 0.29 | 0.69 | 0.999 | 1 |
| 5793 | MIR3074_5P | 133 | 0.3 | 0.69 | 0.98 | 1 |
| 5794 | FUJII_YBX1_TARGETS_DN | 186 | 0.29 | 0.69 | 0.991 | 1 |
| 5795 | CHOI_ATL_STAGE_PREDICTOR | 42 | 0.33 | 0.69 | 0.931 | 1 |
| 5796 | ZNF16_TARGET_GENES | 66 | 0.31 | 0.69 | 0.946 | 1 |
| 5797 | MIR325 | 27 | 0.35 | 0.69 | 0.903 | 1 |
| 5798 | TCTGGAC_MIR198 | 74 | 0.31 | 0.69 | 0.954 | 1 |
| 5799 | PAX5_02 | 16 | 0.39 | 0.69 | 0.888 | 1 |
| 5800 | WP_GLYCOSAMINOGLYCAN_DEGRADATION | 15 | 0.39 | 0.69 | 0.877 | 1 |
| 5801 | MIR449A | 273 | 0.28 | 0.69 | 0.995 | 1 |
| 5802 | MIR5697 | 137 | 0.3 | 0.69 | 0.982 | 1 |
| 5803 | MOOTHA_HUMAN_MITODB_6_2002 | 397 | 0.28 | 0.69 | 0.997 | 1 |
| 5804 | AGCYRWTTC_UNKNOWN | 90 | 0.31 | 0.69 | 0.971 | 1 |
| 5805 | MIR5089_3P | 58 | 0.32 | 0.69 | 0.949 | 1 |

| | | | | | | |
|---|---|---|---|---|---|---|
| 5806 | MIR4750_3P | 36 | 0.34 | 0.69 | 0.939 | 1 |
| 5807 | KYNG_RESPONSE_TO_H2O2_VIA_ERCC6_UP | 32 | 0.34 | 0.69 | 0.941 | 1 |
| 5808 | GARGALOVIC_RESPONSE_TO_OXIDIZED_PHOSPH | 24 | 0.37 | 0.69 | 0.908 | 1 |
| 5809 | NUMATA_CSF3_SIGNALING_VIA_STAT3 | 17 | 0.38 | 0.69 | 0.89 | 1 |
| 5810 | MIR409_3P | 79 | 0.31 | 0.68 | 0.958 | 1 |
| 5811 | MIR5004_5P | 101 | 0.3 | 0.68 | 0.972 | 1 |
| 5812 | CAHOY_OLIGODENDROCUTIC | 82 | 0.31 | 0.68 | 0.97 | 1 |
| 5813 | BOYLAN_MULTIPLE_MYELOMA_D_CLUSTER_DN | 30 | 0.35 | 0.68 | 0.942 | 1 |
| 5814 | ACEVEDO_LIVER_CANCER_WITH_H3K9ME3_UP | 67 | 0.32 | 0.68 | 0.949 | 1 |
| 5815 | MIR3149 | 126 | 0.3 | 0.68 | 0.972 | 1 |
| 5816 | LE_NEURONAL_DIFFERENTIATION_UP | 16 | 0.39 | 0.68 | 0.914 | 1 |
| 5817 | MIR34A_5P | 276 | 0.28 | 0.68 | 0.998 | 1 |
| 5818 | PRMT5_TARGET_GENES | 66 | 0.31 | 0.68 | 0.951 | 1 |
| 5819 | MIR575 | 46 | 0.32 | 0.68 | 0.937 | 1 |
| 5820 | MIR12117 | 73 | 0.31 | 0.68 | 0.968 | 1 |
| 5821 | REACTOME_SIGNALING_BY_FGFR2 | 59 | 0.32 | 0.68 | 0.963 | 1 |
| 5822 | EVI1_03 | 35 | 0.34 | 0.68 | 0.928 | 1 |
| 5823 | AACTGGA_MIR145 | 210 | 0.29 | 0.68 | 0.992 | 1 |
| 5824 | MIR100_3P | 59 | 0.32 | 0.68 | 0.947 | 1 |
| 5825 | MIR380_3P | 179 | 0.29 | 0.68 | 0.99 | 1 |
| 5826 | MIR548BC | 425 | 0.28 | 0.68 | 0.998 | 1 |
| 5827 | ZNF329_TARGET_GENES | 497 | 0.28 | 0.68 | 1 | 1 |
| 5828 | ZNF667_TARGET_GENES | 133 | 0.29 | 0.68 | 0.991 | 1 |
| 5829 | REACTOME_FCGAMMA_RECEPTOR_FCGR_DEPEN | 76 | 0.31 | 0.68 | 0.958 | 1 |
| 5830 | KDM1B_TARGET_GENES | 76 | 0.31 | 0.68 | 0.97 | 1 |
| 5831 | ZNF223_TARGET_GENES | 269 | 0.28 | 0.68 | 0.999 | 1 |
| 5832 | MIR6854_5P | 62 | 0.31 | 0.68 | 0.956 | 1 |
| 5833 | PID_INSULIN_GLUCOSE_PATHWAY | 23 | 0.36 | 0.68 | 0.914 | 1 |
| 5834 | LEIN_MIDBRAIN_MARKERS | 63 | 0.31 | 0.68 | 0.956 | 1 |
| 5835 | MIR451B | 143 | 0.29 | 0.68 | 0.992 | 1 |
| 5836 | MIR101_3P | 406 | 0.28 | 0.68 | 0.999 | 1 |
| 5837 | RASHI_RESPONSE_TO_IONIZING_RADIATION_5 | 106 | 0.3 | 0.68 | 0.978 | 1 |
| 5838 | MIR551B_5P | 445 | 0.28 | 0.68 | 0.999 | 1 |

| | | | | | | |
|---|---|---|---|---|---|---|
| 5839 | MIR3928_5P | 85 | 0.3 | 0.68 | 0.972 | 1 |
| 5840 | MIR548AS_3P | 233 | 0.28 | 0.68 | 0.995 | 1 |
| 5841 | KEGG_NICOTINATE_AND_NICOTINAMIDE_METABC | 15 | 0.39 | 0.68 | 0.876 | 1 |
| 5842 | VANTVEER_BREAST_CANCER_ESR1_UP | 132 | 0.29 | 0.68 | 0.987 | 1 |
| 5843 | MIR509_3P | 47 | 0.33 | 0.68 | 0.956 | 1 |
| 5844 | REACTOME_BUTYRATE_RESPONSE_FACTOR_1_E | 17 | 0.38 | 0.68 | 0.893 | 1 |
| 5845 | AGUIRRE_PANCREATIC_CANCER_COPY_NUMBER_ | 222 | 0.28 | 0.68 | 0.997 | 1 |
| 5846 | YAO_TEMPORAL_RESPONSE_TO_PROGESTERON | 68 | 0.31 | 0.68 | 0.97 | 1 |
| 5847 | MIR1914_5P | 51 | 0.32 | 0.68 | 0.94 | 1 |
| 5848 | KOKKINAKIS_METHIONINE_DEPRIVATION_48HR_D | 55 | 0.32 | 0.68 | 0.964 | 1 |
| 5849 | SOTIRIOU_BREAST_CANCER_GRADE_1_VS_3_UP | 152 | 0.29 | 0.68 | 0.99 | 1 |
| 5850 | MIR6736_5P | 34 | 0.33 | 0.68 | 0.928 | 1 |
| 5851 | MIR4797_5P | 106 | 0.3 | 0.68 | 0.981 | 1 |
| 5852 | PER1_TARGET_GENES | 27 | 0.34 | 0.68 | 0.933 | 1 |
| 5853 | MIR4666A_3P | 482 | 0.28 | 0.68 | 0.999 | 1 |
| 5854 | KEGG_PURINE_METABOLISM | 132 | 0.29 | 0.68 | 0.987 | 1 |
| 5855 | MIR519D_5P | 202 | 0.29 | 0.68 | 0.995 | 1 |
| 5856 | MIR6745 | 77 | 0.31 | 0.68 | 0.969 | 1 |
| 5857 | BOYAULT_LIVER_CANCER_SUBCLASS_G6_UP | 57 | 0.32 | 0.68 | 0.954 | 1 |
| 5858 | MIR497_5P | 468 | 0.28 | 0.68 | 0.999 | 1 |
| 5859 | AMIT_EGF_RESPONSE_480_MCF10A | 39 | 0.33 | 0.68 | 0.942 | 1 |
| 5860 | REACTOME_SIALIC_ACID_METABOLISM | 26 | 0.35 | 0.68 | 0.915 | 1 |
| 5861 | MIR6744_3P | 54 | 0.31 | 0.68 | 0.958 | 1 |
| 5862 | KEGG_TYROSINE_METABOLISM | 23 | 0.36 | 0.68 | 0.919 | 1 |
| 5863 | MIR3938 | 64 | 0.31 | 0.67 | 0.963 | 1 |
| 5864 | MIR5191 | 19 | 0.37 | 0.67 | 0.898 | 1 |
| 5865 | WP_NOTCH1_REGULATION_OF_ENDOTHELIAL_CE | 15 | 0.39 | 0.67 | 0.902 | 1 |
| 5866 | E2F_03 | 235 | 0.28 | 0.67 | 0.997 | 1 |
| 5867 | ZNF707_TARGET_GENES | 33 | 0.34 | 0.67 | 0.931 | 1 |
| 5868 | MIR6851_5P | 91 | 0.3 | 0.67 | 0.978 | 1 |
| 5869 | MIR5685 | 64 | 0.31 | 0.67 | 0.967 | 1 |
| 5870 | WP_TYPE_2_PAPILLARY_RENAL_CELL_CARCINOI | 33 | 0.33 | 0.67 | 0.93 | 1 |
| 5871 | GGCNRNWCTTYS_UNKNOWN | 79 | 0.3 | 0.67 | 0.974 | 1 |

| | | | | | | |
|---|---|---|---|---|---|---|
| 5872 | KEGG_PYRIMIDINE_METABOLISM | 88 | 0.3 | 0.67 | 0.978 | 1 |
| 5873 | MIR6784_3P | 34 | 0.34 | 0.67 | 0.945 | 1 |
| 5874 | SREBP1_01 | 156 | 0.28 | 0.67 | 0.994 | 1 |
| 5875 | MIR3925_3P | 133 | 0.29 | 0.67 | 0.988 | 1 |
| 5876 | PID_RHOA_REG_PATHWAY | 40 | 0.33 | 0.67 | 0.951 | 1 |
| 5877 | BARRIER_COLON_CANCER_RECURRENCE_UP | 33 | 0.34 | 0.67 | 0.931 | 1 |
| 5878 | DUTERTRE_ESTRADIOL_RESPONSE_24HR_UP | 306 | 0.28 | 0.67 | 0.999 | 1 |
| 5879 | MIR4293 | 89 | 0.3 | 0.67 | 0.982 | 1 |
| 5880 | MIR4639_3P | 63 | 0.31 | 0.67 | 0.957 | 1 |
| 5881 | GTTTGTT_MIR495 | 241 | 0.28 | 0.67 | 0.998 | 1 |
| 5882 | CREB3_TARGET_GENES | 343 | 0.28 | 0.67 | 1 | 1 |
| 5883 | RORA2_01 | 126 | 0.29 | 0.67 | 0.988 | 1 |
| 5884 | REACTOME_DEACTIVATION_OF_THE_BETA_CATE | 37 | 0.33 | 0.67 | 0.949 | 1 |
| 5885 | CHEMNITZ_RESPONSE_TO_PROSTAGLANDIN_E2_ | 141 | 0.29 | 0.67 | 0.998 | 1 |
| 5886 | MIR3945 | 34 | 0.34 | 0.67 | 0.944 | 1 |
| 5887 | PID_CDC42_REG_PATHWAY | 28 | 0.34 | 0.67 | 0.941 | 1 |
| 5888 | MIR548E_3P | 424 | 0.27 | 0.67 | 1 | 1 |
| 5889 | MIR3129_3P | 86 | 0.3 | 0.67 | 0.974 | 1 |
| 5890 | MIR5196_3P | 53 | 0.32 | 0.67 | 0.957 | 1 |
| 5891 | VEGF_A_UP.V1_DN | 171 | 0.28 | 0.67 | 0.996 | 1 |
| 5892 | MIR584_3P | 92 | 0.29 | 0.67 | 0.984 | 1 |
| 5893 | ZHENG_FOXP3_TARGETS_IN_THYMUS_UP | 171 | 0.28 | 0.67 | 0.998 | 1 |
| 5894 | WHITEFORD_PEDIATRIC_CANCER_MARKERS | 113 | 0.29 | 0.67 | 0.992 | 1 |
| 5895 | MIR3074_3P | 77 | 0.3 | 0.67 | 0.981 | 1 |
| 5896 | REACTOME_CARGO_CONCENTRATION_IN_THE_EI | 28 | 0.34 | 0.66 | 0.93 | 1 |
| 5897 | MIR4318 | 104 | 0.29 | 0.66 | 0.992 | 1 |
| 5898 | GABRIELY_MIR21_TARGETS | 272 | 0.27 | 0.66 | 1 | 1 |
| 5899 | WP_NAD_BIOSYNTHETIC_PATHWAYS | 17 | 0.37 | 0.66 | 0.914 | 1 |
| 5900 | MIR8084 | 214 | 0.28 | 0.66 | 0.996 | 1 |
| 5901 | MIR548A_3P | 425 | 0.27 | 0.66 | 1 | 1 |
| 5902 | MIR4499 | 98 | 0.29 | 0.66 | 0.986 | 1 |
| 5903 | DARWICHE_PAPILLOMA_RISK_HIGH_DN | 143 | 0.28 | 0.66 | 0.991 | 1 |
| 5904 | TOYOTA_TARGETS_OF_MIR34B_AND_MIR34C | 440 | 0.27 | 0.66 | 1 | 1 |

| | | | | | | |
|---|---|---|---|---|---|---|
| 5905 | MIR548F_3P | 428 | 0.27 | 0.66 | 1 | 1 |
| 5906 | CHIANG_LIVER_CANCER_SUBCLASS_PROLIFERAT | 166 | 0.28 | 0.66 | 0.997 | 1 |
| 5907 | MIR202_5P | 96 | 0.29 | 0.66 | 0.983 | 1 |
| 5908 | CREB_02 | 225 | 0.28 | 0.66 | 0.996 | 1 |
| 5909 | ATCATGA_MIR433 | 105 | 0.29 | 0.66 | 0.989 | 1 |
| 5910 | AACWWCAANK_UNKNOWN | 127 | 0.28 | 0.66 | 0.997 | 1 |
| 5911 | MIR548AZ_3P | 429 | 0.27 | 0.66 | 1 | 1 |
| 5912 | OVOL3_TARGET_GENES | 447 | 0.27 | 0.66 | 1 | 1 |
| 5913 | MIR365A_3P_MIR365B_3P | 101 | 0.29 | 0.66 | 0.994 | 1 |
| 5914 | ZHAN_MULTIPLE_MYELOMA_CD2_UP | 35 | 0.33 | 0.66 | 0.947 | 1 |
| 5915 | MIR4493 | 82 | 0.29 | 0.66 | 0.988 | 1 |
| 5916 | BIOCARTA_PAR1_PATHWAY | 15 | 0.38 | 0.66 | 0.918 | 1 |
| 5917 | MIR4254 | 80 | 0.3 | 0.66 | 0.978 | 1 |
| 5918 | REACTOME_GLUCAGON_SIGNALING_IN_METABOI | 23 | 0.35 | 0.66 | 0.931 | 1 |
| 5919 | MIR4677_5P | 204 | 0.27 | 0.66 | 0.999 | 1 |
| 5920 | MIR1262 | 35 | 0.33 | 0.66 | 0.95 | 1 |
| 5921 | MIR3943 | 34 | 0.33 | 0.66 | 0.945 | 1 |
| 5922 | WEST_ADRENOCORTICAL_TUMOR_MARKERS_UP | 22 | 0.35 | 0.65 | 0.928 | 1 |
| 5923 | TAGCTTT_MIR9 | 211 | 0.28 | 0.65 | 0.999 | 1 |
| 5924 | MMS_MOUSE_LYMPH_HIGH_4HRS_UP | 36 | 0.33 | 0.65 | 0.958 | 1 |
| 5925 | WENDT_COHESIN_TARGETS_UP | 32 | 0.33 | 0.65 | 0.948 | 1 |
| 5926 | MIR3059_3P | 39 | 0.32 | 0.65 | 0.95 | 1 |
| 5927 | REACTOME_CALNEXIN_CALRETICULIN_CYCLE | 26 | 0.34 | 0.65 | 0.912 | 1 |
| 5928 | KEGG_CYSTEINE_AND_METHIONINE_METABOLISI | 26 | 0.34 | 0.65 | 0.934 | 1 |
| 5929 | NAGY_TFTC_COMPONENTS_HUMAN | 18 | 0.36 | 0.65 | 0.919 | 1 |
| 5930 | REACTOME_TP53_REGULATES_TRANSCRIPTION_ | 18 | 0.36 | 0.65 | 0.93 | 1 |
| 5931 | MIR1244 | 81 | 0.29 | 0.65 | 0.989 | 1 |
| 5932 | DICER1_TARGET_GENES | 57 | 0.3 | 0.65 | 0.974 | 1 |
| 5933 | MIR1323 | 279 | 0.27 | 0.65 | 1 | 1 |
| 5934 | AATGGAG_MIR136 | 71 | 0.3 | 0.65 | 0.986 | 1 |
| 5935 | DAVICIONI_RHABDOMYOSARCOMA_PAX_FOXO1_I | 55 | 0.3 | 0.65 | 0.966 | 1 |
| 5936 | MIR6753_5P | 21 | 0.35 | 0.65 | 0.931 | 1 |
| 5937 | MIR3160_3P | 54 | 0.3 | 0.65 | 0.976 | 1 |

| | | | | | | |
|---|---|---|---|---|---|---|
| 5938 | LANDIS_ERBB2_BREAST_TUMORS_324_UP | 139 | 0.28 | 0.65 | 0.991 | 1 |
| 5939 | MIR552_5P | 155 | 0.28 | 0.65 | 0.998 | 1 |
| 5940 | MIR4753_5P | 168 | 0.28 | 0.65 | 0.999 | 1 |
| 5941 | REN_BOUND_BY_E2F | 60 | 0.3 | 0.65 | 0.988 | 1 |
| 5942 | MIR6722_3P | 162 | 0.28 | 0.65 | 0.998 | 1 |
| 5943 | MIR644A | 53 | 0.3 | 0.65 | 0.972 | 1 |
| 5944 | MIR32_3P | 293 | 0.27 | 0.65 | 1 | 1 |
| 5945 | MIR7974 | 78 | 0.29 | 0.65 | 0.986 | 1 |
| 5946 | MIR4701_3P | 37 | 0.32 | 0.65 | 0.947 | 1 |
| 5947 | BIOCARTA_INSULIN_PATHWAY | 20 | 0.35 | 0.65 | 0.922 | 1 |
| 5948 | CHENG_RESPONSE_TO_NICKEL_ACETATE | 42 | 0.31 | 0.65 | 0.967 | 1 |
| 5949 | TGTATGA_MIR4853P | 141 | 0.28 | 0.65 | 0.999 | 1 |
| 5950 | KEGG_AMINOACYL_TRNA_BIOSYNTHESIS | 40 | 0.31 | 0.65 | 0.961 | 1 |
| 5951 | GAJATE_RESPONSE_TO_TRABECTEDIN_DN | 17 | 0.36 | 0.65 | 0.934 | 1 |
| 5952 | USF_01 | 241 | 0.27 | 0.65 | 1 | 1 |
| 5953 | ZNF202_TARGET_GENES | 57 | 0.3 | 0.65 | 0.973 | 1 |
| 5954 | MYBL1_TARGET_GENES | 155 | 0.28 | 0.65 | 0.999 | 1 |
| 5955 | MIR3614_3P | 52 | 0.3 | 0.65 | 0.976 | 1 |
| 5956 | MIR196A_5P_MIR196B_5P | 82 | 0.29 | 0.65 | 0.991 | 1 |
| 5957 | ELVIDGE_HIF1A_TARGETS_UP | 64 | 0.3 | 0.65 | 0.978 | 1 |
| 5958 | VANHARANTA_UTERINE_FIBROID_WITH_7Q_DELE | 35 | 0.33 | 0.64 | 0.944 | 1 |
| 5959 | MIR1271_3P | 61 | 0.3 | 0.64 | 0.985 | 1 |
| 5960 | REACTOME_DEFECTS_IN_VITAMIN_AND_COFACTO | 20 | 0.35 | 0.64 | 0.926 | 1 |
| 5961 | MITSIADES_RESPONSE_TO_APLIDIN_DN | 243 | 0.27 | 0.64 | 1 | 1 |
| 5962 | ELK1_02 | 248 | 0.27 | 0.64 | 1 | 1 |
| 5963 | MIR6772_5P | 26 | 0.33 | 0.64 | 0.938 | 1 |
| 5964 | PID_BARD1_PATHWAY | 29 | 0.33 | 0.64 | 0.961 | 1 |
| 5965 | CAFFAREL_RESPONSE_TO_THC_24HR_5_DN | 52 | 0.3 | 0.64 | 0.971 | 1 |
| 5966 | MIR3175 | 146 | 0.27 | 0.64 | 0.995 | 1 |
| 5967 | ATF1_Q6 | 181 | 0.27 | 0.64 | 0.999 | 1 |
| 5968 | MIR4695_5P | 75 | 0.29 | 0.64 | 0.988 | 1 |
| 5969 | MOOTHA_FFA_OXYDATION | 22 | 0.34 | 0.64 | 0.932 | 1 |
| 5970 | MIR5591_3P | 38 | 0.32 | 0.64 | 0.957 | 1 |

| | | | | | | |
|---|---|---|---|---|---|---|
| 5971 | MIR6814_5P | 92 | 0.28 | 0.64 | 0.996 | 1 |
| 5972 | MIR3199 | 37 | 0.32 | 0.64 | 0.972 | 1 |
| 5973 | MIR3973 | 234 | 0.27 | 0.64 | 1 | 1 |
| 5974 | PELLICCIOTTA_HDAC_IN_ANTIGEN_PRESENTATIC | 62 | 0.3 | 0.64 | 0.977 | 1 |
| 5975 | BARIS_THYROID_CANCER_UP | 24 | 0.34 | 0.64 | 0.943 | 1 |
| 5976 | ALKBH3_TARGET_GENES | 400 | 0.26 | 0.64 | 1 | 1 |
| 5977 | MIR1912_5P | 73 | 0.29 | 0.64 | 0.986 | 1 |
| 5978 | SP1_Q6 | 237 | 0.27 | 0.64 | 1 | 1 |
| 5979 | MIR8052 | 35 | 0.32 | 0.64 | 0.975 | 1 |
| 5980 | MIR6828_3P | 97 | 0.28 | 0.64 | 0.991 | 1 |
| 5981 | POMEROY_MEDULLOBLASTOMA_DESMOPLASIC_ | 53 | 0.3 | 0.64 | 0.989 | 1 |
| 5982 | MIR5582_5P | 344 | 0.26 | 0.64 | 1 | 1 |
| 5983 | REACTOME_RHOH_GTPASE_CYCLE | 35 | 0.32 | 0.63 | 0.964 | 1 |
| 5984 | MORI_IMMATURE_B_LYMPHOCYTE_DN | 86 | 0.28 | 0.63 | 0.994 | 1 |
| 5985 | CACCAGC_MIR138 | 209 | 0.26 | 0.63 | 1 | 1 |
| 5986 | OUYANG_PROSTATE_CANCER_PROGRESSION_UI | 20 | 0.34 | 0.63 | 0.948 | 1 |
| 5987 | REACTOME_VEGFR2_MEDIATED_CELL_PROLIFER | 18 | 0.35 | 0.63 | 0.942 | 1 |
| 5988 | BENPORATH_ES_2 | 27 | 0.33 | 0.63 | 0.956 | 1 |
| 5989 | WP_MTHFR_DEFICIENCY | 24 | 0.34 | 0.63 | 0.937 | 1 |
| 5990 | ZHAN_MULTIPLE_MYELOMA_CD2_DN | 40 | 0.31 | 0.63 | 0.976 | 1 |
| 5991 | MIR485_3P | 188 | 0.27 | 0.63 | 1 | 1 |
| 5992 | MIR539_3P | 188 | 0.27 | 0.63 | 0.999 | 1 |
| 5993 | WP_SPHINGOLIPID_METABOLISM_OVERVIEW | 21 | 0.34 | 0.63 | 0.944 | 1 |
| 5994 | ZFP82_TARGET_GENES | 34 | 0.32 | 0.63 | 0.972 | 1 |
| 5995 | MIR4786_5P | 78 | 0.28 | 0.63 | 0.985 | 1 |
| 5996 | WDHD1_TARGET_GENES | 170 | 0.27 | 0.63 | 0.998 | 1 |
| 5997 | REACTOME_GLUCONEOGENESIS | 26 | 0.33 | 0.63 | 0.952 | 1 |
| 5998 | MIR144_5P | 44 | 0.3 | 0.63 | 0.966 | 1 |
| 5999 | REACTOME_LYSOSOME_VESICLE_BIOGENESIS | 32 | 0.31 | 0.63 | 0.961 | 1 |
| 6000 | MIR1910_5P | 68 | 0.29 | 0.63 | 0.989 | 1 |
| 6001 | WP_MET_IN_TYPE_1_PAPILLARY_RENAL_CELL_C | 57 | 0.29 | 0.63 | 0.978 | 1 |
| 6002 | MIR649 | 61 | 0.29 | 0.63 | 0.983 | 1 |
| 6003 | ZNF85_TARGET_GENES | 281 | 0.26 | 0.63 | 1 | 1 |

| | | | | | | |
|---|---|---|---|---|---|---|
| 6004 | ZNF622_TARGET_GENES | 62 | 0.29 | 0.63 | 0.988 | 1 |
| 6005 | GAGACTG_MIR452 | 83 | 0.28 | 0.63 | 0.996 | 1 |
| 6006 | MIR1285_5P | 57 | 0.29 | 0.63 | 0.981 | 1 |
| 6007 | WALLACE_PROSTATE_CANCER_RACE_DN | 76 | 0.28 | 0.62 | 0.99 | 1 |
| 6008 | MIR302F | 155 | 0.26 | 0.62 | 0.999 | 1 |
| 6009 | MAYBURD_RESPONSE_TO_L663536_DN | 54 | 0.29 | 0.62 | 0.977 | 1 |
| 6010 | MIR20A_3P | 157 | 0.27 | 0.62 | 0.999 | 1 |
| 6011 | LASTOWSKA_NEUROBLASTOMA_COPY_NUMBER_ | 166 | 0.26 | 0.62 | 1 | 1 |
| 6012 | REACTOME_TP53_REGULATES_TRANSCRIPTION_ | 48 | 0.3 | 0.62 | 0.987 | 1 |
| 6013 | KEGG_BETA_ALANINE_METABOLISM | 18 | 0.34 | 0.62 | 0.941 | 1 |
| 6014 | REACTOME_INFLUENZA_INFECTION | 151 | 0.26 | 0.62 | 1 | 1 |
| 6015 | ZMYM2_TARGET_GENES | 114 | 0.27 | 0.62 | 0.994 | 1 |
| 6016 | GRYDER_PAX3FOXO1_TOP_ENHANCERS | 408 | 0.25 | 0.62 | 1 | 1 |
| 6017 | PID_CIRCADIAN_PATHWAY | 15 | 0.35 | 0.62 | 0.932 | 1 |
| 6018 | MIR4778_5P | 153 | 0.27 | 0.62 | 1 | 1 |
| 6019 | REACTOME_MISCELLANEOUS_TRANSPORT_AND_ | 19 | 0.34 | 0.62 | 0.943 | 1 |
| 6020 | ZHANG_BREAST_CANCER_PROGENITORS_UP | 413 | 0.25 | 0.62 | 1 | 1 |
| 6021 | GREENBAUM_E2A_TARGETS_UP | 31 | 0.31 | 0.62 | 0.961 | 1 |
| 6022 | MIR3162_5P | 64 | 0.29 | 0.62 | 0.984 | 1 |
| 6023 | RIZ_ERYTHROID_DIFFERENTIATION | 75 | 0.28 | 0.62 | 0.992 | 1 |
| 6024 | YGTCCTTGR_UNKNOWN | 82 | 0.28 | 0.61 | 0.998 | 1 |
| 6025 | ZSCAN18_TARGET_GENES | 43 | 0.3 | 0.61 | 0.977 | 1 |
| 6026 | MIR609 | 26 | 0.32 | 0.61 | 0.972 | 1 |
| 6027 | MIR4760_3P | 230 | 0.26 | 0.61 | 1 | 1 |
| 6028 | SHEDDEN_LUNG_CANCER_POOR_SURVIVAL_A6 | 431 | 0.25 | 0.61 | 1 | 1 |
| 6029 | MIR6511A_3P_MIR6511B_3P | 53 | 0.29 | 0.61 | 0.983 | 1 |
| 6030 | MIR190B_5P | 93 | 0.27 | 0.61 | 0.997 | 1 |
| 6031 | MIR6767_3P | 18 | 0.34 | 0.61 | 0.951 | 1 |
| 6032 | BOYAULT_LIVER_CANCER_SUBCLASS_G12_UP | 38 | 0.3 | 0.61 | 0.983 | 1 |
| 6033 | RICKMAN_METASTASIS_UP | 326 | 0.25 | 0.61 | 1 | 1 |
| 6034 | MIR550A_3_5P_MIR550A_5P | 63 | 0.28 | 0.61 | 0.994 | 1 |
| 6035 | WP_GENES_RELATED_TO_PRIMARY_CILIUM_DEV | 103 | 0.26 | 0.61 | 0.998 | 1 |
| 6036 | NRF2_01 | 264 | 0.25 | 0.61 | 1 | 1 |

| | | | | | | |
|---|---|---|---|---|---|---|
| 6037 | SGCGSSAAA_E2F1DP2_01 | 160 | 0.26 | 0.61 | 0.999 | 1 |
| 6038 | GRAHAM_CML_DIVIDING_VS_NORMAL_QUIESCEN | 159 | 0.26 | 0.61 | 0.999 | 1 |
| 6039 | BILD_E2F3_ONCOGENIC_SIGNATURE | 235 | 0.25 | 0.61 | 0.999 | 1 |
| 6040 | MIR617 | 56 | 0.28 | 0.61 | 0.986 | 1 |
| 6041 | MIR452_5P | 186 | 0.25 | 0.61 | 1 | 1 |
| 6042 | REACTOME_COOPERATION_OF_PDCL_PHLP1_AN | 29 | 0.31 | 0.61 | 0.97 | 1 |
| 6043 | WP_SPHINGOLIPID_METABOLISM_INTEGRATED_F | 24 | 0.32 | 0.6 | 0.956 | 1 |
| 6044 | MIR190A_5P | 94 | 0.27 | 0.6 | 0.994 | 1 |
| 6045 | MIR5681B | 47 | 0.29 | 0.6 | 0.989 | 1 |
| 6046 | TBX3_TARGET_GENES | 308 | 0.25 | 0.6 | 1 | 1 |
| 6047 | KANG_DOXORUBICIN_RESISTANCE_DN | 19 | 0.33 | 0.6 | 0.951 | 1 |
| 6048 | MIR139_5P | 135 | 0.26 | 0.6 | 0.999 | 1 |
| 6049 | REACTOME_ONCOGENE_INDUCED_SENESCENCE | 33 | 0.3 | 0.6 | 0.979 | 1 |
| 6050 | WP_GLYCEROLIPIDS_AND_GLYCEROPHOSPHOLII | 21 | 0.32 | 0.6 | 0.958 | 1 |
| 6051 | REACTOME_TRANS_GOLGI_NETWORK_VESICLE_I | 68 | 0.27 | 0.6 | 0.992 | 1 |
| 6052 | PUIFFE_INVASION_INHIBITED_BY_ASCITES_UP | 69 | 0.27 | 0.6 | 0.996 | 1 |
| 6053 | WP_SREBF_AND_MIR33_IN_CHOLESTEROL_AND_ | 15 | 0.33 | 0.6 | 0.944 | 1 |
| 6054 | KEGG_PYRUVATE_METABOLISM | 33 | 0.3 | 0.6 | 0.987 | 1 |
| 6055 | KLF14_TARGET_GENES | 227 | 0.25 | 0.59 | 1 | 1 |
| 6056 | MIR1537_5P | 107 | 0.26 | 0.59 | 0.999 | 1 |
| 6057 | PAX3_01 | 16 | 0.33 | 0.59 | 0.957 | 1 |
| 6058 | REACTOME_SIGNALING_BY_CTNNB1_PHOSPHO_S | 15 | 0.34 | 0.59 | 0.939 | 1 |
| 6059 | WHITFIELD_CELL_CYCLE_G2 | 159 | 0.25 | 0.59 | 1 | 1 |
| 6060 | PID_LIS1_PATHWAY | 27 | 0.3 | 0.59 | 0.973 | 1 |
| 6061 | FINETTI_BREAST_CANCERS_KINOME_GRAY | 15 | 0.33 | 0.59 | 0.969 | 1 |
| 6062 | MCM3_TARGET_GENES | 162 | 0.25 | 0.59 | 1 | 1 |
| 6063 | TGAGATT_MIR216 | 93 | 0.26 | 0.59 | 0.998 | 1 |
| 6064 | ZNF582_TARGET_GENES | 59 | 0.27 | 0.59 | 0.989 | 1 |
| 6065 | YGCGYRCGC_UNKNOWN | 305 | 0.24 | 0.59 | 1 | 1 |
| 6066 | WP_METABOLISM_OF_SPINGOLIPIDS_IN_ER_AND | 18 | 0.32 | 0.59 | 0.975 | 1 |
| 6067 | MIR519E_3P | 54 | 0.27 | 0.59 | 0.987 | 1 |
| 6068 | ACAWYAAAG_UNKNOWN | 80 | 0.26 | 0.59 | 0.994 | 1 |
| 6069 | SCHUHMACHER_MYC_TARGETS_UP | 79 | 0.26 | 0.59 | 0.998 | 1 |

| | | | | | | |
|---|---|---|---|---|---|---|
| 6070 | REACTOME_MRNA_DECAY_BY_5_TO_3_EXORIBO | 15 | 0.33 | 0.58 | 0.958 | 1 |
| 6071 | MIR203B_3P | 81 | 0.26 | 0.58 | 1 | 1 |
| 6072 | FAELT_B_CLL_WITH_VH_REARRANGEMENTS_DN | 45 | 0.28 | 0.58 | 0.987 | 1 |
| 6073 | KEGG_GLYCOSAMINOGLYCAN_DEGRADATION | 16 | 0.32 | 0.58 | 0.951 | 1 |
| 6074 | REACTOME_REGULATION_OF_TP53_ACTIVITY_TH | 19 | 0.32 | 0.58 | 0.96 | 1 |
| 6075 | MIR1226_3P | 86 | 0.26 | 0.58 | 0.998 | 1 |
| 6076 | MIR4724_5P | 116 | 0.25 | 0.58 | 1 | 1 |
| 6077 | MIR4764_5P | 64 | 0.27 | 0.58 | 0.998 | 1 |
| 6078 | BOYAULT_LIVER_CANCER_SUBCLASS_G23_UP | 52 | 0.27 | 0.58 | 0.996 | 1 |
| 6079 | MIR4540 | 43 | 0.28 | 0.58 | 0.987 | 1 |
| 6080 | ZNF677_TARGET_GENES | 104 | 0.26 | 0.58 | 1 | 1 |
| 6081 | MIR1180_5P | 28 | 0.3 | 0.58 | 0.982 | 1 |
| 6082 | BIOCARTA_EFP_PATHWAY | 15 | 0.33 | 0.58 | 0.95 | 1 |
| 6083 | MIR1193 | 28 | 0.3 | 0.58 | 0.975 | 1 |
| 6084 | MIR125B_2_3P | 115 | 0.25 | 0.58 | 0.999 | 1 |
| 6085 | MYC_UP.V1_UP | 158 | 0.25 | 0.58 | 1 | 1 |
| 6086 | REACTOME_TBC_RABGAPS | 41 | 0.28 | 0.58 | 0.991 | 1 |
| 6087 | MIR1227_3P | 66 | 0.26 | 0.58 | 0.996 | 1 |
| 6088 | REACTOME_SPHINGOLIPID_DE_NOVO_BIOSYNTH | 38 | 0.28 | 0.58 | 0.982 | 1 |
| 6089 | MIR33B_3P | 54 | 0.27 | 0.58 | 0.995 | 1 |
| 6090 | REACTOME_DEFECTIVE_INTRINSIC_PATHWAY_FC | 22 | 0.31 | 0.58 | 0.976 | 1 |
| 6091 | WP_OXYSTEROLS_DERIVED_FROM_CHOLESTERC | 24 | 0.3 | 0.57 | 0.956 | 1 |
| 6092 | MIR3180_MIR3180_3P | 16 | 0.32 | 0.57 | 0.969 | 1 |
| 6093 | KEGG_OXIDATIVE_PHOSPHORYLATION | 119 | 0.25 | 0.57 | 1 | 1 |
| 6094 | MIR3974 | 79 | 0.26 | 0.57 | 1 | 1 |
| 6095 | MIR2682_3P | 52 | 0.27 | 0.57 | 0.994 | 1 |
| 6096 | ZNF260_TARGET_GENES | 459 | 0.24 | 0.57 | 1 | 1 |
| 6097 | BIOCARTA_EIF_PATHWAY | 16 | 0.32 | 0.57 | 0.966 | 1 |
| 6098 | NEMETH_INFLAMMATORY_RESPONSE_LPS_DN | 25 | 0.3 | 0.57 | 0.981 | 1 |
| 6099 | MIR5003_5P | 166 | 0.24 | 0.57 | 1 | 1 |
| 6100 | REACTOME_SELENOAMINO_ACID_METABOLISM | 108 | 0.25 | 0.57 | 1 | 1 |
| 6101 | MIR10395_3P | 121 | 0.25 | 0.57 | 1 | 1 |
| 6102 | MIR6846_3P | 21 | 0.31 | 0.57 | 0.977 | 1 |

| | | | | | | |
|---|---|---|---|---|---|---|
| 6103 | BLUM_RESPONSE_TO_SALIRASIB_DN | 323 | 0.23 | 0.57 | 1 | 1 |
| 6104 | RORA_TARGET_GENES | 238 | 0.24 | 0.57 | 1 | 1 |
| 6105 | MELLMAN_TUT1_TARGETS_UP | 19 | 0.31 | 0.57 | 0.962 | 1 |
| 6106 | NIKOLSKY_BREAST_CANCER_1Q21_AMPLICON | 31 | 0.29 | 0.57 | 0.987 | 1 |
| 6107 | E2F1_UP.V1_UP | 173 | 0.24 | 0.57 | 1 | 1 |
| 6108 | REACTOME_RESOLUTION_OF_D_LOOP_STRUCTU | 33 | 0.28 | 0.57 | 0.986 | 1 |
| 6109 | RFX7_TARGET_GENES | 462 | 0.23 | 0.57 | 1 | 1 |
| 6110 | REACTOME_CONVERSION_FROM_APC_C_CDC20_ | 20 | 0.31 | 0.57 | 0.973 | 1 |
| 6111 | MIR6831_3P | 51 | 0.27 | 0.56 | 0.995 | 1 |
| 6112 | WP_NSP1_FROM_SARSCOV2_INHIBITS_TRANSLA | 17 | 0.31 | 0.56 | 0.964 | 1 |
| 6113 | MIR5088_5P | 33 | 0.28 | 0.56 | 0.991 | 1 |
| 6114 | MISSIAGLIA_REGULATED_BY_METHYLATION_DN | 118 | 0.24 | 0.56 | 1 | 1 |
| 6115 | MIR1260A_MIR1260B | 91 | 0.25 | 0.56 | 1 | 1 |
| 6116 | PID_MTOR_4PATHWAY | 67 | 0.26 | 0.56 | 0.996 | 1 |
| 6117 | MOREAUX_MULTIPLE_MYELOMA_BY_TACI_DN | 175 | 0.24 | 0.56 | 1 | 1 |
| 6118 | ACTGTAG_MIR139 | 112 | 0.24 | 0.56 | 1 | 1 |
| 6119 | MIR370_5P | 71 | 0.25 | 0.56 | 0.999 | 1 |
| 6120 | LIN_MELANOMA_COPY_NUMBER_DN | 36 | 0.27 | 0.55 | 0.992 | 1 |
| 6121 | WP_TRANSLATION_FACTORS | 53 | 0.26 | 0.55 | 0.997 | 1 |
| 6122 | ZNF250_TARGET_GENES | 459 | 0.23 | 0.55 | 1 | 1 |
| 6123 | ACTWSNACTNY_UNKNOWN | 92 | 0.25 | 0.55 | 1 | 1 |
| 6124 | WEST_ADRENOCORTICAL_TUMOR_UP | 285 | 0.23 | 0.55 | 1 | 1 |
| 6125 | KAUFFMANN_DNA_REPAIR_GENES | 225 | 0.23 | 0.55 | 1 | 1 |
| 6126 | DEN_INTERACT_WITH_LCA5 | 26 | 0.28 | 0.55 | 0.982 | 1 |
| 6127 | GLASS_IGF2BP1_CLIP_TARGETS_KNOCKDOWN_[ | 116 | 0.24 | 0.55 | 0.999 | 1 |
| 6128 | AUNG_GASTRIC_CANCER | 39 | 0.27 | 0.55 | 0.997 | 1 |
| 6129 | GCGNNANTTCC_UNKNOWN | 123 | 0.23 | 0.54 | 1 | 1 |
| 6130 | CHAF1A_TARGET_GENES | 86 | 0.24 | 0.54 | 1 | 1 |
| 6131 | COLDREN_GEFITINIB_RESISTANCE_UP | 83 | 0.24 | 0.54 | 1 | 1 |
| 6132 | REACTOME_SIGNALLING_TO_RAS | 19 | 0.29 | 0.54 | 0.976 | 1 |
| 6133 | RUVBL2_TARGET_GENES | 35 | 0.27 | 0.54 | 0.992 | 1 |
| 6134 | REACTOME_INTRAFLAGELLAR_TRANSPORT | 48 | 0.26 | 0.54 | 0.996 | 1 |
| 6135 | PUJANA_XPRSS_INT_NETWORK | 165 | 0.23 | 0.53 | 1 | 1 |

| | | | | | | |
|---|---|---|---|---|---|---|
| 6136 | MIR6781_3P | 67 | 0.24 | 0.53 | 1 | 1 |
| 6137 | REACTOME_RAF_ACTIVATION | 33 | 0.27 | 0.53 | 0.995 | 1 |
| 6138 | REACTOME_TP53_REGULATES_TRANSCRIPTION_ | 61 | 0.25 | 0.53 | 0.998 | 1 |
| 6139 | ZMYND11_TARGET_GENES | 42 | 0.25 | 0.53 | 0.996 | 1 |
| 6140 | REACTOME_TRNA_PROCESSING | 110 | 0.23 | 0.52 | 1 | 1 |
| 6141 | REACTOME_PINK1_PRKN_MEDIATED_MITOPHAGY | 21 | 0.28 | 0.52 | 0.989 | 1 |
| 6142 | REACTOME_HIV_INFECTION | 223 | 0.22 | 0.52 | 1 | 1 |
| 6143 | MIR3680_5P | 25 | 0.27 | 0.52 | 0.987 | 1 |
| 6144 | ZSCAN23_TARGET_GENES | 73 | 0.24 | 0.52 | 1 | 1 |
| 6145 | SRPK1_TARGET_GENES | 298 | 0.21 | 0.52 | 1 | 1 |
| 6146 | REACTOME_VASOPRESSIN_REGULATES_RENAL_ | 29 | 0.27 | 0.52 | 0.992 | 1 |
| 6147 | YUAN_ZNF143_PARTNERS | 21 | 0.28 | 0.52 | 0.988 | 1 |
| 6148 | SETBP1_TARGET_GENES | 172 | 0.22 | 0.52 | 1 | 1 |
| 6149 | HORIUCHI_WTAP_TARGETS_DN | 300 | 0.21 | 0.52 | 1 | 1 |
| 6150 | OISHI_CHOLANGIOMA_STEM_CELL_LIKE_UP | 319 | 0.21 | 0.51 | 1 | 1 |
| 6151 | E2F4DP1_01 | 229 | 0.21 | 0.51 | 1 | 1 |
| 6152 | HOLLEMAN_VINCRISTINE_RESISTANCE_B_ALL_DI | 15 | 0.29 | 0.51 | 0.979 | 1 |
| 6153 | MIR376A_5P | 43 | 0.24 | 0.51 | 0.996 | 1 |
| 6154 | WELCH_GATA1_TARGETS | 15 | 0.29 | 0.51 | 0.983 | 1 |
| 6155 | GGAANCGGAANY_UNKNOWN | 110 | 0.22 | 0.51 | 1 | 1 |
| 6156 | KEGG_GLYCOLYSIS_GLUCONEOGENESIS | 40 | 0.25 | 0.51 | 0.999 | 1 |
| 6157 | REACTOME_COPII_MEDIATED_VESICLE_TRANSPC | 63 | 0.23 | 0.5 | 1 | 1 |
| 6158 | ACACTAC_MIR1423P | 122 | 0.22 | 0.5 | 1 | 1 |
| 6159 | E2F_02 | 225 | 0.21 | 0.5 | 1 | 1 |
| 6160 | ZNF830_TARGET_GENES | 259 | 0.21 | 0.5 | 1 | 1 |
| 6161 | ACTAYRNNNCCCR_UNKNOWN | 442 | 0.2 | 0.5 | 1 | 1 |
| 6162 | REACTOME_RHO_GTPASES_ACTIVATE_ROCKS | 17 | 0.27 | 0.49 | 0.986 | 1 |
| 6163 | REACTOME_SUMOYLATION_OF_DNA_METHYLATIC | 16 | 0.27 | 0.49 | 0.988 | 1 |
| 6164 | WP_AMINO_ACID_METABOLISM | 69 | 0.22 | 0.49 | 1 | 1 |
| 6165 | EPC1_TARGET_GENES | 199 | 0.2 | 0.49 | 1 | 1 |
| 6166 | PID_P53_REGULATION_PATHWAY | 59 | 0.23 | 0.49 | 1 | 1 |
| 6167 | MANALO_HYPOXIA_DN | 286 | 0.2 | 0.49 | 1 | 1 |
| 6168 | KEGG_VIBRIO_CHOLERAE_INFECTION | 46 | 0.23 | 0.49 | 0.999 | 1 |

| | | | | | | |
|---|---|---|---|---|---|---|
| 6169 | BCAT_GDS748_DN | 37 | 0.24 | 0.48 | 0.999 | 1 |
| 6170 | REACTOME_SYNTHESIS_OF_PA | 24 | 0.26 | 0.48 | 0.99 | 1 |
| 6171 | WP_NUCLEOTIDE_METABOLISM | 19 | 0.26 | 0.48 | 0.993 | 1 |
| 6172 | CHANDRAN_METASTASIS_TOP50_UP | 34 | 0.24 | 0.48 | 0.999 | 1 |
| 6173 | WANG_RESPONSE_TO_ANDROGEN_UP | 26 | 0.25 | 0.48 | 0.996 | 1 |
| 6174 | NIKOLSKY_BREAST_CANCER_6P24_P22_AMPLIC( | 21 | 0.26 | 0.48 | 0.991 | 1 |
| 6175 | REACTOME_CLASS_I_PEROXISOMAL_MEMBRANE | 19 | 0.26 | 0.47 | 0.996 | 1 |
| 6176 | MIR4999_3P | 31 | 0.24 | 0.47 | 0.999 | 1 |
| 6177 | CHIANG_LIVER_CANCER_SUBCLASS_UNANNOTAT | 186 | 0.2 | 0.47 | 1 | 1 |
| 6178 | WP_DNA_IRDOUBLE_STRAND_BREAKS_AND_CELI | 54 | 0.22 | 0.47 | 0.999 | 1 |
| 6179 | TIAN_BHLHA15_TARGETS | 16 | 0.27 | 0.47 | 0.986 | 1 |
| 6180 | E2F4DP2_01 | 225 | 0.2 | 0.47 | 1 | 1 |
| 6181 | E2F1DP1_01 | 225 | 0.2 | 0.47 | 1 | 1 |
| 6182 | PID_ATM_PATHWAY | 34 | 0.24 | 0.47 | 0.999 | 1 |
| 6183 | E2F1DP2_01 | 225 | 0.2 | 0.47 | 1 | 1 |
| 6184 | REACTOME_SIGNALLING_TO_ERKS | 32 | 0.23 | 0.47 | 1 | 1 |
| 6185 | REACTOME_RHO_GTPASES_ACTIVATE_CIT | 17 | 0.26 | 0.46 | 0.991 | 1 |
| 6186 | ACOSTA_PROLIFERATION_INDEPENDENT_MYC_T/ | 81 | 0.21 | 0.46 | 1 | 1 |
| 6187 | MIR4512 | 20 | 0.25 | 0.46 | 0.991 | 1 |
| 6188 | REACTOME_NONSENSE_MEDIATED_DECAY_NMD | 111 | 0.2 | 0.46 | 1 | 1 |
| 6189 | WP_AUTOPHAGY | 28 | 0.23 | 0.46 | 0.999 | 1 |
| 6190 | PITX1_TARGET_GENES | 316 | 0.19 | 0.46 | 1 | 1 |
| 6191 | BIOCARTA_VDR_PATHWAY | 24 | 0.24 | 0.45 | 0.997 | 1 |
| 6192 | REACTOME_DEADENYLATION_OF_MRNA | 25 | 0.24 | 0.45 | 0.997 | 1 |
| 6193 | REACTOME_RESPONSE_OF_EIF2AK4_GCN2_TO_/ | 97 | 0.2 | 0.45 | 1 | 1 |
| 6194 | WANG_METASTASIS_OF_BREAST_CANCER_ESR1_ | 22 | 0.24 | 0.45 | 0.997 | 1 |
| 6195 | WP_CILIARY_LANDSCAPE | 211 | 0.19 | 0.45 | 1 | 1 |
| 6196 | BARRIER_COLON_CANCER_RECURRENCE_DN | 17 | 0.25 | 0.45 | 0.992 | 1 |
| 6197 | REACTOME_ACTIVATION_OF_BAD_AND_TRANSLC | 15 | 0.25 | 0.44 | 0.993 | 1 |
| 6198 | REACTOME_SUMOYLATION_OF_TRANSCRIPTION_ | 41 | 0.22 | 0.44 | 0.999 | 1 |
| 6199 | BURTON_ADIPOGENESIS_12 | 32 | 0.22 | 0.44 | 1 | 1 |
| 6200 | WAKASUGI_HAVE_ZNF143_BINDING_SITES | 54 | 0.2 | 0.43 | 1 | 1 |
| 6201 | REACTOME_FORMATION_OF_TC_NER_PRE_INCISI | 53 | 0.2 | 0.43 | 1 | 1 |

| | | | | | | |
|---|---|---|---|---|---|---|
| 6202 | KEGG_RNA_DEGRADATION | 56 | 0.2 | 0.42 | 1 | 1 |
| 6203 | WP_TCA_CYCLE_AKA_KREBS_OR_CITRIC_ACID_( | 18 | 0.23 | 0.42 | 0.999 | 1 |
| 6204 | MIR4690_5P | 65 | 0.19 | 0.42 | 1 | 1 |
| 6205 | REACTOME_SYNTHESIS_OF_ACTIVE_UBIQUITIN_I | 30 | 0.21 | 0.42 | 0.999 | 1 |
| 6206 | WP_MRNA_PROCESSING | 129 | 0.18 | 0.42 | 1 | 1 |
| 6207 | GOLUB_ALL_VS_AML_UP | 22 | 0.22 | 0.42 | 0.999 | 1 |
| 6208 | MOREAUX_B_LYMPHOCYTE_MATURATION_BY_TA | 76 | 0.19 | 0.41 | 1 | 1 |
| 6209 | REACTOME_MITOCHONDRIAL_TRANSLATION | 96 | 0.18 | 0.41 | 1 | 1 |
| 6210 | REACTOME_ENERGY_DEPENDENT_REGULATION_ | 27 | 0.21 | 0.41 | 0.999 | 1 |
| 6211 | REACTOME_TRANSLATION | 289 | 0.17 | 0.41 | 1 | 1 |
| 6212 | KEGG_PENTOSE_PHOSPHATE_PATHWAY | 20 | 0.22 | 0.41 | 0.999 | 1 |
| 6213 | REACTOME_MTOR_SIGNALLING | 39 | 0.2 | 0.4 | 0.999 | 1 |
| 6214 | REACTOME_HOMOLOGOUS_DNA_PAIRING_AND_S | 42 | 0.2 | 0.4 | 1 | 1 |
| 6215 | REACTOME_REGULATION_OF_PLK1_ACTIVITY_AT | 85 | 0.18 | 0.4 | 1 | 1 |
| 6216 | REACTOME_DISEASES_OF_DNA_REPAIR | 33 | 0.21 | 0.4 | 1 | 1 |
| 6217 | LUI_THYROID_CANCER_CLUSTER_3 | 28 | 0.21 | 0.4 | 0.999 | 1 |
| 6218 | REACTOME_HIV_TRANSCRIPTION_INITIATION | 46 | 0.19 | 0.4 | 1 | 1 |
| 6219 | FINETTI_BREAST_CANCER_KINOME_RED | 16 | 0.22 | 0.4 | 0.997 | 1 |
| 6220 | RAMPON_ENRICHED_LEARNING_ENVIRONMENT_L | 20 | 0.22 | 0.4 | 0.996 | 1 |
| 6221 | WAKABAYASHI_ADIPOGENESIS_PPARG_BOUND_ | 29 | 0.21 | 0.4 | 1 | 1 |
| 6222 | REACTOME_HDR_THROUGH_SINGLE_STRAND_AN | 37 | 0.2 | 0.4 | 1 | 1 |
| 6223 | REACTOME_RESPIRATORY_ELECTRON_TRANSPC | 103 | 0.18 | 0.4 | 1 | 1 |
| 6224 | BORCZUK_MALIGNANT_MESOTHELIOMA_UP | 299 | 0.16 | 0.4 | 1 | 1 |
| 6225 | REACTOME_ERK_MAPK_TARGETS | 21 | 0.21 | 0.39 | 1 | 1 |
| 6226 | HU_ANGIOGENESIS_DN | 34 | 0.19 | 0.39 | 1 | 1 |
| 6227 | BIOCARTA_PYK2_PATHWAY | 27 | 0.2 | 0.38 | 0.999 | 1 |
| 6228 | REACTOME_RESOLUTION_OF_D_LOOP_STRUCTU | 26 | 0.2 | 0.38 | 1 | 1 |
| 6229 | PYEON_CANCER_HEAD_AND_NECK_VS_CERVICA | 184 | 0.16 | 0.37 | 1 | 1 |
| 6230 | PID_IGF1_PATHWAY | 27 | 0.19 | 0.37 | 1 | 1 |
| 6231 | ABDELMOHSEN_ELAVL4_TARGETS | 15 | 0.21 | 0.37 | 0.998 | 1 |
| 6232 | REACTOME_FORMATION_OF_INCISION_COMPLEX | 43 | 0.18 | 0.37 | 1 | 1 |
| 6233 | MIR8086 | 19 | 0.2 | 0.36 | 1 | 1 |
| 6234 | REACTOME_CITRIC_ACID_CYCLE_TCA_CYCLE | 22 | 0.19 | 0.36 | 0.999 | 1 |

| | | | | | | |
|---|---|---|---|---|---|---|
| 6235 | REACTOME_TRNA_PROCESSING_IN_THE_NUCLEU | 59 | 0.17 | 0.36 | 1 | 1 |
| 6236 | REACTOME_ANCHORING_OF_THE_BASAL_BODY_ | 94 | 0.16 | 0.36 | 1 | 1 |
| 6237 | FOXR2_TARGET_GENES | 212 | 0.15 | 0.35 | 1 | 1 |
| 6238 | FARMER_BREAST_CANCER_CLUSTER_5 | 19 | 0.19 | 0.35 | 1 | 1 |
| 6239 | REACTOME_SUMOYLATION_OF_RNA_BINDING_PF | 47 | 0.17 | 0.35 | 1 | 1 |
| 6240 | REACTOME_FORMATION_OF_RNA_POL_II_ELONG | 57 | 0.16 | 0.34 | 1 | 1 |
| 6241 | DING_LUNG_CANCER_EXPRESSION_BY_COPY_NU | 98 | 0.15 | 0.34 | 1 | 1 |
| 6242 | MIR4486 | 28 | 0.18 | 0.34 | 0.999 | 1 |
| 6243 | KEGG_BASE_EXCISION_REPAIR | 33 | 0.17 | 0.34 | 1 | 1 |
| 6244 | NUFIP1_TARGET_GENES | 36 | 0.17 | 0.34 | 1 | 1 |
| 6245 | REACTOME_SUMOYLATION_OF_UBIQUITINYLATIO | 39 | 0.17 | 0.34 | 1 | 1 |
| 6246 | REACTOME_KSRP_KHSRP_BINDS_AND_DESTABIL | 17 | 0.19 | 0.33 | 1 | 1 |
| 6247 | REACTOME_AMINO_ACIDS_REGULATE_MTORC1 | 52 | 0.15 | 0.31 | 1 | 1 |
| 6248 | REACTOME_SEPARATION_OF_SISTER_CHROMATI | 182 | 0.13 | 0.31 | 1 | 1 |
| 6249 | KEGG_CITRATE_CYCLE_TCA_CYCLE | 28 | 0.16 | 0.31 | 1 | 1 |
| 6250 | REACTOME_CELLULAR_RESPONSE_TO_STARVAT | 149 | 0.13 | 0.3 | 1 | 1 |
| 6251 | BIOCARTA_MTOR_PATHWAY | 21 | 0.16 | 0.3 | 1 | 1 |
