## Supplementary dataset 3 for "Polyomavirus ALTOs, but not MTs, downregulate viral early gene expression by activating the NF-κB pathway"

| Gene Symbol | fold change (TID-ALTO/TID) | P-value | log2FC | -log10(P) |
|---|---|---|---|---|
| SH3D19 | 2710.416904 | 0.00342726 | 11.4042991 | 2.46505282 |
| NEO1 | 1890.448668 | 0.00417611 | 10.884513 | 2.37922832 |
| IFIT1 | 960.3714789 | 4.4937E-08 | 9.90744875 | 7.34739121 |
| CDSN | 932.6060952 | 0.01375733 | 9.86512405 | 1.86146574 |
| KCNG3 | 919.740628 | 0.00296068 | 9.84508326 | 2.52860792 |
| PARP12 | 579.6442956 | 5.7408E-07 | 9.17902404 | 6.24102396 |
| AMER1 | 561.7118741 | 5.5052E-07 | 9.13368649 | 6.25922499 |
| AK9 | 532.7603287 | 0.00352048 | 9.05734285 | 2.4533977 |
| NCCRP1 | 529.6844278 | 0.01130681 | 9.04898929 | 1.94665988 |
| KLHL22 | 357.6462051 | 2.2245E-05 | 8.48238932 | 4.65277587 |
| TNFAIP1 | 255.3124321 | 2.4123E-05 | 7.99611998 | 4.61757335 |
| KCTD13 | 245.5565966 | 1.4581E-05 | 7.93991177 | 4.83622067 |
| NDUFAB1 | 233.7439037 | 0.01159316 | 7.86878493 | 1.93579815 |
| LOC122526780 | 224.0702452 | 0.00037974 | 7.80780727 | 3.42050849 |
| CECR2 | 200.1397883 | 0.00020544 | 7.6448642 | 3.68731684 |
| KLHL12 | 198.1090428 | 0.00202311 | 7.63015092 | 2.69398068 |
| ZDHHC8 | 184.1325797 | 0.00253343 | 7.5246011 | 2.59629165 |
| LETMD1 | 137.181306 | 5.8567E-05 | 7.09994009 | 4.23234757 |
| SLC4A7 | 133.349464 | 0.00550983 | 7.05906822 | 2.258862 |
| COL1A2 | 131.1603621 | 0.16784028 | 7.03518798 | 0.7751038 |
| PAK5 | 126.9467241 | 0.00131858 | 6.98807936 | 2.87989463 |
| TCHP | 121.4288045 | 0.0003629 | 6.92396688 | 3.44021099 |
| CPEB3 | 119.5672863 | 1.862E-05 | 6.90167891 | 4.73003197 |
| PLCG1 | 112.1935559 | 1.2795E-05 | 6.809846 | 4.89297622 |
| TMEM263 | 103.4799742 | 1.1113E-05 | 6.69320779 | 4.95415889 |
| FAU | 101.7005191 | 3.626E-07 | 6.66818323 | 6.44057505 |
| MINAR1 | 98.69169973 | 0.01173873 | 6.62485685 | 1.93037876 |
| **TRAF3** | 89.79446164 | 2.3202E-05 | 6.48855456 | 4.63447434 |
| ROBO2 | 88.04655274 | 2.3533E-05 | 6.46019461 | 4.62831956 |
| LGALS3BP | 82.95335549 | 4.4503E-06 | 6.37422843 | 5.35161342 |
| AZGP1 | 78.88575444 | 0.00010292 | 6.30169289 | 3.98749865 |
| PLCG2 | 78.44223011 | 0.00022814 | 6.29355865 | 3.64179462 |
| SH3BP2 | 77.24796892 | 0.15818027 | 6.2714251 | 0.80084768 |
| AHCYL2 | 75.32099158 | 2.812E-06 | 6.23498009 | 5.55098654 |
| GLMN | 71.61839959 | 4.5816E-05 | 6.16225838 | 4.33898265 |
| USP6NL | 69.80860178 | 1.0489E-05 | 6.12533291 | 4.97928092 |
| MB21D2 | 68.59946036 | 8.6402E-07 | 6.10012532 | 6.06347375 |
| MAPK6 | 65.47818701 | 0.00206722 | 6.03294247 | 2.6846143 |
| SLC4A7 | 62.86460269 | 7.0106E-05 | 5.974176 | 4.1542433 |
| RIOX2 | 61.82060045 | 2.9403E-05 | 5.95001576 | 4.53160195 |
| SLC12A6 | 58.26325116 | 1.5016E-06 | 5.8645143 | 5.82343444 |
| CNTRL | 57.73311853 | 0.00657873 | 5.85132725 | 2.18185816 |
| SRCIN1 | 56.08621051 | 1.7899E-05 | 5.8095742 | 4.74716369 |
| CPEB4 | 55.13471745 | 4.6714E-06 | 5.78488914 | 5.33055338 |

| | | | | |
|---|---|---|---|---|
| SCAMP1 | 44.56352114 | 2.6844E-07 | 5.47779133 | 6.57115583 |
| RSAD2 | 43.50027122 | 0.01811354 | 5.44295249 | 1.74199673 |
| DLL3 | 42.0173408 | 0.26073881 | 5.39291295 | 0.58379432 |
| HSD17B13 | 41.69376708 | 3.8665E-05 | 5.38175982 | 4.41267968 |
| SLC35A2 | 41.196263 | 0.0003016 | 5.36444157 | 3.52056414 |
| S100A9 | 41.04134865 | 0.0348012 | 5.35900623 | 1.45840577 |
| FNBP1L | 40.97383419 | 7.436E-07 | 5.356631 | 6.12866259 |
| ITSN1 | 40.28901676 | 1.1456E-05 | 5.33231469 | 4.94095733 |
| SYNRG | 40.11519076 | 1.0067E-05 | 5.32607675 | 4.99711586 |
| USP54 | 39.92391803 | 6.5599E-06 | 5.3191814 | 5.1831036 |
| MRPL24 | 39.59373456 | 1.8326E-05 | 5.30720025 | 4.73693917 |
| MEX3A | 39.51577643 | 1.6508E-06 | 5.30435685 | 5.78231513 |
| MRPL19 | 37.56791385 | 0.00047711 | 5.2314291 | 3.32138582 |
| SEMG2 | 36.16117853 | 0.02862456 | 5.17636979 | 1.54326123 |
| PDZD11 | 36.15677882 | 8.4112E-06 | 5.17619425 | 5.07514451 |
| ADGRL1 | 35.69862123 | 0.00088063 | 5.15779645 | 3.05520597 |
| RAB11FIP2 | 35.45855438 | 1.975E-05 | 5.14806181 | 4.70442488 |
| HRNR | 34.87910013 | 0.00018628 | 5.12429091 | 3.72983454 |
| DCD | 34.75752837 | 5.9655E-05 | 5.11925359 | 4.22435053 |
| **SQSTM1** | 34.51373918 | 3.1017E-06 | 5.10909888 | 5.50839346 |
| KHNYN | 34.29756954 | 0.00024142 | 5.10003444 | 3.61722269 |
| ZC3HAV1 | 33.93347203 | 2.2839E-06 | 5.08463715 | 5.64132712 |
| AHCYL1 | 33.63750773 | 0.00020811 | 5.07199891 | 3.68171235 |
| ZC3H12C | 33.16852356 | 6.1409E-05 | 5.05174289 | 4.21176486 |
| ARG1 | 32.75479772 | 0.00022198 | 5.03363433 | 3.6536809 |
| TNRC6B | 32.5377833 | 0.00022159 | 5.02404406 | 3.65444925 |
| SLC38A1 | 32.48950514 | 0.00033782 | 5.02190186 | 3.47131824 |
| CTTN | 32.39781002 | 9.7018E-06 | 5.01782439 | 5.01314749 |
| TMT1A | 31.27334815 | 1.1464E-05 | 4.96686178 | 4.94065841 |
| RIOX1 | 30.70005206 | 2.3153E-05 | 4.9401692 | 4.6354004 |
| SCYL3 | 30.18036483 | 3.3123E-05 | 4.91553834 | 4.47986497 |
| ZDHHC5 | 30.00444577 | 6.6529E-06 | 4.90710438 | 5.17699186 |
| CLINT1 | 29.71327676 | 6.9598E-07 | 4.89303581 | 6.15740618 |
| KRT6A | 29.68119879 | 0.00376741 | 4.89147746 | 2.423957 |
| GAB1 | 29.51484707 | 1.8248E-05 | 4.88336896 | 4.73878494 |
| SLC30A6 | 28.96704828 | 6.0414E-05 | 4.85634078 | 4.21886404 |
| SORBS2 | 28.19284841 | 0.00093178 | 4.81725734 | 3.03068693 |
| DSG1 | 28.11842347 | 4.879E-05 | 4.8134438 | 4.31166896 |
| ENTR1 | 27.7716172 | 2.4203E-05 | 4.79553929 | 4.61613283 |
| BEGAIN | 27.61839364 | 1.5525E-05 | 4.78755751 | 4.80895545 |
| CHD8 | 26.99798668 | 8.5505E-09 | 4.75477992 | 8.0680095 |
| EPS15L1 | 26.73584414 | 3.6339E-05 | 4.74070332 | 4.43962411 |
| **TRAF2** | 26.68678265 | 6.6625E-06 | 4.73805348 | 5.17636267 |
| EFNB3 | 26.44447031 | 2.4853E-05 | 4.72489417 | 4.6046283 |
| TMEM51 | 26.34918831 | 4.2644E-06 | 4.71968661 | 5.3701402 |

| | | | | |
|---|---|---|---|---|
| GPRC5B | 26.32060253 | 2.8664E-05 | 4.71812061 | 4.5426704 |
| EPS15L1 | 26.27383121 | 9.8953E-06 | 4.71555468 | 5.00457124 |
| KPRP | 25.88456117 | 6.7413E-05 | 4.69401995 | 4.17125912 |
| ZFYVE16 | 25.65656441 | 3.3431E-06 | 4.68125609 | 5.47585268 |
| SCCPDH | 25.3228525 | 0.02581713 | 4.66236802 | 1.58809196 |
| DPH7 | 25.2577805 | 0.07909459 | 4.65865596 | 1.10185323 |
| TGM3 | 25.09674755 | 0.03437734 | 4.6494285 | 1.46372772 |
| SYNGR3 | 25.06432598 | 0.00032304 | 4.64756353 | 3.49074988 |
| CYB5R3 | 24.87668626 | 3.9885E-05 | 4.63672242 | 4.39918516 |
| HSPE1 | 24.75450804 | 4.5747E-06 | 4.62961937 | 5.33963832 |
| NCBP2AS2 | 24.60284037 | 0.0001463 | 4.62075298 | 3.83476162 |
| SBSN | 24.57669785 | 0.00010088 | 4.61921918 | 3.99619685 |
| KRT77 | 24.56412946 | 2.2989E-05 | 4.61848121 | 4.63847245 |
| KLHL9 | 24.45611963 | 0.00447737 | 4.61212361 | 2.34897671 |
| AGFG1 | 24.20936262 | 3.7986E-06 | 4.59749319 | 5.42037942 |
| RAB9A | 24.04072033 | 0.6636458 | 4.58740822 | 0.17806365 |
| KRT25 | 23.955308 | 0.00204294 | 4.58227346 | 2.68974367 |
| RC3H1 | 23.80345047 | 7.5413E-05 | 4.57309881 | 4.12255628 |
| ADISSP | 23.79745053 | 0.04137356 | 4.57273512 | 1.38327716 |
| YTHDF2 | 23.75688245 | 1.2362E-05 | 4.57027362 | 4.90789394 |
| CNNM3 | 23.64714164 | 2.5498E-05 | 4.5635939 | 4.59348999 |
| BIRC2 | 23.62359393 | 0.00010848 | 4.56215656 | 3.96465836 |
| EFNB1 | 23.54980277 | 7.1009E-05 | 4.55764307 | 4.1486839 |
| RAB8A | 23.48659889 | 0.00684145 | 4.55376591 | 2.16485175 |
| AHNAK | 23.36321276 | 1.0131E-05 | 4.54616677 | 4.99432974 |
| SHB | 22.99327121 | 8.2041E-06 | 4.52313982 | 5.08597049 |
| LRIG1 | 22.73496461 | 4.2121E-06 | 4.50684085 | 5.37549689 |
| CYLD | 22.66687413 | 1.8113E-05 | 4.50251355 | 4.7420045 |
| PLEKHA6 | 22.6590157 | 2.6072E-05 | 4.50201329 | 4.58381768 |
| SURF6 | 22.28058116 | 1.7159E-06 | 4.47771496 | 5.76550979 |
| PRKCG | 22.13584216 | 0.00055178 | 4.46831236 | 3.25823672 |
| TROAP | 22.10928155 | 0.00265986 | 4.46658024 | 2.575141 |
| PRKD2 | 22.00142152 | 0.00040631 | 4.45952483 | 3.39113944 |
| SAV1 | 21.61786596 | 0.00017084 | 4.43415221 | 3.76739789 |
| GJC1 | 21.56434825 | 5.2673E-05 | 4.43057621 | 4.27841485 |
| AGO2 | 21.5439078 | 1.1694E-05 | 4.42920806 | 4.93203255 |
| ZBTB14 | 21.37005914 | 0.0082495 | 4.417519 | 2.08357245 |
| ITSN2 | 21.10948994 | 1.785E-05 | 4.39981981 | 4.74836428 |
| AGO3 | 20.96553901 | 2.4255E-05 | 4.38994802 | 4.61519252 |
| CS | 20.79708864 | 3.5901E-05 | 4.37830968 | 4.44489696 |
| EBAG9 | 20.71086205 | 4.8538E-05 | 4.3723157 | 4.31391597 |
| NEBL | 20.50278577 | 6.2809E-05 | 4.35774804 | 4.20198045 |
| SPATA2L | 20.47776459 | 7.7917E-06 | 4.35598633 | 5.10836899 |
| ETFA | 20.47729268 | 0.0006064 | 4.35595308 | 3.21724067 |
| PPP1R21 | 20.45313984 | 0.00015327 | 4.35425043 | 3.81454954 |

| | | | | |
|---|---|---|---|---|
| SYT7 | 20.34480615 | 0.00015434 | 4.34658863 | 3.81152855 |
| KRT16 | 20.2133996 | 0.00354342 | 4.33724008 | 2.45057731 |
| ZCCHC17 | 20.1813319 | 8.2485E-05 | 4.33494949 | 4.08362263 |
| TNRC6B | 20.02310372 | 6.8551E-06 | 4.32359371 | 5.16398533 |
| CALML5 | 19.76115472 | 0.0011407 | 4.30459535 | 2.94282667 |
| STARD3 | 19.71419238 | 0.00039099 | 4.3011627 | 3.40783654 |
| RC3H2 | 19.36763277 | 6.3834E-06 | 4.27557572 | 5.19494895 |
| ALMS1 | 19.30360127 | 2.2382E-05 | 4.27079812 | 4.65009759 |
| ACADM | 18.70400189 | 0.00029987 | 4.22527508 | 3.52306134 |
| TNRC6A | 18.52075047 | 2.5496E-06 | 4.21107065 | 5.5935302 |
| ROBO1 | 18.4249199 | 7.5537E-05 | 4.20358644 | 4.12183803 |
| MACROD1 | 18.28947278 | 0.00171844 | 4.19294158 | 2.76486558 |
| SMAP1 | 18.18821825 | 1.5823E-05 | 4.18493232 | 4.80071512 |
| KRT9 | 18.1635358 | 0.00042093 | 4.18297317 | 3.37578678 |
| ERBIN | 17.97457628 | 2.6165E-06 | 4.16788586 | 5.58227629 |
| KRT10 | 17.94578422 | 1.0057E-05 | 4.16557306 | 4.99753494 |
| SMAP2 | 17.89834979 | 4.3838E-06 | 4.16175467 | 5.35815394 |
| AGO1 | 17.79844766 | 1.3601E-05 | 4.15367951 | 4.86643792 |
| ALTO | 17.73403619 | 1.5876E-05 | 4.14844902 | 4.79924791 |
| AZI2 | 17.65137053 | 0.00121119 | 4.1417083 | 2.91678857 |
| NCK2 | 17.64134355 | 8.4659E-06 | 4.14088853 | 5.07232751 |
| RPL7 | 17.52438936 | 1.7247E-05 | 4.13129227 | 4.7632792 |
| KRT1 | 17.4783736 | 0.00012225 | 4.12749904 | 3.91274222 |
| APBB1 | 17.00450176 | 2.1837E-05 | 4.08784483 | 4.66079961 |
| PDLIM3 | 16.93093506 | 0.00040541 | 4.08158975 | 3.39210223 |
| KIAA1217 | 16.83620605 | 3.956E-06 | 4.07349517 | 5.40274505 |
| MYO1C | 16.8232703 | 0.00014257 | 4.07238628 | 3.84597028 |
| ELP5 | 16.63674064 | 0.00066769 | 4.05630091 | 3.17542423 |
| EPS15 | 16.58586305 | 2.3314E-05 | 4.05188218 | 4.63239073 |
| TRIM37 | 16.49218566 | 0.09525675 | 4.0437107 | 1.02110424 |
| FRS2 | 16.4657583 | 0.07494631 | 4.04139705 | 1.12524975 |
| ODF2 | 16.44253545 | 3.228E-05 | 4.03936088 | 4.49106913 |
| ARGLU1 | 16.42138782 | 0.00026406 | 4.03750415 | 3.57830126 |
| FAM171B | 16.32606099 | 0.00012321 | 4.02910485 | 3.90934012 |
| TMEM74 | 16.222725 | 0.00012202 | 4.01994427 | 3.91355759 |
| ARHGAP32 | 16.21364748 | 1.6906E-05 | 4.01913678 | 4.77197182 |
| MCAM | 16.13616661 | 0.00054179 | 4.01222598 | 3.26616878 |
| RSPRY1 | 16.09486344 | 0.00554856 | 4.00852843 | 2.25581942 |
| YTHDF3 | 15.94509005 | 0.00010649 | 3.99504034 | 3.97269223 |
| MRPL11 | 15.84958101 | 0.00097026 | 3.9863728 | 3.01311121 |
| CCDC15 | 15.79028918 | 0.00010018 | 3.98096569 | 3.99922509 |
| ALDH6A1 | 15.7590992 | 0.00010146 | 3.97811317 | 3.99369218 |
| NTRK3 | 15.65880425 | 4.7162E-05 | 3.96890214 | 4.326412 |
| NHSL2 | 15.56641416 | 3.6083E-05 | 3.96036474 | 4.44270114 |
| DSC1 | 15.3863163 | 0.00021376 | 3.94357597 | 3.67008046 |

| | | | | |
|---|---|---|---|---|
| C1orf21 | 15.32601458 | 8.0451E-05 | 3.93791068 | 4.09446756 |
| KRT78 | 14.73325197 | 7.2152E-05 | 3.881004 | 4.14175363 |
| ANXA1 | 14.58866695 | 0.00010905 | 3.86677616 | 3.96236168 |
| GOPC | 14.58796747 | 1.8648E-05 | 3.86670698 | 4.72936354 |
| PPP2R5E | 14.57562132 | 0.00089317 | 3.86548548 | 3.04906677 |
| CSTA | 14.55150468 | 0.00605379 | 3.86309644 | 2.21797241 |
| FLVCR1 | 14.50530674 | 2.6701E-06 | 3.8585089 | 5.57347919 |
| SNX9 | 14.45526646 | 0.00012169 | 3.8535233 | 3.91473518 |
| DDX3Y | 14.43082913 | 5.0746E-06 | 3.85108229 | 5.2945999 |
| AFTPH | 14.38650608 | 1.9205E-05 | 3.84664436 | 4.71658507 |
| ACOT1 | 14.34717524 | 9.4713E-05 | 3.84269481 | 4.02359131 |
| SLC1A4 | 14.27984624 | 0.00016653 | 3.83590854 | 3.77849587 |
| IMP4 | 14.23209277 | 5.7312E-05 | 3.83107592 | 4.24175218 |
| KRT14 | 14.22091617 | 0.0001935 | 3.82994251 | 3.71330854 |
| CEP164 | 13.83851005 | 8.02E-06 | 3.79061672 | 5.09582603 |
| TNRC6C | 13.81967241 | 9.942E-06 | 3.78865151 | 5.0025264 |
| NUP37 | 13.7747193 | 6.5402E-05 | 3.78395101 | 4.18440964 |
| PDHA1 | 13.75362183 | 0.00026874 | 3.78173968 | 3.57066275 |
| ARFIP2 | 13.35935767 | 2.1195E-05 | 3.73977874 | 4.67377659 |
| PLEKHA5 | 13.34703176 | 6.9361E-06 | 3.73844703 | 5.15888676 |
| RRP36 | 13.31361748 | 0.00011289 | 3.73483072 | 3.94733178 |
| PIP | 13.26522501 | 0.00343474 | 3.72957724 | 2.46410583 |
| EFNB2 | 13.21547367 | 1.8788E-05 | 3.72415623 | 4.72612987 |
| KRT5 | 13.0961472 | 0.00015222 | 3.71107054 | 3.81752788 |
| VAMP2 | 12.94189022 | 0.00024739 | 3.69397644 | 3.60662195 |
| STX6 | 12.89118939 | 0.00012686 | 3.68831347 | 3.89668816 |
| YTHDF1 | 12.83432524 | 6.4424E-06 | 3.68193554 | 5.19095476 |
| ACSL3 | 12.73638148 | 0.01724608 | 3.67088355 | 1.76330962 |
| PATJ | 12.65100899 | 0.00189312 | 3.66118055 | 2.72282192 |
| GPRIN1 | 12.63451143 | 4.3861E-06 | 3.65929797 | 5.35792217 |
| KCNB2 | 12.56880777 | 0.00021786 | 3.6517759 | 3.66181759 |
| RIMKLA | 12.56359049 | 1.4369E-05 | 3.65117692 | 4.84257296 |
| HAUS6 | 12.54992618 | 9.5664E-06 | 3.64960697 | 5.01925081 |
| MDH2 | 12.48919193 | 0.00022439 | 3.64260823 | 3.64899954 |
| GOLGA4 | 12.48323815 | 1.2196E-05 | 3.64192031 | 4.91379717 |
| BAG2 | 12.44527869 | 3.5294E-06 | 3.63752663 | 5.45229536 |
| TOM1L1 | 12.43745981 | 1.6679E-05 | 3.63661996 | 4.77783907 |
| HSPA9 | 12.27795732 | 1.1466E-05 | 3.61799866 | 4.94057018 |
| SLC30A5 | 12.25982862 | 8.1041E-05 | 3.61586691 | 4.09129783 |
| CD2AP | 12.23402481 | 3.7883E-05 | 3.6128272 | 4.42155385 |
| JAG2 | 12.05003213 | 8.2076E-05 | 3.59096509 | 4.0857842 |
| CNBP | 12.0283384 | 7.7872E-06 | 3.58836546 | 5.1086211 |
| CXADR | 11.9893037 | 0.00146975 | 3.58367597 | 2.83275797 |
| BAZ2B | 11.98554745 | 2.9879E-05 | 3.5832239 | 4.52464113 |
| PAK6 | 11.97776806 | 0.00406852 | 3.5822872 | 2.39056401 |

| | | | | |
|---|---|---|---|---|
| ZNF217 | 11.87811643 | 0.0001287 | 3.57023417 | 3.89040839 |
| ARL8A | 11.80818076 | 0.00819834 | 3.56171481 | 2.08627417 |
| SORBS2 | 11.71445708 | 1.489E-05 | 3.55021819 | 4.8270924 |
| HLA-A | 11.68104678 | 5.8803E-05 | 3.54609766 | 4.23059917 |
| ECE1 | 11.66173716 | 9.9427E-05 | 3.54371081 | 4.0024975 |
| JAM3 | 11.65006798 | 0.00054359 | 3.54226647 | 3.26472587 |
| SARS2 | 11.62787918 | 3.234E-05 | 3.53951608 | 4.49026615 |
| NIPSNAP1 | 11.62740943 | 0.00035318 | 3.5394578 | 3.45199946 |
| KRT72 | 11.62423919 | 0.00021335 | 3.53906439 | 3.67091022 |
| ICA1 | 11.62330641 | 0.0002098 | 3.53894862 | 3.67819192 |
| RPL36AL | 11.57517835 | 7.3738E-05 | 3.53296252 | 4.13230602 |
| KRT71 | 11.51984755 | 0.00018661 | 3.52604972 | 3.72905652 |
| PRR11 | 11.47986239 | 0.00026584 | 3.52103344 | 3.57537895 |
| LMTK2 | 11.47619019 | 3.6024E-06 | 3.52057188 | 5.44340405 |
| TOLLIP | 11.45947859 | 8.2118E-06 | 3.5184695 | 5.08556423 |
| KLHL26 | 11.44798752 | 0.00023437 | 3.5170221 | 3.6301046 |
| L2HGDH | 11.43899241 | 0.00012426 | 3.51588807 | 3.90566602 |
| H1-0 | 11.36972287 | 4.114E-05 | 3.50712518 | 4.38573852 |
| IDH3A | 11.348247 | 0.00011464 | 3.50439755 | 3.9406749 |
| EPHX2 | 11.28701405 | 7.9707E-06 | 3.49659197 | 5.09850376 |
| ESD | 11.2691924 | 0.00023694 | 3.49431222 | 3.62536163 |
| **LIMD1** | 11.18740876 | 7.4599E-05 | 3.48380401 | 4.12726551 |
| SREK1IP1 | 11.16566121 | 2.6335E-05 | 3.48099678 | 4.57947251 |
| TRIM25 | 11.08645338 | 4.2847E-05 | 3.47072601 | 4.36807863 |
| FAM171A2 | 11.03987504 | 1.1521E-05 | 3.46465194 | 4.93850546 |
| IGSF9B | 10.94857368 | 0.61229993 | 3.45267103 | 0.21303579 |
| ATAD3B | 10.9468473 | 4.3787E-05 | 3.45244353 | 4.35865877 |
| TENM4 | 10.93050741 | 9.6962E-07 | 3.45028847 | 6.01339894 |
| ACSL4 | 10.83343355 | 0.00076111 | 3.43741866 | 3.11855217 |
| GOLGA5 | 10.79606351 | 1.2924E-05 | 3.43243346 | 4.88861218 |
| COBLL1 | 10.79223107 | 1.5797E-05 | 3.43192124 | 4.80142542 |
| EPB41L1 | 10.78077619 | 0.00016955 | 3.43038915 | 3.7706988 |
| GOLGA4 | 10.77080898 | 0.01387763 | 3.42905471 | 1.85768463 |
| CASP14 | 10.76959407 | 0.00159079 | 3.42889197 | 2.79838796 |
| AKNA | 10.76149079 | 0.00863423 | 3.42780604 | 2.06377624 |
| MTOR | 10.75107818 | 1.1896E-05 | 3.42640944 | 4.924581 |
| ETFB | 10.71227213 | 3.8138E-05 | 3.42119261 | 4.41863811 |
| KRT28 | 10.71077892 | 0.00252911 | 3.4209915 | 2.59703156 |
| **TBK1** | 10.63472686 | 3.7661E-05 | 3.41071108 | 4.42410851 |
| MFSD6 | 10.5516274 | 0.17153937 | 3.39939362 | 0.7656362 |
| MRPS26 | 10.52829581 | 9.6008E-05 | 3.39620002 | 4.01769331 |
| SEMA6A | 10.47096909 | 1.6721E-05 | 3.38832306 | 4.77674068 |
| PCDH19 | 10.41173115 | 0.00711487 | 3.38013806 | 2.147833 |
| ACO2 | 10.37444793 | 1.5694E-05 | 3.37496266 | 4.80426131 |
| TMF1 | 10.30627695 | 2.1912E-05 | 3.36545136 | 4.65931213 |

| | | | | |
|---|---|---|---|---|
| PIEZO2 | 10.27063824 | 3.9095E-05 | 3.36045393 | 4.4078751 |
| CRIP1 | 10.21623927 | 9.7415E-06 | 3.35279231 | 5.01137594 |
| SEC16B | 10.18878955 | 0.00030152 | 3.34891076 | 3.5206808 |
| DSP | 10.16703311 | 0.0019821 | 3.34582684 | 2.70287355 |
| PPP4R3A | 10.1635377 | 0.00127017 | 3.34533075 | 2.89613853 |
| IFT20 | 10.14681464 | 0.0004043 | 3.34295499 | 3.39329893 |
| WDR62 | 10.12320331 | 0.00039328 | 3.33959397 | 3.40529399 |
| KRT27 | 9.972996948 | 0.00045854 | 3.31802711 | 3.33862588 |
| SEC24B | 9.962969945 | 1.7939E-05 | 3.31657587 | 4.74620092 |
| FCHSD2 | 9.95505641 | 1.2004E-05 | 3.31542949 | 4.92067422 |
| CEP85 | 9.84521317 | 0.00010917 | 3.29942244 | 3.96190858 |
| SDR39U1 | 9.814117101 | 0.01464963 | 3.29485849 | 1.83417344 |
| ARFIP1 | 9.804940928 | 8.8565E-06 | 3.29350894 | 5.05273693 |
| HSBP1 | 9.759743976 | 0.02545054 | 3.2868433 | 1.59430308 |
| AFDN | 9.7578387 | 1.1991E-05 | 3.28656164 | 4.92113657 |
| NIPSNAP2 | 9.722201996 | 0.00042993 | 3.28128311 | 3.36659854 |
| GLRX5 | 9.679132565 | 0.00016911 | 3.27487776 | 3.7718367 |
| RAB11FIP5 | 9.61775312 | 4.0473E-05 | 3.26569989 | 4.39283375 |
| RAB12 | 9.591713233 | 2.8928E-05 | 3.26178853 | 4.53868299 |
| RPL30 | 9.5694701 | 0.01526811 | 3.25843904 | 1.81621479 |
| SORBS1 | 9.521674856 | 3.6474E-05 | 3.25121536 | 4.43801399 |
| H2BC21 | 9.514900842 | 4.1621E-05 | 3.25018862 | 4.38069117 |
| ZFYVE9 | 9.503037334 | 0.00093902 | 3.2483887 | 3.02732698 |
| SEC23B | 9.495488393 | 0.00014775 | 3.24724221 | 3.83048201 |
| STEAP3 | 9.478897591 | 5.221E-05 | 3.24471928 | 4.28224968 |
| TTN | 9.463050707 | 0.00207383 | 3.24230536 | 2.68322621 |
| CTNNB1 | 9.434454207 | 7.6349E-05 | 3.23793906 | 4.11719564 |
| EPB41 | 9.426694921 | 9.7607E-05 | 3.23675204 | 4.01051703 |
| WASHC2A | 9.393378915 | 3.7427E-05 | 3.23164421 | 4.4268152 |
| ALG6 | 9.365193696 | 0.00136844 | 3.22730883 | 2.86377367 |
| NUP42 | 9.354226645 | 3.0594E-05 | 3.22561838 | 4.51436029 |
| PCDH9 | 9.251457101 | 3.0957E-05 | 3.20968061 | 4.5092475 |
| SLC29A1 | 9.238903587 | 0.00012275 | 3.20772165 | 3.91097096 |
| BCAR1 | 9.238013928 | 9.7179E-05 | 3.20758272 | 4.01242723 |
| LLPH | 9.231532623 | 2.8161E-05 | 3.20657018 | 4.55035551 |
| HADHB | 9.189678855 | 0.00033105 | 3.20001445 | 3.48011198 |
| FLG | 9.176148082 | 0.00310354 | 3.19788867 | 2.50814265 |
| F5 | 9.172956048 | 7.8071E-05 | 3.19738673 | 4.10750785 |
| CSTB | 9.166216067 | 0.00196829 | 3.19632629 | 2.70591099 |
| PLEKHA6 | 9.159789473 | 5.8128E-06 | 3.19531444 | 5.23561176 |
| STK3 | 9.102367282 | 9.7492E-05 | 3.1862418 | 4.0110293 |
| GCC1 | 9.039054714 | 0.00011292 | 3.17617191 | 3.9472272 |
| NECTIN1 | 8.764807214 | 0.00039055 | 3.13172236 | 3.40832275 |
| PRKCA | 8.754363896 | 0.00012042 | 3.13000235 | 3.91929881 |
| C9orf72 | 8.751008791 | 0.00045759 | 3.12944934 | 3.33952598 |

| | | | | |
|---|---|---|---|---|
| RAB25 | 8.703525424 | 0.00050063 | 3.12159989 | 3.30048281 |
| SCAMP3 | 8.675508766 | 0.00013012 | 3.11694837 | 3.88564835 |
| SNX3 | 8.67091133 | 0.00022771 | 3.11618363 | 3.64260981 |
| LASP1 | 8.666644598 | 0.00019443 | 3.11547354 | 3.71123134 |
| MRPL9 | 8.628972722 | 0.00151669 | 3.10918882 | 2.81910326 |
| JPH1 | 8.608993329 | 6.3504E-06 | 3.10584455 | 5.19720233 |
| AK3 | 8.537366309 | 0.00010317 | 3.09379108 | 3.98646469 |
| PHETA2 | 8.490647479 | 8.0041E-05 | 3.08587457 | 4.09668973 |
| IGF2R | 8.468101459 | 0.01276193 | 3.08203855 | 1.89408359 |
| RPS6KC1 | 8.445933478 | 0.01001496 | 3.07825688 | 1.99935093 |
| STX7 | 8.443247581 | 8.3228E-05 | 3.07779802 | 4.07973058 |
| TLCD3A | 8.434653928 | 0.19456041 | 3.07632888 | 0.71094553 |
| KRT4 | 8.427394188 | 0.00244699 | 3.07508661 | 2.61136721 |
| CDH2 | 8.407525079 | 0.00036938 | 3.07168118 | 3.43252577 |
| ILDR2 | 8.373620919 | 0.00043157 | 3.06585161 | 3.36495086 |
| ILRUN | 8.352602673 | 0.00070849 | 3.06222581 | 3.14966634 |
| SCAMP2 | 8.311111755 | 0.00026722 | 3.05504148 | 3.57313394 |
| PNPLA2 | 8.230958435 | 0.00401628 | 3.04106043 | 2.39617559 |
| BICD1 | 8.226607335 | 3.2838E-05 | 3.04029758 | 4.48362784 |
| ZDHHC3 | 8.219281889 | 0.00094537 | 3.03901235 | 3.02439602 |
| INSR | 8.182118339 | 0.00015643 | 3.0324744 | 3.80568367 |
| MRPL47 | 8.175492569 | 2.6714E-05 | 3.03130565 | 4.57326445 |
| WDR20 | 8.159663454 | 1.2194E-06 | 3.02850965 | 5.91386736 |
| TSPAN9 | 8.134494283 | 0.83200082 | 3.02405266 | 0.07987624 |
| CPD | 8.133628747 | 0.00138542 | 3.02389914 | 2.8584199 |
| STAT5B | 8.129761768 | 5.1407E-06 | 3.02321308 | 5.28897528 |
| TMED10 | 8.120946189 | 0.00012187 | 3.02164783 | 3.91408902 |
| SEC24A | 8.075375294 | 2.9164E-05 | 3.01352931 | 4.53515135 |
| TNIP1 | 8.067095036 | 0.00176181 | 3.01204925 | 2.7540418 |
| ADIRF | 8.065550169 | 0.00029668 | 3.01177295 | 3.52771401 |
| STX10 | 8.060439641 | 0.00026457 | 3.01085853 | 3.57745246 |
| N4BP1 | 8.049405747 | 4.116E-05 | 3.00888228 | 4.38552294 |
| PIK3C2A | 8.024388808 | 6.6299E-05 | 3.00439151 | 4.17849188 |
| AGK | 8.024309669 | 0.00019736 | 3.00437728 | 3.70473213 |
| UBAP1 | 7.999865574 | 3.4303E-05 | 2.99997576 | 4.46466484 |
| ABLIM1 | 7.987785419 | 0.00029401 | 2.99779558 | 3.53163339 |
| SDE2 | 7.977081977 | 4.6697E-05 | 2.9958611 | 4.33071336 |
| PLEKHJ1 | 7.940867775 | 0.00172731 | 2.98929667 | 2.76262918 |
| JUP | 7.939570357 | 0.00050439 | 2.98906094 | 3.29723412 |
| RAB1A | 7.923815667 | 0.00043175 | 2.98619532 | 3.36477051 |
| VTI1B | 7.907043164 | 0.00013893 | 2.9831383 | 3.85721143 |
| CNN3 | 7.895748093 | 0.00103924 | 2.98107596 | 2.98328265 |
| PDLIM5 | 7.885186384 | 4.2249E-06 | 2.97914486 | 5.37417898 |
| ABCB7 | 7.87194226 | 0.01204014 | 2.97671964 | 1.91936863 |
| RPL18 | 7.87026725 | 3.6763E-05 | 2.97641263 | 4.43459169 |

| | | | | |
|---|---|---|---|---|
| TAB3 | 7.763421418 | 0.00161276 | 2.9566926 | 2.79242966 |
| RPA1 | 7.739729205 | 0.00046091 | 2.95228309 | 3.33638033 |
| SMS | 7.733136794 | 5.244E-05 | 2.95105373 | 4.28033664 |
| RPS18 | 7.601799208 | 0.00019987 | 2.92634092 | 3.69925949 |
| LUC7L2 | 7.573561823 | 0.00017478 | 2.92097195 | 3.75752022 |
| SYTL2 | 7.549445878 | 2.5559E-05 | 2.91637076 | 4.59245387 |
| NDUFA10 | 7.537301247 | 6.9341E-05 | 2.91404805 | 4.15900905 |
| ACAT1 | 7.508500447 | 0.00483916 | 2.90852481 | 2.31522967 |
| ANKRD26 | 7.493039417 | 0.00019358 | 2.90555104 | 3.71314556 |
| RACK1 | 7.460599539 | 0.00015704 | 2.89929157 | 3.80399537 |
| GAB2 | 7.456584954 | 3.1863E-05 | 2.89851504 | 4.49671939 |
| DAG1 | 7.448034027 | 0.00050359 | 2.89685966 | 3.29791909 |
| NUP43 | 7.435932245 | 0.00010648 | 2.89451362 | 3.97273155 |
| SDHA | 7.434232231 | 0.00052504 | 2.89418376 | 3.2798099 |
| CEP135 | 7.404066445 | 0.00054027 | 2.88831784 | 3.2673891 |
| SENP1 | 7.332839975 | 0.00368322 | 2.87437206 | 2.43377239 |
| WASHC2C | 7.322105361 | 2.2333E-05 | 2.87225853 | 4.65104978 |
| HIP1 | 7.286922642 | 3.8767E-05 | 2.86530968 | 4.41153767 |
| CRYBG3 | 7.252296857 | 1.9287E-05 | 2.85843798 | 4.71473955 |
| SNX18 | 7.23705644 | 6.8956E-05 | 2.85540302 | 4.16142851 |
| GOSR1 | 7.202930333 | 0.00137058 | 2.84858395 | 2.86309585 |
| ABLIM2 | 7.194349503 | 3.8425E-05 | 2.84686425 | 4.41538656 |
| MTSS2 | 7.172073483 | 0.00021059 | 2.84239027 | 3.6765654 |
| PRDX4 | 7.170335809 | 2.9928E-05 | 2.84204069 | 4.52391922 |
| UBAP2 | 7.11820283 | 1.5535E-05 | 2.83151304 | 4.80868515 |
| YWHAQ | 7.109449374 | 6.1647E-06 | 2.82973783 | 5.21008847 |
| EBNA1BP2 | 7.087824657 | 6.8941E-05 | 2.82534291 | 4.16152151 |
| STON1-GTF2A1L | 7.082384271 | 0.0004956 | 2.82423512 | 3.30486509 |
| FLG2 | 7.078504474 | 0.00880074 | 2.82344458 | 2.055481 |
| SCYL2 | 7.066647429 | 2.8816E-05 | 2.82102593 | 4.54036975 |
| ADCY9 | 7.064553042 | 0.00033285 | 2.82059829 | 3.47774495 |
| GARS1 | 7.062186708 | 0.00120889 | 2.82011496 | 2.91761474 |
| RPL8 | 7.051556123 | 0.00119314 | 2.81794166 | 2.92330831 |
| RUFY2 | 7.047677923 | 0.00266467 | 2.81714799 | 2.57435673 |
| C1QBP | 7.025923269 | 0.01211653 | 2.81268782 | 1.91662185 |
| PTRHD1 | 7.001078744 | 0.00017175 | 2.80757723 | 3.76509171 |
| MRPS15 | 6.99914804 | 0.00013904 | 2.80717932 | 3.85685068 |
| CTNNA1 | 6.995891447 | 6.6724E-05 | 2.8065079 | 4.17571998 |
| AMIGO2 | 6.989931891 | 0.00142041 | 2.8052784 | 2.84758746 |
| PPL | 6.957690972 | 0.00010186 | 2.7986086 | 3.99198533 |
| HADH | 6.953800669 | 0.00239396 | 2.79780171 | 2.62088399 |
| AUP1 | 6.950150116 | 5.1879E-05 | 2.79704414 | 4.28500749 |
| TRIP11 | 6.944731558 | 1.4877E-05 | 2.79591893 | 4.82749206 |
| MTMR9 | 6.92543492 | 0.0001064 | 2.79190468 | 3.97304004 |
| PEX5L | 6.903662234 | 9.0303E-05 | 2.78736188 | 4.04429887 |

| | | | | |
|---|---|---|---|---|
| ANXA2 | 6.8910229 | 0.00021345 | 2.78471815 | 3.6707057 |
| TACC2 | 6.878141874 | 0.00167865 | 2.78201887 | 2.77503891 |
| RPS5 | 6.871102414 | 6.7397E-05 | 2.78054159 | 4.17136164 |
| DDX3X | 6.868919259 | 4.3643E-05 | 2.78008313 | 4.36008504 |
| PDHB | 6.861910573 | 0.00043457 | 2.77861032 | 3.3619439 |
| NDUFAF1 | 6.83496862 | 0.00048806 | 2.77293471 | 3.31152985 |
| STX17 | 6.77521557 | 0.01473925 | 2.76026685 | 1.83152473 |
| RAB23 | 6.726512845 | 3.8893E-05 | 2.74985878 | 4.4101339 |
| TAB2 | 6.71545793 | 0.00072563 | 2.74748578 | 3.13928245 |
| SEMA4C | 6.700256578 | 0.00034555 | 2.74421634 | 3.46148442 |
| RHOG | 6.699152659 | 0.00116973 | 2.74397863 | 2.93191272 |
| BTBD7 | 6.697321968 | 0.00099765 | 2.74358433 | 3.00102358 |
| RAB1B | 6.636118918 | 0.00041273 | 2.73033974 | 3.38433638 |
| PGK1 | 6.627852453 | 4.2857E-05 | 2.72854149 | 4.36797604 |
| GAPDH | 6.60512276 | 6.3976E-05 | 2.72358538 | 4.19398195 |
| MPZL1 | 6.566819887 | 0.0009938 | 2.71519489 | 3.00270277 |
| MTHFD2 | 6.554766747 | 5.2191E-05 | 2.71254444 | 4.28240228 |
| RPS3A | 6.552955226 | 0.00133467 | 2.71214567 | 2.87462577 |
| CDCA3 | 6.54441784 | 0.00039585 | 2.71026486 | 3.40247218 |
| ARFGAP1 | 6.459437641 | 6.4232E-05 | 2.69140857 | 4.19224712 |
| CEP63 | 6.436819528 | 0.00050135 | 2.68634802 | 3.29986027 |
| DARS2 | 6.408392135 | 0.00068095 | 2.67996243 | 3.16688416 |
| NUMB | 6.396857173 | 0.00010045 | 2.67736327 | 3.9980533 |
| ATAD3A | 6.379432468 | 4.7119E-05 | 2.67342808 | 4.32680028 |
| RPL15 | 6.374397281 | 7.5583E-05 | 2.67228894 | 4.12157584 |
| MPST | 6.369048386 | 0.0001731 | 2.67107783 | 3.76170469 |
| BCKDHA2 | 6.368164415 | 0.0002465 | 2.67087758 | 3.60818287 |
| APBB2 | 6.365908661 | 1.4112E-05 | 2.67036646 | 4.8504109 |
| PICALM | 6.333050692 | 3.8715E-05 | 2.66290063 | 4.41212463 |
| CBLB | 6.311906246 | 3.6784E-05 | 2.65807578 | 4.4343396 |
| OCRL | 6.295097139 | 0.00023461 | 2.65422864 | 3.62965074 |
| SEC13 | 6.292755624 | 0.0006336 | 2.65369192 | 3.19818241 |
| LIMK2 | 6.285295931 | 0.04463805 | 2.65198067 | 1.35029479 |
| PRKRIP1 | 6.284151499 | 0.02759996 | 2.65171796 | 1.55909159 |
| TPT1 | 6.28217881 | 0.00644765 | 2.65126501 | 2.19059849 |
| ACBD3 | 6.22826657 | 1.0891E-05 | 2.63883069 | 4.96293684 |
| KIAA0319L | 6.225696901 | 0.00048095 | 2.63823534 | 3.31790222 |
| MRPL28 | 6.206452949 | 0.00760496 | 2.63376899 | 2.11890315 |
| PRUNE2 | 6.196063792 | 0.00016152 | 2.631352 | 3.79176606 |
| POLDIP2 | 6.191083764 | 0.00018303 | 2.63019198 | 3.73747473 |
| PC | 6.187231496 | 4.7622E-05 | 2.62929401 | 4.32219097 |
| ERP44 | 6.183135762 | 6.8518E-05 | 2.62833868 | 4.16419736 |
| FNBP1 | 6.165125436 | 0.00016142 | 2.62413025 | 3.79204317 |
| C1orf198 | 6.16255992 | 4.2209E-05 | 2.62352977 | 4.37459251 |
| TECPR1 | 6.147566153 | 0.0404037 | 2.62001535 | 1.39357889 |

| | | | | |
|---|---|---|---|---|
| BSN | 6.121808763 | 0.00020375 | 2.61395798 | 3.6909036 |
| HSD17B10 | 6.08580265 | 1.6277E-05 | 2.60544755 | 4.7884265 |
| TMPO | 6.058024809 | 2.6025E-05 | 2.59884749 | 4.58461425 |
| UBAP2 | 6.05210621 | 0.00147427 | 2.59743731 | 2.83142297 |
| WDR41 | 6.019009755 | 2.1959E-05 | 2.58952615 | 4.65838866 |
| NADK2 | 6.001873272 | 0.00046089 | 2.58541286 | 3.33640363 |
| PDZD2 | 5.986985471 | 6.8982E-05 | 2.58182977 | 4.16126683 |
| SSBP1 | 5.986750508 | 0.00284007 | 2.58177315 | 2.54667137 |
| HELZ2 | 5.976118576 | 0.00208146 | 2.57920877 | 2.68163152 |
| CPT2 | 5.960233764 | 0.00029573 | 2.57536892 | 3.52911154 |
| LAP3 | 5.934944267 | 0.01079284 | 2.56923448 | 1.96686417 |
| TPPP3 | 5.928932333 | 0.00392771 | 2.56777233 | 2.40586005 |
| CRIP2 | 5.918574519 | 1.2533E-05 | 2.56524975 | 4.90195506 |
| GLS | 5.874651518 | 0.00020248 | 2.55450327 | 3.69362545 |
| FECH | 5.856247569 | 0.00054853 | 2.54997654 | 3.26079781 |
| ISCA2 | 5.844300447 | 0.00119917 | 2.54703035 | 2.92112027 |
| SEMG1 | 5.809465654 | 0.11383693 | 2.53840547 | 0.94371683 |
| UBE2N | 5.807006618 | 0.00025651 | 2.53779468 | 3.59089416 |
| GGCT | 5.801116787 | 0.00050557 | 2.53633066 | 3.2962197 |
| HADHA | 5.795945894 | 3.7778E-05 | 2.53504413 | 4.42276158 |
| MFAP3 | 5.789918313 | 0.00077942 | 2.53354299 | 3.10822732 |
| HYLS1 | 5.786440828 | 0.02914428 | 2.53267624 | 1.53544663 |
| LDB3 | 5.778797605 | 0.0003712 | 2.53076934 | 3.43039599 |
| TENM3 | 5.760190502 | 6.554E-05 | 2.52611653 | 4.18349181 |
| EP400 | 5.749041567 | 0.00012288 | 2.52332146 | 3.91050654 |
| CEP128 | 5.723478619 | 0.00019545 | 2.51689226 | 3.70897223 |
| MRPL37 | 5.711168529 | 0.00024068 | 2.51378596 | 3.6185588 |
| MMTAG2 | 5.69790098 | 0.00024925 | 2.51043055 | 3.60335908 |
| SNX1 | 5.6854819 | 3.9283E-05 | 2.50728264 | 4.40579342 |
| PSMA5 | 5.685119065 | 0.06246686 | 2.50719056 | 1.20435032 |
| CPLX2 | 5.676246299 | 0.00021495 | 2.50493719 | 3.6676699 |
| NUMBL | 5.654120455 | 0.00693438 | 2.49930262 | 2.15899252 |
| PCDH17 | 5.627972392 | 0.1317949 | 2.49261525 | 0.88010138 |
| TRABD | 5.619044373 | 0.00060738 | 2.49032479 | 3.21654051 |
| AARS2 | 5.617701265 | 0.00026822 | 2.48997991 | 3.57151449 |
| RPL21 | 5.592517586 | 0.000126 | 2.48349789 | 3.89964326 |
| MRPL22 | 5.587526284 | 0.00029573 | 2.48220971 | 3.52910226 |
| NUDT14 | 5.581814079 | 0.0015217 | 2.48073407 | 2.81767164 |
| EPB41L5 | 5.578917328 | 0.00211957 | 2.47998517 | 2.67375174 |
| GARRE1 | 5.57737259 | 0.01069452 | 2.47958565 | 1.97083865 |
| PNPT1 | 5.566384197 | 0.00017267 | 2.47674049 | 3.76279562 |
| MTMR6 | 5.563015792 | 0.00019619 | 2.4758672 | 3.7073125 |
| HINT2 | 5.562378166 | 0.03484282 | 2.47570183 | 1.45788666 |
| YWHAE | 5.529885196 | 0.00015547 | 2.46724953 | 3.80834809 |
| LZTS3 | 5.508749454 | 9.706E-05 | 2.46172485 | 4.01295961 |

| | | | | |
|---|---|---|---|---|
| IDH3B | 5.506550867 | 5.3938E-05 | 2.46114894 | 4.26810281 |
| PGRMC1 | 5.490546833 | 0.00070839 | 2.45694984 | 3.14972588 |
| NAA10 | 5.475464113 | 3.0038E-05 | 2.45298126 | 4.52232677 |
| TIAL1 | 5.418283717 | 0.06038888 | 2.43783594 | 1.219043 |
| ECHS1 | 5.411328253 | 0.00089315 | 2.43598276 | 3.04907498 |
| CERT1 | 5.405152687 | 0.00137566 | 2.43433537 | 2.86148946 |
| MMUT | 5.393927444 | 0.15520371 | 2.43133612 | 0.8090979 |
| LUC7L | 5.372649934 | 0.00066669 | 2.42563384 | 3.17607382 |
| DOP1A | 5.364617078 | 0.00260967 | 2.4234752 | 2.58341376 |
| SIPA1L3 | 5.36025002 | 0.00028183 | 2.42230029 | 3.55001143 |
| ACADVL | 5.341589703 | 0.00106788 | 2.41726916 | 2.97147778 |
| SCLT1 | 5.338161036 | 0.00528076 | 2.41634283 | 2.2773034 |
| SHMT2 | 5.318419773 | 6.2304E-06 | 2.41099765 | 5.20548394 |
| TOM1 | 5.310706023 | 4.1751E-05 | 2.40890367 | 4.37933223 |
| PPA2 | 5.304373471 | 0.02843014 | 2.40718236 | 1.54622103 |
| NXN | 5.297981308 | 0.08692961 | 2.40544275 | 1.06083226 |
| ARHGAP8 | 5.272700619 | 0.00206885 | 2.39854208 | 2.68427202 |
| DNAJA3 | 5.262513441 | 0.00012408 | 2.39575201 | 3.90628257 |
| MRPS22 | 5.25588764 | 0.00105876 | 2.39393443 | 2.97520338 |
| RAB3D | 5.254724701 | 3.5843E-05 | 2.39361518 | 4.44559522 |
| ANKS6 | 5.250360645 | 0.20459841 | 2.39241652 | 0.68909775 |
| SUCLG1 | 5.248552435 | 5.4381E-05 | 2.39191958 | 4.26455661 |
| TJP1 | 5.241118382 | 0.00010225 | 2.3898747 | 3.9903238 |
| STX5 | 5.233848032 | 0.00011892 | 2.38787204 | 3.92472822 |
| MRPL1 | 5.230053941 | 0.0016725 | 2.38682583 | 2.776635 |
| ACSF3 | 5.223076673 | 0.01037623 | 2.38489988 | 1.98396032 |
| DDX54 | 5.212837025 | 2.6273E-05 | 2.38206876 | 4.5804949 |
| NDUFAF2 | 5.203589389 | 0.0011763 | 2.37950712 | 2.92948279 |
| PPP1R13L | 5.195916773 | 0.0080527 | 2.37737832 | 2.09405858 |
| POF1B | 5.188497787 | 0.00208669 | 2.3753169 | 2.68054113 |
| FLOT1 | 5.186879462 | 0.00071967 | 2.37486684 | 3.14286812 |
| MRPL39 | 5.183439882 | 0.00094045 | 2.37390983 | 3.02666484 |
| CAPZA1 | 5.181223042 | 0.00027743 | 2.37329269 | 3.55685349 |
| SNAP29 | 5.176172235 | 3.9096E-05 | 2.37188562 | 4.40786451 |
| GPN1 | 5.173279471 | 0.0002397 | 2.37107913 | 3.62032898 |
| RPL37A | 5.169814241 | 0.00142038 | 2.37011244 | 2.84759532 |
| SERPINB4 | 5.146902128 | 0.10328896 | 2.36370435 | 0.98594611 |
| BUD23 | 5.146834993 | 0.00149793 | 2.36368553 | 2.82450738 |
| YWHAH | 5.145136763 | 0.00019005 | 2.36320943 | 3.72113315 |
| NOPCHAP1 | 5.139209395 | 0.00052059 | 2.36154644 | 3.28350459 |
| RPL13A | 5.139069725 | 0.00091128 | 2.36150723 | 3.04034855 |
| H2BC18 | 5.121908856 | 0.00066925 | 2.35668158 | 3.17441442 |
| POLD1 | 5.118748155 | 0.00185791 | 2.35579103 | 2.73097634 |
| PIK3C2B | 5.104794703 | 0.00058269 | 2.35185294 | 3.23456206 |
| PYCR2 | 5.103737134 | 0.00084056 | 2.35155403 | 3.07542909 |

| | | | | |
|---|---|---|---|---|
| HPRT1 | 5.068328119 | 0.00734937 | 2.34150993 | 2.13374963 |
| SH3BP4 | 5.06502191 | 0.00035479 | 2.34056851 | 3.45002627 |
| PMPCB | 5.024673635 | 0.00012024 | 2.32902989 | 3.91996605 |
| ARHGAP18 | 5.023241348 | 2.7585E-05 | 2.32861859 | 4.55932882 |
| ACTA1 | 5.011883395 | 0.00075894 | 2.32535285 | 3.11979343 |
| EPHA4 | 5.006276018 | 0.1871259 | 2.32373784 | 0.72786609 |
| FAIM | 5.005876688 | 0.00072647 | 2.32362275 | 3.13878022 |
| GRPEL1 | 5.005820595 | 0.00156827 | 2.32360659 | 2.80457885 |
| MRPS34 | 4.999851068 | 0.00044027 | 2.32188512 | 3.35627775 |
| MAP2 | 4.999660837 | 0.00070597 | 2.32183023 | 3.15121403 |
| HGS | 4.997376179 | 0.00023203 | 2.32117082 | 3.63445274 |
| ATP5OP | 4.993682775 | 0.00560476 | 2.32010418 | 2.25144273 |
| SYNGR2 | 4.974975302 | 0.0030681 | 2.31468936 | 2.51312988 |
| GRIPAP1 | 4.971020518 | 0.00013097 | 2.31354206 | 3.88281166 |
| ECHDC1 | 4.967091541 | 0.00016298 | 2.31240134 | 3.78786996 |
| USP31 | 4.951413072 | 0.00066877 | 2.30784031 | 3.17472429 |
| LRPPRC | 4.934905523 | 4.8978E-05 | 2.30302247 | 4.31000093 |
| SLC9A1 | 4.909917638 | 0.00146511 | 2.29569882 | 2.83412938 |
| TANC2 | 4.909188333 | 0.00057435 | 2.29548451 | 3.24082348 |
| MEX3B | 4.903936196 | 0.00556605 | 2.29394021 | 2.25445325 |
| AP2M1 | 4.903740857 | 6.9895E-05 | 2.29388274 | 4.15555311 |
| PPIL3 | 4.880371236 | 0.00099769 | 2.28699089 | 3.001004 |
| NSA2 | 4.873169042 | 0.00064193 | 2.28486027 | 3.19251383 |
| EHBP1 | 4.867833476 | 9.096E-05 | 2.28327982 | 4.04115191 |
| RAB14 | 4.864686465 | 0.00045641 | 2.28234682 | 3.34064575 |
| USP32 | 4.859025244 | 0.01041604 | 2.28066693 | 1.98229726 |
| KRT23 | 4.854266922 | 0.00565316 | 2.27925344 | 2.247709 |
| HSPB1 | 4.844723133 | 6.6974E-05 | 2.27641422 | 4.17409054 |
| RAB2A | 4.838475517 | 0.00047838 | 2.27455256 | 3.32022903 |
| PITPNB | 4.827763374 | 0.00040788 | 2.27135497 | 3.38947111 |
| TMEM126B | 4.826227159 | 0.00192895 | 2.27089582 | 2.71467903 |
| STX16 | 4.817663658 | 0.00490336 | 2.26833368 | 2.30950655 |
| NF2 | 4.815097869 | 1.3392E-05 | 2.26756512 | 4.87314082 |
| PRDX2 | 4.81072849 | 3.7711E-05 | 2.26625538 | 4.42353359 |
| DHRS3 | 4.80820942 | 0.00041031 | 2.26549973 | 3.38688329 |
| RPS6 | 4.803997726 | 3.0456E-05 | 2.26423547 | 4.51633421 |
| BMP2K | 4.803708202 | 0.0002506 | 2.26414852 | 3.60102468 |
| ATP5PB | 4.80002789 | 0.01056136 | 2.26304279 | 1.97628015 |
| VPS37A | 4.799426922 | 0.00033639 | 2.26286215 | 3.4731562 |
| RAB21 | 4.791961794 | 0.00227386 | 2.26061641 | 2.64323586 |
| FCHO2 | 4.780578857 | 0.00122777 | 2.25718532 | 2.91088356 |
| RPL36A | 4.77350498 | 0.00051037 | 2.25504896 | 3.29211457 |
| CPNE1 | 4.76931401 | 0.00054079 | 2.25378177 | 3.26697078 |
| APRT | 4.749880858 | 0.00013723 | 2.24789133 | 3.86255707 |
| ADPRS | 4.743309734 | 0.00016818 | 2.24589408 | 3.77421945 |

| | | | | |
|---|---|---|---|---|
| GLUD1 | 4.73077233 | 0.00393455 | 2.24207573 | 2.40510545 |
| PSMD6 | 4.668628264 | 7.3582E-05 | 2.22299872 | 4.13322789 |
| MRPL45 | 4.667394292 | 0.00103781 | 2.22261735 | 2.98388335 |
| OGDH | 4.662375278 | 7.5478E-05 | 2.22106513 | 4.12217999 |
| NECAP2 | 4.659002668 | 0.00060909 | 2.22002116 | 3.21531955 |
| VPS26A | 4.657583426 | 0.00336601 | 2.21958161 | 2.47288438 |
| PARD3 | 4.656366543 | 0.00055755 | 2.21920463 | 3.25371877 |
| CEP78 | 4.638870811 | 0.01947497 | 2.21377367 | 1.7105233 |
| POLE3 | 4.631453337 | 0.00010627 | 2.21146498 | 3.97357987 |
| ACTG1 | 4.624005835 | 2.6306E-05 | 2.20914322 | 4.57994473 |
| CSRP1 | 4.618417824 | 0.00375201 | 2.2073987 | 2.42573548 |
| SYNJ1 | 4.618207385 | 4.4146E-05 | 2.20733296 | 4.35511082 |
| LANCL1 | 4.593982755 | 0.00160447 | 2.19974544 | 2.79466842 |
| SMG7 | 4.588427658 | 0.00013711 | 2.19799986 | 3.8629179 |
| ACAA2 | 4.577232297 | 0.00089132 | 2.19447551 | 3.04996586 |
| RPL23A | 4.575648543 | 0.00021391 | 2.19397624 | 3.66977398 |
| SLC1A5 | 4.574418828 | 0.00034518 | 2.19358846 | 3.46195877 |
| AHNAK2 | 4.569778996 | 4.4917E-05 | 2.1921244 | 4.34759308 |
| MTPN | 4.552341699 | 0.00074664 | 2.18660885 | 3.12688877 |
| CDK4 | 4.547103301 | 0.0009343 | 2.18494778 | 3.02951392 |
| RRS1 | 4.537342884 | 0.00051329 | 2.18184769 | 3.2896333 |
| AP4E1 | 4.533805259 | 0.00056191 | 2.18072242 | 3.25033017 |
| SHC1 | 4.528236244 | 0.00034631 | 2.17894923 | 3.46054006 |
| GAREM1 | 4.512233337 | 0.01344445 | 2.17384167 | 1.87145687 |
| MYO6 | 4.50130868 | 7.8882E-05 | 2.1703445 | 4.10302079 |
| CHP1 | 4.500666056 | 0.00060403 | 2.17013852 | 3.21893837 |
| NHERF2 | 4.495479712 | 0.01096213 | 2.16847507 | 1.96010504 |
| SMCR8 | 4.475202581 | 0.00044379 | 2.16195299 | 3.35282666 |
| NCAM1 | 4.474393806 | 0.01062077 | 2.16169224 | 1.97384389 |
| CCDC124 | 4.467820391 | 0.00196302 | 2.15957119 | 2.70707466 |
| ZDHHC20 | 4.466921591 | 0.18466343 | 2.15928093 | 0.7336191 |
| TFG | 4.461176283 | 0.00010999 | 2.15742416 | 3.95865759 |
| RPS20 | 4.45411089 | 0.00057757 | 2.15513748 | 3.23839681 |
| ATP5PD | 4.433861004 | 0.00287774 | 2.14856354 | 2.54094876 |
| RBM8A | 4.425081963 | 7.1321E-05 | 2.14570418 | 4.14678288 |
| CYB5R1 | 4.424161726 | 0.00037347 | 2.14540412 | 3.42774598 |
| PHACTR4 | 4.421161049 | 0.00020293 | 2.14442529 | 3.69264359 |
| RDH11 | 4.420022852 | 0.00021437 | 2.14405383 | 3.66884251 |
| LSR | 4.393528314 | 0.00022064 | 2.13537999 | 3.65630941 |
| RINT1 | 4.368832711 | 0.0001497 | 2.12724786 | 3.82476583 |
| IFNAR1 | 4.365286727 | 0.00465076 | 2.12607642 | 2.33247591 |
| SMAD4 | 4.364878041 | 0.00125369 | 2.12594134 | 2.90180812 |
| ATP5F1B | 4.360116524 | 0.00016762 | 2.12436669 | 3.77568179 |
| RPS25 | 4.358523848 | 6.3089E-05 | 2.1238396 | 4.20004918 |
| GFM1 | 4.35577168 | 0.00023292 | 2.12292833 | 3.63279391 |

| | | | | |
|---|---|---|---|---|
| PRPF4B | 4.343672174 | 2.8261E-05 | 2.11891522 | 4.54881577 |
| STAM2 | 4.338770703 | 0.0053851 | 2.11728634 | 2.26880641 |
| AP1AR | 4.333655952 | 6.2202E-05 | 2.11558462 | 4.20619912 |
| LRP6 | 4.331734645 | 0.00307382 | 2.11494487 | 2.51232173 |
| PKP3 | 4.327574064 | 4.7127E-05 | 2.11355851 | 4.32673446 |
| RBFOX1 | 4.318034327 | 0.00017843 | 2.11037471 | 3.74853709 |
| RDH14 | 4.300103263 | 0.00463303 | 2.10437131 | 2.33413492 |
| UQCRFS1 | 4.298727088 | 0.00070893 | 2.10390952 | 3.14939375 |
| TSG101 | 4.289208674 | 0.00059036 | 2.10071151 | 3.22887976 |
| USP12 | 4.248794478 | 0.00063057 | 2.08705356 | 3.20026343 |
| HNRNPAB | 4.248116009 | 0.00218869 | 2.08682316 | 2.65981556 |
| LPCAT1 | 4.243439642 | 0.01897127 | 2.08523416 | 1.72190354 |
| CLPP | 4.241796923 | 0.27877207 | 2.08467555 | 0.55475074 |
| UTP23 | 4.239808055 | 0.000874 | 2.08399895 | 3.05848885 |
| RNF121 | 4.23676735 | 0.00706178 | 2.08296391 | 2.15108557 |
| RELL1 | 4.229876003 | 0.00025038 | 2.08061537 | 3.60139213 |
| BSG | 4.226145348 | 0.00010937 | 2.07934239 | 3.9611072 |
| KEAP1 | 4.223024313 | 0.00147368 | 2.07827655 | 2.83159761 |
| RPS24 | 4.220839365 | 0.00126758 | 2.07752992 | 2.89702309 |
| PRDX3 | 4.206424098 | 0.00035511 | 2.07259431 | 3.44963428 |
| RP9 | 4.201700812 | 0.00087795 | 2.07097344 | 3.05653056 |
| ZDHHC17 | 4.200958682 | 0.00050263 | 2.0707186 | 3.2987492 |
| MRPL15 | 4.196846518 | 0.0011705 | 2.0693057 | 2.93162985 |
| TMX1 | 4.192290005 | 8.9541E-05 | 2.06773852 | 4.04797788 |
| COLGALT1 | 4.187067708 | 0.00121388 | 2.06594025 | 2.91582297 |
| MAGED1 | 4.172928062 | 0.00023752 | 2.06106005 | 3.62429275 |
| SCRN1 | 4.147574399 | 0.00327601 | 2.05226786 | 2.48465461 |
| AP1M1 | 4.140507668 | 0.00093306 | 2.04980767 | 3.03009062 |
| DIRAS1 | 4.138706852 | 0.00148866 | 2.04918006 | 2.82720441 |
| UBE2K | 4.137757915 | 0.01602273 | 2.04884924 | 1.79526337 |
| BLTP3B | 4.137414154 | 0.00019806 | 2.04872938 | 3.70319856 |
| COPZ1 | 4.1336043 | 0.00368929 | 2.04740029 | 2.43305673 |
| RCC1 | 4.120026761 | 0.00017774 | 2.04265371 | 3.75022404 |
| RABL6 | 4.117716118 | 0.00016489 | 2.04184437 | 3.7828054 |
| MACROH2A2 | 4.102708356 | 0.00022754 | 2.0365766 | 3.64293276 |
| CCNB2 | 4.097013643 | 0.00056135 | 2.0345727 | 3.25076514 |
| SMG5 | 4.096197107 | 0.00243743 | 2.03428514 | 2.61306837 |
| PSME2 | 4.072035102 | 0.00098237 | 2.02575 | 3.00772475 |
| NOS1AP | 4.068726196 | 0.00016706 | 2.0245772 | 3.77713669 |
| BMPR2 | 4.066254177 | 5.9225E-05 | 2.0237004 | 4.22749799 |
| ECSIT | 4.057649181 | 0.03792179 | 2.02064414 | 1.4211112 |
| FUBP3 | 4.05576962 | 0.00053198 | 2.01997571 | 3.27410245 |
| NDUFA2 | 4.052448379 | 0.00408081 | 2.01879381 | 2.38925314 |
| BRWD3 | 4.050229282 | 0.00110639 | 2.01800358 | 2.95609089 |
| MSI2 | 4.048312454 | 0.15990576 | 2.01732064 | 0.79613589 |

| | | | | |
|---|---|---|---|---|
| ATP9A | 4.041120422 | 0.00466167 | 2.01475534 | 2.33145843 |
| PDXDC1 | 4.036501305 | 0.00010324 | 2.01310536 | 3.9861337 |
| CBFB | 4.032563047 | 0.00162281 | 2.01169709 | 2.78973176 |
| AKAP10 | 4.027732173 | 6.1514E-05 | 2.00996775 | 4.21102664 |
| RPL36 | 4.027265728 | 0.0003734 | 2.00980067 | 3.42782627 |
| LAMTOR3 | 4.015635005 | 0.02053941 | 2.00562814 | 1.68741213 |
| FLOT2 | 4.006638394 | 1.1689E-05 | 2.00239231 | 4.93223752 |
| EIF6 | 4.000346947 | 0.0005647 | 2.00012513 | 3.24818523 |
| RAB35 | 3.999646007 | 0.00209613 | 1.99987232 | 2.67858078 |
| PITRM1 | 3.996312371 | 0.01905746 | 1.99866936 | 1.71993492 |
| SNAP25 | 3.996260587 | 0.000173 | 1.99865066 | 3.76195704 |
| CSNK1D | 3.995486341 | 0.18071745 | 1.99837112 | 0.74299991 |
| BABAM1 | 3.988543442 | 0.00281118 | 1.99586199 | 2.55111132 |
| ABCE1 | 3.982466772 | 0.0011694 | 1.99366232 | 2.93203579 |
| MTCH2 | 3.980070804 | 6.3505E-06 | 1.9927941 | 5.19719114 |
| BLTP3A | 3.978542717 | 0.0004486 | 1.99224009 | 3.34813867 |
| MRPS25 | 3.963872374 | 0.00113074 | 1.98691051 | 2.94663816 |
| SEC61B | 3.962019422 | 0.00577148 | 1.98623595 | 2.23871285 |
| KIF22 | 3.956157427 | 0.07551369 | 1.98409984 | 1.12197431 |
| GNG8 | 3.940886151 | 0.00747256 | 1.97852007 | 2.12653084 |
| FSCN1 | 3.933972014 | 0.00033258 | 1.97598669 | 3.47810472 |
| RPS4X | 3.930284891 | 0.00032921 | 1.97463389 | 3.48252149 |
| GRB2 | 3.929382522 | 0.0003974 | 1.97430262 | 3.40077523 |
| KDSR | 3.923849165 | 0.00211431 | 1.97226958 | 2.67483097 |
| HNRNPH2 | 3.912843436 | 0.04510999 | 1.96821739 | 1.34572726 |
| LSM14A | 3.907976583 | 0.03286012 | 1.96642182 | 1.48333085 |
| DLGAP3 | 3.89644758 | 0.0984482 | 1.96215941 | 1.00679223 |
| JPT1 | 3.886763974 | 7.371E-05 | 1.9585695 | 4.13247486 |
| RBMX | 3.88549831 | 4.7468E-05 | 1.95809963 | 4.32359551 |
| DLG1 | 3.881861597 | 0.00050999 | 1.95674868 | 3.29243717 |
| IDH2 | 3.872321279 | 6.895E-05 | 1.95319866 | 4.16146814 |
| CUEDC1 | 3.864254665 | 0.0006595 | 1.95019017 | 3.1807857 |
| LIN7A | 3.860913145 | 0.00636367 | 1.9489421 | 2.19629222 |
| CHTOP | 3.859392547 | 0.00021796 | 1.94837379 | 3.66162705 |
| ATG9A | 3.854323811 | 0.00091968 | 1.94647778 | 3.03636386 |
| KRT17 | 3.847162118 | 0.00334502 | 1.94379463 | 2.47560093 |
| MRPL50 | 3.846373713 | 0.00037303 | 1.94349894 | 3.42825303 |
| RAB11FIP1 | 3.84434646 | 0.00063739 | 1.94273836 | 3.19559167 |
| GNAO1 | 3.842106672 | 0.00042867 | 1.94189757 | 3.36788003 |
| LLGL1 | 3.838817309 | 0.00384621 | 1.9406619 | 2.41496645 |
| STOML2 | 3.834356704 | 0.00078077 | 1.93898455 | 3.10747938 |
| DBNL | 3.832880383 | 1.8903E-05 | 1.93842898 | 4.72346901 |
| PAK1IP1 | 3.830844412 | 0.0004358 | 1.93766243 | 3.36071076 |
| H1-5 | 3.829348394 | 0.0001384 | 1.93709892 | 3.85885147 |
| DDHD2 | 3.818903326 | 0.18816108 | 1.9331584 | 0.72547021 |

| | | | | |
|---|---|---|---|---|
| RASSF7 | 3.811967197 | 0.00067383 | 1.9305357 | 3.17144987 |
| FHL1 | 3.799271655 | 0.00010556 | 1.92572287 | 3.9765005 |
| NDUFS3 | 3.799181208 | 0.00050659 | 1.92568853 | 3.29534312 |
| H1-10 | 3.79512084 | 0.00066452 | 1.92414582 | 3.1774937 |
| TESC | 3.794867474 | 0.00056745 | 1.9240495 | 3.24607264 |
| FH | 3.794843828 | 0.00050648 | 1.92404051 | 3.29543571 |
| CBLL1 | 3.778043821 | 0.00010992 | 1.91763944 | 3.95893259 |
| IDE | 3.777089357 | 0.00144661 | 1.91727492 | 2.83964744 |
| FARP2 | 3.773926621 | 0.0013027 | 1.91606637 | 2.88515409 |
| RAB6A | 3.771710065 | 0.00148118 | 1.91521878 | 2.82939284 |
| MLF2 | 3.762514127 | 0.00183313 | 1.911697 | 2.73680597 |
| HELZ | 3.751576411 | 0.00077071 | 1.90749694 | 3.11311046 |
| DPP9 | 3.750828908 | 0.05506202 | 1.90720946 | 1.25914785 |
| LYAR | 3.749057683 | 0.00882239 | 1.90652802 | 2.05441367 |
| NOM1 | 3.743581593 | 4.9088E-05 | 1.9044192 | 4.30902597 |
| PAFAH1B3 | 3.741760501 | 0.00046166 | 1.90371722 | 3.33568099 |
| UBE2V2 | 3.734841289 | 0.00422932 | 1.90104694 | 2.37372946 |
| SPICE1 | 3.729052194 | 0.00827692 | 1.89880899 | 2.08213113 |
| AMOTL1 | 3.728927134 | 0.00075086 | 1.89876061 | 3.12443906 |
| MYO1E | 3.723302919 | 0.00234589 | 1.896583 | 2.62969259 |
| MAN1A2 | 3.720798596 | 0.00220951 | 1.8956123 | 2.65570378 |
| SLC7A5 | 3.720049629 | 0.00014179 | 1.89532187 | 3.8483406 |
| UNK | 3.716026083 | 0.0003478 | 1.89376063 | 3.45866866 |
| CBR1 | 3.713162302 | 1.2677E-05 | 1.89264838 | 4.89697531 |
| AP1G1 | 3.71267143 | 0.00098375 | 1.89245764 | 3.00711691 |
| ARL2 | 3.71084055 | 0.01353082 | 1.89174601 | 1.86867601 |
| YWHAZ | 3.704415322 | 0.0003178 | 1.88924586 | 3.49784416 |
| TP53BP2 | 3.701827761 | 5.9547E-05 | 1.88823777 | 4.2251381 |
| NHS | 3.697429392 | 1.2851E-05 | 1.8865226 | 4.89105094 |
| HCN4 | 3.695423742 | 0.00098363 | 1.8857398 | 3.00716987 |
| EPB41L3 | 3.691695049 | 0.00022376 | 1.88428338 | 3.65021369 |
| USP33 | 3.688671831 | 0.00166925 | 1.88310144 | 2.77747859 |
| SEC23IP | 3.687224846 | 0.00032562 | 1.88253539 | 3.48728936 |
| H1-2 | 3.677428703 | 0.00018551 | 1.87869737 | 3.73163763 |
| CDC42BPG | 3.673016115 | 0.00033703 | 1.87696523 | 3.47232654 |
| DVL3 | 3.672088919 | 0.00047511 | 1.87660099 | 3.32320597 |
| CDKL5 | 3.669535558 | 0.00111095 | 1.87559748 | 2.95430544 |
| ATP7A | 3.657592846 | 0.00036587 | 1.87089449 | 3.43667376 |
| AGFG2 | 3.656704743 | 0.0012858 | 1.87054414 | 2.89082788 |
| CNOT4 | 3.647006597 | 0.00059013 | 1.86671281 | 3.22904973 |
| PEA15 | 3.642803577 | 0.04601849 | 1.86504921 | 1.33706768 |
| TMEM63C | 3.6414459 | 0.2207283 | 1.86451141 | 0.65614198 |
| C2CD2L | 3.636953105 | 0.00071603 | 1.86273032 | 3.14506649 |
| HLCS | 3.634425403 | 0.00011437 | 1.86172729 | 3.94170073 |
| ERH | 3.630728917 | 0.00192634 | 1.86025922 | 2.71526811 |

| | | | | |
|---|---|---|---|---|
| MARK2 | 3.629565084 | 1.8384E-06 | 1.85979669 | 5.73555145 |
| MECOM | 3.629400956 | 0.00097017 | 1.85973145 | 3.01315294 |
| ASAP2 | 3.628578344 | 0.00021067 | 1.85940442 | 3.67639083 |
| NUFIP1 | 3.626562329 | 0.70701702 | 1.85860264 | 0.15057013 |
| MRPS27 | 3.606882377 | 0.00180936 | 1.85075238 | 2.74247408 |
| RRP1 | 3.60608862 | 3.706E-05 | 1.85043485 | 4.4310936 |
| SCP2 | 3.604812685 | 0.00017194 | 1.8499243 | 3.76461063 |
| APC | 3.588232612 | 0.00130072 | 1.84327342 | 2.88581501 |
| PDK1 | 3.583108607 | 0.0206011 | 1.84121177 | 1.68610954 |
| MRPS2 | 3.566544477 | 0.00080649 | 1.83452696 | 3.09340072 |
| AK4 | 3.54383034 | 0.00192001 | 1.82530954 | 2.716696 |
| PPFIBP2 | 3.539716989 | 0.06481021 | 1.82363402 | 1.18835657 |
| LETM1 | 3.538377181 | 0.00090953 | 1.82308784 | 3.04118488 |
| FYN | 3.537198433 | 0.00114546 | 1.82260716 | 2.94101876 |
| RAB29 | 3.535082573 | 0.00071158 | 1.82174391 | 3.14777783 |
| BET1L | 3.528141714 | 0.00923379 | 1.81890851 | 2.03461999 |
| ADH5 | 3.516882521 | 0.01193212 | 1.81429714 | 1.92328225 |
| PICK1 | 3.512807222 | 0.00287231 | 1.81262441 | 2.54176803 |
| XRN1 | 3.51204859 | 0.00019868 | 1.8123128 | 3.70184064 |
| NDUFS2 | 3.4946807 | 0.00177961 | 1.80516065 | 2.74967606 |
| CLIC1 | 3.493539081 | 0.00023402 | 1.80468928 | 3.63074842 |
| SIGIRR | 3.492214388 | 0.00468403 | 1.80414213 | 2.32938002 |
| PPWD1 | 3.490390125 | 0.00240701 | 1.8033883 | 2.61852179 |
| EIF4A2 | 3.48836681 | 0.00230394 | 1.80255175 | 2.63752976 |
| HNRNPD | 3.486201249 | 2.0233E-05 | 1.80165585 | 4.69393834 |
| EXOSC6 | 3.479444257 | 0.34917118 | 1.79885689 | 0.45696161 |
| ATP6V0A1 | 3.479032906 | 7.7954E-05 | 1.79868632 | 4.10816002 |
| SLC8A1 | 3.478837035 | 0.00047879 | 1.7986051 | 3.31985172 |
| TNFRSF21 | 3.473406511 | 0.00687817 | 1.79635127 | 2.16252706 |
| GGA3 | 3.470467643 | 0.00648883 | 1.79513008 | 2.18783344 |
| NRDC | 3.464729174 | 0.17010728 | 1.79274259 | 0.76927709 |
| PALS2 | 3.464288869 | 0.00159595 | 1.79255923 | 2.79698093 |
| RPL34 | 3.458755821 | 0.0002432 | 1.79025317 | 3.61404392 |
| SUSD6 | 3.458553136 | 0.0380714 | 1.79016862 | 1.41940117 |
| SH3GL2 | 3.456873051 | 0.00037898 | 1.78946762 | 3.42138781 |
| SPATA2 | 3.454771699 | 0.00011258 | 1.78859038 | 3.94855316 |
| RAP1B | 3.452009161 | 0.00234507 | 1.78743629 | 2.62984468 |
| AMOT | 3.443565703 | 0.00044512 | 1.7839032 | 3.35152184 |
| STAT1 | 3.435049397 | 0.00028647 | 1.78033085 | 3.54291597 |
| EFHD2 | 3.434556966 | 0.00016202 | 1.78012401 | 3.79044264 |
| PAFAH1B2 | 3.433336852 | 0.00061347 | 1.77961141 | 3.21220962 |
| NUDT1 | 3.426471622 | 0.00471609 | 1.77672374 | 2.32641757 |
| SUCLA2 | 3.42520338 | 0.00284922 | 1.77618965 | 2.5452744 |
| PPP1R13B | 3.42370368 | 0.00883105 | 1.77555784 | 2.05398749 |
| CLPB | 3.423323719 | 0.00041715 | 1.77539772 | 3.37971149 |

| | | | | |
|---|---|---|---|---|
| ACAP2 | 3.422562073 | 5.1409E-05 | 1.77507671 | 4.28896265 |
| TEX264 | 3.420976479 | 0.00027381 | 1.77440818 | 3.56254335 |
| RBSN | 3.410498412 | 0.00206262 | 1.76998259 | 2.6855808 |
| CCNB1 | 3.408438092 | 0.00528631 | 1.76911078 | 2.27684727 |
| MCCC2 | 3.404572886 | 7.8789E-05 | 1.76747382 | 4.10353296 |
| NDUFV2 | 3.392793697 | 0.00437272 | 1.76247371 | 2.35924784 |
| SH3PXD2B | 3.385566152 | 0.00018042 | 1.75939711 | 3.74370904 |
| CYBC1 | 3.38418319 | 0.0009441 | 1.75880767 | 3.02498121 |
| RAB11A | 3.376609694 | 7.6342E-05 | 1.75557543 | 4.11723464 |
| YIF1A | 3.371260454 | 0.00112729 | 1.75328809 | 2.94796381 |
| FDXR | 3.366441903 | 0.00410845 | 1.75122457 | 2.38632239 |
| ACAT2 | 3.363813231 | 0.00660046 | 1.75009761 | 2.18042565 |
| STAM | 3.363211964 | 0.00102503 | 1.74983971 | 2.98926345 |
| ATP5F1C | 3.363107463 | 0.00133754 | 1.74979488 | 2.87369296 |
| ZFYVE21 | 3.35346946 | 0.11893005 | 1.74565446 | 0.9247084 |
| FAF2 | 3.350407959 | 0.00258647 | 1.74433677 | 2.58729224 |
| ABCC5 | 3.343639908 | 0.00055237 | 1.74141949 | 3.25776766 |
| C1orf226 | 3.34092612 | 0.00414646 | 1.74024808 | 2.38232226 |
| KSR2 | 3.340834444 | 0.00099624 | 1.74020849 | 3.00163795 |
| DAGLA | 3.334923319 | 0.00687471 | 1.73765359 | 2.16274584 |
| PSMD9 | 3.32462479 | 0.00652837 | 1.73319153 | 2.18519509 |
| ROGDI | 3.323369069 | 0.00019694 | 1.73264652 | 3.70565786 |
| IMPA1 | 3.323138992 | 0.00050292 | 1.73254664 | 3.29850366 |
| SEC24C | 3.3199968 | 0.00040652 | 1.73118185 | 3.39091307 |
| AAK1 | 3.318729664 | 0.00024394 | 1.73063112 | 3.61271506 |
| NUP35 | 3.317009325 | 0.00024453 | 1.72988307 | 3.6116764 |
| RPL35A | 3.303758164 | 0.00090651 | 1.72410808 | 3.04262657 |
| NANP | 3.302403063 | 0.00039707 | 1.72351621 | 3.40113174 |
| CEP68 | 3.296909851 | 0.02969915 | 1.72111444 | 1.52725593 |
| EMD | 3.294579824 | 0.00020514 | 1.72009448 | 3.68794676 |
| ATP6V0D1 | 3.292452979 | 0.00272274 | 1.71916284 | 2.56499326 |
| CAPN7 | 3.290393161 | 0.01470606 | 1.71825998 | 1.83250365 |
| DLST | 3.275782564 | 0.00801761 | 1.7118396 | 2.09595497 |
| KNL1 | 3.266949955 | 0.00056632 | 1.70794435 | 3.24693846 |
| NME7 | 3.262608304 | 0.014113 | 1.70602579 | 1.85038066 |
| CSNK1G1 | 3.255115249 | 0.00057517 | 1.70270862 | 3.24020474 |
| NOP16 | 3.245874905 | 0.0006964 | 1.6986074 | 3.15714275 |
| RPN2 | 3.244353917 | 0.00300835 | 1.69793121 | 2.52167133 |
| ANGEL1 | 3.244333198 | 0.00054038 | 1.69792199 | 3.26730324 |
| CDV3 | 3.242341173 | 0.00024808 | 1.69703591 | 3.60540912 |
| FAF1 | 3.234711309 | 0.0002565 | 1.69363696 | 3.59091182 |
| RPS28 | 3.232642979 | 9.6541E-05 | 1.69271418 | 4.01528782 |
| TMX4 | 3.230964619 | 0.00128464 | 1.69196495 | 2.89121688 |
| CCDC85C | 3.216056728 | 0.0002099 | 1.68529285 | 3.67799628 |
| DHX37 | 3.210139037 | 0.00021202 | 1.68263578 | 3.67362572 |

| | | | | |
|---|---|---|---|---|
| TMEM214 | 3.208585763 | 0.00921558 | 1.68193755 | 2.03547715 |
| CLTA | 3.203411802 | 0.00348981 | 1.67960927 | 2.45719773 |
| BROX | 3.202294508 | 0.09861072 | 1.679106 | 1.00607588 |
| HSPD1 | 3.197859693 | 0.02494862 | 1.67710664 | 1.60295351 |
| PDXP | 3.179916548 | 0.02251991 | 1.6689889 | 1.6474333 |
| RPL18A | 3.177907498 | 8.3851E-05 | 1.66807713 | 4.07649157 |
| CDC42BPA | 3.169576866 | 0.00012603 | 1.66429026 | 3.89954113 |
| TOP1 | 3.169326192 | 0.00091649 | 1.66417615 | 3.0378727 |
| PTRH2 | 3.166938125 | 0.0027084 | 1.66308868 | 2.56728735 |
| PEX5 | 3.165894184 | 0.00657965 | 1.66261304 | 2.1817975 |
| PCMT1 | 3.165129743 | 0.00260556 | 1.66226464 | 2.58409973 |
| KDM2B | 3.16500237 | 0.00330806 | 1.66220658 | 2.48042682 |
| EIF3H | 3.155700573 | 0.00831252 | 1.65796032 | 2.08026729 |
| YWHAB | 3.15403651 | 0.00023343 | 1.65719936 | 3.63183711 |
| ACP1 | 3.149057939 | 0.0040729 | 1.6549203 | 2.3900963 |
| RAB27A | 3.144263105 | 0.00458701 | 1.65272194 | 2.33847076 |
| PYCR1 | 3.133409872 | 0.00131303 | 1.6477335 | 2.88172432 |
| RPS8 | 3.132800095 | 0.00041279 | 1.64745271 | 3.38426967 |
| ATP8B2 | 3.125170065 | 0.27248011 | 1.6439347 | 0.56466519 |
| SIK3 | 3.123710035 | 0.00438175 | 1.64326054 | 2.35835246 |
| SYT13 | 3.122697451 | 0.05498592 | 1.6427928 | 1.25974847 |
| RAB10 | 3.122483085 | 0.00918757 | 1.64269376 | 2.0367993 |
| RPL4 | 3.118621049 | 0.00013203 | 1.64090826 | 3.87934112 |
| TBC1D5 | 3.109631074 | 0.00031536 | 1.63674343 | 3.50118698 |
| PIN1 | 3.104497945 | 0.00113799 | 1.63435998 | 2.94386209 |
| PKP2 | 3.103669263 | 0.00012669 | 1.63397483 | 3.8972533 |
| RPL26 | 3.100657699 | 0.00076412 | 1.63257427 | 3.11684049 |
| CLTB | 3.084080567 | 0.02264577 | 1.62484045 | 1.64501288 |
| FAT1 | 3.082802141 | 0.00683652 | 1.6242423 | 2.16516497 |
| COASY | 3.082201168 | 0.0144327 | 1.62396103 | 1.84065234 |
| DCP1A | 3.075945823 | 0.00021063 | 1.62103009 | 3.67647154 |
| SLC22A5 | 3.069406592 | 0.00359128 | 1.61795977 | 2.44475024 |
| JMY | 3.061580733 | 0.00082964 | 1.61427673 | 3.08111244 |
| MVB12A | 3.055126082 | 0.00043513 | 1.61123192 | 3.36138066 |
| RAB33B | 3.051571406 | 0.0024878 | 1.60955235 | 2.60418401 |
| NAA50 | 3.050948203 | 6.1638E-05 | 1.60925769 | 4.21015407 |
| RALA | 3.050290266 | 0.00576404 | 1.60894654 | 2.23927328 |
| TBRG4 | 3.035391317 | 0.00074941 | 1.60188252 | 3.1252786 |
| AHCTF1 | 3.024138881 | 0.0004302 | 1.5965244 | 3.36633209 |
| ATP2B4 | 3.023628586 | 0.00029557 | 1.59628093 | 3.52933394 |
| ELOC | 3.019530382 | 0.01214079 | 1.59432419 | 1.91575297 |
| UCHL1 | 3.019254649 | 0.00105337 | 1.59419244 | 2.97741945 |
| OPA3 | 3.015516757 | 0.183728 | 1.59240525 | 0.73582465 |
| ATP5MK | 3.005910411 | 0.00027829 | 1.58780201 | 3.55550801 |
| GRK2 | 3.005103229 | 0.00015875 | 1.58741455 | 3.79928912 |

| | | | | |
|---|---|---|---|---|
| UBE2D2 | 3.004236358 | 0.05461191 | 1.58699832 | 1.26271265 |
| GOLGA3 | 3.001862941 | 0.00013906 | 1.58585811 | 3.85679073 |
| RPL3 | 3.001496234 | 0.00024077 | 1.58568186 | 3.61840124 |
| MYO6 | 2.999162884 | 0.01338657 | 1.58455988 | 1.87333078 |
| CEP57 | 2.99043272 | 6.0846E-05 | 1.58035426 | 4.21576471 |
| CAPZA2 | 2.988191123 | 0.0040902 | 1.57927242 | 2.38825518 |
| GRIP1 | 2.984033464 | 0.00587797 | 1.57726371 | 2.23077259 |
| SLITRK4 | 2.983203367 | 0.00857654 | 1.57686233 | 2.06668774 |
| CPLX1 | 2.982368318 | 0.00207703 | 1.57645844 | 2.68255645 |
| NCK1 | 2.981012175 | 0.00096323 | 1.57580227 | 3.01627117 |
| NSDHL | 2.979336697 | 0.00251073 | 1.57499117 | 2.60019951 |
| CEP250 | 2.977902213 | 5.7915E-05 | 1.57429638 | 4.23720748 |
| GMFB | 2.976858448 | 0.00519937 | 1.57379062 | 2.28404891 |
| DNAJB2 | 2.970602966 | 0.01998119 | 1.5707558 | 1.69937874 |
| RPL19 | 2.964623159 | 9.3944E-05 | 1.56784873 | 4.02713296 |
| HNRNPA0 | 2.963141746 | 0.00044739 | 1.56712764 | 3.34931445 |
| KIDINS220 | 2.962017392 | 5.1253E-05 | 1.56658011 | 4.29027785 |
| RNF149 | 2.958615842 | 0.02245809 | 1.56492238 | 1.64862717 |
| EMC2 | 2.957174376 | 0.02227069 | 1.56421932 | 1.6522664 |
| CELF1 | 2.955618616 | 0.00844341 | 1.56346012 | 2.07348199 |
| RAB5C | 2.953169239 | 0.00145769 | 1.56226404 | 2.83633429 |
| TIMM44 | 2.952952887 | 0.00027937 | 1.56215834 | 3.55381797 |
| NUP153 | 2.950157735 | 0.00015631 | 1.56079209 | 3.80600173 |
| DYNLRB2 | 2.941734982 | 0.00051779 | 1.55666728 | 3.28584918 |
| CLPX | 2.940171218 | 0.00037261 | 1.55590017 | 3.42874057 |
| NDUFS4 | 2.934016943 | 0.0015755 | 1.5528772 | 2.80258118 |
| LUC7L3 | 2.932630798 | 0.00116193 | 1.55219546 | 2.93481923 |
| SLC25A19 | 2.931958405 | 0.01976933 | 1.55186464 | 1.704008 |
| FCF1 | 2.930075255 | 0.00725504 | 1.55093772 | 2.13936004 |
| CDC42 | 2.923053664 | 0.0010487 | 1.54747632 | 2.97934706 |
| FTSJ3 | 2.919010956 | 0.00058439 | 1.54547963 | 3.233297 |
| PACSIN2 | 2.912589188 | 0.00036984 | 1.54230223 | 3.43198162 |
| EPB41L4A | 2.908800595 | 0.00012121 | 1.5404244 | 3.91645435 |
| DCXR | 2.904865157 | 0.01953477 | 1.5384712 | 1.70919177 |
| TAGLN2 | 2.894482859 | 0.00105878 | 1.53330561 | 2.97519552 |
| SECTM1 | 2.892199008 | 0.01993437 | 1.53216683 | 1.7003974 |
| PSMA1 | 2.891065192 | 0.00336386 | 1.53160114 | 2.47316273 |
| GNL2 | 2.890120084 | 0.00027047 | 1.53112944 | 3.56788786 |
| RPS3 | 2.889681822 | 0.00030531 | 1.53091065 | 3.51526142 |
| SFT2D3 | 2.888959439 | 0.00130452 | 1.53054995 | 2.88455004 |
| TMEM115 | 2.888380843 | 0.0022877 | 1.53026098 | 2.64060109 |
| CEP192 | 2.888258539 | 0.00057189 | 1.53019989 | 3.24268841 |
| PCNT | 2.884663343 | 0.00061349 | 1.52840296 | 3.21219195 |
| HSD17B4 | 2.880821131 | 0.00841743 | 1.52648009 | 2.0748207 |
| STK4 | 2.880531032 | 0.00026102 | 1.5263348 | 3.58332955 |

| | | | | |
|---|---|---|---|---|
| DNAH8 | 2.87633894 | 0.1974785 | 1.52423369 | 0.70448019 |
| HNRNPH3 | 2.876320035 | 0.00811038 | 1.52422421 | 2.09095881 |
| SFN | 2.871986811 | 0.00603246 | 1.52204912 | 2.21950524 |
| CENPV | 2.869122245 | 0.00013101 | 1.52060944 | 3.88269699 |
| CRKL | 2.867437115 | 0.00093185 | 1.51976185 | 3.03065457 |
| SUMO2 | 2.866126105 | 0.02563097 | 1.51910209 | 1.59123503 |
| GNL3 | 2.85943142 | 0.00296508 | 1.5157283 | 2.52796308 |
| IST1 | 2.859190425 | 0.00016429 | 1.51560671 | 3.78439632 |
| RAC1 | 2.854900837 | 0.00044318 | 1.51344064 | 3.35342292 |
| TUFM | 2.84139143 | 0.00059977 | 1.50659759 | 3.22201293 |
| FAM234A | 2.840182533 | 0.00393932 | 1.50598365 | 2.40457882 |
| LRCH1 | 2.834830754 | 0.00119131 | 1.50326261 | 2.92397702 |
| P3R3URF-PIK3R3 | 2.834676646 | 0.00761646 | 1.50318418 | 2.11824668 |
| GRPEL2 | 2.829850014 | 0.00070096 | 1.50072559 | 3.15430916 |
| LAGE3 | 2.829391152 | 0.00688843 | 1.50049164 | 2.16187978 |
| EPS8 | 2.828442782 | 0.00267973 | 1.50000799 | 2.57190827 |
| SNX30 | 2.826497723 | 0.00418305 | 1.49901553 | 2.37850676 |
| EPCAM | 2.825687786 | 0.01009562 | 1.49860207 | 1.99586688 |
| EFR3A | 2.820161525 | 0.00126863 | 1.4957778 | 2.89666435 |
| RPS13 | 2.819201328 | 0.08839127 | 1.49528651 | 1.05359062 |
| DCUN1D1 | 2.817714651 | 0.0084496 | 1.49452552 | 2.07316364 |
| STX12 | 2.817631075 | 0.00125418 | 1.49448273 | 2.9016407 |
| TRAP1 | 2.81703101 | 0.00023293 | 1.49417544 | 3.63277625 |
| EIF5A | 2.814330108 | 0.00140537 | 1.49279156 | 2.85220953 |
| PALS1 | 2.814282297 | 0.00133699 | 1.49276705 | 2.87387247 |
| CCP110 | 2.811717707 | 0.00040759 | 1.49145176 | 3.38978056 |
| DNAJB6 | 2.810716344 | 0.01212456 | 1.49093787 | 1.91633384 |
| MTHFD1 | 2.809155592 | 0.00127232 | 1.49013653 | 2.89540474 |
| SEC23A | 2.809123698 | 0.00704171 | 1.49012015 | 2.15232207 |
| TMSB4X | 2.808695234 | 0.19345654 | 1.48990009 | 0.71341658 |
| SYNM | 2.808637871 | 0.00125071 | 1.48987062 | 2.90284282 |
| NSFL1C | 2.807084427 | 0.02855679 | 1.48907246 | 1.54429058 |
| FHIP1A | 2.805261113 | 0.00195984 | 1.48813506 | 2.70777929 |
| USP13 | 2.805160188 | 0.00081548 | 1.48808316 | 3.088585 |
| YWHAG | 2.803641508 | 0.00455475 | 1.48730189 | 2.34153568 |
| ST7 | 2.803101384 | 0.01352478 | 1.48702393 | 1.86886983 |
| SPCS3 | 2.801886642 | 0.00110638 | 1.48639859 | 2.95609737 |
| PCGF6 | 2.801459371 | 0.00299303 | 1.48617857 | 2.52388832 |
| LRP8 | 2.797926225 | 0.94959084 | 1.48435792 | 0.02246348 |
| EPB41 | 2.797433104 | 0.00048005 | 1.48410363 | 3.318716 |
| QSOX2 | 2.787700664 | 0.0085919 | 1.47907566 | 2.0659108 |
| LNPEP | 2.785033062 | 0.00230682 | 1.47769445 | 2.63698585 |
| PGD | 2.780445052 | 0.0082307 | 1.47531583 | 2.08456328 |
| CAPZB | 2.780334466 | 0.00060658 | 1.47525845 | 3.21711407 |
| C2orf76 | 2.774929861 | 0.62035061 | 1.47245131 | 0.20736279 |

| | | | | |
|---|---|---|---|---|
| TBCA | 2.774927209 | 0.00116226 | 1.47244993 | 2.93469535 |
| ABL2 | 2.774478456 | 0.00088304 | 1.4722166 | 3.05401799 |
| MYBL2 | 2.772574904 | 0.10461768 | 1.47122644 | 0.98039491 |
| CEP120 | 2.772302686 | 0.00428743 | 1.47108478 | 2.36780336 |
| ARHGAP12 | 2.77000064 | 0.00076078 | 1.46988631 | 3.11874098 |
| RSBN1 | 2.761304421 | 0.00072148 | 1.46534995 | 3.14177831 |
| AP4M1 | 2.759179758 | 5.3326E-05 | 1.46423945 | 4.2730608 |
| VPS45 | 2.755862686 | 0.00503736 | 1.46250401 | 2.29779718 |
| PPP2CB | 2.755519994 | 0.00127558 | 1.4623246 | 2.89429377 |
| UBE2L3 | 2.749564001 | 0.0040412 | 1.45920287 | 2.39348931 |
| PAN2 | 2.749503298 | 0.00085238 | 1.45917102 | 3.06936612 |
| SYN1 | 2.744428947 | 0.00015845 | 1.45650599 | 3.80011354 |
| ATP5MF-PTCD1 | 2.740885127 | 0.00795414 | 1.45464186 | 2.09940657 |
| DAAM1 | 2.74054669 | 0.00073408 | 1.45446371 | 3.13425639 |
| TAF2 | 2.737801454 | 0.00292875 | 1.45301783 | 2.53331839 |
| EML2 | 2.737680742 | 0.00059495 | 1.45295421 | 3.22552234 |
| SLC39A3 | 2.730471623 | 0.05702147 | 1.44915016 | 1.24396156 |
| RDH10 | 2.730105343 | 0.003289 | 1.44895662 | 2.482936 |
| EPN1 | 2.726193971 | 0.00261113 | 1.44688822 | 2.58317158 |
| LRCH4 | 2.725764055 | 0.00764278 | 1.44666069 | 2.11674844 |
| BRIX1 | 2.718154871 | 0.00299694 | 1.44262766 | 2.52332205 |
| TAGLN3 | 2.714453915 | 0.00437052 | 1.44066199 | 2.3594667 |
| KCNQ4 | 2.713500537 | 0.03935462 | 1.44015519 | 1.40500432 |
| PCSK2 | 2.707204239 | 0.10815629 | 1.43680373 | 0.96594823 |
| HAX1 | 2.706161199 | 0.00051536 | 1.43624778 | 3.28788683 |
| CALB2 | 2.703382734 | 0.0257951 | 1.43476578 | 1.5884628 |
| AP2A1 | 2.702095612 | 0.00260593 | 1.43407872 | 2.58403689 |
| ECI2 | 2.697880286 | 0.12662945 | 1.43182633 | 0.89746527 |
| EDC3 | 2.693134121 | 0.00195638 | 1.42928608 | 2.70854681 |
| SBDS | 2.692537051 | 0.00558259 | 1.4289662 | 2.25316431 |
| GABARAPL2 | 2.689806254 | 0.01291989 | 1.42750226 | 1.88874128 |
| CHMP7 | 2.684982445 | 0.00678394 | 1.42491266 | 2.16851818 |
| KCMF1 | 2.683141295 | 0.00744991 | 1.42392303 | 2.12784918 |
| VPS13B | 2.679918133 | 0.01810009 | 1.42218893 | 1.74231924 |
| EIF4B | 2.679121746 | 0.00132486 | 1.42176014 | 2.87783037 |
| SHD | 2.673232037 | 0.04028306 | 1.41858507 | 1.39487749 |
| RPS14 | 2.673209939 | 0.00045005 | 1.41857314 | 3.34674401 |
| RAB2B | 2.671480704 | 0.00095856 | 1.4176396 | 3.01837881 |
| USP11 | 2.67130511 | 0.01422625 | 1.41754477 | 1.84690944 |
| NKAP | 2.660092651 | 0.01631734 | 1.4114765 | 1.78735077 |
| ATPAF2 | 2.657292167 | 0.00364414 | 1.40995686 | 2.43840488 |
| TMEM132A | 2.65711192 | 0.00352007 | 1.409859 | 2.45344886 |
| ALG1 | 2.656614787 | 0.01127016 | 1.40958905 | 1.94806982 |
| RPLP0 | 2.656509087 | 0.00690972 | 1.40953165 | 2.16053966 |
| PPAN-P2RY11 | 2.655863123 | 0.00117838 | 1.4091808 | 2.92871343 |

| | | | | |
|---|---|---|---|---|
| SHISAL2B | 2.655260826 | 6.7851E-05 | 1.40885358 | 4.16844442 |
| ACO1 | 2.652907035 | 0.00457096 | 1.40757412 | 2.33999263 |
| ZWILCH | 2.651723609 | 0.01545288 | 1.40693041 | 1.81099056 |
| TARDBP | 2.643331745 | 0.00010246 | 1.4023575 | 3.9894362 |
| RPSA | 2.640810148 | 0.00500738 | 1.40098059 | 2.30038918 |
| TRAPPC4 | 2.63966963 | 0.08847421 | 1.40035738 | 1.05318329 |
| RNF114 | 2.636281191 | 0.0164798 | 1.39850426 | 1.78304798 |
| GNG4 | 2.631041537 | 0.00164647 | 1.39563402 | 2.78344498 |
| SIPA1L2 | 2.628664172 | 0.01061478 | 1.39432984 | 1.97408903 |
| MRPS23 | 2.626842183 | 0.61919438 | 1.39332953 | 0.208173 |
| FBL | 2.625929712 | 0.00184794 | 1.3928283 | 2.7333119 |
| KAT14 | 2.625634393 | 0.00100066 | 1.39266604 | 2.99971241 |
| TOX2 | 2.625572789 | 0.09203194 | 1.39263219 | 1.03606143 |
| EIPR1 | 2.623855745 | 0.75211547 | 1.39168841 | 0.12371548 |
| LRCH3 | 2.623696239 | 0.02787073 | 1.3916007 | 1.55485171 |
| F11R | 2.618944147 | 0.0269274 | 1.38898529 | 1.56980565 |
| MISP | 2.6167622 | 0.00074494 | 1.38778282 | 3.12787937 |
| SNCG | 2.616617443 | 0.00036323 | 1.38770301 | 3.43982279 |
| CCT7 | 2.614640891 | 0.00143309 | 1.38661281 | 2.84372508 |
| ACOT7 | 2.613665636 | 0.00079341 | 1.38607459 | 3.10050277 |
| CSNK1A1 | 2.607050039 | 0.00479181 | 1.38241827 | 2.3195004 |
| DHCR24 | 2.606397797 | 0.00595218 | 1.38205729 | 2.22532428 |
| ALDOA | 2.602943483 | 0.0008511 | 1.38014399 | 3.07001774 |
| TFRC | 2.601290211 | 0.00075774 | 1.37922736 | 3.12047747 |
| TJAP1 | 2.599487312 | 0.00066267 | 1.37822711 | 3.17870341 |
| AP4B1 | 2.59892613 | 0.00067816 | 1.37791563 | 3.16866615 |
| SLC35B2 | 2.597538382 | 0.00263296 | 1.37714507 | 2.57955625 |
| ITCH | 2.596554467 | 0.00037949 | 1.37659849 | 3.42079612 |
| B3GAT3 | 2.589402575 | 0.03145638 | 1.37261928 | 1.50229124 |
| TFCP2 | 2.585583481 | 0.0028235 | 1.37048989 | 2.54921201 |
| PPP5C | 2.579424987 | 0.00061505 | 1.36704949 | 3.21108621 |
| GNA13 | 2.576968461 | 0.01469739 | 1.36567488 | 1.83275991 |
| SND1 | 2.573526724 | 0.00090998 | 1.36374676 | 3.04096764 |
| RNH1 | 2.572880853 | 0.00187853 | 1.36338465 | 2.72618106 |
| NYAP2 | 2.567894067 | 4.8552E-05 | 1.36058569 | 4.31379457 |
| ABI2 | 2.566646557 | 0.00446585 | 1.35988464 | 2.35009572 |
| IL17RA | 2.565565309 | 0.01390514 | 1.35927675 | 1.85682452 |
| TBC1D22A | 2.560762653 | 0.00068505 | 1.35657354 | 3.16427707 |
| PCNA | 2.553860725 | 0.00237221 | 1.35267985 | 2.62484721 |
| LLGL2 | 2.553468083 | 0.77196474 | 1.35245803 | 0.11240253 |
| CUX1 | 2.548938364 | 0.00021361 | 1.34989649 | 3.67038514 |
| GCDH | 2.547913328 | 0.02895125 | 1.3493162 | 1.53833263 |
| AK2 | 2.547538972 | 0.00878366 | 1.34910422 | 2.05632428 |
| DNAJA2 | 2.54690971 | 0.00022649 | 1.34874782 | 3.64495521 |
| TRIO | 2.542317696 | 0.00148925 | 1.34614433 | 2.82703271 |

| | | | | |
|---|---|---|---|---|
| ALK | 2.536593065 | 0.09446457 | 1.34289209 | 1.02473103 |
| NUDCD2 | 2.535033139 | 0.00291494 | 1.34200461 | 2.53537109 |
| SLC16A1 | 2.532389808 | 0.00108994 | 1.34049949 | 2.96259873 |
| TJP2 | 2.528502816 | 0.00032214 | 1.33828339 | 3.49195035 |
| LONP1 | 2.523545594 | 0.0023001 | 1.33545215 | 2.63825334 |
| PRKAR2A | 2.521210395 | 0.0008408 | 1.33411652 | 3.07530987 |
| ABI1 | 2.520140496 | 0.00056797 | 1.33350417 | 3.24567618 |
| FERMT2 | 2.519681862 | 0.00532081 | 1.33324159 | 2.27402223 |
| AFG3L2 | 2.519620816 | 0.0010873 | 1.33320664 | 2.96365096 |
| H2AFY | 2.518014807 | 0.00602204 | 1.33228677 | 2.2202562 |
| VDAC1 | 2.516620159 | 0.00229571 | 1.33148748 | 2.63908234 |
| KCND2 | 2.511313228 | 0.0005089 | 1.32844198 | 3.29336871 |
| MAP1A | 2.510848891 | 0.02251348 | 1.32817521 | 1.64755731 |
| RPS9 | 2.507368641 | 0.00068923 | 1.32617412 | 3.16163737 |
| PKP1 | 2.504396656 | 0.03880713 | 1.32446308 | 1.41108847 |
| HNRNPDL | 2.500671305 | 0.00456135 | 1.32231544 | 2.34090635 |
| RAB13 | 2.499558656 | 0.00195276 | 1.32167338 | 2.70935024 |
| GPD1L | 2.497776363 | 0.00130156 | 1.32064431 | 2.88553572 |
| VPS13C | 2.496340236 | 0.0016233 | 1.31981458 | 2.78960093 |
| PPP3CB | 2.493527107 | 0.3388449 | 1.31818789 | 0.46999904 |
| VIPAS39 | 2.49276912 | 0.00155722 | 1.31774927 | 2.80764875 |
| OXCT1 | 2.491617729 | 0.00164948 | 1.31708274 | 2.78265245 |
| BCAS3 | 2.488900694 | 0.00148315 | 1.31550867 | 2.828814 |
| HEATR5B | 2.488119755 | 0.00011313 | 1.31505593 | 3.94641785 |
| GANAB | 2.484176548 | 0.00541462 | 1.31276771 | 2.26643168 |
| VANGL2 | 2.483250315 | 0.00505142 | 1.31222969 | 2.29658694 |
| RPS16 | 2.4818163 | 0.00041272 | 1.31139633 | 3.38434114 |
| SFXN4 | 2.479710865 | 0.00150856 | 1.31017191 | 2.82143819 |
| PXN | 2.479224711 | 0.00119204 | 1.30988904 | 2.92370847 |
| CALCOCO2 | 2.475727464 | 0.08433607 | 1.30785251 | 1.07398665 |
| VPS26B | 2.467354824 | 0.0230822 | 1.3029652 | 1.63672272 |
| UQCRC2 | 2.454168578 | 0.00444547 | 1.29523435 | 2.35208271 |
| TXNL1 | 2.453080022 | 0.00074303 | 1.2945943 | 3.12899567 |
| RPL38 | 2.449465619 | 0.00358726 | 1.29246704 | 2.44523678 |
| WASF3 | 2.448968217 | 0.0256614 | 1.29217405 | 1.5907197 |
| SRSF3 | 2.446577957 | 0.02314577 | 1.29076525 | 1.63552835 |
| PSMD8 | 2.445197381 | 0.0022905 | 1.28995093 | 2.64006969 |
| MIB1 | 2.441975612 | 0.00442712 | 1.28804879 | 2.35387914 |
| HNRNPL | 2.437386539 | 0.00024083 | 1.28533506 | 3.61828079 |
| STXBP1 | 2.425020673 | 0.00285914 | 1.27799705 | 2.54376399 |
| MAGED2 | 2.421341788 | 0.01015362 | 1.27580674 | 1.9933791 |
| RPL22L1 | 2.419609633 | 1.7831E-05 | 1.27477431 | 4.74881939 |
| LIMCH1 | 2.419398159 | 0.00041512 | 1.27464821 | 3.38183093 |
| SLC30A7 | 2.416924423 | 0.0201191 | 1.27317236 | 1.69639155 |
| RPL6 | 2.41326834 | 0.00030782 | 1.27098834 | 3.51169748 |

| | | | | |
|---|---|---|---|---|
| SRP9 | 2.402855023 | 0.00951247 | 1.26474961 | 2.02170657 |
| DENR | 2.402555192 | 0.00026486 | 1.26456957 | 3.57698623 |
| COL1A1 | 2.399143903 | 0.11552105 | 1.26251969 | 0.93733886 |
| CEP152 | 2.398180471 | 0.0826666 | 1.26194023 | 1.08266991 |
| UGDH | 2.396679269 | 0.00276423 | 1.26103686 | 2.558426 |
| ALDH7A1 | 2.395230643 | 0.00571812 | 1.26016458 | 2.24274692 |
| NUP88 | 2.387581347 | 0.03666229 | 1.25554989 | 1.43578038 |
| EWSR1 | 2.387065826 | 0.00032874 | 1.25523835 | 3.48314423 |
| MBNL1 | 2.386922655 | 0.02467879 | 1.25515182 | 1.60767612 |
| UQCRC1 | 2.384145908 | 0.01247575 | 1.25347253 | 1.90393343 |
| HBA1; HBA2 | 2.382439094 | 0.00917875 | 1.25243933 | 2.03721662 |
| ADK | 2.38222411 | 0.85139798 | 1.25230914 | 0.06986738 |
| RPL31 | 2.37938451 | 0.00123019 | 1.25058843 | 2.91002652 |
| EXOC3 | 2.378326318 | 0.01368084 | 1.24994667 | 1.86388709 |
| TSFM | 2.37782223 | 0.01331293 | 1.24964086 | 1.87572628 |
| DLD | 2.374219275 | 0.006023 | 1.24745318 | 2.22018696 |
| PPP2R2D | 2.373713406 | 0.03476835 | 1.24714576 | 1.45881594 |
| MAP4K3 | 2.37111812 | 0.00304877 | 1.24556753 | 2.51587472 |
| PLEKHH3 | 2.369501447 | 0.53520755 | 1.24458354 | 0.27147777 |
| FDPS | 2.369478221 | 0.00386269 | 1.2445694 | 2.41311058 |
| RIPK1 | 2.367633343 | 0.00160241 | 1.24344568 | 2.79522607 |
| CNNM2 | 2.365958004 | 0.02085958 | 1.24242447 | 1.68069453 |
| NIBAN2 | 2.365916305 | 0.01612976 | 1.24239904 | 1.79237202 |
| TJP3 | 2.357121678 | 0.00146743 | 1.23702623 | 2.83344211 |
| MAST1 | 2.355897531 | 0.00257396 | 1.23627679 | 2.58939752 |
| TBC1D13 | 2.353443931 | 0.0434895 | 1.23477348 | 1.36161558 |
| TRMT10C | 2.345822169 | 0.00217971 | 1.23009365 | 2.66160144 |
| OTUB1 | 2.345316878 | 0.00168378 | 1.22978286 | 2.7737158 |
| DBT | 2.344688611 | 0.01050788 | 1.22939634 | 1.97848507 |
| SLC7A14 | 2.34270323 | 0.00034655 | 1.22817421 | 3.46022914 |
| PTK7 | 2.340602785 | 0.00144523 | 1.22688012 | 2.84006196 |
| CSNK1E | 2.339873589 | 0.00265559 | 1.22643059 | 2.57583818 |
| NDUFA4 | 2.339414088 | 0.01867529 | 1.22614725 | 1.72873257 |
| SLC3A2 | 2.336323043 | 0.0006013 | 1.22423977 | 3.22090739 |
| MPHOSPH9 | 2.336300699 | 0.00394496 | 1.22422597 | 2.40395689 |
| PDLIM1 | 2.335224834 | 0.0027238 | 1.22356146 | 2.564824 |
| TAB1 | 2.333314949 | 0.00117659 | 1.22238105 | 2.92937566 |
| ATP6V1E1 | 2.333032232 | 0.00327938 | 1.22220624 | 2.4842086 |
| RAD18 | 2.331354941 | 0.00090165 | 1.22116867 | 3.04496049 |
| LSS | 2.330262477 | 0.02128289 | 1.22049247 | 1.67196943 |
| EHBP1L1 | 2.327991843 | 0.00039748 | 1.219086 | 3.40068673 |
| ATP2B3 | 2.32668089 | 0.0116026 | 1.21827335 | 1.93544469 |
| ACTR1A | 2.324940846 | 0.00062607 | 1.21719401 | 3.20337506 |
| TRA2B | 2.322716398 | 0.01508734 | 1.21581301 | 1.8213873 |
| TDRD3 | 2.322312111 | 0.00388998 | 1.21556188 | 2.41005291 |

| | | | | |
|---|---|---|---|---|
| GNB2 | 2.320893079 | 0.00370897 | 1.21468006 | 2.43074693 |
| CLDND1 | 2.320386796 | 0.74980046 | 1.21436531 | 0.1250543 |
| SCYL1 | 2.318665482 | 0.03201534 | 1.2132947 | 1.4946419 |
| PDAP1 | 2.318539299 | 0.00112507 | 1.21321618 | 2.94882172 |
| TRMT2B | 2.318151665 | 0.27270262 | 1.21297496 | 0.56431069 |
| EMC1 | 2.31739395 | 0.01397772 | 1.21250332 | 1.85456351 |
| MRPL27 | 2.313168096 | 0.33211182 | 1.20987011 | 0.47871567 |
| UCHL5 | 2.312721978 | 0.02058724 | 1.20959184 | 1.68640197 |
| SAR1A | 2.311680683 | 0.02417017 | 1.20894213 | 1.61672025 |
| RAP2B | 2.310645555 | 0.01600749 | 1.20829597 | 1.79567677 |
| EIF2B5 | 2.308672976 | 0.03309436 | 1.20706383 | 1.48024601 |
| TOMM40 | 2.305757918 | 0.01206157 | 1.20524105 | 1.91859632 |
| APPL1 | 2.300180877 | 0.00135781 | 1.20174731 | 2.86716041 |
| SOWAHC | 2.29277194 | 0.00819775 | 1.19709286 | 2.0863053 |
| TANK | 2.292222868 | 0.00468721 | 1.19674732 | 2.32908579 |
| BRK1 | 2.289862443 | 0.00596081 | 1.19526094 | 2.22469486 |
| SCD5 | 2.288982052 | 0.03910855 | 1.19470615 | 1.40772832 |
| TTLL12 | 2.288657345 | 0.02060212 | 1.19450148 | 1.68608815 |
| WDR46 | 2.284396362 | 0.00405924 | 1.19181299 | 2.3915551 |
| RBM4 | 2.283291666 | 0.00136233 | 1.19111516 | 2.86571918 |
| ARHGEF10L | 2.281473896 | 0.03411436 | 1.18996615 | 1.46706274 |
| NKAPD1 | 2.28083992 | 0.00356109 | 1.18956519 | 2.44841672 |
| PIP4P1 | 2.276464947 | 0.00063935 | 1.18679524 | 3.19426153 |
| REPS1 | 2.275703031 | 0.00301975 | 1.1863123 | 2.52002851 |
| NEURL4 | 2.27556658 | 0.09537776 | 1.1862258 | 1.02055288 |
| TSC1 | 2.272194611 | 0.00092621 | 1.18408641 | 3.03329228 |
| USP25 | 2.271815329 | 0.0383589 | 1.18384557 | 1.41613389 |
| SLC25A29 | 2.270817181 | 0.00157714 | 1.18321156 | 2.80213084 |
| DLAT | 2.270779947 | 0.00523944 | 1.18318791 | 2.28071476 |
| OIP5 | 2.269611403 | 0.00757456 | 1.1824453 | 2.12064266 |
| DENND1A | 2.265654531 | 2.8981E-07 | 1.17992789 | 6.53788188 |
| ST13 | 2.264312211 | 0.00719043 | 1.1790729 | 2.14324504 |
| VAMP7 | 2.263698212 | 0.00932845 | 1.17868164 | 2.03019047 |
| H4C1; H4C11; H4C | 2.26298696 | 0.00362446 | 1.17822827 | 2.44075619 |
| NIP7 | 2.262349181 | 0.03641479 | 1.17782162 | 1.43872222 |
| PCLO | 2.260449302 | 0.00152233 | 1.17660956 | 2.81749052 |
| RPL32 | 2.260377128 | 0.00317361 | 1.1765635 | 2.49844707 |
| SLC25A3 | 2.257125633 | 0.00408581 | 1.17448672 | 2.38872173 |
| RBM34 | 2.255601965 | 0.00504079 | 1.1735125 | 2.29750132 |
| TMEM41B | 2.254240623 | 0.00055612 | 1.17264152 | 3.25483511 |
| HLA-E | 2.252098983 | 0.00345919 | 1.17127024 | 2.46102506 |
| VPS13A | 2.251870607 | 0.00112042 | 1.17112393 | 2.95061881 |
| XPR1 | 2.250619914 | 0.08322514 | 1.17032243 | 1.07974545 |
| RPN1 | 2.25033969 | 0.00065129 | 1.17014279 | 3.18622553 |
| NCKAP1 | 2.248594243 | 0.00044723 | 1.16902335 | 3.34946818 |

| | | | | |
|---|---|---|---|---|
| AP3S1 | 2.247034773 | 0.00304062 | 1.16802245 | 2.51703789 |
| RPS2 | 2.245037973 | 0.10929599 | 1.16673985 | 0.96139579 |
| RABEP1 | 2.244293499 | 0.00640071 | 1.16626136 | 2.19377193 |
| UBE2M | 2.24268664 | 0.00046077 | 1.16522805 | 3.33651205 |
| SLC37A4 | 2.237267482 | 0.03450916 | 1.16173775 | 1.46206557 |
| PFN1 | 2.236823476 | 0.01045872 | 1.16145141 | 1.9805214 |
| ANKRD40 | 2.227113942 | 0.17824357 | 1.15517537 | 0.74898612 |
| APEX1 | 2.225469551 | 0.00541882 | 1.15410976 | 2.26609512 |
| MKI67 | 2.2241361 | 0.01315879 | 1.15324507 | 1.88078393 |
| SLC20A1 | 2.223138588 | 0.01172069 | 1.15259789 | 1.93104693 |
| GAK | 2.216465972 | 0.00140842 | 1.14826121 | 2.85126789 |
| ABHD17B | 2.214090824 | 0.03405385 | 1.1467144 | 1.46783374 |
| MAP3K7 | 2.21296338 | 0.00182029 | 1.14597958 | 2.73986001 |
| LDHB | 2.212941559 | 0.00096176 | 1.14596535 | 3.01693136 |
| ARF4 | 2.212563157 | 0.00503422 | 1.14571864 | 2.29806756 |
| MAPK1 | 2.211781229 | 0.00275359 | 1.14520869 | 2.56010051 |
| AGPS | 2.208911599 | 0.08096622 | 1.14333568 | 1.09169613 |
| ERAL1 | 2.205042804 | 0.03905615 | 1.14080666 | 1.40831055 |
| TIMM29 | 2.202983969 | 0.04798041 | 1.139459 | 1.31893603 |
| RPL13 | 2.199652448 | 0.00048296 | 1.13727559 | 3.31609033 |
| ATP5F1A | 2.198412779 | 0.00679138 | 1.1364623 | 2.16804216 |
| RPL7A | 2.198302958 | 0.00021638 | 1.13639022 | 3.66478426 |
| RBCK1 | 2.193444611 | 0.00658447 | 1.13319828 | 2.18147945 |
| CNP | 2.192647783 | 0.02335868 | 1.13267408 | 1.63155164 |
| ARF3 | 2.187656208 | 0.0078712 | 1.12938603 | 2.10395925 |
| RAN | 2.18408444 | 0.00234433 | 1.12702863 | 2.6299815 |
| GSTP1 | 2.18168165 | 0.02400131 | 1.1254406 | 1.61976499 |
| SH3GL1 | 2.178408145 | 0.00888414 | 1.12327428 | 2.05138484 |
| TCEB3 | 2.176418537 | 0.00869525 | 1.12195602 | 2.06071787 |
| FGD6 | 2.175864182 | 0.04179208 | 1.12158851 | 1.37890599 |
| YBX1 | 2.175023778 | 0.00032622 | 1.12103117 | 3.48649024 |
| CENPV | 2.174665426 | 0.00434499 | 1.12079346 | 2.36201104 |
| GFM2 | 2.173387237 | 0.12222191 | 1.11994525 | 0.91285094 |
| ST3GAL5 | 2.170212447 | 0.00067611 | 1.11783628 | 3.16997962 |
| CCDC88A | 2.168657893 | 0.00189076 | 1.11680248 | 2.72336451 |
| LCMT2 | 2.163328074 | 0.74048018 | 1.11325247 | 0.13048656 |
| ACAD9 | 2.163059979 | 0.02625528 | 1.11307367 | 1.58078339 |
| RPL35 | 2.159148528 | 0.00070172 | 1.11046249 | 3.15383386 |
| SPAG5 | 2.1571242 | 0.00367197 | 1.10910924 | 2.4351007 |
| ARL6 | 2.155870518 | 0.07398323 | 1.10827053 | 1.13086673 |
| DDX6 | 2.155746316 | 0.01112184 | 1.10818741 | 1.95382326 |
| RPL27A | 2.15419138 | 0.04749577 | 1.10714643 | 1.32334508 |
| ALG13 | 2.153766958 | 0.01120607 | 1.10686216 | 1.95054655 |
| SLC9A6 | 2.151037288 | 0.0456665 | 1.10503253 | 1.34040232 |
| YTHDC2 | 2.148273753 | 0.01099019 | 1.10317785 | 1.95899481 |

| | | | | |
|---|---|---|---|---|
| ANK3 | 2.147615168 | 0.01268152 | 1.1027355 | 1.89682865 |
| CUL3 | 2.146589681 | 0.02063827 | 1.10204645 | 1.68532662 |
| NUDT5 | 2.144256128 | 0.00216093 | 1.10047724 | 2.66535877 |
| HSPA14 | 2.143416189 | 0.05355607 | 1.09991201 | 1.27119128 |
| SEH1L | 2.143321511 | 0.00684589 | 1.09984828 | 2.16457017 |
| PAFAH1B1 | 2.143093269 | 0.01508315 | 1.09969464 | 1.82150788 |
| SLC38A2 | 2.139310119 | 0.01053188 | 1.09714563 | 1.97749418 |
| LRRC59 | 2.13779763 | 0.00566259 | 1.09612529 | 2.24698499 |
| RHEB | 2.137298831 | 0.00266807 | 1.09578864 | 2.57380352 |
| PLEKHA7 | 2.13586901 | 0.00267751 | 1.09482317 | 2.5722691 |
| ENO1 | 2.12860052 | 0.0008505 | 1.08990522 | 3.07032769 |
| CCT4 | 2.127931525 | 0.0061124 | 1.08945173 | 2.21378802 |
| RARS2 | 2.127674314 | 0.02898698 | 1.08927733 | 1.53779698 |
| PHGDH | 2.126614032 | 0.00510822 | 1.08855822 | 2.29173026 |
| NDUFA6 | 2.126490285 | 0.002046 | 1.08847426 | 2.6890938 |
| WWP2 | 2.123913237 | 0.00592375 | 1.08672483 | 2.22740327 |
| TBCEL | 2.121222339 | 0.01155183 | 1.08489585 | 1.9373492 |
| PUSL1 | 2.115944521 | 0.01397283 | 1.0813018 | 1.85471569 |
| SEC24D | 2.115566629 | 0.00076985 | 1.08104412 | 3.11359536 |
| MRPS5 | 2.115267652 | 0.00188385 | 1.08084022 | 2.72495389 |
| SCD | 2.107086147 | 0.0028174 | 1.0752493 | 2.55015124 |
| REPS2 | 2.106369312 | 0.01503711 | 1.07475841 | 1.82283575 |
| PRPS1 | 2.105557367 | 0.03666469 | 1.07420218 | 1.43575197 |
| ESS2 | 2.105263448 | 0.00173522 | 1.07400078 | 2.76064655 |
| PPP2R2A | 2.101963209 | 0.00176232 | 1.07173742 | 2.75391586 |
| VPS8 | 2.101139718 | 0.01528664 | 1.0711721 | 1.81568792 |
| ATP2B1 | 2.099945726 | 0.00105595 | 1.07035204 | 2.97635461 |
| RSL1D1 | 2.098179472 | 0.00078003 | 1.06913809 | 3.10788919 |
| QRICH1 | 2.095537028 | 0.00477973 | 1.06732001 | 2.32059624 |
| SLC7A1 | 2.09447193 | 0.0668315 | 1.06658655 | 1.17501878 |
| CYRIB | 2.092398883 | 0.0207855 | 1.0651579 | 1.68223961 |
| TP53I11 | 2.090790025 | 0.04476758 | 1.06404818 | 1.34903641 |
| ZNF420 | 2.088887923 | 0.18853541 | 1.06273509 | 0.72460707 |
| KCNA4 | 2.088537296 | 0.01994092 | 1.06249291 | 1.70025486 |
| P4HB | 2.088387014 | 0.00906493 | 1.06238909 | 2.0426357 |
| ZDHHC13 | 2.086981763 | 0.07483926 | 1.06141799 | 1.1258705 |
| SAE1 | 2.086525711 | 0.00209102 | 1.0611027 | 2.67964105 |
| OPHN1 | 2.083735982 | 0.00087632 | 1.05917249 | 3.0573378 |
| KCNK13 | 2.07963683 | 0.02775535 | 1.05633161 | 1.55665336 |
| HSP90B1 | 2.076322101 | 0.00252626 | 1.05403027 | 2.59752142 |
| PLEKHO2 | 2.072105224 | 0.77237432 | 1.05109727 | 0.11217217 |
| USP24 | 2.07194846 | 0.04117379 | 1.05098812 | 1.38537918 |
| ALDH9A1 | 2.071565807 | 0.04572889 | 1.05072165 | 1.33980935 |
| EIF4A1 | 2.071505937 | 0.00206299 | 1.05067996 | 2.68550258 |
| DCTN1 | 2.070589193 | 0.00094053 | 1.05004135 | 3.02662739 |

| | | | | |
|---|---|---|---|---|
| TBC1D25 | 2.07004832 | 0.00077279 | 1.04966444 | 3.1119358 |
| RPS26 | 2.069613041 | 0.00511861 | 1.04936105 | 2.29084754 |
| ZNF598 | 2.068406757 | 0.00712656 | 1.04851992 | 2.1471202 |
| RPL27 | 2.067456559 | 0.00347372 | 1.04785702 | 2.45920537 |
| DNAJB11 | 2.067336969 | 0.00663085 | 1.04777356 | 2.17843047 |
| C17orf75 | 2.066504366 | 0.00149635 | 1.04719241 | 2.82496658 |
| PHB1 | 2.058636924 | 0.00178531 | 1.04168941 | 2.74828585 |
| SNX27 | 2.058019826 | 0.01047044 | 1.04125688 | 1.98003513 |
| NPEPPS | 2.057234042 | 0.013657 | 1.04070593 | 1.8646447 |
| PI4K2A | 2.055968568 | 0.01099445 | 1.03981821 | 1.95882659 |
| RNPEP | 2.053599591 | 0.65828592 | 1.03815491 | 0.18158543 |
| LSM14B | 2.051255917 | 0.00587258 | 1.03650749 | 2.23117126 |
| PFDN1 | 2.051004534 | 0.0009354 | 1.03633068 | 3.02900156 |
| PAGR1 | 2.049784451 | 0.13395242 | 1.03547221 | 0.87304944 |
| DOCK7 | 2.049269846 | 0.00356835 | 1.03510997 | 2.44753235 |
| HNRNPA3 | 2.047191436 | 0.00050419 | 1.03364602 | 3.29740312 |
| TDP2 | 2.045995628 | 0.00038761 | 1.03280306 | 3.41160813 |
| SLC25A10 | 2.040666643 | 0.16221722 | 1.02904053 | 0.78990304 |
| FCHO1 | 2.035744256 | 0.08990105 | 1.02555633 | 1.04623522 |
| SLC25A6 | 2.035495015 | 0.0065285 | 1.02537969 | 2.18518648 |
| RPS21 | 2.031809638 | 0.0533803 | 1.02276524 | 1.27261896 |
| SAP25 | 2.02765753 | 0.05799667 | 1.019814 | 1.23659697 |
| DICER1 | 2.027387435 | 0.10624636 | 1.01962182 | 0.97368592 |
| DKFZp686H10254 | 2.025996674 | 0.07655104 | 1.01863181 | 1.11604893 |
| KCNH7 | 2.025770839 | 0.09244415 | 1.01847098 | 1.03412057 |
| PPP3CA | 2.025096475 | 0.00514608 | 1.01799064 | 2.28852306 |
| MYD88 | 2.024822333 | 0.40718602 | 1.01779533 | 0.39020714 |
| INTS3 | 2.023923082 | 0.70051455 | 1.01715446 | 0.15458284 |
| PEAK1 | 2.022538729 | 0.00585899 | 1.01616733 | 2.23217745 |
| DNM2 | 2.022157519 | 0.00408093 | 1.01589538 | 2.38924037 |
| ACVR2A | 2.015212354 | 0.1633114 | 1.01093187 | 0.78698351 |
| LARS2 | 2.014384725 | 0.04269976 | 1.01033925 | 1.36957457 |
| JAK1 | 2.01160721 | 0.67883261 | 1.00834863 | 0.1682373 |
| AP3S2 | 2.010108455 | 0.00446769 | 1.00727334 | 2.34991684 |
| POLR2H | 2.009799852 | 0.00876185 | 1.00705184 | 2.05740413 |
| MED19 | 2.007098796 | 0.23614232 | 1.00511163 | 0.62682617 |
| ASCC3 | 2.004955151 | 0.0670661 | 1.00356997 | 1.17349698 |
| GATD3B | 2.003804017 | 0.02230956 | 1.00274141 | 1.65150908 |
| ARF5 | 2.003100551 | 0.00054841 | 1.00223484 | 3.26089076 |
| GNAQ | 2.002387539 | 0.29557892 | 1.00172122 | 0.52932654 |
| PALLD | 1.999312966 | 0.00045974 | 0.99950432 | 3.33748428 |
| DUSP23 | 1.998008189 | 0.04292778 | 0.9985625 | 1.36726161 |
| TRMT5 | 1.99678014 | 0.00420886 | 0.99767549 | 2.3758356 |
| C11orf98 | 1.99635144 | 0.00120889 | 0.99736572 | 2.91761468 |
| SEC62 | 1.994660866 | 0.00095625 | 0.99614348 | 3.01942639 |

| | | | | |
|---|---|---|---|---|
| STT3A | 1.985661246 | 0.01305134 | 0.98961952 | 1.88434474 |
| HTATIP2 | 1.984972317 | 0.84477212 | 0.98911889 | 0.07326043 |
| UPF1 | 1.982685605 | 0.00119672 | 0.98745593 | 2.92200773 |
| DHFR | 1.980417224 | 0.03593113 | 0.9858044 | 1.4445291 |
| DUSP26 | 1.978167131 | 0.84400287 | 0.98416432 | 0.07365608 |
| KHDRBS2 | 1.977330384 | 0.5381175 | 0.98355395 | 0.26912289 |
| HSDL2 | 1.973213818 | 0.00471328 | 0.9805473 | 2.32667682 |
| LRRC47 | 1.973187071 | 0.01604465 | 0.98052774 | 1.79466976 |
| EIF4A3 | 1.969274868 | 0.00211219 | 0.97766449 | 2.6752669 |
| PALM | 1.96855384 | 0.00136226 | 0.97713617 | 2.86573993 |
| MAP2K3 | 1.967339244 | 0.00840649 | 0.97624575 | 2.07538517 |
| XIAP | 1.964488814 | 0.00577954 | 0.97415395 | 2.23810641 |
| NHERF1 | 1.963823544 | 2.0052E-05 | 0.9736653 | 4.69784916 |
| RNF123 | 1.961535396 | 0.02934596 | 0.97198337 | 1.53245162 |
| TNIK | 1.961081745 | 0.01060348 | 0.97164967 | 1.97455148 |
| CPNE3 | 1.960471845 | 0.03531064 | 0.97120092 | 1.45209445 |
| NRBF2 | 1.958272509 | 0.03712397 | 0.96958154 | 1.43034559 |
| PM20D2 | 1.955770594 | 0.00822697 | 0.96773716 | 2.08476026 |
| CD47 | 1.949024356 | 0.00653706 | 0.96275212 | 2.18461746 |
| NMD3 | 1.944767333 | 0.62187664 | 0.95959757 | 0.20629576 |
| XKR7 | 1.942763865 | 0.0649964 | 0.95811056 | 1.18711071 |
| SCOC | 1.942725993 | 0.26429333 | 0.95808243 | 0.5779138 |
| AXIN1 | 1.942721351 | 0.97788612 | 0.95807899 | 0.00971172 |
| CKB | 1.942677056 | 0.0075092 | 0.95804609 | 2.12440615 |
| IRAK1 | 1.942428234 | 0.03307055 | 0.9578613 | 1.48055855 |
| PRODH | 1.941760237 | 0.00164646 | 0.95736507 | 2.78344818 |
| COQ8A | 1.941040796 | 0.01960349 | 0.95683044 | 1.70766653 |
| WDR1 | 1.939790945 | 0.00395701 | 0.95590118 | 2.40263288 |
| PPP2R1A | 1.937744553 | 0.00297038 | 0.9543784 | 2.52718869 |
| PPP4R4 | 1.936949336 | 0.54778333 | 0.95378622 | 0.26139119 |
| MTX1 | 1.935757144 | 0.10643709 | 0.95289797 | 0.97290701 |
| ARL6IP4 | 1.935540294 | 0.03863661 | 0.95273634 | 1.41300102 |
| RBM4B | 1.932370824 | 0.00056484 | 0.95037198 | 3.24807551 |
| TXN | 1.930369645 | 0.09803181 | 0.94887713 | 1.00863299 |
| GNAI1 | 1.928031223 | 0.07056682 | 0.94712841 | 1.15139947 |
| GNB1 | 1.92785326 | 0.01473182 | 0.94699524 | 1.83174355 |
| SEPTIN8 | 1.927135914 | 3.0591E-05 | 0.94645832 | 4.51440699 |
| LGALS7; LGALS7B | 1.926639961 | 0.00555981 | 0.94608699 | 2.25494022 |
| ARRB2 | 1.923139361 | 0.62023977 | 0.94346331 | 0.20744039 |
| VPS35 | 1.922211568 | 0.03059658 | 0.94276714 | 1.51432712 |
| BCLAF3 | 1.92196754 | 0.08095094 | 0.94258397 | 1.09177811 |
| RPL14 | 1.920022584 | 0.00301594 | 0.94112328 | 2.52057758 |
| NPLOC4 | 1.919521927 | 0.09558112 | 0.94074704 | 1.01962788 |
| CASC3 | 1.918974863 | 0.02566946 | 0.94033581 | 1.5905833 |
| WASL | 1.917134221 | 2.9526E-05 | 0.93895135 | 4.52980216 |

| | | | | |
|---|---|---|---|---|
| PSD3 | 1.917124957 | 0.02129787 | 0.93894437 | 1.67166387 |
| DACT1 | 1.915962454 | 0.02271064 | 0.93806929 | 1.64377072 |
| FKBP4 | 1.914822921 | 0.04098518 | 0.93721098 | 1.38737315 |
| RPL22 | 1.911268917 | 0.00445561 | 0.93453078 | 2.3510933 |
| FBXO3 | 1.907269931 | 0.01387184 | 0.93150904 | 1.85786603 |
| GK3 | 1.906810903 | 0.01184779 | 0.93116178 | 1.92636277 |
| DEPDC7 | 1.904225069 | 0.0271026 | 0.92920401 | 1.56698907 |
| OTULIN | 1.904091127 | 0.61212838 | 0.92910253 | 0.21315748 |
| RPL5 | 1.9037244 | 0.00092903 | 0.92882464 | 3.03196805 |
| PACS1 | 1.90343373 | 0.00481542 | 0.92860434 | 2.3173658 |
| DAZAP1 | 1.896848533 | 0.02095247 | 0.92360448 | 1.67876487 |
| CBX4 | 1.895998508 | 0.35072232 | 0.92295783 | 0.4550366 |
| OTULINL | 1.894411556 | 0.05975393 | 0.92174979 | 1.22363354 |
| LMF2 | 1.893606776 | 0.05516059 | 0.92113677 | 1.25837109 |
| EIF4E | 1.893019513 | 0.0140393 | 0.92068928 | 1.85265469 |
| RUFY1 | 1.891846287 | 0.00780288 | 0.91979487 | 2.10774491 |
| STUB1 | 1.887040354 | 0.06765113 | 0.91612528 | 1.16972492 |
| NDFIP2 | 1.886860257 | 0.02778206 | 0.91598758 | 1.55623554 |
| ELAVL3 | 1.886705809 | 0.00204247 | 0.91586948 | 2.68984519 |
| METTL15 | 1.883954415 | 0.09893643 | 0.91376406 | 1.00464377 |
| IMPDH2 | 1.883560519 | 0.00879207 | 0.91346239 | 2.05590892 |
| PTPRK | 1.882348275 | 0.01154685 | 0.91253358 | 1.93753658 |
| ELAVL2 | 1.882212338 | 0.01424165 | 0.91242939 | 1.84643964 |
| RPL12 | 1.880467772 | 0.00572439 | 0.91109158 | 2.24227047 |
| SLC20A1 | 1.879944364 | 0.0019167 | 0.91068997 | 2.71744479 |
| MAGI1 | 1.87948664 | 0.01387375 | 0.91033866 | 1.85780622 |
| AP2B1 | 1.878922546 | 0.02299285 | 0.9099056 | 1.63840721 |
| STK38L | 1.87693976 | 0.07039681 | 0.90838235 | 1.15244704 |
| NEDD1 | 1.873515488 | 0.00511052 | 0.9057479 | 2.29153483 |
| TMEM97 | 1.862644482 | 0.00381133 | 0.89735234 | 2.4189233 |
| TOMM22 | 1.861227929 | 0.01137363 | 0.89625474 | 1.94410081 |
| SMG9 | 1.860710065 | 0.00100886 | 0.89585327 | 2.99616819 |
| PRKACA | 1.857037419 | 0.02253561 | 0.89300289 | 1.64713071 |
| WIPF2 | 1.85444938 | 0.00040351 | 0.89099089 | 3.39414957 |
| ANKFY1 | 1.853483451 | 0.12169448 | 0.89023923 | 0.91472912 |
| SRSF9 | 1.85293071 | 0.00722042 | 0.88980893 | 2.14143734 |
| NCAM1 | 1.852775354 | 0.0220929 | 0.88968797 | 1.65574722 |
| ARFGAP2 | 1.851465394 | 0.0080094 | 0.88866758 | 2.09639987 |
| CLIC4 | 1.846047013 | 0.06481164 | 0.88443929 | 1.18834696 |
| CAPN1 | 1.84580723 | 0.05250005 | 0.88425189 | 1.27984031 |
| APLP2 | 1.844945305 | 0.03034277 | 0.88357805 | 1.51794478 |
| IP6K1 | 1.844138623 | 0.0285995 | 0.88294711 | 1.5436415 |
| DNAJB1 | 1.842660452 | 0.00214958 | 0.88179025 | 2.66764738 |
| CLMN | 1.84163122 | 0.01208441 | 0.8809842 | 1.91777446 |
| POM121 | 1.838935077 | 0.00899367 | 0.87887055 | 2.0460631 |

| | | | | |
|---|---|---|---|---|
| HNRNPA2B1 | 1.838514175 | 0.01500732 | 0.8785403 | 1.8236968 |
| LBR | 1.838334781 | 0.02772885 | 0.87839952 | 1.55706812 |
| SEC61A1 | 1.837707416 | 0.00798208 | 0.87790709 | 2.09788388 |
| VDAC3 | 1.835751181 | 0.04836035 | 0.87637053 | 1.31551056 |
| PPP1R15B | 1.833086501 | 0.03709881 | 0.87427487 | 1.43064001 |
| TRA2A | 1.832031262 | 0.00426764 | 0.87344412 | 2.36981223 |
| CHMP2B | 1.831885917 | 0.00732953 | 0.87332966 | 2.13492379 |
| TPM3 | 1.829821299 | 0.03340137 | 0.87170276 | 1.47623568 |
| FAR1 | 1.82829069 | 0.03320217 | 0.87049547 | 1.47883348 |
| APOOL | 1.82740387 | 0.00292361 | 0.86979552 | 2.53408109 |
| RUNDC3A | 1.822713607 | 0.07223191 | 0.8660879 | 1.14127091 |
| WDR45B | 1.822532015 | 0.04817522 | 0.86594416 | 1.31717632 |
| RPL28 | 1.822456316 | 0.00801957 | 0.86588423 | 2.09584871 |
| GSK3A | 1.821534519 | 0.03526561 | 0.86515434 | 1.45264865 |
| PRPS2 | 1.820803339 | 0.0477562 | 0.86457511 | 1.32097021 |
| PDHX | 1.819356176 | 0.00121188 | 0.86342801 | 2.91654105 |
| C8orf33 | 1.817448217 | 0.07508068 | 0.86191426 | 1.12447183 |
| SEC22B | 1.815540214 | 0.00012429 | 0.86039889 | 3.90556073 |
| OTUD6B | 1.813851606 | 0.02727683 | 0.85905643 | 1.56420617 |
| MTG1 | 1.811936192 | 0.01179977 | 0.85753215 | 1.92812636 |
| PPP1CC | 1.809495493 | 0.00260791 | 0.85558751 | 2.58370812 |
| HSPA12A | 1.80780201 | 0.31249054 | 0.85423668 | 0.50516313 |
| PDK3 | 1.807484709 | 0.9406198 | 0.85398344 | 0.02658588 |
| NEMP1 | 1.807296966 | 0.0037562 | 0.85383358 | 2.42525181 |
| PPFIBP1 | 1.804737645 | 0.00171555 | 0.85178913 | 2.76559683 |
| MRPS7 | 1.804733526 | 0.00302531 | 0.85178584 | 2.51922942 |
| TNS1 | 1.804513646 | 0.00875621 | 0.85161005 | 2.05768381 |
| FARSA | 1.803461508 | 0.09254919 | 0.85076863 | 1.03362736 |
| CARMIL2 | 1.803164632 | 0.05370226 | 0.85053112 | 1.27000743 |
| STAU2 | 1.801801352 | 0.00119499 | 0.84943996 | 2.92263733 |
| TTC7B | 1.79653335 | 0.05443293 | 0.84521572 | 1.26413825 |
| MTMR4 | 1.796359055 | 0.02901929 | 0.84507574 | 1.53731317 |
| AKR7A2 | 1.793133955 | 0.05909731 | 0.84248327 | 1.22843227 |
| SNRPA1 | 1.790255414 | 0.07269859 | 0.84016543 | 1.13847402 |
| SNX17 | 1.787969699 | 0.01983545 | 0.83832229 | 1.70255798 |
| HNRNPR | 1.786788386 | 0.00478152 | 0.83736878 | 2.32043446 |
| CDC20 | 1.785443768 | 0.06110675 | 0.8362827 | 1.2139108 |
| SGTA | 1.784044241 | 0.09168686 | 0.83515139 | 1.03769291 |
| RHOB | 1.77788285 | 0.60838045 | 0.83016026 | 0.21582475 |
| FAM83H | 1.776786608 | 0.06587221 | 0.82927042 | 1.18129775 |
| ARHGAP1 | 1.776525163 | 0.00620179 | 0.82905812 | 2.207483 |
| TPP2 | 1.776495768 | 0.48379912 | 0.82903425 | 0.31533493 |
| NCOR2 | 1.775832 | 0.01088152 | 0.8284951 | 1.96331063 |
| KPNA4 | 1.775018118 | 0.22422994 | 0.82783375 | 0.6493064 |
| WDR26 | 1.774019928 | 0.0194216 | 0.82702222 | 1.7117151 |

| | | | | |
|---|---|---|---|---|
| PA2G4 | 1.773229684 | 0.01101746 | 0.82637942 | 1.95791852 |
| RPL24 | 1.772264612 | 0.00593941 | 0.82559402 | 2.22625661 |
| RAB7A | 1.770544364 | 0.02236743 | 0.82419299 | 1.65038399 |
| RTCB | 1.767970123 | 0.01040076 | 0.82209389 | 1.98293477 |
| MAP4K5 | 1.767064989 | 0.15706392 | 0.8213551 | 0.80392357 |
| SAC3D1 | 1.764893418 | 0.01008973 | 0.81958106 | 1.99612035 |
| VAT1 | 1.76238726 | 0.71693223 | 0.81753097 | 0.1445219 |
| SLCO3A1 | 1.760656606 | 0.0668609 | 0.81611356 | 1.17482777 |
| ASAP1 | 1.759979501 | 0.01121351 | 0.81555863 | 1.95025852 |
| RPLP2 | 1.758082517 | 0.00390669 | 0.81400279 | 2.40819063 |
| DHCR7 | 1.756041212 | 0.00646384 | 0.8123267 | 2.18950956 |
| RPS27L | 1.755864654 | 0.03055805 | 0.81218164 | 1.51487443 |
| WASH2P | 1.75555598 | 0.19865755 | 0.811928 | 0.70189493 |
| MRI1 | 1.746165924 | 0.00973546 | 0.80419065 | 2.0116436 |
| HSDL1 | 1.745020424 | 0.00277829 | 0.80324392 | 2.55622199 |
| PTGES3 | 1.743129979 | 0.35964473 | 0.80168015 | 0.4441263 |
| DNM1 | 1.741906517 | 0.05551311 | 0.8006672 | 1.25560447 |
| DPM1 | 1.740394809 | 0.00122533 | 0.79941462 | 2.91174603 |
| SNX12 | 1.73925204 | 0.02264902 | 0.79846701 | 1.64495062 |
| NEDD4 | 1.737854716 | 0.12989668 | 0.79730748 | 0.88640196 |
| COTL1 | 1.737640492 | 0.83519846 | 0.79712963 | 0.07821032 |
| SYN3 | 1.735323172 | 0.22052829 | 0.79520436 | 0.65653569 |
| SLC25A12 | 1.734055702 | 0.02221254 | 0.79415024 | 1.65340168 |
| BLVRA | 1.733883528 | 0.00399602 | 0.79400699 | 2.39837277 |
| ARPC3 | 1.732567382 | 0.04902336 | 0.79291146 | 1.30959697 |
| REEP4 | 1.731951282 | 0.74809301 | 0.79239835 | 0.1260444 |
| SLC25A11 | 1.730240737 | 0.0049117 | 0.79097278 | 2.30876823 |
| FASTKD5 | 1.72949597 | 0.03934233 | 0.79035165 | 1.40513997 |
| LCOR | 1.728936516 | 0.00467705 | 0.7898849 | 2.33002797 |
| GCN1 | 1.728140965 | 0.03350717 | 0.7892209 | 1.47486225 |
| GMDS | 1.725878743 | 0.00762872 | 0.78733111 | 2.11754838 |
| SRSF7 | 1.725174215 | 0.04588521 | 0.78674206 | 1.3383273 |
| ATXN10 | 1.724059469 | 0.01385148 | 0.78580954 | 1.85850374 |
| MAPRE1 | 1.721871881 | 0.00032025 | 0.7839778 | 3.49451434 |
| PLEKHG4B | 1.720838722 | 0.03432919 | 0.78311189 | 1.4643364 |
| COG1 | 1.720014748 | 0.04002395 | 0.78242093 | 1.39768007 |
| DTYMK | 1.717313791 | 0.00283513 | 0.78015368 | 2.54742711 |
| RPL11 | 1.717112563 | 0.00819913 | 0.77998462 | 2.08623223 |
| EEPD1 | 1.716824842 | 0.10438739 | 0.77974286 | 0.98135197 |
| PIGG | 1.716621552 | 0.00431002 | 0.77957202 | 2.36552066 |
| ELAVL1 | 1.714055209 | 7.9171E-05 | 0.77741358 | 4.10143418 |
| VAPA | 1.713510664 | 0.02256843 | 0.77695517 | 1.64649861 |
| CSRP2 | 1.711923396 | 0.00248237 | 0.77561815 | 2.60513364 |
| IRS1 | 1.711502341 | 0.00986716 | 0.77526327 | 2.00580794 |
| TMX3 | 1.709439421 | 0.00471444 | 0.7735233 | 2.32657023 |

| | | | | |
|---|---|---|---|---|
| SYT11 | 1.705414935 | 0.12528087 | 0.7701228 | 0.90211524 |
| MOV10 | 1.704362966 | 0.03207694 | 0.76923261 | 1.49380708 |
| PDCL | 1.703654587 | 0.48527165 | 0.76863286 | 0.31401508 |
| HDAC2 | 1.70270899 | 0.03051316 | 0.76783188 | 1.51551284 |
| RAB18 | 1.702254763 | 0.09087189 | 0.76744697 | 1.04157043 |
| SLC7A6OS | 1.701166952 | 0.11064431 | 0.76652473 | 0.95607091 |
| MRPL16 | 1.700917967 | 0.14503407 | 0.76631356 | 0.83852996 |
| COG2 | 1.700414903 | 0.15809252 | 0.76588681 | 0.80108869 |
| CYFIP2 | 1.699041616 | 0.01252735 | 0.76472119 | 1.90214091 |
| PFDN4 | 1.694497764 | 0.06985861 | 0.76085773 | 1.15578003 |
| PEX6 | 1.694103928 | 0.00249303 | 0.76052238 | 2.60327302 |
| IGF2BP3 | 1.693962765 | 0.01358494 | 0.76040216 | 1.86694224 |
| RALBP1 | 1.693205827 | 0.00104171 | 0.75975736 | 2.98225415 |
| ANAPC10 | 1.691782127 | 0.57077654 | 0.75854379 | 0.24353388 |
| BOLA2; BOLA2B | 1.690479418 | 0.01041149 | 0.75743245 | 1.98248695 |
| AP1B1 | 1.687467057 | 0.18116726 | 0.75485934 | 0.74192028 |
| RELT | 1.686937184 | 0.00422707 | 0.75440625 | 2.37396072 |
| HSPA1B | 1.686702732 | 0.00086185 | 0.75420573 | 3.06456621 |
| NFAT5 | 1.686431353 | 0.02816161 | 0.75397359 | 1.55034257 |
| GNAI2 | 1.685358232 | 0.06210821 | 0.75305528 | 1.20685101 |
| LPCAT3 | 1.68515831 | 0.02284649 | 0.75288413 | 1.64118053 |
| VANGL1 | 1.684630335 | 0.20690734 | 0.75243205 | 0.68422411 |
| SHE | 1.682893767 | 0.05125189 | 0.75094411 | 1.29029009 |
| SLC12A2 | 1.681322263 | 0.00970679 | 0.74959628 | 2.01292421 |
| CEP41 | 1.680279065 | 0.01728965 | 0.74870086 | 1.76221381 |
| GPD2 | 1.680217273 | 0.0174343 | 0.7486478 | 1.75859554 |
| INO80 | 1.677745968 | 0.02530418 | 0.74652429 | 1.59680778 |
| TLE5 | 1.674707628 | 0.41007793 | 0.74390925 | 0.38713361 |
| ESYT2 | 1.674190282 | 0.00813733 | 0.74346351 | 2.08951807 |
| OXSR1 | 1.673717574 | 0.17066083 | 0.74305611 | 0.76786615 |
| NBEA | 1.670066768 | 0.02568867 | 0.73990578 | 1.59025844 |
| TCF20 | 1.669671139 | 0.02835006 | 0.73956398 | 1.54744608 |
| PRKAG1 | 1.668812811 | 0.05847735 | 0.73882214 | 1.23301228 |
| GNB5 | 1.667976925 | 0.05619872 | 0.73809933 | 1.25027358 |
| WDR6 | 1.666091645 | 0.01470892 | 0.73646776 | 1.83241927 |
| NCKAP5L | 1.661790132 | 0.02497802 | 0.73273819 | 1.60244197 |
| ABCB10 | 1.66024023 | 0.13240898 | 0.73139201 | 0.87808255 |
| CDC37L1 | 1.659695319 | 0.77361117 | 0.73091842 | 0.11147727 |
| PSMD12 | 1.65902511 | 0.00542619 | 0.73033572 | 2.26550503 |
| TAX1BP1 | 1.657832527 | 0.01772969 | 0.72929827 | 1.75129898 |
| CRNKL1 | 1.654470831 | 0.66550975 | 0.72636986 | 0.17684558 |
| NECAP1 | 1.653486137 | 0.01820843 | 0.72551095 | 1.73972755 |
| ZFP90 | 1.652994128 | 0.27132689 | 0.7250816 | 0.56650716 |
| COQ5 | 1.652661721 | 0.17281571 | 0.72479145 | 0.76241677 |
| WASHC4 | 1.651257677 | 0.00538331 | 0.72356527 | 2.26895054 |

| | | | | |
|---|---|---|---|---|
| HSD17B7 | 1.650262124 | 0.05290448 | 0.7226952 | 1.27650754 |
| RAB33A | 1.650118479 | 0.02063316 | 0.72256961 | 1.68543433 |
| FAM161A | 1.649708386 | 0.19015154 | 0.72221103 | 0.72090014 |
| RASA1 | 1.648057264 | 0.67639797 | 0.72076637 | 0.1697977 |
| MRPS10 | 1.645566267 | 0.78538232 | 0.71858412 | 0.10491888 |
| CANX | 1.641341905 | 0.13483578 | 0.7148758 | 0.87019484 |
| PSMD14 | 1.639681221 | 0.01545178 | 0.71341536 | 1.81102155 |
| NAPA | 1.637122355 | 0.0495005 | 0.71116215 | 1.30539045 |
| ASPSCR1 | 1.63624257 | 0.10663849 | 0.71038664 | 0.97208601 |
| KCNN2 | 1.635163239 | 0.14010039 | 0.70943467 | 0.85356066 |
| VPS37C | 1.634527866 | 0.05322826 | 0.70887397 | 1.27385775 |
| BSDC1 | 1.632849184 | 0.01341938 | 0.70739154 | 1.87226749 |
| PPP1CB | 1.630566963 | 0.00104876 | 0.70537369 | 2.97932349 |
| HACD3 | 1.630533401 | 0.15574962 | 0.70534399 | 0.807573 |
| ITM2C | 1.62940135 | 0.42938847 | 0.70434201 | 0.36714962 |
| RALGAPA2 | 1.627982107 | 0.02011291 | 0.70308484 | 1.69652511 |
| FABP5 | 1.625922069 | 0.05011735 | 0.70125811 | 1.30001188 |
| IARS2 | 1.625129512 | 0.00365033 | 0.7005547 | 2.43766824 |
| SLC25A22 | 1.624870614 | 0.00722818 | 0.70032484 | 2.14097118 |
| TMED8 | 1.624400017 | 0.00677697 | 0.69990695 | 2.16896441 |
| EIF2S1 | 1.623596427 | 0.00184652 | 0.69919307 | 2.7336467 |
| OAT | 1.621724973 | 0.06481989 | 0.69752918 | 1.18829171 |
| RICTOR | 1.621070595 | 0.00214221 | 0.69694692 | 2.6691385 |
| SEPTIN2 | 1.620664108 | 0.09323155 | 0.69658512 | 1.03043708 |
| EIF4H | 1.61834695 | 0.00243663 | 0.69452093 | 2.61321108 |
| AKTIP | 1.618250386 | 0.02234229 | 0.69443485 | 1.65087227 |
| FAM83G | 1.616326842 | 0.02708287 | 0.69271896 | 1.56730529 |
| CDS2 | 1.616114496 | 0.30433532 | 0.69252941 | 0.51664764 |
| HYCC1 | 1.614368626 | 0.08239069 | 0.69097004 | 1.08412186 |
| GAL3ST1 | 1.613806229 | 0.08892243 | 0.69046736 | 1.05098867 |
| CTU1 | 1.611560552 | 0.12856557 | 0.6884584 | 0.8908753 |
| RTN3 | 1.608935033 | 0.01332367 | 0.68610607 | 1.87537619 |
| ATG2B | 1.606275836 | 0.04335088 | 0.68371966 | 1.36300212 |
| TMEM30A | 1.604386794 | 0.08441794 | 0.682022 | 1.07356527 |
| CHCHD3 | 1.60200723 | 0.00645945 | 0.67988066 | 2.18980451 |
| NNT | 1.600681544 | 0.0307975 | 0.67868631 | 1.51148448 |
| GOLGB1 | 1.600369843 | 0.04778996 | 0.67840535 | 1.32066331 |
| UBP1 | 1.599672756 | 0.0369709 | 0.6777768 | 1.43213993 |
| PTPRA | 1.598987495 | 0.03181912 | 0.67715866 | 1.49731179 |
| ASNS | 1.595524613 | 0.32793926 | 0.67403086 | 0.48420659 |
| CCDC43 | 1.595318563 | 0.05469857 | 0.67384454 | 1.26202406 |
| BAIAP2 | 1.594460165 | 0.00087275 | 0.67306805 | 3.05911261 |
| PHYHIPL | 1.590534728 | 0.00675334 | 0.66951187 | 2.17048155 |
| TBL2 | 1.590291777 | 0.04486992 | 0.66929149 | 1.34804475 |
| NT5DC2 | 1.589964386 | 0.37100827 | 0.66899445 | 0.43061641 |

| | | | | |
|---|---|---|---|---|
| NOP53 | 1.58930529 | 0.57365362 | 0.66839628 | 0.24135026 |
| SYNCRIP | 1.588471886 | 0.02453336 | 0.66763956 | 1.61024292 |
| PTPMT1 | 1.588266814 | 0.24072213 | 0.66745329 | 0.61848399 |
| SLC25A1 | 1.586990493 | 0.01948732 | 0.66629349 | 1.71024783 |
| TMEM201 | 1.586615513 | 0.02386855 | 0.66595256 | 1.62217393 |
| PTDSS1 | 1.585147945 | 0.18672549 | 0.6646175 | 0.72879638 |
| WDR89 | 1.582475018 | 0.08542086 | 0.66218272 | 1.06843604 |
| WDR48 | 1.580385706 | 0.01982278 | 0.6602767 | 1.70283551 |
| HMGA1 | 1.578257202 | 0.13208396 | 0.65833233 | 0.87914992 |
| BCL2L2-PABPN1 | 1.575964809 | 0.62292114 | 0.65623532 | 0.20556693 |
| WARS1 | 1.5752265 | 0.01748214 | 0.65555929 | 1.7574054 |
| NUDT16L1 | 1.574377644 | 0.0107068 | 0.65478164 | 1.9703403 |
| TNKS1BP1 | 1.572813756 | 0.131837 | 0.65334784 | 0.87996268 |
| TRAFD1 | 1.572377206 | 0.11288947 | 0.65294735 | 0.94734658 |
| DNAJC13 | 1.569797335 | 0.02504721 | 0.65057832 | 1.60124073 |
| PI4KA | 1.567496765 | 0.02653594 | 0.64846247 | 1.57616553 |
| CRYBG1 | 1.566836811 | 0.02141043 | 0.64785493 | 1.66937464 |
| NETO2 | 1.565917714 | 0.01726734 | 0.6470084 | 1.76277461 |
| TTC7A | 1.563406877 | 0.89779084 | 0.64469329 | 0.04682483 |
| MAP2K1 | 1.561598328 | 0.0748207 | 0.64302341 | 1.12597822 |
| ELFN2 | 1.561519169 | 0.02782245 | 0.64295028 | 1.55560468 |
| RRP7A | 1.560574871 | 0.52918705 | 0.64207757 | 0.27639079 |
| ANP32A | 1.559611117 | 0.46294222 | 0.64118634 | 0.33447321 |
| GOSR2 | 1.557648824 | 0.07003978 | 0.63937001 | 1.15465524 |
| CUL5 | 1.556833302 | 0.04939529 | 0.63861448 | 1.30631445 |
| R3HCC1L | 1.555935755 | 0.83994659 | 0.63778249 | 0.07574833 |
| TGFBR1 | 1.555800557 | 0.05737711 | 0.63765713 | 1.24126132 |
| NHSL1 | 1.555634306 | 0.01698705 | 0.63750296 | 1.7698821 |
| RRAGC | 1.555128205 | 0.11815135 | 0.63703352 | 0.92756131 |
| LDHA | 1.552976029 | 0.83812598 | 0.63503556 | 0.0766907 |
| LMNB2 | 1.552965189 | 0.00165854 | 0.63502549 | 2.78027437 |
| AP2A2 | 1.552405577 | 0.01067412 | 0.63450552 | 1.97166795 |
| OTUD7B | 1.552296404 | 0.51851275 | 0.63440406 | 0.28524056 |
| PHB2 | 1.55171107 | 0.00456237 | 0.63385995 | 2.34080997 |
| CCT6A | 1.550045721 | 0.00729672 | 0.63231077 | 2.13687207 |
| BRD9 | 1.549657742 | 0.01986643 | 0.63194962 | 1.70188025 |
| NEDD4L | 1.54910198 | 0.06757847 | 0.63143212 | 1.17019167 |
| PRDX5 | 1.54901853 | 0.30333649 | 0.6313544 | 0.51807534 |
| DDX19B | 1.54746913 | 0.30038252 | 0.62991063 | 0.52232534 |
| PIGT | 1.547422437 | 0.01401109 | 0.6298671 | 1.85352806 |
| TM9SF4 | 1.542870609 | 0.35756315 | 0.62561708 | 0.44664724 |
| CTDSP2 | 1.542009618 | 0.24001655 | 0.62481176 | 0.61975882 |
| MTMR7 | 1.541120748 | 0.18213535 | 0.6239799 | 0.73960575 |
| RPGRIP1L | 1.540957708 | 0.00668107 | 0.62382727 | 2.17515422 |
| GNAI3 | 1.539512641 | 0.04244896 | 0.62247371 | 1.37213291 |

| | | | | |
|---|---|---|---|---|
| SASS6 | 1.539068768 | 0.03008135 | 0.6220577 | 1.52170261 |
| SNAP47 | 1.537446713 | 0.02998873 | 0.62053641 | 1.52304186 |
| DNAJC11 | 1.537134779 | 0.01636306 | 0.62024367 | 1.78613541 |
| PANK4 | 1.5369297 | 0.39194244 | 0.62005118 | 0.4067777 |
| TPD52L2 | 1.536488709 | 0.17086038 | 0.61963717 | 0.76735862 |
| SCGN | 1.53608413 | 0.02025559 | 0.61925723 | 1.69345517 |
| CLEC16A | 1.534192469 | 0.01668444 | 0.61747948 | 1.77768831 |
| GON7 | 1.533501391 | 0.01689578 | 0.61682947 | 1.77222177 |
| SECISBP2 | 1.533231494 | 0.09101511 | 0.61657554 | 1.04088651 |
| SORD | 1.530644552 | 0.03650536 | 0.6141393 | 1.43764338 |
| RBBP9 | 1.530430837 | 0.27886311 | 0.61393785 | 0.55460894 |
| CC2D1A | 1.528424494 | 0.00441479 | 0.61204528 | 2.35508987 |
| ADSS2 | 1.519831487 | 0.07828212 | 0.60391137 | 1.10633745 |
| POLR2M | 1.518944723 | 0.68794763 | 0.60306937 | 0.16244462 |
| CHEK1 | 1.51820677 | 0.05001186 | 0.60236829 | 1.30092702 |
| YIPF3 | 1.517933523 | 0.13454702 | 0.60210861 | 0.87112593 |
| TUBB8 | 1.517749932 | 0.0724991 | 0.60193411 | 1.13966736 |
| TRAPPC2L | 1.517518147 | 0.12919306 | 0.60171377 | 0.88876082 |
| RAE1 | 1.51665568 | 0.03007662 | 0.60089359 | 1.5217709 |
| UBA1 | 1.513975848 | 0.0271527 | 0.59834219 | 1.56618694 |
| DENND6A | 1.513817165 | 0.62403333 | 0.59819097 | 0.20479221 |
| WDR45 | 1.513363003 | 0.06235236 | 0.59775808 | 1.20514707 |
| RRM1 | 1.513334707 | 0.28268869 | 0.59773111 | 0.54869157 |
| PIGS | 1.511740156 | 0.15400238 | 0.59621018 | 0.81247256 |
| STON2 | 1.509833992 | 0.24820067 | 0.59438993 | 0.60519704 |
| SLC43A3 | 1.508422507 | 0.87237969 | 0.59304058 | 0.05929445 |
| ARL6IP5 | 1.506902168 | 0.29056306 | 0.59158576 | 0.53675959 |
| PAPPA2 | 1.506774626 | 0.36432078 | 0.59146364 | 0.43851605 |
| HSPA2 | 1.506356029 | 0.00636019 | 0.59106279 | 2.19652957 |
| RPS7 | 1.506203862 | 0.04225958 | 0.59091705 | 1.3740748 |
| RPS11 | 1.506110246 | 0.03821411 | 0.59082738 | 1.41777625 |
| NOC3L | 1.505904721 | 0.02192204 | 0.59063049 | 1.65911904 |
| GIGYF1 | 1.505565457 | 0.12476779 | 0.59030543 | 0.90389753 |
| TXNDC5 | 1.504231879 | 0.1366641 | 0.58902698 | 0.86434557 |
| TRIM26 | 1.504019612 | 0.94221058 | 0.58882338 | 0.02585202 |
| GET3 | 1.503554029 | 0.10029327 | 0.58837671 | 0.99872823 |
| GPAT4 | 1.503192376 | 0.07112 | 0.58802965 | 1.14800824 |
| PAIP1 | 1.502319949 | 0.1540339 | 0.5871921 | 0.81238369 |
| NFIC | 1.501310923 | 0.73883092 | 0.58622279 | 0.13145494 |
| MFN2 | 1.498877261 | 0.52848353 | 0.58388225 | 0.27696854 |
| PAWR | 1.498812749 | 0.05310786 | 0.58382015 | 1.2748412 |
| RNASEH1 | 1.498599591 | 0.14560466 | 0.58361496 | 0.83682473 |
| MLEC | 1.495571884 | 0.03622827 | 0.58069725 | 1.44095237 |
| ARID4B | 1.494758304 | 0.07644075 | 0.57991223 | 1.11667505 |
| NUP133 | 1.494053189 | 0.06195232 | 0.57923151 | 1.2079424 |

| | | | | |
|---|---|---|---|---|
| STX3 | 1.49381166 | 0.13159132 | 0.57899826 | 0.88077275 |
| CLPTM1L | 1.492252617 | 0.08549958 | 0.57749178 | 1.068036 |
| RBBP4 | 1.491463839 | 0.03062679 | 0.576729 | 1.51389851 |
| EXOC7 | 1.490709789 | 0.02819609 | 0.57599942 | 1.54981105 |
| MAP1LC3A | 1.489425328 | 0.03001275 | 0.5747558 | 1.52269415 |
| KANK1 | 1.487915612 | 0.028125 | 0.57329271 | 1.55090749 |
| NDUFS1 | 1.486946883 | 0.048892 | 0.57235311 | 1.31076219 |
| TC2N | 1.486897874 | 0.00196879 | 0.57230556 | 2.70580075 |
| GIGYF2 | 1.483684257 | 0.01022092 | 0.5691841 | 1.99050983 |
| RRM2 | 1.482955277 | 0.01053838 | 0.56847509 | 1.97722596 |
| SNCB | 1.48034606 | 0.05840067 | 0.56593447 | 1.23358219 |
| LARP1B | 1.479668771 | 0.01848715 | 0.56527426 | 1.73313004 |
| STARD10 | 1.479525477 | 0.20639891 | 0.56513454 | 0.68529261 |
| TIMM50 | 1.479521217 | 0.0104804 | 0.56513039 | 1.9796221 |
| DAGLB | 1.479347496 | 0.05332848 | 0.56496098 | 1.27304078 |
| HMBS | 1.478627626 | 0.00573232 | 0.56425877 | 2.24166954 |
| KDF1 | 1.473827182 | 0.26883747 | 0.55956737 | 0.5705102 |
| SUMO1 | 1.473380884 | 0.03069406 | 0.55913043 | 1.51294559 |
| RAB5B | 1.473249278 | 0.88984635 | 0.55900156 | 0.05068498 |
| RALY | 1.472029636 | 0.01741153 | 0.55780672 | 1.75916298 |
| GNL1 | 1.471343475 | 0.09955448 | 0.55713407 | 1.0019392 |
| MCCC1 | 1.4707797 | 0.03592098 | 0.55658117 | 1.44465178 |
| MRPS12 | 1.47011119 | 0.16539948 | 0.55592528 | 0.78146587 |
| STOML1 | 1.468956235 | 0.14232182 | 0.55479141 | 0.8467285 |
| UTP15 | 1.4664647 | 0.91249952 | 0.55234234 | 0.03976735 |
| ALDH18A1 | 1.465554236 | 0.03585821 | 0.55144636 | 1.44541133 |
| KCNA3 | 1.463012192 | 0.07981449 | 0.54894179 | 1.09791824 |
| ZFAND6 | 1.4628537 | 0.06683653 | 0.54878549 | 1.17498614 |
| MAPT | 1.461689029 | 0.05447834 | 0.54763641 | 1.26377615 |
| UBE2S | 1.461671983 | 0.34713189 | 0.54761959 | 0.45950549 |
| INPP5B | 1.460571719 | 0.02821118 | 0.5465332 | 1.54957875 |
| RABGEF1 | 1.460479112 | 0.04306125 | 0.54644172 | 1.36591336 |
| ZNF616 | 1.459495337 | 0.77157762 | 0.5454696 | 0.11262038 |
| NBR1 | 1.45865191 | 0.10884666 | 0.54463564 | 0.96318491 |
| TMPO | 1.457221723 | 0.0134095 | 0.54322041 | 1.87258756 |
| AKAP5 | 1.455600128 | 0.89803424 | 0.54161408 | 0.0467071 |
| CNOT10 | 1.453325584 | 0.03029014 | 0.53935794 | 1.51869878 |
| KIAA1671 | 1.452740995 | 0.25000186 | 0.53877751 | 0.60205676 |
| SHROOM2 | 1.450793676 | 0.00039451 | 0.53684236 | 3.40393856 |
| FMR1 | 1.447879433 | 0.11667405 | 0.53394147 | 0.93302572 |
| VDAC2 | 1.447426573 | 0.11206883 | 0.53349016 | 0.95051518 |
| RBX1 | 1.445410711 | 0.07687466 | 0.53147949 | 1.11421677 |
| DCAF7 | 1.443368077 | 0.11781046 | 0.52943925 | 0.92881614 |
| CTDSPL2 | 1.441997642 | 0.11711672 | 0.52806881 | 0.9313811 |
| ZNF579 | 1.440531031 | 0.03178192 | 0.52660074 | 1.49781984 |

| | | | | |
|---|---|---|---|---|
| AHSA1 | 1.440201848 | 0.02965422 | 0.52627102 | 1.52791345 |
| ARHGAP35 | 1.43990978 | 0.01660844 | 0.52597842 | 1.77967128 |
| FAM98A | 1.438400869 | 0.0276639 | 0.5244658 | 1.55808654 |
| ATP1A1 | 1.438264967 | 0.01928317 | 0.52432948 | 1.71482157 |
| LRBA | 1.437144922 | 0.02532985 | 0.52320555 | 1.5963674 |
| MTR | 1.436873027 | 0.84963173 | 0.52293258 | 0.07076928 |
| TLK2 | 1.436371384 | 0.03135377 | 0.52242882 | 1.50371024 |
| HSPH1 | 1.434011078 | 0.10647432 | 0.52005617 | 0.97275512 |
| RPF2 | 1.430812119 | 0.02251872 | 0.51683424 | 1.64745621 |
| GLRX3 | 1.429038977 | 0.16935185 | 0.51504527 | 0.77121006 |
| SH2D3C | 1.428212531 | 0.0919695 | 0.51421068 | 1.03635617 |
| PLD3 | 1.422859285 | 0.0140802 | 0.50879299 | 1.85139132 |
| RNGTT | 1.421132682 | 0.27160817 | 0.50704126 | 0.56605716 |
| AARSD1 | 1.419355626 | 0.03668866 | 0.50523611 | 1.4354682 |
| TWF1 | 1.417670413 | 0.24287619 | 0.50352217 | 0.61461506 |
| PREP | 1.415785405 | 0.08645303 | 0.50160261 | 1.06321979 |
| PRAG1 | 1.414625993 | 0.06849 | 0.50042068 | 1.16437286 |
| | 1.412992697 | 0.21635154 | 0.49875401 | 0.66484 |
| THUMPD1 | 1.412829155 | 0.03424119 | 0.49858702 | 1.46545119 |
| KCTD5 | 1.412078015 | 0.0642417 | 0.4978198 | 1.192183 |
| BLTP1 | 1.411410039 | 0.58974291 | 0.49713718 | 0.22933727 |
| SPINT2 | 1.40991871 | 0.00730323 | 0.49561198 | 2.13648498 |
| HAUS1 | 1.40982251 | 0.04293272 | 0.49551355 | 1.3672116 |
| NOL10 | 1.408155447 | 0.22066912 | 0.4938066 | 0.65625843 |
| CHCHD6 | 1.407920469 | 0.61802266 | 0.49356584 | 0.2089956 |
| LMBRD2 | 1.406624463 | 0.01766025 | 0.49223721 | 1.75300318 |
| PPP3R1 | 1.406097849 | 0.41397261 | 0.49169699 | 0.38302839 |
| STX2 | 1.405131677 | 0.11617548 | 0.49070533 | 0.93488553 |
| RNF31 | 1.404435908 | 0.03698213 | 0.48999079 | 1.43200804 |
| EEF1A2 | 1.403213892 | 0.02214647 | 0.48873494 | 1.65469545 |
| TARS1 | 1.402790879 | 0.02453503 | 0.48829996 | 1.61021345 |
| RPS27 | 1.401291548 | 0.01658291 | 0.48675715 | 1.78033916 |
| NUP58 | 1.400591655 | 0.09203103 | 0.4860364 | 1.03606572 |
| LRRC8B | 1.400135922 | 0.20246113 | 0.48556689 | 0.69365835 |
| ATP2C1 | 1.399944667 | 0.00926854 | 0.48536981 | 2.03298853 |
| NOP56 | 1.398489238 | 0.05572847 | 0.48386915 | 1.25392285 |
| MINK1 | 1.395854465 | 0.4483602 | 0.48114853 | 0.34837295 |
| MPDZ | 1.394338178 | 0.46019536 | 0.47958051 | 0.33705776 |
| SYNPO | 1.394196448 | 0.37031111 | 0.47943386 | 0.43143326 |
| TOR1AIP1 | 1.39077661 | 0.25633032 | 0.47589071 | 0.59120001 |
| ANKRD27 | 1.389283021 | 0.93416897 | 0.47434053 | 0.02957456 |
| UBN1 | 1.387680375 | 0.3289879 | 0.47267531 | 0.48282007 |
| N4BP2 | 1.387560435 | 0.08245051 | 0.47255061 | 1.08380664 |
| VAPB | 1.387047744 | 0.00952249 | 0.47201745 | 2.02124934 |
| MFN1 | 1.385130936 | 0.04012641 | 0.47002236 | 1.39656973 |

| | | | | |
|---|---|---|---|---|
| SNX8 | 1.385006166 | 0.255556 | 0.4698924 | 0.59251391 |
| STMN3 | 1.384872791 | 0.31185925 | 0.46975346 | 0.50604137 |
| PSMA7 | 1.383081956 | 0.13939381 | 0.46788665 | 0.85575652 |
| TCP1 | 1.381423282 | 0.00988424 | 0.46615544 | 2.00505656 |
| KIF26A | 1.38101084 | 0.51789059 | 0.46572464 | 0.28576198 |
| DGCR2 | 1.380537194 | 0.54202878 | 0.46522976 | 0.26597765 |
| SLC39A11 | 1.380497801 | 0.17695715 | 0.46518859 | 0.75213188 |
| SNX2 | 1.379778292 | 0.03940719 | 0.46443647 | 1.40442453 |
| TBC1D17 | 1.377833417 | 0.34532262 | 0.46240147 | 0.46177498 |
| | 1.376765976 | 0.02945133 | 0.46128335 | 1.53089505 |
| CAB39 | 1.376607405 | 0.02891505 | 0.46111718 | 1.53887602 |
| EIF4G2 | 1.374628091 | 0.00779703 | 0.45904135 | 2.10807057 |
| CISD2 | 1.372129917 | 0.03441546 | 0.45641709 | 1.46324645 |
| RPL23 | 1.369925862 | 0.0211194 | 0.45409782 | 1.67531849 |
| HBS1L | 1.369476879 | 0.05307813 | 0.45362491 | 1.2750844 |
| KRT13 | 1.367825286 | 0.29947321 | 0.45188396 | 0.52364202 |
| MRTO4 | 1.367160486 | 0.00642807 | 0.45118261 | 2.19191915 |
| NUP54 | 1.366371056 | 0.0465324 | 0.45034932 | 1.33224458 |
| UBL3 | 1.36617026 | 0.16729626 | 0.45013729 | 0.77651376 |
| ARRDC3 | 1.365651479 | 0.18421182 | 0.44958935 | 0.73468251 |
| PHF6 | 1.365100021 | 0.08288304 | 0.44900666 | 1.08153434 |
| SLC33A1 | 1.364972549 | 0.21770027 | 0.44887194 | 0.66214104 |
| MACO1 | 1.364540086 | 0.02852413 | 0.44841478 | 1.5447876 |
| BUB1 | 1.364394147 | 0.10508315 | 0.44826047 | 0.9784669 |
| PPP4R3B | 1.36257827 | 0.10050406 | 0.4463391 | 0.9978164 |
| HSD17B12 | 1.362449271 | 0.06900895 | 0.44620251 | 1.16109456 |
| SLC4A2 | 1.361223315 | 0.0388187 | 0.44490377 | 1.41095904 |
| TES | 1.360754725 | 0.02554442 | 0.44440705 | 1.59270391 |
| LMNB1 | 1.35964352 | 0.02410889 | 0.44322845 | 1.61782278 |
| RBM14 | 1.359029627 | 0.04563085 | 0.44257691 | 1.34074147 |
| SRA1 | 1.3589662 | 0.79608206 | 0.44250957 | 0.09904216 |
| CCT3 | 1.358772013 | 0.00444644 | 0.44230341 | 2.35198724 |
| AKR1A1 | 1.358273715 | 0.06911218 | 0.44177424 | 1.1604454 |
| FN3KRP | 1.358174238 | 0.01252941 | 0.44166857 | 1.90206934 |
| PKM | 1.357120177 | 0.03949657 | 0.44054848 | 1.4034406 |
| ZBTB11 | 1.354523005 | 0.46298062 | 0.4377849 | 0.33443719 |
| FASTKD1 | 1.3529392 | 0.52316711 | 0.43609701 | 0.28135957 |
| EEIG2 | 1.352798968 | 0.12341606 | 0.43594746 | 0.90862832 |
| LARP1 | 1.351529831 | 0.11065821 | 0.43459336 | 0.95601638 |
| DUT | 1.351483329 | 0.18164954 | 0.43454372 | 0.74076571 |
| NUP160 | 1.350987362 | 0.02130842 | 0.43401418 | 1.67144868 |
| SEPTIN11 | 1.349430704 | 0.18012901 | 0.43235089 | 0.74441634 |
| PSMB5 | 1.349360288 | 0.03038999 | 0.43227561 | 1.51726946 |
| CENPJ | 1.347964879 | 0.03060027 | 0.43078291 | 1.51427471 |
| DNMBP | 1.346620028 | 0.1166617 | 0.42934283 | 0.93307169 |

| | | | | |
|---|---|---|---|---|
| PLEKHA1 | 1.342725333 | 0.21878672 | 0.42516422 | 0.65997904 |
| PTPN23 | 1.342463915 | 0.0138418 | 0.42488331 | 1.85880741 |
| ABT1 | 1.342382122 | 0.52520795 | 0.42479541 | 0.27966871 |
| AURKA | 1.340532883 | 0.06715461 | 0.42280661 | 1.17292419 |
| NCOA7 | 1.338586206 | 0.80667559 | 0.42071005 | 0.09330108 |
| HSP90AB1 | 1.338536216 | 0.04698588 | 0.42065617 | 1.32803261 |
| PPFIA1 | 1.338307391 | 0.3886427 | 0.42040952 | 0.41044949 |
| BRPF1 | 1.336801199 | 0.71565177 | 0.41878493 | 0.14529825 |
| SLC25A4 | 1.335895646 | 0.07561048 | 0.41780732 | 1.12141799 |
| TMEM165 | 1.335633218 | 0.19056048 | 0.41752388 | 0.71996715 |
| FARSB | 1.334662411 | 0.12450541 | 0.41647487 | 0.90481176 |
| TBCD | 1.33077983 | 0.03680232 | 0.41227191 | 1.43412482 |
| STIM1 | 1.33059934 | 0.01511419 | 0.41207622 | 1.82061526 |
| ADO | 1.32883878 | 0.30677472 | 0.41016608 | 0.51318043 |
| CAND1 | 1.32833389 | 0.11276676 | 0.40961783 | 0.94781891 |
| CSDE1 | 1.327957312 | 0.05061518 | 0.40920877 | 1.29571922 |
| LZTS1 | 1.327478389 | 0.0316792 | 0.40868837 | 1.49922579 |
| AARS1 | 1.326207565 | 0.07712965 | 0.40730659 | 1.11277863 |
| ATAT1 | 1.324126552 | 0.65776046 | 0.40504101 | 0.18193224 |
| TM9SF2 | 1.323747167 | 0.08256983 | 0.4046276 | 1.08317859 |
| SLC30A9 | 1.321139523 | 0.10395669 | 0.40178284 | 0.98314754 |
| MYH9 | 1.320219616 | 0.06180423 | 0.40077794 | 1.20898183 |
| SYT3 | 1.320213918 | 0.36769851 | 0.40077171 | 0.43450813 |
| SKP1 | 1.31715576 | 0.21000939 | 0.39742596 | 0.67776128 |
| EXOSC7 | 1.314835474 | 0.47630481 | 0.39488229 | 0.32211504 |
| UCK2 | 1.314303654 | 0.57001911 | 0.39429863 | 0.24411058 |
| STX4 | 1.313598199 | 0.01472968 | 0.39352405 | 1.83180671 |
| CAPNS1 | 1.313199645 | 0.16018898 | 0.39308626 | 0.79536737 |
| RABGGTB | 1.311288997 | 0.14481755 | 0.39098568 | 0.8391788 |
| TALDO1 | 1.309877632 | 0.22284568 | 0.38943204 | 0.65199579 |
| TMEM185B | 1.308216023 | 0.43718565 | 0.38760079 | 0.3593341 |
| ROCK2 | 1.307815244 | 0.32851137 | 0.38715874 | 0.48344959 |
| TMEM126A | 1.306740101 | 0.01466746 | 0.38597223 | 1.83364503 |
| CEP55 | 1.305946529 | 0.08194485 | 0.38509583 | 1.08647832 |
| TMEM87A | 1.305637956 | 0.41417414 | 0.3847549 | 0.38281702 |
| ARHGDIA | 1.30458859 | 0.1041834 | 0.38359492 | 0.98220146 |
| PAN3 | 1.304270688 | 0.11738937 | 0.38324332 | 0.93037124 |
| NDFIP1 | 1.302237404 | 0.37036103 | 0.38099248 | 0.43137472 |
| PSMG1 | 1.301124502 | 0.11454055 | 0.37975902 | 0.94104075 |
| PYGB | 1.30112253 | 0.0672079 | 0.37975683 | 1.1725797 |
| PATL1 | 1.300839759 | 0.03741114 | 0.37944326 | 1.42699903 |
| SUPT16H | 1.300639348 | 0.26579501 | 0.37922098 | 0.57545318 |
| HNRNPK | 1.29733606 | 0.02321207 | 0.37555224 | 1.63428612 |
| DUSP11 | 1.295067042 | 0.40220927 | 0.37302678 | 0.39554792 |
| NUP98 | 1.293449573 | 0.03685192 | 0.37122381 | 1.43353991 |

| | | | | |
|---|---|---|---|---|
| SLC25A5 | 1.292835318 | 0.14822496 | 0.37053852 | 0.82907865 |
| YEATS4 | 1.291242648 | 0.84868127 | 0.36876013 | 0.07125538 |
| UBAC1 | 1.291059808 | 0.26791814 | 0.36855584 | 0.57199788 |
| PHAF1 | 1.290540274 | 0.07668127 | 0.36797516 | 1.11531069 |
| PCBP3 | 1.290055615 | 0.57977084 | 0.36743326 | 0.23674363 |
| GORASP2 | 1.288416675 | 0.21981022 | 0.36559924 | 0.65795212 |
| CACNA1B | 1.286450718 | 0.87319938 | 0.36339619 | 0.05888658 |
| CCT2 | 1.284666393 | 0.06494096 | 0.36139376 | 1.18748129 |
| ALKBH1 | 1.284251292 | 0.8198168 | 0.36092752 | 0.08628319 |
| ISLR2 | 1.282756018 | 0.48419866 | 0.35924679 | 0.31497641 |
| DPM3 | 1.282670703 | 0.16480934 | 0.35915084 | 0.78301818 |
| SRSF5 | 1.282120495 | 0.62258916 | 0.35853185 | 0.20579844 |
| PPA1 | 1.282094569 | 0.21092282 | 0.35850268 | 0.67587643 |
| MYL12B | 1.281913027 | 0.18385996 | 0.35829838 | 0.73551285 |
| MPP1 | 1.281225058 | 0.3739584 | 0.35752392 | 0.4271767 |
| WASF2 | 1.281117331 | 0.0887013 | 0.35740261 | 1.05207004 |
| TMX2 | 1.279364882 | 0.19878246 | 0.35542779 | 0.70162194 |
| SNIP1 | 1.277876153 | 0.08916402 | 0.35374802 | 1.04981038 |
| FKBP5 | 1.27740981 | 0.13396204 | 0.35322143 | 0.87301826 |
| LSM12 | 1.277314637 | 0.10391217 | 0.35311394 | 0.98333359 |
| CCDC88C | 1.274936936 | 0.037583 | 0.35042589 | 1.4250085 |
| IQCB1 | 1.274167074 | 0.63418979 | 0.34955446 | 0.19778075 |
| YKT6 | 1.273696232 | 0.03243425 | 0.34902125 | 1.4889961 |
| TBC1D8B | 1.272246638 | 0.18849103 | 0.34737838 | 0.72470932 |
| ABRAXAS2 | 1.271194738 | 0.05376666 | 0.34618506 | 1.26948692 |
| BUB1B | 1.271157105 | 0.01948972 | 0.34614235 | 1.71019432 |
| SPCS2 | 1.268726232 | 0.00405469 | 0.34338079 | 2.39204279 |
| DDX39A | 1.267457297 | 0.24724114 | 0.34193714 | 0.60687927 |
| COMT | 1.267216731 | 0.40014266 | 0.34166329 | 0.39778515 |
| RPL7L1 | 1.266908995 | 0.73999703 | 0.3413129 | 0.13077002 |
| BECN1 | 1.266462896 | 0.99983878 | 0.34080481 | 7.0022E-05 |
| UBAC2 | 1.265461027 | 0.15294867 | 0.33966308 | 0.8154543 |
| SNRPB2 | 1.264976258 | 0.34669417 | 0.33911031 | 0.46005346 |
| RFX3 | 1.264674105 | 0.09071376 | 0.33876566 | 1.04232685 |
| RABGAP1 | 1.26398606 | 0.04071871 | 0.33798055 | 1.39020596 |
| LEMD3 | 1.263156483 | 0.41874224 | 0.33703337 | 0.37805323 |
| SLC16A7 | 1.26300364 | 0.40783139 | 0.3368588 | 0.38951935 |
| NAV1 | 1.260889891 | 0.42275518 | 0.3344423 | 0.37391106 |
| STX1A | 1.260013284 | 0.06340801 | 0.33343894 | 1.19785589 |
| LPIN1 | 1.259668552 | 0.34464934 | 0.33304418 | 0.46262255 |
| NDUFA9 | 1.258563131 | 0.5670572 | 0.33177759 | 0.24637313 |
| DDX56 | 1.258054584 | 0.01787645 | 0.33119452 | 1.74771861 |
| GPBP1 | 1.25800294 | 0.2100679 | 0.33113529 | 0.67764032 |
| LSM4 | 1.257086902 | 0.76889154 | 0.33008439 | 0.11413492 |
| HSP90AA1 | 1.255998649 | 0.07542453 | 0.32883491 | 1.12248741 |

| | | | | |
|---|---|---|---|---|
| DYNLL2 | 1.255204211 | 0.06578309 | 0.3279221 | 1.1818857 |
| PSMC3 | 1.254715451 | 0.13333771 | 0.32736022 | 0.87504702 |
| NSUN2 | 1.254541486 | 0.43168628 | 0.32716018 | 0.36483175 |
| DDX21 | 1.2544669 | 0.18720378 | 0.3270744 | 0.72768539 |
| NUP107 | 1.25439937 | 0.05098264 | 0.32699674 | 1.2925777 |
| EXOSC2 | 1.253456969 | 0.74980219 | 0.32591247 | 0.1250533 |
| EXOC2 | 1.252463233 | 0.71671525 | 0.32476825 | 0.14465335 |
| DEPDC5 | 1.250743244 | 0.55927994 | 0.32278566 | 0.25237076 |
| UBA2 | 1.250661067 | 0.25385749 | 0.32269087 | 0.59541003 |
| UHRF1 | 1.250639643 | 0.31014264 | 0.32266615 | 0.50843852 |
| CORO1A | 1.249475587 | 0.77193438 | 0.32132271 | 0.11241962 |
| NUCKS1 | 1.248104598 | 0.72361802 | 0.31973884 | 0.14049062 |
| MCM2 | 1.247854886 | 0.05899289 | 0.31945017 | 1.22920036 |
| STAT3 | 1.244030714 | 0.04621764 | 0.31502211 | 1.33519225 |
| WDR47 | 1.243733755 | 0.27922997 | 0.31467768 | 0.55403796 |
| INPPL1 | 1.243614011 | 0.02516905 | 0.31453878 | 1.59913323 |
| TRAPPC10 | 1.242798406 | 0.44717706 | 0.3135923 | 0.34952049 |
| ADIPOR1 | 1.242186053 | 0.97008609 | 0.31288127 | 0.01318972 |
| POLR2E | 1.240304023 | 0.06026943 | 0.3106938 | 1.21990294 |
| SMAD2 | 1.239602618 | 0.33287182 | 0.30987771 | 0.47772297 |
| NRSN2 | 1.238456814 | 0.75905241 | 0.30854356 | 0.11972824 |
| NGDN | 1.234851859 | 0.15413889 | 0.30433798 | 0.81208779 |
| LCLAT1 | 1.234739367 | 0.1650377 | 0.30420655 | 0.78241684 |
| TRNT1 | 1.233000987 | 0.09950547 | 0.30217396 | 1.00215305 |
| PRR12 | 1.232549404 | 0.0257428 | 0.30164548 | 1.58934414 |
| FADS2 | 1.232103785 | 0.22067168 | 0.30112379 | 0.6562534 |
| KIFBP | 1.231923862 | 0.30763807 | 0.30091309 | 0.51195992 |
| RBM45 | 1.231443683 | 0.42155759 | 0.30035065 | 0.37514309 |
| CCDC137 | 1.231081733 | 0.34224676 | 0.29992655 | 0.46566066 |
| RCC2 | 1.230087415 | 0.09905945 | 0.29876084 | 1.00410409 |
| | 1.229214225 | 0.39441489 | 0.29773637 | 0.40404669 |
| ARPC4 | 1.22795319 | 0.15428842 | 0.29625557 | 0.81166668 |
| AGL | 1.227635035 | 0.13115969 | 0.29588172 | 0.88219962 |
| MCM5 | 1.226158607 | 0.04818803 | 0.29414561 | 1.31706081 |
| AP3B1 | 1.222655057 | 0.64502336 | 0.29001744 | 0.19042456 |
| NUDT4 | 1.221260483 | 0.57394346 | 0.28837095 | 0.24113089 |
| LZIC | 1.219198301 | 0.11424911 | 0.2859328 | 0.94214718 |
| SRPRB | 1.217715885 | 0.17278242 | 0.28417757 | 0.76250045 |
| NFX1 | 1.217696449 | 0.92370784 | 0.28415454 | 0.03446537 |
| SYN2 | 1.215795133 | 0.3359929 | 0.28190015 | 0.4736699 |
| SURF2 | 1.214877157 | 0.88216791 | 0.28081044 | 0.05444875 |
| MCM3 | 1.214689282 | 0.10381431 | 0.28058732 | 0.98374277 |
| PFKP | 1.21410652 | 0.27794534 | 0.279895 | 0.55604061 |
| CCDC12 | 1.213248705 | 0.91287935 | 0.27887532 | 0.03958662 |
| YARS1 | 1.212498834 | 0.15215906 | 0.27798336 | 0.81770219 |

| | | | | |
|---|---|---|---|---|
| PFDN2 | 1.211474659 | 0.25795637 | 0.27676423 | 0.58845374 |
| IMMT | 1.210357149 | 0.10815784 | 0.27543282 | 0.96594198 |
| RAB3A | 1.210260558 | 0.80552875 | 0.27531768 | 0.09391895 |
| RPL9; RPL9P7; RPI | 1.209582027 | 0.03200031 | 0.27450861 | 1.49484585 |
| HNRNPH1 | 1.208987801 | 0.11650659 | 0.27379969 | 0.93364951 |
| ATP6V1A | 1.208212222 | 0.15100101 | 0.27287389 | 0.82102014 |
| MAT2A | 1.206335871 | 0.29009646 | 0.27063164 | 0.53745757 |
| PRKAR1B | 1.204914444 | 0.69864343 | 0.26893071 | 0.15574442 |
| LRRC8A | 1.203836466 | 0.65141136 | 0.26763942 | 0.18614467 |
| RABGAP1L | 1.203660454 | 0.18918014 | 0.26742847 | 0.72312446 |
| DIPK2A | 1.202553928 | 0.78433388 | 0.26610159 | 0.10549902 |
| ARL1 | 1.201905737 | 0.20207425 | 0.26532375 | 0.69448902 |
| PNO1 | 1.201722636 | 0.33447409 | 0.26510395 | 0.47563752 |
| MARK3 | 1.200958701 | 0.19428594 | 0.26418654 | 0.71155863 |
| ATL2 | 1.199434773 | 0.138616 | 0.2623547 | 0.85818663 |
| RNF113A | 1.198628521 | 0.13242081 | 0.26138461 | 0.87804376 |
| POLR3A | 1.19798438 | 0.13322625 | 0.2606091 | 0.8754102 |
| KIT | 1.197815807 | 0.04632135 | 0.26040608 | 1.33421878 |
| PPM1A | 1.196804624 | 0.44237284 | 0.25918765 | 0.35421154 |
| AKT3 | 1.196691571 | 0.37148333 | 0.25905137 | 0.43006067 |
| DDX50 | 1.19571147 | 0.27032624 | 0.2578693 | 0.5681118 |
| EIF2B2 | 1.194635165 | 0.13331446 | 0.25657009 | 0.87512273 |
| R3HDM1 | 1.194577267 | 0.02765156 | 0.25650017 | 1.55828029 |
| ITGB1 | 1.193639089 | 0.33906743 | 0.25536669 | 0.46971393 |
| TOMM70 | 1.192559436 | 0.24551421 | 0.25406117 | 0.60992336 |
| PARK7 | 1.191704818 | 0.82725019 | 0.25302693 | 0.08236312 |
| MARF1 | 1.191479388 | 0.13492371 | 0.25275399 | 0.86991173 |
| STIP1 | 1.190972684 | 0.10040092 | 0.25214032 | 0.9982623 |
| STIM2 | 1.190768111 | 0.04785585 | 0.25189249 | 1.32006498 |
| AP3D1 | 1.190219915 | 0.46495923 | 0.25122816 | 0.33258512 |
| BSCL2 | 1.189997856 | 0.27961059 | 0.25095897 | 0.55344639 |
| CEP350 | 1.189723673 | 0.6576986 | 0.25062653 | 0.18197308 |
| SLC25A25 | 1.18924062 | 0.11025047 | 0.25004065 | 0.95761955 |
| PRPSAP2 | 1.188686659 | 0.93247749 | 0.24936847 | 0.03036164 |
| CKAP2 | 1.187567049 | 0.05206638 | 0.24800897 | 1.28344263 |
| RBM39 | 1.18666233 | 0.06474424 | 0.24690947 | 1.18879888 |
| CALCOCO1 | 1.18642062 | 0.92829367 | 0.24661558 | 0.03231461 |
| TTC28 | 1.185287483 | 0.42341079 | 0.24523702 | 0.37323808 |
| HM13 | 1.184012798 | 0.51937928 | 0.24368468 | 0.28451538 |
| HPF1 | 1.182878859 | 0.02347802 | 0.24230233 | 1.62933849 |
| FANCI | 1.18250648 | 0.91811305 | 0.24184809 | 0.03710384 |
| RTN4 | 1.18168036 | 0.10154065 | 0.24083984 | 0.99336007 |
| SLC2A1 | 1.180647417 | 0.01936228 | 0.23957819 | 1.7130435 |
| VCL | 1.18029806 | 0.55831427 | 0.23915123 | 0.25312127 |
| TFDP1 | 1.180061872 | 0.68164903 | 0.2388625 | 0.16643918 |

| | | | | |
|---|---|---|---|---|
| PCF11 | 1.177903143 | 0.90611791 | 0.23622091 | 0.04281529 |
| PLPBP | 1.175812237 | 0.33641844 | 0.2336577 | 0.4731202 |
| KRT19 | 1.174837155 | 0.23899007 | 0.2324608 | 0.62162015 |
| CDC37 | 1.174733616 | 0.85925664 | 0.23233365 | 0.0658771 |
| DNAJA1 | 1.174711385 | 0.8491469 | 0.23230634 | 0.07101717 |
| PSMD11 | 1.173435511 | 0.47956337 | 0.23073856 | 0.31915399 |
| AGAP1 | 1.173413597 | 0.59649685 | 0.23071161 | 0.22439185 |
| PSMC6 | 1.173253185 | 0.92225484 | 0.23051438 | 0.03514906 |
| ARFGAP3 | 1.173223527 | 0.01881777 | 0.23047791 | 1.72543175 |
| ESRP1 | 1.173020483 | 0.368221 | 0.23022821 | 0.43389145 |
| GTPBP2 | 1.172615994 | 0.37447632 | 0.22973064 | 0.42657564 |
| ATP6V1B2 | 1.171410987 | 0.19865557 | 0.22824733 | 0.70189924 |
| TANC2 | 1.171256629 | 0.7805315 | 0.22805721 | 0.10760957 |
| TAOK2 | 1.17124823 | 0.02918872 | 0.22804687 | 1.53478499 |
| NUP155 | 1.170263577 | 0.86540277 | 0.2268335 | 0.06278172 |
| RPL17 | 1.168278496 | 0.14507496 | 0.22438423 | 0.83840753 |
| WDR13 | 1.166526365 | 0.26867734 | 0.22221891 | 0.57076895 |
| DENND4C | 1.166185826 | 0.15294294 | 0.22179769 | 0.81547056 |
| SLC27A3 | 1.164327002 | 0.12044977 | 0.2194963 | 0.91919404 |
| RPRD1A | 1.1639381 | 0.65584658 | 0.21901434 | 0.18319774 |
| TELO2 | 1.163882071 | 0.87311139 | 0.21894489 | 0.05893035 |
| ALDH3A2 | 1.163645811 | 0.83321262 | 0.218652 | 0.07924416 |
| PRKAA1 | 1.163437237 | 0.24879302 | 0.21839338 | 0.60416181 |
| NCDN | 1.161899626 | 0.22468671 | 0.21648544 | 0.64842262 |
| SNX4 | 1.160197149 | 0.47254396 | 0.21436998 | 0.32555778 |
| MICALL2 | 1.159626596 | 0.14845211 | 0.21366033 | 0.82841362 |
| VPS51 | 1.159182811 | 0.92258592 | 0.21310811 | 0.03499318 |
| EIF4ENIF1 | 1.15808553 | 0.90424789 | 0.21174181 | 0.0437125 |
| TANC1 | 1.156839027 | 0.22538318 | 0.21018813 | 0.64707851 |
| VRK1 | 1.156366301 | 0.10267957 | 0.20959847 | 0.98851596 |
| RPP30 | 1.156152798 | 0.85160734 | 0.20933208 | 0.06976061 |
| THOC5 | 1.154249911 | 0.22933597 | 0.20695562 | 0.63952783 |
| RAD23B | 1.153903542 | 0.0774615 | 0.20652263 | 1.11091409 |
| KPNB1 | 1.153779788 | 0.11671625 | 0.2063679 | 0.93286866 |
| SPOUT1 | 1.150697645 | 0.06657559 | 0.2025088 | 1.17668497 |
| ADPGK | 1.148288171 | 0.61296175 | 0.19948474 | 0.21256663 |
| PSMD5 | 1.148182978 | 0.39007149 | 0.19935257 | 0.40885579 |
| KCNH2 | 1.147301545 | 0.66656142 | 0.19824462 | 0.17615983 |
| COPG1 | 1.14553864 | 0.74597511 | 0.19602612 | 0.12727567 |
| HINT1 | 1.141864795 | 0.34828544 | 0.19139184 | 0.45806468 |
| KRT18 | 1.141183564 | 0.10503966 | 0.19053087 | 0.9786467 |
| DRG2 | 1.139918877 | 0.19620919 | 0.18893116 | 0.70728066 |
| GSTCD | 1.139443659 | 0.90325882 | 0.18832959 | 0.04418779 |
| GALK1 | 1.139335375 | 0.70267788 | 0.18819248 | 0.15324372 |
| PTPN11 | 1.138604289 | 0.42730645 | 0.18726644 | 0.36926055 |

| | | | | |
|---|---|---|---|---|
| CLK1 | 1.138476958 | 0.11555449 | 0.18710509 | 0.93721317 |
| NDUFV1 | 1.13728664 | 0.90047366 | 0.18559592 | 0.04552899 |
| PRDX6 | 1.136383457 | 0.09411304 | 0.18444973 | 1.02635019 |
| PIBF1 | 1.135892905 | 0.82954554 | 0.18382682 | 0.08115977 |
| EEF1E1-BLOC1S5 | 1.135764274 | 0.59845973 | 0.18366344 | 0.22296507 |
| R3HDM2 | 1.134733486 | 0.327598 | 0.18235349 | 0.48465876 |
| FPGT | 1.1341836 | 0.35540192 | 0.1816542 | 0.44928023 |
| INPP5F | 1.133149203 | 0.3705022 | 0.18033783 | 0.43120921 |
| PMVK | 1.131764108 | 0.47391383 | 0.17857329 | 0.32430062 |
| ASXL1 | 1.130635197 | 0.3230533 | 0.17713351 | 0.49072582 |
| NTPCR | 1.129207108 | 0.8414913 | 0.17531012 | 0.07495037 |
| ARCN1 | 1.128935802 | 0.51802909 | 0.17496345 | 0.28564585 |
| UBR1 | 1.128562326 | 0.92268238 | 0.1744861 | 0.03494777 |
| VWA5B2 | 1.127638451 | 0.15455446 | 0.17330458 | 0.81091845 |
| PFDN6 | 1.127462389 | 0.10443563 | 0.17307931 | 0.98115133 |
| COPE | 1.127112288 | 0.14236363 | 0.17263125 | 0.84660094 |
| SV2A | 1.127034684 | 0.15383496 | 0.17253191 | 0.81294497 |
| NONO | 1.126569514 | 0.14539306 | 0.17193634 | 0.83745633 |
| EIF5 | 1.126420514 | 0.71342709 | 0.17174551 | 0.1466504 |
| ANKRD50 | 1.124796498 | 0.52962882 | 0.16966401 | 0.27602839 |
| SH3GLB1 | 1.123365538 | 0.17109785 | 0.16782745 | 0.76675546 |
| ATL3 | 1.122253501 | 0.29525862 | 0.1663986 | 0.52979741 |
| GTPBP6 | 1.121930218 | 0.42859016 | 0.16598295 | 0.3679578 |
| WDR5 | 1.11927716 | 0.66737013 | 0.16256733 | 0.17563324 |
| DDX27 | 1.118842251 | 0.21665666 | 0.16200664 | 0.66422796 |
| SRRM1 | 1.116492391 | 0.35580819 | 0.15897342 | 0.44878406 |
| NUP214 | 1.114018022 | 0.14012423 | 0.15577257 | 0.85348675 |
| PSMD13 | 1.112527639 | 0.96194648 | 0.15384118 | 0.01684909 |
| MYL6 | 1.111251262 | 0.98746488 | 0.15218506 | 0.00547834 |
| AGTPBP1 | 1.111196756 | 0.28440734 | 0.15211429 | 0.5460592 |
| PIK3CA | 1.109910446 | 0.31482838 | 0.15044328 | 0.50192612 |
| NFXL1 | 1.109509129 | 0.88369296 | 0.14992154 | 0.0536986 |
| ANKLE2 | 1.109308354 | 0.7754322 | 0.14966045 | 0.11045617 |
| CLTC | 1.107590124 | 0.21847458 | 0.1474241 | 0.66059908 |
| THEM6 | 1.107068065 | 0.17779187 | 0.14674393 | 0.75008809 |
| HNRNPUL2 | 1.105355139 | 0.36078973 | 0.14450997 | 0.44274583 |
| HS1BP3 | 1.10388052 | 0.15420578 | 0.14258403 | 0.81189934 |
| OLA1 | 1.103285663 | 0.34437208 | 0.14180638 | 0.46297207 |
| TNK1 | 1.103200823 | 0.42560247 | 0.14169544 | 0.37099586 |
| RBBP6 | 1.103136648 | 0.90448577 | 0.14161151 | 0.04359826 |
| NCLN | 1.102799572 | 0.13689322 | 0.14117061 | 0.86361807 |
| CSK | 1.101882063 | 0.2632731 | 0.13996982 | 0.57959351 |
| TEX2 | 1.100243035 | 0.30334117 | 0.13782224 | 0.51806865 |
| TBC1D10A | 1.098501972 | 0.37139498 | 0.13553746 | 0.43016397 |
| SYT1 | 1.097703174 | 0.13667641 | 0.13448799 | 0.86430644 |

| | | | | |
|---|---|---|---|---|
| SRSF1 | 1.097647512 | 0.36359983 | 0.13441484 | 0.43937633 |
| RGS12 | 1.097146411 | 0.72523228 | 0.13375606 | 0.13952287 |
| ATP2A2 | 1.096370929 | 0.55747163 | 0.13273598 | 0.25377723 |
| MAEA | 1.096257733 | 0.33827341 | 0.13258702 | 0.47073213 |
| TKFC | 1.096135925 | 0.24783577 | 0.13242671 | 0.60583601 |
| KCTD3 | 1.095159835 | 0.73659995 | 0.13114144 | 0.13276832 |
| ATP1B1 | 1.094519977 | 0.36906753 | 0.13029829 | 0.43289417 |
| CAMLG | 1.093978435 | 0.10231626 | 0.1295843 | 0.99005534 |
| BUD31 | 1.092586755 | 0.64464923 | 0.12774784 | 0.19067653 |
| SLC35E1 | 1.092292237 | 0.59327004 | 0.12735889 | 0.22674758 |
| CIP2A | 1.092013394 | 0.43821192 | 0.12699055 | 0.35831582 |
| CYB5A | 1.09087584 | 0.98361618 | 0.12548691 | 0.00717434 |
| EEF2 | 1.087775856 | 0.35157106 | 0.12138131 | 0.45398688 |
| RHOA | 1.087704475 | 0.64851593 | 0.12128664 | 0.18807935 |
| PSMA4 | 1.087599944 | 0.3181412 | 0.12114798 | 0.49738009 |
| NUP62 | 1.087299175 | 0.32122437 | 0.12074896 | 0.49319152 |
| RPL10 | 1.086550908 | 0.28466941 | 0.11975577 | 0.54565919 |
| VEZT | 1.085415149 | 0.41369046 | 0.11824695 | 0.38332449 |
| SLC25A13 | 1.084827099 | 0.15139636 | 0.11746512 | 0.81988456 |
| STAMBP | 1.084432353 | 0.17189406 | 0.11694006 | 0.76473913 |
| DLGAP5 | 1.084238277 | 0.88615884 | 0.11668184 | 0.05248843 |
| TRAPPC14 | 1.083787477 | 0.91709909 | 0.11608188 | 0.03758374 |
| LRRC40 | 1.08299093 | 0.38028182 | 0.11502116 | 0.41989443 |
| PPP6R2 | 1.081828994 | 0.76816891 | 0.11347247 | 0.11454328 |
| ALKBH5 | 1.080198449 | 0.5378549 | 0.11129638 | 0.26933487 |
| CDK1 | 1.080130538 | 0.48617896 | 0.11120568 | 0.31320384 |
| CNOT1 | 1.07887086 | 0.29278651 | 0.10952219 | 0.53344894 |
| NUP50 | 1.077464341 | 0.5736959 | 0.10764012 | 0.24131825 |
| HAUS5 | 1.076345669 | 0.88385587 | 0.10614147 | 0.05361855 |
| MAP4 | 1.076299592 | 0.16802294 | 0.10607971 | 0.77463142 |
| FBRSL1 | 1.075509108 | 0.62021683 | 0.10501974 | 0.20745645 |
| STIL | 1.0749033 | 0.36844218 | 0.10420688 | 0.43363066 |
| ZC3HAV1L | 1.074457863 | 0.99606669 | 0.10360891 | 0.00171158 |
| MAP2K2 | 1.073456603 | 0.62242064 | 0.10226387 | 0.20591601 |
| HAUS4 | 1.072946291 | 0.77194348 | 0.10157786 | 0.11241449 |
| PARP4 | 1.072708699 | 0.94496042 | 0.10125836 | 0.02458638 |
| POLD2 | 1.072259832 | 0.41829268 | 0.10065454 | 0.37851973 |
| DIS3L2 | 1.072019395 | 0.97314689 | 0.10033101 | 0.0118216 |
| NAPB | 1.071971865 | 0.60223779 | 0.10026704 | 0.22023199 |
| TACC1 | 1.071176223 | 0.3180034 | 0.09919584 | 0.49756824 |
| ALYREF | 1.070783062 | 0.25662825 | 0.09866622 | 0.59069554 |
| CEP97 | 1.070605567 | 0.45297735 | 0.09842706 | 0.34392351 |
| PURA | 1.07041958 | 0.61940474 | 0.09817641 | 0.20802547 |
| CFAP20 | 1.069715566 | 0.2290625 | 0.09722724 | 0.640046 |
| NUP85 | 1.068354528 | 0.58382873 | 0.09539048 | 0.23371454 |

| | | | | |
|---|---|---|---|---|
| TPD52 | 1.067294242 | 0.62786358 | 0.09395797 | 0.20213471 |
| GTF2H3 | 1.066536422 | 0.55644965 | 0.09293323 | 0.25457412 |
| VPS33A | 1.065901103 | 0.21533377 | 0.09207359 | 0.66688786 |
| PSME1 | 1.065681564 | 0.84012684 | 0.09177641 | 0.07565514 |
| RANBP2 | 1.065510544 | 0.45514668 | 0.09154487 | 0.34184862 |
| MAP4K2 | 1.064784163 | 0.74114125 | 0.09056102 | 0.13009902 |
| PSMA6 | 1.06474114 | 0.70310967 | 0.09050272 | 0.15297693 |
| STRAP | 1.064320008 | 0.76468621 | 0.08993199 | 0.11651674 |
| FKBP15 | 1.063930369 | 0.18830914 | 0.08940373 | 0.72512861 |
| AHCY | 1.063182936 | 0.94212914 | 0.08838986 | 0.02588956 |
| BPNT1 | 1.062940502 | 0.79066171 | 0.08806084 | 0.10200929 |
| NUP205 | 1.062623694 | 0.52933046 | 0.08763079 | 0.27627311 |
| ATP1B3 | 1.062496065 | 0.25062184 | 0.0874575 | 0.60098109 |
| NME1-NME2 | 1.062008746 | 0.44755477 | 0.08679565 | 0.34915381 |
| WDR18 | 1.061495295 | 0.58574867 | 0.08609798 | 0.23228869 |
| PCBP1 | 1.06133082 | 0.1712418 | 0.08587442 | 0.76639021 |
| MCRIP2 | 1.059139602 | 0.95149501 | 0.08289276 | 0.02159349 |
| CNOT3 | 1.059083449 | 0.513898 | 0.08281627 | 0.28912308 |
| BZW2 | 1.058500471 | 0.83166498 | 0.08202191 | 0.08005159 |
| U2AF1 | 1.058309173 | 0.58568901 | 0.08176116 | 0.23233292 |
| ATP8A1 | 1.057546555 | 0.70466261 | 0.08072117 | 0.15201877 |
| PSMD1 | 1.057165723 | 0.33689787 | 0.08020155 | 0.47250174 |
| TUBA4A | 1.056881338 | 0.60530954 | 0.07981341 | 0.21802248 |
| SRP68 | 1.056626453 | 0.86747251 | 0.07946543 | 0.06174428 |
| EHD1 | 1.056402252 | 0.65285112 | 0.07915928 | 0.18518584 |
| PAK4 | 1.056040575 | 0.4109295 | 0.07866527 | 0.38623268 |
| CNN2 | 1.055712613 | 0.71042587 | 0.07821716 | 0.14848123 |
| ZNF609 | 1.055234007 | 0.55712825 | 0.07756296 | 0.25404482 |
| MAPT | 1.053398948 | 0.64441094 | 0.07505192 | 0.1908371 |
| RNF19A | 1.052475913 | 0.24729119 | 0.07378722 | 0.60679135 |
| OXA1L | 1.052322594 | 0.26532595 | 0.07357704 | 0.57622027 |
| LANCL2 | 1.052061938 | 0.89362314 | 0.07321964 | 0.04884559 |
| STK39 | 1.051026669 | 0.70041385 | 0.07179928 | 0.15464528 |
| ACTR3 | 1.049072477 | 0.54699907 | 0.06911435 | 0.26201341 |
| ABHD16A | 1.048814026 | 0.41511274 | 0.06875888 | 0.38183394 |
| CHD7 | 1.047061223 | 0.41809088 | 0.0663458 | 0.3787293 |
| ANKS1A | 1.046230858 | 0.71559341 | 0.06520123 | 0.14533367 |
| AK1 | 1.046072191 | 0.90496801 | 0.06498242 | 0.04336677 |
| ARHGAP5 | 1.045688922 | 0.27655921 | 0.06445373 | 0.55821188 |
| CEP290 | 1.044822944 | 0.66934297 | 0.06325848 | 0.17435129 |
| WDR24 | 1.044714748 | 0.51195192 | 0.06310908 | 0.29077082 |
| FAM120B | 1.04379432 | 0.68346475 | 0.06183746 | 0.16528388 |
| ABHD12 | 1.043498489 | 0.42149175 | 0.06142851 | 0.37521092 |
| RPS6KA1 | 1.043365042 | 0.24898467 | 0.061244 | 0.60382739 |
| PSMD3 | 1.04324381 | 0.26445119 | 0.06107636 | 0.57765447 |

| | | | | |
|---|---|---|---|---|
| NOL4 | 1.041760871 | 0.84112809 | 0.05902415 | 0.07513786 |
| ALG5 | 1.039996421 | 0.45283479 | 0.05657856 | 0.34406022 |
| EIF2S3 | 1.038875883 | 0.65100597 | 0.0550233 | 0.18641503 |
| CEP44 | 1.038696496 | 0.74044065 | 0.05477417 | 0.13050974 |
| PPP1R7 | 1.038259499 | 0.3866255 | 0.05416707 | 0.41270951 |
| CLPTM1 | 1.038067233 | 0.97712018 | 0.05389989 | 0.01005202 |
| PEX11B | 1.038066923 | 0.65920365 | 0.05389946 | 0.18098039 |
| HUWE1 | 1.037785244 | 0.69516245 | 0.05350793 | 0.1579137 |
| SNRPG | 1.037470936 | 0.50558424 | 0.05307092 | 0.29620647 |
| WRN | 1.036097645 | 0.97768352 | 0.05115997 | 0.00980171 |
| SH3GLB2 | 1.036093869 | 0.86236841 | 0.05115472 | 0.06430716 |
| PDCD6IP | 1.035798405 | 0.39527111 | 0.05074324 | 0.40310492 |
| DNAJC6 | 1.035739255 | 0.56496897 | 0.05066085 | 0.2479754 |
| SFXN1 | 1.035382199 | 0.86740999 | 0.05016342 | 0.06177558 |
| NT5C3A | 1.034733675 | 0.99173167 | 0.04925949 | 0.00360582 |
| ACTR1B | 1.032443427 | 0.37730175 | 0.04606273 | 0.42331118 |
| SMC4 | 1.032430048 | 0.53777813 | 0.04604404 | 0.26939687 |
| IPO4 | 1.032087446 | 0.45910698 | 0.04556521 | 0.3380861 |
| CHD3 | 1.031178619 | 0.85761222 | 0.04429425 | 0.06670904 |
| RERE | 1.029930894 | 0.74467619 | 0.04254754 | 0.12803253 |
| TPMT | 1.029329794 | 0.98549918 | 0.04170529 | 0.00634373 |
| TRIM62 | 1.025346157 | 0.40198442 | 0.03611105 | 0.39579078 |
| DDX28 | 1.024163886 | 0.33124379 | 0.03444659 | 0.47985225 |
| PCID2 | 1.023959991 | 0.96720969 | 0.03415935 | 0.01447936 |
| NAE1 | 1.023958606 | 0.43132533 | 0.0341574 | 0.36519503 |
| TRMT44 | 1.023616936 | 0.6085371 | 0.03367592 | 0.21571294 |
| STXBP3 | 1.02184231 | 0.98931997 | 0.03117258 | 0.00466323 |
| PIGK | 1.021776919 | 0.79602055 | 0.03108025 | 0.09907572 |
| ATXN3 | 1.021348901 | 0.52732656 | 0.03047579 | 0.27792035 |
| HSPA5 | 1.020963838 | 0.60239422 | 0.02993177 | 0.2201192 |
| CNOT2 | 1.019676545 | 0.69727347 | 0.02811158 | 0.15659686 |
| TBCC | 1.01862677 | 0.84737604 | 0.02662554 | 0.07192382 |
| PTPN2 | 1.018264883 | 0.32626226 | 0.0261129 | 0.48643315 |
| BRSK2 | 1.01793239 | 0.83191354 | 0.02564174 | 0.07992181 |
| PDIA3 | 1.016873772 | 0.80817515 | 0.0241406 | 0.09249451 |
| PFAS | 1.016403708 | 0.40139577 | 0.02347354 | 0.39642721 |
| FLII | 1.015138832 | 0.84772805 | 0.02167705 | 0.07174345 |
| TBCB | 1.014762177 | 0.49724481 | 0.02114165 | 0.30342974 |
| RBM6 | 1.014256703 | 0.8832968 | 0.02042284 | 0.05389334 |
| DCK | 1.013511887 | 0.41299593 | 0.01936301 | 0.38405423 |
| NDUFA12 | 1.013006595 | 0.70843601 | 0.01864357 | 0.14969937 |
| CNOT9 | 1.012841147 | 0.78588291 | 0.01840792 | 0.10464215 |
| SLC25A40 | 1.011968168 | 0.98579592 | 0.01716391 | 0.00621298 |
| ZCCHC14 | 1.011133429 | 0.43309308 | 0.01597339 | 0.36341875 |
| UTP4 | 1.011067374 | 0.40340793 | 0.01587914 | 0.39425557 |

| | | | | |
|---|---|---|---|---|
| ITPA | 1.009729609 | 0.39614144 | 0.01396901 | 0.40214972 |
| DOP1B | 1.009602708 | 0.81381803 | 0.01378769 | 0.08947269 |
| DDX17 | 1.008666595 | 0.99221475 | 0.01244938 | 0.00339432 |
| PTK2 | 1.00791449 | 0.56288885 | 0.01137325 | 0.24957736 |
| EIF3K | 1.007871759 | 0.92445714 | 0.01131208 | 0.03411322 |
| ARAP1 | 1.007230804 | 0.68670383 | 0.01039431 | 0.16323053 |
| ATG4B | 1.00617723 | 0.74862273 | 0.00888445 | 0.12573699 |
| OFD1 | 1.006099353 | 0.69470685 | 0.00877278 | 0.15819842 |
| MON2 | 1.005740132 | 0.62028227 | 0.00825758 | 0.20741063 |
| RB1CC1 | 1.005050518 | 0.5411168 | 0.00726802 | 0.26670898 |
| TRIM27 | 1.003829658 | 0.74600088 | 0.00551448 | 0.12726066 |
| XPO1 | 1.003743714 | 0.64732791 | 0.00539095 | 0.18887567 |
| MAP2K7 | 1.003683437 | 0.47984613 | 0.00530431 | 0.318898 |
| RAD17 | 1.003247985 | 0.61031717 | 0.00467826 | 0.21444441 |
| PRIM1 | 1.002681733 | 0.44110328 | 0.00386374 | 0.35545972 |
| SNTB2 | 1.00233365 | 0.73892504 | 0.00336282 | 0.13139962 |
| SFPQ | 1.001551049 | 0.57569313 | 0.00223596 | 0.23980895 |
| NOP58 | 1.001346106 | 0.85902821 | 0.00194072 | 0.06599258 |
| DDX1 | 1.00073407 | 0.63918573 | 0.00105865 | 0.19437293 |
| CKS2 | 1 | 0.86173586 | 0 | 0.06462583 |
| FAAH | 1 | 0.37390097 | 0 | 0.42724341 |
| DIMT1 | 0.998755157 | 0.77120745 | -0.001797 | 0.11282879 |
| TM9SF3 | 0.998664496 | 0.52110975 | -0.001928 | 0.2830708 |
| TRAPPC11 | 0.99813695 | 0.89559841 | -0.0026903 | 0.04788669 |
| PRKACB | 0.997900475 | 0.82185577 | -0.0030322 | 0.08520439 |
| KRT20 | 0.997344827 | 0.96531148 | -0.0038357 | 0.01533253 |
| FYTTD1 | 0.9972191 | 0.695554 | -0.0040176 | 0.15766915 |
| LMNA | 0.99713383 | 0.35496462 | -0.0041409 | 0.44981493 |
| RING1 | 0.996340277 | 0.51416567 | -0.0052895 | 0.28889693 |
| MAPK8IP3 | 0.99483599 | 0.52903913 | -0.0074694 | 0.2765122 |
| PUM2 | 0.99445424 | 0.25537797 | -0.0080231 | 0.59281657 |
| GOLGA7 | 0.99381714 | 0.7244057 | -0.0089477 | 0.14001814 |
| DNAJC7 | 0.992734322 | 0.48538614 | -0.0105204 | 0.31391263 |
| RANGAP1 | 0.992584677 | 0.94352552 | -0.0107379 | 0.02524635 |
| TOM1L2 | 0.990238552 | 0.31278551 | -0.014152 | 0.50475338 |
| PRKRA | 0.9902109 | 0.35797579 | -0.0141923 | 0.44614635 |
| KNOP1 | 0.990078787 | 0.9640709 | -0.0143848 | 0.01589103 |
| TFAP2A | 0.989447099 | 0.94595432 | -0.0153055 | 0.02412984 |
| CMAS | 0.989148673 | 0.7365879 | -0.0157407 | 0.13277542 |
| MARCKSL1 | 0.987784287 | 0.51526193 | -0.0177321 | 0.28797194 |
| NOP2 | 0.985214717 | 0.37762521 | -0.0214899 | 0.42293902 |
| TIPRL | 0.985198565 | 0.75613737 | -0.0215136 | 0.12139929 |
| OPA1 | 0.984767788 | 0.49347337 | -0.0221445 | 0.30673628 |
| ZNF131 | 0.984439779 | 0.80540733 | -0.0226251 | 0.09398442 |
| PFDN5 | 0.984000723 | 0.85914918 | -0.0232687 | 0.06593142 |

| | | | | |
|---|---|---|---|---|
| CD320 | 0.983597815 | 0.96353821 | -0.0238596 | 0.01613106 |
| MYO1D | 0.983503408 | 0.85327452 | -0.023998 | 0.06891122 |
| RRP12 | 0.983497555 | 0.91347416 | -0.0240066 | 0.03930373 |
| UNC13B | 0.980971691 | 0.48402614 | -0.0277166 | 0.31513118 |
| CTIF | 0.980939413 | 0.61176069 | -0.0277641 | 0.21341844 |
| UTP6 | 0.97950983 | 0.47418803 | -0.0298681 | 0.32404941 |
| CPOX | 0.978819286 | 0.25393176 | -0.0308856 | 0.59528297 |
| CTBP1 | 0.976189595 | 0.28455929 | -0.0347667 | 0.54582724 |
| RNF5 | 0.975418211 | 0.75746603 | -0.0359072 | 0.12063684 |
| ACACA | 0.974579346 | 0.54190466 | -0.0371484 | 0.26607711 |
| CNOT7 | 0.97430318 | 0.77220133 | -0.0375573 | 0.11226945 |
| AASDHPPT | 0.974262768 | 0.99620798 | -0.0376172 | 0.00164998 |
| ZNF184 | 0.973831892 | 0.50072269 | -0.0382553 | 0.30040273 |
| EIF3D | 0.973450486 | 0.91017056 | -0.0388205 | 0.04087721 |
| AIP | 0.969372783 | 0.44836702 | -0.0448765 | 0.34836634 |
| PDIA6 | 0.966949968 | 0.76582962 | -0.0484869 | 0.11586784 |
| LENG1 | 0.966786392 | 0.42801202 | -0.0487309 | 0.36854404 |
| PPM1B | 0.965400856 | 0.38244286 | -0.0508 | 0.41743344 |
| ERCC6L | 0.965183433 | 0.63299061 | -0.0511249 | 0.19860273 |
| PFN2 | 0.965152084 | 0.92979599 | -0.0511718 | 0.03161233 |
| ABL1 | 0.965072357 | 0.85843325 | -0.051291 | 0.06629347 |
| DVL2 | 0.963812128 | 0.63539649 | -0.0531761 | 0.19695519 |
| TRAPPC12 | 0.963212089 | 0.29781025 | -0.0540746 | 0.52606035 |
| SRSF11 | 0.963093468 | 0.87222444 | -0.0542523 | 0.05937175 |
| MBOAT2 | 0.962808265 | 0.45615977 | -0.0546796 | 0.34088302 |
| HOOK1 | 0.962696902 | 0.79213396 | -0.0548464 | 0.10120137 |
| DAP | 0.961563561 | 0.47465038 | -0.0565459 | 0.32362617 |
| PPP1R14B | 0.961294211 | 0.56268992 | -0.05695 | 0.24973087 |
| EIF2A | 0.961087898 | 0.94544552 | -0.0572597 | 0.02436349 |
| EIF2AK2 | 0.958165359 | 0.86872964 | -0.0616534 | 0.06111536 |
| PRKG2 | 0.957526447 | 0.74014765 | -0.0626158 | 0.13068164 |
| DDX47 | 0.957258627 | 0.55765183 | -0.0630193 | 0.25363686 |
| VPS37B | 0.956905657 | 0.58514287 | -0.0635514 | 0.23273808 |
| ELOB | 0.95680261 | 0.81764203 | -0.0637068 | 0.08743679 |
| ARHGAP21 | 0.95670957 | 0.80184569 | -0.0638471 | 0.0959092 |
| AKAP8 | 0.956436474 | 0.90409251 | -0.0642589 | 0.04378713 |
| UBR7 | 0.955426173 | 0.32482805 | -0.0657837 | 0.48834648 |
| PRKAB2 | 0.954396985 | 0.93603523 | -0.0673386 | 0.0287078 |
| CHORDC1 | 0.953985739 | 0.95261764 | -0.0679604 | 0.02108138 |
| ARFGEF2 | 0.953686077 | 0.5168046 | -0.0684136 | 0.28667363 |
| TBC1D2B | 0.953323036 | 0.30300581 | -0.0689629 | 0.51854904 |
| PLK1 | 0.953169051 | 0.88490918 | -0.069196 | 0.0531013 |
| ATAD2B | 0.952592325 | 0.23458821 | -0.0700692 | 0.62969383 |
| COG4 | 0.952563801 | 0.73369304 | -0.0701124 | 0.1344856 |
| MTDH | 0.952419677 | 0.4260648 | -0.0703307 | 0.37052434 |

| | | | | |
|---|---|---|---|---|
| EGLN1 | 0.951221719 | 0.84293532 | -0.0721464 | 0.07420575 |
| NCOA3 | 0.951166839 | 0.17985111 | -0.0722297 | 0.74508687 |
| CDK5RAP2 | 0.950744115 | 0.79524907 | -0.072871 | 0.09949683 |
| TOE1 | 0.950274891 | 0.38746566 | -0.0735832 | 0.41176679 |
| RMI2 | 0.9492537 | 0.69950566 | -0.0751344 | 0.15520877 |
| EIF2B4 | 0.948669427 | 0.98794678 | -0.0760226 | 0.00526645 |
| KLC2 | 0.948534795 | 0.33521372 | -0.0762274 | 0.47467821 |
| INPP1 | 0.9484422 | 0.38745425 | -0.0763682 | 0.41177957 |
| TTK | 0.947700612 | 0.77319967 | -0.0774967 | 0.11170834 |
| IGBP1 | 0.945616785 | 0.18671918 | -0.0806725 | 0.72881106 |
| PI4K2B | 0.944709329 | 0.45437096 | -0.0820576 | 0.34258943 |
| PTBP3 | 0.943732034 | 0.98212697 | -0.0835508 | 0.00783236 |
| GRK6 | 0.942623204 | 0.48825418 | -0.0852469 | 0.31135403 |
| SYF2 | 0.942460832 | 0.41640543 | -0.0854954 | 0.38048361 |
| RRM2B | 0.941744777 | 0.93762757 | -0.086592 | 0.02796963 |
| FMNL2 | 0.941183667 | 0.41953244 | -0.0874518 | 0.37723445 |
| NELFCD | 0.941080499 | 0.68684487 | -0.08761 | 0.16314134 |
| RAP1GAP2 | 0.940933598 | 0.50376043 | -0.0878352 | 0.29777595 |
| TMA16 | 0.940867827 | 0.72840991 | -0.087936 | 0.13762416 |
| NAA15 | 0.940858437 | 0.87990297 | -0.0879504 | 0.05556521 |
| AGPAT1 | 0.940184752 | 0.80984954 | -0.0889838 | 0.09159566 |
| RPAP3 | 0.93928826 | 0.34727411 | -0.0903601 | 0.4593276 |
| UCKL1 | 0.939281686 | 0.85284719 | -0.0903702 | 0.06912878 |
| HSPA8 | 0.938011881 | 0.46893667 | -0.0923219 | 0.32888581 |
| EEF1A1 | 0.937582504 | 0.8322184 | -0.0929824 | 0.07976269 |
| CCDC59 | 0.936971802 | 0.8498596 | -0.0939225 | 0.07065282 |
| PPT1 | 0.936674675 | 0.95704861 | -0.09438 | 0.019066 |
| MICALL1 | 0.936609767 | 0.58385754 | -0.09448 | 0.23369311 |
| STK24 | 0.936166658 | 0.72488158 | -0.0951627 | 0.13973294 |
| RNASEH2C | 0.93586674 | 0.93908825 | -0.095625 | 0.02729359 |
| SPC24 | 0.934671398 | 0.82168505 | -0.0974688 | 0.08529462 |
| C2CD3 | 0.934388404 | 0.63303894 | -0.0979057 | 0.19856958 |
| WDR7 | 0.934288189 | 0.72147962 | -0.0980605 | 0.14177593 |
| UFSP2 | 0.932914924 | 0.796124 | -0.1001826 | 0.09901929 |
| SYNE2 | 0.932822895 | 0.37327274 | -0.1003249 | 0.42797373 |
| CBL | 0.932686622 | 0.50846828 | -0.1005357 | 0.29373614 |
| PUS1 | 0.93243012 | 0.78511002 | -0.1009325 | 0.10506948 |
| KIF5C | 0.93204943 | 0.26291556 | -0.1015216 | 0.58018371 |
| UTP3 | 0.931815888 | 0.3991827 | -0.1018832 | 0.39882829 |
| CLCN5 | 0.93055697 | 0.91364157 | -0.1038336 | 0.03922415 |
| TOP3B | 0.930509701 | 0.91489444 | -0.1039069 | 0.03862901 |
| TUBA1A | 0.928802269 | 0.51283937 | -0.1065566 | 0.29001864 |
| SYNE3 | 0.92798656 | 0.45896956 | -0.1078242 | 0.33821611 |
| STEEP1 | 0.927066482 | 0.49328864 | -0.1092553 | 0.30689889 |
| CCDC22 | 0.926285156 | 0.62233742 | -0.1104717 | 0.20597409 |

| | | | | |
|---|---|---|---|---|
| CLIC6 | 0.926269528 | 0.81665364 | -0.110496 | 0.0879621 |
| PIK3R2 | 0.926166769 | 0.54029508 | -0.1106561 | 0.26736899 |
| GLUL | 0.925880752 | 0.25469719 | -0.1111017 | 0.59397585 |
| ATP13A3 | 0.925641333 | 0.68361309 | -0.1114748 | 0.16518963 |
| PEPD | 0.924839887 | 0.62899374 | -0.1127245 | 0.20135367 |
| ADCK1 | 0.924366645 | 0.46212927 | -0.1134629 | 0.33523652 |
| PTCD3 | 0.924303543 | 0.74353619 | -0.1135614 | 0.12869789 |
| NDC1 | 0.923894215 | 0.6499344 | -0.1142004 | 0.18713048 |
| CRMP1 | 0.923307062 | 0.47415802 | -0.1151176 | 0.3240769 |
| PDCD5 | 0.923002399 | 0.00278219 | -0.1155937 | 2.55561244 |
| OSBP | 0.922696665 | 0.94414745 | -0.1160717 | 0.02496017 |
| GFPT1 | 0.92201946 | 0.605792 | -0.1171309 | 0.21767646 |
| CSNK1G2 | 0.921439152 | 0.46139461 | -0.1180392 | 0.33592748 |
| UBE2I | 0.921320788 | 0.63729303 | -0.1182245 | 0.19566083 |
| NSD2 | 0.920678787 | 0.34949591 | -0.1192302 | 0.4565579 |
| ALX1 | 0.919681125 | 0.77013855 | -0.1207944 | 0.11343114 |
| RO60 | 0.919256598 | 0.46704777 | -0.1214605 | 0.3306387 |
| NISCH | 0.918531324 | 0.76892554 | -0.1225992 | 0.11411571 |
| PSMC4 | 0.917763577 | 0.34997518 | -0.1238055 | 0.45596275 |
| UCK1 | 0.917385417 | 0.8924994 | -0.1244001 | 0.04939207 |
| PPM1F | 0.915529311 | 0.75800785 | -0.127322 | 0.12032629 |
| CTPS2 | 0.91542318 | 0.60037312 | -0.1274893 | 0.22157876 |
| C12orf4 | 0.91218066 | 0.88769947 | -0.1326085 | 0.05173404 |
| INTS12 | 0.911253001 | 0.08516389 | -0.1340764 | 1.06974453 |
| MIS18A | 0.910841376 | 0.11916363 | -0.1347283 | 0.92385628 |
| VPS18 | 0.910651386 | 0.1408061 | -0.1350292 | 0.85137852 |
| EIF2B1 | 0.910622426 | 0.4013853 | -0.1350751 | 0.39643854 |
| CPSF3 | 0.910543612 | 0.34167389 | -0.1352 | 0.46638821 |
| CUL4A | 0.908886764 | 0.14372404 | -0.1378275 | 0.84247059 |
| VASP | 0.908832907 | 0.41621992 | -0.137913 | 0.38067713 |
| ERLIN2 | 0.908132859 | 0.95117115 | -0.1390247 | 0.02174133 |
| CASKIN2 | 0.907365642 | 0.38373492 | -0.1402441 | 0.41596868 |
| SNU13 | 0.907336637 | 0.17172362 | -0.1402902 | 0.76516997 |
| CARHSP1 | 0.906706897 | 0.73770222 | -0.1412918 | 0.13211891 |
| RUVBL2 | 0.905522077 | 0.34906399 | -0.1431783 | 0.45709495 |
| LRSAM1 | 0.905515624 | 0.51385499 | -0.1431886 | 0.28915942 |
| DEK | 0.904696201 | 0.58488629 | -0.1444947 | 0.23292856 |
| EMC8 | 0.903181547 | 0.57110332 | -0.1469121 | 0.24328531 |
| ZNF639 | 0.900824134 | 0.34221486 | -0.1506826 | 0.46570113 |
| ACTN4 | 0.900742748 | 0.89113855 | -0.150813 | 0.05005477 |
| STOM | 0.900027139 | 0.73665344 | -0.1519596 | 0.13273678 |
| MBOAT7 | 0.899287541 | 0.3259215 | -0.1531456 | 0.48688698 |
| DPYSL5 | 0.897222176 | 0.13397673 | -0.1564628 | 0.87297064 |
| VPS35L | 0.896989914 | 0.22106219 | -0.1568363 | 0.65548553 |
| SMNDC1 | 0.89520207 | 0.94909354 | -0.1597147 | 0.02269098 |

| | | | | |
|---|---|---|---|---|
| MCM7 | 0.894933571 | 0.07620656 | -0.1601475 | 1.11800763 |
| FBXO45 | 0.894428551 | 0.80211699 | -0.1609619 | 0.09576228 |
| TUBB3 | 0.894418669 | 0.66477228 | -0.1609778 | 0.1773271 |
| ERP29 | 0.89303292 | 0.36865742 | -0.1632147 | 0.43337702 |
| CDIPT | 0.89292186 | 0.46657064 | -0.1633942 | 0.3310826 |
| GTF2A2 | 0.891138453 | 0.25750578 | -0.1662785 | 0.58921302 |
| DHX15 | 0.891115133 | 0.07238563 | -0.1663163 | 1.14034762 |
| CFL1 | 0.88909377 | 0.61677339 | -0.1695925 | 0.20987437 |
| RNF214 | 0.887441151 | 0.13713597 | -0.1722766 | 0.86284862 |
| TMSB10 | 0.887395014 | 0.13748729 | -0.1723516 | 0.86173745 |
| TMEM59 | 0.886827651 | 0.26737974 | -0.1732743 | 0.5728715 |
| GPATCH4 | 0.886326779 | 0.89584444 | -0.1740894 | 0.0477674 |
| MBD1 | 0.885375885 | 0.12382608 | -0.175638 | 0.90718789 |
| TOMM34 | 0.884906468 | 0.03947395 | -0.1764031 | 1.40368937 |
| DARS1 | 0.88461614 | 0.29570164 | -0.1768765 | 0.52914626 |
| INTS7 | 0.883284152 | 0.23618115 | -0.1790505 | 0.62675477 |
| | 0.882995783 | 0.58544804 | -0.1795215 | 0.23251164 |
| POP1 | 0.882841647 | 0.98595437 | -0.1797734 | 0.00614318 |
| CSE1L | 0.882310709 | 0.84105959 | -0.1806413 | 0.07517323 |
| RAD54B | 0.881493424 | 0.16126422 | -0.1819783 | 0.79246199 |
| SEC31A | 0.88029283 | 0.8413078 | -0.1839446 | 0.07504508 |
| MBNL2 | 0.880143738 | 0.23454708 | -0.1841889 | 0.62976997 |
| THOC6 | 0.879453483 | 0.40697893 | -0.1853208 | 0.39042808 |
| GNA11 | 0.87907913 | 0.80936114 | -0.1859351 | 0.09185765 |
| CTPS1 | 0.87831267 | 0.70996935 | -0.1871935 | 0.1487604 |
| C9orf40 | 0.87826728 | 0.28099954 | -0.187268 | 0.55129439 |
| PTAR1 | 0.877540002 | 0.24022773 | -0.1884632 | 0.61937686 |
| GMPS | 0.875881981 | 0.58678435 | -0.1911916 | 0.23152148 |
| HSPA4L | 0.874112363 | 0.12578722 | -0.1941094 | 0.90036349 |
| RNF169 | 0.873404778 | 0.39775274 | -0.1952777 | 0.40038683 |
| RAB3IP | 0.872241094 | 0.30279894 | -0.1972011 | 0.51884564 |
| ATP11C | 0.871621957 | 0.20717894 | -0.1982256 | 0.68365438 |
| IRAK4 | 0.870559699 | 0.60928848 | -0.1999849 | 0.21517703 |
| CAMSAP1 | 0.870156026 | 0.29519021 | -0.200654 | 0.52989804 |
| ADAR | 0.870095419 | 0.36187097 | -0.2007545 | 0.44144626 |
| RASA4 | 0.869586917 | 0.06212699 | -0.2015979 | 1.20671965 |
| NAV2 | 0.868934694 | 0.42420879 | -0.2026803 | 0.37242033 |
| WASHC5 | 0.868785195 | 0.22782396 | -0.2029286 | 0.6424006 |
| FIBP | 0.868668055 | 0.33217931 | -0.2031231 | 0.47862743 |
| WRNIP1 | 0.8685275 | 0.03603671 | -0.2033566 | 1.44325492 |
| PPP6R3 | 0.868209295 | 0.81996348 | -0.2038852 | 0.08620549 |
| ANP32E | 0.867989394 | 0.20391985 | -0.2042507 | 0.6905405 |
| POU3F1 | 0.867979992 | 0.42345181 | -0.2042663 | 0.37319601 |
| DHX9 | 0.867882326 | 0.54055495 | -0.2044287 | 0.26716015 |
| ACOT8 | 0.867746642 | 0.88281871 | -0.2046542 | 0.05412847 |

| | | | | |
|---|---|---|---|---|
| NF1 | 0.866333147 | 0.19729256 | -0.2070062 | 0.7048893 |
| DHX30 | 0.865119792 | 0.19304035 | -0.2090282 | 0.7143519 |
| PHF20L1 | 0.86498671 | 0.62702816 | -0.2092501 | 0.20271295 |
| TXNIP | 0.864468509 | 0.5051148 | -0.2101147 | 0.29660991 |
| ZCCHC2 | 0.863113949 | 0.43685835 | -0.2123771 | 0.35965936 |
| HNRNPK | 0.863069341 | 0.87995208 | -0.2124516 | 0.05554098 |
| STK38 | 0.856150098 | 0.85817599 | -0.2240643 | 0.06642364 |
| NANS | 0.8554487 | 0.39677596 | -0.2252468 | 0.40145465 |
| SLC27A4 | 0.855200751 | 0.90616834 | -0.225665 | 0.04279112 |
| KRT8 | 0.855083036 | 0.27632619 | -0.2258636 | 0.55857795 |
| CSTF1 | 0.854786918 | 0.42083316 | -0.2263633 | 0.37589005 |
| AIFM1 | 0.853399351 | 0.95551964 | -0.2287071 | 0.01976038 |
| PRKAR1A | 0.852925864 | 0.32528114 | -0.2295077 | 0.48774112 |
| ABCB6 | 0.852795565 | 0.90708369 | -0.2297282 | 0.04235264 |
| TRAPPC8 | 0.852775877 | 0.11280257 | -0.2297615 | 0.94768102 |
| RAD50 | 0.85234135 | 0.25010073 | -0.2304968 | 0.60188504 |
| STXBP2 | 0.852196648 | 0.10441136 | -0.2307417 | 0.98125224 |
| C2CD5 | 0.852196353 | 0.25005408 | -0.2307422 | 0.60196606 |
| DMXL2 | 0.85106503 | 0.32290738 | -0.2326587 | 0.49092203 |
| DYNLL1 | 0.8498941 | 0.96359268 | -0.234645 | 0.01610651 |
| ANKHD1 | 0.849823162 | 0.21162975 | -0.2347654 | 0.67442329 |
| MFHAS1 | 0.849705573 | 0.7735684 | -0.2349651 | 0.11150128 |
| WEE1 | 0.848911701 | 0.71260775 | -0.2363136 | 0.14714946 |
| SRP72 | 0.848434343 | 0.36467696 | -0.2371251 | 0.43809167 |
| PPFIA3 | 0.846009854 | 0.29965715 | -0.2412536 | 0.52337536 |
| PPP4R2 | 0.845428466 | 0.31126785 | -0.2422454 | 0.50686573 |
| NSF | 0.845135634 | 0.08722953 | -0.2427452 | 1.05933646 |
| PSMC2 | 0.841737697 | 0.14737942 | -0.2485574 | 0.83156314 |
| FXR1 | 0.841217033 | 0.09723735 | -0.24945 | 1.01216686 |
| AKAP8L | 0.840802544 | 0.16985107 | -0.2501611 | 0.76993172 |
| SUGP2 | 0.840278223 | 0.46535396 | -0.251061 | 0.33221658 |
| RFC5 | 0.838782603 | 0.40496912 | -0.2536312 | 0.39257809 |
| PRKCI | 0.838736578 | 0.69639339 | -0.2537103 | 0.15714536 |
| CDK2 | 0.838167146 | 0.40034196 | -0.2546901 | 0.39756889 |
| AHDC1 | 0.837705049 | 0.45327219 | -0.2554857 | 0.34364093 |
| PPP1R2 | 0.837664119 | 0.33029076 | -0.2555562 | 0.48110357 |
| ORC4 | 0.836969396 | 0.12840457 | -0.2567532 | 0.89141951 |
| VPS50 | 0.836923386 | 0.17924395 | -0.2568325 | 0.74655548 |
| GGA2 | 0.835654588 | 0.27998844 | -0.2590214 | 0.5528599 |
| WDR76 | 0.83540077 | 0.05198757 | -0.2594596 | 1.28410049 |
| GSN | 0.835361619 | 0.47982347 | -0.2595272 | 0.31891851 |
| TSPYL5 | 0.835356822 | 0.33800757 | -0.2595355 | 0.47107357 |
| CIAPIN1 | 0.834482328 | 0.63803725 | -0.2610466 | 0.19515396 |
| AKAP17A | 0.834047072 | 0.16276027 | -0.2617993 | 0.78845159 |
| POLR2C | 0.83398767 | 0.61033205 | -0.261902 | 0.21443382 |

| | | | | |
|---|---|---|---|---|
| SNAPIN | 0.833571356 | 0.31998602 | -0.2626224 | 0.494869 |
| BRMS1 | 0.833492388 | 0.3689052 | -0.2627591 | 0.43308522 |
| ANKRD17 | 0.832961482 | 0.26917122 | -0.2636783 | 0.56997137 |
| GCC2 | 0.832519464 | 0.35739556 | -0.2644441 | 0.44685085 |
| HDAC6 | 0.832303112 | 0.13928748 | -0.2648191 | 0.85608793 |
| NUP188 | 0.832289662 | 0.17098453 | -0.2648424 | 0.76704318 |
| ARPP19 | 0.831898702 | 0.67077363 | -0.2655202 | 0.17342402 |
| OAS3 | 0.831518799 | 0.71266531 | -0.2661792 | 0.14711438 |
| UNC119B | 0.831439587 | 0.36946004 | -0.2663167 | 0.43243252 |
| SEPTIN7 | 0.831432239 | 0.03130134 | -0.2663294 | 1.50443706 |
| STAT2 | 0.831384596 | 0.11246831 | -0.2664121 | 0.94896983 |
| SCFD1 | 0.830959098 | 0.12829187 | -0.2671506 | 0.89180087 |
| PSMC1 | 0.829376582 | 0.09767428 | -0.2699008 | 1.0102198 |
| IQGAP1 | 0.829197213 | 0.22314622 | -0.2702128 | 0.65141047 |
| GTPBP4 | 0.82901761 | 0.11522148 | -0.2705253 | 0.93846654 |
| SESTD1 | 0.82868597 | 0.09368199 | -0.2711026 | 1.02834388 |
| SMARCAD1 | 0.828653839 | 0.05202388 | -0.2711585 | 1.28379724 |
| DPYSL3 | 0.82811389 | 0.25672428 | -0.2720989 | 0.59053306 |
| FAM98B | 0.825932506 | 0.57795063 | -0.2759042 | 0.23810926 |
| PREB | 0.825612072 | 0.47118208 | -0.276464 | 0.32681124 |
| EXOC4 | 0.824827476 | 0.18618489 | -0.2778357 | 0.73005557 |
| HAUS7 | 0.824444675 | 0.52232679 | -0.2785054 | 0.2820577 |
| HDAC4 | 0.822101368 | 0.55816883 | -0.2826118 | 0.25323442 |
| U2AF2 | 0.821828258 | 0.58675357 | -0.2830912 | 0.23154426 |
| KIF2A | 0.821078988 | 0.5528829 | -0.2844071 | 0.25736684 |
| DSTN | 0.820522064 | 0.23460865 | -0.285386 | 0.62965599 |
| RUVBL1 | 0.82004512 | 0.06781358 | -0.2862248 | 1.1686833 |
| UFL1 | 0.819447779 | 0.4394942 | -0.2872761 | 0.35704685 |
| HNRNPLL | 0.819438514 | 0.47748776 | -0.2872924 | 0.32103775 |
| PDIA4 | 0.818986064 | 0.57610388 | -0.2880892 | 0.2394992 |
| PPME1 | 0.818716023 | 0.05635577 | -0.288565 | 1.24906163 |
| EEFSEC | 0.817986156 | 0.17547837 | -0.2898517 | 0.7557764 |
| XPOT | 0.816682785 | 0.76709039 | -0.2921523 | 0.11515346 |
| SPG7 | 0.816464776 | 0.39333534 | -0.2925374 | 0.40523703 |
| PPP2R5D | 0.816216938 | 0.30991173 | -0.2929754 | 0.50876199 |
| WASF1 | 0.816211567 | 0.25238773 | -0.2929849 | 0.59793177 |
| MAPK3 | 0.816175047 | 0.00886902 | -0.2930495 | 2.05212451 |
| NRDE2 | 0.816071101 | 0.30399668 | -0.2932332 | 0.51713116 |
| PHRF1 | 0.815665271 | 0.12426807 | -0.2939509 | 0.90564046 |
| LYSMD1 | 0.815291176 | 0.81247804 | -0.2946127 | 0.09018837 |
| OTUD3 | 0.815035854 | 0.51959982 | -0.2950646 | 0.28433101 |
| UGCG | 0.814738836 | 0.95297779 | -0.2955904 | 0.02091722 |
| COPA | 0.814713289 | 0.48490368 | -0.2956357 | 0.31434452 |
| DBN1 | 0.812488603 | 0.06379551 | -0.2995805 | 1.19520986 |
| RPS23 | 0.812203844 | 0.39079245 | -0.3000862 | 0.40805383 |

| | | | | |
|---|---|---|---|---|
| PFKL | 0.811495031 | 0.2833401 | -0.3013458 | 0.54769195 |
| SERINC1 | 0.807349866 | 0.4470579 | -0.3087341 | 0.34963622 |
| ISG20L2 | 0.806979288 | 0.21516373 | -0.3093964 | 0.66723094 |
| TAF10 | 0.80653744 | 0.15405854 | -0.3101866 | 0.81231423 |
| NUDC | 0.806435298 | 0.14035119 | -0.3103693 | 0.8527839 |
| RPS12 | 0.806128488 | 0.38919317 | -0.3109183 | 0.40983479 |
| SARS1 | 0.806102626 | 0.0685357 | -0.3109646 | 1.16408315 |
| NRBP2 | 0.805459442 | 0.50674893 | -0.3121161 | 0.29520716 |
| PIR | 0.805447943 | 0.06277377 | -0.3121367 | 1.20222181 |
| GOLGA2 | 0.805271558 | 0.09398972 | -0.3124527 | 1.02691963 |
| CDC23 | 0.804565055 | 0.01515919 | -0.313719 | 1.81932395 |
| PPIL2 | 0.803057191 | 0.01843306 | -0.3164254 | 1.73440257 |
| USP19 | 0.803037445 | 0.2060421 | -0.3164608 | 0.68604403 |
| EFTUD2 | 0.803023915 | 0.00295517 | -0.3164851 | 2.52941794 |
| CHMP4B | 0.802482418 | 0.76400057 | -0.3174583 | 0.11690632 |
| MRPS11 | 0.801941675 | 0.02378343 | -0.3184308 | 1.62372555 |
| CACTIN | 0.800039443 | 0.04485657 | -0.321857 | 1.34817392 |
| PSMD2 | 0.799065163 | 0.04243961 | -0.3236149 | 1.37222862 |
| CHMP5 | 0.798556126 | 0.05878935 | -0.3245343 | 1.23070137 |
| DDX19A | 0.79737524 | 0.26971685 | -0.3266693 | 0.56909192 |
| GPSM1 | 0.796518461 | 0.45831634 | -0.3282203 | 0.33883466 |
| SREK1 | 0.796430316 | 0.00610473 | -0.32838 | 2.2143337 |
| EIF3F | 0.796319141 | 0.04304585 | -0.3285814 | 1.36606874 |
| FRYL | 0.795472965 | 0.15591952 | -0.3301152 | 0.80709952 |
| KPNA2 | 0.79481503 | 0.09866462 | -0.3313089 | 1.00583856 |
| HAUS2 | 0.793949255 | 0.17186727 | -0.3328813 | 0.76480683 |
| MICAL3 | 0.793253989 | 0.45594277 | -0.3341452 | 0.34108967 |
| SSH1 | 0.793053571 | 0.21582774 | -0.3345098 | 0.66589274 |
| TMEM106B | 0.791990931 | 0.59395776 | -0.3364442 | 0.22624444 |
| WIPI2 | 0.791929219 | 0.52870904 | -0.3365566 | 0.27678327 |
| DYRK1A | 0.79053975 | 0.27467061 | -0.3390901 | 0.56118781 |
| NDC80 | 0.789201038 | 0.02340231 | -0.3415352 | 1.63074131 |
| GGA1 | 0.788783722 | 0.18832174 | -0.3422983 | 0.72509954 |
| SRP14 | 0.788645042 | 0.4000305 | -0.342552 | 0.39790689 |
| PRRC2A | 0.788572147 | 0.03691837 | -0.3426853 | 1.43275748 |
| STT3B | 0.78853684 | 0.22206068 | -0.3427499 | 0.65352833 |
| CUL1 | 0.787863096 | 0.14493278 | -0.3439831 | 0.83883339 |
| ABCF2 | 0.787795684 | 0.12099445 | -0.3441066 | 0.91723456 |
| SENP5 | 0.787710816 | 0.53600314 | -0.344262 | 0.27083266 |
| ASPM | 0.787090222 | 0.56743467 | -0.3453991 | 0.24608414 |
| ESF1 | 0.786063186 | 0.03067701 | -0.3472828 | 1.51318701 |
| SLBP | 0.785199771 | 0.21397638 | -0.3488683 | 0.66963417 |
| TENT4B | 0.784618596 | 0.07959378 | -0.3499366 | 1.09912089 |
| ATP5F1D | 0.784426503 | 0.00830143 | -0.3502898 | 2.08084692 |
| CRTC3 | 0.784095717 | 0.05510212 | -0.3508983 | 1.25883172 |

| | | | | |
|---|---|---|---|---|
| BAIAP2L1 | 0.783562285 | 0.07616559 | -0.3518801 | 1.11824121 |
| UBFD1 | 0.783099138 | 0.13727386 | -0.3527331 | 0.86241214 |
| RBM22 | 0.782702031 | 0.00817386 | -0.3534649 | 2.08757283 |
| SWAP70 | 0.782267764 | 0.10267771 | -0.3542656 | 0.98852382 |
| ATL1 | 0.7811075 | 0.12957939 | -0.356407 | 0.88746406 |
| TRIP13 | 0.780996229 | 0.08341622 | -0.3566125 | 1.07874951 |
| RAPGEF6 | 0.780857069 | 0.16522813 | -0.3568696 | 0.78191601 |
| AP3M1 | 0.779935432 | 0.39229857 | -0.3585734 | 0.40638328 |
| PIGX | 0.778152588 | 0.30412896 | -0.361875 | 0.51694223 |
| KBTBD4 | 0.777275719 | 0.17028032 | -0.3635016 | 0.76883553 |
| PPP4C | 0.777266393 | 0.22856062 | -0.363519 | 0.6409986 |
| CLCN7 | 0.776592176 | 0.37085084 | -0.3647709 | 0.43080074 |
| RFC2 | 0.776453613 | 0.08497181 | -0.3650284 | 1.07072516 |
| VCP | 0.775823036 | 0.20026464 | -0.3662005 | 0.69839573 |
| RHOT2 | 0.775609823 | 0.45425729 | -0.366597 | 0.3426981 |
| SOGA1 | 0.775581328 | 0.0434386 | -0.36665 | 1.36212417 |
| TLN1 | 0.774087526 | 0.28446444 | -0.3694314 | 0.54597202 |
| ITM2B | 0.773899749 | 0.70691183 | -0.3697814 | 0.15063475 |
| TXNDC9 | 0.773634083 | 0.10764847 | -0.3702767 | 0.96799214 |
| SETD3 | 0.772731995 | 0.12994322 | -0.37196 | 0.88624636 |
| UBA3 | 0.771681173 | 0.29348264 | -0.3739232 | 0.53241759 |
| MYEF2 | 0.771345095 | 0.05356353 | -0.3745516 | 1.27113078 |
| GMIP | 0.770986011 | 0.09640563 | -0.3752234 | 1.01589759 |
| ARAF | 0.769353788 | 0.02218769 | -0.3782809 | 1.65388789 |
| CDK11B | 0.769264307 | 0.02983251 | -0.3784487 | 1.52531018 |
| MIOS | 0.767912985 | 0.37656126 | -0.3809853 | 0.42416437 |
| PSMC5 | 0.76782683 | 0.27378065 | -0.3811471 | 0.56259725 |
| PDZRN3 | 0.766859942 | 0.12950906 | -0.382965 | 0.88769986 |
| KNSTRN | 0.76670114 | 0.83849161 | -0.3832638 | 0.07650128 |
| FHIP1B | 0.766444801 | 0.35050243 | -0.3837462 | 0.45530897 |
| ADD1 | 0.766395901 | 0.27259612 | -0.3838382 | 0.56448034 |
| FCSK | 0.765793145 | 0.19343004 | -0.3849733 | 0.71347608 |
| SPTLC2 | 0.765147656 | 0.70006217 | -0.3861899 | 0.15486339 |
| UMPS | 0.764433862 | 0.27106444 | -0.3875364 | 0.56692745 |
| CAMK4 | 0.764076429 | 0.03198581 | -0.3882111 | 1.49504265 |
| BAZ1A | 0.763085261 | 0.09593875 | -0.3900838 | 1.01800595 |
| IGHMBP2 | 0.762955373 | 0.37249989 | -0.3903294 | 0.42887385 |
| RBM28 | 0.762928514 | 0.05226235 | -0.3903802 | 1.28181106 |
| ZNHIT6 | 0.762539189 | 0.34018187 | -0.3911166 | 0.46828884 |
| SYAP1 | 0.762301016 | 0.01738586 | -0.3915673 | 1.75980381 |
| RECQL5 | 0.761763276 | 0.03864084 | -0.3925854 | 1.41295348 |
| EIF1 | 0.760656622 | 0.79538304 | -0.3946828 | 0.09942368 |
| SIPA1L1 | 0.760449264 | 0.26232317 | -0.3950761 | 0.58116335 |
| AKR1B1 | 0.760281905 | 0.17284445 | -0.3953936 | 0.76234457 |
| VAC14 | 0.760025153 | 0.06619938 | -0.3958809 | 1.17914609 |

| | | | | |
|---|---|---|---|---|
| YEATS2 | 0.759480225 | 0.03149235 | -0.3969157 | 1.50179488 |
| PHLDB1 | 0.758468026 | 0.72298876 | -0.3988397 | 0.14086845 |
| ZNF800 | 0.758440789 | 0.37503012 | -0.3988915 | 0.42593386 |
| TPI1 | 0.758214015 | 0.23886529 | -0.399323 | 0.62184695 |
| WNK3 | 0.75807982 | 0.27456904 | -0.3995783 | 0.56134843 |
| TUT7 | 0.757763847 | 0.0153288 | -0.4001798 | 1.81449179 |
| MSL1 | 0.757429386 | 0.01240774 | -0.4008167 | 1.90630719 |
| INTS1 | 0.7565358 | 0.33382554 | -0.4025197 | 0.47648044 |
| RASAL1 | 0.756501526 | 0.74903526 | -0.4025851 | 0.12549774 |
| EIF3J | 0.756100907 | 0.03734755 | -0.4033493 | 1.42773784 |
| PIK3R4 | 0.754929266 | 0.2497821 | -0.4055866 | 0.60243869 |
| PPP2R5A | 0.754894651 | 0.40362914 | -0.4056528 | 0.39401749 |
| USO1 | 0.754015157 | 0.14308893 | -0.4073346 | 0.84439396 |
| TMOD3 | 0.753979272 | 0.05921459 | -0.4074032 | 1.22757127 |
| DR1 | 0.753587154 | 0.43983718 | -0.4081537 | 0.35670806 |
| CACYBP | 0.753329386 | 0.71355927 | -0.4086473 | 0.14656995 |
| MSH2 | 0.753204838 | 0.0334402 | -0.4088858 | 1.47573113 |
| TBC1D24 | 0.752700856 | 0.31071843 | -0.4098515 | 0.50763299 |
| INSM1 | 0.751621333 | 0.16091635 | -0.4119221 | 0.79339982 |
| SAMM50 | 0.751597528 | 0.403782 | -0.4119678 | 0.39385304 |
| SPTBN2 | 0.751088944 | 0.11239037 | -0.4129443 | 0.94927092 |
| SPAST | 0.750918122 | 0.18958378 | -0.4132725 | 0.72219883 |
| RRAGA | 0.749323833 | 0.15374576 | -0.4163388 | 0.81319685 |
| KRT80 | 0.749066395 | 0.00065237 | -0.4168345 | 3.18550544 |
| CHMP1A | 0.748990587 | 0.00790615 | -0.4169805 | 2.1020351 |
| CCT5 | 0.748152254 | 0.00481547 | -0.4185962 | 2.3173617 |
| ATP6V1H | 0.74802318 | 0.27549424 | -0.4188451 | 0.55988747 |
| CCDC47 | 0.747480886 | 0.13976286 | -0.4198914 | 0.85460822 |
| MRPL3 | 0.747324274 | 0.97235096 | -0.4201937 | 0.01217695 |
| SSR4 | 0.746897315 | 0.14348292 | -0.4210182 | 0.84319978 |
| MED12 | 0.7465792 | 0.03394042 | -0.4216328 | 1.46928274 |
| SLC39A7 | 0.746177384 | 0.07361642 | -0.4224095 | 1.13302531 |
| PAK2 | 0.746169141 | 0.06339972 | -0.4224254 | 1.19791268 |
| WDR44 | 0.746149558 | 0.49011598 | -0.4224633 | 0.30970113 |
| NXF1 | 0.743714935 | 0.03580075 | -0.4271783 | 1.44610782 |
| CCDC93 | 0.743040597 | 0.19032437 | -0.4284871 | 0.72050559 |
| DCAF10 | 0.742962287 | 0.26198534 | -0.4286391 | 0.58172301 |
| RP2 | 0.741829423 | 0.20210805 | -0.4308406 | 0.69441638 |
| MCC | 0.741560846 | 0.20217284 | -0.431363 | 0.69427718 |
| MRGBP | 0.740195752 | 0.11196925 | -0.4340212 | 0.95090122 |
| NME3 | 0.74012733 | 0.73800865 | -0.4341546 | 0.13193855 |
| MIEF1 | 0.739081648 | 0.3097597 | -0.4361943 | 0.50897508 |
| RAP1GDS1 | 0.738793489 | 0.90625306 | -0.4367569 | 0.04275051 |
| PLAA | 0.737788229 | 0.06944653 | -0.4387213 | 1.15834943 |
| IPO8 | 0.736947512 | 0.15925126 | -0.4403662 | 0.79791713 |

| | | | | |
|---|---|---|---|---|
| G6PD | 0.736812543 | 0.00776913 | -0.4406305 | 2.10962768 |
| LRRC8C | 0.736336419 | 0.01948041 | -0.441563 | 1.71040189 |
| HID1 | 0.736302562 | 0.02329078 | -0.4416294 | 1.63281603 |
| HMGCS1 | 0.736009208 | 0.48476528 | -0.4422043 | 0.31446849 |
| BRD1 | 0.735237016 | 0.33540561 | -0.4437187 | 0.47442968 |
| SPTLC1 | 0.735191714 | 0.54868643 | -0.4438076 | 0.26067578 |
| TECR | 0.734765007 | 0.24143061 | -0.4446452 | 0.61720766 |
| ARHGAP19 | 0.734451745 | 0.41392104 | -0.4452604 | 0.3830825 |
| MTREX | 0.733775267 | 0.03566373 | -0.4465898 | 1.44777323 |
| PRMT3 | 0.733081938 | 0.3584502 | -0.4479536 | 0.44557117 |
| PURB | 0.732927172 | 0.04133611 | -0.4482582 | 1.38367039 |
| TBCE | 0.732906936 | 0.54149433 | -0.4482981 | 0.26640609 |
| STMN1 | 0.732754035 | 0.0039867 | -0.4485991 | 2.39938687 |
| AURKB | 0.732656032 | 0.1435701 | -0.4487921 | 0.84293601 |
| SAAL1 | 0.732249908 | 0.22792624 | -0.449592 | 0.64220567 |
| ZNF668 | 0.731891179 | 0.35659382 | -0.4502989 | 0.44782619 |
| FEN1 | 0.73166543 | 0.03366779 | -0.450744 | 1.47278538 |
| COPG2 | 0.731455773 | 0.03113363 | -0.4511575 | 1.50677026 |
| ZC3H15 | 0.730254709 | 0.02123551 | -0.4535283 | 1.67293731 |
| GPI | 0.729799202 | 0.13839559 | -0.4544285 | 0.85887776 |
| ULK1 | 0.729647201 | 0.44222111 | -0.454729 | 0.35436053 |
| EED | 0.729252983 | 0.01912152 | -0.4555087 | 1.71847767 |
| FUS | 0.728702297 | 0.01970512 | -0.4565986 | 1.70542097 |
| VPS11 | 0.728405162 | 0.0600462 | -0.4571869 | 1.22151449 |
| TTF2 | 0.728278654 | 0.01295792 | -0.4574375 | 1.88746485 |
| RBMX2 | 0.727898966 | 0.10372009 | -0.4581899 | 0.9841371 |
| ZFYVE1 | 0.72736607 | 0.01194911 | -0.4592465 | 1.92266435 |
| PEX19 | 0.727122474 | 0.00585924 | -0.4597297 | 2.23215853 |
| NUFIP2 | 0.726939017 | 0.01375849 | -0.4600938 | 1.86142915 |
| DDI2 | 0.726861687 | 0.83379664 | -0.4602472 | 0.07893986 |
| DGCR8 | 0.726783531 | 0.06646173 | -0.4604024 | 1.17742836 |
| ZNF195 | 0.725365451 | 0.17920951 | -0.4632201 | 0.74663896 |
| UBXN1 | 0.725233812 | 0.39560784 | -0.4634819 | 0.40273511 |
| DPH5 | 0.724683537 | 0.49091655 | -0.464577 | 0.30899233 |
| CUX1 | 0.723861311 | 0.03379966 | -0.4662148 | 1.47108767 |
| MSTO1 | 0.72377603 | 0.21407574 | -0.4663848 | 0.66943254 |
| UBXN4 | 0.722709127 | 0.12232149 | -0.468513 | 0.91249726 |
| MAVS | 0.72182539 | 0.17563663 | -0.4702782 | 0.75538491 |
| CASP3 | 0.721798375 | 0.06299348 | -0.4703322 | 1.2007044 |
| FAM184A | 0.720951887 | 0.15761406 | -0.4720251 | 0.80240504 |
| RFC4 | 0.720528237 | 0.28362821 | -0.4728731 | 0.54725058 |
| IPO9 | 0.719656089 | 0.01476757 | -0.4746205 | 1.8306911 |
| RPS15A | 0.719278939 | 0.28703974 | -0.4753767 | 0.54205797 |
| GART | 0.718986432 | 0.01654458 | -0.4759635 | 1.78134426 |
| SPTBN1 | 0.718719424 | 0.03620033 | -0.4764994 | 1.44128749 |

| | | | | |
|---|---|---|---|---|
| VARS1 | 0.718551428 | 0.04781363 | -0.4768367 | 1.3204483 |
| DDX18 | 0.71850132 | 0.11597959 | -0.4769373 | 0.93561844 |
| MPRIP | 0.718220915 | 0.09945207 | -0.4775004 | 1.00238616 |
| POMGNT1 | 0.718047243 | 0.26305448 | -0.4778493 | 0.57995429 |
| BRCC3 | 0.718039002 | 0.29923552 | -0.4778659 | 0.52398685 |
| ACOX1 | 0.717651144 | 0.27337448 | -0.4786454 | 0.56324203 |
| URB2 | 0.717018434 | 0.01962008 | -0.4799179 | 1.70729914 |
| GTPBP1 | 0.71698139 | 0.03372838 | -0.4799924 | 1.47200448 |
| FAM120A | 0.716908373 | 0.03004539 | -0.4801394 | 1.52222215 |
| DDX39B | 0.716837135 | 0.39164184 | -0.4802827 | 0.40711092 |
| HIP1R | 0.715668106 | 0.093778 | -0.4826374 | 1.02789904 |
| STAU1 | 0.71552375 | 0.14689644 | -0.4829284 | 0.83298872 |
| SURF4 | 0.715436407 | 0.07264817 | -0.4831046 | 1.13877535 |
| SMG8 | 0.715416233 | 0.06860906 | -0.4831452 | 1.16361853 |
| PPP1R8 | 0.715058681 | 0.04205164 | -0.4838665 | 1.37621704 |
| EIF3L | 0.714993801 | 0.00833812 | -0.4839974 | 2.07893178 |
| CAD | 0.71480625 | 0.18618733 | -0.4843758 | 0.73004988 |
| SAMHD1 | 0.713628786 | 0.09433709 | -0.4867543 | 1.02531752 |
| TRMT1 | 0.713589637 | 0.18065275 | -0.4868334 | 0.74315543 |
| RIOK1 | 0.712374868 | 0.041435 | -0.4892915 | 1.38263263 |
| NR3C1 | 0.712028391 | 0.05692316 | -0.4899933 | 1.24471101 |
| SON | 0.711306669 | 0.01250275 | -0.4914564 | 1.90299438 |
| WTAP | 0.711261815 | 0.08071993 | -0.4915474 | 1.09301922 |
| QSER1 | 0.711000942 | 0.02331488 | -0.4920766 | 1.63236673 |
| SRRT | 0.710975586 | 0.16981873 | -0.4921281 | 0.77001441 |
| L3MBTL2 | 0.70962474 | 0.28216549 | -0.4948718 | 0.5494961 |
| DDX23 | 0.709585482 | 0.02025875 | -0.4949516 | 1.69338734 |
| RBM15 | 0.709302982 | 0.00828606 | -0.4955261 | 2.08165171 |
| TAF1 | 0.708791958 | 0.09295132 | -0.4965659 | 1.03174444 |
| TAPT1 | 0.708510292 | 0.05201506 | -0.4971393 | 1.28387088 |
| NEK1 | 0.707897 | 0.10289056 | -0.4983886 | 0.98762447 |
| ZNF160 | 0.707242115 | 0.06139473 | -0.4997239 | 1.21186894 |
| ZNF608 | 0.706659107 | 0.08785675 | -0.5009137 | 1.05622488 |
| TUBGCP3 | 0.706482523 | 0.01314321 | -0.5012742 | 1.88129847 |
| MYO1B | 0.706418004 | 0.00275981 | -0.501406 | 2.55912025 |
| SMPD4 | 0.70472576 | 0.10029929 | -0.5048661 | 0.99870213 |
| PRPF6 | 0.704163465 | 0.05965145 | -0.5060177 | 1.22437898 |
| NAA35 | 0.70395727 | 0.11442125 | -0.5064402 | 0.94149332 |
| ZBTB44 | 0.703697308 | 0.40302164 | -0.5069731 | 0.39467164 |
| PPIB | 0.701933864 | 0.17981646 | -0.510593 | 0.74517056 |
| SMARCD1 | 0.701729667 | 0.09258157 | -0.5110127 | 1.03347547 |
| UVRAG | 0.701046106 | 0.07064742 | -0.5124188 | 1.15090369 |
| DNM3 | 0.700959424 | 0.19053466 | -0.5125972 | 0.72002602 |
| CDKAL1 | 0.700534682 | 0.35594793 | -0.5134716 | 0.44861353 |
| PPP1CA | 0.700031993 | 0.17617914 | -0.5145072 | 0.75404551 |

| | | | | |
|---|---|---|---|---|
| RALGAPA1 | 0.699495782 | 0.03883941 | -0.5156127 | 1.4107274 |
| TRMT61A | 0.698103373 | 0.00544635 | -0.5184874 | 2.26389429 |
| CKAP4 | 0.697986666 | 0.84134963 | -0.5187286 | 0.07502349 |
| CNTROB | 0.697875909 | 0.08226193 | -0.5189576 | 1.08480109 |
| ABCF3 | 0.697577726 | 0.02806482 | -0.5195741 | 1.55183774 |
| SNRPF | 0.696937751 | 0.05420262 | -0.5208983 | 1.26597971 |
| KHDRBS1 | 0.696884293 | 0.41044781 | -0.521009 | 0.38674206 |
| EIF1AD | 0.696589758 | 0.02622199 | -0.5216188 | 1.58133432 |
| ATXN2L | 0.696578616 | 0.01522334 | -0.5216419 | 1.81749017 |
| ITPK1 | 0.696419143 | 0.09647761 | -0.5219722 | 1.01557346 |
| INA | 0.696236027 | 0.2598394 | -0.5223516 | 0.58529499 |
| DGKE | 0.696044119 | 0.00213916 | -0.5227493 | 2.66975648 |
| TBC1D23 | 0.695042557 | 0.32623115 | -0.5248268 | 0.48647457 |
| UBE3A | 0.69442742 | 0.01941487 | -0.5261042 | 1.71186552 |
| KIF14 | 0.694028321 | 0.36581073 | -0.5269336 | 0.43674356 |
| STRADA | 0.693837009 | 0.33543961 | -0.5273313 | 0.47438566 |
| CDYL | 0.691714925 | 0.1842709 | -0.5317505 | 0.73454324 |
| CCDC186 | 0.691549766 | 0.28134129 | -0.532095 | 0.55076652 |
| DMAP1 | 0.691514449 | 0.01895281 | -0.5321687 | 1.72232646 |
| TRIM9 | 0.691109098 | 0.13189655 | -0.5330146 | 0.87976657 |
| KARS1 | 0.690895911 | 0.99572424 | -0.5334597 | 0.00186092 |
| TTC4 | 0.69012185 | 0.25689231 | -0.535077 | 0.5902489 |
| CARS1 | 0.68940756 | 0.01740709 | -0.536571 | 1.75927371 |
| TET2 | 0.68939486 | 0.04223982 | -0.5365976 | 1.37427798 |
| TBL1XR1 | 0.687915139 | 0.03313136 | -0.5396975 | 1.47976076 |
| ZNF324 | 0.687381568 | 0.00329009 | -0.5408169 | 2.48279184 |
| TUBA1C | 0.687280719 | 0.07039232 | -0.5410286 | 1.15247469 |
| SLC25A46 | 0.687110568 | 0.13554964 | -0.5413858 | 0.86790162 |
| CNOT11 | 0.686818032 | 0.04335698 | -0.5420002 | 1.362941 |
| AMN1 | 0.685809288 | 0.00553374 | -0.5441207 | 2.25698087 |
| UBE4A | 0.685632766 | 0.26408813 | -0.544492 | 0.57825112 |
| GTF2H2 | 0.684382967 | 0.12402629 | -0.5471242 | 0.90648625 |
| UBE2Q1 | 0.684282755 | 0.08523627 | -0.5473355 | 1.06937558 |
| PPAT | 0.683109367 | 0.02394564 | -0.5498115 | 1.62077362 |
| CLASRP | 0.682672329 | 0.01787223 | -0.5507348 | 1.74782113 |
| CSNK2A1 | 0.68266663 | 0.04872205 | -0.5507469 | 1.31227445 |
| DCAF13 | 0.682093622 | 0.08785465 | -0.5519583 | 1.05623522 |
| PKN2 | 0.682049314 | 0.06543113 | -0.552052 | 1.1842156 |
| IBTK | 0.681967714 | 0.3182611 | -0.5522247 | 0.49721645 |
| TWF2 | 0.681948553 | 0.03858332 | -0.5522652 | 1.41360037 |
| PJA2 | 0.681438736 | 0.00703737 | -0.5533441 | 2.1525895 |
| RANBP1 | 0.681036359 | 0.00149237 | -0.5541963 | 2.82612239 |
| SRPRA | 0.680804295 | 0.11316465 | -0.554688 | 0.9462892 |
| ZNF629 | 0.680692312 | 0.6511845 | -0.5549253 | 0.18629595 |
| KSR1 | 0.680515819 | 0.32473104 | -0.5552994 | 0.48847619 |

| | | | | |
|---|---|---|---|---|
| FLAD1 | 0.679920082 | 0.50649533 | -0.5565629 | 0.29542456 |
| ZNF512 | 0.679635283 | 0.07287982 | -0.5571673 | 1.13739269 |
| TOR1AIP2 | 0.679047936 | 0.02663594 | -0.5584147 | 1.574532 |
| NOL9 | 0.678790722 | 0.13600564 | -0.5589612 | 0.86644307 |
| CWC25 | 0.678585064 | 0.0717681 | -0.5593984 | 1.14406857 |
| EXOC1 | 0.678457347 | 0.02318749 | -0.55967 | 1.63474622 |
| ZMAT2 | 0.677332211 | 0.03707632 | -0.5620645 | 1.43090338 |
| LYPLA2 | 0.676015875 | 0.02143886 | -0.564871 | 1.66879836 |
| IFT122 | 0.675617881 | 0.05726483 | -0.5657206 | 1.24211201 |
| CBS | 0.674504428 | 0.04253345 | -0.5681002 | 1.3712694 |
| PARP1 | 0.67447257 | 0.01220693 | -0.5681683 | 1.91339339 |
| TRIOBP | 0.674177747 | 0.32255663 | -0.5687991 | 0.49139402 |
| MRPS24 | 0.673655566 | 0.06151508 | -0.569917 | 1.21101841 |
| SREK1 | 0.67320047 | 0.63692444 | -0.5708919 | 0.19591209 |
| TUBG1 | 0.672626229 | 0.03355843 | -0.5721231 | 1.47419837 |
| SRGAP2 | 0.672204462 | 0.03752116 | -0.573028 | 1.42572369 |
| METTL13 | 0.672015265 | 0.01281101 | -0.5734341 | 1.89241648 |
| PCGF1 | 0.671854583 | 0.02255514 | -0.5737791 | 1.64675447 |
| MAP7D1 | 0.671139246 | 0.3148933 | -0.575316 | 0.50183658 |
| RNASEH2B | 0.670709926 | 0.05360968 | -0.5762391 | 1.27075681 |
| KIAA0232 | 0.670652547 | 0.21070571 | -0.5763626 | 0.67632369 |
| RPTOR | 0.670501217 | 0.10809028 | -0.5766881 | 0.96621335 |
| API5 | 0.668254057 | 0.03173288 | -0.5815314 | 1.49849056 |
| PIKFYVE | 0.667945824 | 0.13670322 | -0.582197 | 0.86422125 |
| ZC3H7A | 0.667010859 | 0.00497328 | -0.5842178 | 2.30335728 |
| MOB4 | 0.665989881 | 0.27233185 | -0.5864278 | 0.56490156 |
| PNPLA8 | 0.665974349 | 0.00011958 | -0.5864615 | 3.92235168 |
| ATP8A2 | 0.665476146 | 0.35097541 | -0.5875411 | 0.45472332 |
| PSMD4 | 0.66527736 | 0.12396285 | -0.5879722 | 0.90670846 |
| LTV1 | 0.665184209 | 0.78963642 | -0.5881742 | 0.10257283 |
| COPS4 | 0.665094083 | 0.033737 | -0.5883697 | 1.47189351 |
| WDR11 | 0.664718808 | 0.12984712 | -0.5891839 | 0.88656769 |
| FHIP2A | 0.664653807 | 0.95688429 | -0.589325 | 0.01914058 |
| KAT7 | 0.66359549 | 0.06583729 | -0.591624 | 1.18152805 |
| ADD3 | 0.66211189 | 0.05841486 | -0.5948531 | 1.23347669 |
| HP1BP3 | 0.662063701 | 0.01884549 | -0.5949581 | 1.72479264 |
| NAA38 | 0.662037947 | 0.44130757 | -0.5950142 | 0.35525862 |
| ALKBH4 | 0.658864275 | 0.37348514 | -0.6019468 | 0.42772668 |
| PSMD7 | 0.658771903 | 0.24653583 | -0.6021491 | 0.60811995 |
| NARS1 | 0.658675876 | 0.06880814 | -0.6023594 | 1.16236017 |
| PABPC4 | 0.656196854 | 0.0207112 | -0.6077994 | 1.68379476 |
| CHD2 | 0.656149114 | 0.01044762 | -0.6079044 | 1.98098274 |
| CCHCR1 | 0.655361551 | 0.05709569 | -0.6096371 | 1.24339667 |
| NOC2L | 0.65526488 | 0.00422617 | -0.6098499 | 2.37405293 |
| KRT38 | 0.655028226 | 0.08789888 | -0.610371 | 1.05601667 |

| | | | | |
|---|---|---|---|---|
| STK26 | 0.654526371 | 0.03420322 | -0.6114768 | 1.46593305 |
| ENSA | 0.653339029 | 0.03481172 | -0.6140963 | 1.45827453 |
| PAXBP1 | 0.653137386 | 0.03878512 | -0.6145416 | 1.4113349 |
| MRPS31 | 0.65273777 | 0.00825171 | -0.6154246 | 2.08345612 |
| ZNG1B | 0.652722001 | 0.02781636 | -0.6154594 | 1.5556997 |
| RBPJ | 0.652646422 | 0.05767081 | -0.6156265 | 1.23904395 |
| HSPBP1 | 0.651942919 | 0.60927225 | -0.6171824 | 0.2151886 |
| HK1 | 0.650981344 | 0.1291045 | -0.6193119 | 0.88905864 |
| UTP18 | 0.650603975 | 0.09423609 | -0.6201485 | 1.02578273 |
| LRRC14 | 0.649710218 | 0.48916342 | -0.6221317 | 0.31054603 |
| TBC1D15 | 0.649057483 | 0.03739536 | -0.6235818 | 1.42718226 |
| HYOU1 | 0.648790119 | 0.95558616 | -0.6241762 | 0.01973015 |
| SNRPD1 | 0.648435406 | 0.03431232 | -0.6249652 | 1.46454995 |
| SMAP | 0.648415299 | 0.0106055 | -0.62501 | 1.97446881 |
| SUGT1 | 0.648389751 | 0.13994187 | -0.6250668 | 0.85405233 |
| QNG1 | 0.647993088 | 0.84139905 | -0.6259497 | 0.07499798 |
| CPSF1 | 0.647563317 | 0.00274328 | -0.6269068 | 2.56172917 |
| DDX5 | 0.647112813 | 0.00727366 | -0.6279109 | 2.13824688 |
| ARF6 | 0.647074945 | 0.04140044 | -0.6279953 | 1.38299501 |
| PRPF4 | 0.646641077 | 0.02339575 | -0.6289629 | 1.6308631 |
| SBF1 | 0.646459386 | 0.13354006 | -0.6293684 | 0.87438842 |
| SEPTIN9 | 0.646222323 | 0.01059686 | -0.6298975 | 1.97482262 |
| HMOX2 | 0.646184982 | 0.13543544 | -0.6299809 | 0.86826769 |
| PAX6 | 0.645614789 | 0.0369872 | -0.6312545 | 1.43194859 |
| WDR91 | 0.644705086 | 0.10973371 | -0.6332887 | 0.95965995 |
| NAT14 | 0.644580342 | 0.31468247 | -0.6335679 | 0.50212746 |
| IKBKG | 0.64450007 | 0.38115343 | -0.6337476 | 0.41890016 |
| MEPCE | 0.643204879 | 0.00994442 | -0.6366497 | 2.00242043 |
| MAZ | 0.64262893 | 0.07025129 | -0.6379422 | 1.15334569 |
| EMG1 | 0.642485644 | 0.30757283 | -0.6382639 | 0.51205203 |
| ADNP2 | 0.642222713 | 0.17022632 | -0.6388544 | 0.76897328 |
| OTUD4 | 0.640559082 | 0.03111906 | -0.6425965 | 1.50697359 |
| PIK3R1 | 0.640519733 | 0.33234375 | -0.6426851 | 0.47841249 |
| ALS2 | 0.640223522 | 0.2907637 | -0.6433524 | 0.53645981 |
| ARPC2 | 0.639310579 | 0.33064591 | -0.6454111 | 0.48063684 |
| ZBTB7A | 0.639265194 | 0.25389404 | -0.6455135 | 0.5953475 |
| RPL10A | 0.638131537 | 0.01462141 | -0.6480743 | 1.83501088 |
| COPS8 | 0.638040095 | 0.05058496 | -0.648281 | 1.2959786 |
| PPIP5K2 | 0.637620762 | 0.01006302 | -0.6492295 | 1.99727183 |
| CBX3 | 0.637436289 | 0.00510115 | -0.6496469 | 2.2923315 |
| PEG3 | 0.63700266 | 0.39000985 | -0.6506287 | 0.40892443 |
| GPATCH2L | 0.636892472 | 0.13793003 | -0.6508783 | 0.86034116 |
| LRRC8D | 0.636632724 | 0.07190169 | -0.6514668 | 1.14326087 |
| XAB2 | 0.636367693 | 0.01863386 | -0.6520675 | 1.7296972 |
| AKT2 | 0.636067031 | 0.28933181 | -0.6527493 | 0.53860382 |

| | | | | |
|---|---|---|---|---|
| SLAIN2 | 0.635981274 | 0.03513388 | -0.6529438 | 1.45427394 |
| CERS2 | 0.635859783 | 0.19404985 | -0.6532194 | 0.71208669 |
| PAK1 | 0.63542183 | 0.01704341 | -0.6542134 | 1.7684436 |
| LYSMD2 | 0.635111111 | 0.08222862 | -0.6549191 | 1.08497701 |
| DPYSL2 | 0.634855318 | 0.00538998 | -0.6555003 | 2.2684131 |
| HNRNPU | 0.63468086 | 0.00269406 | -0.6558968 | 2.56959348 |
| CCT8 | 0.6340785 | 0.0045167 | -0.6572666 | 2.34517885 |
| CLK2 | 0.633938027 | 0.49778688 | -0.6575863 | 0.30295655 |
| PRORP | 0.633568943 | 0.02855431 | -0.6584265 | 1.54432829 |
| LUZP1 | 0.633155337 | 0.02136231 | -0.6593686 | 1.67035182 |
| RRP15 | 0.632498674 | 0.00242248 | -0.6608656 | 2.61573971 |
| OSBPL8 | 0.63185182 | 0.01226193 | -0.6623418 | 1.91144116 |
| HDLBP | 0.631683509 | 0.01135554 | -0.6627262 | 1.94479229 |
| FTSJ1 | 0.631476027 | 0.55858277 | -0.6632001 | 0.25291246 |
| EPM2AIP1 | 0.631113277 | 0.11479856 | -0.6640291 | 0.94006357 |
| PAF1 | 0.630758614 | 0.00146399 | -0.6648401 | 2.83446221 |
| HMGXB4 | 0.630116498 | 0.12147465 | -0.6663095 | 0.91551434 |
| BICD2 | 0.628874996 | 0.01656392 | -0.6691548 | 1.78083679 |
| STRA6 | 0.628796659 | 0.03026366 | -0.6693345 | 1.51907856 |
| PTPN1 | 0.62844314 | 0.03596579 | -0.6701459 | 1.44411036 |
| KIF1A | 0.628215633 | 0.00624478 | -0.6706682 | 2.20448306 |
| LSG1 | 0.626933636 | 0.04200322 | -0.6736154 | 1.37671744 |
| FLNA | 0.62661859 | 0.01877532 | -0.6743405 | 1.72641267 |
| NAPG | 0.626043477 | 0.09096137 | -0.6756652 | 1.04114299 |
| PRMT1 | 0.624355457 | 0.06787419 | -0.6795605 | 1.16829536 |
| ESPN | 0.624285759 | 0.01563162 | -0.6797215 | 1.805996 |
| LAMTOR2 | 0.624121289 | 0.21426 | -0.6801017 | 0.6690589 |
| SRGAP1 | 0.62356873 | 0.0689658 | -0.6813795 | 1.1613662 |
| RPS6KB1 | 0.62334042 | 0.17910895 | -0.6819078 | 0.74688271 |
| HEATR1 | 0.622840928 | 0.03362937 | -0.6830643 | 1.47328125 |
| VPS52 | 0.622612395 | 0.10519452 | -0.6835938 | 0.97800687 |
| EIF3E | 0.622531088 | 0.07579197 | -0.6837822 | 1.12037682 |
| ARHGEF12 | 0.622409484 | 0.01232154 | -0.6840641 | 1.90933509 |
| ESYT1 | 0.621985338 | 0.04587187 | -0.6850475 | 1.33845357 |
| DMRTA1 | 0.621621822 | 0.0235739 | -0.6858909 | 1.6275685 |
| HAUS3 | 0.621009081 | 0.07501545 | -0.6873137 | 1.12484929 |
| PDCD11 | 0.620788996 | 0.0368086 | -0.6878251 | 1.43405067 |
| MIA3 | 0.620592897 | 0.0059375 | -0.6882809 | 2.22639651 |
| ATIC | 0.619875742 | 0.05359971 | -0.689949 | 1.27083754 |
| SLK | 0.618934099 | 0.04106447 | -0.6921423 | 1.3865338 |
| POLRMT | 0.618383005 | 0.03044841 | -0.6934274 | 1.51643533 |
| RALGAPB | 0.618243879 | 0.04311772 | -0.693752 | 1.36534416 |
| LAPTM4B | 0.618025537 | 0.91565667 | -0.6942616 | 0.03826734 |
| TRAPPC5 | 0.618003334 | 0.31640823 | -0.6943135 | 0.49975222 |
| PSPC1 | 0.617384178 | 0.00421095 | -0.6957596 | 2.37562001 |

| | | | | |
|---|---|---|---|---|
| EIF2S2 | 0.617032638 | 0.0133606 | -0.6965813 | 1.87417388 |
| UFD1 | 0.616904547 | 0.11822276 | -0.6968808 | 0.9272989 |
| UBL4A | 0.615491014 | 0.13349056 | -0.7001903 | 0.87454944 |
| PTMA | 0.615136083 | 0.14121353 | -0.7010225 | 0.85012369 |
| C17orf80 | 0.615054981 | 0.27168691 | -0.7012127 | 0.56593129 |
| FAM91A1 | 0.614391705 | 0.09547238 | -0.7027694 | 1.02012223 |
| DYNC1I2 | 0.614137384 | 0.03726624 | -0.7033667 | 1.42868443 |
| PAM16 | 0.613725934 | 0.01178606 | -0.7043335 | 1.9286314 |
| ATE1 | 0.612276363 | 0.27415387 | -0.7077451 | 0.56200562 |
| APOB | 0.611777407 | 0.2256506 | -0.7089213 | 0.64656351 |
| NLRX1 | 0.611582146 | 0.36273117 | -0.7093818 | 0.44041512 |
| CORO2A | 0.61156236 | 0.01169829 | -0.7094285 | 1.93187757 |
| DTNB | 0.611183745 | 0.07711892 | -0.7103219 | 1.11283906 |
| SEPTIN5 | 0.610157453 | 0.10979243 | -0.7127465 | 0.95942759 |
| PEBP1 | 0.609541332 | 0.10645186 | -0.714204 | 0.97284675 |
| PDCD6 | 0.608198518 | 0.04307673 | -0.7173858 | 1.36575723 |
| KIF1B | 0.60782312 | 0.09222498 | -0.7182765 | 1.03515143 |
| RTRAF | 0.60665198 | 0.07027917 | -0.721059 | 1.15317338 |
| USP14 | 0.605756483 | 0.15588844 | -0.7231902 | 0.8071861 |
| PRPSAP1 | 0.605346883 | 0.3996574 | -0.724166 | 0.39831214 |
| WDR81 | 0.604904662 | 0.16372731 | -0.7252203 | 0.78587888 |
| LRWD1 | 0.604024923 | 0.04245375 | -0.72732 | 1.3720839 |
| FRG1 | 0.603791504 | 0.02223221 | -0.7278776 | 1.65301741 |
| CDCA5 | 0.603300133 | 0.27299671 | -0.7290522 | 0.56384259 |
| GNL3L | 0.603091199 | 0.30340879 | -0.7295519 | 0.51797185 |
| CNTLN | 0.602386899 | 0.27781547 | -0.7312377 | 0.55624358 |
| ZCCHC7 | 0.601607354 | 0.26842415 | -0.7331059 | 0.57117841 |
| ATXN2 | 0.600521281 | 0.00032278 | -0.7357127 | 3.49108906 |
| ANO8 | 0.6002908 | 0.00713314 | -0.7362665 | 2.14671913 |
| ATG2A | 0.600249709 | 0.28333516 | -0.7363653 | 0.54769954 |
| SESN3 | 0.600109259 | 0.01942148 | -0.7367029 | 1.71171767 |
| EDF1 | 0.599289614 | 0.0337273 | -0.7386747 | 1.47201849 |
| COG3 | 0.599214011 | 0.15743125 | -0.7388567 | 0.80290906 |
| RBM41 | 0.598832074 | 0.26832676 | -0.7397766 | 0.57133601 |
| E2F4 | 0.598695913 | 0.16479167 | -0.7401047 | 0.78306476 |
| PES1 | 0.59740257 | 0.00692599 | -0.7432247 | 2.15951791 |
| DOCK4 | 0.596951835 | 0.18857623 | -0.7443136 | 0.72451306 |
| TCF3 | 0.596585062 | 0.02148612 | -0.7452002 | 1.6678419 |
| RASGRP1 | 0.595737071 | 0.29443997 | -0.7472524 | 0.53100324 |
| NUP93 | 0.595318155 | 0.00235061 | -0.7482672 | 2.62881999 |
| NEFL | 0.594776898 | 0.00944015 | -0.7495795 | 2.02502124 |
| LENG8 | 0.594776171 | 0.17370994 | -0.7495812 | 0.76017532 |
| HECTD1 | 0.594306959 | 0.01541764 | -0.7507198 | 1.81198223 |
| BAG1 | 0.594228957 | 0.13896906 | -0.7509092 | 0.85708189 |
| FBXL18 | 0.593956928 | 0.27471948 | -0.7515698 | 0.56111054 |

| | | | | |
|---|---|---|---|---|
| TSTD2 | 0.593794073 | 0.02446528 | -0.7519654 | 1.61144972 |
| VPS4B | 0.593720096 | 0.0142664 | -0.7521451 | 1.84568565 |
| BAX | 0.593578311 | 0.73567705 | -0.7524897 | 0.13331279 |
| **TRAF6** | 0.593277227 | 0.40441494 | -0.7532217 | 0.39317281 |
| PPIL1 | 0.592724179 | 0.02945604 | -0.7545672 | 1.53082564 |
| PSMD10 | 0.592510361 | 0.01095109 | -0.7550877 | 1.96054269 |
| TRIM24 | 0.59224718 | 0.00434619 | -0.7557287 | 2.36189168 |
| FNBP4 | 0.591771226 | 0.00655849 | -0.7568885 | 2.183196 |
| NACA | 0.591301559 | 0.03208774 | -0.758034 | 1.49366082 |
| CENPF | 0.591301168 | 0.01558595 | -0.758035 | 1.80726668 |
| DHX36 | 0.590467991 | 0.01885027 | -0.7600692 | 1.72468253 |
| RNF213 | 0.58813746 | 0.02168759 | -0.7657747 | 1.66378877 |
| AKT1 | 0.586655509 | 0.0249375 | -0.7694145 | 1.6031471 |
| YTHDC1 | 0.585514635 | 0.00209626 | -0.7722229 | 2.67855501 |
| ZNF84 | 0.584909234 | 0.48916945 | -0.7737153 | 0.31054067 |
| NFKBIB | 0.584695092 | 0.08576356 | -0.7742436 | 1.06669718 |
| ARHGEF2 | 0.584565785 | 0.00922108 | -0.7745627 | 2.03521803 |
| NOB1 | 0.584410971 | 0.095879 | -0.7749448 | 1.0182765 |
| ATP6V0A2 | 0.58285234 | 0.10454233 | -0.7787977 | 0.98070781 |
| WDR77 | 0.580854024 | 0.3217653 | -0.7837525 | 0.4924608 |
| KIF4A | 0.580711058 | 0.01606913 | -0.7841076 | 1.79400756 |
| ADSL | 0.578230764 | 0.00038367 | -0.7902827 | 3.41604181 |
| HECTD4 | 0.577458068 | 0.06646407 | -0.7922119 | 1.17741309 |
| PNP | 0.576901622 | 0.00056313 | -0.7936028 | 3.24938946 |
| DCTPP1 | 0.576210091 | 0.08026594 | -0.7953332 | 1.09546869 |
| USP47 | 0.576078864 | 0.03687817 | -0.7956618 | 1.43323059 |
| PUS3 | 0.575805253 | 0.07400611 | -0.7963471 | 1.1307324 |
| FAM221A | 0.575247033 | 0.04766433 | -0.7977465 | 1.32180648 |
| CSNK2B | 0.575215437 | 0.01540334 | -0.7978257 | 1.81238504 |
| KPNA6 | 0.574964182 | 0.05453596 | -0.798456 | 1.26331707 |
| TEX10 | 0.573378313 | 0.0086423 | -0.8024408 | 2.06337084 |
| CHUK | 0.573257815 | 0.12337108 | -0.802744 | 0.90878664 |
| OSBPL3 | 0.57269769 | 0.00023527 | -0.8041543 | 3.62842624 |
| VRK2 | 0.572520075 | 0.1098061 | -0.8046018 | 0.95937355 |
| RETREG2 | 0.572331427 | 0.00303204 | -0.8050773 | 2.5182646 |
| MYH10 | 0.57229142 | 0.01293026 | -0.8051781 | 1.88839291 |
| KCTD18 | 0.571795095 | 0.03096829 | -0.8064299 | 1.50908279 |
| SBNO1 | 0.571717244 | 0.20969452 | -0.8066263 | 0.67841291 |
| TMOD2 | 0.571508896 | 0.21474783 | -0.8071521 | 0.66807121 |
| PTPN12 | 0.570101288 | 0.0373912 | -0.8107098 | 1.42723062 |
| ACTR2 | 0.568860602 | 0.24655059 | -0.8138529 | 0.60809395 |
| HNRNPA1 | 0.568793207 | 0.00210762 | -0.8140239 | 2.67620686 |
| RUSF1 | 0.568727147 | 0.35580802 | -0.8141914 | 0.44878426 |
| MIA2 | 0.568614738 | 0.07546376 | -0.8144766 | 1.12226157 |
| SP1 | 0.56836976 | 0.32123647 | -0.8150983 | 0.49317515 |

| | | | | |
|---|---|---|---|---|
| ZPR1 | 0.568171955 | 0.29059172 | -0.8156005 | 0.53671676 |
| RAPGEF1 | 0.567941773 | 0.03071315 | -0.8161851 | 1.51267568 |
| COPB2 | 0.567793501 | 0.00218809 | -0.8165618 | 2.65993475 |
| ULK3 | 0.567770142 | 0.17380405 | -0.8166211 | 0.75994012 |
| DHX33 | 0.567545919 | 0.11261073 | -0.817191 | 0.94842021 |
| ABCD3 | 0.566754017 | 0.01188306 | -0.8192054 | 1.92507156 |
| PLEKHG4 | 0.566753174 | 0.17339116 | -0.8192075 | 0.76097304 |
| XPO5 | 0.565482926 | 0.04051429 | -0.8224446 | 1.39239177 |
| NEK7 | 0.565315127 | 0.17946847 | -0.8228728 | 0.74601184 |
| PRMT5 | 0.565199637 | 0.1135982 | -0.8231676 | 0.94462855 |
| ZNF48 | 0.56514762 | 0.52562591 | -0.8233003 | 0.27932324 |
| RPRD1B | 0.564865076 | 0.0005811 | -0.8240218 | 3.23575041 |
| PTMS | 0.564239982 | 0.01427454 | -0.8256192 | 1.84543789 |
| SEPTIN3 | 0.563726724 | 0.00077646 | -0.8269321 | 3.10987948 |
| EXOSC4 | 0.563375708 | 0.01542835 | -0.8278307 | 1.81168045 |
| NUBP1 | 0.563024291 | 0.74276047 | -0.8287309 | 0.12915122 |
| PCCA | 0.563009613 | 0.02498744 | -0.8287685 | 1.60227822 |
| CCZ1B | 0.562967498 | 0.02081335 | -0.8288765 | 1.68165802 |
| PHKG2 | 0.562895268 | 0.00351401 | -0.8290616 | 2.45419699 |
| ALKBH2 | 0.562157784 | 0.10467646 | -0.830953 | 0.98015098 |
| SRPK1 | 0.561692518 | 0.02471815 | -0.8321475 | 1.6069841 |
| PBK | 0.561107127 | 0.01859827 | -0.8336519 | 1.73052749 |
| GTF3C3 | 0.560957901 | 0.02649092 | -0.8340356 | 1.57690292 |
| CREBBP | 0.560411923 | 0.44661911 | -0.8354404 | 0.3500627 |
| MTMR3 | 0.55937634 | 0.00986736 | -0.8381089 | 2.00579885 |
| BUB3 | 0.559214072 | 0.01294691 | -0.8385274 | 1.88783403 |
| CRTC1 | 0.559092112 | 0.00109055 | -0.8388421 | 2.96235616 |
| PLS1 | 0.558368575 | 0.02960686 | -0.8407103 | 1.52860758 |
| TUBB2A | 0.558366796 | 0.02345744 | -0.8407149 | 1.62971934 |
| SUN1 | 0.557522681 | 0.00339433 | -0.8428976 | 2.46924629 |
| CEP170 | 0.557464474 | 0.00148048 | -0.8430482 | 2.82959742 |
| EIF2B3 | 0.556757273 | 0.00132394 | -0.8448796 | 2.87813301 |
| TCF19 | 0.556647267 | 0.04702719 | -0.8451647 | 1.32765096 |
| TIA1 | 0.556117745 | 0.25842061 | -0.8465377 | 0.58767285 |
| URGCP | 0.554905488 | 0.32380517 | -0.849686 | 0.48971622 |
| ARHGAP17 | 0.55358276 | 0.0390234 | -0.8531291 | 1.40867486 |
| PEX14 | 0.55322439 | 0.01049718 | -0.8540633 | 1.9789273 |
| POU4F3 | 0.553073798 | 0.00516633 | -0.8544561 | 2.28681782 |
| FTH1 | 0.552284927 | 0.21924506 | -0.8565153 | 0.65907019 |
| ARPC1A | 0.551988868 | 0.00795845 | -0.8572889 | 2.09917155 |
| SNX29 | 0.550395156 | 0.00126188 | -0.8614603 | 2.89898069 |
| TUFT1 | 0.550070922 | 0.01626048 | -0.8623105 | 1.78886656 |
| DENND4A | 0.548687464 | 0.00027797 | -0.8659435 | 3.55599916 |
| MCM4 | 0.548295751 | 0.00030666 | -0.8669738 | 3.51333804 |
| SH3PXD2A | 0.548051686 | 0.00465931 | -0.8676161 | 2.33167848 |

| | | | | |
|---|---|---|---|---|
| REL | 0.544678521 | 0.00084461 | -0.8765231 | 3.07334455 |
| DMXL1 | 0.544151514 | 0.01656981 | -0.8779197 | 1.78068253 |
| ZCCHC3 | 0.544149449 | 0.11124494 | -0.8779252 | 0.95371973 |
| ORC5 | 0.543465373 | 0.1934991 | -0.87974 | 0.71332106 |
| KIF5B | 0.542761124 | 0.00315735 | -0.8816107 | 2.50067704 |
| NBAS | 0.541865095 | 0.00896794 | -0.8839944 | 2.04730732 |
| INTS4 | 0.541455434 | 0.02256941 | -0.8850855 | 1.64647976 |
| DIP2B | 0.54097186 | 0.02556922 | -0.8863745 | 1.59228245 |
| SHTN1 | 0.540534619 | 0.00352371 | -0.8875411 | 2.45299985 |
| ARID4A | 0.539923043 | 0.00037397 | -0.8891743 | 3.42716231 |
| RAB3GAP1 | 0.539761488 | 0.00709725 | -0.8896061 | 2.14890963 |
| RNF220 | 0.538783612 | 0.26152669 | -0.8922221 | 0.58248399 |
| MYCBP2 | 0.537097761 | 0.00647261 | -0.8967434 | 2.1889207 |
| NAP1L4 | 0.536926925 | 0.00367282 | -0.8972023 | 2.43499985 |
| MDN1 | 0.536708826 | 0.00477259 | -0.8977885 | 2.3212457 |
| LARP7 | 0.536690064 | 0.00415744 | -0.8978389 | 2.38117395 |
| UNC45A | 0.536678425 | 0.006878 | -0.8978702 | 2.16253756 |
| AGPAT5 | 0.536531883 | 0.3125256 | -0.8982642 | 0.50511441 |
| RUBCN | 0.535775728 | 0.03341279 | -0.9002989 | 1.47608721 |
| PPID | 0.535331071 | 0.18567051 | -0.9014967 | 0.73125707 |
| CBX1 | 0.534328265 | 0.18818838 | -0.9042018 | 0.7254072 |
| ATP11A | 0.534318888 | 0.00986963 | -0.9042271 | 2.00569912 |
| PIP5K1C | 0.53420928 | 0.16510716 | -0.9045231 | 0.7822341 |
| CAMSAP3 | 0.533962547 | 0.00911998 | -0.9051895 | 2.0400061 |
| FTO | 0.533525993 | 0.20027836 | -0.9063695 | 0.69836597 |
| DPYSL4 | 0.532714662 | 0.01771997 | -0.9085651 | 1.75153705 |
| TRAPPC6B | 0.53236393 | 0.00051207 | -0.9095153 | 3.29067116 |
| KIF23 | 0.532303677 | 0.01048741 | -0.9096786 | 1.97933179 |
| NAMPT | 0.532287402 | 0.00172479 | -0.9097227 | 2.76326395 |
| BAG5 | 0.531841759 | 4.146E-06 | -0.910931 | 5.38236918 |
| CCDC91 | 0.531705009 | 0.14245442 | -0.911302 | 0.84632406 |
| HMGB1 | 0.53165854 | 0.00154776 | -0.9114281 | 2.81029544 |
| MYO5A | 0.530983105 | 0.01077535 | -0.9132621 | 1.96756871 |
| POLR1C | 0.530892444 | 0.00072304 | -0.9135085 | 3.1408348 |
| ERCC1 | 0.530235809 | 0.09393413 | -0.915294 | 1.02717658 |
| ASF1B | 0.529986265 | 0.00441532 | -0.9159731 | 2.35503788 |
| PPP1R12C | 0.52948463 | 0.00144571 | -0.9173393 | 2.83991847 |
| TNPO3 | 0.529353895 | 0.14602327 | -0.9176955 | 0.83557794 |
| FLYWCH1 | 0.529114332 | 0.19155106 | -0.9183486 | 0.71771545 |
| BORCS7-ASMT | 0.528855895 | 0.02523326 | -0.9190534 | 1.59802668 |
| DNAAF2 | 0.528853987 | 0.0340014 | -0.9190586 | 1.46850315 |
| CENPM | 0.528848343 | 0.30571366 | -0.919074 | 0.51468516 |
| XPO7 | 0.528575669 | 0.00682824 | -0.9198181 | 2.165691 |
| IRGQ | 0.527501894 | 0.14466144 | -0.9227518 | 0.83964721 |
| NCOR1 | 0.526642832 | 0.00131692 | -0.9251032 | 2.88044208 |

| | | | | |
|---|---|---|---|---|
| RAI1 | 0.526033493 | 0.02601112 | -0.9267734 | 1.58484088 |
| PATZ1 | 0.525973326 | 0.0165183 | -0.9269385 | 1.7820347 |
| PFKFB2 | 0.52568802 | 0.03721206 | -0.9277212 | 1.4293163 |
| SARNP | 0.524988881 | 0.02107784 | -0.9296412 | 1.67617392 |
| RNASEH2A | 0.524813852 | 0.01307028 | -0.9301223 | 1.88371518 |
| NUDT12 | 0.524710832 | 0.0435859 | -0.9304055 | 1.36065396 |
| NFKBIE | 0.524241504 | 0.10449712 | -0.9316965 | 0.98089569 |
| SRPK2 | 0.523635045 | 0.0082722 | -0.9333664 | 2.082379 |
| SNX5 | 0.523309688 | 0.00086071 | -0.9342631 | 3.06514435 |
| KLC1 | 0.523268008 | 0.00088277 | -0.934378 | 3.05415305 |
| HEATR6 | 0.523239774 | 0.00680794 | -0.9344559 | 2.1669842 |
| MCM6 | 0.522900509 | 0.02126528 | -0.9353916 | 1.67232885 |
| MDH1 | 0.522714112 | 0.13571773 | -0.935906 | 0.86736342 |
| SNRPA | 0.520437147 | 0.00054947 | -0.9422042 | 3.26005935 |
| PNISR | 0.520204182 | 0.15915783 | -0.9428501 | 0.79817198 |
| GBF1 | 0.519888756 | 0.00100603 | -0.9437251 | 2.99738762 |
| AFAP1 | 0.519505984 | 0.0410542 | -0.9447877 | 1.3866424 |
| PDCL3 | 0.519306162 | 0.04720628 | -0.9453428 | 1.3260002 |
| ACVR1B | 0.518966966 | 0.01142079 | -0.9462854 | 1.94230399 |
| CGGBP1 | 0.518699819 | 0.02663515 | -0.9470282 | 1.57454479 |
| RACGAP1 | 0.518330373 | 0.01461841 | -0.9480562 | 1.83509985 |
| COPB1 | 0.51765329 | 0.00978407 | -0.9499419 | 2.00948054 |
| EDC4 | 0.516969378 | 0.01957376 | -0.9518493 | 1.70832564 |
| HTT | 0.516614213 | 0.05249471 | -0.9528408 | 1.27988444 |
| ZCRB1 | 0.516609153 | 0.84318125 | -0.9528549 | 0.07407906 |
| PRPF40A | 0.515935347 | 0.03756526 | -0.9547378 | 1.42521364 |
| PDE4B | 0.515868407 | 0.00383255 | -0.954925 | 2.41651202 |
| SENP3 | 0.515852117 | 0.02081232 | -0.9549706 | 1.68167942 |
| APAF1 | 0.515596455 | 0.0260804 | -0.9556858 | 1.58368568 |
| NDUFB9 | 0.515176675 | 0.24505796 | -0.9568608 | 0.61073118 |
| DUS2 | 0.514936186 | 0.16981527 | -0.9575344 | 0.77002326 |
| PADI2 | 0.5149157 | 0.01059663 | -0.9575918 | 1.97483227 |
| PRRC2B | 0.513433846 | 0.03184981 | -0.9617497 | 1.49689319 |
| DHX8 | 0.513311822 | 0.11097624 | -0.9620926 | 0.95477 |
| CERK | 0.512872002 | 0.38726475 | -0.9633293 | 0.41199203 |
| RBM15B | 0.512308092 | 0.05437263 | -0.9649164 | 1.26461967 |
| NEFH | 0.511901371 | 0.00198133 | -0.9660622 | 2.70304211 |
| RUNX1T1 | 0.511611146 | 0.04242557 | -0.9668804 | 1.37237233 |
| PABPC1 | 0.510649079 | 0.00925208 | -0.9695959 | 2.03376067 |
| GNPDA1 | 0.510312191 | 0.68368637 | -0.970548 | 0.16514308 |
| ATG3 | 0.510279808 | 0.00619745 | -0.9706395 | 2.20778669 |
| ORC3 | 0.510077149 | 0.10316753 | -0.9712126 | 0.98645697 |
| BCOR | 0.50907562 | 0.00026736 | -0.9740481 | 3.57290784 |
| LYPLAL1 | 0.5087137 | 0.0234303 | -0.9750741 | 1.63022217 |
| ERO1A | 0.508617626 | 0.72255875 | -0.9753466 | 0.14112684 |

| | | | | |
|---|---|---|---|---|
| HSPA4 | 0.50851679 | 0.10517807 | -0.9756327 | 0.97807479 |
| WDR75 | 0.508330115 | 0.00836833 | -0.9761624 | 2.07736144 |
| ELP6 | 0.508317325 | 0.07778654 | -0.9761987 | 1.10909553 |
| EXOSC1 | 0.508199985 | 0.25870966 | -0.9765318 | 0.58718736 |
| ANXA5 | 0.507616605 | 0.30088781 | -0.9781888 | 0.5215954 |
| SMC2 | 0.507428412 | 0.00044585 | -0.9787238 | 3.35081435 |
| PRPF38B | 0.507389538 | 0.01193665 | -0.9788343 | 1.92311745 |
| CHML | 0.50725489 | 0.0843548 | -0.9792172 | 1.07389021 |
| COQ6 | 0.507078889 | 0.07363227 | -0.9797179 | 1.13293183 |
| IPO5 | 0.505994598 | 0.05931206 | -0.9828061 | 1.22685701 |
| MRE11 | 0.50552981 | 0.00458096 | -0.9841319 | 2.33904356 |
| EIF3C | 0.505528638 | 0.00129919 | -0.9841353 | 2.88632697 |
| EEF2K | 0.505340033 | 0.06412616 | -0.9846736 | 1.19296473 |
| UBE2T | 0.503420866 | 0.01589928 | -0.9901631 | 1.7986226 |
| TYK2 | 0.503319892 | 0.05214089 | -0.9904525 | 1.28282159 |
| SH2D5 | 0.50311827 | 0.00071625 | -0.9910305 | 3.14493382 |
| AFG2B | 0.503018295 | 0.00557321 | -0.9913172 | 2.25389487 |
| TOPBP1 | 0.503010095 | 0.08709771 | -0.9913407 | 1.05999325 |
| MYO5B | 0.502989282 | 0.00156895 | -0.9914004 | 2.80439223 |
| AAR2 | 0.502871168 | 0.30327528 | -0.9917393 | 0.51816299 |
| KIF20A | 0.502827674 | 0.00340388 | -0.991864 | 2.46802622 |
| MRPS35 | 0.50240406 | 0.05265929 | -0.99308 | 1.27852498 |
| COPS5 | 0.501868527 | 0.02130662 | -0.9946186 | 1.67148536 |
| NAF1 | 0.501515559 | 0.11869524 | -0.9956336 | 0.92556668 |
| EPG5 | 0.501250429 | 0.15830868 | -0.9963965 | 0.80049527 |
| ANLN | 0.501218072 | 0.07368894 | -0.9964897 | 1.13259771 |
| MAPRE2 | 0.501096818 | 0.01075629 | -0.9968387 | 1.96833759 |
| TMEM63B | 0.500695546 | 0.43699835 | -0.9979945 | 0.3595202 |
| FKBP3 | 0.500124766 | 0.27592573 | -0.99964 | 0.55920779 |
| FXR2 | 0.500111036 | 0.00408665 | -0.9996797 | 2.3886324 |
| PUM3 | 0.499886648 | 0.1244365 | -1.0003271 | 0.90505222 |
| IPO7 | 0.499712249 | 0.023709 | -1.0008305 | 1.62508671 |
| TUBGCP4 | 0.499695353 | 0.05683031 | -1.0008793 | 1.24541995 |
| SMC3 | 0.499178573 | 0.00448719 | -1.0023721 | 2.34802579 |
| PRKDC | 0.49894774 | 0.0066386 | -1.0030394 | 2.17792365 |
| USP42 | 0.498884292 | 0.01338908 | -1.0032228 | 1.87324922 |
| RIMS2 | 0.497350363 | 0.00898002 | -1.0076656 | 2.04672289 |
| PAPSS1 | 0.497308334 | 0.01372283 | -1.0077875 | 1.86255637 |
| CDK13 | 0.496934136 | 0.00076589 | -1.0088734 | 3.11583519 |
| WDR82 | 0.496744482 | 0.02069533 | -1.0094242 | 1.6841277 |
| STAG1 | 0.496532567 | 0.01874012 | -1.0100397 | 1.72722774 |
| MYO10 | 0.495936141 | 0.13112 | -1.0117737 | 0.88233106 |
| SEC63 | 0.495752834 | 0.05797721 | -1.0123071 | 1.2367427 |
| SUN2 | 0.494701353 | 0.01934066 | -1.0153702 | 1.71352881 |
| CAMK2D | 0.494542294 | 0.00731708 | -1.0158342 | 2.13566227 |

| | | | | |
|---|---|---|---|---|
| ARPC5L | 0.494153754 | 0.2476935 | -1.0169681 | 0.60608538 |
| RPAP2 | 0.494142721 | 0.01050961 | -1.0170003 | 1.97841343 |
| ANO6 | 0.493758609 | 0.11024235 | -1.0181222 | 0.95765153 |
| PCNP | 0.493542053 | 0.00113341 | -1.0187551 | 2.94561113 |
| NDRG1 | 0.49319537 | 0.00919048 | -1.0197688 | 2.03666176 |
| TDRKH | 0.492799816 | 0.0011993 | -1.0209264 | 2.92107189 |
| CCNT2 | 0.492478493 | 0.30696317 | -1.0218674 | 0.51291372 |
| PRPF39 | 0.492250618 | 0.00581875 | -1.0225351 | 2.23517015 |
| AP3B2 | 0.492142671 | 0.0012844 | -1.0228515 | 2.89129875 |
| PAICS | 0.492132598 | 0.01136678 | -1.022881 | 1.94436258 |
| GPS2 | 0.491969421 | 0.00503633 | -1.0233594 | 2.29788556 |
| ARHGAP11A | 0.491763548 | 0.02722895 | -1.0239633 | 1.56496913 |
| MSH3 | 0.491330556 | 0.00175988 | -1.0252341 | 2.75451617 |
| UACA | 0.490255031 | 0.03307883 | -1.0283957 | 1.48044992 |
| VMP1 | 0.489695375 | 0.0027072 | -1.0300435 | 2.56748008 |
| USP36 | 0.488884711 | 0.0417643 | -1.0324338 | 1.37919474 |
| VGLL4 | 0.488847624 | 0.01532682 | -1.0325433 | 1.81454789 |
| TLN2 | 0.488831764 | 0.00248227 | -1.0325901 | 2.60515103 |
| FAM76B | 0.488735166 | 0.0031404 | -1.0328752 | 2.50301445 |
| NOP14 | 0.488330649 | 0.00249367 | -1.0340698 | 2.60316185 |
| DSN1 | 0.488211113 | 0.02122162 | -1.034423 | 1.67322144 |
| SCML2 | 0.487582669 | 0.00039543 | -1.0362812 | 3.40293187 |
| URB1 | 0.486567753 | 0.03096875 | -1.0392874 | 1.50907638 |
| ZFAND5 | 0.485355258 | 0.25427365 | -1.042887 | 0.59469864 |
| ANKRD44 | 0.485130791 | 0.18251157 | -1.0435543 | 0.73870959 |
| OSBPL11 | 0.484903982 | 0.0009562 | -1.044229 | 3.01945133 |
| TTC21B | 0.484765016 | 0.10393264 | -1.0446425 | 0.98324805 |
| MED15 | 0.484272369 | 0.00146651 | -1.0461094 | 2.83371358 |
| AQR | 0.483958878 | 0.00842635 | -1.0470436 | 2.07436062 |
| MAGI3 | 0.483895623 | 0.03050276 | -1.0472322 | 1.5156608 |
| UTP11 | 0.483452913 | 0.02600169 | -1.0485527 | 1.58499839 |
| RPS6KA3 | 0.483185864 | 0.04929448 | -1.0493498 | 1.30720171 |
| ACTL6B | 0.482434736 | 0.07534514 | -1.0515943 | 1.12294476 |
| TECPR2 | 0.481829952 | 0.14534568 | -1.053404 | 0.83759787 |
| CDK5 | 0.481683703 | 0.00655729 | -1.053842 | 2.18327572 |
| ILVBL | 0.481615997 | 0.0070888 | -1.0540448 | 2.14942711 |
| ZNF7 | 0.481487646 | 0.1322513 | -1.0544293 | 0.87860006 |
| DCP2 | 0.481250246 | 0.22143416 | -1.0551408 | 0.65475538 |
| MAP7D2 | 0.479997279 | 0.00377495 | -1.0589019 | 2.42308859 |
| NELFB | 0.479570148 | 0.01517786 | -1.0601862 | 1.81878947 |
| H2AC20 | 0.479326824 | 0.28024242 | -1.0609184 | 0.55246613 |
| TTYH3 | 0.479194978 | 0.03837665 | -1.0613153 | 1.41593294 |
| GOLPH3 | 0.479174938 | 0.1499579 | -1.0613756 | 0.82403064 |
| TMEM106C | 0.478443948 | 0.01583977 | -1.0635782 | 1.8002512 |
| CHD3 | 0.478198539 | 0.01735042 | -1.0643184 | 1.76069002 |

| | | | | |
|---|---|---|---|---|
| RPUSD2 | 0.478132546 | 0.00031078 | -1.0645175 | 3.50754055 |
| OGFR | 0.478126704 | 0.01110431 | -1.0645351 | 1.95450831 |
| NINL | 0.478107781 | 0.1661339 | -1.0645922 | 0.77954175 |
| CBX5 | 0.477537953 | 0.02524931 | -1.0663127 | 1.59775045 |
| COPS7B | 0.477397193 | 0.02215374 | -1.066738 | 1.65455294 |
| ZFR | 0.477298636 | 0.0088418 | -1.0670359 | 2.05345932 |
| SMC1A | 0.476716976 | 0.00084096 | -1.0687951 | 3.07522645 |
| DDX51 | 0.476231455 | 0.15902229 | -1.0702652 | 0.79854199 |
| DCX | 0.476226699 | 0.09446471 | -1.0702796 | 1.02473039 |
| QTRT2 | 0.474829209 | 0.06069411 | -1.0745194 | 1.21685347 |
| MORC3 | 0.474789259 | 0.04486207 | -1.0746408 | 1.34812064 |
| KIFC1 | 0.474659838 | 7.3936E-06 | -1.0750341 | 5.13114175 |
| MARS1 | 0.47460032 | 0.0028287 | -1.075215 | 2.54841335 |
| RAB39A | 0.474514417 | 0.05736972 | -1.0754762 | 1.24131729 |
| SRFBP1 | 0.474237699 | 0.01332014 | -1.0763177 | 1.87549111 |
| CTU2 | 0.474151501 | 0.01027164 | -1.07658 | 1.98836003 |
| KAT6A | 0.474017599 | 0.00856823 | -1.0769875 | 2.06710889 |
| PSME3 | 0.473864343 | 0.00098983 | -1.077454 | 3.00443952 |
| NOC4L | 0.473643769 | 0.17084164 | -1.0781257 | 0.76740628 |
| MSH6 | 0.473629178 | 0.00329602 | -1.0781701 | 2.48201013 |
| NCBP1 | 0.473605519 | 0.00355004 | -1.0782422 | 2.4497665 |
| SNRPC | 0.473229068 | 0.01962396 | -1.0793894 | 1.70721337 |
| CLCC1 | 0.473073151 | 0.01177488 | -1.0798648 | 1.92904342 |
| EEF1G | 0.472987736 | 0.00594098 | -1.0801253 | 2.2261421 |
| MYBBP1A | 0.472791886 | 0.0105345 | -1.0807228 | 1.97738588 |
| ETV3 | 0.472464757 | 0.07067714 | -1.0817214 | 1.15072102 |
| NCAPG | 0.470675188 | 0.02699458 | -1.0871963 | 1.56872341 |
| ARHGEF1 | 0.470051389 | 0.00490217 | -1.0891096 | 2.30961189 |
| POR | 0.469964502 | 0.12086111 | -1.0893763 | 0.91771341 |
| SPDL1 | 0.468047534 | 0.03963568 | -1.095273 | 1.40191367 |
| DHX57 | 0.467948602 | 0.04417674 | -1.095578 | 1.35480636 |
| KIF13B | 0.467879836 | 0.00116882 | -1.09579 | 2.93225292 |
| TOMM20 | 0.467471418 | 0.12756042 | -1.0970499 | 0.89428404 |
| PLEC | 0.467070285 | 0.00498402 | -1.0982884 | 2.30242059 |
| OGT | 0.466158751 | 2.0627E-05 | -1.1011067 | 4.68556724 |
| TSSC4 | 0.465663452 | 0.00859102 | -1.1026404 | 2.06595539 |
| MRPS9 | 0.465456439 | 0.12799737 | -1.1032819 | 0.89279894 |
| GTF2A1 | 0.46532973 | 0.04952748 | -1.1036747 | 1.30515381 |
| TAF5 | 0.46452108 | 0.10467794 | -1.106184 | 0.98014484 |
| CLP1 | 0.46285779 | 0.20343771 | -1.1113591 | 0.69156854 |
| PIH1D1 | 0.462763241 | 0.00022524 | -1.1116538 | 3.64734531 |
| ARFGEF3 | 0.462102183 | 0.03624118 | -1.1137162 | 1.44079771 |
| ATM | 0.461541926 | 0.01695838 | -1.1154664 | 1.77061565 |
| LTN1 | 0.461281105 | 0.04803017 | -1.1162819 | 1.3184859 |
| RFC3 | 0.46097197 | 0.03024582 | -1.1172491 | 1.51933469 |

| | | | | |
|---|---|---|---|---|
| TAF15 | 0.45855191 | 0.00345332 | -1.124843 | 2.46176373 |
| HELLS | 0.457528147 | 0.00426006 | -1.1280676 | 2.37058378 |
| CCNL1 | 0.456586234 | 0.00283386 | -1.1310407 | 2.54762146 |
| PGAM5 | 0.456507086 | 0.00019944 | -1.1312908 | 3.70017792 |
| KIF16B | 0.455870632 | 0.00994678 | -1.1333036 | 2.00231742 |
| MCMBP | 0.455430611 | 0.00783865 | -1.1346968 | 2.10575875 |
| TTC33 | 0.455306842 | 0.08659867 | -1.135089 | 1.0624888 |
| TBC1D2 | 0.454886394 | 0.10695417 | -1.1364218 | 0.9708023 |
| ARPC1B | 0.454772507 | 0.00795676 | -1.1367831 | 2.09926378 |
| GTSE1 | 0.454731048 | 0.00642328 | -1.1369146 | 2.19224328 |
| EPC1 | 0.454341394 | 0.00110192 | -1.1381513 | 2.95785117 |
| DPY30 | 0.453999428 | 0.00175179 | -1.1392376 | 2.75651881 |
| PPIG | 0.453844647 | 0.17392471 | -1.1397296 | 0.75963871 |
| ERC1 | 0.451775919 | 0.00441454 | -1.1463207 | 2.3551141 |
| BDP1 | 0.450938319 | 0.07442242 | -1.148998 | 1.1282962 |
| MATR3 | 0.449305728 | 0.00027159 | -1.1542306 | 3.56609104 |
| DIS3L | 0.448613544 | 0.16903367 | -1.1564549 | 0.77202677 |
| MPP2 | 0.448563443 | 0.00640629 | -1.156616 | 2.19339355 |
| NCOA5 | 0.448392265 | 0.00132813 | -1.1571667 | 2.87675799 |
| SNX6 | 0.448173809 | 0.00409157 | -1.1578698 | 2.38810958 |
| FLNB | 0.447825864 | 0.00125325 | -1.1589902 | 2.901961 |
| PIGU | 0.447379103 | 0.08527453 | -1.1604302 | 1.06918067 |
| VPS41 | 0.446954813 | 0.03766335 | -1.1617991 | 1.42408102 |
| CCDC6 | 0.446907367 | 0.0004781 | -1.1619523 | 3.32048502 |
| MAGOH | 0.445998653 | 0.32101517 | -1.1648887 | 0.49347444 |
| ZDBF2 | 0.445642585 | 0.00215366 | -1.166041 | 2.66682319 |
| FAM114A2 | 0.445468863 | 0.01079289 | -1.1666035 | 1.96686224 |
| BZW1 | 0.444882721 | 0.18533914 | -1.168503 | 0.73203286 |
| VBP1 | 0.444835766 | 0.06179057 | -1.1686553 | 1.20907778 |
| PPP1R9B | 0.444585115 | 0.03909282 | -1.1694684 | 1.40790303 |
| NDE1 | 0.443473172 | 0.17697143 | -1.1730813 | 0.75209683 |
| SNRPD2 | 0.443454261 | 0.00146493 | -1.1731428 | 2.83418397 |
| C12orf43 | 0.442217034 | 0.01470518 | -1.1771735 | 1.83252964 |
| PPP6R1 | 0.442155382 | 0.00662514 | -1.1773746 | 2.17880462 |
| ANKRD28 | 0.441822107 | 0.00295023 | -1.1784625 | 2.53014384 |
| RARS1 | 0.441048529 | 0.00089487 | -1.1809907 | 3.04823938 |
| PEG10 | 0.441029509 | 0.00304408 | -1.1810529 | 2.51654346 |
| PWP2 | 0.441021093 | 0.00087289 | -1.1810804 | 3.05904155 |
| FKBP8 | 0.440427966 | 0.00504638 | -1.183022 | 2.29701972 |
| MAD1L1 | 0.440370537 | 0.00050569 | -1.1832101 | 3.29611237 |
| ZNF326 | 0.440264554 | 0.00189608 | -1.1835574 | 2.72214261 |
| TSC2 | 0.439490383 | 0.0025945 | -1.1860965 | 2.58594589 |
| BOD1L1 | 0.439264268 | 5.1075E-05 | -1.1868389 | 4.29178896 |
| MRTFB | 0.439138233 | 3.245E-05 | -1.1872529 | 4.4887834 |
| NUDCD1 | 0.439102762 | 0.07860975 | -1.1873695 | 1.10452359 |

| | | | | |
|---|---|---|---|---|
| POLR1E | 0.438931695 | 0.01765816 | -1.1879316 | 1.75305466 |
| DDX24 | 0.437995705 | 0.00283228 | -1.1910114 | 2.54786338 |
| MKLN1 | 0.437805052 | 0.01172044 | -1.1916395 | 1.93105614 |
| USP10 | 0.43741956 | 0.01073117 | -1.1929104 | 1.96935294 |
| POLR2B | 0.437358507 | 0.00070396 | -1.1931117 | 3.15245309 |
| BOP1 | 0.4372074 | 0.00336431 | -1.1936103 | 2.47310356 |
| ADNP | 0.436908719 | 9.1395E-05 | -1.1945962 | 4.03907851 |
| PCP4 | 0.436421459 | 0.0227103 | -1.1962061 | 1.64377715 |
| BRAF | 0.436334179 | 0.00494851 | -1.1964946 | 2.30552554 |
| BLTP2 | 0.436198871 | 0.05214436 | -1.1969421 | 1.28279263 |
| KIF3A | 0.435571546 | 0.07343194 | -1.1990184 | 1.13411501 |
| CENPH | 0.43525045 | 0.074573 | -1.2000823 | 1.12741838 |
| BRD7 | 0.435210304 | 0.00760544 | -1.2002154 | 2.11887574 |
| VPS16 | 0.435105931 | 0.02322339 | -1.2005614 | 1.63407444 |
| HAUS8 | 0.435018788 | 0.01025935 | -1.2008504 | 1.98888009 |
| C7orf50 | 0.434963125 | 0.00092347 | -1.201035 | 3.0345779 |
| VPS13D | 0.434285896 | 0.11715988 | -1.203283 | 0.93122107 |
| CSNK2A2 | 0.433819771 | 0.03829936 | -1.2048323 | 1.41680853 |
| ZNF771 | 0.433617743 | 0.84050316 | -1.2055043 | 0.07546065 |
| DDX41 | 0.433486632 | 0.02199607 | -1.2059406 | 1.65765485 |
| REXO4 | 0.433174796 | 0.00047991 | -1.2069788 | 3.31883667 |
| ELF2 | 0.43303581 | 0.08798015 | -1.2074418 | 1.05561532 |
| DRG1 | 0.432827812 | 0.01361696 | -1.2081349 | 1.86591981 |
| NKRF | 0.432319006 | 0.00025655 | -1.2098318 | 3.59082556 |
| PTBP1 | 0.432286213 | 0.57170001 | -1.2099413 | 0.2428318 |
| NPM1 | 0.430461069 | 0.03204255 | -1.2160453 | 1.49427294 |
| COG7 | 0.430322588 | 0.06519518 | -1.2165095 | 1.18578449 |
| CROCC | 0.430236625 | 0.25151103 | -1.2167978 | 0.59944296 |
| DKC1 | 0.429895364 | 0.00063968 | -1.2179425 | 3.19403648 |
| MYO9A | 0.429879643 | 0.55996756 | -1.2179953 | 0.25183713 |
| ZNF622 | 0.42986559 | 0.03737951 | -1.2180425 | 1.42736639 |
| HNRNPC | 0.429543184 | 0.00280197 | -1.2191249 | 2.55253673 |
| GTF2H4 | 0.429029181 | 0.04637647 | -1.2208523 | 1.33370231 |
| VPS72 | 0.428840144 | 0.01720405 | -1.2214881 | 1.76436927 |
| ZNF664 | 0.428489447 | 0.07696289 | -1.2226684 | 1.11371861 |
| TMEM209 | 0.428390362 | 0.03300027 | -1.2230021 | 1.48148252 |
| MED17 | 0.428152381 | 0.0859829 | -1.2238037 | 1.06558791 |
| ARHGAP4 | 0.427905319 | 0.01633111 | -1.2246365 | 1.78698417 |
| LAMTOR1 | 0.426766297 | 0.07586492 | -1.2284818 | 1.11995901 |
| BCR | 0.426328539 | 0.01177685 | -1.2299625 | 1.92897082 |
| SNRNP70 | 0.425795886 | 0.00245895 | -1.2317661 | 2.6092497 |
| MARCKS | 0.425672493 | 0.02771154 | -1.2321842 | 1.55733933 |
| DNM1L | 0.425460936 | 0.00171193 | -1.2329014 | 2.76651305 |
| MCM10 | 0.42541067 | 0.07591422 | -1.2330719 | 1.11967686 |
| JAKMIP1 | 0.4252664 | 0.00164075 | -1.2335612 | 2.78495645 |

| | | | | |
|---|---|---|---|---|
| KHDC4 | 0.425026852 | 0.95669164 | -1.2343741 | 0.01922802 |
| PRR5 | 0.424342038 | 0.59485775 | -1.2367005 | 0.22558688 |
| FLYWCH2 | 0.424008126 | 0.00010324 | -1.2378362 | 3.98614328 |
| CC2D1B | 0.423533535 | 0.00065041 | -1.2394519 | 3.18681184 |
| TRIM65 | 0.422847804 | 0.00440286 | -1.2417896 | 2.35626559 |
| CENPI | 0.422827202 | 0.06109819 | -1.2418599 | 1.21397168 |
| POLR3C | 0.422796071 | 0.02076437 | -1.2419661 | 1.68268132 |
| CCDC86 | 0.422543737 | 0.01274245 | -1.2428274 | 1.89474718 |
| DDRGK1 | 0.42237228 | 0.0049462 | -1.2434129 | 2.30572867 |
| PIP4K2B | 0.422203216 | 0.00773431 | -1.2439905 | 2.11157815 |
| KATNA1 | 0.422118707 | 0.08816145 | -1.2442793 | 1.05472126 |
| MEF2D | 0.421569801 | 0.04194583 | -1.2461566 | 1.3773112 |
| TANGO6 | 0.421357422 | 0.22439218 | -1.2468836 | 0.64899228 |
| KATNB1 | 0.42130994 | 0.00663015 | -1.2470461 | 2.1784764 |
| ACTL6A | 0.421085543 | 0.00030868 | -1.2478147 | 3.51049406 |
| EIF4G3 | 0.420942529 | 0.0003355 | -1.2483048 | 3.47430879 |
| NDRG3 | 0.420462452 | 0.10746278 | -1.2499511 | 0.96874194 |
| L3MBTL4 | 0.419893886 | 0.20670293 | -1.2519033 | 0.68465337 |
| NAP1L1 | 0.419815098 | 0.00287128 | -1.252174 | 2.54192395 |
| RMI1 | 0.419499749 | 0.0371082 | -1.2532581 | 1.43053013 |
| DNAJC9 | 0.419484638 | 0.00017261 | -1.2533101 | 3.76294183 |
| CCNK | 0.419397754 | 0.00306187 | -1.253609 | 2.51401339 |
| CTCF | 0.419316228 | 0.00789967 | -1.2538894 | 2.10239132 |
| EIF4G1 | 0.41927982 | 0.00060263 | -1.2540147 | 3.21995208 |
| SF3B3 | 0.419243381 | 0.00042452 | -1.2541401 | 3.37210372 |
| TBC1D10B | 0.419203291 | 0.00503771 | -1.2542781 | 2.29776719 |
| ADAT1 | 0.418933031 | 0.03721098 | -1.2552085 | 1.42932886 |
| PPP6C | 0.418505698 | 0.013072 | -1.2566808 | 1.8836578 |
| THUMPD3 | 0.418025664 | 0.0284135 | -1.2583366 | 1.54647529 |
| BNIP1 | 0.418004704 | 0.01195434 | -1.2584089 | 1.92247443 |
| NAT10 | 0.417762287 | 0.00954139 | -1.2592458 | 2.02038829 |
| CHD6 | 0.417243074 | 0.00099226 | -1.26104 | 3.0033737 |
| ZNF317 | 0.417097734 | 0.05157372 | -1.2615426 | 1.28757154 |
| WDFY1 | 0.416917085 | 0.00031813 | -1.2621676 | 3.49739743 |
| OSBPL10 | 0.41655137 | 0.13179632 | -1.2634337 | 0.88009673 |
| MBD4 | 0.416296955 | 0.01034712 | -1.2643151 | 1.98518035 |
| DYNC1H1 | 0.415963741 | 0.00200682 | -1.2654703 | 2.69749205 |
| ERLIN1 | 0.415953216 | 0.00104805 | -1.2655068 | 2.97961921 |
| GRWD1 | 0.415460904 | 0.05926433 | -1.2672154 | 1.2272066 |
| ATAD2 | 0.415176442 | 0.00139814 | -1.2682035 | 2.85444865 |
| XRCC1 | 0.415028646 | 0.01190737 | -1.2687172 | 1.92418433 |
| MACF1 | 0.414153845 | 0.13641877 | -1.2717613 | 0.86512588 |
| C2CD4C | 0.413528418 | 0.27240309 | -1.2739416 | 0.56478797 |
| ETF1 | 0.412156868 | 0.02037421 | -1.2787346 | 1.69091926 |
| GSPT1 | 0.411959488 | 0.00163268 | -1.2794256 | 2.78709988 |

| | | | | |
|---|---|---|---|---|
| PHIP | 0.411734713 | 0.00013308 | -1.280213 | 3.87589828 |
| FASN | 0.411632546 | 0.00132293 | -1.280571 | 2.87846191 |
| MLH1 | 0.411290221 | 0.00015288 | -1.2817713 | 3.81563698 |
| CIZ1 | 0.411209637 | 0.06071676 | -1.282054 | 1.21669144 |
| USP8 | 0.411088907 | 0.00754581 | -1.2824777 | 2.12229435 |
| STMN2 | 0.410286857 | 0.0025162 | -1.2852952 | 2.59925504 |
| ANAPC1 | 0.410079387 | 0.01750975 | -1.2860249 | 1.75671999 |
| SPATS2 | 0.410048686 | 0.00596867 | -1.2861329 | 2.22412241 |
| BTF3 | 0.40977496 | 0.01832319 | -1.2870963 | 1.73699886 |
| PHLDB3 | 0.409099675 | 0.00853672 | -1.2894757 | 2.06870902 |
| PRC1 | 0.409026052 | 0.0056331 | -1.2897354 | 2.24925231 |
| CBX2 | 0.408695764 | 0.0052431 | -1.2909008 | 2.28041164 |
| ASF1A | 0.408347246 | 0.00643821 | -1.2921316 | 2.19123503 |
| DDX42 | 0.408258493 | 0.00157384 | -1.2924452 | 2.80303853 |
| PPM1G | 0.408229161 | 0.00515108 | -1.2925489 | 2.28810136 |
| KMT2E | 0.408112734 | 8.9814E-05 | -1.2929604 | 4.04665568 |
| IRF3 | 0.407499185 | 0.00270011 | -1.2951309 | 2.56861892 |
| NFIB | 0.407196865 | 0.00453833 | -1.2962016 | 2.34310404 |
| MTFR1L | 0.406989938 | 0.00017634 | -1.296935 | 3.75364391 |
| SOX6 | 0.406407715 | 0.01028598 | -1.2990003 | 1.9877543 |
| WDR59 | 0.405885249 | 0.00229918 | -1.3008562 | 2.63842729 |
| FLCN | 0.405772133 | 0.01282762 | -1.3012583 | 1.89185385 |
| PML | 0.405726477 | 0.00740143 | -1.3014206 | 2.13068441 |
| NCAPH | 0.405289348 | 0.0438739 | -1.3029758 | 1.35779371 |
| RBMXL1 | 0.405253067 | 0.00364468 | -1.303105 | 2.4383411 |
| RRP8 | 0.405179683 | 0.00556743 | -1.3033663 | 2.25434526 |
| PMS1 | 0.404992178 | 0.00347092 | -1.3040341 | 2.45955594 |
| SLC4A1AP | 0.404574109 | 0.00411547 | -1.3055241 | 2.38558077 |
| INF2 | 0.404488762 | 0.00187373 | -1.3058285 | 2.72729379 |
| GAR1 | 0.4037017 | 0.00509859 | -1.3086384 | 2.29255016 |
| SETD2 | 0.403434713 | 0.00336455 | -1.3095929 | 2.473073 |
| RAB3GAP2 | 0.403198826 | 0.00731735 | -1.3104367 | 2.13564632 |
| KNTC1 | 0.402805433 | 0.00302229 | -1.311845 | 2.51966457 |
| ZNF124 | 0.402644318 | 0.00299263 | -1.3124221 | 2.52394683 |
| CLASP1 | 0.402637819 | 0.00158049 | -1.3124454 | 2.80120816 |
| EHMT1 | 0.402128061 | 0.00068844 | -1.3142731 | 3.16213366 |
| MELK | 0.401910198 | 0.02195557 | -1.3150549 | 1.65845536 |
| NKTR | 0.401824391 | 0.00073015 | -1.315363 | 3.13658988 |
| SNRNP40 | 0.401105028 | 0.00114355 | -1.317948 | 2.94174311 |
| ATP1A3 | 0.400459437 | 0.20954907 | -1.320272 | 0.67871425 |
| DDOST | 0.400455651 | 0.29409957 | -1.3202856 | 0.53150562 |
| RDX | 0.400178604 | 9.7373E-05 | -1.3212841 | 4.01156125 |
| SSRP1 | 0.399973862 | 0.00317662 | -1.3220224 | 2.49803406 |
| INTS9 | 0.399748239 | 0.19180185 | -1.3228364 | 0.71714722 |
| DDX59 | 0.399737252 | 0.00988652 | -1.3228761 | 2.00495639 |

| | | | | |
|---|---|---|---|---|
| ZNF646 | 0.3996503 | 0.29469502 | -1.3231899 | 0.5306272 |
| THOC3 | 0.399495041 | 0.11133454 | -1.3237505 | 0.95337007 |
| ZNF711 | 0.398709028 | 0.09302144 | -1.3265918 | 1.03141696 |
| SKIC3 | 0.398036293 | 0.00317437 | -1.3290281 | 2.49834255 |
| RANBP9 | 0.397432255 | 0.02678783 | -1.3312191 | 1.57206246 |
| KRR1 | 0.397183054 | 0.04435519 | -1.332124 | 1.35305557 |
| ABCF1 | 0.396920671 | 0.00053559 | -1.3330774 | 3.27116485 |
| SLAIN1 | 0.396792467 | 0.0328392 | -1.3335435 | 1.48360741 |
| SSH2 | 0.396705825 | 0.00120879 | -1.3338585 | 2.91765001 |
| GLO1 | 0.396611052 | 0.00230125 | -1.3342032 | 2.63803543 |
| CACUL1 | 0.396213436 | 0.00041063 | -1.3356503 | 3.38654917 |
| SF1 | 0.396095694 | 0.00595317 | -1.3360791 | 2.22525183 |
| FIGNL1 | 0.395758949 | 0.00885603 | -1.3373061 | 2.05276097 |
| KIF2C | 0.394818651 | 0.01081161 | -1.340738 | 1.96610945 |
| SRP54 | 0.394675322 | 0.01055402 | -1.3412618 | 1.97658223 |
| SUZ12 | 0.394106808 | 0.01617418 | -1.3433414 | 1.79117782 |
| NEFM | 0.393440469 | 0.00102149 | -1.3457827 | 2.99076793 |
| UTP20 | 0.393331134 | 0.01339701 | -1.3461837 | 1.8729921 |
| NRBP1 | 0.393173845 | 0.00824058 | -1.3467607 | 2.08404236 |
| TRMT2A | 0.393147116 | 0.15620324 | -1.3468588 | 0.80630996 |
| NOL6 | 0.393025789 | 0.00012736 | -1.3473041 | 3.89495015 |
| AVIL | 0.39219958 | 0.00147588 | -1.3503401 | 2.8309479 |
| BIRC6 | 0.392088862 | 0.00252013 | -1.3507474 | 2.5985764 |
| PRPF38A | 0.391444952 | 0.00915656 | -1.3531187 | 2.03826779 |
| RCOR2 | 0.390992788 | 0.28069885 | -1.3547861 | 0.55175937 |
| HNRNPF | 0.38984079 | 0.00074692 | -1.359043 | 3.12672714 |
| PCYT1A | 0.389726712 | 0.00165057 | -1.3594653 | 2.78236489 |
| PGAM1 | 0.389606454 | 0.01549016 | -1.3599105 | 1.80994415 |
| MED14 | 0.388433417 | 0.01309695 | -1.3642608 | 1.88282992 |
| QARS1 | 0.388333278 | 0.00424796 | -1.3646328 | 2.37181916 |
| SMC5 | 0.387983616 | 0.01603912 | -1.3659324 | 1.79481944 |
| MAP4 | 0.387215498 | 0.01586771 | -1.3687914 | 1.79948576 |
| PWWP3A | 0.386905809 | 0.16438038 | -1.3699457 | 0.78415001 |
| ARID1A | 0.386841156 | 0.00184005 | -1.3701868 | 2.73517131 |
| RNPS1 | 0.384511406 | 0.00052798 | -1.3789017 | 3.27738282 |
| LIMA1 | 0.383129455 | 0.00035147 | -1.3840962 | 3.45411567 |
| SAFB | 0.382873149 | 0.00010525 | -1.3850616 | 3.97779372 |
| AGAP3 | 0.382862177 | 0.0246303 | -1.385103 | 1.60853038 |
| CDC42BPB | 0.382461718 | 0.00038248 | -1.3866127 | 3.41739529 |
| CTR9 | 0.382383576 | 0.00079068 | -1.3869075 | 3.1020019 |
| FAM117B | 0.382210343 | 0.01159119 | -1.3875613 | 1.93587181 |
| NFATC2IP | 0.382171459 | 0.01574202 | -1.3877081 | 1.80293964 |
| ZNF721 | 0.381327193 | 0.0121764 | -1.3908987 | 1.91448123 |
| IARS1 | 0.381277144 | 0.00136323 | -1.391088 | 2.86543157 |
| BTAF1 | 0.381175851 | 0.00023153 | -1.3914714 | 3.63538689 |

| | | | | |
|---|---|---|---|---|
| TNFAIP2 | 0.381065728 | 0.00043261 | -1.3918882 | 3.36390169 |
| GPHN | 0.379862649 | 0.00368956 | -1.3964502 | 2.43302519 |
| METTL2B | 0.379709875 | 0.17246647 | -1.3970306 | 0.76329532 |
| USP48 | 0.379580785 | 0.05465779 | -1.3975211 | 1.26234797 |
| BRPF3 | 0.379368739 | 0.08160038 | -1.3983273 | 1.08830784 |
| MED8 | 0.379363431 | 0.06625801 | -1.3983475 | 1.17876161 |
| SRBD1 | 0.378935571 | 0.02436893 | -1.3999755 | 1.61316357 |
| WDR83 | 0.378755661 | 0.00159609 | -1.4006606 | 2.79694275 |
| ZNF644 | 0.377764582 | 0.00111493 | -1.4044406 | 2.9527519 |
| NBN | 0.377725527 | 0.00027165 | -1.4045898 | 3.56598897 |
| PPDPF | 0.377479063 | 0.00427864 | -1.4055315 | 2.36869465 |
| SLTM | 0.377068909 | 0.00023998 | -1.4070999 | 3.61981871 |
| TBC1D16 | 0.377057113 | 0.22063897 | -1.407145 | 0.65631778 |
| GBA2 | 0.375140569 | 0.00704987 | -1.4144968 | 2.15181915 |
| ITPR3 | 0.37501395 | 0.00074116 | -1.4149838 | 3.13008931 |
| NOL8 | 0.374286083 | 0.00531454 | -1.4177867 | 2.27453452 |
| AKAP9 | 0.374112669 | 0.02989188 | -1.4184553 | 1.52444679 |
| SERBP1 | 0.374108454 | 0.00463649 | -1.4184715 | 2.33381061 |
| CARMIL1 | 0.37380504 | 0.00071291 | -1.4196421 | 3.14696436 |
| TTC1 | 0.373552154 | 0.00095279 | -1.4206184 | 3.0210043 |
| ERCC4 | 0.372978969 | 0.03864256 | -1.4228338 | 1.41293413 |
| NCAPG2 | 0.372351687 | 0.01545253 | -1.4252622 | 1.81100037 |
| CCNT1 | 0.372183205 | 0.00137825 | -1.4259151 | 2.86067082 |
| RRP1B | 0.37211328 | 0.00011361 | -1.4261862 | 3.94457984 |
| VAV2 | 0.372110868 | 0.16201803 | -1.4261956 | 0.79043665 |
| HCFC2 | 0.37164502 | 0.00851196 | -1.4280028 | 2.06997022 |
| FAM120C | 0.371590975 | 0.01499891 | -1.4282126 | 1.82394024 |
| ELP4 | 0.37119884 | 0.0028286 | -1.4297359 | 2.54842868 |
| TIMELESS | 0.37099526 | 0.05854488 | -1.4305273 | 1.23251109 |
| CXXC1 | 0.370708787 | 0.1367812 | -1.4316418 | 0.8639736 |
| ZMYM6 | 0.370608544 | 0.12222683 | -1.432032 | 0.91283344 |
| HERC2 | 0.370489751 | 0.00975976 | -1.4324945 | 2.01056108 |
| MAP1B | 0.369959829 | 0.00030372 | -1.4345595 | 3.51752427 |
| WDR36 | 0.369669314 | 0.00198269 | -1.4356928 | 2.70274444 |
| SF3B5 | 0.369507748 | 0.01482442 | -1.4363235 | 1.82902234 |
| NAA25 | 0.369470784 | 0.05994556 | -1.4364678 | 1.22224295 |
| PHKB | 0.369445217 | 0.01352449 | -1.4365676 | 1.86887923 |
| SUPT6H | 0.369083417 | 0.00182736 | -1.4379812 | 2.73817524 |
| VPS53 | 0.368787815 | 0.0015805 | -1.4391371 | 2.8012058 |
| CLSPN | 0.368636222 | 0.009358 | -1.4397303 | 2.02881679 |
| SGF29 | 0.368586508 | 0.20778077 | -1.4399248 | 0.68239465 |
| GLOD4 | 0.368563429 | 0.00775136 | -1.4400152 | 2.11062233 |
| CHTF18 | 0.367378894 | 0.02772767 | -1.4446593 | 1.55708657 |
| SP6 | 0.367204574 | 0.25154741 | -1.4453441 | 0.59938014 |
| GOLPH3L | 0.367153802 | 0.00100605 | -1.4455436 | 2.99737992 |

| | | | | |
|---|---|---|---|---|
| DDHD1 | 0.366599596 | 0.00026544 | -1.4477229 | 3.57603166 |
| VAV3 | 0.365880111 | 0.19676097 | -1.4505571 | 0.70606104 |
| ZRANB3 | 0.365622916 | 0.07092467 | -1.4515716 | 1.14920268 |
| RIOK2 | 0.365361413 | 0.01896555 | -1.4526038 | 1.72203462 |
| LARS1 | 0.364748118 | 6.5626E-05 | -1.4550276 | 4.1829208 |
| HECTD3 | 0.364382091 | 0.1611174 | -1.456476 | 0.79285756 |
| NIPBL | 0.362972241 | 0.00014076 | -1.4620689 | 3.85150763 |
| NBEAL1 | 0.36218845 | 0.13822522 | -1.4651876 | 0.8594127 |
| UBA6 | 0.361994884 | 0.01412589 | -1.4659588 | 1.8499841 |
| PRR14L | 0.361970418 | 0.00551532 | -1.4660563 | 2.25842926 |
| CENPC | 0.361320869 | 0.00034793 | -1.4686475 | 3.45851005 |
| METTL3 | 0.361046893 | 0.00026831 | -1.4697419 | 3.57136114 |
| LRP12 | 0.360967932 | 0.01812906 | -1.4700574 | 1.74162483 |
| SAR1B | 0.358823111 | 0.09998744 | -1.4786553 | 1.00005453 |
| POLE | 0.358514358 | 0.00214562 | -1.4798972 | 2.66844677 |
| CDK5RAP3 | 0.358143631 | 0.29738367 | -1.4813898 | 0.52668288 |
| TUBGCP6 | 0.357339797 | 0.00617418 | -1.4846315 | 2.20942091 |
| INVS | 0.356681309 | 0.04256495 | -1.4872925 | 1.37094791 |
| SIPA1 | 0.356065383 | 0.04023689 | -1.4897859 | 1.39537558 |
| ZC3H8 | 0.3555961 | 0.0605091 | -1.4916886 | 1.2181793 |
| FAM200C | 0.355132652 | 0.00917529 | -1.4935701 | 2.03738017 |
| ABR | 0.354900899 | 0.02429269 | -1.4945119 | 1.6145244 |
| DYNC1LI2 | 0.35473676 | 0.00142173 | -1.4951793 | 2.84718214 |
| ACTR5 | 0.354707647 | 0.04163879 | -1.4952977 | 1.38050189 |
| RELCH | 0.353283767 | 0.00399267 | -1.5011006 | 2.39873625 |
| RAP1GAP | 0.35286808 | 0.04188711 | -1.5027992 | 1.37791957 |
| MMS22L | 0.352808042 | 0.24566242 | -1.5030446 | 0.60966128 |
| USP15 | 0.352581906 | 0.00312698 | -1.5039697 | 2.50487492 |
| DNTTIP2 | 0.352468676 | 0.00571545 | -1.504433 | 2.24294929 |
| ACLY | 0.352180446 | 0.2002707 | -1.5056133 | 0.69838258 |
| IPO11 | 0.35195452 | 0.00991496 | -1.5065391 | 2.00370893 |
| DIP2A | 0.351240897 | 0.00063382 | -1.5094673 | 3.19803319 |
| CGN | 0.350767788 | 0.00011835 | -1.5114118 | 3.92683683 |
| ALG2 | 0.350663482 | 0.00157726 | -1.5118409 | 2.80209784 |
| EXOSC9 | 0.350343409 | 0.03459889 | -1.5131583 | 1.46093781 |
| NFATC3 | 0.349383041 | 0.02648125 | -1.5171185 | 1.57706151 |
| ILF2 | 0.349338535 | 0.00028355 | -1.5173023 | 3.54736829 |
| TBL3 | 0.349286735 | 0.00014359 | -1.5175162 | 3.84287147 |
| BAP1 | 0.349242671 | 0.00649862 | -1.5176983 | 2.18717906 |
| USP4 | 0.349124112 | 0.30143197 | -1.5181881 | 0.52081069 |
| CNDP2 | 0.349025371 | 0.07929227 | -1.5185962 | 1.10076915 |
| SINHCAF | 0.348971466 | 0.01822779 | -1.518819 | 1.73926601 |
| CNST | 0.347969007 | 0.00093338 | -1.5229693 | 3.0299414 |
| PIK3C3 | 0.347868375 | 0.00456506 | -1.5233866 | 2.34055367 |
| ZGPAT | 0.347838886 | 0.06945841 | -1.5235089 | 1.15827515 |

| | | | | |
|---|---|---|---|---|
| PCM1 | 0.347714164 | 0.00056734 | -1.5240263 | 3.24616004 |
| SSX2IP | 0.346394194 | 0.0053434 | -1.5295133 | 2.27218248 |
| DAXX | 0.34629053 | 0.00320659 | -1.5299452 | 2.49395596 |
| RELA | 0.345920497 | 0.00415247 | -1.5314876 | 2.38169329 |
| LARP4B | 0.345084796 | 0.00090887 | -1.5349772 | 3.04149704 |
| CDC16 | 0.344779671 | 0.00692463 | -1.5362534 | 2.15960372 |
| ZZEF1 | 0.344659867 | 0.09924268 | -1.5367548 | 1.00330153 |
| MCRIP1 | 0.344252971 | 0.04589727 | -1.538459 | 1.33821318 |
| BRD8 | 0.344072645 | 0.00154287 | -1.5392149 | 2.8116697 |
| SAP30BP | 0.343571302 | 0.00333272 | -1.5413186 | 2.47720114 |
| GPATCH8 | 0.343394543 | 0.00288853 | -1.542061 | 2.53932252 |
| GPALPP1 | 0.343217625 | 0.00199861 | -1.5428045 | 2.69927261 |
| FAM111B | 0.343144515 | 0.00328138 | -1.5431118 | 2.48394345 |
| LARP4 | 0.343076942 | 0.00125263 | -1.5433959 | 2.90217556 |
| UBR4 | 0.342930132 | 0.00294887 | -1.5440134 | 2.53034435 |
| WDR3 | 0.342795486 | 0.00745346 | -1.54458 | 2.12764236 |
| STRIP2 | 0.342592972 | 0.10991284 | -1.5454325 | 0.95895157 |
| FSD1 | 0.342286039 | 0.00652294 | -1.5467256 | 2.18555653 |
| CASP2 | 0.342208521 | 0.14211871 | -1.5470524 | 0.84734875 |
| DCAF5 | 0.342188948 | 0.00078439 | -1.5471349 | 3.10546521 |
| THADA | 0.341993166 | 0.01588732 | -1.5479606 | 1.79894935 |
| TERF2IP | 0.341570505 | 0.02003513 | -1.5497447 | 1.69820793 |
| RAD21 | 0.34137116 | 0.00595196 | -1.5505869 | 2.2253402 |
| SP2 | 0.341304462 | 0.02028107 | -1.5508688 | 1.69290924 |
| TSR1 | 0.340866509 | 0.01077642 | -1.5527212 | 1.96752552 |
| CUL4B | 0.340811579 | 0.01277537 | -1.5529537 | 1.89362668 |
| PNPLA6 | 0.34047564 | 0.30436117 | -1.5543765 | 0.51661076 |
| MRPS14 | 0.340222987 | 0.01835057 | -1.5554475 | 1.73635047 |
| ARHGAP23 | 0.339917882 | 0.00728061 | -1.5567418 | 2.13783205 |
| TOP3A | 0.339808699 | 0.00267371 | -1.5572053 | 2.5728853 |
| GAMT | 0.339712916 | 0.26569318 | -1.557612 | 0.5756196 |
| PSIP1 | 0.339427374 | 0.00012496 | -1.5588252 | 3.90321738 |
| GPS1 | 0.338264546 | 0.00343523 | -1.5637761 | 2.46404463 |
| MYL6B | 0.337870718 | 0.13257031 | -1.5654568 | 0.87755373 |
| KMT2A | 0.337780942 | 0.00054277 | -1.5658402 | 3.26538553 |
| CCDC14 | 0.337171188 | 0.00249865 | -1.5684468 | 2.60229506 |
| FRA10AC1 | 0.3370037 | 0.00761394 | -1.5691637 | 2.11839077 |
| FEM1A | 0.336277172 | 0.11339078 | -1.5722772 | 0.94542225 |
| PCIF1 | 0.336007503 | 0.00801401 | -1.5734346 | 2.09615023 |
| NMT1 | 0.335987412 | 0.00842251 | -1.5735209 | 2.07455868 |
| TDRD7 | 0.335375116 | 0.01335636 | -1.5761524 | 1.87431175 |
| SNF8 | 0.335067808 | 0.00427443 | -1.577475 | 2.36912227 |
| NUMA1 | 0.334788282 | 0.00136825 | -1.5786791 | 2.86383608 |
| DOCK10 | 0.334555558 | 1.3116E-05 | -1.5796823 | 4.88220616 |
| CPSF7 | 0.333938191 | 0.0004532 | -1.582347 | 3.34371288 |

| | | | | |
|---|---|---|---|---|
| LIN54 | 0.333716924 | 0.01090238 | -1.5833032 | 1.96247872 |
| NSMCE2 | 0.333459564 | 1.3983E-05 | -1.5844163 | 4.8543925 |
| CIAO2B | 0.333398875 | 0.03230116 | -1.5846789 | 1.49078189 |
| UBR3 | 0.333051312 | 6.8258E-05 | -1.5861836 | 4.16584341 |
| CCDC117 | 0.332949375 | 0.16965884 | -1.5866253 | 0.77042352 |
| WDR12 | 0.332680054 | 0.00320812 | -1.5877927 | 2.49374946 |
| HMGB3 | 0.332187156 | 0.00225616 | -1.5899318 | 2.64663019 |
| PI4KB | 0.332121448 | 0.02940353 | -1.5902172 | 1.53160049 |
| TOP2A | 0.33190978 | 0.00096904 | -1.591137 | 3.01365943 |
| ATP13A1 | 0.331772802 | 0.09411021 | -1.5917325 | 1.02636324 |
| SF3B2 | 0.331577304 | 0.0004715 | -1.5925828 | 3.32651915 |
| COPS2 | 0.33157508 | 0.00746638 | -1.5925925 | 2.12688975 |
| PUS7 | 0.33109611 | 0.02185669 | -1.594678 | 1.66041552 |
| TARBP2 | 0.330609464 | 0.00779852 | -1.5968001 | 2.10798757 |
| ZNF250 | 0.330552987 | 0.20114526 | -1.5970465 | 0.6964902 |
| KIAA1522 | 0.329912893 | 4.5448E-05 | -1.5998429 | 4.34248691 |
| KLC4 | 0.329831339 | 0.00110817 | -1.6001996 | 2.95539468 |
| LIMS1 | 0.329570101 | 0.02549503 | -1.6013427 | 1.59354451 |
| POLA1 | 0.329472917 | 5.4018E-05 | -1.6017682 | 4.26746138 |
| NIFK | 0.327775845 | 0.05539954 | -1.6092186 | 1.25649385 |
| BRF1 | 0.327718725 | 0.23388684 | -1.60947 | 0.63099421 |
| MAP1S | 0.327000171 | 0.00044702 | -1.6126367 | 3.349677 |
| DNAAF10 | 0.326933924 | 0.00614198 | -1.612929 | 2.21169151 |
| GKAP1 | 0.326239157 | 0.00025775 | -1.6159981 | 3.5888008 |
| POLD3 | 0.32610974 | 0.00030651 | -1.6165706 | 3.51354884 |
| RIPOR2 | 0.32585678 | 0.16942304 | -1.6176901 | 0.77102752 |
| WNK1 | 0.32474847 | 0.00097021 | -1.6226054 | 3.01313428 |
| UBE3C | 0.324669119 | 0.01677596 | -1.6229579 | 1.77531273 |
| SMARCE1 | 0.324634125 | 0.00078846 | -1.6231134 | 3.10321769 |
| VIRMA | 0.324052279 | 0.00321286 | -1.6257015 | 2.49310844 |
| MEN1 | 0.323033383 | 0.00490691 | -1.6302448 | 2.30919153 |
| UBE2O | 0.322133348 | 0.11726904 | -1.6342701 | 0.93081663 |
| DIAPH3 | 0.321850307 | 2.3824E-05 | -1.6355383 | 4.6229907 |
| CEP170B | 0.321725222 | 0.00358484 | -1.6360991 | 2.44552976 |
| KHSRP | 0.321079359 | 9.7655E-05 | -1.6389982 | 4.01030476 |
| SNRPE | 0.319785627 | 0.01416766 | -1.644823 | 1.84870188 |
| CFL2 | 0.319705049 | 0.00231344 | -1.6451866 | 2.63574199 |
| PIP4K2C | 0.319370173 | 0.00470274 | -1.6466985 | 2.32764937 |
| SNRPB | 0.319253141 | 0.00013795 | -1.6472273 | 3.86028855 |
| BRAP | 0.319051996 | 4.7233E-05 | -1.6481365 | 4.32575394 |
| EVL | 0.318854611 | 0.00041469 | -1.6490293 | 3.38227926 |
| EXOC5 | 0.318770857 | 0.00770235 | -1.6494084 | 2.11337655 |
| TSN | 0.318490449 | 0.02224638 | -1.650678 | 1.65274063 |
| GTF3C4 | 0.3175908 | 0.0001618 | -1.654759 | 3.79103263 |
| RFC1 | 0.317366734 | 0.00106969 | -1.6557772 | 2.97074161 |

| | | | | |
|---|---|---|---|---|
| FUBP1 | 0.317074885 | 0.00057907 | -1.6571045 | 3.23727001 |
| NUDT9 | 0.315658939 | 0.16138092 | -1.6635615 | 0.7921478 |
| SMARCA5 | 0.315573552 | 0.02522154 | -1.6639518 | 1.59822831 |
| EEA1 | 0.315451232 | 0.00413764 | -1.6645111 | 2.38324717 |
| ROCK1 | 0.315208212 | 0.00715665 | -1.665623 | 2.14529009 |
| GIT1 | 0.314398734 | 0.00052451 | -1.6693327 | 3.28024359 |
| NFIA | 0.313980157 | 0.00193587 | -1.6712547 | 2.71312448 |
| LNPK | 0.313362223 | 0.00488887 | -1.6740968 | 2.31079178 |
| TMEM120B | 0.312805464 | 0.21726948 | -1.6766624 | 0.66300127 |
| AIMP1 | 0.312759041 | 0.00057293 | -1.6768765 | 3.24189682 |
| DCP1B | 0.311225015 | 0.00102472 | -1.6839701 | 2.98939681 |
| DDX49 | 0.310702687 | 0.00738986 | -1.6863934 | 2.13136401 |
| RPA2 | 0.310598227 | 0.0003639 | -1.6868785 | 3.43901835 |
| VCPIP1 | 0.310506405 | 0.01341389 | -1.6873051 | 1.8724454 |
| CDK19 | 0.310409017 | 0.44051543 | -1.6877576 | 0.35603888 |
| EXD2 | 0.310307669 | 0.01298585 | -1.6882287 | 1.88652949 |
| ERCC2 | 0.310194365 | 0.08971528 | -1.6887556 | 1.0471336 |
| DCTN3 | 0.310192928 | 0.00067385 | -1.6887623 | 3.17143834 |
| BCAS2 | 0.310166626 | 0.00014145 | -1.6888846 | 3.8493878 |
| PGM2L1 | 0.310154412 | 0.14342759 | -1.6889414 | 0.8433673 |
| ACTMAP | 0.310042506 | 0.04760935 | -1.6894621 | 1.32230778 |
| ZNF808 | 0.309133315 | 0.0302757 | -1.693699 | 1.51890587 |
| ZNF384 | 0.308996942 | 0.0006534 | -1.6943355 | 3.18482184 |
| SOGA3 | 0.308920007 | 0.00047572 | -1.6946948 | 3.32265079 |
| MYO19 | 0.308682563 | 0.0134448 | -1.6958041 | 1.87144578 |
| C1orf50 | 0.308474978 | 0.00178574 | -1.6967746 | 2.74818057 |
| DDX31 | 0.3075883 | 0.00263632 | -1.7009275 | 2.57900185 |
| ORC1 | 0.307344435 | 0.04789732 | -1.7020717 | 1.31968876 |
| RAMAC | 0.307076362 | 0.00099128 | -1.7033306 | 3.0038022 |
| MAPK13 | 0.306324464 | 0.03191089 | -1.7068675 | 1.49606105 |
| RBM27 | 0.306276778 | 0.00029141 | -1.7070921 | 3.53549389 |
| RIC8A | 0.306116468 | 0.00154609 | -1.7078474 | 2.81076543 |
| KATNAL1 | 0.305759888 | 0.00277525 | -1.7095289 | 2.55669723 |
| HJURP | 0.3053996 | 0.00081381 | -1.7112299 | 3.08947433 |
| BCCIP | 0.30535781 | 0.11514746 | -1.7114273 | 0.93874563 |
| ELP1 | 0.304576657 | 0.00177047 | -1.7151227 | 2.75191236 |
| ANKRD11 | 0.30415815 | 0.00081359 | -1.7171064 | 3.08959272 |
| NDRG4 | 0.303947284 | 0.00044136 | -1.718107 | 3.35520782 |
| NEMF | 0.303505145 | 0.00711659 | -1.7202071 | 2.14772789 |
| AGPAT2 | 0.30336764 | 0.10717924 | -1.7208609 | 0.96988934 |
| BRIP1 | 0.302979414 | 0.00587404 | -1.7227083 | 2.23106302 |
| SNW1 | 0.302919881 | 0.0001987 | -1.7229918 | 3.70179358 |
| CTTNBP2NL | 0.30271165 | 0.0144615 | -1.7239839 | 1.83978653 |
| EIF5B | 0.302680412 | 0.00104927 | -1.7241328 | 2.97911202 |
| MED24 | 0.302357088 | 0.07277197 | -1.7256747 | 1.13803584 |

| | | | | |
|---|---|---|---|---|
| CHCHD2P9 | 0.301619656 | 0.16771167 | -1.7291976 | 0.77543672 |
| GTF2B | 0.301581792 | 0.00097252 | -1.7293788 | 3.01210083 |
| ARIH2 | 0.300954613 | 0.08133453 | -1.7323822 | 1.08972506 |
| RBM10 | 0.300655489 | 0.00028415 | -1.7338168 | 3.5464556 |
| USP37 | 0.3006146 | 0.00272291 | -1.734013 | 2.56496725 |
| TAF7 | 0.300573788 | 0.033187 | -1.7342089 | 1.47903205 |
| CCDC174 | 0.300312172 | 0.16234736 | -1.7354651 | 0.78955477 |
| ILF3 | 0.299872778 | 0.00055469 | -1.7375775 | 3.25594726 |
| PHF20 | 0.29974153 | 0.04979591 | -1.7382091 | 1.30280632 |
| PPIP5K1 | 0.299659429 | 0.09131473 | -1.7386043 | 1.03945918 |
| FMNL3 | 0.299585072 | 0.03096358 | -1.7389624 | 1.50914877 |
| MYO5C | 0.299429199 | 0.16853412 | -1.7397132 | 0.77331215 |
| POP4 | 0.299122615 | 0.01017448 | -1.7411911 | 1.99248768 |
| PRPF31 | 0.298675764 | 0.00025348 | -1.7433479 | 3.59605414 |
| SMARCD2 | 0.298515958 | 0.00274239 | -1.74412 | 2.56187077 |
| TEAD1 | 0.298344848 | 0.00033426 | -1.7449472 | 3.47591515 |
| RPA3 | 0.296933642 | 0.00281021 | -1.7517875 | 2.55126103 |
| TTLL5 | 0.296248433 | 0.11733766 | -1.7551206 | 0.93056257 |
| KMT2D | 0.295854022 | 3.1421E-05 | -1.7570426 | 4.50278489 |
| DYNC1LI1 | 0.295705667 | 0.00145314 | -1.7577662 | 2.83769218 |
| AKAP1 | 0.295517208 | 0.00033493 | -1.758686 | 3.47504813 |
| MAPKBP1 | 0.29526137 | 1.6898E-05 | -1.7599355 | 4.77217548 |
| PRRC2C | 0.295092263 | 0.00022562 | -1.760762 | 3.64662869 |
| ATOH1 | 0.294904922 | 7.3976E-05 | -1.7616782 | 4.13090785 |
| MMS19 | 0.294893417 | 0.00645461 | -1.7617345 | 2.1901299 |
| CORO1B | 0.294526401 | 0.00884248 | -1.7635311 | 2.05342616 |
| SKIC2 | 0.294416147 | 0.00097404 | -1.7640713 | 3.01142134 |
| XRN2 | 0.294158662 | 1.8667E-05 | -1.7653336 | 4.72893334 |
| CENPE | 0.294016689 | 0.01439832 | -1.76603 | 1.84168821 |
| CDK7 | 0.293596853 | 0.02003044 | -1.7680916 | 1.69830945 |
| RECQL | 0.293158377 | 0.00033675 | -1.7702478 | 3.47269296 |
| SH3BP5L | 0.293079252 | 0.07593616 | -1.7706373 | 1.11955137 |
| COPS6 | 0.292852505 | 4.8338E-05 | -1.7717539 | 4.31571517 |
| PPP1R10 | 0.291787946 | 8.2584E-05 | -1.7770078 | 4.08310343 |
| CALM3 | 0.291676128 | 0.00011488 | -1.7775608 | 3.93976321 |
| SART1 | 0.291041271 | 0.00023485 | -1.7807043 | 3.62920585 |
| CIC | 0.290837571 | 0.00021041 | -1.7817144 | 3.67693556 |
| SPAG7 | 0.290595294 | 0.00842387 | -1.7829168 | 2.07448859 |
| USP7 | 0.290416352 | 0.00347572 | -1.7838054 | 2.45895557 |
| NCL | 0.290159385 | 0.00107008 | -1.7850825 | 2.97058256 |
| HMGB2 | 0.289239448 | 0.00096928 | -1.7896638 | 3.0135516 |
| TUT4 | 0.288817206 | 0.00138546 | -1.7917714 | 2.85840702 |
| PDE4DIP | 0.288452524 | 0.34167226 | -1.7935942 | 0.46639028 |
| PRMT9 | 0.288268907 | 0.00147597 | -1.7945129 | 2.83092177 |
| RBBP7 | 0.287715689 | 0.00049523 | -1.7972842 | 3.30519046 |

| | | | | |
|---|---|---|---|---|
| MOSPD2 | 0.287687954 | 0.00184738 | -1.7974233 | 2.73344266 |
| PPIH | 0.287560837 | 0.00034378 | -1.7980609 | 3.46371786 |
| HARS1 | 0.287475565 | 0.00838341 | -1.7984888 | 2.07657948 |
| TYW1 | 0.287172945 | 0.02739449 | -1.8000083 | 1.56233673 |
| ECT2 | 0.286664669 | 0.00039306 | -1.802564 | 3.40554309 |
| OCIAD1 | 0.286497514 | 0.14091108 | -1.8034055 | 0.85105485 |
| TNPO2 | 0.286246582 | 0.04359449 | -1.8046696 | 1.36056836 |
| MED11 | 0.286162253 | 0.01207147 | -1.8050947 | 1.9182398 |
| SIRT1 | 0.285504212 | 0.00012667 | -1.8084161 | 3.89733543 |
| RBM12 | 0.284999494 | 0.00044124 | -1.8109687 | 3.35532116 |
| AKAP13 | 0.28396344 | 0.00031873 | -1.8162229 | 3.49658228 |
| PTPA | 0.283775629 | 0.35448248 | -1.8171774 | 0.45040523 |
| ANKRD13A | 0.282964993 | 0.23800275 | -1.8213045 | 0.62341803 |
| DDX55 | 0.282832455 | 0.00140885 | -1.8219804 | 2.85113575 |
| SMG1 | 0.282722036 | 4.553E-06 | -1.8225438 | 5.34170462 |
| KIF5A | 0.282640926 | 0.00107487 | -1.8229577 | 2.96864409 |
| KRAS | 0.282523156 | 0.02974055 | -1.823559 | 1.52665104 |
| GLCCI1 | 0.282513707 | 0.00367637 | -1.8236072 | 2.43458051 |
| HEXIM1 | 0.282388138 | 0.00056883 | -1.8242486 | 3.24501468 |
| GTF2H1 | 0.28205779 | 0.05449934 | -1.8259373 | 1.26360877 |
| SF1 | 0.281923812 | 0.00028254 | -1.8266228 | 3.54891304 |
| WDHD1 | 0.281666584 | 0.0013845 | -1.8279397 | 2.85870809 |
| DDX52 | 0.281436081 | 0.0001474 | -1.8291208 | 3.83150652 |
| PSME3IP1 | 0.281197612 | 2.3623E-05 | -1.8303438 | 4.62666989 |
| SMCHD1 | 0.281042805 | 0.00022313 | -1.8311382 | 3.65143792 |
| MAP3K2 | 0.280757293 | 0.01861869 | -1.8326046 | 1.73005082 |
| PDS5B | 0.280346476 | 0.00014133 | -1.8347172 | 3.84977325 |
| ARPC5 | 0.279764552 | 0.03720603 | -1.8377149 | 1.42938673 |
| CSTF3 | 0.279470419 | 0.00333597 | -1.8392325 | 2.47677734 |
| PKN1 | 0.279042553 | 0.00218912 | -1.841443 | 2.65973004 |
| ZNF567 | 0.278670367 | 0.0157217 | -1.8433685 | 1.80350057 |
| ACTR8 | 0.277308081 | 0.06339554 | -1.8504384 | 1.19794131 |
| ZSCAN31 | 0.276766183 | 0.00060547 | -1.8532604 | 3.21790504 |
| EAF1 | 0.276300839 | 0.02967691 | -1.8556882 | 1.52758127 |
| SUPT7L | 0.276220365 | 0.17795943 | -1.8561084 | 0.74967899 |
| SPIN1 | 0.276175856 | 0.05777711 | -1.8563409 | 1.23824418 |
| TBC1D8 | 0.276012865 | 0.03579614 | -1.8571926 | 1.44616385 |
| PPHLN1 | 0.275870919 | 0.00085234 | -1.8579347 | 3.06938551 |
| WDCP | 0.27549924 | 0.0069429 | -1.8598798 | 2.15845889 |
| RTF1 | 0.275283807 | 0.00020764 | -1.8610083 | 3.68269227 |
| TTF1 | 0.274752943 | 0.03607758 | -1.8637932 | 1.44276261 |
| MTF2 | 0.273277206 | 0.00352222 | -1.871563 | 2.45318398 |
| AGGF1 | 0.272836173 | 0.00107649 | -1.8738932 | 2.96799172 |
| EPRS1 | 0.272623049 | 0.00040732 | -1.8750206 | 3.39006894 |
| CPSF2 | 0.271834693 | 0.000244 | -1.8791985 | 3.61260687 |

| | | | | |
|---|---|---|---|---|
| KIF11 | 0.271805628 | 0.0001439 | -1.8793528 | 3.84193693 |
| CNOT8 | 0.271587719 | 0.01428829 | -1.8805099 | 1.84501985 |
| TCEAL4 | 0.271349308 | 0.02476442 | -1.8817769 | 1.60617189 |
| BMS1 | 0.27125149 | 0.00078772 | -1.882297 | 3.10362854 |
| GFI1 | 0.271157201 | 0.00032042 | -1.8827986 | 3.49427949 |
| ZNF207 | 0.271127448 | 0.01018191 | -1.8829569 | 1.99217079 |
| GTF2F2 | 0.271007883 | 3.1844E-05 | -1.8835933 | 4.4969779 |
| KIF3B | 0.270738124 | 0.03395743 | -1.88503 | 1.46906516 |
| ANK1 | 0.270476472 | 0.00627032 | -1.886425 | 2.20271038 |
| MORC2 | 0.270379431 | 1.1502E-05 | -1.8869427 | 4.93921373 |
| ZNF830 | 0.270214154 | 0.00114563 | -1.8878248 | 2.94095631 |
| TRIP12 | 0.269863309 | 6.1978E-05 | -1.8896993 | 4.20776241 |
| UBN2 | 0.26974866 | 0.015057 | -1.8903123 | 1.82226166 |
| TASOR | 0.269356417 | 0.00252935 | -1.8924117 | 2.59699061 |
| TACC3 | 0.268890529 | 0.00144989 | -1.8949092 | 2.83866616 |
| CDK9 | 0.268870785 | 7.7664E-05 | -1.8950151 | 4.10978072 |
| KPNA3 | 0.268847656 | 0.00666518 | -1.8951392 | 2.17618795 |
| NEK9 | 0.268596022 | 0.00016032 | -1.8964902 | 3.79500091 |
| PCYT2 | 0.268392633 | 0.04833503 | -1.897583 | 1.31573804 |
| NCBP3 | 0.268128391 | 0.00408093 | -1.8990041 | 2.38924041 |
| HAT1 | 0.267768014 | 0.00641368 | -1.9009445 | 2.19289266 |
| FAM193A | 0.267307325 | 9.9252E-05 | -1.9034287 | 4.00325973 |
| TXLNA | 0.267168601 | 0.00010185 | -1.9041776 | 3.99205702 |
| NUDT21 | 0.267154825 | 0.00019723 | -1.904252 | 3.70501804 |
| TASOR2 | 0.267077573 | 0.00335948 | -1.9046693 | 2.4737277 |
| CIT | 0.266108994 | 0.01596032 | -1.9099108 | 1.79695846 |
| DHX35 | 0.265473002 | 0.00295909 | -1.9133629 | 2.52884211 |
| CHAF1A | 0.265420206 | 0.00159631 | -1.9136499 | 2.79688263 |
| TCF4 | 0.265037197 | 0.01088876 | -1.9157332 | 1.96302172 |
| COPS3 | 0.264929738 | 0.00172502 | -1.9163183 | 2.76320629 |
| SACS | 0.264503752 | 0.0001002 | -1.9186399 | 3.99912474 |
| KDM6A | 0.264440879 | 0.00454974 | -1.9189829 | 2.34201296 |
| MED22 | 0.264362467 | 0.00113638 | -1.9194107 | 2.94447658 |
| SMC6 | 0.264046165 | 0.00060612 | -1.9211379 | 3.21744155 |
| EIF3G | 0.264041495 | 7.0487E-05 | -1.9211634 | 4.1518917 |
| TCEA3 | 0.26386641 | 0.00498088 | -1.9221204 | 2.30269434 |
| ZNF782 | 0.263703746 | 0.14490931 | -1.92301 | 0.83890371 |
| TTC9C | 0.26358789 | 0.23381928 | -1.923644 | 0.63111968 |
| SUPT5H | 0.263408844 | 0.0004373 | -1.9246243 | 3.35922046 |
| AMPD2 | 0.263292001 | 0.00050053 | -1.9252644 | 3.30057296 |
| DNMT1 | 0.262633841 | 0.00055716 | -1.9288753 | 3.25402126 |
| ZNF587 | 0.262511464 | 0.11465541 | -1.9295477 | 0.94060543 |
| RNF168 | 0.262457279 | 0.00737382 | -1.9298455 | 2.13230728 |
| ZKSCAN4 | 0.262322345 | 0.00731755 | -1.9305874 | 2.13563439 |
| KIF21A | 0.262317066 | 7.0734E-05 | -1.9306164 | 4.1503728 |

| | | | | |
|---|---|---|---|---|
| PLEKHG3 | 0.261884985 | 0.0012532 | -1.9329947 | 2.90197808 |
| MLLT1 | 0.261884737 | 0.05779578 | -1.9329961 | 1.23810389 |
| SFSWAP | 0.261705384 | 0.00086649 | -1.9339845 | 3.0622384 |
| ERF | 0.261588848 | 0.0003575 | -1.9346271 | 3.44671805 |
| CAMSAP2 | 0.261454594 | 0.00322688 | -1.9353677 | 2.49121735 |
| CHD9 | 0.261204003 | 0.00063553 | -1.9367511 | 3.19686327 |
| TMEM168 | 0.261125791 | 0.30928218 | -1.9371831 | 0.5096451 |
| UBLCP1 | 0.261106045 | 0.02372176 | -1.9372922 | 1.6248531 |
| TGS1 | 0.260929383 | 0.05336859 | -1.9382687 | 1.27271423 |
| BAZ2A | 0.260817277 | 0.00064724 | -1.9388887 | 3.18893577 |
| ERCC3 | 0.26074967 | 0.00023406 | -1.9392627 | 3.63067165 |
| PHKA1 | 0.260355576 | 0.00874237 | -1.9414448 | 2.05837067 |
| CYP20A1 | 0.259340449 | 0.34866692 | -1.9470809 | 0.45758925 |
| DGKH | 0.258933193 | 0.01767188 | -1.9493482 | 1.75271725 |
| UHRF2 | 0.258745654 | 0.06428844 | -1.9503935 | 1.1918671 |
| DHX29 | 0.258445699 | 0.00262535 | -1.9520669 | 2.580812 |
| S100PBP | 0.258202621 | 0.00505608 | -1.9534244 | 2.29618566 |
| TCAF1 | 0.258119265 | 0.00508987 | -1.9538903 | 2.29329328 |
| WNK2 | 0.25676246 | 0.04001485 | -1.9614938 | 1.39777885 |
| ESCO1 | 0.255722343 | 0.05829873 | -1.9673499 | 1.23434089 |
| CASZ1 | 0.255644798 | 0.0017436 | -1.9677874 | 2.75855384 |
| HNRNPUL1 | 0.255452223 | 0.00310739 | -1.9688746 | 2.5076048 |
| RNF25 | 0.254891352 | 0.01284178 | -1.9720457 | 1.89137474 |
| NUMA1 | 0.254448624 | 8.0894E-06 | -1.9745537 | 5.09208232 |
| CAMK2G | 0.254014593 | 9.5349E-05 | -1.9770167 | 4.02068268 |
| RBM12B | 0.253547855 | 0.00052197 | -1.97967 | 3.28235034 |
| EXOC6 | 0.25285909 | 0.02865765 | -1.9835945 | 1.54275939 |
| BAZ1B | 0.252732452 | 0.00038719 | -1.9843172 | 3.41207577 |
| MCF2L | 0.251965477 | 0.22994494 | -1.988702 | 0.63837614 |
| ISY1 | 0.251531154 | 0.00062407 | -1.991191 | 3.20476759 |
| POLR3F | 0.251354084 | 0.0003851 | -1.992207 | 3.41443195 |
| TUBGCP2 | 0.251300122 | 0.00025477 | -1.9925167 | 3.59384898 |
| MAPKAP1 | 0.25044406 | 0.00064472 | -1.9974397 | 3.19062775 |
| YLPM1 | 0.250430199 | 0.00010752 | -1.9975195 | 3.96852816 |
| TIAM1 | 0.250224568 | 0.03660971 | -1.9987046 | 1.43640367 |
| HK2 | 0.250111524 | 0.01259161 | -1.9993566 | 1.89991862 |
| GABPA | 0.249691789 | 0.0216487 | -2.0017797 | 1.66456827 |
| INTS13 | 0.248877698 | 0.00027079 | -2.0064911 | 3.56736903 |
| SMARCA4 | 0.24790699 | 0.0001889 | -2.0121291 | 3.72375892 |
| CARS1 | 0.24759442 | 0.02652331 | -2.0139493 | 1.57637223 |
| SETD1B | 0.247542447 | 0.17792494 | -2.0142522 | 0.74976316 |
| EPPK1 | 0.247502171 | 0.00366144 | -2.0144869 | 2.43634839 |
| PRR36 | 0.24738376 | 0.0081304 | -2.0151773 | 2.08988798 |
| ZC3H13 | 0.247014462 | 3.1986E-05 | -2.0173326 | 4.49504302 |
| MED21 | 0.246927367 | 0.10331961 | -2.0178414 | 0.98581725 |

| | | | | |
|---|---|---|---|---|
| PGLS | 0.246891228 | 0.05808558 | -2.0180525 | 1.23593169 |
| CAPRIN1 | 0.246873643 | 0.00016879 | -2.0181553 | 3.77264736 |
| PARN | 0.246837322 | 0.01461301 | -2.0183675 | 1.83526045 |
| ELP3 | 0.246426079 | 0.00736801 | -2.0207731 | 2.13264983 |
| GSE1 | 0.246383187 | 0.00047007 | -2.0210243 | 3.32783707 |
| POLR2A | 0.246212706 | 1.767E-05 | -2.0220229 | 4.75276918 |
| SSB | 0.246151539 | 0.01006506 | -2.0223813 | 1.99718363 |
| CTDP1 | 0.246106095 | 3.4727E-07 | -2.0226477 | 6.45933066 |
| PRRC1 | 0.245715152 | 0.00053718 | -2.0249413 | 3.26987628 |
| FANCA | 0.24556388 | 0.1325805 | -2.0258297 | 0.87752033 |
| CENPK | 0.245153027 | 0.01091317 | -2.0282455 | 1.96204892 |
| HDAC1 | 0.245070169 | 0.00030385 | -2.0287332 | 3.51734483 |
| NAA30 | 0.244820309 | 0.15959139 | -2.0302049 | 0.79699053 |
| MFF | 0.244696547 | 2.4725E-05 | -2.0309344 | 4.60685673 |
| EEF1B2 | 0.244517248 | 0.00221009 | -2.0319919 | 2.65558963 |
| SCAF1 | 0.244105557 | 0.03801074 | -2.034423 | 1.42009365 |
| EFL1 | 0.243908574 | 0.32377499 | -2.0355876 | 0.4897567 |
| THOC2 | 0.243571371 | 0.00029901 | -2.0375835 | 3.52430839 |
| ANKRD52 | 0.243282119 | 0.102871 | -2.0392978 | 0.98770704 |
| ZNF292 | 0.24271524 | 0.00147109 | -2.0426634 | 2.83236139 |
| SF3B6 | 0.24166131 | 0.00212111 | -2.0489416 | 2.67343625 |
| GPATCH1 | 0.241408861 | 0.03655748 | -2.0504495 | 1.43702373 |
| EIF3B | 0.240355622 | 0.00028731 | -2.0567575 | 3.54165171 |
| DENND3 | 0.239868705 | 0.00410356 | -2.0596831 | 2.38683907 |
| PNN | 0.239647269 | 6.3565E-05 | -2.0610156 | 4.19677871 |
| NHP2 | 0.238487293 | 0.00202758 | -2.0680157 | 2.69302173 |
| ZKSCAN1 | 0.238453538 | 0.00249744 | -2.0682199 | 2.60250481 |
| CDKN2AIP | 0.238132008 | 0.00123319 | -2.0701665 | 2.90897137 |
| STRN | 0.237920256 | 0.00010834 | -2.07145 | 3.96521726 |
| KPNA1 | 0.237499496 | 0.03359197 | -2.0740036 | 1.47376459 |
| GTF3A | 0.237356548 | 0.00616485 | -2.0748722 | 2.21007736 |
| UPF3B | 0.237059041 | 0.00064459 | -2.0766817 | 3.19071794 |
| RBM7 | 0.236576118 | 0.01254919 | -2.0796237 | 1.90138421 |
| ZGRF1 | 0.235882088 | 0.09192985 | -2.0838622 | 1.03654345 |
| PAPOLA | 0.23577117 | 0.00033861 | -2.0845408 | 3.47029933 |
| RETREG1 | 0.23563438 | 0.13820786 | -2.085378 | 0.85946725 |
| CDC45 | 0.235353932 | 0.01717709 | -2.0870961 | 1.76505029 |
| THUMPD2 | 0.2350922 | 0.11380238 | -2.0887014 | 0.94384866 |
| RPAP1 | 0.235050303 | 0.00164751 | -2.0889586 | 2.78317266 |
| HNRNPM | 0.234961372 | 8.8209E-05 | -2.0895045 | 4.05448911 |
| NVL | 0.234891389 | 8.6669E-05 | -2.0899343 | 4.06213721 |
| MAP3K20 | 0.233967357 | 0.02157984 | -2.0956208 | 1.66595169 |
| GIT2 | 0.233837078 | 0.00118931 | -2.0964244 | 2.92470511 |
| POGZ | 0.233826482 | 0.00089961 | -2.0964898 | 3.0459449 |
| MTMR1 | 0.233555636 | 0.00113002 | -2.0981618 | 2.94691323 |

| | | | | |
|---|---|---|---|---|
| DROSHA | 0.233359041 | 0.01081665 | -2.0993767 | 1.96590711 |
| CEP76 | 0.233086274 | 0.12352378 | -2.101064 | 0.90824943 |
| RAVER2 | 0.233055173 | 0.00133475 | -2.1012566 | 2.87459937 |
| BRMS1L | 0.232968661 | 0.05543239 | -2.1017922 | 1.2562364 |
| RNMT | 0.232967795 | 0.0016261 | -2.1017976 | 2.78885365 |
| CCAR2 | 0.23294927 | 1.4303E-05 | -2.1019123 | 4.84455828 |
| CAAP1 | 0.232834725 | 0.00034698 | -2.1026219 | 3.45969622 |
| HOXA10 | 0.231969999 | 0.01260974 | -2.1079899 | 1.89929372 |
| HCFC1 | 0.231966204 | 0.00010446 | -2.1080135 | 3.98103063 |
| USP5 | 0.231719207 | 0.00017284 | -2.1095505 | 3.76234888 |
| CKAP5 | 0.231059757 | 0.00046439 | -2.1136621 | 3.33311368 |
| USP34 | 0.230844679 | 0.00026844 | -2.1150056 | 3.57115824 |
| POLR1A | 0.229926174 | 0.03652524 | -2.1207574 | 1.43740688 |
| RREB1 | 0.229897375 | 0.02284509 | -2.1209381 | 1.64120722 |
| RLIG1 | 0.229779273 | 0.10454938 | -2.1216794 | 0.98067854 |
| DHX38 | 0.228963082 | 0.00163239 | -2.1268131 | 2.78717714 |
| ZNF121 | 0.2289306 | 0.11244967 | -2.1270178 | 0.9490418 |
| ZNF12 | 0.228760155 | 0.00134157 | -2.1280923 | 2.87238666 |
| CCDC9 | 0.228685833 | 0.00182418 | -2.1285611 | 2.73893271 |
| MED28 | 0.228072862 | 0.00075289 | -2.1324333 | 3.12327004 |
| MNAT1 | 0.227863521 | 0.00126066 | -2.1337581 | 2.89940097 |
| C1orf131 | 0.227812978 | 0.00284105 | -2.1340782 | 2.54652124 |
| SPATA5 | 0.227223291 | 0.01119287 | -2.1378174 | 1.95105863 |
| RSBN1L | 0.226627586 | 0.00099514 | -2.1416046 | 3.00211396 |
| PRUNE1 | 0.225985365 | 0.15720697 | -2.1456987 | 0.80352821 |
| CDC42EP4 | 0.225768977 | 0.13264796 | -2.1470808 | 0.87729944 |
| TCF12 | 0.225689751 | 0.00387411 | -2.1475872 | 2.41182856 |
| HLTF | 0.225670662 | 0.01561451 | -2.1477092 | 1.80647161 |
| TLE1 | 0.225618134 | 0.00299369 | -2.1480451 | 2.52379323 |
| CHAMP1 | 0.225283004 | 0.00037631 | -2.1501896 | 3.42445655 |
| EMSY | 0.225260959 | 0.00016427 | -2.1503308 | 3.78443826 |
| PRKAB1 | 0.224659091 | 0.03935329 | -2.1541907 | 1.40501894 |
| NOL11 | 0.224371947 | 0.00354507 | -2.1560358 | 2.45037521 |
| GEMIN4 | 0.22426129 | 0.00029566 | -2.1567475 | 3.52921374 |
| MTBP | 0.224175108 | 0.01357856 | -2.157302 | 1.86714623 |
| POLDIP3 | 0.223653397 | 0.0001023 | -2.1606634 | 3.99014169 |
| C9orf78 | 0.223486817 | 0.00131598 | -2.1617384 | 2.88074928 |
| SAFB2 | 0.223461979 | 0.00062141 | -2.1618987 | 3.20662103 |
| SCAF8 | 0.223269054 | 0.00321531 | -2.1631448 | 2.49277762 |
| IRF2BP2 | 0.223264771 | 0.00157501 | -2.1631725 | 2.80271672 |
| CDC73 | 0.223239969 | 0.00035231 | -2.1633327 | 3.45307184 |
| DPP3 | 0.222740442 | 0.01729245 | -2.1665646 | 1.76214344 |
| CPSF6 | 0.222179369 | 9.0718E-05 | -2.1702032 | 4.04230725 |
| ANAPC7 | 0.222036629 | 0.00702547 | -2.1711304 | 2.15332463 |
| RCL1 | 0.221155157 | 0.08862592 | -2.1768692 | 1.05243927 |

| | | | | |
|---|---|---|---|---|
| MED23 | 0.221110298 | 0.14507086 | -2.1771619 | 0.83841981 |
| DACH1 | 0.220968305 | 0.0010577 | -2.1780886 | 2.97563647 |
| RRBP1 | 0.220565347 | 0.03166877 | -2.1807219 | 1.49936886 |
| RAF1 | 0.219664832 | 0.06054957 | -2.1866242 | 1.21788897 |
| SAMD1 | 0.219175704 | 8.6823E-05 | -2.1898402 | 4.06136518 |
| NOP9 | 0.219154445 | 0.00525959 | -2.1899802 | 2.27904819 |
| ILKAP | 0.218865463 | 0.00288024 | -2.1918838 | 2.54057193 |
| NOSIP | 0.218676334 | 4.281E-05 | -2.193131 | 4.36845424 |
| MYH14 | 0.218164142 | 0.00031062 | -2.1965141 | 3.50777182 |
| EIF3I | 0.218082374 | 0.00148639 | -2.1970549 | 2.82786588 |
| STK11IP | 0.217902768 | 0.22367033 | -2.1982436 | 0.65039161 |
| GLE1 | 0.217580691 | 0.02805382 | -2.2003776 | 1.55200802 |
| ELP2 | 0.217485117 | 0.05547934 | -2.2010114 | 1.25586868 |
| TAF9B | 0.216815592 | 8.3209E-06 | -2.2054596 | 5.07983175 |
| PCCB | 0.21669721 | 0.00036847 | -2.2062475 | 3.43359235 |
| LIN9 | 0.21637628 | 0.04237874 | -2.2083857 | 1.37285199 |
| ZNF280C | 0.216288948 | 0.07686305 | -2.2089681 | 1.11428239 |
| NUB1 | 0.215825928 | 0.1715001 | -2.2120599 | 0.76573562 |
| SUGP1 | 0.215464566 | 0.00841395 | -2.2144775 | 2.07499997 |
| PPP1R12A | 0.214103511 | 2.5988E-05 | -2.2236196 | 4.58521976 |
| NELFA | 0.21294818 | 0.00117407 | -2.2314257 | 2.93030671 |
| ZNF829 | 0.212899371 | 0.02817645 | -2.2317564 | 1.55011368 |
| UBA5 | 0.212692782 | 0.00072386 | -2.233157 | 3.14034564 |
| ZNF362 | 0.21245859 | 0.01402335 | -2.2347464 | 1.85314811 |
| KDM3B | 0.212162874 | 0.00029802 | -2.2367559 | 3.52574994 |
| ENAH | 0.212112759 | 6.1781E-05 | -2.2370967 | 4.20914408 |
| CAP1 | 0.211510058 | 0.00048637 | -2.2412018 | 3.31303179 |
| TSNAX | 0.21146419 | 0.00037864 | -2.2415147 | 3.42177304 |
| ZSCAN25 | 0.211076568 | 0.15691686 | -2.2441617 | 0.80433038 |
| MAPK7 | 0.21086224 | 0.00664781 | -2.2456273 | 2.17732133 |
| TCOF1 | 0.210697931 | 4.9002E-05 | -2.2467519 | 4.30978549 |
| MYT1 | 0.210250955 | 9.4797E-05 | -2.2498157 | 4.02320482 |
| LRRFIP1 | 0.209991062 | 5.0783E-05 | -2.2516002 | 4.29428137 |
| PRDM2 | 0.209422073 | 0.00160242 | -2.2555146 | 2.79522407 |
| C19orf47 | 0.209384412 | 0.0014446 | -2.255774 | 2.84025267 |
| TPX2 | 0.209353571 | 5.5013E-05 | -2.2559866 | 4.25953542 |
| ESRRA | 0.209165023 | 0.16966907 | -2.2572865 | 0.77039732 |
| ING3 | 0.20890251 | 0.00271696 | -2.2590983 | 2.56591742 |
| RRAS2 | 0.208751536 | 0.0944666 | -2.2601413 | 1.02472169 |
| ZC3H14 | 0.208475557 | 2.072E-06 | -2.2620499 | 5.68361491 |
| BCL2L13 | 0.208317763 | 0.06485283 | -2.2631422 | 1.18807104 |
| MORF4L2 | 0.207519247 | 0.20531773 | -2.2686829 | 0.68757354 |
| ZFP2 | 0.207084533 | 8.5015E-06 | -2.2717083 | 5.07050241 |
| PHAX | 0.207084274 | 0.00852534 | -2.2717101 | 2.0692885 |
| ZNFX1 | 0.206918634 | 0.00252684 | -2.2728645 | 2.5974224 |

| | | | | |
|---|---|---|---|---|
| BAG6 | 0.206630352 | 0.0101798 | -2.2748759 | 1.99226081 |
| UPF2 | 0.206530046 | 0.01147078 | -2.2755764 | 1.94040714 |
| DHX16 | 0.206405564 | 0.00126219 | -2.2764462 | 2.89887465 |
| KANSL3 | 0.206399211 | 0.00151926 | -2.2764906 | 2.81836675 |
| FAM50B | 0.206338285 | 0.09138304 | -2.2769166 | 1.03913438 |
| HDGF | 0.205820025 | 9.6598E-05 | -2.2805447 | 4.0150315 |
| URI1 | 0.20565823 | 0.00114322 | -2.2816793 | 2.94187038 |
| MAML1 | 0.205513942 | 0.04652009 | -2.2826918 | 1.33235943 |
| RAVER1 | 0.205109745 | 6.6954E-05 | -2.2855321 | 4.17422047 |
| CDT1 | 0.205094529 | 0.10548971 | -2.2856391 | 0.97678989 |
| G3BP1 | 0.20501361 | 1.1722E-05 | -2.2862084 | 4.93099977 |
| SCAF11 | 0.204280858 | 0.00017325 | -2.2913741 | 3.76133331 |
| SART3 | 0.203788403 | 1.2966E-05 | -2.2948561 | 4.88719617 |
| ZNF536 | 0.203320044 | 0.03302623 | -2.2981756 | 1.48114106 |
| MPHOSPH6 | 0.203126006 | 0.03263406 | -2.2995531 | 1.48632894 |
| EEF1D | 0.202695526 | 0.00027303 | -2.3026138 | 3.56379103 |
| FAM50A | 0.202613911 | 0.00200939 | -2.3031949 | 2.69693503 |
| USP9X | 0.202540218 | 0.00548212 | -2.3037197 | 2.26105127 |
| PANX1 | 0.202425122 | 0.07587681 | -2.3045397 | 1.11989094 |
| RSRC2 | 0.202397081 | 0.0003565 | -2.3047396 | 3.44794341 |
| ZC3H18 | 0.202191682 | 0.00080242 | -2.3062044 | 3.09560093 |
| SVIL | 0.202048492 | 0.08770815 | -2.3072265 | 1.05696004 |
| KRI1 | 0.201121447 | 0.20798755 | -2.3138612 | 0.68196266 |
| HDGFL2 | 0.200950381 | 0.02814468 | -2.3150888 | 1.55060364 |
| BICRA | 0.200807293 | 0.16471464 | -2.3161164 | 0.78326781 |
| SETDB1 | 0.200147533 | 0.00109466 | -2.3208643 | 2.96072022 |
| CDCA8 | 0.1999154 | 0.00451472 | -2.3225385 | 2.34536886 |
| MYCL | 0.199910741 | 0.11243541 | -2.3225721 | 0.94909691 |
| NSMCE4A | 0.199586596 | 0.00091216 | -2.3249133 | 3.03993008 |
| TMCC1 | 0.199169705 | 0.0001629 | -2.3279299 | 3.7880673 |
| ZW10 | 0.198681146 | 0.0005042 | -2.3314731 | 3.29740073 |
| TAF4 | 0.198640805 | 0.0005135 | -2.3317661 | 3.28945974 |
| RBM33 | 0.19860791 | 0.00056937 | -2.332005 | 3.24460437 |
| PINX1 | 0.198462346 | 0.00104926 | -2.3330628 | 2.97911541 |
| ELF2 | 0.198385324 | 0.03893188 | -2.3336228 | 1.40969466 |
| AKAP11 | 0.198324284 | 0.00025015 | -2.3340668 | 3.60180265 |
| POLE2 | 0.198308769 | 0.15480714 | -2.3341796 | 0.81020902 |
| MCM3AP | 0.197778587 | 0.00028723 | -2.3380419 | 3.5417746 |
| OBI1 | 0.197442612 | 0.00019262 | -2.3404947 | 3.71529876 |
| KCTD12 | 0.197401045 | 0.12373925 | -2.3407985 | 0.90749253 |
| NCOA2 | 0.197116921 | 0.006428 | -2.3428765 | 2.19192384 |
| RNF2 | 0.196557905 | 0.00052188 | -2.3469737 | 3.2824257 |
| SNRNP200 | 0.196478987 | 0.0004863 | -2.3475531 | 3.31309569 |
| STRBP | 0.195528395 | 0.00369955 | -2.35455 | 2.43185053 |
| PHF8 | 0.195123894 | 0.01303979 | -2.3575376 | 1.88472949 |

| | | | | |
|---|---|---|---|---|
| CEP131 | 0.193863324 | 0.0013374 | -2.3668882 | 2.87373737 |
| ZFP91 | 0.192230017 | 0.00012958 | -2.3790945 | 3.88744616 |
| PRPF3 | 0.191409399 | 3.2977E-06 | -2.3852664 | 5.48178836 |
| SPECC1L | 0.191083932 | 0.0011981 | -2.3877216 | 2.9215073 |
| MYO9B | 0.190895129 | 5.2725E-06 | -2.3891478 | 5.27798335 |
| NCAPD3 | 0.190635614 | 0.00101937 | -2.3911104 | 2.99166685 |
| SYMPK | 0.190500969 | 2.389E-06 | -2.3921298 | 5.62177884 |
| RTN1 | 0.190448389 | 0.06268196 | -2.392528 | 1.20285746 |
| ZC3H11A | 0.190227201 | 4.9426E-05 | -2.3942045 | 4.30604342 |
| ZNF106 | 0.189043431 | 0.00410612 | -2.4032104 | 2.38656835 |
| UBR5 | 0.188689195 | 0.0305038 | -2.4059163 | 1.515646 |
| JMJD1C | 0.1878251 | 0.00249134 | -2.4125382 | 2.60356776 |
| ARID1B | 0.18781962 | 2.7634E-05 | -2.4125803 | 4.55855098 |
| ICE2 | 0.187038122 | 0.00589418 | -2.4185957 | 2.22957684 |
| DIAPH1 | 0.186995513 | 1.7646E-05 | -2.4189244 | 4.75335213 |
| MED13L | 0.186634477 | 4.2114E-05 | -2.4217126 | 4.37557166 |
| TXLNG | 0.186490928 | 0.00392427 | -2.4228226 | 2.40624069 |
| EML4 | 0.186388313 | 5.4499E-05 | -2.4236167 | 4.2636133 |
| SMARCC1 | 0.186098154 | 6.1518E-05 | -2.4258644 | 4.21099496 |
| ZC3H4 | 0.186054435 | 1.2971E-06 | -2.4262033 | 5.88704323 |
| LMAN1 | 0.185986912 | 0.00096678 | -2.426727 | 3.01467312 |
| DDX10 | 0.185865698 | 0.00058996 | -2.4276676 | 3.22918076 |
| ZFP1 | 0.185722652 | 0.01561684 | -2.4287783 | 1.80640697 |
| SRSF10 | 0.185314742 | 0.0105489 | -2.4319504 | 1.97679287 |
| KCNH6 | 0.185127978 | 0.00041435 | -2.4334051 | 3.38263333 |
| GEMIN5 | 0.18440236 | 0.00366724 | -2.439071 | 2.43566014 |
| SKA3 | 0.184390723 | 0.00013791 | -2.439162 | 3.86041974 |
| TCERG1 | 0.184226752 | 0.00010985 | -2.4404455 | 3.95919345 |
| SNRPD3 | 0.184057153 | 0.0001065 | -2.4417743 | 3.97265673 |
| ZSCAN18 | 0.183836334 | 0.07819544 | -2.4435062 | 1.10681857 |
| BRCA2 | 0.183249558 | 0.10977485 | -2.4481184 | 0.95949714 |
| PARG | 0.183154149 | 0.10301667 | -2.4488697 | 0.9870925 |
| ZZZ3 | 0.183040262 | 0.00038049 | -2.4497671 | 3.4196601 |
| ZNF93 | 0.182870146 | 0.16564251 | -2.4511085 | 0.78082821 |
| BRD4 | 0.182802969 | 0.00082273 | -2.4516386 | 3.08474147 |
| PAXIP1 | 0.182436222 | 1.6157E-05 | -2.4545359 | 4.79163253 |
| TUBGCP5 | 0.181011123 | 0.00644135 | -2.4658497 | 2.19102287 |
| CMTR1 | 0.180992627 | 0.00029396 | -2.4659972 | 3.53170611 |
| MTA2 | 0.180542088 | 0.00194976 | -2.4695929 | 2.71001788 |
| TRMT1L | 0.180481817 | 0.01103118 | -2.4700746 | 1.95737816 |
| CENPQ | 0.178961737 | 0.12256662 | -2.4822769 | 0.91162778 |
| RTF2 | 0.178365139 | 0.00606748 | -2.4870944 | 2.21699181 |
| IRF2BP1 | 0.177904338 | 0.00044409 | -2.4908264 | 3.3525261 |
| GNE | 0.177468122 | 0.00013154 | -2.4943682 | 3.88094575 |
| FBXO38 | 0.177101859 | 0.01512552 | -2.4973487 | 1.82028958 |

| | | | | |
|---|---|---|---|---|
| IRF2BPL | 0.176930916 | 0.00011094 | -2.4987419 | 3.95492321 |
| TKT | 0.176580841 | 0.00328861 | -2.5015993 | 2.48298724 |
| BRCA1 | 0.176573513 | 0.00930039 | -2.5016591 | 2.03149868 |
| ZFP62 | 0.175976784 | 0.01021116 | -2.506543 | 1.99092483 |
| STRIP1 | 0.175900417 | 0.0040848 | -2.5071692 | 2.38882867 |
| SF3B1 | 0.175739641 | 2.3983E-05 | -2.5084884 | 4.6200909 |
| IFT74 | 0.175562443 | 0.00229705 | -2.5099438 | 2.63882902 |
| GRK5 | 0.175436771 | 0.00941537 | -2.5109769 | 2.02616244 |
| PYM1 | 0.175330913 | 0.00427255 | -2.5118477 | 2.36931251 |
| DCTN6 | 0.174835043 | 0.00103149 | -2.5159337 | 2.986537 |
| ING4 | 0.174637427 | 0.01333431 | -2.5175653 | 1.87502961 |
| LEO1 | 0.174637321 | 0.0002897 | -2.5175662 | 3.53804662 |
| OSBPL9 | 0.174578512 | 0.00516681 | -2.5180521 | 2.2867778 |
| GEMIN8 | 0.174216106 | 0.00024792 | -2.5210501 | 3.60568588 |
| RNF40 | 0.173802059 | 9.3943E-06 | -2.5244829 | 5.02713648 |
| RBM17 | 0.173360588 | 0.00422386 | -2.5281521 | 2.37429068 |
| MED1 | 0.172839571 | 0.00025812 | -2.5324945 | 3.58817873 |
| FAM184B | 0.172772017 | 1.3772E-06 | -2.5330585 | 5.86099161 |
| SETX | 0.172277922 | 0.000176 | -2.5371903 | 3.75448027 |
| DCAF1 | 0.172109971 | 0.00122444 | -2.5385974 | 2.9120623 |
| JADE1 | 0.171586687 | 0.0551614 | -2.5429905 | 1.25836472 |
| KDM4B | 0.17135973 | 0.00110891 | -2.5449 | 2.95510457 |
| SRRM2 | 0.171224082 | 2.5886E-05 | -2.5460425 | 4.58693546 |
| TADA1 | 0.171067403 | 0.00130937 | -2.5473632 | 2.88293895 |
| POLR3D | 0.171056665 | 0.00526579 | -2.5474538 | 2.27853686 |
| PRPF19 | 0.170836655 | 1.7707E-05 | -2.5493105 | 4.75185916 |
| ADCK5 | 0.170642904 | 6.7469E-05 | -2.5509477 | 4.17089348 |
| INTS2 | 0.170526107 | 0.06816601 | -2.5519355 | 1.16643212 |
| GLYR1 | 0.170403211 | 9.2325E-05 | -2.5529756 | 4.03468204 |
| KIF20B | 0.16967779 | 0.00065335 | -2.5591304 | 3.18485343 |
| DDX20 | 0.169625205 | 2.2808E-05 | -2.5595775 | 4.64191673 |
| TRIM28 | 0.16909459 | 0.00018489 | -2.5640976 | 3.73309711 |
| GTF3C1 | 0.168904001 | 0.00063405 | -2.5657246 | 3.19787377 |
| KATNIP | 0.168590595 | 0.14869928 | -2.568404 | 0.82769114 |
| RSRC1 | 0.16853985 | 0.00159774 | -2.5688383 | 2.79649316 |
| OSBPL6 | 0.168023574 | 0.0211754 | -2.5732644 | 1.67416838 |
| PHF14 | 0.1678466 | 0.00211461 | -2.5747848 | 2.67476903 |
| DPF2 | 0.167376191 | 0.00188677 | -2.5788338 | 2.72428126 |
| KDM1A | 0.167038611 | 0.00310069 | -2.5817465 | 2.50854161 |
| BUD13 | 0.166811589 | 0.00635177 | -2.5837086 | 2.19710513 |
| CDC5L | 0.166788598 | 0.00011732 | -2.5839074 | 3.93062955 |
| ZNF189 | 0.166665513 | 0.21192313 | -2.5849725 | 0.67382163 |
| SMN1; SMN2 | 0.16658074 | 0.00020539 | -2.5857065 | 3.68742629 |
| BLM | 0.16657141 | 0.00163502 | -2.5857873 | 2.78647787 |
| ZBTB1 | 0.165991512 | 0.16392297 | -2.5908186 | 0.7853602 |

| | | | | |
|---|---|---|---|---|
| ZNF71 | 0.165631703 | 0.04446249 | -2.5939493 | 1.35200619 |
| MAX | 0.165439298 | 0.00363099 | -2.5956261 | 2.43997449 |
| CBX8 | 0.164853918 | 5.0037E-06 | -2.6007399 | 5.30070696 |
| ZNF512B | 0.164514261 | 0.00200477 | -2.6037154 | 2.69793472 |
| CREB1 | 0.164396146 | 0.00064096 | -2.6047516 | 3.19316812 |
| UVSSA | 0.16430321 | 0.1053595 | -2.6055674 | 0.97732631 |
| GPKOW | 0.164188881 | 0.00034962 | -2.6065717 | 3.45640878 |
| SPRYD3 | 0.163719205 | 0.00185349 | -2.6107045 | 2.73200907 |
| YIPF5 | 0.163568005 | 0.07170449 | -2.6120375 | 1.14445364 |
| CRTC2 | 0.163234759 | 5.4516E-06 | -2.6149798 | 5.26347565 |
| GTF3C5 | 0.163172153 | 9.3116E-07 | -2.6155332 | 6.03097723 |
| LAS1L | 0.163038585 | 0.01096931 | -2.6167147 | 1.95982062 |
| BARD1 | 0.162744983 | 0.02294705 | -2.619315 | 1.63927309 |
| ARID2 | 0.162642406 | 1.202E-05 | -2.6202246 | 4.92009859 |
| GTF3C2 | 0.16239852 | 0.00082538 | -2.6223896 | 3.08334355 |
| WBP11 | 0.162368375 | 0.00029642 | -2.6226574 | 3.52809414 |
| SGO2 | 0.162292098 | 0.0176533 | -2.6233353 | 1.75317414 |
| CHN1 | 0.162262984 | 0.0001041 | -2.6235942 | 3.98254364 |
| CABIN1 | 0.162033844 | 0.00082122 | -2.6256329 | 3.08553817 |
| MPND | 0.16201918 | 0.00059347 | -2.6257635 | 3.22659793 |
| CFAP410 | 0.161853219 | 0.19795013 | -2.627242 | 0.7034442 |
| PACS2 | 0.161231523 | 0.00363123 | -2.6327943 | 2.43994587 |
| CIAO3 | 0.16102628 | 6.3676E-05 | -2.6346319 | 4.19602705 |
| MOCS2 | 0.161023819 | 0.00052124 | -2.634654 | 3.28296025 |
| SMARCB1 | 0.161016999 | 0.00234243 | -2.6347151 | 2.63033335 |
| TOP2B | 0.159701226 | 0.00431365 | -2.6465527 | 2.36515469 |
| ANAPC5 | 0.159675338 | 0.00077089 | -2.6467866 | 3.11300772 |
| STRN3 | 0.159562775 | 0.00089165 | -2.647804 | 3.04980684 |
| ZNF318 | 0.159457738 | 0.0062591 | -2.648754 | 2.2034883 |
| HSF1 | 0.158850722 | 0.06884087 | -2.6542565 | 1.16215366 |
| PPIL4 | 0.158004164 | 1.0424E-06 | -2.6619655 | 5.98196092 |
| NACC1 | 0.157356212 | 0.15238384 | -2.667894 | 0.8170611 |
| SUB1 | 0.157336409 | 0.00024872 | -2.6680755 | 3.60428573 |
| CLUH | 0.156920928 | 0.00017766 | -2.6718903 | 3.75040019 |
| UBXN7 | 0.156800374 | 0.00038573 | -2.6729991 | 3.4137189 |
| PLEKHM2 | 0.156254942 | 0.07527776 | -2.6780263 | 1.12333332 |
| DONSON | 0.156137448 | 0.00355974 | -2.6791115 | 2.44858216 |
| FADS1 | 0.155967143 | 0.0012822 | -2.680686 | 2.89204329 |
| EZR | 0.154633788 | 8.8546E-05 | -2.6930725 | 4.05282941 |
| ARHGEF17 | 0.15437165 | 0.02930201 | -2.6955203 | 1.53310258 |
| CDC40 | 0.15428529 | 1.422E-05 | -2.6963276 | 4.84710612 |
| KIN | 0.154062201 | 0.13191478 | -2.6984152 | 0.87970654 |
| BCAP31 | 0.153144157 | 2.0316E-05 | -2.7070378 | 4.69215917 |
| GTF2I | 0.152662284 | 1.8842E-05 | -2.7115844 | 4.72486212 |
| HEBP1 | 0.152556796 | 0.00063688 | -2.7125816 | 3.19594434 |

| | | | | |
|---|---|---|---|---|
| SF3B4 | 0.152422351 | 0.00771517 | -2.7138536 | 2.11265428 |
| TRIR | 0.152053632 | 3.3976E-05 | -2.7173478 | 4.46882497 |
| ZNF532 | 0.151924967 | 0.00068895 | -2.7185691 | 3.16181097 |
| PELO | 0.150621525 | 0.00742377 | -2.7310001 | 2.12937572 |
| PEX1 | 0.150100982 | 0.00339083 | -2.7359947 | 2.46969364 |
| PDCD4 | 0.149778536 | 0.00014392 | -2.7390972 | 3.8418829 |
| SENP6 | 0.149005245 | 0.00057388 | -2.746565 | 3.24118244 |
| RNF20 | 0.148968217 | 0.00012834 | -2.7469235 | 3.89164092 |
| DCTN2 | 0.148945473 | 5.068E-05 | -2.7471438 | 4.2951638 |
| MPHOSPH10 | 0.148706488 | 0.00133926 | -2.7494605 | 2.87313631 |
| L3MBTL3 | 0.148200273 | 0.091239 | -2.75438 | 1.03981949 |
| KDM4A | 0.148094232 | 0.00340665 | -2.7554126 | 2.46767247 |
| PDZD8 | 0.147981342 | 0.00267161 | -2.7565128 | 2.57322726 |
| UBTF | 0.14785658 | 2.19E-05 | -2.7577296 | 4.65954667 |
| RBM25 | 0.147689855 | 0.00012822 | -2.7593574 | 3.89203742 |
| SZRD1 | 0.147654047 | 0.11584535 | -2.7597072 | 0.93612138 |
| PHF21A | 0.14756751 | 9.2082E-07 | -2.760553 | 6.03582578 |
| ERCC5 | 0.146508 | 0.00056511 | -2.7709486 | 3.24786744 |
| PDCD7 | 0.14645582 | 0.00165718 | -2.7714626 | 2.78063107 |
| EVI5L | 0.146430348 | 0.21010198 | -2.7717135 | 0.67756985 |
| TRRAP | 0.146353385 | 0.00171723 | -2.772472 | 2.76517234 |
| TAF6 | 0.146263223 | 0.00056403 | -2.773361 | 3.24869819 |
| NCAPD2 | 0.146151372 | 0.00018182 | -2.7744647 | 3.74034901 |
| THRAP3 | 0.146119608 | 0.00011171 | -2.7747783 | 3.95192125 |
| DCTN4 | 0.14606856 | 0.00895573 | -2.7752824 | 2.04789906 |
| KDM5A | 0.145007584 | 0.00061336 | -2.7857997 | 3.2122863 |
| CCAR1 | 0.144114367 | 2.5292E-05 | -2.7947139 | 4.59702469 |
| HIRA | 0.143998124 | 0.00035938 | -2.7958781 | 3.44445043 |
| KTN1 | 0.143770035 | 2.4571E-06 | -2.7981651 | 5.60957876 |
| BCL11B | 0.143649129 | 8.7666E-06 | -2.7993789 | 5.05716856 |
| BTF3L4 | 0.143337293 | 0.05147595 | -2.8025141 | 1.28839563 |
| DACH2 | 0.143305677 | 0.077162 | -2.8028323 | 1.11259651 |
| DGKZ | 0.143049709 | 0.00265294 | -2.8054115 | 2.57627335 |
| VPS39 | 0.142995333 | 0.03357599 | -2.80596 | 1.47397112 |
| CDC27 | 0.142776455 | 0.00183315 | -2.80817 | 2.7368019 |
| CEBPZ | 0.142750171 | 6.1781E-05 | -2.8084356 | 4.20914236 |
| BORCS6 | 0.142652108 | 0.02127016 | -2.809427 | 1.67222921 |
| GMNN | 0.141959359 | 0.02594057 | -2.8164501 | 1.58602044 |
| TSC22D4 | 0.141319365 | 0.00467994 | -2.8229689 | 2.32976011 |
| BRD3 | 0.141299065 | 0.10336203 | -2.8231762 | 0.98563897 |
| TBC1D22B | 0.141199732 | 0.02636332 | -2.8241907 | 1.57899983 |
| NAB1 | 0.141020754 | 8.6898E-05 | -2.8260206 | 4.06099105 |
| RIF1 | 0.141013458 | 2.2107E-05 | -2.8260952 | 4.65546336 |
| ZNF638 | 0.140725771 | 2.3531E-05 | -2.8290415 | 4.62836766 |
| STRN4 | 0.139466431 | 0.00110943 | -2.8420102 | 2.95489832 |

| | | | | |
|---|---|---|---|---|
| STAG2 | 0.139090596 | 0.00023606 | -2.8459032 | 3.62697713 |
| PRPF8 | 0.138575075 | 1.051E-06 | -2.8512603 | 5.97839879 |
| COMMD2 | 0.138573991 | 0.16571497 | -2.8512716 | 0.78063827 |
| ARMC6 | 0.138412842 | 0.24242095 | -2.8529503 | 0.61542984 |
| PUF60 | 0.138217742 | 0.00243742 | -2.8549853 | 2.61307007 |
| TFIP11 | 0.138057529 | 0.00723836 | -2.8566585 | 2.14036 |
| SYBU | 0.137993329 | 0.13172881 | -2.8573296 | 0.88031924 |
| PPIE | 0.137952486 | 0.00537978 | -2.8577566 | 2.26923582 |
| RNF130 | 0.137857027 | 0.07747985 | -2.8587553 | 1.11081121 |
| G3BP2 | 0.1375181 | 0.00146403 | -2.8623066 | 2.83444898 |
| CLASP2 | 0.13741527 | 0.09069808 | -2.8633858 | 1.04240189 |
| EIF3A | 0.137048402 | 3.9831E-06 | -2.8672426 | 5.39978083 |
| SEPHS1 | 0.136117919 | 0.00534134 | -2.8770711 | 2.27234997 |
| RBBP5 | 0.135899689 | 3.7384E-05 | -2.8793859 | 4.42730869 |
| NCOA1 | 0.134882028 | 0.00195612 | -2.89023 | 2.70860444 |
| ZMYM4 | 0.134524279 | 0.00046755 | -2.8940615 | 3.33017592 |
| SMARCC2 | 0.134450175 | 9.7607E-06 | -2.8948565 | 5.01052081 |
| COIL | 0.134378637 | 0.10119922 | -2.8956243 | 0.99482282 |
| TRMT6 | 0.134075113 | 0.00025705 | -2.8988866 | 3.58997482 |
| RPRD2 | 0.134000444 | 3.4874E-05 | -2.8996903 | 4.45749667 |
| CHAC2 | 0.133527263 | 0.10756295 | -2.9047938 | 0.96833731 |
| OGFOD1 | 0.133483067 | 0.0247429 | -2.9052714 | 1.60654935 |
| POLR1B | 0.133180772 | 0.0381269 | -2.9085423 | 1.4187685 |
| CSTF2T | 0.132982751 | 5.7954E-05 | -2.910689 | 4.2369167 |
| CCM2 | 0.131985655 | 0.00107153 | -2.921547 | 2.96999739 |
| EXO1 | 0.13173866 | 0.00101175 | -2.9242493 | 2.99492765 |
| NASP | 0.131584783 | 4.3994E-05 | -2.9259354 | 4.35660871 |
| EXOSC10 | 0.131561217 | 0.00051005 | -2.9261938 | 3.29238787 |
| ZMYND8 | 0.13155364 | 4.8621E-05 | -2.9262769 | 4.31317377 |
| PHF10 | 0.131525525 | 0.11634461 | -2.9265853 | 0.93425371 |
| PMS2 | 0.131506181 | 0.00028864 | -2.9267975 | 3.53964619 |
| CWC22 | 0.131270081 | 0.00274399 | -2.92939 | 2.56161674 |
| MCM8 | 0.131021412 | 4.8687E-05 | -2.9321255 | 4.31258921 |
| TP53BP1 | 0.129992669 | 5.2111E-06 | -2.9434978 | 5.28307304 |
| GATAD2B | 0.129943518 | 7.6968E-06 | -2.9440434 | 5.11369119 |
| HIRIP3 | 0.129493431 | 0.00728278 | -2.9490492 | 2.13770302 |
| WDR70 | 0.1294574 | 3.0386E-06 | -2.9494507 | 5.51732502 |
| TRIM32 | 0.129417137 | 0.21128262 | -2.9498994 | 0.67513623 |
| KAT8 | 0.129160584 | 0.00013102 | -2.9527622 | 3.88267697 |
| ZFC3H1 | 0.129077539 | 0.00710926 | -2.9536901 | 2.14817557 |
| SCFD2 | 0.12890954 | 2.72E-05 | -2.9555691 | 4.56543686 |
| ZNF451 | 0.128404961 | 4.7134E-06 | -2.9612272 | 5.32666347 |
| SLX4 | 0.128395134 | 0.00086059 | -2.9613376 | 3.06520536 |
| GEMIN6 | 0.127657605 | 0.00244382 | -2.9696486 | 2.61193034 |
| YY1AP1 | 0.126824283 | 0.06105301 | -2.9790971 | 1.21429292 |

| | | | | |
|---|---|---|---|---|
| CUL2 | 0.126674176 | 0.14614956 | -2.9808056 | 0.83520249 |
| CMPK1 | 0.126658605 | 8.1512E-05 | -2.980983 | 4.08877682 |
| UTP14A | 0.126279148 | 0.00060959 | -2.9853117 | 3.21496279 |
| SF3A3 | 0.125788252 | 0.00025089 | -2.9909309 | 3.60051788 |
| TAF6L | 0.125665416 | 0.00048166 | -2.9923404 | 3.31726259 |
| TPR | 0.125526497 | 3.0421E-06 | -2.9939362 | 5.51682075 |
| BYSL | 0.124796334 | 0.0050488 | -3.0023525 | 2.29681188 |
| XRCC6 | 0.124539993 | 0.00013229 | -3.005319 | 3.87845766 |
| JUN | 0.124527429 | 0.05741744 | -3.0054645 | 1.24095615 |
| EHMT2 | 0.123979901 | 0.00063405 | -3.0118218 | 3.19787622 |
| SPEN | 0.123632862 | 0.00139069 | -3.0158658 | 2.85676999 |
| SCAPER | 0.123477965 | 8.17E-05 | -3.0176745 | 4.08777983 |
| BCLAF1 | 0.123035512 | 8.5739E-05 | -3.0228533 | 4.06682326 |
| TTC5 | 0.122963108 | 0.24623861 | -3.0237026 | 0.60864385 |
| STX18 | 0.122802009 | 0.00016037 | -3.0255939 | 3.79488685 |
| NELFE | 0.121882582 | 0.00123417 | -3.0364361 | 2.90862533 |
| ZMYM3 | 0.121763608 | 0.00412618 | -3.0378451 | 2.38445161 |
| PRDM10 | 0.121370851 | 0.06922047 | -3.0425061 | 1.15976548 |
| DEF8 | 0.121345728 | 0.0880383 | -3.0428048 | 1.05532836 |
| ATF7IP | 0.121258411 | 0.00123043 | -3.0438433 | 2.90994452 |
| ACIN1 | 0.121203388 | 3.8277E-05 | -3.0444981 | 4.41706569 |
| SLU7 | 0.120938386 | 0.00205984 | -3.0476559 | 2.68616643 |
| SUPT20H | 0.12088117 | 0.00536983 | -3.0483386 | 2.27003942 |
| KIAA0586 | 0.120871673 | 0.00577258 | -3.0484519 | 2.2386301 |
| PLA2G6 | 0.12078387 | 0.00461837 | -3.0495003 | 2.3355112 |
| EPC2 | 0.120702204 | 0.00022049 | -3.0504761 | 3.65661276 |
| SCAF4 | 0.120647327 | 0.00028 | -3.0511321 | 3.55284117 |
| MECP2 | 0.120470249 | 0.00072574 | -3.0532512 | 3.13921839 |
| DNMT3A | 0.120410541 | 0.00044309 | -3.0539664 | 3.35350901 |
| AIMP2 | 0.120064029 | 0.0026391 | -3.0581241 | 2.57854378 |
| NCBP2 | 0.119778727 | 0.00104044 | -3.0615564 | 2.98278233 |
| NSD3 | 0.119750771 | 2.3756E-05 | -3.0618931 | 4.6242299 |
| AAAS | 0.119663862 | 9.4897E-05 | -3.0629406 | 4.02274611 |
| TTC14 | 0.119627166 | 0.00239593 | -3.063383 | 2.62052501 |
| FIP1L1 | 0.119204022 | 0.00136043 | -3.0684952 | 2.86632523 |
| LIG3 | 0.119055797 | 0.00096217 | -3.0702902 | 3.01674614 |
| XRCC5 | 0.118967319 | 1.1692E-05 | -3.0713628 | 4.93212272 |
| TRIM33 | 0.118767352 | 0.00173954 | -3.0737898 | 2.75956659 |
| ANK2 | 0.118654022 | 0.0165348 | -3.0751671 | 1.78160109 |
| MYO18A | 0.118507886 | 2.8898E-05 | -3.076945 | 4.53912931 |
| VEZF1 | 0.117779667 | 0.01136015 | -3.0858376 | 1.94461594 |
| MED20 | 0.117011487 | 0.00096709 | -3.0952779 | 3.01453352 |
| SRCAP | 0.116846346 | 5.1637E-06 | -3.0973155 | 5.2870353 |
| JUNB | 0.116685685 | 0.08126992 | -3.0993005 | 1.09007019 |
| ZYG11B | 0.116607893 | 0.01647199 | -3.1002627 | 1.78325404 |

| | | | | |
|---|---|---|---|---|
| SUPT3H | 0.116504851 | 0.04849849 | -3.1015381 | 1.31427177 |
| ISL1 | 0.116477477 | 0.00139907 | -3.1018771 | 2.8541599 |
| CADM1 | 0.116375524 | 0.1369901 | -3.1031404 | 0.86331081 |
| TONSL | 0.116265 | 0.01728714 | -3.1045112 | 1.76227685 |
| INCENP | 0.115562519 | 3.4708E-05 | -3.1132545 | 4.45957479 |
| LIG1 | 0.115232501 | 0.00388016 | -3.1173804 | 2.41115055 |
| PHF3 | 0.115166344 | 1.0178E-05 | -3.1182089 | 4.99234135 |
| SLX4IP | 0.115140806 | 0.02568309 | -3.1185289 | 1.59035264 |
| IVNS1ABP | 0.11498837 | 0.01654341 | -3.1204401 | 1.78137506 |
| HMMR | 0.114916146 | 0.00100302 | -3.1213466 | 2.99868855 |
| DBR1 | 0.11338668 | 0.0037587 | -3.1406769 | 2.42496205 |
| BEND5 | 0.112811147 | 0.12202589 | -3.1480185 | 0.913548 |
| ASH1L | 0.11276496 | 8.1193E-05 | -3.1486093 | 4.09048064 |
| DNAJC17 | 0.110763259 | 0.00026988 | -3.1744487 | 3.56883495 |
| ZNF3 | 0.110619209 | 0.00074158 | -3.1763262 | 3.12984085 |
| ATRX | 0.11038667 | 0.00037598 | -3.1793621 | 3.42483573 |
| TOX4 | 0.110067312 | 6.6321E-06 | -3.183542 | 5.17834803 |
| CLN6 | 0.109908544 | 0.00246716 | -3.1856246 | 2.60780316 |
| NUDCD3 | 0.109771863 | 0.00014633 | -3.1874198 | 3.83467336 |
| METTL14 | 0.109294753 | 6.9231E-05 | -3.193704 | 4.15969856 |
| PQBP1 | 0.1091439 | 0.00382302 | -3.1956966 | 2.4175929 |
| CFDP1 | 0.108933614 | 1.8685E-05 | -3.1984789 | 4.72850528 |
| LRIF1 | 0.10856456 | 0.00177116 | -3.2033749 | 2.7517424 |
| HIVEP1 | 0.108488594 | 0.04416336 | -3.2043847 | 1.35493792 |
| ELAC2 | 0.108391932 | 0.07470167 | -3.2056707 | 1.12666967 |
| UIMC1 | 0.108378377 | 6.3771E-06 | -3.2058512 | 5.19537358 |
| SUCO | 0.10820448 | 0.37526758 | -3.2081679 | 0.42565895 |
| CWF19L2 | 0.107894904 | 0.07571949 | -3.2123014 | 1.1207923 |
| FOXJ3 | 0.107488384 | 0.02106135 | -3.2177473 | 1.67651376 |
| PDS5A | 0.107226518 | 0.00014227 | -3.2212663 | 3.84687675 |
| ZRANB2 | 0.107223779 | 0.00047271 | -3.2213032 | 3.3254066 |
| FARP2 | 0.10696467 | 0.09185425 | -3.2247937 | 1.03690072 |
| SEL1L3 | 0.106484571 | 0.12018233 | -3.2312837 | 0.92015938 |
| ZMYM2 | 0.106007346 | 0.0009363 | -3.2377639 | 3.02858267 |
| SIN3A | 0.105588389 | 0.00165786 | -3.2434769 | 2.78045128 |
| RNPC3 | 0.105436425 | 0.00021483 | -3.2455547 | 3.66791034 |
| SAP130 | 0.10531897 | 0.00016225 | -3.2471628 | 3.78982239 |
| GZF1 | 0.105289233 | 0.00020957 | -3.2475702 | 3.67867253 |
| CHERP | 0.105265443 | 4.3181E-05 | -3.2478962 | 4.36471179 |
| NFRKB | 0.104671347 | 4.5622E-05 | -3.2560615 | 4.34082951 |
| STXBP5L | 0.104402423 | 0.00086731 | -3.2597729 | 3.0618254 |
| SDAD1 | 0.104144403 | 0.08706046 | -3.2633428 | 1.06017907 |
| TMCC3 | 0.104062952 | 0.1730505 | -3.2644715 | 0.76182714 |
| MED16 | 0.103795847 | 0.21047878 | -3.2681794 | 0.67679168 |
| ZNF574 | 0.102713199 | 0.08081446 | -3.2833065 | 1.09251091 |

| | | | | |
|---|---|---|---|---|
| SOAT1 | 0.102439511 | 0.00452315 | -3.2871558 | 2.34455931 |
| NR2C2 | 0.102369512 | 0.0407359 | -3.288142 | 1.39002268 |
| NSD1 | 0.102142384 | 0.02718827 | -3.2913465 | 1.56561843 |
| MGA | 0.101934302 | 4.576E-06 | -3.2942885 | 5.33951217 |
| DNAJC21 | 0.101757027 | 0.01507249 | -3.2967997 | 1.82181507 |
| ZNF281 | 0.101409732 | 0.00077317 | -3.301732 | 3.11172239 |
| GON4L | 0.101283459 | 0.00442783 | -3.3035295 | 2.35380899 |
| ARL3 | 0.10035214 | 0.04035126 | -3.3168567 | 1.39414292 |
| NBEAL2 | 0.100313027 | 0.016413 | -3.3174191 | 1.78481204 |
| HOMER3 | 0.100014825 | 0.00357604 | -3.3217142 | 2.44659784 |
| WDR25 | 0.099587651 | 0.00262796 | -3.3278893 | 2.580381 |
| SATB1 | 0.099523313 | 0.00320784 | -3.3288217 | 2.49378703 |
| CFAP298 | 0.099484864 | 0.00141604 | -3.3293791 | 2.84892557 |
| RIMS1 | 0.099481159 | 0.00041409 | -3.3294329 | 3.38290793 |
| SMARCA2 | 0.099437019 | 0.00135809 | -3.3300731 | 2.86707237 |
| PHKA2 | 0.09925297 | 0.00059374 | -3.3327459 | 3.22640572 |
| SETD1A | 0.099237249 | 0.01036273 | -3.3329745 | 1.98452584 |
| RAD51C | 0.09795467 | 0.00360438 | -3.3517419 | 2.44316969 |
| GEMIN2 | 0.097431319 | 0.00330967 | -3.3594706 | 2.48021517 |
| EIF3M | 0.097380685 | 0.00448042 | -3.3602205 | 2.3486817 |
| TAF12 | 0.0972284 | 0.06193011 | -3.3624784 | 1.20809812 |
| JMJD6 | 0.097033884 | 0.00065037 | -3.3653676 | 3.18683941 |
| SATB2 | 0.095765878 | 0.0001709 | -3.3843445 | 3.7672524 |
| XRCC4 | 0.095169782 | 0.146666 | -3.3933526 | 0.83367054 |
| CERS6 | 0.093983484 | 0.0686369 | -3.4114489 | 1.16344236 |
| CWC15 | 0.093775583 | 0.00174743 | -3.4146439 | 2.75760095 |
| CDKN1B | 0.093361154 | 0.00317107 | -3.4210338 | 2.49879435 |
| ECPAS | 0.093197162 | 0.07383 | -3.4235702 | 1.13176716 |
| SSBP2 | 0.092984775 | 0.08322599 | -3.4268617 | 1.07974104 |
| ITPR2 | 0.092952535 | 0.0061458 | -3.427362 | 2.21142185 |
| ARL13B | 0.092871525 | 0.00509703 | -3.4286199 | 2.29268244 |
| FOXK1 | 0.092864783 | 0.0276675 | -3.4287246 | 1.55803006 |
| PRPF18 | 0.092797794 | 0.01204192 | -3.4297657 | 1.91930437 |
| EML1 | 0.092720752 | 0.00030868 | -3.4309639 | 3.51048839 |
| ZNF749 | 0.09213592 | 9.5998E-05 | -3.4400925 | 4.01773822 |
| MTA1 | 0.092013266 | 3.0826E-05 | -3.4420143 | 4.51108235 |
| ZMPSTE24 | 0.092001648 | 0.00179265 | -3.4421965 | 2.74650453 |
| ORC2 | 0.091733148 | 0.0001226 | -3.446413 | 3.91150385 |
| FOXK2 | 0.091607873 | 0.00549804 | -3.4483846 | 2.25979244 |
| RCOR1 | 0.091561572 | 0.00035526 | -3.449114 | 3.4494502 |
| CORO1C | 0.091439934 | 3.355E-06 | -3.4510318 | 5.47430398 |
| RMND5A | 0.091352361 | 0.66645097 | -3.4524142 | 0.17623179 |
| ASH2L | 0.089802193 | 0.00092256 | -3.4771055 | 3.03500409 |
| GMEB1 | 0.089626951 | 0.00118904 | -3.4799236 | 2.92480475 |
| MIDEAS | 0.089541795 | 0.000263 | -3.481295 | 3.58004836 |

| | | | | |
|---|---|---|---|---|
| RCOR3 | 0.089152117 | 3.8315E-05 | -3.4875871 | 4.41662698 |
| PSMA2 | 0.088997015 | 0.05902264 | -3.4900992 | 1.22898138 |
| OSBPL1A | 0.088634123 | 0.06777854 | -3.495994 | 1.1689078 |
| MTPAP | 0.088560869 | 0.03874237 | -3.4971868 | 1.41181379 |
| DIDO1 | 0.088467466 | 2.656E-05 | -3.4987092 | 4.57576933 |
| IK | 0.087048349 | 1.59E-05 | -3.5220393 | 4.79859696 |
| RSF1 | 0.087010019 | 0.00187005 | -3.5226747 | 2.72814718 |
| PPP4R1 | 0.085517041 | 0.00037718 | -3.5476442 | 3.42345336 |
| LDB1 | 0.084609802 | 0.00010382 | -3.5630314 | 3.98372426 |
| PRCC | 0.08449392 | 0.02173466 | -3.5650086 | 1.66284717 |
| ADCY3 | 0.083833303 | 0.02974241 | -3.5763327 | 1.52662392 |
| INTS6 | 0.083564086 | 0.00038574 | -3.5809732 | 3.41371033 |
| CDCA2 | 0.083470474 | 0.00035564 | -3.5825902 | 3.44898926 |
| CAST | 0.083051706 | 0.00019499 | -3.5898464 | 3.70998389 |
| USP38 | 0.082818705 | 0.11645985 | -3.5938996 | 0.93382377 |
| CDK12 | 0.082027087 | 0.00013549 | -3.6077558 | 3.86810249 |
| PRIM2 | 0.081794124 | 0.01181911 | -3.611859 | 1.92741529 |
| MED4 | 0.081793985 | 7.6322E-05 | -3.6118614 | 4.11734799 |
| HMG20B | 0.081676555 | 0.01626942 | -3.6139342 | 1.78862783 |
| DFFA | 0.081570817 | 0.00390883 | -3.6158031 | 2.40795285 |
| CBFA2T2 | 0.081559238 | 1.0913E-05 | -3.6160079 | 4.96205108 |
| USP16 | 0.079265081 | 0.05547728 | -3.6571707 | 1.25588481 |
| CASP8AP2 | 0.079186098 | 0.00536888 | -3.658609 | 2.27011649 |
| MFAP1 | 0.079139446 | 3.2654E-05 | -3.6594592 | 4.48606391 |
| HTATSF1 | 0.078885646 | 0.00015625 | -3.6640934 | 3.80616885 |
| KAT6B | 0.0781092 | 0.04290538 | -3.6783637 | 1.36748825 |
| CCNA2 | 0.0780539 | 0.01623458 | -3.6793855 | 1.7895589 |
| POLR1D | 0.077940822 | 0.07904238 | -3.681477 | 1.10213998 |
| SAMD4B | 0.077788373 | 0.00177764 | -3.6843017 | 2.75015636 |
| KIF18B | 0.07699715 | 0.01646028 | -3.6990511 | 1.78356272 |
| EIF2AK4 | 0.076846784 | 0.00190808 | -3.7018713 | 2.71940314 |
| MED27 | 0.076603241 | 0.00021337 | -3.7064507 | 3.67086985 |
| PHC2 | 0.076110966 | 0.00117365 | -3.7157519 | 2.93046092 |
| C5orf24 | 0.075502484 | 0.01196391 | -3.7273321 | 1.9221268 |
| ZBTB33 | 0.075416804 | 0.07585239 | -3.7289702 | 1.12003075 |
| CSTF2 | 0.075262619 | 0.0001982 | -3.7319227 | 3.70289055 |
| POLR3B | 0.074600129 | 0.00042174 | -3.7446781 | 3.37495973 |
| PHF2 | 0.074528435 | 0.0035075 | -3.7460652 | 2.45500269 |
| HMBOX1 | 0.074266027 | 0.01860005 | -3.7511538 | 1.73048588 |
| ZNF316 | 0.07422224 | 1.1384E-05 | -3.7520046 | 4.94369993 |
| AAGAB | 0.073793757 | 0.03963999 | -3.7603574 | 1.40186645 |
| ZNF197 | 0.073471678 | 0.00076727 | -3.766668 | 3.11505443 |
| COG5 | 0.072976855 | 0.02611495 | -3.7764172 | 1.58311083 |
| EXOSC3 | 0.072963591 | 0.00022358 | -3.7766795 | 3.65056885 |
| CWC27 | 0.072837701 | 0.00040596 | -3.7791708 | 3.39151601 |

| | | | | |
|---|---|---|---|---|
| CPSF4 | 0.072464265 | 0.00017714 | -3.7865865 | 3.75167384 |
| EP300 | 0.072319251 | 0.08829831 | -3.7894765 | 1.05404763 |
| ETAA1 | 0.072154706 | 8.8838E-05 | -3.7927627 | 4.0514013 |
| IMP3 | 0.071177354 | 5.1265E-05 | -3.8124379 | 4.29017581 |
| PLCB3 | 0.071044884 | 0.05429686 | -3.8151254 | 1.26522527 |
| AFF4 | 0.071039506 | 0.00184781 | -3.8152346 | 2.7333435 |
| DNTTIP1 | 0.070836594 | 0.00779786 | -3.8193613 | 2.10802437 |
| RFX1 | 0.070003395 | 0.02005616 | -3.8364313 | 1.6977523 |
| ICE1 | 0.069471843 | 3.6671E-05 | -3.8474278 | 4.43567176 |
| CRCP | 0.068682428 | 0.0042605 | -3.8639152 | 2.37053944 |
| CALR | 0.068002874 | 0.00013068 | -3.8782605 | 3.88378649 |
| PRPF40B | 0.067844303 | 0.007295 | -3.8816285 | 2.13697457 |
| SF3A1 | 0.067796408 | 0.00016077 | -3.8826474 | 3.7937832 |
| SPTB | 0.067768393 | 0.0691984 | -3.8832436 | 1.15990397 |
| CD2BP2 | 0.067652607 | 0.03697751 | -3.8857107 | 1.43206237 |
| RBM48 | 0.06703039 | 0.002121 | -3.8990409 | 2.67345879 |
| INTS10 | 0.066321923 | 0.00056607 | -3.9143704 | 3.24713111 |
| NSMCE3 | 0.065856511 | 0.00371495 | -3.9245301 | 2.43004664 |
| ZNF148 | 0.065180749 | 2.9743E-05 | -3.9394103 | 4.52662118 |
| ZBTB21 | 0.064149551 | 0.01974201 | -3.962417 | 1.70460866 |
| FHOD1 | 0.063714694 | 0.30696072 | -3.9722301 | 0.5129172 |
| MBD3 | 0.063651344 | 8.801E-06 | -3.9736652 | 5.05546866 |
| RBM5 | 0.062934587 | 0.12302827 | -3.9900031 | 0.90999509 |
| FANCM | 0.062461314 | 0.07129534 | -4.0008933 | 1.14693885 |
| KDM5C | 0.062060695 | 0.00066509 | -4.0101763 | 3.17712186 |
| TAF5L | 0.0619167 | 0.00144051 | -4.0135276 | 2.8414838 |
| TCF12 | 0.061896464 | 0.00158988 | -4.0139992 | 2.79863449 |
| ARHGEF18 | 0.061434818 | 0.0006377 | -4.0247997 | 3.1953863 |
| DTNBP1 | 0.06137608 | 0.00596771 | -4.0261797 | 2.22419242 |
| POLA2 | 0.061156707 | 0.0086834 | -4.0313455 | 2.06131029 |
| ATF7 | 0.060877377 | 6.2379E-05 | -4.03795 | 4.20495923 |
| CHD1 | 0.060670711 | 0.03303704 | -4.042856 | 1.48099882 |
| FAN1 | 0.060327786 | 0.01876036 | -4.0510335 | 1.72675889 |
| GATAD2A | 0.060078673 | 8.0796E-05 | -4.0570032 | 4.09260824 |
| RPS6KA5 | 0.060040848 | 6.6439E-05 | -4.0579118 | 4.177578 |
| ZNF174 | 0.059531043 | 2.5469E-05 | -4.070214 | 4.59398349 |
| SETBP1 | 0.058914257 | 0.05327025 | -4.0852394 | 1.27351525 |
| PLRG1 | 0.058107354 | 0.00115915 | -4.1051354 | 2.93586116 |
| BRD2 | 0.058042638 | 0.01383888 | -4.1067431 | 1.85889905 |
| MBOAT1 | 0.057982078 | 0.00040929 | -4.1082491 | 3.38797155 |
| SUPT4H1 | 0.057975231 | 1.6439E-05 | -4.1084195 | 4.78412031 |
| CLNS1A | 0.057923392 | 0.14919066 | -4.1097101 | 0.82625836 |
| ANAPC4 | 0.057873346 | 0.00719558 | -4.1109571 | 2.14293424 |
| CACNB3 | 0.056372769 | 8.3064E-05 | -4.1488578 | 4.08058936 |
| ZNF607 | 0.055819513 | 0.04123931 | -4.1630867 | 1.38468864 |

| | | | | |
|---|---|---|---|---|
| NUSAP1 | 0.055688461 | 0.00022254 | -4.1664778 | 3.65259718 |
| THOC1 | 0.055601548 | 0.00034857 | -4.1687312 | 3.45770634 |
| KDM3A | 0.055373148 | 1.5604E-05 | -4.1746696 | 4.80675645 |
| SPTY2D1 | 0.054376739 | 0.00496745 | -4.2008665 | 2.30386608 |
| SMARCAL1 | 0.054037092 | 0.0394472 | -4.2099062 | 1.40398386 |
| SF3A2 | 0.053933904 | 1.2545E-05 | -4.2126637 | 4.90153603 |
| UNG | 0.053915908 | 0.02173057 | -4.2131452 | 1.6629288 |
| RLF | 0.053902022 | 0.00088582 | -4.2135168 | 3.05265272 |
| ARL14EP | 0.053792967 | 0.01186812 | -4.2164386 | 1.92561789 |
| TCOF1 | 0.053661257 | 0.05628982 | -4.2199754 | 1.24957013 |
| LIN37 | 0.052754299 | 0.01691445 | -4.2445675 | 1.77174201 |
| IWS1 | 0.052174951 | 0.0004587 | -4.2604988 | 3.33847423 |
| ZNF8 | 0.052126696 | 0.01982159 | -4.2618338 | 1.70286149 |
| GATAD1 | 0.051628914 | 0.0002386 | -4.2756769 | 3.62232804 |
| ATAD5 | 0.05058293 | 0.01791189 | -4.3052056 | 1.74685867 |
| WDR33 | 0.050436198 | 3.3377E-05 | -4.3093967 | 4.47654843 |
| ELL | 0.050373527 | 0.00184406 | -4.3111904 | 2.73422566 |
| NPAT | 0.050348821 | 0.05352948 | -4.3118982 | 1.27140697 |
| ZSCAN26 | 0.050064888 | 0.04072133 | -4.320057 | 1.39017801 |
| MED30 | 0.049863022 | 0.08672414 | -4.3258859 | 1.06186002 |
| ZNF428 | 0.049331304 | 0.09001963 | -4.3413528 | 1.04566278 |
| ZCCHC8 | 0.049093578 | 1.1806E-05 | -4.3483219 | 4.92789639 |
| ZSCAN30 | 0.048760811 | 0.1164155 | -4.3581341 | 0.93398918 |
| ZHX3 | 0.047582286 | 0.00394096 | -4.3934316 | 2.40439782 |
| E2F1 | 0.047240359 | 0.11945449 | -4.4038363 | 0.92279752 |
| MDC1 | 0.046926489 | 5.5498E-06 | -4.4134537 | 5.25572161 |
| NCOA6 | 0.046789037 | 8.9564E-05 | -4.4176857 | 4.04786463 |
| ZNF865 | 0.04641677 | 0.36206375 | -4.4292101 | 0.44121495 |
| ZMYND8 | 0.046145137 | 0.03368767 | -4.4376776 | 1.47252896 |
| ILK | 0.04613113 | 5.0104E-06 | -4.4381156 | 5.30012383 |
| TLK1 | 0.046016994 | 0.03781598 | -4.4416894 | 1.42232468 |
| DPF1 | 0.045027804 | 4.1176E-05 | -4.4730401 | 4.38535856 |
| LPIN3 | 0.044117332 | 9.3883E-06 | -4.5025106 | 5.0274111 |
| NCOR1 | 0.043878047 | 0.00030376 | -4.5103569 | 3.51747264 |
| CENPT | 0.043734327 | 0.10454275 | -4.5150901 | 0.98070608 |
| NOLC1 | 0.043694155 | 0.03450092 | -4.5164159 | 1.4621693 |
| POLR3E | 0.043043895 | 0.0272888 | -4.5380476 | 1.56401554 |
| ABCD4 | 0.042671733 | 0.29095022 | -4.5505755 | 0.53618132 |
| CDCA7L | 0.042549522 | 0.0343448 | -4.5547133 | 1.46413899 |
| XPC | 0.042214627 | 0.00696216 | -4.5661132 | 2.15725624 |
| GTF2E1 | 0.04188811 | 0.02557323 | -4.5773154 | 1.5922144 |
| MPHOSPH8 | 0.041457195 | 0.0005981 | -4.5922337 | 3.22322577 |
| TSHZ3 | 0.041316357 | 0.00357985 | -4.5971431 | 2.44613558 |
| KIAA1143 | 0.041287379 | 0.0391721 | -4.5981554 | 1.40702311 |
| ZNF70 | 0.041211896 | 2.1877E-09 | -4.6007954 | 8.66000639 |

| | | | | |
|---|---|---|---|---|
| TADA3 | 0.041087921 | 0.00033471 | -4.6051419 | 3.47533057 |
| MASTL | 0.040794565 | 0.0011026 | -4.6154792 | 2.95758142 |
| EZH2 | 0.040362224 | 0.00017299 | -4.6308505 | 3.76196923 |
| ZNF346 | 0.040073306 | 0.08152307 | -4.6412147 | 1.08871947 |
| SYNE1 | 0.039454836 | 0.04154877 | -4.663654 | 1.38144183 |
| NR0B1 | 0.039294651 | 2.7071E-05 | -4.6695233 | 4.56749022 |
| AFF1 | 0.038169405 | 0.68141693 | -4.7114395 | 0.16658708 |
| PROX1 | 0.037789045 | 0.00030669 | -4.7258881 | 3.51329446 |
| IQGAP3 | 0.036039127 | 2.7797E-05 | -4.7942921 | 4.5560049 |
| TIPIN | 0.03510026 | 0.01610884 | -4.8323745 | 1.79293578 |
| RGS3 | 0.034947717 | 0.04302595 | -4.838658 | 1.36626951 |
| WAC | 0.033136103 | 0.03616334 | -4.9154522 | 1.44173151 |
| SIN3B | 0.033128392 | 0.01324418 | -4.915788 | 1.87797508 |
| RBM19 | 0.033124761 | 0.00606153 | -4.9159462 | 2.21741779 |
| MIGA1 | 0.033070348 | 0.05911189 | -4.918318 | 1.22832518 |
| MCM9 | 0.032778527 | 0.1166131 | -4.9311052 | 0.93325266 |
| FAM8A1 | 0.032709199 | 0.70023267 | -4.9341598 | 0.15475763 |
| ARK2N | 0.032570167 | 0.02232838 | -4.9403051 | 1.65114282 |
| UPRT | 0.032388373 | 8.64E-05 | -4.9483802 | 4.06348819 |
| ZBTB41 | 0.032223671 | 0.07224807 | -4.9557353 | 1.14117372 |
| SRSF2 | 0.031979304 | 1.1161E-07 | -4.9667177 | 6.95228182 |
| LPIN2 | 0.031934993 | 0.08598273 | -4.9687181 | 1.06558877 |
| MROH1 | 0.031543089 | 0.00876687 | -4.9865323 | 2.05715565 |
| NCAPH2 | 0.031073983 | 0.05024128 | -5.008149 | 1.29893931 |
| ZNF263 | 0.030718671 | 0.01598551 | -5.0247404 | 1.79627355 |
| CEP43 | 0.030709605 | 0.03613433 | -5.0251662 | 1.44208001 |
| ZNF182 | 0.030678637 | 0.14233845 | -5.0266218 | 0.84667776 |
| MIER1 | 0.030646019 | 1.1419E-05 | -5.0281565 | 4.94236185 |
| GRHPR | 0.030639316 | 0.00070901 | -5.0284721 | 3.14934844 |
| PITPNM2 | 0.030349297 | 0.07244515 | -5.0421931 | 1.13999069 |
| ZNF687 | 0.029220891 | 0.00017247 | -5.096856 | 3.76327551 |
| GTF2F1 | 0.029169521 | 0.07078298 | -5.0993945 | 1.15007118 |
| ZBTB10 | 0.028094051 | 0.06039142 | -5.1535915 | 1.21902474 |
| PAPOLG | 0.028027203 | 0.00733379 | -5.1570284 | 2.13467139 |
| SNRNP48 | 0.027558329 | 0.02535877 | -5.1813678 | 1.59587184 |
| DOT1L | 0.027207289 | 0.00571505 | -5.199863 | 2.24298019 |
| KANSL2 | 0.026635614 | 0.0467386 | -5.2304996 | 1.33032433 |
| HMCES | 0.026251683 | 0.00015476 | -5.2514463 | 3.81033157 |
| ACTR6 | 0.025598308 | 0.00436496 | -5.2878078 | 2.36001966 |
| AMBRA1 | 0.024924105 | 0.09686267 | -5.3263145 | 1.01384356 |
| HMG20A | 0.024727006 | 0.02049817 | -5.3377686 | 1.68828481 |
| LRRC42 | 0.024249713 | 0.00442092 | -5.3658885 | 2.35448769 |
| SBNO2 | 0.024133674 | 1.4215E-06 | -5.3728086 | 5.8472449 |
| ZC3HC1 | 0.023556656 | 0.00514282 | -5.4077214 | 2.2887986 |
| SCAI | 0.023511793 | 0.02220755 | -5.4104716 | 1.65349937 |

| | | | | |
|---|---|---|---|---|
| APPL2 | 0.023157179 | 0.01113047 | -5.4323967 | 1.95348654 |
| PAAT | 0.022488099 | 0.03546134 | -5.4746944 | 1.45024491 |
| ZNF569 | 0.021805634 | 0.07379103 | -5.5191552 | 1.13199644 |
| TERF2 | 0.021780286 | 3.5702E-05 | -5.5208333 | 4.44730153 |
| ZNF592 | 0.021542846 | 0.00238317 | -5.5366473 | 2.62284488 |
| MVB12B | 0.02141901 | 0.12371457 | -5.5449644 | 0.90757915 |
| XPO6 | 0.02108585 | 3.399E-05 | -5.567581 | 4.4686529 |
| ZBED5 | 0.020931038 | 2.1944E-05 | -5.5782123 | 4.65867458 |
| LIG4 | 0.020095559 | 0.4691371 | -5.6369795 | 0.32870022 |
| MTMR12 | 0.020074034 | 1.2662E-06 | -5.6385256 | 5.89750838 |
| RPF1 | 0.01947646 | 0.10267662 | -5.6821247 | 0.98852842 |
| GCH1 | 0.019447604 | 6.8969E-06 | -5.6842638 | 5.16134853 |
| CCDC149 | 0.019377205 | 0.00050317 | -5.6894957 | 3.29828866 |
| ZNF260 | 0.018545179 | 1.3212E-06 | -5.752812 | 5.87902411 |
| SKOR1 | 0.018347411 | 3.5545E-06 | -5.7682797 | 5.44921808 |
| MLLT6 | 0.018332148 | 0.04810975 | -5.7694803 | 1.31776692 |
| ZNF180 | 0.018235621 | 1.5679E-05 | -5.7770969 | 4.8046796 |
| KANSL1 | 0.01811847 | 0.02433275 | -5.786395 | 1.61380887 |
| SETD5 | 0.018105497 | 1.415E-06 | -5.7874284 | 5.84924641 |
| ZNF672 | 0.018003731 | 0.61744951 | -5.7955603 | 0.20939855 |
| SMTN | 0.017999044 | 0.00094636 | -5.7959359 | 3.02394244 |
| HPCA | 0.01792247 | 0.05645432 | -5.8020867 | 1.24830283 |
| ANKRD12 | 0.017574448 | 0.00869634 | -5.8303768 | 2.06066338 |
| MRTFA | 0.017436438 | 0.11980512 | -5.8417509 | 0.92152463 |
| GTF2E2 | 0.017298441 | 0.01280371 | -5.8532141 | 1.89266429 |
| TCEAL1 | 0.017228506 | 0.02091997 | -5.8590586 | 1.67943891 |
| KIF18A | 0.017164804 | 2.028E-06 | -5.8644028 | 5.69293017 |
| CEP295 | 0.017052587 | 0.11156727 | -5.8738656 | 0.95246318 |
| DNAJC8 | 0.01684037 | 0.01608643 | -5.8919324 | 1.79354036 |
| WDFY3 | 0.016823592 | 5.2093E-06 | -5.8933705 | 5.28322372 |
| SLC35F5 | 0.016472222 | 0.16476312 | -5.923821 | 0.78313998 |
| FOXO6 | 0.016417766 | 0.51487654 | -5.9285983 | 0.2882969 |
| ZNF658 | 0.016355509 | 1.3774E-05 | -5.9340795 | 4.86094265 |
| CLOCK | 0.016348032 | 0.0271017 | -5.9347392 | 1.5670035 |
| CRLF3 | 0.016127182 | 0.01231085 | -5.9543618 | 1.90971203 |
| TRIM45 | 0.015915231 | 0.08012413 | -5.9734481 | 1.09623667 |
| KMT2C | 0.015828217 | 0.00026159 | -5.9813574 | 3.58237817 |
| JADE3 | 0.015795862 | 0.00038701 | -5.9843095 | 3.41227431 |
| ESPL1 | 0.015551371 | 1.7865E-05 | -6.0068144 | 4.74798568 |
| ZNF24 | 0.015494159 | 0.00165564 | -6.0121317 | 2.7810336 |
| WDR43 | 0.015386899 | 0.00393195 | -6.0221537 | 2.40539214 |
| ZBTB2 | 0.015300556 | 1.1201E-06 | -6.0302722 | 5.95075017 |
| GCLC | 0.015241741 | 0.12608187 | -6.0358284 | 0.89934737 |
| PRMT7 | 0.015171585 | 0.03777711 | -6.0424844 | 1.42277123 |
| UAP1L1 | 0.01499858 | 0.0007932 | -6.0590303 | 3.10061961 |

| | | | | |
|---|---|---|---|---|
| TUT1 | 0.014986155 | 0.00225098 | -6.0602259 | 2.64762774 |
| ZNF606 | 0.014978464 | 0.02755635 | -6.0609665 | 1.55977831 |
| AATF | 0.014920707 | 0.00103231 | -6.0665403 | 2.98619194 |
| WAPL | 0.014888653 | 0.00020871 | -6.0696429 | 3.68045453 |
| MED7 | 0.014768407 | 0.00826755 | -6.081342 | 2.08262297 |
| LRRC58 | 0.014674375 | 0.00021822 | -6.0905571 | 3.66110455 |
| EYA3 | 0.014519106 | 0.00975925 | -6.1059036 | 2.0105835 |
| HDAC9 | 0.014049895 | 0.11755938 | -6.1532969 | 0.9297427 |
| ANKRD54 | 0.014032491 | 0.02432755 | -6.155085 | 1.61390158 |
| SMG6 | 0.013950185 | 0.01454646 | -6.1635719 | 1.83724275 |
| ZNF544 | 0.013905429 | 0.00471091 | -6.1682079 | 2.32689503 |
| KMT5B | 0.013897841 | 0.05648991 | -6.1689954 | 1.2480291 |
| DCLRE1A | 0.0138549 | 1.7756E-05 | -6.1734599 | 4.75065156 |
| MSL3 | 0.013854767 | 4.1422E-06 | -6.1734737 | 5.38277178 |
| BTBD10 | 0.01355294 | 0.01736766 | -6.2052503 | 1.76025869 |
| SLC8A2 | 0.01349202 | 0.12843506 | -6.2117499 | 0.89131641 |
| ZNF696 | 0.01328053 | 5.477E-06 | -6.2345434 | 5.26145512 |
| RASAL2 | 0.013169904 | 0.03006877 | -6.2466114 | 1.5218844 |
| MLLT3 | 0.012998153 | 0.00021145 | -6.2655496 | 3.67479636 |
| FOCAD | 0.012979738 | 0.04254054 | -6.2675949 | 1.37119704 |
| LOC122539214 | 0.012722487 | 1.1272E-06 | -6.2964755 | 5.94801002 |
| KAT5 | 0.012605321 | 0.00017034 | -6.3098233 | 3.76867183 |
| ZMYM1 | 0.012480964 | 7.3912E-05 | -6.3241268 | 4.13128519 |
| TAF8 | 0.012436004 | 1.6029E-05 | -6.3293332 | 4.79509614 |
| DOCK1 | 0.012270862 | 0.27684362 | -6.3486196 | 0.55776548 |
| HDAC3 | 0.012179309 | 0.02498287 | -6.359424 | 1.60235773 |
| MBD5 | 0.011893145 | 0.05799468 | -6.3937259 | 1.23661183 |
| SET | 0.011834615 | 0.00325242 | -6.4008434 | 2.48779362 |
| RFX5 | 0.011655192 | 0.06209788 | -6.4228834 | 1.20692326 |
| CCNH | 0.011434859 | 0.09061227 | -6.4504176 | 1.04281299 |
| SLC7A6 | 0.011370137 | 1.2559E-06 | -6.4586065 | 5.90105558 |
| CCDC82 | 0.011341988 | 5.4698E-06 | -6.4621827 | 5.2620313 |
| PARP16 | 0.011137038 | 0.04190747 | -6.4884907 | 1.37770856 |
| WDR74 | 0.010940806 | 1.3528E-08 | -6.5141372 | 7.86877207 |
| ZNF264 | 0.010835133 | 0.01368936 | -6.5281394 | 1.863617 |
| ITPRIPL1 | 0.010673228 | 8.1233E-06 | -6.5498596 | 5.09026984 |
| MIS18BP1 | 0.010660608 | 0.10156137 | -6.5515665 | 0.99327146 |
| ITPRIP | 0.010624187 | 3.7222E-05 | -6.5565038 | 4.42919803 |
| MED9 | 0.010592116 | 1.3532E-05 | -6.5608653 | 4.86864414 |
| AKAP2 | 0.010487498 | 0.6556616 | -6.5751857 | 0.18332025 |
| NSRP1 | 0.010424784 | 0.00449363 | -6.5838387 | 2.34740275 |
| RXFP3 | 0.010174236 | 0.12248383 | -6.6189357 | 0.91192125 |
| FBXL12 | 0.010099394 | 0.0006078 | -6.6295874 | 3.21623825 |
| YY1 | 0.010004441 | 0.0692488 | -6.6432156 | 1.15958772 |
| CDAN1 | 0.009957174 | 0.00073247 | -6.650048 | 3.13521153 |

| | | | | |
|---|---|---|---|---|
| ZNF787 | 0.0099149 | 0.01711679 | -6.656186 | 1.76657771 |
| ZKSCAN8 | 0.00986025 | 0.00280726 | -6.6641601 | 2.55171769 |
| KMT2B | 0.009751259 | 1.8147E-05 | -6.6801958 | 4.74119064 |
| DTNA | 0.009347325 | 0.13562023 | -6.7412307 | 0.86767552 |
| SPNB4 | 0.009312295 | 2.9486E-06 | -6.7466475 | 5.53038099 |
| GCFC2 | 0.008886172 | 0.00245982 | -6.8142222 | 2.60909724 |
| RAD54L2 | 0.008771358 | 4.1622E-07 | -6.8329841 | 6.38067826 |
| DDB1 | 0.008760006 | 0.00539091 | -6.8348524 | 2.26833789 |
| ANGEL2 | 0.008724677 | 1.8998E-05 | -6.8406826 | 4.72129571 |
| ATF2 | 0.00857283 | 3.4622E-05 | -6.8660128 | 4.46064491 |
| ANKHD1 | 0.008543251 | 1.187E-08 | -6.8709991 | 7.925536 |
| TENT4A | 0.008235363 | 0.00011834 | -6.923952 | 3.9268603 |
| ASMTL | 0.00821636 | 0.04005092 | -6.9272849 | 1.39738755 |
| POMP | 0.008208314 | 2.7389E-06 | -6.9286984 | 5.56242627 |
| BORCS5 | 0.008109687 | 2.96E-06 | -6.9461381 | 5.52870265 |
| YJU2 | 0.008106092 | 0.00812668 | -6.9467776 | 2.0900869 |
| GID8 | 0.008024725 | 0.11364028 | -6.9613322 | 0.9444677 |
| ZNF518A | 0.008000306 | 5.6357E-06 | -6.9657291 | 5.24904841 |
| SOX2 | 0.007967012 | 0.06673668 | -6.9717455 | 1.17563543 |
| RBM42 | 0.007897872 | 2.0145E-07 | -6.9843203 | 6.69582745 |
| TP73 | 0.007893318 | 4.9043E-08 | -6.9851523 | 7.3094231 |
| FBXW11 | 0.007745336 | 0.07227977 | -7.0124565 | 1.14098325 |
| RBAK | 0.00760331 | 2.5633E-06 | -7.0391566 | 5.59120793 |
| ZMYND11 | 0.007559165 | 0.13214505 | -7.0475575 | 0.8789491 |
| NAA40 | 0.0075264 | 0.16384711 | -7.0538244 | 0.7855612 |
| SPINDOC | 0.007382871 | 0.04351675 | -7.0816023 | 1.36134357 |
| ZFP3 | 0.007338147 | 4.3033E-06 | -7.0903684 | 5.36619339 |
| ATXN7L1 | 0.007326436 | 0.00212566 | -7.0926727 | 2.67250565 |
| APEX2 | 0.007111414 | 2.9116E-05 | -7.1356478 | 4.53586417 |
| ZEB2 | 0.006890932 | 0.06706932 | -7.1810851 | 1.17347607 |
| FGFR1OP2 | 0.006886151 | 9.1441E-07 | -7.1820865 | 6.03885678 |
| CASP9 | 0.006828184 | 0.0652074 | -7.1942823 | 1.18570311 |
| ETV6 | 0.006811427 | 0.04315624 | -7.1978273 | 1.36495638 |
| EDRF1 | 0.006802226 | 1.9522E-06 | -7.1997773 | 5.7094845 |
| DHX40 | 0.006789986 | 0.17058464 | -7.2023756 | 0.76806009 |
| MTFR1 | 0.006784459 | 0.00100992 | -7.2035504 | 2.99571365 |
| TADA2B | 0.006729738 | 5.2959E-08 | -7.215234 | 7.27605625 |
| RB1 | 0.006727004 | 0.0605123 | -7.2158203 | 1.21815635 |
| DNAJC2 | 0.006712975 | 0.01532442 | -7.218832 | 1.81461601 |
| MED25 | 0.006690414 | 3.5572E-06 | -7.2236888 | 5.44889696 |
| PHC1 | 0.006570726 | 0.05263765 | -7.2497316 | 1.2787035 |
| POLK | 0.006490011 | 0.04318271 | -7.2675635 | 1.3646901 |
| PRDM15 | 0.006450773 | 0.00104031 | -7.2763123 | 2.98283745 |
| THAP12 | 0.006446089 | 0.15844818 | -7.2773602 | 0.80011273 |
| ZNF589 | 0.006411928 | 0.00232665 | -7.2850261 | 2.6332696 |

| | | | | |
|---|---|---|---|---|
| MLLT10 | 0.006407059 | 0.00011102 | -7.286122 | 3.9546175 |
| WDR55 | 0.006348029 | 4.7898E-07 | -7.2994756 | 6.31968587 |
| CHM | 0.006321224 | 1.0706E-05 | -7.3055803 | 4.97038712 |
| DUS3L | 0.006286455 | 0.0412112 | -7.3135376 | 1.38498478 |
| CHKA | 0.006233971 | 0.15992243 | -7.3256327 | 0.79609063 |
| ACBD6 | 0.005997892 | 0.12523827 | -7.3813287 | 0.90226293 |
| DLX3 | 0.005982571 | 7.1831E-05 | -7.3850187 | 4.1436886 |
| NOL4L | 0.005955929 | 0.02890192 | -7.3914576 | 1.5390733 |
| FAM76A | 0.005835004 | 1.9849E-07 | -7.4210507 | 6.70226326 |
| KDM2A | 0.005831868 | 1.329E-06 | -7.4218261 | 5.87648649 |
| ASCC2 | 0.00582979 | 0.00301455 | -7.4223404 | 2.52077803 |
| DTL | 0.005807057 | 0.03446937 | -7.4279772 | 1.46256669 |
| VPS26C | 0.005783963 | 4.3623E-07 | -7.433726 | 6.36028207 |
| SLF1 | 0.005776901 | 0.02820418 | -7.4354886 | 1.54968651 |
| SREBF2 | 0.005741743 | 0.05980575 | -7.4442956 | 1.22325709 |
| VPS35 | 0.005624633 | 0.06104467 | -7.4740254 | 1.21435228 |
| PPP1R12B | 0.005554229 | 9.278E-06 | -7.4921977 | 5.03254715 |
| MED10 | 0.005505092 | 4.3323E-05 | -7.5050175 | 4.36328561 |
| TRMT11 | 0.005468256 | 0.01306139 | -7.5147036 | 1.8840106 |
| ZWINT | 0.005432913 | 0.00836081 | -7.5240584 | 2.07775185 |
| INO80D | 0.005427743 | 2.8983E-06 | -7.5254319 | 5.53785235 |
| KRIT1 | 0.005251245 | 2.6785E-06 | -7.5731248 | 5.57210463 |
| MIER2 | 0.005250639 | 2.5351E-06 | -7.5732912 | 5.59599779 |
| PDPK1 | 0.005098931 | 1.5077E-05 | -7.6155894 | 4.82168829 |
| MSL3P1 | 0.005095724 | 0.00133116 | -7.6164971 | 2.87576919 |
| PSMG3 | 0.00499021 | 6.3198E-07 | -7.6466838 | 6.19929444 |
| CHAF1B | 0.004898197 | 0.02556878 | -7.6735334 | 1.59228993 |
| AVEN | 0.004827328 | 0.00026569 | -7.6945594 | 3.57562691 |
| METAP1 | 0.004824191 | 4.9458E-06 | -7.6954972 | 5.30576146 |
| MED26 | 0.004740055 | 2.1071E-07 | -7.7208805 | 6.6763228 |
| RPP38 | 0.004710499 | 0.0108026 | -7.7299043 | 1.96647166 |
| CYB5B | 0.004689062 | 0.08526558 | -7.7364849 | 1.06922627 |
| HNRNPH1 | 0.004597801 | 0.1164247 | -7.7648403 | 0.93395488 |
| HDAC5 | 0.004573856 | 0.00016984 | -7.7723735 | 3.76996821 |
| COG6 | 0.004562405 | 0.06375278 | -7.7759899 | 1.19550089 |
| SNAPC4 | 0.00439754 | 8.7268E-06 | -7.8290876 | 5.05914576 |
| TK1 | 0.004356992 | 0.06478551 | -7.8424519 | 1.1885221 |
| ZNF483 | 0.004286369 | 0.05951176 | -7.8660283 | 1.22539724 |
| RFX7 | 0.004239366 | 1.8297E-08 | -7.8819358 | 7.73762451 |
| SPECC1 | 0.00419782 | 3.9186E-06 | -7.8961441 | 5.40686996 |
| USE1 | 0.004190267 | 2.4062E-05 | -7.8987421 | 4.61867713 |
| PIAS2 | 0.004148261 | 0.01881976 | -7.9132778 | 1.72538599 |
| AP3D1 | 0.004106214 | 0.01540426 | -7.9279756 | 1.81235916 |
| MREG | 0.004088337 | 0.03355189 | -7.9342703 | 1.47428296 |
| SKI | 0.004062077 | 0.18944607 | -7.9435666 | 0.72251441 |

| | | | | |
|---|---|---|---|---|
| MCRS1 | 0.004061313 | 4.9003E-05 | -7.9438382 | 4.30977824 |
| ATN1 | 0.004058597 | 0.00120443 | -7.9448032 | 2.91921723 |
| EXOC8 | 0.004042775 | 0.06925872 | -7.9504385 | 1.15952551 |
| FER | 0.003951505 | 0.0231915 | -7.983382 | 1.63467113 |
| SPC25 | 0.003950309 | 3.6762E-09 | -7.9838189 | 8.43460021 |
| C2orf49 | 0.003856682 | 1.7449E-06 | -8.018424 | 5.75822503 |
| MIS12 | 0.003797147 | 5.7001E-07 | -8.0408686 | 6.2441155 |
| ATXN7L2 | 0.00376628 | 3.5581E-06 | -8.0526441 | 5.44878579 |
| REXO1 | 0.003764193 | 0.0157639 | -8.0534436 | 1.80233637 |
| COMMD8 | 0.003701166 | 0.01564111 | -8.0778045 | 1.8057325 |
| CKAP2L | 0.003697214 | 1.9434E-06 | -8.0793457 | 5.7114468 |
| PABIR2 | 0.003623551 | 2.1174E-06 | -8.1083803 | 5.67419838 |
| FAM32A | 0.003576559 | 5.5157E-06 | -8.1272121 | 5.25840268 |
| DHX34 | 0.003532349 | 8.4203E-05 | -8.1451565 | 4.0746733 |
| ING5 | 0.003506277 | 3.3554E-06 | -8.1558444 | 5.47425614 |
| PPP1R3F | 0.003490225 | 0.00330163 | -8.1624641 | 2.48127127 |
| ERCC6 | 0.003482019 | 0.00302414 | -8.1658601 | 2.51939842 |
| BNIP3 | 0.003474772 | 0.01392506 | -8.1688662 | 1.85620296 |
| KDM4C | 0.003425476 | 1.6476E-08 | -8.1894797 | 7.7831583 |
| STXBP5 | 0.00339803 | 0.00131655 | -8.2010856 | 2.88056195 |
| MYSM1 | 0.0033788 | 0.03962347 | -8.2092735 | 1.40204752 |
| FOXP4 | 0.003271232 | 0.06500931 | -8.2559501 | 1.18702446 |
| POLR1G | 0.003229955 | 2.5274E-08 | -8.2742704 | 7.59732405 |
| DPF3 | 0.003133821 | 0.27085471 | -8.3178613 | 0.56726361 |
| CLIP2 | 0.003095674 | 0.00869036 | -8.3355307 | 2.06096246 |
| NT5C2 | 0.00300358 | 0.00957953 | -8.3791012 | 2.01865599 |
| CEBPG | 0.002993082 | 0.03408612 | -8.3841524 | 1.46742242 |
| USP3 | 0.002973003 | 3.132E-06 | -8.3938635 | 5.50417181 |
| LPGAT1 | 0.002958807 | 0.00010591 | -8.4007685 | 3.97506343 |
| MAP7D3 | 0.002921835 | 1.2661E-06 | -8.4189098 | 5.897534 |
| ZNF460 | 0.002920536 | 0.01719779 | -8.4195512 | 1.76452727 |
| MPG | 0.002871064 | 0.00713735 | -8.4441987 | 2.14646316 |
| ATAD1 | 0.002862313 | 0.02135475 | -8.4486028 | 1.67050549 |
| E2F6 | 0.002805358 | 7.6545E-09 | -8.4775996 | 8.11608068 |
| NAP1L5 | 0.002777842 | 0.02780735 | -8.4918195 | 1.55584039 |
| MED13 | 0.002730131 | 0.03220549 | -8.516814 | 1.49207003 |
| PABIR1 | 0.002684753 | 0.02465117 | -8.540995 | 1.60816243 |
| TFAP4 | 0.002653617 | 0.00176777 | -8.5578242 | 2.75257514 |
| ZNF630 | 0.002650012 | 0.0042155 | -8.5597854 | 2.37515089 |
| BANP | 0.00263469 | 9.6679E-06 | -8.5681513 | 5.01466669 |
| C12orf57 | 0.002612475 | 3.7886E-05 | -8.5803672 | 4.42152211 |
| ZC3H6 | 0.002586862 | 2.9115E-08 | -8.5945811 | 7.5358892 |
| CHMP3 | 0.002531705 | 0.12916245 | -8.625675 | 0.88886374 |
| E2F3 | 0.002396017 | 0.04946871 | -8.7051459 | 1.3056694 |
| TSC22D2 | 0.002388328 | 2.5674E-06 | -8.7097834 | 5.59051357 |

| | | | | |
|---|---|---|---|---|
| TMEM38B | 0.002322859 | 1.6876E-06 | -8.7498828 | 5.7727429 |
| MED6 | 0.002272318 | 2.9391E-09 | -8.7816198 | 8.53178157 |
| ZFYVE19 | 0.002259905 | 4.8049E-09 | -8.7895221 | 8.31831557 |
| SCNM1 | 0.002257502 | 0.01201184 | -8.7910571 | 1.92039028 |
| ASXL2 | 0.002230172 | 0.0369514 | -8.8086292 | 1.43236912 |
| ACTR10 | 0.002228811 | 0.01015376 | -8.8095103 | 1.99337303 |
| ARNT | 0.002190522 | 0.0002442 | -8.8345095 | 3.61224989 |
| UXT | 0.002150767 | 9.3244E-08 | -8.8609327 | 7.0303777 |
| JUND | 0.002116602 | 3.471E-07 | -8.8840345 | 6.45954242 |
| ZBTB18 | 0.002113136 | 0.02787756 | -8.8863985 | 1.55474524 |
| SLX9 | 0.002050966 | 3.8754E-05 | -8.9294805 | 4.4116801 |
| CWF19L1 | 0.001989187 | 2.3925E-06 | -8.9736052 | 5.62114094 |
| KDM1B | 0.001979691 | 0.02244011 | -8.9805088 | 1.64897503 |
| DDX11L8 | 0.001903798 | 0.04319494 | -9.0369036 | 1.36456717 |
| ZBED4 | 0.001864858 | 1.1605E-06 | -9.0667182 | 5.93537171 |
| CYB5R4 | 0.0018568 | 0.07555243 | -9.072966 | 1.12175159 |
| HERC1 | 0.001856265 | 6.2362E-08 | -9.073382 | 7.2050828 |
| USP39 | 0.001843771 | 0.07900298 | -9.0831247 | 1.1023565 |
| TCEANC2 | 0.001822752 | 1.8518E-05 | -9.0996664 | 4.73241532 |
| HOXD8 | 0.001820589 | 1.8249E-05 | -9.1013792 | 4.73875527 |
| SLMAP | 0.001781459 | 0.06083629 | -9.1327247 | 1.2158373 |
| HMGN5 | 0.00176865 | 3.7243E-06 | -9.1431355 | 5.42895673 |
| DDB2 | 0.001749982 | 0.00040592 | -9.1584445 | 3.39156419 |
| OGA | 0.001737968 | 0.00188109 | -9.168383 | 2.72559066 |
| CENPU | 0.001712734 | 0.04008855 | -9.1894833 | 1.39697969 |
| BEND3 | 0.001709126 | 1.6671E-06 | -9.1925259 | 5.7780262 |
| PCM1 | 0.001662944 | 0.02231941 | -9.2320443 | 1.6513173 |
| CHD1L | 0.001594849 | 9.4021E-07 | -9.2923648 | 6.02677393 |
| POGK | 0.001565486 | 6.5218E-07 | -9.3191733 | 6.1856314 |
| CCDC97 | 0.001484708 | 2.1494E-05 | -9.3956054 | 4.66768737 |
| ZHX1 | 0.001480561 | 0.01928792 | -9.3996401 | 1.71471458 |
| MBD2 | 0.001471333 | 3.091E-08 | -9.4086608 | 7.50990001 |
| RABEP2 | 0.001466865 | 0.00474838 | -9.413048 | 2.32345408 |
| ZNRD2 | 0.001425498 | 1.0796E-07 | -9.4543185 | 6.96673633 |
| TCEA1 | 0.001411099 | 2.5514E-10 | -9.4689653 | 9.59322362 |
| DYNLT3 | 0.001336357 | 8.0241E-08 | -9.5474786 | 7.09560141 |
| HNRNPC | 0.001329326 | 0.04890325 | -9.5550899 | 1.31066232 |
| GMEB2 | 0.001298306 | 4.3618E-07 | -9.5891539 | 6.36033582 |
| ZNF385A | 0.001271852 | 2.6242E-07 | -9.6188534 | 6.58100362 |
| PIAS4 | 0.001233109 | 1.1562E-06 | -9.6634835 | 5.93698343 |
| SENP7 | 0.001209957 | 0.00987576 | -9.6908288 | 2.00542961 |
| MECP2 | 0.001206751 | 0.0236148 | -9.6946565 | 1.62681567 |
| TCP11L1 | 0.001133668 | 0.00345067 | -9.7847861 | 2.46209664 |
| PHF12 | 0.001004364 | 0.0219826 | -9.9595016 | 1.65792102 |
| ZMAT4 | 0.000958875 | 0.00016448 | -10.026369 | 3.78388632 |

| | | | | |
|---|---|---|---|---|
| TAF9 | 0.000862358 | 1.9497E-09 | -10.179425 | 8.71002605 |
| ZNF280D | 0.000862074 | 0.03182066 | -10.179901 | 1.49729087 |
| SUDS3 | 0.000809765 | 0.00542807 | -10.270208 | 2.26535431 |
| INTS14 | 0.000686372 | 4.2143E-07 | -10.508722 | 6.37527805 |
| GPATCH11 | 0.000586383 | 7.0605E-07 | -10.73587 | 6.15116617 |
| TCEAL3 | 0.000584773 | 0.01138064 | -10.739835 | 1.94383345 |
| SALL2 | 0.000525677 | 0.01120757 | -10.893536 | 1.95048856 |
| SETMAR | 0.000492296 | 0.02243254 | -10.988188 | 1.64912161 |
